# Zoospore-derived extracellular vesicles in the flagellated stramenopile *Phytophthora parasitica*

**DOI:** 10.64898/2026.09.11.750887

**Authors:** Marie-Line Kuhn, Carlotta Lupatelli, Blandine Madji Hounoum, Aurélie Seassau, Anne-Sophie Gay, Sophie Pagnotta, François Orange, Ilaria Bassani, Eric Galiana

## Abstract

Zoospores are unicellular, wall-less and flagellated cells produced by a number of eukaryotic microorganisms. They allow microbial dispersion, enable the search and location for new sources of nutrients where they aggregate, initiate pathogen-host interactions and communication within microbiota. Little is known about their ability to release bioactive extracellular vesicles (EVs) supporting these adaptations. Here we used electron microscopy to establish that in the biflagellate zoospores of the heterokont and phytopathogenic species *Phytophthora parasitica,* EV biogenesis occurs from vesicles budding either at cell body plasma membrane or at the front flagellum, in particular from the tubular mastigonemes. Zoospore-conditioned water supernatant was fractionated and characterized by means of morphological, immunochemical, proteomic and lipidomic analyses. Three fractions enriched in cell body, flagella and EVs were obtained by differential ultracentrifugation at 3,000g, 31,000g and 100,000g, respectively, with the EV fraction consisting of small vesicles (100-150 nm). Using mass spectrometry, label-free proteomic analysis of the EV fraction (1,470 proteins) revealed a collection of proteins involved in lipid transport, vesicle membrane and cell wall organization, and tubular mastigoneme architecture, but not in virulence. Label-free quantitative lipidomic analysis (191 lipids) revealed enrichment of EVs in sphingolipids, particularly ceramides compared with the cell body. These findings provide a first hallmark of features characteristic of zoosporic EVs, with ceramides, mastigoneme proteins and the EV-related protein PPTG_13069 containing two tetraspanin MARVELous domains among molecular markers. They define the molecular and cellular principles in understanding zoosporic EV biogenesis within Stramenopiles and intercellular communication between self-aggregating zoospores, with host plant cells or microbiota.

**GRAPHICAL ABSTRACT:** 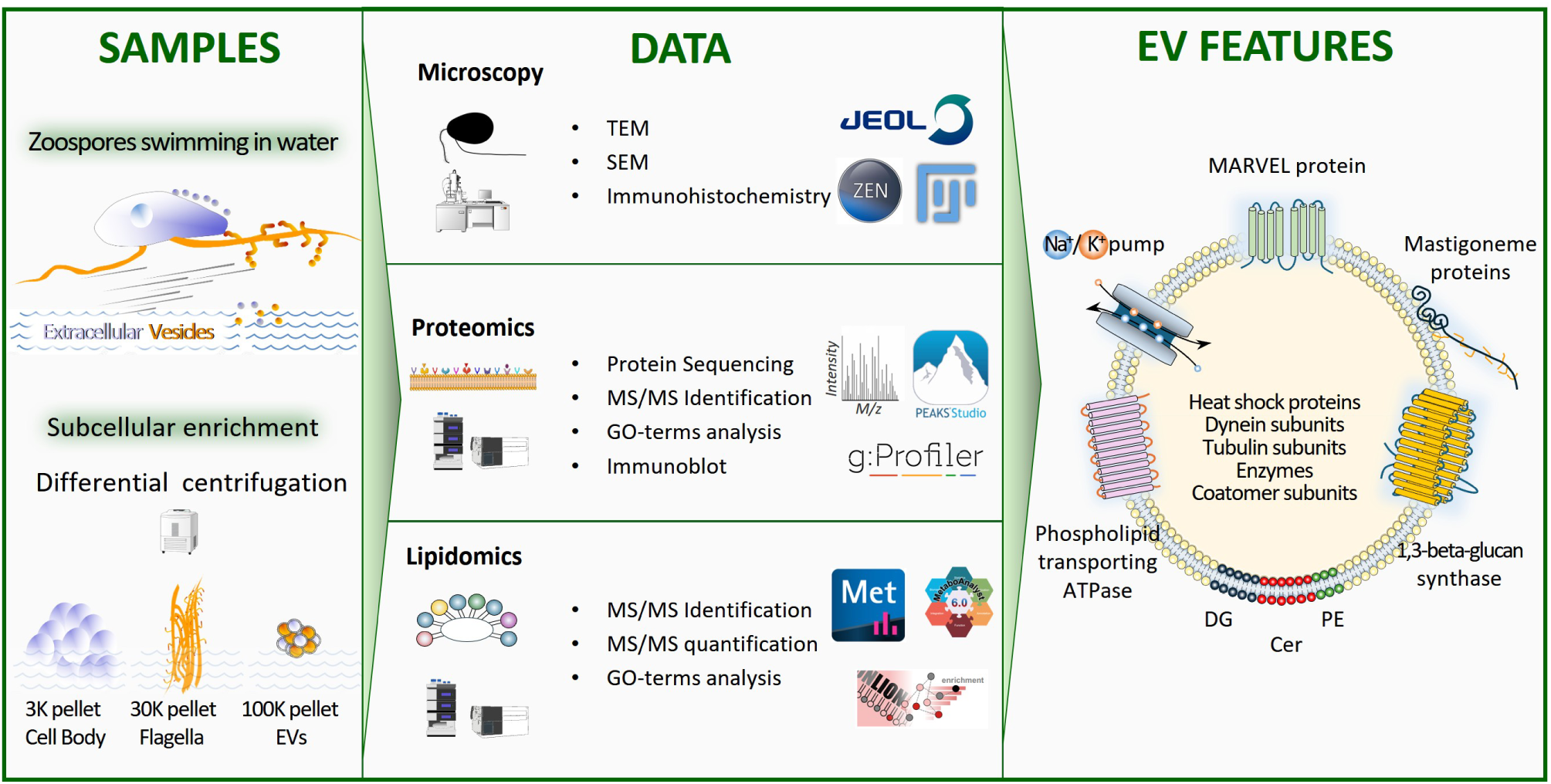

## INTRODUCTION

Zoospores are a category of motile spores produced by stramenopiles, algae and fungi. They are surrounded by a plasma membrane, devoid of cell wall and have one or more flagella, which allow them to swim, explore and disperse microoorganisms to new moist environments. Biflagellate zoospores play important roles in the living cycle of plant pathogen species of the *Phytophthora* genus with regard to dissemination and subsequent infection. During a storm, they are released in soils from sporangia as a high-density, moving inoculum targeting the host plant. Once the root surface is reached, zoospores aggregate, lose the two flagella, discharge adhesive molecules, and synthesize a primary cell wall that will result in the transition to walled-cysts (Bassani *et al*., 2020a; Kasteel *et al*., 2023). The cyst initiates germination and filamentous growth allowing plant penetration and hyphal colonization. Recent studies have indicated the occurrence and the importance of extracellular vesicles (EVs) in *Phytophthora* species *P. capsici* and *P. infestans,* during filamentous and *in planta* growth (Fang *et al*., 2021; Breen *et al*., 2025). Nonetheless, the EVs released by *Phytophthora* zoospores, and more broadly by zoospores of stramenopiles, algae, and fungi, remain poorly characterized.

EVs, such as ectosomes and exosomes, are a mixture of membranous structures discharged by cells. They contain DNA, RNA, metabolites and proteins encased in a phospholipid bilayer (Théry *et al*., 2018, Welsh *et al*., 2024). Exosomes are released during exocytosis from multivesicular bodies (MVBs) filled with intraluminal vesicles (ILVs). Ectosomes are generated by outward budding of plasma membrane followed by pinching off at the base prior extracellular release. In the case of cells harbouring flagella/cilia, EV release may also come from the cell body as from the flagellar/ciliar membranes (Vinay and Belleannée; 2022; Nachury *et al.,* 2019). Cilia-derived EVs (cEVs) biogenesis takes place either at the tip or laterally (Wood and Rosenbaum, 2015; Szempruch *et al*., 2016). In *Chlamydomonas reinhardtii*, flagellar tips shed ectosomes occurring directly from flagellum membrane protrusions (Wood and Rosenbaum, 2015). Bioactive ciliary ectosomes secreted from mating gametes contain all the components for peptidergic signaling (Luxmi and Kin, 2022). Thus, ciliary EVs appear as a functional component of flagella/cilia for sensing and transducing signals (Wang and Barr, 2018).

EV release from donor cells and delivery of their intralumenal content into the cytoplasm of recipient cells regulate various physiological and pathological functions. They are involved in local cell-to-cell communications or systemic signalling through displacement driven by bodily fluid fluxes (O’Brien *et al*., 2020). They also play an important role during host-pathogen interactions. EV release may be used by a host as a strategy to control disease. For example, EVs produced by the host plant *Arabidopsis* transport mRNAs to the *Botrytis cinerea* colonising mycelium where they are translated to compromise infection (Wang *et al*. 2024). EV release may also aid in pathogenesis. *Phytophthora capsici* disrupts plant defense by selectively deploying lipase activity against Arabidopsis EVs (Xu *et al*., 2026). A mix of high concentrated EVs purified a from mycelium and zoospores enhances the virulence of *P. capsici (*Fang *et al*., 2021). In *Trypanosoma brucei*, EVs derived from membranous nanotubes that originate from the flagellum membrane were shown to mediate virulence factor transfer and cause host anemia (Szempruch *et al*. 2016^a,b^).

Zoospores are likely to release EVs from two main sources: the cell body and the flagella. Mastigonemes, filamentous appendages attached to the flagella of many microbial eukaryotes, including stramenopiles and algae (Deflandre, 1934; Liu et al., 2023), may constitute a further, distinct source of EV release. In the biflagellate *C. reinhardtii*, mastigonemes are constitute of a helical structure made of four pairs of anti-parallel mastigoneme-like protein 1 (Mst1) per turn, and of the mastigoneme axial protein (Mstax) as a framework for Mst1 assembly (Wang *et al*., 2023; Huang *et al*., 2024). In biflagellate zoospores of stramenopiles, mastigonemes are tubular and non-tubular and mostly limited to the frontward flagellum, (Bouck *et al*., 1971; Hardham, 1989; Tran *et al*., 2022) while the posterior one, is mainly smooth (Bouck, 1971; Dentler, 1981; Blackman *et al*., 2011; Tran *et al*., 2022). The non-tubular mastigonemes are thin, about 10 nm or less in diameter and of varying length 0.5-1.8 µm. The tubular mastigonemes are tripartite, comprising a basal attachment region, a tubular shaft and generally one or more terminal filaments (Liu *et al*., 2023). They are approximately 50-100 nm in diameter and 0.3-1.5 µm long. In *Phytophthora parasitica (*=*Phytophthora nicotianae)*, the proteins Pnmas1 and Pnmas2 (PPTG_02441 and PPTG_03648) are localised at the tubular shaft of mastigonemes (Blackman *et al*., 2011; Hee *et al*., 2019), while Pnmas3 (PPTG_07238) appears integral to mastigoneme development rather than in mature structure (Hee *et al*., 2019). These proteins are unique to flagellate species within the Stramenopile taxon and have been speculated to participate in mechanosensation and motion regulation (Huang *et al*., 2024). Their sequences harbours three EGF-like domains. Previous proteomic analysis of *P. parasitica* zoospore identified them as plasma membrane-associated proteins, together with receptors, transporters and enzymes (Lupatelli *et al*., 2025).

In the present study we developed a method for the isolation of EVs from zoospores of the phytopathogenic species *Phytophthora parasitica.* We conducted a multi-omics study from appaired samples which enable to dig into EV heterogeneity, biogenesis, and composition (Rai *et al*., 2025; Silva *et al*., 2025). Using electron microscopy we comprehensively imaged EVs and the subcellular location where EV biogenesis occurs (i.e., flagella and cell body). Moreover, we performed comprehensive proteomics and lipidomics profiling, which led to the identification of candidate markers for future investigation into the role of lipid-protein interactions during EV biogenesis. The finding that a subset of EVs are directly generated from the tubular mastigonemes of the front flagellum revealed that mastigonemes are key physical and diffusible structures that may regulate cell-to-cell communication within stramenopiles.

## MATERIAL AND METHODS

### Chemicals and solvents

LC–MS grade solvents and reagents used in the analytical workflow were obtained as follows: methanol (MeOH), 2-propanol, and acetonitrile (ACN) from Biosolve (France). Formic acid (FA), HPLC-grade methyl tert-butyl ether (MTBE), and water (UHPLC grade) were purchased from Carlo Erba Reagents (France). Sodium formate, used for the preparation of the mass spectrometer recalibration solution, and ammonium formate were obtained from Merck (Germany). Sequencing-grade modified trypsin V5111 and the acid-labile detergent RapiGest used in proteomic analyses were purchased from Promega (France) and Waters (France), respectively.

### Preparation of zoospore suspension

Mycelium of *P. parasitica* (isolate 310, Phytophthora INRA collection, Sophia Antipolis) was first cultured for one week in V8 liquid medium (for 1 L: 800 mL distilled water, 200 mL V8 juice, 3 g CaCO_3_) at 24°C under continuous light. After washing and dilaceration, mycelium was incubated for 4 days on water supplemented with 2 % agar to produce sporangia. The zoospores were released in sterile water at ∼10^6^ cells/mL by the following heat shock treatment: incubation at 4°C for 1hr, then incubation in water (4mL per 100mm cell culture dish) at 37°C for 30 minutes.

### Isolation of zoospores-derived EVs

Water containing zoospores was submitted to a three-step procedure of differential ultracentrifugation in order to separate and collect pellet fractions enriched in zoospore cell bodies (CB), flagella (FL) and EVs (EV). At first, flagella were released from cell bodies by vortexing zoospore suspensions (40 mL per sample) for two minutes. Then, the cell bodies fraction (p3K) was pelleted by centrifuging the deflagellated-cell suspension using a Heraeus MegaFuge 1.0 Centrifuge centrifuge (3000g, 10 min, room temperature). The supernatant S1 was subjected to a second centrifugation (3000g) and filtration (10 µm) to remove any cell body contamination. The flagella fractions (p30K) were pelleted from S1 at 31,000g for 30 min at 4 °C (Fu *et al*., 2014; Lupatelli *et al*., 2025) using a Beckman Coulter Avanti J_26 XP centrifuge with a JLA 10.500 rotor. The resulting supernatant S2 were subjected to a filtration (0,45µm) and then to a 100 000g centrifugation in a ThermoFisher WX Ultra 80 centrifuge with a Sorvall TH-641 swinging rotor (1h, 4°C; adjusted K-factor: 306; sedimentation coefficient: s= 5.1 S). The pellet (p100K) enriched in EVs was resuspended in 50 µL of PBS (Phosphate Buffered Saline). Samples were quenched with 200 µL cold MeOH, flash freezed in liquid nitrogen and stored at −20°C till further lipidomic and then proteomic extractions to perform LCMS analyses.

### Lipids extraction and LC-MS analysis

Lipids were extracted using a classical liquid–liquid extraction method based on MTBE/MeOH (10:3, v/v) and 0.1% FA to separate polar and non-polar compounds into distinct phases. Briefly, 615 µL of cold MTBE and 185 µL of cold MeOH were added to each sample, followed by 250 µL of cold water containing 0.1% FA. Samples were vortexed for 45 min at 4°C (1500 rpm) and centrifuged at 14,000 × g for 20 min at 4°C. The upper organic phase, corresponding to the lipid fraction, was carefully collected, evaporated to dryness, and reconstituted in MeOH/H₂O (9:1, v/v) prior to lipidomics analysis. The protein pellet was stored at −80°C before further downstream proteomic analyses.

Lipid separation was performed using an Elute UHPLC system (Bruker Daltonics, Bremen, Germany) equipped with a C18 column from Macherey-Nagel (110 Å, 2.0 × 100 mm, 1.8 µm particle size). The mobile phases consisted of (A) methanol/water (1:1, v/v) and (B) methanol/isopropanol (2:8, v/v), both supplemented with 0.1% FA and 7.5 mM ammonium formate. The column temperature was maintained at 45°C. The gradient program was applied as previously described by Lerner *et al*. (2023) at a flow rate of 0.2 mL/min. A volume of 5 µL of sample was injected in positive ion mode. The autosampler temperature was maintained at 4°C throughout the analysis. Untargeted four-dimensional lipidomics was performed using a trapped ion mobility spectrometry quadrupole time-of-flight mass spectrometer (timsTOF Pro, Bruker Daltonics, Bremen, Germany) equipped with an electrospray ionization (ESI) source operating in positive ion mode. The ESI parameters were set as follows: end plate offset, 500 V; capillary voltage, 4500 V; nebulizer gas (N₂) pressure, 2.2 bar; drying gas (N₂) flow, 10 L/min; and drying temperature, 220°C. For MS/MS acquisition, the instrument was operated in parallel accumulation–serial fragmentation (PASEF) mode with a mass range of m/z 100– 1350 for both MS and MS/MS scans. Ion mobility spectra were recorded over a 1/K₀ range of 0.55–1.90 V·s/cm². The collision energy was set to 30 eV in positive mode. Mass and TIMS calibration were performed in positive mode using sodium formate and the Agilent ESI LC/MS tuning mix, respectively. Data calibration was carried out using a mixture of Agilent ESI LC/MS tuning mix and 0.5 mM sodium formate (1:1, v/v), which was directly infused into the ESI source via a syringe pump, as previously described by Lerner *et al*. (2023).

### Data Processing and lipid Annotation

Data processing was performed using Metaboscape 2025b and DataAnalysis 6.1 (Bruker Daltonics, Bremen, Germany). Metaboscape was used for feature extraction, peak area determination, lipid annotation, and data curation. Feature detection was carried out using an intensity threshold of 500 counts. The filtering rules applied to the 4D extracted features were defined as follows: (1) recursive feature extraction was performed for features detected in at least 2 out of 3 analyses; and (2) features were included in the final bucket table only if present in at least 75% of analyses after recursive feature extraction. Background subtraction was subsequently performed using MeOH/H₂O blanks and elution solvent samples. For enhanced lipid annotation, two complementary tools were employed: the internal Lipid Annotation Tool within MetaboScape 4.0 software (Bruker Daltonics) and the LipidBlast spectral library (Kind *et al*., 2013). The following parameters were applied: mass tolerance of 5.0 ppm, isotope fitting threshold of 250 mSigma, MS/MS spectral matching score up to 400/800, and collision cross section (CCS) tolerance of 3.0%. High-confidence lipid species identification was based on four-dimensional descriptors: accurate mass, retention time (RT), CCS, and MS/MS spectra. The LION web-based ontology enrichment tool was used for identification of enriched lipid-associated terms (Molenaar *et al*., 2019).

### Quality Control and Statistical Analysis

To ensure analytical robustness and reproducibility, pooled quality control (QC) samples (mixture of all analysed samples) were included throughout the analytical sequence. QC samples were injected at the beginning of the run, after every three samples, and at the end of the run. Coefficients of variation were calculated for each lipid species using the formula: CV (%) = (standard deviation / mean) × 100. Lipids exhibiting a CV > 30% in QC samples were excluded from the final dataset.

Statistical analyses were performed using the MetaboAnalyst platform. Univariate analysis was conducted using one-way ANOVA followed by post hoc testing with false discovery rate (FDR) correction, considering an adjusted p-value < 0.05 as statistically significant. Multivariate analysis was performed using principal component analysis (PCA) to assess analytical system performance and detect potential outliers.

### Proteins extraction and LC-MS analysis

Each protein pellet was centrifuged at 17 000g (10min, 4°C), dried after removing the residual supernatant from the lipid extraction step for 25 min and solubilized in 8M urea, 10mM DTT, 0.1% RapiGest (Waters) and 50mM N_4_HCO_3_ pH 7.8 for 1h at 37 °C. Proteins (1µg) were submitted to a brief sonication and then alkylated with 15 mM iodoacetamide in 50mM N_4_HCO_3_ for 30 min at 25°C in the dark. The proteinaceous content was digested overnight at 37 °C in a solution containing 5 mM CaCl_2_, and 12.5 ng/μL sequencing-grade modified trypsin in 25 mM N_4_HCO_3_, pH 7.8. Trypsin and RapiGest were deactivated respectively by reducing the pH to 2 with FA and by incubation of the samples at 37°C for 45 min. The samples were desalted using OMIX C18 pipette tips (100 μL, A57003100, Agilent, Santa Clara, CA, USA), dried and resolubilized in 0.1% FA (20µL) before LCMS analysis.

Nano-High Pressure Liquid Chromatography coupled with High Resolution Mass Spectrometry (Nano-HPLC-HRMS) analysis was performed using a Nano RSLC system (Ultimate 3000, Thermo Fisher Scientific, France) coupled to an Exploris 480 (Thermo Fisher Scientific, France). Each sample (5 μL) was injected and first concentrated on a μ-Precolumn Cartridge AcclaimPepMap 100 C18 (i.d. 5mm, 5 mm, 100 A°; Thermo Fisher Scientific) at a flow rate of 10 µL/min and using solvent containing H2O/ACN/TFA 98%/2%/0.1%. Peptide separation was performed on a 75 µm i.d. × 500 mm (2μm, 100 A°) PepMap RSLC C18 column (Thermo Fisher Scientific). The mobile phase A was 0.1% FA in water, and mobile phase B was 0.1% FA in ACN. The elution gradient started at 2% of B for 3 min and followed the different step t = 103 min, 20% B; t = 123 min, 32% B; t = 125 min 90% B; t = 130 min 90% B. The oven temperature was set at 40°C and the flow rate at 300 nl/min. MS spectra were acquired at a resolution of 120 000 (200 m/z) in a mass range of 375–1500 m/z with an AGC target 3e6 value of and a maximum injection time of 25 ms. The 20 most intense precursor ions were selected and isolated with a window of 2 m/z and fragmented by HCD (Higher energy C-Trap Dissociation) with a normalized collision energy (NCE) of 30%. MS/MS spectra were acquired in the ion trap with an AGC target 5e5 value, the resolution was set at 15,000 at 200 m/z combined with an injection time of 22 ms.

### Data Processing and protein Annotation

The proteomic data were analysed against the *P. parasitica* current proteome containing 26,438 entries (*P. nicotianae*, strain INRA-310, UniProtKB: UP000018817) and with PEAKS Studio (version Xpro, Bioinformatics Solutions) (Ma *et al*., 2003). Digestion mode was set to Trypsin/P specificity, allowing maximum 4 misscleavage with fixed post-translational modifications (PTMs): carbamidomethyl modification of cysteine, and variable modifications of protein N-terminal acetylation and carbamylation, protein C-terminal amidation, methionine oxidation, asparagine and glutamine deamidation, aspartic acid, serine, threonine, tyrosine dehydration. The PTM score was set superior or equal to 20. Parent mass error tolerance was set at 10 ppm and fragment mass tolerance at 0.02 Da. Proteins (n= 4,010) were initially identified from 34911 peptides based on a 1% false discovery rate for both the proteins and peptides, and identification of at least 1 unique peptide in each replicate (Supplementary table1); 1472 proteins were selected for further annotation based on identification of at least 2 unique peptides in each replicate (Supplementary table2).

Protein annotation (names, families, Gene Ontology IDs) was extracted from the UniProt database (The UniProt Consortium, 2023). Protein topology and localization were predicted using Protter (Omasits *et al*., 2014) and DeepTMHMM - 1.0 (Hallgren *et al*., 2022; https://services.healthtech.dtu.dk/services/DeepTMHMM-1.0/) for transmembrane helices in proteins; the NetGPI GPI for GPI-anchors (Gíslason *et al*., 2021); SignalP 5.0 for signal peptides (Almagro Armenteros *et al*., 2019) and DeepLoc 2.1 for subcellular localisation (Ødum *et al*., 2024). Functional enrichment analysis was performed with g:Profiler (Kolberg *et al*., 2023) using g:GOSt, *P. nicotianae* as organism, g:SCS and P<0.001 as significant thresholds.

The proteins were schematically grouped based on their topology and localization as cytosolic proteins or membrane-associated proteins within which we gathered transmembrane and secreted proteins. To avoid misinterpretation due to co-purification of EVs with secreted and soluble proteins other than those functionally associated with EVs, we only considered secreted proteins already identified as associated with the plasma membrane or extracellular anchored-structures in *Phytophthora* species.

### In silico identification of *P. parasitica* zoospore proteins as EV marker candidates

A list of proteins previously characterized in various species as EV markers was established from the literature. The list included transmembrane, GPI-anchored or cytosolic proteins (Welsh *et al*., 2024); proteins exhibiting MARVEL domains identified as markers of *Phytophthora infestans* EVs (Breen *et al*.,2025), and primary cilium proteins identified in other eukaryotes as ciliary EV markers (Vinay and Belleannée, 2022). For each protein sequence, a BlastP search was performed against the 26,438 entries of the *P. parasitica* (strain INRA-310) reference proteome (https://www.uniprot.org/proteomes/UP000018817). *P. parasitica* orthologs were identified and annotated using tools made available by the Consortium UniProt [Universal Protein Resource; [European Bioinformatics Institute (EMBL-EBI)](https://www.ebi.ac.uk/), the [SIB Swiss Institute of Bioinformatics](https://www.sib.swiss/) and the [Protein Information Resource (PIR)](https://proteininformationresource.org/)]. Protein annotation was complemented with transcriptomic and proteomic data from *P. parasitica* zoospore swimming freely (Bassani *et al*., 2020b; Lupatelli *et al*., 2025): mRNA expression measurement (FPKM>2; Fragments Per Kilobase Million); and protein expression measurement (Sum of peak intensities>1000), plus protein relative abundance in flagella versus cell body (0,5<FOLD change>2; FDR <0,01), respectively.

### Transmission electron microscopy and scanning electron microscopy

For analyses of enriched-EVs fractions or zoospore-conditioned water, the samples were prepared using the negative staining method for transmission electron microscopy (TEM). A drop (10 µL) of each sample type was left for 30 min on a TEM copper grid (400 mesh) with a carbon-coated film. The excess liquid was removed with a filter paper. Subsequently, staining was done by adding a drop of 0.5% (w/v) aqueous solution of uranyl acetate on the grid for 1.5 min, followed by removal of excess solution. TEM were also used to analyze thin sections of zoospores mainly as described by Bassani *et al*. (2020b). Zoospores collected by centrifugation were fixed with 1.6 % glutaraldehyde diluted in 0.1 M phosphate buffer (pH 7.4), rinsed in 0.1 M sodium cacodylate buffer (pH 7.4) and post-fixed with 1% osmium tetroxide diluted in 0.1 M phosphate buffer (pH 7.4) reduced with potassium ferrycyanide (1%), for 1 h). After a water wash, cells were dehydrated with several incubations in increasing concentrations of acetone and embedded in epoxy resin (EPON). Ultrathin sections (80 nm) were generated and placed onto (150-mesh) copper grids, stained with uranyl acetate and lead citrate. Scanning Electron Microscopy (SEM) analyses were performed on samples of fixed cells treated as described by Tran *et al*. (2022). Samples were mounted on SEM stubs with silver paint and coated with platinum (3 nm) prior to observing. Electron microscopy observations were carried out at the CCMA center (Centre Commun de Microscopie Appliquée, Université Côte d’Azur) with JEOL JEM-1400 TEM, operating at 100 kV and equipped with an Olympus SIS MORADA camera, and a JEOL JSM-6700F SEM at 3 kV.

For the immunogold detection of Pnmas1 and PnPar in EVs, the samples were initially fixed with 4% PFA and adsorbed onto nickel grids. After blocking with 5% bovine serum albumin, the grids were incubated with a primary antibody (diluted 1:200) against against Pnmas1 (PPTG_02441; Lupatelli *et al*., 2025) or against elicitins (cryptogein, parasiticein-PnPar; Devergne *et al.,* 1992). Following washes, the samples were labeled with a 10 nm colloidal gold-conjugated secondary antibody (diluted 1:200) at room temperature for 1 h. The samples were then post-fixed with 2.5% glutaraldehyde and negatively stained with uranyl acetate.

### Immunolocalization

Zoospores were fixed by mixing cell suspension with 3% glutaraldehyde (1:1) and incubated for 30 min on ice. After centrifugation (5000g for 4 min at 4 °C) and washing with Phosphate-Buffered Saline (PBS pH 7,2), zoospores were spread onto glass slides in 0,1xPBS. Adhesion of cells was achieved by applying a moderate evaporation at 37°C for 10 min. For intracellular detection, cells were permeabilized by adding PBS-Triton (0.1%) for 15 min at room temperature prior incubations with antibodies. The incubations with antibodies were performed as followed. Between 3 washing with PBS, samples were successively incubated: (i) with a blocking solution containing 5% of nonfat dry milk in PBS for 30 min at room temperature (RT); (ii) then with a primary rabbit antibody diluted 1:100 in PBS for 18 hours at 4°C, and (iii) finally with a Fluoprobes 547-conjugated antibody (Interchim, Inc., Montluçon, France), diluted 1:100 in PBS for 30 min at RT. Samples were then mounted in Fluoroshield (Sigma-Aldrich, St. Louis, MO, USA). Image acquisition was performed using a Zeiss LSM 880 confocal laser scanning microscope (Carl Zeiss Microscopy, Germany) on the Microscopy Platform-ISA-INRAE 1355-UNS-CNRS 7254-INRAE PACA-Sophia Antipolis.

### Immunoblot

Samples (1 μg) were separated in 4–20% precast polyacrylamide gels (Bio-Rad), transferred onto a nitrocellulose membrane and stained with Ponceau red. The membranes were incubated and washed sequentially in PBS in the presence of 5% (w/v) nonfat dry milk, a primary rabbit polyclonal antibody (1:2000), a goat peroxidase-conjugated IgG directed against rabbit immunoglobulins (1:2000), and a chemiluminescent substrate for peroxidase. The primary rabbit antibodies used (1:1,000) were raised against Pnmas1, Na^+^/K^+^-ATPase (PPTG_09633, Lupatelli *et al.,* 2025), elicitins, α-tubulin (NB-22-6794, NeoBiotech), Phospholipid-transporting ATPase IA (NB-22-54564, NeoBiotech), and HSP70 (NB-22-6787-200, NeoBiotech). The bound antibodies were detected using the FusionX apparatus (Vilber, Marnela-Vallée, France) and the ECL Western Blotting Substrate protein labelling kit (Promega).

## RESULTS

### Cell body and flagellar membrane protrusions in P. parasitica zoospore

Topological changes of zoospore plasma membrane, observed by TEM and SEM, are evocative of frequent events of vesicle discharge and extend from the front flagellum to the cell body. Figure 1A, B and Supplementary Figure 1 illustrate representative morphotypes of zoospores exhibiting a number of large intracellular vesicles (∼500nm) localised just underneath the curved plasma membrane (Figure 1C). We also captured fusion events of plasma membrane with intracellular vesicles (Figure 1D) or with a multivesicular body filled with a small intraluminal vesicle (Figure 1E, initially detected by Bassani *et al*., 2020b), as well as outward budding of the flagellum membrane (Figure 1F), showing respectively putative exocytosis, exosome and ectosome release by zoospores during zoospore swimming.

**FIGURE 1.**
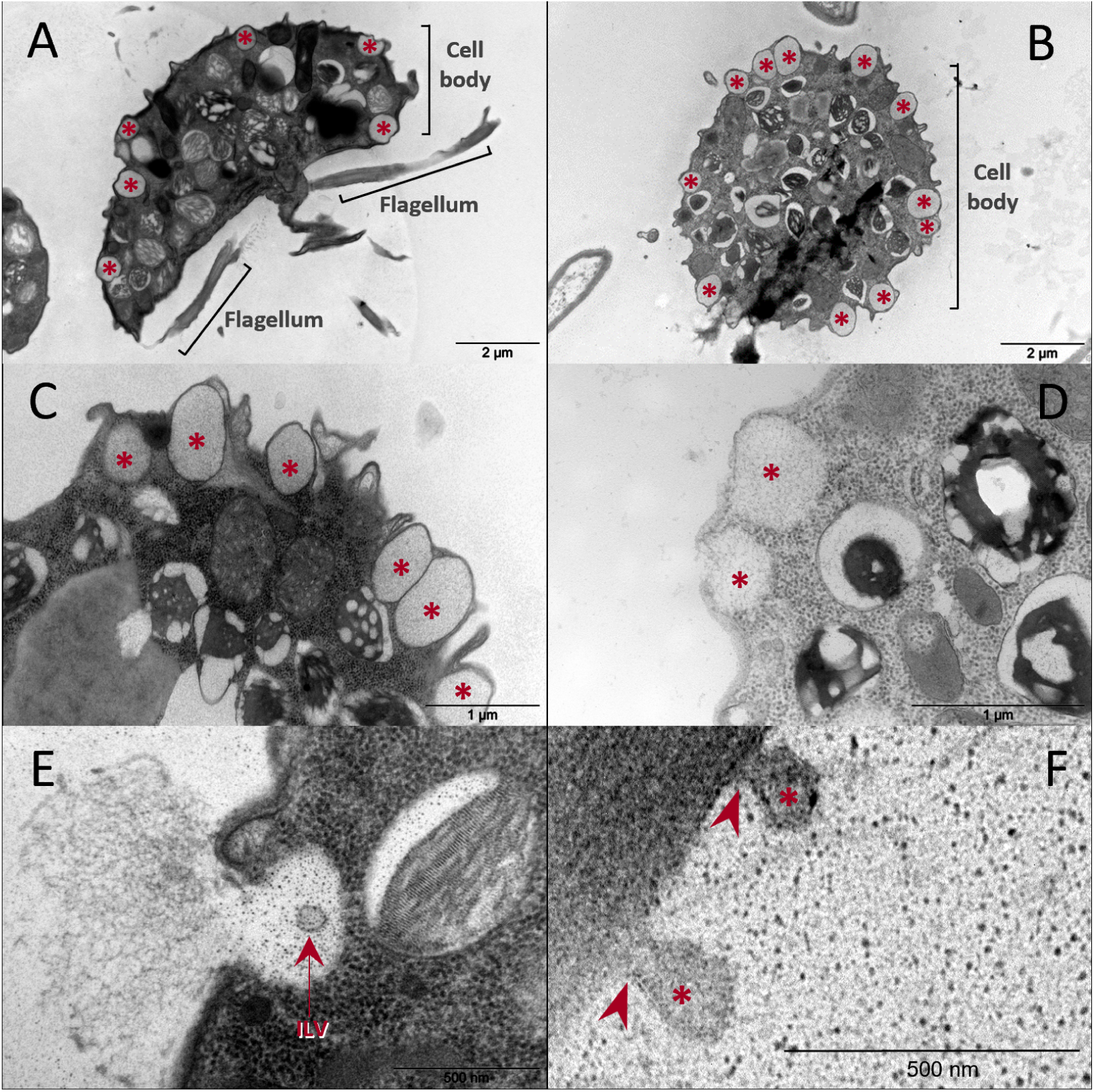
Ultrastructure of zoospore surface. Transmission electron micrographs of thin-sections through *P. parasitica* zoospores. (**A**) Longitudinal section including the reniform cell body and its ventral groove, and the two flagella. (**B**) Transversal section in a plan revealing multiple protrusions of the plasma membrane at the cell body level (**C**) Intracellular vesicles into zoospores just below the plasma membrane. (**D**) Intracellular vesicles merging with the plasma membrane on exocytosis for release content. (**E**) Intralumenal Vesicle (ILV) into a Multivesicular Body (MVB) merged with plasma membrane evocative of exosome release. (**F**) Outward budding from flagellum membrane. Red arrowheads point pinch zones during budding of the mastigoneme membrane. Red stars show vesicles resulting from budding of the plasma membrane (**A**, **B**, **C**) or its flagellar extension (**F**), and fusion of vesicles with the plasma membrane (**D, E**).

A detailed visualisation of the front flagellum showed that the tubular mastigonemes may also contribute to EV biogenesis. TEM and SEM revealed a composition of tubular and vesicular subunits of mastigonemes (Figure 2A-C). The mastigoneme structure elongated by a tubular proximal region (tpr) and then by a distal one composed of vesicles organised in a row (vr). Each vesicle is separated from the others by a short, dense pinch zone (pz) (53.6 ± 12,7 nm, n=14) reminiscent to budding events prerequisite for ectosome release. The diameter of vesicles attached to the mastigonemes was distributed within a range between 40,8 and 182,5nm (mean=110,9±29,9, n=39) (Figure 2D). Ultrathin sections also revealed the membrane nature of the envelope of tubular mastigonemes. When observed in a longitudinal section with aligned vesicular and tubular subunits (Figure 2E), the surrounding membrane shows the typical plasma membrane ultrastructure in TEM: two electron-dense lines separated by a narrow space. Taken together the microscopic observations indicate that zoospore EV release occur from structures having vesicular appearance located at both the cell body and at tubular mastigoneme tips level. Front flagellum tips were also organised in a row of spheroid structures indicating that it could be an additional source of zoospore-derived EVs (Supplementary figure 2).

**FIGURE 2.**
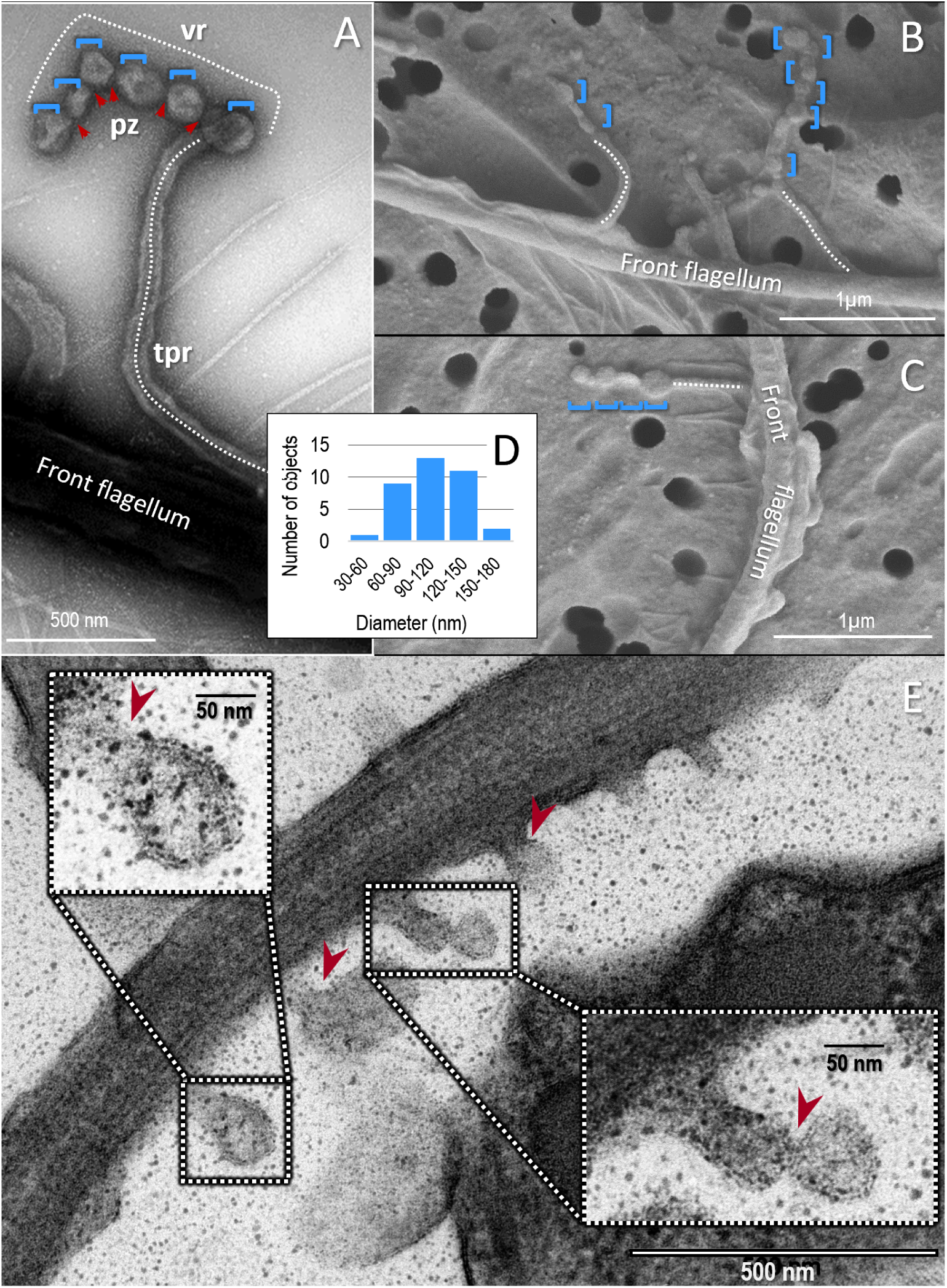
Ultrastructure of tubular mastigonemes. Negative staining TEM (**A**) and SEM (**B, C**) micrographs of front flagellum showing tubular mastigonemes structural organisation: the tubular proximal region (tpr, dotted white line), the distal and terminal one mainly consisting of vesicles rows (vr, blue bracket), a pinch zone (pz; red arrowhead) separating each vesicular form. (**D**) Histogram of diameter distribution of zoospore released-EVs for vesicles attached to the tubular mastigonemes (n=39; increase range 30nm). (**E**) Electron micrograph of an anterior flagellum section including tubular mastigonemes anchored at the flagellar membrane. Inlets show the mastigoneme membrane characterized by two parallel and electron-dense lines with a thin gap between them, which correspond respectively to polar heads of phospholipid layers and to their aliphatic chains. Red arrowheads point pinch zones during budding of the mastigoneme membrane.

### Zoospores swimming in water shed extracellular vesicles

To confirm that zoospores could generate EVs, a further TEM analysis was carried out on extracellular material deposited on a carbon-coated copper grid after zoospores 30-min swimming in water. The deposits mainly consist of tubules and round particles, which had archetypal cupshaped morphology and EVs size range (Figure 3A, B). The diameter of EVs was estimated to range between 40,8 and 182,5 nm (mean =84,6±35,8 nm) (Figure 3C). The concentration of deposited EVs was estimated to 3000-6000 per mL in a 10^6^ zoospore/mL suspension. The release in water of EVs by zoospores allowed us to purify and concentrate EVs derived from zoospores swimming in water using differential ultracentrifugation. Three steps were applied after zoospores deflagellation: a first one at 3,000 x g to pellet cell bodies, a second one at 30,000x g to pellet flagella (Lupatelli *et al*., 2025), and a last one at 100,000g to pellet EVs. EV particles in the 100K fractions observed by TEM microscopy were typical membrane-bound spherical vesicles (Figure 3D, E) with a mean diameter of 91.8 nm whose distribution extended over a range of 30 to 250 nm (Figure 3F). The three fractions recovered by ultracentrifugation were used to characterize the proteinaceous and lipid contents of EVs, as described in the following sections.

**FIGURE 3.**
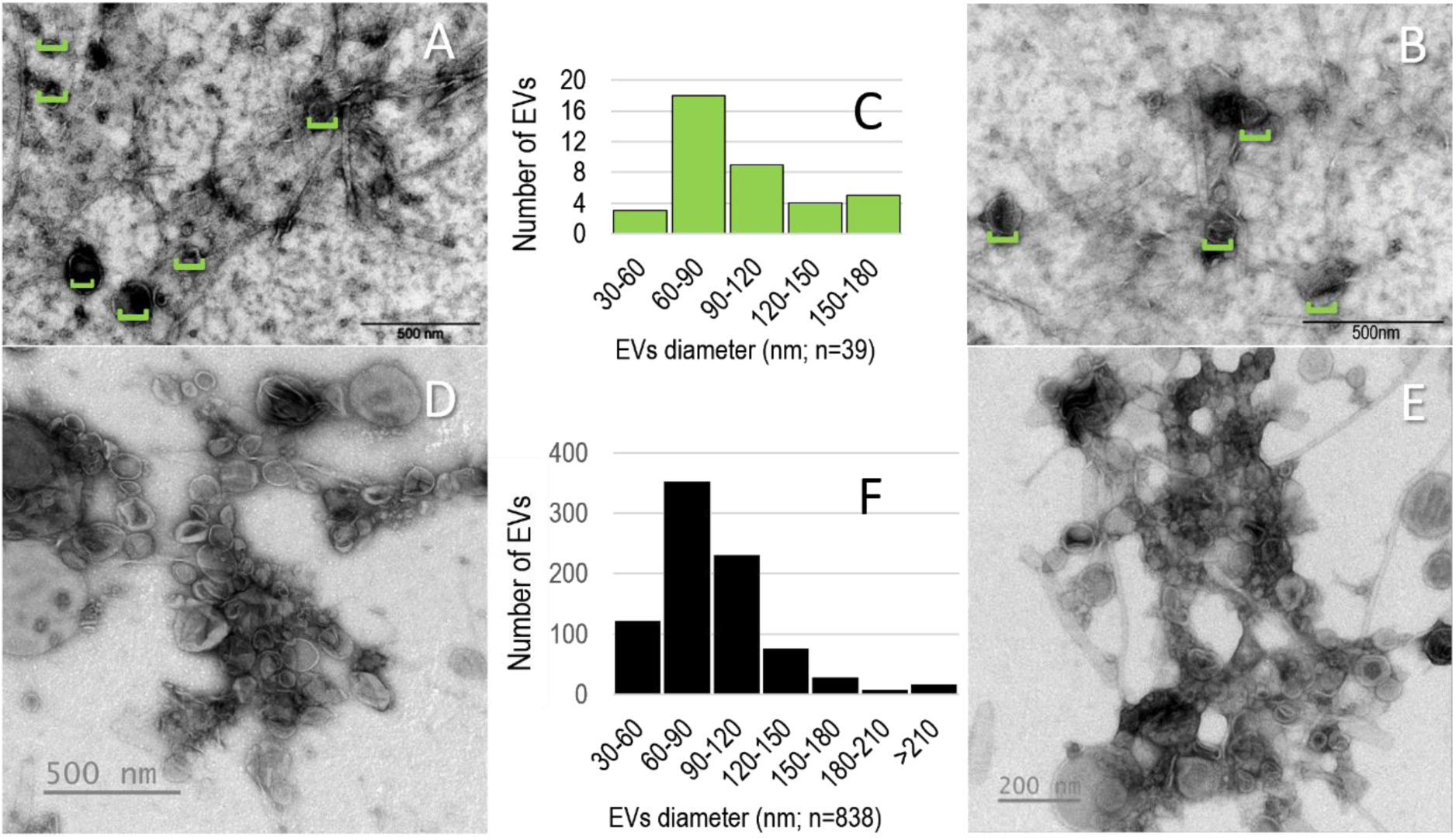
Ultrastructure of EVs shed by zoospores. (**A, B**) TEM negative staining of zoospore extracellular material. Green horizontal brackets indicate positions of extracellular vesicles coated on copper grid that has been explored by zoospores for 30 min. (**C**) Histogram of diameter distribution of zoospore released-EVs (n=69). range of 30nm. (**D, E**) Negative staining of EVs collected in 100k pellet fractions. (**F**) Histogram of diameter distribution (n=838); range of 30nm.

### Proteomics profiling of zoospores extracellular vesicles

To provide a first insight into zoospore EV composition, EVs-enriched samples were at first examined for their proteinaceous content. Three 100K-EV samples were analysed after trypsin digestion, liquid chromatography separation, peptide identification by mass spectrometry and mapping against the *P. parasitica* proteome. Among the 4,010 proteins initially identified (Supplementary table 1), 1 470 met the criterion of ≥ 2 peptides per replicate, were annotated and ranked according to a high to low score (-10log*P*) (Supplementary table 2). We compared the 1470 *P. parasitica* zoospore-EVs proteins to the 1326 high confidence proteins identified investigating the *P. infestans* mycelium-EVs in the same genus (Breen *et al*., 2025). Forty three percent of zoospore-EVs proteins appeared as orthologous proteins (BlastP e-Value<e^−100^) of those included in the *P. infestans* mycelium-EVs list. The percentage reaching 60% within the TOP400 was indicative of good coverage of first-identified proteins from *P. parasitica* zoospores with those characterized from EVs released by *P. infestans* mycelium (Supplementary table 3). Within the list of *P. parasitica* zoospore-EVs proteins we also identified those having similarity (BlastP e-Value<e^−6^) with known EV markers reported in public databases or being members of protein families including EV markers (Chitti *et al*., 2024; Welsh *et al*., 2024; Vinay and Belleannée, 2022; Supplementary table 4). The distribution of identified proteins within the list revealed their positions in the top positions: 20 proteins within the TOP100, 26 proteins within the TOP 400 and 33 within the whole list (Supplementary table 2). Thus, this proteome classification revealed a remarkable richness in putative EV markers among the proteins having the highest score. To proceed to further analyses, we also considered two points. (i) The proteome of EVs derived from a single cell type, as in this case, is necessarily much less diverse than the core proteome (1200 proteins) of a heterogeneous set of EVs generated by many cell types and collected from a bodily fluid such as plasma (Kugeratski *et al*. 2021; Kalluri *et al*., 2024). (ii) Zoospore-derived EVs (∼0.5 10^−3^ µm^3^) would contain far fewer proteins than a whole eukaryotic cell encompassing about 10,000 different proteins (Beck *et al*., 2011), an order of magnitude roughly aligned with the number of distinct transcripts characterized in a zoospore with an approximated volume of 500 µm^3^ (Bassani *et al*., 2020b). Based on these results and quantitative considerations we focused the study on the TOP400 proteins in order to initiate the identification of main candidate EV markers.

### Functional characterization of the identified proteins

GO terms analysis of TOP400 proteins of the 100k EV fractions allowed to identify membrane as the most represented cell compartment and showed enrichment in molecular functions, such as small molecule and nucleic acid binding, catalytic activity, structural molecule activity, phospholipid binding, cytoskeletal motor activity, protein dimerization activity (Figure 4A, Supplementary table 5). Among the 33 proteins without gene ontology annotation, 9 proteins were classified within the TOP20 and 3 are specific of the oomycete lineage.

**FIGURE 4.**
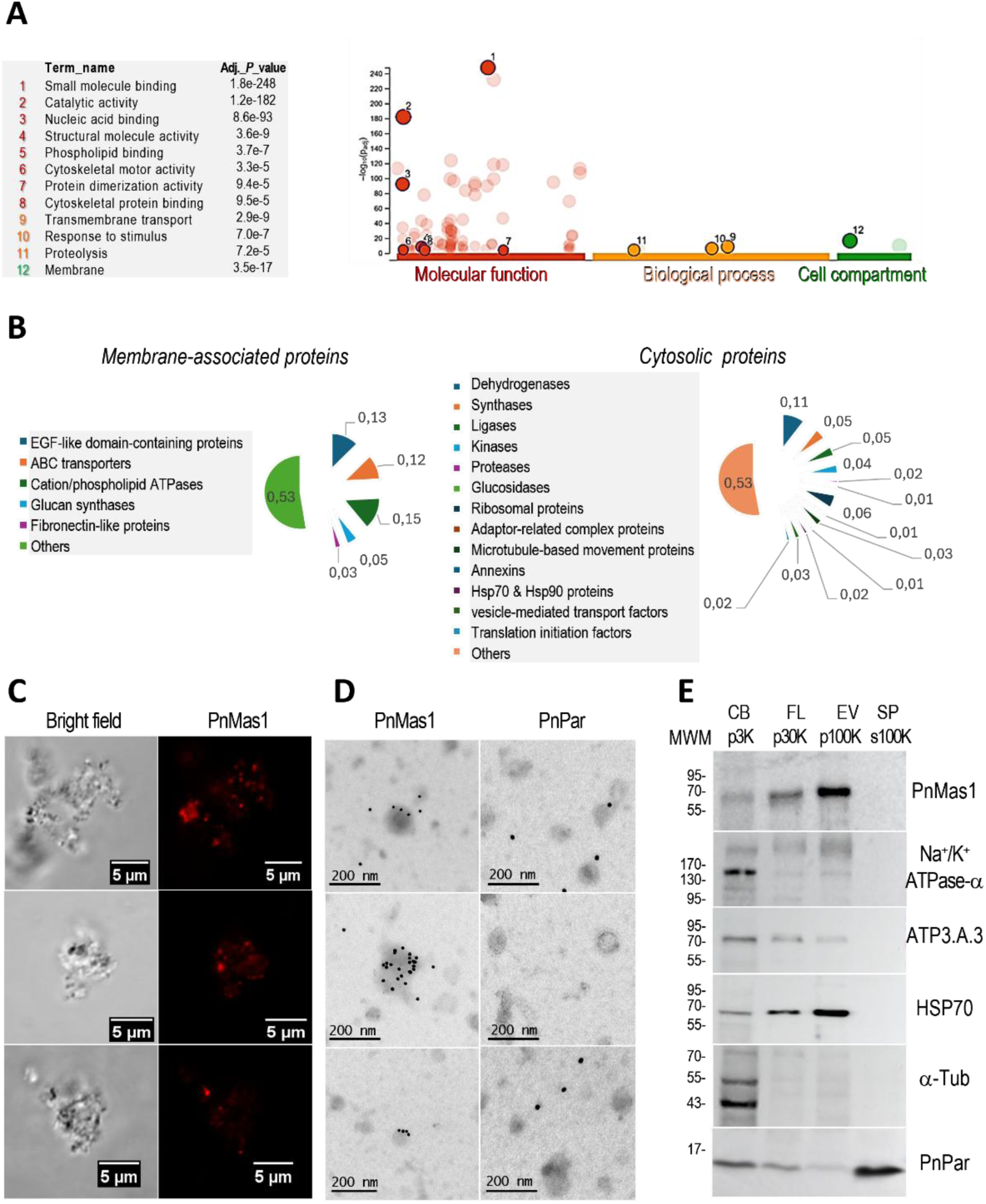
Zoospore EV Proteome. (**A**) Functional enrichment analysis of the TOP 400 proteins identified using g:Profiler. The functional terms organized according to the g:Profiler color code are distributed along the *x*-axis; the enrichment adjusted *p*-values expressed as −log *p*-values are represented on the y-axis. The twelve numbered points and the table indicate the driver terms identified in GO. (**B**) Scatter plots showing the relative abundance of EVs-related proteins families identified as membrane-associated (left part of the panel) or cytoplasmic (right part of the panel) proteins. (**C**) Immunofluorescence and (**D**) immunogold-labeled micrographs of bioparticles collected in EV-p100k pellet fraction stained with an anti-PnMas1 antibody (PnMas1), showing a positive detection for a fraction of EVs. Immunogold-labelling with an antibody against the secreted elicitin parasiticein (PnPar; Devergne *et al.,* 1992) does not decorated EVs. (**E**) Western blot analysis of the CB (p3K), FL (p30K), EVs (p100K) and SP (100K supernatant; s100K) fractions (1µg) using antibodies directed against alpha-tubulin, PnMas1, alpha-Na^+^/K^+^ ATPase, Phospholipid-transporting ATPase A.3, the elicitin PnPar. Molecular weight markers (MWM) are indicated in kilodaltons (kDa).

We further examined the proteome for the features of membrane-associated or cytosolic proteins (Figure 4B), and the pathogenicity factors of *Phytophthora* (Supplementary table 6). The membrane-associated candidates (n=98) clustered into distinct functional groups with ABC transporters, 1,3-beta-glucan synthase (EC 2.4.1.34), EGF-like domain-containing proteins, and cation-transporting ATPases, including phospholipid-transporting ATPases type IV, constituting the major proportions (n=39). The presence of several cation- and phospholipid-transporting ATPases supports their role in outward budding, promoting phospholipid translocation between membrane leaflets and enhancing extracellular vesicle formation (Cocucci and Meldolesi, 2015; Meldolesi, 2018). Additionally, 7 proteins exhibit significant similarities with proteins known as EV markers for others species: PPTG_13069 is the *P. parasitica* ortholog of PiMDP1, one of the MARVELous protein markers for *P. infestans* EVs released from mycelium (Breen *et al*., 2025); PPTG_16242 is the ortholog of *P. cactorum* Prominin, which exhibits similarities with mammals prominins characterized as ciliary EV markers (Vinay and Belleannée, 2022). Four beta-1,3-glucan synthases were also found orthologous to synthases previously reported in yeast (Zhao *et al*., 2019) suggesting that zoospore EVs could contribute to the recruitment of the enzymatic machinery elaborating the primary cell wall just after flagella loss (Wang and Bartnicki-Garcia, 1982). The EV proteome is also composed of cellulases that can help to reduce the thickness of the cell wall of recipient plant cells facilitating EV crossing (Gill *et al*., 2019).

Interestingly, comparison of the EV membrane-associated proteome with that of the *P. parasitica* flagellar membrane (Lupatelli *et al*., 2025) showed substantial overlap, with 11 of 98 membrane-associated EV proteins found among the 75 most abundant proteins in flagella (Supplementary Table 7). This is consistent with a subpopulation of EVs being derived from flagella. Four of these proteins have previously been immunolocalized either on both flagella (the sodium-potassium ATPase PPTG_09633) or specifically on the mastigoneme of the anterior one (the EGF-like domain-containing proteins PPTG_02441, PPTG_07238, PPTG_03648; Hee *et al.,* 2019; Lupatelli *et al*., 2025). Emphasizing the contribution of the flagellum membrane to the EV release, immunodetection of the mastigoneme protein PnMas1 (PPTG_02441; Supplementary Figure 3) established that the EVs enriched fraction included a subset of PnMas1^+^ particles (Figure 4C). Using confocal microscopy, we were able to measure the rate of PnMas1^+^ particles to 0,19 + 0,12 (n=1631), with a size range for particles (diameter) smaller than the one determined for fluorescent microspheres of 500nm (Supplementary Figure 4). Using TEM, the staining of this fraction was characterised by a punctuated immunostaining delimitating vesicles of 100-200 nm (Figure 4D). Immunoblot analysis confirmed that PnMas1 was abundant in the 100K fraction containing zoospores-derived EVs (Figure 4E).

The set of cytosolic candidates (n=300) was highly heterogeneous with certain families of proteins known to be present in greater quantities in EVs (Welsh *et al*., 2024; Vinay and Belleannée, 2022). These include included several proteins clustered in functional groups for microtubule-based movement (n=10) or vesicle-mediated transport (n=11), and also in 2 groups mostly detected within TOP100 including proteins related to EV markers: the proteins of the HSP70 and HSP90 heat shock systems (n=6), and annexins involved in membrane trafficking and that bind to phospholipids (n=4) and also characterized as ciliary EV markers. The set also includes distinct metabolic enzymes, such as dehydrogenases (n=28), synthases (n=11), ligases (n=18), and also ribosomal proteins (n=18).

We also approached the proteomic analysis from the perspective of the pathogenic nature of *Phytophthora* species, which is mediated by an arsenal of effectors (Wang and Jiao, 2019; Boevink *et al*., 2020). Here, prevailing *Phytophthora* effectors such as RxLR effectors targeting plant host functions as negative regulators of immunity (Wang *et al*., 2023), were not identified among the set of proteins initially identified and were therefore not considered as proteins constitutive of zoospore-derived EVs. Few members of the elicitin family triggering the hypersensitive response (HR) and plant defence responses (Derevnina *et al*., 2016) were weakly detected among the 4,010 proteins (Supplementary table1). These results indicated that the EV-zoospore proteome is not enriched in known pathogenesis-related factors.

We then performed SDS-PAGE and immunoblot analyses to compare the content of cell bodies (CB), flagella (FL), EVs (EV) and the 100K supernatant (SP) fractions, the last one being enriched in secreted proteins. In congruence with the identification by LC-MS/MS, bands with the apparent molecular weight of PnMas1, Na^+^/K^+^ ATPase, HSP70, and at least one phospholipid-transporting ATPase (Figure 4E) were detected in EVs, or enriched (as for PnMAs1) in EVs compared with the other fractions, confirming that they are part of the EV composition. Parasiticein (PnPar), a small 10-kDa elicitin highly secreted by *P. parasitica* (Nespoulous *et al*., 1992) and not identified in EVs by proteomics, was characterized as a negative control protein for the immunoblot experiments. As expected, it was the only tested protein absent in the purified EVs preparation p100K while vastly abundant in the secreted protein preparation SP (Figure 4E). Using immunogold, PnPar could no longer be localized in EVs neither as a surface-bound protein nor an internal cytosolic cargo (Figure 4D).

### Comparative lipidomics profiling in cell bodies, flagella, and extracellular vesicles

We next investigated lipid profiling associated with zoospore EVs. We performed a comparative untargeted 4D lipidomic analysis of cell bodies (CB), flagella (FL), and EVs (EV). After quality filtering (CV in QC samples < 30%), 191 unique lipids were obtained for downstream analysis. These lipids belonged to three major classes: sphingolipids (SP), glycerolipids (GL) and glycerophospholipids (GP). Annotated lipids are listed in Supplementary table 8.

At the class level, CB were markedly enriched in glycerophospholipids (GP), accounting for more than 90% of CB lipid profile, with phosphatidylcholine (PC) alone representing approximately 81% (Figure 5A). In contrast, FL and EV exhibited comparable proportions of GP i.e., approximately 54% and 51%, respectively (Figure 5A). Across all sample types, GP represented the class with the highest number of unique annotated lipid species, underscoring their structural diversity.

**FIGURE 5.**
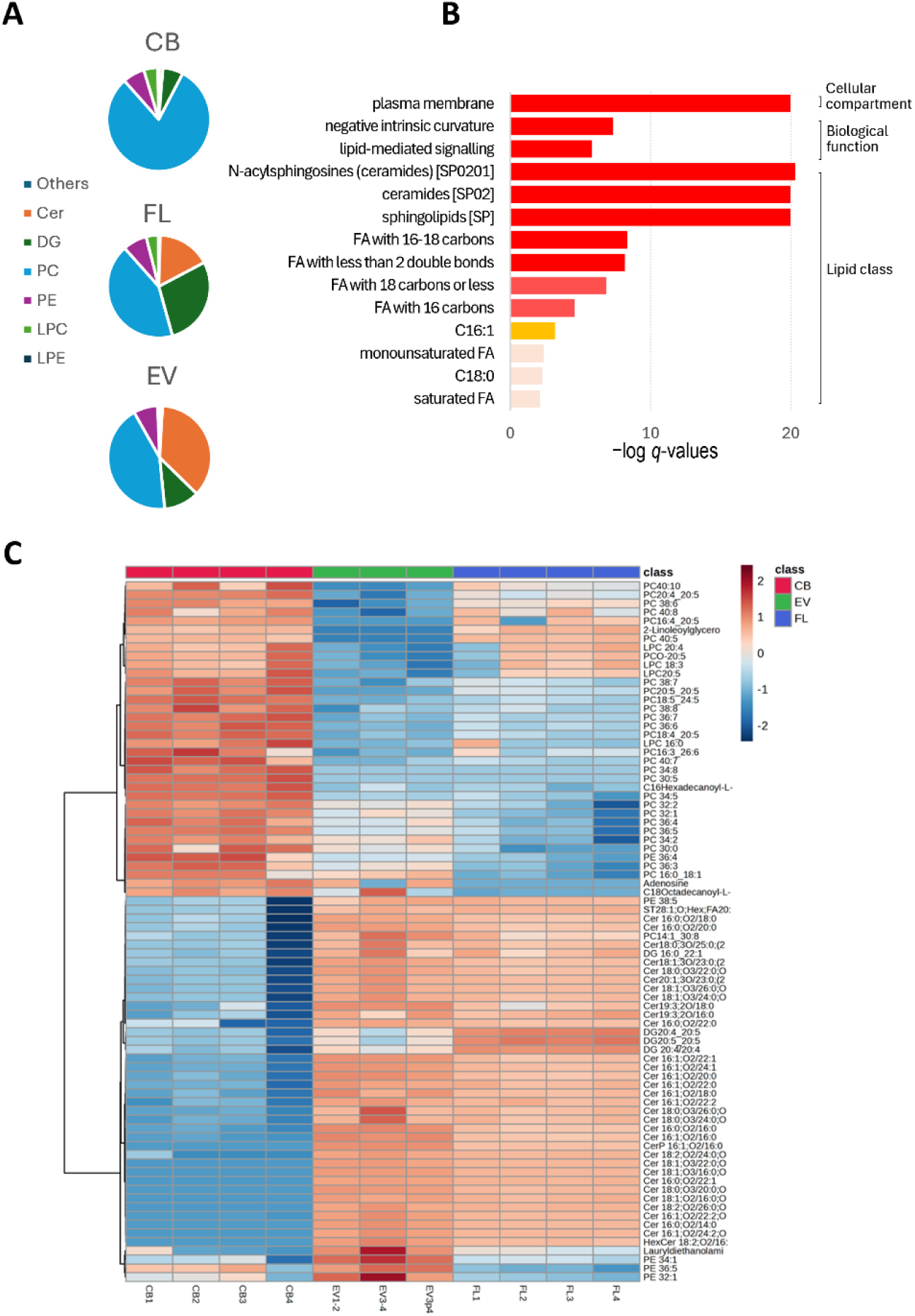
Zoospore EV Lipidome. (**A**) Relative lipid class composition of cell bodies (CB), flagella (FL), and extracellular vesicles (EV). Lipid subclasses include, among the others, ceramides (Cer), diacylglycerols (DG), phosphatidylcholine (PC), phosphatidylethanolamine (PE), lysophosphatidylcholine (LPC), lysophosphatidylethanolamine (LPE). Data are presented as percentages of the total annotated lipids intensities for each sample type. **(B)** LION-term enrichment analysis of the subset EV-related lipids compared to the 191 unique lipids identified using the “target-list mode”. The significant LION-terms are shown (*q* < 0.05). Bar colors are scaled with the enrichment (−log *q*-values). FDR: false-discovery rate. **(C)** Heatmap of differentially expressed lipid species across cell bodies (CB), flagella (FL), and extracellular vesicles (EV). Lipids were selected based on one-way ANOVA with post-hoc tests and a false discovery rate–adjusted p-value<0.05. Abbreviations for lipids: ceramides (Cer), diacylglycerols (DG), phosphatidylcholine (PC), phosphatidylethanolamine (PE), lysophosphatidylcholine (LPC), lysophosphatidylethanolamine (LPE), ether phosphatidylcholine (PCO).

Diacylglycerols (DG), key intermediates in lipid metabolism and membrane curvature regulators, were most abundant in FL (28%) and less represented in EV (11%) and CB (6%). The enrichment of DG in FL is consistent with their known role in promoting changes of physicochemical properties of the membrane, membrane curvature and membrane dynamics (Campomanes *et al*., 2019), which are critical for flagellar membrane extension and structural remodelling. They often act as molecular cues that recruit and activate key protein involved in vesicle formation and membrane trafficking (Tanguy *et al*., 2016).

Interestingly, EV were markedly enriched in sphingolipids (SP), particularly ceramides (Cer), which accounted for 36% of total intensities, compared to 17% in FL and only 1% in CB (Figure 5A). Figure 5B illustrates the LION ontology analysis for enrichment of lipid-associated terms in EVs compared to CB and FL. The top 3 structural and physiological properties enriched in EVs include plasma membrane, negative intrinsic curvature and lipid-mediated signalling associated with ceramides and sphingolipids as the top 2 enriched lipid classes. Ceramides are bioactive lipids known to influence membrane rigidity, curvature, and vesicle budding. Their selective enrichment in EVs supports their involvement in EV biogenesis through the induction and stabilization of membrane curvature (Verderio *et al*., 2018; Dixson *et al*., 2023). A comprehensive summary of lipid terms analysis is provided in Supplemental Table 9.

Similarly, some phosphatidylethanolamines are enriched in EVs (Figure 5C) and, like ceramides and DAG, they are characterized by a small polar headgroup that confers a conical molecular geometry, thereby promoting negative (concave) membrane curvature and budding (Caputo *et al*., 2025, Skotland *et al*., 2017). Together, these findings strongly support the existence of active lipid sorting mechanisms during vesicle formation and highlight the potential contribution of these lipids to EV membrane integrity, structural stability and intercellular signalling.

The heatmap analysis of individual significantly enriched lipids (Figure 5C; Supplementary table 10) revealed clear organelle-specific lipid signatures. EV and FL displayed strong accumulation of 32 ceramide species, whereas these lipids were markedly reduced in CB. In contrast, phosphatidylethanolamine (PE 36:4, PE 34:1, PE 36:5, PE 32:1) and phosphatidylcholine (PC 32:2, PC 32:1, PC 36:4, PC 36:5, PC 34:2, PC 30:0, PC 36:4, PC 36:3, PC 16:0_18:1) molecules exhibited increased intensities in both EV and CB compared to FL. Lysoglycerophospholipids (LPC 20:4, 18:3, 20:5 were markedly enriched in CB and FL. Some diacylglycerol molecules (DG20:4_20:5, DG20:5_20:5, DG20:4_20:4) were strongly enriched in FL and reduced in EV and CB.

Lipidomic data demonstrate that flagella and extracellular vesicles are not simple extension of the cellular membrane but possess highly specialized lipid compositions. EVs are characterized by ceramide enrichment, consistent with membrane stabilization and signalling roles, whereas FL exhibit elevated diacylglycerol content, likely reflecting the dynamic membrane curvature requirements of motility structures. This lipid partitioning suggests active lipid sorting mechanisms during organelle biogenesis in oomycetes and points to potential functional specialization in host–pathogen interaction.

## DISCUSSION

While investigating the stramenopile *P. parasitica*, this study presents the first combined analysis—via microscopy, proteomics, and lipidomics—of the cellular and molecular traits of EVs shed by a zoospore. As expected, release occurs from the cell body as indicated by the spectacular multicurved morphology of the plasma membrane indicative of frequent events of EV biogenesis. Based on budding and extensive vesicularization of the tubular mastigoneme tips, together with the presence of PnMas1 in a subset of EVs, we can affirm that EVs can form and release from the front flagellum, as well. The biogenesis of EVs partly relying on mastigoneme-related processes is reminiscent with EV formation by budding and vesicularization from nanotubes of the flagellar membrane in the unicellular flagellated parasite *Trypanosoma brucei* (Szempruch *et al*., 2016^a,b^). In zoospores, flagella tips also mediate EV biogenesis (Supplementary figure 2). Thus, the formation of EVs may occur not only alongside the flagella but also at their tips as in primary and motile cilia (Vinay and Belleannée, 2022). Their release would result from the gradual tightening of the pinch zone (Figure 2) and would be favoured, during swimming either by (i) external factors such as the friction force generated between the front flagellum and an explored surface (e.g., a root of a host plant, a biofilm formed by zoospores; Bassani *et al*. 2020^b^); (ii) the drag force opposed to the undulating movement of the flagella, which propels the zoospore; (iii) or the water flow in which the zoospore swims, a strong mechanical stress that could also carry away the EVs.

Until now, mastigonemes are mainly considered to amplify the movement generated by flagellar beating. The tubular mastigonemes lead to a reversal of the direction of movement that would normally occur during flagellar beating if the mastigonemes were absent (Bouck, 1971; 1972). Beyond movement regulation according to these biophysical principles, with this study we determine a new function for mastigonemes as a source of EVs, putatively acting in cell–cell communications between zoospores, between zoospores and host plant cells or within the microbiota sharing the same biotope.

In congruence with the study of Breen *et al*. (2025) on EVs derived from *P. infestans* mycelium, we identified a MARVELous protein as a main marker for *P. parasitica* EVs shed by zoospores. These proteins are a constant in the molecular composition of EVs in *Phytophthora*. As the *P. infestans* proteins PiMDP1 and PiMDP2, the *P. parasitica* protein PPTG_13069 exhibits two juxtaposed MARVEL domains having each a four transmembrane-helix architecture. The resolution of the 3D-structure of these proteins should gain insights into the molecular mechanisms that determine how they act during EV biogenesis. Modelling the structure of PPTG_13069 using the SWISS-MODEL program, we identified the best two models (Supplementary figure 5) with regions aligned mostly consisting of amino acids located in the four transmembrane-helix regions of either synaptophysin (in the PPTG_13069 N-term) and synaptogyrin (in the C-term). These two proteins have a tetraspan MARVEL domain and are constitutive of small neurotransmitter containing vesicles (Sánchez-Pulido *et al*., 2002). They contribute to the formation and size regulation of mammal’s synaptic vesicles, the synaptophysin acting as a membrane curvature-promoter (Preobraschenski *et al*., 2025) and the synaptogyrin as a bending entity via its binding to phosphatidylserine (Yu *et al*., 2023).

The three known mastigoneme proteins with several other proteins with EFG-like domains are also included in the EV proteome. We propose the three mastigoneme proteins, in particular PPTG_02441, as markers of a sub set of zoospore EVs (EVs) allowing for the identification of EVs released from the tubular mastigonemes of the front flagellum. EGF-like domain proteins are reported as EV markers as the human membrane protein JAG1 or different subunits of matricellular proteins, such as laminins and integrins (Chitti *et al*., 2024). Some integrin subunits harbouring EGF-like domains are also involved in interactions with EV-related tetraspanins (CD9, CD63, CD81; Bassani and Cingolani, 2012). Nevertheless, it is difficult to speculate beyond on that basis because we have not identified in the proteome of *P. parasitica* any peptides with strong similarities to these human or animal proteins (Supplementary table 4). We can only note that as the flagellar proteome (Lupatelli *et al*., 2025), the zoospore EV proteome includes the three mastigoneme proteins and PPTG_13069 exhibiting similarly to PiMDP1 and PiMDP2 with two tetraspanning MARVEL domains (Supplementary figure 4).

More broadly, the combination of proteomic and lipidomic data provide a window toward a comprehensive understanding about the biology of EVs shed by zoospores.

One aspect is the convergent evidence associating enriched-LION terms (plasma membrane, negative intrinsic curvature, lipid-mediated signalling) with experimental proteomic data. They point out the potential involvement of lipid trafficking in the initiation of membrane curvature, bud formation and EV release. The ABC transporters identified in the EVs samples belongs to the ABCA and ABCG families, previously reported to catalyse lipid translocation across cellular membranes (Alam and Locher, 2023). In addition, Four EV transmembrane proteins are P4-ATPases orthologous to flippases catatalysing the translocation of phospholipids (mainly PC, PE and CER the most abundant lipids in EVs) from the outer to the inner plasma membrane of cell membranes (Pomorski *et al*. 2003; Sakuragi and Nagata, 2023; Kita *et al*., 2024). Accumulating evidence suggests that flippases contribute to the initiation of membrane curvature and EV release (Cocucci and Meldolesi, 2015; Meldolesi, 2018; Best *et al*., 2019) as well as influence extracellular vesicle cargo content as in *Cryptococcus neoformans* (Castelli *et al*., 2025). In *Phytophthora capsici*, deletion of *PcAPT1* encoding a P4-ATPase impairs hyphal growth, phospholipid transport and pathogenicity toward *Capsicum annuum* (Yang *et al*., 2022).

A second aspect refers to the involvement of EVs in the biosynthesis of cell wall polysaccharide biosynthesis that is attested by identification of four 1,3-β-glucan synthases, annexins and Lysophosphatidylethanolamine (LPE) within the EV membrane. This result is consistent with previous reports that LPEs, minor phosphoglycerides found in cell membranes, play a role in the activation of enzymes in isolated plasma membrane vesicles of plants (Palmgren and Sommarin, 1989; Ryu,2004). In addition, in the oomycete *Saprolegnia monoica,* LPE colocalizes with β-1,3-glucan synthases and annexins in lipid rafts, this complex being part of the process which regulate cell wall polysaccharide biosynthesis (Briolay *et al*., 2009). We can speculate that in *P. parasitica* zoospores, following the motile period and during encystment, EVs release could contribute to the fast elaboration and dynamics of the primary cell wall predominantly made of β-1,3-linked glucans (Tokunaga and Bartnicki-Garcia, 1971). It may be questioned in the context of fusion between EVs and recipient plasma membrane during the encystment of a sole cell, or during formation of monospecific biofilm within which recipient cells are zoospores that have co-migrated and aggregated as cysts to the same site on the host surface (Galiana *et al*., 2008).

At first sight we could say this study takes advantages of an investigation on EVs produced by a single cell type, homogenous in size (∼100 nm) and collected in the simplest possible environment: water, in which these swimming cells naturally evolve. This ensures that the study is conducted with zero risk of contamination by the medium, and that it targets a population of EVs few diverse in terms of nature and function compared to studies on EVs collected from body fluids or various forms that a microorganism can present (EVs which are heterogeneous, in size, in nature and have diverse functions). Nevertheless, our results allow us to distinguish at least 2 populations. EVs PnMas1^+^ and EVs PnMas1^−^ that schematically could be EVs released from the front flagellum and those released from the cell body, respectively. Future investigation will be to fractionate the two EVs populations by fluorescence-activated cell sorting in order to delineate their composition (proteins, lipids, nucleic acids, metabolites) that underly any respective role in cell-to-cell communication.

## CONCLUSION

Microorganism-derived EVs are abundant in moist or aquatic environments, but the understanding of their biogenesis and functions is still limited (Schatz and Vardi, 2018; Gill *et al*., 2019). Our data represent a significant step forward for studying EV biology in the extended group of zoosporic microorganisms, in particular the heterokonts harbouring in their biflagellate cell form two different flagella. We establish that EV biogenesis occurs from either the cell body, flagella and tubular mastigonemes. We identified protein and lipid core composition of EVs in *P. parasitica*, characterized by at least two EVs subpopulations, one of which is decorated with a mastigoneme protein.

Another perspective to consider would be to isolate and characterize EVs produced by zoospores from various taxonomic groups within stramenopiles and beyond *Phytophthora*. Building on the finding that zoospore-EV proteomes contain mastigoneme proteins—whose sequence varies across taxa (Hee *et al*., 2019; Harada *et al*., 2026)—investigation on EVs released in aquatic environments Schatz and Vardi, 2018) could provide new insights about roles of these EVs in communication within microbial community.

## ACKNOWLEDGMENTS

The authors thank the PlantBios Collective and Scientific Infrasctucture at Institut Sophia Agrobiotech for access : (i) to microscopy facilities; (ii) to lipidomic and proteomic analysis equipment (Biochemistry Analytical platform : BA Sophia Antipolis from the ISC plantbios https://doi.org/10.15454/qyey-ar89); and (iii) for the access to the PAB-Azur platform at Institut de Pharmacologie Moleculaire et Cellulaire, funded through the SABLES platforms project with support from the European Union and the European Regional Development Fund. The authors also thank the Centre Commun de Microscopie Appliquée (CCMA, Université Côte d’Azur) for access to equipment and expertise in structural characterization using electron microscopy. The authors thank Georges de Sousa for helpful discussions.

## FUNDING

This work was supported by the National Research Agency; project no. ANR22-CE20-0021.

## AUTHORS’ CONTRIBUTIONS

MLK: conceptualization, data curation, formal analysis, investigation, validation, writing original draft.

CL: conceptualization, data curation, formal analysis, investigation, validation, writing original draft

BMH: conceptualization, data curation, formal analysis and validation of lipidomics analysis. Writing original draft.

AS: conceptualization, data curation, formal analysis and validation of proteomic investigation. Writing original draft

ASG: conceptualization, data curation, formal analysis and validation of proteomic investigation

SP: processed the samples and performed the TEM observations.

FO: processed the samples and performed the SEM observations.

IB: conceptualization, data curation, formal analysis, validation, writing original draft.

EG: conceptualization, data curation, formal analysis, investigation, validation, writing original draft. Funding acquisition.

All authors reviewed, revised, and approved the final version of the manuscript.

## DECLARATION OF INTEREST STATEMENT

The authors declare no competing interests.

## TABLES WITH CAPTIONS

**SUPPLEMENTARY TABLE 1**

Proteomics data acquired in positive polarity using the Parallel Accumulation-Serial Fragmentation (PASEF) Data Dependent Acquisition (DDA) mode.

**SUPPLEMENTARY TABLE 2**

Annotations of identified and selected EV protein candidates in terms of being part of a family, putative EV features, and gene ontology.

**SUPPLEMENTARY TABLE 3**

Orthologs of EVs-related proteins between *P.parasitica* and *P. infestans*.

**SUPPLEMENTARY TABLE 4**

List of *P. parasitica* proteins exhibiting or not significant similarity (BlastP e-Value < E^−6^) with known EV markers in other species.

**SUPPLEMENTARY TABLE 5**

GO terms analysis with g:Profiler for enrichment of TOP400 proteins in molecular functions, cellular components, and biological processes.

**SUPPLEMENTARY TABLE 6**

Protein family expansion and subcellular localisation of the TOP400 proteins.

**SUPPLEMENTARY TABLE 7**

An overlap of detected membrane associated proteins between this study and Lupatelli *et al*., 2025.

**SUPPLEMENTARY TABLE 8**

Total number and lipid class distribution of annotated lipids in cell bodies (CB), flagella (FL), and extracellular vesicles (EV).

**SUPPLEMENTARY TABLE 9**

LION-term enrichment analysis of EV samples.

**SUPPLEMENTARY TABLE 10**

Univariate analysis of lipid species quantification (P < 0.05).

---

**SUPPLEMENTARY FIGURE 1.**
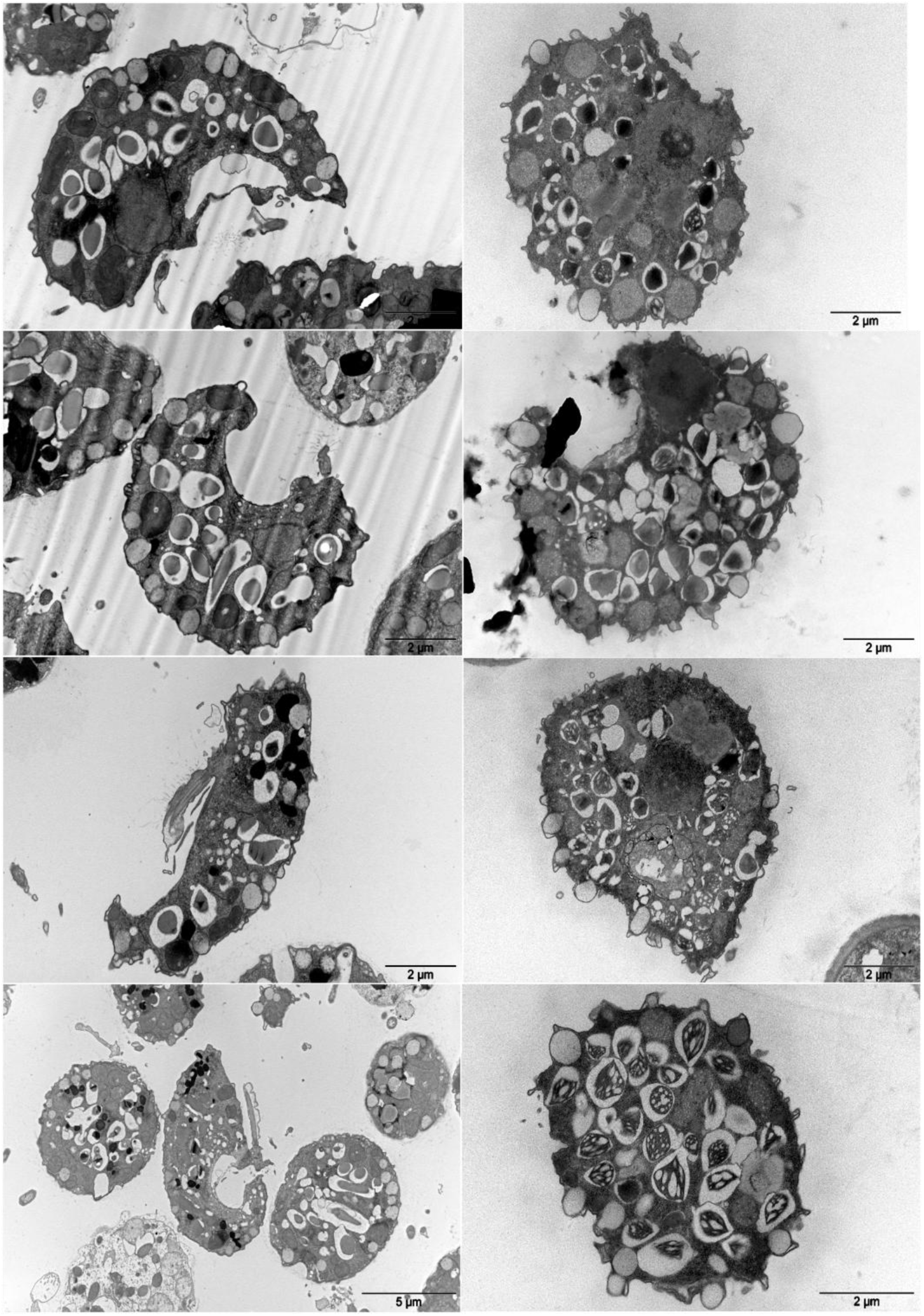
TEM micrographs of zoospore sections. Longitudinal (left micrographs) and transversal (right micrographs) thin-sections of zoospores (80nm) showing cellular morphotypes with several intracellular vesicles bordering plasma membrane and inducing membranous curvature. Zoospores were fixed 5 min after release from sporangia.

**SUPPLEMENTARY FIGURE 2.**
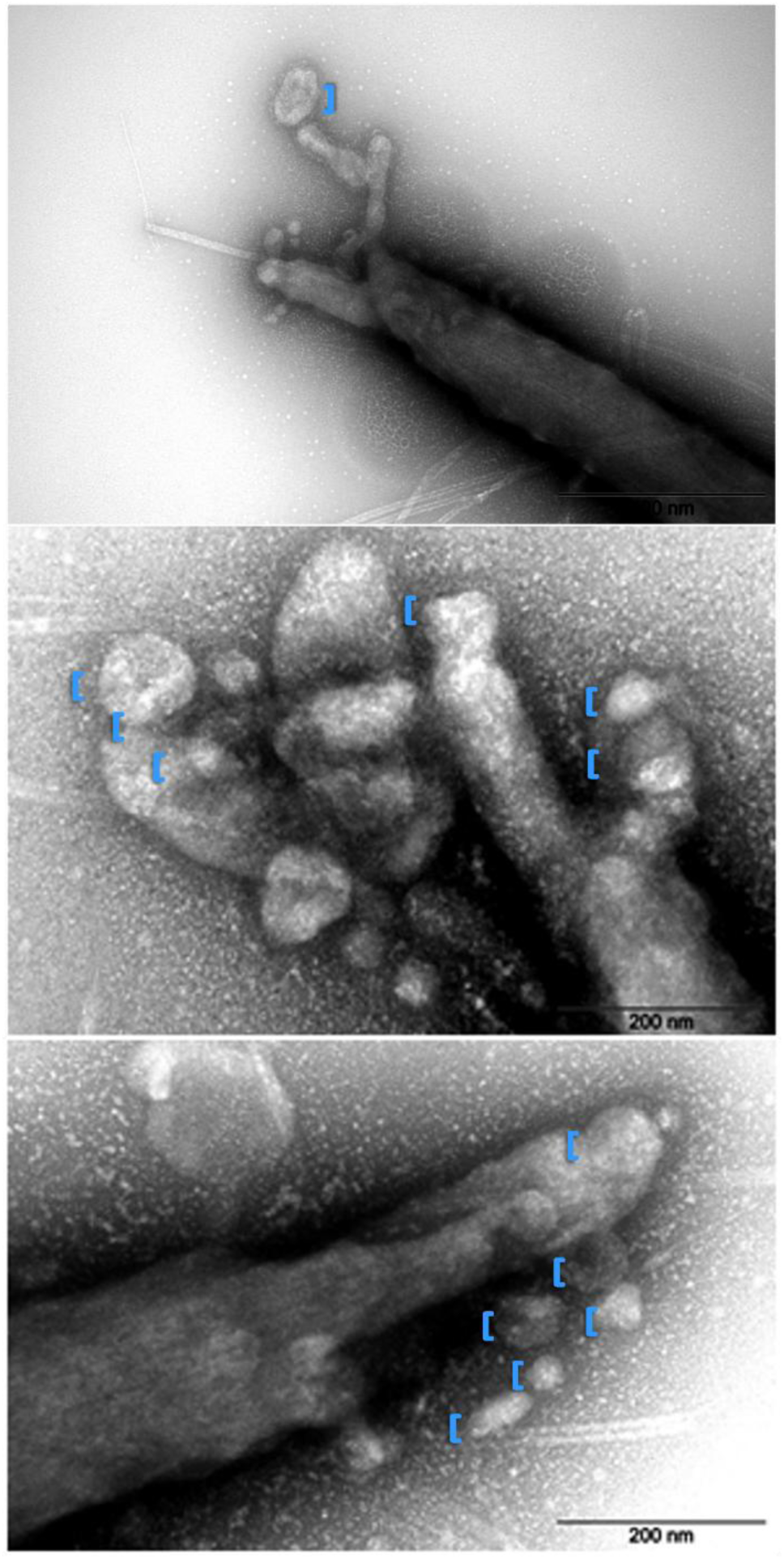
Ultrastructure of the tip of flagella TEM of spheroidal structures emerging at the tip of flagella that are similar in size and shape to those formed at the mastigoneme tip suggesting that EVs release from flagellum may be of lateral and apical origin.

**SUPPLEMENTARY FIGURE 3.**
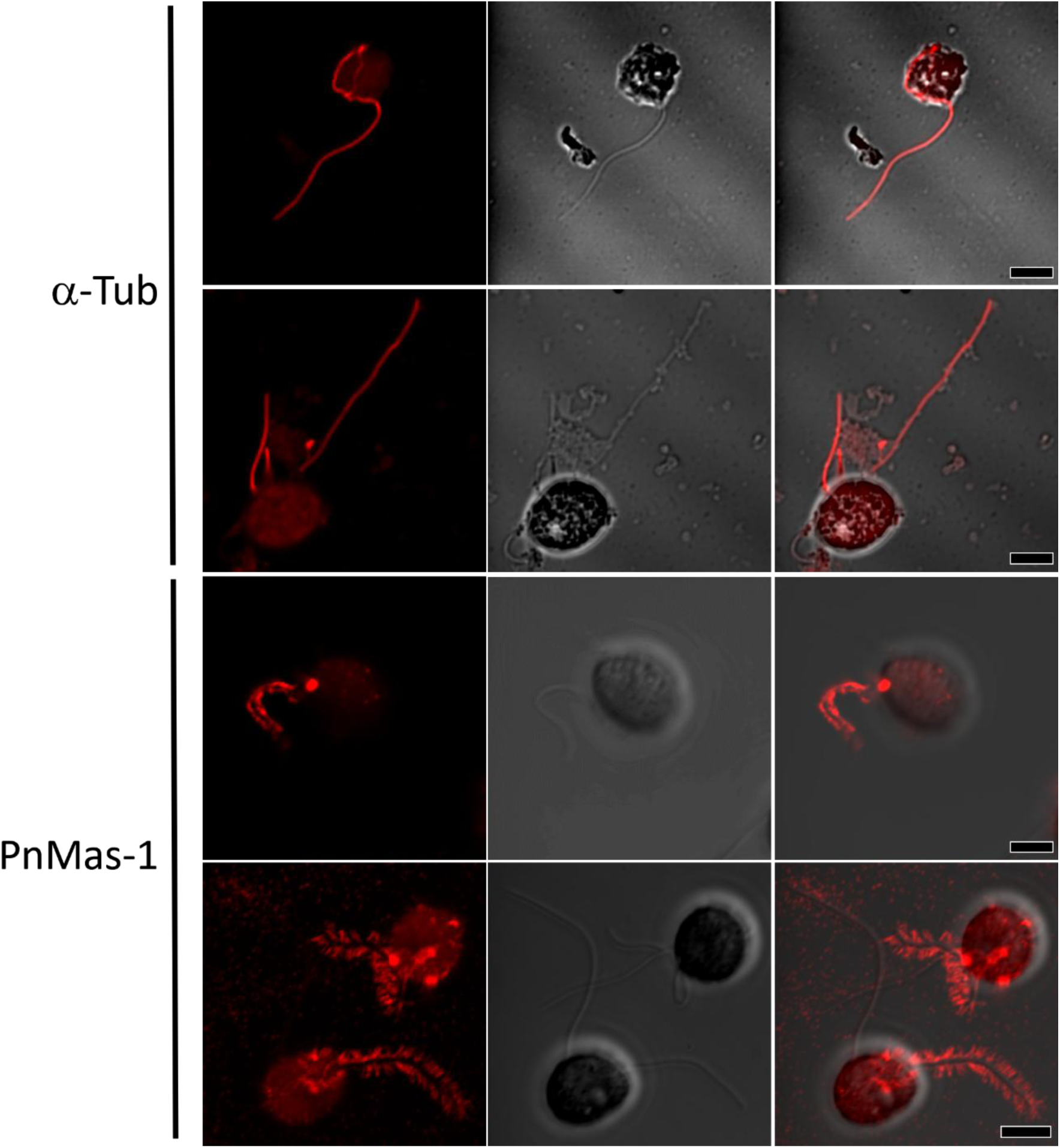
Immunofluorescence assay for α-tubulin and PnMas1 on zoospores. Views of bright field and immunofluorescence merged images acquired with a Microscope Axio Imager Z1 after incubation with rabbit antibodies against α-tubulin (α**Tub**) or against PnMas1. The antibody against α-tubulin decorates the anterior (**a-fl**) and the posterior (**p-fl**) flagellum while the PnMas1 antibody only stains the mastigonemes of the anterior flagellum. Bars: 5µm.

**SUPPLEMENTARY FIGURE 4.**
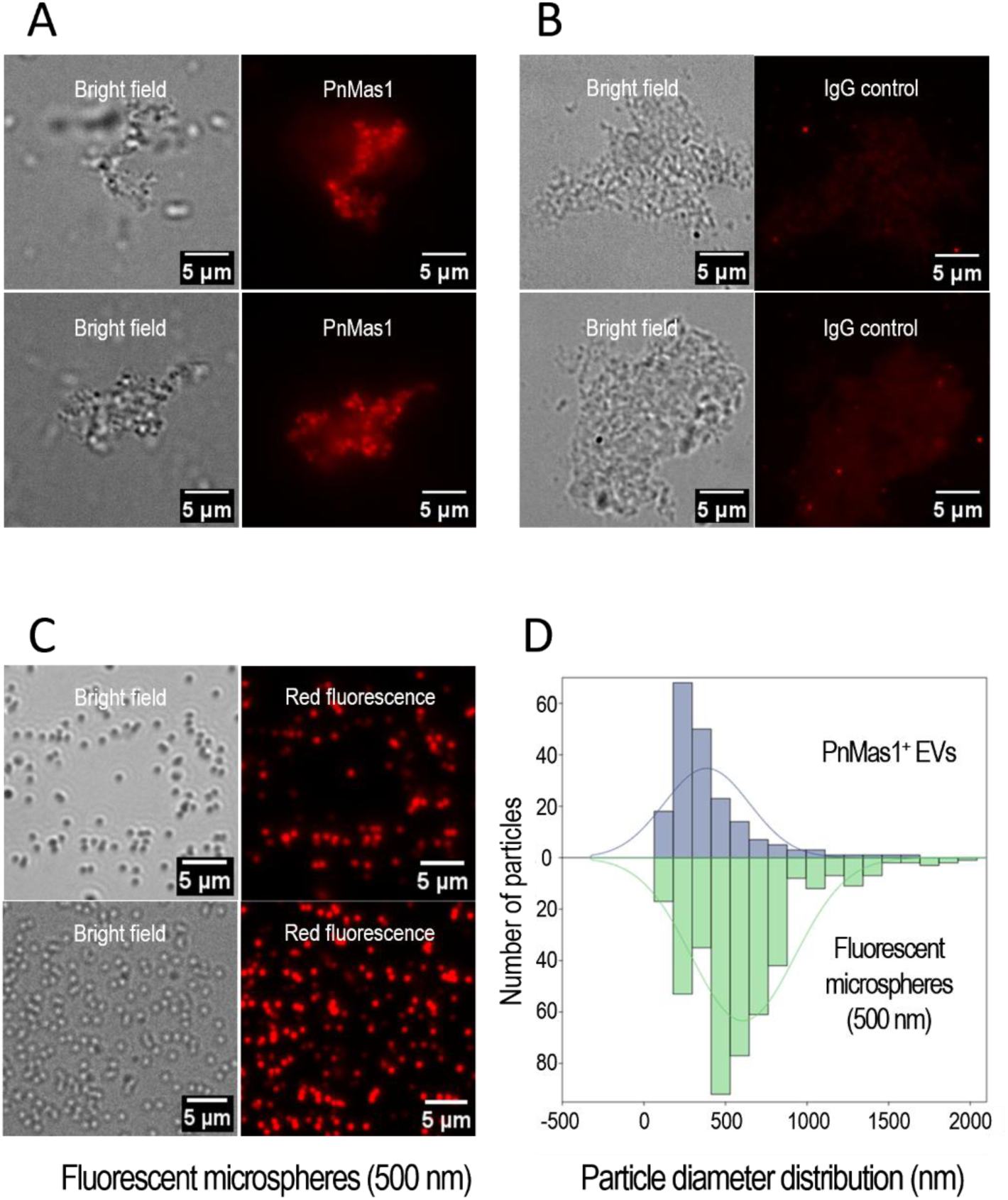
Immunofluorescence and TEM assay for PnMas1 on EVs (**A, B**) Bright field and immunofluorescence of particles collected in EV-p100k pellet fractions stained with an anti-PnMas1 antibody (**A**) or with Immunoglobulin G (igG) purified from non-immune serum (**B**). (**C**) Bright field and red fluorescence of microspheres (diameter=500 nm). (**D**) Bihistogram with normal curves showing the distribution of diameter ranges estimated for PnMas1^+^ EVs and fluorescent microspheres.

**SUPPLEMENTARY FIGURE 5.**
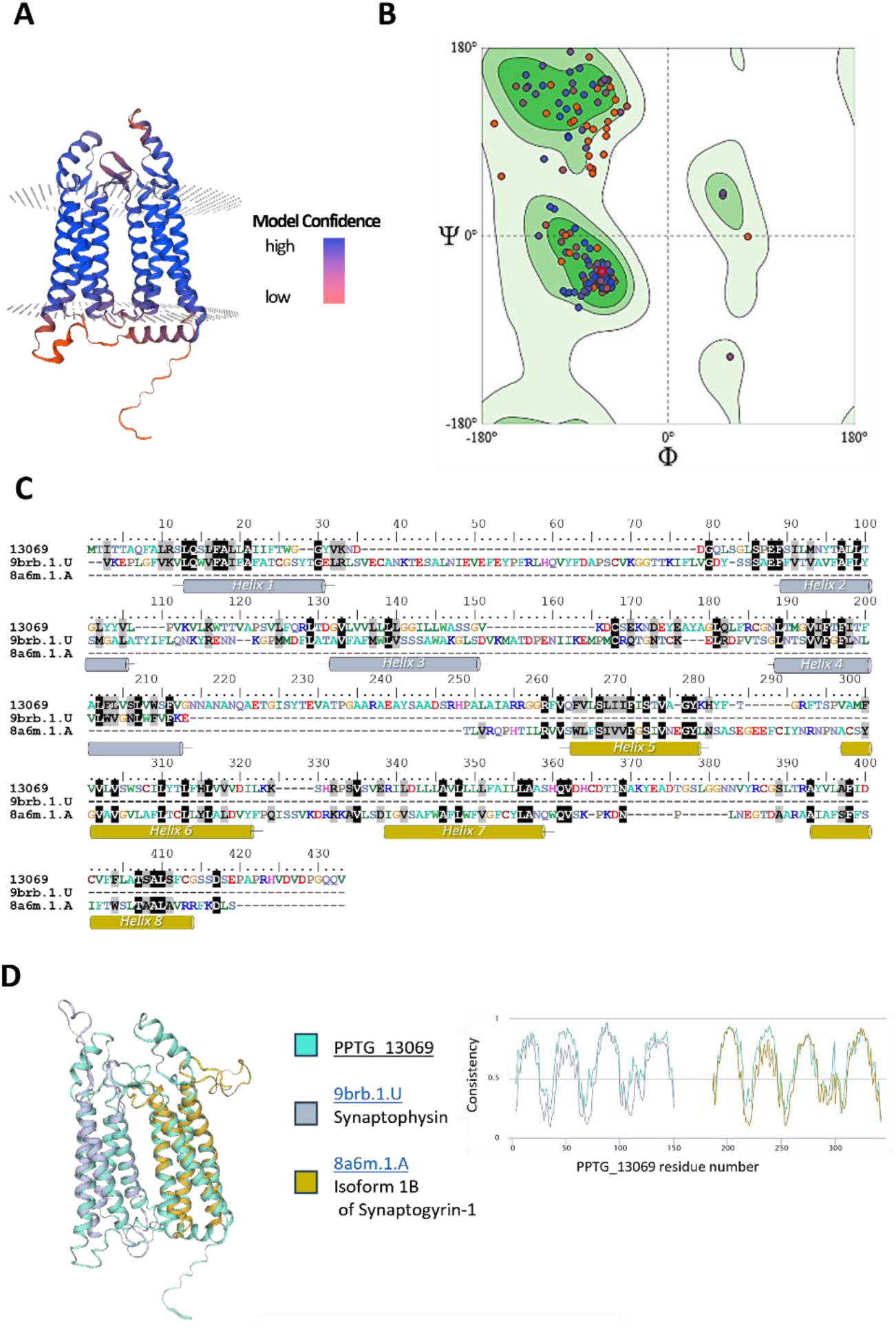
Predicted 3D-structure of PPTG_13069. (**A**) Predicted 3D-structure of PPTG_13069 produced using the SWISS-MODEL server and showing tight reminiscence with the predicted structure of the MARVELous proteins PiMDP1 and PiMDP2 markers for *Phytophthora infestans* EVs (Breen *et al*., 2025). Colors denote the model confidence factor in gradient from purple (high confidence, mainly the 8 transmembrane domains) to orange (low confidence). (**B**) Ramachandran plot showing most points (95.21%) clustered in the core and allowed regions. (**C**) Clustal W alignment of PPTG_13069, the rat Synaptophysin and the human Synaptogyrin-1 protein sequences. Coloring is based upon amino acid properties. The helices set the sequences corresponding to the 8 transmembrane domains of PPG_13069. (**D**) In the left panel, the structural alignment between the predicted 3D-structure of PPTG_13069 (cyan), the crystal structure of Synaptophysin (yellow-brown, PDB: 9brb; QMEANDisCo Global: 0.44 ± 0.07) and the Isoform 1B of Synaptogyrin-1 (grey PDB: 8a6m; QMEANDisCo Global: 0.44 ± 0.07). In the right panel, the consistency score resulting of the structure comparison of the three structures and showing higher scores for the 8 transmembrane domains.

## Notes

### Competing Interest Statement

The authors have declared no competing interest.

