## Supplementary table 1 for "Zoospore-derived extracellular vesicles in the flagellated stramenopile *Phytophthora parasitica*"

**Title: proteomics data acquired in positive polarity using the Parallel Accumulation-Serial Fragmentation (PASEF) Data Dependent Acquisition (DDA) mode**

| Protein Group | Protein ID | Accession | -10logP | Coverage (%) | Coverage (%) EV1-2 | Coverage (%) EV3-4 | Coverage (%) EV3et4 | Area EV1-2 | Area EV3-4 | Area EV3et4 | #Peptides | #Unique | #Spec EV1-2 | #Spec EV3-4 | #Spec EV3et4 | PTM | Avg, Mass | Description |
|---|---|---|---|---|---|---|---|---|---|---|---|---|---|---|---|---|---|---|
| 1 | 11 | PPTG_04587 | 402,1 | 42 | 22 | 26 | 40 | 5,92E+07 | 1,45E+08 | 8,75E+08 | 260 | 260 | 160 | 208 | 392 | Oxidation (M); Carbamidomethylation; Dehydration; Deamidation (NQ); Acetylation (N-term) | 1109158 | \| PPTG_04587 \| Phytophthora parasitica INRA-310 hypothetical protein (10566 aa) |
| 2 | 1 | PPTG_11032 | 378,39 | 75 | 58 | 71 | 73 | 8,27E+08 | 2,81E+09 | 1,06E+10 | 50 | 50 | 96 | 214 | 354 | Oxidation (M); Carbamidomethylation; Dehydration; Deamidation (NQ); Acetylation (N-term); 2 more | 139968 | \| PPTG_11032 \| Phytophthora parasitica INRA-310 hypothetical protein (1373 aa) |
| 8 | 2 | PPTG_06389 | 324,08 | 58 | 46 | 55 | 58 | 4,49E+08 | 1,39E+09 | 4,10E+09 | 39 | 34 | 62 | 107 | 147 | Oxidation (M); Carbamidomethylation; Dehydration; Deamidation (NQ); Acetylation (N-term); 2 more | 94441 | \| PPTG_06389 \| Phytophthora parasitica INRA-310 hypothetical protein (879 aa) |
| 4 | 27 | PPTG_01092 | 323,63 | 35 | 16 | 25 | 34 | 1,47E+07 | 5,98E+07 | 2,98E+08 | 135 | 116 | 65 | 113 | 195 | Oxidation (M); Carbamidomethylation; Dehydration; Deamidation (NQ); Amidation | 607520 | \| PPTG_01092 \| Phytophthora parasitica INRA-310 hypothetical protein (5728 aa) |
| 14 | 18 | PPTG_11936 | 317,04 | 52 | 30 | 46 | 47 | 5,14E+07 | 4,92E+08 | 5,26E+08 | 77 | 77 | 50 | 102 | 111 | Oxidation (M); Carbamidomethylation; Dehydration; Deamidation (NQ); Acetylation (N-term) | 194028 | \| PPTG_11936 \| Phytophthora parasitica INRA-310 clathrin heavy chain (1720 aa) |
| 49 | 6 | PPTG_16264 | 314,02 | 37 | 13 | 29 | 34 | 1,52E+07 | 7,69E+07 | 1,92E+08 | 49 | 49 | 22 | 57 | 66 | Oxidation (M); Carbamidomethylation; Deamidation (NQ); Amidation; Carbamylation | 258251 | \| PPTG_16264 \| Phytophthora parasitica INRA-310 thrombospondin-like protein (2400 aa) |
| 15 | 10 | PPTG_17090 | 301,43 | 61 | 47 | 58 | 55 | 7,35E+07 | 4,82E+08 | 7,95E+08 | 52 | 52 | 49 | 96 | 100 | Oxidation (M); Carbamidomethylation; Dehydration; Deamidation (NQ); Amidation; Carbamylation | 123068 | \| PPTG_17090 \| Phytophthora parasitica INRA-310 phosphoglucomutase (1126 aa) |
| 12 | 40 | PPTG_01091 | 299,77 | 32 | 10 | 18 | 30 | 5,06E+06 | 2,21E+07 | 1,14E+08 | 129 | 110 | 44 | 78 | 150 | Oxidation (M); Carbamidomethylation; Dehydration; Deamidation (NQ) | 630558 | \| PPTG_01091 \| Phytophthora parasitica INRA-310 hypothetical protein (5916 aa) |
| 3 | 3 | PPTG_02441 | 298,82 | 65 | 64 | 64 | 65 | 8,26E+08 | 2,04E+09 | 9,56E+09 | 30 | 30 | 69 | 116 | 251 | Oxidation (M); Carbamidomethylation; Dehydration; Deamidation (NQ); Acetylation (N-term); 2 more | 64040 | \| PPTG_02441 \| Phytophthora parasitica INRA-310 hypothetical protein (598 aa) |
| 23 | 14 | PPTG_09633 | 297,63 | 54 | 38 | 34 | 54 | 6,04E+07 | 5,34E+07 | 9,24E+08 | 59 | 59 | 43 | 45 | 117 | Oxidation (M); Carbamidomethylation; Dehydration; Deamidation (NQ) | 147770 | \| PPTG_09633 \| Phytophthora parasitica INRA-310 sodium/potassium-transporting ATPase subunit alpha (1344 aa) |
| 16 | 19 | PPTG_02434 | 295,49 | 72 | 45 | 67 | 69 | 8,46E+07 | 4,33E+08 | 9,21E+08 | 53 | 53 | 45 | 88 | 106 | Oxidation (M); Carbamidomethylation; Dehydration; Deamidation (NQ); Acetylation (N-term); 2 more | 95440 | \| PPTG_02434 \| Phytophthora parasitica INRA-310 elongation factor 2 (859 aa) |
| 29 | 8 | PPTG_16242 | 292,42 | 45 | 35 | 42 | 44 | 8,17E+07 | 3,39E+08 | 9,64E+08 | 32 | 32 | 28 | 60 | 87 | Oxidation (M); Carbamidomethylation; Deamidation (NQ); Amidation; Carbamylation | 92829 | \| PPTG_16242 \| Phytophthora parasitica INRA-310 hypothetical protein (855 aa) |
| 5 | 13 | PPTG_15107 | 290,83 | 57 | 42 | 57 | 53 | 7,44E+07 | 1,13E+09 | 1,18E+09 | 42 | 41 | 46 | 146 | 132 | Oxidation (M); Carbamidomethylation; Dehydration; Deamidation (NQ); Acetylation (N-term) | 120521 | \| PPTG_15107 \| Phytophthora parasitica INRA-310 hypothetical protein (1080 aa) |
| 6 | 9 | PPTG_13841 | 283,63 | 79 | 63 | 75 | 78 | 3,30E+08 | 9,37E+08 | 4,83E+09 | 28 | 28 | 63 | 86 | 168 | Oxidation (M); Carbamidomethylation; Dehydration; Deamidation (NQ); Acetylation (N-term); Carbam | 53494 | \| PPTG_13841 \| Phytophthora parasitica INRA-310 ATP synthase subunit beta mitochondrial (502 aa) |
| 11 | 151 | PPTG_04733 | 280,93 | 34 | 13 | 17 | 32 | 1,18E+07 | 3,50E+07 | 1,90E+08 | 154 | 154 | 62 | 77 | 156 | Oxidation (M); Carbamidomethylation; Dehydration; Deamidation (NQ); Amidation | 599526 | \| PPTG_04733 \| Phytophthora parasitica INRA-310 hypothetical protein (5309 aa) |
| 9 | 382 | PPTG_03648 | 271,67 | 53 | 51 | 52 | 53 | 3,08E+08 | 8,76E+08 | 3,96E+09 | 19 | 19 | 47 | 64 | 192 | Oxidation (M); Carbamidomethylation; Dehydration; Deamidation (NQ); Acetylation (N-term); 2 more | 42002 | \| PPTG_03648 \| Phytophthora parasitica INRA-310 hypothetical protein (394 aa) |
| 126 | 15 | PPTG_06394 | 265,09 | 44 | 29 | 34 | 42 | 9,58E+06 | 1,18E+07 | 6,92E+07 | 34 | 19 | 28 | 27 | 46 | Oxidation (M); Carbamidomethylation; Deamidation (NQ); Amidation | 123448 | \| PPTG_06394 \| Phytophthora parasitica INRA-310 hypothetical protein (1126 aa) |
| 24 | 160 | PPTG_05265 | 264,3 | 25 | 7 | 19 | 22 | 5,07E+06 | 4,05E+07 | 7,03E+07 | 95 | 95 | 30 | 78 | 96 | Oxidation (M); Carbamidomethylation; Deamidation (NQ); Acetylation (N-term) | 532830 | \| PPTG_05265 \| Phytophthora parasitica INRA-310 dynein heavy chain (4707 aa) |
| 62 | 16 | PPTG_06382 | 260,17 | 38 | 26 | 32 | 37 | 1,86E+07 | 7,48E+07 | 2,08E+08 | 28 | 23 | 26 | 46 | 62 | Oxidation (M); Carbamidomethylation; Dehydration; Deamidation (NQ); Carbamylation | 115701 | \| PPTG_06382 \| Phytophthora parasitica INRA-310 hypothetical protein (1090 aa) |
| 86 | 21 | PPTG_02260 | 259,97 | 40 | 22 | 36 | 38 | 7,26E+06 | 7,11E+07 | 1,36E+08 | 34 | 34 | 21 | 45 | 51 | Oxidation (M); Carbamidomethylation; Dehydration; Deamidation (NQ); Amidation | 123071 | \| PPTG_02260 \| Phytophthora parasitica INRA-310 leucyl-tRNA synthetase (1086 aa) |
| 36 | 80 | PPTG_08772 | 259,32 | 50 | 24 | 44 | 45 | 2,01E+07 | 1,47E+08 | 1,94E+08 | 59 | 58 | 31 | 67 | 65 | Oxidation (M); Carbamidomethylation; Dehydration; Acetylation (N-term) | 163962 | \| PPTG_08772 \| Phytophthora parasitica INRA-310 carbamoyl-phosphate synthase [ammonia] (1500 aa) |
| 22 | 198 | PPTG_14782 | 258,46 | 21 | 4 | 17 | 17 | 2,56E+06 | 3,82E+07 | 5,75E+07 | 106 | 106 | 25 | 91 | 92 | Oxidation (M); Carbamidomethylation; Deamidation (NQ) | 735749 | \| PPTG_14782 \| Phytophthora parasitica INRA-310 regulator-chromosome condensation (RCC1)-like protein (6743 aa) |
| 115 | 31 | PPTG_14544 | 257,61 | 50 | 20 | 46 | 47 | 5,18E+06 | 7,97E+07 | 9,51E+07 | 34 | 34 | 14 | 45 | 45 | Oxidation (M); Carbamidomethylation; Dehydration; Deamidation (NQ); Carbamylation | 97950 | \| PPTG_14544 \| Phytophthora parasitica INRA-310 26S proteasome non-ATPase regulatory subunit 2 (898 aa) |
| 64 | 20 | PPTG_20351 | 256,78 | 67 | 50 | 65 | 67 | 1,84E+07 | 2,84E+08 | 2,95E+08 | 18 | 17 | 24 | 52 | 58 | Oxidation (M); Carbamidomethylation; Deamidation (NQ) | 51969 | \| PPTG_20351 \| Phytophthora parasitica INRA-310 hypothetical protein (466 aa) |
| 54 | 72 | PPTG_06857 | 256,69 | 45 | 19 | 21 | 45 | 1,09E+07 | 1,50E+07 | 1,40E+08 | 61 | 61 | 25 | 29 | 87 | Oxidation (M); Carbamidomethylation; Dehydration; Deamidation (NQ); Carbamylation | 198215 | \| PPTG_06857 \| Phytophthora parasitica INRA-310 hypothetical protein (1765 aa) |
| 30 | 30 | PPTG_02122 | 254,77 | 58 | 40 | 57 | 58 | 4,40E+07 | 3,28E+08 | 4,17E+08 | 33 | 23 | 32 | 71 | 71 | Oxidation (M); Carbamidomethylation; Deamidation (NQ) | 71670 | \| PPTG_02122 \| Phytophthora parasitica INRA-310 hsp70-like protein (658 aa) |
| 17 | 54 | PPTG_05028 | 254,74 | 57 | 51 | 54 | 57 | 1,75E+08 | 3,76E+08 | 2,07E+09 | 40 | 40 | 54 | 66 | 119 | Oxidation (M); Carbamidomethylation; Dehydration; Deamidation (NQ); Carbamylation | 83125 | \| PPTG_05028 \| Phytophthora parasitica INRA-310 acyl-CoA dehydrogenase (765 aa) |
| 35 | 144 | PPTG_14525 | 253,13 | 47 | 19 | 44 | 43 | 2,10E+07 | 1,59E+08 | 2,00E+08 | 55 | 54 | 25 | 70 | 68 | Oxidation (M); Carbamidomethylation; Dehydration; Deamidation (NQ); Carbamylation | 166317 | \| PPTG_14525 \| Phytophthora parasitica INRA-310 carbamoyl-phosphate synthase [ammonia] (1512 aa) |
| 205 | 17 | PPTG_06395 | 250,16 | 34 | 23 | 31 | 33 | 5,19E+06 | 3,21E+06 | 3,16E+07 | 26 | 11 | 22 | 23 | 36 | Oxidation (M); Carbamidomethylation; Deamidation (NQ); Amidation | 119315 | \| PPTG_06395 \| Phytophthora parasitica INRA-310 hypothetical protein (1090 aa) |
| 161 | 38 | PPTG_12938 | 250,14 | 43 | 18 | 22 | 43 | 7,48E+06 | 8,60E+06 | 1,24E+08 | 34 | 25 | 15 | 21 | 54 | Oxidation (M); Carbamidomethylation; Deamidation (NQ) | 115651 | \| PPTG_12938 \| Phytophthora parasitica INRA-310 cellulose synthase catalytic subunit (1017 aa) |
| 74 | 51 | PPTG_12937 | 249,82 | 48 | 29 | 26 | 48 | 2,01E+07 | 2,48E+07 | 2,11E+08 | 40 | 31 | 28 | 28 | 70 | Oxidation (M); Carbamidomethylation; Deamidation (NQ) | 116212 | \| PPTG_12937 \| Phytophthora parasitica INRA-310 cellulose synthase 2 (1027 aa) |
| 136 | 23 | PPTG_02292 | 249,71 | 47 | 21 | 47 | 45 | 6,06E+06 | 1,09E+08 | 1,03E+08 | 32 | 32 | 14 | 45 | 39 | Oxidation (M); Carbamidomethylation; Deamidation (NQ); Acetylation (N-term) | 97060 | \| PPTG_02292 \| Phytophthora parasitica INRA-310 villin-like protein (882 aa) |
| 77 | 41 | PPTG_03714 | 249,4 | 78 | 51 | 76 | 78 | 4,40E+07 | 3,64E+08 | 4,39E+08 | 26 | 26 | 20 | 54 | 49 | Oxidation (M); Carbamidomethylation; Deamidation (NQ) | 43489 | \| PPTG_03714 \| Phytophthora parasitica INRA-310 phosphoglycerate kinase 1 (415 aa) |
| 40 | 26 | PPTG_02565 | 248,99 | 37 | 30 | 36 | 36 | 8,13E+07 | 4,15E+08 | 1,11E+09 | 25 | 25 | 32 | 55 | 70 | Oxidation (M); Carbamidomethylation; Dehydration; Deamidation (NQ); Acetylation (N-term); Amidatic | 85672 | \| PPTG_02565 \| Phytophthora parasitica INRA-310 hypothetical protein (809 aa) |
| 59 | 85 | PPTG_12232 | 248,74 | 32 | 18 | 17 | 30 | 2,16E+07 | 3,17E+07 | 2,72E+08 | 51 | 43 | 31 | 33 | 73 | Oxidation (M); Carbamidomethylation; Deamidation (NQ) | 213194 | \| PPTG_12232 \| Phytophthora parasitica INRA-310 ATP-binding Cassette (ABC) superfamily (1949 aa) |
| 33 | 92 | PPTG_01206 | 246,97 | 50 | 32 | 46 | 47 | 1,79E+07 | 1,82E+08 | 2,39E+08 | 47 | 47 | 32 | 65 | 68 | Oxidation (M); Carbamidomethylation; Dehydration; Deamidation (NQ) | 130208 | \| PPTG_01206 \| Phytophthora parasitica INRA-310 pyruvate carboxylase (1197 aa) |
| 48 | 35 | PPTG_01101 | 246,58 | 83 | 71 | 80 | 83 | 2,42E+07 | 1,46E+08 | 2,22E+08 | 21 | 7 | 31 | 55 | 61 | Oxidation (M); Carbamidomethylation; Deamidation (NQ); Amidation; Carbamylation | 36163 | \| PPTG_01101 \| Phytophthora parasitica INRA-310 glyceraldehyde-3-phosphate dehydrogenase (340 aa) |
| 102 | 43 | PPTG_19375 | 246,52 | 55 | 24 | 48 | 52 | 6,07E+06 | 5,76E+07 | 8,55E+07 | 36 | 36 | 17 | 45 | 46 | Oxidation (M); Carbamidomethylation; Dehydration; Deamidation (NQ); Acetylation (N-term); Amidatic | 89811 | \| PPTG_19375 \| Phytophthora parasitica INRA-310 hypothetical protein (810 aa) |
| 41 | 28 | PPTG_20013 | 246,11 | 70 | 58 | 70 | 65 | 3,48E+07 | 3,74E+08 | 4,17E+08 | 24 | 23 | 25 | 64 | 66 | Oxidation (M); Carbamidomethylation; Deamidation (NQ) | 49046 | \| PPTG_20013 \| Phytophthora parasitica INRA-310 enolase (455 aa) |
| 41 | 29 | PPTG_13544 | 246,11 | 70 | 58 | 70 | 65 | 3,48E+07 | 3,74E+08 | 4,17E+08 | 24 | 23 | 25 | 64 | 66 | Oxidation (M); Carbamidomethylation; Deamidation (NQ) | 49046 | \| PPTG_13544 \| Phytophthora parasitica INRA-310 enolase (455 aa) |
| 487 | 12 | PPTG_06730 | 245,82 | 48 | 16 | 30 | 46 | 1,01E+06 | 5,73E+06 | 3,54E+07 | 19 | 19 | 6 | 13 | 24 | Oxidation (M); Carbamidomethylation; Deamidation (NQ) | 70113 | \| PPTG_06730 \| Phytophthora parasitica INRA-310 hypothetical protein (677 aa) |
| 65 | 146 | PPTG_08579 | 245,01 | 30 | 14 | 14 | 29 | 1,14E+07 | 1,39E+07 | 1,68E+08 | 53 | 53 | 28 | 27 | 74 | Oxidation (M); Carbamidomethylation; Dehydration; Deamidation (NQ) | 250428 | \| PPTG_08579 \| Phytophthora parasitica INRA-310 callose synthase (2229 aa) |
| 50 | 52 | PPTG_07612 | 244,51 | 51 | 24 | 50 | 44 | 8,61E+06 | 8,76E+07 | 1,27E+08 | 40 | 40 | 26 | 60 | 59 | Oxidation (M); Carbamidomethylation; Dehydration; Deamidation (NQ) | 108067 | \| PPTG_07612 \| Phytophthora parasitica INRA-310 AP-2 complex subunit alpha (988 aa) |
| 39 | 50 | PPTG_10603 | 244,39 | 64 | 40 | 64 | 63 | 2,90E+07 | 2,26E+08 | 3,35E+08 | 32 | 32 | 28 | 66 | 63 | Oxidation (M); Carbamidomethylation; Dehydration; Deamidation (NQ); Acetylation (N-term); Carbam | 85646 | \| PPTG_10603 \| Phytophthora parasitica INRA-310 creatine kinase (786 aa) |
| 45 | 59 | PPTG_07345 | 244,2 | 47 | 29 | 26 | 45 | 2,21E+07 | 2,17E+07 | 2,74E+08 | 43 | 37 | 34 | 29 | 84 | Oxidation (M); Carbamidomethylation; Deamidation (NQ); Acetylation (N-term); Carbamylation | 114304 | \| PPTG_07345 \| Phytophthora parasitica INRA-310 P-type ATPase (P-ATPase) superfamily (1049 aa) |
| 51 | 33 | PPTG_03836 | 243 | 59 | 32 | 59 | 55 | 3,45E+07 | 5,16E+08 | 4,95E+08 | 25 | 25 | 23 | 59 | 62 | Oxidation (M); Carbamidomethylation; Dehydration; Deamidation (NQ); Amidation; Carbamylation | 59736 | \| PPTG_03836 \| Phytophthora parasitica INRA-310 secretory protein OPEL (557 aa) |
| 83 | 68 | PPTG_01093 | 242,25 | 44 | 23 | 32 | 43 | 9,69E+06 | 2,58E+07 | 1,51E+08 | 40 | 40 | 26 | 36 | 56 | Oxidation (M); Carbamidomethylation; Deamidation (NQ) | 149030 | \| PPTG_01093 \| Phytophthora parasitica INRA-310 hypothetical protein (1399 aa) |
| 46 | 61 | PPTG_03121 | 241,12 | 57 | 36 | 52 | 56 | 3,69E+07 | 2,37E+08 | 3,78E+08 | 36 | 36 | 31 | 52 | 64 | Oxidation (M); Carbamidomethylation; Deamidation (NQ) | 86664 | \| PPTG_03121 \| Phytophthora parasitica INRA-310 hsp70-like protein (787 aa) |
| 67 | 83 | PPTG_12681 | 240,96 | 42 | 26 | 24 | 42 | 1,03E+07 | 8,20E+06 | 1,86E+08 | 39 | 36 | 29 | 25 | 75 | Oxidation (M); Carbamidomethylation; Dehydration; Deamidation (NQ); Acetylation (N-term); Amidatic | 138952 | \| PPTG_12681 \| Phytophthora parasitica INRA-310 ATP-binding Cassette (ABC) superfamily (1291 aa) |
| 116 | 24 | PPTG_04750 | 240,21 | 80 | 52 | 64 | 80 | 8,15E+07 | 1,84E+08 | 9,87E+08 | 22 | 22 | 24 | 31 | 49 | Oxidation (M); Deamidation (NQ); Amidation | 46344 | \| PPTG_04750 \| Phytophthora parasitica INRA-310 mitochondrial-processing peptidase subunit alpha (448 aa) |
| 208 | 75 | PPTG_19658 | 238,19 | 73 | 47 | 61 | 66 | 1,90E+07 | 1,59E+08 | 1,77E+08 | 24 | 24 | 17 | 29 | 34 | Oxidation (M); Carbamidomethylation; Deamidation (NQ); Amidation | 46906 | \| PPTG_19658 \| Phytophthora parasitica INRA-310 argininosuccinate synthase (422 aa) |
| 208 | 76 | PPTG_02557 | 238,19 | 73 | 47 | 61 | 66 | 1,90E+07 | 1,59E+08 | 1,77E+08 | 24 | 24 | 17 | 29 | 34 | Oxidation (M); Carbamidomethylation; Deamidation (NQ); Amidation | 46906 | \| PPTG_02557 \| Phytophthora parasitica INRA-310 argininosuccinate synthase (422 aa) |
| 19 | 42 | PPTG_01676 | 236,92 | 70 | 69 | 68 | 68 | 3,76E+08 | 1,59E+09 | 3,91E+09 | 20 | 20 | 40 | 76 | 99 | Carbamidomethylation; Dehydration; Deamidation (NQ); Acetylation (N-term); Amidation; Carbamylat | 34949 | \| PPTG_01676 \| Phytophthora parasitica INRA-310 guanine nucleotide-binding protein subunit beta-like protein (318 aa) |
| 103 | 48 | PPTG_16803 | 236,48 | 61 | 35 | 44 | 58 | 2,09E+07 | 6,58E+07 | 2,52E+08 | 34 | 33 | 20 | 34 | 54 | Oxidation (M); Carbamidomethylation; Dehydration; Acetylation (N-term) | 68143 | \| PPTG_16803 \| Phytophthora parasitica INRA-310 V-type proton ATPase catalytic subunit A (619 aa) |
| 142 | 74 | PPTG_13258 | 236,43 | 25 | 10 | 9 | 24 | 4,44E+06 | 8,13E+06 | 8,29E+07 | 43 | 41 | 20 | 17 | 59 | Oxidation (M); Carbamidomethylation; Deamidation (NQ); Carbamylation | 256123 | \| PPTG_13258 \| Phytophthora parasitica INRA-310 callose synthase (2287 aa) |
| 377 | 96 | PPTG_09208 | 235,72 | 49 | 17 | 49 | 49 | 4,66E+06 | 8,05E+07 | 6,62E+07 | 15 | 15 | 9 | 24 | 21 | Oxidation (M); Carbamidomethylation; Dehydration; Deamidation (NQ) | 55628 | \| PPTG_09208 \| Phytophthora parasitica INRA-310 glycoside hydrolase (506 aa) |
| 21 | 36 | PPTG_04703 | 235,49 | 88 | 76 | 75 | 85 | 3,95E+08 | 7,29E+08 | 3,52E+09 | 27 | 27 | 53 | 57 | 99 | Oxidation (M); Carbamidomethylation; Dehydration; Deamidation (NQ); Acetylation (N-term); Carbam | 33753 | \| PPTG_04703 \| Phytophthora parasitica INRA-310 mitochondrial substrate carrier family protein ancA (311 aa) |
| 98 | 34 | PPTG_07368 | 234,9 | 43 | 27 | 39 | 41 | 1,26E+07 | 6,52E+07 | 2,05E+08 | 25 | 25 | 17 | 39 | 54 | Oxidation (M); Carbamidomethylation; Dehydration; Deamidation (NQ); Amidation | 80197 | \| PPTG_07368 \| Phytophthora parasitica INRA-310 trifunctional enzyme subunit alpha (748 aa) |
| 47 | 86 | PPTG_19188 | 233,36 | 51 | 25 | 46 | 40 | 2,48E+07 | 2,98E+08 | 3,59E+08 | 32 | 30 | 18 | 68 | 61 | Oxidation (M); Carbamidomethylation; Dehydration; Deamidation (NQ) | 80429 | \| PPTG_19188 \| Phytophthora parasitica INRA-310 heat shock protein 90-4 (707 aa) |
| 156 | 49 | PPTG_05941 | 233,12 | 38 | 20 | 22 | 38 | 8,18E+06 | 9,44E+06 | 1,53E+08 | 33 | 30 | 20 | 22 | 50 | Oxidation (M); Carbamidomethylation; Dehydration; Deamidation (NQ); Acetylation (N-term) | 117555 | \| PPTG_05941 \| Phytophthora parasitica INRA-310 sodium/potassium-transporting ATPase subunit alpha (1077 aa) |
| 370 | 25 | PPTG_14787 | 232,91 | 47 | 23 | 40 | 45 | 2,40E+06 | 1,92E+07 | 4,13E+07 | 19 | 19 | 8 | 21 | 26 | Oxidation (M); Carbamidomethylation; Deamidation (NQ) | 62832 | \| PPTG_14787 \| Phytophthora parasitica INRA-310 glycoside hydrolase (584 aa) |
| 194 | 84 | PPTG_01216 | 232,64 | 45 | 14 | 42 | 35 | 5,66E+06 | 1,04E+08 | 8,14E+07 | 29 | 29 | 10 | 42 | 31 | Oxidation (M); Carbamidomethylation; Deamidation (NQ); Carbamylation | 118224 | \| PPTG_01216 \| Phytophthora parasitica INRA-310 alpha-glucosidase (1092 aa) |
| 42 | 330 | PPTG_12239 | 231,76 | 22 | 5 | 20 | 15 | 2,07E+06 | 3,97E+07 | 3,68E+07 | 73 | 73 | 18 | 74 | 57 | Oxidation (M); Carbamidomethylation; Deamidation (NQ) | 483850 | \| PPTG_12239 \| Phytophthora parasitica INRA-310 hypothetical protein (4323 aa) |
| 57 | 60 | PPTG_17884 | 230,79 | 77 | 72 | 74 | 77 | 4,62E+05 | 3,29E+06 | 1,20E+08 | 22 | 3 | 36 | 38 | 65 | Oxidation (M); Dehydration; Deamidation (NQ); Acetylation (N-term); Carbamylation | 36179 | \| PPTG_17884 \| Phytophthora parasitica INRA-310 annexin (Annexin) family (329 aa) |
| 53 | 65 | PPTG_17883 | 230,35 | 70 | 55 | 58 | 70 | 4,07E+07 | 5,85E+07 | 3,14E+08 | 22 | 3 | 41 | 37 | 65 | Oxidation (M); Dehydration; Deamidation (NQ); Acetylation (N-term); Carbamylation | 39709 | \| PPTG_17883 \| Phytophthora parasitica INRA-310 annexin (Annexin) family (362 aa) |
| 101 | 112 | PPTG_16401 | 229,83 | 40 | 15 | 36 | 35 | 9,18E+06 | 9,66E+07 | 1,41E+08 | 41 | 41 | 15 | 49 | 44 | Oxidation (M); Carbamidomethylation; Deamidation (NQ); Carbamylation | 131605 | \| PPTG_16401 \| Phytophthora parasitica INRA-310 eukaryotic translation initiation factor 3 subunit A (1147 aa) |
| 26 | 44 | PPTG_02311 | 229,72 | 63 | 50 | 44 | 53 | 2,16E+08 | 3,57E+08 | 2,10E+09 | 23 | 23 | 41 | 46 | 94 | Oxidation (M); Carbamidomethylation; Dehydration; Deamidation (NQ); Acetylation (N-term); Carbam | 47905 | \| PPTG_02311 \| Phytophthora parasitica INRA-310 creatine kinase B-type (436 aa) |
| 143 | 189 | PPTG_10528 | 229,48 | 68 | 42 | 68 | 63 | 1,75E+07 | 2,45E+08 | 2,17E+08 | 16 | 16 | 15 | 44 | 37 | Oxidation (M); Carbamidomethylation; Dehydration; Deamidation (NQ) | 35462 | \| PPTG_10528 \| Phytophthora parasitica INRA-310 malate dehydrogenase mitochondrial (336 aa) |
| 80 | 32 | PPTG_15348 | 229,14 | 71 | 40 | 65 | 65 | 4,36E+07 | 2,14E+08 | 4,56E+08 | 21 | 15 | 20 | 46 | 54 | Oxidation (M); Carbamidomethylation; Deamidation (NQ); Acetylation (N-term); Carbamylation | 41867 | \| PPTG_15348 \| Phytophthora parasitica INRA-310 actin-1 (377 aa) |
| 140 | 55 | PPTG_06442 | 228,1 | 59 | 25 | 47 | 58 | 2,53E+07 | 6,58E+07 | 3,08E+08 | 27 | 27 | 16 | 31 | 49 | Oxidation (M); Carbamidomethylation; Dehydration; Deamidation (NQ); Acetylation (N-term) | 70910 | \| PPTG_06442 \| Phytophthora parasitica INRA-310 long-chain-fatty-acid-CoA ligase (647 aa) |
| 123 | 67 | PPTG_13419 | 227,93 | 48 | 22 | 44 | 42 | 1,12E+07 | 1,39E+08 | 1,38E+08 | 29 | 29 | 13 | 46 | 41 | Oxidation (M); Carbamidomethylation; Dehydration; Deamidation (NQ); Acetylation (N-term); Carbam | 84133 | \| PPTG_13419 \| Phytophthora parasitica INRA-310 aconitate hydratase mitochondrial (787 aa) |
| 69 | 71 | PPTG_08245 | 227,82 | 54 | 42 | 40 | 52 | 1,60E+07 | 4,85E+07 | 2,05E+08 | 30 | 27 | 30 | 38 | 60 | Oxidation (M); Carbamidomethylation; Dehydration; Deamidation (NQ) | 72698 | \| PPTG_08245 \| Phytophthora parasitica INRA-310 luminal-binding protein 5 (659 aa) |
| 137 | 89 | PPTG_18415 | 227,37 | 44 | 20 | 39 | 39 | 5,65E+06 | 5,31E+07 | 8,17E+07 | 34 | 34 | 14 | 41 | 42 | Oxidation (M); Carbamidomethylation; Deamidation (NQ) | 109217 | \| PPTG_18415 \| Phytophthora parasitica INRA-310 coatomer subunit beta (994 aa) |
| 38 | 326 | PPTG_12072 | 227,12 | 21 | 5 | 18 | 14 | 3,75E+06 | 3,91E+07 | 5,10E+07 | 80 | 80 | 21 | 74 | 62 | Oxidation (M); Carbamidomethylation; Deamidation (NQ) | 523990 | \| PPTG_12072 \| Phytophthora parasitica INRA-310 HECT E3 ubiquitin ligase (4811 aa) |
| 149 | 45 | PPTG_11762 | 227,1 | 62 | 44 | 57 | 62 | 2,91E+07 | 2,13E+08 | 3,46E+08 | 24 | 24 | 18 | 36 | 40 | Oxidation (M); Carbamidomethylation; Dehydration; Deamidation (NQ); Acetylation (N-term) | 53274 | \| PPTG_11762 \| Phytophthora parasitica INRA-310 6-phosphogluconate dehydrogenase (decarboxylating) (490 aa) |
| 13 | 62 | PPTG_20451 | 226,55 | 58 | 53 | 52 | 57 | 2,94E+08 | 6,29E+08 | 3,54E+09 | 30 | 30 | 57 | 70 | 142 | Oxidation (M); Carbamidomethylation; Dehydration; Deamidation (NQ); Acetylation (N-term); Carbam | 56147 | \| PPTG_20451 \| Phytophthora parasitica INRA-310 ATP synthase subunit alpha mitochondrial (510 aa) |
| 167 | 46 | PPTG_08188 | 226,48 | 61 | 30 | 58 | 59 | 1,47E+07 | 9,13E+07 | 1,50E+08 | 25 | 25 | 17 | 34 | 38 | Oxidation (M); Carbamidomethylation; Deamidation (NQ); Acetylation (N-term) | 65981 | \| PPTG_08188 \| Phytophthora parasitica INRA-310 phosphoribosylaminoimidazolecarboxamide formyltransferase/IMP cyclohydrolase (617 aa) |
| 61 | 133 | PPTG_13558 | 225,42 | 46 | 21 | 40 | 44 | 2,04E+07 | 1,38E+08 | 2,12E+08 | 36 | 36 | 22 | 51 | 61 | Oxidation (M); Carbamidomethylation; Dehydration; Deamidation (NQ) | 115732 | \| PPTG_13558 \| Phytophthora parasitica INRA-310 elongation factor 3 (1039 aa) |
| 127 | 88 | PPTG_04382 | 225,19 | 49 | 22 | 42 | 45 | 6,53E+06 | 5,07E+07 | 7,97E+07 | 33 | 33 | 17 | 42 | 41 | Oxidation (M); Carbamidomethylation; Dehydration; Deamidation (NQ); Acetylation (N-term) | 98645 | \| PPTG_04382 \| Phytophthora parasitica INRA-310 major vault protein (882 aa) |
| 222 | 143 | PPTG_05087 | 224,84 | 29 | 14 | 15 | 29 | 3,51E+06 | 8,49E+06 | 7,11E+07 | 35 | 35 | 17 | 20 | 41 | Oxidation (M); Carbamidomethylation; Deamidation (NQ) | 178979 | \| PPTG_05087 \| Phytophthora parasitica INRA-310 kinesin-like protein (1618 aa) |
| 147 | 81 | PPTG_01098 | 224,63 | 73 | 54 | 67 | 71 | 1,53E+05 | 1,69E+05 | 6,83E+05 | 18 | 2 | 20 | 36 | 39 | Oxidation (M); Carbamidomethylation; Deamidation (NQ); Acetylation (N-term); Amidation; Carbamyla | 36179 | \| PPTG_01098 \| Phytophthora parasitica INRA-310 glyceraldehyde-3-phosphate dehydrogenase (341 aa) |
| 223 | 125 | PPTG_09519 | 223,85 | 44 | 8 | 36 | 35 | 1,02E+06 | 3,46E+07 | 4,09E+07 | 37 | 37 | 6 | 36 | 36 | Oxidation (M); Carbamidomethylation | 120444 | \| PPTG_09519 \| Phytophthora parasitica INRA-310 ATP-citrate synthase (1106 aa) |
| 92 | 39 | PPTG_15414 | 223,82 | 80 | 61 | 64 | 80 | 2,98E+07 | 8,26E+07 | 5,29E+08 | 23 | 23 | 24 | 35 | 54 | Oxidation (M); Carbamidomethylation; Deamidation (NQ); Amidation | 36771 | \| PPTG_15414 \| Phytophthora parasitica INRA-310 trans-2-enoyl-CoA reductase (349 aa) |
| 66 | 208 | PPTG_01415 | 223,76 | 48 | 17 | 40 | 39 | 4,39E+06 | 5,05E+07 | 8,99E+07 | 49 | 49 | 21 | 49 | 59 | Oxidation (M); Carbamidomethylation; Deamidation (NQ) | 139621 | \| PPTG_01415 \| Phytophthora parasitica INRA-310 coatomer protein complex (1244 aa) |
| 170 | 37 | PPTG_12134 | 223,27 | 68 | 42 | 51 | 68 | 1,72E+07 | 4,92E+07 | 2,16E+08 | 23 | 23 | 17 | 23 | 49 | Oxidation (M); Carbamidomethylation; Dehydration; Deamidation (NQ); Carbamylation | 55147 | \| PPTG_12134 \| Phytophthora parasitica INRA-310 V-type proton ATPase subunit B isoform (496 aa) |
| 157 | 186 | PPTG_08926 | 222,79 | 41 | 21 | 36 | 38 | 6,95E+06 | 4,35E+07 | 7,46E+07 | 34 | 34 | 20 | 34 | 38 | Oxidation (M); Carbamidomethylation; Deamidation (NQ); Carbamylation | 124764 | \| PPTG_08926 \| Phytophthora parasitica INRA-310 DEAD/DEAH box RNA helicase (1095 aa) |
| 190 | 117 | PPTG_18707 | 222,51 | 41 | 17 | 35 | 38 | 5,26E+06 | 4,40E+07 | 6,60E+07 | 37 | 37 | 12 | 31 | 40 | Carbamidomethylation; Deamidation (NQ); Carbamylation | 111269 | \| PPTG_18707 \| Phytophthora parasitica INRA-310 alanyl-tRNA synthetase (1007 aa) |
| 276 | 103 | PPTG_12105 | 222,24 | 18 | 4 | 7 | 18 | 5,55E+05 | 2,25E+06 | 2,42E+07 | 39 | 39 | 6 | 16 | 45 | Deamidation (NQ) | 346604 | \| PPTG_12105 \| Phytophthora parasitica INRA-310 hypothetical protein (3662 aa) |
| 108 | 82 | PPTG_02435 | 222,21 | 71 | 42 | 67 | 71 | 7,62E+06 | 1,02E+08 | 7,08E+07 | 26 | 14 | 20 | 48 | 38 | Oxidation (M); Carbamidomethylation; Dehydration; Deamidation (NQ); Carbamylation | 56708 | \| PPTG_02435 \| Phytophthora parasitica INRA-310 aldehyde dehydrogenase (526 aa) |
| 81 | 215 | PPTG_06139 | 222,03 | 53 | 24 | 49 | 45 | 1,07E+07 | 1,15E+08 | 1,12E+08 | 38 | 34 | 21 | 52 | 46 | Oxidation (M); Carbamidomethylation; Dehydration; Deamidation (NQ); Acetylation (N-term) | 107365 | \| PPTG_06139 \| Phytophthora parasitica INRA-310 hypothetical protein (960 aa) |
| 97 | 142 | PPTG_17398 | 221,83 | 51 | 24 | 48 | 49 | 2,09E+07 | 1,17E+08 | 2,29E+08 | 37 | 37 | 18 | 44 | 49 | Oxidation (M); Carbamidomethylation; Deamidation (NQ); Acetylation (N-term) | 97557 | \| PPTG_17398 \| Phytophthora parasitica INRA-310 glutamyl-tRNA synthetase (877 aa) |
| 188 | 58 | PPTG_05924 | 221,54 | 41 | 19 | 22 | 40 | 9,77E+06 | 1,07E+07 | 1,51E+08 | 28 | 28 | 19 | 21 | 44 | Oxidation (M); Carbamidomethylation; Deamidation (NQ) | 99055 | \| PPTG_05924 \| Phytophthora parasitica INRA-310 vacuolar proton translocating ATPase A subunit (870 aa) |
| 110 | 212 | PPTG_11337 | 221,19 | 30 | 14 | 11 | 28 | 1,57E+07 | 2,28E+07 | 1,39E+08 | 46 | 40 | 25 | 22 | 58 | Oxidation (M); Carbamidomethylation; Deamidation (NQ); Acetylation (N-term) | 220005 | \| PPTG_11337 \| Phytophthora parasitica INRA-310 ATP-binding Cassette (ABC) superfamily (1984 aa) |
| 261 | 121 | PPTG_01995 | 221,18 | 76 | 52 | 70 | 76 | 3,01E+07 | 7,29E+07 | 4,55E+08 | 15 | 15 | 12 | 19 | 39 | Oxidation (M); Carbamidomethylation; Dehydration; Deamidation (NQ); Acetylation (N-term); Amidatic | 32907 | \| PPTG_01995 \| Phytophthora parasitica INRA-310 ATP synthase subunit gamma (305 aa) |
| 191 | 140 | PPTG_02282 | 220,97 | 38 | 17 | 13 | 36 | 5,05E+06 | 4,57E+06 | 7,61E+07 | 38 | 37 | 20 | 14 | 49 | Oxidation (M); Carbamidomethylation; Dehydration; Deamidation (NQ) | 158885 | \| PPTG_02282 \| Phytophthora parasitica INRA-310 phospholipid-transporting ATPase (1430 aa) |
| 130 | 106 | PPTG_12013 | 220,58 | 37 | 11 | 33 | 35 | 2,73E+06 | 3,34E+07 | 5,05E+07 | 35 | 33 | 13 | 38 | 48 | Oxidation (M); Carbamidomethylation; Dehydration; Deamidation (NQ) | 137465 | \| PPTG_12013 \| Phytophthora parasitica INRA-310 Argonaute3 (AGO3) (1269 aa) |
| 158 | 137 | PPTG_00791 | 220,51 | 40 | 18 | 38 | 35 | 3,32E+06 | 3,54E+07 | 4,38E+07 | 31 | 25 | 15 | 40 | 36 | Oxidation (M); Carbamidomethylation; Deamidation (NQ) | 113686 | \| PPTG_00791 \| Phytophthora parasitica INRA-310 metalloprotease family M01 (1009 aa) |
| 34 | 345 | PPTG_15526 | 219,43 | 57 | 47 | 51 | 54 | 2,07E+08 | 9,47E+08 | 2,46E+09 | 18 | 18 | 22 | 53 | 90 | Oxidation (M); Carbamidomethylation; Dehydration; Deamidation (NQ); Acetylation (N-term); Carbam | 34585 | \| PPTG_15526 \| Phytophthora parasitica INRA-310 60S ribosomal protein L5 (302 aa) |
| 169 | 234 | PPTG_13182 | 219,29 | 22 | 7 | 7 | 21 | 4,05E+06 | 8,11E+06 | 7,74E+07 | 45 | 43 | 15 | 15 | 59 | Oxidation (M); Carbamidomethylation; Deamidation (NQ) | 275558 | \| PPTG_13182 \| Phytophthora parasitica INRA-310 callose synthase (2448 aa) |
| 89 | 147 | PPTG_00110 | 219,22 | 60 | 34 | 52 | 52 | 4,08E+07 | 2,57E+08 | 4,06E+08 | 33 | 33 | 24 | 44 | 46 | Carbamidomethylation; Dehydration; Deamidation (NQ); Amidation; Carbamylation | 63275 | \| PPTG_00110 \| Phytophthora parasitica INRA-310 phospholipid hydroperoxide glutathione peroxidase (562 aa) |
| 445 | 353 | PPTG_09984 | 219,22 | 73 | 65 | 52 | 73 | 1,94E+07 | 2,63E+07 | 1,88E+08 | 14 | 14 | 12 | 14 | 20 | Oxidation (M); Carbamidomethylation; Dehydration; Deamidation (NQ) | 27248 | \| PPTG_09984 \| Phytophthora parasitica INRA-310 hypothetical protein (252 aa) |
| 290 | 207 | PPTG_04240 | 219,04 | 60 | 39 | 56 | 52 | 5,37E+06 | 7,72E+07 | 8,05E+07 | 21 | 21 | 9 | 29 | 28 | Oxidation (M); Carbamidomethylation | 41325 | \| PPTG_04240 \| Phytophthora parasitica INRA-310 3-ketoacyl-CoA thiolase mitochondrial (398 aa) |
| 105 | 316 | PPTG_14789 | 218,89 | 18 | 3 | 16 | 11 | 4,09E+05 | 1,98E+07 | 1,50E+07 | 63 | 61 | 9 | 58 | 39 | Oxidation (M); Carbamidomethylation; Dehydration | 467352 | \| PPTG_14789 \| Phytophthora parasitica INRA-310 dynein heavy chain (4161 aa) |

| | | | | | | | | | | | | | | | | | | | |
|---|---|---|---|---|---|---|---|---|---|---|---|---|---|---|---|---|---|---|---|
| 88 | 159 | PPTG_03763 | 218,89 | 57 | 23 | 49 | 53 | 7,39E+06 | 7,84E+07 | 1,05E+08 | 36 | 36 | 22 | 45 | 49 | Oxidation (M); Carbamidomethylation; Dehydration; Deamidation (NQ); Amidation | 89105 | PPTG_03763 | Phytophthora parasitica INRA-310 cell division control protein 48 like E (805 aa) |
| 145 | 102 | PPTG_10576 | 218,74 | 34 | 18 | 24 | 34 | 3,56E+06 | 4,07E+06 | 5,53E+07 | 29 | 16 | 21 | 27 | 47 | Oxidation (M); Carbamidomethylation; Deamidation (NQ); Acetylation (N-term) | 109796 | PPTG_10576 | Phytophthora parasitica INRA-310 ATP-binding Cassette (ABC) superfamily (985 aa) |
| 237 | 77 | PPTG_03165 | 218,68 | 64 | 23 | 50 | 62 | 2,96E+06 | 3,11E+07 | 5,71E+07 | 26 | 26 | 11 | 29 | 35 | Oxidation (M); Carbamidomethylation; Deamidation (NQ); Carbamylation | 60359 | PPTG_03165 | Phytophthora parasitica INRA-310 chaperonin mitochondrial (577 aa) |
| 203 | 128 | PPTG_14992 | 218,68 | 52 | 26 | 46 | 51 | 7,50E+06 | 5,15E+07 | 7,12E+07 | 30 | 30 | 16 | 31 | 34 | Oxidation (M); Carbamidomethylation; Deamidation (NQ) | 84760 | PPTG_14992 | Phytophthora parasitica INRA-310 tyrosyl-tRNA synthetase (766 aa) |
| 172 | 183 | PPTG_06379 | 218,56 | 43 | 18 | 40 | 37 | 3,90E+06 | 3,66E+07 | 6,01E+07 | 35 | 23 | 17 | 36 | 35 | Oxidation (M); Carbamidomethylation; Dehydration | 118081 | PPTG_06379 | Phytophthora parasitica INRA-310 NAD-specific glutamate dehydrogenase (1058 aa) |
| 27 | 101 | PPTG_13414 | 218,49 | 61 | 39 | 53 | 61 | 1,64E+08 | 6,88E+08 | 2,27E+09 | 25 | 25 | 29 | 58 | 91 | Oxidation (M); Carbamidomethylation; Dehydration; Deamidation (NQ); Carbamylation | 44403 | PPTG_13414 | Phytophthora parasitica INRA-310 60S ribosomal protein L3 (390 aa) |
| 297 | 93 | PPTG_06268 | 218,25 | 35 | 13 | 24 | 34 | 8,20E+06 | 4,34E+07 | 1,84E+08 | 24 | 24 | 9 | 22 | 33 | Oxidation (M); Carbamidomethylation; Dehydration; Deamidation (NQ); Amidation | 89977 | PPTG_06268 | Phytophthora parasitica INRA-310 choline transporter-like protein (812 aa) |
| 63 | 78 | PPTG_15113 | 218,06 | 64 | 47 | 57 | 59 | 4,74E+07 | 3,21E+08 | 5,15E+08 | 21 | 21 | 22 | 54 | 58 | Oxidation (M); Carbamidomethylation; Deamidation (NQ) | 50113 | PPTG_15113 | Phytophthora parasitica INRA-310 tubulin beta chain (447 aa) |
| 121 | 738 | PPTG_01682 | 217,92 | 78 | 78 | 78 | 78 | 8,38E+07 | 1,59E+08 | 9,41E+08 | 16 | 16 | 23 | 23 | 56 | Oxidation (M); Deamidation (NQ); Carbamylation | 21006 | PPTG_01682 | Phytophthora parasitica INRA-310 hypothetical protein (184 aa) |
| 311 | 53 | PPTG_00714 | 217,72 | 30 | 15 | 21 | 29 | 1,92E+07 | 4,06E+07 | 2,32E+08 | 20 | 20 | 11 | 18 | 33 | Oxidation (M); Carbamidomethylation; Dehydration; Deamidation (NQ); Acetylation (N-term) | 95841 | PPTG_00714 | Phytophthora parasitica INRA-310 hypothetical protein (892 aa) |
| 111 | 165 | PPTG_09527 | 217,72 | 41 | 22 | 35 | 37 | 7,25E+06 | 8,64E+07 | 9,20E+07 | 31 | 31 | 19 | 43 | 43 | Oxidation (M); Carbamidomethylation; Dehydration; Deamidation (NQ) | 108851 | PPTG_09527 | Phytophthora parasitica INRA-310 26S proteasome non-ATPase regulatory subunit (998 aa) |
| 306 | 56 | PPTG_19590 | 217,31 | 62 | 29 | 50 | 60 | 9,34E+06 | 7,77E+07 | 1,05E+08 | 19 | 19 | 9 | 25 | 29 | Oxidation (M); Carbamidomethylation; Dehydration; Deamidation (NQ); Acetylation (N-term) | 50034 | PPTG_19590 | Phytophthora parasitica INRA-310 calreticulin (443 aa) |
| 225 | 66 | PPTG_08479 | 217,19 | 62 | 20 | 55 | 58 | 3,92E+06 | 4,46E+07 | 4,40E+07 | 21 | 21 | 11 | 31 | 36 | Oxidation (M); Carbamidomethylation; Dehydration; Deamidation (NQ) | 61217 | PPTG_08479 | Phytophthora parasitica INRA-310 methylcrotonoyl-CoA carboxylase beta chain mitochondrial (568 aa) |
| 96 | 155 | PPTG_12389 | 216,78 | 49 | 29 | 43 | 46 | 1,17E+07 | 5,09E+07 | 1,61E+08 | 34 | 34 | 22 | 37 | 53 | Oxidation (M); Carbamidomethylation; Deamidation (NQ) | 101356 | PPTG_12389 | Phytophthora parasitica INRA-310 nuclease (920 aa) |
| 262 | 69 | PPTG_14642 | 216,13 | 41 | 27 | 28 | 41 | 7,38E+06 | 9,56E+06 | 1,04E+08 | 24 | 24 | 16 | 16 | 38 | Oxidation (M); Carbamidomethylation; Deamidation (NQ) | 75352 | PPTG_14642 | Phytophthora parasitica INRA-310 glutamine-fructose-6-phosphate transaminase (isomerizing) (702 aa) |
| 193 | 104 | PPTG_02473 | 215,97 | 68 | 33 | 62 | 53 | 2,11E+06 | 4,13E+07 | 3,51E+07 | 24 | 13 | 16 | 37 | 30 | Oxidation (M); Carbamidomethylation; Dehydration; Deamidation (NQ); Carbamylation | 56698 | PPTG_02473 | Phytophthora parasitica INRA-310 aldehyde dehydrogenase (526 aa) |
| 20 | 109 | PPTG_16532 | 215,96 | 61 | 38 | 55 | 61 | 3,43E+08 | 1,72E+09 | 4,26E+09 | 20 | 20 | 35 | 69 | 105 | Oxidation (M); Carbamidomethylation; Dehydration; Deamidation (NQ); Acetylation (N-term); 2 more | 42656 | PPTG_16532 | Phytophthora parasitica INRA-310 60S ribosomal protein L4 (388 aa) |
| 187 | 57 | PPTG_00859 | 215,93 | 53 | 46 | 49 | 49 | 3,22E+07 | 8,63E+07 | 5,64E+08 | 17 | 17 | 18 | 24 | 42 | Oxidation (M); Dehydration; Deamidation (NQ); Amidation | 63622 | PPTG_00859 | Phytophthora parasitica INRA-310 hypothetical protein (605 aa) |
| 182 | 161 | PPTG_08551 | 215,69 | 61 | 48 | 52 | 57 | 1,70E+07 | 1,11E+08 | 1,43E+08 | 30 | 30 | 18 | 34 | 34 | Oxidation (M); Carbamidomethylation; Deamidation (NQ); Acetylation (N-term) | 59018 | PPTG_08551 | Phytophthora parasitica INRA-310 26S proteasome non-ATPase regulatory subunit 3 (520 aa) |
| 85 | 145 | PPTG_12096 | 215,57 | 30 | 15 | 26 | 27 | 7,10E+06 | 4,98E+07 | 1,06E+08 | 35 | 35 | 19 | 46 | 52 | Oxidation (M); Carbamidomethylation; Deamidation (NQ) | 164314 | PPTG_12096 | Phytophthora parasitica INRA-310 myosin-like protein (1475 aa) |
| 183 | 70 | PPTG_01939 | 215,53 | 32 | 20 | 31 | 31 | 3,08E+07 | 2,20E+08 | 2,73E+08 | 16 | 15 | 18 | 37 | 31 | Oxidation (M); Carbamidomethylation; Deamidation (NQ) | 83275 | PPTG_01939 | Phytophthora parasitica INRA-310 glucan 1 3-beta-glucosidase (751 aa) |
| 216 | 64 | PPTG_10580 | 215,24 | 52 | 28 | 34 | 52 | 6,61E+06 | 9,48E+06 | 1,15E+08 | 20 | 14 | 13 | 19 | 48 | Oxidation (M); Carbamidomethylation; Dehydration; Deamidation (NQ) | 53706 | PPTG_10580 | Phytophthora parasitica INRA-310 ATP-binding Cassette (ABC) superfamily (484 aa) |
| 151 | 148 | PPTG_04350 | 215,23 | 67 | 35 | 67 | 64 | 1,39E+07 | 1,71E+08 | 1,39E+08 | 26 | 25 | 17 | 41 | 36 | Oxidation (M); Carbamidomethylation; Dehydration; Deamidation (NQ) | 47788 | PPTG_04350 | Phytophthora parasitica INRA-310 isocitrate dehydrogenase [NADP] mitochondrial (428 aa) |
| 186 | 105 | PPTG_00228 | 215,22 | 53 | 29 | 51 | 50 | 5,43E+06 | 6,36E+07 | 7,80E+07 | 25 | 25 | 15 | 34 | 35 | Oxidation (M); Carbamidomethylation; Dehydration; Deamidation (NQ) | 66432 | PPTG_00228 | Phytophthora parasitica INRA-310 hypothetical protein (613 aa) |
| 252 | 100 | PPTG_01205 | 215,2 | 33 | 20 | 19 | 33 | 9,97E+06 | 1,91E+07 | 8,08E+07 | 31 | 31 | 18 | 17 | 37 | Oxidation (M); Carbamidomethylation; Dehydration; Deamidation (NQ) | 124086 | PPTG_01205 | Phytophthora parasitica INRA-310 myosin-like protein (1080 aa) |
| 213 | 87 | PPTG_15388 | 214,98 | 65 | 33 | 54 | 61 | 8,64E+06 | 7,91E+07 | 9,90E+07 | 22 | 22 | 15 | 32 | 33 | Oxidation (M); Carbamidomethylation; Deamidation (NQ) | 54290 | PPTG_15388 | Phytophthora parasitica INRA-310 methylmalonate-semialdehyde dehydrogenase (acylating) (506 aa) |
| 201 | 115 | PPTG_15933 | 214,88 | 33 | 14 | 26 | 33 | 5,87E+06 | 3,58E+07 | 6,90E+07 | 27 | 27 | 14 | 31 | 36 | Oxidation (M); Carbamidomethylation; Acetylation (N-term) | 118837 | PPTG_15933 | Phytophthora parasitica INRA-310 valyl-tRNA synthetase (1053 aa) |
| 260 | 157 | PPTG_14406 | 214,11 | 42 | 20 | 20 | 40 | 4,61E+06 | 6,03E+06 | 4,76E+07 | 32 | 32 | 17 | 15 | 38 | Oxidation (M); Carbamidomethylation; Deamidation (NQ) | 115102 | PPTG_14406 | Phytophthora parasitica INRA-310 Pro-apoptotic serine protease nma111-like protein (1051 aa) |
| 305 | 63 | PPTG_00150 | 214,07 | 46 | 20 | 34 | 45 | 7,19E+06 | 2,45E+07 | 1,45E+08 | 22 | 22 | 9 | 17 | 37 | Oxidation (M); Carbamidomethylation; Deamidation (NQ) | 70415 | PPTG_00150 | Phytophthora parasitica INRA-310 succinate dehydrogenase [ubiquinone] flavoprotein subunit mitochondrial (646 aa) |
| 106 | 213 | PPTG_10774 | 213,3 | 44 | 21 | 43 | 39 | 1,82E+07 | 6,96E+07 | 9,42E+07 | 37 | 37 | 18 | 44 | 44 | Oxidation (M); Carbamidomethylation; Deamidation (NQ) | 109186 | PPTG_10774 | Phytophthora parasitica INRA-310 eukaryotic translation initiation factor 3 subunit C (974 aa) |
| 283 | 97 | PPTG_11901 | 212,82 | 58 | 29 | 58 | 45 | 8,85E+06 | 9,45E+07 | 1,42E+08 | 18 | 18 | 12 | 28 | 27 | Oxidation (M); Deamidation (NQ); Carbamylation | 51488 | PPTG_11901 | Phytophthora parasitica INRA-310 citrate synthase (465 aa) |
| 87 | 181 | PPTG_10291 | 212,81 | 46 | 26 | 45 | 44 | 1,47E+07 | 1,43E+08 | 2,45E+08 | 34 | 34 | 20 | 47 | 50 | Oxidation (M); Carbamidomethylation; Dehydration; Deamidation (NQ) | 84973 | PPTG_10291 | Phytophthora parasitica INRA-310 5-methlytetrahydropteroyltriglutamate-homocysteine methyltransferease (768 aa) |
| 70 | 95 | PPTG_20478 | 212,69 | 48 | 36 | 38 | 48 | 9,97E+07 | 1,66E+08 | 8,21E+08 | 22 | 19 | 28 | 33 | 67 | Oxidation (M); Carbamidomethylation; Dehydration; Deamidation (NQ); Carbamylation | 47349 | PPTG_20478 | Phytophthora parasitica INRA-310 annexin (Annexin) family (451 aa) |
| 339 | 73 | PPTG_18730 | 212,59 | 30 | 11 | 8 | 29 | 1,43E+06 | 3,21E+05 | 1,72E+07 | 26 | 17 | 12 | 9 | 37 | Oxidation (M); Carbamidomethylation; Dehydration; Deamidation (NQ); Acetylation (N-term) | 116630 | PPTG_18730 | Phytophthora parasitica INRA-310 P-type ATPase (P-ATPase) superfamily (1060 aa) |
| 43 | 90 | PPTG_11356 | 212,59 | 81 | 68 | 68 | 81 | 1,13E+08 | 5,81E+08 | 1,61E+09 | 19 | 19 | 31 | 42 | 77 | Oxidation (M); Carbamidomethylation; Deamidation (NQ); Acetylation (N-term); Carbamylation | 34408 | PPTG_11356 | Phytophthora parasitica INRA-310 60S acidic ribosomal protein P0 (318 aa) |
| 268 | 263 | PPTG_03117 | 212,29 | 49 | 28 | 49 | 43 | 1,42E+07 | 2,15E+08 | 1,26E+08 | 14 | 11 | 12 | 29 | 28 | Oxidation (M); Carbamidomethylation; Deamidation (NQ); Amidation | 50779 | PPTG_03117 | Phytophthora parasitica INRA-310 serine protease family S08A (487 aa) |
| 163 | 129 | PPTG_13885 | 211 | 54 | 36 | 53 | 48 | 9,47E+06 | 8,22E+07 | 1,13E+08 | 26 | 26 | 17 | 39 | 34 | Oxidation (M); Carbamidomethylation; Acetylation (N-term) | 68832 | PPTG_13885 | Phytophthora parasitica INRA-310 aminotransferase (634 aa) |
| 52 | 91 | PPTG_10972 | 210,97 | 70 | 46 | 48 | 64 | 5,46E+07 | 1,10E+08 | 7,19E+08 | 26 | 26 | 27 | 34 | 82 | Oxidation (M); Carbamidomethylation; Dehydration; Deamidation (NQ); Amidation; Carbamylation | 51893 | PPTG_10972 | Phytophthora parasitica INRA-310 mitochondrial-processing peptidase subunit beta (467 aa) |
| 206 | 120 | PPTG_11759 | 210,83 | 38 | 15 | 22 | 36 | 6,61E+06 | 1,59E+07 | 8,35E+07 | 28 | 28 | 14 | 21 | 46 | Oxidation (M); Carbamidomethylation; Dehydration; Deamidation (NQ) | 114975 | PPTG_11759 | Phytophthora parasitica INRA-310 lysosomal alpha-mannosidase (1026 aa) |
| 211 | 126 | PPTG_17902 | 210,81 | 30 | 16 | 18 | 29 | 1,70E+07 | 2,20E+07 | 2,58E+08 | 28 | 28 | 16 | 19 | 45 | Oxidation (M); Carbamidomethylation; Deamidation (NQ); Amidation | 126885 | PPTG_17902 | Phytophthora parasitica INRA-310 cellulose synthase catalytic subunit [UDP-forming] (1143 aa) |
| 316 | 124 | PPTG_08190 | 210,62 | 37 | 16 | 19 | 35 | 6,91E+05 | 1,03E+06 | 1,07E+07 | 28 | 12 | 15 | 15 | 31 | Carbamidomethylation; Dehydration | 116581 | PPTG_08190 | Phytophthora parasitica INRA-310 phosphatidyl inositol kinase (PIK-B) (1037 aa) |
| 184 | 204 | PPTG_16495 | 210,53 | 38 | 12 | 36 | 36 | 4,03E+06 | 4,61E+07 | 5,76E+07 | 31 | 31 | 12 | 38 | 35 | Oxidation (M); Carbamidomethylation; Dehydration; Amidation; Carbamylation | 126115 | PPTG_16495 | Phytophthora parasitica INRA-310 E1 family ubiquitin activating enzyme (1125 aa) |
| 144 | 251 | PPTG_18870 | 210,31 | 62 | 45 | 49 | 62 | 8,06E+07 | 1,66E+08 | 1,05E+09 | 16 | 16 | 20 | 22 | 54 | Oxidation (M); Carbamidomethylation; Dehydration; Deamidation (NQ); Amidation; Carbamylation | 33936 | PPTG_18870 | Phytophthora parasitica INRA-310 hypothetical protein (309 aa) |
| 135 | 167 | PPTG_08622 | 210,16 | 66 | 35 | 64 | 60 | 1,82E+07 | 1,53E+08 | 2,40E+08 | 27 | 27 | 15 | 41 | 42 | Oxidation (M); Carbamidomethylation; Dehydration; Deamidation (NQ) | 51245 | PPTG_08622 | Phytophthora parasitica INRA-310 serine hydroxymethyltransferase (465 aa) |
| 351 | 1034 | PPTG_01466 | 210,13 | 67 | 60 | 66 | 67 | 3,06E+07 | 1,99E+08 | 3,02E+08 | 10 | 10 | 12 | 21 | 24 | Oxidation (M); Carbamidomethylation; Dehydration; Deamidation (NQ); Acetylation (N-term); Carbam | 17787 | PPTG_01466 | Phytophthora parasitica INRA-310 translation initiation factor eIF-5A (162 aa) |
| 196 | 182 | PPTG_18309 | 210,04 | 55 | 39 | 46 | 55 | 1,74E+07 | 8,65E+07 | 1,65E+08 | 25 | 25 | 19 | 27 | 37 | Oxidation (M); Deamidation (NQ) | 57884 | PPTG_18309 | Phytophthora parasitica INRA-310 protein disulfide-isomerase (522 aa) |
| 224 | 221 | PPTG_08986 | 210,01 | 21 | 7 | 11 | 21 | 4,28E+06 | 8,23E+06 | 4,32E+07 | 34 | 16 | 17 | 20 | 41 | Oxidation (M); Carbamidomethylation; Deamidation (NQ) | 213247 | PPTG_08986 | Phytophthora parasitica INRA-310 ATP-binding Cassette (ABC) superfamily (1946 aa) |
| 241 | 98 | PPTG_08453 | 209,97 | 50 | 24 | 48 | 45 | 2,13E+05 | 2,78E+06 | 5,48E+06 | 17 | 2 | 14 | 26 | 35 | Oxidation (M); Carbamidomethylation; Dehydration; Deamidation (NQ) | 50263 | PPTG_08453 | Phytophthora parasitica INRA-310 tubulin alpha chain (455 aa) |
| 214 | 99 | PPTG_10444 | 209,94 | 55 | 38 | 50 | 49 | 1,51E+07 | 1,14E+08 | 1,77E+08 | 21 | 21 | 20 | 30 | 30 | Oxidation (M); Carbamidomethylation; Dehydration; Deamidation (NQ) | 61035 | PPTG_10444 | Phytophthora parasitica INRA-310 phospholipase D Pi-PLD-like-1 (531 aa) |
| 343 | 434 | PPTG_01189 | 209,88 | 40 | 24 | 29 | 40 | 3,10E+07 | 1,15E+08 | 5,46E+08 | 13 | 13 | 11 | 15 | 32 | Oxidation (M); Carbamidomethylation; Dehydration; Deamidation (NQ); Carbamylation | 48900 | PPTG_01189 | Phytophthora parasitica INRA-310 hypothetical protein (458 aa) |
| 218 | 164 | PPTG_17298 | 209,2 | 48 | 13 | 42 | 48 | 3,61E+06 | 3,71E+07 | 5,36E+07 | 25 | 25 | 8 | 34 | 37 | Oxidation (M); Carbamidomethylation; Dehydration; Deamidation (NQ) | 75448 | PPTG_17298 | Phytophthora parasitica INRA-310 aconitate hydratase (701 aa) |
| 293 | 119 | PPTG_15137 | 209,12 | 46 | 19 | 35 | 41 | 2,98E+06 | 2,30E+07 | 3,94E+07 | 26 | 26 | 11 | 26 | 28 | Oxidation (M); Carbamidomethylation; Deamidation (NQ); Acetylation (N-term) | 78382 | PPTG_15137 | Phytophthora parasitica INRA-310 dynamin-2 (706 aa) |
| 159 | 141 | PPTG_02874 | 208,72 | 62 | 45 | 58 | 59 | 8,40E+06 | 7,61E+07 | 9,32E+07 | 21 | 21 | 17 | 37 | 37 | Oxidation (M); Carbamidomethylation; Dehydration; Deamidation (NQ) | 56242 | PPTG_02874 | Phytophthora parasitica INRA-310 glycerol kinase 1 (513 aa) |
| 195 | 169 | PPTG_01022 | 208,62 | 71 | 25 | 62 | 62 | 1,40E+07 | 1,53E+08 | 1,64E+08 | 27 | 27 | 12 | 38 | 33 | Oxidation (M); Carbamidomethylation; Deamidation (NQ) | 60800 | PPTG_01022 | Phytophthora parasitica INRA-310 glucose-6-phosphate isomerase cytosolic 1 (557 aa) |
| 598 | 107 | PPTG_15898 | 207,7 | 26 | 9 | 5 | 25 | 8,82E+05 | 5,34E+05 | 2,39E+07 | 22 | 22 | 7 | 6 | 23 | Carbamidomethylation; Deamidation (NQ); Acetylation (N-term) | 128994 | PPTG_15898 | Phytophthora parasitica INRA-310 hypothetical protein (1175 aa) |
| 598 | 108 | PPTG_15898 | 207,7 | 26 | 8 | 5 | 24 | 8,82E+05 | 5,34E+05 | 2,39E+07 | 22 | 22 | 7 | 6 | 23 | Carbamidomethylation; Deamidation (NQ); Acetylation (N-term) | 129645 | PPTG_15898 | Phytophthora parasitica INRA-310 hypothetical protein (1181 aa) |
| 90 | 79 | PPTG_07665 | 207,54 | 54 | 38 | 52 | 43 | 2,68E+07 | 4,39E+08 | 4,02E+08 | 21 | 19 | 20 | 54 | 40 | Oxidation (M); Deamidation (NQ); Acetylation (N-term); Amidation; Carbamylation | 60105 | PPTG_07665 | Phytophthora parasitica INRA-310 glucosylceramidase (541 aa) |
| 32 | 572 | PPTG_11031 | 207,34 | 59 | 48 | 40 | 59 | 8,25E+07 | 4,93E+08 | 1,53E+09 | 11 | 11 | 21 | 37 | 112 | Oxidation (M); Carbamidomethylation; Dehydration; Deamidation (NQ); Carbamylation | 33428 | PPTG_11031 | Phytophthora parasitica INRA-310 hypothetical protein (326 aa) |
| 323 | 237 | PPTG_01692 | 207,27 | 72 | 42 | 54 | 72 | 1,44E+07 | 3,43E+07 | 1,76E+08 | 15 | 15 | 9 | 17 | 35 | Oxidation (M); Carbamidomethylation; Dehydration; Acetylation (N-term) | 32939 | PPTG_01692 | Phytophthora parasitica INRA-310 mitochondrial carnitine/acylcarnitine carrier protein (311 aa) |
| 112 | 630 | PPTG_11052 | 207,08 | 34 | 31 | 32 | 34 | 2,10E+08 | 6,93E+08 | 1,73E+09 | 11 | 11 | 25 | 35 | 45 | Oxidation (M); Carbamidomethylation; Dehydration; Deamidation (NQ); Carbamylation | 48587 | PPTG_11052 | Phytophthora parasitica INRA-310 elongation factor 1-alpha (444 aa) |
| 112 | 631 | PPTG_11158 | 207,08 | 34 | 31 | 32 | 34 | 2,10E+08 | 6,93E+08 | 1,73E+09 | 11 | 11 | 25 | 35 | 45 | Oxidation (M); Carbamidomethylation; Dehydration; Deamidation (NQ); Carbamylation | 48587 | PPTG_11158 | Phytophthora parasitica INRA-310 elongation factor 1-alpha (444 aa) |
| 227 | 132 | PPTG_08465 | 207,01 | 51 | 27 | 49 | 45 | 3,32E+06 | 8,40E+06 | 1,94E+07 | 17 | 1 | 15 | 27 | 36 | Oxidation (M); Carbamidomethylation; Dehydration; Deamidation (NQ) | 49919 | PPTG_08465 | Phytophthora parasitica INRA-310 tubulin alpha chain (453 aa) |
| 131 | 136 | PPTG_10588 | 206,91 | 25 | 13 | 11 | 25 | 1,05E+06 | 1,75E+05 | 1,14E+07 | 27 | 11 | 19 | 20 | 60 | Oxidation (M); Carbamidomethylation; Deamidation (NQ) | 151906 | PPTG_10588 | Phytophthora parasitica INRA-310 ATP-binding Cassette (ABC) superfamily (1352 aa) |
| 232 | 94 | PPTG_04978 | 206,69 | 64 | 24 | 53 | 60 | 6,67E+06 | 5,55E+07 | 1,38E+08 | 24 | 24 | 8 | 29 | 39 | Oxidation (M); Carbamidomethylation; Deamidation (NQ) | 51657 | PPTG_04978 | Phytophthora parasitica INRA-310 rab GDP dissociation inhibitor alpha (463 aa) |
| 313 | 110 | PPTG_13413 | 206,39 | 63 | 52 | 30 | 58 | 1,01E+07 | 4,60E+06 | 7,36E+07 | 25 | 25 | 19 | 11 | 32 | Oxidation (M); Carbamidomethylation; Deamidation (NQ) | 52472 | PPTG_13413 | Phytophthora parasitica INRA-310 dolichyl-diphosphooligosaccharide-protein glycosyltransferase subunit (471 aa) |
| 238 | 154 | PPTG_01563 | 206,19 | 46 | 18 | 45 | 46 | 3,64E+06 | 6,67E+07 | 8,04E+07 | 24 | 24 | 11 | 29 | 35 | Oxidation (M); Carbamidomethylation; Deamidation (NQ) | 69507 | PPTG_01563 | Phytophthora parasitica INRA-310 polyadenylate-binding protein 1-B (641 aa) |
| 132 | 130 | PPTG_15102 | 204,81 | 55 | 24 | 50 | 53 | 1,82E+07 | 1,15E+08 | 2,54E+08 | 25 | 25 | 16 | 39 | 44 | Oxidation (M); Carbamidomethylation; Deamidation (NQ); Carbamylation | 62153 | PPTG_15102 | Phytophthora parasitica INRA-310 glucose-6-phosphate dehydrogenase (553 aa) |
| 154 | 396 | PPTG_09470 | 204,64 | 21 | 7 | 19 | 17 | 2,65E+06 | 2,39E+07 | 3,08E+07 | 43 | 43 | 14 | 42 | 37 | Oxidation (M); Carbamidomethylation; Dehydration | 259011 | PPTG_09470 | Phytophthora parasitica INRA-310 acetyl-CoA carboxylase (2327 aa) |
| 363 | 158 | PPTG_14740 | 204,39 | 18 | 7 | 5 | 16 | 2,43E+06 | 1,35E+06 | 3,68E+07 | 30 | 22 | 11 | 8 | 35 | Oxidation (M); Carbamidomethylation; Deamidation (NQ) | 233455 | PPTG_14740 | Phytophthora parasitica INRA-310 callose synthase (2041 aa) |
| 134 | 307 | PPTG_11194 | 203,95 | 19 | 7 | 16 | 15 | 4,96E+06 | 3,06E+07 | 5,24E+07 | 43 | 43 | 18 | 43 | 37 | Deamidation (NQ) | 276413 | PPTG_11194 | Phytophthora parasitica INRA-310 hypothetical protein (2479 aa) |
| 486 | 179 | PPTG_17691 | 203,88 | 36 | 15 | 14 | 36 | 4,36E+06 | 9,50E+06 | 5,35E+07 | 19 | 18 | 7 | 8 | 28 | Oxidation (M); Carbamidomethylation; Deamidation (NQ) | 85970 | PPTG_17691 | Phytophthora parasitica INRA-310 plasma-membrane proton-efflux P-type ATPase (794 aa) |
| 292 | 178 | PPTG_03625 | 203,87 | 35 | 15 | 18 | 33 | 7,25E+06 | 9,03E+06 | 8,45E+07 | 27 | 27 | 13 | 15 | 37 | Oxidation (M); Carbamidomethylation; Dehydration; Deamidation (NQ); Acetylation (N-term) | 114640 | PPTG_03625 | Phytophthora parasitica INRA-310 calcium-transporting ATPase 1 endoplasmic reticulum-type (1046 aa) |
| 114 | 227 | PPTG_02578 | 203,84 | 48 | 14 | 45 | 44 | 4,90E+06 | 3,66E+07 | 6,20E+07 | 33 | 33 | 13 | 42 | 49 | Oxidation (M); Carbamidomethylation; Dehydration; Deamidation (NQ) | 90183 | PPTG_02578 | Phytophthora parasitica INRA-310 arginyl-tRNA synthetase (816 aa) |
| 286 | 116 | PPTG_09888 | 203,74 | 62 | 23 | 62 | 57 | 2,17E+06 | 8,01E+07 | 5,69E+07 | 20 | 19 | 8 | 31 | 27 | Oxidation (M); Carbamidomethylation; Deamidation (NQ) | 48725 | PPTG_09888 | Phytophthora parasitica INRA-310 enolase (458 aa) |
| 287 | 173 | PPTG_14635 | 203,71 | 50 | 25 | 41 | 50 | 4,50E+06 | 6,33E+07 | 7,90E+07 | 18 | 18 | 10 | 27 | 29 | Oxidation (M); Carbamidomethylation; Deamidation (NQ) | 50958 | PPTG_14635 | Phytophthora parasitica INRA-310 aspartyl aminopeptidase (463 aa) |
| 210 | 210 | PPTG_00328 | 203,57 | 35 | 17 | 27 | 29 | 8,36E+06 | 4,14E+07 | 8,74E+07 | 28 | 28 | 14 | 31 | 35 | Oxidation (M); Carbamidomethylation; Dehydration; Deamidation (NQ) | 100907 | PPTG_00328 | Phytophthora parasitica INRA-310 AP-2 complex subunit beta (916 aa) |
| 199 | 123 | PPTG_01949 | 203,49 | 49 | 22 | 49 | 48 | 1,02E+07 | 1,13E+08 | 1,45E+08 | 20 | 20 | 9 | 37 | 36 | Oxidation (M); Carbamidomethylation; Dehydration; Deamidation (NQ) | 52624 | PPTG_01949 | Phytophthora parasitica INRA-310 adenosylhomocysteinase (482 aa) |
| 347 | 275 | PPTG_11937 | 203,41 | 27 | 5 | 7 | 26 | 1,05E+06 | 2,41E+06 | 3,49E+07 | 35 | 35 | 8 | 12 | 37 | Oxidation (M); Carbamidomethylation; Deamidation (NQ); Amidation | 204662 | PPTG_11937 | Phytophthora parasitica INRA-310 dysferlin-like protein (1835 aa) |
| 162 | 172 | PPTG_14774 | 203,06 | 54 | 31 | 52 | 52 | 1,13E+05 | 3,07E+06 | 6,75E+06 | 25 | 1 | 17 | 37 | 36 | Oxidation (M); Carbamidomethylation; Dehydration; Deamidation (NQ); Acetylation (N-term) | 59563 | PPTG_14774 | Phytophthora parasitica INRA-310 malate synthase A (536 aa) |
| 197 | 122 | PPTG_18274 | 202,99 | 43 | 20 | 42 | 41 | 3,50E+06 | 7,17E+07 | 7,09E+07 | 22 | 22 | 11 | 40 | 31 | Oxidation (M); Dehydration; Deamidation (NQ) | 69593 | PPTG_18274 | Phytophthora parasitica INRA-310 chaperone dnaK (649 aa) |
| 181 | 171 | PPTG_20398 | 202,78 | 54 | 29 | 52 | 50 | | 1,61E+06 | | 25 | 1 | 16 | 37 | 33 | Oxidation (M); Carbamidomethylation; Dehydration; Deamidation (NQ); Acetylation (N-term) | 59593 | PPTG_20398 | Phytophthora parasitica INRA-310 malate synthase A (536 aa) |
| 174 | 135 | PPTG_15943 | 202,64 | 42 | 26 | 40 | 39 | 1,41E+07 | 1,38E+08 | 1,58E+08 | 19 | 19 | 17 | 36 | 35 | Oxidation (M); Carbamidomethylation; Dehydration; Deamidation (NQ); Carbamylation | 56487 | PPTG_15943 | Phytophthora parasitica INRA-310 ketol-acid reductoisomerase (516 aa) |
| 524 | 440 | PPTG_04558 | 202,49 | 35 | 24 | 30 | 35 | 4,12E+06 | 2,10E+07 | 5,64E+07 | 12 | 12 | 9 | 16 | 16 | Oxidation (M); Carbamidomethylation; Deamidation (NQ) | 58775 | PPTG_04558 | Phytophthora parasitica INRA-310 5A endo-1 4-betaglucanase (543 aa) |
| 28 | 334 | PPTG_14734 | 202,3 | 57 | 56 | 57 | 57 | 2,85E+08 | 1,31E+09 | 3,22E+09 | 20 | 20 | 29 | 63 | 85 | Oxidation (M); Carbamidomethylation; Dehydration; Deamidation (NQ); Carbamylation | 29659 | PPTG_14734 | Phytophthora parasitica INRA-310 60S ribosomal protein L7a-2 (264 aa) |
| 118 | 113 | PPTG_19127 | 202,21 | 62 | 47 | 57 | 62 | 5,86E+07 | 1,37E+08 | 8,01E+08 | 16 | 16 | 18 | 27 | 58 | Oxidation (M); Carbamidomethylation; Dehydration; Deamidation (NQ); Carbamylation | 36747 | PPTG_19127 | Phytophthora parasitica INRA-310 mitochondrial Carrier (MC) family (346 aa) |
| 495 | 1041 | PPTG_09250 | 202,17 | 55 | 36 | 55 | 55 | 1,01E+07 | 1,13E+08 | 1,26E+08 | 9 | 9 | 6 | 17 | 20 | Oxidation (M); Dehydration; Deamidation (NQ) | 20919 | PPTG_09250 | Phytophthora parasitica INRA-310 NAD(P)H:quinone oxidoreductase type IV (201 aa) |
| 120 | 388 | PPTG_02536 | 202,07 | 20 | 7 | 16 | 15 | 3,59E+06 | 2,84E+07 | 4,76E+07 | 48 | 48 | 17 | 43 | 42 | Oxidation (M); Carbamidomethylation; Dehydration; Deamidation (NQ) | 299654 | PPTG_02536 | Phytophthora parasitica INRA-310 translational activator GCN1 (2760 aa) |
| 150 | 177 | PPTG_00853 | 201,96 | 52 | 32 | 51 | 48 | 2,21E+07 | 2,10E+08 | 2,57E+08 | 24 | 24 | 16 | 41 | 37 | Oxidation (M); Carbamidomethylation; Deamidation (NQ); Acetylation (N-term) | 57146 | PPTG_00853 | Phytophthora parasitica INRA-310 inositol-3-phosphate synthase (518 aa) |
| 93 | 259 | PPTG_07764 | 201,85 | 67 | 39 | 59 | 59 | 2,18E+08 | 8,52E+08 | 2,17E+09 | 22 | 22 | 15 | 35 | 62 | Oxidation (M); Carbamidomethylation; Dehydration; Deamidation (NQ); Carbamylation | 29587 | PPTG_07764 | Phytophthora parasitica INRA-310 40S ribosomal protein S3a (262 aa) |
| 279 | 249 | PPTG_08768 | 201,54 | 22 | 5 | 11 | 21 | 7,84E+05 | 3,64E+06 | 2,89E+07 | 36 | 21 | 11 | 18 | 38 | Oxidation (M); Carbamidomethylation; Deamidation (NQ) | 218385 | PPTG_08768 | Phytophthora parasitica INRA-310 ATP-binding Cassette (ABC) superfamily (1970 aa) |
| 141 | 127 | PPTG_11971 | 201,53 | 63 | 41 | 43 | 63 | 2,08E+07 | 3,38E+07 | 2,81E+08 | 23 | 23 | 17 | 24 | 55 | Oxidation (M); Carbamidomethylation; Dehydration; Deamidation (NQ); Acetylation (N-term); Carbam | 51655 | PPTG_11971 | Phytophthora parasitica INRA-310 hypothetical protein (497 aa) |
| 230 | 153 | PPTG_13000 | 201,52 | 60 | 51 | 52 | 60 | 3,04E+07 | 1,79E+08 | 3,20E+08 | 20 | 20 | 17 | 26 | 34 | Oxidation (M); Carbamidomethylation; Dehydration; Deamidation (NQ); Carbamylation | 40839 | PPTG_13000 | Phytophthora parasitica INRA-310 dihydrolipoyl dehydrogenase 1 mitochondrial (388 aa) |
| 394 | 244 | PPTG_11916 | 201,43 | 71 | 24 | 69 | 65 | 2,25E+06 | 3,15E+07 | 4,18E+07 | 16 | 13 | 6 | 22 | 24 | Oxidation (M); Carbamidomethylation; Deamidation (NQ); Acetylation (N-term) | 36455 | PPTG_11916 | Phytophthora parasitica INRA-310 fructose-1 6-bisphosphatase (334 aa) |
| 202 | 340 | PPTG_00673 | 200,91 | 35 | 9 | 28 | 28 | 1,79E+06 | 2,84E+07 | 3,82E+07 | 36 | 36 | 10 | 35 | 36 | Oxidation (M); Carbamidomethylation; Dehydration; Deamidation (NQ) | 134949 | PPTG_00673 | Phytophthora parasitica INRA-310 Cullin-associated NEDD8-dissociated protein (1223 aa) |
| 321 | 217 | PPTG_09177 | 200,83 | 59 | 15 | 39 | 55 | 9,72E+06 | 3,84E+07 | 1,48E+08 | 24 | 24 | 8 | 20 | 33 | Oxidation (M); Carbamidomethylation; Deamidation (NQ); Amidation | 48005 | PPTG_09177 | Phytophthora parasitica INRA-310 hypothetical protein (427 aa) |
| 212 | 206 | PPTG_04610 | 200,62 | 38 | 8 | 34 | 32 | 2,78E+06 | 2,55E+07 | 4,49E+07 | 31 | 31 | 9 | 34 | 37 | Oxidation (M); Carbamidomethylation; Amidation | 120200 | PPTG_04610 | Phytophthora parasitica INRA-310 coatomer subunit beta' (1095 aa) |
| 139 | 285 | PPTG_12917 | 200,59 | 38 | 14 | 33 | 32 | 8,96E+06 | 5,47E+07 | 7,79E+07 | 40 | 40 | 16 | 41 | 40 | Oxidation (M); Carbamidomethylation; Deamidation (NQ) | 132898 | PPTG_12917 | Phytophthora parasitica INRA-310 isoleucyl-tRNA synthetase (1176 aa) |
| 264 | 460 | PPTG_17187 | 200,46 | 42 | 26 | 42 | 42 | 2,38E+07 | 3,67E+08 | 3,07E+08 | 13 | 13 | 9 | 33 | 28 | Oxidation (M); Carbamidomethylation; Deamidation (NQ); Acetylation (N-term); Carbamylation | 48688 | PPTG_17187 | Phytophthora parasitica INRA-310 glycosyl transferase (459 aa) |
| 307 | 114 | PPTG_18619 | 200,3 | 58 | 34 | 47 | 58 | 8,52E+06 | 2,35E+07 | 1,33E+08 | 20 | 20 | 12 | 17 | 34 | Oxidation (M); Carbamidomethylation; Deamidation (NQ) | 55114 | PPTG_18619 | Phytophthora parasitica INRA-310 NADH dehydrogenase flavoprotein 1 (500 aa) |
| 192 | 236 | PPTG_19136 | 200,29 | 58 | 21 | 50 | 58 | 3,94E+06 | 5,72E+07 | 7,82E+07 | 30 | 30 | 10 | 34 | 39 | Oxidation (M); Carbamidomethylation; Deamidation (NQ) | 65643 | PPTG_19136 | Phytophthora parasitica INRA-310 acyl-CoA dehydrogenase family member 9 (611 aa) |
| 340 | 223 | PPTG_16241 | 200,16 | 36 | 5 | 32 | 26 | 1,27E+06 | 4,12E+07 | 4,26E+07 | 29 | 29 | 5 | 28 | 25 | Oxidation (M); Carbamidomethylation; Dehydration; Deamidation (NQ) | 117884 | PPTG_16241 | Phytophthora parasitica INRA-310 oxoglutarate dehydrogenase (succinyl-transferring) E1 component (1044 aa) |
| 374 | 211 | PPTG_03132 | 199,92 | 31 | 6 | 29 | 28 | 3,83E+05 | 2,17E+07 | 2,03E+07 | 24 | 24 | 5 | 26 | 23 | Carbamidomethylation | 116058 | PPTG_03132 | Phytophthora parasitica INRA-310 Glu/Leu/Phe/Val dehydrogenase family protein (1033 aa) |
| 454 | 184 | PPTG_15793 | 199,89 | 37 | 18 | 14 | 37 | 2,13E+06 | 4,80E+06 | 3,29E+07 | 20 | 20 | 9 | 9 | 27 | Oxidation (M); Carbamidomethylation; Dehydration; Deamidation (NQ); Acetylation (N-term); Carbam | 70816 | PPTG_15793 | Phytophthora parasitica INRA-310 guanylate-binding protein (631 aa) |
| 148 | 156 | PPTG_17946 | 199,8 | 56 | 44 | 48 | 55 | 2,96E+07 | 1,54E+08 | 3,44E+08 | 21 | 19 | 22 | 35 | 38 | Oxidation (M); Carbamidomethylation; Deamidation (NQ); Carbamylation | 47145 | PPTG_17946 | Phytophthora parasitica INRA-310 ATP-dependent RNA helicase eIF4A (412 aa) |
| 296 | 193 | PPTG_13850 | 199,58 | 50 | 26 | 45 | 49 | 7,53E+06 | 7,92E+07 | 8,18E+07 | 22 | 22 | 12 | 26 | 26 | Oxidation (M); Carbamidomethylation | 56620 | PPTG_13850 | Phytophthora parasitica INRA-310 fumarate hydratase mitochondrial (529 aa) |
| 189 | 1308 | PPTG_07238 | 199,45 | 41 | 38 | 41 | 41 | 1,45E+08 | 4,68E+08 | 2,53E+09 | 9 | 9 | 12 | 24 | 48 | Oxidation (M); Carbamidomethylation; Dehydration; Deamidation (NQ); Amidation; Carbamylation | 24143 | PPTG_07238 | Phytophthora parasitica INRA-310 hypothetical protein (227 aa) |

| | | | | | | | | | | | | | | | | | | | |
|---|---|---|---|---|---|---|---|---|---|---|---|---|---|---|---|---|---|---|---|
| 318 | 218 | PPTG_16640 | 199,29 | 44 | 12 | 44 | 43 | 3,65E+06 | 8,64E+07 | 7,08E+07 | 19 | 19 | 5 | 30 | 26 | Oxidation (M); Carbamidomethylation; Deamidation (NQ) | 60142 | PPTG_16640 | Phytophthora parasitica INRA-310 aldehyde dehydrogenase (548 aa) |
| 348 | 202 | PPTG_19659 | 199,2 | 54 | 22 | 49 | 49 | 3,45E+06 | 4,97E+07 | 6,84E+07 | 18 | 18 | 7 | 25 | 25 | Oxidation (M); Carbamidomethylation; Amidation | 41848 | PPTG_19659 | Phytophthora parasitica INRA-310 alcohol dehydrogenase class-3 chain H (388 aa) |
| 348 | 203 | PPTG_02558 | 199,2 | 54 | 22 | 49 | 49 | 3,45E+06 | 4,97E+07 | 6,84E+07 | 18 | 18 | 7 | 25 | 25 | Oxidation (M); Carbamidomethylation; Amidation | 41848 | PPTG_02558 | Phytophthora parasitica INRA-310 alcohol dehydrogenase class-3 chain H (388 aa) |
| 254 | 453 | PPTG_11465 | 198,92 | 56 | 16 | 50 | 53 | 1,70E+07 | 4,87E+08 | 2,64E+08 | 18 | 18 | 3 | 36 | 33 | Oxidation (M); Carbamidomethylation; Dehydration; Deamidation (NQ); Carbamylation | 37550 | PPTG_11465 | Phytophthora parasitica INRA-310 hypothetical protein (336 aa) |
| 349 | 240 | PPTG_13436 | 198,67 | 27 | 5 | 22 | 25 | 1,88E+06 | 1,77E+07 | 2,94E+07 | 27 | 27 | 6 | 24 | 27 | Oxidation (M); Carbamidomethylation; Deamidation (NQ) | 133667 | PPTG_13436 | Phytophthora parasitica INRA-310 protein transporter Sec31A (1256 aa) |
| 265 | 252 | PPTG_07472 | 198,57 | 42 | 12 | 36 | 37 | 1,56E+06 | 1,84E+07 | 2,24E+07 | 31 | 20 | 9 | 29 | 31 | Oxidation (M); Carbamidomethylation; Deamidation (NQ) | 110580 | PPTG_07472 | Phytophthora parasitica INRA-310 tripeptidyl-peptidase (1002 aa) |
| 173 | 239 | PPTG_10119 | 198,56 | 39 | 17 | 31 | 37 | 5,64E+06 | 6,27E+07 | 1,34E+08 | 28 | 28 | 12 | 35 | 41 | Oxidation (M); Carbamidomethylation; Dehydration; Deamidation (NQ) | 100864 | PPTG_10119 | Phytophthora parasitica INRA-310 coatomer subunit gamma (911 aa) |
| 165 | 243 | PPTG_01922 | 198,34 | 59 | 22 | 57 | 49 | 5,32E+06 | 1,07E+08 | 9,14E+07 | 28 | 28 | 10 | 40 | 40 | Oxidation (M); Carbamidomethylation; Deamidation (NQ) | 63019 | PPTG_01922 | Phytophthora parasitica INRA-310 phosphoenolpyruvate carboxykinase (ATP) (570 aa) |
| 294 | 131 | PPTG_14608 | 198,27 | 49 | 22 | 37 | 49 | 7,41E+06 | 2,41E+07 | 1,04E+08 | 22 | 22 | 11 | 21 | 33 | Oxidation (M); Carbamidomethylation; Deamidation (NQ) | 67064 | PPTG_14608 | Phytophthora parasitica INRA-310 electron transfer flavoprotein-ubiquinone oxidoreductase (611 aa) |
| 319 | 195 | PPTG_10900 | 198,15 | 31 | 22 | 21 | 31 | 6,44E+06 | 6,90E+06 | 1,01E+08 | 24 | 24 | 18 | 15 | 28 | Oxidation (M); Deamidation (NQ) | 94064 | PPTG_10900 | Phytophthora parasitica INRA-310 N-acetylated-alpha-linked acidic dipeptidase-like protein (869 aa) |
| 278 | 201 | PPTG_03857 | 198,12 | 41 | 16 | 24 | 41 | 3,85E+06 | 1,60E+07 | 6,80E+07 | 27 | 27 | 12 | 18 | 37 | Oxidation (M); Carbamidomethylation; Deamidation (NQ) | 84097 | PPTG_03857 | Phytophthora parasitica INRA-310 vesicle-fusing ATPase (766 aa) |
| 767 | 111 | PPTG_00165 | 197,84 | 41 | 9 | 4 | 40 | 8,56E+05 | 5,14E+05 | 1,90E+07 | 19 | 18 | 4 | 2 | 23 | Oxidation (M); Carbamidomethylation; Dehydration; Deamidation (NQ) | 78749 | PPTG_00165 | Phytophthora parasitica INRA-310 beta-glucan synthesis-associated protein (724 aa) |
| 220 | 261 | PPTG_16899 | 197,65 | 56 | 28 | 54 | 54 | 2,88E+07 | 1,68E+08 | 2,69E+08 | 19 | 19 | 12 | 32 | 35 | Oxidation (M); Carbamidomethylation; Deamidation (NQ); Acetylation (N-term) | 43481 | PPTG_16899 | Phytophthora parasitica INRA-310 proliferation-associated protein metalloprotease family M24X (394 aa) |
| 251 | 294 | PPTG_14388 | 197,36 | 10 | 1 | 8 | 8 | 2,12E+05 | 1,16E+07 | 2,07E+07 | 36 | 36 | 4 | 32 | 36 | Oxidation (M); Carbamidomethylation; Dehydration; Deamidation (NQ) | 451188 | PPTG_14388 | Phytophthora parasitica INRA-310 HECT E3 ubiquitin ligase (4150 aa) |
| 461 | 187 | PPTG_16875 | 197,28 | 29 | 5 | 7 | 28 | 1,54E+06 | 3,25E+06 | 3,85E+07 | 28 | 28 | 7 | 8 | 30 | Oxidation (M); Carbamidomethylation | 144454 | PPTG_16875 | Phytophthora parasitica INRA-310 P-type ATPase (P-ATPase) superfamily (1340 aa) |
| 200 | 847 | PPTG_15783 | 196,89 | 66 | 61 | 57 | 65 | 7,17E+07 | 3,64E+08 | 8,50E+08 | 15 | 15 | 19 | 28 | 35 | Dehydration; Deamidation (NQ); Carbamylation | 21559 | PPTG_15783 | Phytophthora parasitica INRA-310 30S ribosomal protein S7e (191 aa) |
| 560 | 920 | PPTG_14116 | 196,72 | 60 | 59 | 60 | 60 | 7,41E+06 | 6,17E+07 | 7,15E+07 | 9 | 9 | 11 | 14 | 14 | Oxidation (M); Carbamidomethylation; Dehydration; Deamidation (NQ) | 26794 | PPTG_14116 | Phytophthora parasitica INRA-310 hypothetical protein (248 aa) |
| 56 | 524 | PPTG_11874 | 196,67 | 55 | 51 | 54 | 55 | 1,92E+08 | 7,09E+08 | 1,86E+09 | 15 | 15 | 34 | 46 | 60 | Oxidation (M); Carbamidomethylation; Dehydration; Deamidation (NQ); Amidation; Carbamylation | 28295 | PPTG_11874 | Phytophthora parasitica INRA-310 40S ribosomal protein S2 (262 aa) |
| 527 | 152 | PPTG_15328 | 195,86 | 25 | 5 | 9 | 24 | 1,03E+06 | 1,42E+06 | 3,27E+07 | 25 | 22 | 5 | 8 | 27 | Oxidation (M); Carbamidomethylation; Deamidation (NQ) | 154030 | PPTG_15328 | Phytophthora parasitica INRA-310 phospholipid-transporting ATPase (1392 aa) |
| 275 | 139 | PPTG_01838 | 195,48 | 54 | 14 | 47 | 54 | 2,16E+06 | 2,88E+07 | 4,56E+07 | 19 | 19 | 6 | 32 | 30 | Oxidation (M); Carbamidomethylation; Deamidation (NQ) | 62980 | PPTG_01838 | Phytophthora parasitica INRA-310 phosphoribosylaminoimidazole carboxylase (596 aa) |
| 357 | 214 | PPTG_18811 | 195,47 | 51 | 29 | 51 | 44 | 4,96E+06 | 7,39E+07 | 7,81E+07 | 14 | 14 | 10 | 24 | 22 | Oxidation (M); Carbamidomethylation; Deamidation (NQ) | 46032 | PPTG_18811 | Phytophthora parasitica INRA-310 medium-chain specific acyl-CoA dehydrogenase mitochondrial (421 aa) |
| 330 | 1917 | PPTG_11033 | 195,29 | 54 | 34 | 54 | 54 | 7,70E+07 | 3,45E+08 | 9,29E+08 | 5 | 5 | 10 | 16 | 34 | Carbamidomethylation; Dehydration; Deamidation (NQ); Carbamylation | 21959 | PPTG_11033 | Phytophthora parasitica INRA-310 hypothetical protein (212 aa) |
| 168 | 196 | PPTG_16504 | 194,8 | 51 | 24 | 42 | 50 | 1,51E+07 | 1,16E+08 | 2,69E+08 | 21 | 21 | 15 | 32 | 42 | Oxidation (M); Carbamidomethylation; Dehydration; Deamidation (NQ); Amidation | 59241 | PPTG_16504 | Phytophthora parasitica INRA-310 isocitrate lyase (538 aa) |
| 453 | 250 | PPTG_13500 | 194,8 | 33 | 13 | 28 | 32 | 1,94E+06 | 1,97E+07 | 2,37E+07 | 19 | 19 | 8 | 17 | 20 | Oxidation (M); Carbamidomethylation; Deamidation (NQ) | 77874 | PPTG_13500 | Phytophthora parasitica INRA-310 cysteinyl-tRNA synthetase (689 aa) |
| 133 | 174 | PPTG_14253 | 194,17 | 35 | 25 | 31 | 35 | 3,05E+07 | 1,72E+08 | 3,86E+08 | 19 | 18 | 20 | 37 | 42 | Oxidation (M); Carbamidomethylation; Deamidation (NQ) | 69763 | PPTG_14253 | Phytophthora parasitica INRA-310 hypothetical protein (632 aa) |
| 266 | 185 | PPTG_06616 | 193,99 | 47 | 26 | 29 | 46 | 1,53E+07 | 2,46E+07 | 1,84E+08 | 21 | 21 | 15 | 19 | 35 | Oxidation (M); Carbamidomethylation; Deamidation (NQ); Acetylation (N-term) | 60202 | PPTG_06616 | Phytophthora parasitica INRA-310 adenylosuccinate synthetase (555 aa) |
| 355 | 220 | PPTG_09817 | 193,89 | 44 | 19 | 38 | 44 | 5,44E+06 | 1,43E+07 | 1,05E+08 | 17 | 17 | 10 | 14 | 32 | Oxidation (M); Carbamidomethylation; Deamidation (NQ) | 53696 | PPTG_09817 | Phytophthora parasitica INRA-310 endoplasmic reticulum-Golgi intermediate compartment protein (470 aa) |
| 337 | 701 | PPTG_11914 | 193,79 | 58 | 32 | 41 | 58 | 5,27E+07 | 2,06E+08 | 4,37E+08 | 14 | 11 | 11 | 20 | 28 | Deamidation (NQ); Carbamylation | 24982 | PPTG_11914 | Phytophthora parasitica INRA-310 EF-1 guanine nucleotide exchange domain-containing protein (229 aa) |
| 342 | 235 | PPTG_15972 | 193,17 | 34 | 7 | 29 | 30 | 1,14E+06 | 1,56E+07 | 1,67E+07 | 25 | 23 | 7 | 26 | 25 | Oxidation (M); Carbamidomethylation | 100368 | PPTG_15972 | Phytophthora parasitica INRA-310 puromycin-sensitive aminopeptidase (885 aa) |
| 259 | 329 | PPTG_03240 | 193,16 | 22 | 9 | 21 | 18 | 1,89E+06 | 1,41E+07 | 3,24E+07 | 30 | 30 | 12 | 30 | 28 | Oxidation (M); Carbamidomethylation | 173444 | PPTG_03240 | Phytophthora parasitica INRA-310 ferredoxin-dependent glutamate synthase (1583 aa) |
| 335 | 162 | PPTG_11537 | 193,04 | 57 | 36 | 41 | 57 | 3,77E+07 | 1,26E+08 | 2,70E+08 | 17 | 17 | 16 | 19 | 24 | Oxidation (M); Carbamidomethylation; Dehydration; Deamidation (NQ) | 51895 | PPTG_11537 | Phytophthora parasitica INRA-310 lipase (480 aa) |
| 388 | 200 | PPTG_18378 | 192,71 | 73 | 43 | 50 | 73 | 1,48E+07 | 3,37E+07 | 2,38E+08 | 14 | 14 | 10 | 14 | 29 | Oxidation (M); Carbamidomethylation; Dehydration; Deamidation (NQ); Amidation | 37347 | PPTG_18378 | Phytophthora parasitica INRA-310 guanine nucleotide-binding protein G(I)/G(S)/G(T) subunit beta (345 aa) |
| 341 | 255 | PPTG_13570 | 192,62 | 52 | 33 | 44 | 44 | 9,45E+06 | 5,41E+07 | 8,00E+07 | 22 | 22 | 13 | 22 | 23 | Oxidation (M); Carbamidomethylation; Dehydration; Deamidation (NQ) | 53535 | PPTG_13570 | Phytophthora parasitica INRA-310 hypothetical protein (495 aa) |
| 291 | 531 | PPTG_09098 | 192,36 | 11 | 1 | 10 | 6 | 9,41E+04 | 1,06E+07 | 5,45E+06 | 44 | 42 | 3 | 42 | 20 | Carbamidomethylation | 517144 | PPTG_09098 | Phytophthora parasitica INRA-310 sporangia induced dynein heavy chain (4552 aa) |
| 176 | 188 | PPTG_10590 | 192,26 | 16 | 9 | 10 | 16 | | | 3,64E+06 | 21 | 1 | 18 | 21 | 48 | Oxidation (M); Carbamidomethylation; Deamidation (NQ) | 152085 | PPTG_10590 | Phytophthora parasitica INRA-310 ATP-binding Cassette (ABC) superfamily (1359 aa) |
| 270 | 253 | PPTG_15229 | 192,07 | 37 | 15 | 28 | 36 | 4,25E+06 | 1,97E+07 | 4,10E+07 | 27 | 26 | 12 | 25 | 31 | Oxidation (M); Carbamidomethylation; Deamidation (NQ); Acetylation (N-term); Carbamylation | 97308 | PPTG_15229 | Phytophthora parasitica INRA-310 hypothetical protein (872 aa) |
| 171 | 149 | PPTG_08506 | 191,83 | 49 | 27 | 43 | 47 | 9,29E+06 | 8,68E+07 | 1,15E+08 | 17 | 7 | 15 | 35 | 39 | Oxidation (M); Carbamidomethylation; Dehydration; Deamidation (NQ); Carbamylation | 49928 | PPTG_08506 | Phytophthora parasitica INRA-310 tubulin alpha chain (454 aa) |
| 171 | 150 | PPTG_08498 | 191,83 | 49 | 27 | 43 | 47 | 9,29E+06 | 8,68E+07 | 1,15E+08 | 17 | 7 | 15 | 35 | 39 | Oxidation (M); Carbamidomethylation; Dehydration; Deamidation (NQ); Carbamylation | 49898 | PPTG_08498 | Phytophthora parasitica INRA-310 tubulin alpha chain (454 aa) |
| 255 | 138 | PPTG_02328 | 191,76 | 71 | 41 | 55 | 69 | 1,20E+07 | 2,53E+07 | 1,49E+08 | 19 | 19 | 16 | 22 | 33 | Oxidation (M); Carbamidomethylation; Dehydration; Deamidation (NQ); Acetylation (N-term) | 38356 | PPTG_02328 | Phytophthora parasitica INRA-310 NADH dehydrogenase [ubiquinone] 1 alpha subcomplex subunit (345 aa) |
| 410 | 278 | PPTG_14857 | 191,75 | 21 | 5 | 18 | 19 | 2,47E+06 | 2,15E+07 | 3,12E+07 | 21 | 21 | 6 | 21 | 22 | Oxidation (M); Carbamidomethylation | 120588 | PPTG_14857 | Phytophthora parasitica INRA-310 AP-4 complex subunit epsilon (1104 aa) |
| 198 | 268 | PPTG_02398 | 191,43 | 50 | 34 | 48 | 50 | 9,53E+06 | 7,54E+07 | 1,19E+08 | 17 | 13 | 14 | 32 | 36 | Oxidation (M); Carbamidomethylation; Deamidation (NQ); Amidation; Carbamylation | 45969 | PPTG_02398 | Phytophthora parasitica INRA-310 elongation factor 1-gamma (407 aa) |
| 248 | 176 | PPTG_17524 | 191,22 | 57 | 37 | 53 | 55 | 1,89E+07 | 1,12E+08 | 2,23E+08 | 17 | 16 | 15 | 27 | 31 | Oxidation (M); Carbamidomethylation; Dehydration; Deamidation (NQ); Carbamylation | 41765 | PPTG_17524 | Phytophthora parasitica INRA-310 thioredoxin-dependent peroxide reductase (378 aa) |
| 295 | 163 | PPTG_18809 | 191,07 | 36 | 18 | 31 | 36 | 4,01E+06 | 2,70E+07 | 6,19E+07 | 22 | 22 | 11 | 24 | 29 | Oxidation (M); Carbamidomethylation; Dehydration; Deamidation (NQ) | 80269 | PPTG_18809 | Phytophthora parasitica INRA-310 prolyl-tRNA synthetase (712 aa) |
| 397 | 604 | PPTG_16306 | 191,03 | 72 | 36 | 65 | 72 | 1,57E+06 | 3,63E+07 | 4,49E+07 | 12 | 12 | 5 | 23 | 24 | Oxidation (M); Deamidation (NQ) | 24751 | PPTG_16306 | Phytophthora parasitica INRA-310 hypothetical protein (231 aa) |
| 207 | 170 | PPTG_02800 | 190,79 | 61 | 45 | 57 | 60 | 2,48E+07 | 2,25E+08 | 2,91E+08 | 19 | 19 | 16 | 30 | 35 | Oxidation (M); Carbamidomethylation; Deamidation (NQ); Acetylation (N-term) | 36669 | PPTG_02800 | Phytophthora parasitica INRA-310 transaldolase (335 aa) |
| 352 | 348 | PPTG_16142 | 190,79 | 50 | 43 | 50 | 50 | 1,04E+07 | 7,70E+07 | 9,58E+07 | 19 | 19 | 13 | 23 | 21 | Oxidation (M); Carbamidomethylation | 47212 | PPTG_16142 | Phytophthora parasitica INRA-310 glutathione S-transferase (417 aa) |
| 249 | 293 | PPTG_16510 | 190,74 | 55 | 17 | 52 | 50 | 1,30E+07 | 1,43E+08 | 1,83E+08 | 23 | 23 | 8 | 30 | 35 | Oxidation (M); Carbamidomethylation; Dehydration; Deamidation (NQ); Acetylation (N-term) | 54855 | PPTG_16510 | Phytophthora parasitica INRA-310 inosine-5'-monophosphate dehydrogenase (509 aa) |
| 383 | 238 | PPTG_14664 | 190,72 | 44 | 17 | 35 | 43 | 2,53E+06 | 2,24E+07 | 3,37E+07 | 19 | 19 | 8 | 22 | 23 | Oxidation (M); Carbamidomethylation; Deamidation (NQ); Acetylation (N-term) | 63777 | PPTG_14664 | Phytophthora parasitica INRA-310 serine/threonine-protein phosphatase 2A regulatory subunit A (578 aa) |
| 492 | 199 | PPTG_20006 | 190,44 | 66 | 36 | 45 | 62 | 3,32E+06 | 7,55E+06 | 5,40E+07 | 15 | 15 | 9 | 13 | 21 | Oxidation (M); Carbamidomethylation; Dehydration; Deamidation (NQ) | 32963 | PPTG_20006 | Phytophthora parasitica INRA-310 alpha-soluble NSF attachment protein (294 aa) |
| 217 | 219 | PPTG_00098 | 190,42 | 74 | 63 | 69 | 72 | 5,60E+07 | 1,46E+08 | 4,07E+08 | 22 | 22 | 17 | 27 | 35 | Oxidation (M); Deamidation (NQ) | 34207 | PPTG_00098 | Phytophthora parasitica INRA-310 hypothetical protein (318 aa) |
| 104 | 521 | PPTG_13005 | 190,35 | 67 | 52 | 53 | 67 | 9,63E+07 | 4,55E+08 | 1,46E+09 | 14 | 14 | 18 | 39 | 51 | Oxidation (M); Carbamidomethylation; Dehydration; Deamidation (NQ); Carbamylation | 25688 | PPTG_13005 | Phytophthora parasitica INRA-310 50S ribosomal protein L1 (230 aa) |
| 354 | 376 | PPTG_13491 | 190,08 | 23 | 14 | 12 | 23 | 5,85E+06 | 6,65E+06 | 5,49E+07 | 26 | 26 | 16 | 12 | 28 | Oxidation (M); Carbamidomethylation | 126887 | PPTG_13491 | Phytophthora parasitica INRA-310 phospholipase D Pi-PXPH-PLD (1118 aa) |
| 329 | 166 | PPTG_03845 | 189,77 | 37 | 17 | 24 | 37 | 9,73E+06 | 6,97E+06 | 7,28E+07 | 19 | 17 | 9 | 16 | 35 | Oxidation (M); Carbamidomethylation; Deamidation (NQ); Acetylation (N-term) | 64468 | PPTG_03845 | Phytophthora parasitica INRA-310 cell 5A endo-1 4-betaglucanase (583 aa) |
| 229 | 256 | PPTG_06869 | 189,53 | 33 | 24 | 33 | 28 | 1,30E+07 | 7,45E+07 | 1,17E+08 | 22 | 21 | 20 | 30 | 27 | Oxidation (M); Carbamidomethylation; Carbamylation | 83906 | PPTG_06869 | Phytophthora parasitica INRA-310 threonyl-tRNA synthetase (745 aa) |
| 239 | 190 | PPTG_18357 | 189,26 | 18 | 9 | 10 | 17 | | 8,98E+04 | 1,50E+05 | 21 | 3 | 14 | 17 | 44 | Oxidation (M); Carbamidomethylation; Deamidation (NQ) | 143109 | PPTG_18357 | Phytophthora parasitica INRA-310 ATP-binding Cassette (ABC) superfamily (1277 aa) |
| 72 | 279 | PPTG_05552 | 189,09 | 64 | 40 | 56 | 62 | 2,09E+07 | 2,33E+08 | 2,61E+08 | 16 | 16 | 23 | 53 | 51 | Oxidation (M); Carbamidomethylation; Dehydration; Deamidation (NQ); Carbamylation | 39403 | PPTG_05552 | Phytophthora parasitica INRA-310 fructose-bisphosphate aldolase (359 aa) |
| 72 | 280 | PPTG_05553 | 189,09 | 64 | 40 | 56 | 62 | 2,09E+07 | 2,33E+08 | 2,61E+08 | 16 | 16 | 23 | 53 | 51 | Oxidation (M); Carbamidomethylation; Dehydration; Deamidation (NQ); Carbamylation | 39461 | PPTG_05553 | Phytophthora parasitica INRA-310 fructose-bisphosphate aldolase (359 aa) |
| 333 | 277 | PPTG_02824 | 189 | 48 | 7 | 43 | 43 | 9,53E+05 | 3,02E+07 | 2,49E+07 | 24 | 24 | 4 | 29 | 26 | Oxidation (M); Carbamidomethylation; Deamidation (NQ) | 69658 | PPTG_02824 | Phytophthora parasitica INRA-310 choline/Carnitine O-acyltransferase (622 aa) |
| 792 | 913 | PPTG_12292 | 188,48 | 36 | 6 | 32 | 36 | 5,72E+05 | 1,28E+07 | 1,37E+07 | 10 | 10 | 2 | 13 | 14 | Oxidation (M); Carbamidomethylation | 51478 | PPTG_12292 | Phytophthora parasitica INRA-310 hypothetical protein (480 aa) |
| 470 | 191 | PPTG_10113 | 188,42 | 28 | 4 | 24 | 23 | 5,23E+05 | 1,43E+07 | 9,23E+06 | 19 | 19 | 3 | 21 | 20 | Oxidation (M); Carbamidomethylation; Dehydration; Deamidation (NQ) | 117248 | PPTG_10113 | Phytophthora parasitica INRA-310 importin-like protein (1080 aa) |
| 314 | 180 | PPTG_14687 | 188,39 | 58 | 38 | 49 | 55 | 8,71E+06 | 4,67E+07 | 1,09E+08 | 17 | 17 | 13 | 23 | 26 | Oxidation (M); Carbamidomethylation; Deamidation (NQ) | 46702 | PPTG_14687 | Phytophthora parasitica INRA-310 homoserine dehydrogenase (437 aa) |
| 381 | 205 | PPTG_00180 | 188,39 | 54 | 32 | 44 | 50 | 4,66E+06 | 3,64E+07 | 7,84E+07 | 20 | 20 | 11 | 21 | 21 | Oxidation (M); Carbamidomethylation; Deamidation (NQ) | 52482 | PPTG_00180 | Phytophthora parasitica INRA-310 UDP-glucose 6-dehydrogenase (476 aa) |
| 273 | 168 | PPTG_07998 | 188,28 | 51 | 33 | 43 | 48 | 5,47E+06 | 2,59E+07 | 9,94E+07 | 20 | 20 | 14 | 21 | 33 | Oxidation (M); Carbamidomethylation; Dehydration; Deamidation (NQ) | 52225 | PPTG_07998 | Phytophthora parasitica INRA-310 V-type proton ATPase subunit H (464 aa) |
| 728 | 355 | PPTG_06649 | 188,16 | 27 | 8 | 7 | 26 | 1,39E+06 | 1,77E+06 | 2,17E+07 | 13 | 13 | 6 | 7 | 18 | Oxidation (M); Carbamidomethylation; Deamidation (NQ) | 76842 | PPTG_06649 | Phytophthora parasitica INRA-310 hypothetical protein (720 aa) |
| 528 | 1017 | PPTG_00284 | 187,99 | 53 | 41 | 53 | 50 | 7,27E+06 | 3,38E+07 | 2,32E+08 | 11 | 11 | 7 | 14 | 19 | Oxidation (M); Deamidation (NQ) | 23556 | PPTG_00284 | Phytophthora parasitica INRA-310 cytochrome b-c1 complex subunit Rieske (216 aa) |
| 458 | 216 | PPTG_14739 | 187,96 | 29 | 14 | 12 | 25 | 3,34E+06 | 4,33E+06 | 3,65E+07 | 21 | 13 | 11 | 11 | 23 | Oxidation (M); Carbamidomethylation; Dehydration | 103057 | PPTG_14739 | Phytophthora parasitica INRA-310 hypothetical protein (896 aa) |
| 331 | 276 | PPTG_02671 | 187,68 | 52 | 45 | 49 | 52 | 1,02E+07 | 6,67E+07 | 1,06E+08 | 18 | 18 | 15 | 21 | 24 | Oxidation (M); Carbamidomethylation; Deamidation (NQ) | 40841 | PPTG_02671 | Phytophthora parasitica INRA-310 glutathione S-transferase omega-like protein (355 aa) |
| 242 | 262 | PPTG_20479 | 187,65 | 58 | 39 | 44 | 58 | 1,56E+07 | 4,77E+07 | 2,27E+08 | 17 | 17 | 16 | 22 | 36 | Oxidation (M); Carbamidomethylation; Deamidation (NQ) | 45034 | PPTG_20479 | Phytophthora parasitica INRA-310 NADH-ubiquinone oxidoreductase 49 kDa subunit (393 aa) |
| 378 | 304 | PPTG_02226 | 187,37 | 31 | 12 | 13 | 25 | 9,72E+05 | 1,11E+06 | 9,21E+06 | 25 | 16 | 13 | 11 | 29 | Oxidation (M); Carbamidomethylation; Acetylation (N-term); Amidation; Carbamylation | 116571 | PPTG_02226 | Phytophthora parasitica INRA-310 P-type ATPase (P-ATPase) superfamily (1068 aa) |
| 280 | 209 | PPTG_11055 | 187,06 | 42 | 26 | 28 | 42 | 5,38E+06 | 1,06E+07 | 8,58E+07 | 22 | 22 | 15 | 16 | 36 | Oxidation (M); Carbamidomethylation | 68180 | PPTG_11055 | Phytophthora parasitica INRA-310 glycerol-3-phosphate dehydrogenase (616 aa) |
| 301 | 282 | PPTG_16969 | 186,93 | 61 | 38 | 58 | 59 | 7,40E+06 | 5,49E+07 | 6,85E+07 | 19 | 19 | 15 | 25 | 24 | Oxidation (M); Dehydration; Deamidation (NQ); Acetylation (N-term) | 40055 | PPTG_16969 | Phytophthora parasitica INRA-310 carbohydrate-binding protein (370 aa) |
| 344 | 291 | PPTG_02785 | 186,9 | 22 | 16 | 17 | 22 | 2,08E+07 | 1,62E+07 | 1,69E+08 | 16 | 16 | 15 | 14 | 29 | Oxidation (M); Carbamidomethylation; Deamidation (NQ) | 80562 | PPTG_02785 | Phytophthora parasitica INRA-310 V-type H(+)-translocating pyrophosphatase (769 aa) |
| 312 | 248 | PPTG_13257 | 186,83 | 42 | 25 | 31 | 38 | 6,87E+06 | 2,69E+07 | 6,24E+07 | 19 | 19 | 13 | 21 | 28 | Oxidation (M); Carbamidomethylation; Deamidation (NQ) | 55075 | PPTG_13257 | Phytophthora parasitica INRA-310 serine hydroxymethyltransferase (503 aa) |
| 459 | 226 | PPTG_07317 | 186,66 | 57 | 26 | 43 | 53 | 2,93E+06 | 2,35E+07 | 4,25E+07 | 16 | 16 | 7 | 15 | 23 | Oxidation (M); Carbamidomethylation; Deamidation (NQ); Acetylation (N-term) | 49590 | PPTG_07317 | Phytophthora parasitica INRA-310 adenylyl cyclase-associated protein (462 aa) |
| 382 | 230 | PPTG_06269 | 186,34 | 55 | 28 | 52 | 46 | 6,23E+06 | 7,06E+07 | 7,68E+07 | 19 | 19 | 9 | 25 | 19 | Oxidation (M); Carbamidomethylation; Acetylation (N-term) | 47031 | PPTG_06269 | Phytophthora parasitica INRA-310 succinyl-CoA ligase beta-chain (438 aa) |
| 367 | 222 | PPTG_03926 | 186,29 | 56 | 22 | 45 | 55 | 8,16E+06 | 5,45E+07 | 8,12E+07 | 23 | 23 | 9 | 20 | 26 | Oxidation (M); Carbamidomethylation; Carbamylation | 58109 | PPTG_03926 | Phytophthora parasitica INRA-310 aspartyl-tRNA synthetase (515 aa) |
| 250 | 1220 | PPTG_17667 | 185,69 | 45 | 37 | 40 | 45 | 1,22E+07 | 1,40E+08 | 1,37E+08 | 7 | 6 | 8 | 32 | 33 | Oxidation (M); Carbamidomethylation; Dehydration; Deamidation (NQ); Acetylation (N-term); Carbam | 30399 | PPTG_17667 | Phytophthora parasitica INRA-310 chitinase (277 aa) |
| 384 | 175 | PPTG_03931 | 185,33 | 32 | 18 | 20 | 32 | 5,49E+06 | 1,50E+07 | 7,49E+07 | 18 | 18 | 12 | 14 | 27 | Oxidation (M); Carbamidomethylation; Carbamylation | 75107 | PPTG_03931 | Phytophthora parasitica INRA-310 annexin (Annexin) family (691 aa) |
| 109 | 748 | PPTG_04112 | 185,29 | 62 | 43 | 62 | 62 | 7,13E+07 | 8,63E+08 | 8,22E+08 | 12 | 12 | 13 | 45 | 48 | Oxidation (M); Deamidation (NQ); Carbamylation | 28169 | PPTG_04112 | Phytophthora parasitica INRA-310 manganese superoxide dismutase (251 aa) |
| 541 | 662 | PPTG_15407 | 185,27 | 25 | 17 | 23 | 25 | 2,03E+06 | 2,73E+07 | 2,60E+07 | 10 | 10 | 8 | 15 | 17 | Oxidation (M); Carbamidomethylation; Deamidation (NQ); Acetylation (N-term) | 72766 | PPTG_15407 | Phytophthora parasitica INRA-310 hypothetical protein (683 aa) |
| 119 | 582 | PPTG_16072 | 185,23 | 57 | 57 | 49 | 57 | 1,24E+08 | 4,67E+08 | 1,26E+09 | 14 | 9 | 23 | 32 | 48 | Dehydration; Deamidation (NQ); Carbamylation | 24212 | PPTG_16072 | Phytophthora parasitica INRA-310 60S ribosomal protein L6 (221 aa) |
| 621 | 228 | PPTG_15489 | 185,2 | 39 | 12 | 20 | 35 | 9,26E+05 | 4,37E+06 | 1,30E+07 | 16 | 14 | 5 | 10 | 20 | Oxidation (M) | 68270 | PPTG_15489 | Phytophthora parasitica INRA-310 hsp70-like protein (609 aa) |
| 503 | 269 | PPTG_12608 | 184,97 | 31 | 12 | 9 | 31 | 2,61E+06 | 2,68E+06 | 4,26E+07 | 18 | 18 | 9 | 6 | 27 | Oxidation (M); Carbamidomethylation | 95371 | PPTG_12608 | Phytophthora parasitica INRA-310 hypothetical protein (846 aa) |
| 468 | 264 | PPTG_19677 | 184,96 | 36 | 16 | 17 | 34 | 2,22E+05 | | 7,72E+06 | 21 | 6 | 12 | 10 | 22 | Carbamidomethylation; Acetylation (N-term) | 93342 | PPTG_19677 | Phytophthora parasitica INRA-310 phosphatidyl inositol kinase (PIK-B) (837 aa) |
| 327 | 510 | PPTG_18605 | 184,94 | 12 | 1 | 1 | 12 | 2,99E+05 | 9,48E+05 | 1,54E+07 | 47 | 47 | 6 | 7 | 47 | Carbamidomethylation | 585533 | PPTG_18605 | Phytophthora parasitica INRA-310 hypothetical protein (5449 aa) |
| 518 | 295 | PPTG_13495 | 184,92 | 42 | 16 | 39 | 37 | 1,87E+06 | 2,15E+07 | 3,08E+07 | 16 | 16 | 8 | 17 | 16 | Oxidation (M); Deamidation (NQ); Acetylation (N-term) | 53117 | PPTG_13495 | Phytophthora parasitica INRA-310 inosine-5'-monophosphate dehydrogenase (500 aa) |
| 420 | 339 | PPTG_02400 | 184,88 | 55 | 24 | 42 | 52 | 7,78E+06 | 5,08E+07 | 9,20E+07 | 23 | 22 | 7 | 18 | 24 | Oxidation (M); Carbamidomethylation | 39658 | PPTG_02400 | Phytophthora parasitica INRA-310 thioredoxin-like protein (363 aa) |
| 472 | 241 | PPTG_08136 | 184,8 | 24 | 13 | 16 | 24 | 4,71E+06 | 4,56E+06 | 5,62E+07 | 16 | 16 | 10 | 13 | 21 | Oxidation (M); Carbamidomethylation | 97911 | PPTG_08136 | Phytophthora parasitica INRA-310 dolichyl-diphosphooligosaccharide-protein glycosyltransferase subunit STT3A (886 aa) |
| 364 | 381 | PPTG_09870 | 184,67 | 16 | 2 | 5 | 15 | 1,29E+07 | 2,48E+07 | 2,49E+07 | 37 | 37 | 5 | 12 | 38 | Oxidation (M); Carbamidomethylation; Dehydration; Deamidation (NQ) | 317105 | PPTG_09870 | Phytophthora parasitica INRA-310 hypothetical protein (2874 aa) |
| 502 | 359 | PPTG_16844 | 184,4 | 61 | 24 | 52 | 61 | 2,72E+06 | 2,42E+07 | 3,76E+07 | 15 | 15 | 5 | 17 | 20 | Oxidation (M) | 33923 | PPTG_16844 | Phytophthora parasitica INRA-310 2 4-dienoyl-CoA reductase (323 aa) |
| 457 | 375 | PPTG_00768 | 184,26 | 35 | 24 | 33 | 35 | 4,93E+06 | 2,46E+07 | 4,84E+07 | 14 | 14 | 9 | 16 | 20 | Oxidation (M); Carbamidomethylation; Deamidation (NQ) | 55072 | PPTG_00768 | Phytophthora parasitica INRA-310 coronin-like protein (502 aa) |
| 160 | 921 | PPTG_19399 | 184,25 | 77 | 49 | 77 | 77 | 5,41E+07 | 3,71E+08 | 6,81E+08 | 11 | 10 | 13 | 33 | 45 | Oxidation (M); Carbamidomethylation; Dehydration; Deamidation (NQ); Carbamylation | 18291 | PPTG_19399 | Phytophthora parasitica INRA-310 peptidyl-prolyl cis-trans isomerase (172 aa) |
| 308 | 246 | PPTG_13453 | 183,84 | 57 | 32 | 35 | 57 | 1,79E+07 | 2,09E+07 | 2,32E+08 | 18 | 18 | 13 | 13 | 37 | Oxidation (M); Carbamidomethylation; Dehydration; Deamidation (NQ) | 40812 | PPTG_13453 | Phytophthora parasitica INRA-310 G-protein alpha subunit 1 (356 aa) |
| 390 | 258 | PPTG_11336 | 183,7 | 30 | 10 | 22 | 28 | 1,52E+06 | 1,82E+07 | 2,39E+07 | 20 | 20 | 8 | 19 | 25 | Oxidation (M); Carbamidomethylation | 93404 | PPTG_11336 | Phytophthora parasitica INRA-310 AP-1 complex subunit beta (842 aa) |
| 366 | 362 | PPTG_05088 | 183,61 | 28 | 5 | 24 | 17 | 4,95E+05 | 2,02E+07 | 1,18E+07 | 29 | 29 | 5 | 30 | 20 | Oxidation (M); Carbamidomethylation; Deamidation (NQ) | 140132 | PPTG_05088 | Phytophthora parasitica INRA-310 5-oxoprolinase (1293 aa) |
| 493 | 936 | PPTG_15134 | 183,55 | 81 | 42 | 81 | 80 | 5,47E+07 | 2,86E+08 | 6,34E+08 | 11 | 11 | 7 | 18 | 18 | Oxidation (M); Carbamidomethylation; Acetylation (N-term); Carbamylation | 15102 | PPTG_15134 | Phytophthora parasitica INRA-310 40S ribosomal protein S12 (139 aa) |
| 285 | 287 | PPTG_14749 | 183,53 | 62 | 26 | 43 | 58 | 4,52E+06 | 2,74E+07 | 5,17E+07 | 23 | 23 | 11 | 23 | 32 | Oxidation (M); Carbamidomethylation | 51753 | PPTG_14749 | Phytophthora parasitica INRA-310 seryl-tRNA synthetase (455 aa) |
| 324 | 458 | PPTG_05429 | 183,35 | 41 | 20 | 34 | 41 | 6,40E+06 | 6,07E+07 | 1,49E+08 | 14 | 14 | 9 | 22 | 30 | Oxidation (M); Carbamidomethylation; Dehydration; Deamidation (NQ); Acetylation (N-term); Amidatic | 43829 | PPTG_05429 | Phytophthora parasitica INRA-310 cAMP-dependent protein kinase regulatory subunit (395 aa) |
| 406 | 371 | PPTG_02700 | 183,35 | 66 | 31 | 65 | 57 | 2,78E+06 | 3,65E+07 | 3,19E+07 | 15 | 15 | 9 | 23 | 18 | Oxidation (M); Carbamidomethylation | 34331 | PPTG_02700 | Phytophthora parasitica INRA-310 carbon-nitrogen hydrolase (316 aa) |
| 267 | 406 | PPTG_04052 | 183,32 | 39 | 9 | 26 | 38 | 1,78E+06 | 2,47E+07 | 3,74E+07 | 31 | 31 | 9 | 28 | 32 | Oxidation (M); Carbamidomethylation; Deamidation (NQ); Acetylation (N-term) | 120641 | PPTG_04052 | Phytophthora parasitica INRA-310 phosphoribosylglycinamide synthetase (1144 aa) |
| 231 | 477 | PPTG_12075 | 183,31 | 14 | 4 | 13 | 10 | 8,30E+05 | 1,24E+07 | 1,47E+07 | 37 | 37 | 9 | 35 | 32 | Oxidation (M); Carbamidomethylation | 324746 | PPTG_12075 | Phytophthora parasitica INRA-310 ubiquitin-specific protease (2914 aa) |
| 615 | 506 | PPTG_16811 | 183,27 | 61 | 15 | 55 | 61 | 6,29E+05 | 3,60E+07 | 1,42E+07 | 13 | 13 | 3 | 18 | 15 | Oxidation (M) | 32878 | PPTG_16811 | Phytophthora parasitica INRA-310 aldose 1-epimerase (301 aa) |
| 379 | 331 | PPTG_16755 | 182,93 | 33 | 9 | 27 | 25 | 1,66E+06 | 1,43E+07 | 1,81E+07 | 27 | 27 | 10 | 22 | 21 | Oxidation (M); Carbamidomethylation | 116898 | PPTG_16755 | Phytophthora parasitica INRA-310 alpha-actinin-1 (1035 aa) |
| 361 | 305 | PPTG_19423 | 182,87 | 59 | 19 | 29 | 57 | 3,63E+06 | 7,68E+06 | 1,03E+08 | 23 | 23 | 8 | 13 | 35 | Oxidation (M); Carbamidomethylation; Dehydration; Deamidation (NQ) | 63310 | PPTG_19423 | Phytophthora parasitica INRA-310 hypothetical protein (576 aa) |
| 396 | 254 | PPTG_01358 | 182,84 | 44 | 28 | 42 | 42 | 1,43E+06 | 2,13E+07 | 2,23E+07 | 17 | 17 | 10 | 19 | 23 | Oxidation (M); Carbamidomethylation; Deamidation (NQ); Acetylation (N-term) | 57599 | PPTG_01358 | Phytophthora parasitica INRA-310 glyceraldehyde-3-phosphate dehydrogenase (525 aa) |
| 309 | 421 | PPTG_02403 | 182,79 | 16 | 4 | 11 | 14 | 1,28E+06 | 9,38E+06 | 1,78E+07 | 33 | 33 | 9 | 24 | 29 | Oxidation (M); Carbamidomethylation | 275276 | PPTG_02403 | Phytophthora parasitica INRA-310 pre-mRNA-processing-splicing factor 8 (2360 aa) |

| | | | | | | | | | | | | | | | | | | | |
|---|---|---|---|---|---|---|---|---|---|---|---|---|---|---|---|---|---|---|---|
| 625 | 545 | PPTG_09838 | 182,76 | 36 | 15 | 36 | 36 | 1,81E+06 | 3,20E+07 | 3,42E+07 | 12 | 12 | 5 | 15 | 15 | Oxidation (M); Carbamidomethylation | 50724 | PPTG_09838 | Phytophthora parasitica INRA-310 programmed cell death protein (470 aa) |
| 228 | 194 | PPTG_08466 | 182,76 | 47 | 28 | 42 | 44 | 1,74E+05 | 9,60E+05 | 8,92E+05 | 17 | 3 | 15 | 27 | 36 | Oxidation (M); Carbamidomethylation; Dehydration; Deamidation (NQ) | 49941 | PPTG_08466 | Phytophthora parasitica INRA-310 tubulin alpha chain (453 aa) |
| 389 | 365 | PPTG_00417 | 182,66 | 25 | 11 | 7 | 24 | 3,32E+06 | 2,82E+06 | 3,56E+07 | 26 | 24 | 13 | 9 | 30 | Oxidation (M); Carbamidomethylation; Deamidation (NQ) | 148130 | PPTG_00417 | Phytophthora parasitica INRA-310 P-type ATPase (P-ATPase) superfamily (1325 aa) |
| 626 | 403 | PPTG_02851 | 182,6 | 25 | 5 | 25 | 15 | 5,63E+05 | 1,20E+07 | 5,86E+06 | 18 | 18 | 4 | 20 | 11 | Carbamidomethylation | 114937 | PPTG_02851 | Phytophthora parasitica INRA-310 E1 family ubiquitin activating enzyme (1036 aa) |
| 626 | 404 | PPTG_02851 | 182,6 | 25 | 4 | 25 | 15 | 5,63E+05 | 1,20E+07 | 5,86E+06 | 18 | 18 | 4 | 20 | 11 | Carbamidomethylation | 117830 | PPTG_02851 | Phytophthora parasitica INRA-310 E1 family ubiquitin activating enzyme (1064 aa) |
| 274 | 225 | PPTG_12936 | 182,58 | 28 | 16 | 27 | 28 | 2,02E+06 | 1,30E+07 | 3,89E+07 | 17 | 17 | 10 | 23 | 35 | Oxidation (M); Carbamidomethylation; Acetylation (N-term) | 96482 | PPTG_12936 | Phytophthora parasitica INRA-310 protein transport Sec23/Sec24-like protein (862 aa) |
| 95 | 605 | PPTG_05975 | 182,47 | 55 | 41 | 55 | 55 | 3,81E+07 | 7,14E+08 | 5,96E+08 | 13 | 13 | 12 | 57 | 43 | Oxidation (M); Carbamidomethylation; Dehydration; Deamidation (NQ); Carbamylation | 30601 | PPTG_05975 | Phytophthora parasitica INRA-310 carbonic anhydrase (273 aa) |
| 490 | 288 | PPTG_02370 | 182,3 | 29 | 12 | 28 | 20 | 8,90E+06 | 7,01E+07 | 7,51E+07 | 14 | 14 | 7 | 19 | 17 | Oxidation (M); Carbamidomethylation; Deamidation (NQ) | 83906 | PPTG_02370 | Phytophthora parasitica INRA-310 endo-1 3(4)-beta-glucanase (751 aa) |
| 180 | 580 | PPTG_18566 | 182,21 | 15 | 1 | 11 | 10 | 6,38E+05 | 1,10E+07 | 1,51E+07 | 55 | 55 | 6 | 41 | 39 | Oxidation (M); Carbamidomethylation; Dehydration; Deamidation (NQ) | 495119 | PPTG_18566 | Phytophthora parasitica INRA-310 dynein heavy chain (4397 aa) |
| 519 | 298 | PPTG_06281 | 182,17 | 34 | 12 | 31 | 26 | 2,22E+06 | 1,81E+07 | 3,95E+07 | 19 | 19 | 7 | 17 | 17 | Carbamidomethylation | 83795 | PPTG_06281 | Phytophthora parasitica INRA-310 propionyl coenzyme A carboxylase (Pi-PCC1) (765 aa) |
| 404 | 495 | PPTG_09222 | 182,13 | 45 | 29 | 34 | 45 | 9,60E+06 | 1,80E+07 | 1,80E+08 | 14 | 13 | 9 | 14 | 27 | Oxidation (M); Carbamidomethylation; Amidation | 43035 | PPTG_09222 | Phytophthora parasitica INRA-310 calcium/calmodulin-dependent protein kinase (386 aa) |
| 277 | 349 | PPTG_08673 | 181,99 | 21 | 5 | 14 | 18 | 1,48E+06 | 1,19E+07 | 2,68E+07 | 32 | 32 | 10 | 24 | 33 | Oxidation (M); Carbamidomethylation; Carbamylation | 224059 | PPTG_08673 | Phytophthora parasitica INRA-310 brefeldin A-inhibited guanine nucleotide-exchange protein (2047 aa) |
| 411 | 197 | PPTG_00390 | 181,96 | 45 | 28 | 27 | 45 | 3,89E+06 | 8,47E+06 | 4,86E+07 | 17 | 17 | 12 | 13 | 24 | Oxidation (M); Dehydration; Deamidation (NQ) | 60703 | PPTG_00390 | Phytophthora parasitica INRA-310 apoptosis-inducing factor 1 (560 aa) |
| 55 | 257 | PPTG_06192 | 181,92 | 75 | 62 | 75 | 75 | 2,63E+08 | 1,20E+09 | 3,06E+09 | 17 | 17 | 35 | 45 | 60 | Oxidation (M); Carbamidomethylation; Dehydration; Deamidation (NQ); Carbamylation | 27053 | PPTG_06192 | Phytophthora parasitica INRA-310 40S ribosomal protein (247 aa) |
| 512 | 468 | PPTG_10783 | 181,75 | 17 | 13 | 13 | 15 | 8,79E+06 | 4,23E+07 | 1,12E+08 | 11 | 11 | 12 | 16 | 14 | Carbamidomethylation; Deamidation (NQ) | 112828 | PPTG_10783 | Phytophthora parasitica INRA-310 hypothetical protein (1076 aa) |
| 702 | 485 | PPTG_18587 | 181,72 | 32 | 6 | 30 | 32 | 8,96E+05 | 1,72E+07 | 2,52E+07 | 13 | 13 | 3 | 13 | 16 | Carbamidomethylation; Deamidation (NQ) | 55777 | PPTG_18587 | Phytophthora parasitica INRA-310 exoglucanase 1 (522 aa) |
| 969 | 350 | PPTG_16769 | 181,62 | 19 | 8 | 12 | 19 | 2,79E+05 | 1,35E+06 | 8,00E+06 | 12 | 12 | 4 | 6 | 13 | Carbamidomethylation | 112981 | PPTG_16769 | Phytophthora parasitica INRA-310 tenascin-like protein (1061 aa) |
| 414 | 283 | PPTG_02259 | 181,54 | 49 | 16 | 42 | 47 | 3,82E+06 | 5,42E+07 | 9,31E+07 | 19 | 19 | 5 | 20 | 24 | Oxidation (M); Carbamidomethylation; Dehydration | 50557 | PPTG_02259 | Phytophthora parasitica INRA-310 eukaryotic translation initiation factor 2 subunit 3 (463 aa) |
| 418 | 424 | PPTG_08401 | 181,16 | 65 | 31 | 59 | 65 | 5,92E+06 | 6,35E+07 | 7,53E+07 | 15 | 14 | 8 | 19 | 22 | Oxidation (M); Carbamidomethylation; Dehydration; Acetylation (N-term) | 29868 | PPTG_08401 | Phytophthora parasitica INRA-310 26S proteasome non-ATPase regulatory subunit (265 aa) |
| 477 | 412 | PPTG_12440 | 180,84 | 58 | 29 | 45 | 58 | 2,15E+06 | 1,74E+07 | 5,60E+07 | 14 | 14 | 7 | 13 | 24 | Oxidation (M); Carbamidomethylation; Deamidation (NQ) | 35408 | PPTG_12440 | Phytophthora parasitica INRA-310 hypothetical protein (330 aa) |
| 529 | 539 | PPTG_00500 | 180,74 | 52 | 28 | 51 | 44 | 3,81E+06 | 5,75E+07 | 5,13E+07 | 15 | 15 | 6 | 20 | 14 | Oxidation (M); Carbamidomethylation | 27089 | PPTG_00500 | Phytophthora parasitica INRA-310 electron transfer flavoprotein subunit beta (252 aa) |
| 437 | 323 | PPTG_13924 | 180,6 | 42 | 5 | 40 | 35 | 3,86E+05 | 2,15E+07 | 2,14E+07 | 23 | 23 | 2 | 24 | 20 | Oxidation (M); Carbamidomethylation | 84539 | PPTG_13924 | Phytophthora parasitica INRA-310 prolyl endopeptidase (753 aa) |
| 716 | 229 | PPTG_09717 | 180,51 | 25 | 9 | 3 | 23 | 1,40E+06 | 9,50E+05 | 1,95E+07 | 19 | 19 | 7 | 3 | 21 | Oxidation (M); Carbamidomethylation; Deamidation (NQ) | 111809 | PPTG_09717 | Phytophthora parasitica INRA-310 ATP-binding Cassette (ABC) superfamily (1012 aa) |
| 245 | 315 | PPTG_07447 | 180,4 | 32 | 8 | 21 | 31 | 2,24E+06 | 1,56E+07 | 3,81E+07 | 26 | 26 | 8 | 26 | 39 | Oxidation (M); Carbamidomethylation; Dehydration; Deamidation (NQ) | 103597 | PPTG_07447 | Phytophthora parasitica INRA-310 Argonaute1 (AGO1) (931 aa) |
| 564 | 352 | PPTG_06246 | 180,31 | 37 | 10 | 12 | 36 | 1,36E+06 | 4,15E+06 | 2,89E+07 | 18 | 17 | 8 | 8 | 22 | Oxidation (M); Carbamidomethylation; Deamidation (NQ) | 74090 | PPTG_06246 | Phytophthora parasitica INRA-310 protein kinase (675 aa) |
| 37 | 620 | PPTG_14424 | 180,3 | 67 | 60 | 60 | 67 | 2,93E+08 | 1,18E+09 | 2,87E+09 | 15 | 15 | 30 | 52 | 77 | Oxidation (M); Carbamidomethylation; Dehydration; Deamidation (NQ); Acetylation (N-term); 2 more | 21905 | PPTG_14424 | Phytophthora parasitica INRA-310 60S ribosomal protein L9 (190 aa) |
| 350 | 265 | PPTG_02597 | 180,18 | 26 | 9 | 24 | 25 | 1,65E+06 | 2,11E+07 | 2,38E+07 | 19 | 19 | 7 | 27 | 23 | Oxidation (M); Carbamidomethylation; Dehydration; Deamidation (NQ); Acetylation (N-term) | 93943 | PPTG_02597 | Phytophthora parasitica INRA-310 importin subunit beta (859 aa) |
| 365 | 626 | PPTG_06597 | 179,98 | 35 | 25 | 27 | 31 | 2,71E+07 | 1,03E+08 | 3,27E+08 | 15 | 15 | 12 | 19 | 24 | Oxidation (M); Carbamidomethylation; Dehydration; Deamidation (NQ); Carbamylation | 44825 | PPTG_06597 | Phytophthora parasitica INRA-310 hypothetical protein (427 aa) |
| 992 | 598 | PPTG_14738 | 179,81 | 26 | 8 | 8 | 26 | 1,24E+06 | 1,53E+06 | 2,95E+07 | 11 | 11 | 4 | 4 | 14 | Carbamidomethylation; Deamidation (NQ) | 61753 | PPTG_14738 | Phytophthora parasitica INRA-310 hypothetical protein (554 aa) |
| 465 | 290 | PPTG_18479 | 179,8 | 71 | 17 | 71 | 63 | 2,62E+06 | 4,34E+07 | 4,98E+07 | 15 | 15 | 5 | 21 | 19 | Oxidation (M); Carbamidomethylation; Dehydration; Deamidation (NQ); Amidation | 38743 | PPTG_18479 | Phytophthora parasitica INRA-310 hypothetical protein (361 aa) |
| 71 | 366 | PPTG_10646 | 179,75 | 58 | 45 | 58 | 58 | 1,44E+08 | 7,55E+08 | 1,91E+09 | 18 | 18 | 23 | 51 | 53 | Oxidation (M); Carbamidomethylation; Dehydration; Deamidation (NQ); Acetylation (N-term); Carbam | 29669 | PPTG_10646 | Phytophthora parasitica INRA-310 40S ribosomal protein S4 (262 aa) |
| 658 | 687 | PPTG_19949 | 179,71 | 42 | 6 | 42 | 34 | | 7,01E+06 | 3,29E+06 | 10 | 7 | 2 | 20 | 12 | Oxidation (M); Carbamidomethylation; Deamidation (NQ) | 50909 | PPTG_19949 | Phytophthora parasitica INRA-310 serine protease family S08A (488 aa) |
| 628 | 354 | PPTG_16088 | 179,68 | 47 | 14 | 38 | 42 | 9,20E+05 | 1,62E+07 | 1,64E+07 | 15 | 15 | 4 | 15 | 16 | Oxidation (M); Carbamidomethylation; Deamidation (NQ) | 44525 | PPTG_16088 | Phytophthora parasitica INRA-310 pyruvate dehydrogenase E1 component subunit alpha (403 aa) |
| 257 | 462 | PPTG_13499 | 179,62 | 58 | 42 | 58 | 58 | 1,28E+06 | 1,36E+07 | 3,48E+07 | 13 | 8 | 14 | 23 | 34 | Oxidation (M); Dehydration; Deamidation (NQ) | 26190 | PPTG_13499 | Phytophthora parasitica INRA-310 60S ribosomal protein L6 (234 aa) |
| 596 | 308 | PPTG_11926 | 179,61 | 37 | 10 | 11 | 35 | 1,33E+06 | 1,50E+06 | 1,41E+07 | 22 | 22 | 6 | 6 | 24 | Dehydration; Deamidation (NQ) | 88653 | PPTG_11926 | Phytophthora parasitica INRA-310 hypothetical protein (795 aa) |
| 971 | 937 | PPTG_00140 | 179,55 | 27 | 5 | 27 | 27 | 3,70E+05 | 1,04E+07 | 6,45E+06 | 9 | 8 | 2 | 11 | 10 | Carbamidomethylation | 48820 | PPTG_00140 | Phytophthora parasitica INRA-310 glycoside hydrolase (461 aa) |
| 405 | 266 | PPTG_09673 | 179,55 | 73 | 35 | 47 | 68 | 2,53E+06 | 1,21E+07 | 4,44E+07 | 18 | 18 | 8 | 15 | 27 | Oxidation (M); Carbamidomethylation; Deamidation (NQ); Acetylation (N-term) | 36793 | PPTG_09673 | Phytophthora parasitica INRA-310 hypothetical protein (343 aa) |
| 100 | 1164 | PPTG_13069 | 179,41 | 25 | 22 | 22 | 25 | 6,20E+08 | 7,26E+08 | 3,58E+09 | 10 | 10 | 37 | 25 | 47 | Oxidation (M); Carbamidomethylation; Dehydration; Deamidation (NQ); Acetylation (N-term); Carbam | 38891 | PPTG_13069 | Phytophthora parasitica INRA-310 hypothetical protein (358 aa) |
| 413 | 416 | PPTG_11437 | 179,13 | 58 | 22 | 58 | 45 | 6,04E+06 | 6,60E+07 | 8,42E+07 | 16 | 16 | 7 | 22 | 20 | Oxidation (M); Carbamidomethylation; Dehydration; Deamidation (NQ) | 36482 | PPTG_11437 | Phytophthora parasitica INRA-310 malate dehydrogenase (337 aa) |
| 441 | 245 | PPTG_04024 | 179,07 | 40 | 9 | 31 | 37 | 8,05E+05 | 8,48E+06 | 2,07E+07 | 15 | 15 | 4 | 19 | 23 | Oxidation (M); Carbamidomethylation; Deamidation (NQ) | 61147 | PPTG_04024 | Phytophthora parasitica INRA-310 protein kinase (556 aa) |
| 538 | 498 | PPTG_09495 | 178,94 | 34 | 21 | 32 | 28 | 2,84E+06 | 1,97E+07 | 2,70E+07 | 12 | 12 | 8 | 16 | 16 | Oxidation (M); Carbamidomethylation | 54302 | PPTG_09495 | Phytophthora parasitica INRA-310 NADP-specific glutamate dehydrogenase (495 aa) |
| 431 | 267 | PPTG_02716 | 178,92 | 57 | 31 | 27 | 52 | 4,26E+06 | 4,68E+06 | 7,46E+07 | 20 | 20 | 10 | 9 | 28 | Oxidation (M); Carbamidomethylation; Deamidation (NQ) | 47051 | PPTG_02716 | Phytophthora parasitica INRA-310 ribose-phosphate pyrophosphokinase (426 aa) |
| 544 | 341 | PPTG_03875 | 178,79 | 9 | 3 | 3 | 9 | 1,14E+06 | 2,55E+06 | 1,32E+07 | 23 | 23 | 9 | 8 | 22 | Deamidation (NQ) | 325544 | PPTG_03875 | Phytophthora parasitica INRA-310 hypothetical protein (2896 aa) |
| 685 | 273 | PPTG_10537 | 178,59 | 23 | 6 | 1 | 23 | 2,99E+05 | 0 | 1,18E+07 | 21 | 20 | 6 | 1 | 25 | Oxidation (M); Carbamidomethylation | 143376 | PPTG_10537 | Phytophthora parasitica INRA-310 P-type ATPase (P-ATPase) superfamily (1292 aa) |
| 244 | 661 | PPTG_00340 | 178,57 | 11 | 1 | 9 | 7 | 3,25E+05 | 9,20E+06 | 8,48E+06 | 43 | 42 | 5 | 39 | 29 | Oxidation (M); Carbamidomethylation; Deamidation (NQ) | 528140 | PPTG_00340 | Phytophthora parasitica INRA-310 dynein heavy chain (4623 aa) |
| 539 | 758 | PPTG_03645 | 178,44 | 44 | 28 | 31 | 44 | 5,42E+06 | 1,91E+07 | 8,35E+07 | 12 | 12 | 9 | 12 | 19 | Carbamidomethylation; Deamidation (NQ); Amidation | 40630 | PPTG_03645 | Phytophthora parasitica INRA-310 hypothetical protein (380 aa) |
| 368 | 224 | PPTG_17319 | 178,23 | 37 | 13 | 35 | 37 | 4,07E+06 | 2,67E+07 | 5,52E+07 | 18 | 18 | 8 | 22 | 25 | Oxidation (M); Carbamidomethylation; Dehydration | 62488 | PPTG_17319 | Phytophthora parasitica INRA-310 eukaryotic translation initiation factor 3 (552 aa) |
| 636 | 318 | PPTG_12256 | 178,16 | 19 | 6 | 13 | 17 | 1,11E+06 | 7,38E+06 | 1,81E+07 | 17 | 17 | 5 | 11 | 18 | Oxidation (M); Carbamidomethylation | 120865 | PPTG_12256 | Phytophthora parasitica INRA-310 protein transporter Sec24 (1121 aa) |
| 326 | 579 | PPTG_11360 | 177,93 | 27 | 10 | 27 | 27 | 3,96E+06 | 5,16E+07 | 4,48E+07 | 10 | 10 | 9 | 29 | 23 | Oxidation (M); Carbamidomethylation; Deamidation (NQ) | 68017 | PPTG_11360 | Phytophthora parasitica INRA-310 carbohydrate esterase (664 aa) |
| 1205 | 456 | PPTG_08833 | 177,88 | 25 | 5 | 2 | 23 | 1,98E+05 | 3,82E+05 | 1,06E+07 | 11 | 11 | 2 | 1 | 15 | Carbamidomethylation | 71494 | PPTG_08833 | Phytophthora parasitica INRA-310 tenascin-like protein (680 aa) |
| 785 | 303 | PPTG_15911 | 177,87 | 46 | 16 | 24 | 46 | 6,19E+05 | 4,30E+06 | 2,12E+07 | 16 | 16 | 4 | 9 | 16 | Carbamidomethylation; Deamidation (NQ); Amidation | 61963 | PPTG_15911 | Phytophthora parasitica INRA-310 calnexin (558 aa) |
| 128 | 557 | PPTG_19077 | 177,87 | 75 | 37 | 68 | 54 | 1,75E+08 | 6,44E+08 | 1,77E+09 | 15 | 15 | 16 | 35 | 49 | Oxidation (M); Carbamidomethylation; Dehydration; Deamidation (NQ); Acetylation (N-term); Carbam | 30757 | PPTG_19077 | Phytophthora parasitica INRA-310 40S ribosomal protein SA (283 aa) |
| 356 | 270 | PPTG_07737 | 177,83 | 42 | 18 | 40 | 38 | 3,29E+06 | 2,14E+07 | 3,64E+07 | 18 | 18 | 8 | 24 | 24 | Oxidation (M); Carbamidomethylation; Dehydration; Deamidation (NQ) | 56817 | PPTG_07737 | Phytophthora parasitica INRA-310 T-complex protein 1 subunit beta (527 aa) |
| 858 | 804 | PPTG_13082 | 177,78 | 59 | 23 | 28 | 59 | 3,23E+06 | 1,08E+07 | 6,47E+07 | 10 | 10 | 4 | 6 | 17 | Oxidation (M); Carbamidomethylation | 30318 | PPTG_13082 | Phytophthora parasitica INRA-310 NADH dehydrogenase flavoprotein 2 (272 aa) |
| 571 | 516 | PPTG_15631 | 177,78 | 63 | 46 | 34 | 63 | 1,53E+07 | 1,02E+07 | 1,23E+08 | 12 | 10 | 11 | 11 | 16 | Carbamidomethylation; Deamidation (NQ) | 29949 | PPTG_15631 | Phytophthora parasitica INRA-310 voltage-dependent anion-selective channel protein (283 aa) |
| 516 | 466 | PPTG_12116 | 177,75 | 56 | 28 | 34 | 52 | 7,15E+06 | 1,24E+07 | 1,16E+08 | 13 | 13 | 9 | 12 | 21 | Oxidation (M); Carbamidomethylation; Deamidation (NQ) | 44376 | PPTG_12116 | Phytophthora parasitica INRA-310 ATP synthase (C/AC39) subunit (395 aa) |
| 358 | 300 | PPTG_00991 | 177,73 | 42 | 10 | 35 | 33 | 2,94E+06 | 4,22E+07 | 5,49E+07 | 20 | 20 | 7 | 25 | 23 | Oxidation (M); Carbamidomethylation; Deamidation (NQ) | 65616 | PPTG_00991 | Phytophthora parasitica INRA-310 dihydroxy-acid dehydratase (617 aa) |
| 282 | 260 | PPTG_10595 | 177,66 | 15 | 6 | 8 | 13 | | 2,08E+05 | 9,08E+05 | 17 | 5 | 12 | 14 | 41 | Oxidation (M); Carbamidomethylation; Deamidation (NQ) | 152648 | PPTG_10595 | Phytophthora parasitica INRA-310 ATP-binding Cassette (ABC) superfamily (1355 aa) |
| 480 | 281 | PPTG_13968 | 177,59 | 49 | 26 | 37 | 46 | 2,27E+06 | 1,28E+07 | 1,77E+07 | 15 | 15 | 8 | 17 | 19 | Oxidation (M); Carbamidomethylation | 51754 | PPTG_13968 | Phytophthora parasitica INRA-310 ubiquinone biosynthesis monooxygenase (483 aa) |
| 552 | 344 | PPTG_08403 | 177,48 | 39 | 12 | 35 | 32 | 2,27E+06 | 2,98E+07 | 3,26E+07 | 13 | 13 | 4 | 17 | 18 | Oxidation (M); Carbamidomethylation; Carbamylation | 51264 | PPTG_08403 | Phytophthora parasitica INRA-310 xylose isomerase 1 (453 aa) |
| 376 | 306 | PPTG_18230 | 177,15 | 47 | 20 | 42 | 31 | 2,01E+06 | 2,72E+07 | 3,62E+07 | 22 | 22 | 9 | 25 | 20 | Oxidation (M); Carbamidomethylation; Dehydration; Deamidation (NQ); Acetylation (N-term); Carbam | 57303 | PPTG_18230 | Phytophthora parasitica INRA-310 T-complex protein 1 subunit delta (534 aa) |
| 403 | 302 | PPTG_03709 | 177,1 | 43 | 14 | 31 | 40 | 1,96E+06 | 2,48E+07 | 5,25E+07 | 17 | 12 | 6 | 19 | 24 | Oxidation (M); Carbamidomethylation; Dehydration; Deamidation (NQ); Acetylation (N-term) | 54531 | PPTG_03709 | Phytophthora parasitica INRA-310 pyruvate kinase (506 aa) |
| 690 | 711 | PPTG_16236 | 176,87 | 24 | 9 | 24 | 20 | 4,37E+06 | 3,25E+07 | 8,55E+07 | 11 | 11 | 4 | 13 | 15 | Oxidation (M); Carbamidomethylation; Deamidation (NQ) | 57650 | PPTG_16236 | Phytophthora parasitica INRA-310 transglutaminase elicitor M81D (531 aa) |
| 565 | 296 | PPTG_02106 | 176,86 | 42 | 14 | 15 | 39 | 2,13E+06 | 2,63E+06 | 3,68E+07 | 21 | 21 | 8 | 8 | 22 | Oxidation (M); Carbamidomethylation | 83316 | PPTG_02106 | Phytophthora parasitica INRA-310 serine protease family S15 (741 aa) |
| 322 | 446 | PPTG_06377 | 176,75 | 40 | 6 | 29 | 36 | 5,51E+05 | 1,26E+07 | 1,72E+07 | 29 | 27 | 5 | 27 | 29 | Oxidation (M); Carbamidomethylation; Deamidation (NQ) | 102026 | PPTG_06377 | Phytophthora parasitica INRA-310 metalloprotease family M01 (903 aa) |
| 371 | 289 | PPTG_11378 | 176,65 | 43 | 19 | 41 | 38 | 5,10E+06 | 4,88E+07 | 6,87E+07 | 18 | 18 | 8 | 25 | 22 | Oxidation (M); Carbamidomethylation | 56100 | PPTG_11378 | Phytophthora parasitica INRA-310 eukaryotic translation initiation factor 3 subunit E-interacting protein (491 aa) |
| 449 | 364 | PPTG_02894 | 176,63 | 62 | 45 | 62 | 62 | 1,13E+07 | 1,61E+08 | 1,35E+08 | 13 | 13 | 9 | 20 | 17 | Oxidation (M); Carbamidomethylation; Deamidation (NQ) | 30014 | PPTG_02894 | Phytophthora parasitica INRA-310 enoyl-CoA hydratase (280 aa) |
| 122 | 499 | PPTG_03669 | 176,54 | 65 | 37 | 55 | 61 | 2,20E+08 | 7,67E+08 | 2,10E+09 | 14 | 14 | 22 | 30 | 49 | Oxidation (M); Carbamidomethylation; Dehydration; Deamidation (NQ); Acetylation (N-term); Carbam | 21852 | PPTG_03669 | Phytophthora parasitica INRA-310 40S ribosomal protein (197 aa) |
| 284 | 379 | PPTG_06725 | 176,51 | 21 | 10 | 9 | 20 | 4,66E+06 | 1,47E+06 | 5,05E+07 | 26 | 16 | 14 | 14 | 38 | Oxidation (M); Carbamidomethylation; Dehydration; Deamidation (NQ) | 151504 | PPTG_06725 | Phytophthora parasitica INRA-310 ATP-binding Cassette (ABC) superfamily (1350 aa) |
| 385 | 377 | PPTG_04949 | 176,34 | 62 | 31 | 59 | 62 | 3,98E+06 | 3,63E+07 | 5,47E+07 | 13 | 13 | 6 | 23 | 24 | Oxidation (M); Carbamidomethylation; Deamidation (NQ) | 28778 | PPTG_04949 | Phytophthora parasitica INRA-310 hypothetical protein (273 aa) |
| 463 | 233 | PPTG_17401 | 176,31 | 50 | 39 | 44 | 50 | 5,05E+06 | 1,67E+07 | 5,62E+07 | 15 | 15 | 12 | 15 | 18 | Carbamidomethylation | 45145 | PPTG_17401 | Phytophthora parasitica INRA-310 hypothetical protein (417 aa) |
| 429 | 400 | PPTG_18213 | 176,14 | 35 | 1 | 27 | 24 | 6,70E+04 | 1,25E+07 | 1,63E+07 | 27 | 27 | 1 | 24 | 22 | Oxidation (M); Carbamidomethylation | 123320 | PPTG_18213 | Phytophthora parasitica INRA-310 CRM1 C terminal Exportin 1-like protein (1077 aa) |
| 713 | 1361 | PPTG_07817 | 176,11 | 28 | 11 | 28 | 27 | 1,55E+06 | 3,64E+07 | 3,22E+07 | 8 | 8 | 4 | 17 | 11 | Oxidation (M); Carbamidomethylation | 52583 | PPTG_07817 | Phytophthora parasitica INRA-310 hypothetical protein (507 aa) |
| 246 | 515 | PPTG_18731 | 176,08 | 14 | 4 | 8 | 13 | 2,03E+06 | 1,33E+07 | 2,79E+07 | 35 | 35 | 13 | 23 | 37 | Oxidation (M); Carbamidomethylation; Dehydration | 317612 | PPTG_18731 | Phytophthora parasitica INRA-310 hypothetical protein (2886 aa) |
| 177 | 1252 | PPTG_07085 | 175,88 | 56 | 56 | 56 | 56 | 9,57E+07 | 4,35E+08 | 1,26E+09 | 11 | 11 | 14 | 31 | 42 | Oxidation (M); Carbamidomethylation; Dehydration; Deamidation (NQ); Carbamylation | 15217 | PPTG_07085 | Phytophthora parasitica INRA-310 ribosomal S17 (132 aa) |
| 497 | 418 | PPTG_11500 | 175,84 | 31 | 15 | 27 | 31 | 1,51E+07 | 3,45E+07 | 1,19E+08 | 14 | 14 | 10 | 16 | 17 | Oxidation (M); Carbamidomethylation | 62429 | PPTG_11500 | Phytophthora parasitica INRA-310 hypothetical protein (564 aa) |
| 392 | 360 | PPTG_02443 | 175,82 | 41 | 9 | 32 | 37 | 1,26E+06 | 1,40E+07 | 2,11E+07 | 24 | 24 | 7 | 22 | 23 | Carbamidomethylation; Dehydration; Deamidation (NQ) | 88755 | PPTG_02443 | Phytophthora parasitica INRA-310 dipeptidyl peptidase (802 aa) |
| 392 | 361 | PPTG_02443 | 175,82 | 41 | 9 | 32 | 37 | 1,26E+06 | 1,40E+07 | 2,11E+07 | 24 | 24 | 7 | 22 | 23 | Carbamidomethylation; Dehydration; Deamidation (NQ) | 88755 | PPTG_02443 | Phytophthora parasitica INRA-310 dipeptidyl peptidase (802 aa) |
| 517 | 317 | PPTG_15878 | 175,8 | 35 | 8 | 18 | 33 | 2,07E+06 | 1,90E+07 | 2,94E+07 | 21 | 21 | 6 | 14 | 21 | Oxidation (M); Carbamidomethylation; Deamidation (NQ) | 89402 | PPTG_15878 | Phytophthora parasitica INRA-310 ribonucleoside-diphosphate reductase large subunit (793 aa) |
| 271 | 320 | PPTG_11894 | 175,73 | 51 | 29 | 49 | 44 | 6,43E+06 | 5,61E+07 | 5,33E+07 | 22 | 21 | 11 | 29 | 28 | Oxidation (M); Carbamidomethylation; Deamidation (NQ) | 48469 | PPTG_11894 | Phytophthora parasitica INRA-310 26S protease regulatory subunit 7 (439 aa) |
| 269 | 272 | PPTG_16668 | 175,42 | 51 | 38 | 51 | 44 | 1,24E+07 | 7,58E+07 | 8,08E+07 | 17 | 17 | 18 | 28 | 23 | Oxidation (M); Carbamidomethylation; Deamidation (NQ) | 43880 | PPTG_16668 | Phytophthora parasitica INRA-310 26S proteasome non-ATPase regulatory subunit 13 (384 aa) |
| 570 | 471 | PPTG_00631 | 175,33 | 43 | 27 | 42 | 43 | 3,04E+06 | 3,48E+07 | 4,30E+07 | 12 | 12 | 7 | 14 | 17 | Carbamidomethylation; Deamidation (NQ) | 45778 | PPTG_00631 | Phytophthora parasitica INRA-310 hypothetical protein (419 aa) |
| 1101 | 899 | PPTG_06749 | 175,1 | 55 | 5 | 55 | 31 | 2,76E+05 | 1,68E+07 | 9,72E+06 | 9 | 9 | 1 | 13 | 6 | Oxidation (M); Carbamidomethylation; Deamidation (NQ) | 36290 | PPTG_06749 | Phytophthora parasitica INRA-310 ribonuclease (342 aa) |
| 386 | 286 | PPTG_12038 | 174,88 | 66 | 23 | 52 | 66 | 1,97E+06 | 1,75E+07 | 3,90E+07 | 18 | 17 | 9 | 20 | 24 | Oxidation (M); Carbamidomethylation; Dehydration; Deamidation (NQ) | 36635 | PPTG_12038 | Phytophthora parasitica INRA-310 serine/threonine-protein phosphatase PP2A (320 aa) |
| 576 | 309 | PPTG_05659 | 174,75 | 34 | 5 | 32 | 24 | 1,95E+05 | 1,20E+07 | 6,44E+06 | 15 | 15 | 3 | 20 | 15 | Oxidation (M); Carbamidomethylation; Deamidation (NQ) | 82764 | PPTG_05659 | Phytophthora parasitica INRA-310 carbohydrate-binding protein (782 aa) |
| 646 | 588 | PPTG_12822 | 174,7 | 35 | 16 | 33 | 33 | 5,71E+05 | 1,63E+07 | 1,28E+07 | 13 | 13 | 4 | 18 | 12 | Carbamidomethylation; Dehydration; Deamidation (NQ) | 48198 | PPTG_12822 | Phytophthora parasitica INRA-310 hypothetical protein (458 aa) |
| 446 | 374 | PPTG_13120 | 174,49 | 53 | 30 | 47 | 45 | 3,34E+06 | 2,38E+07 | 1,35E+08 | 17 | 17 | 8 | 16 | 22 | Oxidation (M); Deamidation (NQ) | 36313 | PPTG_13120 | Phytophthora parasitica INRA-310 annexin (Annexin) family (330 aa) |
| 469 | 747 | PPTG_02295 | 174,22 | 43 | 25 | 43 | 40 | 4,92E+06 | 4,85E+07 | 5,45E+07 | 10 | 10 | 10 | 17 | 17 | Carbamidomethylation; Dehydration; Deamidation (NQ) | 36759 | PPTG_02295 | Phytophthora parasitica INRA-310 26S proteasome non-ATPase regulatory subunit 7 (325 aa) |
| 790 | 1454 | PPTG_01302 | 174,2 | 22 | 13 | 22 | 18 | 6,20E+06 | 5,47E+07 | 7,38E+07 | 6 | 6 | 5 | 12 | 12 | Oxidation (M); Carbamidomethylation; Deamidation (NQ) | 54193 | PPTG_01302 | Phytophthora parasitica INRA-310 hypothetical protein (513 aa) |
| 635 | 274 | PPTG_07760 | 174,2 | 35 | 15 | 30 | 30 | 9,58E+05 | 1,30E+07 | 2,12E+07 | 16 | 16 | 6 | 13 | 15 | Oxidation (M); Carbamidomethylation; Deamidation (NQ) | 61993 | PPTG_07760 | Phytophthora parasitica INRA-310 phosphoacetylglucosamine mutase (562 aa) |
| 325 | 369 | PPTG_05022 | 174,07 | 48 | 23 | 40 | 46 | 1,02E+07 | 8,86E+07 | 1,38E+08 | 21 | 21 | 9 | 22 | 29 | Oxidation (M); Carbamidomethylation; Deamidation (NQ) | 46799 | PPTG_05022 | Phytophthora parasitica INRA-310 aspartate aminotransferase (427 aa) |
| 603 | 778 | PPTG_09874 | 173,98 | 40 | 18 | 40 | 40 | 5,38E+06 | 7,58E+07 | 1,24E+08 | 9 | 9 | 5 | 15 | 16 | Oxidation (M); Carbamidomethylation | 39474 | PPTG_09874 | Phytophthora parasitica INRA-310 glutamine synthetase (358 aa) |
| 622 | 332 | PPTG_02839 | 173,95 | 32 | 11 | 28 | 27 | 9,47E+05 | 9,96E+06 | 1,19E+07 | 16 | 16 | 6 | 15 | 14 | Carbamidomethylation | 68672 | PPTG_02839 | Phytophthora parasitica INRA-310 replication protein A DNA-binding subunit (629 aa) |
| 443 | 479 | PPTG_00924 | 173,9 | 40 | 10 | 30 | 36 | 4,98E+06 | 4,14E+07 | 5,97E+07 | 21 | 21 | 6 | 17 | 23 | Oxidation (M); Carbamidomethylation | 76003 | PPTG_00924 | Phytophthora parasitica INRA-310 glycyl-tRNA synthetase (682 aa) |
| 851 | 1086 | PPTG_16233 | 173,73 | 13 | 8 | 13 | 9 | 8,84E+05 | 2,00E+07 | 1,49E+07 | 7 | 7 | 5 | 13 | 9 | Oxidation (M); Carbamidomethylation; Dehydration; Deamidation (NQ) | 94266 | PPTG_16233 | Phytophthora parasitica INRA-310 transglutaminase elicitor (911 aa) |
| 592 | 508 | PPTG_18330 | 173,71 | 48 | 10 | 44 | 48 | 1,35E+06 | 2,14E+07 | 3,17E+07 | 14 | 14 | 4 | 15 | 18 | Oxidation (M); Carbamidomethylation | 41508 | PPTG_18330 | Phytophthora parasitica INRA-310 nucleosome assembly protein 1 (369 aa) |
| 535 | 489 | PPTG_08378 | 173,69 | 38 | 16 | 31 | 38 | 3,22E+06 | 3,40E+07 | 4,31E+07 | 13 | 13 | 6 | 15 | 19 | Oxidation (M); Carbamidomethylation | 46619 | PPTG_08378 | Phytophthora parasitica INRA-310 26S proteasome non-ATPase regulatory subunit 6 (417 aa) |
| 586 | 310 | PPTG_12430 | 173,55 | 41 | 11 | 14 | 41 | 8,11E+05 | 1,44E+06 | 2,47E+07 | 19 | 19 | 6 | 7 | 24 | Carbamidomethylation; Deamidation (NQ) | 72166 | PPTG_12430 | Phytophthora parasitica INRA-310 hypothetical protein (638 aa) |
| 315 | 292 | PPTG_05337 | 173,53 | 50 | 20 | 43 | 50 | 5,89E+06 | 4,72E+07 | 6,87E+07 | 17 | 17 | 8 | 27 | 27 | Oxidation (M); Carbamidomethylation; Dehydration; Acetylation (N-term) | 55004 | PPTG_05337 | Phytophthora parasitica INRA-310 glutamate decarboxylase (494 aa) |
| 451 | 536 | PPTG_03719 | 173,35 | 20 | 5 | 9 | 18 | 2,25E+06 | 1,42E+07 | 3,60E+07 | 27 | 27 | 8 | 12 | 25 | Oxidation (M); Carbamidomethylation | 173057 | PPTG_03719 | Phytophthora parasitica INRA-310 myosin-like protein (1536 aa) |
| 666 | 691 | PPTG_15739 | 173,13 | 44 | 19 | 33 | 41 | 1,58E+06 | 7,78E+06 | 1,61E+07 | 13 | 13 | 6 | 11 | 16 | Oxidation (M); Deamidation (NQ) | 35081 | PPTG_15739 | Phytophthora parasitica INRA-310 30S ribosomal protein S5 (313 aa) |
| 474 | 335 | PPTG_08167 | 173,06 | 61 | 20 | 40 | 57 | 3,14E+06 | 2,67E+07 | 3,66E+07 | 17 | 17 | 6 | 15 | 23 | Oxidation (M); Carbamidomethylation; Dehydration; Deamidation (NQ); Amidation | 39406 | PPTG_08167 | Phytophthora parasitica INRA-310 AhpC/TSA family Redoxin (368 aa) |
| 434 | 481 | PPTG_03886 | 173,05 | 67 | 47 | 57 | 56 | 7,56E+06 | 6,53E+07 | 6,80E+07 | 13 | 13 | 10 | 19 | 18 | Oxidation (M); Carbamidomethylation; Deamidation (NQ); Acetylation (N-term) | 26607 | PPTG_03886 | Phytophthora parasitica INRA-310 proteasome subunit alpha type-5 (245 aa) |
| 896 | 457 | PPTG_05574 | 172,95 | 52 | 7 | 9 | 52 | 5,22E+05 | 9,68E+05 | 3,54E+07 | 14 | 14 | 3 | 3 | 19 | Oxidation (M) | 40963 | PPTG_05574 | Phytophthora parasitica INRA-310 stomatin-like protein (377 aa) |
| 597 | 299 | PPTG_16051 | 172,9 | 15 | 2 | 12 | 13 | 3,31E+05 | 4,21E+06 | 8,85E+06 | 18 | 18 | 3 | 16 | 17 | Carbamidomethylation; Deamidation (NQ); Acetylation (N-term) | 168478 | PPTG_16051 | Phytophthora parasitica INRA-310 intraflagellar transporter 140 (1508 aa) |
| 412 | 402 | PPTG_05308 | 172,87 | 49 | 13 | 40 | 46 | 2,18E+06 | 1,93E+07 | 2,24E+07 | 21 | 21 | 6 | 21 | 22 | Oxidation (M); Carbamidomethylation; Deamidation (NQ); Acetylation (N-term) | 59180 | PPTG_05308 | Phytophthora parasitica INRA-310 T-complex protein 1 subunit alpha (547 aa) |

| | | | | | | | | | | | | | | | | | | | |
|---|---|---|---|---|---|---|---|---|---|---|---|---|---|---|---|---|---|---|---|
| 272 | 443 | PPTG_05779 | 172,7 | 43 | 12 | 32 | 40 | 2,21E+06 | 1,98E+07 | 3,99E+07 | 27 | 27 | 9 | 26 | 33 | Oxidation (M); Carbamidomethylation; Dehydration; Deamidation (NQ) | 89415 | PPTG_05779 | Phytophthora parasitica INRA-310 acyl-CoA dehydrogenase (802 aa) |
| 179 | 627 | PPTG_04484 | 172,52 | 57 | 48 | 55 | 57 | 4,94E+07 | 4,16E+08 | 5,69E+08 | 13 | 13 | 16 | 36 | 35 | Oxidation (M); Deamidation (NQ); Acetylation (N-term); Carbamylation | 28470 | PPTG_04484 | Phytophthora parasitica INRA-310 14-3-3-like protein (250 aa) |
| 506 | 398 | PPTG_14511 | 172,2 | 27 | 6 | 23 | 17 | 4,88E+05 | 9,76E+06 | 9,08E+06 | 23 | 23 | 5 | 22 | 15 | Oxidation (M); Carbamidomethylation | 123513 | PPTG_14511 | Phytophthora parasitica INRA-310 importin-like protein (1130 aa) |
| 336 | 327 | PPTG_11181 | 172,14 | 51 | 39 | 42 | 51 | 1,33E+07 | 1,99E+07 | 1,19E+08 | 14 | 13 | 14 | 17 | 28 | Oxidation (M); Carbamidomethylation | 39622 | PPTG_11181 | Phytophthora parasitica INRA-310 protein kinase (360 aa) |
| 310 | 425 | PPTG_08153 | 172,13 | 47 | 21 | 45 | 44 | 1,27E+06 | 2,44E+07 | 4,02E+07 | 18 | 17 | 8 | 26 | 27 | Oxidation (M); Carbamidomethylation; Amidation | 44886 | PPTG_08153 | Phytophthora parasitica INRA-310 26S protease regulatory subunit 8 like A (402 aa) |
| 599 | 297 | PPTG_07818 | 172,05 | 25 | 7 | 12 | 24 | 9,23E+05 | 4,24E+06 | 1,68E+07 | 18 | 18 | 6 | 9 | 20 | Carbamidomethylation; Deamidation (NQ) | 115249 | PPTG_07818 | Phytophthora parasitica INRA-310 neutral alpha-glucosidase (1023 aa) |
| 504 | 430 | PPTG_01506 | 171,99 | 27 | 6 | 21 | 21 | 8,44E+05 | 1,47E+07 | 1,81E+07 | 22 | 22 | 5 | 18 | 19 | Carbamidomethylation; Dehydration; Deamidation (NQ) | 114230 | PPTG_01506 | Phytophthora parasitica INRA-310 aspartate aminotransferase (1029 aa) |
| 694 | 301 | PPTG_02078 | 171,91 | 46 | 30 | 19 | 44 | 4,65E+06 | 3,58E+06 | 3,27E+07 | 14 | 14 | 10 | 7 | 15 | Oxidation (M); Carbamidomethylation | 50028 | PPTG_02078 | Phytophthora parasitica INRA-310 dolichyl-diphosphooligosaccharide-protein glycosyltransferase 48kD subunit (449 aa) |
| 577 | 455 | PPTG_09099 | 171,89 | 51 | 15 | 51 | 44 | 1,09E+06 | 3,18E+07 | 2,72E+07 | 13 | 13 | 4 | 21 | 13 | Oxidation (M); Deamidation (NQ) | 39015 | PPTG_09099 | Phytophthora parasitica INRA-310 ornithine carbamoyltransferase (354 aa) |
| 556 | 415 | PPTG_10597 | 171,69 | 63 | 25 | 62 | 59 | 1,23E+06 | 2,80E+07 | 2,31E+07 | 15 | 11 | 5 | 17 | 17 | Carbamidomethylation; Acetylation (N-term) | 28105 | PPTG_10597 | Phytophthora parasitica INRA-310 glutathione S-transferase (252 aa) |
| 345 | 1128 | PPTG_04475 | 171,68 | 61 | 54 | 61 | 61 | 2,30E+07 | 1,99E+08 | 4,80E+08 | 7 | 7 | 11 | 19 | 28 | Oxidation (M); Dehydration; Deamidation (NQ); Amidation | 20615 | PPTG_04475 | Phytophthora parasitica INRA-310 hypothetical protein (192 aa) |
| 590 | 426 | PPTG_00941 | 171,68 | 48 | 24 | 44 | 38 | 2,22E+06 | 2,15E+07 | 2,02E+07 | 13 | 13 | 7 | 19 | 11 | Carbamidomethylation; Deamidation (NQ) | 43556 | PPTG_00941 | Phytophthora parasitica INRA-310 acetyl-CoA acetyltransferase (413 aa) |
| 400 | 394 | PPTG_16147 | 171,59 | 48 | 26 | 42 | 35 | 6,73E+06 | 6,26E+07 | 7,00E+07 | 19 | 19 | 13 | 19 | 19 | Oxidation (M); Carbamylation | 46511 | PPTG_16147 | Phytophthora parasitica INRA-310 glutathione S-transferase (418 aa) |
| 941 | 529 | PPTG_15403 | 171,59 | 53 | 17 | 12 | 50 | 1,90E+06 | 1,47E+06 | 2,92E+07 | 13 | 13 | 6 | 4 | 14 | Carbamidomethylation | 44086 | PPTG_15403 | Phytophthora parasitica INRA-310 hypothetical protein (395 aa) |
| 398 | 338 | PPTG_02533 | 171,55 | 65 | 32 | 63 | 63 | 3,54E+06 | 4,76E+07 | 5,14E+07 | 16 | 16 | 8 | 23 | 21 | Oxidation (M); Carbamidomethylation; Deamidation (NQ) | 31079 | PPTG_02533 | Phytophthora parasitica INRA-310 dihydrodipicolinate synthase (293 aa) |
| 511 | 702 | PPTG_12671 | 171,5 | 60 | 39 | 59 | 52 | 6,95E+06 | 5,60E+07 | 5,43E+07 | 12 | 12 | 11 | 16 | 15 | Oxidation (M); Deamidation (NQ); Acetylation (N-term) | 26079 | PPTG_12671 | Phytophthora parasitica INRA-310 proteasome subunit alpha type-2 (237 aa) |
| 886 | 1653 | PPTG_12747 | 171,39 | 20 | 13 | 20 | 18 | 3,51E+06 | 7,73E+07 | 5,16E+07 | 5 | 5 | 5 | 10 | 11 | Oxidation (M); Carbamidomethylation | 50806 | PPTG_12747 | Phytophthora parasitica INRA-310 hypothetical protein (502 aa) |
| 425 | 319 | PPTG_06713 | 171,35 | 42 | 18 | 41 | 40 | 4,36E+06 | 2,59E+07 | 5,69E+07 | 18 | 16 | 7 | 19 | 22 | Oxidation (M) | 58813 | PPTG_06713 | Phytophthora parasitica INRA-310 catalase (523 aa) |
| 460 | 452 | PPTG_05211 | 171,23 | 50 | 9 | 36 | 37 | 2,30E+06 | 2,32E+07 | 3,65E+07 | 25 | 24 | 4 | 18 | 23 | Oxidation (M); Carbamidomethylation | 58653 | PPTG_05211 | Phytophthora parasitica INRA-310 T-complex protein 1 theta subunit (546 aa) |
| 369 | 389 | PPTG_05481 | 171,11 | 39 | 11 | 34 | 32 | 1,80E+06 | 1,78E+07 | 2,48E+07 | 25 | 25 | 7 | 25 | 23 | Oxidation (M); Deamidation (NQ) | 86036 | PPTG_05481 | Phytophthora parasitica INRA-310 Cullin family protein (749 aa) |
| 338 | 929 | PPTG_17323 | 170,96 | 68 | 40 | 68 | 68 | 3,08E+07 | 1,61E+08 | 2,71E+08 | 13 | 13 | 10 | 21 | 28 | Oxidation (M); Carbamidomethylation; Deamidation (NQ) | 20156 | PPTG_17323 | Phytophthora parasitica INRA-310 hypothetical protein (178 aa) |
| 234 | 322 | PPTG_08306 | 170,89 | 67 | 20 | 66 | 67 | 1,66E+06 | 5,67E+07 | 7,51E+07 | 17 | 17 | 6 | 34 | 36 | Oxidation (M); Carbamidomethylation; Carbamylation | 33074 | PPTG_08306 | Phytophthora parasitica INRA-310 pyridoxal biosynthesis lyase pdxS (311 aa) |
| 1718 | 2306 | PPTG_04194 | 170,88 | 25 | 11 | 25 | 25 | 1,42E+05 | 5,71E+06 | 5,67E+06 | 4 | 4 | 1 | 6 | 4 | Carbamidomethylation | 35050 | PPTG_04194 | Phytophthora parasitica INRA-310 hypothetical protein (341 aa) |
| 755 | 386 | PPTG_18839 | 170,78 | 26 | 10 | 17 | 26 | 5,05E+05 | 2,85E+06 | 1,52E+07 | 13 | 13 | 4 | 9 | 17 | Oxidation (M); Carbamidomethylation; Dehydration; Acetylation (N-term) | 72372 | PPTG_18839 | Phytophthora parasitica INRA-310 hypothetical protein (689 aa) |
| 438 | 414 | PPTG_00911 | 170,66 | 30 | 13 | 26 | 23 | 1,46E+07 | 9,13E+07 | 1,40E+08 | 19 | 19 | 8 | 19 | 19 | Carbamidomethylation | 75583 | PPTG_00911 | Phytophthora parasitica INRA-310 transketolase (697 aa) |
| 647 | 752 | PPTG_17585 | 170,61 | 45 | 19 | 45 | 45 | 2,33E+06 | 2,93E+07 | 4,09E+07 | 10 | 10 | 4 | 14 | 16 | Oxidation (M); Carbamidomethylation; Deamidation (NQ) | 36668 | PPTG_17585 | Phytophthora parasitica INRA-310 26S proteasome non-ATPase regulatory subunit (357 aa) |
| 391 | 397 | PPTG_02266 | 170,21 | 32 | 11 | 24 | 31 | 1,47E+06 | 8,26E+06 | 3,02E+07 | 23 | 23 | 9 | 18 | 25 | Oxidation (M); Carbamidomethylation | 90006 | PPTG_02266 | Phytophthora parasitica INRA-310 cGMP-dependent protein kinase (811 aa) |
| 548 | 312 | PPTG_17912 | 170,1 | 46 | 5 | 36 | 38 | 5,40E+05 | 1,44E+07 | 2,00E+07 | 18 | 18 | 3 | 16 | 20 | Oxidation (M); Carbamidomethylation | 62220 | PPTG_17912 | Phytophthora parasitica INRA-310 hypothetical protein (563 aa) |
| 784 | 980 | PPTG_10741 | 169,92 | 23 | 6 | 21 | 18 | 5,73E+05 | 2,99E+07 | 2,64E+07 | 10 | 10 | 2 | 15 | 12 | Oxidation (M); Carbamidomethylation; Deamidation (NQ) | 55310 | PPTG_10741 | Phytophthora parasitica INRA-310 calcineurin-like phosphoesterase (504 aa) |
| 256 | 475 | PPTG_14168 | 169,91 | 38 | 15 | 28 | 31 | 7,59E+06 | 4,18E+07 | 9,01E+07 | 24 | 24 | 11 | 27 | 33 | Oxidation (M); Carbamidomethylation | 80441 | PPTG_14168 | Phytophthora parasitica INRA-310 eukaryotic translation initiation factor 3 subunit (697 aa) |
| 692 | 372 | PPTG_09054 | 169,82 | 41 | 15 | 34 | 41 | 2,40E+06 | 1,52E+07 | 4,24E+07 | 12 | 12 | 5 | 12 | 15 | Oxidation (M); Carbamidomethylation; Dehydration; Deamidation (NQ) | 45556 | PPTG_09054 | Phytophthora parasitica INRA-310 eukaryotic peptide chain release factor subunit 1 (408 aa) |
| 320 | 337 | PPTG_07024 | 169,82 | 66 | 48 | 48 | 60 | 3,03E+07 | 8,33E+07 | 4,76E+08 | 13 | 13 | 14 | 15 | 32 | Oxidation (M); Dehydration; Deamidation (NQ); Carbamylation | 24885 | PPTG_07024 | Phytophthora parasitica INRA-310 ATP synthase subunit O (232 aa) |
| 415 | 325 | PPTG_10267 | 169,73 | 37 | 30 | 35 | 37 | 9,79E+06 | 7,15E+07 | 9,25E+07 | 15 | 15 | 11 | 20 | 18 | Oxidation (M); Carbamidomethylation; Deamidation (NQ) | 50223 | PPTG_10267 | Phytophthora parasitica INRA-310 hypothetical protein (447 aa) |
| 607 | 427 | PPTG_04806 | 169,61 | 51 | 29 | 42 | 47 | 1,13E+06 | 8,13E+06 | 2,58E+07 | 14 | 13 | 7 | 14 | 15 | Carbamidomethylation; Acetylation (N-term) | 36814 | PPTG_04806 | Phytophthora parasitica INRA-310 serine/threonine-protein phosphatase PP1 (324 aa) |
| 672 | 314 | PPTG_06393 | 169,51 | 31 | 7 | 25 | 31 | 1,04E+06 | 5,90E+06 | 1,57E+07 | 15 | 15 | 4 | 12 | 17 | Carbamidomethylation | 82310 | PPTG_06393 | Phytophthora parasitica INRA-310 hypothetical protein (755 aa) |
| 683 | 1431 | PPTG_02872 | 169,36 | 42 | 15 | 39 | 39 | 1,53E+06 | 3,59E+07 | 2,88E+07 | 8 | 8 | 6 | 15 | 12 | Oxidation (M); Carbamidomethylation | 28921 | PPTG_02872 | Phytophthora parasitica INRA-310 hypothetical protein (267 aa) |
| 432 | 370 | PPTG_17688 | 169,35 | 33 | 7 | 28 | 32 | 1,04E+06 | 1,59E+07 | 2,46E+07 | 19 | 19 | 4 | 20 | 23 | Oxidation (M); Carbamidomethylation; Amidation | 80276 | PPTG_17688 | Phytophthora parasitica INRA-310 hypothetical protein (753 aa) |
| 440 | 343 | PPTG_14636 | 169,3 | 40 | 12 | 32 | 40 | 2,52E+06 | 3,38E+07 | 3,24E+07 | 17 | 17 | 7 | 18 | 21 | Oxidation (M); Carbamidomethylation | 59912 | PPTG_14636 | Phytophthora parasitica INRA-310 propionyl-CoA carboxylase beta chain mitochondrial (543 aa) |
| 582 | 324 | PPTG_05533 | 169,22 | 37 | 15 | 31 | 37 | 1,39E+06 | 9,41E+06 | 2,35E+07 | 14 | 13 | 6 | 13 | 18 | Oxidation (M); Carbamidomethylation | 54964 | PPTG_05533 | Phytophthora parasitica INRA-310 multiple inositol polyphosphate phosphatase (489 aa) |
| 473 | 961 | PPTG_15376 | 169,22 | 31 | 16 | 25 | 29 | 4,69E+05 | | 5,04E+05 | 8 | 2 | 7 | 17 | 20 | Oxidation (M); Carbamidomethylation; Deamidation (NQ); Amidation; Carbamylation | 41733 | PPTG_15376 | Phytophthora parasitica INRA-310 actin-2 (376 aa) |
| 426 | 555 | PPTG_09844 | 169,04 | 31 | 15 | 30 | 29 | 6,56E+05 | 1,73E+07 | 1,32E+07 | 12 | 9 | 5 | 22 | 21 | Oxidation (M); Carbamidomethylation; Deamidation (NQ) | 54758 | PPTG_09844 | Phytophthora parasitica INRA-310 1 3-beta-glucanosyltransferase (503 aa) |
| 44 | 658 | PPTG_04186 | 169,03 | 47 | 46 | 47 | 46 | 3,36E+08 | 1,25E+09 | 3,19E+09 | 13 | 13 | 35 | 50 | 64 | Oxidation (M); Dehydration; Deamidation (NQ); Acetylation (N-term); Carbamylation | 27692 | PPTG_04186 | Phytophthora parasitica INRA-310 60S ribosomal protein L7 (245 aa) |
| 489 | 825 | PPTG_10299 | 168,75 | 43 | 40 | 40 | 43 | 1,31E+07 | 1,40E+08 | 1,06E+08 | 12 | 12 | 10 | 18 | 15 | Deamidation (NQ) | 26671 | PPTG_10299 | Phytophthora parasitica INRA-310 hypothetical protein (244 aa) |
| 401 | 347 | PPTG_18768 | 168,52 | 52 | 32 | 49 | 49 | 4,70E+06 | 2,43E+07 | 3,36E+07 | 18 | 18 | 12 | 19 | 20 | Oxidation (M); Carbamidomethylation; Deamidation (NQ); Acetylation (N-term) | 50725 | PPTG_18768 | Phytophthora parasitica INRA-310 AP-4 complex subunit mu-1 (448 aa) |
| 416 | 384 | PPTG_04013 | 168,43 | 26 | 10 | 16 | 25 | 2,44E+06 | 1,24E+07 | 3,59E+07 | 21 | 21 | 9 | 16 | 24 | Oxidation (M); Carbamidomethylation; Deamidation (NQ) | 111112 | PPTG_04013 | Phytophthora parasitica INRA-310 protein transporter Sec24 (1025 aa) |
| 540 | 392 | PPTG_05029 | 168,41 | 68 | 26 | 61 | 61 | 6,64E+05 | 8,30E+06 | 1,01E+07 | 14 | 14 | 6 | 16 | 18 | Oxidation (M); Carbamidomethylation; Dehydration; Deamidation (NQ) | 33461 | PPTG_05029 | Phytophthora parasitica INRA-310 YjeF (318 aa) |
| 456 | 399 | PPTG_09645 | 168,38 | 42 | 4 | 38 | 35 | 2,56E+06 | 3,17E+07 | 4,66E+07 | 18 | 18 | 2 | 22 | 21 | Oxidation (M); Carbamidomethylation; Deamidation (NQ) | 65008 | PPTG_09645 | Phytophthora parasitica INRA-310 triosephosphate isomerase/glyceraldehyde-3-phosphate dehydrogenase (603 aa) |
| 233 | 790 | PPTG_17956 | 168,34 | 47 | 34 | 44 | 47 | 1,73E+08 | 7,02E+08 | 1,73E+09 | 12 | 12 | 12 | 25 | 39 | Oxidation (M); Carbamidomethylation; Dehydration; Deamidation (NQ); Carbamylation | 23696 | PPTG_17956 | Phytophthora parasitica INRA-310 60S ribosomal protein L10 (209 aa) |
| 542 | 593 | PPTG_18089 | 168,25 | 40 | 11 | 34 | 40 | 1,92E+06 | 1,74E+07 | 3,16E+07 | 12 | 12 | 4 | 15 | 21 | Oxidation (M); Carbamidomethylation; Dehydration; Deamidation (NQ) | 46886 | PPTG_18089 | Phytophthora parasitica INRA-310 dihydrolipoyllysine-residue acetyltransferase component-pyruvate dehydrogenase complex (438 aa) |
| 526 | 383 | PPTG_00539 | 168,17 | 32 | 17 | 29 | 27 | 2,39E+06 | 1,61E+07 | 2,20E+07 | 15 | 15 | 8 | 16 | 16 | Oxidation (M); Carbamidomethylation; Deamidation (NQ) | 70860 | PPTG_00539 | Phytophthora parasitica INRA-310 choline/Carnitine O-acyltransferase (641 aa) |
| 704 | 927 | PPTG_13332 | 168,15 | 23 | 10 | 21 | 21 | 2,83E+06 | 1,76E+07 | 3,79E+07 | 9 | 9 | 6 | 12 | 14 | Carbamidomethylation; Deamidation (NQ) | 58453 | PPTG_13332 | Phytophthora parasitica INRA-310 cysteine protease family C01A (536 aa) |
| 691 | 378 | PPTG_00044 | 168,14 | 41 | 7 | 36 | 36 | 8,83E+04 | 8,31E+06 | 1,10E+07 | 14 | 13 | 3 | 15 | 14 | | 51805 | PPTG_00044 | Phytophthora parasitica INRA-310 calcium/calmodulin-dependent protein kinase II (472 aa) |
| 450 | 464 | PPTG_16444 | 168,11 | 66 | 35 | 64 | 50 | 8,86E+06 | 7,34E+07 | 9,49E+07 | 13 | 13 | 9 | 20 | 17 | Deamidation (NQ) | 29336 | PPTG_16444 | Phytophthora parasitica INRA-310 2 3-bisphosphoglycerate-dependent phosphoglycerate mutase (261 aa) |
| 967 | 395 | PPTG_01765 | 167,96 | 24 | 6 | 4 | 24 | 3,53E+05 | 3,87E+05 | 1,09E+07 | 13 | 12 | 4 | 3 | 16 | Oxidation (M); Carbamidomethylation; Dehydration; Deamidation (NQ) | 85542 | PPTG_01765 | Phytophthora parasitica INRA-310 transmembrane protein (773 aa) |
| 462 | 346 | PPTG_07933 | 167,92 | 48 | 22 | 41 | 45 | 2,06E+06 | 2,66E+07 | 3,61E+07 | 16 | 11 | 6 | 19 | 20 | Oxidation (M); Carbamidomethylation | 39712 | PPTG_07933 | Phytophthora parasitica INRA-310 voltage-gated potassium channel subunit beta (356 aa) |
| 786 | 328 | PPTG_11925 | 167,91 | 37 | 12 | 19 | 35 | 1,52E+06 | 4,27E+06 | 2,16E+07 | 15 | 15 | 6 | 8 | 15 | Carbamidomethylation; Deamidation (NQ) | 62173 | PPTG_11925 | Phytophthora parasitica INRA-310 hypothetical protein (565 aa) |
| 718 | 486 | PPTG_00307 | 167,69 | 39 | 8 | 30 | 34 | 4,29E+05 | 7,76E+06 | 8,80E+06 | 13 | 8 | 3 | 14 | 14 | Oxidation (M); Carbamidomethylation; Acetylation (N-term) | 47894 | PPTG_00307 | Phytophthora parasitica INRA-310 hypothetical protein (438 aa) |
| 660 | 642 | PPTG_03063 | 167,66 | 36 | 15 | 31 | 31 | 1,03E+06 | 3,33E+07 | 2,37E+07 | 11 | 10 | 4 | 18 | 12 | Oxidation (M); Carbamidomethylation; Dehydration | 44890 | PPTG_03063 | Phytophthora parasitica INRA-310 hypothetical protein (411 aa) |
| 536 | 356 | PPTG_00731 | 167,65 | 47 | 7 | 45 | 42 | 4,36E+05 | 1,08E+07 | 8,07E+06 | 14 | 14 | 3 | 19 | 18 | Oxidation (M); Carbamidomethylation; Deamidation (NQ) | 54165 | PPTG_00731 | Phytophthora parasitica INRA-310 succinate semialdehyde dehydrogenase (508 aa) |
| 455 | 358 | PPTG_20079 | 167,63 | 36 | 17 | 27 | 36 | 9,04E+05 | 7,88E+06 | 1,30E+07 | 17 | 6 | 8 | 16 | 21 | Carbamidomethylation; Dehydration; Deamidation (NQ) | 67133 | PPTG_20079 | Phytophthora parasitica INRA-310 hypothetical protein (604 aa) |
| 507 | 432 | PPTG_11779 | 167,52 | 35 | 10 | 31 | 25 | 9,90E+05 | 1,37E+07 | 1,94E+07 | 18 | 18 | 6 | 20 | 16 | Oxidation (M); Carbamidomethylation; Acetylation (N-term) | 71183 | PPTG_11779 | Phytophthora parasitica INRA-310 choline/Carnitine O-acyltransferase (634 aa) |
| 483 | 949 | PPTG_15106 | 167,46 | 76 | 25 | 76 | 68 | 5,20E+06 | 1,88E+08 | 1,30E+08 | 10 | 10 | 6 | 21 | 17 | Oxidation (M); Carbamidomethylation; Deamidation (NQ) | 18865 | PPTG_15106 | Phytophthora parasitica INRA-310 hypothetical protein (168 aa) |
| 353 | 537 | PPTG_03934 | 167,38 | 45 | 25 | 45 | 43 | 1,09E+07 | 1,48E+08 | 1,54E+08 | 14 | 14 | 9 | 25 | 23 | Oxidation (M); Carbamidomethylation; Deamidation (NQ); Acetylation (N-term); Carbamylation | 37146 | PPTG_03934 | Phytophthora parasitica INRA-310 annexin (Annexin) family (335 aa) |
| 778 | 422 | PPTG_06662 | 167,34 | 46 | 10 | 41 | 46 | 2,17E+05 | 1,11E+07 | 1,41E+07 | 13 | 13 | 2 | 12 | 15 | Carbamidomethylation | 43707 | PPTG_06662 | Phytophthora parasitica INRA-310 histidinol dehydrogenase (422 aa) |
| 642 | 509 | PPTG_11134 | 167,13 | 25 | 8 | 16 | 24 | 8,50E+05 | 6,21E+06 | 1,36E+07 | 17 | 17 | 5 | 11 | 18 | Oxidation (M); Carbamidomethylation | 100807 | PPTG_11134 | Phytophthora parasitica INRA-310 hypothetical protein (917 aa) |
| 579 | 488 | PPTG_08520 | 167,07 | 23 | 4 | 16 | 13 | 9,32E+05 | 9,20E+06 | 1,27E+07 | 22 | 22 | 5 | 18 | 14 | Carbamidomethylation | 132299 | PPTG_08520 | Phytophthora parasitica INRA-310 pre-mRNA-splicing factor RSE1 (1198 aa) |
| 970 | 693 | PPTG_07827 | 166,99 | 34 | 9 | 24 | 34 | 1,04E+06 | 3,88E+06 | 2,21E+07 | 10 | 10 | 3 | 8 | 12 | Carbamidomethylation; Deamidation (NQ) | 47023 | PPTG_07827 | Phytophthora parasitica INRA-310 lectin (415 aa) |
| 547 | 502 | PPTG_19695 | 166,99 | 24 | 16 | 22 | 24 | 3,09E+06 | 7,16E+07 | 5,85E+07 | 13 | 13 | 7 | 16 | 16 | Carbamidomethylation; Acetylation (N-term) | 74414 | PPTG_19695 | Phytophthora parasitica INRA-310 hypothetical protein (685 aa) |
| 422 | 454 | PPTG_09068 | 166,94 | 5 | 1 | 4 | 4 | 2,80E+05 | 5,52E+06 | 6,09E+06 | 28 | 28 | 5 | 22 | 21 | Carbamidomethylation | 766862 | PPTG_09068 | Phytophthora parasitica INRA-310 transmembrane protein (7034 aa) |
| 684 | 1365 | PPTG_08708 | 166,74 | 33 | 3 | 33 | 25 | 1,10E+06 | 5,73E+07 | 3,56E+07 | 6 | 6 | 1 | 18 | 14 | Oxidation (M); Carbamidomethylation; Deamidation (NQ); Acetylation (N-term); Carbamylation | 40238 | PPTG_08708 | Phytophthora parasitica INRA-310 hypothetical protein (385 aa) |
| 467 | 532 | PPTG_19359 | 166,51 | 15 | 4 | 13 | 9 | 1,01E+06 | 1,17E+07 | 1,34E+07 | 23 | 23 | 7 | 20 | 17 | Carbamidomethylation; Amidation | 204707 | PPTG_19359 | Phytophthora parasitica INRA-310 DNA-directed RNA polymerase II largest subunit (1860 aa) |
| 76 | 636 | PPTG_02269 | 166,46 | 63 | 47 | 61 | 63 | 2,28E+08 | 9,00E+08 | 2,28E+09 | 12 | 12 | 27 | 38 | 59 | Carbamidomethylation; Dehydration; Deamidation (NQ); Carbamylation | 21480 | PPTG_02269 | Phytophthora parasitica INRA-310 60S ribosomal protein (193 aa) |
| 703 | 926 | PPTG_14777 | 166,45 | 11 | 6 | 11 | 11 | 3,54E+06 | 4,74E+07 | 6,56E+07 | 8 | 8 | 6 | 13 | 13 | Oxidation (M); Carbamidomethylation; Deamidation (NQ) | 109268 | PPTG_14777 | Phytophthora parasitica INRA-310 hypothetical protein (1082 aa) |
| 399 | 411 | PPTG_05019 | 166,32 | 48 | 24 | 48 | 46 | 1,91E+06 | 2,46E+07 | 2,76E+07 | 16 | 16 | 9 | 22 | 20 | Oxidation (M); Carbamidomethylation; Dehydration; Deamidation (NQ) | 43792 | PPTG_05019 | Phytophthora parasitica INRA-310 isovaleryl-CoA dehydrogenase (400 aa) |
| 525 | 401 | PPTG_10204 | 166,16 | 47 | 22 | 26 | 47 | 8,03E+06 | 1,89E+07 | 1,27E+08 | 14 | 14 | 8 | 11 | 22 | Oxidation (M); Carbamidomethylation; Deamidation (NQ) | 40499 | PPTG_10204 | Phytophthora parasitica INRA-310 ribose-phosphate pyrophosphokinase (379 aa) |
| 757 | 807 | PPTG_03523 | 166 | 59 | 28 | 53 | 59 | 3,84E+06 | 3,96E+07 | 5,37E+07 | 10 | 10 | 5 | 11 | 14 | Oxidation (M) | 25774 | PPTG_03523 | Phytophthora parasitica INRA-310 GrxB family glutaredoxin (228 aa) |
| 476 | 413 | PPTG_13439 | 165,96 | 42 | 17 | 29 | 42 | 6,28E+06 | 3,62E+07 | 5,90E+07 | 19 | 19 | 7 | 16 | 21 | Oxidation (M); Carbamidomethylation; Deamidation (NQ) | 55043 | PPTG_13439 | Phytophthora parasitica INRA-310 6-phosphofructokinase (505 aa) |
| 488 | 435 | PPTG_04138 | 165,57 | 54 | 21 | 49 | 45 | 4,36E+06 | 4,37E+07 | 7,51E+07 | 14 | 14 | 6 | 19 | 18 | Oxidation (M); Carbamidomethylation; Deamidation (NQ) | 32862 | PPTG_04138 | Phytophthora parasitica INRA-310 rRNA 2'-O-methyltransferase fibrillarin 2 (319 aa) |
| 643 | 533 | PPTG_06991 | 165,56 | 35 | 8 | 27 | 35 | 8,76E+05 | 9,67E+06 | 1,99E+07 | 14 | 14 | 4 | 13 | 17 | Oxidation (M); Carbamidomethylation; Deamidation (NQ); Acetylation (N-term) | 63440 | PPTG_06991 | Phytophthora parasitica INRA-310 inorganic pyrophosphatase (558 aa) |
| 496 | 554 | PPTG_01960 | 165,39 | 84 | 57 | 77 | 78 | 8,37E+06 | 7,77E+07 | 1,04E+08 | 12 | 12 | 8 | 17 | 18 | Oxidation (M); Carbamidomethylation; Deamidation (NQ) | 18776 | PPTG_01960 | Phytophthora parasitica INRA-310 hypothetical protein (168 aa) |
| 448 | 368 | PPTG_12168 | 165,39 | 59 | 26 | 45 | 55 | 7,24E+06 | 3,54E+07 | 6,40E+07 | 16 | 16 | 8 | 19 | 19 | Oxidation (M); Carbamidomethylation | 45216 | PPTG_12168 | Phytophthora parasitica INRA-310 elongation factor Tu (417 aa) |
| 714 | 947 | PPTG_04619 | 165,34 | 49 | 32 | 35 | 46 | 3,09E+06 | 5,78E+07 | 4,54E+07 | 10 | 10 | 7 | 14 | 11 | Carbamidomethylation; Deamidation (NQ) | 29518 | PPTG_04619 | Phytophthora parasitica INRA-310 hypothetical protein (277 aa) |
| 766 | 472 | PPTG_04276 | 165,29 | 22 | 12 | 14 | 22 | 2,27E+06 | 2,13E+06 | 3,40E+07 | 11 | 11 | 6 | 7 | 16 | Oxidation (M); Carbamidomethylation; Dehydration; Deamidation (NQ); Amidation | 83674 | PPTG_04276 | Phytophthora parasitica INRA-310 hypothetical protein (764 aa) |
| 591 | 408 | PPTG_09278 | 165,16 | 21 | 4 | 17 | 18 | 4,07E+05 | 7,76E+06 | 9,08E+06 | 18 | 18 | 4 | 16 | 17 | Carbamidomethylation | 137275 | PPTG_09278 | Phytophthora parasitica INRA-310 WD repeat protein 35 (1219 aa) |
| 569 | 437 | PPTG_06297 | 165,11 | 50 | 28 | 44 | 43 | 5,51E+06 | 4,80E+07 | 7,19E+07 | 15 | 13 | 8 | 14 | 16 | Oxidation (M); Carbamidomethylation | 38123 | PPTG_06297 | Phytophthora parasitica INRA-310 mannitol dehydrogenase (355 aa) |
| 605 | 357 | PPTG_19366 | 165,1 | 39 | 13 | 32 | 38 | 7,40E+05 | 1,19E+07 | 1,04E+07 | 14 | 10 | 6 | 17 | 13 | Oxidation (M); Carbamidomethylation; Deamidation (NQ) | 58917 | PPTG_19366 | Phytophthora parasitica INRA-310 NADP-dependent malic enzyme (539 aa) |
| 494 | 442 | PPTG_04879 | 165,08 | 79 | 17 | 79 | 76 | 2,11E+06 | 3,07E+07 | 3,97E+07 | 14 | 14 | 4 | 20 | 19 | Oxidation (M); Deamidation (NQ) | 25363 | PPTG_04879 | Phytophthora parasitica INRA-310 hypothetical protein (230 aa) |
| 478 | 439 | PPTG_06519 | 164,98 | 36 | 12 | 31 | 34 | 1,03E+06 | 9,78E+06 | 1,21E+07 | 20 | 20 | 9 | 16 | 19 | Carbamidomethylation | 71872 | PPTG_06519 | Phytophthora parasitica INRA-310 metalloprotease family M01 (639 aa) |
| 595 | 391 | PPTG_08490 | 164,98 | 12 | 2 | 9 | 10 | 8,98E+05 | 3,98E+06 | 1,12E+07 | 17 | 17 | 5 | 14 | 17 | Oxidation (M); Carbamidomethylation | 162726 | PPTG_08490 | Phytophthora parasitica INRA-310 myosin-like protein (1465 aa) |
| 334 | 467 | PPTG_01365 | 164,93 | 47 | 24 | 33 | 45 | 2,70E+06 | 1,83E+07 | 3,13E+07 | 23 | 23 | 12 | 21 | 26 | Oxidation (M); Carbamidomethylation | 58268 | PPTG_01365 | Phytophthora parasitica INRA-310 T-complex protein 1 subunit gamma (531 aa) |
| 906 | 675 | PPTG_06818 | 164,9 | 58 | 8 | 42 | 58 | 3,54E+05 | 5,89E+06 | 1,51E+07 | 12 | 12 | 2 | 9 | 14 | Oxidation (M); Carbamidomethylation | 23505 | PPTG_06818 | Phytophthora parasitica INRA-310 methyltransferase (207 aa) |
| 1288 | 1340 | PPTG_00479 | 164,84 | 28 | 0 | 28 | 18 | | 6,52E+06 | 5,34E+06 | 6 | 6 | 0 | 9 | 7 | Carbamidomethylation | 41494 | PPTG_00479 | Phytophthora parasitica INRA-310 cellulose binding elicitor lectin (CBEL) (385 aa) |
| 499 | 351 | PPTG_16449 | 164,68 | 39 | 17 | 29 | 39 | 2,64E+06 | 2,13E+07 | 3,07E+07 | 17 | 17 | 8 | 15 | 19 | Oxidation (M); Carbamidomethylation | 57483 | PPTG_16449 | Phytophthora parasitica INRA-310 pyruvate kinase (523 aa) |
| 675 | 904 | PPTG_17103 | 164,64 | 34 | 18 | 32 | 30 | 1,72E+07 | 7,09E+07 | 1,30E+08 | 12 | 12 | 6 | 14 | 13 | Oxidation (M) | 39530 | PPTG_17103 | Phytophthora parasitica INRA-310 urate oxidase (362 aa) |
| 727 | 1385 | PPTG_15650 | 164,61 | 49 | 23 | 39 | 49 | 1,32E+07 | 4,31E+07 | 9,26E+07 | 6 | 6 | 6 | 8 | 17 | Oxidation (M); Carbamidomethylation; Deamidation (NQ); Amidation | 20226 | PPTG_15650 | Phytophthora parasitica INRA-310 hypothetical protein (181 aa) |
| 857 | 522 | PPTG_02615 | 164,36 | 23 | 6 | 7 | 23 | 1,07E+06 | 1,58E+06 | 2,00E+07 | 14 | 14 | 5 | 5 | 17 | Carbamidomethylation; Dehydration; Deamidation (NQ) | 90875 | PPTG_02615 | Phytophthora parasitica INRA-310 hypothetical protein (812 aa) |
| 505 | 438 | PPTG_19557 | 164,32 | 34 | 6 | 30 | 30 | 1,81E+06 | 2,10E+07 | 3,06E+07 | 18 | 18 | 4 | 21 | 17 | Oxidation (M); Carbamidomethylation; Dehydration; Deamidation (NQ) | 81805 | PPTG_19557 | Phytophthora parasitica INRA-310 methionyl-tRNA synthetase (728 aa) |
| 725 | 628 | PPTG_06783 | 164,07 | 44 | 14 | 36 | 44 | 3,35E+05 | 1,36E+08 | 1,76E+08 | 10 | 10 | 3 | 12 | 16 | Carbamidomethylation; Carbamylation | 38131 | PPTG_06783 | Phytophthora parasitica INRA-310 hypothetical protein (381 aa) |
| 417 | 342 | PPTG_14256 | 164,06 | 37 | 12 | 30 | 35 | 2,15E+06 | 2,41E+07 | 2,27E+07 | 16 | 15 | 7 | 21 | 21 | Oxidation (M); Carbamidomethylation; Deamidation (NQ); Amidation | 70483 | PPTG_14256 | Phytophthora parasitica INRA-310 hypothetical protein (650 aa) |
| 491 | 550 | PPTG_10870 | 163,97 | 40 | 17 | 30 | 31 | 2,12E+06 | 2,58E+07 | 2,98E+07 | 20 | 20 | 9 | 15 | 19 | Oxidation (M); Carbamidomethylation; Deamidation (NQ) | 68971 | PPTG_10870 | Phytophthora parasitica INRA-310 phenylalanyl-tRNA synthetase beta subunit (615 aa) |
| 697 | 962 | PPTG_16154 | 163,85 | 45 | 14 | 35 | 39 | 2,09E+06 | 4,06E+07 | 3,93E+07 | 11 | 11 | 3 | 15 | 14 | Oxidation (M); Carbamidomethylation; Deamidation (NQ) | 32013 | PPTG_16154 | Phytophthora parasitica INRA-310 proteasome subunit beta type-7 (296 aa) |
| 537 | 423 | PPTG_09863 | 163,75 | 18 | 7 | 4 | 18 | 1,67E+06 | 5,57E+05 | 2,03E+07 | 18 | 15 | 9 | 5 | 26 | Oxidation (M); Carbamidomethylation; Deamidation (NQ) | 150517 | PPTG_09863 | Phytophthora parasitica INRA-310 ATP-binding Cassette (ABC) superfamily (1353 aa) |
| 770 | 637 | PPTG_02668 | 163,75 | 33 | 13 | 26 | 27 | 4,94E+05 | 7,23E+06 | 6,00E+06 | 14 | 14 | 6 | 11 | 12 | Deamidation (NQ); Carbamylation | 58133 | PPTG_02668 | Phytophthora parasitica INRA-310 hypothetical protein (519 aa) |
| 754 | 1095 | PPTG_04900 | 163,75 | 58 | 29 | 29 | 58 | 6,45E+06 | 9,89E+06 | 8,06E+07 | 10 | 10 | 6 | 6 | 18 | Oxidation (M); Carbamidomethylation; Dehydration; Deamidation (NQ) | 22310 | PPTG_04900 | Phytophthora parasitica INRA-310 Ras protein let-60 (197 aa) |
| 662 | 507 | PPTG_02607 | 163,71 | 5 | 1 | 0 | 5 | 1,38E+05 | 1,52E+04 | 8,56E+06 | 28 | 28 | 4 | 1 | 28 | Carbamidomethylation | 737778 | PPTG_02607 | Phytophthora parasitica INRA-310 hypothetical protein (6717 aa) |
| 60 | 1116 | PPTG_01105 | 163,66 | 70 | 69 | 67 | 67 | 9,60E+07 | 5,35E+08 | 1,75E+09 | 10 | 10 | 21 | 41 | 74 | Oxidation (M); Carbamidomethylation; Dehydration; Deamidation (NQ); Carbamylation | 14939 | PPTG_01105 | Phytophthora parasitica INRA-310 60S ribosomal protein L23 (142 aa) |

| | | | | | | | | | | | | | | | | | | | |
|---|---|---|---|---|---|---|---|---|---|---|---|---|---|---|---|---|---|---|---|
| 545 | 367 | PPTG_00178 | 163,61 | 31 | 16 | 12 | 31 | 2,98E+06 | 3,78E+06 | 3,65E+07 | 19 | 19 | 10 | 8 | 21 | Carbamidomethylation; Deamidation (NQ) | 101196 | PPTG_00178 | Phytophthora parasitica INRA-310 adenylate kinase (926 aa) |
| 1060 | 824 | PPTG_11108 | 163,5 | 12 | 0 | 12 | 8 | | 5,76E+06 | 8,26E+06 | 10 | 10 | 0 | 11 | 10 | Oxidation (M); Carbamidomethylation | 163014 | PPTG_11108 | Phytophthora parasitica INRA-310 hypothetical protein (1535 aa) |
| 555 | 336 | PPTG_00091 | 163,38 | 48 | 20 | 30 | 48 | 1,11E+06 | 3,30E+06 | 2,82E+07 | 15 | 13 | 6 | 11 | 22 | Oxidation (M); Carbamidomethylation | 45707 | PPTG_00091 | Phytophthora parasitica INRA-310 coproporphyrinogen III oxidase (403 aa) |
| 765 | 380 | PPTG_12649 | 163,36 | 45 | 11 | 5 | 45 | 6,95E+05 | 8,55E+05 | 1,57E+07 | 17 | 14 | 5 | 3 | 21 | Oxidation (M); Carbamidomethylation; Deamidation (NQ); Acetylation (N-term) | 53392 | PPTG_12649 | Phytophthora parasitica INRA-310 syntaxin-binding protein (475 aa) |
| 523 | 591 | PPTG_01826 | 163,35 | 59 | 43 | 49 | 55 | 2,97E+06 | 7,63E+06 | 3,84E+07 | 11 | 9 | 11 | 13 | 17 | Oxidation (M); Dehydration | 24173 | PPTG_01826 | Phytophthora parasitica INRA-310 Ras-like protein Rab-11B (219 aa) |
| 387 | 596 | PPTG_15085 | 163,32 | 30 | 6 | 26 | 27 | 1,08E+06 | 9,82E+06 | 2,82E+07 | 27 | 27 | 6 | 22 | 25 | Oxidation (M); Carbamidomethylation; Deamidation (NQ) | 106244 | PPTG_15085 | Phytophthora parasitica INRA-310 hypothetical protein (980 aa) |
| 612 | 487 | PPTG_08927 | 163,29 | 41 | 20 | 34 | 41 | 1,86E+06 | 4,13E+06 | 3,00E+07 | 14 | 14 | 6 | 11 | 19 | Oxidation (M); Carbamidomethylation | 42189 | PPTG_08927 | Phytophthora parasitica INRA-310 ribose-phosphate pyrophosphokinase (392 aa) |
| 611 | 448 | PPTG_05778 | 163,24 | 43 | 14 | 43 | 30 | 8,74E+05 | 2,20E+07 | 2,37E+07 | 14 | 14 | 4 | 17 | 15 | Oxidation (M); Carbamidomethylation; Deamidation (NQ) | 44381 | PPTG_05778 | Phytophthora parasitica INRA-310 short-chain specific acyl-CoA dehydrogenase (414 aa) |
| 428 | 373 | PPTG_00719 | 163,17 | 19 | 5 | 14 | 18 | 1,02E+06 | 7,02E+06 | 2,00E+07 | 20 | 18 | 6 | 16 | 25 | Oxidation (M); Carbamidomethylation; Acetylation (N-term) | 141690 | PPTG_00719 | Phytophthora parasitica INRA-310 myosin-like protein (1258 aa) |
| 856 | 1258 | PPTG_12003 | 162,95 | 37 | 18 | 34 | 28 | 1,70E+06 | 2,27E+07 | 2,90E+07 | 9 | 9 | 4 | 13 | 10 | Oxidation (M); Carbamidomethylation; Deamidation (NQ) | 31865 | PPTG_12003 | Phytophthora parasitica INRA-310 proteasome subunit beta type-5 (289 aa) |
| 855 | 964 | PPTG_11311 | 162,91 | 45 | 19 | 45 | 23 | 1,10E+06 | 1,01E+08 | 9,06E+07 | 8 | 8 | 4 | 13 | 10 | Oxidation (M); Carbamidomethylation; Deamidation (NQ) | 37943 | PPTG_11311 | Phytophthora parasitica INRA-310 cysteine protease family C01A (356 aa) |
| 855 | 965 | PPTG_11313 | 162,91 | 45 | 19 | 45 | 23 | 1,10E+06 | 1,01E+08 | 9,06E+07 | 8 | 8 | 4 | 13 | 10 | Oxidation (M); Carbamidomethylation; Deamidation (NQ) | 37943 | PPTG_11313 | Phytophthora parasitica INRA-310 cysteine protease family C01A (356 aa) |
| 654 | 553 | PPTG_15540 | 162,91 | 49 | 13 | 47 | 44 | 5,83E+05 | 1,42E+07 | 1,93E+07 | 13 | 13 | 3 | 15 | 16 | Carbamidomethylation; Deamidation (NQ) | 39349 | PPTG_15540 | Phytophthora parasitica INRA-310 dihydroorotase (359 aa) |
| 557 | 433 | PPTG_15776 | 162,78 | 35 | 19 | 21 | 35 | 3,02E+06 | 7,30E+06 | 4,36E+07 | 15 | 15 | 9 | 9 | 21 | Carbamidomethylation | 53946 | PPTG_15776 | Phytophthora parasitica INRA-310 NADH-ubiquinone oxidoreductase (481 aa) |
| 393 | 473 | PPTG_00961 | 162,76 | 40 | 19 | 35 | 40 | 2,81E+06 | 2,39E+07 | 4,27E+07 | 14 | 14 | 6 | 21 | 25 | Oxidation (M); Carbamidomethylation | 48254 | PPTG_00961 | Phytophthora parasitica INRA-310 26S protease regulatory subunit 6A (434 aa) |
| 645 | 735 | PPTG_13432 | 162,61 | 40 | 20 | 40 | 40 | 1,75E+06 | 3,94E+07 | 3,55E+07 | 9 | 9 | 6 | 13 | 15 | Oxidation (M); Carbamidomethylation; Deamidation (NQ) | 38618 | PPTG_13432 | Phytophthora parasitica INRA-310 hypothetical protein (365 aa) |
| 209 | 736 | PPTG_04728 | 162,55 | 52 | 36 | 43 | 50 | 1,54E+08 | 6,22E+08 | 1,71E+09 | 13 | 13 | 17 | 30 | 34 | Oxidation (M); Dehydration; Deamidation (NQ); Acetylation (N-term); Carbamylation | 23034 | PPTG_04728 | Phytophthora parasitica INRA-310 50S ribosomal protein L13e (206 aa) |
| 842 | 777 | PPTG_09333 | 162,33 | 53 | 40 | 39 | 53 | 2,85E+06 | 6,04E+06 | 3,55E+07 | 9 | 8 | 6 | 7 | 14 | Oxidation (M); Acetylation (N-term) | 22550 | PPTG_09333 | Phytophthora parasitica INRA-310 Rab18 family GTPase (203 aa) |
| 1139 | 612 | PPTG_12682 | 162,32 | 22 | 2 | 22 | 8 0 | | 5,36E+06 | 1,04E+06 | 11 | 9 | 1 | 12 | 6 | Oxidation (M) | 84668 | PPTG_12682 | Phytophthora parasitica INRA-310 beta glucosidase (776 aa) |
| 794 | 410 | PPTG_02166 | 162,16 | 19 | 6 | 4 | 19 | 7,68E+05 | 2,24E+05 | 1,67E+07 | 15 | 14 | 5 | 3 | 20 | Oxidation (M); Carbamidomethylation; Deamidation (NQ) | 108446 | PPTG_02166 | Phytophthora parasitica INRA-310 anoctamin-like protein (935 aa) |
| 640 | 390 | PPTG_08089 | 162,11 | 59 | 10 | 42 | 52 | 9,23E+05 | 1,82E+07 | 2,23E+07 | 14 | 14 | 3 | 16 | 15 | Oxidation (M); Carbamidomethylation | 36984 | PPTG_08089 | Phytophthora parasitica INRA-310 carbonic anhydrase (325 aa) |
| 810 | 1182 | PPTG_09295 | 162,08 | 21 | 13 | 17 | 21 | 1,93E+06 | 9,07E+06 | 2,48E+07 | 7 | 7 | 5 | 10 | 13 | Oxidation (M); Carbamidomethylation; Deamidation (NQ) | 56703 | PPTG_09295 | Phytophthora parasitica INRA-310 hypothetical protein (543 aa) |
| 562 | 363 | PPTG_09326 | 161,79 | 40 | 19 | 28 | 28 | 2,49E+06 | 1,38E+07 | 2,18E+07 | 17 | 17 | 9 | 15 | 14 | Oxidation (M); Carbamidomethylation | 59501 | PPTG_09326 | Phytophthora parasitica INRA-310 tryptophanyl-tRNA synthetase (538 aa) |
| 549 | 493 | PPTG_04562 | 161,75 | 30 | 7 | 25 | 28 | 1,03E+06 | 1,06E+07 | 1,85E+07 | 17 | 17 | 5 | 16 | 18 | Oxidation (M); Carbamidomethylation | 76166 | PPTG_04562 | Phytophthora parasitica INRA-310 glutaminyl-tRNA synthetase (670 aa) |
| 359 | 451 | PPTG_10943 | 161,75 | 46 | 21 | 41 | 41 | 2,57E+06 | 3,38E+07 | 3,89E+07 | 16 | 15 | 7 | 23 | 26 | Oxidation (M); Carbamidomethylation; Deamidation (NQ) | 44188 | PPTG_10943 | Phytophthora parasitica INRA-310 26S protease regulatory subunit 10B like A (395 aa) |
| 423 | 552 | PPTG_14471 | 161,72 | 57 | 29 | 49 | 51 | 4,38E+06 | 6,81E+07 | 6,92E+07 | 14 | 14 | 6 | 21 | 21 | Oxidation (M); Carbamidomethylation; Deamidation (NQ); Acetylation (N-term); Amidation | 29147 | PPTG_14471 | Phytophthora parasitica INRA-310 proteasome subunit alpha type-3 (259 aa) |
| 421 | 444 | PPTG_07011 | 161,66 | 62 | 22 | 56 | 52 | 4,02E+06 | 5,44E+07 | 4,62E+07 | 18 | 14 | 6 | 21 | 22 | Oxidation (M); Carbamidomethylation; Deamidation (NQ) | 35291 | PPTG_07011 | Phytophthora parasitica INRA-310 glutathione S-transferase (316 aa) |
| 594 | 469 | PPTG_05351 | 161,61 | 53 | 25 | 44 | 47 | 2,11E+06 | 2,12E+07 | 3,04E+07 | 12 | 11 | 6 | 16 | 15 | Carbamidomethylation; Deamidation (NQ) | 30983 | PPTG_05351 | Phytophthora parasitica INRA-310 peroxisomal multifunctional enzyme (290 aa) |
| 655 | 641 | PPTG_18320 | 161,52 | 49 | 23 | 49 | 42 | 1,91E+06 | 2,19E+07 | 2,97E+07 | 11 | 11 | 6 | 15 | 13 | Oxidation (M); Carbamidomethylation; Dehydration; Deamidation (NQ) | 30180 | PPTG_18320 | Phytophthora parasitica INRA-310 hypothetical protein (283 aa) |
| 375 | 431 | PPTG_18967 | 161,5 | 35 | 21 | 35 | 34 | 3,76E+06 | 3,66E+07 | 4,29E+07 | 16 | 16 | 9 | 22 | 23 | Oxidation (M); Carbamidomethylation; Dehydration | 52518 | PPTG_18967 | Phytophthora parasitica INRA-310 peptidyl-prolyl cis-trans isomerase (479 aa) |
| 302 | 573 | PPTG_08523 | 161,35 | 68 | 58 | 68 | 66 | 9,53E+06 | 7,61E+07 | 7,95E+07 | 11 | 11 | 15 | 27 | 22 | Oxidation (M); Carbamidomethylation; Deamidation (NQ); Acetylation (N-term) | 22240 | PPTG_08523 | Phytophthora parasitica INRA-310 proteasome subunit beta type-2 (198 aa) |
| 870 | 1118 | PPTG_09291 | 161,27 | 30 | 14 | 11 | 30 | 4,56E+06 | 5,63E+06 | 4,69E+07 | 7 | 7 | 7 | 5 | 14 | Oxidation (M); Carbamidomethylation; Deamidation (NQ) | 44855 | PPTG_09291 | Phytophthora parasitica INRA-310 hypothetical protein (396 aa) |
| 919 | 393 | PPTG_05391 | 161,24 | 23 | 6 | 19 | 18 | 2,96E+05 | 6,33E+06 | 5,53E+06 | 13 | 12 | 3 | 11 | 10 | Oxidation (M) | 89836 | PPTG_05391 | Phytophthora parasitica INRA-310 hypothetical protein (804 aa) |
| 983 | 407 | PPTG_11162 | 160,87 | 25 | 4 | 6 | 24 | 3,42E+05 | 2,00E+06 | 1,47E+07 | 15 | 15 | 3 | 3 | 16 | Carbamidomethylation; Deamidation (NQ) | 93169 | PPTG_11162 | Phytophthora parasitica INRA-310 hypothetical protein (835 aa) |
| 482 | 1599 | PPTG_17268 | 160,83 | 62 | 61 | 51 | 62 | 1,70E+07 | 2,95E+07 | 2,06E+08 | 11 | 11 | 11 | 11 | 22 | Oxidation (M); Deamidation (NQ); Carbamylation | 15060 | PPTG_17268 | Phytophthora parasitica INRA-310 hypothetical protein (134 aa) |
| 380 | 445 | PPTG_00034 | 160,76 | 41 | 16 | 33 | 41 | 4,00E+06 | 4,39E+07 | 5,03E+07 | 17 | 17 | 6 | 21 | 26 | Oxidation (M); Deamidation (NQ) | 50800 | PPTG_00034 | Phytophthora parasitica INRA-310 argininosuccinate lyase (462 aa) |
| 430 | 599 | PPTG_09542 | 160,56 | 13 | 3 | 11 | 11 | 1,09E+06 | 1,12E+07 | 1,61E+07 | 24 | 24 | 6 | 21 | 20 | Oxidation (M); Carbamidomethylation | 216251 | PPTG_09542 | Phytophthora parasitica INRA-310 hypothetical protein (1916 aa) |
| 215 | 982 | PPTG_06371 | 160,25 | 72 | 45 | 59 | 62 | 1,01E+08 | 4,94E+08 | 1,35E+09 | 11 | 11 | 12 | 22 | 46 | Carbamidomethylation; Dehydration; Deamidation (NQ); Acetylation (N-term); Carbamylation | 17505 | PPTG_06371 | Phytophthora parasitica INRA-310 60S ribosomal protein L12 (166 aa) |
| 129 | 817 | PPTG_16130 | 160,11 | 58 | 44 | 45 | 58 | 2,06E+08 | 6,12E+08 | 1,87E+09 | 15 | 15 | 22 | 29 | 49 | Oxidation (M); Carbamidomethylation; Dehydration; Deamidation (NQ); Carbamylation | 16494 | PPTG_16130 | Phytophthora parasitica INRA-310 60S ribosomal protein L27 (145 aa) |
| 533 | 513 | PPTG_01517 | 159,88 | 32 | 20 | 27 | 32 | 7,25E+06 | 2,51E+07 | 8,84E+07 | 12 | 12 | 8 | 13 | 19 | Dehydration; Deamidation (NQ) | 47876 | PPTG_01517 | Phytophthora parasitica INRA-310 trifunctional enzyme subunit beta (447 aa) |
| 509 | 503 | PPTG_02652 | 159,85 | 20 | 11 | 14 | 19 | 1,37E+06 | 5,37E+06 | 2,00E+07 | 16 | 14 | 8 | 14 | 20 | Oxidation (M); Dehydration; Deamidation (NQ) | 94780 | PPTG_02652 | Phytophthora parasitica INRA-310 heat shock protein 90 (836 aa) |
| 125 | 874 | PPTG_15537 | 159,72 | 55 | 54 | 54 | 55 | 1,80E+08 | 8,34E+08 | 2,21E+09 | 12 | 12 | 25 | 33 | 43 | Oxidation (M); Carbamidomethylation; Dehydration; Deamidation (NQ); Acetylation (N-term); Carbam | 20306 | PPTG_15537 | Phytophthora parasitica INRA-310 60S ribosomal protein (179 aa) |
| 862 | 1170 | PPTG_09814 | 159,67 | 29 | 27 | 27 | 29 | 3,33E+06 | 7,35E+06 | 2,11E+07 | 7 | 7 | 7 | 9 | 11 | Carbamidomethylation | 34782 | PPTG_09814 | Phytophthora parasitica INRA-310 hypothetical protein (321 aa) |
| 898 | 558 | PPTG_14746 | 159,56 | 56 | 16 | 50 | 49 | 2,16E+05 | 9,65E+06 | 7,31E+06 | 12 | 12 | 2 | 12 | 11 | Carbamidomethylation; Deamidation (NQ) | 29351 | PPTG_14746 | Phytophthora parasitica INRA-310 carbohydrate esterase (283 aa) |
| 328 | 835 | PPTG_11404 | 159,27 | 54 | 36 | 54 | 54 | 1,71E+07 | 1,53E+08 | 2,92E+08 | 15 | 15 | 9 | 24 | 27 | Oxidation (M); Carbamidomethylation; Dehydration; Deamidation (NQ) | 23491 | PPTG_11404 | Phytophthora parasitica INRA-310 peroxiredoxin-2 (209 aa) |
| 762 | 1151 | PPTG_01586 | 159,1 | 23 | 18 | 23 | 23 | 3,21E+06 | 3,27E+07 | 3,67E+07 | 8 | 8 | 8 | 10 | 12 | Oxidation (M); Carbamidomethylation | 47183 | PPTG_01586 | Phytophthora parasitica INRA-310 lipase (427 aa) |
| 584 | 704 | PPTG_04494 | 159,09 | 49 | 24 | 43 | 43 | 6,78E+06 | 2,45E+07 | 8,29E+07 | 11 | 11 | 8 | 12 | 17 | Oxidation (M); Carbamidomethylation | 32394 | PPTG_04494 | Phytophthora parasitica INRA-310 NADH-cytochrome b5 reductase (297 aa) |
| 99 | 729 | PPTG_05682 | 159,01 | 57 | 44 | 50 | 57 | 2,11E+08 | 8,38E+08 | 2,23E+09 | 14 | 14 | 26 | 39 | 45 | Oxidation (M); Dehydration; Deamidation (NQ); Carbamylation | 20688 | PPTG_05682 | Phytophthora parasitica INRA-310 60S ribosomal protein L18a (177 aa) |
| 888 | 483 | PPTG_08132 | 158,94 | 30 | 2 | 23 | 19 | 6,15E+04 | 7,63E+06 | 4,42E+06 | 17 | 17 | 1 | 12 | 12 | Oxidation (M); Carbamidomethylation | 90145 | PPTG_08132 | Phytophthora parasitica INRA-310 urease (843 aa) |
| 1024 | 1225 | PPTG_10455 | 158,65 | 51 | 16 | 34 | 51 | 1,05E+07 | 4,97E+07 | 8,42E+07 | 7 | 4 | 3 | 7 | 12 | Amidation | 20912 | PPTG_10455 | Phytophthora parasitica INRA-310 EF-1 guanine nucleotide exchange domain-containing protein (193 aa) |
| 530 | 490 | PPTG_08989 | 158,32 | 10 | 3 | 6 | 9 | | | 2,55E+05 | 18 | 1 | 8 | 11 | 21 | Oxidation (M); Carbamidomethylation; Deamidation (NQ) | 196059 | PPTG_08989 | Phytophthora parasitica INRA-310 ATP-binding Cassette (ABC) superfamily (1783 aa) |
| 567 | 461 | PPTG_14633 | 158,29 | 45 | 6 | 42 | 30 | 1,23E+06 | 1,52E+07 | 2,84E+07 | 16 | 16 | 3 | 18 | 17 | Oxidation (M); Carbamidomethylation; Deamidation (NQ) | 48056 | PPTG_14633 | Phytophthora parasitica INRA-310 3-hydroxyisobutyryl-CoA hydrolase (428 aa) |
| 221 | 1801 | PPTG_00353 | 158,19 | 46 | 39 | 45 | 41 | 6,28E+07 | 2,43E+08 | 7,72E+08 | 10 | 10 | 12 | 29 | 38 | Oxidation (M); Carbamidomethylation; Dehydration; Deamidation (NQ); Acetylation (N-term); Carbam | 16065 | PPTG_00353 | Phytophthora parasitica INRA-310 40S ribosomal protein S14 (151 aa) |
| 639 | 795 | PPTG_06980 | 158,18 | 68 | 27 | 44 | 68 | 3,29E+06 | 7,44E+06 | 8,04E+07 | 13 | 13 | 4 | 9 | 20 | Oxidation (M); Dehydration | 22592 | PPTG_06980 | Phytophthora parasitica INRA-310 hypothetical protein (199 aa) |
| 637 | 463 | PPTG_06424 | 158,09 | 26 | 16 | 12 | 22 | 1,42E+06 | 2,70E+06 | 2,74E+07 | 15 | 15 | 9 | 9 | 16 | Oxidation (M); Carbamidomethylation; Dehydration | 78439 | PPTG_06424 | Phytophthora parasitica INRA-310 chloride Channel (ClC) family (706 aa) |
| 688 | 441 | PPTG_08929 | 158,06 | 35 | 18 | 29 | 31 | 6,95E+06 | 2,74E+07 | 6,51E+07 | 12 | 12 | 7 | 11 | 14 | Oxidation (M); Carbamidomethylation | 51355 | PPTG_08929 | Phytophthora parasitica INRA-310 CBL-interacting serine/threonine-protein kinase (465 aa) |
| 407 | 774 | PPTG_06270 | 158 | 75 | 38 | 64 | 70 | 6,01E+06 | 5,33E+07 | 8,89E+07 | 11 | 11 | 10 | 18 | 22 | Oxidation (M); Dehydration; Deamidation (NQ) | 20654 | PPTG_06270 | Phytophthora parasitica INRA-310 ADP-ribosylation factor-like protein 3 (184 aa) |
| 436 | 492 | PPTG_17617 | 157,79 | 34 | 19 | 34 | 32 | 2,54E+06 | 3,09E+07 | 3,83E+07 | 13 | 13 | 6 | 20 | 21 | Oxidation (M); Carbamidomethylation | 51419 | PPTG_17617 | Phytophthora parasitica INRA-310 eukaryotic translation initiation factor 3 (448 aa) |
| 372 | 484 | PPTG_04043 | 157,56 | 30 | 14 | 25 | 28 | 1,79E+07 | 4,02E+07 | 1,62E+08 | 19 | 19 | 8 | 17 | 30 | Oxidation (M); Carbamidomethylation; Dehydration; Deamidation (NQ) | 78359 | PPTG_04043 | Phytophthora parasitica INRA-310 copine (722 aa) |
| 872 | 650 | PPTG_06413 | 157,53 | 70 | 40 | 61 | 60 | 1,34E+06 | 1,06E+07 | 1,31E+07 | 10 | 10 | 5 | 10 | 11 | Oxidation (M); Carbamidomethylation; Acetylation (N-term) | 23507 | PPTG_06413 | Phytophthora parasitica INRA-310 demethylmenaquinone methyltransferase (225 aa) |
| 442 | 723 | PPTG_12286 | 157,52 | 44 | 41 | 41 | 44 | 1,18E+07 | 1,22E+08 | 1,32E+08 | 10 | 10 | 12 | 16 | 18 | Oxidation (M); Carbamidomethylation; Dehydration; Deamidation (NQ); Amidation | 27145 | PPTG_12286 | Phytophthora parasitica INRA-310 triosephosphate isomerase (251 aa) |
| 844 | 766 | PPTG_03968 | 157,34 | 51 | 19 | 47 | 45 | 1,82E+05 | 3,99E+06 | 3,70E+06 | 10 | 4 | 4 | 11 | 12 | Oxidation (M); Carbamidomethylation | 33694 | PPTG_03968 | Phytophthora parasitica INRA-310 biotin-protein ligase (312 aa) |
| 346 | 551 | PPTG_20487 | 157,24 | 53 | 30 | 41 | 53 | 5,52E+06 | 1,88E+07 | 8,82E+07 | 14 | 11 | 12 | 17 | 29 | Oxidation (M); Carbamylation | 27588 | PPTG_20487 | Phytophthora parasitica INRA-310 hypothetical protein (251 aa) |
| 466 | 624 | PPTG_07079 | 157,23 | 51 | 34 | 51 | 51 | 2,01E+05 | 4,63E+06 | 5,69E+06 | 10 | 4 | 8 | 16 | 21 | Oxidation (M); Carbamidomethylation; Deamidation (NQ) | 31076 | PPTG_07079 | Phytophthora parasitica INRA-310 cysteine synthase (293 aa) |
| 693 | 813 | PPTG_08135 | 157,17 | 51 | 21 | 51 | 35 | 6,92E+06 | 6,81E+07 | 7,09E+07 | 11 | 11 | 5 | 15 | 12 | Oxidation (M); Carbamidomethylation | 27172 | PPTG_08135 | Phytophthora parasitica INRA-310 proteasome subunit alpha type-7 threonine protease family T01A (250 aa) |
| 708 | 1760 | PPTG_09268 | 157,17 | 56 | 25 | 43 | 56 | 6,70E+06 | 8,89E+07 | 8,97E+07 | 6 | 6 | 4 | 15 | 13 | Oxidation (M); Carbamidomethylation; Deamidation (NQ) | 18383 | PPTG_09268 | Phytophthora parasitica INRA-310 endoribonuclease L-PSP (173 aa) |
| 520 | 450 | PPTG_17620 | 157,03 | 57 | 21 | 44 | 42 | 2,25E+06 | 1,44E+07 | 2,62E+07 | 17 | 17 | 9 | 17 | 15 | Oxidation (M); Carbamidomethylation; Dehydration; Deamidation (NQ) | 46427 | PPTG_17620 | Phytophthora parasitica INRA-310 peroxisomal multifunctional enzyme (422 aa) |
| 424 | 629 | PPTG_14823 | 156,77 | 61 | 41 | 53 | 51 | 1,24E+07 | 8,10E+06 | 5,54E+07 | 11 | 8 | 12 | 15 | 21 | Oxidation (M); Acetylation (N-term); Carbamylation | 22293 | PPTG_14823 | Phytophthora parasitica INRA-310 Rab1 family GTPase (PiYpt1) (202 aa) |
| 889 | 578 | PPTG_12673 | 156,71 | 42 | 10 | 22 | 42 | 9,49E+05 | 1,70E+06 | 2,54E+07 | 12 | 10 | 3 | 7 | 15 | Oxidation (M) | 41191 | PPTG_12673 | Phytophthora parasitica INRA-310 calcium/calmodulin dependent protein kinase (371 aa) |
| 775 | 482 | PPTG_14768 | 156,63 | 35 | 17 | 29 | 33 | 1,16E+06 | 6,93E+06 | 1,36E+07 | 11 | 11 | 7 | 10 | 12 | Oxidation (M); Carbamidomethylation | 49777 | PPTG_14768 | Phytophthora parasitica INRA-310 RuvB-like 1 (455 aa) |
| 820 | 571 | PPTG_09986 | 156,27 | 24 | 8 | 17 | 24 | 6,37E+05 | 7,57E+06 | 1,73E+07 | 14 | 14 | 4 | 10 | 13 | Carbamidomethylation | 70158 | PPTG_09986 | Phytophthora parasitica INRA-310 ABC transporter E family member 2 (627 aa) |
| 574 | 519 | PPTG_01994 | 156,23 | 38 | 18 | 17 | 36 | 4,04E+06 | 8,07E+06 | 5,21E+07 | 19 | 19 | 9 | 9 | 20 | Oxidation (M); Carbamidomethylation | 55054 | PPTG_01994 | Phytophthora parasitica INRA-310 mitochondrial Carrier (MC) family (500 aa) |
| 865 | 646 | PPTG_08075 | 156,13 | 19 | 8 | 8 | 17 | 2,24E+06 | 1,95E+06 | 1,58E+07 | 15 | 15 | 6 | 6 | 14 | Oxidation (M); Carbamidomethylation; Deamidation (NQ) | 106026 | PPTG_08075 | Phytophthora parasitica INRA-310 plasma-membrane proton-efflux P-type ATPase (966 aa) |
| 1054 | 1061 | PPTG_10776 | 156,07 | 30 | 3 | 26 | 27 | 4,20E+05 | 1,30E+07 | 1,33E+07 | 9 | 9 | 1 | 10 | 10 | Oxidation (M); Carbamidomethylation | 47594 | PPTG_10776 | Phytophthora parasitica INRA-310 cathepsin B cysteine protease family C01A (426 aa) |
| 532 | 512 | PPTG_03960 | 156,05 | 40 | 13 | 32 | 38 | 1,88E+06 | 1,57E+07 | 3,02E+07 | 14 | 14 | 5 | 16 | 19 | Oxidation (M); Carbamidomethylation | 49602 | PPTG_03960 | Phytophthora parasitica INRA-310 26S proteasome regulatory subunit 4 like B (446 aa) |
| 798 | 542 | PPTG_16547 | 155,94 | 51 | 10 | 49 | 48 | 3,22E+05 | 5,99E+06 | 9,04E+06 | 11 | 11 | 2 | 13 | 13 | Oxidation (M); Carbamidomethylation | 35386 | PPTG_16547 | Phytophthora parasitica INRA-310 aspartate carbamoyltransferase (322 aa) |
| 561 | 538 | PPTG_18963 | 155,87 | 7 | 1 | 5 | 4 | 6,39E+04 | 4,58E+06 | 5,70E+06 | 25 | 25 | 2 | 18 | 18 | Carbamidomethylation | 505929 | PPTG_18963 | Phytophthora parasitica INRA-310 HECT E3 ubiquitin ligase (4562 aa) |
| 600 | 501 | PPTG_19379 | 155,87 | 23 | 6 | 18 | 20 | 9,91E+05 | 9,64E+06 | 1,44E+07 | 16 | 16 | 5 | 13 | 18 | Oxidation (M); Carbamidomethylation; Dehydration; Deamidation (NQ) | 96337 | PPTG_19379 | Phytophthora parasitica INRA-310 histidyl-tRNA synthetase (886 aa) |
| 747 | 409 | PPTG_20004 | 155,85 | 34 | 4 | 30 | 32 | 3,96E+05 | 9,73E+06 | 8,09E+06 | 14 | 14 | 2 | 14 | 14 | Carbamidomethylation | 62553 | PPTG_20004 | Phytophthora parasitica INRA-310 hypothetical protein (588 aa) |
| 986 | 799 | PPTG_08537 | 155,75 | 31 | 18 | 20 | 31 | 1,61E+06 | 8,64E+06 | 1,52E+07 | 9 | 9 | 5 | 6 | 11 | Oxidation (M) | 42494 | PPTG_08537 | Phytophthora parasitica INRA-310 protein phosphatase 2C (396 aa) |
| 943 | 621 | PPTG_03687 | 155,6 | 22 | 4 | 21 | 13 | 3,31E+05 | 1,89E+07 | 1,37E+07 | 12 | 12 | 2 | 13 | 9 | Oxidation (M); Deamidation (NQ) | 86590 | PPTG_03687 | Phytophthora parasitica INRA-310 glycosyl hydrolase (806 aa) |
| 910 | 1591 | PPTG_00563 | 155,55 | 48 | 39 | 48 | 48 | 2,10E+05 | 1,87E+05 | 3,09E+06 | 5 | 1 | 4 | 10 | 11 | Oxidation (M); Carbamidomethylation; Acetylation (N-term) | 19548 | PPTG_00563 | Phytophthora parasitica INRA-310 SCP-like extracellular protein (184 aa) |
| 890 | 797 | PPTG_14118 | 155,55 | 23 | 11 | 22 | 19 | 2,65E+06 | 1,26E+07 | 2,07E+07 | 9 | 9 | 6 | 10 | 9 | Oxidation (M); Carbamidomethylation | 54333 | PPTG_14118 | Phytophthora parasitica INRA-310 H/ACA ribonucleoprotein complex subunit 4 (485 aa) |
| 698 | 500 | PPTG_03917 | 155,39 | 40 | 22 | 30 | 40 | 2,66E+06 | 8,08E+06 | 3,49E+07 | 14 | 14 | 6 | 10 | 16 | Oxidation (M); Acetylation (N-term) | 46780 | PPTG_03917 | Phytophthora parasitica INRA-310 H - or Na -translocating F-type V-type and A-type ATPase (F-ATPase) superfamily (416 aa) |
| 531 | 564 | PPTG_13404 | 155,39 | 19 | 3 | 14 | 16 | 2,64E+05 | 6,68E+06 | 1,22E+07 | 23 | 23 | 3 | 18 | 19 | Carbamidomethylation | 156129 | PPTG_13404 | Phytophthora parasitica INRA-310 hypothetical protein (1422 aa) |
| 648 | 459 | PPTG_14247 | 155,37 | 28 | 8 | 18 | 28 | 1,81E+06 | 5,88E+06 | 3,20E+07 | 14 | 14 | 6 | 10 | 18 | Oxidation (M); Carbamidomethylation; Deamidation (NQ) | 69837 | PPTG_14247 | Phytophthora parasitica INRA-310 phosphatidylinositide phosphatase -like protein (612 aa) |
| 588 | 511 | PPTG_03627 | 155,29 | 49 | 7 | 36 | 46 | 6,05E+05 | 8,92E+06 | 1,35E+07 | 18 | 18 | 3 | 14 | 20 | Oxidation (M); Carbamidomethylation; Deamidation (NQ) | 44081 | PPTG_03627 | Phytophthora parasitica INRA-310 actin-like protein 2 (390 aa) |
| 395 | 494 | PPTG_05618 | 155,18 | 45 | 33 | 45 | 42 | 7,08E+06 | 5,96E+07 | 1,25E+08 | 12 | 12 | 10 | 20 | 22 | Oxidation (M) | 34922 | PPTG_05618 | Phytophthora parasitica INRA-310 3 2-trans-enoyl-CoA isomerase (315 aa) |
| 515 | 717 | PPTG_13487 | 155,18 | 62 | 53 | 49 | 58 | 5,25E+06 | 8,72E+06 | 4,83E+07 | 11 | 9 | 13 | 12 | 17 | Oxidation (M); Carbamidomethylation; Acetylation (N-term) | 22673 | PPTG_13487 | Phytophthora parasitica INRA-310 Ras-like protein Rab-8A (203 aa) |
| 563 | 449 | PPTG_02695 | 154,96 | 36 | 11 | 34 | 24 | 1,21E+06 | 1,56E+07 | 2,22E+07 | 13 | 13 | 5 | 19 | 14 | Oxidation (M); Carbamidomethylation; Deamidation (NQ) | 57806 | PPTG_02695 | Phytophthora parasitica INRA-310 nucleolar protein Nop56 (519 aa) |
| 956 | 842 | PPTG_11354 | 154,93 | 41 | 24 | 38 | 32 | 1,01E+06 | 8,50E+06 | 9,43E+06 | 10 | 10 | 5 | 10 | 8 | | 33133 | PPTG_11354 | Phytophthora parasitica INRA-310 regulator-microtubule dynamics protein (297 aa) |
| 911 | 1937 | PPTG_18664 | 154,87 | 41 | 9 | 41 | 30 | 1,52E+06 | 4,08E+07 | 3,80E+07 | 6 | 6 | 2 | 11 | 12 | Carbamidomethylation; Deamidation (NQ); Carbamylation | 20492 | PPTG_18664 | Phytophthora parasitica INRA-310 hypothetical protein (208 aa) |
| 827 | 590 | PPTG_04844 | 154,83 | 41 | 13 | 28 | 36 | 1,79E+06 | 1,96E+07 | 1,90E+07 | 12 | 12 | 3 | 11 | 13 | Oxidation (M); Carbamidomethylation; Dehydration; Deamidation (NQ) | 40533 | PPTG_04844 | Phytophthora parasitica INRA-310 eukaryotic translation initiation factor 3 subunit H (367 aa) |
| 909 | 750 | PPTG_08288 | 154,8 | 28 | 8 | 21 | 28 | 1,23E+06 | 4,32E+06 | 3,48E+07 | 10 | 10 | 3 | 6 | 16 | Oxidation (M); Carbamidomethylation | 46002 | PPTG_08288 | Phytophthora parasitica INRA-310 protein phosphatase 2C (422 aa) |
| 743 | 556 | PPTG_07329 | 154,77 | 38 | 22 | 22 | 36 | 2,64E+06 | 4,44E+06 | 4,15E+07 | 13 | 12 | 7 | 6 | 17 | Carbamidomethylation; Deamidation (NQ) | 40565 | PPTG_07329 | Phytophthora parasitica INRA-310 thioredoxin-like protein (364 aa) |
| 776 | 436 | PPTG_01070 | 154,66 | 43 | 22 | 27 | 39 | 1,98E+06 | 2,77E+06 | 2,23E+07 | 13 | 13 | 7 | 8 | 14 | | 45101 | PPTG_01070 | Phytophthora parasitica INRA-310 dihydroorotate oxidase (425 aa) |
| 178 | 963 | PPTG_17588 | 154,51 | 33 | 24 | 33 | 32 | 1,42E+08 | 6,13E+08 | 1,65E+09 | 14 | 14 | 14 | 32 | 41 | Carbamidomethylation; Dehydration; Deamidation (NQ); Carbamylation | 33057 | PPTG_17588 | Phytophthora parasitica INRA-310 40S ribosomal protein S11 (291 aa) |
| 749 | 1253 | PPTG_05057 | 154,4 | 26 | 20 | 26 | 22 | 6,34E+06 | 5,16E+07 | 6,05E+07 | 8 | 8 | 5 | 14 | 11 | Oxidation (M); Deamidation (NQ) | 43090 | PPTG_05057 | Phytophthora parasitica INRA-310 hypothetical protein (398 aa) |
| 303 | 830 | PPTG_02563 | 154,4 | 26 | 21 | 24 | 24 | 8,25E+07 | 2,15E+08 | 9,70E+08 | 9 | 9 | 14 | 18 | 32 | Carbamidomethylation; Deamidation (NQ); Carbamylation | 43053 | PPTG_02563 | Phytophthora parasitica INRA-310 hypothetical protein (405 aa) |
| 500 | 700 | PPTG_09997 | 154,16 | 15 | 3 | 11 | 15 | 4,54E+05 | 8,60E+06 | 9,36E+06 | 20 | 20 | 4 | 17 | 21 | Oxidation (M); Carbamidomethylation | 167240 | PPTG_09997 | Phytophthora parasitica INRA-310 sulfite reductase [NADPH] subunit beta (1504 aa) |
| 663 | 657 | PPTG_03707 | 154,13 | 32 | 8 | 24 | 30 | 3,06E+05 | 2,43E+06 | 5,02E+06 | 13 | 7 | 3 | 12 | 17 | Oxidation (M); Carbamidomethylation; Dehydration; Acetylation (N-term) | 54397 | PPTG_03707 | Phytophthora parasitica INRA-310 pyruvate kinase (506 aa) |
| 258 | 548 | PPTG_05621 | 154,05 | 33 | 18 | 28 | 32 | 2,05E+07 | 6,69E+07 | 1,32E+08 | 20 | 19 | 14 | 24 | 33 | Oxidation (M); Carbamidomethylation; Dehydration; Deamidation (NQ) | 69267 | PPTG_05621 | Phytophthora parasitica INRA-310 DEAD/DEAH box RNA helicase (639 aa) |
| 936 | 603 | PPTG_04070 | 154,01 | 23 | 4 | 13 | 23 | 2,42E+05 | 1,71E+06 | 1,19E+07 | 13 | 13 | 2 | 7 | 15 | | 76680 | PPTG_04070 | Phytophthora parasitica INRA-310 ATP-binding Cassette (ABC) superfamily (698 aa) |
| 651 | 417 | PPTG_03675 | 153,91 | 46 | 18 | 38 | 46 | 7,35E+05 | 6,07E+06 | 1,07E+07 | 13 | 13 | 6 | 13 | 15 | Oxidation (M); Carbamidomethylation; Acetylation (N-term); Carbamylation | 49917 | PPTG_03675 | Phytophthora parasitica INRA-310 glutaredoxin (449 aa) |
| 1057 | 2203 | PPTG_13222 | 153,85 | 46 | 31 | 31 | 46 | 2,09E+06 | 7,20E+06 | 4,36E+07 | 4 | 4 | 5 | 6 | 10 | Oxidation (M); Dehydration; Deamidation (NQ) | 17231 | PPTG_13222 | Phytophthora parasitica INRA-310 hypothetical protein (170 aa) |
| 1057 | 2204 | PPTG_13222 | 153,85 | 46 | 31 | 31 | 46 | 2,09E+06 | 7,20E+06 | 4,36E+07 | 4 | 4 | 5 | 6 | 10 | Oxidation (M); Dehydration; Deamidation (NQ) | 17231 | PPTG_13222 | Phytophthora parasitica INRA-310 hypothetical protein (170 aa) |

| | | | | | | | | | | | | | | | | | | | |
|---|---|---|---|---|---|---|---|---|---|---|---|---|---|---|---|---|---|---|---|
| 289 | 644 | PPTG_06420 | 153,75 | 63 | 36 | 53 | 63 | 9,10E+07 | 2,24E+08 | 8,49E+08 | 11 | 11 | 12 | 22 | 32 | Oxidation (M); Carbamidomethylation; Dehydration; Carbamylation | 23912 | PPTG_06420 | Phytophthora parasitica INRA-310 nucleoside diphosphate kinase B (226 aa) |
| 1555 | 1076 | PPTG_08967 | 153,61 | 28 | 4 | 1 | 25 | 2,84E+05 | 3,53E+05 | 4,01E+06 | 9 | 9 | 2 | 1 | 9 | Oxidation (M); Carbamidomethylation | 55191 | PPTG_08967 | Phytophthora parasitica INRA-310 gamma-soluble NSF attachment protein (509 aa) |
| 479 | 568 | PPTG_03938 | 153,56 | 31 | 11 | 25 | 28 | 2,67E+06 | 2,26E+07 | 3,05E+07 | 20 | 20 | 7 | 18 | 19 | Oxidation (M); Carbamidomethylation; Deamidation (NQ) | 80892 | PPTG_03938 | Phytophthora parasitica INRA-310 delta-1-pyrroline-5-carboxylate synthetase (758 aa) |
| 836 | 1124 | PPTG_01107 | 153,55 | 37 | 23 | 26 | 37 | 1,62E+07 | 2,55E+07 | 1,77E+08 | 10 | 10 | 5 | 6 | 16 | Oxidation (M); Deamidation (NQ) | 25552 | PPTG_01107 | Phytophthora parasitica INRA-310 hypothetical protein (237 aa) |
| 602 | 474 | PPTG_14719 | 153,53 | 22 | 5 | 21 | 12 | 6,10E+05 | 9,84E+06 | 1,34E+07 | 17 | 17 | 4 | 18 | 14 | Oxidation (M); Carbamidomethylation; Deamidation (NQ) | 109870 | PPTG_14719 | Phytophthora parasitica INRA-310 exportin-2-like protein (970 aa) |
| 746 | 447 | PPTG_12276 | 153,5 | 14 | 5 | 3 | 14 | 3,16E+05 | 3,44E+04 | 8,25E+06 | 15 | 10 | 6 | 4 | 20 | Oxidation (M); Carbamidomethylation | 150616 | PPTG_12276 | Phytophthora parasitica INRA-310 ATP-binding Cassette (ABC) superfamily (1354 aa) |
| 686 | 517 | PPTG_09168 | 153,34 | 25 | 7 | 20 | 25 | 3,68E+05 | 4,34E+06 | 8,01E+06 | 16 | 16 | 4 | 12 | 16 | Carbamidomethylation; Dehydration | 93497 | PPTG_09168 | Phytophthora parasitica INRA-310 cGMP-dependent 3' 5'-cGMP phosphodiesterase (850 aa) |
| 550 | 541 | PPTG_04886 | 153,31 | 25 | 14 | 18 | 25 | 3,95E+06 | 1,12E+07 | 4,16E+07 | 12 | 12 | 9 | 11 | 19 | Oxidation (M); Carbamidomethylation | 70981 | PPTG_04886 | Phytophthora parasitica INRA-310 tubby-like protein (633 aa) |
| 1172 | 917 | PPTG_15080 | 153,26 | 34 | 10 | 18 | 34 | 1,84E+05 | 7,43E+05 | 5,81E+06 | 8 | 8 | 2 | 5 | 11 | Oxidation (M); Carbamidomethylation | 48980 | PPTG_15080 | Phytophthora parasitica INRA-310 copine-like protein (435 aa) |
| 952 | 681 | PPTG_13264 | 153,04 | 25 | 9 | 23 | 19 | 1,19E+06 | 9,97E+06 | 1,36E+07 | 12 | 12 | 3 | 12 | 8 | Carbamidomethylation; Deamidation (NQ); Carbamylation | 66955 | PPTG_13264 | Phytophthora parasitica INRA-310 eukaryotic translation initiation factor 2A (620 aa) |
| 1127 | 955 | PPTG_07724 | 152,86 | 41 | 22 | 41 | 25 | 7,35E+05 | 1,61E+07 | 9,87E+06 | 8 | 8 | 4 | 10 | 5 | Carbamidomethylation | 33354 | PPTG_07724 | Phytophthora parasitica INRA-310 glycoside hydrolase (299 aa) |
| 630 | 632 | PPTG_14474 | 152,72 | 68 | 24 | 52 | 68 | 9,89E+05 | 4,72E+06 | 2,65E+07 | 11 | 10 | 5 | 11 | 19 | Oxidation (M); Carbamidomethylation; Carbamylation | 23285 | PPTG_14474 | Phytophthora parasitica INRA-310 Ras-like protein Rab-7a (208 aa) |
| 332 | 1313 | PPTG_17566 | 152,69 | 56 | 42 | 48 | 51 | 3,11E+07 | 1,52E+08 | 4,49E+08 | 10 | 10 | 10 | 18 | 31 | Oxidation (M); Carbamidomethylation; Dehydration; Deamidation (NQ); Carbamylation | 15834 | PPTG_17566 | Phytophthora parasitica INRA-310 50S ribosomal protein L32 (136 aa) |
| 1365 | 1020 | PPTG_09286 | 152,62 | 14 | 5 | 9 | 14 | 1,79E+05 | 5,63E+05 | 6,59E+06 | 8 | 6 | 3 | 5 | 7 | Carbamidomethylation | 83438 | PPTG_09286 | Phytophthora parasitica INRA-310 hypothetical protein (812 aa) |
| 452 | 600 | PPTG_03990 | 152,56 | 10 | 4 | 6 | 9 | 8,98E+05 | 8,03E+06 | 1,18E+07 | 24 | 24 | 9 | 15 | 21 | Oxidation (M) | 297006 | PPTG_03990 | Phytophthora parasitica INRA-310 phosphatidylinositol kinase (PIK-L2) (2654 aa) |
| 821 | 491 | PPTG_05288 | 152,51 | 21 | 4 | 17 | 20 | 3,01E+05 | 9,12E+06 | 1,14E+07 | 13 | 13 | 2 | 12 | 13 | Carbamidomethylation; Acetylation (N-term) | 93180 | PPTG_05288 | Phytophthora parasitica INRA-310 ubiquitin carboxyl-terminal hydrolase (852 aa) |
| 772 | 429 | PPTG_19371 | 152,48 | 16 | 2 | 7 | 16 | 3,60E+05 | 1,75E+06 | 8,46E+06 | 15 | 14 | 4 | 7 | 17 | Oxidation (M); Carbamidomethylation; Dehydration; Deamidation (NQ) | 130627 | PPTG_19371 | Phytophthora parasitica INRA-310 mitogen-activated protein kinase (1196 aa) |
| 802 | 789 | PPTG_17571 | 152,47 | 45 | 28 | 25 | 45 | 1,24E+06 | 2,90E+06 | 1,87E+07 | 9 | 9 | 7 | 6 | 15 | Deamidation (NQ) | 35080 | PPTG_17571 | Phytophthora parasitica INRA-310 hypothetical protein (320 aa) |
| 589 | 696 | PPTG_17616 | 152,36 | 22 | 16 | 22 | 22 | 2,07E+06 | 2,85E+07 | 2,65E+07 | 11 | 11 | 7 | 16 | 14 | Carbamidomethylation; Dehydration; Deamidation (NQ) | 65734 | PPTG_17616 | Phytophthora parasitica INRA-310 hypothetical protein (578 aa) |
| 668 | 843 | PPTG_05149 | 152,31 | 32 | 16 | 26 | 32 | 1,26E+06 | 1,80E+07 | 3,92E+07 | 10 | 9 | 5 | 12 | 16 | Oxidation (M); Deamidation (NQ) | 37400 | PPTG_05149 | Phytophthora parasitica INRA-310 cAMP-dependent protein kinase catalytic subunit (331 aa) |
| 649 | 561 | PPTG_19704 | 152,27 | 57 | 11 | 57 | 36 | 5,54E+05 | 7,77E+06 | 7,41E+06 | 11 | 7 | 4 | 15 | 15 | Oxidation (M); Carbamidomethylation; Deamidation (NQ) | 36170 | PPTG_19704 | Phytophthora parasitica INRA-310 cysteine synthase A (342 aa) |
| 546 | 528 | PPTG_14362 | 152,14 | 10 | 3 | 7 | 8 | 5,20E+05 | 4,80E+06 | 9,55E+06 | 20 | 20 | 6 | 16 | 17 | Oxidation (M); Carbamidomethylation | 280482 | PPTG_14362 | Phytophthora parasitica INRA-310 hypothetical protein (2504 aa) |
| 756 | 525 | PPTG_00739 | 151,87 | 25 | 3 | 21 | 25 | 2,41E+05 | 6,49E+06 | 9,20E+06 | 11 | 11 | 1 | 13 | 16 | Oxidation (M); Carbamidomethylation | 66291 | PPTG_00739 | Phytophthora parasitica INRA-310 hypothetical protein (594 aa) |
| 899 | 526 | PPTG_09530 | 151,87 | 28 | 11 | 8 | 25 | 6,30E+05 | 3,32E+06 | 2,42E+07 | 13 | 13 | 5 | 5 | 15 | Oxidation (M); Acetylation (N-term) | 69175 | PPTG_09530 | Phytophthora parasitica INRA-310 vacuolar protein sorting-associated protein (626 aa) |
| 606 | 946 | PPTG_09215 | 151,77 | 20 | 13 | 19 | 20 | 2,03E+06 | 1,29E+08 | 4,71E+07 | 8 | 8 | 5 | 15 | 16 | Oxidation (M) | 65814 | PPTG_09215 | Phytophthora parasitica INRA-310 glucosylceramidase (604 aa) |
| 669 | 567 | PPTG_13582 | 151,76 | 31 | 12 | 23 | 27 | 1,94E+06 | 1,97E+07 | 2,79E+07 | 11 | 11 | 6 | 13 | 14 | Oxidation (M); Carbamidomethylation; Deamidation (NQ) | 62479 | PPTG_13582 | Phytophthora parasitica INRA-310 2-isopropylmalate synthase (577 aa) |
| 940 | 749 | PPTG_19098 | 151,64 | 24 | 4 | 22 | 21 | 2,77E+05 | 1,17E+07 | 1,06E+07 | 10 | 10 | 2 | 11 | 11 | Oxidation (M); Carbamidomethylation | 64355 | PPTG_19098 | Phytophthora parasitica INRA-310 hypothetical protein (613 aa) |
| 580 | 775 | PPTG_10213 | 151,57 | 10 | 2 | 2 | 10 | 2,53E+05 | 5,28E+05 | 1,31E+07 | 25 | 25 | 5 | 6 | 26 | Carbamidomethylation | 353151 | PPTG_10213 | Phytophthora parasitica INRA-310 hypothetical protein (3329 aa) |
| 580 | 776 | PPTG_10213 | 151,57 | 10 | 2 | 2 | 10 | 2,53E+05 | 5,28E+05 | 1,31E+07 | 25 | 25 | 5 | 6 | 26 | Carbamidomethylation | 353151 | PPTG_10213 | Phytophthora parasitica INRA-310 hypothetical protein (3329 aa) |
| 674 | 803 | PPTG_08413 | 151,54 | 54 | 29 | 45 | 50 | 4,09E+06 | 7,26E+06 | 5,60E+07 | 10 | 10 | 6 | 9 | 18 | Oxidation (M); Carbamidomethylation; Dehydration | 22381 | PPTG_08413 | Phytophthora parasitica INRA-310 synaptobrevin-like protein (200 aa) |
| 758 | 478 | PPTG_06879 | 151,52 | 40 | 13 | 32 | 35 | 1,41E+06 | 1,18E+07 | 1,56E+07 | 13 | 13 | 4 | 13 | 13 | Oxidation (M); Carbamidomethylation; Deamidation (NQ) | 46059 | PPTG_06879 | Phytophthora parasitica INRA-310 hypothetical protein (408 aa) |
| 1363 | 771 | PPTG_15291 | 151,44 | 27 | 5 | 0 | 27 | | | 2,14E+06 | 10 | 4 | 2 | 0 | 13 | Carbamidomethylation; Deamidation (NQ) | 60561 | PPTG_15291 | Phytophthora parasitica INRA-310 transmembrane protein (536 aa) |
| 661 | 1397 | PPTG_19125 | 151,44 | 47 | 28 | 29 | 47 | 7,99E+06 | 1,65E+07 | 1,42E+08 | 6 | 6 | 5 | 8 | 21 | Oxidation (M); Carbamidomethylation; Dehydration; Deamidation (NQ); Carbamylation | 29469 | PPTG_19125 | Phytophthora parasitica INRA-310 cytochrome c1 (271 aa) |
| 618 | 1014 | PPTG_07145 | 151,28 | 54 | 42 | 42 | 54 | 1,00E+07 | 5,22E+07 | 8,49E+07 | 9 | 9 | 8 | 13 | 15 | Oxidation (M); Carbamidomethylation; Dehydration; Deamidation (NQ) | 25290 | PPTG_07145 | Phytophthora parasitica INRA-310 crinkler (CRN) family protein pseudogene (230 aa) |
| 1075 | 1102 | PPTG_13946 | 151,06 | 38 | 15 | 24 | 38 | 6,39E+05 | 7,40E+06 | 9,74E+06 | 9 | 9 | 3 | 6 | 11 | Carbamidomethylation | 33805 | PPTG_13946 | Phytophthora parasitica INRA-310 microtubule -associated protein EB1 (310 aa) |
| 447 | 1284 | PPTG_01700 | 150,99 | 65 | 43 | 48 | 65 | 3,33E+07 | 1,99E+08 | 3,70E+08 | 7 | 7 | 9 | 19 | 18 | Deamidation (NQ); Acetylation (N-term); Amidation | 17175 | PPTG_01700 | Phytophthora parasitica INRA-310 transcription factor BTF3-like protein (163 aa) |
| 409 | 1349 | PPTG_01954 | 150,8 | 52 | 42 | 51 | 51 | 8,51E+07 | 4,64E+08 | 8,66E+08 | 8 | 8 | 7 | 20 | 22 | Oxidation (M); Carbamidomethylation; Deamidation (NQ); Acetylation (N-term); Carbamylation | 17913 | PPTG_01954 | Phytophthora parasitica INRA-310 ubiquitin-40S ribosomal protein S27a (157 aa) |
| 763 | 549 | PPTG_01436 | 150,8 | 11 | 3 | 7 | 9 | 5,38E+05 | 3,18E+06 | 6,00E+06 | 16 | 16 | 5 | 10 | 14 | Carbamidomethylation | 199421 | PPTG_01436 | Phytophthora parasitica INRA-310 intraflagellar transporter 172 (1772 aa) |
| 638 | 689 | PPTG_12624 | 150,8 | 29 | 2 | 23 | 19 | 6,62E+05 | 1,56E+07 | 2,22E+07 | 18 | 18 | 2 | 16 | 16 | Oxidation (M); Carbamidomethylation; Dehydration; Deamidation (NQ) | 108241 | PPTG_12624 | Phytophthora parasitica INRA-310 glycine dehydrogenase (999 aa) |
| 866 | 570 | PPTG_16129 | 150,79 | 21 | 3 | 0 | 21 | 2,73E+05 | | 1,37E+07 | 22 | 22 | 3 | 0 | 23 | Carbamidomethylation | 151389 | PPTG_16129 | Phytophthora parasitica INRA-310 hypothetical protein (1330 aa) |
| 578 | 908 | PPTG_15127 | 150,77 | 55 | 23 | 55 | 38 | 2,84E+06 | 3,20E+07 | 3,79E+07 | 12 | 12 | 6 | 17 | 15 | Oxidation (M); Carbamidomethylation; Dehydration; Deamidation (NQ) | 27529 | PPTG_15127 | Phytophthora parasitica INRA-310 proteasome subunit alpha type-4 (251 aa) |
| 553 | 609 | PPTG_04830 | 150,61 | 18 | 2 | 14 | 16 | 2,44E+05 | 3,66E+06 | 1,02E+07 | 18 | 18 | 3 | 17 | 19 | Oxidation (M); Carbamidomethylation; Dehydration; Deamidation (NQ) | 144925 | PPTG_04830 | Phytophthora parasitica INRA-310 L-aminoadipate-semialdehyde dehydrogenase large subunit (1312 aa) |
| 879 | 1109 | PPTG_06464 | 150,54 | 43 | 19 | 43 | 43 | 5,92E+06 | 1,05E+08 | 9,53E+07 | 9 | 9 | 3 | 10 | 13 | Oxidation (M); Dehydration | 26648 | PPTG_06464 | Phytophthora parasitica INRA-310 hypothetical protein (248 aa) |
| 1108 | 1936 | PPTG_12760 | 150,34 | 8 | 7 | 8 | 8 | 3,73E+06 | 6,86E+07 | 1,03E+08 | 4 | 4 | 4 | 8 | 8 | Carbamidomethylation; Deamidation (NQ) | 79356 | PPTG_12760 | Phytophthora parasitica INRA-310 hypothetical protein (799 aa) |
| 773 | 730 | PPTG_06756 | 150,31 | 57 | 21 | 54 | 54 | 5,71E+05 | 2,99E+06 | 5,39E+06 | 11 | 1 | 4 | 11 | 14 | Oxidation (M); Amidation | 26393 | PPTG_06756 | Phytophthora parasitica INRA-310 short chain dehydrogenase (243 aa) |
| 719 | 504 | PPTG_02151 | 150,19 | 34 | 10 | 24 | 34 | 1,50E+06 | 9,80E+06 | 1,85E+07 | 14 | 14 | 4 | 11 | 16 | Oxidation (M); Carbamidomethylation | 55114 | PPTG_02151 | Phytophthora parasitica INRA-310 DnaJ subfamily C protein (502 aa) |
| 732 | 718 | PPTG_15545 | 150 | 30 | 22 | 22 | 27 | 3,76E+06 | 6,47E+06 | 2,47E+07 | 10 | 8 | 9 | 9 | 13 | Oxidation (M); Carbamidomethylation; Deamidation (NQ) | 45009 | PPTG_15545 | Phytophthora parasitica INRA-310 hypothetical protein (410 aa) |
| 833 | 953 | PPTG_14334 | 149,99 | 36 | 13 | 33 | 33 | 3,80E+06 | 1,50E+07 | 4,03E+07 | 9 | 9 | 4 | 12 | 11 | | 34985 | PPTG_14334 | Phytophthora parasitica INRA-310 hydroxyacyl-coenzyme A dehydrogenase (325 aa) |
| 839 | 865 | PPTG_04856 | 149,96 | 33 | 18 | 31 | 33 | 2,18E+06 | 1,79E+07 | 2,23E+07 | 9 | 9 | 6 | 9 | 12 | Oxidation (M) | 41949 | PPTG_04856 | Phytophthora parasitica INRA-310 hypothetical protein (382 aa) |
| 955 | 514 | PPTG_04854 | 149,96 | 29 | 13 | 16 | 26 | 5,48E+05 | 2,35E+06 | 6,17E+06 | 12 | 12 | 5 | 7 | 11 | Oxidation (M); Carbamidomethylation | 64746 | PPTG_04854 | Phytophthora parasitica INRA-310 bardet-Biedl syndrome 1 family protein (590 aa) |
| 1454 | 1181 | PPTG_14842 | 149,85 | 16 | 7 | 5 | 14 | 2,78E+06 | 6,32E+06 | 2,71E+07 | 8 | 8 | 4 | 2 | 8 | Oxidation (M); Carbamidomethylation | 73212 | PPTG_14842 | Phytophthora parasitica INRA-310 endomembrane protein 70-like protein (646 aa) |
| 613 | 560 | PPTG_02161 | 149,76 | 63 | 40 | 46 | 63 | 1,09E+07 | 2,00E+07 | 1,18E+08 | 10 | 10 | 7 | 11 | 18 | Oxidation (M); Carbamidomethylation; Deamidation (NQ); Acetylation (N-term); Carbamylation | 23244 | PPTG_02161 | Phytophthora parasitica INRA-310 Ras-like protein Rab-2-A (211 aa) |
| 990 | 569 | PPTG_02090 | 149,76 | 26 | 13 | 11 | 26 | 1,49E+06 | 1,62E+06 | 1,55E+07 | 11 | 11 | 6 | 4 | 12 | Carbamidomethylation | 60130 | PPTG_02090 | Phytophthora parasitica INRA-310 aldehyde dehydrogenase (542 aa) |
| 673 | 653 | PPTG_15632 | 149,76 | 51 | 26 | 37 | 40 | 5,74E+06 | 6,10E+06 | 3,89E+07 | 12 | 10 | 7 | 10 | 16 | Oxidation (M); Deamidation (NQ) | 30549 | PPTG_15632 | Phytophthora parasitica INRA-310 voltage-dependent anion-selective channel protein (283 aa) |
| 427 | 1093 | PPTG_02396 | 149,67 | 38 | 28 | 38 | 38 | 1,09E+07 | 4,92E+07 | 8,46E+07 | 7 | 3 | 9 | 20 | 19 | Deamidation (NQ) | 25213 | PPTG_02396 | Phytophthora parasitica INRA-310 hypothetical protein (230 aa) |
| 676 | 607 | PPTG_03260 | 149,65 | 34 | 17 | 31 | 34 | 2,24E+06 | 2,76E+07 | 3,13E+07 | 14 | 14 | 5 | 14 | 14 | Oxidation (M); Carbamidomethylation; Deamidation (NQ) | 48074 | PPTG_03260 | Phytophthora parasitica INRA-310 acetylornithine aminotransferase (440 aa) |
| 521 | 699 | PPTG_16923 | 149,55 | 30 | 22 | 30 | 30 | 1,19E+07 | 6,88E+07 | 1,03E+08 | 10 | 7 | 8 | 15 | 18 | Oxidation (M); Dehydration; Deamidation (NQ) | 52964 | PPTG_16923 | Phytophthora parasitica INRA-310 glutathione transferase theta class (466 aa) |
| 485 | 583 | PPTG_09690 | 149,43 | 31 | 12 | 29 | 25 | 1,47E+06 | 2,58E+07 | 2,61E+07 | 11 | 11 | 5 | 20 | 18 | Oxidation (M); Carbamidomethylation; Deamidation (NQ) | 59688 | PPTG_09690 | Phytophthora parasitica INRA-310 methylmalonate-semialdehyde dehydrogenase (acylating) (551 aa) |
| 724 | 668 | PPTG_06388 | 149,35 | 25 | 7 | 22 | 23 | 1,10E+06 | 1,16E+07 | 9,06E+06 | 11 | 8 | 4 | 14 | 13 | Oxidation (M); Carbamidomethylation; Dehydration; Deamidation (NQ); Amidation | 58355 | PPTG_06388 | Phytophthora parasitica INRA-310 hypothetical protein (525 aa) |
| 780 | 741 | PPTG_10122 | 149,12 | 27 | 12 | 27 | 19 | 4,12E+06 | 3,08E+07 | 3,96E+07 | 11 | 11 | 7 | 13 | 9 | Oxidation (M); Carbamidomethylation; Deamidation (NQ) | 61554 | PPTG_10122 | Phytophthora parasitica INRA-310 hsp70-like protein (570 aa) |
| 164 | 1531 | PPTG_01761 | 149,11 | 45 | 44 | 45 | 45 | 3,95E+07 | 4,34E+08 | 5,73E+08 | 6 | 6 | 13 | 42 | 35 | Oxidation (M); Carbamidomethylation; Dehydration; Acetylation (N-term); Carbamylation | 16990 | PPTG_01761 | Phytophthora parasitica INRA-310 nucleoside diphosphate kinase B (152 aa) |
| 1143 | 1232 | PPTG_08347 | 149,06 | 48 | 17 | 48 | 45 | 6,72E+05 | 4,38E+06 | 3,77E+07 | 7 | 7 | 2 | 7 | 10 | Deamidation (NQ) | 26255 | PPTG_08347 | Phytophthora parasitica INRA-310 NADH ubiquinone oxidoreductase subunit (227 aa) |
| 219 | 1007 | PPTG_14112 | 148,93 | 39 | 24 | 39 | 33 | 1,11E+08 | 4,55E+08 | 1,26E+09 | 10 | 10 | 13 | 31 | 35 | Oxidation (M); Carbamidomethylation; Deamidation (NQ); Carbamylation | 27782 | PPTG_14112 | Phytophthora parasitica INRA-310 60S ribosomal protein L8 (257 aa) |
| 831 | 665 | PPTG_18243 | 148,89 | 40 | 3 | 39 | 22 | 9,95E+04 | 6,59E+06 | 5,09E+06 | 14 | 14 | 1 | 17 | 9 | Oxidation (M); Carbamidomethylation; Deamidation (NQ) | 55822 | PPTG_18243 | Phytophthora parasitica INRA-310 4-aminobutyrate aminotransferase (504 aa) |
| 653 | 547 | PPTG_09562 | 148,85 | 30 | 9 | 12 | 28 | 1,73E+06 | 2,60E+06 | 2,85E+07 | 17 | 17 | 6 | 9 | 19 | Oxidation (M); Carbamidomethylation | 81701 | PPTG_09562 | Phytophthora parasitica INRA-310 phosphatidylinositide phosphatase (742 aa) |
| 761 | 666 | PPTG_14139 | 148,83 | 47 | 19 | 43 | 47 | 3,47E+06 | 2,64E+07 | 3,39E+07 | 11 | 11 | 6 | 11 | 13 | Oxidation (M) | 34627 | PPTG_14139 | Phytophthora parasitica INRA-310 hypothetical protein (311 aa) |
| 659 | 780 | PPTG_03631 | 148,68 | 49 | 33 | 49 | 49 | 2,79E+06 | 2,70E+07 | 2,82E+07 | 9 | 9 | 6 | 15 | 13 | Oxidation (M); Carbamidomethylation; Deamidation (NQ); Acetylation (N-term) | 30381 | PPTG_03631 | Phytophthora parasitica INRA-310 eukaryotic translation initiation factor 3 subunit F (280 aa) |
| 1493 | 866 | PPTG_11136 | 148,5 | 26 | 0 | 23 | 15 | | 2,71E+06 | 1,52E+06 | 9 | 9 | 0 | 8 | 5 | Carbamidomethylation | 62057 | PPTG_11136 | Phytophthora parasitica INRA-310 serine protease family S10 (555 aa) |
| 513 | 470 | PPTG_16288 | 148,49 | 58 | 20 | 55 | 42 | 3,26E+06 | 3,84E+07 | 4,40E+07 | 13 | 13 | 6 | 18 | 18 | Oxidation (M); Carbamidomethylation; Deamidation (NQ); Acetylation (N-term) | 32770 | PPTG_16288 | Phytophthora parasitica INRA-310 nicotinate-nucleotide diphosphorylase (carboxylating) (302 aa) |
| 801 | 584 | PPTG_02623 | 148,44 | 27 | 8 | 9 | 25 | 1,45E+06 | 1,52E+06 | 1,75E+07 | 14 | 14 | 4 | 4 | 20 | Oxidation (M); Carbamidomethylation | 63248 | PPTG_02623 | Phytophthora parasitica INRA-310 galactonolactone dehydrogenase (561 aa) |
| 1023 | 905 | PPTG_14269 | 148,26 | 20 | 7 | 19 | 13 | 5,12E+05 | 1,83E+07 | 1,38E+07 | 10 | 10 | 3 | 10 | 9 | Oxidation (M); Carbamidomethylation | 63107 | PPTG_14269 | Phytophthora parasitica INRA-310 hypothetical protein (571 aa) |
| 854 | 497 | PPTG_12599 | 148,25 | 36 | 11 | 5 | 35 | 1,05E+06 | 1,08E+06 | 1,46E+07 | 16 | 16 | 6 | 3 | 18 | Oxidation (M); Carbamidomethylation | 66584 | PPTG_12599 | Phytophthora parasitica INRA-310 glutamyl-tRNA(Gln) amidotransferase subunit A (619 aa) |
| 435 | 772 | PPTG_06138 | 148,23 | 51 | 18 | 47 | 34 | 2,50E+06 | 2,71E+07 | 3,93E+07 | 21 | 21 | 8 | 22 | 17 | Oxidation (M); Carbamidomethylation; Deamidation (NQ) | 51387 | PPTG_06138 | Phytophthora parasitica INRA-310 hypothetical protein (469 aa) |
| 566 | 639 | PPTG_06287 | 148,21 | 7 | 3 | 5 | 7 | | | 1,90E+05 | 15 | 1 | 9 | 10 | 19 | Oxidation (M); Carbamidomethylation; Deamidation (NQ) | 214095 | PPTG_06287 | Phytophthora parasitica INRA-310 ATP-binding Cassette (ABC) superfamily (1935 aa) |
| 774 | 518 | PPTG_00525 | 148,19 | 27 | 5 | 18 | 27 | 5,58E+05 | 2,15E+06 | 1,34E+07 | 15 | 15 | 3 | 9 | 17 | Carbamidomethylation | 70633 | PPTG_00525 | Phytophthora parasitica INRA-310 atlastin-like protein (621 aa) |
| 657 | 595 | PPTG_10954 | 148,05 | 40 | 26 | 34 | 26 | 1,11E+07 | 1,16E+08 | 9,72E+07 | 13 | 13 | 8 | 14 | 12 | Carbamidomethylation; Dehydration; Deamidation (NQ) | 41828 | PPTG_10954 | Phytophthora parasitica INRA-310 glucan 1 3-beta-glucosidase (395 aa) |
| 634 | 659 | PPTG_18968 | 148,05 | 11 | 3 | 8 | 10 | 7,91E+05 | 6,25E+06 | 1,06E+07 | 17 | 16 | 4 | 14 | 16 | Oxidation (M); Carbamidomethylation | 202680 | PPTG_18968 | Phytophthora parasitica INRA-310 DNA-directed RNA polymerase I subunit RPA1 (1812 aa) |
| 510 | 496 | PPTG_05082 | 148,04 | 30 | 20 | 30 | 30 | 5,79E+06 | 4,70E+07 | 5,79E+07 | 12 | 12 | 10 | 17 | 15 | Oxidation (M); Carbamidomethylation | 49971 | PPTG_05082 | Phytophthora parasitica INRA-310 26S proteasome non-ATPase regulatory subunit 12 (438 aa) |
| 166 | 974 | PPTG_15753 | 147,86 | 47 | 40 | 43 | 46 | 1,61E+08 | 8,27E+08 | 2,17E+09 | 11 | 11 | 22 | 27 | 40 | Oxidation (M); Dehydration; Deamidation (NQ); Carbamylation | 22629 | PPTG_15753 | Phytophthora parasitica INRA-310 30S ribosomal protein S4 (193 aa) |
| 619 | 1926 | PPTG_14025 | 147,69 | 66 | 41 | 66 | 66 | 4,73E+07 | 2,47E+08 | 4,83E+08 | 5 | 5 | 8 | 13 | 15 | Carbamidomethylation; Deamidation (NQ) | 11364 | PPTG_14025 | Phytophthora parasitica INRA-310 hypothetical protein (112 aa) |
| 1502 | 643 | PPTG_13761 | 147,68 | 20 | 2 | 0 | 20 | 1,86E+05 | | 5,51E+06 | 11 | 11 | 1 | 0 | 12 | Carbamidomethylation | 97792 | PPTG_13761 | Phytophthora parasitica INRA-310 voltage-gated Ion Channel (VIC) superfamily (884 aa) |
| 796 | 480 | PPTG_16174 | 147,61 | 23 | 11 | 11 | 23 | 2,43E+06 | 4,37E+06 | 2,79E+07 | 13 | 13 | 7 | 7 | 14 | Oxidation (M); Carbamidomethylation | 78197 | PPTG_16174 | Phytophthora parasitica INRA-310 hypothetical protein (694 aa) |
| 891 | 523 | PPTG_02205 | 147,58 | 20 | 4 | 2 | 20 | 3,46E+05 | 2,63E+05 | 9,26E+06 | 18 | 16 | 4 | 2 | 19 | Oxidation (M); Carbamidomethylation; Deamidation (NQ) | 123964 | PPTG_02205 | Phytophthora parasitica INRA-310 phospholipid-transporting ATPase (1108 aa) |
| 823 | 610 | PPTG_06550 | 147,53 | 17 | 4 | 8 | 17 | 9,79E+05 | 3,12E+06 | 1,28E+07 | 16 | 16 | 4 | 7 | 16 | Oxidation (M); Carbamidomethylation | 131429 | PPTG_06550 | Phytophthora parasitica INRA-310 myosin-like protein (1158 aa) |
| 800 | 530 | PPTG_04831 | 147,47 | 28 | 2 | 22 | 26 | 9,30E+04 | 5,40E+07 | 1,11E+07 | 16 | 16 | 1 | 13 | 14 | Carbamidomethylation | 77392 | PPTG_04831 | Phytophthora parasitica INRA-310 methylcrotonoyl-CoA carboxylase subunit alpha (703 aa) |
| 1109 | 540 | PPTG_00384 | 147,33 | 9 | 4 | 2 | 7 | 4,61E+05 | 8,96E+05 | 6,77E+06 | 14 | 14 | 5 | 4 | 10 | Oxidation (M) | 221808 | PPTG_00384 | Phytophthora parasitica INRA-310 myoferlin-like protein (1974 aa) |
| 699 | 640 | PPTG_19368 | 147,14 | 44 | 17 | 17 | 42 | 3,75E+06 | 6,50E+06 | 4,29E+07 | 12 | 12 | 7 | 8 | 17 | Oxidation (M); Dehydration | 37838 | PPTG_19368 | Phytophthora parasitica INRA-310 mitochondrial Carrier (MC) family (348 aa) |
| 1022 | 848 | PPTG_17386 | 147,14 | 26 | 8 | 24 | 23 | 1,23E+06 | 1,59E+07 | 2,86E+07 | 9 | 9 | 3 | 10 | 9 | Carbamidomethylation | 57594 | PPTG_17386 | Phytophthora parasitica INRA-310 hypothetical protein (546 aa) |
| 583 | 559 | PPTG_02222 | 147,12 | 20 | 8 | 16 | 17 | 2,87E+06 | 2,27E+07 | 3,92E+07 | 15 | 15 | 6 | 15 | 16 | Oxidation (M); Carbamidomethylation; Deamidation (NQ) | 92172 | PPTG_02222 | Phytophthora parasitica INRA-310 AP-1 complex subunit gamma-1 (850 aa) |
| 419 | 1596 | PPTG_13130 | 147,06 | 64 | 40 | 53 | 64 | 3,54E+06 | 9,73E+07 | 9,52E+07 | 7 | 7 | 5 | 24 | 20 | Oxidation (M); Dehydration; Deamidation (NQ); Acetylation (N-term) | 16825 | PPTG_13130 | Phytophthora parasitica INRA-310 calmodulin (150 aa) |
| 829 | 534 | PPTG_02310 | 147,06 | 32 | 3 | 30 | 21 | 1,04E+06 | 1,61E+07 | 3,29E+07 | 14 | 14 | 2 | 14 | 11 | Oxidation (M); Carbamidomethylation | 62635 | PPTG_02310 | Phytophthora parasitica INRA-310 T-complex protein 1 subunit eta (581 aa) |
| 1076 | 938 | PPTG_13124 | 146,99 | 25 | 10 | 13 | 24 | 7,01E+05 | 5,64E+06 | 1,07E+07 | 11 | 11 | 3 | 6 | 11 | Oxidation (M); Carbamidomethylation | 58821 | PPTG_13124 | Phytophthora parasitica INRA-310 T-complex protein 1 subunit epsilon (537 aa) |
| 867 | 1112 | PPTG_16034 | 146,97 | 36 | 19 | 29 | 36 | 2,67E+06 | 2,26E+07 | 3,91E+07 | 8 | 8 | 5 | 10 | 11 | Oxidation (M); Dehydration; Deamidation (NQ) | 30101 | PPTG_16034 | Phytophthora parasitica INRA-310 hypothetical protein (279 aa) |
| 880 | 976 | PPTG_07334 | 146,84 | 16 | 5 | 12 | 16 | 9,07E+05 | 4,21E+07 | 4,46E+07 | 8 | 8 | 3 | 11 | 12 | Oxidation (M); Carbamidomethylation | 69695 | PPTG_07334 | Phytophthora parasitica INRA-310 hypothetical protein (664 aa) |
| 1016 | 1592 | PPTG_00570 | 146,83 | 41 | 21 | 41 | 41 | | 7,08E+04 | 7,13E+05 | 5 | 1 | 3 | 9 | 10 | Oxidation (M); Carbamidomethylation; Acetylation (N-term) | 22648 | PPTG_00570 | Phytophthora parasitica INRA-310 SCP-like extracellular protein (213 aa) |
| 1016 | 1593 | PPTG_00570 | 146,83 | 41 | 21 | 41 | 41 | | 7,08E+04 | 7,13E+05 | 5 | 1 | 3 | 9 | 10 | Oxidation (M); Carbamidomethylation; Acetylation (N-term) | 22648 | PPTG_00570 | Phytophthora parasitica INRA-310 SCP-like extracellular protein (213 aa) |
| 585 | 581 | PPTG_14524 | 146,83 | 27 | 7 | 21 | 26 | 9,08E+05 | 1,31E+07 | 1,86E+07 | 17 | 17 | 4 | 16 | 17 | Oxidation (M); Carbamidomethylation | 76785 | PPTG_14524 | Phytophthora parasitica INRA-310 hypothetical protein (718 aa) |
| 825 | 585 | PPTG_10325 | 146,79 | 31 | 11 | 16 | 29 | 1,31E+06 | 1,00E+07 | 1,78E+07 | 14 | 14 | 5 | 8 | 14 | Oxidation (M); Carbamidomethylation | 57044 | PPTG_10325 | Phytophthora parasitica INRA-310 phenylalanyl-tRNA synthetase alpha chain (503 aa) |
| 751 | 698 | PPTG_05245 | 146,69 | 35 | 4 | 18 | 33 | 7,04E+05 | 9,07E+06 | 2,32E+07 | 17 | 17 | 2 | 10 | 18 | Oxidation (M); Carbamidomethylation | 59354 | PPTG_05245 | Phytophthora parasitica INRA-310 T-complex protein 1 zeta subunit (540 aa) |
| 871 | 594 | PPTG_11884 | 146,67 | 9 | 2 | 1 | 9 | 2,83E+05 | 7,16E+05 | 1,09E+07 | 19 | 19 | 4 | 3 | 19 | Carbamidomethylation | 251630 | PPTG_11884 | Phytophthora parasitica INRA-310 hypothetical protein (2276 aa) |
| 608 | 592 | PPTG_08060 | 146,49 | 32 | 11 | 18 | 32 | 1,18E+06 | 7,13E+06 | 2,02E+07 | 19 | 19 | 6 | 11 | 19 | Carbamidomethylation | 76277 | PPTG_08060 | Phytophthora parasitica INRA-310 2 4-dienoyl-CoA reductase (704 aa) |
| 828 | 902 | PPTG_13260 | 146,44 | 43 | 16 | 23 | 40 | 2,18E+06 | 5,06E+06 | 3,55E+07 | 10 | 10 | 5 | 8 | 14 | Carbamidomethylation; Deamidation (NQ) | 33425 | PPTG_13260 | Phytophthora parasitica INRA-310 hypothetical protein (308 aa) |
| 155 | 978 | PPTG_00988 | 146,43 | 44 | 37 | 39 | 43 | 2,59E+08 | 1,02E+09 | 2,55E+09 | 12 | 12 | 19 | 31 | 43 | Oxidation (M); Carbamidomethylation; Dehydration; Carbamylation | 23158 | PPTG_00988 | Phytophthora parasitica INRA-310 50S ribosomal protein L13 (204 aa) |
| 921 | 834 | PPTG_00112 | 146,41 | 40 | 18 | 26 | 40 | 2,30E+06 | 4,87E+06 | 2,86E+07 | 10 | 10 | 5 | 6 | 13 | Carbamidomethylation; Deamidation (NQ) | 32191 | PPTG_00112 | Phytophthora parasitica INRA-310 mitochondrial 2-oxoglutarate/malate carrier protein (307 aa) |
| 760 | 1347 | PPTG_12927 | 146,36 | 72 | 35 | 72 | 52 | 1,44E+07 | 1,32E+08 | 1,36E+08 | 6 | 6 | 5 | 15 | 10 | Oxidation (M); Carbamidomethylation; Dehydration; Deamidation (NQ); Acetylation (N-term) | 15949 | PPTG_12927 | Phytophthora parasitica INRA-310 actin-depolymerizing factor (144 aa) |
| 670 | 589 | PPTG_02509 | 146,23 | 34 | 8 | 31 | 28 | 2,71E+06 | 3,18E+07 | 3,68E+07 | 16 | 16 | 4 | 13 | 16 | Carbamidomethylation; Deamidation (NQ) | 54137 | PPTG_02509 | Phytophthora parasitica INRA-310 adenylosuccinate lyase (481 aa) |

| | | | | | | | | | | | | | | | | | | | |
|---|---|---|---|---|---|---|---|---|---|---|---|---|---|---|---|---|---|---|---|
| 68 | 956 | PPTG_11897 | 146,23 | 47 | 38 | 44 | 47 | 1,58E+08 | 6,28E+08 | 1,81E+09 | 9 | 9 | 24 | 39 | 66 | Carbamidomethylation; Dehydration; Deamidation (NQ); Carbamylation | 23499 | PPTG_11897 | Phytophthora parasitica INRA-310 40S ribosomal protein S8 (206 aa) |
| 797 | 535 | PPTG_12407 | 146,21 | 17 | 4 | 14 | 17 | 2,82E+05 | 4,03E+06 | 7,34E+06 | 14 | 14 | 3 | 12 | 13 | Carbamidomethylation; Carbamylation | 117197 | PPTG_12407 | Phytophthora parasitica INRA-310 nonsense transcript regulator (1053 aa) |
| 985 | 614 | PPTG_14134 | 146,18 | 19 | 7 | 4 | 16 | 6,47E+05 | 5,50E+05 | 6,90E+06 | 15 | 15 | 5 | 3 | 14 | Oxidation (M); Carbamidomethylation; Dehydration; Deamidation (NQ); Carbamylation | 108408 | PPTG_14134 | Phytophthora parasitica INRA-310 hypothetical protein (981 aa) |
| 107 | 836 | PPTG_02042 | 146,05 | 56 | 50 | 50 | 56 | 2,53E+08 | 9,90E+08 | 2,44E+09 | 10 | 10 | 26 | 36 | 44 | Carbamidomethylation; Dehydration; Deamidation (NQ); Carbamylation | 18286 | PPTG_02042 | Phytophthora parasitica INRA-310 40S ribosomal protein S16 (164 aa) |
| 726 | 739 | PPTG_12191 | 145,96 | 26 | 12 | 14 | 24 | 3,69E+06 | 4,32E+06 | 4,76E+07 | 12 | 12 | 8 | 7 | 16 | Oxidation (M); Carbamidomethylation | 68832 | PPTG_12191 | Phytophthora parasitica INRA-310 hypothetical protein (630 aa) |
| 908 | 1919 | PPTG_13482 | 145,92 | 22 | 14 | 22 | 22 | 1,64E+06 | 5,23E+07 | 4,17E+07 | 4 | 4 | 3 | 11 | 11 | Carbamidomethylation; Dehydration; Deamidation (NQ); Acetylation (N-term) | 36135 | PPTG_13482 | Phytophthora parasitica INRA-310 cellulose binding elicitor lectin (CBEL) (340 aa) |
| 665 | 663 | PPTG_10070 | 145,86 | 16 | 5 | 12 | 10 | 4,21E+05 | 4,20E+06 | 8,19E+06 | 18 | 18 | 6 | 14 | 13 | Carbamidomethylation | 144669 | PPTG_10070 | Phytophthora parasitica INRA-310 intraflagellar transporter 122 (1288 aa) |
| 738 | 740 | PPTG_07910 | 145,8 | 56 | 33 | 42 | 51 | 1,08E+07 | 6,76E+07 | 9,73E+07 | 12 | 12 | 8 | 10 | 13 | Carbamidomethylation; Deamidation (NQ) | 23469 | PPTG_07910 | Phytophthora parasitica INRA-310 proteasome subunit beta type-6 (219 aa) |
| 362 | 1154 | PPTG_10908 | 145,75 | 58 | 53 | 40 | 58 | 1,18E+08 | 4,46E+08 | 1,26E+09 | 8 | 8 | 11 | 17 | 28 | Oxidation (M); Dehydration; Carbamylation | 16473 | PPTG_10908 | Phytophthora parasitica INRA-310 40S ribosomal protein S10 (145 aa) |
| 501 | 722 | PPTG_17135 | 145,75 | 12 | 2 | 9 | 8 | 7,01E+05 | 6,48E+06 | 1,25E+07 | 25 | 25 | 4 | 18 | 19 | Carbamidomethylation | 258438 | PPTG_17135 | Phytophthora parasitica INRA-310 CCR4-NOT transcription complex subunit (2373 aa) |
| 942 | 587 | PPTG_01560 | 145,67 | 31 | 5 | 29 | 25 | 1,86E+05 | 5,48E+06 | 9,01E+06 | 11 | 11 | 3 | 11 | 10 | Oxidation (M); Carbamidomethylation; Acetylation (N-term) | 61392 | PPTG_01560 | Phytophthora parasitica INRA-310 chorismate mutase (559 aa) |
| 1070 | 844 | PPTG_13586 | 145,57 | 15 | 2 | 3 | 14 | 3,97E+05 | 4,37E+05 | 1,30E+07 | 15 | 15 | 3 | 3 | 14 | Carbamidomethylation | 133488 | PPTG_13586 | Phytophthora parasitica INRA-310 hypothetical protein (1181 aa) |
| 573 | 816 | PPTG_08286 | 145,19 | 35 | 17 | 29 | 35 | 2,00E+06 | 1,03E+07 | 3,45E+07 | 11 | 11 | 6 | 12 | 20 | Oxidation (M) | 37539 | PPTG_08286 | Phytophthora parasitica INRA-310 cytochrome c peroxidase (338 aa) |
| 735 | 941 | PPTG_12903 | 145,19 | 68 | 50 | 58 | 68 | 6,77E+06 | 1,44E+07 | 1,16E+08 | 10 | 10 | 6 | 11 | 14 | Oxidation (M); Carbamidomethylation | 16848 | PPTG_12903 | Phytophthora parasitica INRA-310 hypothetical protein (149 aa) |
| 1062 | 1179 | PPTG_06552 | 145,06 | 52 | 17 | 52 | 38 | 7,15E+05 | 9,36E+06 | 1,25E+07 | 7 | 7 | 3 | 10 | 8 | Oxidation (M); Carbamidomethylation; Dehydration; Deamidation (NQ) | 26304 | PPTG_06552 | Phytophthora parasitica INRA-310 eukaryotic translation initiation factor 6 (246 aa) |
| 715 | 574 | PPTG_02620 | 145,03 | 33 | 8 | 26 | 26 | 3,34E+06 | 3,32E+07 | 3,81E+07 | 16 | 16 | 4 | 13 | 14 | Carbamidomethylation; Deamidation (NQ); Carbamylation | 57730 | PPTG_02620 | Phytophthora parasitica INRA-310 aldehyde dehydrogenase family 7 member A1 (536 aa) |
| 1196 | 1921 | PPTG_10186 | 145 | 28 | 14 | 28 | 28 | 9,84E+05 | 1,88E+07 | 1,66E+07 | 4 | 4 | 2 | 8 | 8 | Oxidation (M); Carbamidomethylation; Dehydration | 29263 | PPTG_10186 | Phytophthora parasitica INRA-310 hypothetical protein (271 aa) |
| 681 | 1315 | PPTG_04295 | 144,96 | 40 | 23 | 40 | 40 | 1,27E+07 | 1,48E+08 | 1,26E+08 | 10 | 10 | 5 | 16 | 12 | Dehydration; Deamidation (NQ); Carbamylation | 24542 | PPTG_04295 | Phytophthora parasitica INRA-310 superoxide dismutase [Mn] mitochondrial (221 aa) |
| 894 | 690 | PPTG_09270 | 144,83 | 45 | 12 | 38 | 34 | 1,09E+06 | 7,79E+06 | 1,18E+07 | 11 | 11 | 4 | 11 | 10 | | 35516 | PPTG_09270 | Phytophthora parasitica INRA-310 hypothetical protein (320 aa) |
| 824 | 601 | PPTG_01706 | 144,57 | 26 | 11 | 26 | 21 | 6,27E+05 | 1,36E+07 | 1,87E+07 | 12 | 12 | 4 | 13 | 10 | Carbamidomethylation | 52287 | PPTG_01706 | Phytophthora parasitica INRA-310 orotidine 5'-phosphate decarboxylase (495 aa) |
| 360 | 1115 | PPTG_16727 | 144,55 | 50 | 41 | 45 | 36 | | 1,72E+06 | | 10 | 1 | 13 | 21 | 22 | Oxidation (M); Deamidation (NQ); Acetylation (N-term) | 23685 | PPTG_16727 | Phytophthora parasitica INRA-310 hypothetical protein (218 aa) |
| 1034 | 672 | PPTG_08316 | 144,49 | 41 | 24 | 15 | 41 | 3,95E+06 | 6,13E+06 | 4,45E+07 | 10 | 10 | 5 | 4 | 12 | Oxidation (M); Carbamidomethylation | 31100 | PPTG_08316 | Phytophthora parasitica INRA-310 mitochondrial Carrier (MC) family (293 aa) |
| 402 | 751 | PPTG_18250 | 144,36 | 25 | 23 | 24 | 25 | 1,08E+07 | 5,65E+07 | 1,29E+08 | 11 | 11 | 12 | 18 | 21 | Oxidation (M); Carbamidomethylation; Deamidation (NQ) | 46318 | PPTG_18250 | Phytophthora parasitica INRA-310 casein kinase (403 aa) |
| 923 | 520 | PPTG_11450 | 144,35 | 31 | 9 | 4 | 29 | 5,74E+05 | 2,01E+05 | 2,14E+07 | 14 | 14 | 5 | 2 | 17 | Oxidation (M); Carbamidomethylation | 66401 | PPTG_11450 | Phytophthora parasitica INRA-310 GAF domain-containing protein (598 aa) |
| 508 | 623 | PPTG_06467 | 144,32 | 29 | 4 | 24 | 26 | 3,51E+06 | 2,98E+07 | 5,88E+07 | 17 | 17 | 3 | 20 | 19 | Oxidation (M); Carbamidomethylation; Deamidation (NQ); Acetylation (N-term) | 72771 | PPTG_06467 | Phytophthora parasitica INRA-310 lysyl-tRNA synthetase (653 aa) |
| 769 | 719 | PPTG_06758 | 144,27 | 61 | 14 | 50 | 57 | | 2,46E+05 | 1,04E+06 | 12 | 2 | 3 | 11 | 15 | Oxidation (M); Deamidation (NQ); Amidation | 26245 | PPTG_06758 | Phytophthora parasitica INRA-310 short chain dehydrogenase (243 aa) |
| 980 | 1612 | PPTG_12759 | 144,2 | 12 | 12 | 12 | 4 | 3,90E+06 | 3,78E+07 | 3,87E+07 | 6 | 6 | 7 | 10 | 6 | Carbamidomethylation; Deamidation (NQ) | 103871 | PPTG_12759 | Phytophthora parasitica INRA-310 mucin-like protein (1047 aa) |
| 620 | 725 | PPTG_09393 | 144,1 | 12 | 4 | 9 | 10 | 1,47E+06 | 4,01E+06 | 1,01E+07 | 16 | 15 | 6 | 14 | 15 | Oxidation (M); Carbamidomethylation; Deamidation (NQ) | 161974 | PPTG_09393 | Phytophthora parasitica INRA-310 DEAD/DEAH box RNA helicase (1465 aa) |
| 554 | 615 | PPTG_14103 | 144,02 | 56 | 34 | 48 | 56 | 4,45E+06 | 2,62E+07 | 6,26E+07 | 11 | 11 | 7 | 12 | 20 | Oxidation (M); Deamidation (NQ) | 26146 | PPTG_14103 | Phytophthora parasitica INRA-310 hypothetical protein (250 aa) |
| 1005 | 864 | PPTG_02850 | 143,97 | 41 | 12 | 30 | 30 | 5,23E+05 | 4,36E+06 | 1,08E+07 | 11 | 11 | 3 | 9 | 10 | Carbamidomethylation | 45441 | PPTG_02850 | Phytophthora parasitica INRA-310 hypothetical protein (402 aa) |
| 235 | 745 | PPTG_11464 | 143,83 | 57 | 42 | 57 | 54 | 8,52E+07 | 9,67E+08 | 7,69E+08 | 13 | 13 | 13 | 36 | 27 | Oxidation (M); Carbamidomethylation; Deamidation (NQ); Carbamylation | 30457 | PPTG_11464 | Phytophthora parasitica INRA-310 hypothetical protein (280 aa) |
| 679 | 546 | PPTG_09918 | 143,78 | 49 | 14 | 49 | 49 | 1,09E+06 | 1,05E+07 | 2,32E+07 | 11 | 11 | 4 | 14 | 15 | Oxidation (M); Carbamidomethylation; Deamidation (NQ) | 39321 | PPTG_09918 | Phytophthora parasitica INRA-310 mannose-1-phosphate guanyltransferase beta (360 aa) |
| 764 | 746 | PPTG_16221 | 143,74 | 12 | 4 | 6 | 11 | 5,10E+05 | 4,10E+06 | 1,05E+07 | 16 | 15 | 5 | 9 | 15 | Carbamidomethylation; Deamidation (NQ); Carbamylation | 153709 | PPTG_16221 | Phytophthora parasitica INRA-310 myosin-like protein (1364 aa) |
| 706 | 694 | PPTG_11383 | 143,67 | 43 | 34 | 37 | 43 | 1,00E+07 | 1,28E+07 | 9,63E+07 | 10 | 10 | 8 | 9 | 15 | | 30608 | PPTG_11383 | Phytophthora parasitica INRA-310 prohibitin (276 aa) |
| 514 | 1607 | PPTG_10318 | 143,59 | 60 | 41 | 54 | 60 | 9,34E+07 | 5,50E+08 | 1,44E+09 | 7 | 7 | 8 | 13 | 21 | Carbamidomethylation; Deamidation (NQ); Amidation; Carbamylation | 14324 | PPTG_10318 | Phytophthora parasitica INRA-310 60S ribosomal protein L22 (126 aa) |
| 1448 | 1582 | PPTG_00652 | 143,56 | 61 | 48 | 12 | 61 | 2,59E+06 | 3,54E+06 | 2,76E+07 | 7 | 7 | 4 | 2 | 8 | Carbamidomethylation | 21069 | PPTG_00652 | Phytophthora parasitica INRA-310 hypothetical protein (191 aa) |
| 1115 | 602 | PPTG_07419 | 143,48 | 14 | 1 | 2 | 14 | 5,09E+04 | 3,12E+05 | 6,92E+06 | 15 | 15 | 1 | 3 | 15 | Carbamidomethylation | 137484 | PPTG_07419 | Phytophthora parasitica INRA-310 hypothetical protein (1227 aa) |
| 304 | 1790 | PPTG_11004 | 143,45 | 50 | 23 | 39 | 50 | 8,83E+07 | 4,36E+08 | 1,22E+09 | 9 | 9 | 14 | 19 | 31 | Oxidation (M); Dehydration; Deamidation (NQ); Acetylation (N-term); Carbamylation | 15667 | PPTG_11004 | Phytophthora parasitica INRA-310 40S ribosomal protein (138 aa) |
| 947 | 649 | PPTG_03815 | 143,45 | 40 | 7 | 33 | 27 | 1,43E+05 | 9,22E+06 | 6,98E+06 | 12 | 12 | 2 | 12 | 10 | | 45714 | PPTG_03815 | Phytophthora parasitica INRA-310 glycine amidinotransferase (397 aa) |
| 787 | 767 | PPTG_09934 | 143,42 | 22 | 3 | 22 | 15 | | 5,73E+05 | 3,48E+04 | 12 | 2 | 2 | 17 | 10 | Carbamidomethylation; Dehydration; Deamidation (NQ) | 68895 | PPTG_09934 | Phytophthora parasitica INRA-310 glycoside hydrolase (621 aa) |
| 1026 | 1645 | PPTG_17633 | 143,42 | 35 | 20 | 35 | 35 | 5,64E+05 | 8,29E+06 | 1,31E+07 | 8 | 8 | 4 | 9 | 9 | | 27944 | PPTG_17633 | Phytophthora parasitica INRA-310 50S ribosomal protein L25 (252 aa) |
| 705 | 733 | PPTG_19057 | 143,31 | 43 | 23 | 38 | 40 | 1,35E+06 | 1,20E+07 | 1,09E+07 | 11 | 11 | 5 | 15 | 11 | Oxidation (M); Carbamidomethylation | 42538 | PPTG_19057 | Phytophthora parasitica INRA-310 glycine cleavage system T protein (392 aa) |
| 682 | 544 | PPTG_18216 | 143,05 | 27 | 6 | 20 | 27 | 6,36E+05 | 1,01E+07 | 1,97E+07 | 13 | 13 | 3 | 13 | 17 | Oxidation (M); Carbamidomethylation; Deamidation (NQ) | 76543 | PPTG_18216 | Phytophthora parasitica INRA-310 hypothetical protein (703 aa) |
| 901 | 761 | PPTG_05195 | 142,99 | 18 | 12 | 11 | 18 | 1,47E+06 | 1,87E+06 | 2,20E+07 | 11 | 11 | 6 | 6 | 13 | Oxidation (M); Carbamidomethylation | 69401 | PPTG_05195 | Phytophthora parasitica INRA-310 translocation protein SEC63 (607 aa) |
| 667 | 606 | PPTG_11829 | 142,97 | 15 | 3 | 10 | 13 | 3,54E+05 | 5,12E+06 | 1,04E+07 | 17 | 17 | 4 | 12 | 17 | Oxidation (M); Carbamidomethylation | 145811 | PPTG_11829 | Phytophthora parasitica INRA-310 hypothetical protein (1288 aa) |
| 809 | 619 | PPTG_08040 | 142,96 | 30 | 16 | 23 | 21 | 1,93E+06 | 1,33E+07 | 2,58E+07 | 13 | 12 | 7 | 10 | 11 | Carbamidomethylation; Dehydration; Deamidation (NQ) | 54500 | PPTG_08040 | Phytophthora parasitica INRA-310 acetyl-coenzyme A synthetase (487 aa) |
| 741 | 850 | PPTG_12094 | 142,94 | 18 | 4 | 18 | 12 | 6,17E+05 | 1,54E+07 | 1,15E+07 | 14 | 14 | 4 | 15 | 12 | Carbamidomethylation; Dehydration; Deamidation (NQ) | 106242 | PPTG_12094 | Phytophthora parasitica INRA-310 beta-mannosidase (938 aa) |
| 736 | 543 | PPTG_18748 | 142,9 | 29 | 11 | 7 | 28 | 1,37E+06 | 2,16E+06 | 2,26E+07 | 15 | 15 | 7 | 6 | 18 | Oxidation (M); Carbamidomethylation; Dehydration | 83991 | PPTG_18748 | Phytophthora parasitica INRA-310 hypothetical protein (741 aa) |
| 840 | 1352 | PPTG_10656 | 142,87 | 15 | 8 | 15 | 11 | 2,16E+06 | 3,46E+07 | 2,91E+07 | 6 | 6 | 5 | 12 | 10 | Oxidation (M); Carbamidomethylation; Dehydration; Deamidation (NQ); Carbamylation | 68718 | PPTG_10656 | Phytophthora parasitica INRA-310 elicitor-like transglutaminase (655 aa) |
| 1017 | 563 | PPTG_02675 | 142,87 | 21 | 5 | 17 | 16 | 6,37E+05 | 7,93E+06 | 8,18E+06 | 12 | 12 | 3 | 10 | 9 | Carbamidomethylation | 79758 | PPTG_02675 | Phytophthora parasitica INRA-310 glutamine-dependent NAD(+) synthetase (716 aa) |
| 759 | 1138 | PPTG_15910 | 142,82 | 50 | 23 | 50 | 48 | 3,94E+06 | 5,98E+07 | 6,52E+07 | 8 | 8 | 3 | 14 | 13 | Oxidation (M); Carbamidomethylation; Deamidation (NQ) | 26240 | PPTG_15910 | Phytophthora parasitica INRA-310 proteasome subunit beta (232 aa) |
| 481 | 608 | PPTG_06210 | 142,76 | 25 | 12 | 24 | 22 | 2,06E+06 | 1,44E+07 | 2,51E+07 | 14 | 14 | 9 | 16 | 19 | Oxidation (M); Carbamidomethylation; Deamidation (NQ) | 75043 | PPTG_06210 | Phytophthora parasitica INRA-310 acyl-CoA synthetase short-chain family member (680 aa) |
| 1208 | 840 | PPTG_06410 | 142,75 | 9 | 0 | 3 | 8 | | 1,01E+06 | 6,84E+06 | 10 | 10 | 0 | 5 | 12 | Carbamidomethylation | 166502 | PPTG_06410 | Phytophthora parasitica INRA-310 leishmanolysin-like peptidase (1537 aa) |
| 997 | 779 | PPTG_05918 | 142,61 | 22 | 6 | 18 | 15 | 7,20E+05 | 8,06E+06 | 1,48E+07 | 11 | 11 | 4 | 9 | 9 | Oxidation (M); Carbamidomethylation | 76436 | PPTG_05918 | Phytophthora parasitica INRA-310 3-isopropylmalate dehydratase large subunit (710 aa) |
| 978 | 1418 | PPTG_02297 | 142,53 | 55 | 23 | 49 | 55 | 1,84E+06 | 3,36E+07 | 3,18E+07 | 6 | 6 | 3 | 10 | 10 | Oxidation (M); Carbamidomethylation | 16281 | PPTG_02297 | Phytophthora parasitica INRA-310 hypothetical protein (153 aa) |
| 830 | 680 | PPTG_11571 | 142,49 | 27 | 4 | 24 | 17 | 1,11E+06 | 7,98E+06 | 1,22E+07 | 14 | 14 | 3 | 13 | 11 | Oxidation (M); Carbamidomethylation; Deamidation (NQ) | 77452 | PPTG_11571 | Phytophthora parasitica INRA-310 transcriptional repressor TUP1-like protein (726 aa) |
| 1085 | 731 | PPTG_15123 | 142,48 | 27 | 7 | 11 | 27 | 1,55E+06 | 5,60E+06 | 2,82E+07 | 10 | 10 | 3 | 5 | 12 | Carbamidomethylation | 60313 | PPTG_15123 | Phytophthora parasitica INRA-310 vacuolar-sorting receptor (547 aa) |
| 933 | 634 | PPTG_09469 | 142,42 | 27 | 6 | 7 | 27 | 5,36E+05 | 2,04E+06 | 2,01E+07 | 14 | 14 | 4 | 3 | 17 | Oxidation (M); Carbamidomethylation | 81095 | PPTG_09469 | Phytophthora parasitica INRA-310 glycerol-3-phosphate O-acyltransferase (729 aa) |
| 117 | 1146 | PPTG_15525 | 142,36 | 52 | 50 | 51 | 52 | 2,80E+08 | 1,02E+09 | 2,63E+09 | 9 | 9 | 21 | 37 | 46 | Dehydration; Deamidation (NQ); Carbamylation | 15186 | PPTG_15525 | Phytophthora parasitica INRA-310 60S ribosomal protein L14 (137 aa) |
| 806 | 1161 | PPTG_16775 | 142,32 | 31 | 16 | 23 | 24 | 2,43E+06 | 1,12E+07 | 2,16E+07 | 8 | 8 | 6 | 9 | 13 | Oxidation (M); Carbamidomethylation; Deamidation (NQ) | 39729 | PPTG_16775 | Phytophthora parasitica INRA-310 adenosine kinase (369 aa) |
| 822 | 875 | PPTG_13862 | 142,23 | 53 | 18 | 27 | 53 | 3,24E+06 | 8,44E+06 | 4,63E+07 | 13 | 13 | 5 | 7 | 15 | Deamidation (NQ) | 28919 | PPTG_13862 | Phytophthora parasitica INRA-310 V-type ATPase D subunit (261 aa) |
| 572 | 566 | PPTG_07617 | 142,22 | 26 | 13 | 23 | 21 | 2,15E+06 | 1,76E+07 | 1,61E+07 | 14 | 11 | 7 | 15 | 16 | Oxidation (M); Carbamidomethylation; Dehydration; Deamidation (NQ); Carbamylation | 61261 | PPTG_07617 | Phytophthora parasitica INRA-310 hypothetical protein (545 aa) |
| 1069 | 562 | PPTG_13731 | 142,2 | 20 | 4 | 6 | 19 | 1,92E+05 | 8,10E+05 | 1,08E+07 | 14 | 14 | 3 | 4 | 13 | Carbamidomethylation | 101280 | PPTG_13731 | Phytophthora parasitica INRA-310 phosphatidylinositol kinase (PIK-H1) (907 aa) |
| 627 | 1364 | PPTG_18717 | 142,11 | 47 | 42 | 42 | 47 | 2,26E+07 | 9,06E+07 | 2,74E+08 | 7 | 6 | 8 | 10 | 17 | Oxidation (M); Carbamidomethylation | 16065 | PPTG_18717 | Phytophthora parasitica INRA-310 40S ribosomal protein S23 (146 aa) |
| 1042 | 715 | PPTG_03967 | 142,1 | 32 | 7 | 21 | 28 | 6,73E+05 | 4,29E+05 | 4,36E+06 | 12 | 6 | 3 | 7 | 11 | Carbamidomethylation | 52862 | PPTG_03967 | Phytophthora parasitica INRA-310 50S ribosomal protein L1 (493 aa) |
| 1567 | 1588 | PPTG_09242 | 142,01 | 40 | 7 | 7 | 40 | 5,79E+05 | 8,12E+05 | 1,26E+07 | 5 | 5 | 1 | 1 | 10 | Oxidation (M); Carbamidomethylation; Deamidation (NQ) | 27081 | PPTG_09242 | Phytophthora parasitica INRA-310 NADH dehydrogenase [ubiquinone] iron-sulfur protein 4 (243 aa) |
| 700 | 710 | PPTG_15752 | 141,98 | 39 | 30 | 34 | 39 | 9,01E+06 | 7,12E+06 | 6,07E+07 | 10 | 10 | 9 | 10 | 13 | Oxidation (M); Dehydration; Deamidation (NQ) | 33237 | PPTG_15752 | Phytophthora parasitica INRA-310 prohibitin-2 (300 aa) |
| 957 | 667 | PPTG_11140 | 141,9 | 13 | 3 | 12 | 11 | 1,36E+05 | 4,55E+06 | 4,00E+06 | 12 | 12 | 2 | 11 | 10 | Oxidation (M); Carbamidomethylation | 125431 | PPTG_11140 | Phytophthora parasitica INRA-310 exportin-like protein (1115 aa) |
| 1174 | 1127 | PPTG_16175 | 141,89 | 28 | 11 | 10 | 28 | 3,05E+06 | 2,84E+06 | 2,63E+07 | 10 | 10 | 4 | 3 | 11 | Oxidation (M); Carbamidomethylation | 40263 | PPTG_16175 | Phytophthora parasitica INRA-310 hypothetical protein (358 aa) |
| 999 | 575 | PPTG_11732 | 141,74 | 36 | 5 | 29 | 24 | 8,86E+04 | 4,31E+06 | 4,72E+06 | 13 | 13 | 2 | 12 | 8 | Carbamidomethylation; Dehydration; Deamidation (NQ) | 57716 | PPTG_11732 | Phytophthora parasitica INRA-310 diaminopimelate decarboxylase (528 aa) |
| 961 | 716 | PPTG_15500 | 141,74 | 40 | 16 | 32 | 37 | 2,87E+05 | 9,57E+06 | 8,75E+06 | 11 | 11 | 4 | 9 | 10 | Carbamidomethylation | 36761 | PPTG_15500 | Phytophthora parasitica INRA-310 WD domain-containing protein (342 aa) |
| 152 | 1035 | PPTG_09132 | 141,68 | 56 | 37 | 48 | 56 | 1,56E+08 | 6,93E+08 | 1,74E+09 | 11 | 11 | 18 | 33 | 43 | Oxidation (M); Carbamidomethylation; Dehydration; Deamidation (NQ); Carbamylation | 17247 | PPTG_09132 | Phytophthora parasitica INRA-310 40S ribosomal protein S13 (152 aa) |
| 1170 | 1004 | PPTG_16307 | 141,56 | 38 | 6 | 6 | 38 | 1,20E+06 | 1,96E+06 | 2,03E+07 | 11 | 11 | 2 | 2 | 14 | Carbamidomethylation; Dehydration | 39172 | PPTG_16307 | Phytophthora parasitica INRA-310 hypothetical protein (340 aa) |
| 94 | 881 | PPTG_16153 | 141,5 | 62 | 61 | 57 | 62 | 2,72E+08 | 1,15E+09 | 2,64E+09 | 14 | 14 | 24 | 36 | 52 | Oxidation (M); Carbamidomethylation; Dehydration; Deamidation (NQ); Acetylation (N-term); Carbam | 17579 | PPTG_16153 | Phytophthora parasitica INRA-310 40S ribosomal protein S18 (153 aa) |
| 860 | 979 | PPTG_19298 | 141,49 | 37 | 18 | 34 | 37 | 5,52E+06 | 2,63E+07 | 4,84E+07 | 9 | 9 | 5 | 9 | 13 | Oxidation (M) | 34762 | PPTG_19298 | Phytophthora parasitica INRA-310 26S proteasome non-ATPase regulatory subunit 14 (312 aa) |
| 1162 | 664 | PPTG_15285 | 141,36 | 15 | 4 | 1 | 15 | 2,87E+05 | 2,71E+05 | 6,01E+06 | 12 | 6 | 3 | 1 | 14 | Carbamidomethylation; Deamidation (NQ) | 100482 | PPTG_15285 | Phytophthora parasitica INRA-310 transmembrane protein (886 aa) |
| 729 | 616 | PPTG_02483 | 141,28 | 44 | 17 | 32 | 42 | 2,70E+06 | 1,29E+07 | 2,84E+07 | 13 | 13 | 6 | 11 | 14 | Oxidation (M); Carbamidomethylation; Deamidation (NQ) | 41702 | PPTG_02483 | Phytophthora parasitica INRA-310 hypothetical protein (380 aa) |
| 581 | 808 | PPTG_05693 | 141,23 | 11 | 1 | 8 | 8 | 4,20E+05 | 6,17E+06 | 1,42E+07 | 22 | 22 | 3 | 16 | 18 | | 257715 | PPTG_05693 | Phytophthora parasitica INRA-310 pre-mRNA-splicing helicase BRR2 (2293 aa) |
| 124 | 1927 | PPTG_17494 | 141,15 | 53 | 36 | 53 | 53 | 6,58E+07 | 3,20E+08 | 1,20E+09 | 6 | 6 | 16 | 28 | 57 | Oxidation (M); Carbamidomethylation; Dehydration; Deamidation (NQ); Acetylation (N-term); Carbam | 12261 | PPTG_17494 | Phytophthora parasitica INRA-310 60S ribosomal protein L30 (113 aa) |
| 1064 | 1939 | PPTG_03256 | 141,09 | 69 | 30 | 62 | 69 | 3,96E+06 | 1,82E+07 | 7,24E+07 | 8 | 8 | 2 | 6 | 13 | Carbamidomethylation | 9911 | PPTG_03256 | Phytophthora parasitica INRA-310 hypothetical protein (85 aa) |
| 926 | 651 | PPTG_18612 | 141,04 | 27 | 1 | 24 | 18 | 5,73E+04 | 4,37E+06 | 4,15E+06 | 13 | 13 | 1 | 12 | 11 | | 71557 | PPTG_18612 | Phytophthora parasitica INRA-310 trehalase (639 aa) |
| 696 | 769 | PPTG_08373 | 141,02 | 36 | 12 | 33 | 36 | 3,05E+06 | 2,08E+07 | 4,05E+07 | 11 | 11 | 4 | 12 | 16 | Oxidation (M) | 38007 | PPTG_08373 | Phytophthora parasitica INRA-310 eukaryotic translation initiation factor 2 subunit alpha (333 aa) |
| 723 | 586 | PPTG_03695 | 140,98 | 23 | 6 | 15 | 22 | 6,10E+05 | 4,07E+06 | 5,82E+07 | 15 | 15 | 4 | 10 | 17 | Oxidation (M); Carbamidomethylation | 83097 | PPTG_03695 | Phytophthora parasitica INRA-310 hypothetical protein (724 aa) |
| 900 | 682 | PPTG_10236 | 140,96 | 28 | 5 | 9 | 26 | 5,08E+05 | 9,23E+05 | 2,10E+07 | 16 | 16 | 3 | 5 | 17 | Oxidation (M); Carbamidomethylation; Deamidation (NQ) | 79844 | PPTG_10236 | Phytophthora parasitica INRA-310 fatty-acid-CoA ligase (736 aa) |
| 900 | 683 | PPTG_10236 | 140,96 | 27 | 5 | 9 | 26 | 5,08E+05 | 9,23E+05 | 2,10E+07 | 16 | 16 | 3 | 5 | 17 | Oxidation (M); Carbamidomethylation; Deamidation (NQ) | 82118 | PPTG_10236 | Phytophthora parasitica INRA-310 fatty-acid-CoA ligase (756 aa) |
| 1135 | 1092 | PPTG_15937 | 140,89 | 24 | 2 | 19 | 24 | 3,71E+05 | 1,42E+07 | 1,20E+07 | 7 | 6 | 1 | 9 | 9 | Oxidation (M); Carbamidomethylation | 56713 | PPTG_15937 | Phytophthora parasitica INRA-310 glycoside hydrolase (518 aa) |
| 1056 | 1389 | PPTG_02290 | 140,88 | 77 | 31 | 50 | 77 | 3,31E+06 | 3,64E+06 | 3,68E+07 | 7 | 7 | 2 | 6 | 13 | Oxidation (M); Dehydration; Deamidation (NQ); Acetylation (N-term) | 11796 | PPTG_02290 | Phytophthora parasitica INRA-310 hypothetical protein (105 aa) |
| 995 | 846 | PPTG_08145 | 140,81 | 33 | 11 | 31 | 19 | 9,58E+04 | 9,92E+06 | 9,21E+06 | 11 | 11 | 3 | 11 | 8 | Oxidation (M); Carbamidomethylation | 44892 | PPTG_08145 | Phytophthora parasitica INRA-310 4-aminobutyrate aminotransferase (421 aa) |
| 629 | 565 | PPTG_10143 | 140,72 | 18 | 8 | 12 | 16 | 1,38E+06 | 8,10E+06 | 1,27E+07 | 13 | 13 | 7 | 12 | 16 | Oxidation (M); Carbamidomethylation | 110937 | PPTG_10143 | Phytophthora parasitica INRA-310 alpha-aminoadipic semialdehyde synthase (1015 aa) |
| 834 | 1009 | PPTG_00708 | 140,69 | 23 | 12 | 22 | 18 | 9,76E+05 | 9,08E+06 | 1,30E+07 | 8 | 8 | 6 | 10 | 11 | Oxidation (M); Carbamidomethylation | 58520 | PPTG_00708 | Phytophthora parasitica INRA-310 importin alpha-2 subunit (533 aa) |
| 791 | 742 | PPTG_09327 | 140,66 | 48 | 14 | 43 | 41 | 8,76E+05 | 1,08E+07 | 9,45E+06 | 11 | 11 | 4 | 12 | 13 | Oxidation (M); Carbamidomethylation | 36356 | PPTG_09327 | Phytophthora parasitica INRA-310 delta-aminolevulinic acid dehydratase (328 aa) |
| 1438 | 1664 | PPTG_00418 | 140,62 | 23 | 3 | 23 | 15 | 1,66E+05 | 8,94E+06 | 2,89E+06 | 5 | 5 | 1 | 7 | 6 | Carbamidomethylation | 45631 | PPTG_00418 | Phytophthora parasitica INRA-310 hypothetical protein (439 aa) |
| 644 | 576 | PPTG_00608 | 140,6 | 28 | 10 | 24 | 23 | 1,26E+06 | 1,08E+07 | 1,52E+07 | 12 | 12 | 5 | 15 | 14 | Oxidation (M); Carbamidomethylation; Deamidation (NQ) | 63182 | PPTG_00608 | Phytophthora parasitica INRA-310 asparagine synthetase (564 aa) |
| 838 | 660 | PPTG_16326 | 140,6 | 24 | 10 | 23 | 18 | 6,86E+05 | 7,66E+06 | 7,64E+06 | 12 | 12 | 5 | 12 | 10 | Carbamidomethylation | 60604 | PPTG_16326 | Phytophthora parasitica INRA-310 tryptophan biosynthesis protein TRP1 (562 aa) |
| 963 | 611 | PPTG_18241 | 140,53 | 26 | 12 | 9 | 26 | 7,67E+05 | 9,42E+05 | 1,48E+07 | 12 | 12 | 6 | 4 | 13 | Carbamidomethylation; Deamidation (NQ) | 78971 | PPTG_18241 | Phytophthora parasitica INRA-310 phosphatidylinositol-3 4 5-trisphosphate 5-phosphatase (714 aa) |
| 720 | 950 | PPTG_16234 | 140,51 | 17 | 6 | 15 | 17 | 3,13E+06 | 4,23E+07 | 4,22E+07 | 9 | 7 | 4 | 15 | 12 | Oxidation (M); Dehydration | 81201 | PPTG_16234 | Phytophthora parasitica INRA-310 transglutaminase elicitor (769 aa) |
| 1396 | 3176 | PPTG_19310 | 140,51 | 36 | 36 | 36 | 36 | 3,48E+06 | 2,34E+07 | 1,35E+08 | 5 | 5 | 4 | 6 | 5 | | 13017 | PPTG_19310 | Phytophthora parasitica INRA-310 hypothetical protein (119 aa) |
| 925 | 638 | PPTG_06097 | 140,48 | 29 | 0 | 19 | 27 | | 8,97E+06 | 1,45E+07 | 14 | 14 | 0 | 9 | 15 | Deamidation (NQ) | 73469 | PPTG_06097 | Phytophthora parasitica INRA-310 peptidase (663 aa) |
| 587 | 781 | PPTG_06606 | 140,45 | 34 | 12 | 28 | 32 | 4,65E+06 | 2,90E+07 | 5,40E+07 | 12 | 12 | 6 | 14 | 17 | Oxidation (M); Carbamidomethylation; Deamidation (NQ) | 42902 | PPTG_06606 | Phytophthora parasitica INRA-310 hypothetical protein (392 aa) |
| 789 | 826 | PPTG_13151 | 140,44 | 17 | 12 | 14 | 17 | 4,97E+06 | 1,38E+07 | 6,13E+07 | 9 | 9 | 7 | 10 | 12 | Oxidation (M); Carbamidomethylation; Deamidation (NQ) | 72439 | PPTG_13151 | Phytophthora parasitica INRA-310 endomembrane protein 70-like protein (644 aa) |
| 522 | 1350 | PPTG_14216 | 140,38 | 44 | 37 | 44 | 44 | 1,57E+06 | 2,28E+07 | 3,23E+07 | 6 | 6 | 5 | 19 | 17 | Oxidation (M); Carbamidomethylation; Deamidation (NQ); Acetylation (N-term) | 22637 | PPTG_14216 | Phytophthora parasitica INRA-310 proteasome subunit beta type-3 (205 aa) |
| 948 | 784 | PPTG_03220 | 140,34 | 25 | 11 | 21 | 23 | 1,77E+06 | 8,00E+06 | 1,83E+07 | 10 | 10 | 5 | 8 | 11 | | 57655 | PPTG_03220 | Phytophthora parasitica INRA-310 glycoside hydrolase (537 aa) |
| 744 | 669 | PPTG_13960 | 140,24 | 14 | 3 | 10 | 13 | 6,46E+05 | 4,12E+06 | 9,69E+06 | 12 | 12 | 3 | 11 | 16 | Oxidation (M); Deamidation (NQ) | 111841 | PPTG_13960 | Phytophthora parasitica INRA-310 AP-3 complex subunit delta (1005 aa) |
| 601 | 1166 | PPTG_02092 | 140,18 | 59 | 10 | 59 | 59 | 7,73E+04 | 1,35E+07 | 2,49E+07 | 8 | 8 | 1 | 15 | 20 | Oxidation (M); Carbamidomethylation; Deamidation (NQ); Amidation | 22101 | PPTG_02092 | Phytophthora parasitica INRA-310 peptidyl-prolyl cis-trans isomerase B (207 aa) |
| 745 | 622 | PPTG_02293 | 140,13 | 32 | 2 | 32 | 16 | 1,59E+05 | 8,90E+06 | 7,21E+06 | 14 | 14 | 1 | 18 | 11 | Oxidation (M); Carbamidomethylation; Deamidation (NQ) | 68817 | PPTG_02293 | Phytophthora parasitica INRA-310 NAD-dependent malic enzyme (623 aa) |
| 938 | 818 | PPTG_04798 | 140,11 | 42 | 3 | 24 | 42 | 2,00E+05 | 3,61E+06 | 2,37E+07 | 11 | 11 | 1 | 7 | 16 | Oxidation (M); Carbamidomethylation | 42289 | PPTG_04798 | Phytophthora parasitica INRA-310 pyridine nucleotide-disulphide oxidoreductase (395 aa) |

| | | | | | | | | | | | | | | | | | | | |
|---|---|---|---|---|---|---|---|---|---|---|---|---|---|---|---|---|---|---|---|
| 938 | 819 | PPTG_04800 | 140,11 | 42 | 3 | 24 | 42 | 2,00E+05 | 3,61E+06 | 2,37E+07 | 11 | 11 | 1 | 7 | 16 | Oxidation (M); Carbamidomethylation | 42303 | PPTG_04800 | Phytophthora parasitica INRA-310 pyridine nucleotide-disulphide oxidoreductase (395 aa) |
| 807 | 721 | PPTG_02740 | 140,04 | 48 | 31 | 44 | 48 | 6,72E+06 | 1,04E+07 | 8,02E+07 | 10 | 10 | 6 | 10 | 12 | Oxidation (M); Carbamidomethylation; Deamidation (NQ) | 25429 | PPTG_02740 | Phytophthora parasitica INRA-310 hypothetical protein (231 aa) |
| 1351 | 1524 | PPTG_02823 | 140,03 | 23 | 5 | 16 | 23 | 5,87E+05 | 1,21E+07 | 2,80E+07 | 6 | 6 | 1 | 5 | 10 | Oxidation (M); Carbamidomethylation; Acetylation (N-term) | 41609 | PPTG_02823 | Phytophthora parasitica INRA-310 GPI-anchored serine rich tenascin-like glycoprotein (408 aa) |
| 848 | 712 | PPTG_16771 | 140,02 | 33 | 4 | 33 | 28 | 7,45E+05 | 9,64E+06 | 6,77E+06 | 13 | 11 | 2 | 15 | 10 | Carbamidomethylation | 55979 | PPTG_16771 | Phytophthora parasitica INRA-310 cystathionine beta-synthase (514 aa) |
| 912 | 673 | PPTG_00422 | 140,01 | 33 | 7 | 27 | 32 | 4,35E+05 | 7,53E+06 | 1,41E+07 | 11 | 11 | 2 | 10 | 12 | Oxidation (M); Carbamidomethylation; Acetylation (N-term) | 47910 | PPTG_00422 | Phytophthora parasitica INRA-310 DnaJ subfamily C member 2 protein (421 aa) |
| 779 | 678 | PPTG_02494 | 139,99 | 53 | 19 | 53 | 46 | 1,23E+06 | 1,32E+07 | 1,40E+07 | 10 | 10 | 5 | 13 | 11 | Oxidation (M); Carbamidomethylation | 31578 | PPTG_02494 | Phytophthora parasitica INRA-310 3-hydroxyisobutyrate dehydrogenase (301 aa) |
| 1021 | 1121 | PPTG_17678 | 139,83 | 16 | 11 | 14 | 16 | 5,45E+06 | 9,71E+06 | 4,33E+07 | 7 | 7 | 5 | 6 | 11 | Carbamidomethylation | 78212 | PPTG_17678 | Phytophthora parasitica INRA-310 croquemort-like mating protein M82 (725 aa) |
| 1110 | 727 | PPTG_08439 | 139,82 | 6 | 2 | 1 | 6 | 4,48E+05 | 0 | 7,96E+06 | 14 | 13 | 5 | 1 | 13 | Carbamidomethylation | 293933 | PPTG_08439 | Phytophthora parasitica INRA-310 phospholipid-transporting ATPase (2629 aa) |
| 882 | 838 | PPTG_00131 | 139,71 | 36 | 18 | 32 | 36 | 1,53E+06 | 9,41E+06 | 1,75E+07 | 9 | 9 | 6 | 9 | 11 | | 34381 | PPTG_00131 | Phytophthora parasitica INRA-310 hypothetical protein (298 aa) |
| 1277 | 726 | PPTG_18713 | 139,6 | 14 | 6 | 2 | 13 | 3,76E+05 | 9,73E+04 | 6,04E+06 | 10 | 10 | 4 | 1 | 11 | Oxidation (M); Carbamidomethylation | 96861 | PPTG_18713 | Phytophthora parasitica INRA-310 cAMP-specific 3' 5'-cyclic phosphodiesterase (856 aa) |
| 671 | 713 | PPTG_04233 | 139,58 | 13 | 4 | 5 | 13 | 7,84E+05 | 3,49E+06 | 1,15E+07 | 18 | 18 | 5 | 8 | 20 | Carbamidomethylation; Deamidation (NQ) | 186488 | PPTG_04233 | Phytophthora parasitica INRA-310 UDP-glucose:glycoprotein glucosyltransferase (1658 aa) |
| 373 | 1238 | PPTG_15670 | 139,48 | 51 | 31 | 39 | 51 | 6,77E+07 | 3,10E+08 | 9,19E+08 | 10 | 10 | 8 | 18 | 29 | Oxidation (M); Dehydration; Deamidation (NQ); Carbamylation | 17399 | PPTG_15670 | Phytophthora parasitica INRA-310 30S ribosomal protein S19e (155 aa) |
| 887 | 720 | PPTG_11812 | 139,37 | 7 | 4 | 5 | 6 | 6,66E+05 | 3,56E+06 | 7,09E+06 | 12 | 12 | 6 | 8 | 11 | Carbamidomethylation | 241892 | PPTG_11812 | Phytophthora parasitica INRA-310 chromodomain protein (2157 aa) |
| 1171 | 896 | PPTG_01724 | 139,3 | 18 | 6 | 7 | 18 | 6,33E+05 | 1,69E+06 | 9,13E+06 | 9 | 9 | 3 | 5 | 10 | Carbamidomethylation | 77529 | PPTG_01724 | Phytophthora parasitica INRA-310 protein kinase (716 aa) |
| 920 | 810 | PPTG_09026 | 139,21 | 41 | 15 | 23 | 39 | 2,54E+06 | 7,65E+06 | 4,87E+07 | 10 | 10 | 4 | 7 | 13 | Oxidation (M); Carbamidomethylation; Deamidation (NQ) | 32753 | PPTG_09026 | Phytophthora parasitica INRA-310 mitochondrial Carrier (MC) family (305 aa) |
| 805 | 645 | PPTG_15024 | 139,18 | 27 | 6 | 23 | 22 | 1,58E+06 | 1,58E+07 | 1,69E+07 | 13 | 13 | 5 | 11 | 12 | Carbamidomethylation; Deamidation (NQ) | 66958 | PPTG_15024 | Phytophthora parasitica INRA-310 lysosomal Pro-X carboxypeptidase (603 aa) |
| 717 | 625 | PPTG_02253 | 139,13 | 13 | 7 | 10 | 12 | | 2,37E+05 | 8,38E+05 | 14 | 2 | 8 | 11 | 12 | Carbamidomethylation; Dehydration; Carbamylation | 117434 | PPTG_02253 | Phytophthora parasitica INRA-310 NAD-specific glutamate dehydrogenase (1055 aa) |
| 731 | 705 | PPTG_04743 | 139,12 | 16 | 5 | 13 | 10 | 1,06E+07 | 5,14E+07 | 5,52E+07 | 14 | 14 | 6 | 14 | 11 | Oxidation (M); Carbamidomethylation; Dehydration; Deamidation (NQ) | 125550 | PPTG_04743 | Phytophthora parasitica INRA-310 hypothetical protein (1135 aa) |
| 1123 | 759 | PPTG_04076 | 139,08 | 37 | 5 | 19 | 37 | 7,79E+05 | 2,35E+06 | 9,05E+06 | 10 | 10 | 2 | 5 | 12 | Oxidation (M); Carbamidomethylation | 48823 | PPTG_04076 | Phytophthora parasitica INRA-310 hypothetical protein (447 aa) |
| 1297 | 652 | PPTG_14756 | 139 | 22 | 3 | 1 | 20 | 2,74E+05 | 5,47E+04 | 4,95E+06 | 14 | 14 | 3 | 1 | 12 | Carbamidomethylation | 91083 | PPTG_14756 | Phytophthora parasitica INRA-310 hypothetical protein (839 aa) |
| 1041 | 617 | PPTG_17882 | 138,95 | 32 | 10 | 4 | 32 | 5,70E+05 | 1,28E+05 | 8,84E+06 | 12 | 12 | 5 | 2 | 14 | Oxidation (M); Dehydration; Deamidation (NQ) | 58432 | PPTG_17882 | Phytophthora parasitica INRA-310 hypothetical protein (528 aa) |
| 1126 | 800 | PPTG_09467 | 138,92 | 49 | 16 | 22 | 43 | 3,57E+05 | 9,38E+05 | 4,41E+06 | 10 | 9 | 4 | 5 | 10 | Carbamidomethylation; Acetylation (N-term) | 35089 | PPTG_09467 | Phytophthora parasitica INRA-310 serine/threonine-protein phosphatase PP1-alpha catalytic subunit (310 aa) |
| 847 | 732 | PPTG_08321 | 138,86 | 59 | 25 | 38 | 56 | 2,42E+06 | 2,02E+07 | 1,96E+07 | 12 | 12 | 4 | 10 | 13 | Oxidation (M); Deamidation (NQ) | 28550 | PPTG_08321 | Phytophthora parasitica INRA-310 hypothetical protein (264 aa) |
| 873 | 648 | PPTG_13691 | 138,78 | 13 | 3 | 13 | 9 | 5,24E+05 | 1,12E+07 | 8,26E+06 | 13 | 13 | 3 | 13 | 10 | Carbamidomethylation | 132972 | PPTG_13691 | Phytophthora parasitica INRA-310 endothelin-converting enzyme 1 metalloprotease family M13 (1180 aa) |
| 710 | 942 | PPTG_02703 | 138,74 | 48 | 25 | 41 | 48 | 4,16E+06 | 2,48E+07 | 4,88E+07 | 8 | 8 | 4 | 13 | 15 | Oxidation (M); Carbamidomethylation; Dehydration; Deamidation (NQ) | 26856 | PPTG_02703 | Phytophthora parasitica INRA-310 3-hydroxyacyl-CoA dehydrogenase type-2 (257 aa) |
| 1512 | 1651 | PPTG_16216 | 138,71 | 13 | 0 | 13 | 8 | | 2,12E+06 | 1,57E+06 | 5 | 5 | 0 | 8 | 5 | Oxidation (M); Carbamidomethylation; Deamidation (NQ) | 94970 | PPTG_16216 | Phytophthora parasitica INRA-310 hypothetical protein (910 aa) |
| 944 | 782 | PPTG_12683 | 138,7 | 20 | 10 | 18 | 15 | 1,09E+06 | 1,35E+07 | 1,19E+07 | 10 | 10 | 5 | 11 | 8 | Carbamidomethylation; Deamidation (NQ) | 66538 | PPTG_12683 | Phytophthora parasitica INRA-310 crinkler (CRN) family protein pseudogene (595 aa) |
| 803 | 692 | PPTG_07242 | 138,62 | 61 | 7 | 55 | 48 | 8,64E+05 | 1,90E+07 | 2,14E+07 | 12 | 12 | 2 | 12 | 14 | Oxidation (M); Deamidation (NQ) | 33990 | PPTG_07242 | Phytophthora parasitica INRA-310 electron transfer flavoprotein subunit alpha (331 aa) |
| 737 | 577 | PPTG_11618 | 138,61 | 44 | 16 | 20 | 42 | 3,19E+06 | 1,36E+06 | 2,72E+07 | 13 | 13 | 5 | 7 | 19 | Oxidation (M); Carbamidomethylation | 48008 | PPTG_11618 | Phytophthora parasitica INRA-310 hypothetical protein (442 aa) |
| 905 | 1129 | PPTG_18786 | 138,59 | 36 | 17 | 29 | 30 | 2,52E+05 | 5,47E+06 | 6,30E+06 | 7 | 7 | 3 | 12 | 10 | Oxidation (M); Carbamidomethylation | 30928 | PPTG_18786 | Phytophthora parasitica INRA-310 hypothetical protein (280 aa) |
| 1182 | 985 | PPTG_06692 | 138,58 | 47 | 14 | 21 | 47 | 1,18E+06 | 2,32E+06 | 1,37E+07 | 8 | 8 | 3 | 4 | 11 | Oxidation (M) | 23982 | PPTG_06692 | Phytophthora parasitica INRA-310 hypothetical protein (211 aa) |
| 543 | 1113 | PPTG_09412 | 138,51 | 27 | 12 | 17 | 27 | 4,88E+06 | 9,51E+06 | 5,78E+07 | 9 | 7 | 9 | 10 | 21 | Oxidation (M); Deamidation (NQ) | 55676 | PPTG_09412 | Phytophthora parasitica INRA-310 major facilitator superfamily transporter (MFS) (512 aa) |
| 656 | 851 | PPTG_02446 | 138,49 | 61 | 24 | 58 | 51 | 8,39E+06 | 5,13E+07 | 5,58E+07 | 13 | 13 | 4 | 17 | 13 | Oxidation (M); Carbamidomethylation; Dehydration; Deamidation (NQ) | 28031 | PPTG_02446 | Phytophthora parasitica INRA-310 proteasome subunit alpha (254 aa) |
| 624 | 734 | PPTG_15671 | 138,44 | 30 | 6 | 23 | 24 | 1,69E+05 | 1,06E+07 | 1,35E+07 | 11 | 11 | 2 | 16 | 17 | Oxidation (M); Deamidation (NQ); Acetylation (N-term) | 54794 | PPTG_15671 | Phytophthora parasitica INRA-310 nucleolar protein NOP5 (497 aa) |
| 974 | 1286 | PPTG_12288 | 138,41 | 22 | 22 | 22 | 22 | 9,64E+06 | 6,77E+07 | 7,49E+07 | 6 | 6 | 7 | 8 | 8 | Carbamidomethylation | 44456 | PPTG_12288 | Phytophthora parasitica INRA-310 hypothetical protein (417 aa) |
| 226 | 996 | PPTG_05319 | 138,39 | 47 | 26 | 39 | 47 | 1,81E+08 | 6,33E+08 | 1,64E+09 | 11 | 11 | 17 | 26 | 35 | Oxidation (M); Dehydration; Deamidation (NQ); Carbamylation | 24489 | PPTG_05319 | Phytophthora parasitica INRA-310 60S ribosomal protein (207 aa) |
| 965 | 597 | PPTG_05362 | 138,27 | 27 | 7 | 23 | 24 | 8,26E+05 | 1,00E+07 | 1,79E+07 | 12 | 12 | 3 | 10 | 10 | Carbamidomethylation | 64118 | PPTG_05362 | Phytophthora parasitica INRA-310 CTP synthase (581 aa) |
| 935 | 998 | PPTG_17182 | 138,25 | 31 | 9 | 29 | 25 | 7,06E+05 | 9,56E+06 | 1,01E+07 | 8 | 8 | 2 | 10 | 12 | Oxidation (M); Carbamidomethylation; Acetylation (N-term) | 38359 | PPTG_17182 | Phytophthora parasitica INRA-310 sorbitol dehydrogenase (360 aa) |
| 815 | 654 | PPTG_05388 | 138,06 | 41 | 22 | 32 | 39 | 1,52E+06 | 1,12E+07 | 1,67E+07 | 11 | 11 | 6 | 10 | 12 | Oxidation (M); Carbamidomethylation | 41597 | PPTG_05388 | Phytophthora parasitica INRA-310 pyruvate dehydrogenase (381 aa) |
| 1141 | 1920 | PPTG_07843 | 138,05 | 20 | 6 | 20 | 20 | 7,63E+05 | 9,06E+06 | 8,54E+06 | 4 | 3 | 2 | 9 | 8 | Oxidation (M); Carbamidomethylation; Deamidation (NQ) | 37347 | PPTG_07843 | Phytophthora parasitica INRA-310 hypothetical protein (360 aa) |
| 989 | 1103 | PPTG_15023 | 138,05 | 18 | 11 | 13 | 18 | 1,89E+06 | 5,17E+06 | 1,79E+07 | 7 | 7 | 6 | 6 | 10 | Oxidation (M); Carbamidomethylation; Dehydration; Deamidation (NQ) | 54511 | PPTG_15023 | Phytophthora parasitica INRA-310 UTP-glucose-1-phosphate uridylyltransferase (482 aa) |
| 950 | 1168 | PPTG_07406 | 138 | 31 | 12 | 28 | 31 | 1,99E+06 | 2,49E+07 | 2,89E+07 | 8 | 8 | 4 | 10 | 10 | Oxidation (M); Deamidation (NQ); Acetylation (N-term) | 31805 | PPTG_07406 | Phytophthora parasitica INRA-310 proteasomal ubiquitin receptor ADRM1-like protein (306 aa) |
| 768 | 760 | PPTG_00887 | 137,98 | 17 | 5 | 14 | 10 | 8,31E+05 | 6,25E+06 | 5,69E+06 | 15 | 15 | 6 | 14 | 9 | Carbamidomethylation; Acetylation (N-term) | 126390 | PPTG_00887 | Phytophthora parasitica INRA-310 DNA damage-binding protein (1149 aa) |
| 1003 | 966 | PPTG_14718 | 137,92 | 21 | 7 | 15 | 17 | 7,25E+05 | 5,52E+06 | 8,20E+06 | 10 | 10 | 3 | 10 | 9 | Oxidation (M); Carbamidomethylation; Deamidation (NQ) | 58020 | PPTG_14718 | Phytophthora parasitica INRA-310 amidophosphoribosyltransferase (520 aa) |
| 1124 | 867 | PPTG_10266 | 137,69 | 36 | 19 | 22 | 36 | 5,57E+05 | 9,57E+05 | 7,53E+06 | 9 | 9 | 4 | 5 | 10 | Carbamidomethylation | 37849 | PPTG_10266 | Phytophthora parasitica INRA-310 hypothetical protein (334 aa) |
| 1131 | 1059 | PPTG_15563 | 137,69 | 33 | 9 | 33 | 22 | 1,60E+05 | 6,04E+06 | 3,65E+06 | 7 | 7 | 3 | 9 | 7 | Carbamidomethylation | 40745 | PPTG_15563 | Phytophthora parasitica INRA-310 glycoside hydrolase (385 aa) |
| 1053 | 852 | PPTG_16453 | 137,67 | 35 | 12 | 15 | 35 | 4,29E+05 | 2,17E+06 | 1,06E+07 | 11 | 11 | 3 | 6 | 12 | Carbamidomethylation | 43899 | PPTG_16453 | Phytophthora parasitica INRA-310 hypothetical protein (396 aa) |
| 1146 | 2336 | PPTG_18886 | 137,57 | 21 | 5 | 21 | 21 | 5,49E+04 | 8,30E+06 | 1,19E+07 | 6 | 6 | 1 | 8 | 10 | Oxidation (M) | 31605 | PPTG_18886 | Phytophthora parasitica INRA-310 fructokinase (295 aa) |
| 1071 | 1263 | PPTG_15025 | 137,54 | 22 | 15 | 10 | 22 | 1,46E+06 | 1,50E+06 | 2,80E+07 | 7 | 7 | 4 | 4 | 12 | Deamidation (NQ) | 42488 | PPTG_15025 | Phytophthora parasitica INRA-310 hypothetical protein (394 aa) |
| 1088 | 857 | PPTG_10392 | 137,3 | 17 | 1 | 14 | 11 | 0 | 6,17E+06 | 2,80E+06 | 13 | 13 | 1 | 12 | 7 | | 99538 | PPTG_10392 | Phytophthora parasitica INRA-310 nuclear cap-binding protein subunit (894 aa) |
| 804 | 1110 | PPTG_14382 | 137,22 | 26 | 18 | 21 | 26 | 5,31E+06 | 1,50E+07 | 6,04E+07 | 7 | 7 | 7 | 8 | 13 | Oxidation (M); Dehydration; Deamidation (NQ) | 41408 | PPTG_14382 | Phytophthora parasitica INRA-310 hypothetical protein (384 aa) |
| 1002 | 755 | PPTG_08786 | 137 | 20 | 7 | 3 | 17 | 1,00E+06 | 8,27E+05 | 1,32E+07 | 16 | 16 | 5 | 3 | 14 | Oxidation (M) | 109119 | PPTG_08786 | Phytophthora parasitica INRA-310 hypothetical protein (995 aa) |
| 1321 | 1260 | PPTG_09275 | 136,84 | 53 | 18 | 43 | 40 | 5,94E+05 | 4,31E+06 | 1,28E+07 | 8 | 8 | 3 | 6 | 7 | Oxidation (M); Deamidation (NQ) | 20816 | PPTG_09275 | Phytophthora parasitica INRA-310 NAD(P)H:quinone oxidoreductase type IV (201 aa) |
| 937 | 676 | PPTG_17513 | 136,71 | 35 | 12 | 24 | 35 | 1,09E+06 | 1,11E+07 | 2,99E+07 | 11 | 11 | 4 | 8 | 12 | Carbamidomethylation | 46928 | PPTG_17513 | Phytophthora parasitica INRA-310 DEAD-box ATP-dependent RNA helicase 8 (412 aa) |
| 1044 | 703 | PPTG_02681 | 136,7 | 19 | 4 | 19 | 10 | 1,45E+05 | 4,50E+06 | 2,41E+06 | 11 | 11 | 2 | 12 | 7 | Oxidation (M); Carbamidomethylation; Deamidation (NQ) | 93753 | PPTG_02681 | Phytophthora parasitica INRA-310 alpha-amylase (866 aa) |
| 1006 | 870 | PPTG_06782 | 136,69 | 32 | 2 | 27 | 20 | 2,22E+05 | 4,89E+06 | 5,41E+06 | 11 | 11 | 1 | 12 | 9 | Deamidation (NQ) | 56987 | PPTG_06782 | Phytophthora parasitica INRA-310 hypothetical protein (558 aa) |
| 1011 | 831 | PPTG_04127 | 136,66 | 21 | 4 | 2 | 19 | 3,65E+05 | 1,40E+06 | 9,48E+06 | 17 | 16 | 4 | 2 | 16 | Oxidation (M); Carbamidomethylation; Dehydration; Carbamylation | 105817 | PPTG_04127 | Phytophthora parasitica INRA-310 protein kinase (953 aa) |
| 846 | 677 | PPTG_09787 | 136,62 | 36 | 6 | 33 | 23 | 2,85E+05 | 9,12E+06 | 1,17E+07 | 12 | 12 | 2 | 13 | 12 | Oxidation (M); Carbamidomethylation | 53619 | PPTG_09787 | Phytophthora parasitica INRA-310 glutathione reductase (494 aa) |
| 852 | 878 | PPTG_02066 | 136,51 | 34 | 8 | 22 | 34 | 9,45E+05 | 3,93E+06 | 2,67E+07 | 9 | 8 | 4 | 7 | 16 | Oxidation (M); Carbamidomethylation; Deamidation (NQ) | 33640 | PPTG_02066 | Phytophthora parasitica INRA-310 hypothetical protein (306 aa) |
| 748 | 655 | PPTG_09121 | 136,48 | 28 | 9 | 25 | 27 | 8,55E+05 | 9,23E+06 | 1,67E+07 | 13 | 13 | 4 | 12 | 14 | Carbamidomethylation; Deamidation (NQ) | 56530 | PPTG_09121 | Phytophthora parasitica INRA-310 pyridine nucleotide-disulphide oxidoreductase (521 aa) |
| 631 | 670 | PPTG_08095 | 136,47 | 45 | 9 | 42 | 42 | 1,57E+06 | 2,44E+07 | 3,22E+07 | 12 | 12 | 3 | 14 | 18 | Oxidation (M); Carbamidomethylation | 36297 | PPTG_08095 | Phytophthora parasitica INRA-310 carbonic anhydrase (318 aa) |
| 813 | 1053 | PPTG_13316 | 136,46 | 64 | 25 | 43 | 64 | 3,47E+06 | 9,90E+06 | 7,22E+07 | 7 | 7 | 4 | 8 | 16 | Oxidation (M); Deamidation (NQ) | 14412 | PPTG_13316 | Phytophthora parasitica INRA-310 hypothetical protein (129 aa) |
| 1051 | 743 | PPTG_11616 | 136,22 | 28 | 8 | 8 | 26 | 5,88E+05 | 5,37E+05 | 1,45E+07 | 10 | 10 | 3 | 4 | 14 | Oxidation (M); Carbamidomethylation | 57110 | PPTG_11616 | Phytophthora parasitica INRA-310 tubby-like protein (518 aa) |
| 1539 | 2337 | PPTG_07985 | 136,16 | 31 | 14 | 31 | 31 | 1,19E+06 | 1,81E+07 | 1,54E+07 | 3 | 2 | 1 | 7 | 5 | Oxidation (M); Carbamidomethylation | 21355 | PPTG_07985 | Phytophthora parasitica INRA-310 hypothetical protein (205 aa) |
| 1385 | 871 | PPTG_00778 | 136,06 | 34 | 4 | 9 | 34 | 2,90E+05 | 1,13E+06 | 1,50E+07 | 9 | 9 | 1 | 3 | 11 | Carbamidomethylation | 37572 | PPTG_00778 | Phytophthora parasitica INRA-310 alcohol dehydrogenase (345 aa) |
| 475 | 1210 | PPTG_00066 | 136,04 | 25 | 20 | 23 | 22 | 6,68E+06 | 1,02E+08 | 1,32E+08 | 7 | 7 | 9 | 19 | 16 | Oxidation (M); Carbamidomethylation; Deamidation (NQ) | 34535 | PPTG_00066 | Phytophthora parasitica INRA-310 hypothetical protein (330 aa) |
| 1242 | 952 | PPTG_05956 | 136,01 | 44 | 5 | 34 | 39 | 1,70E+05 | 8,94E+06 | 1,13E+07 | 9 | 9 | 1 | 8 | 8 | Oxidation (M); Carbamidomethylation | 28181 | PPTG_05956 | Phytophthora parasitica INRA-310 proliferating cell nuclear antigen (pcna) (260 aa) |
| 1119 | 697 | PPTG_03693 | 136 | 11 | 3 | 4 | 11 | 4,71E+05 | 1,38E+06 | 8,76E+06 | 12 | 12 | 3 | 4 | 12 | Carbamidomethylation | 123327 | PPTG_03693 | Phytophthora parasitica INRA-310 cGMP-dependent protein kinase (1106 aa) |
| 1558 | 1511 | PPTG_05343 | 135,88 | 31 | 14 | 25 | 22 | 1,09E+06 | 5,04E+06 | 3,07E+06 | 6 | 6 | 3 | 5 | 4 | Oxidation (M) | 27520 | PPTG_05343 | Phytophthora parasitica INRA-310 hypothetical protein (252 aa) |
| 1326 | 1310 | PPTG_13440 | 135,73 | 22 | 6 | 22 | 19 | | 3,21E+06 | 4,32E+05 | 6 | 2 | 2 | 9 | 5 | Oxidation (M); Carbamidomethylation; Amidation | 55122 | PPTG_13440 | Phytophthora parasitica INRA-310 alanine aminotransferase 2 (500 aa) |
| 1160 | 633 | PPTG_09762 | 135,68 | 15 | 3 | 0 | 14 | 1,59E+05 | | 6,48E+06 | 13 | 13 | 4 | 0 | 14 | Carbamidomethylation | 135825 | PPTG_09762 | Phytophthora parasitica INRA-310 hypothetical protein (1211 aa) |
| 818 | 884 | PPTG_13259 | 135,57 | 4 | 0 | 4 | 2 | | 3,38E+06 | 4,64E+06 | 18 | 18 | 0 | 16 | 11 | Oxidation (M); Carbamidomethylation | 551262 | PPTG_13259 | Phytophthora parasitica INRA-310 HECT E3 ubiquitin ligase (5010 aa) |
| 1209 | 837 | PPTG_02677 | 135,53 | 17 | 4 | 1 | 15 | 2,57E+05 | 3,38E+05 | 7,69E+06 | 15 | 15 | 3 | 1 | 13 | Carbamidomethylation | 114033 | PPTG_02677 | Phytophthora parasitica INRA-310 calcium-transporting ATPase (1041 aa) |
| 1019 | 783 | PPTG_04373 | 135,46 | 44 | 18 | 38 | 44 | 2,61E+05 | 3,53E+06 | 6,59E+06 | 10 | 10 | 4 | 8 | 10 | Carbamidomethylation; Deamidation (NQ) | 33461 | PPTG_04373 | Phytophthora parasitica INRA-310 hypothetical protein (309 aa) |
| 915 | 935 | PPTG_13565 | 135,42 | 29 | 14 | 29 | 29 | 6,86E+06 | 3,58E+07 | 3,60E+07 | 9 | 8 | 4 | 10 | 10 | Carbamidomethylation | 39471 | PPTG_13565 | Phytophthora parasitica INRA-310 voltage-gated potassium channel subunit beta (350 aa) |
| 861 | 1050 | PPTG_18069 | 135,41 | 46 | 25 | 46 | 42 | 1,72E+06 | 1,35E+07 | 2,20E+07 | 8 | 8 | 5 | 11 | 11 | Oxidation (M); Carbamidomethylation; Dehydration; Deamidation (NQ) | 22345 | PPTG_18069 | Phytophthora parasitica INRA-310 hypothetical protein (195 aa) |
| 609 | 1356 | PPTG_05458 | 135,36 | 42 | 40 | 28 | 42 | 5,94E+07 | 1,76E+08 | 4,74E+08 | 8 | 8 | 9 | 11 | 16 | Oxidation (M); Carbamylation | 16739 | PPTG_05458 | Phytophthora parasitica INRA-310 60S ribosomal protein L23a (151 aa) |
| 617 | 888 | PPTG_18551 | 135,28 | 21 | 5 | 21 | 18 | 2,43E+06 | 1,86E+07 | 2,64E+07 | 16 | 13 | 4 | 16 | 16 | Oxidation (M); Dehydration; Deamidation (NQ) | 90632 | PPTG_18551 | Phytophthora parasitica INRA-310 N-terminal acetyltransferase A complex subunit nat1-like protein (786 aa) |
| 835 | 1360 | PPTG_03134 | 135,27 | 16 | 11 | 15 | 16 | 4,81E+06 | 2,54E+07 | 4,13E+07 | 8 | 8 | 5 | 10 | 12 | Carbamidomethylation; Deamidation (NQ) | 50123 | PPTG_03134 | Phytophthora parasitica INRA-310 dihydrolipoyllysine-residue acetyltransferase component-pyruvate dehydrogenase complex (481 aa) |
| 869 | 798 | PPTG_15401 | 134,99 | 30 | 10 | 29 | 23 | 1,20E+06 | 8,94E+06 | 1,29E+07 | 11 | 11 | 5 | 11 | 10 | Carbamidomethylation | 51698 | PPTG_15401 | Phytophthora parasitica INRA-310 cysteine desulfurase mitochondrial (473 aa) |
| 1273 | 1106 | PPTG_07468 | 134,93 | 18 | 10 | 10 | 18 | 6,30E+05 | 1,54E+06 | 5,22E+06 | 8 | 8 | 4 | 4 | 8 | Oxidation (M); Carbamidomethylation; Carbamylation | 59248 | PPTG_07468 | Phytophthora parasitica INRA-310 protein kinase (526 aa) |
| 1279 | 671 | PPTG_16075 | 134,88 | 12 | 1 | 1 | 12 | | 6,21E+03 | 5,98E+06 | 12 | 10 | 1 | 1 | 14 | Oxidation (M); Carbamidomethylation | 133943 | PPTG_16075 | Phytophthora parasitica INRA-310 phospholipid-transporting ATPase (1202 aa) |
| 984 | 737 | PPTG_14405 | 134,87 | 14 | 1 | 8 | 13 | 5,77E+04 | 1,46E+06 | 3,29E+06 | 14 | 14 | 1 | 8 | 13 | Carbamidomethylation | 146095 | PPTG_14405 | Phytophthora parasitica INRA-310 hypothetical protein (1326 aa) |
| 1116 | 973 | PPTG_06596 | 134,77 | 19 | 2 | 17 | 11 | 1,37E+05 | 7,31E+06 | 2,87E+06 | 11 | 11 | 1 | 11 | 7 | | 86392 | PPTG_06596 | Phytophthora parasitica INRA-310 acylamino-acid-releasing enzyme (786 aa) |
| 795 | 788 | PPTG_03858 | 134,76 | 25 | 5 | 15 | 22 | 7,52E+05 | 3,53E+06 | 9,34E+06 | 17 | 17 | 3 | 11 | 14 | Carbamidomethylation | 80123 | PPTG_03858 | Phytophthora parasitica INRA-310 thimet oligopeptidase (707 aa) |
| 1181 | 882 | PPTG_13152 | 134,76 | 31 | 11 | 13 | 31 | 3,28E+05 | 9,25E+05 | 7,47E+06 | 9 | 9 | 4 | 4 | 10 | Carbamidomethylation | 44572 | PPTG_13152 | Phytophthora parasitica INRA-310 hypothetical protein (401 aa) |
| 975 | 1390 | PPTG_01646 | 134,68 | 66 | 28 | 58 | 66 | 1,31E+06 | 1,79E+07 | 2,13E+07 | 6 | 6 | 4 | 9 | 10 | Oxidation (M); Carbamidomethylation | 12303 | PPTG_01646 | Phytophthora parasitica INRA-310 peptidyl-prolyl cis-trans isomerase Pin1 (114 aa) |
| 897 | 762 | PPTG_05104 | 134,68 | 14 | 5 | 6 | 13 | 2,11E+06 | 4,92E+06 | 1,98E+07 | 12 | 12 | 5 | 7 | 13 | Oxidation (M); Carbamidomethylation | 129710 | PPTG_05104 | Phytophthora parasitica INRA-310 P-type ATPase (P-ATPase) superfamily (1202 aa) |
| 1258 | 1355 | PPTG_00758 | 134,68 | 31 | 8 | 31 | 23 | 8,50E+05 | 4,72E+07 | 3,27E+07 | 7 | 7 | 2 | 9 | 6 | Carbamidomethylation | 37812 | PPTG_00758 | Phytophthora parasitica INRA-310 hypothetical protein (341 aa) |
| 812 | 900 | PPTG_05278 | 134,66 | 18 | 7 | 15 | 18 | 7,43E+06 | 1,19E+07 | 1,01E+08 | 10 | 10 | 5 | 9 | 14 | Oxidation (M); Deamidation (NQ); Acetylation (N-term) | 61695 | PPTG_05278 | Phytophthora parasitica INRA-310 Ca2:Cation Antiporter (CaCA) family (564 aa) |
| 1000 | 679 | PPTG_00969 | 134,55 | 19 | 6 | 17 | 13 | 2,00E+06 | 1,13E+07 | 1,57E+07 | 11 | 11 | 4 | 10 | 8 | Oxidation (M); Carbamidomethylation | 82528 | PPTG_00969 | Phytophthora parasitica INRA-310 methylmalonyl-CoA mutase mitochondrial (754 aa) |
| 968 | 709 | PPTG_02501 | 134,52 | 27 | 8 | 18 | 20 | 1,68E+06 | 1,65E+07 | 2,57E+07 | 12 | 12 | 5 | 8 | 10 | Oxidation (M); Carbamidomethylation; Deamidation (NQ) | 51054 | PPTG_02501 | Phytophthora parasitica INRA-310 hypothetical protein (460 aa) |
| 927 | 753 | PPTG_13814 | 134,49 | 17 | 13 | 12 | 17 | 1,73E+06 | 5,38E+06 | 1,71E+07 | 9 | 9 | 7 | 7 | 10 | Oxidation (M); Carbamidomethylation; Deamidation (NQ) | 73252 | PPTG_13814 | Phytophthora parasitica INRA-310 long-chain-fatty-acid-CoA ligase (667 aa) |
| 734 | 868 | PPTG_09261 | 134,45 | 38 | 15 | 35 | 38 | 2,20E+06 | 1,44E+07 | 2,14E+07 | 10 | 10 | 5 | 13 | 13 | Carbamidomethylation; Dehydration; Deamidation (NQ) | 30739 | PPTG_09261 | Phytophthora parasitica INRA-310 hypothetical protein (279 aa) |
| 1118 | 1137 | PPTG_16135 | 134,36 | 38 | 9 | 34 | 33 | 4,95E+04 | 2,48E+06 | 2,55E+06 | 8 | 8 | 2 | 9 | 8 | Carbamidomethylation; Acetylation (N-term) | 29797 | PPTG_16135 | Phytophthora parasitica INRA-310 hypothetical protein (264 aa) |
| 1256 | 1216 | PPTG_15564 | 134,33 | 27 | 0 | 24 | 19 | | 7,70E+06 | 8,90E+06 | 8 | 8 | 0 | 9 | 8 | Oxidation (M) | 40091 | PPTG_15564 | Phytophthora parasitica INRA-310 hypothetical protein (367 aa) |
| 928 | 856 | PPTG_09146 | 134,2 | 36 | 8 | 18 | 32 | 2,90E+05 | 2,45E+06 | 7,14E+06 | 14 | 14 | 3 | 7 | 14 | Carbamidomethylation; Deamidation (NQ); Acetylation (N-term) | 47239 | PPTG_09146 | Phytophthora parasitica INRA-310 dual specificity protein phosphatase (423 aa) |
| 1207 | 812 | PPTG_04333 | 134,11 | 9 | 1 | 4 | 8 | 1,22E+05 | 1,31E+06 | 4,51E+06 | 11 | 11 | 2 | 5 | 10 | | 167497 | PPTG_04333 | Phytophthora parasitica INRA-310 nuclear pore complex protein Nup155 (1519 aa) |
| 913 | 714 | PPTG_14242 | 134,02 | 36 | 3 | 34 | 20 | 3,45E+05 | 9,49E+06 | 5,63E+06 | 11 | 11 | 2 | 13 | 9 | Carbamidomethylation | 53823 | PPTG_14242 | Phytophthora parasitica INRA-310 hypothetical protein (489 aa) |
| 1151 | 757 | PPTG_13847 | 133,98 | 15 | 9 | 4 | 15 | 3,60E+05 | 2,93E+05 | 3,43E+07 | 10 | 10 | 5 | 2 | 11 | Carbamidomethylation | 90501 | PPTG_13847 | Phytophthora parasitica INRA-310 hypothetical protein (813 aa) |
| 616 | 763 | PPTG_09264 | 133,97 | 38 | 4 | 27 | 35 | 1,19E+06 | 1,03E+07 | 1,65E+07 | 18 | 18 | 3 | 15 | 18 | Oxidation (M); Carbamidomethylation; Deamidation (NQ) | 71949 | PPTG_09264 | Phytophthora parasitica INRA-310 choline/Carnitine O-acyltransferase (635 aa) |
| 814 | 1368 | PPTG_13910 | 133,96 | 36 | 25 | 29 | 36 | 4,39E+06 | 1,05E+07 | 6,23E+07 | 8 | 8 | 7 | 9 | 12 | Oxidation (M); Carbamidomethylation; Deamidation (NQ) | 23690 | PPTG_13910 | Phytophthora parasitica INRA-310 NADH dehydrogenase iron-sulfur protein 8 (212 aa) |
| 1004 | 1264 | PPTG_06372 | 133,66 | 16 | 11 | 16 | 16 | 1,92E+06 | 1,96E+07 | 3,12E+07 | 7 | 7 | 4 | 9 | 9 | Carbamidomethylation | 58540 | PPTG_06372 | Phytophthora parasitica INRA-310 calcyphosin-like protein (538 aa) |
| 1407 | 706 | PPTG_06696 | 133,47 | 20 | 2 | 2 | 20 | 1,60E+05 | 0 | 5,48E+06 | 12 | 11 | 1 | 1 | 12 | Oxidation (M) | 80881 | PPTG_06696 | Phytophthora parasitica INRA-310 transmembrane protein (714 aa) |
| 1183 | 1058 | PPTG_16595 | 133,37 | 37 | 11 | 37 | 30 | 2,23E+05 | 1,08E+07 | 7,10E+06 | 8 | 8 | 2 | 9 | 7 | Carbamidomethylation | 32954 | PPTG_16595 | Phytophthora parasitica INRA-310 fumarylacetoacetate (FAA) hydrolase (306 aa) |
| 1227 | 695 | PPTG_11756 | 133,33 | 35 | 7 | 9 | 30 | 2,14E+06 | 3,37E+06 | 2,12E+07 | 12 | 12 | 3 | 4 | 10 | Carbamidomethylation | 58607 | PPTG_11756 | Phytophthora parasitica INRA-310 serine palmitoyltransferase (534 aa) |

| | | | | | | | | | | | | | | | | | | | |
|---|---|---|---|---|---|---|---|---|---|---|---|---|---|---|---|---|---|---|---|
| 832 | 793 | PPTG_12099 | 133,15 | 26 | 5 | 18 | 24 | 1,28E+06 | 5,53E+06 | 1,28E+07 | 11 | 11 | 3 | 11 | 13 | Oxidation (M); Carbamidomethylation; Deamidation (NQ); Amidation | 57755 | PPTG_12099 | Phytophthora parasitica INRA-310 CBL-interacting serine/threonine-protein kinase (520 aa) |
| 1161 | 656 | PPTG_18781 | 133,08 | 31 | 6 | 4 | 30 | 3,43E+05 | 2,92E+05 | 1,20E+07 | 13 | 12 | 3 | 2 | 13 | Oxidation (M); Carbamidomethylation; Deamidation (NQ) | 67620 | PPTG_18781 | Phytophthora parasitica INRA-310 ATP-binding Cassette (ABC) superfamily (614 aa) |
| 988 | 1148 | PPTG_02730 | 132,84 | 24 | 9 | 20 | 24 | 1,75E+06 | 2,01E+07 | 1,97E+07 | 8 | 8 | 3 | 9 | 10 | Oxidation (M); Carbamidomethylation | 48351 | PPTG_02730 | Phytophthora parasitica INRA-310 hypothetical protein (446 aa) |
| 1136 | 754 | PPTG_16799 | 132,84 | 31 | 2 | 10 | 28 | 6,01E+05 | 3,11E+06 | 1,98E+07 | 13 | 13 | 1 | 4 | 14 | Carbamidomethylation; Amidation | 56644 | PPTG_16799 | Phytophthora parasitica INRA-310 thioredoxin-like protein (497 aa) |
| 1423 | 1809 | PPTG_12536 | 132,75 | 43 | 21 | 43 | 43 | 9,99E+05 | 1,04E+07 | 1,01E+07 | 4 | 4 | 2 | 7 | 5 | Oxidation (M); Carbamidomethylation; Dehydration; Deamidation (NQ) | 16219 | PPTG_12536 | Phytophthora parasitica INRA-310 PpiC-type peptidyl-prolyl cis-trans isomerase (149 aa) |
| 753 | 708 | PPTG_01690 | 132,67 | 47 | 23 | 35 | 37 | 1,31E+06 | 7,75E+06 | 1,35E+07 | 14 | 14 | 7 | 12 | 11 | Carbamidomethylation; Acetylation (N-term) | 38927 | PPTG_01690 | Phytophthora parasitica INRA-310 phosphoribosylaminoimidazole-succinocarboxamide synthase (342 aa) |
| 1168 | 1200 | PPTG_06370 | 132,65 | 37 | 15 | 34 | 37 | 1,51E+06 | 6,67E+06 | 1,52E+07 | 7 | 7 | 3 | 7 | 8 | Carbamidomethylation | 23258 | PPTG_06370 | Phytophthora parasitica INRA-310 hypothetical protein (221 aa) |
| 1228 | 1028 | PPTG_19149 | 132,46 | 41 | 18 | 24 | 41 | 4,35E+05 | 2,57E+06 | 1,73E+07 | 8 | 8 | 3 | 5 | 9 | Carbamidomethylation | 27209 | PPTG_19149 | Phytophthora parasitica INRA-310 dolichol-phosphate mannosyltransferase (241 aa) |
| 709 | 786 | PPTG_00875 | 132,46 | 21 | 2 | 19 | 17 | 2,65E+05 | 3,76E+06 | 7,95E+06 | 17 | 17 | 2 | 15 | 15 | Carbamidomethylation; Deamidation (NQ); Carbamylation | 113688 | PPTG_00875 | Phytophthora parasitica INRA-310 hypothetical protein (1035 aa) |
| 782 | 686 | PPTG_19189 | 132,24 | 15 | 2 | 14 | 13 | 5,53E+05 | 1,10E+07 | 1,61E+07 | 11 | 11 | 2 | 14 | 13 | Oxidation (M); Carbamidomethylation; Acetylation (N-term) | 115718 | PPTG_19189 | Phytophthora parasitica INRA-310 hypothetical protein (1031 aa) |
| 1236 | 806 | PPTG_01496 | 132,22 | 22 | 4 | 11 | 22 | 7,82E+05 | 1,05E+06 | 2,19E+07 | 10 | 10 | 2 | 5 | 10 | | 63464 | PPTG_01496 | Phytophthora parasitica INRA-310 hypothetical protein (562 aa) |
| 843 | 1167 | PPTG_00442 | 132,12 | 45 | 17 | 42 | 34 | 2,97E+06 | 8,18E+07 | 6,28E+07 | 7 | 7 | 4 | 13 | 10 | Carbamidomethylation; Deamidation (NQ) | 24335 | PPTG_00442 | Phytophthora parasitica INRA-310 hypothetical protein (225 aa) |
| 1049 | 855 | PPTG_17984 | 132,11 | 42 | 9 | 35 | 34 | 1,44E+06 | 1,90E+07 | 2,38E+07 | 10 | 10 | 3 | 9 | 9 | Oxidation (M); Carbamidomethylation; Deamidation (NQ); Carbamylation | 39248 | PPTG_17984 | Phytophthora parasitica INRA-310 pyruvate dehydrogenase E1 component subunit beta mitochondrial (360 aa) |
| 1138 | 1377 | PPTG_13362 | 132,03 | 36 | 7 | 36 | 25 | 2,96E+05 | 3,45E+06 | 2,49E+06 | 7 | 7 | 2 | 9 | 8 | Oxidation (M) | 35711 | PPTG_13362 | Phytophthora parasitica INRA-310 RNA polymerase Rpb3/Rpb11 dimerisation domain-containing protein (320 aa) |
| 881 | 839 | PPTG_08473 | 132,03 | 25 | 11 | 13 | 25 | 1,06E+06 | 2,62E+06 | 1,22E+07 | 11 | 10 | 6 | 7 | 13 | Carbamidomethylation | 53208 | PPTG_08473 | Phytophthora parasitica INRA-310 protein kinase (474 aa) |
| 853 | 1367 | PPTG_06915 | 131,84 | 58 | 25 | 58 | 52 | 3,70E+06 | 3,49E+07 | 3,54E+07 | 8 | 8 | 5 | 13 | 9 | Carbamidomethylation | 15886 | PPTG_06915 | Phytophthora parasitica INRA-310 copper/zinc superoxide dismutase (156 aa) |
| 1072 | 1032 | PPTG_07301 | 131,84 | 34 | 23 | 30 | 30 | 1,30E+06 | 6,97E+06 | 9,93E+06 | 7 | 7 | 5 | 7 | 8 | | 31557 | PPTG_07301 | Phytophthora parasitica INRA-310 peroxisomal 2 4-dienoyl-CoA reductase (298 aa) |
| 1274 | 707 | PPTG_06943 | 131,82 | 9 | 0 | 1 | 8 | | 4,22E+05 | 6,26E+06 | 13 | 13 | 0 | 2 | 14 | Carbamidomethylation | 211484 | PPTG_06943 | Phytophthora parasitica INRA-310 hypothetical protein (1893 aa) |
| 902 | 879 | PPTG_07225 | 131,78 | 28 | 10 | 20 | 25 | 1,75E+06 | 1,63E+07 | 2,17E+07 | 11 | 11 | 4 | 9 | 12 | Oxidation (M); Acetylation (N-term); Carbamylation | 55254 | PPTG_07225 | Phytophthora parasitica INRA-310 asparaginyl-tRNA synthetase (496 aa) |
| 884 | 1157 | PPTG_19545 | 131,77 | 55 | 34 | 46 | 55 | 7,08E+06 | 2,10E+07 | 6,89E+07 | 7 | 7 | 6 | 8 | 12 | Oxidation (M); Deamidation (NQ) | 19942 | PPTG_19545 | Phytophthora parasitica INRA-310 hypothetical protein (185 aa) |
| 849 | 853 | PPTG_10533 | 131,6 | 60 | 15 | 45 | 40 | 1,48E+06 | 2,01E+07 | 1,97E+07 | 14 | 14 | 4 | 12 | 11 | Oxidation (M); Carbamidomethylation | 36036 | PPTG_10533 | Phytophthora parasitica INRA-310 eukaryotic translation initiation factor 3 subunit (328 aa) |
| 1156 | 820 | PPTG_08766 | 131,59 | 18 | 8 | 5 | 18 | 2,92E+05 | 1,10E+06 | 7,97E+06 | 10 | 9 | 4 | 3 | 11 | Carbamidomethylation | 74579 | PPTG_08766 | Phytophthora parasitica INRA-310 mitogen-activated protein kinase (664 aa) |
| 1253 | 1044 | PPTG_05903 | 131,55 | 48 | 14 | 31 | 44 | 1,73E+06 | 8,03E+06 | 1,70E+07 | 8 | 8 | 3 | 5 | 9 | Oxidation (M); Carbamidomethylation; Acetylation (N-term) | 26125 | PPTG_05903 | Phytophthora parasitica INRA-310 catechol O-methyltransferase (242 aa) |
| 1729 | 1158 | PPTG_02084 | 131,54 | 38 | 12 | 3 | 38 | 3,39E+05 | 8,24E+05 | 1,20E+07 | 7 | 7 | 2 | 1 | 8 | Carbamidomethylation | 30547 | PPTG_02084 | Phytophthora parasitica INRA-310 hypothetical protein (281 aa) |
| 958 | 785 | PPTG_14972 | 131,38 | 21 | 6 | 18 | 21 | 1,35E+05 | 2,55E+06 | 5,72E+06 | 11 | 11 | 3 | 9 | 11 | Carbamidomethylation | 65400 | PPTG_14972 | Phytophthora parasitica INRA-310 hypothetical protein (593 aa) |
| 959 | 872 | PPTG_13317 | 131,31 | 16 | 9 | 13 | 16 | 8,15E+05 | 3,80E+06 | 1,44E+07 | 9 | 9 | 5 | 8 | 10 | Carbamidomethylation | 72006 | PPTG_13317 | Phytophthora parasitica INRA-310 hypothetical protein (665 aa) |
| 783 | 688 | PPTG_01338 | 131,3 | 72 | 42 | 68 | 67 | 4,56E+06 | 2,07E+07 | 2,51E+07 | 10 | 10 | 7 | 11 | 11 | Oxidation (M); Carbamidomethylation; Acetylation (N-term) | 19725 | PPTG_01338 | Phytophthora parasitica INRA-310 hypothetical protein (177 aa) |
| 740 | 1345 | PPTG_10200 | 131,27 | 48 | 38 | 48 | 48 | 2,60E+06 | 4,40E+07 | 4,11E+07 | 6 | 6 | 6 | 12 | 13 | Oxidation (M); Carbamidomethylation; Deamidation (NQ) | 23021 | PPTG_10200 | Phytophthora parasitica INRA-310 isochorismatase (210 aa) |
| 1045 | 892 | PPTG_14788 | 131,19 | 33 | 15 | 31 | 27 | 2,84E+06 | 1,72E+07 | 3,76E+07 | 9 | 9 | 3 | 9 | 9 | Carbamidomethylation | 43628 | PPTG_14788 | Phytophthora parasitica INRA-310 GTP-binding protein YchF (391 aa) |
| 1125 | 876 | PPTG_18280 | 131,16 | 31 | 12 | 8 | 31 | 3,73E+05 | 8,91E+05 | 6,46E+06 | 9 | 7 | 3 | 3 | 13 | Oxidation (M); Dehydration; Deamidation (NQ) | 42101 | PPTG_18280 | Phytophthora parasitica INRA-310 hypothetical protein (385 aa) |
| 1134 | 916 | PPTG_12069 | 131,14 | 32 | 10 | 15 | 32 | 7,41E+05 | 2,67E+06 | 8,26E+06 | 10 | 10 | 3 | 5 | 11 | Oxidation (M); Carbamidomethylation | 53366 | PPTG_12069 | Phytophthora parasitica INRA-310 serine protease family S10 (478 aa) |
| 931 | 814 | PPTG_09767 | 130,87 | 33 | 10 | 27 | 31 | 3,58E+05 | 6,13E+06 | 1,03E+07 | 11 | 11 | 3 | 10 | 11 | Carbamidomethylation; Deamidation (NQ) | 46427 | PPTG_09767 | Phytophthora parasitica INRA-310 hypothetical protein (419 aa) |
| 247 | 1420 | PPTG_16014 | 130,87 | 38 | 19 | 38 | 32 | 8,57E+07 | 4,41E+08 | 1,16E+09 | 6 | 6 | 7 | 26 | 40 | Oxidation (M); Deamidation (NQ); Carbamylation | 20952 | PPTG_16014 | Phytophthora parasitica INRA-310 60S ribosomal protein (191 aa) |
| 730 | 1932 | PPTG_10319 | 130,86 | 65 | 64 | 65 | 65 | 2,19E+07 | 1,06E+08 | 2,36E+08 | 6 | 6 | 4 | 12 | 15 | Deamidation (NQ); Carbamylation | 11448 | PPTG_10319 | Phytophthora parasitica INRA-310 60S acidic ribosomal protein P2 (114 aa) |
| 907 | 1451 | PPTG_08251 | 130,81 | 56 | 12 | 51 | 46 | 2,98E+06 | 7,94E+07 | 5,45E+07 | 8 | 8 | 2 | 12 | 11 | Oxidation (M); Deamidation (NQ) | 16618 | PPTG_08251 | Phytophthora parasitica INRA-310 nucleoredoxin (147 aa) |
| 1193 | 1452 | PPTG_15152 | 130,76 | 24 | 16 | 21 | 23 | 1,24E+06 | 5,53E+06 | 9,61E+06 | 7 | 7 | 4 | 7 | 7 | | 40756 | PPTG_15152 | Phytophthora parasitica INRA-310 casein kinase II subunit alpha-2 (349 aa) |
| 1087 | 1654 | PPTG_18704 | 130,74 | 59 | 17 | 47 | 59 | 9,24E+06 | 3,96E+07 | 2,53E+08 | 5 | 5 | 2 | 6 | 12 | Deamidation (NQ) | 17563 | PPTG_18704 | Phytophthora parasitica INRA-310 ATP synthase subunit delta' (170 aa) |
| 826 | 724 | PPTG_08911 | 130,63 | 34 | 8 | 34 | 21 | 4,83E+05 | 8,47E+06 | 8,92E+06 | 11 | 9 | 3 | 14 | 10 | Oxidation (M); Carbamidomethylation | 47693 | PPTG_08911 | Phytophthora parasitica INRA-310 crinkler (CRN) family protein (421 aa) |
| 1353 | 928 | PPTG_00928 | 130,59 | 27 | 2 | 27 | 14 | 4,79E+04 | 4,96E+06 | 3,61E+06 | 9 | 9 | 1 | 9 | 5 | Carbamidomethylation | 49109 | PPTG_00928 | Phytophthora parasitica INRA-310 chorismate synthase (457 aa) |
| 876 | 898 | PPTG_06939 | 130,52 | 29 | 26 | 23 | 27 | 1,16E+06 | 5,16E+06 | 8,41E+06 | 9 | 8 | 9 | 8 | 9 | Carbamidomethylation | 45921 | PPTG_06939 | Phytophthora parasitica INRA-310 short/branched chain specific acyl-CoA dehydrogenase (424 aa) |
| 1014 | 877 | PPTG_10094 | 130,42 | 44 | 12 | 31 | 37 | 7,12E+05 | 8,61E+06 | 1,18E+07 | 10 | 9 | 4 | 8 | 10 | Oxidation (M); Carbamidomethylation; Dehydration | 35198 | PPTG_10094 | Phytophthora parasitica INRA-310 serine/threonine-protein phosphatase PP-X isozyme 2 (309 aa) |
| 652 | 863 | PPTG_00108 | 130,39 | 44 | 20 | 36 | 44 | 2,48E+06 | 1,00E+07 | 3,64E+07 | 12 | 8 | 6 | 11 | 17 | Oxidation (M); Carbamidomethylation; Dehydration; Deamidation (NQ) | 32483 | PPTG_00108 | Phytophthora parasitica INRA-310 phospholipid hydroperoxide glutathione peroxidase (289 aa) |
| 994 | 770 | PPTG_06221 | 130,38 | 44 | 13 | 13 | 41 | 7,59E+05 | 1,34E+06 | 1,80E+07 | 12 | 12 | 4 | 4 | 14 | Oxidation (M); Deamidation (NQ) | 34704 | PPTG_06221 | Phytophthora parasitica INRA-310 Rab11 family GTPase (316 aa) |
| 1008 | 744 | PPTG_04178 | 130,32 | 25 | 5 | 12 | 25 | 1,32E+06 | 5,37E+06 | 2,34E+07 | 10 | 10 | 3 | 6 | 13 | Oxidation (M); Carbamidomethylation | 59428 | PPTG_04178 | Phytophthora parasitica INRA-310 exopolysaccharide phosphotransferase (516 aa) |
| 771 | 756 | PPTG_11749 | 130,25 | 9 | 2 | 7 | 9 | 2,61E+05 | 3,00E+06 | 7,67E+06 | 15 | 15 | 3 | 12 | 14 | Oxidation (M) | 195415 | PPTG_11749 | Phytophthora parasitica INRA-310 vacuolar protein sorting-associated protein 8 (1776 aa) |
| 793 | 981 | PPTG_14070 | 130,22 | 14 | 3 | 7 | 13 | 8,77E+05 | 1,16E+07 | 1,44E+07 | 16 | 12 | 3 | 9 | 16 | Carbamidomethylation | 149128 | PPTG_14070 | Phytophthora parasitica INRA-310 ATP-binding Cassette (ABC) superfamily (1346 aa) |
| 1059 | 821 | PPTG_01494 | 130,18 | 31 | 6 | 22 | 23 | 1,04E+06 | 8,90E+06 | 1,39E+07 | 12 | 12 | 3 | 8 | 10 | Oxidation (M) | 55690 | PPTG_01494 | Phytophthora parasitica INRA-310 hypothetical protein (502 aa) |
| 1059 | 822 | PPTG_19733 | 130,18 | 31 | 6 | 22 | 23 | 1,04E+06 | 8,90E+06 | 1,39E+07 | 12 | 12 | 3 | 8 | 10 | Oxidation (M) | 55690 | PPTG_19733 | Phytophthora parasitica INRA-310 hypothetical protein (502 aa) |
| 1397 | 1311 | PPTG_18914 | 130,15 | 18 | 7 | 9 | 18 | | 5,08E+05 | 2,31E+06 | 8 | 2 | 3 | 4 | 8 | Carbamidomethylation | 54488 | PPTG_18914 | Phytophthora parasitica INRA-310 hypothetical protein (489 aa) |
| 904 | 986 | PPTG_12028 | 130,14 | 29 | 10 | 25 | 29 | 1,19E+06 | 6,66E+06 | 1,69E+07 | 8 | 5 | 3 | 11 | 11 | Oxidation (M); Carbamidomethylation | 38416 | PPTG_12028 | Phytophthora parasitica INRA-310 D-isomer specific 2-hydroxyacid dehydrogenase (353 aa) |
| 1294 | 792 | PPTG_09959 | 130,1 | 23 | 3 | 2 | 23 | 1,38E+05 | 1,52E+05 | 5,13E+06 | 12 | 12 | 2 | 1 | 13 | Carbamidomethylation | 68460 | PPTG_09959 | Phytophthora parasitica INRA-310 hypothetical protein (612 aa) |
| 1080 | 845 | PPTG_08368 | 130,07 | 31 | 6 | 24 | 26 | 2,28E+05 | 4,61E+06 | 4,47E+06 | 10 | 10 | 2 | 9 | 9 | Oxidation (M); Carbamidomethylation | 53142 | PPTG_08368 | Phytophthora parasitica INRA-310 betaine aldehyde dehydrogenase (492 aa) |
| 850 | 933 | PPTG_01003 | 129,74 | 37 | 11 | 33 | 34 | 3,97E+06 | 2,20E+07 | 4,55E+07 | 11 | 11 | 4 | 10 | 13 | Oxidation (M); Carbamidomethylation | 34309 | PPTG_01003 | Phytophthora parasitica INRA-310 actin-like protein 2/3 complex subunit (304 aa) |
| 1169 | 1231 | PPTG_18442 | 129,71 | 19 | 6 | 19 | 17 | 1,08E+05 | 1,03E+07 | 1,01E+07 | 6 | 6 | 1 | 10 | 7 | Oxidation (M); Carbamidomethylation | 46904 | PPTG_18442 | Phytophthora parasitica INRA-310 UV excision repair protein RAD23 (468 aa) |
| 701 | 728 | PPTG_00246 | 129,59 | 32 | 11 | 22 | 28 | 5,55E+06 | 2,91E+07 | 5,73E+07 | 14 | 14 | 5 | 11 | 16 | Oxidation (M); Carbamidomethylation; Acetylation (N-term) | 49796 | PPTG_00246 | Phytophthora parasitica INRA-310 AP-2 complex subunit mu (438 aa) |
| 722 | 889 | PPTG_17096 | 129,58 | 29 | 12 | 18 | 29 | 2,79E+06 | 8,93E+06 | 6,53E+07 | 12 | 10 | 6 | 9 | 16 | Oxidation (M); Carbamidomethylation | 49733 | PPTG_17096 | Phytophthora parasitica INRA-310 protein kinase (446 aa) |
| 1271 | 1443 | PPTG_05079 | 129,52 | 23 | 4 | 23 | 13 | 3,74E+05 | 1,19E+07 | 7,40E+06 | 6 | 6 | 2 | 9 | 6 | Carbamidomethylation | 47130 | PPTG_05079 | Phytophthora parasitica INRA-310 hypothetical protein (448 aa) |
| 1009 | 791 | PPTG_14299 | 129,5 | 65 | 17 | 47 | 59 | 5,54E+05 | 4,56E+06 | 1,33E+07 | 11 | 11 | 3 | 8 | 11 | Oxidation (M); Carbamidomethylation; Acetylation (N-term) | 24340 | PPTG_14299 | Phytophthora parasitica INRA-310 oxidoreductase htatip2-like protein (224 aa) |
| 185 | 1414 | PPTG_08299 | 129,47 | 11 | 6 | 11 | 11 | 4,03E+07 | 1,51E+08 | 5,53E+08 | 10 | 10 | 13 | 28 | 44 | Oxidation (M); Dehydration; Deamidation (NQ); Carbamylation | 69602 | PPTG_08299 | Phytophthora parasitica INRA-310 Ser/Thr kinase (619 aa) |
| 1232 | 989 | PPTG_13136 | 129,4 | 18 | 1 | 1 | 18 | 4,59E+05 | 4,42E+05 | 1,34E+07 | 12 | 12 | 1 | 1 | 15 | Oxidation (M); Carbamidomethylation | 94748 | PPTG_13136 | Phytophthora parasitica INRA-310 voltage-gated Ion Channel (VIC) superfamily (862 aa) |
| 677 | 895 | PPTG_02717 | 129,32 | 31 | 5 | 30 | 26 | 4,04E+05 | 7,72E+06 | 5,72E+06 | 17 | 17 | 3 | 16 | 14 | Carbamidomethylation; Carbamylation | 76738 | PPTG_02717 | Phytophthora parasitica INRA-310 hsp75-like protein (688 aa) |
| 680 | 829 | PPTG_19449 | 129,32 | 25 | 13 | 15 | 25 | 9,88E+06 | 1,29E+07 | 8,82E+07 | 12 | 12 | 6 | 9 | 18 | Oxidation (M); Deamidation (NQ) | 52454 | PPTG_19449 | Phytophthora parasitica INRA-310 protein transporter Sec61 subunit alpha (475 aa) |
| 895 | 954 | PPTG_02437 | 129,32 | 33 | 9 | 25 | 33 | 3,47E+05 | 1,03E+07 | 7,90E+06 | 9 | 9 | 3 | 10 | 12 | Oxidation (M); Carbamidomethylation; Amidation | 36610 | PPTG_02437 | Phytophthora parasitica INRA-310 protein phosphatase 2C (345 aa) |
| 845 | 794 | PPTG_09868 | 129,09 | 19 | 4 | 16 | 16 | 1,86E+06 | 1,10E+07 | 1,83E+07 | 14 | 14 | 3 | 13 | 11 | | 92412 | PPTG_09868 | Phytophthora parasitica INRA-310 endo-1 3(4)-beta-glucanase (825 aa) |
| 1096 | 787 | PPTG_04696 | 129,05 | 28 | 3 | 16 | 25 | 7,43E+04 | 5,04E+06 | 5,91E+06 | 13 | 13 | 1 | 8 | 11 | Carbamidomethylation | 70438 | PPTG_04696 | Phytophthora parasitica INRA-310 xaa-Pro aminopeptidase (634 aa) |
| 1120 | 1022 | PPTG_19282 | 129,04 | 27 | 7 | 19 | 22 | 5,57E+05 | 4,48E+06 | 9,77E+06 | 11 | 11 | 2 | 7 | 10 | Oxidation (M) | 51552 | PPTG_19282 | Phytophthora parasitica INRA-310 hypothetical protein (474 aa) |
| 993 | 809 | PPTG_06658 | 128,99 | 11 | 1 | 2 | 11 | 1,39E+05 | 3,76E+05 | 1,17E+07 | 16 | 16 | 1 | 4 | 17 | Oxidation (M); Carbamidomethylation | 179571 | PPTG_06658 | Phytophthora parasitica INRA-310 hypothetical protein (1593 aa) |
| 1307 | 858 | PPTG_06981 | 128,92 | 35 | 4 | 15 | 35 | 3,41E+04 | 1,25E+06 | 7,42E+06 | 10 | 10 | 1 | 4 | 11 | Carbamidomethylation; Acetylation (N-term) | 42796 | PPTG_06981 | Phytophthora parasitica INRA-310 hypothetical protein (370 aa) |
| 991 | 796 | PPTG_02605 | 128,83 | 11 | 1 | 6 | 10 | 6,49E+04 | 2,46E+06 | 4,39E+06 | 14 | 14 | 1 | 8 | 13 | Oxidation (M); Carbamidomethylation | 175460 | PPTG_02605 | Phytophthora parasitica INRA-310 hypothetical protein (1534 aa) |
| 1052 | 2338 | PPTG_05233 | 128,69 | 61 | 43 | 61 | 52 | 2,40E+05 | 2,95E+06 | 2,73E+06 | 6 | 6 | 3 | 9 | 9 | Carbamidomethylation; Deamidation (NQ); Acetylation (N-term) | 12115 | PPTG_05233 | Phytophthora parasitica INRA-310 eukaryotic translation initiation factor 1b (110 aa) |
| 650 | 911 | PPTG_02739 | 128,58 | 48 | 34 | 48 | 48 | 5,44E+06 | 3,57E+07 | 5,16E+07 | 9 | 9 | 8 | 13 | 13 | Oxidation (M) | 22269 | PPTG_02739 | Phytophthora parasitica INRA-310 glutathione S-transferase (201 aa) |
| 243 | 1934 | PPTG_15683 | 128,29 | 77 | 77 | 77 | 77 | 2,17E+07 | 7,39E+07 | 2,27E+08 | 4 | 4 | 15 | 20 | 39 | Oxidation (M); Dehydration; Deamidation (NQ); Acetylation (N-term); Carbamylation | 8616 | PPTG_15683 | Phytophthora parasitica INRA-310 30S ribosomal protein S21e (80 aa) |
| 953 | 811 | PPTG_05330 | 128,26 | 11 | 2 | 2 | 10 | 2,29E+06 | 8,30E+06 | 2,85E+07 | 16 | 16 | 4 | 4 | 15 | Oxidation (M); Carbamidomethylation | 193033 | PPTG_05330 | Phytophthora parasitica INRA-310 hypothetical protein (1704 aa) |
| 1132 | 919 | PPTG_04097 | 128,24 | 20 | 6 | 3 | 20 | 4,83E+05 | 3,18E+05 | 1,03E+07 | 13 | 12 | 4 | 2 | 13 | | 77780 | PPTG_04097 | Phytophthora parasitica INRA-310 hypothetical protein (697 aa) |
| 742 | 1130 | PPTG_13095 | 127,99 | 7 | 2 | 3 | 5 | 8,92E+05 | 3,37E+06 | 5,80E+06 | 18 | 18 | 6 | 9 | 15 | Oxidation (M) | 334614 | PPTG_13095 | Phytophthora parasitica INRA-310 ubiquitin-specific protease (2985 aa) |
| 1293 | 1583 | PPTG_05122 | 127,91 | 42 | 8 | 25 | 42 | 3,14E+05 | 1,07E+07 | 8,90E+06 | 6 | 6 | 1 | 8 | 7 | Carbamidomethylation; Dehydration; Deamidation (NQ) | 21352 | PPTG_05122 | Phytophthora parasitica INRA-310 hypothetical protein (198 aa) |
| 1380 | 1120 | PPTG_12093 | 127,9 | 41 | 14 | 19 | 41 | 6,79E+05 | 8,44E+05 | 7,10E+06 | 7 | 7 | 3 | 4 | 8 | | 34310 | PPTG_12093 | Phytophthora parasitica INRA-310 syntaxin-like protein (302 aa) |
| 1164 | 1006 | PPTG_00371 | 127,85 | 25 | 10 | 19 | 25 | 3,34E+05 | 2,67E+06 | 3,75E+06 | 8 | 4 | 3 | 7 | 8 | | 40530 | PPTG_00371 | Phytophthora parasitica INRA-310 voltage-gated potassium channel subunit beta (364 aa) |
| 1001 | 833 | PPTG_15675 | 127,84 | 33 | 6 | 30 | 25 | 4,20E+05 | 9,15E+06 | 7,14E+06 | 11 | 11 | 2 | 12 | 8 | Oxidation (M); Carbamidomethylation | 52098 | PPTG_15675 | Phytophthora parasitica INRA-310 hypothetical protein (471 aa) |
| 712 | 988 | PPTG_06654 | 127,73 | 53 | 27 | 32 | 46 | 5,21E+06 | 1,83E+07 | 7,71E+07 | 12 | 10 | 8 | 8 | 16 | Oxidation (M); Carbamidomethylation; Dehydration; Acetylation (N-term); Carbamylation | 26394 | PPTG_06654 | Phytophthora parasitica INRA-310 adenylate kinase (242 aa) |
| 892 | 880 | PPTG_15477 | 127,52 | 25 | 9 | 21 | 23 | 1,94E+06 | 1,04E+07 | 3,19E+07 | 9 | 9 | 3 | 11 | 11 | Carbamidomethylation; Deamidation (NQ); Carbamylation | 53028 | PPTG_15477 | Phytophthora parasitica INRA-310 hypothetical protein (479 aa) |
| 1100 | 957 | PPTG_03066 | 127,45 | 36 | 2 | 34 | 31 | 1,70E+05 | 3,04E+06 | 1,10E+07 | 10 | 10 | 1 | 9 | 10 | Oxidation (M) | 42677 | PPTG_03066 | Phytophthora parasitica INRA-310 C2 domain-containing protein (383 aa) |
| 1040 | 1087 | PPTG_15304 | 127,31 | 36 | 14 | 36 | 33 | 5,50E+05 | 7,29E+06 | 8,51E+06 | 7 | 7 | 3 | 9 | 9 | Carbamidomethylation; Deamidation (NQ) | 29278 | PPTG_15304 | Phytophthora parasitica INRA-310 glycerol-3-phosphate dehydrogenase (270 aa) |
| 750 | 886 | PPTG_02025 | 127,25 | 50 | 27 | 31 | 50 | 6,25E+06 | 2,42E+07 | 5,33E+07 | 12 | 12 | 8 | 8 | 14 | Carbamidomethylation | 29333 | PPTG_02025 | Phytophthora parasitica INRA-310 hypothetical protein (264 aa) |
| 885 | 1399 | PPTG_00954 | 127,05 | 31 | 24 | 24 | 31 | 1,24E+07 | 1,63E+07 | 1,15E+08 | 9 | 9 | 6 | 6 | 14 | | 22706 | PPTG_00954 | Phytophthora parasitica INRA-310 protein disulfide-isomerase (211 aa) |
| 1263 | 1700 | PPTG_19262 | 126,88 | 17 | 0 | 17 | 12 | | 2,09E+07 | 1,44E+07 | 7 | 7 | 0 | 10 | 7 | Carbamidomethylation | 59845 | PPTG_19262 | Phytophthora parasitica INRA-310 hypothetical protein (553 aa) |
| 1268 | 1379 | PPTG_13508 | 126,87 | 47 | 25 | 32 | 47 | 9,13E+05 | 1,76E+06 | 1,93E+07 | 7 | 7 | 3 | 4 | 10 | Oxidation (M); Carbamidomethylation | 17798 | PPTG_13508 | Phytophthora parasitica INRA-310 hypothetical protein (172 aa) |
| 1218 | 801 | PPTG_09255 | 126,84 | 13 | 1 | 7 | 11 | 2,17E+05 | 2,68E+06 | 4,62E+06 | 11 | 11 | 1 | 6 | 10 | Oxidation (M); Carbamidomethylation; Deamidation (NQ) | 114143 | PPTG_09255 | Phytophthora parasitica INRA-310 vacuolar protein sorting-associated protein (1009 aa) |
| 1907 | 1720 | PPTG_17901 | 126,78 | 25 | 7 | 4 | 25 | 1,43E+06 | 8,95E+05 | 1,49E+07 | 5 | 5 | 2 | 1 | 6 | Oxidation (M); Carbamidomethylation; Deamidation (NQ) | 39111 | PPTG_17901 | Phytophthora parasitica INRA-310 hypothetical protein (360 aa) |
| 1097 | 1047 | PPTG_00641 | 126,67 | 7 | 1 | 4 | 6 | 8,00E+04 | 1,28E+06 | 1,61E+06 | 13 | 13 | 2 | 9 | 9 | Carbamidomethylation | 258491 | PPTG_00641 | Phytophthora parasitica INRA-310 microtubule-associated protein (2400 aa) |
| 875 | 869 | PPTG_04185 | 126,62 | 14 | 2 | 7 | 14 | 1,29E+05 | 2,11E+06 | 5,05E+06 | 16 | 16 | 2 | 9 | 15 | Carbamidomethylation | 165029 | PPTG_04185 | Phytophthora parasitica INRA-310 hypothetical protein (1515 aa) |
| 707 | 1023 | PPTG_00550 | 126,6 | 18 | 1 | 6 | 18 | 1,83E+05 | 2,86E+06 | 2,05E+07 | 22 | 22 | 1 | 9 | 22 | Oxidation (M); Carbamidomethylation | 156448 | PPTG_00550 | Phytophthora parasitica INRA-310 hypothetical protein (1377 aa) |
| 1739 | 1635 | PPTG_11053 | 126,43 | 47 | 23 | 47 | 43 | 1,09E+06 | 3,36E+07 | 1,42E+07 | 5 | 5 | 2 | 5 | 4 | Carbamidomethylation | 19150 | PPTG_11053 | Phytophthora parasitica INRA-310 hypothetical protein (181 aa) |
| 739 | 915 | PPTG_01937 | 126,41 | 48 | 37 | 23 | 48 | 1,01E+07 | 1,82E+07 | 1,19E+08 | 11 | 11 | 9 | 6 | 16 | Oxidation (M); Carbamidomethylation | 25861 | PPTG_01937 | Phytophthora parasitica INRA-310 ATP synthase (E/31 kDa) subunit (227 aa) |
| 1033 | 940 | PPTG_14107 | 126,4 | 7 | 2 | 6 | 4 | 1,41E+05 | 1,80E+06 | 2,81E+06 | 12 | 12 | 3 | 11 | 7 | Carbamidomethylation | 233978 | PPTG_14107 | Phytophthora parasitica INRA-310 hypothetical protein (2107 aa) |
| 1251 | 1395 | PPTG_02660 | 126,37 | 61 | 12 | 61 | 61 | 1,25E+05 | 9,03E+06 | 7,65E+06 | 8 | 8 | 1 | 8 | 8 | Oxidation (M) | 15371 | PPTG_02660 | Phytophthora parasitica INRA-310 hypothetical protein (137 aa) |
| 1079 | 773 | PPTG_05866 | 126,37 | 32 | 3 | 29 | 20 | 1,55E+05 | 3,68E+06 | 2,26E+06 | 11 | 9 | 1 | 11 | 8 | Oxidation (M) | 47884 | PPTG_05866 | Phytophthora parasitica INRA-310 crinkler (CRN) family protein (422 aa) |
| 1366 | 1534 | PPTG_01055 | 126,19 | 21 | 12 | 18 | 21 | 1,87E+05 | 5,01E+06 | 5,99E+06 | 5 | 5 | 2 | 6 | 7 | Carbamidomethylation | 41475 | PPTG_01055 | Phytophthora parasitica INRA-310 hypothetical protein (367 aa) |
| 808 | 1111 | PPTG_08494 | 126,15 | 49 | 18 | 32 | 49 | 2,84E+06 | 7,03E+06 | 4,90E+07 | 11 | 11 | 4 | 8 | 16 | Carbamidomethylation; Deamidation (NQ) | 22123 | PPTG_08494 | Phytophthora parasitica INRA-310 hypothetical protein (204 aa) |
| 859 | 861 | PPTG_18270 | 126,15 | 31 | 4 | 31 | 21 | 1,30E+06 | 2,85E+07 | 2,92E+07 | 13 | 13 | 2 | 16 | 9 | Oxidation (M); Carbamidomethylation; Deamidation (NQ) | 52688 | PPTG_18270 | Phytophthora parasitica INRA-310 glycoside hydrolase (465 aa) |
| 960 | 1015 | PPTG_02405 | 126,11 | 22 | 13 | 18 | 22 | 6,21E+05 | 1,08E+07 | 9,61E+06 | 9 | 9 | 4 | 9 | 10 | Carbamidomethylation; Deamidation (NQ); Carbamylation | 53741 | PPTG_02405 | Phytophthora parasitica INRA-310 Ras GTPase-activating protein-binding protein (501 aa) |
| 1036 | 841 | PPTG_07323 | 126,01 | 11 | 2 | 1 | 11 | 3,22E+05 | 2,31E+05 | 1,32E+07 | 14 | 14 | 3 | 2 | 16 | Carbamidomethylation | 201768 | PPTG_07323 | Phytophthora parasitica INRA-310 hypothetical protein (1801 aa) |
| 1155 | 951 | PPTG_16825 | 125,85 | 9 | 0 | 5 | 9 | | 2,41E+06 | 8,70E+06 | 11 | 11 | 0 | 6 | 12 | Carbamidomethylation | 148184 | PPTG_16825 | Phytophthora parasitica INRA-310 myosin-like protein (1304 aa) |
| 1215 | 805 | PPTG_06806 | 125,77 | 18 | 5 | 3 | 18 | 4,89E+05 | 4,14E+05 | 1,00E+07 | 12 | 12 | 3 | 2 | 12 | Carbamidomethylation; Deamidation (NQ) | 90013 | PPTG_06806 | Phytophthora parasitica INRA-310 hypothetical protein (810 aa) |
| 1612 | 1265 | PPTG_08695 | 125,76 | 38 | 0 | 38 | 25 | | 4,20E+06 | 4,74E+06 | 7 | 7 | 0 | 7 | 5 | Carbamidomethylation | 30252 | PPTG_08695 | Phytophthora parasitica INRA-310 replication protein A 32 kDa subunit (281 aa) |
| 1827 | 1946 | PPTG_01465 | 125,71 | 45 | 5 | 45 | 45 | 1,16E+05 | 2,75E+06 | 4,45E+06 | 5 | 5 | 1 | 4 | 5 | Carbamidomethylation | 18312 | PPTG_01465 | Phytophthora parasitica INRA-310 hypothetical protein (166 aa) |

| | | | | | | | | | | | | | | | | | | | |
|---|---|---|---|---|---|---|---|---|---|---|---|---|---|---|---|---|---|---|---|
| 1048 | 1071 | PPTG_06104 | 125,68 | 26 | 10 | 15 | 26 | 3,07E+06 | 8,30E+06 | 4,52E+07 | 10 | 10 | 4 | 6 | 10 | Carbamidomethylation | 45579 | PPTG_06104 | Phytophthora parasitica INRA-310 proline iminopeptidase (404 aa) |
| 752 | 1942 | PPTG_08333 | 125,62 | 48 | 36 | 48 | 48 | 7,76E+07 | 3,11E+08 | 8,39E+08 | 6 | 6 | 5 | 10 | 15 | Deamidation (NQ); Carbamylation | 10994 | PPTG_08333 | Phytophthora parasitica INRA-310 hypothetical protein (99 aa) |
| 1213 | 1068 | PPTG_09698 | 125,61 | 14 | 4 | 14 | 12 | 6,03E+04 | 3,28E+06 | 2,99E+06 | 7 | 7 | 2 | 8 | 7 | Carbamidomethylation | 79974 | PPTG_09698 | Phytophthora parasitica INRA-310 phospholipase A-2-activating protein (749 aa) |
| 863 | 2314 | PPTG_07986 | 125,58 | 19 | 6 | 15 | 19 | 1,32E+06 | 2,52E+07 | 3,08E+07 | 3 | 3 | 4 | 11 | 12 | Oxidation (M); Carbamidomethylation; Acetylation (N-term) | 31994 | PPTG_07986 | Phytophthora parasitica INRA-310 hypothetical protein (292 aa) |
| 883 | 802 | PPTG_17400 | 125,45 | 30 | 6 | 18 | 30 | 6,63E+05 | 6,07E+06 | 1,79E+07 | 10 | 10 | 3 | 8 | 15 | Oxidation (M) | 48770 | PPTG_17400 | Phytophthora parasitica INRA-310 AP-1 complex subunit mu-1 (426 aa) |
| 998 | 1077 | PPTG_00686 | 125,44 | 21 | 5 | 19 | 19 | 3,99E+05 | 4,86E+06 | 9,16E+06 | 9 | 9 | 2 | 10 | 10 | Carbamidomethylation | 58492 | PPTG_00686 | Phytophthora parasitica INRA-310 threonine synthase (533 aa) |
| 1411 | 969 | PPTG_06672 | 125,41 | 11 | 4 | 4 | 11 | 2,98E+05 | 6,78E+05 | 1,10E+07 | 8 | 8 | 3 | 3 | 8 | Carbamidomethylation | 98061 | PPTG_06672 | Phytophthora parasitica INRA-310 hypothetical protein (891 aa) |
| 1498 | 862 | PPTG_09306 | 125,41 | 14 | 3 | 3 | 14 | 1,25E+05 | 2,94E+05 | 4,09E+06 | 9 | 9 | 2 | 2 | 9 | Carbamidomethylation | 93332 | PPTG_09306 | Phytophthora parasitica INRA-310 hypothetical protein (837 aa) |
| 1102 | 1001 | PPTG_02890 | 125,38 | 33 | 19 | 27 | 31 | 7,71E+05 | 5,32E+06 | 1,26E+07 | 8 | 8 | 4 | 8 | 8 | Carbamidomethylation | 40426 | PPTG_02890 | Phytophthora parasitica INRA-310 protein kinase (367 aa) |
| 1296 | 1040 | PPTG_18123 | 125,18 | 17 | 5 | 10 | 17 | 3,02E+05 | 1,41E+06 | 6,65E+06 | 8 | 8 | 2 | 5 | 9 | | 65430 | PPTG_18123 | Phytophthora parasitica INRA-310 sulfite oxidase (588 aa) |
| 1167 | 1065 | PPTG_03585 | 125,12 | 24 | 9 | 24 | 17 | 3,90E+05 | 5,35E+06 | 4,66E+06 | 8 | 8 | 3 | 9 | 6 | Oxidation (M); Carbamidomethylation; Dehydration; Deamidation (NQ) | 47776 | PPTG_03585 | Phytophthora parasitica INRA-310 ornithine aminotransferase mitochondrial (433 aa) |
| 917 | 912 | PPTG_15689 | 125,11 | 17 | 3 | 12 | 11 | 1,05E+05 | 8,36E+06 | 5,65E+06 | 15 | 15 | 2 | 11 | 11 | Carbamidomethylation | 109730 | PPTG_15689 | Phytophthora parasitica INRA-310 hypothetical protein (981 aa) |
| 1458 | 1072 | PPTG_11307 | 125,01 | 46 | 5 | 42 | 15 | 9,42E+04 | 5,58E+06 | 2,70E+06 | 10 | 10 | 1 | 9 | 4 | Carbamidomethylation | 35317 | PPTG_11307 | Phytophthora parasitica INRA-310 SUMO-activating enzyme (SAE) (321 aa) |
| 987 | 883 | PPTG_11789 | 124,89 | 11 | 2 | 9 | 7 | 3,70E+05 | 3,01E+06 | 2,86E+06 | 13 | 13 | 3 | 10 | 9 | Carbamidomethylation; Deamidation (NQ) | 164186 | PPTG_11789 | Phytophthora parasitica INRA-310 hypothetical protein (1473 aa) |
| 695 | 1314 | PPTG_04051 | 124,85 | 55 | 25 | 32 | 55 | 6,69E+07 | 2,88E+08 | 8,10E+08 | 8 | 8 | 5 | 10 | 17 | Oxidation (M); Carbamidomethylation; Carbamylation | 13481 | PPTG_04051 | Phytophthora parasitica INRA-310 40S ribosomal protein S20 (121 aa) |
| 893 | 1000 | PPTG_06888 | 124,82 | 21 | 2 | 11 | 17 | 2,43E+05 | 4,38E+06 | 1,23E+07 | 17 | 17 | 2 | 8 | 15 | Carbamidomethylation | 111399 | PPTG_06888 | Phytophthora parasitica INRA-310 hypothetical protein (1016 aa) |
| 1428 | 1673 | PPTG_13126 | 124,58 | 14 | 8 | 12 | 14 | 4,60E+05 | 9,52E+05 | 1,04E+07 | 5 | 5 | 3 | 4 | 7 | Carbamidomethylation | 61157 | PPTG_13126 | Phytophthora parasitica INRA-310 hypothetical protein (545 aa) |
| 1028 | 914 | PPTG_06837 | 124,51 | 31 | 6 | 23 | 24 | 2,40E+05 | 6,35E+06 | 5,56E+06 | 12 | 12 | 2 | 9 | 10 | Oxidation (M); Carbamidomethylation | 57822 | PPTG_06837 | Phytophthora parasitica INRA-310 hypothetical protein (524 aa) |
| 1324 | 1132 | PPTG_13472 | 124,48 | 25 | 7 | 25 | 13 | 2,20E+05 | 6,65E+06 | 5,12E+06 | 7 | 7 | 2 | 9 | 5 | Carbamidomethylation; Deamidation (NQ) | 51363 | PPTG_13472 | Phytophthora parasitica INRA-310 hypothetical protein (465 aa) |
| 934 | 894 | PPTG_02241 | 124,33 | 24 | 0 | 22 | 22 | | 1,48E+07 | 2,41E+07 | 11 | 11 | 0 | 11 | 13 | Carbamidomethylation | 58332 | PPTG_02241 | Phytophthora parasitica INRA-310 coatomer subunit delta (537 aa) |
| 1191 | 827 | PPTG_05501 | 124,27 | 20 | 0 | 10 | 20 | | 2,05E+06 | 5,14E+06 | 11 | 11 | 0 | 6 | 12 | Oxidation (M); Carbamidomethylation | 70916 | PPTG_05501 | Phytophthora parasitica INRA-310 acetolactate synthase large subunit biosynthetic type (649 aa) |
| 1488 | 1639 | PPTG_12273 | 124,19 | 45 | 10 | 34 | 45 | 1,15E+06 | 8,02E+06 | 1,37E+07 | 6 | 6 | 1 | 5 | 7 | Oxidation (M); Deamidation (NQ) | 19537 | PPTG_12273 | Phytophthora parasitica INRA-310 caltractin (173 aa) |
| 1615 | 1498 | PPTG_16550 | 124,19 | 33 | 5 | 26 | 17 | 1,52E+05 | 6,02E+06 | 3,37E+06 | 6 | 6 | 1 | 7 | 4 | Carbamidomethylation | 42626 | PPTG_16550 | Phytophthora parasitica INRA-310 hypothetical protein (416 aa) |
| 1315 | 1048 | PPTG_13535 | 124,19 | 24 | 7 | 17 | 21 | 1,63E+05 | 2,87E+06 | 9,54E+06 | 8 | 8 | 3 | 6 | 7 | Carbamidomethylation | 48989 | PPTG_13535 | Phytophthora parasitica INRA-310 disulfide-isomerase (447 aa) |
| 1684 | 1278 | PPTG_02871 | 124,14 | 19 | 7 | 0 | 17 | 6,79E+05 | | 6,40E+06 | 7 | 7 | 3 | 0 | 8 | Carbamidomethylation | 55614 | PPTG_02871 | Phytophthora parasitica INRA-310 K+-dependent Na+/Ca+ exchanger (516 aa) |
| 1472 | 2820 | PPTG_14724 | 124,11 | 50 | 40 | 40 | 50 | 1,00E+07 | 5,66E+07 | 1,04E+08 | 4 | 4 | 4 | 4 | 6 | | 11869 | PPTG_14724 | Phytophthora parasitica INRA-310 hypothetical protein (118 aa) |
| 1111 | 823 | PPTG_18087 | 124,11 | 21 | 0 | 17 | 18 | | 1,03E+06 | 2,17E+06 | 11 | 11 | 0 | 9 | 10 | Acetylation (N-term) | 80689 | PPTG_18087 | Phytophthora parasitica INRA-310 intraflagellar transporter 81 (703 aa) |
| 1055 | 1277 | PPTG_10402 | 123,97 | 43 | 23 | 38 | 43 | 1,25E+06 | 1,06E+07 | 1,41E+07 | 7 | 7 | 4 | 8 | 9 | Oxidation (M); Carbamidomethylation | 24846 | PPTG_10402 | Phytophthora parasitica INRA-310 hypothetical protein (228 aa) |
| 1086 | 1617 | PPTG_10282 | 123,95 | 41 | 22 | 32 | 41 | 5,56E+05 | 4,36E+06 | 2,38E+07 | 6 | 6 | 2 | 7 | 11 | Oxidation (M); Deamidation (NQ) | 23014 | PPTG_10282 | Phytophthora parasitica INRA-310 ran-specific GTPase-activating protein (205 aa) |
| 788 | 901 | PPTG_18912 | 123,95 | 14 | 5 | 5 | 13 | 5,71E+05 | 8,30E+05 | 3,82E+06 | 16 | 1 | 7 | 6 | 16 | Oxidation (M); Carbamidomethylation | 157869 | PPTG_18912 | Phytophthora parasitica INRA-310 P-type ATPase (P-ATPase) superfamily (1403 aa) |
| 288 | 1324 | PPTG_16818 | 123,87 | 56 | 48 | 48 | 56 | 1,29E+08 | 4,96E+08 | 1,38E+09 | 6 | 6 | 12 | 20 | 34 | Oxidation (M); Dehydration; Deamidation (NQ); Carbamylation | 12326 | PPTG_16818 | Phytophthora parasitica INRA-310 S25 ribosomal protein (112 aa) |
| 153 | 1744 | PPTG_10118 | 123,83 | 47 | 30 | 46 | 47 | 1,29E+08 | 4,18E+08 | 1,23E+09 | 8 | 8 | 21 | 32 | 41 | Oxidation (M); Dehydration; Deamidation (NQ); Acetylation (N-term); Carbamylation | 12611 | PPTG_10118 | Phytophthora parasitica INRA-310 histone H2B (116 aa) |
| 153 | 1745 | PPTG_13452 | 123,83 | 46 | 30 | 45 | 46 | 1,29E+08 | 4,18E+08 | 1,23E+09 | 8 | 8 | 21 | 32 | 41 | Oxidation (M); Dehydration; Deamidation (NQ); Acetylation (N-term); Carbamylation | 12796 | PPTG_13452 | Phytophthora parasitica INRA-310 histone H2B (118 aa) |
| 1449 | 1705 | PPTG_10079 | 123,74 | 17 | 6 | 15 | 17 | | | 2,23E+05 | 5 | 1 | 2 | 7 | 5 | Oxidation (M); Carbamidomethylation; Amidation | 54123 | PPTG_10079 | Phytophthora parasitica INRA-310 alanine aminotransferase 2 (492 aa) |
| 1077 | 958 | PPTG_18702 | 123,73 | 37 | 12 | 30 | 16 | 6,35E+05 | 6,01E+06 | 3,86E+06 | 10 | 10 | 4 | 9 | 7 | Carbamidomethylation; Dehydration; Deamidation (NQ) | 41356 | PPTG_18702 | Phytophthora parasitica INRA-310 peroxisomal (S)-2-hydroxy-acid oxidase (383 aa) |
| 976 | 923 | PPTG_18295 | 123,67 | 44 | 6 | 40 | 36 | 5,78E+05 | 8,50E+06 | 8,20E+06 | 11 | 11 | 2 | 11 | 10 | Oxidation (M); Carbamidomethylation | 40612 | PPTG_18295 | Phytophthora parasitica INRA-310 protein MYG1 (366 aa) |
| 1219 | 828 | PPTG_07315 | 123,67 | 36 | 3 | 12 | 36 | 3,70E+05 | 1,13E+06 | 1,32E+07 | 11 | 11 | 1 | 4 | 12 | Oxidation (M); Carbamidomethylation; Deamidation (NQ); Acetylation (N-term) | 44659 | PPTG_07315 | Phytophthora parasitica INRA-310 hypothetical protein (408 aa) |
| 918 | 918 | PPTG_00975 | 123,6 | 5 | 0 | 4 | 5 | | 2,38E+06 | 5,84E+06 | 14 | 14 | 0 | 11 | 13 | Carbamidomethylation | 363808 | PPTG_00975 | Phytophthora parasitica INRA-310 hypothetical protein (3319 aa) |
| 973 | 972 | PPTG_07498 | 123,58 | 35 | 10 | 24 | 35 | 1,07E+06 | 1,49E+07 | 1,16E+07 | 9 | 9 | 4 | 9 | 10 | Oxidation (M); Carbamidomethylation; Deamidation (NQ); Acetylation (N-term) | 39016 | PPTG_07498 | Phytophthora parasitica INRA-310 3-isopropylmalate dehydrogenase (362 aa) |
| 817 | 832 | PPTG_11731 | 123,53 | 38 | 9 | 19 | 38 | 2,64E+06 | 6,23E+06 | 3,45E+07 | 12 | 12 | 4 | 8 | 16 | Oxidation (M); Carbamidomethylation | 40885 | PPTG_11731 | Phytophthora parasitica INRA-310 hypothetical protein (370 aa) |
| 1243 | 984 | PPTG_04179 | 123,5 | 23 | 4 | 14 | 23 | 1,83E+05 | 1,41E+06 | 6,53E+06 | 9 | 9 | 2 | 5 | 10 | Carbamidomethylation | 53932 | PPTG_04179 | Phytophthora parasitica INRA-310 exopolysaccharide phosphotransferase (478 aa) |
| 1374 | 1043 | PPTG_01569 | 123,47 | 19 | 5 | 12 | 18 | 3,05E+05 | 3,52E+06 | 8,85E+06 | 8 | 8 | 2 | 6 | 7 | Carbamidomethylation | 50315 | PPTG_01569 | Phytophthora parasitica INRA-310 eukaryotic translation initiation factor 5 (446 aa) |
| 1248 | 1619 | PPTG_11388 | 123,47 | 24 | 12 | 24 | 24 | 2,97E+05 | 1,50E+06 | 8,83E+06 | 6 | 6 | 3 | 6 | 8 | Carbamidomethylation; Deamidation (NQ) | 30713 | PPTG_11388 | Phytophthora parasitica INRA-310 copine-like protein (280 aa) |
| 1333 | 1907 | PPTG_18395 | 123,43 | 19 | 9 | 19 | 9 | 1,40E+05 | 3,68E+06 | 1,32E+06 | 5 | 4 | 3 | 8 | 5 | Carbamidomethylation; Deamidation (NQ) | 43365 | PPTG_18395 | Phytophthora parasitica INRA-310 hypothetical protein (412 aa) |
| 1246 | 1203 | PPTG_04003 | 123,35 | 55 | 14 | 55 | 55 | 6,05E+04 | 9,24E+06 | 8,31E+06 | 7 | 7 | 1 | 8 | 8 | Acetylation (N-term) | 19169 | PPTG_04003 | Phytophthora parasitica INRA-310 hypothetical protein (175 aa) |
| 1074 | 1242 | PPTG_19383 | 123,33 | 44 | 17 | 41 | 33 | 2,23E+06 | 1,26E+07 | 1,91E+07 | 7 | 7 | 4 | 9 | 7 | Carbamidomethylation | 24833 | PPTG_19383 | Phytophthora parasitica INRA-310 ATP:cob(I)alamin adenosyltransferase (225 aa) |
| 1090 | 968 | PPTG_18258 | 123,32 | 22 | 8 | 20 | 18 | 4,77E+05 | 3,65E+06 | 5,58E+06 | 10 | 10 | 3 | 9 | 8 | Carbamidomethylation | 59358 | PPTG_18258 | Phytophthora parasitica INRA-310 imidazole glycerol phosphate synthase hisHF (549 aa) |
| 1557 | 1008 | PPTG_07422 | 123,3 | 8 | 3 | 0 | 7 | 9,01E+04 | | 2,82E+06 | 10 | 5 | 3 | 0 | 9 | Oxidation (M); Carbamidomethylation; Deamidation (NQ) | 164347 | PPTG_07422 | Phytophthora parasitica INRA-310 resistance-Nodulation-Cell Division (RND) superfamily (1493 aa) |
| 1364 | 977 | PPTG_16766 | 123,27 | 36 | 5 | 34 | 26 | 7,21E+05 | 1,44E+07 | 5,62E+06 | 9 | 9 | 1 | 8 | 6 | Carbamidomethylation | 32961 | PPTG_16766 | Phytophthora parasitica INRA-310 peroxisomal hydratase-dehydrogenase-epimerase (299 aa) |
| 1195 | 1757 | PPTG_08738 | 123,2 | 42 | 10 | 42 | 32 | 1,51E+05 | 2,88E+07 | 2,29E+07 | 6 | 5 | 1 | 10 | 7 | Carbamidomethylation; Deamidation (NQ) | 18708 | PPTG_08738 | Phytophthora parasitica INRA-310 SCP-like extracellular protein (174 aa) |
| 1375 | 1052 | PPTG_01811 | 123,18 | 23 | 5 | 23 | 15 | 1,84E+05 | 4,77E+06 | 4,22E+06 | 8 | 8 | 2 | 8 | 5 | Carbamidomethylation | 51064 | PPTG_01811 | Phytophthora parasitica INRA-310 histidine biosynthesis trifunctional protein (478 aa) |
| 1018 | 1207 | PPTG_02424 | 123,16 | 41 | 8 | 27 | 38 | 1,15E+06 | 1,31E+07 | 5,80E+07 | 8 | 8 | 2 | 7 | 13 | Oxidation (M); Carbamidomethylation | 31040 | PPTG_02424 | Phytophthora parasitica INRA-310 succinate dehydrogenase [ubiquinone] iron-sulfur subunit mitochondrial (272 aa) |
| 1150 | 1661 | PPTG_19378 | 122,94 | 37 | 16 | 26 | 37 | 3,90E+06 | 1,15E+07 | 4,34E+07 | 5 | 5 | 3 | 7 | 9 | Carbamidomethylation; Deamidation (NQ) | 25545 | PPTG_19378 | Phytophthora parasitica INRA-310 endoglucanase (239 aa) |
| 1175 | 1147 | PPTG_02217 | 122,94 | 24 | 5 | 24 | 18 | 8,41E+05 | 1,38E+07 | 1,54E+07 | 7 | 7 | 2 | 9 | 7 | Oxidation (M); Carbamidomethylation; Deamidation (NQ) | 45589 | PPTG_02217 | Phytophthora parasitica INRA-310 glutaryl-CoA dehydrogenase mitochondrial (421 aa) |
| 1198 | 948 | PPTG_04264 | 122,87 | 50 | 7 | 17 | 50 | 5,88E+05 | 9,90E+05 | 1,14E+07 | 10 | 10 | 2 | 3 | 13 | Oxidation (M); Carbamidomethylation; Acetylation (N-term) | 35689 | PPTG_04264 | Phytophthora parasitica INRA-310 sideroflexin-1-like protein (326 aa) |
| 868 | 1046 | PPTG_06564 | 122,77 | 7 | 2 | 6 | 5 | 4,89E+05 | 3,33E+06 | 3,32E+06 | 15 | 15 | 5 | 12 | 9 | Carbamidomethylation | 256461 | PPTG_06564 | Phytophthora parasitica INRA-310 hypothetical protein (2377 aa) |
| 1405 | 1516 | PPTG_07614 | 122,65 | 25 | 18 | 21 | 22 | 8,54E+05 | 6,54E+06 | 2,26E+07 | 7 | 7 | 4 | 5 | 6 | | 31636 | PPTG_07614 | Phytophthora parasitica INRA-310 3-oxo-5-alpha-steroid 4-dehydrogenase (283 aa) |
| 1073 | 860 | PPTG_14384 | 122,59 | 16 | 2 | 10 | 13 | 3,55E+04 | 2,04E+06 | 5,10E+06 | 13 | 13 | 1 | 8 | 11 | | 110425 | PPTG_14384 | Phytophthora parasitica INRA-310 kinesin-like protein (977 aa) |
| 1047 | 1029 | PPTG_16603 | 122,26 | 20 | 3 | 15 | 17 | 5,39E+05 | 6,24E+06 | 1,46E+07 | 12 | 12 | 2 | 9 | 10 | Carbamidomethylation | 72131 | PPTG_16603 | Phytophthora parasitica INRA-310 hypothetical protein (631 aa) |
| 1529 | 1205 | PPTG_16493 | 122,11 | 32 | 3 | 26 | 23 | 6,34E+04 | 4,63E+06 | 2,57E+06 | 7 | 7 | 1 | 6 | 6 | Carbamidomethylation; Acetylation (N-term) | 36510 | PPTG_16493 | Phytophthora parasitica INRA-310 serine/threonine-protein phosphatase 6 catalytic subunit (321 aa) |
| 1197 | 897 | PPTG_13449 | 122,05 | 29 | 5 | 21 | 27 | 2,11E+05 | 4,54E+06 | 4,54E+06 | 9 | 9 | 2 | 7 | 9 | Carbamidomethylation; Acetylation (N-term) | 55226 | PPTG_13449 | Phytophthora parasitica INRA-310 serine/threonine-protein phosphatase 5 (495 aa) |
| 1379 | 1036 | PPTG_07816 | 122,02 | 37 | 13 | 19 | 32 | 5,34E+05 | 8,32E+05 | 6,56E+06 | 8 | 8 | 3 | 4 | 8 | Carbamidomethylation | 30927 | PPTG_07816 | Phytophthora parasitica INRA-310 Rab21/Rab5 family GTPase (278 aa) |
| 1092 | 849 | PPTG_11612 | 121,85 | 27 | 5 | 6 | 27 | 4,55E+05 | 1,29E+06 | 1,55E+07 | 11 | 11 | 3 | 3 | 14 | Carbamidomethylation | 64176 | PPTG_11612 | Phytophthora parasitica INRA-310 calcium-dependent protein kinase (582 aa) |
| 966 | 1002 | PPTG_17147 | 121,84 | 45 | 7 | 37 | 30 | 9,86E+05 | 9,40E+06 | 1,47E+07 | 13 | 13 | 2 | 12 | 9 | Oxidation (M); Carbamidomethylation; Acetylation (N-term) | 41008 | PPTG_17147 | Phytophthora parasitica INRA-310 hypothetical protein (373 aa) |
| 1867 | 1135 | PPTG_15953 | 121,73 | 25 | 3 | 0 | 25 | 7,09E+04 | | 4,60E+06 | 7 | 7 | 1 | 0 | 8 | Carbamidomethylation | 43606 | PPTG_15953 | Phytophthora parasitica INRA-310 hypothetical protein (386 aa) |
| 1257 | 1236 | PPTG_06099 | 121,58 | 41 | 9 | 33 | 36 | 4,33E+05 | 8,46E+06 | 8,95E+06 | 7 | 7 | 2 | 7 | 8 | Carbamidomethylation | 31248 | PPTG_06099 | Phytophthora parasitica INRA-310 S-formylglutathione hydrolase (283 aa) |
| 945 | 906 | PPTG_17439 | 121,47 | 41 | 12 | 32 | 41 | 4,54E+05 | 8,28E+06 | 1,24E+07 | 11 | 11 | 3 | 9 | 12 | Carbamidomethylation | 35518 | PPTG_17439 | Phytophthora parasitica INRA-310 NmrA-like family protein (318 aa) |
| 1856 | 1240 | PPTG_05462 | 121,34 | 7 | 1 | 0 | 7 | 4,71E+04 | | 2,33E+06 | 7 | 7 | 1 | 0 | 8 | | 170786 | PPTG_05462 | Phytophthora parasitica INRA-310 hypothetical protein (1541 aa) |
| 951 | 991 | PPTG_19332 | 121,31 | 14 | 11 | 11 | 14 | 1,90E+06 | 3,72E+06 | 2,52E+07 | 8 | 8 | 6 | 7 | 11 | Carbamidomethylation; Dehydration | 81048 | PPTG_19332 | Phytophthora parasitica INRA-310 choline transporter-like protein (746 aa) |
| 1431 | 1075 | PPTG_07828 | 121,3 | 27 | 2 | 11 | 27 | 9,01E+04 | 1,04E+06 | 1,01E+07 | 8 | 8 | 1 | 3 | 10 | Carbamidomethylation | 46993 | PPTG_07828 | Phytophthora parasitica INRA-310 lectin (418 aa) |
| 964 | 910 | PPTG_11903 | 121,29 | 24 | 2 | 15 | 24 | 2,42E+05 | 4,52E+06 | 1,01E+07 | 10 | 10 | 1 | 7 | 15 | Oxidation (M); Carbamidomethylation | 51510 | PPTG_11903 | Phytophthora parasitica INRA-310 RuvB-like 2 (464 aa) |
| 1252 | 1622 | PPTG_13854 | 121,29 | 44 | 23 | 44 | 44 | 8,03E+05 | 8,73E+06 | 2,15E+07 | 6 | 6 | 2 | 7 | 8 | Oxidation (M); Carbamidomethylation | 16757 | PPTG_13854 | Phytophthora parasitica INRA-310 ubiquitin-conjugating enzyme E2 35 (150 aa) |
| 1206 | 1270 | PPTG_05322 | 121,27 | 54 | 15 | 54 | 39 | 5,75E+05 | 1,08E+07 | 1,32E+07 | 7 | 7 | 2 | 8 | 8 | Deamidation (NQ); Acetylation (N-term) | 22582 | PPTG_05322 | Phytophthora parasitica INRA-310 hypothetical protein (197 aa) |
| 1235 | 1566 | PPTG_08558 | 121,26 | 28 | 22 | 21 | 25 | 1,31E+06 | 2,80E+06 | 2,02E+07 | 7 | 7 | 5 | 5 | 7 | Oxidation (M); Deamidation (NQ) | 29391 | PPTG_08558 | Phytophthora parasitica INRA-310 hypothetical protein (276 aa) |
| 1224 | 1217 | PPTG_00147 | 121,25 | 25 | 10 | 25 | 23 | 2,07E+06 | 1,98E+07 | 2,82E+07 | 7 | 7 | 3 | 8 | 6 | Carbamidomethylation | 35651 | PPTG_00147 | Phytophthora parasitica INRA-310 alcohol dehydrogenase (328 aa) |
| 924 | 983 | PPTG_11462 | 121,24 | 27 | 17 | 25 | 24 | 2,14E+06 | 2,12E+07 | 2,53E+07 | 10 | 10 | 6 | 9 | 9 | Carbamidomethylation | 43436 | PPTG_11462 | Phytophthora parasitica INRA-310 branched-chain amino acid aminotransferase (391 aa) |
| 874 | 1609 | PPTG_00848 | 120,98 | 38 | 16 | 29 | 38 | 1,23E+06 | 3,92E+07 | 2,59E+07 | 5 | 4 | 4 | 11 | 11 | Oxidation (M); Carbamidomethylation; Dehydration; Deamidation (NQ); Carbamylation | 21740 | PPTG_00848 | Phytophthora parasitica INRA-310 peptidyl-prolyl cis-trans isomerase 1 (204 aa) |
| 1035 | 907 | PPTG_05032 | 120,89 | 11 | 1 | 10 | 8 | 1,00E+05 | 5,62E+06 | 3,32E+06 | 12 | 12 | 1 | 12 | 8 | Carbamidomethylation | 157848 | PPTG_05032 | Phytophthora parasitica INRA-310 xanthine dehydrogenase (1449 aa) |
| 1231 | 887 | PPTG_03089 | 120,88 | 20 | 4 | 11 | 18 | 4,90E+05 | 2,47E+06 | 6,58E+06 | 10 | 10 | 2 | 6 | 9 | Carbamidomethylation | 62721 | PPTG_03089 | Phytophthora parasitica INRA-310 hypothetical protein (568 aa) |
| 1192 | 1074 | PPTG_08068 | 120,86 | 21 | 5 | 19 | 17 | 7,16E+05 | 6,60E+06 | 1,01E+07 | 9 | 9 | 2 | 8 | 8 | Carbamidomethylation | 52079 | PPTG_08068 | Phytophthora parasitica INRA-310 hypothetical protein (471 aa) |
| 1479 | 1412 | PPTG_12648 | 120,84 | 23 | 11 | 13 | 23 | 1,08E+06 | 1,46E+06 | 2,00E+07 | 6 | 6 | 3 | 3 | 7 | | 36209 | PPTG_12648 | Phytophthora parasitica INRA-310 hypothetical protein (327 aa) |
| 1163 | 854 | PPTG_15104 | 120,82 | 18 | 1 | 7 | 15 | 1,00E+05 | 9,96E+05 | 3,31E+06 | 12 | 12 | 1 | 6 | 11 | Carbamidomethylation | 98174 | PPTG_15104 | Phytophthora parasitica INRA-310 programmed cell death 6-interacting protein (884 aa) |
| 1113 | 1297 | PPTG_00604 | 120,62 | 12 | 2 | 8 | 6 | 2,96E+05 | 2,72E+06 | 2,83E+06 | 11 | 11 | 3 | 8 | 7 | Carbamidomethylation; Dehydration; Deamidation (NQ) | 135142 | PPTG_00604 | Phytophthora parasitica INRA-310 DNA-directed RNA polymerase II subunit RPB2 (1203 aa) |
| 1289 | 1177 | PPTG_15284 | 120,61 | 21 | 0 | 15 | 19 | | 4,09E+06 | 8,42E+06 | 7 | 7 | 0 | 7 | 9 | Carbamidomethylation; Dehydration; Deamidation (NQ) | 43214 | PPTG_15284 | Phytophthora parasitica INRA-310 hypothetical protein (392 aa) |
| 996 | 987 | PPTG_15508 | 120,59 | 12 | 2 | 9 | 11 | 5,68E+05 | 8,08E+06 | 1,55E+07 | 12 | 12 | 2 | 9 | 11 | | 118696 | PPTG_15508 | Phytophthora parasitica INRA-310 AP-3 complex subunit beta (1080 aa) |
| 1029 | 1067 | PPTG_18060 | 120,34 | 34 | 12 | 30 | 29 | 1,18E+06 | 2,56E+07 | 2,13E+07 | 9 | 9 | 3 | 10 | 9 | Carbamidomethylation | 28279 | PPTG_18060 | Phytophthora parasitica INRA-310 phosphomannomutase (248 aa) |
| 922 | 873 | PPTG_05912 | 120,3 | 42 | 8 | 24 | 39 | 1,35E+06 | 1,71E+07 | 1,93E+07 | 12 | 12 | 2 | 8 | 14 | Oxidation (M); Carbamidomethylation; Dehydration | 35355 | PPTG_05912 | Phytophthora parasitica INRA-310 tetrahydrofolate dehydrogenase (318 aa) |
| 1117 | 975 | PPTG_06085 | 120,3 | 14 | 5 | 11 | 10 | 6,28E+04 | 7,15E+05 | 8,78E+05 | 10 | 6 | 4 | 8 | 7 | Carbamidomethylation | 89361 | PPTG_06085 | Phytophthora parasitica INRA-310 pyridoxal phosphate phosphatase (794 aa) |
| 1356 | 1304 | PPTG_03164 | 120,27 | 15 | 6 | 9 | 13 | 2,69E+05 | 3,89E+05 | 6,57E+06 | 8 | 8 | 3 | 4 | 8 | Oxidation (M); Carbamidomethylation | 74718 | PPTG_03164 | Phytophthora parasitica INRA-310 hypothetical protein (675 aa) |
| 1695 | 1594 | PPTG_07067 | 120,24 | 35 | 9 | 35 | 31 | | 2,41E+05 | 4,90E+05 | 5 | 1 | 1 | 6 | 4 | Carbamidomethylation; Deamidation (NQ); Acetylation (N-term) | 31449 | PPTG_07067 | Phytophthora parasitica INRA-310 protein transporter SEC13 (286 aa) |
| 678 | 934 | PPTG_05414 | 120,22 | 31 | 5 | 13 | 28 | 4,65E+04 | 2,29E+06 | 6,37E+06 | 15 | 14 | 2 | 11 | 20 | Oxidation (M); Carbamidomethylation | 58495 | PPTG_05414 | Phytophthora parasitica INRA-310 ATP-dependent RNA helicase DBP2 (544 aa) |
| 1122 | 891 | PPTG_10426 | 120,18 | 23 | 3 | 9 | 21 | 2,62E+05 | 1,90E+06 | 7,14E+06 | 12 | 12 | 2 | 5 | 12 | Acetylation (N-term) | 87730 | PPTG_10426 | Phytophthora parasitica INRA-310 hypothetical protein (784 aa) |
| 929 | 1097 | PPTG_12299 | 120,17 | 17 | 4 | 12 | 15 | 1,24E+06 | 7,28E+06 | 1,39E+07 | 12 | 12 | 3 | 11 | 10 | Oxidation (M); Carbamidomethylation | 101364 | PPTG_12299 | Phytophthora parasitica INRA-310 fimbrin-like protein (932 aa) |
| 1420 | 1178 | PPTG_08876 | 120,17 | 20 | 7 | 17 | 15 | 6,45E+05 | 4,49E+06 | 3,48E+06 | 7 | 7 | 2 | 6 | 6 | Oxidation (M); Carbamidomethylation | 53571 | PPTG_08876 | Phytophthora parasitica INRA-310 succinyl-CoA:3-ketoacid-coenzyme A transferase mitochondrial (500 aa) |
| 1486 | 1169 | PPTG_13677 | 120,14 | 25 | 4 | 20 | 21 | 6,18E+04 | 1,59E+06 | 3,03E+06 | 7 | 7 | 1 | 6 | 6 | Carbamidomethylation | 44105 | PPTG_13677 | Phytophthora parasitica INRA-310 hypothetical protein (393 aa) |
| 1308 | 1165 | PPTG_00486 | 120,12 | 22 | 2 | 18 | 22 | 4,08E+05 | 5,28E+06 | 7,22E+06 | 8 | 6 | 1 | 6 | 9 | Carbamidomethylation | 48551 | PPTG_00486 | Phytophthora parasitica INRA-310 alcohol dehydrogenase (450 aa) |
| 1505 | 1143 | PPTG_06719 | 120,09 | 14 | 1 | 0 | 14 | 4,01E+04 | | 4,01E+06 | 11 | 11 | 1 | 0 | 12 | Carbamidomethylation | 107594 | PPTG_06719 | Phytophthora parasitica INRA-310 hypothetical protein (960 aa) |
| 1176 | 1163 | PPTG_17429 | 120,07 | 17 | 2 | 8 | 13 | 5,71E+05 | 2,34E+06 | 6,04E+06 | 12 | 12 | 2 | 6 | 10 | Oxidation (M); Carbamidomethylation; Acetylation (N-term) | 103000 | PPTG_17429 | Phytophthora parasitica INRA-310 hypothetical protein (937 aa) |
| 1335 | 1569 | PPTG_07053 | 120,06 | 49 | 43 | 25 | 49 | 3,20E+06 | 6,03E+06 | 3,63E+07 | 5 | 5 | 4 | 4 | 8 | Oxidation (M) | 14039 | PPTG_07053 | Phytophthora parasitica INRA-310 CSH (127 aa) |
| 1058 | 967 | PPTG_04280 | 120,02 | 24 | 0 | 22 | 15 | | 5,20E+06 | 4,59E+06 | 13 | 12 | 0 | 12 | 9 | Oxidation (M); Carbamidomethylation | 74586 | PPTG_04280 | Phytophthora parasitica INRA-310 catalase-peroxidase (691 aa) |
| 1445 | 1413 | PPTG_08092 | 120,02 | 21 | 0 | 19 | 19 | | 2,65E+06 | 3,80E+06 | 6 | 6 | 0 | 7 | 7 | Oxidation (M); Carbamidomethylation | 58606 | PPTG_08092 | Phytophthora parasitica INRA-310 transcription factor (542 aa) |
| 1153 | 932 | PPTG_09748 | 119,96 | 9 | 1 | 8 | 8 | 8,91E+04 | 2,41E+06 | 3,55E+06 | 10 | 10 | 1 | 8 | 9 | | 132899 | PPTG_09748 | Phytophthora parasitica INRA-310 hypothetical protein (1195 aa) |
| 1266 | 1049 | PPTG_08041 | 119,96 | 56 | 34 | 43 | 39 | 1,14E+06 | 8,53E+06 | 1,29E+07 | 9 | 8 | 5 | 6 | 6 | Carbamidomethylation | 24347 | PPTG_08041 | Phytophthora parasitica INRA-310 hypothetical protein (219 aa) |
| 1812 | 1954 | PPTG_11595 | 119,93 | 41 | 16 | 34 | 41 | 2,27E+05 | 2,88E+06 | 4,72E+06 | 4 | 4 | 1 | 5 | 4 | | 17159 | PPTG_11595 | Phytophthora parasitica INRA-310 hypothetical protein (165 aa) |
| 1039 | 999 | PPTG_09057 | 119,89 | 22 | 4 | 17 | 17 | 2,50E+05 | 2,90E+06 | 2,71E+06 | 11 | 11 | 2 | 10 | 9 | Oxidation (M); Carbamidomethylation | 62623 | PPTG_09057 | Phytophthora parasitica INRA-310 autophagy-like protein 7 (561 aa) |
| 1415 | 944 | PPTG_18680 | 119,8 | 8 | 0 | 1 | 8 | | 2,11E+05 | 1,15E+07 | 13 | 13 | 0 | 1 | 13 | Deamidation (NQ) | 219481 | PPTG_18680 | Phytophthora parasitica INRA-310 carbohydrate-binding protein (1998 aa) |

| | | | | | | | | | | | | | | | | | | | |
|---|---|---|---|---|---|---|---|---|---|---|---|---|---|---|---|---|---|---|---|
| 1032 | 992 | PPTG_09944 | 119,68 | 10 | 1 | 6 | 9 | 4,63E+05 | 6,30E+06 | 1,43E+07 | 12 | 12 | 1 | 9 | 11 | Carbamidomethylation | 157287 | PPTG_09944 | Phytophthora parasitica INRA-310 eukaryotic translation initiation factor 4 gamma (1425 aa) |
| 1217 | 1303 | PPTG_13996 | 119,68 | 29 | 14 | 25 | 29 | 8,13E+05 | 7,05E+06 | 1,20E+07 | 6 | 6 | 3 | 7 | 7 | Carbamidomethylation | 26946 | PPTG_13996 | Phytophthora parasitica INRA-310 50S ribosomal protein L15 (252 aa) |
| 1148 | 1085 | PPTG_19075 | 119,68 | 39 | 12 | 35 | 29 | 9,81E+04 | 4,19E+06 | 3,06E+06 | 9 | 9 | 3 | 9 | 7 | Carbamidomethylation | 37606 | PPTG_19075 | Phytophthora parasitica INRA-310 twinfilin-like protein (341 aa) |
| 1359 | 1011 | PPTG_05936 | 119,68 | 12 | 1 | 11 | 8 | 1,18E+05 | 3,35E+06 | 3,60E+06 | 9 | 8 | 1 | 8 | 6 | Carbamidomethylation | 101882 | PPTG_05936 | Phytophthora parasitica INRA-310 ATP-dependent protease La (916 aa) |
| 1214 | 1450 | PPTG_09259 | 119,67 | 27 | 20 | 27 | 20 | 5,88E+05 | 8,84E+06 | 5,28E+06 | 6 | 6 | 4 | 7 | 6 | | 33041 | PPTG_09259 | Phytophthora parasitica INRA-310 2 3-bisphosphoglycerate-dependent phosphoglycerate mutase (293 aa) |
| 1200 | 1149 | PPTG_03299 | 119,66 | 43 | 35 | 29 | 32 | 3,11E+06 | 6,52E+06 | 3,36E+07 | 8 | 8 | 6 | 5 | 7 | Oxidation (M); Carbamidomethylation; Deamidation (NQ) | 23137 | PPTG_03299 | Phytophthora parasitica INRA-310 hypothetical protein (209 aa) |
| 1491 | 890 | PPTG_09277 | 119,6 | 12 | 3 | 0 | 12 | 3,36E+05 | | 6,79E+06 | 9 | 9 | 3 | 0 | 10 | Carbamidomethylation; Deamidation (NQ) | 98908 | PPTG_09277 | Phytophthora parasitica INRA-310 hypothetical protein (877 aa) |
| 1210 | 1341 | PPTG_08475 | 119,59 | 17 | 11 | 12 | 16 | 6,31E+05 | 2,89E+06 | 5,79E+06 | 7 | 7 | 5 | 5 | 7 | Dehydration | 51969 | PPTG_08475 | Phytophthora parasitica INRA-310 hypothetical protein (477 aa) |
| 1154 | 1142 | PPTG_11247 | 119,56 | 10 | 6 | 10 | 7 | 1,08E+06 | 5,44E+06 | 9,85E+06 | 8 | 8 | 4 | 8 | 6 | Carbamidomethylation | 108839 | PPTG_11247 | Phytophthora parasitica INRA-310 eukaryotic translation initiation factor 5B (974 aa) |
| 1466 | 1159 | PPTG_19151 | 119,51 | 14 | 3 | 12 | 9 | 1,41E+05 | 2,00E+06 | 2,20E+06 | 7 | 7 | 2 | 6 | 6 | Carbamidomethylation | 82734 | PPTG_19151 | Phytophthora parasitica INRA-310 translation initiation factor eIF-2B subunit epsilon (742 aa) |
| 1262 | 1631 | PPTG_15073 | 119,51 | 22 | 13 | 22 | 13 | 1,06E+07 | 5,26E+07 | 1,17E+08 | 6 | 6 | 4 | 7 | 6 | Carbamidomethylation | 38781 | PPTG_15073 | Phytophthora parasitica INRA-310 sporangia induced protein (379 aa) |
| 1355 | 1267 | PPTG_09418 | 119,49 | 6 | 1 | 1 | 6 | | | 4,84E+05 | 7 | 2 | 2 | 2 | 11 | Oxidation (M); Carbamidomethylation | 151262 | PPTG_09418 | Phytophthora parasitica INRA-310 ATP-binding Cassette (ABC) superfamily (1364 aa) |
| 1355 | 1276 | PPTG_09420 | 119,49 | 8 | 1 | 1 | 8 | | | 4,84E+05 | 7 | 2 | 2 | 2 | 11 | Oxidation (M); Carbamidomethylation | 122580 | PPTG_09420 | Phytophthora parasitica INRA-310 ATP-binding Cassette (ABC) superfamily (1102 aa) |
| 78 | 2147 | PPTG_04251 | 119,48 | 36 | 28 | 32 | 32 | 1,07E+07 | 2,03E+08 | 5,83E+08 | 5 | 5 | 11 | 50 | 60 | Oxidation (M); Dehydration; Deamidation (NQ); Acetylation (N-term); Amidation; Carbamylation | 17700 | PPTG_04251 | Phytophthora parasitica INRA-310 40S ribosomal protein S15 (156 aa) |
| 1332 | 1407 | PPTG_13875 | 119,36 | 42 | 26 | 42 | 42 | 1,42E+06 | 1,30E+07 | 1,89E+07 | 6 | 6 | 3 | 6 | 7 | | 19531 | PPTG_13875 | Phytophthora parasitica INRA-310 hypothetical protein (179 aa) |
| 1370 | 1334 | PPTG_08965 | 119,22 | 19 | 8 | 17 | 19 | 1,11E+05 | 1,95E+06 | 3,34E+06 | 6 | 6 | 2 | 6 | 7 | Carbamidomethylation | 46292 | PPTG_08965 | Phytophthora parasitica INRA-310 hypothetical protein (434 aa) |
| 949 | 1125 | PPTG_14832 | 119,13 | 22 | 11 | 11 | 22 | 7,02E+05 | 1,69E+06 | 1,12E+07 | 8 | 8 | 5 | 6 | 12 | Carbamidomethylation; Dehydration; Deamidation (NQ) | 46333 | PPTG_14832 | Phytophthora parasitica INRA-310 lectin (416 aa) |
| 1455 | 1318 | PPTG_05877 | 119,12 | 18 | 0 | 18 | 12 | | 3,69E+06 | 2,25E+06 | 7 | 6 | 0 | 9 | 5 | | 86734 | PPTG_05877 | Phytophthora parasitica INRA-310 glycoside hydrolase (789 aa) |
| 1459 | 1283 | PPTG_04788 | 119,05 | 21 | 2 | 21 | 21 | 1,44E+05 | 6,20E+06 | 4,99E+06 | 6 | 6 | 1 | 7 | 6 | Carbamidomethylation | 50650 | PPTG_04788 | Phytophthora parasitica INRA-310 arginine biosynthesis ArgJ mitochondrial (466 aa) |
| 1358 | 1348 | PPTG_00515 | 119,03 | 7 | 3 | 4 | 7 | 2,19E+05 | 4,16E+05 | 7,34E+06 | 7 | 7 | 2 | 3 | 10 | Oxidation (M) | 119172 | PPTG_00515 | Phytophthora parasitica INRA-310 hypothetical protein (1045 aa) |
| 1285 | 960 | PPTG_13113 | 118,92 | 11 | 2 | 9 | 8 | 3,07E+05 | 2,64E+06 | 3,53E+06 | 10 | 10 | 2 | 7 | 7 | Carbamidomethylation; Dehydration; Amidation; Carbamylation | 121995 | PPTG_13113 | Phytophthora parasitica INRA-310 DNA polymerase delta catalytic subunit (1090 aa) |
| 1089 | 924 | PPTG_01231 | 118,88 | 28 | 10 | 24 | 22 | 7,20E+05 | 3,62E+06 | 5,88E+06 | 9 | 9 | 3 | 9 | 8 | Oxidation (M); Carbamidomethylation | 44166 | PPTG_01231 | Phytophthora parasitica INRA-310 AMP-binding enzyme (407 aa) |
| 1668 | 1230 | PPTG_14629 | 118,73 | 27 | 0 | 0 | 27 | | | 1,53E+07 | 7 | 7 | 0 | 0 | 11 | Carbamidomethylation; Deamidation (NQ) | 44535 | PPTG_14629 | Phytophthora parasitica INRA-310 hypothetical protein (395 aa) |
| 1977 | 1292 | PPTG_00742 | 118,69 | 12 | 0 | 4 | 12 | | 4,93E+05 | 3,30E+06 | 6 | 6 | 0 | 2 | 6 | Carbamidomethylation | 81486 | PPTG_00742 | Phytophthora parasitica INRA-310 hypothetical protein (797 aa) |
| 1185 | 885 | PPTG_08947 | 118,62 | 15 | 3 | 6 | 15 | 6,99E+04 | 1,08E+06 | 6,42E+06 | 11 | 11 | 2 | 5 | 11 | | 107960 | PPTG_08947 | Phytophthora parasitica INRA-310 hypothetical protein (956 aa) |
| 1179 | 1107 | PPTG_01013 | 118,62 | 25 | 2 | 21 | 19 | 1,37E+05 | 6,76E+06 | 2,13E+06 | 9 | 9 | 1 | 11 | 6 | Carbamidomethylation; Deamidation (NQ) | 49695 | PPTG_01013 | Phytophthora parasitica INRA-310 homogentisate 1 2-dioxygenase (441 aa) |
| 1147 | 995 | PPTG_01711 | 118,6 | 32 | 14 | 25 | 28 | 1,69E+06 | 1,34E+07 | 1,70E+07 | 9 | 9 | 4 | 7 | 8 | Carbamidomethylation | 32148 | PPTG_01711 | Phytophthora parasitica INRA-310 ATP phosphoribosyltransferase (300 aa) |
| 1094 | 1070 | PPTG_14623 | 118,45 | 12 | 1 | 11 | 12 | 2,32E+05 | 3,18E+06 | 3,59E+07 | 8 | 8 | 1 | 8 | 11 | Oxidation (M); Carbamidomethylation | 89230 | PPTG_14623 | Phytophthora parasitica INRA-310 glycoside hydrolase (814 aa) |
| 1237 | 1063 | PPTG_03736 | 118,41 | 24 | 11 | 14 | 20 | 1,66E+06 | 4,38E+06 | 2,22E+07 | 8 | 8 | 4 | 5 | 8 | | 42315 | PPTG_03736 | Phytophthora parasitica INRA-310 hypothetical protein (385 aa) |
| 962 | 1480 | PPTG_03562 | 118,39 | 30 | 14 | 23 | 23 | 5,89E+05 | 4,92E+06 | 2,73E+07 | 8 | 8 | 3 | 9 | 11 | Oxidation (M); Carbamidomethylation; Deamidation (NQ) | 27325 | PPTG_03562 | Phytophthora parasitica INRA-310 polysaccharide lyase (262 aa) |
| 1091 | 1025 | PPTG_14270 | 118,38 | 20 | 3 | 18 | 14 | 3,22E+05 | 6,90E+06 | 7,87E+06 | 9 | 9 | 2 | 10 | 8 | Oxidation (M); Carbamidomethylation; Deamidation (NQ) | 60769 | PPTG_14270 | Phytophthora parasitica INRA-310 hypothetical protein (547 aa) |
| 1091 | 1026 | PPTG_14270 | 118,38 | 20 | 3 | 18 | 14 | 3,22E+05 | 6,90E+06 | 7,87E+06 | 9 | 9 | 2 | 10 | 8 | Oxidation (M); Carbamidomethylation; Deamidation (NQ) | 60769 | PPTG_14270 | Phytophthora parasitica INRA-310 hypothetical protein (547 aa) |
| 1091 | 1027 | PPTG_14270T2 | 118,38 | 20 | 3 | 18 | 14 | 3,22E+05 | 6,90E+06 | 7,87E+06 | 9 | 9 | 2 | 10 | 8 | Oxidation (M); Carbamidomethylation; Deamidation (NQ) | 60769 | PPTG_14270 | Phytophthora parasitica INRA-310 hypothetical protein (547 aa) |
| 711 | 1429 | PPTG_07051 | 118,37 | 69 | 35 | 58 | 69 | 1,39E+06 | 2,78E+07 | 1,40E+08 | 7 | 7 | 3 | 10 | 19 | Oxidation (M); Dehydration; Deamidation (NQ); Acetylation (N-term); Amidation | 12060 | PPTG_07051 | Phytophthora parasitica INRA-310 cytochrome c (113 aa) |
| 1061 | 922 | PPTG_14386 | 118,15 | 16 | 9 | 14 | 13 | 6,18E+05 | 6,26E+06 | 7,63E+06 | 9 | 9 | 5 | 8 | 8 | Oxidation (M); Carbamidomethylation | 84438 | PPTG_14386 | Phytophthora parasitica INRA-310 beta-D-xylosidase (773 aa) |
| 1140 | 893 | PPTG_12471 | 118,07 | 19 | 5 | 6 | 19 | 2,01E+05 | 6,23E+05 | 5,83E+06 | 11 | 9 | 4 | 4 | 11 | Carbamidomethylation | 83646 | PPTG_12471 | Phytophthora parasitica INRA-310 beta-glucan synthesis-associated protein (755 aa) |
| 1344 | 1708 | PPTG_17152 | 118,04 | 55 | 27 | 55 | 49 | 1,92E+06 | 1,65E+07 | 1,65E+07 | 5 | 5 | 3 | 7 | 6 | Oxidation (M); Carbamidomethylation | 13832 | PPTG_17152 | Phytophthora parasitica INRA-310 ribonucleoprotein-associated protein (129 aa) |
| 1575 | 1080 | PPTG_06171 | 118,03 | 9 | 1 | 2 | 9 | 1,87E+05 | 1,23E+06 | 9,07E+06 | 8 | 8 | 1 | 2 | 9 | Oxidation (M); Carbamidomethylation | 119708 | PPTG_06171 | Phytophthora parasitica INRA-310 protein kinase (1116 aa) |
| 1426 | 1299 | PPTG_00308 | 118,02 | 19 | 2 | 12 | 17 | | 7,79E+05 | 1,82E+06 | 8 | 3 | 1 | 6 | 7 | Oxidation (M); Carbamidomethylation | 48634 | PPTG_00308 | Phytophthora parasitica INRA-310 hypothetical protein (444 aa) |
| 1247 | 1437 | PPTG_14391 | 118,01 | 10 | 2 | 10 | 10 | 9,33E+04 | 2,83E+06 | 1,93E+06 | 7 | 4 | 1 | 9 | 7 | Oxidation (M) | 84400 | PPTG_14391 | Phytophthora parasitica INRA-310 glycoside hydrolase (775 aa) |
| 1487 | 1465 | PPTG_14802 | 117,98 | 32 | 10 | 21 | 32 | 1,14E+06 | 6,06E+06 | 9,09E+06 | 6 | 6 | 2 | 5 | 6 | Oxidation (M); Deamidation (NQ) | 26009 | PPTG_14802 | Phytophthora parasitica INRA-310 acetolactate synthase small subunit (242 aa) |
| 1239 | 1114 | PPTG_04184 | 117,98 | 10 | 1 | 8 | 10 | 1,45E+05 | 2,14E+06 | 4,97E+06 | 9 | 9 | 1 | 7 | 9 | Carbamidomethylation | 127800 | PPTG_04184 | Phytophthora parasitica INRA-310 hypothetical protein (1185 aa) |
| 1050 | 1089 | PPTG_09348 | 117,98 | 26 | 11 | 14 | 25 | 1,23E+06 | 2,39E+06 | 1,98E+07 | 12 | 12 | 4 | 6 | 11 | Carbamidomethylation; Carbamylation | 47941 | PPTG_09348 | Phytophthora parasitica INRA-310 hypothetical protein (424 aa) |
| 1180 | 2154 | PPTG_10374 | 117,87 | 53 | 47 | 47 | 46 | 6,67E+06 | 1,11E+07 | 5,13E+07 | 4 | 4 | 6 | 6 | 6 | Oxidation (M); Deamidation (NQ) | 11712 | PPTG_10374 | Phytophthora parasitica INRA-310 hypothetical protein (104 aa) |
| 1736 | 1738 | PPTG_19740 | 117,84 | 20 | 7 | 17 | 20 | 1,19E+06 | 6,14E+06 | 6,59E+06 | 5 | 5 | 2 | 4 | 5 | Carbamidomethylation | 39521 | PPTG_19740 | Phytophthora parasitica INRA-310 lipase (362 aa) |
| 1142 | 930 | PPTG_17649 | 117,75 | 12 | 1 | 6 | 12 | 1,11E+05 | 1,43E+06 | 6,92E+06 | 12 | 12 | 1 | 6 | 12 | Carbamidomethylation | 138799 | PPTG_17649 | Phytophthora parasitica INRA-310 hypothetical protein (1290 aa) |
| 903 | 1139 | PPTG_11061 | 117,68 | 31 | 11 | 31 | 31 | 3,67E+05 | 3,53E+06 | 6,14E+06 | 8 | 7 | 4 | 9 | 12 | Oxidation (M); Carbamidomethylation; Dehydration; Deamidation (NQ) | 35756 | PPTG_11061 | Phytophthora parasitica INRA-310 serine/threonine-protein phosphatase 2A catalytic subunit alpha isoform (313 aa) |
| 1031 | 1104 | PPTG_07092 | 117,6 | 17 | 5 | 17 | 17 | 5,75E+05 | 7,12E+06 | 1,04E+07 | 9 | 9 | 2 | 10 | 10 | Deamidation (NQ) | 60991 | PPTG_07092 | Phytophthora parasitica INRA-310 importin subunit alpha (560 aa) |
| 1114 | 1214 | PPTG_16032 | 117,5 | 7 | 1 | 0 | 6 | 1,69E+05 | 9,82E+04 | 4,25E+06 | 18 | 18 | 2 | 1 | 16 | Carbamidomethylation | 299319 | PPTG_16032 | Phytophthora parasitica INRA-310 callose synthase (2648 aa) |
| 841 | 1956 | PPTG_04970 | 117,4 | 51 | 44 | 30 | 51 | 4,39E+06 | 5,82E+07 | 9,43E+07 | 5 | 4 | 4 | 11 | 12 | Oxidation (M); Carbamidomethylation; Deamidation (NQ); Carbamylation | 14839 | PPTG_04970 | Phytophthora parasitica INRA-310 histone H2A type 1 (141 aa) |
| 1523 | 1171 | PPTG_15078 | 117,3 | 30 | 3 | 10 | 30 | 1,05E+05 | 5,78E+05 | 7,89E+06 | 8 | 8 | 1 | 2 | 10 | Oxidation (M) | 36102 | PPTG_15078 | Phytophthora parasitica INRA-310 estradiol 17-beta-dehydrogenase (329 aa) |
| 1347 | 2079 | PPTG_04665 | 117,23 | 67 | 45 | 67 | 67 | 5,90E+05 | 9,37E+06 | 1,55E+07 | 4 | 4 | 3 | 6 | 7 | | 11018 | PPTG_04665 | Phytophthora parasitica INRA-310 glutaredoxin (105 aa) |
| 1554 | 1173 | PPTG_05471 | 117,16 | 6 | 2 | 2 | 6 | 1,00E+05 | 4,51E+05 | 3,15E+06 | 7 | 7 | 2 | 3 | 7 | Carbamidomethylation | 172453 | PPTG_05471 | Phytophthora parasitica INRA-310 brefeldin A-inhibited guanine nucleotide-exchange protein (1551 aa) |
| 1194 | 1644 | PPTG_16150 | 117,13 | 53 | 53 | 53 | 47 | 1,28E+06 | 1,10E+07 | 1,06E+07 | 6 | 6 | 5 | 7 | 6 | Carbamidomethylation | 13655 | PPTG_16150 | Phytophthora parasitica INRA-310 hypothetical protein (121 aa) |
| 864 | 1162 | PPTG_09516 | 117,04 | 61 | 21 | 57 | 61 | 7,83E+06 | 3,95E+07 | 7,64E+07 | 9 | 9 | 5 | 10 | 12 | Carbamidomethylation; Acetylation (N-term) | 24263 | PPTG_09516 | Phytophthora parasitica INRA-310 GTP-binding nuclear protein Ran (215 aa) |
| 1469 | 1195 | PPTG_00774 | 117,04 | 31 | 14 | 12 | 31 | 4,11E+05 | 1,72E+06 | 5,41E+06 | 7 | 7 | 3 | 3 | 8 | | 44529 | PPTG_00774 | Phytophthora parasitica INRA-310 actin-like protein (404 aa) |
| 1706 | 1273 | PPTG_00206 | 117,03 | 45 | 6 | 6 | 45 | 1,33E+06 | 2,60E+06 | 2,82E+07 | 7 | 7 | 1 | 1 | 9 | Oxidation (M); Carbamidomethylation | 19509 | PPTG_00206 | Phytophthora parasitica INRA-310 hypothetical protein (182 aa) |
| 1591 | 1243 | PPTG_18654 | 117 | 22 | 5 | 12 | 17 | 6,17E+05 | 1,59E+06 | 4,46E+06 | 7 | 7 | 2 | 4 | 6 | Oxidation (M) | 46329 | PPTG_18654 | Phytophthora parasitica INRA-310 protein disulfide-isomerase (423 aa) |
| 1781 | 1807 | PPTG_02889 | 116,85 | 21 | 9 | 19 | 14 | 4,97E+05 | 7,27E+06 | 7,40E+06 | 5 | 5 | 2 | 4 | 4 | | 52462 | PPTG_02889 | Phytophthora parasitica INRA-310 hypothetical protein (500 aa) |
| 1595 | 1964 | PPTG_11245 | 116,72 | 48 | 16 | 48 | 48 | 6,12E+05 | 1,13E+07 | 1,43E+07 | 4 | 4 | 2 | 5 | 5 | | 13480 | PPTG_11245 | Phytophthora parasitica INRA-310 hypothetical protein (126 aa) |
| 1046 | 1033 | PPTG_01321 | 116,69 | 43 | 3 | 34 | 36 | 1,14E+05 | 1,16E+07 | 5,72E+06 | 13 | 13 | 1 | 10 | 10 | Carbamidomethylation; Deamidation (NQ); Acetylation (N-term) | 41299 | PPTG_01321 | Phytophthora parasitica INRA-310 hypothetical protein (354 aa) |
| 1190 | 1088 | PPTG_19559 | 116,67 | 11 | 5 | 7 | 11 | 3,93E+05 | 2,17E+06 | 6,15E+06 | 9 | 9 | 3 | 5 | 10 | Oxidation (M) | 84831 | PPTG_19559 | Phytophthora parasitica INRA-310 myosin-like protein (775 aa) |
| 1483 | 1269 | PPTG_00569 | 116,66 | 19 | 8 | 6 | 19 | 7,80E+04 | 2,56E+05 | 3,18E+06 | 7 | 7 | 3 | 2 | 8 | Carbamidomethylation | 54318 | PPTG_00569 | Phytophthora parasitica INRA-310 hypothetical protein (487 aa) |
| 1694 | 1564 | PPTG_03020 | 116,55 | 38 | 10 | 38 | 32 | 7,51E+05 | 4,97E+06 | 3,04E+06 | 5 | 5 | 2 | 5 | 4 | Oxidation (M) | 20993 | PPTG_03020 | Phytophthora parasitica INRA-310 NAD(P)H:quinone oxidoreductase type IV (201 aa) |
| 1694 | 1565 | PPTG_03020 | 116,55 | 38 | 10 | 38 | 32 | 7,51E+05 | 4,97E+06 | 3,04E+06 | 5 | 5 | 2 | 5 | 4 | Oxidation (M) | 20993 | PPTG_03020 | Phytophthora parasitica INRA-310 NAD(P)H:quinone oxidoreductase type IV (201 aa) |
| 1376 | 1069 | PPTG_00045 | 116,53 | 21 | 6 | 6 | 21 | 5,65E+05 | 8,44E+05 | 8,85E+06 | 8 | 8 | 3 | 3 | 9 | Oxidation (M); Carbamidomethylation; Dehydration; Deamidation (NQ) | 55569 | PPTG_00045 | Phytophthora parasitica INRA-310 hypothetical protein (520 aa) |
| 1440 | 943 | PPTG_02518 | 116,51 | 27 | 7 | 2 | 27 | 8,35E+04 | 1,10E+05 | 4,49E+06 | 10 | 8 | 3 | 1 | 10 | Carbamidomethylation | 49333 | PPTG_02518 | Phytophthora parasitica INRA-310 CorA Metal Ion transporter (MIT) family (442 aa) |
| 1580 | 1354 | PPTG_03859 | 116,5 | 48 | 0 | 32 | 35 | | 6,39E+06 | 8,48E+06 | 6 | 6 | 0 | 5 | 7 | Oxidation (M); Deamidation (NQ) | 26402 | PPTG_03859 | Phytophthora parasitica INRA-310 6-phosphogluconolactonase (246 aa) |
| 1508 | 1553 | PPTG_06866 | 116,45 | 17 | 4 | 17 | 13 | | 8,26E+05 | 2,41E+05 | 5 | 3 | 1 | 7 | 5 | | 57685 | PPTG_06866 | Phytophthora parasitica INRA-310 catalase (516 aa) |
| 464 | 2277 | PPTG_01694 | 116,41 | 19 | 14 | 19 | 19 | 3,50E+07 | 1,49E+08 | 4,56E+08 | 4 | 4 | 7 | 14 | 24 | Oxidation (M); Dehydration; Deamidation (NQ); Acetylation (N-term); Carbamylation | 21881 | PPTG_01694 | Phytophthora parasitica INRA-310 50S ribosomal protein L19e (186 aa) |
| 610 | 1462 | PPTG_01865 | 116,38 | 5 | 3 | 5 | 5 | 5,09E+06 | 4,95E+07 | 5,62E+07 | 6 | 6 | 6 | 17 | 13 | Oxidation (M); Carbamidomethylation; Dehydration; Deamidation (NQ) | 152968 | PPTG_01865 | Phytophthora parasitica INRA-310 mucin-like protein (1494 aa) |
| 1360 | 1244 | PPTG_13874 | 116,36 | 19 | 2 | 17 | 9 | 1,12E+05 | 2,55E+06 | 3,01E+06 | 9 | 9 | 1 | 9 | 5 | Acetylation (N-term) | 71903 | PPTG_13874 | Phytophthora parasitica INRA-310 signal recognition particle (656 aa) |
| 1260 | 1057 | PPTG_02812 | 116,26 | 13 | 0 | 11 | 12 | | 2,69E+06 | 2,91E+06 | 9 | 9 | 0 | 9 | 8 | | 101560 | PPTG_02812 | Phytophthora parasitica INRA-310 hypothetical protein (908 aa) |
| 1381 | 1198 | PPTG_04177 | 116,25 | 31 | 8 | 15 | 29 | 7,07E+05 | 2,61E+06 | 9,51E+06 | 8 | 8 | 3 | 5 | 7 | Oxidation (M); Carbamidomethylation | 42101 | PPTG_04177 | Phytophthora parasitica INRA-310 protein kinase (378 aa) |
| 1203 | 1145 | PPTG_08508 | 116,22 | 15 | 7 | 10 | 15 | 9,65E+05 | 4,93E+06 | 9,78E+06 | 8 | 8 | 3 | 5 | 10 | Oxidation (M); Carbamidomethylation | 75958 | PPTG_08508 | Phytophthora parasitica INRA-310 glucosylceramidase (674 aa) |
| 877 | 939 | PPTG_02167 | 116,21 | 21 | 5 | 17 | 19 | 5,78E+05 | 5,25E+06 | 9,06E+06 | 12 | 12 | 4 | 10 | 12 | Deamidation (NQ) | 75825 | PPTG_02167 | Phytophthora parasitica INRA-310 nucleolar GTP-binding protein 1 (665 aa) |
| 1760 | 1526 | PPTG_07734 | 116,19 | 11 | 5 | 3 | 10 | 1,78E+05 | 2,07E+05 | 4,90E+06 | 6 | 5 | 2 | 2 | 6 | Oxidation (M) | 66469 | PPTG_07734 | Phytophthora parasitica INRA-310 protein kinase (626 aa) |
| 1157 | 1141 | PPTG_15854 | 116,18 | 30 | 2 | 23 | 24 | 1,67E+05 | 1,85E+07 | 9,46E+06 | 8 | 8 | 1 | 9 | 8 | Oxidation (M) | 35336 | PPTG_15854 | Phytophthora parasitica INRA-310 FKBP-type peptidyl-prolyl cis-trans isomerase (327 aa) |
| 1369 | 1184 | PPTG_13548 | 116,13 | 24 | 0 | 9 | 24 | | 5,81E+05 | 1,52E+07 | 11 | 11 | 0 | 4 | 11 | | 51339 | PPTG_13548 | Phytophthora parasitica INRA-310 hypothetical protein (459 aa) |
| 1295 | 1016 | PPTG_01312 | 116,07 | 9 | 4 | 2 | 9 | 3,02E+05 | 3,12E+05 | 2,20E+06 | 10 | 6 | 4 | 3 | 9 | | 147796 | PPTG_01312 | Phytophthora parasitica INRA-310 ATP-binding Cassette (ABC) superfamily (1339 aa) |
| 1152 | 1206 | PPTG_01305 | 116,01 | 17 | 5 | 13 | 15 | 4,89E+05 | 3,40E+06 | 1,09E+07 | 9 | 9 | 2 | 6 | 10 | | 57616 | PPTG_01305 | Phytophthora parasitica INRA-310 hypothetical protein (515 aa) |
| 1468 | 1643 | PPTG_01148 | 116 | 23 | 4 | 23 | 23 | 4,45E+05 | 1,48E+07 | 1,36E+07 | 5 | 5 | 1 | 7 | 6 | Oxidation (M); Carbamidomethylation | 31240 | PPTG_01148 | Phytophthora parasitica INRA-310 hypothetical protein (296 aa) |
| 2155 | 1140 | PPTG_16931 | 115,92 | 13 | 0 | 0 | 13 | | | 2,02E+06 | 7 | 7 | 0 | 0 | 7 | Carbamidomethylation | 81807 | PPTG_16931 | Phytophthora parasitica INRA-310 hypothetical protein (720 aa) |
| 1310 | 1288 | PPTG_15232 | 115,89 | 27 | 15 | 27 | 27 | 5,29E+05 | 6,51E+06 | 5,28E+06 | 6 | 6 | 4 | 6 | 6 | | 36146 | PPTG_15232 | Phytophthora parasitica INRA-310 aldose 1-epimerase (331 aa) |
| 1478 | 2353 | PPTG_11267 | 115,84 | 18 | 11 | 18 | 18 | 4,55E+05 | 4,79E+07 | 3,75E+07 | 3 | 3 | 3 | 5 | 6 | Oxidation (M); Deamidation (NQ); Acetylation (N-term) | 31045 | PPTG_11267 | Phytophthora parasitica INRA-310 hypothetical protein (293 aa) |
| 1416 | 959 | PPTG_18987 | 115,82 | 11 | 1 | 7 | 5 | 9,10E+03 | 9,46E+05 | 1,21E+06 | 11 | 11 | 1 | 7 | 6 | Oxidation (M); Carbamidomethylation | 144708 | PPTG_18987 | Phytophthora parasitica INRA-310 tubulin-specific chaperone D (1297 aa) |
| 2047 | 1779 | PPTG_00440 | 115,63 | 32 | 0 | 27 | 11 | | 3,16E+06 | 1,61E+06 | 5 | 5 | 0 | 6 | 2 | Carbamidomethylation | 25748 | PPTG_00440 | Phytophthora parasitica INRA-310 hypothetical protein (245 aa) |
| 1744 | 1342 | PPTG_03835 | 115,63 | 5 | 1 | 3 | 4 | | | 3,58E+05 | 6 | 1 | 2 | 3 | 5 | | 139280 | PPTG_03835 | Phytophthora parasitica INRA-310 ATP-binding Cassette (ABC) superfamily (1286 aa) |
| 1264 | 1003 | PPTG_15925 | 115,61 | 25 | 4 | 16 | 24 | 2,81E+05 | 3,47E+06 | 5,89E+06 | 9 | 9 | 2 | 6 | 9 | Oxidation (M) | 65722 | PPTG_15925 | Phytophthora parasitica INRA-310 choline dehydrogenase (588 aa) |
| 1107 | 1255 | PPTG_18935 | 115,57 | 22 | 13 | 22 | 18 | 1,23E+06 | 4,82E+06 | 1,65E+07 | 7 | 6 | 4 | 9 | 7 | Carbamidomethylation; Acetylation (N-term) | 43698 | PPTG_18935 | Phytophthora parasitica INRA-310 glucokinase (399 aa) |
| 1540 | 1366 | PPTG_07606 | 115,47 | 31 | 14 | 9 | 31 | 1,43E+06 | 1,91E+06 | 1,58E+07 | 6 | 6 | 3 | 2 | 8 | | 36672 | PPTG_07606 | Phytophthora parasitica INRA-310 transmembrane protein (321 aa) |
| 1362 | 1012 | PPTG_04973 | 115,43 | 14 | 1 | 12 | 12 | 7,90E+04 | 3,06E+06 | 3,06E+06 | 8 | 8 | 1 | 7 | 7 | Carbamidomethylation | 82058 | PPTG_04973 | Phytophthora parasitica INRA-310 dynein intermediate chain 2 (751 aa) |
| 1573 | 1704 | PPTG_09002 | 115,26 | 25 | 8 | 10 | 25 | 3,93E+05 | 5,89E+05 | 6,66E+06 | 6 | 6 | 2 | 3 | 7 | Oxidation (M); Carbamidomethylation | 33216 | PPTG_09002 | Phytophthora parasitica INRA-310 hypothetical protein (298 aa) |
| 1225 | 1247 | PPTG_12231 | 115,24 | 22 | 2 | 22 | 20 | 1,12E+05 | 4,39E+06 | 3,90E+06 | 7 | 7 | 1 | 9 | 7 | Oxidation (M) | 46917 | PPTG_12231 | Phytophthora parasitica INRA-310 diaminopimelate decarboxylase (423 aa) |
| 1651 | 1241 | PPTG_05598 | 115,23 | 24 | 7 | 3 | 24 | 8,08E+05 | 6,59E+05 | 1,13E+07 | 7 | 7 | 2 | 1 | 8 | Oxidation (M) | 38542 | PPTG_05598 | Phytophthora parasitica INRA-310 phosphatidylinositol-3 4 5-trisphosphate 3-phosphatase (336 aa) |
| 1078 | 1155 | PPTG_04169 | 115,16 | 9 | 1 | 5 | 9 | 2,91E+05 | 1,47E+06 | 5,97E+06 | 12 | 11 | 2 | 7 | 11 | Carbamidomethylation | 155876 | PPTG_04169 | Phytophthora parasitica INRA-310 phosphatidylinositol kinase (1374 aa) |
| 1130 | 1054 | PPTG_18422 | 115,13 | 15 | 8 | 13 | 13 | 7,68E+05 | 2,54E+06 | 7,98E+06 | 8 | 8 | 4 | 7 | 8 | Carbamidomethylation | 77526 | PPTG_18422 | Phytophthora parasitica INRA-310 hsp70-like protein (703 aa) |
| 1707 | 1380 | PPTG_11457 | 115,13 | 27 | 0 | 21 | 21 | | 3,31E+06 | 2,21E+06 | 6 | 6 | 0 | 6 | 5 | | 45728 | PPTG_11457 | Phytophthora parasitica INRA-310 hypothetical protein (411 aa) |
| 1513 | 1662 | PPTG_01546 | 115,11 | 18 | 7 | 12 | 18 | 9,13E+05 | 1,62E+06 | 1,90E+07 | 5 | 5 | 3 | 4 | 6 | Oxidation (M) | 43383 | PPTG_01546 | Phytophthora parasitica INRA-310 hypothetical protein (387 aa) |
| 1098 | 1233 | PPTG_15717 | 115,11 | 60 | 20 | 46 | 60 | 5,77E+06 | 1,36E+07 | 9,62E+07 | 7 | 7 | 3 | 6 | 11 | Oxidation (M); Carbamidomethylation | 18550 | PPTG_15717 | Phytophthora parasitica INRA-310 hypothetical protein (163 aa) |
| 1730 | 1640 | PPTG_04091 | 115,09 | 24 | 10 | 24 | 21 | 1,13E+05 | 2,30E+06 | 1,52E+06 | 5 | 5 | 2 | 5 | 4 | | 35978 | PPTG_04091 | Phytophthora parasitica INRA-310 gamma-glutamyl hydrolase (318 aa) |
| 1184 | 1101 | PPTG_14741 | 115,02 | 9 | 1 | 6 | 7 | 2,41E+05 | 3,17E+06 | 6,06E+06 | 11 | 11 | 2 | 8 | 8 | Carbamidomethylation | 154772 | PPTG_14741 | Phytophthora parasitica INRA-310 hypothetical protein (1399 aa) |
| 1769 | 1078 | PPTG_18531 | 114,97 | 21 | 0 | 4 | 21 | | 3,18E+05 | 4,76E+06 | 8 | 8 | 0 | 2 | 8 | Carbamidomethylation; Acetylation (N-term) | 56618 | PPTG_18531 | Phytophthora parasitica INRA-310 hypothetical protein (502 aa) |
| 1352 | 997 | PPTG_20156 | 114,92 | 38 | 0 | 8 | 38 | | 1,81E+05 | 7,25E+06 | 8 | 5 | 0 | 3 | 12 | Oxidation (M); Dehydration | 36971 | PPTG_20156 | Phytophthora parasitica INRA-310 syntaxin-binding protein (342 aa) |
| 1858 | 1626 | PPTG_02905 | 114,89 | 28 | 0 | 3 | 28 | | 1,59E+05 | 5,60E+06 | 6 | 6 | 0 | 1 | 8 | Oxidation (M); Carbamidomethylation | 35394 | PPTG_02905 | Phytophthora parasitica INRA-310 syntaxin (317 aa) |
| 1880 | 1606 | PPTG_11081 | 114,88 | 17 | 7 | 6 | 17 | 2,77E+05 | 1,60E+05 | 5,13E+06 | 5 | 5 | 2 | 2 | 5 | | 46061 | PPTG_11081 | Phytophthora parasitica INRA-310 hypothetical protein (417 aa) |

| | | | | | | | | | | | | | | | | | | | |
|---|---|---|---|---|---|---|---|---|---|---|---|---|---|---|---|---|---|---|---|
| 1708 | 1432 | PPTG_02580 | 114,87 | 16 | 2 | 6 | 16 | 1,53E+05 | 9,32E+05 | 5,10E+06 | 6 | 6 | 1 | 3 | 7 | Carbamidomethylation; Deamidation (NQ) | 50361 | PPTG_02580 | Phytophthora parasitica INRA-310 transmembrane protein (445 aa) |
| 777 | 2030 | PPTG_00630 | 114,82 | 31 | 30 | 30 | 25 | 1,41E+07 | 8,53E+07 | 3,18E+08 | 5 | 5 | 6 | 9 | 14 | Oxidation (M); Carbamidomethylation; Deamidation (NQ); Carbamylation | 14056 | PPTG_00630 | Phytophthora parasitica INRA-310 60S ribosomal protein L31 (121 aa) |
| 1144 | 1325 | PPTG_13102 | 114,54 | 8 | 5 | 7 | 8 | 7,15E+05 | 5,52E+06 | 7,70E+06 | 7 | 7 | 4 | 7 | 8 | Carbamidomethylation | 101302 | PPTG_13102 | Phytophthora parasitica INRA-310 transportin-like protein (904 aa) |
| 816 | 1381 | PPTG_06127 | 114,49 | 38 | 22 | 22 | 38 | 2,01E+06 | 7,78E+06 | 4,11E+07 | 6 | 3 | 6 | 8 | 14 | Oxidation (M); Carbamidomethylation; Deamidation (NQ) | 22321 | PPTG_06127 | Phytophthora parasitica INRA-310 NADH ubiquinone oxidoreductase (206 aa) |
| 816 | 1382 | PPTG_18759 | 114,49 | 38 | 22 | 22 | 38 | 2,01E+06 | 7,78E+06 | 4,11E+07 | 6 | 3 | 6 | 8 | 14 | Oxidation (M); Carbamidomethylation; Deamidation (NQ) | 22321 | PPTG_18759 | Phytophthora parasitica INRA-310 NADH ubiquinone oxidoreductase (206 aa) |
| 1037 | 1271 | PPTG_13276 | 114,33 | 4 | 1 | 3 | 4 | | 0 | | 9 | 1 | 3 | 6 | 12 | Oxidation (M); Carbamidomethylation; Deamidation (NQ) | 213438 | PPTG_13276 | Phytophthora parasitica INRA-310 ATP-binding Cassette (ABC) superfamily (1934 aa) |
| 1421 | 1468 | PPTG_02754 | 114,23 | 31 | 17 | 14 | 31 | 7,25E+05 | 7,69E+05 | 6,78E+06 | 7 | 7 | 3 | 3 | 8 | Acetylation (N-term) | 24631 | PPTG_02754 | Phytophthora parasitica INRA-310 hypothetical protein (220 aa) |
| 1345 | 1430 | PPTG_15143 | 114,19 | 15 | 11 | 15 | 11 | 8,24E+05 | 9,71E+06 | 9,22E+06 | 6 | 6 | 5 | 6 | 5 | Carbamidomethylation | 56536 | PPTG_15143 | Phytophthora parasitica INRA-310 hypothetical protein (507 aa) |
| 1038 | 1212 | PPTG_17118 | 114,16 | 39 | 8 | 27 | 26 | 1,29E+06 | 1,08E+07 | 3,76E+06 | 10 | 10 | 2 | 9 | 10 | Oxidation (M); Carbamidomethylation; Dehydration; Acetylation (N-term) | 45653 | PPTG_17118 | Phytophthora parasitica INRA-310 26S protease regulatory subunit 6B (406 aa) |
| 1020 | 1030 | PPTG_13520 | 114,16 | 34 | 2 | 29 | 23 | 1,36E+05 | 3,77E+06 | 8,80E+06 | 11 | 11 | 1 | 9 | 12 | Oxidation (M); Carbamidomethylation | 45198 | PPTG_13520 | Phytophthora parasitica INRA-310 2-amino-3-ketobutyrate coenzyme A ligase mitochondrial (414 aa) |
| 1007 | 1279 | PPTG_19393 | 114,16 | 15 | 6 | 14 | 13 | 1,12E+06 | 4,02E+06 | 5,48E+06 | 8 | 8 | 4 | 9 | 9 | Oxidation (M) | 56689 | PPTG_19393 | Phytophthora parasitica INRA-310 dihydrolipoyllysine-residue succinyltransferase component-2-oxoglutarate dehydrogenase complex (545 aa) |
| 1159 | 1394 | PPTG_09216 | 114,04 | 15 | 6 | 13 | 10 | 5,60E+05 | 5,77E+06 | 3,54E+06 | 8 | 7 | 4 | 8 | 6 | Deamidation (NQ) | 60672 | PPTG_09216 | Phytophthora parasitica INRA-310 glucosylceramidase (543 aa) |
| 1158 | 1373 | PPTG_09260 | 114,01 | 29 | 16 | 25 | 29 | 1,10E+06 | 5,97E+06 | 1,25E+07 | 7 | 7 | 3 | 7 | 8 | Dehydration; Deamidation (NQ); Acetylation (N-term) | 27352 | PPTG_09260 | Phytophthora parasitica INRA-310 hypothetical protein (244 aa) |
| 1618 | 1119 | PPTG_00519 | 113,99 | 14 | 0 | 7 | 14 | | 6,13E+05 | 7,27E+06 | 8 | 8 | 0 | 3 | 9 | | 85529 | PPTG_00519 | Phytophthora parasitica INRA-310 endothelin-converting enzyme metalloprotease family M13 (779 aa) |
| 1325 | 1208 | PPTG_05916 | 113,99 | 33 | 13 | 28 | 33 | 1,63E+05 | 3,23E+06 | 5,92E+06 | 7 | 7 | 3 | 6 | 7 | Carbamidomethylation | 32908 | PPTG_05916 | Phytophthora parasitica INRA-310 F-actin-capping protein subunit alpha (289 aa) |
| 1389 | 1073 | PPTG_02506 | 113,97 | 28 | 12 | 2 | 28 | 2,00E+05 | 3,11E+05 | 2,85E+06 | 10 | 8 | 4 | 1 | 10 | Oxidation (M); Carbamidomethylation | 51176 | PPTG_02506 | Phytophthora parasitica INRA-310 CorA Metal Ion transporter (MIT) family (457 aa) |
| 1286 | 1213 | PPTG_19427 | 113,95 | 15 | 4 | 11 | 15 | 2,91E+05 | 2,26E+06 | 6,11E+06 | 7 | 7 | 2 | 6 | 8 | Oxidation (M); Carbamidomethylation | 60053 | PPTG_19427 | Phytophthora parasitica INRA-310 FACT complex subunit SSRP1 (534 aa) |
| 932 | 1677 | PPTG_12022 | 113,92 | 18 | 12 | 18 | 15 | 9,08E+04 | 3,66E+05 | 8,62E+05 | 5 | 2 | 6 | 10 | 8 | Oxidation (M); Carbamidomethylation; Deamidation (NQ) | 40584 | PPTG_12022 | Phytophthora parasitica INRA-310 D-isomer specific 2-hydroxyacid dehydrogenase (372 aa) |
| 1261 | 2320 | PPTG_16172 | 113,9 | 25 | 6 | 25 | 25 | 2,84E+06 | 5,20E+07 | 4,75E+07 | 5 | 5 | 1 | 8 | 8 | Oxidation (M); Carbamidomethylation; Deamidation (NQ) | 30072 | PPTG_16172 | Phytophthora parasitica INRA-310 hypothetical protein (274 aa) |
| 2172 | 1963 | PPTG_19428 | 113,86 | 12 | 3 | 12 | 6 | 2,68E+05 | 4,70E+06 | 2,56E+06 | 4 | 4 | 1 | 4 | 2 | Carbamidomethylation | 49536 | PPTG_19428 | Phytophthora parasitica INRA-310 GPI-anchored leucine-rich lipoprotein (454 aa) |
| 1229 | 1105 | PPTG_17926 | 113,8 | 25 | 8 | 15 | 20 | 1,37E+06 | 3,23E+06 | 1,45E+07 | 9 | 9 | 3 | 6 | 8 | Carbamidomethylation | 48053 | PPTG_17926 | Phytophthora parasitica INRA-310 3-deoxy-7-phosphoheptulonate synthase (442 aa) |
| 1012 | 1031 | PPTG_00934 | 113,78 | 35 | 14 | 18 | 31 | 8,57E+05 | 3,37E+06 | 1,19E+07 | 9 | 9 | 4 | 7 | 11 | Oxidation (M); Carbamidomethylation | 32330 | PPTG_00934 | Phytophthora parasitica INRA-310 protein phosphatase 2C (298 aa) |
| 1309 | 1316 | PPTG_03992 | 113,76 | 29 | 8 | 9 | 27 | 1,00E+06 | 1,33E+06 | 1,74E+07 | 8 | 8 | 3 | 3 | 10 | Oxidation (M); Carbamidomethylation | 33498 | PPTG_03992 | Phytophthora parasitica INRA-310 NADH-cytochrome b5 reductase (301 aa) |
| 1605 | 1152 | PPTG_05790 | 113,75 | 27 | 0 | 23 | 19 | | 8,75E+06 | 3,91E+06 | 7 | 7 | 0 | 7 | 5 | Carbamidomethylation | 37605 | PPTG_05790 | Phytophthora parasitica INRA-310 hypothetical protein (333 aa) |
| 1566 | 1287 | PPTG_16268 | 113,67 | 8 | 2 | 3 | 8 | 2,34E+05 | 7,79E+05 | 1,03E+07 | 6 | 6 | 2 | 2 | 8 | Carbamidomethylation | 115527 | PPTG_16268 | Phytophthora parasitica INRA-310 cellulose synthase 4 (1020 aa) |
| 1402 | 1189 | PPTG_04098 | 113,27 | 16 | 3 | 3 | 16 | 1,14E+06 | 4,01E+05 | 1,29E+07 | 9 | 9 | 3 | 2 | 10 | Carbamidomethylation | 77903 | PPTG_04098 | Phytophthora parasitica INRA-310 hypothetical protein (684 aa) |
| 1081 | 1038 | PPTG_07911 | 113,1 | 8 | 3 | 8 | 8 | 2,50E+05 | 2,87E+06 | 1,99E+06 | 8 | 8 | 3 | 8 | 9 | Carbamidomethylation | 130742 | PPTG_07911 | Phytophthora parasitica INRA-310 hypothetical protein (1164 aa) |
| 1871 | 1199 | PPTG_01133 | 113,01 | 9 | 1 | 0 | 9 | 9,99E+04 | | 4,11E+06 | 8 | 8 | 1 | 0 | 8 | Carbamidomethylation | 124863 | PPTG_01133 | Phytophthora parasitica INRA-310 P-type ATPase (P-ATPase) superfamily (1115 aa) |
| 1320 | 1064 | PPTG_13977 | 113 | 20 | 3 | 15 | 15 | 1,17E+05 | 2,07E+06 | 6,98E+06 | 9 | 9 | 1 | 8 | 7 | Deamidation (NQ) | 55765 | PPTG_13977 | Phytophthora parasitica INRA-310 intraflagellar transporter 52 (502 aa) |
| 1188 | 971 | PPTG_04242 | 112,83 | 14 | 0 | 10 | 12 | | 1,66E+06 | 4,06E+06 | 11 | 11 | 0 | 8 | 10 | | 116261 | PPTG_04242 | Phytophthora parasitica INRA-310 hypothetical protein (1026 aa) |
| 1673 | 1228 | PPTG_07651 | 112,82 | 30 | 2 | 25 | 6 | 2,00E+05 | 9,48E+05 | 6,72E+05 | 9 | 9 | 1 | 7 | 3 | Oxidation (M); Carbamidomethylation; Deamidation (NQ) | 51224 | PPTG_07651 | Phytophthora parasitica INRA-310 hypothetical protein (464 aa) |
| 1912 | 1688 | PPTG_13825 | 112,79 | 17 | 4 | 3 | 17 | 7,38E+05 | 2,80E+05 | 1,11E+07 | 5 | 5 | 2 | 1 | 6 | Carbamidomethylation | 47120 | PPTG_13825 | Phytophthora parasitica INRA-310 protein kinase (426 aa) |
| 1461 | 1793 | PPTG_12106 | 112,76 | 17 | 10 | 14 | 17 | 8,39E+05 | 3,14E+06 | 8,33E+06 | 5 | 5 | 3 | 5 | 6 | | 45944 | PPTG_12106 | Phytophthora parasitica INRA-310 hypothetical protein (505 aa) |
| 1656 | 1083 | PPTG_11458 | 112,7 | 9 | 1 | 0 | 9 | 8,93E+04 | | 5,80E+06 | 9 | 9 | 1 | 0 | 10 | Oxidation (M); Carbamidomethylation; Deamidation (NQ) | 112920 | PPTG_11458 | Phytophthora parasitica INRA-310 hypothetical protein (1003 aa) |
| 1967 | 1254 | PPTG_11171 | 112,63 | 25 | 0 | 6 | 21 | | 1,74E+05 | 4,05E+06 | 7 | 7 | 0 | 2 | 6 | Carbamidomethylation | 39240 | PPTG_11171 | Phytophthora parasitica INRA-310 myosin-like protein (366 aa) |
| 2023 | 1933 | PPTG_02406 | 112,63 | 38 | 18 | 7 | 38 | 1,49E+05 0 | | 4,82E+06 | 4 | 4 | 2 | 1 | 5 | | 24190 | PPTG_02406 | Phytophthora parasitica INRA-310 hypothetical protein (224 aa) |
| 1860 | 1150 | PPTG_04139 | 112,52 | 6 | 0 | 0 | 6 | 5,89E+04 | | 4,06E+06 | 9 | 9 | 1 | 0 | 8 | | 207297 | PPTG_04139 | Phytophthora parasitica INRA-310 hypothetical protein (1938 aa) |
| 1334 | 1056 | PPTG_08625 | 112,48 | 27 | 2 | 22 | 19 | 1,64E+05 | 2,74E+06 | 3,52E+06 | 11 | 9 | 1 | 8 | 7 | Carbamidomethylation | 53026 | PPTG_08625 | Phytophthora parasitica INRA-310 cystathionine beta-synthase (489 aa) |
| 1216 | 1248 | PPTG_11447 | 112,39 | 24 | 11 | 17 | 24 | 1,36E+06 | 3,02E+06 | 1,52E+07 | 8 | 8 | 3 | 5 | 9 | Oxidation (M); Carbamidomethylation; Dehydration; Deamidation (NQ); Carbamylation | 47119 | PPTG_11447 | Phytophthora parasitica INRA-310 serine protease family S33 (424 aa) |
| 2251 | 2146 | PPTG_19415 | 112,21 | 13 | 0 | 10 | 13 | | 1,49E+06 | 1,17E+06 | 4 | 3 | 0 | 3 | 4 | Carbamidomethylation | 57460 | PPTG_19415 | Phytophthora parasitica INRA-310 hypothetical protein (548 aa) |
| 1753 | 1234 | PPTG_17308 | 112,1 | 14 | 7 | 0 | 14 | 2,84E+05 | | 4,46E+06 | 7 | 7 | 3 | 0 | 7 | | 71817 | PPTG_17308 | Phytophthora parasitica INRA-310 hypothetical protein (660 aa) |
| 1121 | 1433 | PPTG_01819 | 112,05 | 34 | 11 | 30 | 28 | 2,38E+06 | 2,21E+07 | 3,48E+07 | 7 | 7 | 3 | 7 | 9 | Carbamidomethylation | 27169 | PPTG_01819 | Phytophthora parasitica INRA-310 proteasome subunit beta (248 aa) |
| 1515 | 1193 | PPTG_07200 | 112,02 | 24 | 4 | 7 | 22 | 1,57E+07 | 2,65E+05 | 5,13E+06 | 8 | 8 | 2 | 2 | 9 | Oxidation (M); Carbamidomethylation; Dehydration; Acetylation (N-term); Amidation; Carbamylation | 37149 | PPTG_07200 | Phytophthora parasitica INRA-310 mitochondrial Carrier (MC) family (338 aa) |
| 1453 | 1131 | PPTG_19709 | 112,01 | 18 | 0 | 15 | 15 | | 3,49E+06 | 4,85E+06 | 8 | 6 | 0 | 7 | 7 | Carbamidomethylation | 66610 | PPTG_19709 | Phytophthora parasitica INRA-310 glycoside hydrolase (590 aa) |
| 1532 | 993 | PPTG_17411 | 111,98 | 33 | 3 | 15 | 25 | 8,10E+04 | 1,88E+06 | 6,05E+06 | 9 | 9 | 1 | 5 | 7 | Carbamidomethylation | 41776 | PPTG_17411 | Phytophthora parasitica INRA-310 hypothetical protein (390 aa) |
| 1145 | 1679 | PPTG_08066 | 111,95 | 44 | 27 | 34 | 44 | 2,29E+06 | 3,52E+07 | 2,41E+07 | 5 | 5 | 3 | 8 | 8 | Oxidation (M); Dehydration; Acetylation (N-term) | 16951 | PPTG_08066 | Phytophthora parasitica INRA-310 hypothetical protein (154 aa) |
| 1165 | 1024 | PPTG_13613 | 111,9 | 8 | 3 | 5 | 8 | 4,71E+05 | 1,82E+06 | 6,51E+06 | 9 | 9 | 4 | 5 | 9 | Carbamidomethylation | 171933 | PPTG_13613 | Phytophthora parasitica INRA-310 hypothetical protein (1562 aa) |
| 1341 | 1188 | PPTG_10401 | 111,78 | 31 | 6 | 9 | 29 | 8,82E+05 | 9,59E+05 | 1,20E+07 | 9 | 9 | 3 | 3 | 10 | Deamidation (NQ) | 35023 | PPTG_10401 | Phytophthora parasitica INRA-310 hypothetical protein (318 aa) |
| 1911 | 1676 | PPTG_12908 | 111,76 | 11 | 2 | 9 | 10 | 6,11E+04 | 9,17E+05 | 1,27E+06 | 5 | 5 | 1 | 4 | 4 | | 60354 | PPTG_12908 | Phytophthora parasitica INRA-310 serine protease family S28 (544 aa) |
| 1082 | 1332 | PPTG_01130 | 111,56 | 14 | 0 | 13 | 10 | | 3,16E+06 | 7,06E+06 | 12 | 12 | 0 | 10 | 10 | Oxidation (M) | 106254 | PPTG_01130 | Phytophthora parasitica INRA-310 serine/threonine-protein phosphatase (936 aa) |
| 1484 | 1573 | PPTG_15118 | 111,54 | 15 | 7 | 10 | 15 | 1,05E+06 | 9,25E+06 | 1,42E+07 | 5 | 5 | 3 | 4 | 6 | | 40501 | PPTG_15118 | Phytophthora parasitica INRA-310 hypothetical protein (356 aa) |
| 1686 | 1335 | PPTG_02655 | 111,54 | 26 | 0 | 7 | 26 | | 7,50E+05 | 1,14E+07 | 6 | 6 | 0 | 2 | 9 | Oxidation (M) | 38689 | PPTG_02655 | Phytophthora parasitica INRA-310 non-selective Cation Channel-2 (NSCC2) family (335 aa) |
| 1606 | 1369 | PPTG_13672 | 111,5 | 34 | 0 | 34 | 12 | | 8,22E+06 | 5,89E+06 | 7 | 7 | 0 | 9 | 3 | Carbamidomethylation; Acetylation (N-term) | 43955 | PPTG_13672 | Phytophthora parasitica INRA-310 cystathionine gamma-lyase (409 aa) |
| 1507 | 1425 | PPTG_05094 | 111,38 | 32 | 0 | 28 | 24 | | 3,12E+06 | 4,48E+06 | 6 | 6 | 0 | 7 | 6 | | 41957 | PPTG_05094 | Phytophthora parasitica INRA-310 hypothetical protein (386 aa) |
| 1223 | 1333 | PPTG_19880 | 111,38 | 22 | 2 | 13 | 16 | 4,45E+05 | 7,00E+06 | 4,93E+06 | 11 | 11 | 1 | 8 | 8 | Oxidation (M); Carbamidomethylation | 64681 | PPTG_19880 | Phytophthora parasitica INRA-310 protein phosphatase 2 (584 aa) |
| 1644 | 1404 | PPTG_18716 | 111,37 | 11 | 3 | 10 | 3 | 1,38E+05 | 1,88E+06 | 1,19E+06 | 7 | 7 | 2 | 7 | 2 | Deamidation (NQ) | 84143 | PPTG_18716 | Phytophthora parasitica INRA-310 hypothetical protein (756 aa) |
| 1807 | 1768 | PPTG_07062 | 111,37 | 37 | 11 | 37 | 25 | | 1,80E+05 | | 5 | 1 | 1 | 6 | 3 | Carbamidomethylation; Deamidation (NQ) | 26936 | PPTG_07062 | Phytophthora parasitica INRA-310 protein transporter SEC13 (247 aa) |
| 1276 | 1144 | PPTG_11270 | 111,28 | 7 | 1 | 2 | 7 | 1,27E+05 | 5,33E+05 | 3,39E+06 | 11 | 11 | 2 | 3 | 11 | | 202372 | PPTG_11270 | Phytophthora parasitica INRA-310 calcium/calmodulin-dependent protein kinase kinase (1852 aa) |
| 1317 | 1204 | PPTG_20054 | 111,24 | 37 | 0 | 25 | 37 | | 8,08E+06 | 1,45E+07 | 9 | 9 | 0 | 7 | 9 | Carbamidomethylation; Acetylation (N-term) | 41281 | PPTG_20054 | Phytophthora parasitica INRA-310 hypothetical protein (368 aa) |
| 1477 | 2031 | PPTG_13262 | 111,21 | 23 | 4 | 23 | 23 | 2,20E+05 | 3,03E+06 | 1,37E+07 | 5 | 5 | 1 | 5 | 8 | | 29931 | PPTG_13262 | Phytophthora parasitica INRA-310 enoyl-CoA hydratase/isomerase family protein (270 aa) |
| 1189 | 1037 | PPTG_15656 | 111,15 | 32 | 4 | 25 | 26 | 2,81E+05 | 4,05E+06 | 9,18E+06 | 8 | 8 | 1 | 8 | 9 | Oxidation (M); Carbamidomethylation | 35158 | PPTG_15656 | Phytophthora parasitica INRA-310 hypothetical protein (317 aa) |
| 1291 | 1156 | PPTG_17510 | 111,12 | 15 | 2 | 12 | 15 | 6,70E+04 | 3,20E+06 | 3,03E+06 | 8 | 8 | 1 | 6 | 9 | Carbamidomethylation | 68077 | PPTG_17510 | Phytophthora parasitica INRA-310 hypothetical protein (634 aa) |
| 1395 | 1878 | PPTG_17122 | 111,08 | 37 | 22 | 37 | 33 | 3,60E+05 | 1,42E+07 | 8,60E+06 | 4 | 4 | 2 | 7 | 6 | Oxidation (M); Carbamidomethylation; Deamidation (NQ) | 22655 | PPTG_17122 | Phytophthora parasitica INRA-310 hypothetical protein (205 aa) |
| 1625 | 1967 | PPTG_10740 | 110,93 | 32 | 0 | 32 | 32 | | 5,66E+06 | 3,82E+06 | 4 | 4 | 0 | 5 | 7 | Oxidation (M); Deamidation (NQ); Acetylation (N-term) | 28097 | PPTG_10740 | Phytophthora parasitica INRA-310 CUG-BP- and ETR-3-like factor (263 aa) |
| 1770 | 1383 | PPTG_08097 | 110,65 | 15 | 1 | 0 | 14 | 1,55E+05 | | 6,81E+06 | 10 | 10 | 1 | 0 | 9 | Oxidation (M) | 83998 | PPTG_08097 | Phytophthora parasitica INRA-310 hypothetical protein (744 aa) |
| 2126 | 1590 | PPTG_09481 | 110,55 | 19 | 0 | 7 | 19 | | 4,39E+05 | 2,48E+06 | 5 | 5 | 0 | 2 | 5 | | 36253 | PPTG_09481 | Phytophthora parasitica INRA-310 hypothetical protein (317 aa) |
| 1689 | 1581 | PPTG_08464 | 110,44 | 33 | 3 | 12 | 33 | 2,12E+05 | 1,40E+06 | 1,69E+07 | 6 | 6 | 1 | 2 | 8 | | 26469 | PPTG_08464 | Phytophthora parasitica INRA-310 hypothetical protein (234 aa) |
| 1568 | 1215 | PPTG_01600 | 110,44 | 9 | 1 | 4 | 7 | 6,74E+04 | 7,02E+05 | 1,43E+06 | 9 | 9 | 1 | 4 | 7 | Oxidation (M); Amidation | 146464 | PPTG_01600 | Phytophthora parasitica INRA-310 glycoside hydrolase (1337 aa) |
| 1576 | 1224 | PPTG_11855 | 110,35 | 23 | 7 | 4 | 23 | 1,93E+05 | 2,43E+05 | 5,17E+06 | 7 | 7 | 2 | 1 | 9 | Oxidation (M); Carbamidomethylation | 39587 | PPTG_11855 | Phytophthora parasitica INRA-310 Ras family GTPase (352 aa) |
| 1292 | 1470 | PPTG_10539 | 110,35 | 17 | 7 | 12 | 17 | 4,72E+05 | 3,11E+06 | 4,70E+06 | 6 | 6 | 3 | 6 | 7 | Oxidation (M); Deamidation (NQ) | 59581 | PPTG_10539 | Phytophthora parasitica INRA-310 translation initiation factor eIF-2B subunit delta (539 aa) |
| 1412 | 1045 | PPTG_06152 | 110,35 | 10 | 0 | 8 | 10 | | 1,80E+06 | 3,19E+06 | 8 | 8 | 0 | 6 | 8 | | 115054 | PPTG_06152 | Phytophthora parasitica INRA-310 insulin-degrading-like enzyme metalloprotease family M16A (1007 aa) |
| 1254 | 1222 | PPTG_05144 | 110,25 | 45 | 6 | 37 | 31 | 9,21E+04 | 4,37E+06 | 3,21E+06 | 9 | 9 | 1 | 9 | 7 | Carbamidomethylation; Deamidation (NQ); Acetylation (N-term) | 31061 | PPTG_05144 | Phytophthora parasitica INRA-310 aldo/keto reductase family protein (278 aa) |
| 575 | 2581 | PPTG_09688 | 110,22 | 45 | 17 | 33 | 45 | 1,79E+07 | 1,44E+08 | 5,63E+08 | 5 | 5 | 5 | 11 | 22 | Oxidation (M); Carbamidomethylation; Deamidation (NQ); Carbamylation | 9060 | PPTG_09688 | Phytophthora parasitica INRA-310 40S ribosomal protein S27 (83 aa) |
| 977 | 1021 | PPTG_09435 | 110,19 | 18 | 7 | 11 | 18 | 6,23E+05 | 1,19E+06 | 8,30E+06 | 10 | 10 | 5 | 7 | 11 | Carbamidomethylation; Dehydration | 71810 | PPTG_09435 | Phytophthora parasitica INRA-310 hypothetical protein (654 aa) |
| 2094 | 3173 | PPTG_10199 | 110,08 | 15 | 0 | 15 | 13 | | 6,65E+06 | 3,37E+06 | 3 | 3 | 0 | 5 | 3 | Carbamidomethylation | 30845 | PPTG_10199 | Phytophthora parasitica INRA-310 hypothetical protein (298 aa) |
| 819 | 1160 | PPTG_02795 | 110,04 | 8 | 2 | 3 | 8 | 3,58E+06 | 3,14E+06 | 9,89E+06 | 14 | 14 | 5 | 7 | 15 | Carbamidomethylation; Dehydration; Deamidation (NQ); Amidation | 211859 | PPTG_02795 | Phytophthora parasitica INRA-310 myosin-like protein (1862 aa) |
| 1327 | 1636 | PPTG_18767 | 109,99 | 29 | 13 | 29 | 24 | 1,01E+05 | 1,13E+07 | 9,59E+06 | 6 | 6 | 2 | 8 | 6 | Oxidation (M); Carbamidomethylation; Deamidation (NQ) | 25722 | PPTG_18767 | Phytophthora parasitica INRA-310 hypothetical protein (223 aa) |
| 1346 | 1952 | PPTG_18525 | 109,91 | 38 | 28 | 38 | 38 | 1,53E+06 | 1,44E+07 | 2,07E+07 | 5 | 5 | 3 | 7 | 6 | Carbamidomethylation | 19583 | PPTG_18525 | Phytophthora parasitica INRA-310 hypothetical protein (178 aa) |
| 1680 | 1084 | PPTG_17174 | 109,91 | 10 | 2 | 1 | 10 | 1,45E+05 | 5,91E+04 | 3,34E+06 | 8 | 8 | 2 | 1 | 8 | Oxidation (M); Carbamidomethylation | 159713 | PPTG_17174 | Phytophthora parasitica INRA-310 transmembrane protein (1455 aa) |
| 1068 | 1133 | PPTG_01288 | 109,76 | 20 | 7 | 17 | 16 | 1,07E+06 | 5,04E+06 | 1,30E+07 | 10 | 10 | 4 | 8 | 9 | | 59271 | PPTG_01288 | Phytophthora parasitica INRA-310 nicotinate phosphoribosyltransferase (534 aa) |
| 1524 | 1608 | PPTG_17496 | 109,75 | 19 | 2 | 19 | 14 | 9,82E+04 | 3,91E+06 | 3,15E+06 | 5 | 5 | 1 | 8 | 4 | Oxidation (M); Carbamidomethylation; Deamidation (NQ); Acetylation (N-term) | 58475 | PPTG_17496 | Phytophthora parasitica INRA-310 GPI anchor protein (568 aa) |
| 1592 | 1416 | PPTG_03239 | 109,75 | 24 | 0 | 19 | 16 | | 5,36E+05 | 2,90E+06 | 7 | 7 | 0 | 5 | 7 | Oxidation (M); Carbamidomethylation | 41737 | PPTG_03239 | Phytophthora parasitica INRA-310 cysteine protease family C01A (378 aa) |
| 1793 | 1460 | PPTG_07347 | 109,74 | 32 | 0 | 29 | 15 | | 3,05E+06 | 6,71E+05 | 8 | 8 | 0 | 7 | 3 | Carbamidomethylation; Deamidation (NQ) | 43318 | PPTG_07347 | Phytophthora parasitica INRA-310 hypothetical protein (422 aa) |
| 1025 | 1136 | PPTG_16802 | 109,72 | 20 | 10 | 5 | 16 | 1,36E+06 | 3,35E+06 | 1,76E+07 | 12 | 12 | 5 | 4 | 13 | Oxidation (M); Carbamidomethylation | 86708 | PPTG_16802 | Phytophthora parasitica INRA-310 calcineurin-like phosphoesterase (773 aa) |
| 1199 | 1090 | PPTG_14661 | 109,7 | 17 | 2 | 17 | 15 | 3,35E+04 | 2,23E+06 | 3,52E+06 | 8 | 8 | 1 | 8 | 9 | Oxidation (M); Carbamidomethylation | 67151 | PPTG_14661 | Phytophthora parasitica INRA-310 hypothetical protein (598 aa) |
| 1750 | 1280 | PPTG_08182 | 109,68 | 11 | 0 | 10 | 5 | | 1,52E+06 | 6,07E+05 | 8 | 8 | 0 | 6 | 4 | | 91109 | PPTG_08182 | Phytophthora parasitica INRA-310 hypothetical protein (828 aa) |
| 1642 | 1306 | PPTG_09875 | 109,66 | 12 | 2 | 6 | 12 | 6,48E+05 | 1,46E+06 | 8,67E+06 | 6 | 6 | 1 | 3 | 7 | Carbamidomethylation; Deamidation (NQ) | 73468 | PPTG_09875 | Phytophthora parasitica INRA-310 long-chain fatty acid transporter (657 aa) |
| 1413 | 1307 | PPTG_08785 | 109,64 | 8 | 2 | 6 | 8 | 4,23E+05 | 2,80E+06 | 1,53E+07 | 7 | 7 | 1 | 5 | 8 | Carbamidomethylation | 103125 | PPTG_08785 | Phytophthora parasitica INRA-310 hypothetical protein (945 aa) |
| 1015 | 1185 | PPTG_15547 | 109,6 | 19 | 15 | 19 | 19 | 2,23E+06 | 5,54E+06 | 2,03E+07 | 7 | 5 | 6 | 8 | 8 | Oxidation (M); Carbamidomethylation; Deamidation (NQ) | 44266 | PPTG_15547 | Phytophthora parasitica INRA-310 hypothetical protein (409 aa) |
| 1095 | 1321 | PPTG_13569 | 109,59 | 13 | 4 | 8 | 13 | 5,81E+05 | 2,98E+06 | 8,28E+06 | 8 | 8 | 3 | 5 | 12 | Oxidation (M); Carbamidomethylation | 98598 | PPTG_13569 | Phytophthora parasitica INRA-310 pyruvate phosphate dikinase (901 aa) |
| 1409 | 1250 | PPTG_01292 | 109,54 | 22 | 3 | 16 | 22 0 | | 2,04E+06 | 6,42E+06 | 7 | 7 | 1 | 5 | 8 | | 39865 | PPTG_01292 | Phytophthora parasitica INRA-310 protein phosphatase 1 regulatory subunit (354 aa) |
| 1528 | 1055 | PPTG_06643 | 109,53 | 29 | 0 | 17 | 29 | | 1,78E+06 | 4,87E+06 | 8 | 8 | 0 | 5 | 8 | Carbamidomethylation | 41624 | PPTG_06643 | Phytophthora parasitica INRA-310 ferrochelatase (370 aa) |
| 2115 | 1446 | PPTG_18636 | 109,52 | 12 | 0 | 1 | 12 | | 2,71E+05 | 2,41E+06 | 6 | 6 | 0 | 1 | 6 | Carbamidomethylation | 86825 | PPTG_18636 | Phytophthora parasitica INRA-310 hypothetical protein (782 aa) |
| 689 | 1508 | PPTG_18599 | 109,48 | 38 | 24 | 31 | 32 | 1,46E+08 | 5,29E+08 | 1,45E+09 | 8 | 8 | 7 | 12 | 13 | Carbamidomethylation; Carbamylation | 16487 | PPTG_18599 | Phytophthora parasitica INRA-310 60S ribosomal protein L27a (149 aa) |
| 1762 | 1474 | PPTG_01118 | 109,35 | 5 | 2 | 3 | 5 | 3,47E+04 | 1,37E+06 | 1,40E+06 | 6 | 6 | 1 | 3 | 6 | | 169788 | PPTG_01118 | Phytophthora parasitica INRA-310 COPII coat assembly protein SEC16 (1622 aa) |
| 1394 | 1305 | PPTG_18019 | 109,28 | 12 | 1 | 10 | 10 | 1,67E+05 | 1,61E+06 | 3,33E+06 | 9 | 8 | 1 | 7 | 7 | Carbamidomethylation; Deamidation (NQ) | 96880 | PPTG_18019 | Phytophthora parasitica INRA-310 protein kinase (875 aa) |
| 1093 | 1042 | PPTG_09030 | 109,19 | 12 | 4 | 10 | 11 | 3,79E+05 | 4,73E+06 | 1,03E+07 | 10 | 10 | 3 | 8 | 9 | Carbamidomethylation | 98306 | PPTG_09030 | Phytophthora parasitica INRA-310 bifunctional aspartokinase/homoserine dehydrogenase (894 aa) |
| 939 | 1663 | PPTG_18756 | 109,15 | 38 | 27 | 27 | 33 | 6,99E+05 | 9,49E+05 | 6,65E+06 | 5 | 2 | 5 | 7 | 12 | Oxidation (M); Carbamidomethylation; Deamidation (NQ); Carbamylation | 24316 | PPTG_18756 | Phytophthora parasitica INRA-310 NADH ubiquinone oxidoreductase (219 aa) |
| 1607 | 1406 | PPTG_01488 | 108,94 | 13 | 3 | 11 | 13 | 2,04E+05 | 8,78E+05 | 4,16E+06 | 6 | 6 | 1 | 5 | 6 | | 56440 | PPTG_01488 | Phytophthora parasitica INRA-310 signal recognition particle (512 aa) |
| 1357 | 1652 | PPTG_06136 | 108,88 | 75 | 31 | 73 | 75 | | 6,94E+05 | 3,66E+05 | 6 | 2 | 2 | 7 | 6 | Oxidation (M) | 9105 | PPTG_06136 | Phytophthora parasitica INRA-310 hypothetical protein (78 aa) |
| 1921 | 1613 | PPTG_16777 | 108,7 | 12 | 4 | 4 | 12 | 7,69E+05 | 1,68E+06 | 6,75E+06 | 5 | 5 | 2 | 2 | 5 | | 55119 | PPTG_16777 | Phytophthora parasitica INRA-310 hypothetical protein (496 aa) |
| 1128 | 1117 | PPTG_10101 | 108,63 | 25 | 3 | 18 | 23 | 6,06E+05 | 2,95E+07 | 2,43E+07 | 9 | 9 | 1 | 9 | 9 | Oxidation (M); Carbamidomethylation | 48967 | PPTG_10101 | Phytophthora parasitica INRA-310 DEAD-box ATP-dependent RNA helicase 56 (429 aa) |
| 1832 | 1239 | PPTG_12321 | 108,56 | 29 | 0 | 23 | 19 | | 1,16E+06 | 9,50E+05 | 7 | 7 | 0 | 5 | 5 | | 33404 | PPTG_12321 | Phytophthora parasitica INRA-310 geranylgeranyl pyrophosphate synthetase (297 aa) |
| 1301 | 1186 | PPTG_11582 | 108,56 | 4 | 0 | 3 | 2 | | 9,92E+05 | 6,08E+05 | 11 | 6 | 0 | 9 | 7 | Carbamidomethylation | 385368 | PPTG_11582 | Phytophthora parasitica INRA-310 axonemal dynein heavy chain (3448 aa) |
| 2330 | 2357 | PPTG_18389 | 108,56 | 31 | 0 | 31 | 18 | | 5,36E+06 | 1,97E+06 | 3 | 3 | 0 | 4 | 2 | Oxidation (M); Carbamidomethylation | 15205 | PPTG_18389 | Phytophthora parasitica INRA-310 hypothetical protein (137 aa) |
| 1490 | 1153 | PPTG_15462 | 108,54 | 7 | 0 | 5 | 6 | | 2,46E+06 | 1,70E+06 | 9 | 9 | 0 | 6 | 7 | Carbamidomethylation | 174962 | PPTG_15462 | Phytophthora parasitica INRA-310 hypothetical protein (1586 aa) |

| | | | | | | | | | | | | | | | | | | | |
|---|---|---|---|---|---|---|---|---|---|---|---|---|---|---|---|---|---|---|---|
| 1284 | 1372 | PPTG_00783 | 108,29 | 15 | 10 | 15 | 10 | 1,33E+05 | 1,63E+06 | 1,66E+06 | 6 | 1 | 4 | 7 | 5 | Carbamidomethylation; Deamidation (NQ) | 51843 | PPTG_00783 | Phytophthora parasitica INRA-310 hypothetical protein (465 aa) |
| 1811 | 1940 | PPTG_10959 | 108,28 | 42 | 17 | 34 | 42 | 2,59E+05 | 2,50E+06 | 4,97E+06 | 5 | 5 | 1 | 4 | 5 | Oxidation (M) | 17792 | PPTG_10959 | Phytophthora parasitica INRA-310 hypothetical protein (159 aa) |
| 1339 | 1227 | PPTG_02654 | 108,24 | 25 | 5 | 15 | 25 | 3,20E+05 | 3,07E+06 | 8,38E+06 | 8 | 8 | 2 | 5 | 9 | Carbamidomethylation; Deamidation (NQ); Acetylation (N-term) | 39146 | PPTG_02654 | Phytophthora parasitica INRA-310 hypothetical protein (363 aa) |
| 1368 | 1180 | PPTG_10769 | 108,23 | 31 | 7 | 31 | 18 | 6,00E+05 | 3,80E+06 | 5,44E+06 | 8 | 8 | 2 | 8 | 5 | Carbamidomethylation; Deamidation (NQ) | 38893 | PPTG_10769 | Phytophthora parasitica INRA-310 ribosome maturation protein SBDS (350 aa) |
| 1099 | 1066 | PPTG_00702 | 108,22 | 23 | 0 | 23 | 17 | | 1,85E+07 | 1,97E+07 | 9 | 9 | 0 | 10 | 10 | Oxidation (M); Carbamidomethylation; Deamidation (NQ) | 58780 | PPTG_00702 | Phytophthora parasitica INRA-310 GMP synthase [glutamine-hydrolyzing] (529 aa) |
| 1874 | 1604 | PPTG_06256 | 108,2 | 19 | 0 | 19 | 16 | | 3,90E+06 | 2,33E+06 | 5 | 4 | 0 | 5 | 4 | Carbamidomethylation | 32852 | PPTG_06256 | Phytophthora parasitica INRA-310 hypothetical protein (319 aa) |
| 1973 | 1374 | PPTG_15785 | 108,19 | 9 | 1 | 2 | 9 | 5,37E+04 | 5,04E+04 | 1,55E+06 | 6 | 6 | 1 | 1 | 6 | Carbamidomethylation | 110741 | PPTG_15785 | Phytophthora parasitica INRA-310 hypothetical protein (990 aa) |
| 91 | 1674 | PPTG_10116 | 108,16 | 53 | 42 | 52 | 53 | 3,31E+08 | 1,06E+09 | 2,13E+09 | 8 | 7 | 25 | 40 | 48 | Oxidation (M); Dehydration; Deamidation (NQ); Acetylation (N-term); Carbamylation | 11411 | PPTG_10116 | Phytophthora parasitica INRA-310 histone H4 (104 aa) |
| 91 | 1675 | PPTG_02835 | 108,16 | 53 | 42 | 52 | 53 | 3,31E+08 | 1,06E+09 | 2,13E+09 | 8 | 7 | 25 | 40 | 48 | Oxidation (M); Dehydration; Deamidation (NQ); Acetylation (N-term); Carbamylation | 11411 | PPTG_02835 | Phytophthora parasitica INRA-310 histone H4 (104 aa) |
| 1556 | 1548 | PPTG_11020 | 108,16 | 5 | 2 | 4 | 4 | 2,13E+05 | 2,27E+06 | 3,16E+06 | 6 | 6 | 2 | 5 | 5 | | 176756 | PPTG_11020 | Phytophthora parasitica INRA-310 paired amphipathic helix protein Sin3 (1599 aa) |
| 1177 | 1005 | PPTG_10296 | 108,16 | 21 | 1 | 8 | 21 | 0 | 8,97E+05 | 5,08E+06 | 10 | 10 | 1 | 5 | 12 | Carbamidomethylation; Deamidation (NQ) | 82804 | PPTG_10296 | Phytophthora parasitica INRA-310 hypothetical protein (749 aa) |
| 2269 | 1978 | PPTG_15051 | 108,05 | 16 | 0 | 0 | 16 | | | 5,96E+06 | 4 | 4 | 0 | 0 | 6 | Oxidation (M) | 44208 | PPTG_15051 | Phytophthora parasitica INRA-310 voltage-gated Ion Channel (VIC) superfamily (402 aa) |
| 1371 | 1391 | PPTG_11045 | 108,03 | 16 | 6 | 16 | 13 | 3,00E+05 | 4,71E+06 | 3,71E+06 | 6 | 5 | 4 | 6 | 5 | Carbamidomethylation | 44927 | PPTG_11045 | Phytophthora parasitica INRA-310 acyl-CoA dehydrogenase (412 aa) |
| 2333 | 2540 | PPTG_00602 | 107,97 | 14 | 0 | 2 | 14 | | 3,24E+05 | 6,85E+06 | 4 | 4 | 0 | 1 | 5 | Oxidation (M); Carbamidomethylation | 43054 | PPTG_00602 | Phytophthora parasitica INRA-310 hypothetical protein (402 aa) |
| 1354 | 1176 | PPTG_02143 | 107,92 | 7 | 1 | 3 | 6 | 3,24E+05 | 6,00E+05 | 4,41E+06 | 9 | 9 | 1 | 4 | 10 | Oxidation (M); Carbamidomethylation | 159421 | PPTG_02143 | Phytophthora parasitica INRA-310 P-type ATPase (P-ATPase) superfamily (1445 aa) |
| 946 | 1711 | PPTG_01894 | 107,89 | 41 | 32 | 41 | 41 | 1,06E+07 | 1,54E+07 | 1,45E+08 | 6 | 6 | 4 | 7 | 13 | Oxidation (M); Deamidation (NQ) | 16221 | PPTG_01894 | Phytophthora parasitica INRA-310 hypothetical protein (143 aa) |
| 1255 | 1175 | PPTG_00771 | 107,87 | 22 | 6 | 12 | 22 | 3,22E+05 | 5,86E+05 | 5,92E+06 | 7 | 6 | 2 | 5 | 10 | Carbamidomethylation; Deamidation (NQ) | 53012 | PPTG_00771 | Phytophthora parasitica INRA-310 serine palmitoyltransferase (484 aa) |
| 1187 | 1447 | PPTG_11225 | 107,86 | 8 | 1 | 6 | 6 | 0 | 2,35E+06 | 5,12E+06 | 11 | 11 | 1 | 7 | 10 | Oxidation (M) | 157252 | PPTG_11225 | Phytophthora parasitica INRA-310 hypothetical protein (1375 aa) |
| 1457 | 1559 | PPTG_14825 | 107,85 | 21 | 12 | 21 | 18 | 2,24E+05 | 1,75E+06 | 2,35E+06 | 5 | 5 | 3 | 6 | 5 | | 42991 | PPTG_14825 | Phytophthora parasitica INRA-310 WD domain-containing protein (393 aa) |
| 1662 | 1483 | PPTG_01392 | 107,84 | 33 | 4 | 9 | 33 | 1,03E+05 | 3,74E+05 | 8,10E+06 | 7 | 7 | 1 | 2 | 8 | | 36267 | PPTG_01392 | Phytophthora parasitica INRA-310 hypothetical protein (326 aa) |
| 1489 | 1108 | PPTG_03305 | 107,79 | 10 | 0 | 9 | 3 | | 1,25E+06 | 1,87E+06 | 10 | 10 | 0 | 8 | 5 | Carbamidomethylation | 177739 | PPTG_03305 | Phytophthora parasitica INRA-310 5'-3' exoribonuclease 1 (1595 aa) |
| 1646 | 1455 | PPTG_04732 | 107,69 | 17 | 0 | 14 | 14 | | 2,10E+06 | 1,35E+06 | 6 | 6 | 0 | 6 | 5 | Carbamidomethylation | 56413 | PPTG_04732 | Phytophthora parasitica INRA-310 cytosol aminopeptidase (520 aa) |
| 1646 | 1456 | PPTG_04732 | 107,69 | 17 | 0 | 14 | 14 | | 2,10E+06 | 1,35E+06 | 6 | 6 | 0 | 6 | 5 | Carbamidomethylation | 56413 | PPTG_04732 | Phytophthora parasitica INRA-310 cytosol aminopeptidase (520 aa) |
| 1316 | 1081 | PPTG_09819 | 107,66 | 16 | 1 | 14 | 12 | 3,72E+05 | 3,46E+06 | 5,59E+06 | 9 | 9 | 1 | 8 | 7 | Carbamidomethylation | 75446 | PPTG_09819 | Phytophthora parasitica INRA-310 hypothetical protein (677 aa) |
| 1520 | 1884 | PPTG_04396 | 107,61 | 29 | 8 | 20 | 29 | 3,60E+06 | 8,60E+06 | 5,50E+07 | 4 | 4 | 2 | 5 | 6 | | 22389 | PPTG_04396 | Phytophthora parasitica INRA-310 hypothetical protein (207 aa) |
| 1746 | 1773 | PPTG_18146 | 107,5 | 5 | 3 | 4 | 3 | 9,88E+05 | 2,72E+06 | 7,89E+06 | 5 | 1 | 3 | 4 | 3 | Dehydration; Acetylation (N-term) | 114447 | PPTG_18146 | Phytophthora parasitica INRA-310 ATP-binding Cassette (ABC) superfamily (1045 aa) |
| 1565 | 1256 | PPTG_00987 | 107,49 | 9 | 4 | 3 | 8 | 2,59E+05 | 4,51E+05 | 3,93E+06 | 7 | 7 | 3 | 3 | 6 | Carbamidomethylation | 101461 | PPTG_00987 | Phytophthora parasitica INRA-310 protein kinase (929 aa) |
| 1616 | 1998 | PPTG_15231 | 107,48 | 33 | 0 | 33 | 33 | | 1,24E+07 | 1,31E+07 | 6 | 6 | 0 | 5 | 7 | Oxidation (M) | 25376 | PPTG_15231 | Phytophthora parasitica INRA-310 NPP1-like protein (238 aa) |
| 1245 | 1094 | PPTG_19697 | 107,39 | 23 | 5 | 20 | 23 | 6,16E+05 | 6,97E+06 | 1,09E+07 | 8 | 8 | 2 | 7 | 8 | Carbamidomethylation | 46809 | PPTG_19697 | Phytophthora parasitica INRA-310 fumarylacetoacetase (426 aa) |
| 1173 | 1123 | PPTG_13738 | 107,39 | 7 | 1 | 6 | 5 | 2,41E+05 | 3,16E+06 | 3,47E+06 | 10 | 9 | 2 | 8 | 8 | Carbamidomethylation | 170836 | PPTG_13738 | Phytophthora parasitica INRA-310 DNA-directed RNA polymerase III largest subunit (1535 aa) |
| 1305 | 1338 | PPTG_00662 | 107,34 | 8 | 1 | 8 | 6 | 5,39E+05 | 4,66E+06 | 6,25E+06 | 7 | 7 | 1 | 8 | 7 | Oxidation (M) | 143193 | PPTG_00662 | Phytophthora parasitica INRA-310 eukaryotic translation initiation factor 3 subunit (1305 aa) |
| 1443 | 1343 | PPTG_13532 | 107,32 | 17 | 4 | 17 | 13 | 8,91E+04 | 1,67E+06 | 1,14E+06 | 7 | 7 | 2 | 7 | 5 | Carbamidomethylation | 56087 | PPTG_13532 | Phytophthora parasitica INRA-310 hypothetical protein (509 aa) |
| 1201 | 1187 | PPTG_01026 | 107,3 | 15 | 7 | 11 | 15 | 4,07E+05 | 3,72E+06 | 6,05E+06 | 8 | 8 | 4 | 6 | 8 | Carbamidomethylation | 64360 | PPTG_01026 | Phytophthora parasitica INRA-310 hypothetical protein (561 aa) |
| 1661 | 1202 | PPTG_06992 | 107,26 | 17 | 2 | 2 | 15 | 1,84E+05 | 0 | 4,69E+06 | 9 | 9 | 1 | 1 | 9 | Oxidation (M); Carbamidomethylation | 72660 | PPTG_06992 | Phytophthora parasitica INRA-310 phosphatidylinositol-4-phosphate-5-kinase (PIPK-C) (651 aa) |
| 1424 | 1134 | PPTG_05312 | 107,25 | 10 | 2 | 1 | 10 | 2,52E+05 | 0 | 5,16E+06 | 9 | 9 | 3 | 1 | 10 | | 119671 | PPTG_05312 | Phytophthora parasitica INRA-310 hypothetical protein (1060 aa) |
| 1314 | 1039 | PPTG_02401 | 107,23 | 25 | 4 | 10 | 23 | 3,32E+05 | 3,94E+06 | 2,04E+07 | 9 | 8 | 3 | 4 | 9 | Oxidation (M); Carbamidomethylation | 45227 | PPTG_02401 | Phytophthora parasitica INRA-310 protein kinase (401 aa) |
| 1562 | 1183 | PPTG_13428 | 107,16 | 5 | 0 | 0 | 5 | | | 3,56E+06 | 12 | 12 | 0 | 0 | 12 | Carbamidomethylation | 313955 | PPTG_13428 | Phytophthora parasitica INRA-310 hypothetical protein (2847 aa) |
| 1267 | 1201 | PPTG_03813 | 107,13 | 16 | 4 | 9 | 16 | 1,35E+06 | 3,44E+06 | 1,38E+07 | 9 | 8 | 3 | 5 | 9 | Oxidation (M); Carbamidomethylation | 77202 | PPTG_03813 | Phytophthora parasitica INRA-310 HECT E3 ubiquitin ligase (684 aa) |
| 1318 | 1294 | PPTG_17805 | 107,08 | 84 | 21 | 76 | 84 | 1,27E+06 | 3,05E+07 | 2,97E+07 | 7 | 7 | 2 | 7 | 7 | | 10557 | PPTG_17805 | Phytophthora parasitica INRA-310 chaperonin (100 aa) |
| 498 | 1735 | PPTG_01387 | 107,05 | 33 | 21 | 26 | 33 | 7,95E+07 | 2,88E+08 | 7,68E+08 | 7 | 7 | 9 | 13 | 21 | Carbamidomethylation; Dehydration; Deamidation (NQ); Amidation; Carbamylation | 17143 | PPTG_01387 | Phytophthora parasitica INRA-310 60S ribosomal protein L24 (152 aa) |
| 1533 | 1448 | PPTG_19290 | 107,03 | 21 | 10 | 17 | 21 | 6,07E+05 | 3,43E+06 | 6,94E+06 | 6 | 6 | 3 | 4 | 6 | Carbamidomethylation | 41313 | PPTG_19290 | Phytophthora parasitica INRA-310 3-methyl-2-oxobutanoate hydroxymethyltransferase (377 aa) |
| 1870 | 3091 | PPTG_18068 | 107,01 | 57 | 24 | 57 | 24 | 1,00E+05 | 1,10E+06 | 8,49E+05 | 2 | 2 | 1 | 5 | 3 | Carbamidomethylation | 9951 | PPTG_18068 | Phytophthora parasitica INRA-310 hypothetical protein (88 aa) |
| 1300 | 1122 | PPTG_01276 | 106,99 | 16 | 2 | 11 | 15 | 2,86E+05 | 6,00E+06 | 7,71E+06 | 9 | 9 | 1 | 7 | 8 | Oxidation (M); Carbamidomethylation | 83617 | PPTG_01276 | Phytophthora parasitica INRA-310 serine/threonine-protein phosphatase 6 regulatory subunit (763 aa) |
| 1563 | 1568 | PPTG_05217 | 106,97 | 26 | 6 | 6 | 26 | 2,70E+05 | 6,78E+05 | 1,36E+07 | 5 | 5 | 1 | 3 | 8 | Oxidation (M); Deamidation (NQ) | 36376 | PPTG_05217 | Phytophthora parasitica INRA-310 3-ketodihydrosphingosine reductase (338 aa) |
| 1473 | 1262 | PPTG_04952 | 106,96 | 46 | 5 | 25 | 46 | 1,15E+06 | 8,33E+06 | 2,06E+07 | 7 | 7 | 1 | 5 | 8 | Carbamidomethylation | 20242 | PPTG_04952 | Phytophthora parasitica INRA-310 hypothetical protein (177 aa) |
| 1535 | 1532 | PPTG_18841 | 106,93 | 28 | 9 | 12 | 26 | 1,27E+06 | 1,91E+06 | 1,25E+07 | 6 | 6 | 3 | 3 | 7 | Oxidation (M) | 37521 | PPTG_18841 | Phytophthora parasitica INRA-310 hypothetical protein (343 aa) |
| 1104 | 1082 | PPTG_16625 | 106,91 | 19 | 7 | 12 | 19 | 4,44E+05 | 1,65E+06 | 7,77E+06 | 10 | 10 | 4 | 6 | 10 | Oxidation (M); Carbamidomethylation | 65597 | PPTG_16625 | Phytophthora parasitica INRA-310 hypothetical protein (590 aa) |
| 1302 | 1191 | PPTG_02638 | 106,86 | 8 | 1 | 7 | 5 | 2,74E+05 | 1,83E+06 | 3,47E+06 | 10 | 10 | 2 | 8 | 6 | Oxidation (M); Carbamidomethylation; Acetylation (N-term) | 139010 | PPTG_02638 | Phytophthora parasitica INRA-310 hypothetical protein (1268 aa) |
| 1441 | 1192 | PPTG_09093 | 106,85 | 32 | 0 | 16 | 32 | | 1,74E+06 | 6,21E+06 | 9 | 9 | 0 | 5 | 9 | Acetylation (N-term) | 37868 | PPTG_09093 | Phytophthora parasitica INRA-310 autophagocytosis associated protein (338 aa) |
| 1083 | 1930 | PPTG_17690 | 106,83 | 10 | 7 | 7 | 10 | 4,77E+06 | 6,89E+06 | 4,13E+07 | 4 | 3 | 4 | 5 | 10 | Oxidation (M); Deamidation (NQ); Carbamylation | 52000 | PPTG_17690 | Phytophthora parasitica INRA-310 hypothetical protein (474 aa) |
| 1506 | 1266 | PPTG_11613 | 106,77 | 19 | 2 | 13 | 19 | 4,69E+04 | 2,97E+06 | 3,96E+06 | 7 | 7 | 1 | 5 | 7 | | 61512 | PPTG_11613 | Phytophthora parasitica INRA-310 intraflagellar transporter 72/74 (536 aa) |
| 1734 | 1329 | PPTG_12918 | 106,74 | 33 | 7 | 7 | 33 | 3,35E+05 | 7,39E+05 | 4,86E+06 | 7 | 7 | 2 | 2 | 7 | Carbamidomethylation; Dehydration | 33365 | PPTG_12918 | Phytophthora parasitica INRA-310 mitochondrial Carrier (MC) family (308 aa) |
| 1043 | 1537 | PPTG_12021 | 106,74 | 19 | 7 | 14 | 19 | 0 | 2,22E+06 | 3,33E+06 | 5 | 2 | 4 | 8 | 9 | Oxidation (M); Carbamidomethylation; Deamidation (NQ) | 36085 | PPTG_12021 | Phytophthora parasitica INRA-310 D-isomer specific 2-hydroxyacid dehydrogenase (334 aa) |
| 1452 | 1099 | PPTG_00628 | 106,62 | 44 | 9 | 21 | 44 | 2,79E+05 | 6,13E+06 | 9,13E+06 | 7 | 7 | 2 | 5 | 7 | Oxidation (M) | 31948 | PPTG_00628 | Phytophthora parasitica INRA-310 hypothetical protein (301 aa) |
| 1690 | 1589 | PPTG_15160 | 106,6 | 14 | 0 | 12 | 11 | | 2,88E+06 | 2,37E+06 | 5 | 5 | 0 | 5 | 6 | Oxidation (M); Carbamidomethylation | 56775 | PPTG_15160 | Phytophthora parasitica INRA-310 pre-mRNA-processing factor 19 (526 aa) |
| 2062 | 1507 | PPTG_09272 | 106,52 | 29 | 0 | 11 | 24 | | 3,76E+06 | 6,24E+06 | 6 | 6 | 0 | 3 | 5 | | 33165 | PPTG_09272 | Phytophthora parasitica INRA-310 hypothetical protein (304 aa) |
| 1619 | 1363 | PPTG_15088 | 106,49 | 19 | 2 | 17 | 9 | 1,29E+05 | 2,69E+06 | 1,83E+06 | 7 | 7 | 1 | 7 | 4 | Carbamidomethylation | 52217 | PPTG_15088 | Phytophthora parasitica INRA-310 COP9 signalosome complex subunit 3 (466 aa) |
| 1250 | 1261 | PPTG_02622 | 106,48 | 37 | 10 | 33 | 37 | 2,43E+06 | 3,50E+07 | 3,58E+07 | 7 | 7 | 2 | 7 | 8 | Carbamidomethylation; Acetylation (N-term) | 27053 | PPTG_02622 | Phytophthora parasitica INRA-310 hypothetical protein (252 aa) |
| 1312 | 1730 | PPTG_03369 | 106,45 | 11 | 5 | 5 | 11 | 2,73E+05 | 3,34E+06 | 4,69E+06 | 5 | 5 | 3 | 6 | 7 | Carbamidomethylation; Dehydration; Acetylation (N-term); Carbamylation | 60642 | PPTG_03369 | Phytophthora parasitica INRA-310 glucosylceramidase (540 aa) |
| 1063 | 1330 | PPTG_10317 | 106,41 | 28 | 7 | 14 | 26 | 6,35E+05 | 7,78E+06 | 1,34E+07 | 12 | 12 | 3 | 7 | 11 | Oxidation (M); Carbamidomethylation; Acetylation (N-term) | 54459 | PPTG_10317 | Phytophthora parasitica INRA-310 3-phosphoinositide-dependent protein kinase (486 aa) |
| 1722 | 1671 | PPTG_18933 | 106,41 | 20 | 8 | 16 | 20 | 8,40E+04 | 1,96E+06 | 2,97E+06 | 5 | 4 | 2 | 4 | 5 | Carbamidomethylation | 38560 | PPTG_18933 | Phytophthora parasitica INRA-310 glucokinase (352 aa) |
| 1722 | 1672 | PPTG_18927 | 106,41 | 20 | 8 | 16 | 20 | 8,40E+04 | 1,96E+06 | 2,97E+06 | 5 | 4 | 2 | 4 | 5 | Carbamidomethylation | 38649 | PPTG_18927 | Phytophthora parasitica INRA-310 glucokinase (353 aa) |
| 1474 | 1336 | PPTG_15364 | 106,36 | 14 | 4 | 14 | 9 | 1,92E+05 | 4,82E+06 | 1,67E+06 | 7 | 7 | 2 | 8 | 4 | Carbamidomethylation | 64267 | PPTG_15364 | Phytophthora parasitica INRA-310 phenylalanine and histidine ammonia-lyase (600 aa) |
| 1392 | 1770 | PPTG_03064 | 106,33 | 13 | 6 | 13 | 11 | 6,30E+05 | 8,14E+06 | 6,21E+06 | 5 | 4 | 2 | 8 | 5 | Oxidation (M); Carbamidomethylation | 47979 | PPTG_03064 | Phytophthora parasitica INRA-310 hypothetical protein (447 aa) |
| 1610 | 1434 | PPTG_02204 | 106,21 | 32 | 11 | 21 | 32 | 4,10E+05 | 2,41E+06 | 8,55E+06 | 6 | 6 | 2 | 4 | 6 | Oxidation (M); Carbamidomethylation | 27215 | PPTG_02204 | Phytophthora parasitica INRA-310 hypothetical protein (247 aa) |
| 1204 | 1467 | PPTG_12102 | 106,15 | 12 | 3 | 12 | 12 | 2,07E+05 | 5,37E+06 | 7,74E+06 | 6 | 4 | 2 | 8 | 8 | Oxidation (M); Carbamidomethylation | 64201 | PPTG_12102 | Phytophthora parasitica INRA-310 lactase-phlorizin hydrolase (574 aa) |
| 1759 | 1351 | PPTG_07417 | 106,12 | 9 | 2 | 0 | 8 | | | 1,66E+06 | 9 | 4 | 2 | 0 | 8 | Oxidation (M); Carbamidomethylation; Dehydration; Deamidation (NQ) | 163054 | PPTG_07417 | Phytophthora parasitica INRA-310 resistance-Nodulation-Cell Division (RND) superfamily (1477 aa) |
| 1238 | 1098 | PPTG_12242 | 106,04 | 15 | 5 | 13 | 13 | 5,85E+05 | 3,72E+06 | 8,33E+06 | 8 | 8 | 3 | 7 | 7 | | 65972 | PPTG_12242 | Phytophthora parasitica INRA-310 D-3-phosphoglycerate dehydrogenase (604 aa) |
| 1602 | 1375 | PPTG_17994 | 105,91 | 17 | 7 | 12 | 13 | 2,17E+05 | 2,33E+06 | 2,84E+06 | 6 | 6 | 2 | 4 | 6 | Oxidation (M) | 47657 | PPTG_17994 | Phytophthora parasitica INRA-310 hypothetical protein (469 aa) |
| 1984 | 1894 | PPTG_13689 | 105,64 | 8 | 4 | 4 | 8 | 1,85E+05 | 2,68E+05 | 3,62E+06 | 4 | 4 | 2 | 2 | 4 | | 77324 | PPTG_13689 | Phytophthora parasitica INRA-310 hypothetical protein (718 aa) |
| 2233 | 1945 | PPTG_10517 | 105,63 | 34 | 7 | 14 | 34 | 5,22E+05 | 2,99E+06 | 7,56E+06 | 4 | 4 | 1 | 2 | 4 | | 18977 | PPTG_10517 | Phytophthora parasitica INRA-310 caltractin (167 aa) |
| 1220 | 1249 | PPTG_18898 | 105,61 | 8 | 1 | 6 | 7 | 7,10E+04 | 1,37E+06 | 8,90E+06 | 10 | 10 | 1 | 7 | 9 | Oxidation (M); Carbamidomethylation | 172701 | PPTG_18898 | Phytophthora parasitica INRA-310 RAC family serine/threonine-protein kinase (1560 aa) |
| 1329 | 1694 | PPTG_11742 | 105,59 | 28 | 23 | 23 | 28 | 1,30E+06 | 5,09E+06 | 8,84E+06 | 5 | 5 | 4 | 5 | 7 | Oxidation (M) | 25677 | PPTG_11742 | Phytophthora parasitica INRA-310 hypothetical protein (244 aa) |
| 1933 | 1798 | PPTG_11824 | 105,5 | 22 | 8 | 16 | 22 | 6,05E+04 | 2,90E+06 | 3,96E+06 | 5 | 5 | 1 | 3 | 5 | Carbamidomethylation | 37062 | PPTG_11824 | Phytophthora parasitica INRA-310 hypothetical protein (349 aa) |
| 1593 | 1519 | PPTG_09033 | 105,5 | 13 | 10 | 5 | 13 | 3,41E+05 | 5,29E+05 | 6,38E+06 | 6 | 6 | 4 | 2 | 6 | Oxidation (M) | 55111 | PPTG_09033 | Phytophthora parasitica INRA-310 CAAX prenyl protease 1 (485 aa) |
| 1669 | 1469 | PPTG_04171 | 105,48 | 9 | 3 | 5 | 9 | 4,16E+05 | 7,52E+05 | 4,53E+06 | 6 | 6 | 2 | 3 | 6 | | 76897 | PPTG_04171 | Phytophthora parasitica INRA-310 SEC14 cytosolic factor (704 aa) |
| 300 | 2186 | PPTG_06043 | 105,44 | 38 | 37 | 37 | 38 | 8,29E+07 | 3,34E+08 | 8,61E+08 | 4 | 4 | 9 | 23 | 32 | Carbamidomethylation; Deamidation (NQ); Carbamylation | 10453 | PPTG_06043 | Phytophthora parasitica INRA-310 60S ribosomal protein L37a (96 aa) |
| 1178 | 1766 | PPTG_15678 | 105,37 | 44 | 29 | 38 | 44 | 7,97E+06 | 1,79E+07 | 9,06E+07 | 5 | 5 | 4 | 6 | 8 | Deamidation (NQ) | 13394 | PPTG_15678 | Phytophthora parasitica INRA-310 hypothetical protein (117 aa) |
| 1634 | 1666 | PPTG_20426 | 105,33 | 20 | 10 | 14 | 20 | 6,68E+06 | 1,08E+07 | 5,03E+07 | 5 | 5 | 3 | 4 | 5 | | 34268 | PPTG_20426 | Phytophthora parasitica INRA-310 cytochrome b (297 aa) |
| 1643 | 1357 | PPTG_10290 | 105,33 | 6 | 1 | 5 | 5 | 1,25E+04 | 2,16E+06 | 1,04E+06 | 7 | 7 | 1 | 5 | 5 | Carbamidomethylation | 142388 | PPTG_10290 | Phytophthora parasitica INRA-310 hypothetical protein (1282 aa) |
| 1137 | 1172 | PPTG_01253 | 105,23 | 15 | 4 | 14 | 7 | 4,66E+05 | 4,56E+06 | 4,18E+06 | 11 | 11 | 3 | 10 | 6 | Carbamidomethylation | 109518 | PPTG_01253 | Phytophthora parasitica INRA-310 U5 small nuclear ribonucleoprotein component (987 aa) |
| 1919 | 3182 | PPTG_08421 | 105,18 | 28 | 23 | 23 | 28 | 6,69E+06 | 1,08E+07 | 4,79E+07 | 3 | 3 | 2 | 3 | 4 | Carbamidomethylation; Deamidation (NQ); Acetylation (N-term) | 17255 | PPTG_08421 | Phytophthora parasitica INRA-310 PLAC8 family protein (161 aa) |
| 1422 | 1331 | PPTG_17908 | 105,18 | 9 | 5 | 8 | 6 | 5,14E+05 | 2,84E+06 | 5,38E+06 | 6 | 6 | 3 | 6 | 5 | Dehydration; Deamidation (NQ) | 80796 | PPTG_17908 | Phytophthora parasitica INRA-310 hypothetical protein (713 aa) |
| 1879 | 1423 | PPTG_11868 | 105 | 25 | 7 | 4 | 25 | 2,36E+05 | 3,61E+04 | 4,38E+06 | 6 | 6 | 2 | 1 | 6 | Oxidation (M) | 32818 | PPTG_11868 | Phytophthora parasitica INRA-310 mitochondrial Carrier (MC) family (304 aa) |
| 2141 | 2065 | PPTG_11871 | 104,96 | 25 | 0 | 25 | 15 | | 2,10E+06 | 6,78E+05 | 4 | 4 | 0 | 4 | 3 | Oxidation (M) | 31005 | PPTG_11871 | Phytophthora parasitica INRA-310 hypothetical protein (280 aa) |
| 1675 | 1502 | PPTG_08002 | 104,88 | 21 | 3 | 7 | 18 | 8,81E+04 | 0 | 3,86E+06 | 7 | 7 | 1 | 2 | 8 | Deamidation (NQ) | 53760 | PPTG_08002 | Phytophthora parasitica INRA-310 hypothetical protein (493 aa) |
| 1419 | 1678 | PPTG_19369 | 104,87 | 16 | 4 | 10 | 16 | | | 3,96E+04 | 5 | 1 | 2 | 6 | 6 | Oxidation (M); Carbamidomethylation; Carbamylation | 65129 | PPTG_19369 | Phytophthora parasitica INRA-310 NADP-dependent malic enzyme (597 aa) |
| 972 | 1126 | PPTG_05510 | 104,85 | 16 | 4 | 12 | 16 | 4,50E+05 | 5,44E+06 | 9,87E+06 | 10 | 10 | 2 | 10 | 11 | Oxidation (M); Carbamidomethylation | 68483 | PPTG_05510 | Phytophthora parasitica INRA-310 ribosomal RNA methyltransferase nop2-like protein (609 aa) |
| 1553 | 2104 | PPTG_07371 | 104,83 | 4 | 3 | 3 | 4 | | | 1,44E+05 | 4 | 1 | 3 | 4 | 5 | | 137375 | PPTG_07371 | Phytophthora parasitica INRA-310 ATP-binding Cassette (ABC) superfamily (1269 aa) |
| 1105 | 1696 | PPTG_19683 | 104,82 | 46 | 13 | 46 | 46 | 4,80E+06 | 4,10E+07 | 3,92E+07 | 6 | 6 | 2 | 9 | 9 | Oxidation (M); Carbamidomethylation; Deamidation (NQ) | 16658 | PPTG_19683 | Phytophthora parasitica INRA-310 proteasome subunit alpha type-6 (150 aa) |
| 1905 | 3455 | PPTG_14695 | 104,54 | 23 | 0 | 0 | 23 | | | 3,61E+07 | 4 | 4 | 0 | 0 | 9 | Oxidation (M) | 23478 | PPTG_14695 | Phytophthora parasitica INRA-310 hypothetical protein (218 aa) |
| 1833 | 1777 | PPTG_01441 | 104,49 | 18 | 9 | 13 | 16 | 5,32E+05 | 5,07E+06 | 5,34E+06 | 5 | 5 | 2 | 4 | 4 | | 38427 | PPTG_01441 | Phytophthora parasitica INRA-310 urate catabolism protein (341 aa) |
| 1387 | 1237 | PPTG_05377 | 104,48 | 38 | 6 | 34 | 33 | 2,61E+05 | 5,88E+06 | 8,71E+06 | 8 | 8 | 1 | 7 | 7 | Carbamidomethylation | 26809 | PPTG_05377 | Phytophthora parasitica INRA-310 hypothetical protein (241 aa) |
| 1822 | 1795 | PPTG_15677 | 104,35 | 51 | 13 | 39 | 51 | 3,48E+05 | 2,28E+06 | 4,76E+06 | 5 | 5 | 1 | 3 | 6 | Acetylation (N-term) | 14126 | PPTG_15677 | Phytophthora parasitica INRA-310 hypothetical protein (123 aa) |
| 981 | 1229 | PPTG_06146 | 104,34 | 24 | 9 | 21 | 24 | 1,63E+06 | 8,73E+06 | 1,47E+07 | 8 | 8 | 4 | 10 | 9 | Oxidation (M); Carbamidomethylation | 51035 | PPTG_06146 | Phytophthora parasitica INRA-310 ATP-dependent RNA helicase DBP5 (459 aa) |
| 1186 | 1259 | PPTG_08964 | 104,25 | 20 | 1 | 16 | 10 | 7,42E+04 | 2,92E+06 | 3,42E+06 | 11 | 11 | 1 | 10 | 7 | Carbamidomethylation | 77426 | PPTG_08964 | Phytophthora parasitica INRA-310 hypothetical protein (699 aa) |
| 1778 | 1471 | PPTG_19041 | 104,24 | 17 | 0 | 17 | 12 | | 4,05E+06 | 1,94E+06 | 6 | 6 | 0 | 6 | 4 | Carbamidomethylation | 45776 | PPTG_19041 | Phytophthora parasitica INRA-310 acetate kinase (429 aa) |
| 1761 | 1268 | PPTG_10899 | 104,2 | 6 | 1 | 1 | 6 | 3,14E+05 | 4,73E+05 | 4,68E+06 | 8 | 8 | 2 | 1 | 7 | Oxidation (M); Carbamidomethylation | 154925 | PPTG_10899 | Phytophthora parasitica INRA-310 ryanodine-Inositol 1 4 5-triphosphate Receptor Ca2 Channel (RIR-CaC) family (1387 aa) |
| 1447 | 1579 | PPTG_09548 | 104,18 | 26 | 3 | 24 | 26 | 8,64E+04 | 2,69E+06 | 4,64E+06 | 6 | 6 | 1 | 6 | 7 | Oxidation (M); Carbamidomethylation | 27575 | PPTG_09548 | Phytophthora parasitica INRA-310 small nuclear ribonucleoprotein-associated protein (263 aa) |
| 1373 | 1623 | PPTG_17414 | 104,17 | 16 | 9 | 12 | 16 | 8,98E+05 | 3,59E+06 | 1,19E+07 | 5 | 5 | 3 | 5 | 7 | Oxidation (M) | 45919 | PPTG_17414 | Phytophthora parasitica INRA-310 hypothetical protein (445 aa) |
| 1546 | 1554 | PPTG_16503 | 104,06 | 39 | 5 | 39 | 39 | 6,13E+05 | 5,63E+06 | 7,95E+06 | 6 | 6 | 1 | 6 | 6 | Carbamidomethylation | 27424 | PPTG_16503 | Phytophthora parasitica INRA-310 peptidyl-prolyl cis-trans isomerase (257 aa) |
| 1476 | 1538 | PPTG_05609 | 104,03 | 18 | 6 | 15 | 18 | 5,78E+05 | 3,92E+06 | 7,23E+06 | 6 | 6 | 2 | 6 | 6 | Oxidation (M); Carbamidomethylation; Deamidation (NQ) | 40345 | PPTG_05609 | Phytophthora parasitica INRA-310 Ras-like GTP-binding protein D (356 aa) |
| 1649 | 1096 | PPTG_04797 | 103,97 | 25 | 0 | 14 | 22 | | 4,60E+05 | 5,44E+06 | 8 | 5 | 0 | 4 | 7 | Oxidation (M) | 42584 | PPTG_04797 | Phytophthora parasitica INRA-310 pyridine nucleotide-disulphide oxidoreductase (394 aa) |
| 1494 | 1562 | PPTG_05256 | 103,96 | 8 | 3 | 6 | 6 | 5,32E+05 | 2,02E+06 | 8,98E+06 | 6 | 6 | 2 | 6 | 4 | Oxidation (M) | 88906 | PPTG_05256 | Phytophthora parasitica INRA-310 LETM1 and EF-hand domain-containing protein 1 (796 aa) |
| 1480 | 1473 | PPTG_04751 | 103,95 | 3 | 0 | 2 | 2 | 0 | 1,07E+06 | 1,86E+06 | 9 | 9 | 1 | 6 | 6 | Carbamidomethylation | 379196 | PPTG_04751 | Phytophthora parasitica INRA-310 hypothetical protein (3343 aa) |
| 1940 | 1571 | PPTG_12540 | 103,91 | 22 | 0 | 18 | 13 | | 6,30E+06 | 2,83E+06 | 6 | 6 | 0 | 5 | 4 | Carbamidomethylation; Deamidation (NQ) | 39580 | PPTG_12540 | Phytophthora parasitica INRA-310 hypothetical protein (385 aa) |

| | | | | | | | | | | | | | | | | | | | |
|---|---|---|---|---|---|---|---|---|---|---|---|---|---|---|---|---|---|---|---|
| 1510 | 1458 | PPTG_13651 | 103,81 | 31 | 3 | 28 | 31 | 2,48E+05 | 5,09E+06 | 6,17E+06 | 6 | 6 | 1 | 6 | 6 | Deamidation (NQ); Acetylation (N-term) | 32402 | PPTG_13651 | Phytophthora parasitica INRA-310 coatomer subunit epsilon (293 aa) |
| 811 | 2016 | PPTG_10197 | 103,76 | 60 | 40 | 60 | 60 | 3,57E+07 | 1,35E+08 | 3,65E+08 | 6 | 6 | 3 | 10 | 15 | Carbamidomethylation; Deamidation (NQ); Carbamylation | 7809 | PPTG_10197 | Phytophthora parasitica INRA-310 30S ribosomal protein S28e (68 aa) |
| 1221 | 1415 | PPTG_16312 | 103,72 | 17 | 0 | 12 | 17 | | 5,65E+06 | 9,75E+06 | 8 | 8 | 0 | 7 | 10 | Oxidation (M); Carbamidomethylation | 62019 | PPTG_16312 | Phytophthora parasitica INRA-310 translation elongation factor 1-alpha (564 aa) |
| 1290 | 1302 | PPTG_18989 | 103,68 | 7 | 3 | 6 | 5 | 1,01E+06 | 3,17E+06 | 6,10E+06 | 7 | 7 | 3 | 6 | 7 | Dehydration; Deamidation (NQ) | 134528 | PPTG_18989 | Phytophthora parasitica INRA-310 inositol-3 4-bisphosphate 4-phosphatase (1214 aa) |
| 1306 | 1561 | PPTG_06494 | 103,66 | 27 | 11 | 22 | 27 | 2,27E+05 | 2,74E+06 | 8,31E+06 | 5 | 5 | 2 | 6 | 8 | | 28825 | PPTG_06494 | Phytophthora parasitica INRA-310 hypothetical protein (271 aa) |
| 1541 | 1747 | PPTG_18900 | 103,63 | 30 | 0 | 30 | 30 | | 6,37E+06 | 9,41E+06 | 5 | 5 | 0 | 6 | 7 | Oxidation (M); Carbamidomethylation | 26659 | PPTG_18900 | Phytophthora parasitica INRA-310 ubiquitin carboxyl-terminal hydrolase (240 aa) |
| 1338 | 3357 | PPTG_00858 | 103,58 | 44 | 26 | 44 | 41 | 1,38E+06 | 2,93E+06 | 3,10E+07 | 4 | 4 | 3 | 6 | 7 | Oxidation (M) | 8013 | PPTG_00858 | Phytophthora parasitica INRA-310 hypothetical protein (74 aa) |
| 2217 | 3374 | PPTG_10233 | 103,56 | 21 | 0 | 21 | 21 | | 7,79E+06 | 1,02E+07 | 3 | 3 | 0 | 3 | 4 | Carbamidomethylation | 15490 | PPTG_10233 | Phytophthora parasitica INRA-310 hypothetical protein (147 aa) |
| 1499 | 1388 | PPTG_13578 | 103,49 | 14 | 3 | 11 | 9 | 9,41E+05 | 2,19E+06 | 2,28E+06 | 8 | 8 | 2 | 6 | 5 | | 76678 | PPTG_13578 | Phytophthora parasitica INRA-310 hypothetical protein (691 aa) |
| 2107 | 1977 | PPTG_19674 | 103,48 | 9 | 1 | 7 | 6 | | 6,78E+05 | 3,41E+05 | 4 | 2 | 1 | 3 | 3 | Carbamidomethylation | 62762 | PPTG_19674 | Phytophthora parasitica INRA-310 transglutaminase elicitor M81C (578 aa) |
| 558 | 1517 | PPTG_04926 | 103,47 | 45 | 20 | 39 | 45 | 5,12E+07 | 2,15E+08 | 5,48E+08 | 10 | 10 | 4 | 15 | 20 | Dehydration; Deamidation (NQ); Acetylation (N-term); Carbamylation | 15595 | PPTG_04926 | Phytophthora parasitica INRA-310 ribosomal protein L28e (140 aa) |
| 1577 | 1371 | PPTG_13556 | 103,46 | 6 | 2 | 4 | 6 | 4,49E+05 | 3,04E+06 | 5,24E+06 | 6 | 6 | 2 | 4 | 6 | Carbamidomethylation | 133722 | PPTG_13556 | Phytophthora parasitica INRA-310 strumpellin (1163 aa) |
| 1650 | 1196 | PPTG_06328 | 103,43 | 10 | 2 | 3 | 10 | 2,33E+04 | 7,26E+05 | 3,03E+06 | 7 | 7 | 1 | 2 | 8 | Carbamidomethylation; Dehydration; Deamidation (NQ) | 88071 | PPTG_06328 | Phytophthora parasitica INRA-310 VAC14 family protein (789 aa) |
| 1710 | 1624 | PPTG_10842 | 103,41 | 14 | 2 | 4 | 14 | 4,60E+05 | 5,73E+05 | 7,34E+06 | 6 | 6 | 1 | 2 | 8 | Oxidation (M); Carbamidomethylation | 57400 | PPTG_10842 | Phytophthora parasitica INRA-310 sodium/hydrogen exchanger 3 (518 aa) |
| 1608 | 1408 | PPTG_01294 | 103,37 | 25 | 3 | 20 | 25 | 1,88E+05 | 2,49E+06 | 5,88E+06 | 7 | 7 | 1 | 5 | 6 | Carbamidomethylation | 42864 | PPTG_01294 | Phytophthora parasitica INRA-310 mitogen-activated protein kinase (375 aa) |
| 1442 | 1235 | PPTG_15779 | 103,31 | 17 | 2 | 13 | 11 | 7,64E+04 | 2,57E+06 | 2,46E+06 | 9 | 9 | 1 | 7 | 6 | | 68814 | PPTG_15779 | Phytophthora parasitica INRA-310 signal recognition particle (615 aa) |
| 1340 | 1504 | PPTG_02908 | 103,29 | 21 | 11 | 17 | 18 | 2,59E+05 | 2,78E+06 | 7,25E+06 | 6 | 6 | 3 | 6 | 7 | Carbamidomethylation | 41410 | PPTG_02908 | Phytophthora parasitica INRA-310 developmentally-regulated GTP-binding protein 1 (370 aa) |
| 1641 | 1600 | PPTG_15692 | 103,29 | 6 | 1 | 4 | 4 | 5,76E+04 | 1,41E+06 | 2,25E+06 | 7 | 7 | 1 | 5 | 5 | | 130280 | PPTG_15692 | Phytophthora parasitica INRA-310 hypothetical protein (1175 aa) |
| 2072 | 2164 | PPTG_14255 | 103,28 | 18 | 0 | 18 | 18 | | 2,38E+06 | 1,75E+06 | 4 | 4 | 0 | 4 | 4 | Carbamidomethylation | 29360 | PPTG_14255 | Phytophthora parasitica INRA-310 hypothetical protein (269 aa) |
| 1377 | 1190 | PPTG_07069 | 103,25 | 25 | 7 | 18 | 25 | 1,13E+06 | 6,41E+06 | 1,91E+07 | 7 | 7 | 2 | 5 | 8 | Carbamidomethylation | 35836 | PPTG_07069 | Phytophthora parasitica INRA-310 30S ribosomal protein S9 (323 aa) |
| 2129 | 1715 | PPTG_02357 | 103,24 | 25 | 0 | 25 | 10 | | 1,68E+06 | 6,51E+05 | 5 | 5 | 0 | 5 | 2 | | 35429 | PPTG_02357 | Phytophthora parasitica INRA-310 hypothetical protein (312 aa) |
| 979 | 1422 | PPTG_11928 | 103,24 | 29 | 4 | 26 | 29 | 6,41E+05 | 2,28E+07 | 3,45E+07 | 7 | 7 | 1 | 11 | 11 | Oxidation (M); Carbamidomethylation | 33178 | PPTG_11928 | Phytophthora parasitica INRA-310 succinyl-CoA ligase subunit alpha (319 aa) |
| 1386 | 1211 | PPTG_11770 | 103,2 | 43 | 18 | 16 | 40 | 1,38E+06 | 4,01E+06 | 1,16E+07 | 8 | 8 | 3 | 4 | 8 | Oxidation (M); Carbamidomethylation; Deamidation (NQ) | 36300 | PPTG_11770 | Phytophthora parasitica INRA-310 hypothetical protein (329 aa) |
| 1838 | 1481 | PPTG_13437 | 103,09 | 17 | 0 | 15 | 14 | | 2,38E+06 | 2,47E+06 | 6 | 6 | 0 | 5 | 5 | Carbamidomethylation | 53319 | PPTG_13437 | Phytophthora parasitica INRA-310 hypothetical protein (481 aa) |
| 1841 | 1778 | PPTG_17517 | 103,02 | 38 | 4 | 30 | 38 | 2,01E+05 | 2,32E+06 | 2,82E+06 | 5 | 5 | 1 | 4 | 5 | Oxidation (M) | 21062 | PPTG_17517 | Phytophthora parasitica INRA-310 hypothetical protein (191 aa) |
| 1202 | 1300 | PPTG_08457 | 103,01 | 29 | 0 | 21 | 29 | | 3,44E+06 | 1,05E+07 | 8 | 8 | 0 | 7 | 11 | Oxidation (M); Carbamidomethylation | 43146 | PPTG_08457 | Phytophthora parasitica INRA-310 3-ketoacyl-CoA thiolase B peroxisomal (412 aa) |
| 1303 | 1320 | PPTG_12597 | 102,96 | 10 | 1 | 7 | 6 | 2,05E+05 | 3,21E+06 | 4,34E+06 | 12 | 12 | 1 | 8 | 7 | Oxidation (M); Carbamidomethylation; Deamidation (NQ) | 138834 | PPTG_12597 | Phytophthora parasitica INRA-310 calpain-like protein (1273 aa) |
| 1658 | 1713 | PPTG_12781 | 102,92 | 9 | 2 | 1 | 8 | 2,15E+05 | 1,36E+05 | 4,98E+06 | 8 | 8 | 2 | 1 | 8 | Oxidation (M); Carbamidomethylation | 122966 | PPTG_12781 | Phytophthora parasitica INRA-310 cGMP-dependent protein kinase (1103 aa) |
| 1582 | 1251 | PPTG_05384 | 102,9 | 12 | 1 | 12 | 6 | 9,88E+04 | 1,85E+06 | 2,52E+06 | 7 | 6 | 1 | 7 | 4 | Oxidation (M) | 89328 | PPTG_05384 | Phytophthora parasitica INRA-310 hypothetical protein (790 aa) |
| 1281 | 1219 | PPTG_11486 | 102,89 | 3 | 0 | 2 | 3 | | 1,25E+06 | 2,06E+07 | 10 | 10 | 0 | 6 | 10 | Deamidation (NQ) | 355472 | PPTG_11486 | Phytophthora parasitica INRA-310 phosphatidylinositol kinase (PIK-L1) (3243 aa) |
| 1586 | 1495 | PPTG_01179 | 102,85 | 15 | 3 | 15 | 15 | 7,66E+04 | 1,44E+06 | 1,85E+06 | 5 | 5 | 1 | 6 | 5 | Acetylation (N-term) | 48320 | PPTG_01179 | Phytophthora parasitica INRA-310 dynein 1 light intermediate chain (436 aa) |
| 1875 | 1610 | PPTG_10723 | 102,81 | 8 | 3 | 0 | 8 | 5,15E+04 | | 4,03E+06 | 5 | 4 | 2 | 0 | 7 | Oxidation (M); Deamidation (NQ) | 89209 | PPTG_10723 | Phytophthora parasitica INRA-310 ATP-binding Cassette (ABC) superfamily protein (826 aa) |
| 1103 | 1985 | PPTG_13081 | 102,78 | 79 | 40 | 79 | 79 | 1,67E+06 | 1,54E+07 | 5,56E+07 | 4 | 4 | 3 | 8 | 9 | Oxidation (M); Carbamidomethylation; Deamidation (NQ); Acetylation (N-term) | 7554 | PPTG_13081 | Phytophthora parasitica INRA-310 hypothetical protein (68 aa) |
| 1674 | 1438 | PPTG_03685 | 102,74 | 6 | 0 | 2 | 6 | | 4,54E+05 | 3,24E+06 | 8 | 8 | 0 | 3 | 8 | | 188483 | PPTG_03685 | Phytophthora parasitica INRA-310 hypothetical protein (1721 aa) |
| 1013 | 2325 | PPTG_04035 | 102,71 | 32 | 32 | 32 | 32 | 1,67E+07 | 4,77E+07 | 2,09E+08 | 3 | 3 | 6 | 7 | 9 | Oxidation (M); Carbamylation | 16234 | PPTG_04035 | Phytophthora parasitica INRA-310 ADP-ribosylation factor (144 aa) |
| 1425 | 1275 | PPTG_18224 | 102,68 | 24 | 7 | 10 | 24 | 2,54E+05 | 2,59E+06 | 7,41E+06 | 7 | 7 | 2 | 4 | 8 | Carbamidomethylation | 46752 | PPTG_18224 | Phytophthora parasitica INRA-310 WD repeat-containing protein 69 (418 aa) |
| 1645 | 1358 | PPTG_18700 | 102,6 | 5 | 0 | 4 | 3 | | 2,55E+06 | 2,02E+06 | 8 | 8 | 0 | 6 | 5 | Acetylation (N-term) | 208915 | PPTG_18700 | Phytophthora parasitica INRA-310 nuclear mitotic apparatus protein (1894 aa) |
| 1772 | 1428 | PPTG_01192 | 102,49 | 16 | 0 | 6 | 14 | | 4,51E+05 | 6,36E+06 | 8 | 8 | 0 | 3 | 7 | | 62190 | PPTG_01192 | Phytophthora parasitica INRA-310 calcineurin subunit B (590 aa) |
| 982 | 1823 | PPTG_08290 | 102,45 | 66 | 22 | 66 | 61 | 2,07E+06 | 4,03E+07 | 2,53E+07 | 6 | 6 | 2 | 13 | 8 | Oxidation (M); Acetylation (N-term) | 9628 | PPTG_08290 | Phytophthora parasitica INRA-310 hypothetical protein (89 aa) |
| 1758 | 1703 | PPTG_11875 | 102,4 | 6 | 2 | 2 | 6 | 1,82E+05 | 3,26E+05 | 4,13E+06 | 5 | 5 | 2 | 2 | 6 | | 95273 | PPTG_11875 | Phytophthora parasitica INRA-310 cell division protease ftsH (874 aa) |
| 1850 | 3179 | PPTG_11034 | 102,34 | 50 | 13 | 50 | 50 | 1,07E+07 | 4,74E+07 | 4,89E+07 | 2 | 2 | 2 | 4 | 4 | Carbamidomethylation | 17486 | PPTG_11034 | Phytophthora parasitica INRA-310 hypothetical protein (169 aa) |
| 1241 | 1938 | PPTG_08250 | 102,32 | 25 | 11 | 25 | 25 | 9,90E+05 | 5,65E+06 | 1,14E+07 | 4 | 4 | 2 | 5 | 10 | Oxidation (M); Carbamidomethylation | 22320 | PPTG_08250 | Phytophthora parasitica INRA-310 hypothetical protein (202 aa) |
| 2145 | 2322 | PPTG_17872 | 102,31 | 20 | 5 | 20 | 20 | 2,72E+05 | 4,53E+06 | 4,93E+06 | 3 | 3 | 1 | 3 | 3 | | 23751 | PPTG_17872 | Phytophthora parasitica INRA-310 hypothetical protein (207 aa) |
| 1212 | 1500 | PPTG_09950 | 102,19 | 7 | 0 | 5 | 7 | | 2,69E+06 | 3,58E+06 | 11 | 11 | 0 | 7 | 10 | Carbamidomethylation | 192737 | PPTG_09950 | Phytophthora parasitica INRA-310 hypothetical protein (1712 aa) |
| 1682 | 1463 | PPTG_09135 | 102,17 | 12 | 0 | 6 | 11 | | 1,40E+06 | 2,59E+06 | 7 | 7 | 0 | 4 | 7 | | 67613 | PPTG_09135 | Phytophthora parasitica INRA-310 phosphoglucomutase (610 aa) |
| 1622 | 1767 | PPTG_12930 | 102,1 | 23 | 6 | 11 | 23 | 3,65E+05 | 3,25E+05 | 7,49E+06 | 6 | 6 | 2 | 3 | 7 | Oxidation (M); Carbamidomethylation; Deamidation (NQ) | 34515 | PPTG_12930 | Phytophthora parasitica INRA-310 hypothetical protein (329 aa) |
| 1609 | 1421 | PPTG_10878 | 101,83 | 31 | 0 | 27 | 24 | | 5,01E+06 | 5,23E+06 | 7 | 7 | 0 | 6 | 6 | Carbamidomethylation | 32587 | PPTG_10878 | Phytophthora parasitica INRA-310 hypothetical protein (304 aa) |
| 1496 | 1378 | PPTG_09573 | 101,78 | 19 | 3 | 17 | 14 | 1,22E+05 | 4,00E+06 | 3,24E+06 | 8 | 8 | 1 | 7 | 5 | Carbamidomethylation; Deamidation (NQ); Amidation | 45442 | PPTG_09573 | Phytophthora parasitica INRA-310 hypothetical protein (405 aa) |
| 1678 | 2153 | PPTG_00139 | 101,77 | 20 | 8 | 12 | 16 | 8,82E+05 | 8,25E+06 | 7,98E+06 | 4 | 4 | 2 | 4 | 5 | Deamidation (NQ); Amidation | 36160 | PPTG_00139 | Phytophthora parasitica INRA-310 alcohol dehydrogenase (333 aa) |
| 1653 | 1772 | PPTG_11476 | 101,68 | 17 | 7 | 9 | 11 | 3,38E+05 | 1,67E+06 | 3,09E+06 | 7 | 7 | 3 | 4 | 4 | Carbamidomethylation | 51545 | PPTG_11476 | Phytophthora parasitica INRA-310 hypothetical protein (473 aa) |
| 1319 | 1642 | PPTG_08193 | 101,64 | 21 | 2 | 19 | 17 | 3,23E+05 | 7,57E+06 | 8,58E+06 | 8 | 8 | 1 | 6 | 9 | Oxidation (M); Carbamidomethylation | 45756 | PPTG_08193 | Phytophthora parasitica INRA-310 uroporphyrinogen decarboxylase (409 aa) |
| 1233 | 1298 | PPTG_18647 | 101,55 | 18 | 5 | 18 | 15 | 1,39E+06 | 1,08E+07 | 2,12E+07 | 7 | 7 | 2 | 8 | 7 | Oxidation (M); Dehydration | 47074 | PPTG_18647 | Phytophthora parasitica INRA-310 glycylpeptide N-tetradecanoyltransferase 2 (409 aa) |
| 1579 | 1223 | PPTG_18888 | 101,53 | 17 | 4 | 8 | 13 | 1,36E+05 | 2,29E+06 | 6,54E+06 | 8 | 8 | 2 | 4 | 6 | Carbamidomethylation | 70514 | PPTG_18888 | Phytophthora parasitica INRA-310 vacuolar protein sorting-associated protein 33A (630 aa) |
| 1633 | 2399 | PPTG_01356 | 101,46 | 23 | 23 | 23 | 23 | 1,37E+06 | 7,67E+05 | 6,95E+06 | 3 | 3 | 4 | 4 | 4 | | 22942 | PPTG_01356 | Phytophthora parasitica INRA-310 6 7-dimethyl-8-ribityllumazine synthase (216 aa) |
| 1868 | 1542 | PPTG_02902 | 101,36 | 25 | 0 | 25 | 15 | | 5,59E+05 | 1,23E+06 | 5 | 5 | 0 | 5 | 4 | Oxidation (M); Carbamidomethylation | 31864 | PPTG_02902 | Phytophthora parasitica INRA-310 hypothetical protein (277 aa) |
| 1620 | 1392 | PPTG_16621 | 101,23 | 29 | 8 | 3 | 29 | 1,65E+05 | 3,80E+05 | 6,06E+06 | 8 | 8 | 2 | 1 | 9 | Oxidation (M); Deamidation (NQ) | 32765 | PPTG_16621 | Phytophthora parasitica INRA-310 mitochondrial Carrier (MC) family (303 aa) |
| 1313 | 1851 | PPTG_12174 | 101,14 | 13 | 7 | 13 | 13 | 8,24E+05 | 7,53E+06 | 5,75E+07 | 5 | 5 | 2 | 5 | 9 | Oxidation (M); Carbamidomethylation | 36027 | PPTG_12174 | Phytophthora parasitica INRA-310 hypothetical protein (320 aa) |
| 1777 | 1194 | PPTG_14016 | 101,1 | 14 | 1 | 1 | 14 | 5,20E+05 | 2,07E+05 | 9,00E+06 | 8 | 8 | 1 | 1 | 8 | Carbamidomethylation | 88560 | PPTG_14016 | Phytophthora parasitica INRA-310 hypothetical protein (804 aa) |
| 1572 | 1520 | PPTG_01821 | 101,1 | 30 | 13 | 16 | 17 | 9,82E+05 | 1,15E+06 | 3,09E+06 | 8 | 8 | 4 | 4 | 4 | Acetylation (N-term) | 27957 | PPTG_01821 | Phytophthora parasitica INRA-310 hypothetical protein (255 aa) |
| 1660 | 1949 | PPTG_11095 | 101,09 | 26 | 0 | 14 | 26 | | 7,55E+06 | 2,68E+07 | 6 | 6 | 0 | 4 | 7 | Carbamidomethylation; Deamidation (NQ) | 27059 | PPTG_11095 | Phytophthora parasitica INRA-310 polysaccharide lyase (250 aa) |
| 1660 | 1950 | PPTG_11096 | 101,09 | 26 | 0 | 14 | 26 | | 7,55E+06 | 2,68E+07 | 6 | 6 | 0 | 4 | 7 | Carbamidomethylation; Deamidation (NQ) | 27059 | PPTG_11096 | Phytophthora parasitica INRA-310 polysaccharide lyase (250 aa) |
| 1836 | 2339 | PPTG_06763 | 101,01 | 13 | 3 | 13 | 13 | 3,95E+05 | 2,48E+06 | 1,15E+07 | 3 | 3 | 1 | 4 | 5 | Oxidation (M); Carbamidomethylation | 42462 | PPTG_06763 | Phytophthora parasitica INRA-310 Solute:Sodium Symporter (SSS) family (397 aa) |
| 1724 | 1756 | PPTG_05390 | 100,97 | 32 | 0 | 14 | 32 | | 2,01E+06 | 1,82E+07 | 5 | 5 | 0 | 4 | 7 | Carbamidomethylation | 38727 | PPTG_05390 | Phytophthora parasitica INRA-310 hypothetical protein (343 aa) |
| 1785 | 1245 | PPTG_08933 | 100,95 | 13 | 3 | 1 | 13 | 2,38E+05 | 2,44E+04 | 9,41E+06 | 7 | 7 | 2 | 1 | 7 | Carbamidomethylation | 75229 | PPTG_08933 | Phytophthora parasitica INRA-310 hypothetical protein (671 aa) |
| 1956 | 1993 | PPTG_13987 | 100,87 | 36 | 11 | 11 | 36 | 6,19E+05 | 7,88E+05 | 6,76E+06 | 4 | 4 | 2 | 2 | 5 | Oxidation (M); Acetylation (N-term) | 22716 | PPTG_13987 | Phytophthora parasitica INRA-310 hypothetical protein (209 aa) |
| 1581 | 1578 | PPTG_13455 | 100,84 | 12 | 4 | 10 | 7 | 3,90E+05 | 1,96E+06 | 6,61E+06 | 6 | 6 | 2 | 5 | 5 | Carbamidomethylation | 66826 | PPTG_13455 | Phytophthora parasitica INRA-310 signal recognition particle receptor subunit alpha (602 aa) |
| 1657 | 1499 | PPTG_06084 | 100,84 | 10 | 4 | 10 | 8 | | 1,34E+05 | 4,44E+05 | 5 | 1 | 2 | 5 | 4 | Carbamidomethylation | 58114 | PPTG_06084 | Phytophthora parasitica INRA-310 hypothetical protein (519 aa) |
| 1670 | 1514 | PPTG_05921 | 100,76 | 19 | 0 | 12 | 19 | | 3,92E+06 | 5,65E+06 | 6 | 6 | 0 | 5 | 6 | Oxidation (M); Carbamidomethylation | 38167 | PPTG_05921 | Phytophthora parasitica INRA-310 dimeric dihydrodiol dehydrogenase (348 aa) |
| 1614 | 1436 | PPTG_01919 | 100,69 | 37 | 19 | 11 | 37 | 1,21E+06 | 2,09E+06 | 1,16E+07 | 7 | 7 | 3 | 2 | 7 | | 21168 | PPTG_01919 | Phytophthora parasitica INRA-310 ADP-ribosylation factor family protein (189 aa) |
| 298 | 1634 | PPTG_18995 | 100,62 | 35 | 30 | 35 | 35 | 1,18E+08 | 4,51E+08 | 1,25E+09 | 5 | 5 | 17 | 23 | 24 | Oxidation (M); Carbamidomethylation; Dehydration; Deamidation (NQ); Carbamylation | 14981 | PPTG_18995 | Phytophthora parasitica INRA-310 40S ribosomal protein S15a (131 aa) |
| 1954 | 1947 | PPTG_00833 | 100,56 | 10 | 5 | 5 | 10 | 4,64E+05 | 1,99E+06 | 8,04E+06 | 4 | 4 | 2 | 2 | 5 | | 62748 | PPTG_00833 | Phytophthora parasitica INRA-310 hypothetical protein (605 aa) |
| 1804 | 1576 | PPTG_14609 | 100,56 | 19 | 0 | 19 | 15 | | 1,12E+06 | 1,33E+06 | 5 | 5 | 0 | 6 | 4 | Oxidation (M); Carbamidomethylation; Acetylation (N-term); Carbamylation | 34775 | PPTG_14609 | Phytophthora parasitica INRA-310 dnaJ heat shock protein (308 aa) |
| 1265 | 1339 | PPTG_04759 | 100,55 | 22 | 6 | 17 | 22 | 2,61E+05 | 4,10E+06 | 9,27E+06 | 8 | 8 | 2 | 6 | 9 | Carbamidomethylation | 45561 | PPTG_04759 | Phytophthora parasitica INRA-310 translation factor pelota (405 aa) |
| 1534 | 1488 | PPTG_11133 | 100,48 | 32 | 0 | 16 | 29 | | 1,31E+06 | 4,30E+06 | 8 | 8 | 0 | 4 | 9 | Oxidation (M) | 32274 | PPTG_11133 | Phytophthora parasitica INRA-310 hypothetical protein (288 aa) |
| 1259 | 1883 | PPTG_08090 | 100,45 | 22 | 19 | 19 | 22 | 3,87E+05 | 8,04E+05 | 3,59E+06 | 4 | 2 | 6 | 5 | 6 | | 22523 | PPTG_08090 | Phytophthora parasitica INRA-310 GTP-binding protein YPTC1 (204 aa) |
| 2118 | 1875 | PPTG_10410 | 100,44 | 27 | 6 | 11 | 27 | 2,52E+05 | 1,46E+06 | 3,61E+06 | 4 | 4 | 1 | 2 | 4 | | 19596 | PPTG_10410 | Phytophthora parasitica INRA-310 hypothetical protein (190 aa) |
| 1464 | 2470 | PPTG_04433 | 100,4 | 42 | 0 | 33 | 42 | | 3,68E+05 | 9,84E+06 | 3 | 3 | 0 | 2 | 12 | Oxidation (M); Dehydration; Deamidation (NQ) | 12953 | PPTG_04433 | Phytophthora parasitica INRA-310 V-type ATPase G subunit (114 aa) |
| 2250 | 2068 | PPTG_12538 | 100,28 | 21 | 4 | 18 | 12 | 1,73E+05 | 4,81E+06 | 4,28E+06 | 4 | 3 | 1 | 3 | 3 | Carbamidomethylation | 29983 | PPTG_12538 | Phytophthora parasitica INRA-310 hypothetical protein (284 aa) |
| 1957 | 1997 | PPTG_17123 | 100,22 | 23 | 7 | 23 | 12 | 1,57E+05 | 6,55E+06 | 3,37E+06 | 4 | 4 | 1 | 5 | 3 | Carbamidomethylation | 23771 | PPTG_17123 | Phytophthora parasitica INRA-310 hypothetical protein (216 aa) |
| 1635 | 1951 | PPTG_10232 | 100,12 | 16 | 7 | 16 | 16 | 1,14E+05 | 6,86E+06 | 3,89E+06 | 4 | 4 | 2 | 5 | 5 | | 35039 | PPTG_10232 | Phytophthora parasitica INRA-310 hypothetical protein (328 aa) |
| 559 | 2366 | PPTG_04153 | 100,08 | 31 | 24 | 31 | 25 | 6,02E+07 | 1,98E+08 | 5,33E+08 | 5 | 5 | 8 | 13 | 18 | Dehydration; Deamidation (NQ); Carbamylation | 14345 | PPTG_04153 | Phytophthora parasitica INRA-310 60S ribosomal protein L35 (124 aa) |
| 1791 | 1743 | PPTG_08714 | 100,07 | 13 | 0 | 2 | 13 | 0 | | 4,06E+06 | 5 | 5 | 0 | 1 | 9 | Oxidation (M); Carbamidomethylation; Dehydration; Deamidation (NQ) | 58203 | PPTG_08714 | Phytophthora parasitica INRA-310 hypothetical protein (520 aa) |
| 2459 | 1871 | PPTG_12121 | 99,97 | 11 | 0 | 0 | 11 | | | 2,25E+06 | 4 | 4 | 0 | 0 | 5 | | 56122 | PPTG_12121 | Phytophthora parasitica INRA-310 zinc transporter (506 aa) |
| 2310 | 1849 | PPTG_10276 | 99,94 | 10 | 4 | 0 | 10 | 2,79E+05 | | 2,97E+06 | 4 | 4 | 2 | 0 | 4 | | 70951 | PPTG_10276 | Phytophthora parasitica INRA-310 cell division protease ftsH (655 aa) |
| 1298 | 1732 | PPTG_13236 | 99,94 | 12 | 3 | 10 | 12 | 8,55E+04 | 2,17E+06 | 2,50E+07 | 6 | 6 | 2 | 5 | 9 | Oxidation (M); Deamidation (NQ) | 59153 | PPTG_13236 | Phytophthora parasitica INRA-310 hypothetical protein (526 aa) |
| 1742 | 2132 | PPTG_18125 | 99,9 | 16 | 0 | 16 | 14 | | 2,46E+06 | 1,21E+06 | 5 | 4 | 0 | 6 | 5 | Carbamidomethylation; Deamidation (NQ) | 46592 | PPTG_18125 | Phytophthora parasitica INRA-310 endoglucanase (442 aa) |
| 1382 | 1291 | PPTG_01665 | 99,89 | 6 | 2 | 6 | 5 | 1,40E+05 | 1,46E+06 | 3,91E+06 | 7 | 7 | 2 | 7 | 6 | Oxidation (M) | 152965 | PPTG_01665 | Phytophthora parasitica INRA-310 hypothetical protein (1363 aa) |
| 1270 | 2373 | PPTG_09123 | 99,86 | 49 | 49 | 49 | 49 | 1,03E+07 | 1,44E+07 | 9,63E+07 | 3 | 3 | 4 | 5 | 8 | Oxidation (M) | 8259 | PPTG_09123 | Phytophthora parasitica INRA-310 hypothetical protein (77 aa) |
| 1808 | 1784 | PPTG_02765 | 99,85 | 29 | 0 | 23 | 25 | | 6,37E+06 | 4,53E+06 | 6 | 6 | 0 | 5 | 5 | Carbamidomethylation; Acetylation (N-term) | 29509 | PPTG_02765 | Phytophthora parasitica INRA-310 anamorsin (277 aa) |
| 1272 | 1370 | PPTG_10277 | 99,66 | 25 | 5 | 18 | 25 | 1,53E+06 | 7,40E+06 | 1,37E+07 | 7 | 7 | 2 | 7 | 8 | Oxidation (M) | 36588 | PPTG_10277 | Phytophthora parasitica INRA-310 hypothetical protein (332 aa) |
| 1864 | 1387 | PPTG_09625 | 99,65 | 9 | 1 | 5 | 7 | 1,54E+05 | 1,13E+06 | 1,89E+06 | 6 | 6 | 1 | 3 | 5 | Carbamidomethylation | 100464 | PPTG_09625 | Phytophthora parasitica INRA-310 heat shock protein 101 (895 aa) |
| 1802 | 1400 | PPTG_11935 | 99,65 | 44 | 4 | 44 | 10 0 | | 3,45E+06 | 2,76E+06 | 7 | 7 | 1 | 7 | 2 | Carbamidomethylation | 30493 | PPTG_11935 | Phytophthora parasitica INRA-310 hypothetical protein (281 aa) |
| 1806 | 1712 | PPTG_05894 | 99,61 | 35 | 4 | 32 | 30 0 | | 4,82E+06 | 1,81E+06 | 5 | 5 | 1 | 4 | 5 | Carbamidomethylation; Deamidation (NQ) | 26420 | PPTG_05894 | Phytophthora parasitica INRA-310 catechol O-methyltransferase (239 aa) |
| 2110 | 1506 | PPTG_10955 | 99,54 | 4 | 0 | 0 | 4 | | | 1,90E+06 | 7 | 7 | 0 | 0 | 7 | | 227633 | PPTG_10955 | Phytophthora parasitica INRA-310 hypothetical protein (2064 aa) |
| 1583 | 1410 | PPTG_03706 | 99,54 | 10 | 1 | 6 | 5 | 1,49E+05 | 2,55E+06 | 1,06E+06 | 10 | 10 | 1 | 7 | 4 | Carbamidomethylation | 137925 | PPTG_03706 | Phytophthora parasitica INRA-310 trehalose-phosphatase (1247 aa) |
| 1444 | 1218 | PPTG_19212 | 99,45 | 24 | 9 | 10 | 24 | 5,99E+05 | 9,97E+05 | 8,24E+06 | 8 | 8 | 3 | 3 | 8 | Oxidation (M) | 39019 | PPTG_19212 | Phytophthora parasitica INRA-310 hypothetical protein (358 aa) |
| 1570 | 1601 | PPTG_06470 | 99,45 | 9 | 5 | 8 | 6 | 4,27E+05 | 2,04E+06 | 2,92E+06 | 5 | 5 | 3 | 4 | 5 | Carbamidomethylation; Deamidation (NQ) | 84430 | PPTG_06470 | Phytophthora parasitica INRA-310 translation elongation factor G (763 aa) |
| 1560 | 1405 | PPTG_09604 | 99,44 | 5 | 1 | 2 | 4 | 4,12E+04 | 2,61E+05 | 6,72E+07 | 10 | 10 | 1 | 3 | 8 | Carbamidomethylation; Amidation; Carbamylation | 246490 | PPTG_09604 | Phytophthora parasitica INRA-310 hypothetical protein (2237 aa) |
| 1521 | 2044 | PPTG_10860 | 99,44 | 33 | 9 | 27 | 32 | 1,40E+05 | 3,56E+06 | 1,09E+07 | 6 | 6 | 1 | 5 | 7 | Oxidation (M); Carbamidomethylation | 20155 | PPTG_10860 | Phytophthora parasitica INRA-310 hypothetical protein (185 aa) |
| 2645 | 3340 | PPTG_13153 | 99,41 | 11 | 7 | 11 | 11 | 4,82E+05 | 4,55E+06 | 6,59E+06 | 2 | 2 | 1 | 2 | 2 | Carbamidomethylation | 38916 | PPTG_13153 | Phytophthora parasitica INRA-310 hypothetical protein (381 aa) |
| 1872 | 1424 | PPTG_12058 | 99,37 | 22 | 0 | 9 | 22 | | 1,69E+06 | 7,21E+06 | 6 | 6 | 0 | 2 | 7 | Carbamidomethylation | 42137 | PPTG_12058 | Phytophthora parasitica INRA-310 hypothetical protein (392 aa) |
| 1517 | 1558 | PPTG_07141 | 99,36 | 26 | 6 | 26 | 26 | 1,23E+05 | 3,91E+06 | 3,95E+06 | 6 | 6 | 1 | 7 | 5 | Carbamidomethylation; Deamidation (NQ) | 33350 | PPTG_07141 | Phytophthora parasitica INRA-310 phytanoyl-CoA dioxygenase domain-containing protein (296 aa) |
| 1439 | 2379 | PPTG_18670 | 99,35 | 30 | 16 | 23 | 29 | 9,10E+05 | 8,59E+06 | 8,68E+06 | 4 | 4 | 2 | 7 | 5 | Oxidation (M) | 13138 | PPTG_18670 | Phytophthora parasitica INRA-310 hypothetical protein (117 aa) |
| 1410 | 1503 | PPTG_18965 | 99,34 | 6 | 1 | 2 | 4 | 5,95E+04 | 2,51E+05 | 2,75E+06 | 9 | 8 | 3 | 4 | 7 | | 188423 | PPTG_18965 | Phytophthora parasitica INRA-310 kinesin-like protein (1702 aa) |
| 1924 | 1722 | PPTG_19116 | 99,28 | 36 | 0 | 29 | 30 | | 2,73E+06 | 1,45E+06 | 5 | 5 | 0 | 5 | 4 | Carbamidomethylation | 22419 | PPTG_19116 | Phytophthora parasitica INRA-310 hypothetical protein (200 aa) |
| 1691 | 1698 | PPTG_05904 | 99,27 | 23 | 9 | 18 | 11 | 3,75E+05 | 1,76E+06 | 1,07E+06 | 7 | 7 | 3 | 5 | 3 | Carbamidomethylation; Acetylation (N-term) | 34967 | PPTG_05904 | Phytophthora parasitica INRA-310 translation initiation factor eIF-2B subunit alpha (321 aa) |

| | | | | | | | | | | | | | | | | | | |
|---|---|---|---|---|---|---|---|---|---|---|---|---|---|---|---|---|---|---|
| 1816 | 1209 | PPTG_05777 | 99,26 | 19 | 0 | 0 | 19 | | | 2,38E+06 | 8 | 8 | 0 | 0 | 10 | Acetylation (N-term) | 79909 | \| PPTG_05777 \| Phytophthora parasitica INRA-310 receptor kinase (714 aa) |
| 1732 | 2410 | PPTG_08722 | 99,25 | 17 | 0 | 17 | 17 | | 1,26E+07 | 1,01E+07 | 4 | 4 | 0 | 5 | 6 | Oxidation (M); Carbamidomethylation | 49630 | \| PPTG_08722 \| Phytophthora parasitica INRA-310 SCP-like extracellular protein (458 aa) |
| 1672 | 1689 | PPTG_15665 | 99,15 | 25 | 9 | 15 | 25 | 5,41E+05 | 1,76E+06 | 6,06E+06 | 5 | 5 | 2 | 3 | 6 | Oxidation (M); Carbamidomethylation; Acetylation (N-term) | 28134 | \| PPTG_15665 \| Phytophthora parasitica INRA-310 hypothetical protein (263 aa) |
| 1460 | 1727 | PPTG_16395 | 99,09 | 21 | 5 | 5 | 21 | 5,06E+05 | 6,64E+05 | 5,50E+06 | 5 | 5 | 2 | 2 | 10 | Oxidation (M); Carbamidomethylation | 45833 | \| PPTG_16395 \| Phytophthora parasitica INRA-310 hypothetical protein (420 aa) |
| 1745 | 1680 | PPTG_00794 | 99 | 5 | 2 | 1 | 5 | | | 1,71E+05 | 5 | 3 | 3 | 1 | 6 | Carbamidomethylation | 144878 | \| PPTG_00794 \| Phytophthora parasitica INRA-310 myosin-like protein (1278 aa) |
| 1372 | 1309 | PPTG_18630 | 98,83 | 9 | 0 | 8 | 7 | | 2,71E+06 | 3,05E+06 | 9 | 9 | 0 | 8 | 7 | Carbamidomethylation | 132430 | \| PPTG_18630 \| Phytophthora parasitica INRA-310 trehalose-phosphatase (1179 aa) |
| 1446 | 1459 | PPTG_08325 | 98,83 | 14 | 8 | 14 | 11 | 8,94E+05 | 4,46E+06 | 7,73E+06 | 6 | 6 | 3 | 6 | 5 | Oxidation (M); Carbamidomethylation | 52883 | \| PPTG_08325 \| Phytophthora parasitica INRA-310 xylulose kinase (494 aa) |
| 1817 | 1493 | PPTG_19303 | 98,82 | 30 | 0 | 15 | 22 | | 1,55E+06 | 2,55E+06 | 7 | 7 | 0 | 5 | 5 | Oxidation (M); Acetylation (N-term) | 42782 | \| PPTG_19303 \| Phytophthora parasitica INRA-310 hsp90-like protein (399 aa) |
| 1738 | 2116 | PPTG_02280 | 98,75 | 35 | 6 | 35 | 35 | 3,16E+05 | 9,18E+06 | 1,29E+07 | 4 | 4 | 1 | 5 | 5 | | 15327 | \| PPTG_02280 \| Phytophthora parasitica INRA-310 HIT domain-containing protein (145 aa) |
| 1530 | 1272 | PPTG_11511 | 98,73 | 36 | 3 | 31 | 26 | 6,94E+04 | 3,75E+06 | 1,38E+06 | 7 | 7 | 1 | 7 | 5 | | 32466 | \| PPTG_11511 \| Phytophthora parasitica INRA-310 F-actin-capping protein subunit beta (288 aa) |
| 1894 | 1955 | PPTG_06558 | 98,66 | 34 | 0 | 28 | 28 | | 1,38E+06 | 5,93E+05 | 5 | 5 | 0 | 5 | 4 | | 17482 | \| PPTG_06558 \| Phytophthora parasitica INRA-310 hypothetical protein (156 aa) |
| 1786 | 1353 | PPTG_02351 | 98,64 | 29 | 0 | 25 | 23 | | 1,25E+06 | 1,41E+06 | 6 | 6 | 0 | 5 | 5 | | 30200 | \| PPTG_02351 \| Phytophthora parasitica INRA-310 guanylate kinase (272 aa) |
| 1775 | 1891 | PPTG_13091 | 98,63 | 18 | 0 | 18 | 11 | | 4,59E+06 | 2,54E+06 | 5 | 5 | 0 | 7 | 3 | Carbamidomethylation; Deamidation (NQ) | 39993 | \| PPTG_13091 \| Phytophthora parasitica INRA-310 hypothetical protein (369 aa) |
| 1861 | 1546 | PPTG_05645 | 98,5 | 6 | 1 | 6 | 4 | 3,88E+05 | 2,92E+06 | 2,33E+06 | 5 | 5 | 1 | 5 | 3 | Oxidation (M); Carbamidomethylation | 119478 | \| PPTG_05645 \| Phytophthora parasitica INRA-310 ubiquitin conjugation factor E4 (1068 aa) |
| 914 | 1632 | PPTG_16133 | 98,49 | 45 | 38 | 39 | 30 | 7,63E+07 | 2,41E+08 | 3,17E+08 | 8 | 8 | 7 | 10 | 7 | Oxidation (M) | 12453 | \| PPTG_16133 \| Phytophthora parasitica INRA-310 50S ribosomal protein L35Ae (112 aa) |
| 2539 | 2371 | PPTG_03207 | 98,47 | 32 | 12 | 22 | 22 | 7,98E+04 | 1,07E+06 | 3,02E+06 | 3 | 3 | 1 | 2 | 2 | Carbamidomethylation | 18183 | \| PPTG_03207 \| Phytophthora parasitica INRA-310 hypothetical protein (162 aa) |
| 1790 | 1647 | PPTG_07988 | 98,27 | 32 | 0 | 32 | 32 | | 5,65E+06 | 6,57E+06 | 5 | 5 | 0 | 5 | 5 | Carbamidomethylation | 23407 | \| PPTG_07988 \| Phytophthora parasitica INRA-310 hypothetical protein (213 aa) |
| 2238 | 2380 | PPTG_09285 | 98,19 | 49 | 31 | 49 | 31 | | 1,46E+05 | | 3 | 1 | 2 | 3 | 2 | | 8717 | \| PPTG_09285 \| Phytophthora parasitica INRA-310 hypothetical protein (87 aa) |
| 2453 | 2365 | PPTG_08139 | 98,18 | 8 | 0 | 8 | 4 | | 1,02E+06 | 6,20E+05 | 3 | 3 | 0 | 3 | 2 | Carbamidomethylation | 60544 | \| PPTG_08139 \| Phytophthora parasitica INRA-310 hypothetical protein (572 aa) |
| 1597 | 1724 | PPTG_09351 | 98,08 | 12 | 0 | 10 | 12 | | 9,29E+05 | 1,10E+06 | 5 | 5 | 0 | 5 | 7 | Oxidation (M); Deamidation (NQ) | 82687 | \| PPTG_09351 \| Phytophthora parasitica INRA-310 hypothetical protein (754 aa) |
| 1328 | 1649 | PPTG_11597 | 98,06 | 16 | 7 | 14 | 16 | 1,03E+06 | 8,43E+06 | 1,19E+07 | 5 | 5 | 3 | 6 | 7 | | 44521 | \| PPTG_11597 \| Phytophthora parasitica INRA-310 hypothetical protein (403 aa) |
| 1401 | 1968 | PPTG_08552 | 98,05 | 47 | 35 | 35 | 47 | 3,92E+06 | 5,82E+06 | 5,47E+07 | 4 | 4 | 3 | 4 | 8 | Oxidation (M); Deamidation (NQ) | 14785 | \| PPTG_08552 \| Phytophthora parasitica INRA-310 hypothetical protein (130 aa) |
| 1961 | 2657 | PPTG_12308 | 98,04 | 17 | 6 | 13 | 17 | 1,63E+05 | 6,31E+06 | 4,21E+06 | 3 | 3 | 1 | 4 | 4 | Oxidation (M); Carbamidomethylation | 40826 | \| PPTG_12308 \| Phytophthora parasitica INRA-310 elicitin-like protein (388 aa) |
| 1222 | 1281 | PPTG_09490 | 98,02 | 11 | 4 | 9 | 11 | 1,85E+06 | 5,21E+06 | 2,61E+07 | 6 | 5 | 2 | 6 | 9 | Oxidation (M); Carbamidomethylation | 61962 | \| PPTG_09490 \| Phytophthora parasitica INRA-310 major facilitator superfamily transporter (MFS) (564 aa) |
| 1659 | 1490 | PPTG_13805 | 97,89 | 20 | 0 | 15 | 20 | | 2,32E+06 | 1,32E+06 | 6 | 6 | 0 | 5 | 6 | | 46526 | \| PPTG_13805 \| Phytophthora parasitica INRA-310 hypothetical protein (434 aa) |
| 1348 | 1274 | PPTG_18421 | 97,86 | 46 | 3 | 12 | 46 | 1,89E+05 | 1,29E+06 | 1,12E+07 | 9 | 9 | 1 | 3 | 12 | Oxidation (M); Carbamidomethylation; Acetylation (N-term) | 28518 | \| PPTG_18421 \| Phytophthora parasitica INRA-310 C2 domain-containing protein (251 aa) |
| 1951 | 1714 | PPTG_08806 | 97,74 | 27 | 3 | 16 | 27 | 4,28E+05 | 1,40E+06 | 6,79E+06 | 5 | 5 | 1 | 3 | 5 | Carbamidomethylation | 33349 | \| PPTG_08806 \| Phytophthora parasitica INRA-310 axin interactor (298 aa) |
| 1920 | 3529 | PPTG_08778 | 97,72 | 41 | 0 | 41 | 20 | | 8,01E+06 | 6,00E+05 | 2 | 2 | 0 | 7 | 2 | Deamidation (NQ) | 14874 | \| PPTG_08778 \| Phytophthora parasitica INRA-310 hypothetical protein (136 aa) |
| 1585 | 1476 | PPTG_17627 | 97,65 | 22 | 7 | 4 | 20 | 8,57E+05 | 5,05E+05 | 6,83E+06 | 7 | 7 | 3 | 2 | 7 | Carbamidomethylation; Acetylation (N-term) | 43861 | \| PPTG_17627 \| Phytophthora parasitica INRA-310 hypothetical protein (388 aa) |
| 1429 | 1751 | PPTG_06691 | 97,63 | 30 | 19 | 23 | 30 | 5,57E+05 | 2,07E+06 | 1,26E+07 | 5 | 5 | 3 | 4 | 7 | Oxidation (M); Carbamidomethylation | 22452 | \| PPTG_06691 \| Phytophthora parasitica INRA-310 hypothetical protein (202 aa) |
| 1904 | 2327 | PPTG_19485 | 97,56 | 12 | 4 | 12 | 12 | 1,98E+05 | 4,64E+06 | 5,32E+06 | 3 | 3 | 1 | 3 | 5 | Carbamidomethylation; Dehydration; Deamidation (NQ) | 35652 | \| PPTG_19485 \| Phytophthora parasitica INRA-310 carbohydrate-binding protein (342 aa) |
| 2116 | 1873 | PPTG_03752 | 97,55 | 9 | 1 | 6 | 9 | 5,79E+04 | 3,92E+05 | 1,81E+06 | 4 | 4 | 1 | 2 | 4 | | 58869 | \| PPTG_03752 \| Phytophthora parasitica INRA-310 hypothetical protein (537 aa) |
| 2325 | 1959 | PPTG_10872 | 97,54 | 16 | 4 | 16 | 2 0 | | 1,60E+06 | 3,42E+05 | 4 | 4 | 1 | 4 | 1 | | 47029 | \| PPTG_10872 \| Phytophthora parasitica INRA-310 3-oxoacyl-[acyl-carrier-protein] synthase (446 aa) |
| 1511 | 1494 | PPTG_19849 | 97,49 | 11 | 1 | 9 | 8 | 1,19E+05 | 2,39E+06 | 4,25E+06 | 8 | 8 | 1 | 6 | 6 | | 81197 | \| PPTG_19849 \| Phytophthora parasitica INRA-310 vacuolar protein sorting-associated protein 18 (723 aa) |
| 1417 | 1296 | PPTG_15981 | 97,46 | 38 | 10 | 27 | 26 | 2,13E+06 | 9,09E+06 | 9,25E+06 | 7 | 7 | 3 | 6 | 5 | Oxidation (M); Carbamidomethylation; Deamidation (NQ); Acetylation (N-term); Amidation | 29248 | \| PPTG_15981 \| Phytophthora parasitica INRA-310 glycoside hydrolase (280 aa) |
| 1623 | 1989 | PPTG_08112 | 97,43 | 27 | 7 | 14 | 27 | 3,15E+05 | 4,00E+06 | 1,37E+07 | 6 | 6 | 1 | 4 | 7 | Dehydration | 23599 | \| PPTG_08112 \| Phytophthora parasitica INRA-310 hypothetical protein (208 aa) |
| 1711 | 1633 | PPTG_00696 | 97,38 | 15 | 2 | 11 | 12 | 5,08E+04 | 2,66E+06 | 4,29E+06 | 6 | 6 | 1 | 5 | 5 | | 62005 | \| PPTG_00696 \| Phytophthora parasitica INRA-310 malonyl-CoA decarboxylase (MCD) (551 aa) |
| 1797 | 2036 | PPTG_20500 | 97,38 | 19 | 5 | 16 | 19 | 1,51E+05 | 3,19E+06 | 1,08E+07 | 4 | 4 | 1 | 4 | 5 | Oxidation (M) | 29342 | \| PPTG_20500 \| Phytophthora parasitica INRA-310 hypothetical protein (262 aa) |
| 2388 | 2273 | PPTG_13879 | 97,33 | 21 | 8 | 14 | 14 | 4,35E+05 | 5,89E+06 | 2,10E+06 | 3 | 3 | 1 | 3 | 2 | Carbamidomethylation | 33754 | \| PPTG_13879 \| Phytophthora parasitica INRA-310 hypothetical protein (323 aa) |
| 1543 | 2009 | PPTG_13652 | 97,3 | 41 | 14 | 41 | 41 | 4,48E+05 | 7,08E+06 | 1,07E+07 | 4 | 4 | 1 | 6 | 6 | Carbamidomethylation; Deamidation (NQ) | 19506 | \| PPTG_13652 \| Phytophthora parasitica INRA-310 actin-like protein 2/3 complex subunit 3 (175 aa) |
| 1917 | 2161 | PPTG_09243 | 97,27 | 22 | 3 | 8 | 22 | 9,37E+05 | 2,47E+06 | 1,97E+07 | 4 | 4 | 1 | 2 | 6 | Oxidation (M) | 27746 | \| PPTG_09243 \| Phytophthora parasitica INRA-310 hypothetical protein (255 aa) |
| 2329 | 2334 | PPTG_04627 | 97,25 | 9 | 0 | 3 | 9 | | 5,59E+04 | 1,60E+06 | 3 | 3 | 0 | 1 | 5 | Oxidation (M); Carbamidomethylation; Acetylation (N-term); Carbamylation | 70714 | \| PPTG_04627 \| Phytophthora parasitica INRA-310 dicarboxylate/Amino Acid:Cation (Na or H) Symporter (DAACS) family (660 aa) |
| 1611 | 1475 | PPTG_00740 | 97,21 | 24 | 7 | 15 | 20 | 1,48E+05 | 1,24E+06 | 4,20E+06 | 6 | 6 | 2 | 4 | 6 | Carbamidomethylation; Acetylation (N-term) | 42498 | \| PPTG_00740 \| Phytophthora parasitica INRA-310 hypothetical protein (379 aa) |
| 1654 | 1403 | PPTG_19141 | 97,19 | 8 | 1 | 8 | 4 | 3,38E+05 | 3,16E+06 | 3,77E+06 | 7 | 7 | 1 | 7 | 3 | | 122701 | \| PPTG_19141 \| Phytophthora parasitica INRA-310 hypothetical protein (1098 aa) |
| 1399 | 1668 | PPTG_11217 | 97,14 | 16 | 7 | 10 | 16 | 8,23E+05 | 1,94E+06 | 1,02E+07 | 5 | 5 | 2 | 4 | 9 | Deamidation (NQ) | 43709 | \| PPTG_11217 \| Phytophthora parasitica INRA-310 hypothetical protein (390 aa) |
| 1821 | 1737 | PPTG_17150 | 97,13 | 16 | 2 | 14 | 13 | 8,62E+05 | 4,15E+06 | 6,70E+06 | 5 | 5 | 1 | 4 | 5 | Oxidation (M) | 48889 | \| PPTG_17150 \| Phytophthora parasitica INRA-310 hypothetical protein (438 aa) |
| 1896 | 1411 | PPTG_10791 | 97,13 | 11 | 1 | 3 | 10 | 1,77E+05 | 2,66E+05 | 3,04E+06 | 7 | 7 | 1 | 2 | 6 | Carbamidomethylation | 66279 | \| PPTG_10791 \| Phytophthora parasitica INRA-310 sphingosine-1-phosphate lyase (608 aa) |
| 1451 | 1984 | PPTG_12838 | 97,12 | 19 | 12 | 19 | 12 | 2,61E+06 | 3,33E+07 | 2,25E+07 | 6 | 6 | 3 | 7 | 4 | Carbamidomethylation | 29635 | \| PPTG_12838 \| Phytophthora parasitica INRA-310 hypothetical protein (263 aa) |
| 1601 | 1362 | PPTG_18780 | 97,07 | 19 | 6 | 2 | 15 | 2,67E+05 | | 4,02E+06 | 9 | 8 | 3 | 1 | 8 | Oxidation (M); Carbamidomethylation | 67402 | \| PPTG_18780 \| Phytophthora parasitica INRA-310 ATP-binding Cassette (ABC) superfamily (617 aa) |
| 1343 | 1523 | PPTG_12040 | 96,93 | 16 | 3 | 14 | 15 | 3,05E+04 | 1,77E+06 | 1,70E+06 | 7 | 7 | 1 | 7 | 8 | Carbamidomethylation | 61167 | \| PPTG_12040 \| Phytophthora parasitica INRA-310 choline/Carnitine O-acyltransferase (540 aa) |
| 1737 | 1971 | PPTG_13998 | 96,89 | 28 | 22 | 28 | 22 | 2,91E+06 | 4,00E+06 | 1,88E+07 | 4 | 4 | 3 | 4 | 4 | Carbamidomethylation | 18621 | \| PPTG_13998 \| Phytophthora parasitica INRA-310 hypothetical protein (163 aa) |
| 1737 | 1972 | PPTG_13998 | 96,89 | 28 | 22 | 28 | 22 | 2,91E+06 | 4,00E+06 | 1,88E+07 | 4 | 4 | 3 | 4 | 4 | Carbamidomethylation | 18621 | \| PPTG_13998 \| Phytophthora parasitica INRA-310 hypothetical protein (163 aa) |
| 1839 | 1695 | PPTG_15714 | 96,89 | 26 | 0 | 11 | 26 | | 1,81E+06 | 8,73E+06 | 6 | 6 | 0 | 3 | 7 | | 35312 | \| PPTG_15714 \| Phytophthora parasitica INRA-310 arabinan endo-1 5-alpha-L-arabinosidase A (323 aa) |
| 1211 | 1489 | PPTG_10108 | 96,85 | 3 | 0 | 2 | 3 | | 1,66E+06 | 3,04E+06 | 11 | 11 | 0 | 7 | 10 | Carbamidomethylation | 420923 | \| PPTG_10108 \| Phytophthora parasitica INRA-310 hypothetical protein (3816 aa) |
| 2312 | 1944 | PPTG_15380 | 96,83 | 40 | 0 | 27 | 40 | | 1,97E+06 | 4,05E+06 | 4 | 4 | 0 | 2 | 4 | | 15444 | \| PPTG_15380 \| Phytophthora parasitica INRA-310 50S ribosomal protein L11 (145 aa) |
| 1234 | 1496 | PPTG_02467 | 96,75 | 11 | 1 | 10 | 5 | 3,79E+04 | 3,05E+06 | 3,20E+06 | 11 | 11 | 1 | 10 | 6 | Carbamidomethylation | 123051 | \| PPTG_02467 \| Phytophthora parasitica INRA-310 MMS19 nucleotide excision repair protein (1137 aa) |
| 1406 | 1896 | PPTG_14599 | 96,73 | 9 | 4 | 4 | 9 0 | | 7,22E+04 | 5,18E+05 | 5 | 2 | 3 | 3 | 8 | Oxidation (M) | 69631 | \| PPTG_14599 \| Phytophthora parasitica INRA-310 ATP-binding Cassette (ABC) superfamily (621 aa) |
| 1629 | 1788 | PPTG_10420 | 96,69 | 26 | 14 | 23 | 26 | 3,31E+05 | 2,06E+06 | 2,13E+06 | 5 | 5 | 3 | 4 | 5 | | 25652 | \| PPTG_10420 \| Phytophthora parasitica INRA-310 YggS family pyridoxal phosphate enzyme (235 aa) |
| 2015 | 2284 | PPTG_13049 | 96,63 | 13 | 7 | 13 | 13 | 9,57E+05 | 3,25E+06 | 9,95E+06 | 3 | 3 | 2 | 3 | 3 | Carbamidomethylation | 36078 | \| PPTG_13049 \| Phytophthora parasitica INRA-310 serine protease family S33 (328 aa) |
| 1883 | 2238 | PPTG_02109 | 96,63 | 11 | 11 | 11 | 11 | 4,54E+05 | 4,03E+06 | 3,12E+06 | 3 | 3 | 3 | 3 | 3 | Deamidation (NQ) | 47228 | \| PPTG_02109 \| Phytophthora parasitica INRA-310 hypothetical protein (431 aa) |
| 2201 | 1290 | PPTG_14266 | 96,57 | 7 | 0 | 0 | 7 | | | 1,60E+06 | 7 | 7 | 0 | 0 | 7 | Carbamidomethylation | 174553 | \| PPTG_14266 \| Phytophthora parasitica INRA-310 fibronectin type III domain-containing protein (1637 aa) |
| 1481 | 1417 | PPTG_05816 | 96,57 | 6 | 1 | 6 | 4 0 | | 1,67E+06 | 1,51E+06 | 7 | 7 | 1 | 7 | 5 | Carbamidomethylation | 148067 | \| PPTG_05816 \| Phytophthora parasitica INRA-310 SpoU rRNA methylase family protein (1315 aa) |
| 2065 | 1759 | PPTG_14505 | 96,55 | 17 | 0 | 5 | 17 | | 4,00E+06 | 1,68E+07 | 5 | 5 | 0 | 2 | 6 | Dehydration; Deamidation (NQ) | 43027 | \| PPTG_14505 \| Phytophthora parasitica INRA-310 hypothetical protein (388 aa) |
| 1767 | 1615 | PPTG_17541 | 96,48 | 8 | 4 | 4 | 8 | 2,79E+05 | 6,24E+05 | 5,04E+06 | 5 | 5 | 2 | 2 | 6 | Carbamidomethylation | 79201 | \| PPTG_17541 \| Phytophthora parasitica INRA-310 hypothetical protein (706 aa) |
| 1545 | 1478 | PPTG_13304 | 96,47 | 10 | 1 | 4 | 10 | 3,50E+05 | 2,68E+06 | 7,87E+06 | 6 | 6 | 1 | 3 | 9 | Oxidation (M); Deamidation (NQ) | 83585 | \| PPTG_13304 \| Phytophthora parasitica INRA-310 autophagy-like protein (742 aa) |
| 1755 | 1796 | PPTG_00372 | 96,45 | 13 | 5 | 13 | 13 | 1,11E+05 | 1,22E+06 | 1,12E+06 | 4 | 2 | 2 | 4 | 4 | | 39580 | \| PPTG_00372 \| Phytophthora parasitica INRA-310 voltage-gated potassium channel subunit beta (354 aa) |
| 1547 | 1832 | PPTG_15938 | 96,44 | 6 | 0 | 6 | 5 | | 3,26E+05 | | 6 | 1 | 0 | 8 | 5 | Oxidation (M) | 84946 | \| PPTG_15938 \| Phytophthora parasitica INRA-310 beta glucosidase (782 aa) |
| 1696 | 1637 | PPTG_00084 | 96,37 | 35 | 18 | 29 | 26 | 3,45E+05 | 3,53E+06 | 4,00E+06 | 5 | 5 | 2 | 5 | 4 | | 16556 | \| PPTG_00084 \| Phytophthora parasitica INRA-310 endoribonuclease L-PSP (161 aa) |
| 1578 | 1557 | PPTG_10800 | 96,37 | 11 | 0 | 6 | 9 | | 3,64E+06 | 3,85E+06 | 8 | 7 | 0 | 5 | 7 | Oxidation (M); Carbamidomethylation; Carbamylation | 106972 | \| PPTG_10800 \| Phytophthora parasitica INRA-310 haustorium-specific membrane protein (1017 aa) |
| 1390 | 1492 | PPTG_14115 | 96,31 | 18 | 8 | 12 | 18 | 1,03E+06 | 6,92E+06 | 9,50E+06 | 6 | 6 | 3 | 4 | 8 | Oxidation (M); Carbamidomethylation; Dehydration | 35771 | \| PPTG_14115 \| Phytophthora parasitica INRA-310 dihydroxyacetone kinase (348 aa) |
| 733 | 1899 | PPTG_08955 | 96,3 | 12 | 7 | 12 | 12 | 1,41E+08 | 5,00E+08 | 1,06E+09 | 5 | 5 | 6 | 10 | 15 | Dehydration; Carbamylation | 32792 | \| PPTG_08955 \| Phytophthora parasitica INRA-310 hypothetical protein (308 aa) |
| 1720 | 1541 | PPTG_00772 | 96,29 | 17 | 0 | 8 | 17 | | 6,07E+04 | 2,01E+06 | 6 | 5 | 0 | 2 | 9 | Carbamidomethylation; Deamidation (NQ) | 53353 | \| PPTG_00772 \| Phytophthora parasitica INRA-310 serine palmitoyltransferase (485 aa) |
| 1648 | 1775 | PPTG_20306 | 96,21 | 18 | 4 | 18 | 15 | 3,54E+05 | 5,12E+06 | 5,51E+06 | 6 | 6 | 1 | 6 | 4 | | 36914 | \| PPTG_20306 \| Phytophthora parasitica INRA-310 S-adenosylmethionine synthase 2 (336 aa) |
| 1648 | 1776 | PPTG_09226 | 96,21 | 15 | 4 | 15 | 13 | 3,54E+05 | 5,12E+06 | 5,51E+06 | 6 | 6 | 1 | 6 | 4 | | 42934 | \| PPTG_09226 \| Phytophthora parasitica INRA-310 S-adenosylmethionine synthase 2 (392 aa) |
| 1531 | 1337 | PPTG_07219 | 96,19 | 10 | 3 | 5 | 9 | 1,49E+05 | 1,19E+06 | 2,50E+06 | 8 | 8 | 2 | 4 | 7 | | 94509 | \| PPTG_07219 \| Phytophthora parasitica INRA-310 hypothetical protein (859 aa) |
| 1500 | 1409 | PPTG_07731 | 96,12 | 27 | 0 | 13 | 27 | | 2,17E+07 | 3,52E+07 | 9 | 9 | 0 | 4 | 9 | Carbamidomethylation | 39177 | \| PPTG_07731 \| Phytophthora parasitica INRA-310 calcium-binding protein (347 aa) |
| 1795 | 1656 | PPTG_14399 | 96,06 | 22 | 7 | 13 | 19 | 4,95E+05 | 2,38E+06 | 5,90E+06 | 5 | 5 | 2 | 4 | 4 | Carbamidomethylation | 43306 | \| PPTG_14399 \| Phytophthora parasitica INRA-310 actin-like protein 2/3 complex subunit (392 aa) |
| 1820 | 1731 | PPTG_19390 | 96,06 | 18 | 4 | 14 | 18 | 1,91E+04 | 1,68E+06 | 3,36E+06 | 5 | 5 | 1 | 4 | 5 | Acetylation (N-term) | 39322 | \| PPTG_19390 \| Phytophthora parasitica INRA-310 hypothetical protein (348 aa) |
| 1766 | 1376 | PPTG_12460 | 95,99 | 11 | 2 | 1 | 11 | | | 2,37E+06 | 7 | 5 | 2 | 1 | 7 | Carbamidomethylation | 84425 | \| PPTG_12460 \| Phytophthora parasitica INRA-310 hypothetical protein (763 aa) |
| 2180 | 2479 | PPTG_17441 | 95,91 | 25 | 0 | 25 | 16 | | 2,46E+06 | 2,93E+06 | 3 | 3 | 0 | 4 | 3 | Oxidation (M); Carbamidomethylation | 36270 | \| PPTG_17441 \| Phytophthora parasitica INRA-310 hypothetical protein (341 aa) |
| 1463 | 2212 | PPTG_09929 | 95,88 | 8 | 1 | 8 | 6 | | 2,89E+05 | | 5 | 1 | 1 | 8 | 5 | Carbamidomethylation | 70751 | \| PPTG_09929 \| Phytophthora parasitica INRA-310 glycoside hydrolase (637 aa) |
| 1613 | 1401 | PPTG_19561 | 95,79 | 8 | 0 | 6 | 6 | | 2,14E+05 | 4,99E+05 | 7 | 2 | 0 | 6 | 6 | Carbamidomethylation | 125917 | \| PPTG_19561 \| Phytophthora parasitica INRA-310 hypothetical protein (1109 aa) |
| 1663 | 1530 | PPTG_03846 | 95,79 | 13 | 2 | 10 | 8 | | 6,43E+05 | 2,69E+05 | 6 | 2 | 1 | 5 | 5 | Oxidation (M); Carbamidomethylation | 64540 | \| PPTG_03846 \| Phytophthora parasitica INRA-310 cell 5A endo-1 4-betaglucanase (583 aa) |
| 1304 | 1328 | PPTG_12006 | 95,79 | 13 | 1 | 12 | 9 | | 3,78E+06 | 3,69E+06 | 8 | 6 | 1 | 8 | 7 | Oxidation (M); Carbamidomethylation | 64488 | \| PPTG_12006 \| Phytophthora parasitica INRA-310 glycoside hydrolase (572 aa) |
| 1342 | 1449 | PPTG_10452 | 95,77 | 24 | 14 | 17 | 24 | 4,02E+05 | 1,80E+06 | 4,58E+06 | 6 | 4 | 3 | 6 | 7 | Oxidation (M); Carbamidomethylation | 34763 | \| PPTG_10452 \| Phytophthora parasitica INRA-310 3-hydroxyisobutyrate dehydrogenase (324 aa) |
| 1400 | 1697 | PPTG_15673 | 95,75 | 28 | 28 | 17 | 28 | 1,24E+06 | 7,94E+05 | 8,64E+06 | 5 | 5 | 5 | 3 | 7 | Deamidation (NQ) | 21722 | \| PPTG_15673 \| Phytophthora parasitica INRA-310 hypothetical protein (199 aa) |
| 1818 | 1497 | PPTG_04212 | 95,73 | 26 | 0 | 13 | 23 | | 1,07E+06 | 1,25E+06 | 7 | 7 | 0 | 4 | 6 | | 39120 | \| PPTG_04212 \| Phytophthora parasitica INRA-310 hypothetical protein (352 aa) |
| 1596 | 1527 | PPTG_16429 | 95,71 | 17 | 2 | 12 | 11 | 4,04E+04 | 1,55E+06 | 2,63E+06 | 7 | 7 | 1 | 5 | 6 | Carbamidomethylation | 60633 | \| PPTG_16429 \| Phytophthora parasitica INRA-310 glutamate synthase (543 aa) |
| 1166 | 1289 | PPTG_00670 | 95,7 | 14 | 1 | 13 | 13 | 4,93E+05 | 4,56E+06 | 9,46E+06 | 8 | 8 | 1 | 7 | 10 | Oxidation (M); Carbamidomethylation; Deamidation (NQ) | 69817 | \| PPTG_00670 \| Phytophthora parasitica INRA-310 ATP-binding cassette sub-family F member 2 (615 aa) |
| 1398 | 1442 | PPTG_02802 | 95,66 | 24 | 5 | 16 | 24 | 2,27E+05 | 1,33E+06 | 3,53E+06 | 8 | 8 | 2 | 5 | 8 | Carbamidomethylation | 45775 | \| PPTG_02802 \| Phytophthora parasitica INRA-310 GTPase (408 aa) |
| 2319 | 2354 | PPTG_15512 | 95,62 | 15 | 0 | 15 | 15 | | 3,84E+06 | 3,95E+06 | 3 | 3 | 0 | 3 | 3 | Oxidation (M) | 34655 | \| PPTG_15512 \| Phytophthora parasitica INRA-310 SCP-like extracellular protein (335 aa) |
| 1497 | 1461 | PPTG_19264 | 95,58 | 9 | 4 | 4 | 9 | 3,60E+05 | 4,27E+05 | 4,43E+06 | 7 | 7 | 3 | 3 | 7 | Oxidation (M) | 90486 | \| PPTG_19264 \| Phytophthora parasitica INRA-310 choline/Carnitine O-acyltransferase (795 aa) |
| 1514 | 1762 | PPTG_11599 | 95,57 | 11 | 7 | 5 | 11 | 4,35E+05 | 9,00E+05 | 6,01E+06 | 5 | 5 | 3 | 2 | 8 | Oxidation (M) | 61542 | \| PPTG_11599 \| Phytophthora parasitica INRA-310 hypothetical protein (556 aa) |
| 1698 | 1876 | PPTG_17055 | 95,54 | 14 | 5 | 8 | 6 0 | | 3,57E+05 0 | | 6 | 4 | 2 | 5 | 4 | | 46325 | \| PPTG_17055 \| Phytophthora parasitica INRA-310 eukaryotic initiation factor 4A-III (407 aa) |
| 1278 | 2014 | PPTG_01568 | 95,41 | 15 | 8 | 13 | 10 | | 9,59E+04 | | 5 | 1 | 4 | 7 | 5 | Oxidation (M); Carbamylation | 46398 | \| PPTG_01568 \| Phytophthora parasitica INRA-310 aldehyde dehydrogenase (431 aa) |
| 1278 | 2017 | PPTG_20220 | 95,41 | 13 | 6 | 11 | 8 | | 9,59E+04 | | 5 | 1 | 4 | 7 | 5 | Oxidation (M); Carbamylation | 56198 | \| PPTG_20220 \| Phytophthora parasitica INRA-310 aldehyde dehydrogenase (521 aa) |
| 1685 | 1326 | PPTG_15791 | 95,4 | 22 | 11 | 3 | 22 | 4,24E+05 | 2,26E+05 | 4,74E+06 | 7 | 7 | 3 | 1 | 7 | | 40902 | \| PPTG_15791 \| Phytophthora parasitica INRA-310 hypothetical protein (368 aa) |
| 1927 | 2061 | PPTG_17695 | 95,36 | 23 | 8 | 10 | 18 | 3,05E+04 | 9,69E+05 | 2,67E+06 | 6 | 6 | 2 | 2 | 5 | | 31453 | \| PPTG_17695 \| Phytophthora parasitica INRA-310 urease accessory protein ureG (288 aa) |
| 1970 | 1535 | PPTG_02606 | 95,36 | 7 | 2 | 0 | 6 | 1,14E+05 | | 1,59E+06 | 7 | 7 | 2 | 0 | 6 | Carbamidomethylation | 125358 | \| PPTG_02606 \| Phytophthora parasitica INRA-310 hypothetical protein (1144 aa) |
| 2080 | 1457 | PPTG_06584 | 95,33 | 13 | 1 | 1 | 12 | | 3,63E+05 | 4,89E+06 | 7 | 3 | 1 | 1 | 6 | Carbamidomethylation | 74458 | \| PPTG_06584 \| Phytophthora parasitica INRA-310 transmembrane protein (686 aa) |
| 2218 | 1453 | PPTG_18560 | 95,33 | 14 | 0 | 0 | 14 | | | 4,62E+06 | 6 | 6 | 0 | 0 | 7 | Carbamidomethylation | 59087 | \| PPTG_18560 \| Phytophthora parasitica INRA-310 calcium-dependent protein kinase (524 aa) |
| 1269 | 1485 | PPTG_15292 | 95,15 | 40 | 18 | 19 | 40 | 1,21E+06 | 1,65E+06 | 1,54E+07 | 6 | 6 | 3 | 5 | 9 | Oxidation (M); Dehydration | 21672 | \| PPTG_15292 \| Phytophthora parasitica INRA-310 Rab5 family GTPase (201 aa) |
| 1544 | 1293 | PPTG_02140 | 95,03 | 18 | 7 | 4 | 18 | 4,51E+05 | 6,64E+05 | 7,37E+06 | 7 | 7 | 3 | 2 | 8 | Carbamidomethylation | 57037 | \| PPTG_02140 \| Phytophthora parasitica INRA-310 aldehyde dehydrogenase (515 aa) |
| 2196 | 2612 | PPTG_16975 | 94,99 | 21 | 9 | 9 | 21 | 9,92E+05 | 4,20E+06 | 8,77E+06 | 3 | 3 | 1 | 2 | 4 | | 21477 | \| PPTG_16975 \| Phytophthora parasitica INRA-310 hypothetical protein (193 aa) |
| 1503 | 1544 | PPTG_00182 | 94,98 | 6 | 1 | 4 | 4 | 1,03E+05 | 1,15E+06 | 1,66E+06 | 8 | 8 | 1 | 6 | 6 | Carbamidomethylation | 187882 | \| PPTG_00182 \| Phytophthora parasitica INRA-310 adenylate kinase (1727 aa) |
| 1516 | 1522 | PPTG_05184 | 94,96 | 18 | 4 | 18 | 15 | 3,73E+05 | 3,50E+06 | 3,81E+06 | 6 | 6 | 2 | 6 | 5 | | 45193 | \| PPTG_05184 \| Phytophthora parasitica INRA-310 hypothetical protein (407 aa) |

| | | | | | | | | | | | | | | | | | |
|---|---|---|---|---|---|---|---|---|---|---|---|---|---|---|---|---|---|
| 1462 | 1969 | PPTG_07610 | 94,93 | 40 | 18 | 39 | 40 | 7,90E+05 | 7,82E+06 | 1,16E+07 | 6 | 6 | 2 | 6 | 6 | | 13477 \| PPTG_07610 \| Phytophthora parasitica INRA-310 hypothetical protein (120 aa) |
| 1828 | 1992 | PPTG_02041 | 94,87 | 47 | 12 | 25 | 47 | 4,86E+04 | 9,27E+05 | 1,39E+06 | 4 | 4 | 1 | 3 | 6 | Oxidation (M); Carbamidomethylation | 14468 \| PPTG_02041 \| Phytophthora parasitica INRA-310 hypothetical protein (134 aa) |
| 1885 | 1482 | PPTG_15122 | 94,8 | 11 | 1 | 2 | 11 | 5,93E+04 | 9,90E+04 | 2,57E+06 | 7 | 7 | 1 | 1 | 7 | | 92142 \| PPTG_15122 \| Phytophthora parasitica INRA-310 hypothetical protein (835 aa) |
| 2630 | 3341 | PPTG_02053 | 94,75 | 31 | 0 | 31 | 16 | | 2,24E+06 | 7,81E+05 | 2 | 2 | 0 | 4 | 1 | Carbamidomethylation | 13720 \| PPTG_02053 \| Phytophthora parasitica INRA-310 protease inhibitor EpiC3 (128 aa) |
| 1590 | 2206 | PPTG_05357 | 94,75 | 20 | 12 | 16 | 17 | 8,03E+05 | 2,81E+06 | 8,09E+06 | 4 | 4 | 3 | 4 | 5 | Oxidation (M); Carbamidomethylation; Deamidation (NQ); Carbamylation | 29029 \| PPTG_05357 \| Phytophthora parasitica INRA-310 hypothetical protein (267 aa) |
| 1683 | 2022 | PPTG_19266 | 94,75 | 55 | 20 | 55 | 35 | 1,82E+05 | 2,95E+06 | 2,25E+06 | 4 | 4 | 1 | 7 | 3 | Oxidation (M) | 10789 \| PPTG_19266 \| Phytophthora parasitica INRA-310 U6 snRNA-associated Sm-like protein LSm2 (97 aa) |
| 1681 | 1445 | PPTG_07997 | 94,75 | 38 | 5 | 24 | 38 | 9,82E+05 | 9,44E+06 | 1,57E+07 | 6 | 6 | 1 | 4 | 6 | Oxidation (M) | 22046 \| PPTG_07997 \| Phytophthora parasitica INRA-310 flagellar adenylate kinase (198 aa) |
| 2106 | 2583 | PPTG_18184 | 94,74 | 45 | 12 | 45 | 45 | 7,77E+05 | 8,03E+06 | 1,48E+07 | 3 | 3 | 1 | 4 | 3 | Carbamidomethylation | 10366 \| PPTG_18184 \| Phytophthora parasitica INRA-310 dynein light chain 2 cytoplasmic (90 aa) |
| 1436 | 1295 | PPTG_00862 | 94,64 | 17 | 0 | 15 | 12 | | 3,74E+06 | 3,16E+06 | 8 | 8 | 0 | 8 | 6 | Oxidation (M); Carbamidomethylation | 55555 \| PPTG_00862 \| Phytophthora parasitica INRA-310 anthranilate synthase component I (505 aa) |
| 2444 | 2505 | PPTG_00978 | 94,64 | 9 | 0 | 0 | 9 | | | 2,11E+06 | 4 | 4 | 0 | 0 | 5 | Deamidation (NQ) | 67768 \| PPTG_00978 \| Phytophthora parasitica INRA-310 homeodomain transcription factor (609 aa) |
| 1603 | 1472 | PPTG_11046 | 94,62 | 11 | 1 | 4 | 9 | 5,82E+04 | 5,69E+05 | 6,59E+06 | 8 | 8 | 1 | 3 | 8 | Carbamidomethylation | 84243 \| PPTG_11046 \| Phytophthora parasitica INRA-310 H -translocating Pyrophosphatase (H -PPase) family (802 aa) |
| 1624 | 1426 | PPTG_14666 | 94,59 | 16 | 2 | 9 | 16 | 3,67E+05 | 1,27E+06 | 1,41E+06 | 7 | 7 | 1 | 4 | 7 | Carbamidomethylation | 56984 \| PPTG_14666 \| Phytophthora parasitica INRA-310 hypothetical protein (530 aa) |
| 1230 | 1486 | PPTG_05410 | 94,57 | 9 | 1 | 7 | 8 | 4,40E+04 | 2,31E+06 | 3,87E+06 | 10 | 10 | 1 | 7 | 9 | | 141738 \| PPTG_05410 \| Phytophthora parasitica INRA-310 hypothetical protein (1258 aa) |
| 1972 | 1257 | PPTG_15095 | 94,47 | 8 | 0 | 0 | 8 | | | 1,84E+06 | 8 | 8 | 0 | 0 | 8 | Carbamidomethylation | 207803 \| PPTG_15095 \| Phytophthora parasitica INRA-310 hypothetical protein (1925 aa) |
| 1934 | 1914 | PPTG_06253 | 94,47 | 19 | 4 | 19 | 19 | 1,53E+05 | 5,38E+06 | 3,83E+06 | 4 | 3 | 1 | 4 | 4 | Carbamidomethylation | 31573 \| PPTG_06253 \| Phytophthora parasitica INRA-310 hypothetical protein (304 aa) |
| 1337 | 3199 | PPTG_08951 | 94,47 | 15 | 15 | 15 | 15 | 1,75E+07 | 8,16E+07 | 1,94E+08 | 2 | 2 | 3 | 5 | 8 | Carbamidomethylation; Carbamylation | 18564 \| PPTG_08951 \| Phytophthora parasitica INRA-310 hypothetical protein (184 aa) |
| 1133 | 1402 | PPTG_18906 | 94,37 | 43 | 17 | 43 | 36 | 1,63E+06 | 1,43E+07 | 1,77E+07 | 8 | 8 | 4 | 8 | 7 | Oxidation (M); Carbamidomethylation; Acetylation (N-term) | 19780 \| PPTG_18906 \| Phytophthora parasitica INRA-310 actin-like protein 2/3 complex subunit 4 (170 aa) |
| 1065 | 2405 | PPTG_12754 | 94,35 | 54 | 38 | 54 | 54 | 5,04E+07 | 2,05E+08 | 5,13E+08 | 4 | 4 | 4 | 7 | 10 | Carbamidomethylation; Deamidation (NQ) | 8084 \| PPTG_12754 \| Phytophthora parasitica INRA-310 60S ribosomal protein L38 (70 aa) |
| 2514 | 1897 | PPTG_16729 | 94,33 | 27 | 0 | 0 | 27 | | | 3,29E+06 | 4 | 4 | 0 | 0 | 5 | Oxidation (M) | 28780 \| PPTG_16729 \| Phytophthora parasitica INRA-310 hypothetical protein (256 aa) |
| 1383 | 1552 | PPTG_16515 | 94,31 | 37 | 7 | 26 | 21 | 3,71E+05 | 3,70E+06 | 8,30E+06 | 9 | 9 | 2 | 7 | 6 | Carbamidomethylation | 35130 \| PPTG_16515 \| Phytophthora parasitica INRA-310 hypothetical protein (306 aa) |
| 1955 | 1982 | PPTG_19377 | 94,31 | 21 | 10 | 16 | 21 | 7,71E+05 | 1,93E+06 | 7,13E+06 | 4 | 4 | 2 | 3 | 4 | Deamidation (NQ) | 25537 \| PPTG_19377 \| Phytophthora parasitica INRA-310 endoglucanase (242 aa) |
| 1873 | 1464 | PPTG_07340 | 94,3 | 12 | 0 | 7 | 9 | | 8,02E+05 | 9,12E+05 | 7 | 7 | 0 | 4 | 5 | | 78135 \| PPTG_07340 \| Phytophthora parasitica INRA-310 U3 small nucleolar RNA-associated protein (713 aa) |
| 1584 | 1439 | PPTG_12011 | 94,25 | 13 | 0 | 10 | 9 | | 6,07E+05 | 2,20E+06 | 7 | 4 | 0 | 6 | 6 | Oxidation (M); Carbamidomethylation | 65190 \| PPTG_12011 \| Phytophthora parasitica INRA-310 beta-glucosidase (580 aa) |
| 1470 | 1748 | PPTG_11488 | 94,22 | 15 | 4 | 4 | 15 | 3,01E+06 | 3,56E+06 | 2,74E+07 | 5 | 5 | 3 | 4 | 7 | Oxidation (M) | 53111 \| PPTG_11488 \| Phytophthora parasitica INRA-310 amino Acid/Auxin Permease (AAAP) family (489 aa) |
| 1923 | 1685 | PPTG_15215 | 94,21 | 30 | 0 | 30 | 24 | | 2,08E+06 | 1,52E+06 | 5 | 5 | 0 | 5 | 4 | Carbamidomethylation | 25155 \| PPTG_15215 \| Phytophthora parasitica INRA-310 hypoxanthine-guanine phosphoribosyltransferase (223 aa) |
| 2445 | 1513 | PPTG_14668 | 94,21 | 3 | 0 | 0 | 3 | | | 1,59E+06 | 5 | 5 | 0 | 0 | 5 | | 264916 \| PPTG_14668 \| Phytophthora parasitica INRA-310 hypothetical protein (2440 aa) |
| 1993 | 1477 | PPTG_15646 | 94,06 | 25 | 0 | 22 | 8 | | 2,23E+06 | 1,16E+06 | 7 | 7 | 0 | 6 | 2 | Oxidation (M); Carbamidomethylation | 38532 \| PPTG_15646 \| Phytophthora parasitica INRA-310 purine nucleoside phosphorylase 1 (353 aa) |
| 1299 | 2010 | PPTG_01099 | 94,02 | 62 | 18 | 37 | 54 | | | 3,74E+05 | 5 | 1 | 1 | 7 | 8 | Oxidation (M); Deamidation (NQ); Acetylation (N-term); Carbamylation | 11464 \| PPTG_01099 \| Phytophthora parasitica INRA-310 hypothetical protein (108 aa) |
| 1805 | 1699 | PPTG_16244 | 93,94 | 14 | 2 | 12 | 14 | 6,33E+04 | 2,98E+06 | 2,60E+06 | 5 | 5 | 1 | 4 | 5 | Carbamidomethylation | 50657 \| PPTG_16244 \| Phytophthora parasitica INRA-310 glucan 1 3-beta-glucosidase (461 aa) |
| 1756 | 1344 | PPTG_17988 | 93,88 | 12 | 0 | 9 | 9 | | 8,53E+05 | 2,71E+06 | 6 | 6 | 0 | 5 | 5 | Oxidation (M); Carbamidomethylation | 67085 \| PPTG_17988 \| Phytophthora parasitica INRA-310 ATP-dependent RNA helicase (604 aa) |
| 2007 | 1669 | PPTG_13915 | 93,85 | 29 | 5 | 9 | 29 | 1,28E+05 | 6,38E+05 | 6,07E+06 | 5 | 5 | 1 | 2 | 5 | Carbamidomethylation; Acetylation (N-term) | 33971 \| PPTG_13915 \| Phytophthora parasitica INRA-310 hypothetical protein (300 aa) |
| 1855 | 2037 | PPTG_09287 | 93,85 | 6 | 2 | 6 | 6 | 7,19E+04 | 3,84E+05 | 6,31E+05 | 4 | 4 | 1 | 4 | 4 | | 111869 \| PPTG_09287 \| Phytophthora parasitica INRA-310 hypothetical protein (1005 aa) |
| 1418 | 1359 | PPTG_17947 | 93,78 | 20 | 0 | 10 | 20 | | 1,62E+06 | 7,28E+06 | 8 | 8 | 0 | 4 | 10 | Oxidation (M) | 46823 \| PPTG_17947 \| Phytophthora parasitica INRA-310 hypothetical protein (422 aa) |
| 2338 | 1872 | PPTG_19100 | 93,65 | 28 | 0 | 20 | 16 | | 8,99E+05 | 1,12E+06 | 4 | 4 | 0 | 3 | 3 | Carbamidomethylation | 30427 \| PPTG_19100 \| Phytophthora parasitica INRA-310 hypothetical protein (280 aa) |
| 1869 | 1584 | PPTG_02022 | 93,64 | 7 | 1 | 5 | 6 | 3,18E+05 | 2,19E+06 | 3,04E+06 | 5 | 5 | 1 | 4 | 4 | | 94408 \| PPTG_02022 \| Phytophthora parasitica INRA-310 hypothetical protein (866 aa) |
| 1852 | 2127 | PPTG_09137 | 93,63 | 63 | 27 | 54 | 63 | 8,98E+04 | 1,76E+06 | 1,93E+06 | 4 | 4 | 1 | 4 | 5 | Carbamidomethylation; Deamidation (NQ); Acetylation (N-term) | 11315 \| PPTG_09137 \| Phytophthora parasitica INRA-310 hypothetical protein (105 aa) |
| 1826 | 1901 | PPTG_08802 | 93,61 | 20 | 8 | 20 | 17 | 8,63E+04 | 1,33E+06 | 1,82E+06 | 4 | 4 | 1 | 5 | 4 | Deamidation (NQ) | 39649 \| PPTG_08802 \| Phytophthora parasitica INRA-310 translation initiation factor eIF-2B subunit beta (364 aa) |
| 1771 | 1614 | PPTG_14263 | 93,59 | 6 | 0 | 5 | 5 | | 1,55E+06 | 4,11E+06 | 6 | 6 | 0 | 5 | 5 | | 129806 \| PPTG_14263 \| Phytophthora parasitica INRA-310 DNA-directed RNA polymerase III subunit polypeptide (1167 aa) |
| 1773 | 1466 | PPTG_13442 | 93,52 | 12 | 2 | 5 | 12 | 6,36E+04 | 5,03E+05 | 3,16E+06 | 6 | 6 | 1 | 3 | 6 | | 79772 \| PPTG_13442 \| Phytophthora parasitica INRA-310 bardet-Biedl syndrome 2 protein (724 aa) |
| 2140 | 1948 | PPTG_19961 | 93,38 | 19 | 0 | 6 | 19 | | 7,81E+05 | 3,70E+06 | 4 | 4 | 0 | 2 | 5 | Carbamidomethylation; Dehydration; Deamidation (NQ) | 34591 \| PPTG_19961 \| Phytophthora parasitica INRA-310 nuclear LIM factor interactor-interacting protein cleavage-specific form (308 aa) |
| 1322 | 1693 | PPTG_02504 | 93,37 | 22 | 12 | 14 | 22 | 2,60E+05 | 7,42E+05 | 3,20E+06 | 6 | 5 | 3 | 6 | 7 | | 33821 \| PPTG_02504 \| Phytophthora parasitica INRA-310 cyclin-dependent kinase 5 (298 aa) |
| 1898 | 1549 | PPTG_18187 | 93,34 | 14 | 0 | 8 | 14 | | 1,61E+05 | 4,08E+06 | 6 | 6 | 0 | 3 | 6 | Carbamidomethylation | 55699 \| PPTG_18187 \| Phytophthora parasitica INRA-310 hypothetical protein (498 aa) |
| 1819 | 1681 | PPTG_01622 | 93,25 | 37 | 7 | 30 | 37 | 2,26E+05 | 2,80E+06 | 5,28E+06 | 5 | 5 | 1 | 4 | 5 | Carbamidomethylation | 16140 \| PPTG_01622 \| Phytophthora parasitica INRA-310 hypothetical protein (146 aa) |
| 1430 | 1813 | PPTG_03756 | 93,24 | 67 | 22 | 42 | 67 | 3,54E+06 | 9,94E+06 | 6,18E+07 | 7 | 7 | 2 | 4 | 8 | Oxidation (M); Carbamidomethylation; Acetylation (N-term) | 9311 \| PPTG_03756 \| Phytophthora parasitica INRA-310 hypothetical protein (82 aa) |
| 1621 | 1736 | PPTG_01109 | 93,16 | 35 | 9 | 35 | 35 | 6,67E+04 | 1,58E+06 | 5,11E+06 | 6 | 6 | 1 | 5 | 6 | Carbamidomethylation | 20637 \| PPTG_01109 \| Phytophthora parasitica INRA-310 Ras family GTPase (185 aa) |
| 2135 | 2041 | PPTG_08903 | 93,16 | 17 | 0 | 6 | 17 | | 1,20E+06 | 5,95E+06 | 4 | 4 | 0 | 2 | 5 | | 43538 \| PPTG_08903 \| Phytophthora parasitica INRA-310 endoplasmic reticulum-Golgi intermediate compartment protein (394 aa) |
| 2045 | 2713 | PPTG_04071 | 93,11 | 10 | 2 | 10 | 8 | 7,26E+05 | 1,39E+07 | 9,98E+06 | 3 | 3 | 1 | 5 | 2 | Oxidation (M); Carbamidomethylation | 33468 \| PPTG_04071 \| Phytophthora parasitica INRA-310 hypothetical protein (304 aa) |
| 1617 | 2305 | PPTG_03643 | 92,99 | 24 | 15 | 24 | 24 | 6,59E+05 | 1,80E+06 | 2,81E+07 | 4 | 4 | 2 | 3 | 7 | Oxidation (M); Carbamidomethylation | 16971 \| PPTG_03643 \| Phytophthora parasitica INRA-310 hypothetical protein (146 aa) |
| 1537 | 1889 | PPTG_08070 | 92,98 | 15 | 2 | 13 | 14 | 1,75E+05 | 3,84E+06 | 2,97E+06 | 5 | 5 | 1 | 6 | 6 | Carbamidomethylation | 44277 \| PPTG_08070 \| Phytophthora parasitica INRA-310 hypothetical protein (424 aa) |
| 1475 | 1484 | PPTG_14762 | 92,89 | 12 | 5 | 11 | 12 | 5,16E+05 | 6,92E+06 | 3,74E+06 | 6 | 6 | 3 | 5 | 6 | | 69450 \| PPTG_14762 \| Phytophthora parasitica INRA-310 hypothetical protein (617 aa) |
| 1897 | 1536 | PPTG_03656 | 92,76 | 23 | 7 | 7 | 20 | 6,63E+05 | 5,53E+05 | 1,36E+07 | 6 | 6 | 2 | 2 | 5 | Carbamidomethylation | 29731 \| PPTG_03656 \| Phytophthora parasitica INRA-310 hypothetical protein (269 aa) |
| 2341 | 2011 | PPTG_00769 | 92,66 | 18 | 0 | 18 | 11 | | 3,93E+06 | 3,63E+06 | 4 | 4 | 0 | 4 | 2 | Carbamidomethylation | 26628 \| PPTG_00769 \| Phytophthora parasitica INRA-310 hypothetical protein (233 aa) |
| 1888 | 1319 | PPTG_00327 | 92,65 | 26 | 0 | 9 | 21 | | 4,34E+05 | 4,59E+06 | 8 | 8 | 0 | 2 | 7 | Carbamidomethylation | 39380 \| PPTG_00327 \| Phytophthora parasitica INRA-310 bardet-Biedl syndrome 5 protein (349 aa) |
| 2171 | 1787 | PPTG_09293 | 92,59 | 9 | 0 | 2 | 9 | | 1,73E+05 | 3,50E+06 | 5 | 5 | 0 | 1 | 6 | Carbamidomethylation; Acetylation (N-term) | 94808 \| PPTG_09293 \| Phytophthora parasitica INRA-310 hypothetical protein (862 aa) |
| 2064 | 1667 | PPTG_06680 | 92,55 | 27 | 4 | 23 | 8 | 3,64E+05 | 4,22E+06 | 3,25E+06 | 6 | 6 | 1 | 5 | 2 | Oxidation (M); Carbamidomethylation | 31850 \| PPTG_06680 \| Phytophthora parasitica INRA-310 cell differentiation protein RCD1 (287 aa) |
| 2019 | 1701 | PPTG_15961 | 92,52 | 14 | 0 | 9 | 14 | | 5,85E+05 | 2,66E+06 | 5 | 5 | 0 | 3 | 5 | Carbamidomethylation | 49163 \| PPTG_15961 \| Phytophthora parasitica INRA-310 sporangia induced Bardet-Biedl syndrome 4 protein (429 aa) |
| 1518 | 1665 | PPTG_16559 | 92,51 | 40 | 0 | 34 | 22 | | 3,96E+06 | 1,41E+06 | 7 | 7 | 0 | 8 | 5 | Oxidation (M); Deamidation (NQ) | 27107 \| PPTG_16559 \| Phytophthora parasitica INRA-310 oxidoreductase (259 aa) |
| 1467 | 1563 | PPTG_02308 | 92,49 | 18 | 8 | 14 | 18 | 2,70E+05 | 2,13E+06 | 4,38E+06 | 6 | 6 | 3 | 5 | 6 | Carbamidomethylation | 43594 \| PPTG_02308 \| Phytophthora parasitica INRA-310 hypothetical protein (381 aa) |
| 1066 | 1957 | PPTG_02141 | 92,41 | 34 | 12 | 12 | 34 | 8,89E+05 | 2,89E+06 | 2,19E+07 | 4 | 4 | 4 | 5 | 12 | Oxidation (M); Dehydration; Acetylation (N-term); Carbamylation | 23317 \| PPTG_02141 \| Phytophthora parasitica INRA-310 hypothetical protein (215 aa) |
| 1664 | 1545 | PPTG_13309 | 92,41 | 5 | 1 | 4 | 5 | 1,67E+05 | 1,68E+06 | 4,74E+06 | 5 | 5 | 1 | 4 | 6 | Oxidation (M) | 123697 \| PPTG_13309 \| Phytophthora parasitica INRA-310 hypothetical protein (1089 aa) |
| 1889 | 1789 | PPTG_08183 | 92,38 | 18 | 3 | 15 | 8 | 8,35E+04 | 1,33E+06 | 6,19E+05 | 6 | 6 | 1 | 5 | 3 | Carbamidomethylation | 46597 \| PPTG_08183 \| Phytophthora parasitica INRA-310 imidazolonepropionase (433 aa) |
| 1587 | 1725 | PPTG_07762 | 92,38 | 13 | 4 | 11 | 9 | 4,13E+04 | 1,96E+06 | 3,41E+06 | 6 | 6 | 2 | 6 | 4 | Carbamidomethylation | 55237 \| PPTG_07762 \| Phytophthora parasitica INRA-310 hypothetical protein (503 aa) |
| 1857 | 1441 | PPTG_11972 | 92,37 | 11 | 1 | 0 | 11 | 5,74E+04 | | 4,72E+06 | 8 | 8 | 1 | 0 | 8 | | 99780 \| PPTG_11972 \| Phytophthora parasitica INRA-310 protein kinase (895 aa) |
| 2138 | 2311 | PPTG_10779 | 92,35 | 11 | 4 | 11 | 11 | 2,23E+05 | 2,58E+06 | 2,37E+06 | 3 | 3 | 1 | 3 | 3 | | 36352 \| PPTG_10779 \| Phytophthora parasitica INRA-310 hypothetical protein (329 aa) |
| 1604 | 1491 | PPTG_20185 | 92,35 | 17 | 4 | 10 | 15 | 3,41E+05 | 4,88E+05 | 1,09E+07 | 7 | 7 | 2 | 4 | 6 | Carbamidomethylation | 47430 \| PPTG_20185 \| Phytophthora parasitica INRA-310 hypothetical protein (436 aa) |
| 1712 | 1657 | PPTG_00522 | 92,33 | 21 | 3 | 19 | 16 | 2,02E+05 | 3,44E+06 | 3,57E+06 | 6 | 6 | 1 | 5 | 5 | Oxidation (M); Carbamidomethylation | 43172 \| PPTG_00522 \| Phytophthora parasitica INRA-310 12-oxophytodienoate reductase (397 aa) |
| 1712 | 1659 | PPTG_00521 | 92,33 | 21 | 3 | 19 | 16 | 2,02E+05 | 3,44E+06 | 3,57E+06 | 6 | 6 | 1 | 5 | 5 | Oxidation (M); Carbamidomethylation | 43142 \| PPTG_00521 \| Phytophthora parasitica INRA-310 12-oxophytodienoate reductase (397 aa) |
| 1561 | 1586 | PPTG_01647 | 92,27 | 6 | 2 | 3 | 5 | 1,86E+05 | 2,57E+06 | 2,68E+06 | 7 | 7 | 2 | 4 | 6 | Carbamidomethylation | 129716 \| PPTG_01647 \| Phytophthora parasitica INRA-310 myosin-like protein (1153 aa) |
| 1561 | 1587 | PPTG_01647 | 92,27 | 6 | 2 | 3 | 5 | 1,86E+05 | 2,57E+06 | 2,68E+06 | 7 | 7 | 2 | 4 | 6 | Carbamidomethylation | 129845 \| PPTG_01647 \| Phytophthora parasitica INRA-310 myosin-like protein (1154 aa) |
| 1859 | 1301 | PPTG_13378 | 92,23 | 2 | 0 | 0 | 2 | | | 1,45E+06 | 9 | 9 | 0 | 0 | 9 | Carbamidomethylation; Deamidation (NQ) | 764433 \| PPTG_13378 \| Phytophthora parasitica INRA-310 hypothetical protein (7170 aa) |
| 2170 | 1630 | PPTG_16103 | 92,17 | 18 | 3 | 3 | 18 | 1,45E+05 | 3,00E+05 | 3,75E+06 | 5 | 5 | 1 | 1 | 5 | | 33722 \| PPTG_16103 \| Phytophthora parasitica INRA-310 hypothetical protein (303 aa) |
| 1942 | 1953 | PPTG_09677 | 92,12 | 22 | 5 | 22 | 16 | 1,32E+05 | 3,50E+06 | 2,60E+06 | 4 | 4 | 1 | 4 | 4 | Acetylation (N-term) | 24028 \| PPTG_09677 \| Phytophthora parasitica INRA-310 deoxyribose-phosphate aldolase (228 aa) |
| 1834 | 2193 | PPTG_08764 | 92,11 | 30 | 8 | 13 | 30 | 1,63E+05 | 1,51E+06 | 2,43E+06 | 4 | 4 | 1 | 3 | 6 | Deamidation (NQ) | 23963 \| PPTG_08764 \| Phytophthora parasitica INRA-310 DNA-directed RNA polymerase I II and III subunit RPABC1 (207 aa) |
| 1787 | 1386 | PPTG_04943 | 92,05 | 11 | 2 | 1 | 11 | 1,60E+05 | 1,74E+05 | 3,62E+06 | 7 | 7 | 2 | 1 | 7 | | 103205 \| PPTG_04943 \| Phytophthora parasitica INRA-310 phosphatidylinositol kinase (PIK-H2) (926 aa) |
| 2050 | 1859 | PPTG_02876 | 92,01 | 26 | 0 | 7 | 26 | | 3,44E+05 | 3,30E+06 | 5 | 5 | 0 | 2 | 6 | Oxidation (M); Carbamidomethylation | 36930 \| PPTG_02876 \| Phytophthora parasitica INRA-310 mitochondrial Carrier (MC) family (339 aa) |
| 1350 | 2326 | PPTG_19706 | 91,91 | 41 | 41 | 41 | 41 | 3,33E+05 | 1,61E+06 | 2,26E+06 | 3 | 1 | 4 | 6 | 6 | Oxidation (M) | 10937 \| PPTG_19706 \| Phytophthora parasitica INRA-310 hypothetical protein (103 aa) |
| 2277 | 2274 | PPTG_07600 | 91,89 | 28 | 0 | 28 | 10 | | 4,57E+06 | 2,90E+06 | 3 | 3 | 0 | 4 | 2 | Carbamidomethylation; Deamidation (NQ) | 22354 \| PPTG_07600 \| Phytophthora parasitica INRA-310 hypothetical protein (201 aa) |
| 1391 | 1709 | PPTG_10880 | 91,82 | 49 | 18 | 49 | 49 | 4,27E+05 | 6,12E+06 | 8,78E+06 | 5 | 5 | 2 | 6 | 7 | Oxidation (M); Carbamidomethylation; Acetylation (N-term) | 13568 \| PPTG_10880 \| Phytophthora parasitica INRA-310 hypothetical protein (126 aa) |
| 2033 | 2418 | PPTG_10398 | 91,8 | 14 | 14 | 14 | 11 | 4,19E+05 | 1,34E+06 | 2,69E+06 | 3 | 3 | 3 | 3 | 2 | | 29545 \| PPTG_10398 \| Phytophthora parasitica INRA-310 hypothetical protein (263 aa) |
| 1952 | 1765 | PPTG_18329 | 91,74 | 14 | 0 | 14 | 10 | | 1,72E+06 | 2,83E+06 | 5 | 5 | 0 | 5 | 4 | Oxidation (M) | 50299 \| PPTG_18329 \| Phytophthora parasitica INRA-310 histone deacetylase (448 aa) |
| 2266 | 2520 | PPTG_04165 | 91,54 | 3 | 0 | 1 | 3 | | | 3,68E+05 | 4 | 2 | 0 | 1 | 5 | Carbamidomethylation | 152436 \| PPTG_04165 \| Phytophthora parasitica INRA-310 phosphatidylinositol kinase (1351 aa) |
| 2123 | 1983 | PPTG_17670 | 91,49 | 14 | 2 | 14 | 5 | 1,66E+05 | 1,44E+06 | 1,70E+06 | 4 | 4 | 1 | 4 | 2 | Oxidation (M); Carbamidomethylation | 55096 \| PPTG_17670 \| Phytophthora parasitica INRA-310 E1 family ubiquitin activating enzyme (504 aa) |
| 2167 | 2466 | PPTG_01474 | 91,48 | 20 | 0 | 12 | 17 | | 8,20E+05 | 1,00E+06 | 5 | 5 | 0 | 3 | 4 | Oxidation (M) | 40552 \| PPTG_01474 \| Phytophthora parasitica INRA-310 hypothetical protein (369 aa) |
| 1966 | 1835 | PPTG_12171 | 91,39 | 8 | 5 | 1 | 6 | 3,25E+05 | 4,34E+05 | 4,24E+06 | 5 | 5 | 3 | 1 | 4 | | 85734 \| PPTG_12171 \| Phytophthora parasitica INRA-310 hypothetical protein (753 aa) |
| 1740 | 2000 | PPTG_01559 | 91,37 | 19 | 3 | 5 | 17 | 1,89E+05 | 3,32E+05 | 5,49E+06 | 5 | 5 | 2 | 2 | 7 | Oxidation (M); Deamidation (NQ) | 37587 \| PPTG_01559 \| Phytophthora parasitica INRA-310 hypothetical protein (345 aa) |
| 2261 | 2054 | PPTG_07089 | 91,28 | 30 | 0 | 23 | 30 | | 2,59E+06 | 3,43E+06 | 4 | 4 | 0 | 3 | 4 | Carbamidomethylation | 28589 \| PPTG_07089 \| Phytophthora parasitica INRA-310 NPP1-like protein (256 aa) |
| 2150 | 1979 | PPTG_09540 | 91,23 | 14 | 3 | 14 | 7 | 4,53E+04 | 1,99E+06 | 6,63E+05 | 4 | 4 | 1 | 4 | 2 | | 41854 \| PPTG_09540 \| Phytophthora parasitica INRA-310 hypothetical protein (398 aa) |
| 1789 | 1595 | PPTG_16291 | 91,22 | 20 | 4 | 20 | 16 | 2,39E+05 | 2,29E+06 | 2,68E+06 | 5 | 5 | 1 | 5 | 4 | | 32394 \| PPTG_16291 \| Phytophthora parasitica INRA-310 3-mercaptopyruvate sulfurtransferase (292 aa) |
| 2040 | 2106 | PPTG_02949 | 91,21 | 40 | 0 | 29 | 33 | | 2,81E+06 | 2,34E+06 | 4 | 3 | 0 | 4 | 4 | Carbamidomethylation | 15384 \| PPTG_02949 \| Phytophthora parasitica INRA-310 polysaccharide lyase (146 aa) |
| 1844 | 1987 | PPTG_15405 | 91,17 | 17 | 3 | 12 | 17 | 6,35E+04 | 1,54E+06 | 7,07E+06 | 4 | 4 | 1 | 3 | 6 | Oxidation (M); Carbamidomethylation | 43867 \| PPTG_15405 \| Phytophthora parasitica INRA-310 hypothetical protein (393 aa) |
| 2290 | 2092 | PPTG_15482 | 91,16 | 23 | 5 | 0 | 23 | 8,65E+04 | | 1,54E+06 | 4 | 4 | 1 | 0 | 5 | | 25438 \| PPTG_15482 \| Phytophthora parasitica INRA-310 transmembrane protein (230 aa) |
| 1677 | 1687 | PPTG_00103 | 91,12 | 19 | 4 | 4 | 19 | | | 2,24E+05 | 6 | 2 | 1 | 2 | 8 | | 34494 \| PPTG_00103 \| Phytophthora parasitica INRA-310 phospholipid hydroperoxide glutathione peroxidase (309 aa) |
| 1799 | 2118 | PPTG_12755 | 90,98 | 20 | 3 | 17 | 20 | 3,77E+05 | 1,07E+07 | 1,53E+07 | 5 | 5 | 1 | 4 | 5 | | 26603 \| PPTG_12755 \| Phytophthora parasitica INRA-310 pre-mRNA-splicing factor SF2-like protein (239 aa) |
| 1647 | 1510 | PPTG_10821 | 90,94 | 19 | 2 | 17 | 17 | 4,37E+04 | 2,33E+06 | 1,25E+06 | 6 | 6 | 1 | 5 | 5 | Carbamidomethylation; Acetylation (N-term) | 45941 \| PPTG_10821 \| Phytophthora parasitica INRA-310 COP9 signalosome complex subunit 1 (403 aa) |
| 1709 | 1521 | PPTG_04860 | 90,84 | 23 | 0 | 17 | 23 | | 1,02E+06 | 2,58E+06 | 6 | 6 | 0 | 4 | 7 | Carbamidomethylation | 37547 \| PPTG_04860 \| Phytophthora parasitica INRA-310 hypothetical protein (337 aa) |
| 2036 | 1487 | PPTG_14716 | 90,8 | 10 | 0 | 3 | 8 | | 1,86E+05 | 3,02E+06 | 7 | 7 | 0 | 2 | 6 | Carbamidomethylation | 109440 \| PPTG_14716 \| Phytophthora parasitica INRA-310 hypothetical protein (1021 aa) |
| 878 | 2417 | PPTG_13080 | 90,78 | 34 | 31 | 34 | 34 | 1,05E+08 | 2,70E+08 | 5,33E+08 | 4 | 4 | 4 | 10 | 12 | Carbamidomethylation; Dehydration; Deamidation (NQ); Carbamylation | 12437 \| PPTG_13080 \| Phytophthora parasitica INRA-310 30S ribosomal protein S26e (108 aa) |
| 2134 | 2033 | PPTG_16655 | 90,77 | 14 | 3 | 6 | 14 | 8,82E+04 | 2,33E+05 | 2,76E+06 | 4 | 4 | 1 | 2 | 4 | Carbamidomethylation; Acetylation (N-term) | 37019 \| PPTG_16655 \| Phytophthora parasitica INRA-310 mitoferrin-like protein (343 aa) |
| 1757 | 1501 | PPTG_11090 | 90,73 | 10 | 0 | 5 | 10 | | 4,85E+05 | 1,99E+06 | 7 | 7 | 0 | 3 | 7 | | 86419 \| PPTG_11090 \| Phytophthora parasitica INRA-310 hypothetical protein (797 aa) |
| 2239 | 2389 | PPTG_06974 | 90,73 | 18 | 0 | 0 | 18 | | | 3,85E+06 | 3 | 3 | 0 | 0 | 7 | Carbamidomethylation; Deamidation (NQ) | 28613 \| PPTG_06974 \| Phytophthora parasitica INRA-310 cytochrome b ascorbate-dependent protein (268 aa) |
| 1414 | 1638 | PPTG_09583 | 90,69 | 5 | 0 | 4 | 3 | 6,51E+04 | 1,57E+06 | 1,98E+06 | 10 | 10 | 1 | 7 | 6 | Carbamidomethylation | 212816 \| PPTG_09583 \| Phytophthora parasitica INRA-310 rRNA biogenesis protein rrp5 (1935 aa) |
| 2535 | 1991 | PPTG_08165 | 90,68 | 13 | 3 | 0 | 13 | 4,18E+05 | | 4,82E+06 | 4 | 4 | 1 | 0 | 4 | Carbamidomethylation | 67003 \| PPTG_08165 \| Phytophthora parasitica INRA-310 cathepsin cysteine protease family C01A (599 aa) |
| 1564 | 1999 | PPTG_01645 | 90,64 | 14 | 6 | 12 | 6 | 6,33E+05 | 3,21E+06 | 3,14E+06 | 7 | 7 | 3 | 6 | 3 | | 65931 \| PPTG_01645 \| Phytophthora parasitica INRA-310 hypothetical protein (593 aa) |
| 2326 | 1981 | PPTG_01053 | 90,49 | 28 | 0 | 20 | 23 | | 9,36E+05 | 2,80E+06 | 4 | 3 | 0 | 3 | 3 | Carbamidomethylation | 19313 \| PPTG_01053 \| Phytophthora parasitica INRA-310 hypothetical protein (174 aa) |
| 1569 | 1440 | PPTG_00938 | 90,45 | 14 | 0 | 14 | 12 | | 3,89E+06 | 7,92E+06 | 6 | 6 | 0 | 6 | 6 | Oxidation (M) | 60177 \| PPTG_00938 \| Phytophthora parasitica INRA-310 soluble pyridine nucleotide transhydrogenase (551 aa) |
| 1754 | 1547 | PPTG_06694 | 90,43 | 6 | 0 | 1 | 5 | | | 3,00E+06 | 9 | 6 | 1 | 1 | 8 | Oxidation (M); Carbamidomethylation; Deamidation (NQ) | 169192 \| PPTG_06694 \| Phytophthora parasitica INRA-310 phospholipid-transporting ATPase (1521 aa) |

| | | | | | | | | | | | | | | | | | | | |
|---|---|---|---|---|---|---|---|---|---|---|---|---|---|---|---|---|---|---|---|
| 1731 | 1764 | PPTG_11219 | 90,35 | 9 | 2 | 7 | 9 | 2,08E+05 | 1,11E+06 | 5,19E+06 | 6 | 6 | 1 | 4 | 6 | Carbamidomethylation | 70549 | PPTG_11219 | Phytophthora parasitica INRA-310 protein kinase (647 aa) |
| 2566 | 2288 | PPTG_05364 | 90,34 | 8 | 0 | 3 | 7 | | 1,80E+06 | 2,78E+06 | 4 | 4 | 0 | 2 | 3 | Oxidation (M) | 73715 | PPTG_05364 | Phytophthora parasitica INRA-310 aspartyl protease family A01B (685 aa) |
| 2024 | 2067 | PPTG_06690 | 90,3 | 28 | 8 | 8 | 28 | 6,79E+04 | 3,68E+05 | 5,56E+06 | 4 | 4 | 1 | 2 | 5 | Carbamidomethylation | 23828 | PPTG_06690 | Phytophthora parasitica INRA-310 hypothetical protein (212 aa) |
| 2803 | 2704 | PPTG_07806 | 90,23 | 12 | 0 | 4 | 12 | | 4,08E+05 | 6,19E+06 | 3 | 3 | 0 | 1 | 3 | Oxidation (M) | 37994 | PPTG_07806 | Phytophthora parasitica INRA-310 hypothetical protein (353 aa) |
| 2002 | 1820 | PPTG_02038 | 90,23 | 10 | 0 | 6 | 10 | | 4,01E+05 | 1,39E+06 | 5 | 5 | 0 | 3 | 5 | | 68163 | PPTG_02038 | Phytophthora parasitica INRA-310 thioredoxin-like protein (607 aa) |
| 1950 | 1707 | PPTG_15633 | 90,21 | 27 | 0 | 14 | 27 | | 9,54E+05 | 1,05E+06 | 6 | 6 | 0 | 3 | 6 | Oxidation (M); Acetylation (N-term) | 35270 | PPTG_15633 | Phytophthora parasitica INRA-310 nuclear migration protein nudC (308 aa) |
| 1434 | 1597 | PPTG_16831 | 90,19 | 22 | 4 | 22 | 22 | 5,47E+05 | 5,16E+06 | 7,13E+06 | 7 | 7 | 1 | 7 | 6 | Carbamidomethylation | 38369 | PPTG_16831 | Phytophthora parasitica INRA-310 hypothetical protein (339 aa) |
| 1631 | 2043 | PPTG_01000 | 90,06 | 29 | 15 | 29 | 29 | 4,60E+05 | 7,09E+06 | 5,94E+06 | 4 | 4 | 2 | 5 | 5 | Carbamidomethylation | 17481 | PPTG_01000 | Phytophthora parasitica INRA-310 hypothetical protein (151 aa) |
| 1433 | 1577 | PPTG_02472 | 90,06 | 9 | 2 | 4 | 7 | 1,93E+05 | 1,56E+06 | 2,79E+06 | 9 | 9 | 2 | 4 | 8 | Oxidation (M); Carbamidomethylation; Acetylation (N-term) | 135417 | PPTG_02472 | Phytophthora parasitica INRA-310 hypothetical protein (1231 aa) |
| 2276 | 1877 | PPTG_12921 | 90,04 | 11 | 0 | 8 | 7 | | 1,19E+06 | 2,16E+06 | 4 | 4 | 0 | 3 | 3 | | 67567 | PPTG_12921 | Phytophthora parasitica INRA-310 hypothetical protein (608 aa) |
| 1728 | 2358 | PPTG_13018 | 89,93 | 15 | 3 | 15 | 3 | 7,82E+04 | 1,34E+07 | 1,11E+07 | 3 | 3 | 1 | 7 | 3 | Carbamidomethylation; Deamidation (NQ) | 44184 | PPTG_13018 | Phytophthora parasitica INRA-310 hypothetical protein (421 aa) |
| 1693 | 1929 | PPTG_16158 | 89,92 | 13 | 4 | 13 | 10 | 1,19E+05 | 3,76E+06 | 3,99E+06 | 5 | 5 | 1 | 5 | 5 | | 38496 | PPTG_16158 | Phytophthora parasitica INRA-310 hypothetical protein (347 aa) |
| 1495 | 1726 | PPTG_04180 | 89,9 | 25 | 0 | 12 | 25 | | 9,57E+05 | 4,15E+06 | 8 | 8 | 0 | 4 | 9 | Oxidation (M); Carbamidomethylation | 41355 | PPTG_04180 | Phytophthora parasitica INRA-310 hypothetical protein (368 aa) |
| 2796 | 2411 | PPTG_04803 | 89,84 | 15 | 0 | 15 | 3 | | 1,77E+06 | 7,87E+05 | 3 | 3 | 0 | 3 | 1 | Carbamidomethylation | 31928 | PPTG_04803 | Phytophthora parasitica INRA-310 hypothetical protein (300 aa) |
| 2796 | 2412 | PPTG_04803 | 89,84 | 15 | 0 | 15 | 3 | | 1,77E+06 | 7,87E+05 | 3 | 3 | 0 | 3 | 1 | Carbamidomethylation | 31928 | PPTG_04803 | Phytophthora parasitica INRA-310 hypothetical protein (300 aa) |
| 1574 | 1533 | PPTG_09841 | 89,82 | 5 | 0 | 4 | 3 | | 1,50E+06 | 5,48E+05 | 7 | 5 | 1 | 6 | 5 | Oxidation (M); Carbamidomethylation; Dehydration | 200693 | PPTG_09841 | Phytophthora parasitica INRA-310 histone-lysine N-methyltransferase (1777 aa) |
| 1788 | 1539 | PPTG_01184 | 89,81 | 6 | 0 | 6 | 4 | | 2,05E+06 | 3,74E+05 | 6 | 6 | 0 | 6 | 4 | Carbamidomethylation | 120769 | PPTG_01184 | Phytophthora parasitica INRA-310 enhancer-mRNA-decapping protein (1117 aa) |
| 2182 | 1836 | PPTG_07599 | 89,78 | 37 | 0 | 29 | 21 | | 1,49E+06 | 1,99E+06 | 5 | 5 | 0 | 4 | 3 | Carbamidomethylation; Acetylation (N-term) | 19046 | PPTG_07599 | Phytophthora parasitica INRA-310 hypothetical protein (174 aa) |
| 2133 | 1802 | PPTG_02243 | 89,69 | 6 | 0 | 5 | 4 | | 9,18E+05 | 6,07E+05 | 5 | 5 | 0 | 4 | 3 | Carbamidomethylation | 93261 | PPTG_02243 | Phytophthora parasitica INRA-310 hypothetical protein (850 aa) |
| 1929 | 2110 | PPTG_18765 | 89,69 | 10 | 0 | 10 | 8 | | 1,72E+06 | 7,41E+05 | 4 | 4 | 0 | 5 | 4 | Carbamidomethylation | 67523 | PPTG_18765 | Phytophthora parasitica INRA-310 cell 5A endo-1 4-betaglucanase (636 aa) |
| 1930 | 2396 | PPTG_14068 | 89,67 | 38 | 26 | 26 | 38 | 3,82E+06 | 9,28E+06 | 3,95E+07 | 3 | 3 | 2 | 2 | 5 | | 11375 | PPTG_14068 | Phytophthora parasitica INRA-310 hypothetical protein (102 aa) |
| 2231 | 1782 | PPTG_04094 | 89,66 | 9 | 2 | 0 | 9 | 1,20E+05 | | 2,39E+06 | 6 | 5 | 1 | 0 | 6 | Carbamidomethylation | 76855 | PPTG_04094 | Phytophthora parasitica INRA-310 hypothetical protein (693 aa) |
| 1525 | 1611 | PPTG_13865 | 89,63 | 40 | 14 | 36 | 33 | 1,19E+06 | 8,18E+06 | 2,49E+07 | 6 | 6 | 2 | 5 | 6 | Oxidation (M); Carbamidomethylation | 20140 | PPTG_13865 | Phytophthora parasitica INRA-310 ADP-ribosylation factor family protein (180 aa) |
| 2480 | 2091 | PPTG_14994 | 89,59 | 22 | 9 | 9 | 22 | 3,84E+04 | 4,45E+05 | 2,26E+07 | 3 | 3 | 1 | 1 | 3 | Carbamidomethylation | 21035 | PPTG_14994 | Phytophthora parasitica INRA-310 hypothetical protein (187 aa) |
| 1863 | 1718 | PPTG_08652 | 89,58 | 10 | 0 | 8 | 9 | | 8,10E+05 | 1,43E+06 | 5 | 5 | 0 | 5 | 4 | | 62737 | PPTG_08652 | Phytophthora parasitica INRA-310 tubulin-specific chaperone E (567 aa) |
| 2013 | 2144 | PPTG_03156 | 89,57 | 10 | 0 | 9 | 10 | | 6,25E+06 | 5,34E+06 | 4 | 4 | 0 | 3 | 5 | Carbamidomethylation | 44976 | PPTG_03156 | Phytophthora parasitica INRA-310 hypothetical protein (446 aa) |
| 1741 | 2075 | PPTG_10433 | 89,51 | 27 | 9 | 27 | 15 | 2,24E+05 | 2,28E+06 | 1,25E+06 | 5 | 5 | 2 | 6 | 3 | Oxidation (M); Carbamidomethylation | 27216 | PPTG_10433 | Phytophthora parasitica INRA-310 hypothetical protein (235 aa) |
| 3027 | 2651 | PPTG_05385 | 89,46 | 21 | 0 | 0 | 21 | | | 3,75E+05 | 3 | 3 | 0 | 0 | 3 | Carbamidomethylation; Acetylation (N-term) | 23953 | PPTG_05385 | Phytophthora parasitica INRA-310 hypothetical protein (210 aa) |
| 1666 | 2084 | PPTG_09639 | 89,44 | 11 | 0 | 8 | 10 | | 5,88E+06 | 5,74E+06 | 7 | 7 | 0 | 5 | 6 | Oxidation (M); Carbamidomethylation | 61827 | PPTG_09639 | Phytophthora parasitica INRA-310 lysosomal Pro-X carboxypeptidase (566 aa) |
| 1688 | 1572 | PPTG_12865 | 89,4 | 26 | 7 | 26 | 11 | 1,02E+06 | 1,91E+07 | 1,01E+07 | 6 | 5 | 2 | 6 | 3 | Oxidation (M); Carbamidomethylation | 26310 | PPTG_12865 | Phytophthora parasitica INRA-310 hypothetical protein (252 aa) |
| 1980 | 2053 | PPTG_16235 | 89,31 | 9 | 1 | 8 | 6 | | 1,17E+06 | 5,13E+05 | 4 | 2 | 1 | 4 | 3 | | 61868 | PPTG_16235 | Phytophthora parasitica INRA-310 transglutaminase elicitor-like protein (578 aa) |
| 2458 | 1528 | PPTG_03734 | 89,31 | 5 | 0 | 0 | 5 | | | 2,06E+06 | 5 | 5 | 0 | 0 | 5 | | 165102 | PPTG_03734 | Phytophthora parasitica INRA-310 sterol 3-beta-glucosyltransferase (1493 aa) |
| 1814 | 2109 | PPTG_07025 | 89,27 | 21 | 11 | 16 | 21 | 4,00E+05 | 5,45E+05 | 4,59E+06 | 5 | 5 | 2 | 3 | 5 | | 24184 | PPTG_07025 | Phytophthora parasitica INRA-310 Rab21/Rab5 family GTPase (225 aa) |
| 1551 | 2178 | PPTG_02353 | 89,25 | 33 | 16 | 33 | 33 | 8,03E+05 | 2,30E+06 | 7,45E+06 | 4 | 4 | 3 | 5 | 5 | Oxidation (M); Carbamidomethylation; Dehydration; Deamidation (NQ); Acetylation (N-term) | 19537 | PPTG_02353 | Phytophthora parasitica INRA-310 hypothetical protein (179 aa) |
| 1435 | 1797 | PPTG_06799 | 89,21 | 16 | 7 | 16 | 14 | 1,49E+05 | 2,97E+06 | 5,64E+06 | 5 | 4 | 2 | 6 | 6 | Carbamidomethylation | 40141 | PPTG_06799 | Phytophthora parasitica INRA-310 cAMP-dependent protein kinase catalytic subunit (354 aa) |
| 2950 | 3641 | PPTG_20433 | 89,06 | 7 | 6 | 6 | 7 | 1,38E+06 | 4,91E+06 | 1,19E+07 | 2 | 2 | 1 | 1 | 2 | | 53294 | PPTG_20433 | Phytophthora parasitica INRA-310 hypothetical protein (606 aa) |
| 1598 | 1856 | PPTG_04259 | 89,03 | 26 | 0 | 22 | 19 | | 1,93E+06 | 1,45E+06 | 7 | 7 | 0 | 6 | 6 | Oxidation (M); Deamidation (NQ) | 42533 | PPTG_04259 | Phytophthora parasitica INRA-310 2-amino-3-carboxymuconate-6-semialdehyde decarboxylase (381 aa) |
| 1526 | 1806 | PPTG_15684 | 89,01 | 58 | 0 | 46 | 58 | | 4,24E+06 | 4,96E+06 | 5 | 5 | 0 | 6 | 7 | Acetylation (N-term) | 12922 | PPTG_15684 | Phytophthora parasitica INRA-310 prefoldin subunit 2 (113 aa) |
| 1751 | 1723 | PPTG_14422 | 88,91 | 1 | 0 | 1 | 1 | | 6,16E+05 | 1,31E+06 | 6 | 5 | 0 | 4 | 6 | | 516584 | PPTG_14422 | Phytophthora parasitica INRA-310 dynein heavy chain (4557 aa) |
| 1890 | 1792 | PPTG_14826 | 88,88 | 22 | 3 | 11 | 16 | 1,24E+05 | 2,05E+06 | 1,81E+06 | 6 | 6 | 1 | 4 | 4 | Oxidation (M); Carbamidomethylation | 37467 | PPTG_14826 | Phytophthora parasitica INRA-310 replication factor C subunit 4 (340 aa) |
| 1723 | 1690 | PPTG_14196 | 88,79 | 7 | 2 | 6 | 5 | 1,92E+05 | 2,30E+06 | 2,11E+06 | 5 | 5 | 2 | 5 | 4 | Carbamidomethylation | 88247 | PPTG_14196 | Phytophthora parasitica INRA-310 adenosylmethionine-8-amino-7-oxononanoate aminotransferase (805 aa) |
| 1403 | 1755 | PPTG_16763 | 88,77 | 30 | 4 | 30 | 30 | 6,39E+05 | 1,15E+07 | 1,65E+07 | 5 | 5 | 1 | 7 | 7 | Oxidation (M) | 24089 | PPTG_16763 | Phytophthora parasitica INRA-310 hypothetical protein (226 aa) |
| 2025 | 2078 | PPTG_07189 | 88,69 | 17 | 0 | 17 | 17 | | 1,63E+06 | 1,89E+06 | 4 | 4 | 0 | 4 | 4 | | 35264 | PPTG_07189 | Phytophthora parasitica INRA-310 N-carbamoylputrescine amidase (321 aa) |
| 1713 | 1684 | PPTG_10386 | 88,64 | 26 | 4 | 16 | 21 | 6,07E+05 | 1,58E+06 | 5,94E+06 | 7 | 6 | 1 | 4 | 6 | | 30893 | PPTG_10386 | Phytophthora parasitica INRA-310 peroxiredoxin-4 (276 aa) |
| 1906 | 1716 | PPTG_04124 | 88,56 | 9 | 3 | 2 | 9 | 1,91E+05 | 6,48E+04 | 2,69E+06 | 5 | 5 | 2 | 1 | 6 | | 76870 | PPTG_04124 | Phytophthora parasitica INRA-310 hypothetical protein (677 aa) |
| 1906 | 1717 | PPTG_04124 | 88,56 | 8 | 3 | 2 | 8 | 1,91E+05 | 6,48E+04 | 2,69E+06 | 5 | 5 | 2 | 1 | 6 | | 79264 | PPTG_04124 | Phytophthora parasitica INRA-310 hypothetical protein (698 aa) |
| 2222 | 2198 | PPTG_14218 | 88,51 | 22 | 10 | 14 | 18 | 4,44E+05 | 5,46E+05 | 5,04E+06 | 4 | 4 | 2 | 2 | 3 | | 20713 | PPTG_14218 | Phytophthora parasitica INRA-310 hypothetical protein (182 aa) |
| 2156 | 1842 | PPTG_15132 | 88,51 | 21 | 0 | 3 | 21 | | 3,64E+05 | 1,70E+06 | 6 | 6 | 0 | 1 | 6 | | 43355 | PPTG_15132 | Phytophthora parasitica INRA-310 allantoicase (393 aa) |
| 1735 | 1585 | PPTG_18631 | 88,46 | 19 | 6 | 11 | 16 | 1,77E+05 | 2,84E+06 | 1,78E+06 | 6 | 6 | 2 | 4 | 5 | | 39156 | PPTG_18631 | Phytophthora parasitica INRA-310 hypothetical protein (354 aa) |
| 2670 | 1973 | PPTG_02614 | 88,34 | 10 | 0 | 0 | 10 | | | 1,50E+06 | 4 | 4 | 0 | 0 | 4 | Carbamidomethylation | 64760 | PPTG_02614 | Phytophthora parasitica INRA-310 hypothetical protein (580 aa) |
| 2336 | 1815 | PPTG_03559 | 88,33 | 16 | 0 | 16 | 3 | | 1,41E+06 | 6,81E+05 | 5 | 5 | 0 | 5 | 1 | Carbamidomethylation | 48861 | PPTG_03559 | Phytophthora parasitica INRA-310 hypothetical protein (458 aa) |
| 1589 | 1868 | PPTG_14075 | 88,32 | 14 | 9 | 5 | 14 | 5,69E+05 | 2,95E+05 | 9,95E+06 | 6 | 6 | 3 | 2 | 7 | Deamidation (NQ) | 40491 | PPTG_14075 | Phytophthora parasitica INRA-310 hypothetical protein (383 aa) |
| 1882 | 1841 | PPTG_03844 | 88,3 | 9 | 2 | 7 | 6 | | 1,36E+05 | | 5 | 1 | 1 | 4 | 4 | Oxidation (M); Carbamidomethylation | 64069 | PPTG_03844 | Phytophthora parasitica INRA-310 cell 5A endo-1 4-betaglucanase (583 aa) |
| 1504 | 3192 | PPTG_18856 | 88,26 | 12 | 12 | 12 | 12 | 8,50E+06 | 7,12E+07 | 5,97E+07 | 2 | 2 | 3 | 4 | 6 | Carbamidomethylation; Deamidation (NQ) | 30232 | PPTG_18856 | Phytophthora parasitica INRA-310 protease inhibitor EpiC4 (281 aa) |
| 2187 | 2152 | PPTG_07022 | 88,19 | 10 | 0 | 10 | 8 | | 1,41E+06 | 1,16E+06 | 4 | 4 | 0 | 4 | 3 | Acetylation (N-term) | 56309 | PPTG_07022 | Phytophthora parasitica INRA-310 exosome complex exonuclease RRP45-like protein (520 aa) |
| 1986 | 1670 | PPTG_02404 | 88,06 | 9 | 0 | 5 | 8 | | 1,21E+06 | 2,69E+06 | 6 | 6 | 0 | 3 | 5 | Carbamidomethylation | 78124 | PPTG_02404 | Phytophthora parasitica INRA-310 CCR4-NOT transcription complex subunit (709 aa) |
| 2081 | 1840 | PPTG_01983 | 88,01 | 14 | 0 | 5 | 14 | | 1,86E+06 | 6,67E+06 | 5 | 5 | 0 | 3 | 5 | Carbamidomethylation | 54328 | PPTG_01983 | Phytophthora parasitica INRA-310 hypothetical protein (500 aa) |
| 1437 | 1505 | PPTG_07366 | 87,88 | 38 | 10 | 13 | 38 | 3,01E+06 | 3,85E+06 | 4,31E+07 | 7 | 7 | 2 | 3 | 9 | Oxidation (M); Carbamidomethylation | 22226 | PPTG_07366 | Phytophthora parasitica INRA-310 Ras-like protein Rac1 (199 aa) |
| 2317 | 2271 | PPTG_13086 | 87,88 | 16 | 0 | 9 | 16 | | 5,18E+05 | 2,54E+06 | 3 | 3 | 0 | 2 | 4 | Carbamidomethylation | 25912 | PPTG_13086 | Phytophthora parasitica INRA-310 hypothetical protein (235 aa) |
| 1794 | 1556 | PPTG_09044 | 87,79 | 13 | 2 | 7 | 13 | 1,16E+05 | 4,07E+05 | 4,82E+06 | 6 | 6 | 1 | 3 | 6 | | 57729 | PPTG_09044 | Phytophthora parasitica INRA-310 hypothetical protein (500 aa) |
| 2364 | 2114 | PPTG_18158 | 87,74 | 15 | 3 | 11 | 9 | 4,67E+04 | 4,97E+05 | 9,89E+05 | 4 | 4 | 1 | 3 | 2 | Oxidation (M) | 41058 | PPTG_18158 | Phytophthora parasitica INRA-310 2-nitropropane dioxygenase (387 aa) |
| 1594 | 1560 | PPTG_13614 | 87,73 | 34 | 0 | 15 | 31 | | 7,43E+06 | 9,88E+06 | 7 | 7 | 0 | 5 | 7 | | 30444 | PPTG_13614 | Phytophthora parasitica INRA-310 50S ribosomal protein L4 (278 aa) |
| 1925 | 1818 | PPTG_02718 | 87,73 | 12 | 2 | 10 | 9 | 3,57E+05 | 2,69E+06 | 4,68E+06 | 5 | 5 | 1 | 4 | 4 | Oxidation (M) | 52801 | PPTG_02718 | Phytophthora parasitica INRA-310 D-lactate dehydrogenase (493 aa) |
| 2000 | 2297 | PPTG_05774 | 87,7 | 19 | 0 | 19 | 9 | | 1,16E+06 | 1,22E+06 | 3 | 3 | 0 | 6 | 2 | Carbamidomethylation | 49990 | PPTG_05774 | Phytophthora parasitica INRA-310 serine protease family S08A (478 aa) |
| 2964 | 2263 | PPTG_05098 | 87,68 | 6 | 0 | 0 | 6 | | | 1,32E+06 | 3 | 3 | 0 | 0 | 3 | | 76653 | PPTG_05098 | Phytophthora parasitica INRA-310 hypothetical protein (683 aa) |
| 2119 | 2047 | PPTG_16404 | 87,62 | 5 | 4 | 0 | 5 | 2,69E+05 | | 3,30E+06 | 4 | 4 | 3 | 0 | 4 | Oxidation (M); Carbamidomethylation | 101990 | PPTG_16404 | Phytophthora parasitica INRA-310 hypothetical protein (902 aa) |
| 2154 | 3038 | PPTG_00855 | 87,61 | 13 | 0 | 13 | 13 | | 1,96E+06 | 1,11E+07 | 3 | 3 | 0 | 3 | 4 | Carbamidomethylation | 19078 | PPTG_00855 | Phytophthora parasitica INRA-310 hypothetical protein (171 aa) |
| 1803 | 1575 | PPTG_13826 | 87,61 | 46 | 0 | 15 | 46 | | 4,46E+05 | 6,07E+06 | 6 | 6 | 0 | 2 | 8 | Acetylation (N-term) | 22025 | PPTG_13826 | Phytophthora parasitica INRA-310 hypothetical protein (199 aa) |
| 2117 | 1763 | PPTG_03676 | 87,55 | 10 | 0 | 4 | 10 | | 5,50E+05 | 1,32E+06 | 5 | 5 | 0 | 2 | 5 | | 71839 | PPTG_03676 | Phytophthora parasitica INRA-310 translation elongation factor 1-alpha (661 aa) |
| 2159 | 2093 | PPTG_10067 | 87,49 | 21 | 0 | 21 | 11 | | 1,10E+06 | 1,04E+06 | 4 | 4 | 0 | 4 | 2 | Carbamidomethylation | 27541 | PPTG_10067 | Phytophthora parasitica INRA-310 hypothetical protein (253 aa) |
| 2095 | 1924 | PPTG_09281 | 87,39 | 22 | 0 | 17 | 19 | | 1,01E+06 | 4,56E+06 | 5 | 5 | 0 | 4 | 4 | Oxidation (M); Carbamidomethylation | 34900 | PPTG_09281 | Phytophthora parasitica INRA-310 hypothetical protein (321 aa) |
| 1891 | 1825 | PPTG_00715 | 87,34 | 5 | 2 | 5 | 4 | 1,81E+05 | 1,48E+06 | 8,78E+05 | 4 | 4 | 2 | 4 | 3 | | 120659 | PPTG_00715 | Phytophthora parasitica INRA-310 hypothetical protein (1092 aa) |
| 1349 | 2209 | PPTG_20203 | 87,33 | 43 | 43 | 43 | 43 | 3,37E+06 | 5,86E+07 | 5,03E+07 | 3 | 3 | 3 | 5 | 7 | | 9227 | PPTG_20203 | Phytophthora parasitica INRA-310 FK506-binding protein 1 (84 aa) |
| 2538 | 2300 | PPTG_02786 | 87,32 | 11 | 0 | 2 | 11 | | | 4,35E+06 | 3 | 2 | 0 | 1 | 4 | Oxidation (M); Carbamidomethylation | 42238 | PPTG_02786 | Phytophthora parasitica INRA-310 UDP-glucose 4-epimerase (384 aa) |
| 1687 | 1570 | PPTG_03131 | 87,25 | 40 | 8 | 40 | 34 | 1,29E+05 | 6,82E+06 | 7,30E+06 | 5 | 5 | 1 | 6 | 4 | Carbamidomethylation | 16238 | PPTG_03131 | Phytophthora parasitica INRA-310 mitochondrial 39-S ribosomal protein L47 (145 aa) |
| 2177 | 2323 | PPTG_01920 | 87,21 | 19 | 5 | 13 | 19 | 2,81E+05 | 1,06E+06 | 3,57E+06 | 3 | 3 | 1 | 2 | 4 | | 24404 | PPTG_01920 | Phytophthora parasitica INRA-310 hypothetical protein (210 aa) |
| 2409 | 2376 | PPTG_03082 | 87,15 | 15 | 0 | 15 | 11 | | 3,17E+06 | 1,04E+06 | 3 | 3 | 0 | 4 | 2 | | 31633 | PPTG_03082 | Phytophthora parasitica INRA-310 chitin-binding protein (305 aa) |
| 1830 | 2045 | PPTG_00611 | 87,14 | 43 | 6 | 14 | 43 | 3,38E+06 | 1,64E+07 | 4,44E+07 | 5 | 5 | 1 | 2 | 7 | | 17267 | PPTG_00611 | Phytophthora parasitica INRA-310 hypothetical protein (161 aa) |
| 1630 | 1843 | PPTG_12014 | 87,11 | 16 | 6 | 13 | 16 | 3,13E+06 | 7,25E+06 | 3,15E+07 | 6 | 6 | 2 | 5 | 5 | Acetylation (N-term) | 33216 | PPTG_12014 | Phytophthora parasitica INRA-310 hypothetical protein (291 aa) |
| 1909 | 1990 | PPTG_11690 | 87,1 | 18 | 3 | 14 | 15 | 1,90E+05 | 1,83E+06 | 3,07E+06 | 5 | 5 | 1 | 4 | 4 | Oxidation (M); Carbamidomethylation | 48223 | PPTG_11690 | Phytophthora parasitica INRA-310 hypothetical protein (434 aa) |
| 1989 | 1916 | PPTG_10714 | 87,03 | 6 | 2 | 4 | 4 | 1,64E+05 | 1,89E+05 | 9,71E+05 | 5 | 1 | 2 | 3 | 3 | Carbamidomethylation; Dehydration; Acetylation (N-term) | 78516 | PPTG_10714 | Phytophthora parasitica INRA-310 ATP-binding Cassette (ABC) superfamily (718 aa) |
| 1249 | 2032 | PPTG_20175 | 87,01 | 8 | 2 | 7 | 8 | | 4,27E+05 | 3,23E+05 | 4 | 1 | 1 | 8 | 8 | Carbamidomethylation; Dehydration; Deamidation (NQ) | 53826 | PPTG_20175 | Phytophthora parasitica INRA-310 1 3-beta-glucanosyltransferase (490 aa) |
| 1926 | 1922 | PPTG_02912 | 86,96 | 26 | 5 | 19 | 13 | 1,13E+05 | 9,51E+06 | 1,14E+06 | 6 | 6 | 1 | 5 | 3 | Carbamidomethylation | 32956 | PPTG_02912 | Phytophthora parasitica INRA-310 COP9 signalosome complex subunit (292 aa) |
| 2111 | 1660 | PPTG_18747 | 86,89 | 8 | 0 | 5 | 7 | | 5,70E+05 | 9,54E+05 | 5 | 5 | 0 | 3 | 4 | | 96636 | PPTG_18747 | Phytophthora parasitica INRA-310 inositol polyphosphate 5-phosphatase (877 aa) |
| 2253 | 2243 | PPTG_05596 | 86,81 | 43 | 0 | 34 | 43 | | 2,44E+06 | 2,78E+06 | 4 | 4 | 0 | 3 | 4 | Acetylation (N-term) | 15509 | PPTG_05596 | Phytophthora parasitica INRA-310 hypothetical protein (144 aa) |
| 2301 | 2099 | PPTG_13376 | 86,81 | 17 | 4 | 9 | 13 | 1,72E+05 | 2,70E+05 | 1,71E+06 | 4 | 4 | 1 | 2 | 3 | | 28786 | PPTG_13376 | Phytophthora parasitica INRA-310 transmembrane protein (274 aa) |
| 2037 | 1629 | PPTG_06776 | 86,8 | 11 | 0 | 1 | 11 | | 2,35E+05 | 2,84E+06 | 7 | 7 | 0 | 1 | 7 | | 86246 | PPTG_06776 | Phytophthora parasitica INRA-310 hypothetical protein (782 aa) |
| 1700 | 1905 | PPTG_08344 | 86,78 | 17 | 8 | 17 | 17 | | 1,49E+05 | 4,33E+05 | 4 | 1 | 2 | 5 | 4 | Oxidation (M) | 25425 | PPTG_08344 | Phytophthora parasitica INRA-310 hypothetical protein (225 aa) |
| 1779 | 1603 | PPTG_18242 | 86,76 | 5 | 1 | 3 | 5 | 7,66E+04 | 1,36E+06 | 3,51E+06 | 5 | 5 | 1 | 4 | 5 | | 124297 | PPTG_18242 | Phytophthora parasitica INRA-310 hypothetical protein (1120 aa) |
| 2653 | 2199 | PPTG_18143 | 86,75 | 7 | 0 | 4 | 7 | | 0 | | 3 | 1 | 0 | 1 | 3 | | 53183 | PPTG_18143 | Phytophthora parasitica INRA-310 ATP-binding Cassette (ABC) superfamily (486 aa) |
| 1492 | 1529 | PPTG_07870 | 86,73 | 11 | 5 | 7 | 11 | 5,09E+05 | 3,31E+06 | 6,65E+06 | 6 | 6 | 3 | 4 | 6 | | 73735 | PPTG_07870 | Phytophthora parasitica INRA-310 long-chain-fatty-acid-CoA ligase (667 aa) |
| 2279 | 2210 | PPTG_10963 | 86,72 | 12 | 0 | 12 | 6 | | 2,93E+06 | 4,08E+06 | 4 | 4 | 0 | 4 | 2 | Oxidation (M) | 47790 | PPTG_10963 | Phytophthora parasitica INRA-310 hypothetical protein (435 aa) |
| 2248 | 3349 | PPTG_01293 | 86,71 | 13 | 0 | 9 | 13 | | 2,03E+05 | 2,03E+06 | 2 | 2 | 0 | 1 | 6 | Oxidation (M); Deamidation (NQ); Acetylation (N-term) | 31321 | PPTG_01293 | Phytophthora parasitica INRA-310 hypothetical protein (289 aa) |
| 1456 | 1444 | PPTG_19699 | 86,68 | 13 | 1 | 13 | 11 | 6,08E+04 | 1,72E+06 | 2,28E+06 | 7 | 7 | 1 | 7 | 6 | Carbamidomethylation; Acetylation (N-term) | 78450 | PPTG_19699 | Phytophthora parasitica INRA-310 TRAF3-interacting protein (733 aa) |
| 2052 | 1966 | PPTG_04096 | 86,65 | 9 | 4 | 0 | 7 | 1,37E+05 | | 2,26E+06 | 6 | 6 | 2 | 0 | 6 | | 79884 | PPTG_04096 | Phytophthora parasitica INRA-310 transmembrane protein (707 aa) |
| 1776 | 2058 | PPTG_05050 | 86,63 | 37 | 0 | 25 | 37 | | 7,96E+06 | 1,54E+07 | 6 | 6 | 0 | 4 | 6 | Oxidation (M); Carbamidomethylation | 15843 | PPTG_05050 | Phytophthora parasitica INRA-310 FK506-binding protein 2 (143 aa) |
| 2335 | 1555 | PPTG_05363 | 86,62 | 10 | 0 | 0 | 10 | | | 1,64E+06 | 6 | 6 | 0 | 0 | 6 | Carbamidomethylation | 86125 | PPTG_05363 | Phytophthora parasitica INRA-310 hypothetical protein (787 aa) |
| 1992 | 2403 | PPTG_10715 | 86,61 | 14 | 5 | 8 | 11 | 3,48E+05 | 3,31E+05 | 3,18E+06 | 4 | 2 | 2 | 2 | 4 | Carbamidomethylation; Dehydration; Acetylation (N-term) | 34099 | PPTG_10715 | Phytophthora parasitica INRA-310 ATP-binding Cassette (ABC) superfamily (314 aa) |
| 1815 | 3272 | PPTG_15682 | 86,52 | 35 | 22 | 35 | 35 | 3,31E+06 | 1,56E+07 | 3,32E+07 | 3 | 3 | 1 | 5 | 4 | Oxidation (M) | 9412 | PPTG_15682 | Phytophthora parasitica INRA-310 mitochondrial protein Translocase (MPT) family (84 aa) |
| 2328 | 2324 | PPTG_15676 | 86,43 | 34 | 9 | 22 | 34 | 1,56E+05 | 3,97E+06 | 2,35E+07 | 3 | 3 | 1 | 2 | 3 | Carbamidomethylation | 13421 | PPTG_15676 | Phytophthora parasitica INRA-310 hypothetical protein (117 aa) |
| 1899 | 1602 | PPTG_17953 | 86,42 | 24 | 0 | 14 | 24 | | 1,94E+06 | 1,06E+07 | 5 | 5 | 0 | 3 | 6 | Carbamidomethylation; Deamidation (NQ) | 30986 | PPTG_17953 | Phytophthora parasitica INRA-310 endoplasmic reticulum-Golgi intermediate compartment protein (279 aa) |
| 2021 | 1865 | PPTG_03759 | 86,39 | 6 | 2 | 1 | 6 | 2,93E+05 | 3,94E+05 | 3,17E+06 | 5 | 5 | 2 | 1 | 5 | | 97002 | PPTG_03759 | Phytophthora parasitica INRA-310 guanylate-binding protein (882 aa) |
| 1938 | 2291 | PPTG_18253 | 86,37 | 16 | 7 | 9 | 14 | 2,95E+05 | 3,03E+05 | 2,29E+06 | 4 | 4 | 2 | 2 | 5 | Oxidation (M); Carbamidomethylation | 36003 | PPTG_18253 | Phytophthora parasitica INRA-310 hypothetical protein (310 aa) |
| 2046 | 2933 | PPTG_04102 | 86,24 | 18 | 0 | 18 | 16 | | 3,17E+06 | 1,29E+06 | 3 | 3 | 0 | 5 | 3 | | 24556 | PPTG_04102 | Phytophthora parasitica INRA-310 hypothetical protein (220 aa) |
| 2249 | 3383 | PPTG_17779 | 86,24 | 18 | 12 | 18 | 18 | 5,62E+05 | 3,84E+06 | 5,52E+06 | 2 | 2 | 1 | 3 | 3 | Oxidation (M); Carbamidomethylation | 19041 | PPTG_17779 | Phytophthora parasitica INRA-310 elicitin-like protein (198 aa) |
| 1809 | 1853 | PPTG_03703 | 86,19 | 14 | 7 | 11 | 11 | 1,32E+06 | 4,64E+06 | 1,12E+07 | 5 | 5 | 2 | 4 | 4 | Carbamylation | 42264 | PPTG_03703 | Phytophthora parasitica INRA-310 hypothetical protein (389 aa) |

| | | | | | | | | | | | | | | | | | | |
|---|---|---|---|---|---|---|---|---|---|---|---|---|---|---|---|---|---|---|
| 2032 | 2332 | PPTG_09574 | 86,15 | 19 | 6 | 19 | 19 | 2,50E+05 | 2,98E+06 | 5,69E+06 | 3 | 3 | 1 | 4 | 3 | Oxidation (M) | 24660 | PPTG_09574 \| Phytophthora parasitica INRA-310 hypothetical protein (211 aa) |
| 1699 | 1895 | PPTG_09165 | 86,14 | 3 | 0 | 3 | 3 | 9,57E+04 | 1,65E+06 | 2,19E+06 | 5 | 5 | 1 | 5 | 5 | | 162929 | PPTG_09165 \| Phytophthora parasitica INRA-310 hypothetical protein (1469 aa) |
| 2597 | 2659 | PPTG_15690 | 86,11 | 23 | 0 | 19 | 23 | | 3,84E+05 | 1,72E+06 | 3 | 3 | 0 | 2 | 3 | Carbamidomethylation | 27606 | PPTG_15690 \| Phytophthora parasitica INRA-310 SCO1 family protein (248 aa) |
| 2691 | 2390 | PPTG_19535 | 86,09 | 19 | 0 | 8 | 19 | | 9,36E+04 | 2,79E+05 | 3 | 3 | 0 | 1 | 3 | | 24231 | PPTG_19535 \| Phytophthora parasitica INRA-310 hypothetical protein (212 aa) |
| 1509 | 1628 | PPTG_03761 | 86,04 | 3 | 1 | 3 | 3 | 2,18E+05 | 1,50E+06 | 2,76E+06 | 6 | 6 | 2 | 5 | 6 | | 217954 | PPTG_03761 \| Phytophthora parasitica INRA-310 phosphoinositide 3-kinase regulatory subunit (2017 aa) |
| 1106 | 1761 | PPTG_01351 | 85,89 | 26 | 18 | 14 | 26 | 0 | 0 | 9,46E+06 | 6 | 3 | 6 | 5 | 9 | Carbamylation | 23271 | PPTG_01351 \| Phytophthora parasitica INRA-310 Ras-like protein RAB1BV (208 aa) |
| 2219 | 1974 | PPTG_01848 | 85,64 | 20 | 4 | 16 | 12 | 1,06E+05 | 1,25E+06 | 1,35E+06 | 4 | 4 | 1 | 3 | 3 | Oxidation (M) | 35570 | PPTG_01848 \| Phytophthora parasitica INRA-310 hypothetical protein (316 aa) |
| 1714 | 1733 | PPTG_09643 | 85,61 | 13 | 2 | 11 | 10 | 5,42E+04 | 1,93E+06 | 3,40E+06 | 6 | 6 | 1 | 5 | 5 | Carbamidomethylation | 55821 | PPTG_09643 \| Phytophthora parasitica INRA-310 60S ribosomal export protein nmd3 (499 aa) |
| 1842 | 1803 | PPTG_09511 | 85,59 | 21 | 9 | 5 | 21 | 2,61E+05 | | 4,74E+06 | 5 | 4 | 3 | 2 | 5 | | 26677 | PPTG_09511 \| Phytophthora parasitica INRA-310 Ras-like protein Rab-6A (243 aa) |
| 1997 | 2049 | PPTG_02852 | 85,57 | 8 | 2 | 6 | 8 | 0 | 6,96E+05 | 1,52E+06 | 4 | 4 | 1 | 3 | 4 | | 69387 | PPTG_02852 \| Phytophthora parasitica INRA-310 hypothetical protein (626 aa) |
| 1895 | 2220 | PPTG_19564 | 85,55 | 16 | 9 | 16 | 12 | 3,82E+05 | 5,94E+06 | 4,85E+06 | 3 | 3 | 2 | 4 | 3 | | 33051 | PPTG_19564 \| Phytophthora parasitica INRA-310 hypothetical protein (304 aa) |
| 1550 | 2015 | PPTG_01420 | 85,52 | 39 | 32 | 32 | 39 | 2,23E+06 | 5,55E+06 | 2,34E+07 | 4 | 4 | 3 | 5 | 5 | | 12364 | PPTG_01420 \| Phytophthora parasitica INRA-310 Rab5 family GTPase (113 aa) |
| 2504 | 2333 | PPTG_06870 | 85,52 | 6 | 2 | 6 | 2 | 0 | 3,48E+05 | 3,12E+05 | 3 | 3 | 1 | 3 | 1 | Carbamidomethylation | 76693 | PPTG_06870 \| Phytophthora parasitica INRA-310 methylenetetrahydrofolate reductase (678 aa) |
| 1893 | 1870 | PPTG_15433 | 85,48 | 9 | 2 | 5 | 8 | 4,99E+04 | 7,66E+05 | 2,39E+06 | 6 | 6 | 1 | 3 | 5 | Carbamidomethylation | 96761 | PPTG_15433 \| Phytophthora parasitica INRA-310 regulator-G protein signaling (861 aa) |
| 2121 | 1683 | PPTG_10512 | 85,44 | 9 | 0 | 6 | 5 | | 8,62E+05 | 6,85E+05 | 6 | 6 | 0 | 4 | 3 | | 82667 | PPTG_10512 \| Phytophthora parasitica INRA-310 hypothetical protein (736 aa) |
| 2010 | 1906 | PPTG_09378 | 85,42 | 4 | 0 | 3 | 4 | | 1,75E+06 | 2,82E+06 | 5 | 5 | 0 | 3 | 5 | Carbamidomethylation | 138231 | PPTG_09378 \| Phytophthora parasitica INRA-310 splicing factor 3B subunit 1 (1235 aa) |
| 1748 | 1540 | PPTG_07947 | 85,35 | 6 | 1 | 2 | 6 | 9,12E+04 | 1,01E+06 | 3,68E+06 | 6 | 6 | 1 | 2 | 7 | Oxidation (M) | 146504 | PPTG_07947 \| Phytophthora parasitica INRA-310 hypothetical protein (1317 aa) |
| 1336 | 2465 | PPTG_08061 | 85,32 | 27 | 27 | 26 | 27 | 1,62E+07 | 2,37E+07 | 1,49E+08 | 4 | 4 | 5 | 4 | 7 | Deamidation (NQ) | 12036 | PPTG_08061 \| Phytophthora parasitica INRA-310 hypothetical protein (108 aa) |
| 2235 | 2128 | PPTG_07850 | 85,3 | 39 | 8 | 39 | 24 | 1,39E+05 | 9,81E+05 | 2,92E+05 | 4 | 4 | 1 | 4 | 2 | | 13057 | PPTG_07850 \| Phytophthora parasitica INRA-310 hypothetical protein (115 aa) |
| 2039 | 2076 | PPTG_16421 | 85,18 | 6 | 2 | 6 | 5 | 8,30E+04 | 1,34E+06 | 2,00E+06 | 4 | 4 | 1 | 4 | 3 | Carbamidomethylation | 80317 | PPTG_16421 \| Phytophthora parasitica INRA-310 hypothetical protein (742 aa) |
| 2274 | 2310 | PPTG_04170 | 85,13 | 13 | 4 | 13 | 9 | 2,61E+04 | 2,11E+06 | 3,15E+06 | 3 | 3 | 1 | 3 | 2 | | 34965 | PPTG_04170 \| Phytophthora parasitica INRA-310 hypothetical protein (319 aa) |
| 2342 | 2046 | PPTG_16176 | 85,13 | 5 | 2 | 2 | 5 | 0 | 0 | 2,42E+06 | 4 | 4 | 1 | 1 | 4 | | 100957 | PPTG_16176 \| Phytophthora parasitica INRA-310 mannosyl-oligosaccharide glucosidase (893 aa) |
| 2122 | 1909 | PPTG_15126 | 85,09 | 12 | 0 | 9 | 12 | | 7,79E+05 | 1,38E+06 | 4 | 4 | 0 | 3 | 4 | Carbamidomethylation | 50300 | PPTG_15126 \| Phytophthora parasitica INRA-310 hypothetical protein (467 aa) |
| 1637 | 2501 | PPTG_07773 | 85,09 | 11 | 0 | 11 | 7 | | 3,59E+06 | 3,50E+06 | 3 | 3 | 0 | 7 | 5 | Oxidation (M); Carbamidomethylation | 54250 | PPTG_07773 \| Phytophthora parasitica INRA-310 hypothetical protein (514 aa) |
| 2767 | 2515 | PPTG_18029 | 85,08 | 5 | 0 | 5 | 1 | | 1,56E+06 | 5,51E+05 | 3 | 3 | 0 | 3 | 1 | Carbamidomethylation | 141696 | PPTG_18029 \| Phytophthora parasitica INRA-310 tripeptidyl-peptidase (1342 aa) |
| 1697 | 1710 | PPTG_19827 | 85,06 | 14 | 3 | 10 | 14 | 0 | 4,53E+06 | 7,62E+06 | 6 | 6 | 1 | 4 | 6 | Carbamidomethylation; Acetylation (N-term) | 49231 | PPTG_19827 \| Phytophthora parasitica INRA-310 imidazoleglycerol-phosphate dehydratase (465 aa) |
| 1676 | 1518 | PPTG_14584 | 85,04 | 36 | 6 | 21 | 36 | 6,42E+05 | 8,00E+06 | 9,66E+06 | 6 | 6 | 1 | 4 | 6 | Deamidation (NQ) | 21981 | PPTG_14584 \| Phytophthora parasitica INRA-310 small COPII coat GTPase (192 aa) |
| 1747 | 1746 | PPTG_02596 | 84,97 | 2 | 0 | 2 | 1 | | 7,35E+05 | 1,07E+06 | 8 | 8 | 0 | 6 | 4 | | 446782 | PPTG_02596 \| Phytophthora parasitica INRA-310 DNA-dependent protein kinase catalytic subunit (4000 aa) |
| 1542 | 1814 | PPTG_18304 | 84,93 | 22 | 0 | 13 | 22 | | 5,61E+06 | 7,41E+06 | 5 | 5 | 0 | 6 | 7 | Oxidation (M) | 30497 | PPTG_18304 \| Phytophthora parasitica INRA-310 dihydrolipoamide succinyltransferase (287 aa) |
| 2220 | 2105 | PPTG_04609 | 84,93 | 27 | 9 | 27 | 18 | 8,49E+04 | 2,27E+06 | 1,53E+06 | 4 | 4 | 1 | 4 | 2 | Carbamidomethylation | 20296 | PPTG_04609 \| Phytophthora parasitica INRA-310 hypothetical protein (194 aa) |
| 2236 | 2221 | PPTG_00610 | 84,71 | 8 | 0 | 7 | 4 | | 8,50E+05 | 1,23E+05 | 5 | 4 | 0 | 4 | 3 | | 81125 | PPTG_00610 \| Phytophthora parasitica INRA-310 protein kinase (749 aa) |
| 2976 | 2763 | PPTG_18794 | 84,7 | 11 | 0 | 11 | 0 | | 6,80E+05 | | 3 | 3 | 0 | 3 | 0 | | 38681 | PPTG_18794 \| Phytophthora parasitica INRA-310 arabinogalactan endo-1 4-beta-galactosidase (359 aa) |
| 2469 | 2429 | PPTG_10910 | 84,65 | 16 | 0 | 9 | 10 | | 9,95E+05 | 1,96E+06 | 4 | 4 | 0 | 2 | 3 | | 36779 | PPTG_10910 \| Phytophthora parasitica INRA-310 Delta(3 5)-Delta(2 4)-dienoyl-CoA isomerase (336 aa) |
| 1774 | 1758 | PPTG_08016 | 84,61 | 3 | 1 | 2 | 2 | 2,08E+05 | 7,94E+05 | 1,87E+06 | 6 | 6 | 2 | 3 | 5 | | 231710 | PPTG_08016 \| Phytophthora parasitica INRA-310 hypothetical protein (2084 aa) |
| 2183 | 1900 | PPTG_12354 | 84,57 | 19 | 4 | 4 | 19 | 1,38E+05 | 0 | 4,91E+06 | 5 | 5 | 1 | 1 | 5 | Acetylation (N-term) | 39701 | PPTG_12354 \| Phytophthora parasitica INRA-310 mitochondrial Carrier (MC) family (362 aa) |
| 3251 | 3434 | PPTG_03489 | 84,57 | 30 | 0 | 21 | 30 | | 6,79E+05 | 2,27E+05 | 2 | 2 | 0 | 1 | 2 | Carbamidomethylation | 13696 | PPTG_03489 \| Phytophthora parasitica INRA-310 protease inhibitor EpiC1 (127 aa) |
| 2234 | 2028 | PPTG_15637 | 84,54 | 36 | 0 | 26 | 28 | | 1,53E+06 | 2,68E+06 | 4 | 4 | 0 | 4 | 3 | Deamidation (NQ) | 14899 | PPTG_15637 \| Phytophthora parasitica INRA-310 hypothetical protein (131 aa) |
| 1240 | 1620 | PPTG_12010 | 84,49 | 13 | 3 | 9 | 10 | 2,48E+05 | 4,19E+06 | 1,47E+07 | 8 | 6 | 2 | 6 | 9 | Oxidation (M); Carbamidomethylation | 65499 | PPTG_12010 \| Phytophthora parasitica INRA-310 beta-glucosidase (583 aa) |
| 2018 | 1574 | PPTG_09872 | 84,49 | 14 | 3 | 3 | 14 | 4,34E+05 | 1,88E+06 | 8,36E+06 | 6 | 6 | 1 | 1 | 6 | Carbamidomethylation; Dehydration; Acetylation (N-term) | 51183 | PPTG_09872 \| Phytophthora parasitica INRA-310 hypothetical protein (474 aa) |
| 1953 | 1858 | PPTG_02271 | 84,48 | 16 | 0 | 16 | 12 | | 2,06E+06 | 3,09E+06 | 5 | 5 | 0 | 5 | 4 | | 49158 | PPTG_02271 \| Phytophthora parasitica INRA-310 mannose-1-phosphate guanyltransferase alpha (445 aa) |
| 2287 | 1706 | PPTG_10228 | 84,46 | 2 | 0 | 0 | 2 | | | 2,00E+06 | 6 | 6 | 0 | 0 | 6 | | 343759 | PPTG_10228 \| Phytophthora parasitica INRA-310 hypothetical protein (3250 aa) |
| 2309 | 2746 | PPTG_20356 | 84,45 | 21 | 8 | 15 | 21 | 4,67E+04 | 5,56E+05 | 7,39E+05 | 3 | 3 | 1 | 2 | 3 | Oxidation (M); Deamidation (NQ) | 25065 | PPTG_20356 \| Phytophthora parasitica INRA-310 hypothetical protein (227 aa) |
| 2041 | 2120 | PPTG_16959 | 84,4 | 8 | 2 | 4 | 8 | 4,11E+05 | 1,17E+06 | 4,59E+06 | 4 | 4 | 1 | 2 | 5 | Oxidation (M); Carbamidomethylation | 70051 | PPTG_16959 \| Phytophthora parasitica INRA-310 hypothetical protein (644 aa) |
| 1959 | 2453 | PPTG_04846 | 84,36 | 17 | 0 | 13 | 17 | | 3,67E+06 | 2,53E+06 | 4 | 4 | 0 | 4 | 5 | Oxidation (M); Carbamidomethylation | 28596 | PPTG_04846 \| Phytophthora parasitica INRA-310 ribulose-phosphate 3-epimerase (264 aa) |
| 2451 | 2141 | PPTG_10687 | 84,23 | 12 | 0 | 10 | 4 | | 1,93E+05 | 6,58E+05 | 4 | 4 | 0 | 3 | 2 | Carbamidomethylation | 42477 | PPTG_10687 \| Phytophthora parasitica INRA-310 formate dehydrogenase (387 aa) |
| 1978 | 1752 | PPTG_11445 | 84,18 | 9 | 1 | 0 | 9 | 4,87E+04 | | 2,79E+06 | 6 | 6 | 1 | 0 | 7 | Oxidation (M) | 86416 | PPTG_11445 \| Phytophthora parasitica INRA-310 hypothetical protein (781 aa) |
| 2258 | 3351 | PPTG_10259 | 84,17 | 19 | 19 | 19 | 19 | 4,07E+05 | 3,67E+05 | 1,95E+06 | 2 | 2 | 2 | 2 | 3 | Carbamidomethylation | 24681 | PPTG_10259 \| Phytophthora parasitica INRA-310 phosphatidylinositol synthase (PIS) (218 aa) |
| 2733 | 1866 | PPTG_14643 | 84,08 | 7 | 0 | 0 | 7 | | | 2,61E+06 | 4 | 4 | 0 | 0 | 4 | Oxidation (M); Carbamidomethylation | 81235 | PPTG_14643 \| Phytophthora parasitica INRA-310 protein kinase (762 aa) |
| 1752 | 1740 | PPTG_06261 | 84,04 | 3 | 1 | 2 | 3 | 6,80E+04 | 1,07E+06 | 2,35E+06 | 5 | 5 | 1 | 4 | 5 | | 241736 | PPTG_06261 \| Phytophthora parasitica INRA-310 DEAD/DEAH box RNA helicase (2168 aa) |
| 1881 | 1774 | PPTG_10833 | 84,04 | 7 | 1 | 5 | 7 | 3,18E+04 | 3,15E+05 | 1,93E+06 | 5 | 5 | 1 | 3 | 5 | | 97613 | PPTG_10833 \| Phytophthora parasitica INRA-310 hypothetical protein (923 aa) |
| 1974 | 1650 | PPTG_02206 | 83,94 | 8 | 0 | 5 | 5 | | 8,99E+05 | 2,19E+06 | 7 | 7 | 0 | 4 | 4 | | 99693 | PPTG_02206 \| Phytophthora parasitica INRA-310 pre-mRNA-splicing factor SYF1-like protein (865 aa) |
| 1536 | 1833 | PPTG_15071 | 83,93 | 27 | 4 | 27 | 27 | 1,99E+05 | 3,58E+06 | 6,40E+06 | 5 | 5 | 1 | 6 | 6 | Carbamidomethylation; Deamidation (NQ) | 22128 | PPTG_15071 \| Phytophthora parasitica INRA-310 hypothetical protein (212 aa) |
| 2516 | 2182 | PPTG_11043 | 83,91 | 13 | 3 | 3 | 10 | 1,10E+05 | 3,43E+05 | 3,39E+06 | 4 | 4 | 1 | 1 | 3 | Carbamidomethylation | 47357 | PPTG_11043 \| Phytophthora parasitica INRA-310 transmembrane protein (432 aa) |
| 1886 | 1692 | PPTG_14667 | 83,9 | 3 | 0 | 0 | 3 | | 7,30E+04 | 2,32E+06 | 8 | 8 | 0 | 1 | 8 | | 297402 | PPTG_14667 \| Phytophthora parasitica INRA-310 hypothetical protein (2667 aa) |
| 2288 | 1741 | PPTG_16983 | 83,89 | 8 | 2 | 0 | 6 | 1,78E+04 | | 1,94E+06 | 6 | 6 | 1 | 0 | 5 | Deamidation (NQ) | 120477 | PPTG_16983 \| Phytophthora parasitica INRA-310 hypothetical protein (1085 aa) |
| 1914 | 1729 | PPTG_12611 | 83,89 | 8 | 1 | 5 | 8 | 6,54E+05 | 3,06E+06 | 9,38E+06 | 5 | 5 | 1 | 3 | 5 | | 86816 | PPTG_12611 \| Phytophthora parasitica INRA-310 hypothetical protein (812 aa) |
| 1726 | 1845 | PPTG_02659 | 83,83 | 8 | 5 | 3 | 8 | 1,03E+06 | 8,84E+05 | 1,24E+07 | 5 | 5 | 3 | 2 | 6 | Carbamidomethylation | 73334 | PPTG_02659 \| Phytophthora parasitica INRA-310 hypothetical protein (675 aa) |
| 2429 | 2482 | PPTG_03823 | 83,79 | 23 | 0 | 23 | 23 | | 1,32E+06 | 1,95E+06 | 3 | 3 | 0 | 3 | 3 | Carbamidomethylation; Acetylation (N-term) | 28418 | PPTG_03823 \| Phytophthora parasitica INRA-310 actin-like protein (258 aa) |
| 2562 | 1819 | PPTG_09234 | 83,74 | 12 | 0 | 0 | 12 | | | 1,59E+06 | 5 | 5 | 0 | 0 | 5 | Carbamidomethylation | 50433 | PPTG_09234 \| Phytophthora parasitica INRA-310 hypothetical protein (467 aa) |
| 2545 | 2564 | PPTG_12907 | 83,73 | 27 | 0 | 14 | 20 | | 1,14E+06 | 3,66E+06 | 4 | 4 | 0 | 2 | 3 | Carbamidomethylation | 22971 | PPTG_12907 \| Phytophthora parasitica INRA-310 hypothetical protein (213 aa) |
| 1471 | 2395 | PPTG_15628 | 83,71 | 39 | 21 | 39 | 39 | 9,49E+05 | 5,90E+06 | 9,21E+06 | 4 | 4 | 3 | 6 | 5 | Deamidation (NQ) | 12210 | PPTG_15628 \| Phytophthora parasitica INRA-310 small nuclear ribonucleoprotein Sm D1 (113 aa) |
| 1432 | 1550 | PPTG_09861 | 83,7 | 8 | 3 | 8 | 8 | | 5,78E+05 | 9,77E+05 | 6 | 1 | 2 | 6 | 6 | Oxidation (M); Carbamidomethylation | 80260 | PPTG_09861 \| Phytophthora parasitica INRA-310 galactokinase (731 aa) |
| 2157 | 1855 | PPTG_01352 | 83,69 | 10 | 0 | 10 | 8 | | 1,05E+06 | 7,98E+05 | 4 | 4 | 0 | 4 | 3 | | 57723 | PPTG_01352 \| Phytophthora parasitica INRA-310 arginine n-methyltransferase (507 aa) |
| 1323 | 1799 | PPTG_13220 | 83,69 | 4 | 1 | 3 | 3 | 3,22E+05 | 2,72E+06 | 5,17E+06 | 8 | 8 | 2 | 7 | 7 | Carbamidomethylation | 234322 | PPTG_13220 \| Phytophthora parasitica INRA-310 hypothetical protein (2145 aa) |
| 1796 | 1864 | PPTG_18406 | 83,63 | 22 | 9 | 17 | 19 | 6,13E+05 | 2,89E+06 | 7,15E+06 | 5 | 5 | 2 | 4 | 4 | | 31216 | PPTG_18406 \| Phytophthora parasitica INRA-310 peroxisomal trans-2-enoyl-CoA reductase (293 aa) |
| 1701 | 1962 | PPTG_09020 | 83,61 | 12 | 0 | 10 | 9 | | 1,44E+06 | 6,20E+05 | 4 | 4 | 0 | 7 | 4 | Carbamidomethylation; Deamidation (NQ) | 40019 | PPTG_09020 \| Phytophthora parasitica INRA-310 hypothetical protein (356 aa) |
| 1965 | 1822 | PPTG_12758 | 83,55 | 2 | 0 | 1 | 1 | | 1,42E+05 | 3,00E+05 | 7 | 5 | 0 | 5 | 3 | | 535673 | PPTG_12758 \| Phytophthora parasitica INRA-310 dynein heavy chain outer arm (4752 aa) |
| 2100 | 2344 | PPTG_04847 | 83,53 | 42 | 30 | 42 | 42 | 6,27E+05 | 5,40E+06 | 7,65E+06 | 3 | 3 | 2 | 3 | 3 | Acetylation (N-term) | 10214 | PPTG_04847 \| Phytophthora parasitica INRA-310 hypothetical protein (91 aa) |
| 2189 | 2342 | PPTG_00052 | 83,53 | 14 | 10 | 10 | 14 | 4,52E+05 | 1,75E+05 | 4,16E+06 | 3 | 3 | 2 | 2 | 3 | Oxidation (M) | 31387 | PPTG_00052 \| Phytophthora parasitica INRA-310 retinol dehydrogenase (284 aa) |
| 1892 | 1831 | PPTG_00633 | 83,39 | 18 | 3 | 6 | 18 | 1,14E+05 | 5,19E+05 | 9,80E+06 | 6 | 6 | 1 | 2 | 6 | Carbamidomethylation; Dehydration; Deamidation (NQ) | 42304 | PPTG_00633 \| Phytophthora parasitica INRA-310 XRP2-like protein (384 aa) |
| 1798 | 2070 | PPTG_09015 | 83,39 | 13 | 6 | 11 | 11 | 6,58E+04 | 7,64E+05 | 8,48E+06 | 4 | 4 | 3 | 3 | 4 | | 48798 | PPTG_09015 \| Phytophthora parasitica INRA-310 transmembrane protein (433 aa) |
| 1939 | 2293 | PPTG_00576 | 83,37 | 32 | 8 | 32 | 25 | 4,01E+05 | 4,65E+06 | 4,02E+06 | 4 | 4 | 1 | 5 | 3 | Carbamidomethylation | 19552 | PPTG_00576 \| Phytophthora parasitica INRA-310 hypothetical protein (180 aa) |
| 1792 | 1994 | PPTG_17697 | 83,37 | 6 | 2 | 6 | 6 | 2,04E+05 | 1,72E+06 | 2,33E+06 | 4 | 4 | 1 | 5 | 4 | Carbamidomethylation | 77595 | PPTG_17697 \| Phytophthora parasitica INRA-310 choline/Carnitine O-acyltransferase (694 aa) |
| 1913 | 1728 | PPTG_02399 | 83,37 | 6 | 0 | 4 | 5 | | 1,31E+06 | 2,48E+06 | 6 | 6 | 0 | 4 | 5 | | 129380 | PPTG_02399 \| Phytophthora parasitica INRA-310 hypothetical protein (1163 aa) |
| 1831 | 2073 | PPTG_01944 | 83,34 | 18 | 11 | 14 | 12 | 2,73E+06 | 2,23E+07 | 1,63E+07 | 5 | 5 | 3 | 4 | 3 | | 28294 | PPTG_01944 \| Phytophthora parasitica INRA-310 hypothetical protein (250 aa) |
| 1862 | 1618 | PPTG_10896 | 83,31 | 11 | 3 | 6 | 10 | 7,05E+05 | 9,08E+06 | 9,23E+06 | 5 | 5 | 1 | 4 | 4 | Oxidation (M); Carbamidomethylation; Dehydration | 55958 | PPTG_10896 \| Phytophthora parasitica INRA-310 ubiquitin carboxyl-terminal hydrolase (505 aa) |
| 2791 | 2148 | PPTG_15954 | 83,24 | 18 | 0 | 14 | 4 | | 6,93E+05 | 0 | 4 | 4 | 0 | 3 | 1 | | 31778 | PPTG_15954 \| Phytophthora parasitica INRA-310 phosphoglycerate mutase family protein (280 aa) |
| 1703 | 2024 | PPTG_01705 | 83,23 | 43 | 7 | 37 | 35 | 8,18E+05 | 7,01E+06 | 1,11E+07 | 5 | 5 | 1 | 5 | 5 | Deamidation (NQ) | 17689 | PPTG_01705 \| Phytophthora parasitica INRA-310 hypothetical protein (155 aa) |
| 1851 | 3415 | PPTG_03073 | 83,23 | 20 | 0 | 15 | 20 | | 1,31E+06 | 1,17E+07 | 2 | 2 | 0 | 3 | 7 | Deamidation (NQ); Acetylation (N-term) | 18918 | PPTG_03073 \| Phytophthora parasitica INRA-310 transmembrane protein (176 aa) |
| 1878 | 2145 | PPTG_15119 | 83,19 | 12 | 7 | 9 | 12 | 2,89E+05 | 2,29E+06 | 4,32E+06 | 4 | 4 | 2 | 3 | 4 | | 38484 | PPTG_15119 \| Phytophthora parasitica INRA-310 hypothetical protein (341 aa) |
| 2843 | 2384 | PPTG_13869 | 83,13 | 15 | 0 | 0 | 15 | | | 6,85E+05 | 3 | 3 | 0 | 0 | 4 | Carbamidomethylation | 38766 | PPTG_13869 \| Phytophthora parasitica INRA-310 hypothetical protein (345 aa) |
| 1655 | 1742 | PPTG_08452 | 83,12 | 11 | 0 | 8 | 9 | | 1,66E+06 | 3,88E+06 | 7 | 7 | 0 | 5 | 6 | | 70291 | PPTG_08452 \| Phytophthora parasitica INRA-310 choline dehydrogenase (642 aa) |
| 2311 | 1882 | PPTG_07791 | 83,08 | 23 | 0 | 23 | 7 | | 1,08E+06 | 7,83E+05 | 4 | 4 | 0 | 4 | 2 | Carbamidomethylation | 33591 | PPTG_07791 \| Phytophthora parasitica INRA-310 hypothetical protein (314 aa) |
| 2615 | 1861 | PPTG_06586 | 82,97 | 8 | 1 | 0 | 7 | | | 4,65E+05 | 5 | 1 | 1 | 0 | 4 | Carbamidomethylation | 75480 | PPTG_06586 \| Phytophthora parasitica INRA-310 transmembrane protein (696 aa) |
| 2360 | 1844 | PPTG_00865 | 82,89 | 14 | 0 | 11 | 5 | | 4,33E+05 | 3,06E+05 | 5 | 5 | 0 | 4 | 2 | Carbamidomethylation | 68095 | PPTG_00865 \| Phytophthora parasitica INRA-310 hypothetical protein (606 aa) |
| 2199 | 2834 | PPTG_11596 | 82,87 | 15 | 11 | 15 | 11 | 7,79E+04 | 1,41E+06 | 4,81E+05 | 2 | 2 | 1 | 4 | 2 | Deamidation (NQ) | 33437 | PPTG_11596 \| Phytophthora parasitica INRA-310 hypothetical protein (301 aa) |
| 2587 | 2361 | PPTG_16542 | 82,86 | 19 | 0 | 12 | 19 | | 1,61E+05 | 2,52E+06 | 3 | 3 | 0 | 2 | 3 | Carbamidomethylation | 32637 | PPTG_16542 \| Phytophthora parasitica INRA-310 stromal cell-derived factor 2 (296 aa) |
| 1367 | 1627 | PPTG_05140 | 82,82 | 9 | 0 | 7 | 7 | | 1,42E+06 | 3,70E+06 | 9 | 9 | 0 | 7 | 8 | Oxidation (M); Carbamidomethylation | 131925 | PPTG_05140 \| Phytophthora parasitica INRA-310 DNA-directed RNA polymerase I subunit RPA2 (1180 aa) |
| 2454 | 2433 | PPTG_17993 | 82,8 | 5 | 0 | 5 | 3 | | 4,16E+05 | 5,77E+05 | 3 | 3 | 0 | 3 | 2 | | 77686 | PPTG_17993 \| Phytophthora parasitica INRA-310 hypothetical protein (719 aa) |
| 1727 | 2217 | PPTG_02283 | 82,6 | 13 | 4 | 13 | 13 | 4,83E+04 | 1,49E+06 | 3,43E+06 | 4 | 4 | 1 | 5 | 5 | Oxidation (M) | 34734 | PPTG_02283 \| Phytophthora parasitica INRA-310 hypothetical protein (312 aa) |
| 2174 | 2066 | PPTG_10990 | 82,57 | 12 | 3 | 7 | 12 | 1,27E+05 | 2,06E+06 | 2,77E+06 | 4 | 4 | 1 | 2 | 4 | Oxidation (M) | 46047 | PPTG_10990 \| Phytophthora parasitica INRA-310 hypothetical protein (400 aa) |
| 2387 | 2246 | PPTG_13052 | 82,5 | 29 | 20 | 11 | 29 | 1,76E+06 | 2,14E+06 | 1,21E+07 | 3 | 3 | 2 | 1 | 3 | | 13579 | PPTG_13052 \| Phytophthora parasitica INRA-310 hypothetical protein (123 aa) |
| 2986 | 3087 | PPTG_14230 | 82,48 | 17 | 0 | 0 | 17 | | | 3,01E+06 | 3 | 3 | 0 | 0 | 3 | | 21618 | PPTG_14230 \| Phytophthora parasitica INRA-310 hypothetical protein (199 aa) |
| 1427 | 1567 | PPTG_09865 | 82,48 | 13 | 5 | 13 | 13 | | 6,48E+05 | 1,67E+06 | 6 | 1 | 2 | 6 | 6 | Oxidation (M); Carbamidomethylation | 45345 | PPTG_09865 \| Phytophthora parasitica INRA-310 galactokinase (420 aa) |
| 1782 | 2050 | PPTG_16685 | 82,46 | 28 | 0 | 23 | 15 | | 8,72E+05 | 2,84E+06 | 7 | 3 | 0 | 6 | 4 | Carbamidomethylation | 38251 | PPTG_16685 \| Phytophthora parasitica INRA-310 alcohol dehydrogenase (360 aa) |
| 2289 | 1913 | PPTG_09203 | 82,44 | 22 | 0 | 13 | 22 | | 5,47E+05 | 2,93E+06 | 4 | 4 | 0 | 2 | 4 | Oxidation (M) | 23289 | PPTG_09203 \| Phytophthora parasitica INRA-310 hypothetical protein (212 aa) |
| 2051 | 1869 | PPTG_06444 | 82,43 | 8 | 1 | 3 | 8 | 7,94E+04 | 6,18E+05 | 3,04E+06 | 5 | 5 | 1 | 2 | 5 | | 82406 | PPTG_06444 \| Phytophthora parasitica INRA-310 hypothetical protein (732 aa) |
| 1854 | 2042 | PPTG_09290 | 82,4 | 24 | 0 | 24 | 24 | | 1,14E+06 | 6,83E+05 | 4 | 4 | 0 | 4 | 6 | Oxidation (M); Deamidation (NQ) | 28880 | PPTG_09290 \| Phytophthora parasitica INRA-310 hypothetical protein (262 aa) |
| 1522 | 2525 | PPTG_01934 | 82,11 | 32 | 15 | 22 | 32 | 1,18E+06 | 1,51E+07 | 2,43E+07 | 3 | 3 | 1 | 5 | 7 | Carbamidomethylation; Deamidation (NQ) | 11904 | PPTG_01934 \| Phytophthora parasitica INRA-310 small cysteine rich protein SCR108 (111 aa) |
| 1908 | 1791 | PPTG_05438 | 82,01 | 6 | 1 | 4 | 6 | 7,41E+04 | 8,24E+05 | 1,71E+06 | 5 | 5 | 1 | 3 | 5 | Carbamidomethylation | 130762 | PPTG_05438 \| Phytophthora parasitica INRA-310 hypothetical protein (1182 aa) |
| 1721 | 1648 | PPTG_00698 | 81,97 | 23 | 0 | 14 | 23 | | 2,48E+06 | 6,89E+06 | 6 | 6 | 0 | 4 | 7 | Oxidation (M) | 33784 | PPTG_00698 \| Phytophthora parasitica INRA-310 Ras-like GTP-binding protein A (294 aa) |
| 1902 | 2157 | PPTG_11880 | 81,95 | 12 | 0 | 7 | 12 | | 6,51E+05 | 3,51E+06 | 6 | 6 | 0 | 3 | 6 | Carbamidomethylation | 53600 | PPTG_11880 \| Phytophthora parasitica INRA-310 calcium-binding atopy-like autoantigen-like protein (462 aa) |
| 2403 | 1847 | PPTG_18120 | 81,91 | 12 | 0 | 11 | 4 | | 1,25E+06 | 2,59E+05 | 5 | 5 | 0 | 4 | 2 | | 54051 | PPTG_18120 \| Phytophthora parasitica INRA-310 hypothetical protein (500 aa) |
| 2392 | 2374 | PPTG_01445 | 81,89 | 28 | 13 | 21 | 28 | 2,43E+05 | 3,87E+06 | 8,21E+06 | 3 | 3 | 1 | 2 | 3 | Carbamidomethylation | 13157 | PPTG_01445 \| Phytophthora parasitica INRA-310 hypothetical protein (119 aa) |
| 1922 | 1682 | PPTG_06545 | 81,89 | 24 | 0 | 19 | 17 | | 1,27E+06 | 2,04E+06 | 6 | 6 | 0 | 5 | 4 | | 40725 | PPTG_06545 \| Phytophthora parasitica INRA-310 D-cysteine desulfhydrase (371 aa) |
| 1330 | 1848 | PPTG_08324 | 81,88 | 17 | 11 | 17 | 15 | 1,83E+05 | 5,50E+06 | 5,75E+06 | 5 | 5 | 3 | 8 | 5 | Oxidation (M) | 42927 | PPTG_08324 \| Phytophthora parasitica INRA-310 lipoamide acyltransferase component-branched-chain alpha-keto acid dehydrogenase complex (400 aa) |

| | | | | | | | | | | | | | | | | | | | |
|---|---|---|---|---|---|---|---|---|---|---|---|---|---|---|---|---|---|---|---|
| 2569 | 2407 | PPTG_08017 | 81,87 | 7 | 0 | 5 | 7 | | 5,27E+05 | 2,10E+06 | 3 | 3 | 0 | 2 | 3 | | 63718 | PPTG_08017 | Phytophthora parasitica INRA-310 NPP1-like protein (617 aa) |
| 1636 | 2398 | PPTG_01812 | 81,86 | 43 | 10 | 10 | 43 | 3,09E+06 | 3,36E+06 | 6,77E+07 | 3 | 3 | 3 | 2 | 7 | Oxidation (M) | 13268 | PPTG_01812 | Phytophthora parasitica INRA-310 hypothetical protein (117 aa) |
| 2026 | 2103 | PPTG_03206 | 81,84 | 5 | 2 | 5 | 4 | 6,95E+04 | 1,26E+06 | 1,48E+06 | 4 | 4 | 1 | 4 | 3 | | 93092 | PPTG_03206 | Phytophthora parasitica INRA-310 hypothetical protein (829 aa) |
| 2001 | 1543 | PPTG_14669 | 81,84 | 5 | 1 | 1 | 5 | 2,23E+04 | 1,18E+05 | 2,36E+06 | 6 | 6 | 1 | 1 | 6 | Carbamidomethylation | 198401 | PPTG_14669 | Phytophthora parasitica INRA-310 hypothetical protein (1833 aa) |
| 2257 | 3184 | PPTG_18384 | 81,8 | 6 | 0 | 6 | 6 | | 2,23E+06 | 1,24E+06 | 2 | 1 | 0 | 4 | 3 | Oxidation (M); Carbamidomethylation | 55711 | PPTG_18384 | Phytophthora parasitica INRA-310 hypothetical protein (542 aa) |
| 1743 | 1810 | PPTG_15736 | 81,73 | 20 | 5 | 16 | 14 | 2,75E+05 | 2,61E+06 | 1,98E+06 | 6 | 6 | 2 | 5 | 4 | Carbamidomethylation | 35088 | PPTG_15736 | Phytophthora parasitica INRA-310 ubiquitin carboxyl-terminal hydrolase (314 aa) |
| 1665 | 1960 | PPTG_05943 | 81,73 | 8 | 2 | 3 | 8 | | | 5,37E+05 | 5 | 2 | 3 | 2 | 6 | Oxidation (M); Deamidation (NQ) | 64109 | PPTG_05943 | Phytophthora parasitica INRA-310 sodium/potassium-transporting ATPase subunit alpha (588 aa) |
| 2247 | 3191 | PPTG_03490 | 81,68 | 21 | 0 | 21 | 10 | | 2,59E+06 | 1,57E+06 | 3 | 3 | 0 | 5 | 2 | Oxidation (M) | 16598 | PPTG_03490 | Phytophthora parasitica INRA-310 hypothetical protein (147 aa) |
| 2734 | 1904 | PPTG_10034 | 81,67 | 8 | 0 | 0 | 8 | | | 1,11E+06 | 4 | 4 | 0 | 0 | 4 | | 97290 | PPTG_10034 | Phytophthora parasitica INRA-310 exocyst complex component (877 aa) |
| 1813 | 2002 | PPTG_08008 | 81,64 | 13 | 2 | 13 | 4 | 1,13E+05 | 2,14E+06 | 6,36E+05 | 6 | 6 | 1 | 7 | 2 | Carbamidomethylation | 59114 | PPTG_08008 | Phytophthora parasitica INRA-310 gamma-glutamyltransferase (557 aa) |
| 2703 | 1860 | PPTG_16378 | 81,61 | 5 | 0 | 0 | 5 | | | 1,02E+06 | 4 | 4 | 0 | 0 | 4 | | 92136 | PPTG_16378 | Phytophthora parasitica INRA-310 hypothetical protein (829 aa) |
| 2720 | 2855 | PPTG_05370 | 81,57 | 14 | 0 | 14 | 3 | | 9,39E+05 | 5,75E+05 | 3 | 3 | 0 | 3 | 1 | Carbamidomethylation | 45746 | PPTG_05370 | Phytophthora parasitica INRA-310 nucleolin (442 aa) |
| 2210 | 2495 | PPTG_04563 | 81,55 | 22 | 8 | 16 | 18 | 1,22E+05 | 1,91E+06 | 2,92E+06 | 4 | 4 | 1 | 3 | 3 | Carbamidomethylation | 25567 | PPTG_04563 | Phytophthora parasitica INRA-310 U1 small nuclear ribonucleoprotein A (234 aa) |
| 1404 | 2378 | PPTG_11387 | 81,52 | 13 | 10 | 13 | 13 | 4,83E+06 | 2,59E+07 | 3,39E+07 | 3 | 1 | 4 | 6 | 5 | Oxidation (M); Carbamidomethylation; Carbamylation | 23046 | PPTG_11387 | Phytophthora parasitica INRA-310 hypothetical protein (225 aa) |
| 2509 | 2600 | PPTG_15759 | 81,49 | 15 | 0 | 10 | 15 | | 1,58E+06 | 2,09E+06 | 3 | 3 | 0 | 2 | 3 | Carbamidomethylation | 30746 | PPTG_15759 | Phytophthora parasitica INRA-310 hypothetical protein (288 aa) |
| 2204 | 2136 | PPTG_17833 | 81,44 | 15 | 2 | 15 | 7 | 8,59E+04 | 1,02E+06 | 9,74E+05 | 4 | 4 | 1 | 4 | 2 | | 38723 | PPTG_17833 | Phytophthora parasitica INRA-310 tRNA (adenine-N(1)-)-methyltransferase catalytic subunit (349 aa) |
| 2076 | 2415 | PPTG_05234 | 81,28 | 29 | 12 | 29 | 29 | 4,62E+05 | 5,64E+06 | 7,78E+06 | 3 | 3 | 1 | 4 | 3 | | 15204 | PPTG_05234 | Phytophthora parasitica INRA-310 hypothetical protein (130 aa) |
| 1679 | 2187 | PPTG_12253 | 81,21 | 10 | 4 | 9 | 10 | 1,13E+05 | 6,34E+05 | 1,61E+06 | 5 | 5 | 2 | 4 | 5 | | 56228 | PPTG_12253 | Phytophthora parasitica INRA-310 GTP-binding protein (515 aa) |
| 1987 | 1753 | PPTG_18911 | 81,21 | 25 | 0 | 25 | 15 | | 1,44E+06 | 1,38E+06 | 5 | 5 | 0 | 5 | 3 | | 35377 | PPTG_18911 | Phytophthora parasitica INRA-310 translation elongation factor Ts (332 aa) |
| 2327 | 2140 | PPTG_06266 | 81,16 | 32 | 0 | 6 | 32 | | 6,72E+05 | 3,50E+06 | 5 | 5 | 0 | 1 | 5 | Carbamidomethylation; Acetylation (N-term) | 21197 | PPTG_06266 | Phytophthora parasitica INRA-310 hypothetical protein (187 aa) |
| 1027 | 3345 | PPTG_19861 | 81,13 | 38 | 38 | 38 | 38 | 4,03E+06 | 5,71E+07 | 5,42E+07 | 2 | 2 | 3 | 9 | 10 | Oxidation (M); Carbamidomethylation; Deamidation (NQ) | 12336 | PPTG_19861 | Phytophthora parasitica INRA-310 elicitin (119 aa) |
| 1027 | 3346 | PPTG_09075 | 81,13 | 38 | 38 | 38 | 38 | 4,03E+06 | 5,71E+07 | 5,42E+07 | 2 | 2 | 3 | 9 | 10 | Oxidation (M); Carbamidomethylation; Deamidation (NQ) | 12336 | PPTG_09075 | Phytophthora parasitica INRA-310 elicitin (119 aa) |
| 2173 | 1965 | PPTG_12659 | 81,12 | 18 | 3 | 3 | 18 | 1,14E+05 0 | | 2,12E+06 | 5 | 5 | 1 | 1 | 5 | | 34719 | PPTG_12659 | Phytophthora parasitica INRA-310 NADH-cytochrome b5 reductase (316 aa) |
| 2585 | 2129 | PPTG_02564 | 81,1 | 15 | 0 | 15 | 3 | | 1,93E+06 0 | | 4 | 4 | 0 | 4 | 1 | Carbamylation | 42381 | PPTG_02564 | Phytophthora parasitica INRA-310 hypothetical protein (390 aa) |
| 2561 | 1786 | PPTG_13077 | 81,05 | 6 | 0 | 0 | 6 | | | 1,23E+06 | 5 | 5 | 0 | 0 | 5 | | 138446 | PPTG_13077 | Phytophthora parasitica INRA-310 phosphatidylinositol-4-phosphate-5-kinase (PIPK-D7/GPCR-PIPK) (1239 aa) |
| 1824 | 1828 | PPTG_19139 | 81,04 | 30 | 0 | 27 | 20 | | 4,23E+06 | 5,32E+06 | 6 | 6 | 0 | 5 | 5 | Carbamidomethylation; Acetylation (N-term) | 30383 | PPTG_19139 | Phytophthora parasitica INRA-310 radial spoke protein (269 aa) |
| 1824 | 1829 | PPTG_19139 | 81,04 | 30 | 0 | 27 | 20 | | 4,23E+06 | 5,32E+06 | 6 | 6 | 0 | 5 | 5 | Carbamidomethylation; Acetylation (N-term) | 30383 | PPTG_19139 | Phytophthora parasitica INRA-310 radial spoke protein (269 aa) |
| 1988 | 1817 | PPTG_13557 | 81,02 | 15 | 0 | 15 | 15 | | 9,74E+05 | 2,12E+06 | 4 | 4 | 0 | 4 | 4 | | 41663 | PPTG_13557 | Phytophthora parasitica INRA-310 serine/threonine-protein phosphatase 2A regulatory subunit B' (373 aa) |
| 2112 | 1686 | PPTG_07437 | 80,99 | 6 | 0 | 1 | 6 | | 7,99E+04 | 4,04E+06 | 6 | 4 | 0 | 1 | 6 | Carbamidomethylation; Acetylation (N-term) | 133604 | PPTG_07437 | Phytophthora parasitica INRA-310 cleavage induced protein kinase (1193 aa) |
| 1519 | 1750 | PPTG_14065 | 80,94 | 9 | 2 | 7 | 7 | 6,27E+04 | 1,27E+06 | 2,24E+06 | 7 | 7 | 1 | 6 | 6 | Carbamidomethylation | 87562 | PPTG_14065 | Phytophthora parasitica INRA-310 Cullin family protein (762 aa) |
| 1829 | 2020 | PPTG_05794 | 80,89 | 11 | 0 | 11 | 6 | | 6,13E+06 | 3,90E+06 | 4 | 4 | 0 | 6 | 4 | Carbamidomethylation; Deamidation (NQ) | 59417 | PPTG_05794 | Phytophthora parasitica INRA-310 beta-glucosidase (534 aa) |
| 2069 | 1961 | PPTG_13512 | 80,85 | 12 | 0 | 12 | 12 | | 1,91E+06 | 3,05E+06 | 4 | 4 | 0 | 4 | 4 | | 46617 | PPTG_13512 | Phytophthora parasitica INRA-310 kynurenine-oxoglutarate transaminase (416 aa) |
| 1916 | 1881 | PPTG_11398 | 80,83 | 6 | 2 | 2 | 6 | 1,53E+05 | 8,37E+05 | 3,40E+06 | 5 | 5 | 2 | 2 | 5 | Carbamidomethylation | 89908 | PPTG_11398 | Phytophthora parasitica INRA-310 hypothetical protein (802 aa) |
| 2517 | 2316 | PPTG_09597 | 80,83 | 11 | 0 | 11 | 7 | | 8,86E+05 | 7,29E+05 | 3 | 3 | 0 | 3 | 2 | | 38855 | PPTG_09597 | Phytophthora parasitica INRA-310 D-isomer specific 2-hydroxyacid dehydrogenase (353 aa) |
| 1639 | 1625 | PPTG_16999 | 80,8 | 3 | 1 | 1 | 3 | 2,04E+05 | 4,05E+05 | 1,79E+06 | 6 | 6 | 2 | 3 | 6 | Carbamidomethylation | 283529 | PPTG_16999 | Phytophthora parasitica INRA-310 hypothetical protein (2534 aa) |
| 3152 | 3344 | PPTG_14727 | 80,76 | 24 | 0 | 24 | 13 | | 5,06E+06 | 2,22E+06 | 2 | 2 | 0 | 2 | 1 | Carbamidomethylation | 12225 | PPTG_14727 | Phytophthora parasitica INRA-310 hypothetical protein (113 aa) |
| 1733 | 2427 | PPTG_18860 | 80,74 | 38 | 12 | 38 | 38 | 2,89E+05 | 2,85E+06 | 6,70E+06 | 3 | 3 | 1 | 5 | 5 | Oxidation (M); Carbamidomethylation | 11233 | PPTG_18860 | Phytophthora parasitica INRA-310 mitochondrial protein Translocase (MPT) family (104 aa) |
| 1837 | 2813 | PPTG_14822 | 80,65 | 28 | 13 | 28 | 28 | 2,85E+05 | 1,40E+06 | 2,69E+06 | 4 | 4 | 2 | 4 | 4 | Carbamidomethylation | 13460 | PPTG_14822 | Phytophthora parasitica INRA-310 autophagy-like protein (117 aa) |
| 2406 | 2262 | PPTG_15412 | 80,62 | 10 | 0 | 10 | 2 | | 9,12E+05 | 2,73E+05 | 5 | 5 | 0 | 5 | 1 | Carbamidomethylation | 70837 | PPTG_15412 | Phytophthora parasitica INRA-310 hypothetical protein (663 aa) |
| 2284 | 2526 | PPTG_05469 | 80,59 | 7 | 2 | 7 | 2 | 1,59E+05 | 2,14E+06 | 1,41E+06 | 4 | 4 | 1 | 4 | 1 | | 82303 | PPTG_05469 | Phytophthora parasitica INRA-310 oligopeptidase A (733 aa) |
| 2130 | 2197 | PPTG_15985 | 80,56 | 19 | 0 | 6 | 19 | | 5,02E+05 | 1,68E+06 | 4 | 3 | 0 | 2 | 5 | Oxidation (M); Acetylation (N-term) | 29084 | PPTG_15985 | Phytophthora parasitica INRA-310 hypothetical protein (277 aa) |
| 1915 | 1780 | PPTG_04944 | 80,53 | 5 | 0 | 3 | 4 | | 6,26E+05 | 2,37E+06 | 6 | 6 | 0 | 3 | 6 | Oxidation (M) | 150637 | PPTG_04944 | Phytophthora parasitica INRA-310 hypothetical protein (1356 aa) |
| 3248 | 3385 | PPTG_04881 | 80,5 | 8 | 3 | 0 | 8 | 4,86E+05 | | 5,33E+06 | 2 | 2 | 1 | 0 | 2 | | 31566 | PPTG_04881 | Phytophthora parasitica INRA-310 hypothetical protein (298 aa) |
| 2127 | 1958 | PPTG_03889 | 80,5 | 13 | 1 | 4 | 10 | 5,77E+04 | 1,77E+05 | 1,41E+06 | 6 | 6 | 1 | 2 | 4 | | 65980 | PPTG_03889 | Phytophthora parasitica INRA-310 ATPase family AAA domain-containing protein 3A (585 aa) |
| 1840 | 1769 | PPTG_09552 | 80,45 | 13 | 6 | 2 | 13 | 7,84E+05 | 2,14E+05 | 5,02E+06 | 6 | 6 | 3 | 1 | 6 | Oxidation (M) | 60355 | PPTG_09552 | Phytophthora parasitica INRA-310 solute carrier family facilitated glucose transporter (556 aa) |
| 2464 | 2201 | PPTG_02391 | 80,41 | 8 | 0 | 1 | 8 | | 2,38E+05 | 2,88E+06 | 4 | 4 | 0 | 1 | 4 | Carbamidomethylation | 74919 | PPTG_02391 | Phytophthora parasitica INRA-310 choline transporter-like protein (689 aa) |
| 2063 | 1621 | PPTG_02656 | 80,38 | 12 | 1 | 0 | 12 | 1,62E+05 | | 2,13E+06 | 6 | 6 | 2 | 0 | 6 | Deamidation (NQ) | 68839 | PPTG_02656 | Phytophthora parasitica INRA-310 calcineurin-like phosphoesterase (613 aa) |
| 2460 | 2630 | PPTG_18144 | 80,36 | 11 | 2 | 2 | 11 | | | 1,19E+05 | 3 | 1 | 1 | 1 | 3 | Oxidation (M) | 57839 | PPTG_18144 | Phytophthora parasitica INRA-310 ATP-binding Cassette (ABC) superfamily (529 aa) |
| 2014 | 2181 | PPTG_11817 | 80,27 | 10 | 0 | 9 | 7 | | 8,39E+05 | 6,97E+05 | 5 | 5 | 0 | 4 | 4 | | 68606 | PPTG_11817 | Phytophthora parasitica INRA-310 NAD-dependent epimerase/dehydratase (623 aa) |
| 1715 | 1812 | PPTG_18350 | 80,27 | 30 | 6 | 30 | 20 | 1,84E+05 | 2,57E+06 | 2,83E+06 | 6 | 6 | 1 | 6 | 4 | | 23386 | PPTG_18350 | Phytophthora parasitica INRA-310 hypothetical protein (210 aa) |
| 2339 | 1923 | PPTG_16605 | 80,25 | 8 | 0 | 4 | 6 | | 4,11E+05 | 8,28E+05 | 4 | 4 | 0 | 2 | 4 | | 73606 | PPTG_16605 | Phytophthora parasitica INRA-310 SEC1 family transporter SLY1 (659 aa) |
| 2082 | 1925 | PPTG_05014 | 80,12 | 13 | 0 | 13 | 7 | | 2,81E+06 | 3,46E+06 | 5 | 5 | 0 | 5 | 3 | Carbamidomethylation | 55340 | PPTG_05014 | Phytophthora parasitica INRA-310 D-2-hydroxyglutarate dehydrogenase (502 aa) |
| 1825 | 1887 | PPTG_02088 | 80,1 | 19 | 4 | 14 | 19 | 8,22E+05 | 6,59E+06 | 1,21E+07 | 4 | 4 | 1 | 4 | 5 | Oxidation (M); Carbamidomethylation | 26235 | PPTG_02088 | Phytophthora parasitica INRA-310 hypothetical protein (241 aa) |
| 2191 | 2347 | PPTG_19546 | 80,1 | 10 | 3 | 10 | 10 | 1,95E+05 | 2,36E+06 | 3,34E+06 | 3 | 3 | 1 | 3 | 3 | Acetylation (N-term) | 43365 | PPTG_19546 | Phytophthora parasitica INRA-310 hypothetical protein (415 aa) |
| 1638 | 3342 | PPTG_01442 | 80,07 | 26 | 26 | 26 | 26 | 4,55E+06 | 3,47E+06 | 4,36E+07 | 2 | 2 | 3 | 3 | 6 | Oxidation (M); Acetylation (N-term) | 11401 | PPTG_01442 | Phytophthora parasitica INRA-310 hypothetical protein (106 aa) |
| 2163 | 2207 | PPTG_00097 | 80,07 | 7 | 3 | 3 | 6 | 2,64E+05 | 6,60E+05 | 2,37E+06 | 4 | 4 | 2 | 2 | 3 | | 65235 | PPTG_00097 | Phytophthora parasitica INRA-310 ATP-binding Cassette (ABC) superfamily (589 aa) |
| 2097 | 2089 | PPTG_14858 | 80,04 | 16 | 0 | 12 | 12 | | 7,44E+05 | 1,59E+06 | 5 | 5 | 0 | 4 | 4 | Acetylation (N-term) | 39359 | PPTG_14858 | Phytophthora parasitica INRA-310 hypothetical protein (355 aa) |
| 2109 | 1605 | PPTG_16085 | 79,95 | 1 | 0 | 0 | 1 | | 1,19E+05 | 1,23E+06 | 6 | 6 | 0 | 1 | 6 | | 690078 | PPTG_16085 | Phytophthora parasitica INRA-310 hypothetical protein (6449 aa) |
| 2070 | 2019 | PPTG_10432 | 79,9 | 11 | 5 | 11 | 5 | 5,89E+05 | 4,41E+06 | 4,75E+06 | 4 | 4 | 2 | 4 | 2 | | 45293 | PPTG_10432 | Phytophthora parasitica INRA-310 hypothetical protein (425 aa) |
| 2730 | 3177 | PPTG_17186 | 79,84 | 8 | 0 | 8 | 8 | | 7,56E+05 | 4,48E+05 | 2 | 2 | 0 | 2 | 2 | Carbamidomethylation | 43678 | PPTG_17186 | Phytophthora parasitica INRA-310 glycoside hydrolase (418 aa) |
| 1884 | 2456 | PPTG_00090 | 79,77 | 10 | 7 | 7 | 10 | 2,00E+05 | 7,20E+05 | 1,92E+06 | 3 | 1 | 3 | 3 | 3 | Oxidation (M) | 39602 | PPTG_00090 | Phytophthora parasitica INRA-310 coproporphyrinogen III oxidase aerobic (345 aa) |
| 2550 | 2737 | PPTG_04719 | 79,75 | 13 | 0 | 13 | 11 | | 2,91E+06 | 1,23E+06 | 3 | 3 | 0 | 3 | 2 | | 38376 | PPTG_04719 | Phytophthora parasitica INRA-310 arginase (356 aa) |
| 1361 | 1854 | PPTG_05007 | 79,73 | 6 | 0 | 4 | 3 | | 1,48E+06 | 2,88E+06 | 10 | 10 | 0 | 8 | 7 | Oxidation (M); Carbamidomethylation | 206016 | PPTG_05007 | Phytophthora parasitica INRA-310 hypothetical protein (1852 aa) |
| 3146 | 3193 | PPTG_01683 | 79,7 | 44 | 0 | 0 | 44 | | | 9,37E+05 | 2 | 2 | 0 | 0 | 3 | Oxidation (M); Carbamidomethylation; Acetylation (N-term) | 9387 | PPTG_01683 | Phytophthora parasitica INRA-310 hypothetical protein (87 aa) |
| 1702 | 2023 | PPTG_05095 | 79,58 | 25 | 4 | 20 | 25 | 7,06E+04 | 1,28E+06 | 3,13E+06 | 5 | 5 | 1 | 4 | 6 | Oxidation (M) | 28271 | PPTG_05095 | Phytophthora parasitica INRA-310 C2 domain-containing protein (249 aa) |
| 1823 | 1827 | PPTG_04021 | 79,57 | 13 | 0 | 13 | 13 | | 7,38E+05 | 7,79E+05 | 5 | 5 | 0 | 5 | 5 | | 60097 | PPTG_04021 | Phytophthora parasitica INRA-310 hypothetical protein (542 aa) |
| 2195 | 2551 | PPTG_07273 | 79,46 | 11 | 0 | 6 | 11 | | 2,54E+06 | 4,23E+06 | 4 | 4 | 0 | 3 | 4 | Carbamidomethylation; Deamidation (NQ) | 39046 | PPTG_07273 | Phytophthora parasitica INRA-310 GMP reductase (363 aa) |
| 2761 | 1943 | PPTG_14040 | 79,45 | 5 | 0 | 0 | 5 | | | 2,96E+06 | 4 | 4 | 0 | 0 | 4 | | 125080 | PPTG_14040 | Phytophthora parasitica INRA-310 hypothetical protein (1100 aa) |
| 1671 | 1616 | PPTG_11249 | 79,44 | 21 | 6 | 10 | 21 | 7,40E+05 | 4,73E+06 | 1,07E+07 | 5 | 5 | 2 | 3 | 6 | Deamidation (NQ) | 34085 | PPTG_11249 | Phytophthora parasitica INRA-310 hypothetical protein (313 aa) |
| 2203 | 2034 | PPTG_08960 | 79,38 | 9 | 0 | 9 | 8 | | 2,08E+06 | 7,59E+05 | 4 | 4 | 0 | 4 | 3 | Carbamidomethylation; Acetylation (N-term) | 65857 | PPTG_08960 | Phytophthora parasitica INRA-310 dynein-like protein (590 aa) |
| 2283 | 2488 | PPTG_15982 | 79,31 | 14 | 0 | 6 | 14 | | 9,78E+05 | 1,87E+06 | 3 | 2 | 0 | 2 | 4 | Oxidation (M) | 29011 | PPTG_15982 | Phytophthora parasitica INRA-310 carbohydrate esterase (280 aa) |
| 1628 | 1702 | PPTG_15050 | 79,31 | 10 | 3 | 7 | 10 | 3,87E+05 | 4,37E+05 | 3,68E+06 | 6 | 6 | 2 | 4 | 6 | Carbamidomethylation | 80133 | PPTG_15050 | Phytophthora parasitica INRA-310 copine-like protein (725 aa) |
| 1900 | 1749 | PPTG_15519 | 79,27 | 12 | 3 | 5 | 12 | 2,02E+05 | 6,13E+05 | 7,67E+06 | 5 | 5 | 1 | 2 | 6 | Oxidation (M); Carbamidomethylation | 77395 | PPTG_15519 | Phytophthora parasitica INRA-310 protein kinase (704 aa) |
| 2066 | 1805 | PPTG_09204 | 79,26 | 14 | 3 | 5 | 14 | 5,97E+04 | 6,34E+05 | 4,91E+06 | 5 | 5 | 1 | 2 | 5 | | 42188 | PPTG_09204 | Phytophthora parasitica INRA-310 mitochondrial Carrier (MC) family (387 aa) |
| 1784 | 2681 | PPTG_00813 | 79,08 | 30 | 6 | 14 | 30 | 3,56E+05 | 6,41E+05 | 8,25E+06 | 6 | 6 | 1 | 2 | 7 | | 14797 | PPTG_00813 | Phytophthora parasitica INRA-310 hypothetical protein (133 aa) |
| 2367 | 2413 | PPTG_05325 | 79 | 20 | 3 | 10 | 20 | 4,13E+05 | 1,89E+06 | 8,74E+06 | 3 | 3 | 1 | 2 | 3 | Oxidation (M) | 27079 | PPTG_05325 | Phytophthora parasitica INRA-310 C2 domain-containing protein (241 aa) |
| 1962 | 2188 | PPTG_16761 | 78,98 | 15 | 7 | 7 | 15 | 4,18E+05 | 1,45E+06 | 8,14E+06 | 4 | 4 | 3 | 2 | 4 | | 35516 | PPTG_16761 | Phytophthora parasitica INRA-310 mitochondrial protein Translocase (MPT) family (323 aa) |
| 1853 | 3212 | PPTG_17175 | 78,95 | 28 | 13 | 28 | 28 | 9,77E+05 | 1,09E+07 | 8,93E+06 | 3 | 3 | 2 | 4 | 4 | | 10582 | PPTG_17175 | Phytophthora parasitica INRA-310 hypothetical protein (96 aa) |
| 2390 | 2302 | PPTG_00667 | 78,86 | 23 | 7 | 17 | 23 | 4,20E+04 | 4,36E+05 | 1,14E+06 | 3 | 3 | 1 | 2 | 3 | | 20832 | PPTG_00667 | Phytophthora parasitica INRA-310 ADP-ribosylation factor-like protein 2 (187 aa) |
| 1985 | 2980 | PPTG_14129 | 78,85 | 6 | 2 | 6 | 6 | 7,54E+04 | 9,75E+05 | 3,35E+06 | 2 | 2 | 1 | 3 | 4 | Oxidation (M) | 47290 | PPTG_14129 | Phytophthora parasitica INRA-310 hypothetical protein (426 aa) |
| 2048 | 1821 | PPTG_14747 | 78,82 | 23 | 0 | 23 | 5 | | 1,85E+06 | 3,74E+05 | 5 | 5 | 0 | 6 | 2 | Oxidation (M); Deamidation (NQ) | 34431 | PPTG_14747 | Phytophthora parasitica INRA-310 cysteine protease family C26 (315 aa) |
| 2316 | 2185 | PPTG_16125 | 78,78 | 9 | 0 | 7 | 7 | | 2,88E+06 | 1,69E+06 | 4 | 4 | 0 | 3 | 3 | | 65827 | PPTG_16125 | Phytophthora parasitica INRA-310 hypothetical protein (596 aa) |
| 1949 | 3343 | PPTG_18572 | 78,73 | 17 | 17 | 17 | 17 | 3,73E+06 | 6,00E+06 | 3,84E+07 | 2 | 2 | 2 | 3 | 4 | Carbamidomethylation | 17352 | PPTG_18572 | Phytophthora parasitica INRA-310 transmembrane protein (166 aa) |
| 2185 | 2072 | PPTG_10841 | 78,72 | 6 | 1 | 5 | 2 | 7,48E+04 | 7,19E+05 | 6,67E+05 | 5 | 5 | 1 | 4 | 2 | Carbamidomethylation | 120592 | PPTG_10841 | Phytophthora parasitica INRA-310 dihydroflavonol-4-reductase (1089 aa) |
| 1331 | 2463 | PPTG_08961 | 78,68 | 43 | 32 | 43 | 43 | 3,49E+06 | 6,58E+06 | 4,14E+07 | 3 | 3 | 3 | 7 | 6 | Oxidation (M) | 8779 | PPTG_08961 | Phytophthora parasitica INRA-310 hypothetical protein (78 aa) |
| 1627 | 3384 | PPTG_09434 | 78,63 | 7 | 7 | 7 | 3 | 2,37E+05 | 1,87E+05 | | 3 | 1 | 4 | 4 | 4 | Oxidation (M); Carbamylation | 55909 | PPTG_09434 | Phytophthora parasitica INRA-310 sugar transporter (512 aa) |
| 2340 | 2005 | PPTG_08124 | 78,61 | 8 | 1 | 5 | 4 | 1,22E+05 | 9,94E+05 | 7,90E+05 | 4 | 4 | 1 | 3 | 2 | Carbamidomethylation | 73600 | PPTG_08124 | Phytophthora parasitica INRA-310 hypothetical protein (654 aa) |
| 2340 | 2007 | PPTG_08124 | 78,61 | 7 | 1 | 5 | 3 | 1,22E+05 | 9,94E+05 | 7,90E+05 | 4 | 4 | 1 | 3 | 2 | Carbamidomethylation | 77178 | PPTG_08124 | Phytophthora parasitica INRA-310 hypothetical protein (687 aa) |
| 2588 | 2386 | PPTG_18095 | 78,57 | 41 | 0 | 41 | 31 | | 2,23E+06 | 5,90E+05 | 3 | 3 | 0 | 3 | 2 | Oxidation (M); Acetylation (N-term) | 12157 | PPTG_18095 | Phytophthora parasitica INRA-310 hypothetical protein (114 aa) |
| 1450 | 1783 | PPTG_12167 | 78,53 | 12 | 5 | 4 | 12 | 5,37E+05 | 4,14E+05 | 4,66E+06 | 7 | 7 | 3 | 3 | 8 | Oxidation (M); Deamidation (NQ) | 68146 | PPTG_12167 | Phytophthora parasitica INRA-310 ATP-binding Cassette (ABC) superfamily (614 aa) |
| 1901 | 2018 | PPTG_17605 | 78,5 | 12 | 5 | 12 | 10 | 2,73E+05 | 2,55E+06 | 2,18E+06 | 4 | 4 | 2 | 4 | 3 | | 51835 | PPTG_17605 | Phytophthora parasitica INRA-310 adenosine deaminase (456 aa) |
| 2087 | 2195 | PPTG_07848 | 78,45 | 18 | 2 | 16 | 9 | 9,29E+05 | 1,05E+07 | 7,80E+06 | 4 | 4 | 1 | 4 | 3 | | 41345 | PPTG_07848 | Phytophthora parasitica INRA-310 hypothetical protein (384 aa) |
| 2344 | 2095 | PPTG_11282 | 78,41 | 34 | 0 | 8 | 34 | | 2,53E+05 | 3,76E+06 | 5 | 5 | 0 | 1 | 5 | | 27505 | PPTG_11282 | Phytophthora parasitica INRA-310 hypothetical protein (251 aa) |
| 1749 | 1852 | PPTG_08759 | 78,39 | 6 | 0 | 4 | 4 | | 1,88E+06 | 3,38E+06 | 7 | 7 | 0 | 6 | 4 | | 141177 | PPTG_08759 | Phytophthora parasitica INRA-310 hypothetical protein (1272 aa) |
| 1692 | 1918 | PPTG_11915 | 78,38 | 14 | 2 | 6 | 14 | | | 1,42E+05 | 4 | 1 | 1 | 3 | 7 | Oxidation (M); Carbamidomethylation | 37123 | PPTG_11915 | Phytophthora parasitica INRA-310 fructose-1 6-bisphosphatase (337 aa) |
| 2404 | 2117 | PPTG_17559 | 78,36 | 14 | 0 | 14 | 8 | | 1,26E+06 | 7,20E+05 | 4 | 4 | 0 | 4 | 2 | | 40222 | PPTG_17559 | Phytophthora parasitica INRA-310 hypothetical protein (372 aa) |
| 2506 | 2539 | PPTG_09605 | 78,27 | 16 | 0 | 14 | 2 | | 6,58E+05 | 2,87E+05 | 4 | 4 | 0 | 4 | 1 | Deamidation (NQ) | 43980 | PPTG_09605 | Phytophthora parasitica INRA-310 hypothetical protein (398 aa) |
| 1969 | 2455 | PPTG_02110 | 78,13 | 2 | 2 | 2 | 2 | 1,22E+05 | 9,87E+05 | 2,24E+06 | 3 | 3 | 2 | 3 | 3 | | 145910 | PPTG_02110 | Phytophthora parasitica INRA-310 inositol-3 4-bisphosphate 4-phosphatase (1322 aa) |
| 2027 | 2115 | PPTG_15082 | 78,1 | 13 | 0 | 13 | 7 |  | 1,55E+06 | 2,92E+06 | 5 | 5 | 0 | 4 | 4 |  | 45477 | PPTG_15082 | Phytophthora parasitica INRA-310 phosphoglycerate kinase chloroplastic (429 aa) |
| 2208 | 2451 | PPTG_16748 | 78,07 | 8 | 2 | 8 | 8 | 1,01E+05 | 1,39E+06 | 1,72E+06 | 3 | 3 | 1 | 3 | 3 | | 43949 | PPTG_16748 | Phytophthora parasitica INRA-310 cysteine desulfurase (411 aa) |
| 1941 | 1721 | PPTG_05091 | 78,03 | 28 | 4 | 4 | 28 | 3,85E+05 | 2,83E+05 | 6,31E+06 | 6 | 6 | 1 | 1 | 7 | Oxidation (M); Carbamidomethylation | 28670 | PPTG_05091 | Phytophthora parasitica INRA-310 hypothetical protein (256 aa) |
| 2108 | 1863 | PPTG_00428 | 78,02 | 2 | 0 | 1 | 2 | | 2,90E+05 | 1,54E+06 | 5 | 5 | 0 | 2 | 5 | | 355462 | PPTG_00428 | Phytophthora parasitica INRA-310 vacuolar protein sorting-associated protein (3248 aa) |
| 2417 | 2808 | PPTG_04036 | 77,96 | 49 | 30 | 21 | 39 | 1,02E+06 | 1,65E+06 | 5,84E+06 | 4 | 4 | 2 | 1 | 3 |  | 12555 | PPTG_04036 | Phytophthora parasitica INRA-310 V-type ATPase F subunit (116 aa) |
| 2622 | 2671 | PPTG_18386 | 77,94 | 8 | 3 | 6 | 5 | | 1,74E+06 | | 3 | 1 | 1 | 2 | 2 | Carbamidomethylation | 46700 | PPTG_18386 | Phytophthora parasitica INRA-310 carbohydrate-binding protein (446 aa) |
| 2320 | 2356 | PPTG_04838 | 77,92 | 12 | 0 | 9 | 9 | | 7,70E+05 | 9,46E+05 | 4 | 4 | 0 | 3 | 3 | Oxidation (M); Carbamidomethylation | 45309 | PPTG_04838 | Phytophthora parasitica INRA-310 angio-associated migratory cell protein (419 aa) |
| 2149 | 1879 | PPTG_16585 | 77,91 | 12 | 0 | 12 | 9 | | 9,40E+05 | 9,43E+05 | 4 | 4 | 0 | 4 | 3 | Carbamidomethylation | 43247 | PPTG_16585 | Phytophthora parasitica INRA-310 hypothetical protein (397 aa) |

| | | | | | | | | | | | | | | | | | | | |
|---|---|---|---|---|---|---|---|---|---|---|---|---|---|---|---|---|---|---|---|
| 2357 | 2914 | PPTG_19719 | 77,85 | 11 | 0 | 11 | 11 | | 1,15E+06 | 1,46E+06 | 2 | 2 | 0 | 3 | 3 | Oxidation (M); Carbamidomethylation | 32601 | PPTG_19719 | Phytophthora parasitica INRA-310 hypothetical protein (323 aa) |
| 2679 | 2574 | PPTG_18416 | 77,85 | 11 | 3 | 0 | 9 | 1,08E+05 | | 1,44E+06 | 4 | 4 | 1 | 0 | 3 | | 47516 | PPTG_18416 | Phytophthora parasitica INRA-310 hypothetical protein (432 aa) |
| 2380 | 3167 | PPTG_10453 | 77,82 | 16 | 8 | 16 | 15 | 2,19E+04 | 8,57E+05 | 6,15E+05 | 3 | 3 | 1 | 3 | 2 | Carbamidomethylation | 20921 | PPTG_10453 | Phytophthora parasitica INRA-310 hypothetical protein (189 aa) |
| 2708 | 2279 | PPTG_02906 | 77,8 | 10 | 0 | 7 | 8 | | 2,99E+05 | 1,83E+05 | 3 | 3 | 0 | 2 | 2 | | 43800 | PPTG_02906 | Phytophthora parasitica INRA-310 5'-AMP-activated protein kinase subunit gamma (394 aa) |
| 2271 | 2253 | PPTG_13430 | 77,8 | 6 | 0 | 4 | 3 | | 6,17E+05 | 6,58E+05 | 5 | 4 | 0 | 3 | 3 | Carbamidomethylation | 122726 | PPTG_13430 | Phytophthora parasitica INRA-310 N-acetyltransferase 10 (1096 aa) |
| 1717 | 1986 | PPTG_14264 | 77,75 | 20 | 2 | 17 | 8 | 2,89E+05 | 1,92E+07 | 3,60E+06 | 7 | 7 | 1 | 6 | 4 | Oxidation (M) | 37201 | PPTG_14264 | Phytophthora parasitica INRA-310 hypothetical protein (336 aa) |
| 1725 | 1838 | PPTG_11003 | 77,74 | 5 | 2 | 3 | 5 | 1,52E+05 | 8,23E+05 | 2,64E+06 | 5 | 5 | 2 | 3 | 6 | Oxidation (M) | 99599 | PPTG_11003 | Phytophthora parasitica INRA-310 ATP-binding cassette sub-family F member 1 (881 aa) |
| 2175 | 2071 | PPTG_19350 | 77,69 | 12 | 0 | 9 | 12 | | 5,65E+06 | 1,33E+07 | 4 | 4 | 0 | 3 | 4 | | 43590 | PPTG_19350 | Phytophthora parasitica INRA-310 protein kinase (387 aa) |
| 2792 | 2158 | PPTG_11440 | 77,69 | 6 | 0 | 0 | 6 | | | 1,43E+06 | 4 | 4 | 0 | 0 | 4 | | 86648 | PPTG_11440 | Phytophthora parasitica INRA-310 hypothetical protein (782 aa) |
| 2294 | 2251 | PPTG_00376 | 77,68 | 14 | 0 | 10 | 5 | | 8,45E+05 | 1,44E+05 | 5 | 4 | 0 | 4 | 2 | Carbamidomethylation | 47592 | PPTG_00376 | Phytophthora parasitica INRA-310 isocitrate dehydrogenase [NADP] (423 aa) |
| 1800 | 2276 | PPTG_01635 | 77,65 | 37 | 7 | 28 | 29 | 7,83E+05 | 1,08E+07 | 1,06E+07 | 4 | 4 | 1 | 3 | 6 | Oxidation (M); Deamidation (NQ) | 12094 | PPTG_01635 | Phytophthora parasitica INRA-310 peptidyl-prolyl isomerase FKBP12 (113 aa) |
| 3010 | 2513 | PPTG_05219 | 77,61 | 15 | 0 | 0 | 15 | | | 9,18E+05 | 3 | 3 | 0 | 0 | 3 | Carbamidomethylation | 32896 | PPTG_05219 | Phytophthora parasitica INRA-310 maspardin-like protein (289 aa) |
| 1843 | 1910 | PPTG_11184 | 77,61 | 7 | 1 | 5 | 7 | 5,92E+04 | 5,98E+05 | 1,10E+06 | 5 | 4 | 1 | 4 | 5 | Carbamidomethylation | 77234 | PPTG_11184 | Phytophthora parasitica INRA-310 glucan 1 3-beta-glucosidase (681 aa) |
| 2268 | 2214 | PPTG_16019 | 77,59 | 5 | 0 | 3 | 4 | | 6,43E+05 | 3,74E+05 | 4 | 4 | 0 | 3 | 3 | Carbamidomethylation | 132175 | PPTG_16019 | Phytophthora parasitica INRA-310 RNA polymerase-associated protein CTR9 (1164 aa) |
| 2690 | 1995 | PPTG_08437 | 77,55 | 6 | 0 | 0 | 6 | | | 1,74E+06 | 4 | 4 | 0 | 0 | 4 | | 88207 | PPTG_08437 | Phytophthora parasitica INRA-310 hypothetical protein (782 aa) |
| 2426 | 2218 | PPTG_18368 | 77,51 | 16 | 0 | 16 | 6 | | 9,76E+05 | 3,37E+05 | 4 | 4 | 0 | 4 | 2 | | 45512 | PPTG_18368 | Phytophthora parasitica INRA-310 mannan endo-1 4-beta-mannosidase (420 aa) |
| 2124 | 2008 | PPTG_12140 | 77,31 | 5 | 0 | 5 | 2 | | 2,26E+06 | 7,10E+05 | 5 | 5 | 0 | 5 | 2 | | 132237 | PPTG_12140 | Phytophthora parasitica INRA-310 hypothetical protein (1212 aa) |
| 2038 | 2025 | PPTG_14022 | 77,27 | 12 | 4 | 4 | 9 | 1,62E+05 | 2,92E+05 | 1,62E+06 | 5 | 5 | 2 | 2 | 4 | | 50501 | PPTG_14022 | Phytophthora parasitica INRA-310 soluble calcium-activated nucleotidase (457 aa) |
| 1845 | 2059 | PPTG_05444 | 77,27 | 46 | 0 | 46 | 38 | | 5,61E+06 | 6,04E+06 | 4 | 4 | 0 | 6 | 4 | Oxidation (M) | 12860 | PPTG_05444 | Phytophthora parasitica INRA-310 hypothetical protein (107 aa) |
| 1845 | 2060 | PPTG_05446 | 77,27 | 46 | 0 | 46 | 38 | | 5,61E+06 | 6,04E+06 | 4 | 4 | 0 | 6 | 4 | Oxidation (M) | 12860 | PPTG_05446 | Phytophthora parasitica INRA-310 hypothetical protein (107 aa) |
| 2383 | 1931 | PPTG_10089 | 77,26 | 20 | 3 | 7 | 12 | 5,08E+04 | 9,21E+04 | 6,19E+06 | 5 | 5 | 1 | 1 | 4 | Carbamidomethylation | 41929 | PPTG_10089 | Phytophthora parasitica INRA-310 serine protease family S01B (393 aa) |
| 1588 | 1781 | PPTG_11891 | 77,24 | 12 | 4 | 8 | 10 | 4,19E+05 | 2,42E+06 | 3,11E+06 | 6 | 6 | 2 | 4 | 6 | Oxidation (M); Carbamidomethylation | 63282 | PPTG_11891 | Phytophthora parasitica INRA-310 N(2) N(2)-dimethylguanosine tRNA methyltransferase (568 aa) |
| 2202 | 1912 | PPTG_13183 | 77,13 | 19 | 0 | 7 | 17 | | 1,28E+05 | 2,49E+06 | 5 | 5 | 0 | 2 | 5 | Carbamidomethylation | 34984 | PPTG_13183 | Phytophthora parasitica INRA-310 hypothetical protein (331 aa) |
| 2407 | 2362 | PPTG_16055 | 77,12 | 6 | 2 | 5 | 6 | 1,14E+05 | 9,35E+05 | 1,10E+06 | 3 | 3 | 1 | 2 | 3 | | 72448 | PPTG_16055 | Phytophthora parasitica INRA-310 peptidase (652 aa) |
| 2020 | 1824 | PPTG_11777 | 77,12 | 7 | 0 | 3 | 6 | | 2,69E+05 | 1,90E+06 | 6 | 6 | 0 | 3 | 5 | Carbamidomethylation | 90143 | PPTG_11777 | Phytophthora parasitica INRA-310 alpha-N-acetylglucosaminidase (NAGLU) (783 aa) |
| 2618 | 2422 | PPTG_05250 | 77,12 | 20 | 5 | 20 | 5 | 7,34E+04 | 2,98E+06 | 2,77E+05 | 3 | 3 | 1 | 3 | 1 | Carbamidomethylation | 26217 | PPTG_05250 | Phytophthora parasitica INRA-310 elicitin-like mating protein M25 (236 aa) |
| 2616 | 2126 | PPTG_12228 | 77,07 | 26 | 0 | 0 | 26 | | | 1,05E+06 | 4 | 4 | 0 | 0 | 5 | | 36203 | PPTG_12228 | Phytophthora parasitica INRA-310 hypothetical protein (341 aa) |
| 2147 | 2382 | PPTG_00230 | 77,04 | 8 | 3 | 8 | 8 | 1,03E+05 | 8,09E+05 | 1,31E+06 | 3 | 3 | 1 | 3 | 3 | Carbamidomethylation | 52823 | PPTG_00230 | Phytophthora parasitica INRA-310 hypothetical protein (496 aa) |
| 1944 | 2040 | PPTG_05488 | 77,04 | 14 | 7 | 11 | 14 | 2,12E+05 | 9,26E+05 | 1,60E+06 | 4 | 4 | 2 | 3 | 4 | | 41978 | PPTG_05488 | Phytophthora parasitica INRA-310 hypothetical protein (372 aa) |
| 2463 | 3149 | PPTG_19721 | 77,04 | 8 | 3 | 8 | 5 | 2,94E+05 | 4,83E+06 | 2,10E+06 | 2 | 2 | 1 | 3 | 1 | Carbamidomethylation | 43648 | PPTG_19721 | Phytophthora parasitica INRA-310 carbohydrate esterase (396 aa) |
| 1945 | 2097 | PPTG_08156 | 77 | 7 | 0 | 5 | 7 | | 2,22E+06 | 3,32E+06 | 4 | 4 | 0 | 4 | 5 | Carbamidomethylation | 70127 | PPTG_08156 | Phytophthora parasitica INRA-310 hypothetical protein (645 aa) |
| 2280 | 2258 | PPTG_16972 | 76,8 | 3 | 0 | 3 | 3 | | 1,18E+06 | 2,15E+06 | 3 | 3 | 0 | 3 | 3 | | 127628 | PPTG_16972 | Phytophthora parasitica INRA-310 nucleolar protein 6 (1129 aa) |
| 1393 | 2294 | PPTG_13477 | 76,78 | 46 | 37 | 37 | 37 | 7,79E+06 | 2,00E+07 | 1,60E+08 | 4 | 4 | 5 | 4 | 6 | Oxidation (M) | 8323 | PPTG_13477 | Phytophthora parasitica INRA-310 H - or Na -translocating F-type V-type and A-type ATPase (F-ATPase) superfamily (77 aa) |
| 2503 | 2175 | PPTG_06403 | 76,72 | 30 | 0 | 0 | 30 | | | 1,92E+06 | 4 | 4 | 0 | 0 | 5 | Carbamidomethylation; Deamidation (NQ) | 17451 | PPTG_06403 | Phytophthora parasitica INRA-310 hypothetical protein (161 aa) |
| 2349 | 2590 | PPTG_19079 | 76,65 | 24 | 0 | 24 | 6 | | 2,09E+06 | 2,53E+05 | 3 | 3 | 0 | 5 | 1 | Oxidation (M) | 18817 | PPTG_19079 | Phytophthora parasitica INRA-310 hypothetical protein (177 aa) |
| 2096 | 2085 | PPTG_11355 | 76,56 | 10 | 4 | 4 | 9 | 3,64E+05 | 3,20E+07 | 2,43E+06 | 5 | 5 | 2 | 2 | 4 | Oxidation (M); Carbamidomethylation | 54222 | PPTG_11355 | Phytophthora parasitica INRA-310 ethanolamine-phosphate cytidylyltransferase (471 aa) |
| 2053 | 2012 | PPTG_05619 | 76,52 | 15 | 6 | 7 | 15 | 4,95E+05 | 6,84E+05 | 3,73E+06 | 4 | 4 | 2 | 2 | 4 | Carbamidomethylation | 42352 | PPTG_05619 | Phytophthora parasitica INRA-310 aspartate-semialdehyde dehydrogenase (397 aa) |
| 2345 | 2166 | PPTG_11056 | 76,49 | 6 | 0 | 1 | 6 | | 2,29E+05 | 2,70E+06 | 5 | 5 | 0 | 1 | 5 | Carbamidomethylation | 134881 | PPTG_11056 | Phytophthora parasitica INRA-310 tenascin-like protein (1243 aa) |
| 2532 | 3115 | PPTG_00296 | 76,45 | 23 | 9 | 23 | 23 | 2,09E+05 | 1,18E+06 | 1,36E+06 | 2 | 2 | 1 | 2 | 2 | | 14023 | PPTG_00296 | Phytophthora parasitica INRA-310 pterin-4-alpha-carbinolamine dehydratase (122 aa) |
| 2049 | 1830 | PPTG_02738 | 76,44 | 15 | 0 | 13 | 3 | | 8,66E+05 | 7,15E+05 | 6 | 5 | 0 | 6 | 2 | Carbamidomethylation; Deamidation (NQ) | 74075 | PPTG_02738 | Phytophthora parasitica INRA-310 catalase/peroxidase HPI (688 aa) |
| 2137 | 2223 | PPTG_02947 | 76,43 | 8 | 0 | 8 | 6 | | 2,80E+06 | 2,27E+06 | 4 | 3 | 0 | 4 | 3 | Carbamidomethylation | 75055 | PPTG_02947 | Phytophthora parasitica INRA-310 polysaccharide lyase (711 aa) |
| 1964 | 1691 | PPTG_10274 | 76,37 | 2 | 0 | 1 | 1 | | 8,67E+05 | 1,90E+06 | 6 | 6 | 0 | 3 | 5 | | 483731 | PPTG_10274 | Phytophthora parasitica INRA-310 sporangia induced dynein heavy chain (4262 aa) |
| 2461 | 2801 | PPTG_10767 | 76,3 | 9 | 0 | 9 | 9 | | 9,30E+05 | 6,36E+05 | 2 | 2 | 0 | 3 | 2 | Carbamidomethylation | 41523 | PPTG_10767 | Phytophthora parasitica INRA-310 hypothetical protein (384 aa) |
| 2005 | 2423 | PPTG_08953 | 76,25 | 9 | 2 | 9 | 9 | 3,95E+06 | 4,04E+07 | 8,43E+07 | 3 | 3 | 1 | 4 | 3 | Oxidation (M); Carbamidomethylation | 36365 | PPTG_08953 | Phytophthora parasitica INRA-310 hypothetical protein (342 aa) |
| 2489 | 2048 | PPTG_16052 | 76,17 | 7 | 0 | 3 | 6 | | 5,54E+05 | 3,61E+05 | 4 | 4 | 0 | 2 | 3 | | 63157 | PPTG_16052 | Phytophthora parasitica INRA-310 hypothetical protein (565 aa) |
| 2844 | 2409 | PPTG_11896 | 76,15 | 8 | 0 | 3 | 8 | | 0 | 3,84E+05 | 3 | 3 | 0 | 1 | 3 | | 49100 | PPTG_11896 | Phytophthora parasitica INRA-310 DNA primase eukaryotic-type small subunit (429 aa) |
| 2083 | 1996 | PPTG_18189 | 76,12 | 23 | 4 | 15 | 16 | 1,43E+05 | 6,91E+05 | 1,91E+07 | 5 | 5 | 1 | 3 | 4 | Carbamidomethylation; Deamidation (NQ) | 28500 | PPTG_18189 | Phytophthora parasitica INRA-310 hypothetical protein (251 aa) |
| 1552 | 2637 | PPTG_11386 | 76,12 | 15 | 5 | 15 | 8 | | 1,24E+05 | | 3 | 1 | 3 | 6 | 4 | Oxidation (M); Carbamidomethylation; Acetylation (N-term); Carbamylation | 25602 | PPTG_11386 | Phytophthora parasitica INRA-310 hypothetical protein (242 aa) |
| 1937 | 2027 | PPTG_02694 | 76,08 | 11 | 2 | 2 | 11 | 1,42E+05 | 3,41E+05 | 6,13E+06 | 5 | 5 | 1 | 1 | 7 | Oxidation (M); Carbamidomethylation; Deamidation (NQ) | 58456 | PPTG_02694 | Phytophthora parasitica INRA-310 hypothetical protein (530 aa) |
| 2262 | 2454 | PPTG_14912 | 76,08 | 17 | 7 | 14 | 8 | 1,05E+06 | 4,54E+06 | 6,04E+06 | 4 | 4 | 2 | 3 | 2 | Carbamidomethylation | 32946 | PPTG_14912 | Phytophthora parasitica INRA-310 hypothetical protein (289 aa) |
| 2400 | 3350 | PPTG_07488 | 76,02 | 17 | 7 | 17 | 7 | 3,65E+06 | 2,56E+07 | 3,04E+07 | 2 | 2 | 1 | 3 | 2 | Carbamidomethylation; Deamidation (NQ) | 20004 | PPTG_07488 | Phytophthora parasitica INRA-310 hypothetical protein (194 aa) |
| 2800 | 2550 | PPTG_02395 | 76,02 | 4 | 1 | 0 | 4 | 1,52E+05 | | 9,27E+05 | 3 | 3 | 1 | 0 | 3 | Oxidation (M) | 105818 | PPTG_02395 | Phytophthora parasitica INRA-310 phosphatidylinositol-4-phosphate-5-kinase (Pi-PIPK-D2) (935 aa) |
| 2178 | 2345 | PPTG_08320 | 75,97 | 16 | 4 | 16 | 16 | 6,16E+05 | 2,97E+06 | 4,97E+06 | 3 | 3 | 1 | 3 | 3 | | 26596 | PPTG_08320 | Phytophthora parasitica INRA-310 dihydrolipoyllysine-residue acetyltransferase component-pyruvate dehydrogenase complex (252 aa) |
| 2839 | 1892 | PPTG_06087 | 75,93 | 13 | 0 | 0 | 13 | | | 2,38E+06 | 4 | 4 | 0 | 0 | 4 | | 37144 | PPTG_06087 | Phytophthora parasitica INRA-310 dolichyl-phosphate beta-glucosyltransferase (335 aa) |
| 2197 | 2672 | PPTG_14483 | 75,89 | 4 | 0 | 4 | 1 | | 4,17E+05 | | 4 | 2 | 0 | 5 | 2 | Oxidation (M) | 88923 | PPTG_14483 | Phytophthora parasitica INRA-310 glycoside hydrolase (812 aa) |
| 2343 | 2062 | PPTG_16111 | 75,85 | 17 | 0 | 17 | 11 | | 1,39E+06 | 6,01E+05 | 4 | 4 | 0 | 4 | 2 | | 38248 | PPTG_16111 | Phytophthora parasitica INRA-310 ATP-binding Cassette (ABC) superfamily (348 aa) |
| 2337 | 1850 | PPTG_18574 | 75,8 | 8 | 0 | 2 | 8 | | 1,11E+05 | 1,54E+06 | 5 | 5 | 0 | 1 | 5 | | 70296 | PPTG_18574 | Phytophthora parasitica INRA-310 protein kinase (637 aa) |
| 2563 | 1888 | PPTG_14363 | 75,77 | 5 | 0 | 0 | 5 | | | 1,40E+06 | 5 | 5 | 0 | 0 | 5 | Carbamidomethylation | 134722 | PPTG_14363 | Phytophthora parasitica INRA-310 P-type ATPase (P-ATPase) superfamily (1212 aa) |
| 2058 | 2272 | PPTG_04241 | 75,77 | 28 | 9 | 23 | 28 | 9,39E+04 | 2,09E+06 | 3,42E+06 | 4 | 4 | 1 | 3 | 4 | Oxidation (M); Carbamidomethylation | 16940 | PPTG_04241 | Phytophthora parasitica INRA-310 AP-4 complex subunit sigma (145 aa) |
| 2567 | 2363 | PPTG_20189 | 75,73 | 4 | 1 | 4 | 1 | 7,34E+04 | 5,16E+05 | 4,28E+05 | 3 | 3 | 1 | 3 | 1 | Carbamidomethylation | 116171 | PPTG_20189 | Phytophthora parasitica INRA-310 glycerophosphodiesterase (1044 aa) |
| 2567 | 2364 | PPTG_20189 | 75,73 | 4 | 1 | 4 | 1 | 7,34E+04 | 5,16E+05 | 4,28E+05 | 3 | 3 | 1 | 3 | 1 | Carbamidomethylation | 116171 | PPTG_20189 | Phytophthora parasitica INRA-310 glycerophosphodiesterase (1044 aa) |
| 2009 | 1880 | PPTG_08885 | 75,7 | 13 | 0 | 9 | 11 | | 3,32E+06 | 3,87E+06 | 5 | 5 | 0 | 4 | 4 | | 49770 | PPTG_08885 | Phytophthora parasitica INRA-310 kynurenine-oxoglutarate transaminase (449 aa) |
| 2528 | 2783 | PPTG_08206 | 75,67 | 10 | 4 | 10 | 4 | 1,16E+05 | 1,38E+06 | 5,94E+05 | 3 | 3 | 1 | 3 | 1 | | 38483 | PPTG_08206 | Phytophthora parasitica INRA-310 ubiquinone biosynthesis protein COQ9 (361 aa) |
| 2179 | 2397 | PPTG_02213 | 75,66 | 9 | 2 | 9 | 9 | 1,98E+05 | 6,93E+05 | 1,85E+06 | 3 | 3 | 1 | 3 | 3 | | 54688 | PPTG_02213 | Phytophthora parasitica INRA-310 hypothetical protein (494 aa) |
| 2465 | 2208 | PPTG_08930 | 75,63 | 9 | 0 | 6 | 9 | | 8,59E+05 | 1,46E+06 | 3 | 3 | 0 | 2 | 3 | | 43819 | PPTG_08930 | Phytophthora parasitica INRA-310 cobalamin synthesis protein (394 aa) |
| 3109 | 3194 | PPTG_00430 | 75,54 | 8 | 5 | 0 | 8 | 9,75E+04 | | 7,32E+05 | 2 | 2 | 1 | 0 | 2 | | 29647 | PPTG_00430 | Phytophthora parasitica INRA-310 hypothetical protein (284 aa) |
| 2318 | 2289 | PPTG_06147 | 75,53 | 9 | 0 | 4 | 9 | | 1,69E+06 | 3,12E+06 | 4 | 4 | 0 | 2 | 4 | | 46794 | PPTG_06147 | Phytophthora parasitica INRA-310 hypothetical protein (427 aa) |
| 2361 | 1911 | PPTG_05700 | 75,46 | 7 | 1 | 0 | 7 | 6,21E+04 | | 1,05E+06 | 5 | 5 | 1 | 0 | 5 | | 83254 | PPTG_05700 | Phytophthora parasitica INRA-310 inactive ubiquitin carboxyl-terminal hydrolase (745 aa) |
| 2347 | 2176 | PPTG_16940 | 75,44 | 22 | 5 | 13 | 14 | | 1,12E+05 | 9,01E+05 | 4 | 3 | 1 | 2 | 3 | | 25452 | PPTG_16940 | Phytophthora parasitica INRA-310 hypothetical protein (223 aa) |
| 2741 | 2638 | PPTG_01831 | 75,44 | 15 | 0 | 11 | 11 | | 1,11E+06 | 1,04E+06 | 3 | 3 | 0 | 2 | 2 | | 32972 | PPTG_01831 | Phytophthora parasitica INRA-310 thioredoxin H-type (297 aa) |
| 2456 | 2524 | PPTG_07042 | 75,36 | 18 | 0 | 5 | 18 | | 1,85E+05 | 7,23E+06 | 3 | 3 | 0 | 1 | 4 | | 31214 | PPTG_07042 | Phytophthora parasitica INRA-310 hypothetical protein (297 aa) |
| 2673 | 2381 | PPTG_12065 | 75,27 | 3 | 0 | 2 | 3 | | 1,00E+05 | 2,03E+05 | 3 | 1 | 0 | 2 | 2 | Carbamidomethylation | 114086 | PPTG_12065 | Phytophthora parasitica INRA-310 Argonaute5 (AGO5) (1028 aa) |
| 2092 | 3110 | PPTG_13937 | 75,27 | 9 | 9 | 9 | 9 | 1,31E+05 | 0 | 1,36E+06 | 3 | 1 | 3 | 3 | 2 | | 28589 | PPTG_13937 | Phytophthora parasitica INRA-310 choline-phosphate cytidylyltransferase B (252 aa) |
| 1599 | 1903 | PPTG_13843 | 75,23 | 12 | 4 | 10 | 10 | 3,22E+05 | 2,43E+06 | 1,35E+07 | 5 | 5 | 2 | 4 | 6 | Dehydration; Deamidation (NQ) | 56668 | PPTG_13843 | Phytophthora parasitica INRA-310 protein phosphatase 2A regulatory B subunit (495 aa) |
| 2068 | 1928 | PPTG_16782 | 75,21 | 14 | 2 | 12 | 6 | 6,46E+04 | 1,34E+06 | 1,02E+06 | 5 | 5 | 1 | 4 | 3 | Carbamidomethylation | 53422 | PPTG_16782 | Phytophthora parasitica INRA-310 glutathione synthetase (483 aa) |
| 1626 | 2155 | PPTG_17701 | 75,21 | 22 | 0 | 18 | 17 | | 2,42E+06 | 3,42E+06 | 6 | 6 | 0 | 6 | 6 | | 37301 | PPTG_17701 | Phytophthora parasitica INRA-310 NAD-dependent glycerol-3-phosphate dehydrogenase (349 aa) |
| 3467 | 2885 | PPTG_15377 | 75,16 | 16 | 0 | 0 | 16 | | | 4,51E+05 | 2 | 2 | 0 | 0 | 2 | Carbamidomethylation | 34011 | PPTG_15377 | Phytophthora parasitica INRA-310 hypothetical protein (303 aa) |
| 1928 | 2101 | PPTG_07810 | 75,13 | 18 | 0 | 14 | 11 | | 1,65E+06 | 1,58E+06 | 6 | 6 | 0 | 5 | 4 | Carbamidomethylation | 45999 | PPTG_07810 | Phytophthora parasitica INRA-310 4-hydroxyphenylpyruvate dioxygenase (419 aa) |
| 1928 | 2102 | PPTG_07812 | 75,13 | 18 | 0 | 14 | 11 | | 1,65E+06 | 1,58E+06 | 6 | 6 | 0 | 5 | 4 | Carbamidomethylation | 46113 | PPTG_07812 | Phytophthora parasitica INRA-310 4-hydroxyphenylpyruvate dioxygenase (421 aa) |
| 1948 | 3252 | PPTG_18312 | 75,1 | 20 | 8 | 20 | 20 | 5,16E+06 | 1,36E+07 | 7,51E+07 | 2 | 2 | 1 | 3 | 5 | Deamidation (NQ) | 17736 | PPTG_18312 | Phytophthora parasitica INRA-310 hypothetical protein (164 aa) |
| 2462 | 2499 | PPTG_08221 | 75,03 | 26 | 0 | 17 | 17 | | 1,85E+06 | 2,13E+06 | 3 | 3 | 0 | 3 | 2 | Oxidation (M); Acetylation (N-term) | 17326 | PPTG_08221 | Phytophthora parasitica INRA-310 nucleoredoxin (151 aa) |
| 1971 | 1826 | PPTG_06155 | 74,86 | 2 | 0 | 0 | 2 | | 1,70E+05 | 2,37E+06 | 7 | 7 | 0 | 1 | 7 | | 392171 | PPTG_06155 | Phytophthora parasitica INRA-310 hypothetical protein (3555 aa) |
| 3180 | 2530 | PPTG_03257 | 74,84 | 9 | 0 | 9 | 0 | | 9,65E+05 | | 3 | 3 | 0 | 3 | 0 | Carbamidomethylation | 45654 | PPTG_03257 | Phytophthora parasitica INRA-310 queuine tRNA-ribosyltransferase (410 aa) |
| 2351 | 2610 | PPTG_03513 | 74,84 | 8 | 0 | 4 | 8 | | 4,23E+05 | 1,07E+06 | 4 | 4 | 0 | 2 | 4 | | 75290 | PPTG_03513 | Phytophthora parasitica INRA-310 hypothetical protein (654 aa) |
| 2384 | 2074 | PPTG_02530 | 74,79 | 22 | 0 | 11 | 22 | | 1,06E+06 | 8,72E+06 | 4 | 4 | 0 | 2 | 4 | | 21814 | PPTG_02530 | Phytophthora parasitica INRA-310 hypothetical protein (204 aa) |
| 2527 | 2750 | PPTG_17668 | 74,77 | 18 | 0 | 15 | 10 | | 4,61E+06 | 0 | 3 | 2 | 0 | 3 | 2 | Deamidation (NQ) | 25011 | PPTG_17668 | Phytophthora parasitica INRA-310 chitinase (230 aa) |
| 1960 | 2614 | PPTG_17170 | 74,68 | 15 | 0 | 15 | 9 | | 1,52E+06 | 3,80E+05 | 5 | 5 | 0 | 6 | 3 | | 49185 | PPTG_17170 | Phytophthora parasitica INRA-310 hypothetical protein (495 aa) |
| 1538 | 2001 | PPTG_12009 | 74,67 | 11 | 0 | 5 | 9 | | 1,34E+05 | 1,34E+06 | 6 | 4 | 0 | 5 | 8 | Oxidation (M); Carbamidomethylation | 64894 | PPTG_12009 | Phytophthora parasitica INRA-310 glycoside hydrolase (574 aa) |
| 1780 | 1804 | PPTG_07128 | 74,63 | 5 | 0 | 3 | 4 | | 1,23E+06 | 1,56E+06 | 6 | 6 | 0 | 4 | 6 | | 140554 | PPTG_07128 | Phytophthora parasitica INRA-310 regulator-nonsense transcripts 2 (1251 aa) |
| 1810 | 1867 | PPTG_18910 | 74,58 | 13 | 4 | 13 | 11 | 1,72E+05 | 2,86E+06 | 2,75E+06 | 4 | 4 | 1 | 5 | 4 | Oxidation (M) | 37148 | PPTG_18910 | Phytophthora parasitica INRA-310 spermine synthase (333 aa) |
| 2486 | 2608 | PPTG_03933 | 74,58 | 7 | 0 | 4 | 7 | | 5,11E+05 | 8,00E+05 | 3 | 3 | 0 | 2 | 3 | | 62080 | PPTG_03933 | Phytophthora parasitica INRA-310 hypothetical protein (556 aa) |
| 2486 | 2609 | PPTG_03933 | 74,58 | 7 | 0 | 4 | 7 | | 5,11E+05 | 8,00E+05 | 3 | 3 | 0 | 2 | 3 | | 62080 | PPTG_03933 | Phytophthora parasitica INRA-310 hypothetical protein (556 aa) |
| 1549 | 3388 | PPTG_09080 | 74,56 | 18 | 13 | 18 | 18 | 2,50E+06 | 5,90E+07 | 4,33E+07 | 2 | 2 | 1 | 7 | 5 | Oxidation (M); Carbamidomethylation; Carbamylation | 18056 | PPTG_09080 | Phytophthora parasitica INRA-310 elicitin-like protein (184 aa) |
| 3153 | 3362 | PPTG_05081 | 74,55 | 14 | 0 | 14 | 9 | | 2,66E+05 | 9,16E+04 | 2 | 2 | 0 | 2 | 1 | Acetylation (N-term) | 21801 | PPTG_05081 | Phytophthora parasitica INRA-310 hypothetical protein (205 aa) |
| 2313 | 2082 | PPTG_04231 | 74,44 | 9 | 2 | 3 | 6 | 9,83E+04 | 2,00E+05 | 1,14E+06 | 5 | 5 | 1 | 1 | 4 | Carbamidomethylation | 70828 | PPTG_04231 | Phytophthora parasitica INRA-310 protein kinase (652 aa) |
| 2067 | 1902 | PPTG_14751 | 74,36 | 7 | 2 | 1 | 7 | 1,86E+05 | 1,53E+05 | 1,96E+06 | 5 | 5 | 2 | 1 | 5 | Carbamidomethylation | 98249 | PPTG_14751 | Phytophthora parasitica INRA-310 hypothetical protein (884 aa) |
| 1652 | 2693 | PPTG_18090 | 74,33 | 3 | 1 | 1 | 3 | | | 0 | 4 | 1 | 2 | 2 | 7 | Oxidation (M) | 151459 | PPTG_18090 | Phytophthora parasitica INRA-310 ATP-binding Cassette (ABC) superfamily protein (1356 aa) |
| 2281 | 2259 | PPTG_12227 | 74,31 | 26 | 0 | 11 | 21 | | 8,29E+05 | 3,02E+06 | 5 | 5 | 0 | 2 | 4 | Carbamidomethylation | 20011 | PPTG_12227 | Phytophthora parasitica INRA-310 hypothetical protein (176 aa) |
| 2536 | 2230 | PPTG_09681 | 74,31 | 7 | 0 | 2 | 7 | | 2,44E+05 | 2,10E+06 | 4 | 4 | 0 | 1 | 4 | | 76844 | PPTG_09681 | Phytophthora parasitica INRA-310 hypothetical protein (703 aa) |
| 1996 | 2021 | PPTG_17418 | 74,25 | 11 | 0 | 3 | 11 | | 4,02E+05 | 1,84E+06 | 6 | 5 | 0 | 2 | 6 | | 63969 | PPTG_17418 | Phytophthora parasitica INRA-310 hypothetical protein (563 aa) |
| 781 | 3450 | PPTG_13425 | 74,23 | 24 | 13 | 14 | 24 | 1,40E+07 | 6,45E+07 | 2,43E+08 | 3 | 3 | 5 | 9 | 15 | Oxidation (M); Deamidation (NQ) | 9246 | PPTG_13425 | Phytophthora parasitica INRA-310 ubiquitin-like protein/ribosomal protein (87 aa) |
| 2198 | 2709 | PPTG_04822 | 74,22 | 12 | 9 | 12 | 9 | 5,20E+05 | 3,78E+06 | 5,01E+06 | 3 | 3 | 2 | 3 | 2 | Carbamidomethylation | 25947 | PPTG_04822 | Phytophthora parasitica INRA-310 ribose 5-phosphate isomerase A (242 aa) |
| 2467 | 2267 | PPTG_11417 | 74,14 | 19 | 0 | 11 | 19 | | 9,86E+05 | 2,07E+06 | 3 | 3 | 0 | 2 | 3 | | 21925 | PPTG_11417 | Phytophthora parasitica INRA-310 peroxiredoxin-2 (199 aa) |
| 2467 | 2268 | PPTG_11416 | 74,14 | 19 | 0 | 11 | 19 | | 9,86E+05 | 2,07E+06 | 3 | 3 | 0 | 2 | 3 | | 21941 | PPTG_11416 | Phytophthora parasitica INRA-310 peroxiredoxin-2 (199 aa) |
| 1935 | 1980 | PPTG_02746 | 74,12 | 11 | 0 | 8 | 11 | | 1,35E+06 | 2,60E+06 | 5 | 5 | 0 | 4 | 5 | | 69013 | PPTG_02746 | Phytophthora parasitica INRA-310 DNA polymerase subunit alpha B (625 aa) |
| 2165 | 2283 | PPTG_19295 | 74,08 | 38 | 7 | 26 | 31 | 1,82E+05 | 1,98E+06 | 3,91E+06 | 4 | 4 | 1 | 3 | 3 | | 12412 | PPTG_19295 | Phytophthora parasitica INRA-310 hypothetical protein (112 aa) |

| | | | | | | | | | | | | | | | | | | | | |
|---|---|---|---|---|---|---|---|---|---|---|---|---|---|---|---|---|---|---|---|---|
| 2078 | 3207 | PPTG_13939 | 74,08 | 7 | 7 | 7 | 7 | | 3,55E+05 | 1,33E+06 | 2,62E+06 | 3 | 1 | 3 | 3 | 2 | | 35849 | PPTG_13939 | Phytophthora parasitica INRA-310 choline-phosphate cytidylyltransferase B (312 aa) |
| 2428 | 2445 | PPTG_12098 | 74,07 | 9 | 0 | 9 | 9 | | | 7,02E+05 | 1,07E+06 | 3 | 3 | 0 | 3 | 3 | | 56786 | PPTG_12098 | Phytophthora parasitica INRA-310 pre-mRNA-splicing factor prp46 (522 aa) |
| 1979 | 1908 | PPTG_13109 | 74,06 | 7 | 0 | 7 | 4 | | | 9,67E+05 | 1,54E+06 | 4 | 4 | 0 | 5 | 3 | | 81285 | PPTG_13109 | Phytophthora parasitica INRA-310 ATP-binding cassette protein (723 aa) |
| 2449 | 2038 | PPTG_08094 | 74,02 | 5 | 0 | 1 | 5 | | | 1,24E+05 | 1,60E+06 | 4 | 4 | 0 | 1 | 4 | | 108772 | PPTG_08094 | Phytophthora parasitica INRA-310 hypothetical protein (959 aa) |
| 2900 | 2768 | PPTG_18018 | 73,96 | 11 | 6 | 0 | 11 | | 2,71E+04 | | 1,28E+06 | 3 | 3 | 1 | 0 | 3 | Acetylation (N-term) | 38154 | PPTG_18018 | Phytophthora parasitica INRA-310 hypothetical protein (341 aa) |
| 1943 | 1976 | PPTG_09484 | 73,93 | 7 | 0 | 7 | 5 | | | 9,04E+05 | 1,22E+06 | 5 | 5 | 0 | 5 | 4 | | 101871 | PPTG_09484 | Phytophthora parasitica INRA-310 ubiquitin-specific protease (906 aa) |
| 2762 | 2111 | PPTG_16781 | 73,86 | 4 | 0 | 0 | 4 | | | | 3,06E+06 | 4 | 4 | 0 | 0 | 4 | Carbamidomethylation | 136828 | PPTG_16781 | Phytophthora parasitica INRA-310 hypothetical protein (1217 aa) |
| 2762 | 2113 | PPTG_16781 | 73,86 | 4 | 0 | 0 | 4 | | | | 3,06E+06 | 4 | 4 | 0 | 0 | 4 | Carbamidomethylation | 140238 | PPTG_16781 | Phytophthora parasitica INRA-310 hypothetical protein (1249 aa) |
| 1982 | 2006 | PPTG_08516 | 73,85 | 10 | 1 | 0 | 10 | | | | 5,19E+06 | 7 | 6 | 1 | 0 | 7 | Carbamidomethylation | 83690 | PPTG_08516 | Phytophthora parasitica INRA-310 protein kinase (755 aa) |
| 3056 | 2528 | PPTG_18866 | 73,79 | 5 | 0 | 0 | 5 | | | | 8,54E+05 | 3 | 3 | 0 | 0 | 3 | Carbamidomethylation | 73563 | PPTG_18866 | Phytophthora parasitica INRA-310 hypothetical protein (696 aa) |
| 2022 | 1885 | PPTG_11215 | 73,72 | 8 | 1 | 6 | 4 | | 4,26E+05 | 2,42E+06 | 8,89E+05 | 5 | 5 | 1 | 4 | 3 | Carbamidomethylation | 86448 | PPTG_11215 | Phytophthora parasitica INRA-310 hypothetical protein (772 aa) |
| 4270 | 4086 | PPTG_10047 | 73,68 | 15 | 0 | 15 | 0 | | | 4,52E+05 | | 1 | 1 | 0 | 1 | 0 | Carbamidomethylation | 26853 | PPTG_10047 | Phytophthora parasitica INRA-310 hypothetical protein (266 aa) |
| 2823 | 3245 | PPTG_05352 | 73,54 | 13 | 0 | 13 | 13 | | | 7,64E+05 | 1,57E+06 | 2 | 2 | 0 | 2 | 2 | | 23303 | PPTG_05352 | Phytophthora parasitica INRA-310 hypothetical protein (211 aa) |
| 1716 | 1970 | PPTG_13465 | 73,54 | 12 | 6 | 10 | 12 | | 2,95E+05 | 2,13E+06 | 5,18E+06 | 5 | 5 | 2 | 4 | 5 | Carbamidomethylation | 46668 | PPTG_13465 | Phytophthora parasitica INRA-310 histidinol-phosphate aminotransferase (422 aa) |
| 1763 | 1941 | PPTG_04901 | 73,5 | 9 | 1 | 2 | 9 | | 1,62E+05 | 7,42E+05 | 3,11E+06 | 7 | 7 | 1 | 2 | 7 | Oxidation (M) | 100072 | PPTG_04901 | Phytophthora parasitica INRA-310 protein kinase (898 aa) |
| 1910 | 2004 | PPTG_08824 | 73,49 | 11 | 2 | 10 | 10 | 0 | | 1,15E+06 | 2,69E+06 | 5 | 5 | 1 | 4 | 4 | | 50757 | PPTG_08824 | Phytophthora parasitica INRA-310 hypothetical protein (472 aa) |
| 2411 | 2460 | PPTG_08739 | 73,49 | 6 | 0 | 6 | 6 | | | 6,44E+05 | 6,83E+05 | 3 | 3 | 0 | 3 | 3 | | 73941 | PPTG_08739 | Phytophthora parasitica INRA-310 acyl-CoA dehydrogenase/oxidase (662 aa) |
| 2056 | 2194 | PPTG_09144 | 73,47 | 4 | 1 | 4 | 2 | | 1,20E+05 | 5,93E+05 | 1,15E+06 | 4 | 4 | 1 | 4 | 3 | | 150000 | PPTG_09144 | Phytophthora parasitica INRA-310 hypothetical protein (1315 aa) |
| 3324 | 2905 | PPTG_10638 | 73,44 | 3 | 0 | 0 | 3 | | | | 4,19E+05 | 2 | 2 | 0 | 0 | 2 | Carbamidomethylation | 104397 | PPTG_10638 | Phytophthora parasitica INRA-310 phosphatidylinositol-4-phosphate-5-kinase (PIPK-D6/GPCR-PIPK) (939 aa) |
| 2008 | 1839 | PPTG_16210 | 73,42 | 8 | 1 | 5 | 6 | | 1,01E+05 | 9,06E+05 | 1,64E+06 | 5 | 5 | 1 | 3 | 4 | | 104410 | PPTG_16210 | Phytophthora parasitica INRA-310 hypothetical protein (934 aa) |
| 2016 | 2319 | PPTG_03774 | 73,42 | 12 | 4 | 12 | 12 | | 6,06E+04 | 2,58E+06 | 3,05E+06 | 3 | 3 | 1 | 4 | 3 | Carbamidomethylation; Acetylation (N-term) | 33828 | PPTG_03774 | Phytophthora parasitica INRA-310 carboxyvinyl-carboxyphosphonate phosphorylmutase (313 aa) |
| 2158 | 1886 | PPTG_07411 | 73,34 | 14 | 0 | 3 | 14 | | | 6,85E+05 | 2,77E+06 | 6 | 6 | 0 | 1 | 6 | | 44582 | PPTG_07411 | Phytophthora parasitica INRA-310 hypothetical protein (412 aa) |
| 2188 | 2192 | PPTG_13867 | 73,33 | 8 | 3 | 8 | 8 | | 3,68E+05 | 1,35E+06 | 4,07E+06 | 3 | 3 | 1 | 3 | 3 | Carbamidomethylation | 52309 | PPTG_13867 | Phytophthora parasitica INRA-310 hypothetical protein (478 aa) |
| 1975 | 1857 | PPTG_13300 | 73,31 | 5 | 0 | 3 | 5 | | | 1,36E+06 | 1,89E+06 | 5 | 5 | 0 | 3 | 5 | | 120146 | PPTG_13300 | Phytophthora parasitica INRA-310 hypothetical protein (1085 aa) |
| 2707 | 2275 | PPTG_08411 | 73,27 | 7 | 0 | 0 | 7 | | | | 1,22E+06 | 4 | 4 | 0 | 0 | 4 | | 103131 | PPTG_08411 | Phytophthora parasitica INRA-310 polycystin Cation Channel (PCC) family (932 aa) |
| 2302 | 2159 | PPTG_05008 | 73,25 | 9 | 0 | 7 | 7 | | | 1,28E+06 | 2,00E+06 | 4 | 4 | 0 | 3 | 3 | | 57472 | PPTG_05008 | Phytophthora parasitica INRA-310 heat shock protein HslVU ATPase subunit HslU (514 aa) |
| 2232 | 1862 | PPTG_13585 | 73,23 | 6 | 0 | 0 | 6 | | | | 1,31E+06 | 7 | 7 | 0 | 0 | 7 | Carbamidomethylation | 178719 | PPTG_13585 | Phytophthora parasitica INRA-310 P-type ATPase (P-ATPase) superfamily (1598 aa) |
| 1931 | 2523 | PPTG_05974 | 73,22 | 22 | 0 | 22 | 11 | | | 4,72E+06 | 2,15E+06 | 4 | 4 | 0 | 7 | 2 | Oxidation (M); Carbamidomethylation; Deamidation (NQ) | 32095 | PPTG_05974 | Phytophthora parasitica INRA-310 carbonic anhydrase (282 aa) |
| 2928 | 4210 | PPTG_15237 | 73,2 | 21 | 0 | 21 | 21 | | | 3,22E+06 | 1,89E+05 | 1 | 1 | 0 | 2 | 2 | Carbamidomethylation | 16067 | PPTG_15237 | Phytophthora parasitica INRA-310 elicitin-like protein (161 aa) |
| 2420 | 3201 | PPTG_10243 | 73,17 | 29 | 0 | 14 | 29 | | | 8,50E+05 | 5,98E+06 | 3 | 3 | 0 | 2 | 4 | Oxidation (M); Carbamidomethylation | 9333 | PPTG_10243 | Phytophthora parasitica INRA-310 hypothetical protein (85 aa) |
| 2401 | 3363 | PPTG_10227 | 73,12 | 37 | 23 | 37 | 37 | | 8,99E+05 | 9,68E+06 | 6,52E+07 | 2 | 2 | 1 | 2 | 3 | | 9145 | PPTG_10227 | Phytophthora parasitica INRA-310 hypothetical protein (87 aa) |
| 2391 | 2367 | PPTG_10327 | 73,08 | 11 | 3 | 8 | 11 | 0 | | 4,89E+05 | 4,27E+06 | 3 | 3 | 1 | 2 | 3 | | 33128 | PPTG_10327 | Phytophthora parasitica INRA-310 mitochondrial Carrier (MC) family (308 aa) |
| 1976 | 2436 | PPTG_13412 | 73,07 | 21 | 0 | 21 | 21 | | | 2,30E+06 | 5,39E+06 | 3 | 3 | 0 | 4 | 4 | Oxidation (M) | 18869 | PPTG_13412 | Phytophthora parasitica INRA-310 50S ribosomal protein L7/L12 (179 aa) |
| 1887 | 1837 | PPTG_08476 | 73,04 | 22 | 18 | 13 | 18 | | 8,96E+05 | 6,35E+06 | 7,66E+06 | 4 | 4 | 3 | 3 | 3 | Carbamidomethylation | 27955 | PPTG_08476 | Phytophthora parasitica INRA-310 hypothetical protein (248 aa) |
| 1983 | 1811 | PPTG_13045 | 72,98 | 8 | 0 | 3 | 8 | | | 3,91E+05 | 3,36E+06 | 6 | 6 | 0 | 2 | 6 | | 75227 | PPTG_13045 | Phytophthora parasitica INRA-310 leukotriene A-4 hydrolase-like protein (671 aa) |
| 2089 | 2372 | PPTG_09835 | 72,96 | 7 | 0 | 7 | 7 | | | 5,39E+06 | 3,87E+06 | 3 | 2 | 0 | 4 | 4 | Carbamidomethylation | 54214 | PPTG_09835 | Phytophthora parasitica INRA-310 1 3-beta-glucanosyltransferase (495 aa) |
| 2212 | 2517 | PPTG_09671 | 72,92 | 10 | 5 | 7 | 10 | | 5,87E+04 | 1,04E+06 | 2,39E+06 | 3 | 3 | 1 | 2 | 4 | Oxidation (M); Deamidation (NQ) | 41401 | PPTG_09671 | Phytophthora parasitica INRA-310 hypothetical protein (379 aa) |
| 2085 | 2094 | PPTG_12786 | 72,9 | 12 | 3 | 12 | 10 | | 5,13E+04 | 1,87E+06 | 9,08E+05 | 4 | 4 | 1 | 4 | 3 | | 35982 | PPTG_12786 | Phytophthora parasitica INRA-310 ATP-binding Cassette (ABC) superfamily (325 aa) |
| 1600 | 2282 | PPTG_13696 | 72,87 | 14 | 4 | 14 | 9 | | | 2,12E+05 | 2,63E+05 | 3 | 2 | 1 | 6 | 5 | Deamidation (NQ) | 31357 | PPTG_13696 | Phytophthora parasitica INRA-310 peroxisomal multifunctional enzyme type 2 (287 aa) |
| 2042 | 2142 | PPTG_02526 | 72,82 | 25 | 3 | 22 | 11 | | | 1,35E+06 | 2,27E+06 | 5 | 4 | 1 | 4 | 3 | Carbamidomethylation; Acetylation (N-term) | 25015 | PPTG_02526 | Phytophthora parasitica INRA-310 maleylacetoacetate isomerase (220 aa) |
| 2640 | 4087 | PPTG_16238 | 72,78 | 12 | 12 | 12 | 12 | | 5,00E+04 | 1,54E+06 | 1,47E+06 | 1 | 1 | 1 | 2 | 2 | | 18813 | PPTG_16238 | Phytophthora parasitica INRA-310 hypothetical protein (184 aa) |
| 2658 | 3066 | PPTG_18820 | 72,72 | 4 | 0 | 4 | 4 | | | 3,20E+05 | 5,06E+05 | 2 | 2 | 0 | 2 | 2 | | 86421 | PPTG_18820 | Phytophthora parasitica INRA-310 hypothetical protein (750 aa) |
| 1877 | 2087 | PPTG_15538 | 72,67 | 6 | 1 | 4 | 6 | | 1,12E+05 | 1,26E+06 | 3,07E+06 | 5 | 5 | 1 | 3 | 5 | | 89440 | PPTG_15538 | Phytophthora parasitica INRA-310 TFIIH basal transcription factor complex helicase subunit (790 aa) |
| 2358 | 3142 | PPTG_11374 | 72,53 | 16 | 11 | 16 | 16 | | 1,63E+06 | 2,97E+06 | 1,29E+07 | 2 | 2 | 1 | 2 | 3 | | 15252 | PPTG_11374 | Phytophthora parasitica INRA-310 zinc finger CDGSH domain-containing protein 1 (135 aa) |
| 2795 | 2393 | PPTG_09559 | 72,48 | 7 | 0 | 0 | 7 | | | | 1,11E+06 | 4 | 4 | 0 | 0 | 4 | | 67851 | PPTG_09559 | Phytophthora parasitica INRA-310 hypothetical protein (609 aa) |
| 1968 | 2394 | PPTG_06803 | 72,45 | 3 | 0 | 2 | 2 | | | 1,19E+06 | 2,45E+06 | 5 | 5 | 0 | 4 | 4 | | 231266 | PPTG_06803 | Phytophthora parasitica INRA-310 hypothetical protein (2087 aa) |
| 2369 | 2484 | PPTG_16437 | 72,45 | 9 | 3 | 3 | 9 | | 1,15E+05 | 2,98E+05 | 2,57E+06 | 4 | 4 | 1 | 1 | 4 | Carbamidomethylation | 43155 | PPTG_16437 | Phytophthora parasitica INRA-310 hypothetical protein (394 aa) |
| 2735 | 2244 | PPTG_16433 | 72,42 | 15 | 0 | 5 | 15 | | | 5,00E+05 | 4,77E+06 | 3 | 3 | 0 | 1 | 3 | | 27761 | PPTG_16433 | Phytophthora parasitica INRA-310 isoamyl acetate-hydrolyzing esterase 1 (248 aa) |
| 2214 | 2613 | PPTG_10485 | 72,42 | 28 | 0 | 22 | 23 | | | 2,76E+06 | 4,08E+06 | 4 | 4 | 0 | 4 | 3 | | 14281 | PPTG_10485 | Phytophthora parasitica INRA-310 hypothetical protein (129 aa) |
| 2408 | 2369 | PPTG_01535 | 72,35 | 7 | 2 | 7 | 5 | | 9,65E+04 | 1,29E+06 | 1,57E+06 | 3 | 3 | 1 | 3 | 2 | Acetylation (N-term) | 55337 | PPTG_01535 | Phytophthora parasitica INRA-310 sperm-associated antigen 6 (510 aa) |
| 2164 | 2278 | PPTG_08890 | 72,29 | 15 | 4 | 6 | 15 | 0 | | 1,98E+06 | 5,37E+06 | 4 | 4 | 1 | 2 | 4 | | 28504 | PPTG_08890 | Phytophthora parasitica INRA-310 hypothetical protein (251 aa) |
| 2540 | 2401 | PPTG_08576 | 72,25 | 46 | 0 | 29 | 46 | | | 1,26E+06 | 3,72E+06 | 3 | 3 | 0 | 2 | 3 | | 11633 | PPTG_08576 | Phytophthora parasitica INRA-310 30S ribosomal protein S16 (103 aa) |
| 3192 | 3254 | PPTG_19730 | 72,25 | 9 | 0 | 0 | 9 | | | | 3,54E+06 | 2 | 2 | 0 | 0 | 3 | | 47633 | PPTG_19730 | Phytophthora parasitica INRA-310 amino Acid/Auxin Permease (AAAP) family (440 aa) |
| 2272 | 2675 | PPTG_03864 | 72,23 | 8 | 0 | 4 | 6 | | | 2,31E+05 | 1,35E+06 | 4 | 4 | 0 | 2 | 4 | Oxidation (M) | 85148 | PPTG_03864 | Phytophthora parasitica INRA-310 serine protease family S09A (744 aa) |
| 2142 | 2179 | PPTG_17661 | 72,19 | 9 | 3 | 9 | 5 | | 5,23E+04 | 7,63E+05 | 7,43E+05 | 4 | 4 | 1 | 4 | 2 | | 51799 | PPTG_17661 | Phytophthora parasitica INRA-310 COP9 signalosome complex subunit 2 (447 aa) |
| 2377 | 2797 | PPTG_14520 | 72,19 | 9 | 2 | 2 | 9 | | 1,75E+06 | 2,73E+06 | 3,11E+07 | 3 | 3 | 1 | 1 | 4 | Oxidation (M); Acetylation (N-term) | 52129 | PPTG_14520 | Phytophthora parasitica INRA-310 hypothetical protein (485 aa) |
| 2090 | 2489 | PPTG_04063 | 72,19 | 38 | 29 | 29 | 38 | | 6,30E+06 | 1,17E+07 | 5,72E+07 | 3 | 3 | 2 | 2 | 4 | Carbamidomethylation | 9148 | PPTG_04063 | Phytophthora parasitica INRA-310 hypothetical protein (78 aa) |
| 2548 | 2696 | PPTG_08096 | 72,17 | 15 | 5 | 5 | 15 | | 9,70E+04 | 1,06E+06 | 1,30E+06 | 3 | 3 | 1 | 1 | 3 | Carbamidomethylation | 41008 | PPTG_08096 | Phytophthora parasitica INRA-310 carbonic anhydrase (363 aa) |
| 2850 | 2535 | PPTG_13386 | 72,17 | 10 | 0 | 10 | 2 | | | 4,16E+05 | | 3 | 1 | 0 | 3 | 1 | | 41961 | PPTG_13386 | Phytophthora parasitica INRA-310 alkaline phosphatase (379 aa) |
| 2693 | 2549 | PPTG_16725 | 72,15 | 9 | 0 | 0 | 9 | | | | 7,86E+05 | 4 | 3 | 0 | 0 | 4 | | 63480 | PPTG_16725 | Phytophthora parasitica INRA-310 exocyst complex component (565 aa) |
| 2722 | 2876 | PPTG_16025 | 72,03 | 6 | 3 | 3 | 6 | | 3,39E+05 | 6,89E+04 | 1,72E+06 | 2 | 2 | 1 | 1 | 2 | | 50545 | PPTG_16025 | Phytophthora parasitica INRA-310 sorting and assembly machinery component 50 (464 aa) |
| 2084 | 2080 | PPTG_08666 | 72,02 | 20 | 3 | 16 | 6 | | 1,41E+05 | 3,29E+06 | 2,17E+06 | 6 | 6 | 1 | 5 | 2 | Carbamidomethylation | 52050 | PPTG_08666 | Phytophthora parasitica INRA-310 hypothetical protein (501 aa) |
| 2184 | 2052 | PPTG_15741 | 72,01 | 10 | 0 | 10 | 4 | | | 1,06E+06 | 7,85E+05 | 4 | 4 | 0 | 4 | 3 | Carbamidomethylation; Deamidation (NQ) | 49518 | PPTG_15741 | Phytophthora parasitica INRA-310 methionine aminopeptidase (451 aa) |
| 2768 | 2527 | PPTG_06750 | 71,98 | 27 | 0 | 14 | 22 | | | 1,49E+06 | 1,41E+06 | 3 | 3 | 0 | 2 | 2 | Carbamidomethylation | 18719 | PPTG_06750 | Phytophthora parasitica INRA-310 S-phase kinase-associated protein 1A (167 aa) |
| 1932 | 3371 | PPTG_18620 | 71,97 | 7 | 7 | 7 | 7 | | 1,13E+07 | 3,87E+07 | 1,60E+08 | 2 | 2 | 2 | 2 | 5 | Carbamidomethylation; Deamidation (NQ) | 34858 | PPTG_18620 | Phytophthora parasitica INRA-310 hypothetical protein (339 aa) |
| 2586 | 2249 | PPTG_15902 | 71,94 | 9 | 0 | 3 | 9 | | | 1,92E+05 | 3,78E+06 | 4 | 4 | 0 | 1 | 4 | | 46644 | PPTG_15902 | Phytophthora parasitica INRA-310 hypothetical protein (407 aa) |
| 2586 | 2250 | PPTG_15902 | 71,94 | 9 | 0 | 3 | 9 | | | 1,92E+05 | 3,78E+06 | 4 | 4 | 0 | 1 | 4 | | 46644 | PPTG_15902 | Phytophthora parasitica INRA-310 hypothetical protein (407 aa) |
| 1958 | 2245 | PPTG_11344 | 71,93 | 3 | 0 | 3 | 3 | | | 2,99E+06 | 1,72E+06 | 4 | 3 | 0 | 5 | 4 | Oxidation (M); Carbamidomethylation | 127616 | PPTG_11344 | Phytophthora parasitica INRA-310 hypothetical protein (1120 aa) |
| 1876 | 2063 | PPTG_06224 | 71,92 | 13 | 2 | 10 | 7 | | 4,50E+04 | 1,67E+06 | 3,10E+06 | 6 | 6 | 1 | 4 | 4 | Carbamidomethylation | 57452 | PPTG_06224 | Phytophthora parasitica INRA-310 hypothetical protein (526 aa) |
| 2617 | 2248 | PPTG_00092 | 71,91 | 13 | 0 | 13 | 3 | | | 1,26E+06 | 2,67E+05 | 4 | 4 | 0 | 4 | 1 | | 46259 | PPTG_00092 | Phytophthora parasitica INRA-310 hypothetical protein (427 aa) |
| 3610 | 4211 | PPTG_14175 | 71,83 | 5 | 0 | 5 | 5 | | | 5,01E+05 | 3,47E+05 | 1 | 1 | 0 | 1 | 1 | | 40429 | PPTG_14175 | Phytophthora parasitica INRA-310 carbohydrate-binding protein (384 aa) |
| 2419 | 3196 | PPTG_15794 | 71,81 | 30 | 30 | 30 | 30 | | 1,17E+06 | 6,21E+06 | 8,02E+06 | 2 | 2 | 2 | 2 | 2 | | 9912 | PPTG_15794 | Phytophthora parasitica INRA-310 hypothetical protein (88 aa) |
| 2568 | 2377 | PPTG_01550 | 71,77 | 13 | 9 | 0 | 13 | | 3,29E+04 | | 1,19E+06 | 3 | 3 | 2 | 0 | 3 | | 31546 | PPTG_01550 | Phytophthora parasitica INRA-310 hypothetical protein (285 aa) |
| 2671 | 2169 | PPTG_06566 | 71,71 | 30 | 0 | 0 | 30 | | | | 5,80E+05 | 4 | 4 | 0 | 0 | 4 | | 22847 | PPTG_06566 | Phytophthora parasitica INRA-310 hypothetical protein (206 aa) |
| 1010 | 2171 | PPTG_19308 | 71,71 | 33 | 26 | 26 | 33 | | 4,65E+07 | 1,65E+08 | 4,82E+08 | 4 | 4 | 3 | 8 | 11 | Oxidation (M); Dehydration; Acetylation (N-term); Carbamylation | 11991 | PPTG_19308 | Phytophthora parasitica INRA-310 50S ribosomal protein L36e (108 aa) |
| 2365 | 2124 | PPTG_16855 | 71,7 | 15 | 4 | 4 | 15 | | | | 4,86E+05 | 4 | 3 | 1 | 1 | 4 | Carbamidomethylation | 34470 | PPTG_16855 | Phytophthora parasitica INRA-310 Rab11 family GTPase (311 aa) |
| 2704 | 2057 | PPTG_18088 | 71,67 | 10 | 0 | 0 | 10 | | | | 9,89E+05 | 4 | 4 | 0 | 0 | 4 | Carbamidomethylation | 47585 | PPTG_18088 | Phytophthora parasitica INRA-310 protein kinase (441 aa) |
| 2478 | 3017 | PPTG_16313 | 71,67 | 7 | 4 | 7 | 7 | | 5,21E+04 | 8,94E+05 | 1,56E+06 | 2 | 2 | 1 | 2 | 2 | Carbamidomethylation | 39803 | PPTG_16313 | Phytophthora parasitica INRA-310 replication factor C subunit 5 (354 aa) |
| 1947 | 2680 | PPTG_07082 | 71,63 | 42 | 12 | 26 | 42 | | 2,93E+05 | 3,05E+06 | 5,08E+06 | 4 | 4 | 1 | 3 | 5 | | 8438 | PPTG_07082 | Phytophthora parasitica INRA-310 DNA-directed RNA polymerase I II and III subunit RPABC5 (73 aa) |
| 2493 | 2281 | PPTG_02341 | 71,59 | 8 | 0 | 0 | 8 | | | | 2,02E+06 | 4 | 4 | 0 | 0 | 5 | | 50270 | PPTG_02341 | Phytophthora parasitica INRA-310 hypothetical protein (458 aa) |
| 1783 | 2165 | PPTG_06042 | 71,55 | 16 | 0 | 14 | 7 | | | 1,61E+06 | 1,29E+06 | 7 | 7 | 0 | 7 | 3 | Oxidation (M) | 50928 | PPTG_06042 | Phytophthora parasitica INRA-310 D-3-phosphoglycerate dehydrogenase (468 aa) |
| 2385 | 2160 | PPTG_02223 | 71,55 | 18 | 0 | 18 | 8 | | | 6,13E+05 | 4,03E+05 | 4 | 4 | 0 | 4 | 2 | | 34525 | PPTG_02223 | Phytophthora parasitica INRA-310 protein farnesyltransferase/geranylgeranyltransferase (300 aa) |
| 2764 | 2340 | PPTG_17682 | 71,53 | 19 | 7 | 0 | 19 | | 1,15E+05 | | 2,88E+06 | 3 | 3 | 1 | 0 | 3 | Acetylation (N-term) | 20331 | PPTG_17682 | Phytophthora parasitica INRA-310 signal peptidase I (181 aa) |
| 2241 | 2432 | PPTG_10675 | 71,35 | 18 | 7 | 18 | 18 | | 2,42E+05 | 3,33E+06 | 5,89E+06 | 3 | 3 | 1 | 3 | 3 | | 19595 | PPTG_10675 | Phytophthora parasitica INRA-310 hypothetical protein (170 aa) |
| 1998 | 2077 | PPTG_02846 | 71,32 | 5 | 1 | 5 | 3 | | 2,15E+05 | 1,88E+06 | 2,28E+06 | 4 | 4 | 1 | 4 | 3 | | 119979 | PPTG_02846 | Phytophthora parasitica INRA-310 transcription elongation factor SPT5 (1103 aa) |
| 2718 | 2796 | PPTG_00834 | 71,29 | 8 | 0 | 6 | 6 | | | 2,06E+05 | 4,67E+05 | 3 | 3 | 0 | 2 | 2 | Carbamidomethylation | 53693 | PPTG_00834 | Phytophthora parasitica INRA-310 hypothetical protein (498 aa) |
| 2237 | 2239 | PPTG_14625 | 71,25 | 22 | 0 | 18 | 22 | | | 1,38E+06 | 1,02E+06 | 4 | 4 | 0 | 3 | 4 | Carbamidomethylation | 24805 | PPTG_14625 | Phytophthora parasitica INRA-310 uracil phosphoribosyltransferase (228 aa) |
| 2433 | 3059 | PPTG_11291 | 71,17 | 7 | 0 | 7 | 7 | | | 7,02E+05 | 9,66E+05 | 2 | 2 | 0 | 3 | 3 | | 53264 | PPTG_11291 | Phytophthora parasitica INRA-310 hypothetical protein (501 aa) |
| 2153 | 2566 | PPTG_07798 | 71,12 | 3 | 1 | 2 | 2 | | 1,75E+05 | 5,14E+05 | 2,07E+06 | 5 | 5 | 1 | 3 | 3 | | 192663 | PPTG_07798 | Phytophthora parasitica INRA-310 nuclear pore complex protein (1827 aa) |
| 2028 | 2119 | PPTG_10001 | 71,12 | 31 | 0 | 31 | 22 | | | 1,98E+06 | 1,64E+06 | 4 | 4 | 0 | 5 | 3 | Oxidation (M); Carbamidomethylation | 21286 | PPTG_10001 | Phytophthora parasitica INRA-310 maltose O-acetyltransferase (195 aa) |
| 2672 | 2360 | PPTG_09594 | 71,1 | 4 | 0 | 3 | 1 | | | 2,84E+05 | 2,06E+05 | 4 | 4 | 0 | 3 | 1 | | 158547 | PPTG_09594 | Phytophthora parasitica INRA-310 hypothetical protein (1453 aa) |
| 2706 | 2247 | PPTG_00511 | 71,09 | 13 | 0 | 0 | 13 | | | | 2,08E+06 | 4 | 4 | 0 | 0 | 4 | Carbamidomethylation | 37075 | PPTG_00511 | Phytophthora parasitica INRA-310 hypothetical protein (339 aa) |
| 2240 | 2430 | PPTG_10451 | 71,04 | 14 | 5 | 10 | 14 | | | 1,10E+06 | 1,08E+06 | 3 | 1 | 1 | 3 | 3 | Carbamidomethylation | 32790 | PPTG_10451 | Phytophthora parasitica INRA-310 3-hydroxyisobutyrate dehydrogenase (312 aa) |
| 2629 | 3237 | PPTG_07283 | 71,02 | 12 | 0 | 9 | 12 | | | 1,68E+06 | 2,84E+06 | 3 | 3 | 0 | 2 | 3 | | 20172 | PPTG_07283 | Phytophthora parasitica INRA-310 H/ACA ribonucleoprotein complex subunit (204 aa) |
| 2086 | 2098 | PPTG_13857 | 70,94 | 8 | 2 | 8 | 6 | | 9,99E+04 | 4,60E+05 | 1,01E+06 | 4 | 4 | 1 | 4 | 3 | Carbamidomethylation | 59287 | PPTG_13857 | Phytophthora parasitica INRA-310 tRNA nucleotidyltransferase (533 aa) |
| 2482 | 2343 | PPTG_13057 | 70,94 | 22 | 0 | 11 | 18 | | | 3,50E+05 | 1,17E+06 | 4 | 4 | 0 | 2 | 3 | Carbamidomethylation | 32712 | PPTG_13057 | Phytophthora parasitica INRA-310 glucanase inhibitor protein (314 aa) |
| 2252 | 2232 | PPTG_06402 | 70,94 | 4 | 0 | 4 | 3 | | | 9,50E+05 | 1,00E+06 | 4 | 4 | 0 | 4 | 3 | | 107084 | PPTG_06402 | Phytophthora parasitica INRA-310 hypothetical protein (981 aa) |
| 2490 | 2055 | PPTG_02592 | 70,91 | 6 | 0 | 2 | 6 | | | 1,12E+05 | 6,69E+05 | 4 | 4 | 0 | 1 | 4 | | 81505 | PPTG_02592 | Phytophthora parasitica INRA-310 hypothetical protein (725 aa) |
| 1527 | 2782 | PPTG_10422 | 70,89 | 20 | 15 | 15 | 20 | | 2,06E+07 | 7,95E+07 | 3,53E+08 | 3 | 3 | 2 | 4 | 7 | Oxidation (M) | 18423 | PPTG_10422 | Phytophthora parasitica INRA-310 60S ribosomal protein (161 aa) |
| 2363 | 2090 | PPTG_13325 | 70,77 | 12 | 0 | 6 | 10 | | | 6,16E+05 | 6,02E+05 | 4 | 4 | 0 | 3 | 3 | | 53693 | PPTG_13325 | Phytophthora parasitica INRA-310 UDP-N-acetylhexosamine pyrophosphorylase (494 aa) |
| 2991 | 2461 | PPTG_18763 | 70,74 | 6 | 0 | 6 | 0 | | | 1,86E+06 | | 3 | 3 | 0 | 3 | 0 | | 77599 | PPTG_18763 | Phytophthora parasitica INRA-310 metalloprotease family M13 (707 aa) |
| 2894 | 2516 | PPTG_04115 | 70,73 | 10 | 0 | 3 | 10 | | | 8,41E+05 | 1,34E+06 | 3 | 3 | 0 | 1 | 3 | Carbamidomethylation | 42963 | PPTG_04115 | Phytophthora parasitica INRA-310 hypothetical protein (431 aa) |
| 2073 | 2254 | PPTG_00083 | 70,71 | 13 | 3 | 13 | 10 | | 1,91E+05 | 2,54E+06 | 2,58E+06 | 4 | 4 | 1 | 4 | 3 | Carbamidomethylation | 28956 | PPTG_00083 | Phytophthora parasitica INRA-310 carbonic anhydrase (270 aa) |
| 2291 | 2135 | PPTG_05238 | 70,63 | 9 | 2 | 0 | 9 | 0 | | | 1,50E+06 | 5 | 4 | 1 | 0 | 5 | Carbamidomethylation | 80078 | PPTG_05238 | Phytophthora parasitica INRA-310 HECT E3 ubiquitin ligase (720 aa) |
| 1632 | 2081 | PPTG_10222 | 70,61 | 19 | 13 | 13 | 19 | | 1,13E+06 | 1,62E+06 | 1,30E+07 | 4 | 4 | 3 | 4 | 5 | Oxidation (M) | 29131 | PPTG_10222 | Phytophthora parasitica INRA-310 hypothetical protein (263 aa) |
| 1994 | 1874 | PPTG_16982 | 70,54 | 4 | 0 | 2 | 3 | | | 3,17E+05 | 1,26E+06 | 6 | 6 | 0 | 3 | 5 | Oxidation (M); Carbamidomethylation | 175534 | PPTG_16982 | Phytophthora parasitica INRA-310 DNA polymerase alpha catalytic subunit (1563 aa) |

| | | | | | | | | | | | | | | | | | | | |
|---|---|---|---|---|---|---|---|---|---|---|---|---|---|---|---|---|---|---|---|
| 2044 | 2443 | PPTG_10761 | 70,45 | 17 | 10 | 17 | 17 | 5,18E+05 | 1,78E+06 | 4,88E+06 | 3 | 3 | 2 | 3 | 3 | Acetylation (N-term) | 24031 | PPTG_10761 | Phytophthora parasitica INRA-310 rho GDP-dissociation inhibitor (211 aa) |
| 2452 | 2303 | PPTG_04795 | 70,38 | 12 | 0 | 6 | 9 | | | 5,33E+05 | 4 | 1 | 0 | 2 | 3 | | 43033 | PPTG_04795 | Phytophthora parasitica INRA-310 pyridine nucleotide-disulphide oxidoreductase (394 aa) |
| 2055 | 2177 | PPTG_04405 | 70,33 | 14 | 4 | 14 | 10 | 6,15E+04 | 1,01E+06 | 6,51E+05 | 4 | 4 | 1 | 4 | 3 | | 35946 | PPTG_04405 | Phytophthora parasitica INRA-310 vacuolar protein sorting-associated protein 26B-A (314 aa) |
| 2709 | 2290 | PPTG_12241 | 70,24 | 8 | 0 | 0 | 8 | | | 3,00E+06 | 4 | 4 | 0 | 0 | 4 | | 59190 | PPTG_12241 | Phytophthora parasitica INRA-310 hypothetical protein (526 aa) |
| 2793 | 2163 | PPTG_03896 | 70,21 | 9 | 0 | 0 | 9 | | | 1,91E+06 | 4 | 4 | 0 | 0 | 4 | | 57191 | PPTG_03896 | Phytophthora parasitica INRA-310 hypothetical protein (513 aa) |
| 3181 | 2688 | PPTG_02089 | 70,16 | 9 | 0 | 0 | 9 | | | 6,40E+06 | 3 | 3 | 0 | 0 | 3 | | 44551 | PPTG_02089 | Phytophthora parasitica INRA-310 phosphatidylserine decarboxylase (398 aa) |
| 2694 | 2626 | PPTG_11007 | 70,15 | 9 | 0 | 2 | 6 | | 1,04E+06 | 2,50E+06 | 4 | 4 | 0 | 1 | 3 | Carbamidomethylation | 55807 | PPTG_11007 | Phytophthora parasitica INRA-310 Gpr7 transmembrane protein (497 aa) |
| 2598 | 2684 | PPTG_00403 | 70,13 | 15 | 0 | 12 | 15 | | 3,38E+05 | 1,01E+06 | 3 | 3 | 0 | 2 | 3 | | 25874 | PPTG_00403 | Phytophthora parasitica INRA-310 vesicle-associated membrane protein (224 aa) |
| 2603 | 2756 | PPTG_03812 | 70,08 | 7 | 2 | 4 | 4 | 9,01E+04 | 1,16E+05 | 1,82E+06 | 3 | 3 | 1 | 2 | 2 | Carbamidomethylation | 61248 | PPTG_03812 | Phytophthora parasitica INRA-310 aspartyl protease family A01A (560 aa) |
| 2603 | 2757 | PPTG_03812 | 70,08 | 7 | 2 | 4 | 4 | 9,01E+04 | 1,16E+05 | 1,82E+06 | 3 | 3 | 1 | 2 | 2 | Carbamidomethylation | 61335 | PPTG_03812 | Phytophthora parasitica INRA-310 aspartyl protease family A01A (561 aa) |
| 1995 | 1915 | PPTG_13603 | 70,04 | 8 | 6 | 2 | 8 | 3,08E+05 | 1,92E+05 | 3,32E+06 | 4 | 4 | 3 | 1 | 4 | Carbamidomethylation | 64852 | PPTG_13603 | Phytophthora parasitica INRA-310 ATPase (586 aa) |
| 2842 | 2312 | PPTG_01456 | 69,99 | 7 | 0 | 0 | 7 | | | 1,56E+06 | 4 | 4 | 0 | 0 | 4 | | 85049 | PPTG_01456 | Phytophthora parasitica INRA-310 hypothetical protein (761 aa) |
| 2399 | 3314 | PPTG_00748 | 69,97 | 30 | 20 | 30 | 20 | 2,30E+05 | 1,34E+06 | 2,62E+06 | 2 | 2 | 1 | 3 | 2 | | 10620 | PPTG_00748 | Phytophthora parasitica INRA-310 hypothetical protein (90 aa) |
| 1990 | 2088 | PPTG_13216 | 69,97 | 2 | 1 | 2 | 2 | 8,54E+04 | 3,08E+05 | 7,21E+05 | 4 | 4 | 1 | 3 | 4 | | 232078 | PPTG_13216 | Phytophthora parasitica INRA-310 hypothetical protein (2131 aa) |
| 2207 | 2450 | PPTG_01385 | 69,92 | 10 | 6 | 6 | 10 | 3,00E+05 | 4,75E+05 | 2,96E+06 | 3 | 3 | 2 | 2 | 3 | | 36656 | PPTG_01385 | Phytophthora parasitica INRA-310 tricarboxylate transporter (335 aa) |
| 2656 | 2511 | PPTG_16314 | 69,83 | 14 | 0 | 8 | 10 | | 3,32E+05 | 8,44E+05 | 3 | 3 | 0 | 2 | 2 | | 29918 | PPTG_16314 | Phytophthora parasitica INRA-310 hypothetical protein (283 aa) |
| 2297 | 2719 | PPTG_01755 | 69,78 | 25 | 5 | 14 | 20 | 2,92E+05 | 1,28E+06 | 2,55E+06 | 4 | 4 | 1 | 2 | 3 | | 19121 | PPTG_01755 | Phytophthora parasitica INRA-310 hypothetical protein (163 aa) |
| 2853 | 2563 | PPTG_00040 | 69,74 | 27 | 0 | 27 | 11 | | 9,69E+05 | 2,94E+05 | 3 | 3 | 0 | 3 | 1 | | 16647 | PPTG_00040 | Phytophthora parasitica INRA-310 iron-sulfur cluster assembly protein (151 aa) |
| 2011 | 2029 | PPTG_10874 | 69,73 | 6 | 1 | 1 | 6 | 2,29E+04 | 3,54E+05 | 2,45E+06 | 6 | 6 | 1 | 1 | 6 | | 121869 | PPTG_10874 | Phytophthora parasitica INRA-310 alpha-1 3-mannosyltransferase (1083 aa) |
| 2659 | 2577 | PPTG_20155 | 69,71 | 35 | 0 | 8 | 35 | | 9,91E+05 | 3,81E+06 | 3 | 3 | 0 | 1 | 3 | Carbamidomethylation | 13036 | PPTG_20155 | Phytophthora parasitica INRA-310 hypothetical protein (121 aa) |
| 1705 | 3658 | PPTG_06290 | 69,69 | 24 | 0 | 24 | 24 | | 8,05E+06 | 5,96E+06 | 1 | 1 | 0 | 6 | 5 | Oxidation (M); Carbamidomethylation; Deamidation (NQ) | 11942 | PPTG_06290 | Phytophthora parasitica INRA-310 small cysteine rich protein SCR108 (111 aa) |
| 2502 | 3165 | PPTG_05430 | 69,59 | 11 | 6 | 11 | 11 | 6,72E+05 | 8,13E+06 | 8,52E+06 | 2 | 2 | 1 | 2 | 2 | | 24145 | PPTG_05430 | Phytophthora parasitica INRA-310 ATP-dependent protease hslV (225 aa) |
| 2332 | 2497 | PPTG_18983 | 69,56 | 9 | 0 | 9 | 7 | | 5,07E+06 | 4,92E+06 | 3 | 3 | 0 | 3 | 3 | Carbamylation | 49841 | PPTG_18983 | Phytophthora parasitica INRA-310 glycoside hydrolase (444 aa) |
| 2103 | 3005 | PPTG_08478 | 69,53 | 29 | 29 | 29 | 29 | 1,81E+06 | 2,63E+06 | 1,37E+07 | 2 | 2 | 3 | 2 | 3 | Carbamidomethylation; Deamidation (NQ) | 12089 | PPTG_08478 | Phytophthora parasitica INRA-310 hypothetical protein (108 aa) |
| 2984 | 2975 | PPTG_16779 | 69,52 | 40 | 21 | 40 | 0 | 1,04E+05 | 7,57E+05 | | 2 | 2 | 1 | 2 | 0 | | 13083 | PPTG_16779 | Phytophthora parasitica INRA-310 hypothetical protein (122 aa) |
| 2223 | 2321 | PPTG_04349 | 69,49 | 6 | 1 | 1 | 6 | 2,47E+05 | 8,08E+05 | 4,76E+06 | 5 | 5 | 1 | 1 | 5 | Carbamidomethylation | 100625 | PPTG_04349 | Phytophthora parasitica INRA-310 ribosomal protein S6 kinase (906 aa) |
| 2564 | 2150 | PPTG_09247 | 69,48 | 10 | 0 | 6 | 7 | | 3,03E+05 | 7,96E+05 | 4 | 4 | 0 | 2 | 3 | Carbamidomethylation | 50617 | PPTG_09247 | Phytophthora parasitica INRA-310 hypothetical protein (466 aa) |
| 2763 | 2112 | PPTG_18899 | 69,46 | 6 | 0 | 0 | 6 | | | 5,90E+05 | 4 | 4 | 0 | 0 | 4 | Carbamidomethylation | 105560 | PPTG_18899 | Phytophthora parasitica INRA-310 hypothetical protein (980 aa) |
| 3041 | 3117 | PPTG_07631 | 69,45 | 8 | 0 | 0 | 8 | | | 3,93E+06 | 2 | 2 | 0 | 0 | 3 | Carbamidomethylation | 46050 | PPTG_07631 | Phytophthora parasitica INRA-310 ATP-binding Cassette (ABC) superfamily (409 aa) |
| 2102 | 2548 | PPTG_08045 | 69,36 | 40 | 0 | 37 | 40 | | 1,26E+06 | 1,86E+06 | 4 | 4 | 0 | 4 | 4 | Oxidation (M) | 11866 | PPTG_08045 | Phytophthora parasitica INRA-310 thioredoxin (106 aa) |
| 2427 | 2335 | PPTG_15888 | 69,33 | 12 | 3 | 10 | 6 | 3,72E+04 | 6,34E+05 | 4,37E+05 | 4 | 4 | 1 | 3 | 2 | Carbamidomethylation | 43862 | PPTG_15888 | Phytophthora parasitica INRA-310 polycomb protein (396 aa) |
| 2518 | 2402 | PPTG_13913 | 69,23 | 21 | 0 | 16 | 21 | | 8,58E+05 | 1,88E+06 | 3 | 3 | 0 | 2 | 3 | Oxidation (M); Carbamidomethylation | 26206 | PPTG_13913 | Phytophthora parasitica INRA-310 eukaryotic translation initiation factor 2 subunit beta (234 aa) |
| 2525 | 2622 | PPTG_15641 | 69,19 | 39 | 0 | 16 | 39 | | 4,52E+05 | 2,50E+06 | 4 | 4 | 0 | 1 | 4 | Deamidation (NQ) | 12137 | PPTG_15641 | Phytophthora parasitica INRA-310 hypothetical protein (107 aa) |
| 1835 | 2265 | PPTG_19283 | 69,19 | 32 | 0 | 32 | 24 | | 3,73E+06 | 4,94E+06 | 4 | 4 | 0 | 5 | 5 | | 14650 | PPTG_19283 | Phytophthora parasitica INRA-310 hypothetical protein (130 aa) |
| 2511 | 2711 | PPTG_03624 | 69,14 | 11 | 0 | 6 | 9 | | 1,91E+05 | 1,84E+06 | 4 | 4 | 0 | 2 | 3 | | 58210 | PPTG_03624 | Phytophthora parasitica INRA-310 hypothetical protein (513 aa) |
| 1465 | 2634 | PPTG_05264 | 69,09 | 19 | 12 | 14 | 19 | 9,08E+05 | 2,49E+06 | 5,08E+06 | 3 | 2 | 3 | 5 | 6 | Deamidation (NQ); Carbamylation | 14569 | PPTG_05264 | Phytophthora parasitica INRA-310 histone H2A (137 aa) |
| 3466 | 2865 | PPTG_09632 | 69,05 | 5 | 0 | 0 | 5 | | | 1,72E+06 | 2 | 2 | 0 | 0 | 2 | | 55916 | PPTG_09632 | Phytophthora parasitica INRA-310 hypothetical protein (504 aa) |
| 2278 | 3029 | PPTG_09284 | 69,04 | 6 | 6 | 6 | 6 | 6,48E+06 | 1,68E+07 | 4,77E+07 | 2 | 2 | 2 | 2 | 2 | | 40661 | PPTG_09284 | Phytophthora parasitica INRA-310 hypothetical protein (405 aa) |
| 2200 | 3140 | PPTG_14010 | 69 | 10 | 7 | 10 | 10 | 6,69E+04 | 2,94E+06 | 1,97E+06 | 2 | 2 | 1 | 3 | 3 | | 24295 | PPTG_14010 | Phytophthora parasitica INRA-310 ATP-dependent Clp protease proteolytic subunit (223 aa) |
| 2299 | 1988 | PPTG_11712 | 68,96 | 14 | 0 | 14 | 2 | | 1,47E+06 | 3,28E+05 | 5 | 5 | 0 | 5 | 1 | Carbamidomethylation | 53959 | PPTG_11712 | Phytophthora parasitica INRA-310 Pro-apoptotic serine protease nma111-like protein (491 aa) |
| 2774 | 2773 | PPTG_10898 | 68,95 | 7 | 2 | 0 | 7 | 8,48E+04 | | 5,87E+05 | 3 | 3 | 1 | 0 | 3 | | 58911 | PPTG_10898 | Phytophthora parasitica INRA-310 Iron(III)-zinc(II) purple acid phosphatase (522 aa) |
| 2820 | 3198 | PPTG_09680 | 68,88 | 11 | 0 | 11 | 11 | | 4,44E+06 | 5,75E+06 | 2 | 2 | 0 | 2 | 2 | Carbamidomethylation | 33483 | PPTG_09680 | Phytophthora parasitica INRA-310 ribonuclease (316 aa) |
| 2541 | 2452 | PPTG_08507 | 68,87 | 6 | 0 | 6 | 4 | | 9,62E+05 | 1,99E+05 | 3 | 3 | 0 | 3 | 2 | Carbamidomethylation | 76603 | PPTG_08507 | Phytophthora parasitica INRA-310 glucosylceramidase (681 aa) |
| 2300 | 2039 | PPTG_08275 | 68,86 | 19 | 5 | 5 | 19 | 2,34E+05 | 3,92E+05 | 2,50E+06 | 4 | 4 | 1 | 1 | 4 | Carbamidomethylation | 24522 | PPTG_08275 | Phytophthora parasitica INRA-310 Rab32 family GTPase (219 aa) |
| 2696 | 2807 | PPTG_20424 | 68,82 | 8 | 5 | 3 | 8 | 1,39E+05 | | 1,23E+06 | 2 | 1 | 1 | 1 | 2 | | 28800 | PPTG_20424 | Phytophthora parasitica INRA-310 UDP-glucose 4-epimerase (263 aa) |
| 2847 | 2507 | PPTG_00288 | 68,78 | 7 | 0 | 0 | 7 | | | 1,57E+06 | 4 | 4 | 0 | 0 | 4 | Carbamidomethylation | 84181 | PPTG_00288 | Phytophthora parasitica INRA-310 hypothetical protein (743 aa) |
| 1548 | 3358 | PPTG_20376 | 68,77 | 25 | 0 | 25 | 25 | | 2,34E+07 | 2,08E+07 | 2 | 2 | 0 | 8 | 5 | Oxidation (M); Carbamidomethylation; Acetylation (N-term) | 20231 | PPTG_20376 | Phytophthora parasitica INRA-310 hypothetical protein (187 aa) |
| 2088 | 2219 | PPTG_17569 | 68,76 | 12 | 8 | 12 | 12 | 6,55E+05 | 6,06E+06 | 7,60E+06 | 3 | 3 | 2 | 3 | 3 | | 33437 | PPTG_17569 | Phytophthora parasitica INRA-310 spermidine synthase (296 aa) |
| 2423 | 3392 | PPTG_17477 | 68,75 | 12 | 0 | 12 | 12 | | 9,92E+06 | 1,04E+07 | 2 | 2 | 0 | 3 | 3 | Oxidation (M); Carbamidomethylation | 42306 | PPTG_17477 | Phytophthora parasitica INRA-310 carbohydrate-binding protein (414 aa) |
| 2913 | 3361 | PPTG_02755 | 68,7 | 12 | 0 | 12 | 12 | | 1,54E+06 | 1,29E+06 | 2 | 2 | 0 | 2 | 2 | | 28719 | PPTG_02755 | Phytophthora parasitica INRA-310 hypothetical protein (264 aa) |
| 2798 | 2467 | PPTG_17412 | 68,62 | 20 | 0 | 7 | 20 | | 2,40E+05 | 2,61E+05 | 3 | 3 | 0 | 1 | 3 | | 18014 | PPTG_17412 | Phytophthora parasitica INRA-310 hypothetical protein (159 aa) |
| 2912 | 3356 | PPTG_06567 | 68,59 | 31 | 11 | 11 | 31 | 7,77E+05 | 2,55E+06 | 7,38E+06 | 2 | 2 | 1 | 1 | 2 | | 10652 | PPTG_06567 | Phytophthora parasitica INRA-310 mitochondrial protein Translocase (MPT) family (89 aa) |
| 2446 | 2475 | PPTG_15111 | 68,51 | 2 | 0 | 2 | 2 | | 3,22E+05 | 3,80E+05 | 3 | 3 | 0 | 2 | 3 | | 170018 | PPTG_15111 | Phytophthora parasitica INRA-310 kinesin-like protein (1506 aa) |
| 3252 | 3437 | PPTG_01301 | 68,47 | 6 | 3 | 3 | 3 | 1,53E+05 | 1,15E+06 | 2,13E+05 | 2 | 2 | 1 | 1 | 1 | Carbamidomethylation | 51965 | PPTG_01301 | Phytophthora parasitica INRA-310 hypothetical protein (488 aa) |
| 1936 | 2026 | PPTG_03983 | 68,44 | 16 | 3 | 10 | 16 | 9,65E+04 | 2,44E+06 | 3,58E+06 | 5 | 4 | 1 | 3 | 5 | Carbamidomethylation | 41737 | PPTG_03983 | Phytophthora parasitica INRA-310 methionine aminopeptidase (379 aa) |
| 2397 | 2875 | PPTG_17687 | 68,43 | 12 | 3 | 12 | 7 | 1,72E+05 | 1,21E+06 | 1,39E+06 | 3 | 3 | 1 | 3 | 2 | | 39626 | PPTG_17687 | Phytophthora parasitica INRA-310 hypothetical protein (366 aa) |
| 2654 | 2867 | PPTG_01592 | 68,31 | 14 | 0 | 9 | 9 | | 2,57E+05 | 4,28E+05 | 3 | 3 | 0 | 2 | 2 | Acetylation (N-term) | 33016 | PPTG_01592 | Phytophthora parasitica INRA-310 hypothetical protein (298 aa) |
| 2491 | 2224 | PPTG_11861 | 68,16 | 10 | 0 | 0 | 10 | | | 4,94E+06 | 5 | 5 | 0 | 0 | 5 | Carbamidomethylation | 73161 | PPTG_11861 | Phytophthora parasitica INRA-310 signal peptide peptidase-like aspartyl protease family A22B (676 aa) |
| 2296 | 2560 | PPTG_18044 | 68 | 6 | 0 | 6 | 2 | | 1,23E+05 | 1,95E+05 | 4 | 4 | 0 | 4 | 2 | | 116865 | PPTG_18044 | Phytophthora parasitica INRA-310 metalloprotease family M16C (1048 aa) |
| 2450 | 2151 | PPTG_05296 | 67,97 | 2 | 0 | 2 | 1 | | 7,45E+05 | 5,84E+05 | 5 | 5 | 0 | 4 | 1 | | 291435 | PPTG_05296 | Phytophthora parasitica INRA-310 ubiquitin-specific protease (2684 aa) |
| 2303 | 2234 | PPTG_02371 | 67,97 | 12 | 0 | 5 | 12 | | 1,12E+06 | 7,87E+06 | 4 | 4 | 0 | 2 | 4 | Carbamidomethylation | 46532 | PPTG_02371 | Phytophthora parasitica INRA-310 glycogen Synthase Kinase 3 beta (413 aa) |
| 2415 | 2741 | PPTG_06513 | 67,96 | 5 | 2 | 4 | 5 | 1,08E+05 | 3,97E+05 | 2,18E+06 | 3 | 3 | 1 | 2 | 3 | | 68186 | PPTG_06513 | Phytophthora parasitica INRA-310 endomembrane protein 70-like protein (599 aa) |
| 2389 | 2285 | PPTG_19076 | 67,95 | 7 | 0 | 5 | 4 | | 7,02E+05 | 4,41E+05 | 4 | 4 | 0 | 3 | 3 | Oxidation (M) | 81049 | PPTG_19076 | Phytophthora parasitica INRA-310 hypothetical protein (736 aa) |
| 2314 | 2096 | PPTG_16120 | 67,92 | 19 | 0 | 19 | 9 | | 1,41E+06 | 1,35E+06 | 4 | 4 | 0 | 4 | 2 | Carbamidomethylation | 23120 | PPTG_16120 | Phytophthora parasitica INRA-310 hypothetical protein (213 aa) |
| 3179 | 2435 | PPTG_02647 | 67,9 | 5 | 0 | 0 | 5 | | | 1,00E+06 | 3 | 3 | 0 | 0 | 3 | | 102568 | PPTG_02647 | Phytophthora parasitica INRA-310 hypothetical protein (920 aa) |
| 2496 | 2615 | PPTG_06497 | 67,9 | 12 | 0 | 10 | 8 | | 1,33E+06 | 1,18E+06 | 4 | 4 | 0 | 2 | 3 | | 30314 | PPTG_06497 | Phytophthora parasitica INRA-310 iron-sulfur cluster scaffold protein Nfu-like protein (273 aa) |
| 1801 | 2498 | PPTG_01987 | 67,79 | 2 | 1 | 2 | 2 | | | 8,36E+05 | 3 | 1 | 2 | 3 | 5 | | 131960 | PPTG_01987 | Phytophthora parasitica INRA-310 protein kinase (1216 aa) |
| 2648 | 4213 | PPTG_02532 | 67,78 | 10 | 0 | 10 | 10 | | 1,91E+06 | 1,39E+06 | 1 | 1 | 0 | 3 | 2 | Carbamidomethylation | 68057 | PPTG_02532 | Phytophthora parasitica INRA-310 hypothetical protein (662 aa) |
| 2071 | 2069 | PPTG_01670 | 67,78 | 14 | 4 | 11 | 2 | 2,78E+05 | 4,48E+06 | 3,52E+05 | 5 | 5 | 2 | 5 | 1 | Dehydration; Deamidation (NQ) | 50186 | PPTG_01670 | Phytophthora parasitica INRA-310 serine protease family S10 (457 aa) |
| 2381 | 3228 | PPTG_00663 | 67,77 | 20 | 12 | 20 | 20 | 9,11E+04 | 1,77E+06 | 1,65E+06 | 2 | 2 | 1 | 3 | 2 | | 17692 | PPTG_00663 | Phytophthora parasitica INRA-310 zinc-binding protein (164 aa) |
| 2244 | 2640 | PPTG_09853 | 67,72 | 6 | 1 | 4 | 3 | 7,94E+04 | 5,30E+05 | 1,21E+06 | 4 | 3 | 1 | 3 | 3 | Carbamidomethylation | 82573 | PPTG_09853 | Phytophthora parasitica INRA-310 protein kinase (762 aa) |
| 2161 | 2139 | PPTG_09480 | 67,72 | 4 | 1 | 1 | 4 | | | 2,35E+06 | 5 | 4 | 1 | 1 | 5 | Carbamidomethylation | 121649 | PPTG_09480 | Phytophthora parasitica INRA-310 mitogen-activated protein kinase (1098 aa) |
| 3231 | 4214 | PPTG_03148 | 67,7 | 10 | 10 | 10 | 10 0 | | 1,27E+06 | 7,45E+05 | 1 | 1 | 1 | 1 | 1 | Carbamidomethylation | 30212 | PPTG_03148 | Phytophthora parasitica INRA-310 hypothetical protein (284 aa) |
| 2508 | 2587 | PPTG_18884 | 67,68 | 3 | 2 | 2 | 3 | 7,69E+04 | 3,12E+05 | 1,41E+06 | 3 | 3 | 1 | 1 | 3 | | 88717 | PPTG_18884 | Phytophthora parasitica INRA-310 hypothetical protein (812 aa) |
| 2849 | 2533 | PPTG_11780 | 67,63 | 25 | 0 | 6 | 25 | | 8,11E+04 | 1,42E+06 | 3 | 3 | 0 | 1 | 3 | | 18488 | PPTG_11780 | Phytophthora parasitica INRA-310 hypothetical protein (172 aa) |
| 2221 | 2168 | PPTG_04397 | 67,59 | 9 | 2 | 7 | 5 | 6,67E+04 | 1,26E+06 | 9,35E+05 | 5 | 5 | 1 | 4 | 2 | Dehydration; Deamidation (NQ); Acetylation (N-term) | 67391 | PPTG_04397 | Phytophthora parasitica INRA-310 hypothetical protein (599 aa) |
| 2186 | 2130 | PPTG_14050T2 | 67,58 | 5 | 0 | 5 | 4 | | 7,74E+05 | 1,00E+06 | 4 | 4 | 0 | 4 | 3 | | 133512 | PPTG_14050 | Phytophthora parasitica INRA-310 hypothetical protein (1214 aa) |
| 2186 | 2131 | PPTG_14050 | 67,58 | 5 | 0 | 5 | 4 | | 7,74E+05 | 1,00E+06 | 4 | 4 | 0 | 4 | 3 | | 140702 | PPTG_14050 | Phytophthora parasitica INRA-310 hypothetical protein (1277 aa) |
| 2324 | 2778 | PPTG_11232 | 67,54 | 6 | 0 | 4 | 6 | | 5,45E+05 | 7,00E+05 | 3 | 3 | 0 | 2 | 4 | Deamidation (NQ) | 97191 | PPTG_11232 | Phytophthora parasitica INRA-310 hypothetical protein (891 aa) |
| 2362 | 2035 | PPTG_14598 | 67,46 | 14 | 5 | 0 | 11 | | | 1,58E+06 | 5 | 3 | 2 | 0 | 4 | Oxidation (M) | 39301 | PPTG_14598 | Phytophthora parasitica INRA-310 hypothetical protein (349 aa) |
| 2193 | 2476 | PPTG_00209 | 67,46 | 4 | 2 | 4 | 2 | 1,81E+05 | 1,07E+06 | 1,33E+06 | 3 | 3 | 2 | 3 | 2 | | 87433 | PPTG_00209 | Phytophthora parasitica INRA-310 Cullin family protein (756 aa) |
| 1865 | 2173 | PPTG_02637 | 67,42 | 8 | 2 | 8 | 8 | 7,80E+04 | 1,62E+06 | 1,76E+06 | 4 | 4 | 1 | 4 | 4 | Carbamidomethylation; Acetylation (N-term) | 69303 | PPTG_02637 | Phytophthora parasitica INRA-310 ubiquitin-activating enzyme (E1) (625 aa) |
| 2114 | 2083 | PPTG_12866 | 67,4 | 3 | 0 | 2 | 2 | | 4,98E+05 | 7,78E+05 | 5 | 5 | 0 | 3 | 4 | | 223639 | PPTG_12866 | Phytophthora parasitica INRA-310 hypothetical protein (2046 aa) |
| 2273 | 2898 | PPTG_01168 | 67,4 | 4 | 0 | 2 | 4 | | 4,90E+05 | 3,37E+06 | 3 | 3 | 0 | 2 | 4 | Oxidation (M) | 67596 | PPTG_01168 | Phytophthora parasitica INRA-310 hypothetical protein (654 aa) |
| 2520 | 2473 | PPTG_01554 | 67,39 | 14 | 0 | 10 | 14 | | 7,18E+05 | 2,99E+06 | 3 | 3 | 0 | 2 | 3 | | 23455 | PPTG_01554 | Phytophthora parasitica INRA-310 hypothetical protein (212 aa) |
| 2101 | 2474 | PPTG_05692 | 67,34 | 29 | 0 | 29 | 29 | | 6,25E+06 | 5,86E+06 | 3 | 3 | 0 | 4 | 4 | Oxidation (M) | 12837 | PPTG_05692 | Phytophthora parasitica INRA-310 hypothetical protein (120 aa) |
| 2132 | 2421 | PPTG_19425 | 67,29 | 26 | 0 | 17 | 15 | | 1,28E+06 | 1,29E+06 | 5 | 5 | 0 | 3 | 3 | Deamidation (NQ) | 24374 | PPTG_19425 | Phytophthora parasitica INRA-310 short chain dehydrogenase (232 aa) |
| 2612 | 3214 | PPTG_07123 | 67,25 | 11 | 0 | 11 | 11 | | 5,12E+05 | 5,95E+05 | 2 | 2 | 0 | 3 | 2 | Carbamidomethylation | 53476 | PPTG_07123 | Phytophthora parasitica INRA-310 hypothetical protein (492 aa) |
| 2437 | 3368 | PPTG_12305 | 67,23 | 9 | 0 | 9 | 9 | | 2,01E+06 | 2,48E+06 | 2 | 2 | 0 | 3 | 3 | Carbamidomethylation; Deamidation (NQ) | 31912 | PPTG_12305 | Phytophthora parasitica INRA-310 GP-anchored serine-rich elicitin INL13-like protein (311 aa) |
| 2448 | 2996 | PPTG_06148 | 67,23 | 2 | 1 | 2 | 2 | 1,28E+05 | 1,09E+06 | 2,04E+06 | 2 | 2 | 1 | 2 | 2 | | 120009 | PPTG_06148 | Phytophthora parasitica INRA-310 DEAD/DEAH box RNA helicase (1067 aa) |
| 2206 | 2441 | PPTG_00245 | 67,21 | 10 | 2 | 6 | 8 | 2,93E+05 | 2,71E+06 | 2,05E+06 | 4 | 4 | 1 | 3 | 3 | | 45691 | PPTG_00245 | Phytophthora parasitica INRA-310 tryptophan synthase beta chain (425 aa) |
| 2483 | 2428 | PPTG_09092 | 67,21 | 25 | 6 | 6 | 18 | 3,25E+05 | 2,21E+05 | 2,99E+06 | 4 | 4 | 1 | 1 | 3 | | 23402 | PPTG_09092 | Phytophthora parasitica INRA-310 Rab2 family GTPase (208 aa) |
| 2305 | 2385 | PPTG_16370 | 67,16 | 7 | 2 | 5 | 3 | 9,63E+04 | 8,20E+05 | 1,23E+06 | 4 | 4 | 1 | 3 | 2 | Carbamidomethylation | 74637 | PPTG_16370 | Phytophthora parasitica INRA-310 hypothetical protein (682 aa) |
| 2933 | 2881 | PPTG_12934 | 67,09 | 7 | 0 | 7 | 2 | | 6,42E+05 | 5,98E+05 | 3 | 3 | 0 | 3 | 1 | | 59219 | PPTG_12934 | Phytophthora parasitica INRA-310 hypothetical protein (540 aa) |
| 2168 | 2664 | PPTG_14008 | 67,06 | 17 | 0 | 4 | 14 | | 7,31E+05 | 1,32E+06 | 6 | 6 | 0 | 2 | 5 | Carbamidomethylation; Acetylation (N-term) | 44777 | PPTG_14008 | Phytophthora parasitica INRA-310 phosphomannose isomerase type I (404 aa) |
| 1846 | 3269 | PPTG_14554 | 67,04 | 10 | 4 | 4 | 10 | | | 2,35E+05 | 3 | 1 | 2 | 2 | 6 | Carbamylation | 25073 | PPTG_14554 | Phytophthora parasitica INRA-310 adenylate kinase (229 aa) |
| 2012 | 2137 | PPTG_08783 | 66,98 | 20 | 0 | 12 | 20 | | 7,15E+05 | 1,18E+06 | 5 | 5 | 0 | 3 | 5 | Carbamidomethylation; Acetylation (N-term) | 28904 | PPTG_08783 | Phytophthora parasitica INRA-310 hypothetical protein (257 aa) |
| 2594 | 2554 | PPTG_10083 | 66,88 | 3 | 0 | 3 | 2 | | 7,36E+05 | 8,15E+05 | 3 | 3 | 0 | 3 | 2 | | 120806 | PPTG_10083 | Phytophthora parasitica INRA-310 inositol hexakisphosphate and diphosphoinositol-pentakisphosphate kinase (1067 aa) |
| 2004 | 2235 | PPTG_06916 | 66,86 | 1 | 0 | 1 | 1 | | 2,63E+05 | 1,69E+05 | 5 | 3 | 0 | 5 | 3 | | 559030 | PPTG_06916 | Phytophthora parasitica INRA-310 dynein heavy chain (4980 aa) |
| 2004 | 2236 | PPTG_06916 | 66,86 | 1 | 0 | 1 | 1 | | 2,63E+05 | 1,69E+05 | 5 | 3 | 0 | 5 | 3 | | 559030 | PPTG_06916 | Phytophthora parasitica INRA-310 dynein heavy chain (4980 aa) |
| 2526 | 2649 | PPTG_17632 | 66,77 | 6 | 0 | 6 | 4 | | 5,80E+05 | 6,23E+05 | 3 | 3 | 0 | 3 | 2 | Carbamidomethylation | 82725 | PPTG_17632 | Phytophthora parasitica INRA-310 cleavage and polyadenylation specificity factor subunit (737 aa) |
| 3782 | 4212 | PPTG_01614 | 66,72 | 26 | 0 | 0 | 26 | | | 1,53E+06 | 1 | 1 | 0 | 0 | 2 | Acetylation (N-term) | 15976 | PPTG_01614 | Phytophthora parasitica INRA-310 hypothetical protein (142 aa) |
| 3111 | 3234 | PPTG_05465 | 66,71 | 11 | 0 | 7 | 11 | | 1,45E+05 | 5,41E+05 | 2 | 2 | 0 | 1 | 2 | | 30833 | PPTG_05465 | Phytophthora parasitica INRA-310 serine protease family S01A (291 aa) |
| 2515 | 2133 | PPTG_11089 | 66,69 | 5 | 1 | 0 | 5 | 4,29E+04 | | 2,92E+06 | 4 | 4 | 1 | 0 | 4 | | 89299 | PPTG_11089 | Phytophthora parasitica INRA-310 hypothetical protein (797 aa) |
| 2515 | 2134 | PPTG_11089 | 66,69 | 5 | 1 | 0 | 5 | 4,29E+04 | | 2,92E+06 | 4 | 4 | 1 | 0 | 4 | | 90715 | PPTG_11089 | Phytophthora parasitica INRA-310 hypothetical protein (810 aa) |
| 2350 | 2594 | PPTG_08482 | 66,62 | 4 | 1 | 3 | 4 | 2,77E+04 | 5,60E+05 | 9,89E+05 | 3 | 3 | 1 | 2 | 3 | Carbamidomethylation | 77415 | PPTG_08482 | Phytophthora parasitica INRA-310 protein kinase (695 aa) |

| | | | | | | | | | | | | | | | | | |
|---|---|---|---|---|---|---|---|---|---|---|---|---|---|---|---|---|---|
| 2431 | 2710 | PPTG_14791 | 66,6 | 26 | 7 | 7 | 26 | 1,33E+06 | 5,26E+06 | 1,19E+07 | 4 | 4 | 1 | 1 | 4 | | 15947 \| PPTG_14791 \| Phytophthora parasitica INRA-310 50S ribosomal protein L13 (141 aa) |
| 2845 | 2438 | PPTG_05519 | 66,58 | 5 | 0 | 0 | 5 | | | 1,26E+06 | 4 | 4 | 0 | 0 | 4 | | 91713 \| PPTG_05519 \| Phytophthora parasitica INRA-310 protein kinase (811 aa) |
| 2591 | 2494 | PPTG_20397 | 66,51 | 8 | 0 | 5 | 8 | | 6,15E+05 | 3,21E+06 | 3 | 3 | 0 | 2 | 3 | | 40929 \| PPTG_20397 \| Phytophthora parasitica INRA-310 calcineurin-like phosphoesterase (354 aa) |
| 1991 | 2215 | PPTG_11497 | 66,48 | 7 | 0 | 3 | 7 | | 1,60E+06 | 1,03E+07 | 4 | 4 | 0 | 2 | 6 | | 57537 \| PPTG_11497 \| Phytophthora parasitica INRA-310 hypothetical protein (523 aa) |
| 1991 | 2216 | PPTG_11497 | 66,48 | 7 | 0 | 3 | 7 | | 1,60E+06 | 1,03E+07 | 4 | 4 | 0 | 2 | 6 | | 57537 \| PPTG_11497 \| Phytophthora parasitica INRA-310 hypothetical protein (523 aa) |
| 2476 | 2641 | PPTG_07653 | 66,45 | 28 | 0 | 8 | 28 | | 1,67E+06 | 3,36E+06 | 3 | 3 | 0 | 1 | 4 | Carbamidomethylation | 20992 \| PPTG_07653 \| Phytophthora parasitica INRA-310 cleavage induced protein (200 aa) |
| 2943 | 3488 | PPTG_03499 | 66,39 | 6 | 0 | 6 | 6 | | 9,63E+05 | 1,04E+06 | 2 | 2 | 0 | 2 | 2 | | 52437 \| PPTG_03499 \| Phytophthora parasitica INRA-310 mannan endo-1 4-beta-mannosidase (493 aa) |
| 2054 | 2064 | PPTG_19158 | 66,27 | 8 | 0 | 4 | 8 | | 6,79E+05 | 1,80E+06 | 5 | 5 | 0 | 3 | 5 | Oxidation (M) | 83476 \| PPTG_19158 \| Phytophthora parasitica INRA-310 6-phosphofructo-2-kinase/fructose-2 6-biphosphatase (742 aa) |
| 2745 | 2669 | PPTG_10326 | 66,23 | 5 | 0 | 3 | 3 | | 3,10E+05 | 3,76E+05 | 3 | 3 | 0 | 2 | 2 | Carbamidomethylation | 89851 \| PPTG_10326 \| Phytophthora parasitica INRA-310 pseudouridylate synthase (810 aa) |
| 2136 | 2143 | PPTG_19168 | 66,23 | 10 | 0 | 10 | 10 | | 1,37E+06 | 3,26E+06 | 3 | 3 | 0 | 4 | 3 | Dehydration; Deamidation (NQ) | 38470 \| PPTG_19168 \| Phytophthora parasitica INRA-310 arabinogalactan endo-1 4-beta-galactosidase (359 aa) |
| 2840 | 2255 | PPTG_11869 | 66,2 | 6 | 0 | 0 | 6 | | | 8,23E+05 | 4 | 4 | 0 | 0 | 4 | | 87774 \| PPTG_11869 \| Phytophthora parasitica INRA-310 hypothetical protein (788 aa) |
| 2176 | 2138 | PPTG_16827 | 66,15 | 11 | 2 | 10 | 6 | 8,79E+04 | 1,51E+06 | 1,75E+05 | 5 | 5 | 1 | 4 | 2 | Carbamidomethylation | 59746 \| PPTG_16827 \| Phytophthora parasitica INRA-310 hypothetical protein (542 aa) |
| 2546 | 2644 | PPTG_04012 | 66,14 | 8 | 0 | 6 | 8 | | 6,85E+05 | 1,84E+06 | 3 | 3 | 0 | 2 | 3 | Carbamidomethylation | 60643 \| PPTG_04012 \| Phytophthora parasitica INRA-310 lysosomal Pro-X carboxypeptidase (543 aa) |
| 1719 | 2666 | PPTG_18298 | 66,07 | 4 | 2 | 1 | 4 | 4,72E+04 | | 2,74E+05 | 3 | 1 | 3 | 2 | 6 | Oxidation (M) | 81900 \| PPTG_18298 \| Phytophthora parasitica INRA-310 ATP-binding Cassette (ABC) superfamily (730 aa) |
| 2113 | 2056 | PPTG_10911 | 65,98 | 4 | 0 | 3 | 3 | | 4,76E+05 | 2,53E+06 | 5 | 5 | 0 | 3 | 4 | Carbamidomethylation | 159175 \| PPTG_10911 \| Phytophthora parasitica INRA-310 hypothetical protein (1438 aa) |
| 1764 | 1898 | PPTG_13721 | 65,96 | 5 | 0 | 4 | 4 | | 1,70E+06 | 3,22E+06 | 5 | 5 | 0 | 5 | 5 | Oxidation (M) | 123405 \| PPTG_13721 \| Phytophthora parasitica INRA-310 FACT complex subunit SPT16 (1079 aa) |
| 2873 | 3242 | PPTG_05231 | 65,92 | 14 | 6 | 6 | 14 | 1,83E+05 | 2,15E+05 | 2,63E+06 | 2 | 2 | 1 | 1 | 2 | | 24162 \| PPTG_05231 \| Phytophthora parasitica INRA-310 hypothetical protein (217 aa) |
| 2873 | 3243 | PPTG_05231 | 65,92 | 14 | 6 | 6 | 14 | 1,83E+05 | 2,15E+05 | 2,63E+06 | 2 | 2 | 1 | 1 | 2 | | 24162 \| PPTG_05231 \| Phytophthora parasitica INRA-310 hypothetical protein (217 aa) |
| 3019 | 3133 | PPTG_08459 | 65,92 | 5 | 0 | 3 | 5 | | 1,16E+05 | 6,54E+05 | 2 | 2 | 0 | 1 | 2 | Carbamidomethylation | 61327 \| PPTG_08459 \| Phytophthora parasitica INRA-310 xaa-Pro dipeptidase metalloprotease family M24B (554 aa) |
| 2125 | 2426 | PPTG_05586 | 65,92 | 1 | 0 | 1 | 0 | | 8,76E+04 | 2,50E+05 | 4 | 3 | 1 | 4 | 2 | | 331607 \| PPTG_05586 \| Phytophthora parasitica INRA-310 U3 small nucleolar RNA-associated protein (2969 aa) |
| 2190 | 2346 | PPTG_10665 | 65,89 | 9 | 4 | 5 | 8 | 2,84E+04 | 3,92E+05 | 1,80E+06 | 4 | 4 | 2 | 2 | 3 | Deamidation (NQ) | 58317 \| PPTG_10665 \| Phytophthora parasitica INRA-310 serine protease family S08A (533 aa) |
| 2881 | 3364 | PPTG_03649 | 65,86 | 31 | 0 | 31 | 31 | | 8,38E+05 | 4,66E+06 | 2 | 2 | 0 | 2 | 2 | Oxidation (M) | 11588 \| PPTG_03649 \| Phytophthora parasitica INRA-310 hypothetical protein (111 aa) |
| 2034 | 2588 | PPTG_01498 | 65,82 | 17 | 0 | 13 | 14 | | 1,35E+06 | 3,06E+06 | 5 | 5 | 0 | 4 | 4 | Carbamidomethylation | 35697 \| PPTG_01498 \| Phytophthora parasitica INRA-310 eukaryotic translation initiation factor 3 subunit G (326 aa) |
| 2346 | 2174 | PPTG_05073 | 65,76 | 11 | 0 | 4 | 8 | | 5,06E+05 | 3,73E+06 | 5 | 5 | 0 | 2 | 4 | Oxidation (M) | 58282 \| PPTG_05073 \| Phytophthora parasitica INRA-310 polysaccharide lyase (545 aa) |
| 2851 | 2553 | PPTG_00746 | 65,7 | 16 | 0 | 16 | 3 | | 1,02E+06 | 1,50E+06 | 3 | 3 | 0 | 3 | 1 | | 25071 \| PPTG_00746 \| Phytophthora parasitica INRA-310 cGMP 3' 5'-cyclic phosphodiesterase subunit delta (218 aa) |
| 2893 | 2205 | PPTG_17663 | 65,69 | 9 | 0 | 0 | 9 | | | 4,89E+05 | 4 | 2 | 0 | 0 | 4 | | 57998 \| PPTG_17663 \| Phytophthora parasitica INRA-310 dicarboxylate/Amino Acid:Cation (Na or H) Symporter (DAACS) family protein (539 aa) |
| 2935 | 3365 | PPTG_15934 | 65,67 | 20 | 11 | 11 | 20 | 2,36E+05 | 4,08E+05 | 3,30E+06 | 2 | 2 | 1 | 1 | 2 | | 13006 \| PPTG_15934 \| Phytophthora parasitica INRA-310 hypothetical protein (124 aa) |
| 2841 | 2307 | PPTG_14163 | 65,66 | 5 | 0 | 0 | 5 | | | 6,91E+05 | 4 | 4 | 0 | 0 | 4 | | 133155 \| PPTG_14163 \| Phytophthora parasitica INRA-310 transient Receptor Ca2 Channel (Trp-CC) family (1198 aa) |
| 1704 | 3025 | PPTG_07664 | 65,61 | 11 | 11 | 11 | 11 | 2,07E+06 | 5,86E+06 | 1,71E+07 | 2 | 2 | 3 | 4 | 4 | Oxidation (M) | 27101 \| PPTG_07664 \| Phytophthora parasitica INRA-310 NPP1-like protein (246 aa) |
| 2436 | 3258 | PPTG_01341 | 65,6 | 3 | 3 | 3 | 3 | 1,16E+05 | 2,83E+05 | 5,89E+05 | 2 | 1 | 2 | 2 | 2 | Carbamidomethylation | 82375 \| PPTG_01341 \| Phytophthora parasitica INRA-310 threonyl-tRNA synthetase (724 aa) |
| 2869 | 3208 | PPTG_13979 | 65,59 | 27 | 14 | 27 | 14 | 7,19E+04 | 1,04E+06 | 1,14E+06 | 2 | 2 | 1 | 2 | 1 | | 11637 \| PPTG_13979 \| Phytophthora parasitica INRA-310 transcription elongation factor B polypeptide 1 (106 aa) |
| 2359 | 3190 | PPTG_06711 | 65,5 | 13 | 0 | 13 | 13 | | 1,82E+06 | 1,66E+06 | 3 | 3 | 0 | 3 | 3 | Acetylation (N-term) | 24661 \| PPTG_06711 \| Phytophthora parasitica INRA-310 hypothetical protein (227 aa) |
| 2315 | 2125 | PPTG_01429 | 65,43 | 6 | 0 | 4 | 3 | | 3,52E+05 | 4,42E+04 | 5 | 5 | 0 | 4 | 2 | Carbamidomethylation | 123984 \| PPTG_01429 \| Phytophthora parasitica INRA-310 integrator complex subunit 3 (1136 aa) |
| 2753 | 3111 | PPTG_16843 | 65,42 | 7 | 4 | 4 | 7 | 2,62E+04 | 1,11E+05 | 1,28E+06 | 2 | 2 | 1 | 1 | 2 | | 35520 \| PPTG_16843 \| Phytophthora parasitica INRA-310 hypothetical protein (310 aa) |
| 2940 | 3417 | PPTG_00615 | 65,4 | 27 | 0 | 27 | 27 | | 7,93E+05 | 1,27E+06 | 2 | 2 | 0 | 2 | 2 | | 12240 \| PPTG_00615 \| Phytophthora parasitica INRA-310 hypothetical protein (117 aa) |
| 2570 | 2477 | PPTG_10223 | 65,39 | 6 | 0 | 3 | 5 | | 1,52E+05 | 7,34E+05 | 4 | 4 | 0 | 2 | 3 | | 92201 \| PPTG_10223 \| Phytophthora parasitica INRA-310 beta-galactosidase (820 aa) |
| 2992 | 2491 | PPTG_01816 | 65,34 | 4 | 0 | 2 | 2 | | 1,71E+06 | 3,81E+05 | 3 | 3 | 0 | 2 | 1 | | 117023 \| PPTG_01816 \| Phytophthora parasitica INRA-310 unc-45 family protein (1074 aa) |
| 2098 | 2149 | PPTG_07999 | 65,31 | 21 | 7 | 3 | 21 | 6,75E+05 | 2,93E+05 | 8,82E+06 | 5 | 5 | 2 | 1 | 5 | Oxidation (M) | 27448 \| PPTG_07999 \| Phytophthora parasitica INRA-310 hypothetical protein (244 aa) |
| 2143 | 2222 | PPTG_08330 | 65,3 | 7 | 0 | 5 | 7 | | 7,37E+05 | 7,56E+05 | 4 | 4 | 0 | 3 | 4 | Carbamidomethylation | 78777 \| PPTG_08330 \| Phytophthora parasitica INRA-310 tRNA (cytosine-5-)-methyltransferase NSUN2 (702 aa) |
| 2682 | 2211 | PPTG_12737 | 65,3 | 13 | 0 | 9 | 8 | | 1,01E+05 | 4,94E+05 | 3 | 3 | 0 | 2 | 2 | | 37651 \| PPTG_12737 \| Phytophthora parasitica INRA-310 epoxide hydrolase (337 aa) |
| 2967 | 2892 | PPTG_14759 | 65,18 | 4 | 0 | 3 | 4 | | 4,21E+05 | 1,41E+06 | 2 | 2 | 0 | 1 | 2 | | 92777 \| PPTG_14759 \| Phytophthora parasitica INRA-310 hypothetical protein (826 aa) |
| 2421 | 3231 | PPTG_08420 | 65,18 | 23 | 13 | 23 | 23 | 3,74E+05 | 1,16E+06 | 3,16E+06 | 2 | 2 | 1 | 2 | 3 | Carbamidomethylation; Acetylation (N-term); Carbamylation | 16574 \| PPTG_08420 \| Phytophthora parasitica INRA-310 PLAC8 family protein (154 aa) |
| 1903 | 2237 | PPTG_01268 | 65,15 | 8 | 2 | 6 | 8 | | | 3,29E+05 | 4 | 1 | 1 | 3 | 5 | Oxidation (M) | 49739 \| PPTG_01268 \| Phytophthora parasitica INRA-310 hypothetical protein (435 aa) |
| 2574 | 2691 | PPTG_13398 | 65,12 | 11 | 0 | 9 | 11 | | 2,14E+05 | 2,71E+05 | 3 | 3 | 0 | 2 | 3 | | 29495 \| PPTG_13398 \| Phytophthora parasitica INRA-310 hypothetical protein (256 aa) |
| 2227 | 2712 | PPTG_06299 | 65,04 | 15 | 5 | 11 | 9 | | 1,16E+05 | | 3 | 1 | 2 | 3 | 2 | Oxidation (M) | 23512 \| PPTG_06299 \| Phytophthora parasitica INRA-310 mannitol dehydrogenase (222 aa) |
| 3243 | 2459 | PPTG_13218 | 65,03 | 15 | 0 | 0 | 15 | | | 1,60E+06 | 3 | 3 | 0 | 0 | 3 | | 28432 \| PPTG_13218 \| Phytophthora parasitica INRA-310 hypothetical protein (266 aa) |
| 4409 | 4217 | PPTG_20350 | 65 | 24 | 0 | 24 | 0 | | 1,29E+06 | | 1 | 1 | 0 | 1 | 0 | Carbamidomethylation | 16582 \| PPTG_20350 \| Phytophthora parasitica INRA-310 hypothetical protein (153 aa) |
| 2166 | 2287 | PPTG_05491 | 64,97 | 17 | 3 | 17 | 9 | 1,54E+05 | 1,48E+06 | 1,76E+06 | 4 | 4 | 1 | 4 | 2 | Carbamidomethylation | 35406 \| PPTG_05491 \| Phytophthora parasitica INRA-310 U2 small nuclear ribonucleoprotein A' (318 aa) |
| 2666 | 2522 | PPTG_12501 | 64,95 | 5 | 0 | 2 | 5 | | 3,33E+05 | 9,30E+05 | 3 | 3 | 0 | 1 | 3 | | 66561 \| PPTG_12501 \| Phytophthora parasitica INRA-310 ribosomal RNA processing protein 1 (584 aa) |
| 2228 | 2740 | PPTG_07284 | 64,91 | 57 | 16 | 57 | 47 | 2,87E+05 | 2,59E+06 | 2,38E+06 | 3 | 3 | 1 | 4 | 2 | Oxidation (M); Carbamidomethylation | 8619 \| PPTG_07284 \| Phytophthora parasitica INRA-310 small nuclear ribonucleoprotein G (77 aa) |
| 2512 | 2806 | PPTG_13112 | 64,84 | 7 | 0 | 4 | 5 | | 7,52E+05 | 4,67E+05 | 3 | 3 | 0 | 3 | 2 | | 69591 \| PPTG_13112 \| Phytophthora parasitica INRA-310 hypothetical protein (641 aa) |
| 2641 | 2483 | PPTG_05188 | 64,78 | 10 | 0 | 10 | 7 | | 7,65E+05 | 1,64E+05 | 3 | 3 | 0 | 3 | 2 | | 43798 \| PPTG_05188 \| Phytophthora parasitica INRA-310 actin-like protein (393 aa) |
| 2410 | 2447 | PPTG_18784 | 64,75 | 10 | 3 | 6 | 10 | 1,27E+05 | 2,06E+05 | 1,70E+06 | 3 | 3 | 1 | 2 | 3 | | 36982 \| PPTG_18784 \| Phytophthora parasitica INRA-310 alternative oxidase (326 aa) |
| 2519 | 2440 | PPTG_01999 | 64,71 | 7 | 0 | 2 | 7 | | 2,40E+05 | 7,27E+05 | 4 | 4 | 0 | 1 | 4 | | 59012 \| PPTG_01999 \| Phytophthora parasitica INRA-310 glucosidase (524 aa) |
| 2205 | 2317 | PPTG_12477 | 64,69 | 8 | 2 | 5 | 8 | 9,89E+04 | 4,39E+05 | 1,85E+06 | 3 | 3 | 1 | 2 | 4 | Oxidation (M) | 46118 \| PPTG_12477 \| Phytophthora parasitica INRA-310 cysteine protease family C54 (413 aa) |
| 2160 | 2108 | PPTG_06108 | 64,69 | 8 | 0 | 5 | 6 | | 5,57E+05 | 2,17E+06 | 5 | 5 | 0 | 3 | 4 | | 94524 \| PPTG_06108 \| Phytophthora parasitica INRA-310 hypothetical protein (882 aa) |
| 2144 | 2225 | PPTG_01393 | 64,66 | 2 | 0 | 1 | 2 | | 7,74E+05 | 1,23E+06 | 4 | 4 | 0 | 3 | 4 | | 242041 \| PPTG_01393 \| Phytophthora parasitica INRA-310 hypothetical protein (2196 aa) |
| 1946 | 2506 | PPTG_18420 | 64,65 | 27 | 6 | 27 | 27 | 4,85E+05 | 6,63E+06 | 1,38E+07 | 3 | 3 | 1 | 4 | 4 | Oxidation (M); Deamidation (NQ) | 14678 \| PPTG_18420 \| Phytophthora parasitica INRA-310 C2 domain-containing protein (132 aa) |
| 2675 | 3135 | PPTG_00720 | 64,59 | 5 | 0 | 5 | 5 | | 2,96E+05 | 6,49E+05 | 2 | 2 | 0 | 2 | 2 | | 55410 \| PPTG_00720 \| Phytophthora parasitica INRA-310 NADPH:adrenodoxin oxidoreductase (506 aa) |
| 2162 | 2172 | PPTG_14250 | 64,58 | 7 | 5 | 5 | 7 | 4,50E+05 | 3,42E+05 | 4,66E+06 | 3 | 3 | 2 | 2 | 3 | | 51217 \| PPTG_14250 \| Phytophthora parasitica INRA-310 hypothetical protein (462 aa) |
| 2031 | 2315 | PPTG_11234 | 64,57 | 4 | 1 | 3 | 4 | 8,22E+04 | 4,62E+05 | 2,54E+06 | 4 | 4 | 1 | 3 | 4 | Oxidation (M); Carbamidomethylation | 123472 \| PPTG_11234 \| Phytophthora parasitica INRA-310 hypothetical protein (1103 aa) |
| 2398 | 3102 | PPTG_03860 | 64,54 | 12 | 0 | 12 | 12 | | 2,62E+06 | 1,31E+06 | 2 | 2 | 0 | 3 | 3 | Carbamidomethylation; Deamidation (NQ) | 35305 \| PPTG_03860 \| Phytophthora parasitica INRA-310 hypothetical protein (328 aa) |
| 2481 | 2231 | PPTG_18491 | 64,53 | 17 | 0 | 5 | 17 | | 3,11E+05 | 2,22E+06 | 4 | 4 | 0 | 1 | 4 | | 28134 \| PPTG_18491 \| Phytophthora parasitica INRA-310 hypothetical protein (253 aa) |
| 2455 | 2359 | PPTG_09921 | 64,46 | 7 | 0 | 5 | 3 | | 1,31E+06 | 7,92E+05 | 4 | 4 | 0 | 3 | 2 | | 71140 \| PPTG_09921 \| Phytophthora parasitica INRA-310 importin subunit alpha (650 aa) |
| 2494 | 2296 | PPTG_19456 | 64,46 | 10 | 0 | 10 | 2 | | 1,04E+06 | | 4 | 2 | 0 | 4 | 1 | Oxidation (M); Carbamidomethylation | 55857 \| PPTG_19456 \| Phytophthora parasitica INRA-310 alkaline phosphatase (511 aa) |
| 2555 | 2872 | PPTG_05691 | 64,45 | 28 | 0 | 28 | 21 | | 3,37E+06 | 2,39E+06 | 3 | 3 | 0 | 3 | 2 | | 12281 \| PPTG_05691 \| Phytophthora parasitica INRA-310 hypothetical protein (117 aa) |
| 2711 | 2416 | PPTG_01757 | 64,44 | 4 | 1 | 0 | 4 | 7,38E+04 | | 8,22E+05 | 3 | 2 | 1 | 0 | 3 | | 82636 \| PPTG_01757 \| Phytophthora parasitica INRA-310 transmembrane protein (751 aa) |
| 2565 | 2202 | PPTG_10320 | 64,43 | 20 | 0 | 5 | 20 | | 2,03E+05 | 3,13E+06 | 4 | 4 | 0 | 1 | 4 | | 25479 \| PPTG_10320 \| Phytophthora parasitica INRA-310 Ras family GTPase (227 aa) |
| 2003 | 2123 | PPTG_08216 | 64,35 | 5 | 0 | 5 | 3 | | 1,78E+06 | 1,67E+06 | 3 | 3 | 0 | 5 | 3 | Deamidation (NQ) | 102454 \| PPTG_08216 \| Phytophthora parasitica INRA-310 oxoglutarate dehydrogenase (succinyl-transferring) E1 component (923 aa) |
| 3025 | 2547 | PPTG_18035 | 64,32 | 6 | 0 | 0 | 6 | | | 5,20E+05 | 3 | 3 | 0 | 0 | 3 | Carbamidomethylation | 64258 \| PPTG_18035 \| Phytophthora parasitica INRA-310 aspartokinase (597 aa) |
| 2766 | 2486 | PPTG_15029 | 64,29 | 8 | 0 | 8 | 0 | | 8,52E+05 | | 4 | 4 | 0 | 4 | 0 | | 85214 \| PPTG_15029 \| Phytophthora parasitica INRA-310 hypothetical protein (765 aa) |
| 2794 | 2328 | PPTG_08223 | 64,28 | 7 | 0 | 4 | 4 | | 8,78E+05 | 1,23E+06 | 3 | 3 | 0 | 2 | 2 | Carbamidomethylation | 67994 \| PPTG_08223 \| Phytophthora parasitica INRA-310 formate-tetrahydrofolate ligase (639 aa) |
| 2895 | 2545 | PPTG_04008 | 64,27 | 4 | 2 | 0 | 4 | 5,40E+04 | | 8,23E+05 | 3 | 3 | 1 | 0 | 3 | | 82379 \| PPTG_04008 \| Phytophthora parasitica INRA-310 anoctamin-like protein (707 aa) |
| 2405 | 2196 | PPTG_06079 | 64,21 | 15 | 0 | 13 | 9 | | 5,93E+05 | 1,43E+06 | 4 | 4 | 0 | 3 | 3 | Carbamidomethylation | 52310 \| PPTG_06079 \| Phytophthora parasitica INRA-310 hypothetical protein (474 aa) |
| 3190 | 3241 | PPTG_20064 | 64,17 | 10 | 0 | 10 | 4 | | 4,41E+05 | | 2 | 1 | 0 | 2 | 1 | Carbamidomethylation | 32324 \| PPTG_20064 \| Phytophthora parasitica INRA-310 hypothetical protein (301 aa) |
| 2043 | 2233 | PPTG_11281 | 64,03 | 17 | 0 | 17 | 10 | | 7,82E+05 | 6,50E+05 | 5 | 5 | 0 | 5 | 3 | | 39976 \| PPTG_11281 \| Phytophthora parasitica INRA-310 hypothetical protein (364 aa) |
| 2636 | 3474 | PPTG_04172 | 64,02 | 6 | 4 | 6 | 6 | 1,23E+05 | 7,47E+05 | 1,83E+06 | 2 | 2 | 1 | 2 | 2 | | 48643 \| PPTG_04172 \| Phytophthora parasitica INRA-310 threonine dehydratase (451 aa) |
| 2386 | 2227 | PPTG_16688 | 63,93 | 10 | 1 | 1 | 10 | 5,41E+05 | 4,91E+05 | 4,52E+06 | 4 | 4 | 1 | 1 | 4 | | 59945 \| PPTG_16688 \| Phytophthora parasitica INRA-310 solute carrier family facilitated glucose transporter (552 aa) |
| 2769 | 2542 | PPTG_02790 | 63,84 | 6 | 0 | 1 | 6 | | 9,73E+04 | 3,11E+06 | 3 | 3 | 0 | 1 | 3 | Carbamidomethylation | 61989 \| PPTG_02790 \| Phytophthora parasitica INRA-310 hypothetical protein (574 aa) |
| 2298 | 3232 | PPTG_09179 | 63,84 | 11 | 4 | 11 | 4 | 3,67E+05 | 3,26E+06 | 5,01E+06 | 2 | 2 | 1 | 3 | 2 | Oxidation (M) | 42812 \| PPTG_09179 \| Phytophthora parasitica INRA-310 hypothetical protein (388 aa) |
| 2099 | 2226 | PPTG_18718 | 63,82 | 31 | 0 | 27 | 25 | | 1,40E+06 | 1,70E+06 | 5 | 5 | 0 | 4 | 4 | Carbamidomethylation | 28517 \| PPTG_18718 \| Phytophthora parasitica INRA-310 hydroxyacylglutathione hydrolase (260 aa) |
| 2151 | 2264 | PPTG_03093 | 63,79 | 5 | 1 | 2 | 4 | 1,26E+05 | 5,51E+05 | 1,55E+06 | 5 | 5 | 1 | 2 | 4 | | 115152 \| PPTG_03093 \| Phytophthora parasitica INRA-310 hypothetical protein (1048 aa) |
| 3090 | 2437 | PPTG_11827 | 63,79 | 5 | 0 | 0 | 5 | | | 3,56E+05 | 3 | 3 | 0 | 0 | 3 | Carbamidomethylation | 72281 \| PPTG_11827 \| Phytophthora parasitica INRA-310 hypothetical protein (644 aa) |
| 2230 | 3311 | PPTG_17561 | 63,76 | 26 | 0 | 26 | 18 | | 1,74E+06 | 3,95E+05 | 2 | 2 | 0 | 5 | 2 | Deamidation (NQ) | 17241 \| PPTG_17561 \| Phytophthora parasitica INRA-310 glycine cleavage system H protein (161 aa) |
| 2882 | 3370 | PPTG_13013 | 63,74 | 5 | 0 | 5 | 5 | | 1,63E+06 | 1,07E+06 | 2 | 2 | 0 | 2 | 2 | Carbamidomethylation | 52092 \| PPTG_13013 \| Phytophthora parasitica INRA-310 hypothetical protein (507 aa) |
| 2855 | 2673 | PPTG_00335 | 63,73 | 23 | 0 | 23 | 6 | | 3,16E+06 0 | | 3 | 3 | 0 | 3 | 1 | Carbamidomethylation | 19409 \| PPTG_00335 \| Phytophthora parasitica INRA-310 hypothetical protein (171 aa) |
| 2817 | 3145 | PPTG_13562 | 63,71 | 9 | 4 | 4 | 9 | 8,63E+04 | 1,90E+05 | 8,50E+05 | 2 | 2 | 1 | 1 | 2 | | 44494 \| PPTG_13562 \| Phytophthora parasitica INRA-310 hypothetical protein (413 aa) |
| 2608 | 3033 | PPTG_05431 | 63,7 | 5 | 3 | 5 | 5 | 5,63E+04 | 6,04E+05 | 1,19E+06 | 2 | 2 | 1 | 2 | 2 | | 48586 \| PPTG_05431 \| Phytophthora parasitica INRA-310 hypothetical protein (453 aa) |
| 2771 | 2631 | PPTG_17151 | 63,7 | 5 | 0 | 5 | 3 | | 7,33E+05 | 8,57E+04 | 3 | 3 | 0 | 3 | 1 | | 66968 \| PPTG_17151 \| Phytophthora parasitica INRA-310 mitochondrial import receptor subunit TOM70 (598 aa) |
| 2259 | 3487 | PPTG_17798 | 63,68 | 12 | 12 | 12 | 7 | 5,31E+05 | 5,78E+06 | 4,91E+06 | 2 | 1 | 2 | 3 | 2 | | 17025 \| PPTG_17798 \| Phytophthora parasitica INRA-310 hypothetical protein (166 aa) |
| 3552 | 3222 | PPTG_10032 | 63,66 | 7 | 0 | 7 | 0 | | 2,66E+05 | | 2 | 2 | 0 | 2 | 0 | | 43312 \| PPTG_10032 \| Phytophthora parasitica INRA-310 hypothetical protein (369 aa) |
| 2896 | 2573 | PPTG_11461 | 63,65 | 21 | 0 | 7 | 21 | | 1,26E+06 | 2,98E+06 | 3 | 3 | 0 | 1 | 3 | | 25428 \| PPTG_11461 \| Phytophthora parasitica INRA-310 hypothetical protein (230 aa) |
| 2430 | 2619 | PPTG_08642 | 63,63 | 22 | 6 | 9 | 22 | 8,84E+04 | 3,23E+05 | 4,36E+06 | 3 | 3 | 1 | 1 | 4 | | 15488 \| PPTG_08642 \| Phytophthora parasitica INRA-310 hypothetical protein (135 aa) |
| 3763 | 3377 | PPTG_19530 | 63,56 | 19 | 0 | 0 | 19 | | | 3,43E+05 | 2 | 2 | 0 | 0 | 2 | | 26056 \| PPTG_19530 \| Phytophthora parasitica INRA-310 transmembrane protein (247 aa) |
| 2029 | 2229 | PPTG_03651 | 63,55 | 12 | 2 | 7 | 6 | 5,38E+04 | 5,46E+05 | 1,36E+06 | 6 | 6 | 1 | 3 | 4 | Oxidation (M); Carbamidomethylation | 60758 \| PPTG_03651 \| Phytophthora parasitica INRA-310 6-phosphofructo-2-kinase/fructose-2 6-biphosphatase (543 aa) |
| 2006 | 2531 | PPTG_06540 | 63,51 | 4 | 1 | 4 | 3 | 9,61E+04 | 9,67E+05 | 1,58E+06 | 4 | 4 | 1 | 4 | 3 | | 100217 \| PPTG_06540 \| Phytophthora parasitica INRA-310 U3 small nucleolar RNA-associated protein (915 aa) |
| 2485 | 2570 | PPTG_01280 | 63,5 | 4 | 0 | 2 | 4 | | 3,88E+05 | 1,89E+05 | 3 | 3 | 0 | 2 | 3 | | 114662 \| PPTG_01280 \| Phytophthora parasitica INRA-310 hypothetical protein (1018 aa) |
| 2744 | 2661 | PPTG_15161 | 63,47 | 4 | 0 | 0 | 4 | | | 1,97E+06 | 3 | 3 | 0 | 0 | 4 | Carbamidomethylation | 98285 \| PPTG_15161 \| Phytophthora parasitica INRA-310 meiosis-specific nuclear structural protein 1 (844 aa) |
| 2846 | 2468 | PPTG_10968 | 63,45 | 15 | 0 | 7 | 15 | | 2,98E+04 | 6,66E+05 | 3 | 3 | 0 | 1 | 3 | Carbamidomethylation | 40153 \| PPTG_10968 \| Phytophthora parasitica INRA-310 hypothetical protein (364 aa) |
| 2728 | 3041 | PPTG_17157 | 63,33 | 6 | 0 | 6 | 3 | | 1,65E+06 | 3,59E+05 | 2 | 2 | 0 | 3 | 1 | Oxidation (M) | 42966 \| PPTG_17157 \| Phytophthora parasitica INRA-310 arginine N-methyltransferase (378 aa) |
| 2293 | 2200 | PPTG_14459 | 63,33 | 5 | 0 | 3 | 2 | | 8,61E+05 | 9,59E+05 | 5 | 5 | 0 | 3 | 3 | | 143767 \| PPTG_14459 \| Phytophthora parasitica INRA-310 hypothetical protein (1336 aa) |
| 3400 | 3152 | PPTG_10093 | 63,23 | 4 | 0 | 0 | 4 | | | 5,25E+05 | 2 | 2 | 0 | 0 | 2 | Oxidation (M) | 88352 \| PPTG_10093 \| Phytophthora parasitica INRA-310 anoctamin-like protein (775 aa) |
| 2131 | 2301 | PPTG_06726 | 63,15 | 10 | 1 | 5 | 5 | 3,42E+05 | 1,58E+06 | 1,28E+06 | 6 | 5 | 1 | 3 | 3 | | 82710 \| PPTG_06726 \| Phytophthora parasitica INRA-310 phosphate acetyltransferase (761 aa) |
| 2544 | 2544 | PPTG_11018 | 63,14 | 5 | 0 | 3 | 5 | | 5,02E+05 | 1,09E+06 | 3 | 3 | 0 | 2 | 3 | Oxidation (M) | 78114 \| PPTG_11018 \| Phytophthora parasitica INRA-310 acyl-CoA dehydrogenase/oxidase (703 aa) |
| 2537 | 2240 | PPTG_03691 | 63,13 | 6 | 0 | 5 | 2 | | 6,36E+05 | 1,10E+06 | 4 | 4 | 0 | 3 | 2 | | 108187 \| PPTG_03691 \| Phytophthora parasitica INRA-310 hypothetical protein (977 aa) |
| 3131 | 2355 | PPTG_05150 | 63,11 | 7 | 0 | 0 | 7 | | | 1,91E+06 | 3 | 3 | 0 | 0 | 3 | | 65811 \| PPTG_05150 \| Phytophthora parasitica INRA-310 hypothetical protein (582 aa) |

| | | | | | | | | | | | | | | | | | | | |
|---|---|---|---|---|---|---|---|---|---|---|---|---|---|---|---|---|---|---|---|
| 2440 | 2502 | PPTG_04804 | 63,09 | 16 | 0 | 6 | 16 | | 4,86E+05 | 1,01E+06 | 3 | 3 | 0 | 2 | 4 | Carbamidomethylation | 31893 | PPTG_04804 | Phytophthora parasitica INRA-310 bifunctional folD protein (299 aa) |
| 3404 | 3389 | PPTG_18528 | 63,08 | 3 | 0 | 0 | 3 | | | 4,84E+05 | 2 | 2 | 0 | 0 | 2 | Carbamidomethylation | 113356 | PPTG_18528 | Phytophthora parasitica INRA-310 phosphatidylinositol-4-phosphate-5-kinase (PIPK-D1/GPCR-PIPK) (1015 aa) |
| 2432 | 2827 | PPTG_16502 | 63,08 | 8 | 3 | 5 | 8 | 1,00E+05 | 7,26E+05 | 2,37E+06 | 3 | 3 | 1 | 2 | 3 | | 40345 | PPTG_16502 | Phytophthora parasitica INRA-310 palmitoyl-protein thioesterase 1 (356 aa) |
| 2413 | 2601 | PPTG_13255 | 63 | 17 | 11 | 6 | 17 | 4,30E+05 | 4,35E+05 | 5,40E+06 | 3 | 3 | 2 | 1 | 3 | | 20243 | PPTG_13255 | Phytophthora parasitica INRA-310 hypothetical protein (179 aa) |
| 2578 | 2777 | PPTG_01973 | 62,99 | 28 | 9 | 18 | 19 | 2,52E+05 | 2,14E+06 | 1,71E+06 | 3 | 3 | 1 | 2 | 2 | | 16587 | PPTG_01973 | Phytophthora parasitica INRA-310 prefoldin alpha subunit (149 aa) |
| 2533 | 3229 | PPTG_18863 | 62,95 | 3 | 2 | 3 | 3 | 5,74E+04 | 6,53E+05 | 9,95E+05 | 2 | 2 | 1 | 2 | 2 | | 86033 | PPTG_18863 | Phytophthora parasitica INRA-310 hypothetical protein (796 aa) |
| 3647 | 3380 | PPTG_07161 | 62,92 | 14 | 0 | 14 | 0 | | 7,38E+05 | | 2 | 2 | 0 | 2 | 0 | | 21378 | PPTG_07161 | Phytophthora parasitica INRA-310 hypothetical protein (192 aa) |
| 2304 | 2280 | PPTG_14479 | 62,91 | 9 | 0 | 6 | 4 | | 1,92E+05 | 8,33E+05 | 5 | 5 | 0 | 3 | 3 | Carbamidomethylation | 75624 | PPTG_14479 | Phytophthora parasitica INRA-310 indigoidine synthase A family protein (711 aa) |
| 2292 | 2162 | PPTG_18896 | 62,9 | 6 | 1 | 5 | 3 | 4,45E+04 | 6,75E+05 | 5,52E+05 | 4 | 4 | 1 | 3 | 2 | Carbamidomethylation | 92155 | PPTG_18896 | Phytophthora parasitica INRA-310 TFIIH basal transcription factor complex helicase XPB subunit (821 aa) |
| 2209 | 2464 | PPTG_07086 | 62,82 | 32 | 0 | 32 | 32 | | 8,06E+05 | 8,33E+05 | 3 | 3 | 0 | 3 | 4 | | 15819 | PPTG_07086 | Phytophthora parasitica INRA-310 hypothetical protein (147 aa) |
| 2577 | 2776 | PPTG_04839 | 62,8 | 18 | 0 | 13 | 11 | | 1,37E+06 | 1,39E+06 | 4 | 4 | 0 | 3 | 2 | | 25352 | PPTG_04839 | Phytophthora parasitica INRA-310 eukaryotic initiation factor 4E (223 aa) |
| 2829 | 3332 | PPTG_00129 | 62,77 | 11 | 6 | 11 | 6 | 1,96E+05 | 9,50E+05 | 1,63E+06 | 2 | 2 | 1 | 2 | 1 | | 25389 | PPTG_00129 | Phytophthora parasitica INRA-310 hypothetical protein (236 aa) |
| 2060 | 2400 | PPTG_09046 | 62,71 | 22 | 0 | 22 | 21 | | 7,23E+06 | 6,24E+06 | 4 | 4 | 0 | 5 | 3 | | 19338 | PPTG_09046 | Phytophthora parasitica INRA-310 hypothetical protein (171 aa) |
| 2872 | 3230 | PPTG_18625 | 62,71 | 9 | 6 | 9 | 4 | 2,90E+05 | 1,75E+06 | 1,51E+06 | 2 | 2 | 1 | 2 | 1 | | 24679 | PPTG_18625 | Phytophthora parasitica INRA-310 eukaryotic initiation factor 4E (217 aa) |
| 2981 | 2700 | PPTG_10493 | 62,67 | 11 | 0 | 0 | 11 | | | 1,14E+06 | 3 | 3 | 0 | 0 | 3 | Carbamidomethylation | 43602 | PPTG_10493 | Phytophthora parasitica INRA-310 hypothetical protein (394 aa) |
| 1963 | 2304 | PPTG_19374 | 62,57 | 34 | 9 | 25 | 34 | 6,17E+05 | 2,87E+06 | 4,69E+06 | 4 | 4 | 2 | 3 | 4 | Deamidation (NQ) | 12558 | PPTG_19374 | Phytophthora parasitica INRA-310 small nuclear ribonucleoprotein Sm D2 (111 aa) |
| 2907 | 3260 | PPTG_18195 | 62,56 | 18 | 0 | 7 | 18 | | 1,15E+06 | 3,75E+07 | 2 | 2 | 0 | 1 | 3 | | 12159 | PPTG_18195 | Phytophthora parasitica INRA-310 60S ribosomal protein L44 (105 aa) |
| 2499 | 2866 | PPTG_10534 | 62,51 | 10 | 4 | 10 | 6 | 2,62E+05 | 3,90E+06 | 1,69E+06 | 2 | 2 | 2 | 2 | 1 | Carbamidomethylation | 40477 | PPTG_10534 | Phytophthora parasitica INRA-310 hypothetical protein (366 aa) |
| 2285 | 2552 | PPTG_01743 | 62,5 | 34 | 0 | 27 | 22 | | 4,59E+06 | 1,40E+07 | 4 | 4 | 0 | 3 | 3 | | 11273 | PPTG_01743 | Phytophthora parasitica INRA-310 small ubiquitin-like modifier (SUMO) (102 aa) |
| 2683 | 2471 | PPTG_19022 | 62,5 | 5 | 0 | 3 | 2 | | 2,46E+06 | 1,39E+06 | 3 | 3 | 0 | 2 | 2 | | 86971 | PPTG_19022 | Phytophthora parasitica INRA-310 serine protease (809 aa) |
| 2542 | 2457 | PPTG_06495 | 62,43 | 4 | 0 | 4 | 3 | | 7,16E+05 | 1,35E+06 | 3 | 3 | 0 | 3 | 2 | | 88592 | PPTG_06495 | Phytophthora parasitica INRA-310 hook domain-containing protein (795 aa) |
| 2074 | 2309 | PPTG_12890 | 62,42 | 15 | 2 | 6 | 12 | | 1,23E+06 | 6,60E+05 | 4 | 3 | 1 | 3 | 4 | | 36740 | PPTG_12890 | Phytophthora parasitica INRA-310 calcium/calmodulin dependent protein kinase (328 aa) |
| 3061 | 2847 | PPTG_16166 | 62,36 | 8 | 0 | 6 | 3 | | 2,93E+05 | 2,57E+05 | 3 | 3 | 0 | 2 | 1 | Carbamidomethylation | 54528 | PPTG_16166 | Phytophthora parasitica INRA-310 hypothetical protein (479 aa) |
| 2492 | 2241 | PPTG_06861 | 62,35 | 6 | 0 | 1 | 6 | | 6,89E+04 | 7,60E+05 | 4 | 4 | 0 | 1 | 4 | | 113893 | PPTG_06861 | Phytophthora parasitica INRA-310 APS kinase/ATP sulfurlyase/pyrophosphatase fusion protein (1017 aa) |
| 2414 | 2695 | PPTG_03080 | 62,3 | 5 | 2 | 5 | 4 | 4,51E+04 | 1,29E+06 | 6,55E+05 | 3 | 3 | 1 | 3 | 2 | | 72496 | PPTG_03080 | Phytophthora parasitica INRA-310 peptidase (662 aa) |
| 3117 | 3399 | PPTG_15751 | 62,26 | 17 | 0 | 0 | 17 | | | 1,08E+06 | 2 | 2 | 0 | 0 | 3 | | 21511 | PPTG_15751 | Phytophthora parasitica INRA-310 mitochondrial protein Translocase (MPT) family (207 aa) |
| 3250 | 3431 | PPTG_01163 | 62,26 | 18 | 0 | 18 | 8 | | 8,82E+05 | 4,63E+05 | 2 | 2 | 0 | 2 | 1 | | 33085 | PPTG_01163 | Phytophthora parasitica INRA-310 hypothetical protein (308 aa) |
| 2245 | 2718 | PPTG_15731 | 62,24 | 27 | 8 | 27 | 27 | 7,76E+05 | 8,56E+06 | 1,03E+07 | 3 | 3 | 1 | 3 | 3 | | 12921 | PPTG_15731 | Phytophthora parasitica INRA-310 small nuclear ribonucleoprotein Sm D3 (117 aa) |
| 2620 | 2568 | PPTG_06554 | 62,18 | 11 | 3 | 7 | 9 | 1,89E+05 | 9,85E+05 | 8,40E+05 | 3 | 3 | 1 | 2 | 2 | | 31240 | PPTG_06554 | Phytophthora parasitica INRA-310 eukaryotic initiation factor 4E (272 aa) |
| 2988 | 3157 | PPTG_11483 | 62,12 | 10 | 0 | 10 | 5 | | 7,84E+05 0 | | 2 | 2 | 0 | 2 | 1 | | 29606 | PPTG_11483 | Phytophthora parasitica INRA-310 sporangia induced inner dynein arm light chain (264 aa) |
| 2674 | 3010 | PPTG_13914 | 62,1 | 10 | 0 | 10 | 10 | | 2,37E+06 | 2,78E+06 | 2 | 2 | 0 | 2 | 2 | | 18706 | PPTG_13914 | Phytophthora parasitica INRA-310 39S ribosomal protein L24 mitochondrial (167 aa) |
| 2776 | 2851 | PPTG_16152 | 62,1 | 11 | 0 | 9 | 9 | | 4,13E+05 | 5,44E+05 | 3 | 3 | 0 | 2 | 2 | | 35857 | PPTG_16152 | Phytophthora parasitica INRA-310 hypothetical protein (323 aa) |
| 3434 | 3160 | PPTG_13123 | 62,01 | 4 | 0 | 0 | 4 | | | 1,27E+06 | 2 | 2 | 0 | 0 | 2 | Carbamidomethylation | 89700 | PPTG_13123 | Phytophthora parasitica INRA-310 hypothetical protein (828 aa) |
| 3149 | 3303 | PPTG_12715 | 61,97 | 11 | 0 | 0 | 11 | | | 7,13E+05 | 2 | 2 | 0 | 0 | 3 | | 33515 | PPTG_12715 | Phytophthora parasitica INRA-310 hypothetical protein (299 aa) |
| 2470 | 2446 | PPTG_05459 | 61,96 | 9 | 0 | 5 | 7 | | 1,33E+05 | 5,04E+05 | 4 | 4 | 0 | 2 | 3 | | 64513 | PPTG_05459 | Phytophthora parasitica INRA-310 striatin-3 family protein (594 aa) |
| 3170 | 3822 | PPTG_00062 | 61,91 | 13 | 13 | 13 | 13 | 1,58E+05 | 6,95E+06 | 3,89E+06 | 1 | 1 | 1 | 1 | 1 | Carbamidomethylation | 18370 | PPTG_00062 | Phytophthora parasitica INRA-310 hypothetical protein (180 aa) |
| 2714 | 2653 | PPTG_17495 | 61,82 | 14 | 0 | 9 | 7 | | 4,45E+05 | 6,45E+05 | 3 | 3 | 0 | 2 | 2 | | 40128 | PPTG_17495 | Phytophthora parasitica INRA-310 2OG-Fe(II) oxygenase superfamily protein (349 aa) |
| 2146 | 2352 | PPTG_11785 | 61,81 | 9 | 0 | 6 | 5 | | 8,33E+05 | 1,53E+06 | 6 | 6 | 0 | 4 | 3 | Oxidation (M); Carbamidomethylation | 86949 | PPTG_11785 | Phytophthora parasitica INRA-310 hypothetical protein (776 aa) |
| 2822 | 3244 | PPTG_17117 | 61,81 | 15 | 9 | 9 | 15 | 7,95E+04 | 2,01E+05 | 1,22E+06 | 2 | 2 | 1 | 1 | 2 | Carbamidomethylation | 19651 | PPTG_17117 | Phytophthora parasitica INRA-310 hypothetical protein (171 aa) |
| 2593 | 2546 | PPTG_18318 | 61,77 | 17 | 0 | 17 | 11 | | 6,60E+05 | 2,30E+05 | 3 | 3 | 0 | 3 | 2 | | 27205 | PPTG_18318 | Phytophthora parasitica INRA-310 N(6)-adenine-specific DNA methyltransferase 2 (235 aa) |
| 2374 | 2720 | PPTG_11433 | 61,68 | 3 | 1 | 2 | 3 | 7,17E+04 | 4,58E+05 | 1,06E+06 | 3 | 3 | 1 | 2 | 3 | | 109191 | PPTG_11433 | Phytophthora parasitica INRA-310 hypothetical protein (976 aa) |
| 2275 | 2508 | PPTG_03586 | 61,62 | 5 | 0 | 2 | 5 | | 1,07E+06 | 3,56E+06 | 4 | 4 | 0 | 2 | 4 | Oxidation (M) | 85505 | PPTG_03586 | Phytophthora parasitica INRA-310 AMP deaminase 2 (742 aa) |
| 2870 | 3218 | PPTG_16435 | 61,61 | 6 | 3 | 3 | 6 | 1,48E+05 | 2,44E+05 | 1,30E+06 | 2 | 2 | 1 | 1 | 2 | | 43387 | PPTG_16435 | Phytophthora parasitica INRA-310 hypothetical protein (393 aa) |
| 2785 | 3209 | PPTG_15035 | 61,54 | 2 | 0 | 2 | 2 | | 3,78E+05 | 5,01E+05 | 2 | 2 | 0 | 2 | 2 | | 154748 | PPTG_15035 | Phytophthora parasitica INRA-310 2' 5'-phosphodiesterase (1364 aa) |
| 2755 | 3202 | PPTG_11997 | 61,53 | 6 | 0 | 6 | 6 | | 1,30E+07 | 1,21E+07 | 2 | 2 | 0 | 2 | 2 | | 36844 | PPTG_11997 | Phytophthora parasitica INRA-310 hypothetical protein (340 aa) |
| 2871 | 3224 | PPTG_20158 | 61,51 | 11 | 6 | 6 | 11 | 1,02E+05 | 5,87E+05 | 9,71E+05 | 2 | 2 | 1 | 1 | 2 | | 25797 | PPTG_20158 | Phytophthora parasitica INRA-310 hypothetical protein (218 aa) |
| 2848 | 2510 | PPTG_10391 | 61,5 | 7 | 0 | 0 | 7 | | | 2,14E+06 | 4 | 4 | 0 | 0 | 4 | | 64724 | PPTG_10391 | Phytophthora parasitica INRA-310 hypothetical protein (574 aa) |
| 2379 | 2888 | PPTG_08497 | 61,43 | 13 | 3 | 8 | 8 | | 2,35E+05 | 4,72E+05 | 3 | 2 | 1 | 3 | 2 | | 33651 | PPTG_08497 | Phytophthora parasitica INRA-310 cyclin-dependent kinase (305 aa) |
| 2422 | 3268 | PPTG_10775 | 61,4 | 7 | 7 | 7 | 7 | 4,20E+05 | 2,86E+06 | 4,21E+06 | 2 | 2 | 2 | 2 | 2 | Deamidation (NQ) | 36178 | PPTG_10775 | Phytophthora parasitica INRA-310 aspartyl protease family A01B (333 aa) |
| 2368 | 2478 | PPTG_10695 | 61,39 | 4 | 1 | 2 | 2 | 6,03E+04 | 3,72E+05 | 1,17E+06 | 4 | 4 | 1 | 2 | 3 | | 137056 | PPTG_10695 | Phytophthora parasitica INRA-310 hypothetical protein (1268 aa) |
| 2059 | 2368 | PPTG_08962 | 61,38 | 22 | 7 | 22 | 22 | 3,60E+05 | 4,39E+06 | 9,25E+06 | 3 | 3 | 1 | 3 | 4 | Carbamidomethylation | 16117 | PPTG_08962 | Phytophthora parasitica INRA-310 ribosomal protein L7Ae/L30e/S12e/Gadd4 (148 aa) |
| 2366 | 2167 | PPTG_18846 | 61,35 | 5 | 0 | 4 | 3 | | 7,81E+05 | 1,47E+06 | 4 | 4 | 0 | 3 | 3 | | 100923 | PPTG_18846 | Phytophthora parasitica INRA-310 periodic tryptophan protein 2 (913 aa) |
| 3764 | 3379 | PPTG_13420 | 61,3 | 5 | 0 | 0 | 5 | | | 3,95E+05 | 2 | 2 | 0 | 0 | 2 | | 57037 | PPTG_13420 | Phytophthora parasitica INRA-310 hypothetical protein (522 aa) |
| 3391 | 2897 | PPTG_20489 | 61,3 | 18 | 0 | 18 | 0 | | 1,78E+07 | | 2 | 2 | 0 | 2 | 0 | Oxidation (M) | 23601 | PPTG_20489 | Phytophthora parasitica INRA-310 hypothetical protein (222 aa) |
| 2530 | 2791 | PPTG_18328 | 61,29 | 3 | 0 | 2 | 3 | | 4,01E+05 | 3,43E+06 | 3 | 3 | 0 | 2 | 3 | | 119224 | PPTG_18328 | Phytophthora parasitica INRA-310 hypothetical protein (1046 aa) |
| 2373 | 2627 | PPTG_13887 | 61,26 | 6 | 2 | 4 | 6 | 1,99E+05 | 1,15E+06 | 1,37E+06 | 3 | 3 | 1 | 2 | 3 | | 61907 | PPTG_13887 | Phytophthora parasitica INRA-310 hypothetical protein (557 aa) |
| 2579 | 2784 | PPTG_09704 | 61,16 | 9 | 0 | 9 | 5 | | 6,85E+05 | 8,79E+05 | 3 | 3 | 0 | 3 | 2 | Carbamidomethylation | 47665 | PPTG_09704 | Phytophthora parasitica INRA-310 homoserine O-acetyltransferase (426 aa) |
| 2371 | 2579 | PPTG_05034 | 61,12 | 2 | 0 | 2 | 1 | | 9,39E+04 | 1,35E+05 | 3 | 2 | 0 | 3 | 3 | Oxidation (M) | 161694 | PPTG_05034 | Phytophthora parasitica INRA-310 chromatin-remodeling complex ATPase chain (1395 aa) |
| 2321 | 2462 | PPTG_09811 | 61,07 | 10 | 0 | 5 | 10 | | 1,35E+05 | 1,46E+06 | 4 | 4 | 0 | 2 | 4 | | 46189 | PPTG_09811 | Phytophthora parasitica INRA-310 transmembrane protein (411 aa) |
| 1999 | 2184 | PPTG_12169 | 61,05 | 8 | 2 | 3 | 8 | 3,00E+05 | 7,20E+05 | 3,15E+06 | 5 | 5 | 1 | 2 | 5 | Carbamidomethylation | 59045 | PPTG_12169 | Phytophthora parasitica INRA-310 protein kinase (532 aa) |
| 2128 | 2299 | PPTG_15645 | 61,05 | 8 | 2 | 6 | 6 | 7,04E+04 | 2,83E+05 | 2,46E+06 | 4 | 4 | 1 | 3 | 3 | Carbamidomethylation | 60641 | PPTG_15645 | Phytophthora parasitica INRA-310 hypothetical protein (544 aa) |
| 2495 | 2565 | PPTG_19135 | 61,02 | 5 | 0 | 2 | 4 | | 2,38E+05 | 4,03E+05 | 4 | 4 | 0 | 2 | 3 | | 106918 | PPTG_19135 | Phytophthora parasitica INRA-310 pre-mRNA-processing factor (962 aa) |
| 2224 | 2387 | PPTG_05267 | 60,88 | 3 | 1 | 3 | 3 | 1,34E+05 | 1,02E+06 | 1,25E+06 | 3 | 3 | 1 | 3 | 3 | | 116689 | PPTG_05267 | Phytophthora parasitica INRA-310 hypothetical protein (1056 aa) |
| 2899 | 2685 | PPTG_17094 | 60,83 | 6 | 0 | 5 | 3 | | 1,17E+06 | 1,81E+06 | 3 | 3 | 0 | 2 | 2 | | 76092 | PPTG_17094 | Phytophthora parasitica INRA-310 hypothetical protein (676 aa) |
| 3144 | 3137 | PPTG_13978 | 60,81 | 23 | 0 | 16 | 23 | | 1,07E+05 | 6,54E+05 | 2 | 2 | 0 | 1 | 2 | Carbamidomethylation | 12253 | PPTG_13978 | Phytophthora parasitica INRA-310 protein enhancer-rudimentary (107 aa) |
| 2765 | 2458 | PPTG_07428 | 60,75 | 9 | 0 | 3 | 9 | | 1,74E+05 | 1,18E+06 | 3 | 3 | 0 | 1 | 3 | Carbamidomethylation | 48695 | PPTG_07428 | Phytophthora parasitica INRA-310 hypothetical protein (425 aa) |
| 3254 | 3441 | PPTG_20469 | 60,69 | 11 | 0 | 8 | 11 | | 2,29E+06 | 5,93E+06 | 2 | 2 | 0 | 1 | 2 | | 40475 | PPTG_20469 | Phytophthora parasitica INRA-310 hypothetical protein (462 aa) |
| 2370 | 2537 | PPTG_07036 | 60,65 | 7 | 2 | 4 | 7 0 | | 1,13E+06 | 2,88E+06 | 3 | 3 | 1 | 2 | 3 | | 52078 | PPTG_07036 | Phytophthora parasitica INRA-310 hypothetical protein (468 aa) |
| 3036 | 3040 | PPTG_00153 | 60,62 | 9 | 0 | 6 | 9 | 0 | | 4,00E+05 | 2 | 2 | 0 | 1 | 2 | | 36609 | PPTG_00153 | Phytophthora parasitica INRA-310 hypothetical protein (343 aa) |
| 2396 | 2848 | PPTG_08784 | 60,6 | 43 | 9 | 14 | 34 | 1,32E+05 | 2,00E+06 | 6,58E+06 | 3 | 3 | 1 | 2 | 3 | Oxidation (M) | 11209 | PPTG_08784 | Phytophthora parasitica INRA-310 hypothetical protein (100 aa) |
| 2725 | 2969 | PPTG_13470 | 60,57 | 7 | 0 | 7 | 7 | | 4,68E+05 | 5,43E+05 | 2 | 2 | 0 | 2 | 2 | Carbamidomethylation | 42953 | PPTG_13470 | Phytophthora parasitica INRA-310 flap endonuclease 1 (391 aa) |
| 3093 | 2732 | PPTG_01004 | 60,55 | 13 | 0 | 0 | 13 | | | 7,44E+05 | 3 | 3 | 0 | 0 | 3 | Carbamidomethylation | 42582 | PPTG_01004 | Phytophthora parasitica INRA-310 hypothetical protein (384 aa) |
| 3093 | 2733 | PPTG_01004 | 60,55 | 12 | 0 | 0 | 12 | | | 7,44E+05 | 3 | 3 | 0 | 0 | 3 | Carbamidomethylation | 44509 | PPTG_01004 | Phytophthora parasitica INRA-310 hypothetical protein (402 aa) |
| 3203 | 3504 | PPTG_19167 | 60,54 | 7 | 0 | 4 | 7 | | 6,63E+05 | 5,60E+05 | 2 | 2 | 0 | 1 | 2 | | 39258 | PPTG_19167 | Phytophthora parasitica INRA-310 arabinogalactan endo-1 4-beta-galactosidase (363 aa) |
| 2030 | 2242 | PPTG_04950 | 60,53 | 7 | 0 | 6 | 5 | | 1,06E+06 | 8,89E+05 | 5 | 5 | 0 | 4 | 4 | Carbamidomethylation | 88554 | PPTG_04950 | Phytophthora parasitica INRA-310 hypothetical protein (804 aa) |
| 3558 | 3393 | PPTG_10839 | 60,45 | 19 | 0 | 0 | 19 | | | 1,69E+06 | 2 | 2 | 0 | 0 | 2 | | 18234 | PPTG_10839 | Phytophthora parasitica INRA-310 hypothetical protein (162 aa) |
| 2941 | 3429 | PPTG_09988 | 60,39 | 7 | 0 | 7 | 2 | | 1,08E+06 | 2,66E+05 | 2 | 2 | 0 | 3 | 1 | | 57190 | PPTG_09988 | Phytophthora parasitica INRA-310 hypothetical protein (538 aa) |
| 1768 | 2811 | PPTG_06157 | 60,33 | 2 | 1 | 1 | 2 | | | 5,05E+04 | 3 | 1 | 2 | 2 | 6 | Oxidation (M) | 151584 | PPTG_06157 | Phytophthora parasitica INRA-310 ATP-binding Cassette (ABC) superfamily (1360 aa) |
| 2810 | 2857 | PPTG_10293 | 60,31 | 7 | 0 | 4 | 7 | | 8,92E+05 | 2,45E+05 | 2 | 2 | 0 | 1 | 3 | | 35360 | PPTG_10293 | Phytophthora parasitica INRA-310 hypothetical protein (335 aa) |
| 2684 | 2900 | PPTG_08163 | 60,3 | 3 | 0 | 3 | 3 | | 4,17E+05 | 3,81E+05 | 2 | 2 | 0 | 2 | 2 | | 108643 | PPTG_08163 | Phytophthora parasitica INRA-310 hypothetical protein (977 aa) |
| 2581 | 2836 | PPTG_03930 | 60,28 | 18 | 0 | 18 | 5 | | 2,07E+06 | 2,97E+05 | 3 | 3 | 0 | 4 | 1 | Oxidation (M); Acetylation (N-term) | 20020 | PPTG_03930 | Phytophthora parasitica INRA-310 peptidyl-prolyl cis-trans isomerase H (183 aa) |
| 2687 | 3124 | PPTG_04345 | 60,2 | 5 | 2 | 2 | 5 | 1,39E+05 | 4,13E+05 | 1,02E+06 | 2 | 2 | 1 | 1 | 2 | | 67029 | PPTG_04345 | Phytophthora parasitica INRA-310 Poly(A)-specific ribonuclease PARN-like protein (591 aa) |
| 2797 | 2424 | PPTG_18842 | 60,19 | 15 | 0 | 0 | 15 | | | 1,06E+06 | 4 | 4 | 0 | 0 | 4 | | 44725 | PPTG_18842 | Phytophthora parasitica INRA-310 hypothetical protein (395 aa) |
| 2077 | 2618 | PPTG_01533 | 60,18 | 19 | 4 | 8 | 19 | 1,22E+05 | 1,02E+06 | 4,54E+06 | 5 | 5 | 1 | 2 | 5 | | 25224 | PPTG_01533 | Phytophthora parasitica INRA-310 transmembrane protein (227 aa) |
| 2713 | 2607 | PPTG_17419 | 60,11 | 5 | 0 | 3 | 3 | | 1,57E+05 | 5,28E+05 | 3 | 2 | 0 | 2 | 2 | | 68628 | PPTG_17419 | Phytophthora parasitica INRA-310 hypothetical protein (614 aa) |
| 2973 | 2654 | PPTG_12348 | 60,09 | 2 | 0 | 0 | 2 | | | 7,32E+05 | 3 | 3 | 0 | 0 | 3 | Carbamidomethylation | 210784 | PPTG_12348 | Phytophthora parasitica INRA-310 myosin-like protein (1871 aa) |
| 2308 | 2621 | PPTG_14758 | 60,08 | 3 | 1 | 1 | 2 | 1,25E+05 | | 1,07E+05 | 3 | 2 | 1 | 2 | 3 | Carbamidomethylation | 127907 | PPTG_14758 | Phytophthora parasitica INRA-310 phosphatidyl inositol kinase (PIK-G1) (1134 aa) |
| 3352 | 3163 | PPTG_07332 | 60,06 | 3 | 0 | 0 | 3 | | | 1,54E+05 | 2 | 2 | 0 | 0 | 2 | | 119902 | PPTG_07332 | Phytophthora parasitica INRA-310 peroxisome biogenesis factor (1104 aa) |
| 2267 | 2742 | PPTG_01014 | 60,06 | 1 | 0 | 1 | 0 | | 3,08E+05 | | 4 | 2 | 1 | 4 | 1 | | 477154 | PPTG_01014 | Phytophthora parasitica INRA-310 dynein heavy chain (4251 aa) |
| 3132 | 2557 | PPTG_05092 | 60,01 | 13 | 0 | 2 | 11 | | 2,37E+05 | 1,05E+06 | 3 | 3 | 0 | 1 | 2 | | 37524 | PPTG_05092 | Phytophthora parasitica INRA-310 hypothetical protein (342 aa) |
| 2590 | 2485 | PPTG_02791 | 59,96 | 8 | 2 | 5 | 5 | 1,10E+05 | 5,71E+05 | 9,85E+05 | 3 | 3 | 1 | 2 | 2 | Carbamidomethylation | 49685 | PPTG_02791 | Phytophthora parasitica INRA-310 hypothetical protein (454 aa) |
| 2665 | 2286 | PPTG_05216 | 59,95 | 4 | 0 | 2 | 2 | | 6,41E+05 | 5,04E+05 | 4 | 4 | 0 | 2 | 2 | | 134715 | PPTG_05216 | Phytophthora parasitica INRA-310 ATP-dependent RNA helicase DHX8 (1186 aa) |
| 3195 | 3403 | PPTG_08424 | 59,95 | 10 | 3 | 3 | 6 | 5,60E+05 | 1,17E+06 | 2,85E+05 | 2 | 2 | 1 | 1 | 1 | | 36992 | PPTG_08424 | Phytophthora parasitica INRA-310 drug/Metabolite transporter (DMT) superfamily (332 aa) |
| 2255 | 2639 | PPTG_19213 | 59,94 | 23 | 7 | 13 | 23 | 4,99E+04 | 4,32E+05 | 1,93E+06 | 4 | 4 | 1 | 2 | 4 | Carbamidomethylation | 16500 | PPTG_19213 | Phytophthora parasitica INRA-310 hypothetical protein (151 aa) |
| 2061 | 2793 | PPTG_08137 | 59,93 | 44 | 15 | 15 | 44 | 2,76E+07 | 5,06E+07 | 1,65E+08 | 3 | 3 | 2 | 2 | 4 | Carbamidomethylation; Acetylation (N-term) | 6581 | PPTG_08137 | Phytophthora parasitica INRA-310 40S ribosomal protein S29 (55 aa) |
| 3055 | 2519 | PPTG_08104 | 59,89 | 3 | 0 | 0 | 3 | | | 1,57E+06 | 3 | 3 | 0 | 0 | 3 | | 112422 | PPTG_08104 | Phytophthora parasitica INRA-310 hypothetical protein (991 aa) |
| 2611 | 3195 | PPTG_01552 | 59,87 | 29 | 0 | 29 | 29 | | 5,61E+05 | 1,70E+06 | 2 | 2 | 0 | 2 | 3 | Carbamidomethylation; Deamidation (NQ) | 12387 | PPTG_01552 | Phytophthora parasitica INRA-310 hypothetical protein (113 aa) |
| 3390 | 2883 | PPTG_05743 | 59,82 | 3 | 0 | 0 | 3 | | | 9,21E+05 | 2 | 2 | 0 | 0 | 2 | | 83099 | PPTG_05743 | Phytophthora parasitica INRA-310 transmembrane protein (753 aa) |
| 2547 | 2670 | PPTG_16477 | 59,78 | 17 | 6 | 6 | 17 | 2,20E+05 | 6,73E+05 | 2,91E+06 | 3 | 3 | 1 | 1 | 3 | Carbamidomethylation | 27297 | PPTG_16477 | Phytophthora parasitica INRA-310 isoamyl acetate-hydrolyzing esterase 1 (245 aa) |
| 2295 | 2538 | PPTG_12655 | 59,73 | 34 | 0 | 23 | 34 | | 3,89E+05 | 1,73E+06 | 3 | 3 | 0 | 2 | 4 | Dehydration; Deamidation (NQ) | 12488 | PPTG_12655 | Phytophthora parasitica INRA-310 peptide-methionine (S)-S-oxide reductase (108 aa) |
| 2625 | 2825 | PPTG_13075 | 59,7 | 12 | 0 | 9 | 12 | | 5,90E+05 | 9,71E+05 | 3 | 3 | 0 | 2 | 3 | | 30686 | PPTG_13075 | Phytophthora parasitica INRA-310 hypothetical protein (266 aa) |
| 2729 | 3175 | PPTG_09937 | 59,69 | 6 | 0 | 6 | 6 | | 1,19E+06 | 1,70E+06 | 2 | 2 | 0 | 2 | 2 | | 37179 | PPTG_09937 | Phytophthora parasitica INRA-310 rRNA-processing protein EBP2 (335 aa) |
| 2225 | 2536 | PPTG_09440 | 59,68 | 17 | 5 | 13 | 12 | 6,83E+04 | 1,61E+06 | 2,03E+06 | 4 | 4 | 1 | 3 | 3 | Carbamidomethylation | 32694 | PPTG_09440 | Phytophthora parasitica INRA-310 ubiquinone/menaquinone biosynthesis methyltransferase ubiE (293 aa) |
| 2818 | 3172 | PPTG_02056 | 59,63 | 12 | 0 | 12 | 12 | | 5,13E+05 | 4,28E+06 | 2 | 2 | 0 | 2 | 2 | Carbamidomethylation | 21671 | PPTG_02056 | Phytophthora parasitica INRA-310 hypothetical protein (199 aa) |
| 2880 | 3335 | PPTG_20110 | 59,62 | 21 | 0 | 21 | 15 | | 1,33E+07 | 4,69E+06 | 2 | 2 | 0 | 3 | 1 | Carbamidomethylation | 16029 | PPTG_20110 | Phytophthora parasitica INRA-310 S-phase kinase-associated protein 1A (141 aa) |
| 2457 | 2541 | PPTG_04041 | 59,6 | 7 | 0 | 0 | 7 | | | 9,00E+05 | 5 | 5 | 0 | 0 | 5 | Carbamidomethylation | 87110 | PPTG_04041 | Phytophthora parasitica INRA-310 cAMP-dependent protein kinase catalytic subunit (780 aa) |

| | | | | | | | | | | | | | | | | | | | |
|---|---|---|---|---|---|---|---|---|---|---|---|---|---|---|---|---|---|---|---|
| 2936 | 3378 | PPTG_00234 | 59,59 | 7 | 3 | 3 | 7 | 1,84E+05 | 6,48E+04 | 1,52E+06 | 2 | 2 | 1 | 1 | 2 | Oxidation (M) | 49925 | PPTG_00234 | Phytophthora parasitica INRA-310 drug/Metabolite transporter (DMT) superfamily (444 aa) |
| 2705 | 2190 | PPTG_14743 | 59,58 | 9 | 0 | 0 | 9 | | | 6,52E+05 | 4 | 4 | 0 | 0 | 4 | | 57778 | PPTG_14743 | Phytophthora parasitica INRA-310 endoplasmic oxidoreductin (513 aa) |
| 3249 | 3412 | PPTG_19638 | 59,48 | 17 | 0 | 17 | 9 | | 1,47E+06 | 1,47E+05 | 2 | 2 | 0 | 2 | 1 | | 20782 | PPTG_19638 | Phytophthora parasitica INRA-310 geranylgeranyl transferase type-2 subunit alpha (179 aa) |
| 2091 | 2575 | PPTG_15190 | 59,47 | 4 | 1 | 3 | 4 | 6,94E+04 | 1,69E+06 | 1,14E+06 | 4 | 4 | 1 | 3 | 4 | | 90045 | PPTG_15190 | Phytophthora parasitica INRA-310 hypothetical protein (804 aa) |
| 2211 | 2500 | PPTG_09784 | 59,47 | 9 | 0 | 9 | 7 | | 1,69E+06 | 1,36E+06 | 4 | 4 | 0 | 4 | 3 | Oxidation (M); Carbamidomethylation | 52385 | PPTG_09784 | Phytophthora parasitica INRA-310 CUG-BP- and ETR-3-like factor (496 aa) |
| 2575 | 2702 | PPTG_02352 | 59,45 | 8 | 0 | 5 | 8 | | 1,52E+05 | 3,87E+05 | 3 | 3 | 0 | 2 | 3 | | 41358 | PPTG_02352 | Phytophthora parasitica INRA-310 hypothetical protein (393 aa) |
| 2216 | 3210 | PPTG_10135 | 59,42 | 5 | 2 | 5 | 5 | 2,82E+05 | 5,69E+05 | 2,30E+06 | 2 | 1 | 1 | 3 | 3 | Oxidation (M); Carbamidomethylation | 54307 | PPTG_10135 | Phytophthora parasitica INRA-310 major facilitator superfamily transporter (MFS) (501 aa) |
| 2395 | 2705 | PPTG_02924 | 59,42 | 34 | 13 | 25 | 34 | 2,08E+05 | 2,43E+06 | 3,84E+06 | 3 | 3 | 1 | 2 | 3 | Acetylation (N-term) | 10558 | PPTG_02924 | Phytophthora parasitica INRA-310 U6 snRNA-associated Sm-like protein LSm3 (93 aa) |
| 2867 | 3127 | PPTG_04658 | 59,4 | 14 | 8 | 8 | 14 | 1,07E+05 | 2,71E+04 | 1,67E+06 | 2 | 2 | 1 | 1 | 2 | Carbamidomethylation | 21454 | PPTG_04658 | Phytophthora parasitica INRA-310 hypothetical protein (182 aa) |
| 2213 | 2569 | PPTG_14221 | 59,39 | 17 | 3 | 13 | 14 | | 6,92E+06 | 1,96E+06 | 3 | 2 | 1 | 3 | 3 | Oxidation (M); Carbamidomethylation; Acetylation (N-term); Carbamylation | 29302 | PPTG_14221 | Phytophthora parasitica INRA-310 hypothetical protein (276 aa) |
| 2595 | 2635 | PPTG_14030 | 59,39 | 13 | 0 | 10 | 13 | | 5,82E+05 | 1,70E+06 | 3 | 3 | 0 | 2 | 3 | Carbamidomethylation; Acetylation (N-term) | 35277 | PPTG_14030 | Phytophthora parasitica INRA-310 hypothetical protein (326 aa) |
| 2970 | 2809 | PPTG_01350 | 59,38 | 2 | 0 | 1 | 1 | | 3,33E+05 | 7,27E+05 | 3 | 3 | 0 | 2 | 1 | | 157947 | PPTG_01350 | Phytophthora parasitica INRA-310 hypothetical protein (1415 aa) |
| 3110 | 3226 | PPTG_01154 | 59,28 | 9 | 0 | 9 | 0 | | 1,18E+05 | | 2 | 2 | 0 | 3 | 0 | | 29176 | PPTG_01154 | Phytophthora parasitica INRA-310 enoyl-[acyl-carrier-protein] reductase [NADH] (278 aa) |
| 2468 | 2318 | PPTG_10912 | 59,25 | 4 | 0 | 4 | 3 | | 8,86E+05 | 1,37E+06 | 3 | 3 | 0 | 3 | 2 | | 102178 | PPTG_10912 | Phytophthora parasitica INRA-310 nuclear pore complex protein Nup93-like protein (916 aa) |
| 2256 | 2715 | PPTG_01475 | 59,24 | 13 | 3 | 13 | 13 | 7,85E+04 | 1,91E+06 | 1,45E+06 | 3 | 3 | 1 | 3 | 3 | | 37329 | PPTG_01475 | Phytophthora parasitica INRA-310 hypothetical protein (351 aa) |
| 3212 | 3636 | PPTG_04357 | 59,24 | 5 | 0 | 5 | 3 | | 2,96E+05 | 9,37E+04 | 2 | 2 | 0 | 2 | 1 | | 63871 | PPTG_04357 | Phytophthora parasitica INRA-310 aspartyl/glutamyl-tRNA(Asn/Gln) amidotransferase subunit B (575 aa) |
| 3419 | 2894 | PPTG_05040 | 59,14 | 5 | 0 | 0 | 5 | | | 6,52E+05 | 2 | 2 | 0 | 0 | 2 | | 59196 | PPTG_05040 | Phytophthora parasitica INRA-310 acetyl-coenzyme A transporter (535 aa) |
| 2667 | 2699 | PPTG_10847 | 59,12 | 4 | 0 | 4 | 1 | | 3,26E+05 | 1,03E+05 | 3 | 3 | 0 | 3 | 1 | | 122272 | PPTG_10847 | Phytophthora parasitica INRA-310 hypothetical protein (1109 aa) |
| 2638 | 3635 | PPTG_05409 | 59,09 | 21 | 12 | 12 | 21 | 1,24E+05 | 1,01E+05 | 1,54E+06 | 2 | 2 | 1 | 1 | 3 | Oxidation (M) | 12481 | PPTG_05409 | Phytophthora parasitica INRA-310 hypothetical protein (114 aa) |
| 2192 | 2439 | PPTG_05937 | 59,09 | 19 | 4 | 4 | 19 | 3,01E+05 | 5,43E+05 | 6,66E+06 | 4 | 4 | 1 | 1 | 5 | Oxidation (M) | 25677 | PPTG_05937 | Phytophthora parasitica INRA-310 hypothetical protein (224 aa) |
| 2678 | 2406 | PPTG_09183 | 59,09 | 2 | 0 | 1 | 1 | | 4,52E+05 | 6,56E+05 | 3 | 3 | 0 | 2 | 2 | Carbamidomethylation | 209332 | PPTG_09183 | Phytophthora parasitica INRA-310 histone deacetylase (1923 aa) |
| 2974 | 2785 | PPTG_15721 | 59,07 | 2 | 0 | 0 | 2 | | | 9,66E+05 | 3 | 3 | 0 | 0 | 3 | | 194124 | PPTG_15721 | Phytophthora parasitica INRA-310 ATP-binding Cassette (ABC) superfamily (1774 aa) |
| 3389 | 2840 | PPTG_08402 | 59,07 | 8 | 0 | 0 | 8 | | | 5,72E+05 | 2 | 2 | 0 | 0 | 2 | | 41859 | PPTG_08402 | Phytophthora parasitica INRA-310 cytosolic 5'-nucleotidase III-like protein (370 aa) |
| 2075 | 2375 | PPTG_04611 | 59,03 | 29 | 12 | 29 | 10 | 3,04E+06 | 1,71E+07 | 2,21E+07 | 4 | 4 | 2 | 4 | 2 | | 18767 | PPTG_04611 | Phytophthora parasitica INRA-310 AP-2 complex subunit sigma (157 aa) |
| 2573 | 2682 | PPTG_13162 | 59,01 | 3 | 1 | 2 | 2 | 1,50E+05 | 6,96E+05 | 1,05E+06 | 3 | 3 | 1 | 2 | 2 | | 135624 | PPTG_13162 | Phytophthora parasitica INRA-310 hypothetical protein (1239 aa) |
| 2748 | 2774 | PPTG_01577 | 58,98 | 7 | 0 | 4 | 5 | | 1,62E+05 | 4,96E+05 | 3 | 3 | 0 | 2 | 2 | | 47899 | PPTG_01577 | Phytophthora parasitica INRA-310 hypothetical protein (431 aa) |
| 2748 | 2775 | PPTG_01577 | 58,98 | 7 | 0 | 4 | 5 | | 1,62E+05 | 4,96E+05 | 3 | 3 | 0 | 2 | 2 | | 47899 | PPTG_01577 | Phytophthora parasitica INRA-310 hypothetical protein (431 aa) |
| 2813 | 2874 | PPTG_07579 | 58,94 | 10 | 0 | 10 | 3 | | 4,70E+05 | 1,72E+05 | 3 | 3 | 0 | 3 | 1 | | 36424 | PPTG_07579 | Phytophthora parasitica INRA-310 flagellar radial spoke protein (327 aa) |
| 2677 | 2213 | PPTG_13786 | 58,89 | 7 | 0 | 0 | 7 | | | 2,54E+06 | 4 | 4 | 0 | 0 | 4 | | 68860 | PPTG_13786 | Phytophthora parasitica INRA-310 hypothetical protein (643 aa) |
| 2488 | 2981 | PPTG_16180 | 58,87 | 7 | 4 | 7 | 7 | 8,54E+04 | 1,86E+06 | 3,58E+06 | 2 | 2 | 1 | 2 | 2 | Carbamidomethylation | 43428 | PPTG_16180 | Phytophthora parasitica INRA-310 histidine acid phosphatase (381 aa) |
| 2605 | 2829 | PPTG_13971 | 58,86 | 7 | 0 | 7 | 5 | | 2,55E+06 | 1,74E+06 | 3 | 3 | 0 | 3 | 2 | | 39176 | PPTG_13971 | Phytophthora parasitica INRA-310 hypothetical protein (347 aa) |
| 2354 | 2787 | PPTG_16026 | 58,84 | 6 | 3 | 4 | 6 | 9,87E+04 | 4,50E+05 | 1,20E+06 | 3 | 3 | 1 | 2 | 3 | | 50835 | PPTG_16026 | Phytophthora parasitica INRA-310 hypothetical protein (470 aa) |
| 2799 | 2480 | PPTG_13994 | 58,69 | 5 | 2 | 0 | 5 | 4,86E+04 | | 1,11E+06 | 3 | 3 | 1 | 0 | 3 | | 60602 | PPTG_13994 | Phytophthora parasitica INRA-310 adenylate kinase (547 aa) |
| 2985 | 3077 | PPTG_18033 | 58,66 | 7 | 0 | 7 | 3 | | 6,69E+05 | 1,96E+05 | 2 | 2 | 0 | 2 | 1 | Carbamidomethylation | 53958 | PPTG_18033 | Phytophthora parasitica INRA-310 glutathione reductase (501 aa) |
| 2466 | 2257 | PPTG_13377 | 58,64 | 1 | 0 | 1 | 1 | | 5,06E+05 | 1,35E+06 | 3 | 3 | 0 | 2 | 3 | | 290512 | PPTG_13377 | Phytophthora parasitica INRA-310 hypothetical protein (2584 aa) |
| 2477 | 2650 | PPTG_09917 | 58,62 | 7 | 0 | 4 | 7 | | 2,04E+05 | 6,23E+05 | 3 | 3 | 0 | 2 | 3 | | 70967 | PPTG_09917 | Phytophthora parasitica INRA-310 leishmanolysin-like peptidase metalloprotease family M08 (649 aa) |
| 2510 | 2633 | PPTG_04657 | 58,57 | 6 | 0 | 3 | 5 | | 6,06E+05 | 1,70E+06 | 4 | 4 | 0 | 2 | 3 | | 77507 | PPTG_04657 | Phytophthora parasitica INRA-310 serine/threonine protein kinase (708 aa) |
| 2472 | 2503 | PPTG_17698 | 58,47 | 2 | 0 | 0 | 2 | | | 1,48E+06 | 5 | 5 | 0 | 0 | 5 | | 299132 | PPTG_17698 | Phytophthora parasitica INRA-310 3'5'-cyclic nucleotide phosphodiesterase (2717 aa) |
| 2334 | 2558 | PPTG_18447 | 58,44 | 9 | 3 | 9 | 5 | 7,14E+05 | 2,84E+06 | 4,68E+06 | 3 | 2 | 1 | 3 | 2 | | 33624 | PPTG_18447 | Phytophthora parasitica INRA-310 hypothetical protein (303 aa) |
| 2724 | 2901 | PPTG_14408 | 58,42 | 3 | 1 | 0 | 2 | 1,76E+05 | | 5,99E+05 | 3 | 3 | 1 | 0 | 3 | Oxidation (M) | 105599 | PPTG_14408 | Phytophthora parasitica INRA-310 exocyst complex component 4 (949 aa) |
| 3148 | 3247 | PPTG_05666 | 58,41 | 15 | 0 | 15 | 6 | | 3,64E+05 | 1,67E+05 | 2 | 2 | 0 | 2 | 1 | Carbamidomethylation | 35883 | PPTG_05666 | Phytophthora parasitica INRA-310 hypothetical protein (338 aa) |
| 2394 | 2481 | PPTG_00622 | 58,34 | 8 | 0 | 3 | 8 | | 1,20E+05 | 3,01E+05 | 3 | 3 | 0 | 2 | 4 | Deamidation (NQ) | 78636 | PPTG_00622 | Phytophthora parasitica INRA-310 hypothetical protein (732 aa) |
| 2808 | 2824 | PPTG_05300 | 58,15 | 5 | 0 | 4 | 4 | | 2,30E+05 | 2,51E+05 | 3 | 3 | 0 | 2 | 2 | Carbamidomethylation | 76065 | PPTG_05300 | Phytophthora parasitica INRA-310 tRNA-dihydrouridine synthase 3 (680 aa) |
| 2093 | 3132 | PPTG_06475 | 58,14 | 14 | 14 | 14 | 14 | 8,94E+05 | 5,28E+06 | 6,96E+06 | 2 | 2 | 2 | 3 | 3 | Carbamidomethylation | 27129 | PPTG_06475 | Phytophthora parasitica INRA-310 hypothetical protein (239 aa) |
| 3147 | 3217 | PPTG_12287 | 58,13 | 11 | 0 | 4 | 11 | | 7,33E+05 | 1,38E+06 | 2 | 2 | 0 | 1 | 2 | Oxidation (M) | 28133 | PPTG_12287 | Phytophthora parasitica INRA-310 cysteine protease family C01A (256 aa) |
| 2861 | 2906 | PPTG_06089 | 58,11 | 7 | 0 | 7 | 7 | | 3,79E+06 | 5,41E+06 | 2 | 2 | 0 | 2 | 2 | Carbamidomethylation | 38580 | PPTG_06089 | Phytophthora parasitica INRA-310 hypothetical protein (357 aa) |
| 2246 | 2837 | PPTG_05209 | 58,08 | 13 | 7 | 9 | 9 | 4,91E+04 | 2,57E+05 | 1,15E+06 | 3 | 3 | 2 | 2 | 3 | Deamidation (NQ) | 25924 | PPTG_05209 | Phytophthora parasitica INRA-310 hypothetical protein (232 aa) |
| 2874 | 3255 | PPTG_04933 | 58,08 | 6 | 1 | 6 | 1 | | 1,41E+05 | | 2 | 1 | 1 | 2 | 1 | | 56529 | PPTG_04933 | Phytophthora parasitica INRA-310 glycoside hydrolase (501 aa) |
| 3118 | 3402 | PPTG_17416 | 58,07 | 8 | 0 | 4 | 8 | 0 | | 6,02E+05 | 2 | 2 | 0 | 1 | 2 | Carbamidomethylation | 46435 | PPTG_17416 | Phytophthora parasitica INRA-310 hypothetical protein (442 aa) |
| 3543 | 3007 | PPTG_14806 | 58,06 | 5 | 0 | 0 | 5 | | | 6,66E+05 | 2 | 2 | 0 | 0 | 2 | | 52619 | PPTG_14806 | Phytophthora parasitica INRA-310 hypothetical protein (466 aa) |
| 2017 | 2518 | PPTG_18558 | 58,03 | 4 | 0 | 4 | 4 | | 5,27E+05 | 7,23E+05 | 3 | 3 | 0 | 4 | 4 | Deamidation (NQ) | 98681 | PPTG_18558 | Phytophthora parasitica INRA-310 bifunctional aspartokinase/homoserine dehydrogenase (912 aa) |
| 2057 | 2270 | PPTG_01425 | 58,02 | 6 | 1 | 4 | 6 | 1,45E+05 | 6,67E+05 | 1,60E+06 | 4 | 4 | 1 | 3 | 4 | | 83134 | PPTG_01425 | Phytophthora parasitica INRA-310 vacuolar protein sorting-associated protein (725 aa) |
| 2628 | 3236 | PPTG_15170 | 57,98 | 5 | 3 | 5 | 5 | 5,15E+05 | 9,06E+05 | 6,83E+06 | 2 | 1 | 1 | 2 | 2 | | 41770 | PPTG_15170 | Phytophthora parasitica INRA-310 polygalacturonase (398 aa) |
| 2181 | 2605 | PPTG_14571 | 57,95 | 6 | 1 | 5 | 4 | 1,03E+05 | 3,12E+05 | 2,04E+06 | 4 | 4 | 1 | 3 | 3 | | 79803 | PPTG_14571 | Phytophthora parasitica INRA-310 hypothetical protein (701 aa) |
| 2148 | 2504 | PPTG_19140 | 57,95 | 5 | 0 | 5 | 4 | | 4,49E+05 | 1,14E+06 | 4 | 4 | 0 | 4 | 3 | Carbamidomethylation | 80229 | PPTG_19140 | Phytophthora parasitica INRA-310 dicer-like (719 aa) |
| 3684 | 4093 | PPTG_19803 | 57,87 | 16 | 0 | 16 | 16 | | 3,69E+06 | 3,26E+06 | 1 | 1 | 0 | 1 | 1 | Carbamidomethylation | 10479 | PPTG_19803 | Phytophthora parasitica INRA-310 hypothetical protein (98 aa) |
| 2393 | 2469 | PPTG_18443 | 57,87 | 12 | 0 | 7 | 12 | | 3,14E+05 | 1,56E+06 | 4 | 4 | 0 | 2 | 4 | | 41456 | PPTG_18443 | Phytophthora parasitica INRA-310 extracellular signal-regulated kinase 2 (366 aa) |
| 3030 | 2869 | PPTG_13015 | 57,84 | 6 | 0 | 6 | 0 | | 4,08E+05 | | 2 | 2 | 0 | 3 | 0 | Carbamidomethylation; Deamidation (NQ) | 58650 | PPTG_13015 | Phytophthora parasitica INRA-310 hypothetical protein (525 aa) |
| 2286 | 3092 | PPTG_12406 | 57,82 | 9 | 9 | 9 | 9 | 1,42E+05 | 1,77E+05 | 1,68E+06 | 1 | 1 | 2 | 1 | 3 | Deamidation (NQ) | 20125 | PPTG_12406 | Phytophthora parasitica INRA-310 hypothetical protein (180 aa) |
| 2626 | 2877 | PPTG_13517 | 57,76 | 8 | 5 | 5 | 8 | 1,10E+05 | 5,99E+05 | 1,00E+06 | 2 | 2 | 1 | 2 | 2 | Oxidation (M) | 33208 | PPTG_13517 | Phytophthora parasitica INRA-310 bifunctional folD protein (308 aa) |
| 2352 | 2628 | PPTG_12383 | 57,72 | 11 | 0 | 11 | 11 | | 1,78E+06 | 1,82E+06 | 3 | 3 | 0 | 3 | 3 | Carbamidomethylation | 44894 | PPTG_12383 | Phytophthora parasitica INRA-310 RecA protein (424 aa) |
| 3544 | 3049 | PPTG_15761 | 57,7 | 32 | 0 | 0 | 32 | | | 5,89E+05 | 2 | 2 | 0 | 0 | 2 | Carbamidomethylation | 14580 | PPTG_15761 | Phytophthora parasitica INRA-310 hypothetical protein (137 aa) |
| 2425 | 4218 | PPTG_06766 | 57,65 | 9 | 9 | 9 | 9 | 1,03E+06 | 2,13E+06 | 1,03E+07 | 1 | 1 | 1 | 2 | 3 | Oxidation (M) | 19786 | PPTG_06766 | Phytophthora parasitica INRA-310 Solute:Sodium Symporter (SSS) family (186 aa) |
| 3262 | 3560 | PPTG_01661 | 57,65 | 27 | 0 | 13 | 27 | | 1,69E+05 | 1,64E+06 | 2 | 2 | 0 | 1 | 2 | Carbamidomethylation | 12169 | PPTG_01661 | Phytophthora parasitica INRA-310 complement control module protein (116 aa) |
| 2993 | 2769 | PPTG_01839 | 57,62 | 10 | 0 | 0 | 10 | | | 2,87E+06 | 3 | 3 | 0 | 0 | 3 | | 63418 | PPTG_01839 | Phytophthora parasitica INRA-310 hypothetical protein (574 aa) |
| 2601 | 2707 | PPTG_17656 | 57,62 | 10 | 5 | 3 | 10 | 9,34E+04 | 2,85E+05 | 2,70E+06 | 3 | 3 | 1 | 1 | 3 | Carbamidomethylation | 38107 | PPTG_17656 | Phytophthora parasitica INRA-310 hypothetical protein (341 aa) |
| 2962 | 2959 | PPTG_12395 | 57,59 | 0 | 0 | 0 | 0 | | 1,48E+05 | 2,43E+05 | 2 | 2 | 0 | 2 | 1 | | 686216 | PPTG_12395 | Phytophthora parasitica INRA-310 midasin-like protein (6123 aa) |
| 2642 | 2556 | PPTG_16209 | 57,5 | 9 | 0 | 9 | 6 | | 1,08E+06 | 9,71E+05 | 3 | 3 | 0 | 3 | 2 | | 37679 | PPTG_16209 | Phytophthora parasitica INRA-310 GPN-loop GTPase 1 (346 aa) |
| 2229 | 2994 | PPTG_02312 | 57,48 | 24 | 18 | 18 | 18 | 7,06E+04 | 2,83E+06 | 5,77E+06 | 3 | 3 | 2 | 2 | 3 | Oxidation (M) | 13215 | PPTG_02312 | Phytophthora parasitica INRA-310 50S ribosomal protein L20 (116 aa) |
| 3011 | 2645 | PPTG_00762 | 57,45 | 3 | 0 | 0 | 3 | | | 9,08E+05 | 3 | 3 | 0 | 0 | 3 | | 113263 | PPTG_00762 | Phytophthora parasitica INRA-310 hypothetical protein (1016 aa) |
| 2786 | 3291 | PPTG_12650 | 57,44 | 9 | 5 | 9 | 5 | 1,09E+05 | 7,01E+05 | 9,09E+05 | 2 | 2 | 1 | 2 | 1 | | 31417 | PPTG_12650 | Phytophthora parasitica INRA-310 hypothetical protein (269 aa) |
| 2443 | 3396 | PPTG_15693 | 57,43 | 22 | 12 | 11 | 22 | 3,42E+05 | 1,19E+06 | 1,63E+07 | 2 | 2 | 1 | 1 | 4 | Oxidation (M) | 9417 | PPTG_15693 | Phytophthora parasitica INRA-310 hypothetical protein (95 aa) |
| 3346 | 2917 | PPTG_00741 | 57,42 | 6 | 0 | 6 | 0 | | 4,37E+05 | | 2 | 2 | 0 | 2 | 0 | | 61668 | PPTG_00741 | Phytophthora parasitica INRA-310 amino-acid acetyltransferase (564 aa) |
| 2710 | 2308 | PPTG_03976 | 57,38 | 12 | 0 | 6 | 6 | | 3,08E+05 | 5,26E+05 | 4 | 3 | 0 | 2 | 2 | Carbamidomethylation | 42219 | PPTG_03976 | Phytophthora parasitica INRA-310 methionine aminopeptidase (384 aa) |
| 2742 | 2646 | PPTG_14430 | 57,37 | 7 | 0 | 7 | 3 | | 7,53E+05 | 9,06E+04 | 3 | 3 | 0 | 3 | 1 | | 64585 | PPTG_14430 | Phytophthora parasitica INRA-310 hypothetical protein (583 aa) |
| 3651 | 3430 | PPTG_19580 | 57,33 | 17 | 0 | 0 | 17 | | | 6,19E+05 | 2 | 2 | 0 | 0 | 2 | | 14562 | PPTG_19580 | Phytophthora parasitica INRA-310 hypothetical protein (129 aa) |
| 2282 | 2404 | PPTG_01417 | 57,25 | 3 | 0 | 1 | 3 | | 4,93E+05 | 1,75E+06 | 4 | 4 | 0 | 2 | 4 | | 154759 | PPTG_01417 | Phytophthora parasitica INRA-310 hypothetical protein (1410 aa) |
| 2260 | 3540 | PPTG_10450 | 57,23 | 16 | 9 | 16 | 16 | 4,38E+05 | 2,61E+06 | 5,93E+06 | 2 | 2 | 1 | 3 | 3 | Oxidation (M) | 15030 | PPTG_10450 | Phytophthora parasitica INRA-310 nuclear transcription factor Y subunit (129 aa) |
| 2270 | 2663 | PPTG_13150 | 57,13 | 1 | 0 | 1 | 1 | 1,70E+04 | 1,74E+05 | 4,24E+05 | 3 | 3 | 1 | 3 | 2 | | 362204 | PPTG_13150 | Phytophthora parasitica INRA-310 myosin-like protein (3296 aa) |
| 2908 | 3262 | PPTG_04371 | 57,09 | 5 | 0 | 5 | 5 | | 1,99E+05 | 3,19E+06 | 2 | 2 | 0 | 2 | 2 | | 55740 | PPTG_04371 | Phytophthora parasitica INRA-310 lysophospholipid acyltransferase (494 aa) |
| 2152 | 2266 | PPTG_15470 | 57,04 | 12 | 0 | 9 | 8 | | 1,27E+06 | 3,38E+06 | 4 | 4 | 0 | 3 | 4 | Oxidation (M); Carbamidomethylation | 35657 | PPTG_15470 | Phytophthora parasitica INRA-310 replication factor C subunit 2 (328 aa) |
| 2931 | 2731 | PPTG_02287 | 56,99 | 10 | 0 | 4 | 10 | | 5,75E+05 | 2,39E+06 | 3 | 3 | 0 | 1 | 3 | Carbamidomethylation | 43226 | PPTG_02287 | Phytophthora parasitica INRA-310 hypothetical protein (394 aa) |
| 2139 | 3653 | PPTG_15686 | 56,99 | 15 | 15 | 15 | 15 | 2,13E+06 | 1,10E+07 | 4,33E+07 | 1 | 1 | 1 | 2 | 4 | Oxidation (M) | 13169 | PPTG_15686 | Phytophthora parasitica INRA-310 hypothetical protein (119 aa) |
| 2589 | 2408 | PPTG_11192 | 56,97 | 13 | 0 | 13 | 5 | | 2,03E+06 | 1,09E+06 | 3 | 3 | 0 | 3 | 2 | | 40624 | PPTG_11192 | Phytophthora parasitica INRA-310 hypothetical protein (373 aa) |
| 2784 | 3180 | PPTG_14021 | 56,95 | 15 | 9 | 6 | 15 | 1,48E+05 | 3,44E+05 | 3,45E+06 | 2 | 2 | 1 | 1 | 2 | | 20627 | PPTG_14021 | Phytophthora parasitica INRA-310 ADP-ribosylation factor family protein (184 aa) |
| 2787 | 3418 | PPTG_18038 | 56,85 | 9 | 6 | 6 | 4 | 0 | 5,16E+05 | 6,78E+05 | 2 | 2 | 2 | 1 | 1 | Oxidation (M); Carbamidomethylation | 37982 | PPTG_18038 | Phytophthora parasitica INRA-310 WD domain-containing protein (346 aa) |
| 2787 | 3419 | PPTG_18038 | 56,85 | 9 | 5 | 5 | 4 | 0 | 5,16E+05 | 6,78E+05 | 2 | 2 | 2 | 1 | 1 | Oxidation (M); Carbamidomethylation | 38474 | PPTG_18038 | Phytophthora parasitica INRA-310 WD domain-containing protein (351 aa) |
| 2892 | 4151 | PPTG_10229 | 56,81 | 5 | 0 | 5 | 5 | | 1,53E+06 | 0 | 2 | 2 | 0 | 3 | 1 | | 39236 | PPTG_10229 | Phytophthora parasitica INRA-310 hypothetical protein (371 aa) |
| 2500 | 2953 | PPTG_02554 | 56,8 | 4 | 2 | 4 | 4 | 8,56E+04 | 9,63E+05 | 6,45E+05 | 2 | 2 | 1 | 2 | 2 | | 65877 | PPTG_02554 | Phytophthora parasitica INRA-310 methylenetetrahydrofolate reductase (592 aa) |
| 2637 | 3527 | PPTG_09335 | 56,78 | 40 | 0 | 22 | 40 | | 3,74E+06 | 3,93E+06 | 3 | 3 | 0 | 2 | 3 | Carbamidomethylation | 9360 | PPTG_09335 | Phytophthora parasitica INRA-310 hypothetical protein (87 aa) |
| 2824 | 3259 | PPTG_12735 | 56,72 | 9 | 0 | 4 | 9 | | 1,70E+05 | 1,54E+06 | 2 | 2 | 0 | 1 | 3 | Carbamidomethylation; Deamidation (NQ) | 29855 | PPTG_12735 | Phytophthora parasitica INRA-310 SNARE associated Golgi protein (273 aa) |
| 2910 | 3321 | PPTG_16064 | 56,66 | 6 | 0 | 3 | 6 | | 4,74E+05 | 5,24E+05 | 2 | 2 | 0 | 2 | 2 | | 57318 | PPTG_16064 | Phytophthora parasitica INRA-310 U4/U6 small nuclear ribonucleoprotein Prp31 (528 aa) |
| 3162 | 3587 | PPTG_16255 | 56,65 | 25 | 0 | 9 | 25 | | 6,36E+05 | 1,11E+06 | 2 | 2 | 0 | 1 | 2 | | 12814 | PPTG_16255 | Phytophthora parasitica INRA-310 dynein light chain (118 aa) |
| 2543 | 2529 | PPTG_10636 | 56,64 | 12 | 4 | 12 | 4 | 1,67E+05 | 2,81E+06 | 1,01E+05 | 3 | 3 | 1 | 3 | 1 | Carbamidomethylation | 25418 | PPTG_10636 | Phytophthora parasitica INRA-310 hypothetical protein (230 aa) |
| 2688 | 3128 | PPTG_13401 | 56,64 | 2 | 0 | 2 | 2 | | 4,71E+05 | 1,69E+06 | 2 | 2 | 0 | 2 | 2 | | 102852 | PPTG_13401 | Phytophthora parasitica INRA-310 RAC family serine/threonine-protein kinase (928 aa) |
| 2688 | 3129 | PPTG_13401 | 56,64 | 2 | 0 | 2 | 2 | | 4,71E+05 | 1,69E+06 | 2 | 2 | 0 | 2 | 2 | | 102852 | PPTG_13401 | Phytophthora parasitica INRA-310 RAC family serine/threonine-protein kinase (928 aa) |
| 3091 | 2602 | PPTG_05763 | 56,63 | 5 | 0 | 0 | 5 | | | 1,43E+06 | 3 | 3 | 0 | 0 | 3 | | 76564 | PPTG_05763 | Phytophthora parasitica INRA-310 sulfate Permease (SulP) family (701 aa) |
| 2475 | 2292 | PPTG_10394 | 56,62 | 21 | 5 | 0 | 21 | 7,31E+05 | | 1,09E+07 | 4 | 4 | 1 | 0 | 4 | Oxidation (M) | 28723 | PPTG_10394 | Phytophthora parasitica INRA-310 hypothetical protein (259 aa) |
| 2830 | 3398 | PPTG_20359 | 56,6 | 23 | 12 | 23 | 12 | 6,33E+05 | 4,79E+06 | 4,86E+06 | 2 | 2 | 1 | 2 | 1 | | 11074 | PPTG_20359 | Phytophthora parasitica INRA-310 hypothetical protein (97 aa) |
| 2576 | 2708 | PPTG_13952 | 56,58 | 3 | 0 | 2 | 3 | | 5,01E+05 | 7,29E+05 | 3 | 3 | 0 | 2 | 3 | | 111290 | PPTG_13952 | Phytophthora parasitica INRA-310 hypothetical protein (1004 aa) |
| 3092 | 2676 | PPTG_05946 | 56,57 | 17 | 0 | 0 | 17 | | | 1,65E+06 | 3 | 3 | 0 | 0 | 3 | | 25634 | PPTG_05946 | Phytophthora parasitica INRA-310 hypothetical protein (225 aa) |
| 2754 | 3185 | PPTG_16624 | 56,54 | 3 | 0 | 3 | 3 | | 5,05E+05 | 7,59E+05 | 2 | 2 | 0 | 2 | 2 | Carbamidomethylation | 56820 | PPTG_16624 | Phytophthora parasitica INRA-310 regulator-chromosome condensation (RCC1)-like protein (529 aa) |
| 2607 | 2992 | PPTG_13033 | 56,52 | 11 | 6 | 11 | 11 | 4,51E+04 | 8,09E+05 | 1,84E+06 | 2 | 2 | 1 | 2 | 2 | Oxidation (M) | 20515 | PPTG_13033 | Phytophthora parasitica INRA-310 phenol acid carboxylase (179 aa) |
| 2938 | 3397 | PPTG_09565 | 56,43 | 6 | 0 | 6 | 6 | | 4,05E+05 | 5,37E+05 | 2 | 2 | 0 | 2 | 2 | | 44500 | PPTG_09565 | Phytophthora parasitica INRA-310 hypothetical protein (414 aa) |
| 2331 | 2370 | PPTG_02847 | 56,4 | 12 | 0 | 4 | 10 | | 4,03E+05 | 9,75E+05 | 5 | 5 | 0 | 2 | 4 | | 58115 | PPTG_02847 | Phytophthora parasitica INRA-310 hypothetical protein (523 aa) |
| 2243 | 2576 | PPTG_05504 | 56,36 | 3 | 1 | 3 | 3 | 6,97E+04 | 1,11E+06 | 1,62E+06 | 3 | 3 | 1 | 3 | 3 | | 93602 | PPTG_05504 | Phytophthora parasitica INRA-310 hypothetical protein (835 aa) |

| | | | | | | | | | | | | | | | | | | | |
|---|---|---|---|---|---|---|---|---|---|---|---|---|---|---|---|---|---|---|---|
| 2534 | 3271 | PPTG_17134 | 56,29 | 6 | 6 | 2 | 6 | 1,29E+05 | 1,10E+05 | 7,42E+05 | 2 | 2 | 2 | 1 | 2 | | 41035 | PPTG_17134 | Phytophthora parasitica INRA-310 3-phosphoinositide-dependent protein kinase (364 aa) |
| 2534 | 3273 | PPTG_17134 | 56,29 | 5 | 5 | 2 | 5 | 1,29E+05 | 1,10E+05 | 7,42E+05 | 2 | 2 | 2 | 1 | 2 | | 51984 | PPTG_17134 | Phytophthora parasitica INRA-310 3-phosphoinositide-dependent protein kinase (460 aa) |
| 2680 | 3074 | PPTG_19805 | 56,29 | 2 | 1 | 1 | 2 | | | 5,10E+05 | 2 | 1 | 1 | 1 | 2 | Carbamidomethylation | 105604 | PPTG_19805 | Phytophthora parasitica INRA-310 anoctamin-like protein (909 aa) |
| 2680 | 3075 | PPTG_02163 | 56,29 | 2 | 1 | 1 | 2 | | | 5,10E+05 | 2 | 1 | 1 | 1 | 2 | Carbamidomethylation | 110936 | PPTG_02163 | Phytophthora parasitica INRA-310 anoctamin-like protein (956 aa) |
| 2939 | 3413 | PPTG_04446 | 56,28 | 15 | 0 | 15 | 15 | | 6,97E+05 | 1,17E+06 | 2 | 2 | 0 | 2 | 2 | Carbamidomethylation | 22273 | PPTG_04446 | Phytophthora parasitica INRA-310 hypothetical protein (198 aa) |
| 3155 | 3428 | PPTG_07064 | 56,27 | 4 | 0 | 4 | 2 | | 5,95E+05 | 0 | 2 | 2 | 0 | 2 | 1 | Oxidation (M); Carbamidomethylation | 58774 | PPTG_07064 | Phytophthora parasitica INRA-310 hypothetical protein (549 aa) |
| 3335 | 2891 | PPTG_15698 | 56,25 | 9 | 0 | 0 | 9 | | | 1,36E+06 | 2 | 2 | 0 | 0 | 2 | | 28768 | PPTG_15698 | Phytophthora parasitica INRA-310 nitroreductase family protein (264 aa) |
| 3040 | 3103 | PPTG_17512 | 56,24 | 3 | 0 | 2 | 3 | | 2,15E+05 | 4,75E+05 | 2 | 2 | 0 | 1 | 2 | Oxidation (M); Carbamidomethylation | 88080 | PPTG_17512 | Phytophthora parasitica INRA-310 hypothetical protein (802 aa) |
| 2592 | 2496 | PPTG_01027 | 56,16 | 3 | 1 | 1 | 3 | 4,90E+04 | 5,82E+04 | 8,48E+05 | 3 | 3 | 1 | 1 | 3 | | 108687 | PPTG_01027 | Phytophthora parasitica INRA-310 alpha alpha-trehalose-phosphate synthase [UDP-forming] (958 aa) |
| 2779 | 2977 | PPTG_13701 | 56,16 | 4 | 0 | 4 | 2 | | 1,43E+06 | 1,71E+06 | 3 | 3 | 0 | 3 | 1 | | 78703 | PPTG_13701 | Phytophthora parasitica INRA-310 hypothetical protein (699 aa) |
| 2306 | 2567 | PPTG_11943 | 56,11 | 2 | 0 | 2 | 1 | | 7,16E+05 | 3,59E+05 | 5 | 5 | 0 | 4 | 2 | | 271618 | PPTG_11943 | Phytophthora parasitica INRA-310 integrator complex subunit (2446 aa) |
| 2522 | 2591 | PPTG_03522 | 56,03 | 4 | 0 | 2 | 3 | | 6,59E+05 | 7,23E+05 | 4 | 4 | 0 | 2 | 3 | Oxidation (M) | 114572 | PPTG_03522 | Phytophthora parasitica INRA-310 hypothetical protein (1031 aa) |
| 2918 | 3497 | PPTG_11009 | 55,99 | 7 | 4 | 4 | 7 | 1,43E+05 | 5,33E+05 | 1,80E+06 | 2 | 2 | 1 | 1 | 2 | | 32192 | PPTG_11009 | Phytophthora parasitica INRA-310 phosphatidylinositol transfer protein SFH5 (288 aa) |
| 2613 | 3283 | PPTG_17473 | 55,91 | 14 | 6 | 14 | 14 | 2,30E+05 | 1,62E+06 | 1,28E+06 | 2 | 2 | 1 | 2 | 2 | Acetylation (N-term); Carbamylation | 23804 | PPTG_17473 | Phytophthora parasitica INRA-310 prefoldin subunit 3 (212 aa) |
| 2689 | 3170 | PPTG_11484 | 55,88 | 4 | 0 | 4 | 4 | | 2,48E+05 | 1,46E+06 | 2 | 2 | 0 | 2 | 2 | | 76188 | PPTG_11484 | Phytophthora parasitica INRA-310 leukotriene A-4 hydrolase (677 aa) |
| 3183 | 2694 | PPTG_20144 | 55,86 | 11 | 0 | 0 | 11 | | | 6,68E+05 | 3 | 1 | 0 | 0 | 3 | | 32969 | PPTG_20144 | Phytophthora parasitica INRA-310 dicarboxylate/Amino Acid:Cation (Na or H) Symporter (DAACS) family protein (308 aa) |
| 2716 | 2745 | PPTG_04737 | 55,82 | 7 | 0 | 7 | 3 | | 3,62E+05 | 2,09E+05 | 3 | 3 | 0 | 3 | 1 | | 45220 | PPTG_04737 | Phytophthora parasitica INRA-310 regulator-chromosome condensation (RCC1)-like protein (425 aa) |
| 2621 | 2624 | PPTG_14610 | 55,79 | 5 | 0 | 5 | 2 | | 9,52E+05 | 1,33E+06 | 3 | 3 | 0 | 3 | 2 | | 73480 | PPTG_14610 | Phytophthora parasitica INRA-310 hypothetical protein (659 aa) |
| 2717 | 2751 | PPTG_04972 | 55,79 | 1 | 0 | 1 | 0 | | 3,02E+05 | 2,65E+05 | 3 | 3 | 0 | 3 | 1 | | 283448 | PPTG_04972 | Phytophthora parasitica INRA-310 hypothetical protein (2599 aa) |
| 2447 | 2636 | PPTG_14323 | 55,75 | 3 | 0 | 2 | 2 | | 1,75E+05 | 9,19E+05 | 3 | 3 | 0 | 2 | 3 | | 140924 | PPTG_14323 | Phytophthora parasitica INRA-310 RNA polymerase II-associated protein 1 (1272 aa) |
| 2802 | 2698 | PPTG_12067 | 55,69 | 7 | 0 | 0 | 7 | | | 6,60E+05 | 4 | 4 | 0 | 0 | 4 | Oxidation (M) | 70804 | PPTG_12067 | Phytophthora parasitica INRA-310 protein kinase (622 aa) |
| 1918 | 2739 | PPTG_14380 | 55,62 | 5 | 2 | 2 | 5 | | | 8,24E+05 | 3 | 2 | 2 | 2 | 5 | | 50945 | PPTG_14380 | Phytophthora parasitica INRA-310 protein kinase (453 aa) |
| 2937 | 3395 | PPTG_14033 | 55,61 | 31 | 0 | 31 | 31 | | 7,56E+06 | 7,89E+06 | 2 | 2 | 0 | 2 | 2 | Carbamidomethylation; Acetylation (N-term) | 10772 | PPTG_14033 | Phytophthora parasitica INRA-310 small nuclear ribonucleoprotein F (99 aa) |
| 2989 | 3308 | PPTG_14745 | 55,57 | 6 | 0 | 6 | 4 | | 4,21E+05 | 4,42E+05 | 2 | 2 | 0 | 2 | 1 | | 52394 | PPTG_14745 | Phytophthora parasitica INRA-310 aldehyde dehydrogenase (486 aa) |
| 2169 | 2668 | PPTG_09403 | 55,53 | 5 | 1 | 0 | 5 | 3,19E+05 | | 8,51E+06 | 5 | 5 | 1 | 0 | 6 | | 126949 | PPTG_09403 | Phytophthora parasitica INRA-310 hypothetical protein (1130 aa) |
| 837 | 3473 | PPTG_02741 | 55,48 | 20 | 10 | 20 | 20 | | 0 | 3,05E+07 | 2 | 1 | 7 | 9 | 11 | Oxidation (M); Dehydration; Deamidation (NQ); Acetylation (N-term); Carbamylation | 13734 | PPTG_02741 | Phytophthora parasitica INRA-310 histone H4 (127 aa) |
| 3345 | 2744 | PPTG_06450 | 55,47 | 2 | 0 | 0 | 2 | | | 3,86E+05 | 2 | 2 | 0 | 0 | 2 | | 106351 | PPTG_06450 | Phytophthora parasitica INRA-310 chloride Channel (ClC) family (958 aa) |
| 2356 | 2823 | PPTG_04918 | 55,45 | 7 | 7 | 7 | 7 | 7,86E+05 | 2,79E+06 | 5,64E+06 | 2 | 2 | 2 | 2 | 2 | | 36084 | PPTG_04918 | Phytophthora parasitica INRA-310 alcohol dehydrogenase (333 aa) |
| 2505 | 2431 | PPTG_13605 | 55,41 | 20 | 0 | 0 | 20 | | | 2,11E+06 | 5 | 5 | 0 | 0 | 5 | Carbamidomethylation | 41525 | PPTG_13605 | Phytophthora parasitica INRA-310 hypothetical protein (380 aa) |
| 2982 | 2727 | PPTG_11151 | 55,3 | 5 | 0 | 0 | 5 | | | 1,73E+06 | 3 | 2 | 0 | 0 | 3 | | 69727 | PPTG_11151 | Phytophthora parasitica INRA-310 ATP-binding Cassette (ABC) superfamily (623 aa) |
| 2348 | 2449 | PPTG_07433 | 55,28 | 16 | 4 | 7 | 12 | 1,61E+05 | 7,72E+05 | 2,11E+06 | 4 | 4 | 1 | 2 | 3 | Carbamidomethylation | 40315 | PPTG_07433 | Phytophthora parasitica INRA-310 replication factor C subunit 3 (353 aa) |
| 2242 | 2532 | PPTG_02567 | 55,27 | 21 | 8 | 21 | 21 | 6,08E+04 | 1,39E+06 | 2,03E+06 | 3 | 3 | 1 | 3 | 3 | | 15821 | PPTG_02567 | Phytophthora parasitica INRA-310 hypothetical protein (144 aa) |
| 2886 | 3521 | PPTG_06213 | 55,26 | 7 | 3 | 7 | 3 | 1,08E+05 | 9,60E+05 | 1,26E+06 | 2 | 2 | 1 | 2 | 1 | | 44698 | PPTG_06213 | Phytophthora parasitica INRA-310 hypothetical protein (405 aa) |
| 2558 | 3249 | PPTG_00419 | 55,23 | 2 | 2 | 2 | 2 | 2,81E+04 | 7,80E+05 | 1,52E+06 | 2 | 2 | 1 | 2 | 2 | | 95868 | PPTG_00419 | Phytophthora parasitica INRA-310 histone deacetylase (863 aa) |
| 2484 | 2493 | PPTG_05697 | 55,22 | 8 | 0 | 0 | 8 | | | 1,91E+06 | 5 | 5 | 0 | 0 | 5 | | 82894 | PPTG_05697 | Phytophthora parasitica INRA-310 hypothetical protein (751 aa) |
| 3244 | 2674 | PPTG_17603 | 55,21 | 6 | 0 | 0 | 6 | | | 1,49E+06 | 3 | 3 | 0 | 0 | 3 | | 71751 | PPTG_17603 | Phytophthora parasitica INRA-310 chloride Channel (ClC) family (649 aa) |
| 3096 | 2810 | PPTG_12047 | 55,08 | 6 | 0 | 4 | 2 | | 1,98E+05 | 6,51E+04 | 3 | 2 | 0 | 2 | 1 | | 62232 | PPTG_12047 | Phytophthora parasitica INRA-310 DEAD/DEAH box RNA helicase (562 aa) |
| 2559 | 3400 | PPTG_13840 | 55,06 | 6 | 0 | 5 | 6 | | 1,49E+06 | 3,20E+06 | 3 | 3 | 0 | 2 | 3 | | 50283 | PPTG_13840 | Phytophthora parasitica INRA-310 serine/threonine protein phosphatase 2A (437 aa) |
| 2554 | 2864 | PPTG_03729 | 55 | 24 | 0 | 24 | 9 | | 1,62E+06 | 1,60E+06 | 3 | 3 | 0 | 3 | 2 | | 27454 | PPTG_03729 | Phytophthora parasitica INRA-310 hypothetical protein (261 aa) |
| 2471 | 2472 | PPTG_03840 | 54,97 | 2 | 0 | 1 | 2 | | 1,90E+05 | 3,52E+05 | 4 | 4 | 0 | 1 | 4 | | 183860 | PPTG_03840 | Phytophthora parasitica INRA-310 hypothetical protein (1652 aa) |
| 2263 | 2603 | PPTG_13393 | 54,97 | 10 | 4 | 10 | 10 | 6,43E+04 | 6,04E+05 | 1,43E+06 | 3 | 3 | 1 | 3 | 3 | Carbamidomethylation | 33839 | PPTG_13393 | Phytophthora parasitica INRA-310 phosphoglycerate mutase family protein (301 aa) |
| 1847 | 4220 | PPTG_10579 | 54,96 | 9 | 9 | 9 | 9 | 3,87E+05 | 0 | 1,43E+07 | 1 | 1 | 2 | 1 | 7 | Oxidation (M); Dehydration; Deamidation (NQ) | 22020 | PPTG_10579 | Phytophthora parasitica INRA-310 hypothetical protein (192 aa) |
| 3260 | 3538 | PPTG_09073 | 54,95 | 30 | 0 | 17 | 30 | | 1,62E+05 | 1,53E+06 | 2 | 2 | 0 | 1 | 2 | Carbamidomethylation | 12771 | PPTG_09073 | Phytophthora parasitica INRA-310 beta-elicitin cryptogein (121 aa) |
| 2412 | 2592 | PPTG_08536 | 54,91 | 32 | 0 | 32 | 32 | | 9,79E+05 | 7,44E+05 | 3 | 3 | 0 | 3 | 3 | Oxidation (M) | 14091 | PPTG_08536 | Phytophthora parasitica INRA-310 hypothetical protein (121 aa) |
| 2631 | 3405 | PPTG_11993 | 54,89 | 6 | 0 | 6 | 6 | | 2,30E+06 | 5,22E+05 | 2 | 2 | 0 | 3 | 2 | Deamidation (NQ) | 56354 | PPTG_11993 | Phytophthora parasitica INRA-310 ubiquitin family protein (542 aa) |
| 3098 | 2841 | PPTG_12889 | 54,86 | 6 | 0 | 3 | 3 | | 2,63E+05 | 1,72E+06 | 3 | 3 | 0 | 1 | 2 | Carbamidomethylation; Acetylation (N-term) | 61168 | PPTG_12889 | Phytophthora parasitica INRA-310 hypothetical protein (558 aa) |
| 2969 | 2895 | PPTG_05074 | 54,83 | 7 | 0 | 4 | 3 | | 3,45E+05 | 3,04E+05 | 3 | 3 | 0 | 2 | 1 | | 64148 | PPTG_05074 | Phytophthora parasitica INRA-310 polysaccharide lyase (587 aa) |
| 2975 | 2677 | PPTG_08805 | 54,79 | 5 | 0 | 0 | 5 | | | 1,26E+06 | 3 | 3 | 0 | 0 | 3 | | 89742 | PPTG_08805 | Phytophthora parasitica INRA-310 hypothetical protein (795 aa) |
| 2875 | 3275 | PPTG_03224 | 54,78 | 7 | 0 | 7 | 7 | | 3,64E+05 | 7,63E+05 | 2 | 2 | 0 | 2 | 2 | | 36924 | PPTG_03224 | Phytophthora parasitica INRA-310 chromatin modification-like protein EAF3 (324 aa) |
| 2623 | 2690 | PPTG_02159 | 54,77 | 34 | 0 | 34 | 26 | | 1,64E+06 | 0 | 3 | 3 | 0 | 3 | 2 | | 9573 | PPTG_02159 | Phytophthora parasitica INRA-310 hypothetical protein (83 aa) |
| 3044 | 3297 | PPTG_18586 | 54,75 | 4 | 0 | 2 | 4 | | 4,28E+05 | 1,30E+06 | 2 | 2 | 0 | 1 | 2 | | 56085 | PPTG_18586 | Phytophthora parasitica INRA-310 quinolinate synthetase A protein (518 aa) |
| 3359 | 3838 | PPTG_17893 | 54,72 | 4 | 0 | 4 | 4 | | 1,08E+05 | 6,10E+05 | 1 | 1 | 0 | 1 | 1 | | 52110 | PPTG_17893 | Phytophthora parasitica INRA-310 hypothetical protein (475 aa) |
| 3065 | 2970 | PPTG_08015 | 54,7 | 11 | 0 | 11 | 4 | | 7,91E+05 | 4,54E+05 | 2 | 2 | 0 | 2 | 1 | Carbamidomethylation | 35750 | PPTG_08015 | Phytophthora parasitica INRA-310 hydroxymethylglutaryl-CoA lyase (341 aa) |
| 2264 | 3469 | PPTG_11396 | 54,7 | 8 | 0 | 5 | 8 | | 2,31E+06 | 3,74E+06 | 2 | 2 | 0 | 3 | 4 | Oxidation (M); Carbamidomethylation | 20499 | PPTG_11396 | Phytophthora parasitica INRA-310 hypothetical protein (195 aa) |
| 2609 | 3063 | PPTG_04941 | 54,66 | 32 | 23 | 0 | 32 | 2,97E+05 | | 3,66E+06 | 2 | 2 | 1 | 0 | 4 | Acetylation (N-term) | 7852 | PPTG_04941 | Phytophthora parasitica INRA-310 protein translocase SEC61 complex gamma subunit (72 aa) |
| 2254 | 2571 | PPTG_15044 | 54,65 | 19 | 0 | 19 | 19 | | 9,46E+05 | 3,08E+06 | 3 | 3 | 0 | 3 | 4 | Oxidation (M); Carbamidomethylation | 18767 | PPTG_15044 | Phytophthora parasitica INRA-310 hypothetical protein (159 aa) |
| 3074 | 3404 | PPTG_08382 | 54,63 | 15 | 0 | 9 | 15 | | 0 | 1,24E+06 | 2 | 2 | 0 | 1 | 2 | | 16238 | PPTG_08382 | Phytophthora parasitica INRA-310 hypothetical protein (144 aa) |
| 2353 | 2721 | PPTG_10865 | 54,56 | 5 | 1 | 3 | 5 | 7,01E+04 | 3,40E+05 | 1,34E+06 | 3 | 3 | 1 | 2 | 3 | Carbamidomethylation | 66923 | PPTG_10865 | Phytophthora parasitica INRA-310 hypothetical protein (591 aa) |
| 3332 | 3873 | PPTG_19981 | 54,52 | 4 | 0 | 4 | 4 | | 6,36E+05 | 3,19E+05 | 1 | 1 | 0 | 1 | 1 | | 41633 | PPTG_19981 | Phytophthora parasitica INRA-310 elicitor-like transglutaminase (379 aa) |
| 2897 | 2582 | PPTG_08493 | 54,48 | 5 | 0 | 3 | 3 | | 6,74E+05 | 6,22E+05 | 3 | 3 | 0 | 2 | 2 | | 85806 | PPTG_08493 | Phytophthora parasitica INRA-310 glutamate-cysteine ligase catalytic subunit (766 aa) |
| 2884 | 3483 | PPTG_00685 | 54,45 | 6 | 6 | 3 | 3 | 5,50E+04 | | | 2 | 1 | 2 | 1 | 1 | | 39935 | PPTG_00685 | Phytophthora parasitica INRA-310 Rab11 family GTPase (365 aa) |
| 3134 | 2828 | PPTG_19693 | 54,42 | 10 | 0 | 0 | 10 | | | 1,33E+06 | 3 | 2 | 0 | 0 | 3 | | 31006 | PPTG_19693 | Phytophthora parasitica INRA-310 hypothetical protein (277 aa) |
| 3026 | 2593 | PPTG_19418 | 54,38 | 11 | 0 | 6 | 4 | | 0 | 8,40E+04 | 3 | 3 | 0 | 2 | 1 | | 32105 | PPTG_19418 | Phytophthora parasitica INRA-310 polysaccharide lyase (294 aa) |
| 2105 | 4228 | PPTG_19868 | 54,38 | 19 | 19 | 19 | 19 | 8,37E+05 | 1,10E+07 | 3,69E+05 | 1 | 1 | 2 | 4 | 2 | Carbamidomethylation | 19264 | PPTG_19868 | Phytophthora parasitica INRA-310 elicitin INF2A-like protein (195 aa) |
| 2676 | 3146 | PPTG_02664 | 54,37 | 4 | 0 | 4 | 4 | | 4,01E+05 | 4,34E+05 | 2 | 2 | 0 | 2 | 2 | Carbamidomethylation | 107174 | PPTG_02664 | Phytophthora parasitica INRA-310 hypothetical protein (970 aa) |
| 2572 | 2629 | PPTG_04269 | 54,29 | 10 | 0 | 10 | 2 | | 5,59E+05 | 2,10E+05 | 4 | 4 | 0 | 4 | 1 | Carbamidomethylation | 53212 | PPTG_04269 | Phytophthora parasitica INRA-310 serine protease family S10 (480 aa) |
| 3199 | 3440 | PPTG_10273 | 54,27 | 5 | 0 | 2 | 5 | | 1,90E+05 | 1,76E+06 | 2 | 2 | 0 | 1 | 2 | | 76865 | PPTG_10273 | Phytophthora parasitica INRA-310 hypothetical protein (681 aa) |
| 3256 | 3443 | PPTG_17486 | 54,26 | 24 | 0 | 14 | 24 | | 1,09E+05 | 4,44E+05 | 2 | 2 | 0 | 1 | 2 | Deamidation (NQ) | 11954 | PPTG_17486 | Phytophthora parasitica INRA-310 hypothetical protein (112 aa) |
| 2777 | 2854 | PPTG_09238 | 54,25 | 11 | 0 | 3 | 11 | | 5,13E+05 | 1,14E+06 | 3 | 3 | 0 | 1 | 3 | | 46591 | PPTG_09238 | Phytophthora parasitica INRA-310 hypothetical protein (410 aa) |
| 2781 | 3035 | PPTG_07487 | 54,24 | 4 | 2 | 4 | 2 | 0 | 1,05E+06 | 2,55E+06 | 2 | 2 | 1 | 2 | 1 | Oxidation (M) | 61526 | PPTG_07487 | Phytophthora parasitica INRA-310 zinc (Zn2)-Iron (Fe2) Permease (ZIP) family (579 aa) |
| 2852 | 2555 | PPTG_03539 | 54,2 | 10 | 0 | 7 | 5 | | 9,91E+05 | 9,57E+05 | 3 | 3 | 0 | 2 | 2 | | 42410 | PPTG_03539 | Phytophthora parasitica INRA-310 hypothetical protein (386 aa) |
| 3157 | 3498 | PPTG_17795 | 54,16 | 15 | 5 | 15 | 0 | | 3,10E+05 | | 2 | 1 | 1 | 2 | 0 | | 12942 | PPTG_17795 | Phytophthora parasitica INRA-310 hypothetical protein (129 aa) |
| 2790 | 3886 | PPTG_06563 | 54,13 | 8 | 0 | 8 | 8 | | 2,46E+05 | 6,65E+05 | 1 | 1 | 0 | 2 | 2 | | 26222 | PPTG_06563 | Phytophthora parasitica INRA-310 hypothetical protein (233 aa) |
| 2700 | 2997 | PPTG_16905 | 54,11 | 6 | 0 | 6 | 6 | | 1,81E+05 | 7,40E+05 | 2 | 2 | 0 | 2 | 2 | Carbamidomethylation | 41099 | PPTG_16905 | Phytophthora parasitica INRA-310 hypothetical protein (369 aa) |
| 2513 | 2930 | PPTG_09188 | 54,11 | 6 | 4 | 4 | 6 | 4,67E+05 | 6,24E+05 | 4,06E+06 | 2 | 2 | 1 | 1 | 3 | | 39479 | PPTG_09188 | Phytophthora parasitica INRA-310 major facilitator superfamily transporter (MFS) (355 aa) |
| 3028 | 2660 | PPTG_02613 | 54,11 | 10 | 0 | 0 | 10 | | | 9,30E+05 | 3 | 3 | 0 | 0 | 3 | Oxidation (M) | 44072 | PPTG_02613 | Phytophthora parasitica INRA-310 hypothetical protein (399 aa) |
| 2736 | 2434 | PPTG_01320 | 54,1 | 3 | 0 | 0 | 3 | | | 1,13E+06 | 4 | 4 | 0 | 0 | 4 | | 146644 | PPTG_01320 | Phytophthora parasitica INRA-310 hypothetical protein (1323 aa) |
| 2376 | 2759 | PPTG_18735 | 54,08 | 3 | 0 | 3 | 3 | | 6,23E+05 | 1,25E+06 | 3 | 3 | 0 | 3 | 3 | | 113225 | PPTG_18735 | Phytophthora parasitica INRA-310 hypothetical protein (1018 aa) |
| 2521 | 2512 | PPTG_14632 | 54,08 | 4 | 0 | 2 | 4 | | 5,95E+05 | 1,44E+06 | 3 | 3 | 0 | 2 | 3 | | 94740 | PPTG_14632 | Phytophthora parasitica INRA-310 cGMP-dependent protein kinase (847 aa) |
| 2743 | 2655 | PPTG_03888 | 54,08 | 9 | 0 | 9 | 2 | | 6,38E+05 | 5,33E+05 | 3 | 3 | 0 | 3 | 1 | | 47796 | PPTG_03888 | Phytophthora parasitica INRA-310 NPL4-like protein (429 aa) |
| 2806 | 2789 | PPTG_12277 | 54,06 | 7 | 0 | 2 | 7 | | 2,84E+05 | 3,19E+06 | 3 | 3 | 0 | 1 | 3 | | 48818 | PPTG_12277 | Phytophthora parasitica INRA-310 7-dehydrocholesterol reductase (436 aa) |
| 2627 | 3147 | PPTG_18715 | 54 | 24 | 13 | 24 | 24 | 3,48E+05 | 2,32E+06 | 2,62E+06 | 2 | 2 | 1 | 2 | 2 | | 10080 | PPTG_18715 | Phytophthora parasitica INRA-310 hypothetical protein (88 aa) |
| 2879 | 3310 | PPTG_07160 | 53,95 | 5 | 0 | 5 | 5 | | 1,88E+06 | 1,40E+06 | 2 | 2 | 0 | 2 | 2 | | 58779 | PPTG_07160 | Phytophthora parasitica INRA-310 glutamate carboxypeptidase (538 aa) |
| 2721 | 2859 | PPTG_18082 | 53,87 | 3 | 1 | 1 | 2 | 0 | 0 | | 3 | 1 | 1 | 1 | 2 | Carbamidomethylation; Acetylation (N-term) | 137368 | PPTG_18082 | Phytophthora parasitica INRA-310 cleavage induced protein kinase (1223 aa) |
| 3562 | 3476 | PPTG_08616 | 53,82 | 6 | 0 | 0 | 6 | | | 8,15E+05 | 2 | 2 | 0 | 0 | 2 | Carbamidomethylation | 53650 | PPTG_08616 | Phytophthora parasitica INRA-310 phosphatidylserine synthase (466 aa) |
| 3394 | 2950 | PPTG_04911 | 53,79 | 6 | 0 | 0 | 6 | | | 1,19E+06 | 2 | 2 | 0 | 0 | 2 | Carbamidomethylation | 47490 | PPTG_04911 | Phytophthora parasitica INRA-310 hypothetical protein (420 aa) |
| 2946 | 3585 | PPTG_01270 | 53,79 | 6 | 0 | 6 | 6 | | 5,52E+05 | 6,01E+05 | 2 | 2 | 0 | 2 | 2 | Carbamidomethylation | 65708 | PPTG_01270 | Phytophthora parasitica INRA-310 hypothetical protein (624 aa) |
| 3510 | 3758 | PPTG_13326 | 53,79 | 10 | 0 | 10 | 10 | | 1,11E+05 | 0 | 1 | 1 | 0 | 1 | 1 | Carbamidomethylation | 19838 | PPTG_13326 | Phytophthora parasitica INRA-310 hypothetical protein (178 aa) |
| 3510 | 3986 | PPTG_13326 | 53,79 | 11 | 0 | 11 | 11 | | 1,11E+05 | 0 | 1 | 1 | 0 | 1 | 1 | Carbamidomethylation | 18998 | PPTG_13326 | Phytophthora parasitica INRA-310 hypothetical protein (170 aa) |
| 3113 | 3277 | PPTG_03963 | 53,68 | 4 | 0 | 4 | 2 | | 6,58E+05 | 1,94E+05 | 2 | 2 | 0 | 2 | 1 | | 58379 | PPTG_03963 | Phytophthora parasitica INRA-310 hypothetical protein (539 aa) |
| 2751 | 2938 | PPTG_15112 | 53,67 | 5 | 3 | 5 | 2 | 4,68E+04 | 9,93E+05 | 3,43E+05 | 2 | 2 | 1 | 2 | 1 | Carbamidomethylation | 46319 | PPTG_15112 | Phytophthora parasitica INRA-310 hypothetical protein (438 aa) |
| 2194 | 2534 | PPTG_01978 | 53,65 | 8 | 5 | 3 | 8 | 1,67E+05 | | 1,61E+06 | 3 | 2 | 2 | 2 | 3 | | 35481 | PPTG_01978 | Phytophthora parasitica INRA-310 cyclin-dependent kinase 1 (310 aa) |
| 3175 | 4108 | PPTG_05277 | 53,65 | 8 | 0 | 7 | 8 | | 3,19E+05 | 4,96E+05 | 2 | 2 | 0 | 1 | 2 | | 24189 | PPTG_05277 | Phytophthora parasitica INRA-310 hypothetical protein (209 aa) |
| 2551 | 2771 | PPTG_16540 | 53,6 | 4 | 0 | 3 | 4 | | 8,65E+05 | 1,95E+06 | 3 | 3 | 0 | 2 | 3 | | 114118 | PPTG_16540 | Phytophthora parasitica INRA-310 5'-3' exoribonuclease (1012 aa) |
| 2549 | 2722 | PPTG_02458 | 53,59 | 5 | 0 | 3 | 5 | | 1,08E+06 | 2,07E+06 | 3 | 3 | 0 | 2 | 3 | | 81587 | PPTG_02458 | Phytophthora parasitica INRA-310 hypothetical protein (708 aa) |
| 3553 | 3287 | PPTG_15969 | 53,57 | 5 | 0 | 0 | 5 | | | 8,36E+05 | 2 | 2 | 0 | 0 | 2 | | 50479 | PPTG_15969 | Phytophthora parasitica INRA-310 hypothetical protein (457 aa) |
| 3059 | 2804 | PPTG_00297 | 53,53 | 31 | 9 | 0 | 21 | 6,65E+04 | | 1,42E+05 | 3 | 3 | 1 | 0 | 2 | | 12271 | PPTG_00297 | Phytophthora parasitica INRA-310 hypothetical protein (108 aa) |
| 2479 | 3082 | PPTG_13590 | 53,5 | 11 | 7 | 11 | 7 | 2,83E+04 | 5,54E+05 | 5,38E+05 | 2 | 2 | 1 | 2 | 2 | Carbamidomethylation | 25229 | PPTG_13590 | Phytophthora parasitica INRA-310 hypothetical protein (219 aa) |
| 2479 | 3083 | PPTG_13590 | 53,5 | 11 | 7 | 11 | 7 | 2,83E+04 | 5,54E+05 | 5,38E+05 | 2 | 2 | 1 | 2 | 2 | Carbamidomethylation | 25229 | PPTG_13590 | Phytophthora parasitica INRA-310 hypothetical protein (219 aa) |
| 2698 | 2890 | PPTG_11555 | 53,43 | 7 | 0 | 7 | 7 | | 8,22E+05 | 8,29E+05 | 2 | 2 | 0 | 2 | 2 | Carbamidomethylation | 36629 | PPTG_11555 | Phytophthora parasitica INRA-310 hypothetical protein (325 aa) |
| 3173 | 4001 | PPTG_16297 | 53,41 | 17 | 17 | 17 | 17 | 3,33E+04 | 4,08E+05 | 3,78E+05 | 1 | 1 | 1 | 1 | 1 | | 10176 | PPTG_16297 | Phytophthora parasitica INRA-310 hypothetical protein (90 aa) |
| 2507 | 2584 | PPTG_06727 | 53,39 | 4 | 0 | 3 | 4 | | 4,14E+05 | 9,58E+05 | 3 | 2 | 0 | 2 | 3 | | 82870 | PPTG_06727 | Phytophthora parasitica INRA-310 phosphate acetyltransferase (760 aa) |
| 3340 | 3154 | PPTG_01313 | 53,35 | 4 | 0 | 0 | 4 | | | 1,26E+06 | 2 | 2 | 0 | 0 | 2 | | 59889 | PPTG_01313 | Phytophthora parasitica INRA-310 cytochrome P450 (541 aa) |
| 3014 | 2941 | PPTG_09296 | 53,33 | 3 | 0 | 3 | 1 | | 6,21E+05 | 1,94E+05 | 2 | 2 | 0 | 2 | 1 | Carbamidomethylation | 67829 | PPTG_09296 | Phytophthora parasitica INRA-310 hypothetical protein (626 aa) |
| 2740 | 2632 | PPTG_01396 | 53,26 | 3 | 0 | 2 | 2 | | 2,08E+05 | 4,37E+05 | 3 | 3 | 0 | 2 | 2 | Carbamidomethylation | 127732 | PPTG_01396 | Phytophthora parasitica INRA-310 DEAD/DEAH box RNA helicase (1156 aa) |
| 3075 | 3467 | PPTG_06666 | 53,23 | 3 | 0 | 2 | 3 | | 1,62E+05 | 4,66E+05 | 2 | 2 | 0 | 1 | 2 | | 101977 | PPTG_06666 | Phytophthora parasitica INRA-310 IQ and AAA domain-containing protein (923 aa) |

| | | | | | | | | | | | | | | | | | | | |
|---|---|---|---|---|---|---|---|---|---|---|---|---|---|---|---|---|---|---|---|
| 2372 | 2595 | PPTG_12233 | 53,22 | 3 | 0 | 3 | 3 | | 7,44E+05 | 1,52E+06 | 3 | 3 | 0 | 3 | 3 | | 100116 | PPTG_12233 | Phytophthora parasitica INRA-310 translation initiation factor IF-2 (917 aa) |
| 2812 | 2871 | PPTG_02799 | 53,17 | 4 | 0 | 4 | 4 | | 5,15E+05 | 9,25E+05 | 2 | 2 | 0 | 2 | 2 | | 63623 | PPTG_02799 | Phytophthora parasitica INRA-310 hypothetical protein (559 aa) |
| 3068 | 3188 | PPTG_04099 | 53,12 | 4 | 0 | 2 | 4 | | 5,69E+05 | 6,07E+05 | 2 | 2 | 0 | 1 | 2 | | 65164 | PPTG_04099 | Phytophthora parasitica INRA-310 hypothetical protein (614 aa) |
| 4603 | 4221 | PPTG_13339 | 53,12 | 19 | 0 | 0 | 19 | | | 2,12E+05 | 1 | 1 | 0 | 0 | 1 | Carbamidomethylation | 7784 | PPTG_13339 | Phytophthora parasitica INRA-310 hypothetical protein (75 aa) |
| 2977 | 2835 | PPTG_07381 | 53,11 | 3 | 1 | 0 | 2 | 7,35E+04 | | 1,38E+05 | 3 | 3 | 1 | 0 | 2 | | 105524 | PPTG_07381 | Phytophthora parasitica INRA-310 ATP-binding Cassette (ABC) superfamily (987 aa) |
| 2868 | 3174 | PPTG_13909 | 53,1 | 9 | 0 | 4 | 9 | | 5,68E+05 | 1,16E+06 | 2 | 2 | 0 | 1 | 3 | | 29876 | PPTG_13909 | Phytophthora parasitica INRA-310 hypothetical protein (270 aa) |
| 3182 | 2689 | PPTG_05407 | 53,1 | 8 | 0 | 0 | 8 | | | 1,92E+06 | 3 | 3 | 0 | 0 | 3 | | 47310 | PPTG_05407 | Phytophthora parasitica INRA-310 hypothetical protein (423 aa) |
| 2915 | 3422 | PPTG_05172 | 53,09 | 27 | 0 | 27 | 9 | | 1,50E+06 | 0 | 2 | 2 | 0 | 3 | 1 | | 13795 | PPTG_05172 | Phytophthora parasitica INRA-310 hypothetical protein (126 aa) |
| 3069 | 3197 | PPTG_13665 | 53,09 | 5 | 0 | 5 | 3 | | 5,38E+05 | 4,38E+05 | 2 | 2 | 0 | 2 | 1 | | 53284 | PPTG_13665 | Phytophthora parasitica INRA-310 hypothetical protein (470 aa) |
| 2906 | 3107 | PPTG_04395 | 53,08 | 5 | 0 | 5 | 5 | | 1,83E+06 | 1,45E+06 | 2 | 2 | 0 | 2 | 2 | Carbamidomethylation | 63294 | PPTG_04395 | Phytophthora parasitica INRA-310 serine protease family S33 (578 aa) |
| 2523 | 2611 | PPTG_05554 | 53,04 | 15 | 0 | 2 | 14 | | 3,20E+05 | 1,11E+06 | 5 | 5 | 0 | 1 | 4 | Oxidation (M) | 50764 | PPTG_05554 | Phytophthora parasitica INRA-310 hypothetical protein (481 aa) |
| 2773 | 2728 | PPTG_06062 | 53,04 | 5 | 0 | 0 | 5 | | | 7,29E+05 | 4 | 4 | 0 | 0 | 4 | Carbamidomethylation | 73493 | PPTG_06062 | Phytophthora parasitica INRA-310 hypothetical protein (658 aa) |
| 2498 | 2714 | PPTG_07157 | 53,03 | 5 | 0 | 3 | 5 | | 7,28E+05 | 1,32E+06 | 3 | 3 | 0 | 2 | 3 | Oxidation (M); Carbamidomethylation | 69802 | PPTG_07157 | Phytophthora parasitica INRA-310 crinkler (CRN) family protein (618 aa) |
| 2816 | 3094 | PPTG_13443 | 53 | 14 | 0 | 14 | 14 | | 1,10E+06 | 1,12E+06 | 2 | 2 | 0 | 2 | 2 | | 20941 | PPTG_13443 | Phytophthora parasitica INRA-310 Ham1 family protein (192 aa) |
| 2619 | 2442 | PPTG_12515 | 52,97 | 10 | 3 | 3 | 10 | 1,68E+05 | 2,72E+05 | 3,74E+06 | 3 | 3 | 1 | 1 | 3 | Oxidation (M) | 28929 | PPTG_12515 | Phytophthora parasitica INRA-310 PpiC-type peptidyl-prolyl cis-trans isomerase (266 aa) |
| 2524 | 2617 | PPTG_00861 | 52,93 | 3 | 1 | 2 | 2 | 1,06E+05 | 8,23E+05 | 8,95E+05 | 3 | 3 | 1 | 2 | 2 | | 114124 | PPTG_00861 | Phytophthora parasitica INRA-310 hypothetical protein (1027 aa) |
| 2596 | 2643 | PPTG_08093 | 52,92 | 3 | 0 | 3 | 2 | | 9,09E+05 | 8,38E+05 | 3 | 3 | 0 | 3 | 2 | | 122718 | PPTG_08093 | Phytophthora parasitica INRA-310 hypothetical protein (1178 aa) |
| 3202 | 3466 | PPTG_02663 | 52,87 | 3 | 0 | 3 | 1 | | 1,48E+06 | 7,37E+05 | 2 | 2 | 0 | 2 | 1 | | 69704 | PPTG_02663 | Phytophthora parasitica INRA-310 polycomb protein (612 aa) |
| 3542 | 3006 | PPTG_03894 | 52,87 | 3 | 0 | 0 | 3 | | | 9,46E+05 | 2 | 2 | 0 | 0 | 2 | | 100998 | PPTG_03894 | Phytophthora parasitica INRA-310 calmodulin-like protein (920 aa) |
| 4054 | 3872 | PPTG_13299 | 52,85 | 6 | 0 | 0 | 6 | | | 7,23E+05 | 1 | 1 | 0 | 0 | 1 | | 39913 | PPTG_13299 | Phytophthora parasitica INRA-310 drug/Metabolite transporter (DMT) superfamily (366 aa) |
| 2890 | 3944 | PPTG_02032 | 52,8 | 8 | 0 | 8 | 8 | | 1,36E+06 | 3,48E+06 | 1 | 1 | 0 | 2 | 2 | | 19458 | PPTG_02032 | Phytophthora parasitica INRA-310 hypothetical protein (174 aa) |
| 2835 | 3598 | PPTG_06081 | 52,78 | 7 | 3 | 7 | 3 | 1,00E+05 | 7,32E+05 | 7,06E+05 | 2 | 2 | 1 | 2 | 1 | | 42321 | PPTG_06081 | Phytophthora parasitica INRA-310 hypothetical protein (377 aa) |
| 2783 | 3139 | PPTG_18297 | 52,78 | 12 | 6 | 6 | 12 | 2,90E+05 | 5,10E+05 | 2,86E+06 | 2 | 2 | 1 | 1 | 2 | Carbamidomethylation | 21320 | PPTG_18297 | Phytophthora parasitica INRA-310 hypothetical protein (200 aa) |
| 3071 | 3263 | PPTG_16897 | 52,78 | 3 | 0 | 3 | 1 | | 4,99E+05 | 1,24E+05 | 2 | 2 | 0 | 2 | 1 | | 76134 | PPTG_16897 | Phytophthora parasitica INRA-310 pescadillo-like protein (662 aa) |
| 2602 | 2743 | PPTG_01522 | 52,71 | 4 | 1 | 3 | 2 | | 4,33E+05 | 1,08E+06 | 3 | 2 | 1 | 2 | 2 | Carbamidomethylation | 77320 | PPTG_01522 | Phytophthora parasitica INRA-310 mitogen-activated protein kinase (708 aa) |
| 2663 | 2838 | PPTG_02081 | 52,71 | 6 | 0 | 4 | 5 | | 3,08E+05 | 6,66E+04 | 3 | 3 | 0 | 2 | 2 | | 85835 | PPTG_02081 | Phytophthora parasitica INRA-310 adenylate kinase (781 aa) |
| 2610 | 3085 | PPTG_09907 | 52,66 | 7 | 3 | 7 | 7 | 6,11E+04 | 3,06E+05 | 5,59E+05 | 2 | 2 | 1 | 2 | 2 | | 38813 | PPTG_09907 | Phytophthora parasitica INRA-310 S-methyl-5-thioribose-1-phosphate isomerase (363 aa) |
| 3275 | 4223 | PPTG_02433 | 52,64 | 16 | 0 | 16 | 16 | | 1,40E+05 | 6,34E+05 | 1 | 1 | 0 | 2 | 1 | Carbamidomethylation | 15936 | PPTG_02433 | Phytophthora parasitica INRA-310 hypothetical protein (154 aa) |
| 2898 | 2656 | PPTG_17784 | 52,6 | 6 | 0 | 6 | 1 | | 1,18E+06 | 6,98E+05 | 3 | 3 | 0 | 3 | 1 | | 64694 | PPTG_17784 | Phytophthora parasitica INRA-310 hypothetical protein (606 aa) |
| 2633 | 3420 | PPTG_07559 | 52,58 | 29 | 17 | 17 | 29 | 0 | 5,74E+05 | 2,51E+06 | 2 | 2 | 1 | 2 | 2 | Dehydration; Deamidation (NQ) | 10593 | PPTG_07559 | Phytophthora parasitica INRA-310 hypothetical protein (94 aa) |
| 2633 | 3421 | PPTG_17767 | 52,58 | 23 | 14 | 14 | 23 | 0 | 5,74E+05 | 2,51E+06 | 2 | 2 | 1 | 2 | 2 | Dehydration; Deamidation (NQ) | 12924 | PPTG_17767 | Phytophthora parasitica INRA-310 hypothetical protein (116 aa) |
| 2719 | 2817 | PPTG_06942 | 52,57 | 23 | 6 | 6 | 17 | 1,29E+05 | 3,08E+05 | 2,60E+06 | 3 | 3 | 1 | 1 | 2 | | 17888 | PPTG_06942 | Phytophthora parasitica INRA-310 hypothetical protein (162 aa) |
| 3142 | 3119 | PPTG_19099 | 52,56 | 23 | 0 | 9 | 23 | | 1,03E+06 | 1,65E+06 | 2 | 2 | 0 | 1 | 2 | | 14288 | PPTG_19099 | Phytophthora parasitica INRA-310 hypothetical protein (128 aa) |
| 3783 | 4224 | PPTG_15905 | 52,55 | 10 | 0 | 10 | 10 | | 4,60E+05 | 2,60E+06 | 1 | 1 | 0 | 1 | 1 | | 23061 | PPTG_15905 | Phytophthora parasitica INRA-310 hypothetical protein (208 aa) |
| 3214 | 3639 | PPTG_06231 | 52,52 | 11 | 0 | 6 | 4 | | 1,98E+05 | 4,91E+05 | 2 | 2 | 0 | 2 | 1 | | 38247 | PPTG_06231 | Phytophthora parasitica INRA-310 pectinesterase (349 aa) |
| 3253 | 3438 | PPTG_00033 | 52,48 | 22 | 0 | 12 | 22 | | 6,93E+05 | 1,32E+06 | 2 | 2 | 0 | 1 | 2 | | 11912 | PPTG_00033 | Phytophthora parasitica INRA-310 hypothetical protein (103 aa) |
| 3775 | 3667 | PPTG_11881 | 52,48 | 10 | 0 | 10 | 0 | | 1,40E+05 | | 2 | 2 | 0 | 2 | 0 | | 35753 | PPTG_11881 | Phytophthora parasitica INRA-310 hypothetical protein (314 aa) |
| 3057 | 2701 | PPTG_06380 | 52,39 | 8 | 0 | 0 | 8 | | | 4,03E+05 | 3 | 3 | 0 | 0 | 3 | | 47394 | PPTG_06380 | Phytophthora parasitica INRA-310 hypothetical protein (414 aa) |
| 2809 | 2845 | PPTG_13350 | 52,34 | 9 | 0 | 6 | 7 | | 4,11E+05 | 1,31E+06 | 3 | 3 | 0 | 2 | 2 | | 40388 | PPTG_13350 | Phytophthora parasitica INRA-310 S-adenosyl-L-methionine-dependent methyltransferase (359 aa) |
| 3602 | 4096 | PPTG_10185 | 52,34 | 4 | 4 | 0 | 4 | 4,77E+04 | | 2,95E+05 | 1 | 1 | 1 | 0 | 1 | | 40318 | PPTG_10185 | Phytophthora parasitica INRA-310 cation Diffusion Facilitator (CDF) family (364 aa) |
| 2801 | 2687 | PPTG_11135 | 52,25 | 8 | 0 | 6 | 5 | | 5,38E+05 | 8,02E+05 | 3 | 3 | 0 | 2 | 2 | Carbamidomethylation | 59973 | PPTG_11135 | Phytophthora parasitica INRA-310 serine protease family S10 (547 aa) |
| 2442 | 2832 | PPTG_15883 | 52,22 | 11 | 8 | 8 | 7 | 1,47E+05 | 5,00E+05 | 2,38E+06 | 3 | 2 | 2 | 2 | 2 | Carbamidomethylation | 38048 | PPTG_15883 | Phytophthora parasitica INRA-310 ATP-binding Cassette (ABC) superfamily protein (334 aa) |
| 3641 | 3186 | PPTG_05591 | 52,2 | 21 | 0 | 0 | 21 | | | 8,86E+05 | 2 | 2 | 0 | 0 | 2 | Oxidation (M) | 19488 | PPTG_05591 | Phytophthora parasitica INRA-310 hypothetical protein (180 aa) |
| 2604 | 2795 | PPTG_01273 | 52,15 | 33 | 0 | 33 | 24 | | 1,79E+06 | 2,31E+06 | 3 | 3 | 0 | 3 | 2 | Carbamidomethylation | 9214 | PPTG_01273 | Phytophthora parasitica INRA-310 U6 snRNA-associated Sm-like protein LSm6 (84 aa) |
| 3012 | 2725 | PPTG_16534 | 52,13 | 6 | 0 | 0 | 6 | | | 8,56E+05 | 3 | 3 | 0 | 0 | 3 | Carbamidomethylation | 66707 | PPTG_16534 | Phytophthora parasitica INRA-310 glucosidase 2 subunit beta (594 aa) |
| 3012 | 2726 | PPTG_16534 | 52,13 | 6 | 0 | 0 | 6 | | | 8,56E+05 | 3 | 3 | 0 | 0 | 3 | Carbamidomethylation | 66821 | PPTG_16534 | Phytophthora parasitica INRA-310 glucosidase 2 subunit beta (595 aa) |
| 2775 | 2843 | PPTG_13243 | 52,13 | 9 | 0 | 2 | 9 | | 2,68E+06 | 1,53E+07 | 3 | 3 | 0 | 1 | 3 | Oxidation (M) | 45226 | PPTG_13243 | Phytophthora parasitica INRA-310 hypothetical protein (404 aa) |
| 2929 | 4226 | PPTG_03241 | 52,08 | 15 | 0 | 0 | 15 | | | 7,85E+06 | 1 | 1 | 0 | 0 | 4 | Oxidation (M); Deamidation (NQ) | 18393 | PPTG_03241 | Phytophthora parasitica INRA-310 hypothetical protein (172 aa) |
| 2606 | 2893 | PPTG_11711 | 52,05 | 15 | 0 | 15 | 0 | | 1,15E+06 | | 5 | 5 | 0 | 5 | 0 | | 48709 | PPTG_11711 | Phytophthora parasitica INRA-310 hypothetical protein (442 aa) |
| 3094 | 2753 | PPTG_01602 | 52,04 | 16 | 0 | 0 | 16 | | | 2,94E+06 | 3 | 3 | 0 | 0 | 3 | | 24581 | PPTG_01602 | Phytophthora parasitica INRA-310 transmembrane protein (230 aa) |
| 2079 | 3448 | PPTG_15169 | 52 | 6 | 0 | 6 | 6 | | 3,70E+05 | 7,56E+06 | 2 | 1 | 0 | 3 | 5 | | 39400 | PPTG_15169 | Phytophthora parasitica INRA-310 polygalacturonase (381 aa) |
| 2770 | 2598 | PPTG_16726 | 51,95 | 9 | 0 | 0 | 9 | | | 3,00E+06 | 4 | 4 | 0 | 0 | 4 | | 50192 | PPTG_16726 | Phytophthora parasitica INRA-310 hypothetical protein (456 aa) |
| 3485 | 3317 | PPTG_11834 | 51,91 | 2 | 0 | 1 | 1 | | 2,41E+05 | 1,99E+05 | 2 | 2 | 0 | 1 | 1 | | 114706 | PPTG_11834 | Phytophthora parasitica INRA-310 exportin-T (1031 aa) |
| 3018 | 3114 | PPTG_09983 | 51,86 | 1 | 0 | 1 | 1 | | 9,32E+04 | 5,11E+05 | 2 | 2 | 0 | 1 | 2 | | 186176 | PPTG_09983 | Phytophthora parasitica INRA-310 hypothetical protein (1652 aa) |
| 3104 | 3015 | PPTG_05780 | 51,81 | 4 | 2 | 0 | 4 | 2,12E+05 | | 2,16E+06 | 2 | 2 | 1 | 0 | 2 | | 66321 | PPTG_05780 | Phytophthora parasitica INRA-310 alpha mannosyltransferase (588 aa) |
| 3201 | 3462 | PPTG_00842 | 51,8 | 8 | 0 | 4 | 8 | | 6,52E+05 | 1,61E+06 | 2 | 2 | 0 | 1 | 2 | | 35244 | PPTG_00842 | Phytophthora parasitica INRA-310 hydroxyacylglutathione hydrolase (316 aa) |
| 2355 | 2805 | PPTG_16683 | 51,76 | 18 | 0 | 18 | 2 | | 5,43E+04 | | 5 | 1 | 0 | 5 | 1 | Carbamidomethylation | 43326 | PPTG_16683 | Phytophthora parasitica INRA-310 alcohol dehydrogenase (408 aa) |
| 3549 | 3122 | PPTG_17981 | 51,71 | 3 | 0 | 0 | 3 | | | 0 | 2 | 2 | 0 | 0 | 2 | | 75504 | PPTG_17981 | Phytophthora parasitica INRA-310 hypothetical protein (669 aa) |
| 2580 | 2792 | PPTG_12039 | 51,67 | 6 | 0 | 5 | 6 | | 2,73E+05 | 1,24E+06 | 3 | 3 | 0 | 2 | 3 | Carbamidomethylation | 57133 | PPTG_12039 | Phytophthora parasitica INRA-310 sporangia induced Trp protein (510 aa) |
| 3637 | 2952 | PPTG_08496 | 51,65 | 6 | 0 | 0 | 6 | | | 5,19E+05 | 2 | 2 | 0 | 0 | 2 | | 65340 | PPTG_08496 | Phytophthora parasitica INRA-310 dicarboxylate/Amino Acid:Cation (Na or H) Symporter (DAACS) family (614 aa) |
| 2738 | 2623 | PPTG_08410 | 51,64 | 12 | 0 | 2 | 10 | | 1,51E+05 | 4,61E+05 | 4 | 4 | 0 | 1 | 3 | | 40390 | PPTG_08410 | Phytophthora parasitica INRA-310 cytosolic 5'-nucleotidase III-like protein (359 aa) |
| 2889 | 3716 | PPTG_09025 | 51,63 | 5 | 2 | 5 | 2 | 4,03E+04 | 2,50E+05 | 5,98E+05 | 2 | 2 | 1 | 2 | 1 | | 52202 | PPTG_09025 | Phytophthora parasitica INRA-310 hypothetical protein (466 aa) |
| 3482 | 3279 | PPTG_08082 | 51,59 | 5 | 0 | 0 | 5 | | | 2,63E+05 | 2 | 2 | 0 | 0 | 2 | | 55960 | PPTG_08082 | Phytophthora parasitica INRA-310 serine protease family S09X (509 aa) |
| 3650 | 3425 | PPTG_14026 | 51,58 | 14 | 0 | 0 | 14 | | | 4,17E+05 | 2 | 2 | 0 | 0 | 2 | | 18927 | PPTG_14026 | Phytophthora parasitica INRA-310 hypothetical protein (169 aa) |
| 3097 | 2812 | PPTG_16851 | 51,56 | 13 | 0 | 7 | 13 | | 9,10E+04 | 4,60E+05 | 2 | 2 | 0 | 1 | 2 | | 27489 | PPTG_16851 | Phytophthora parasitica INRA-310 NPP1-like protein (250 aa) |
| 2942 | 3456 | PPTG_16021 | 51,55 | 4 | 2 | 2 | 4 | 1,65E+05 | 3,26E+05 | 1,22E+06 | 2 | 2 | 1 | 1 | 2 | | 46233 | PPTG_16021 | Phytophthora parasitica INRA-310 hypothetical protein (416 aa) |
| 2226 | 2580 | PPTG_09676 | 51,53 | 12 | 8 | 4 | 12 | 2,08E+05 | 1,36E+05 | 1,18E+06 | 3 | 3 | 2 | 1 | 4 | | 26426 | PPTG_09676 | Phytophthora parasitica INRA-310 riboflavin synthase alpha subunit (244 aa) |
| 2826 | 3299 | PPTG_01478 | 51,47 | 15 | 0 | 4 | 15 | | 3,06E+06 | 3,84E+06 | 2 | 2 | 0 | 2 | 2 | | 27066 | PPTG_01478 | Phytophthora parasitica INRA-310 hypothetical protein (236 aa) |
| 2815 | 3065 | PPTG_15697 | 51,47 | 11 | 0 | 11 | 11 | | 4,72E+05 | 1,55E+06 | 2 | 2 | 0 | 2 | 2 | Oxidation (M) | 26204 | PPTG_15697 | Phytophthora parasitica INRA-310 nitroreductase family protein (239 aa) |
| 3662 | 3583 | PPTG_12703 | 51,47 | 5 | 2 | 0 | 3 | 1,08E+05 | | 3,26E+05 | 2 | 2 | 1 | 0 | 1 | | 55719 | PPTG_12703 | Phytophthora parasitica INRA-310 glycoside-Pentoside-Hexuronide (GPH):Cation Symporter family (512 aa) |
| 3784 | 4233 | PPTG_05823 | 51,46 | 4 | 4 | 4 | 0 | 5,25E+04 | 3,07E+05 | | 1 | 1 | 1 | 1 | 0 | | 26977 | PPTG_05823 | Phytophthora parasitica INRA-310 hypothetical protein (249 aa) |
| 3191 | 3250 | PPTG_09102 | 51,39 | 31 | 0 | 20 | 31 | | 1,43E+05 | 2,23E+05 | 2 | 2 | 0 | 1 | 2 | Carbamidomethylation | 10711 | PPTG_09102 | Phytophthora parasitica INRA-310 hypothetical protein (96 aa) |
| 3393 | 2920 | PPTG_02502 | 51,33 | 6 | 0 | 0 | 6 | | | 4,30E+05 | 2 | 2 | 0 | 0 | 2 | Carbamidomethylation | 47929 | PPTG_02502 | Phytophthora parasitica INRA-310 P-3 complex subunit mu (426 aa) |
| 2322 | 2662 | PPTG_02827 | 51,29 | 2 | 1 | 2 | 2 | 1,02E+05 | 5,98E+05 | 1,50E+06 | 3 | 3 | 1 | 2 | 3 | | 114237 | PPTG_02827 | Phytophthora parasitica INRA-310 vacuolar protein sorting-associated protein (1025 aa) |
| 2712 | 2586 | PPTG_19327 | 51,28 | 4 | 0 | 2 | 3 | | 9,01E+05 | 1,68E+06 | 3 | 3 | 0 | 2 | 2 | | 83901 | PPTG_19327 | Phytophthora parasitica INRA-310 hypothetical protein (758 aa) |
| 3029 | 2678 | PPTG_07162 | 51,26 | 9 | 0 | 6 | 3 | | 1,04E+06 | 0 | 3 | 3 | 0 | 2 | 1 | | 34871 | PPTG_07162 | Phytophthora parasitica INRA-310 carbohydrate esterase (333 aa) |
| 3255 | 3442 | PPTG_07948 | 51,25 | 3 | 0 | 3 | 2 | | 8,14E+04 | 0 | 2 | 2 | 0 | 2 | 1 | | 109501 | PPTG_07948 | Phytophthora parasitica INRA-310 hypothetical protein (995 aa) |
| 2557 | 3064 | PPTG_02060 | 51,21 | 4 | 2 | 4 | 4 | 6,65E+04 | 3,38E+05 | 9,05E+05 | 2 | 2 | 1 | 2 | 2 | | 54617 | PPTG_02060 | Phytophthora parasitica INRA-310 hypothetical protein (514 aa) |
| 2856 | 2758 | PPTG_13048 | 51,19 | 3 | 0 | 1 | 3 | | 4,91E+05 | 1,29E+06 | 3 | 3 | 0 | 1 | 3 | Oxidation (M) | 101677 | PPTG_13048 | Phytophthora parasitica INRA-310 eukaryotic translation initiation factor (945 aa) |
| 3172 | 3908 | PPTG_01542 | 51,19 | 28 | 28 | 28 | 28 | 3,13E+05 | 3,74E+05 | 8,48E+05 | 1 | 1 | 1 | 1 | 1 | Carbamidomethylation | 10379 | PPTG_01542 | Phytophthora parasitica INRA-310 hypothetical protein (101 aa) |
| 3171 | 3896 | PPTG_11123 | 51,11 | 5 | 0 | 5 | 5 | | 1,06E+05 | 2,25E+05 | 1 | 1 | 0 | 1 | 2 | | 35415 | PPTG_11123 | Phytophthora parasitica INRA-310 retinol dehydrogenase (330 aa) |
| 3554 | 3301 | PPTG_11625 | 51,11 | 14 | 0 | 6 | 8 | | 1,92E+05 | 2,15E+05 | 2 | 2 | 0 | 1 | 1 | Carbamidomethylation | 26312 | PPTG_11625 | Phytophthora parasitica INRA-310 protein-L-isoaspartate O-methyltransferase (242 aa) |
| 3058 | 2706 | PPTG_12594 | 51,11 | 6 | 0 | 0 | 6 | | | 8,91E+05 | 3 | 3 | 0 | 0 | 3 | | 77331 | PPTG_12594 | Phytophthora parasitica INRA-310 hypothetical protein (685 aa) |
| 2624 | 2734 | PPTG_11853 | 51,11 | 7 | 0 | 1 | 7 | | 2,35E+05 | 2,89E+06 | 4 | 4 | 0 | 1 | 4 | Carbamidomethylation | 82986 | PPTG_11853 | Phytophthora parasitica INRA-310 transmembrane protein (740 aa) |
| 3138 | 2974 | PPTG_17293 | 51,11 | 5 | 0 | 5 | 3 | | 1,84E+05 | 1,33E+06 | 2 | 1 | 0 | 2 | 1 | Dehydration | 45330 | PPTG_17293 | Phytophthora parasitica INRA-310 protein kinase (418 aa) |
| 3020 | 3246 | PPTG_19888 | 51 | 5 | 0 | 5 | 3 | | 3,11E+05 | 0 | 2 | 2 | 0 | 2 | 1 | Carbamidomethylation | 49549 | PPTG_19888 | Phytophthora parasitica INRA-310 hypothetical protein (454 aa) |
| 3412 | 3924 | PPTG_04656 | 50,99 | 4 | 4 | 0 | 4 | 5,26E+04 | | 8,59E+05 | 1 | 1 | 1 | 0 | 1 | | 60831 | PPTG_04656 | Phytophthora parasitica INRA-310 hypothetical protein (561 aa) |
| 1866 | 2729 | PPTG_10117 | 50,96 | 34 | 5 | 5 | 34 | 3,44E+07 | 8,85E+07 | 1,78E+08 | 3 | 3 | 2 | 2 | 5 | Carbamidomethylation | 15340 | PPTG_10117 | Phytophthora parasitica INRA-310 histone H3 (137 aa) |
| 1866 | 2730 | PPTG_02834 | 50,96 | 34 | 5 | 5 | 34 | 3,44E+07 | 8,85E+07 | 1,78E+08 | 3 | 3 | 2 | 2 | 5 | Carbamidomethylation | 15340 | PPTG_02834 | Phytophthora parasitica INRA-310 histone H3 (137 aa) |
| 3561 | 3472 | PPTG_06054 | 50,92 | 20 | 0 | 20 | 0 | | 8,52E+05 | | 2 | 2 | 0 | 2 | 0 | | 16627 | PPTG_06054 | Phytophthora parasitica INRA-310 riboflavin kinase (150 aa) |
| 2497 | 2642 | PPTG_01428 | 50,9 | 1 | 0 | 1 | 0 | 2,71E+04 | 5,33E+05 | 2,07E+05 | 3 | 3 | 1 | 3 | 1 | | 509209 | PPTG_01428 | Phytophthora parasitica INRA-310 hect-domain containing peotein (4614 aa) |
| 2571 | 2492 | PPTG_11432 | 50,9 | 5 | 1 | 2 | 5 | 5,18E+04 | 3,74E+05 | 2,29E+06 | 3 | 3 | 1 | 1 | 3 | | 73625 | PPTG_11432 | Phytophthora parasitica INRA-310 GAF domain-containing protein (665 aa) |
| 2782 | 3051 | PPTG_04745 | 50,85 | 8 | 3 | 8 | 3 | 5,79E+04 | 9,79E+05 | 4,35E+05 | 2 | 2 | 1 | 2 | 1 | | 33094 | PPTG_04745 | Phytophthora parasitica INRA-310 inositol polyphosphate multikinase (290 aa) |
| 3392 | 2908 | PPTG_13103 | 50,85 | 8 | 0 | 0 | 8 | | | 1,18E+06 | 2 | 2 | 0 | 0 | 2 | | 36126 | PPTG_13103 | Phytophthora parasitica INRA-310 nucleotide-binding protein 1 (344 aa) |
| 3013 | 2760 | PPTG_00705 | 50,85 | 6 | 0 | 0 | 6 | | | 1,96E+06 | 3 | 3 | 0 | 0 | 3 | | 51661 | PPTG_00705 | Phytophthora parasitica INRA-310 vacuolar protein sorting-associating protein (461 aa) |
| 2749 | 2830 | PPTG_00713 | 50,79 | 5 | 0 | 5 | 5 | | 5,99E+05 | 1,58E+06 | 2 | 2 | 0 | 2 | 2 | Oxidation (M) | 53500 | PPTG_00713 | Phytophthora parasitica INRA-310 protoporphyrinogen oxidase (493 aa) |
| 2901 | 2802 | PPTG_01659 | 50,74 | 6 | 0 | 2 | 6 | | 2,23E+05 | 2,65E+06 | 3 | 3 | 0 | 1 | 3 | | 55548 | PPTG_01659 | Phytophthora parasitica INRA-310 lysophospholipid acyltransferase (493 aa) |
| 3483 | 3280 | PPTG_18182 | 50,7 | 5 | 0 | 0 | 5 | | | 4,20E+05 | 2 | 2 | 0 | 0 | 2 | Carbamidomethylation | 58271 | PPTG_18182 | Phytophthora parasitica INRA-310 hypothetical protein (513 aa) |
| 2737 | 2589 | PPTG_00598 | 50,66 | 5 | 0 | 4 | 3 | | 4,87E+05 | 1,22E+06 | 3 | 3 | 0 | 2 | 2 | | 73133 | PPTG_00598 | Phytophthora parasitica INRA-310 serine protease family S33 (669 aa) |
| 2418 | 2948 | PPTG_01948 | 50,64 | 15 | 8 | 8 | 15 | 6,69E+05 | 8,67E+05 | 6,47E+06 | 2 | 2 | 1 | 2 | 3 | Dehydration; Deamidation (NQ); Acetylation (N-term) | 13158 | PPTG_01948 | Phytophthora parasitica INRA-310 hypothetical protein (119 aa) |
| 2104 | 3475 | PPTG_14169 | 50,64 | 3 | 1 | 3 | 1 | | 2,81E+05 | | 2 | 1 | 2 | 3 | 3 | | 75960 | PPTG_14169 | Phytophthora parasitica INRA-310 protein kinase (688 aa) |
| 2378 | 2800 | PPTG_08043 | 50,64 | 1 | 0 | 1 | 1 | | 3,53E+05 | 1,51E+06 | 3 | 3 | 0 | 2 | 3 | | 245102 | PPTG_08043 | Phytophthora parasitica INRA-310 myosin-like protein (2217 aa) |
| 2862 | 2934 | PPTG_07629 | 50,57 | 4 | 2 | 2 | 4 | 9,62E+05 | 9,09E+05 | 9,76E+06 | 2 | 2 | 1 | 1 | 2 | Carbamidomethylation; Amidation | 68585 | PPTG_07629 | Phytophthora parasitica INRA-310 ATP-binding Cassette (ABC) superfamily (613 aa) |
| 3095 | 2781 | PPTG_00179 | 50,51 | 10 | 0 | 0 | 10 | | | 1,14E+06 | 3 | 3 | 0 | 0 | 3 | | 31079 | PPTG_00179 | Phytophthora parasitica INRA-310 hypothetical protein (276 aa) |
| 3540 | 2927 | PPTG_07771 | 50,49 | 7 | 0 | 0 | 7 | | | 5,81E+05 | 2 | 2 | 0 | 0 | 2 | | 46909 | PPTG_07771 | Phytophthora parasitica INRA-310 protein phosphatase 2C (429 aa) |
| 2903 | 2887 | PPTG_08212 | 50,46 | 10 | 0 | 2 | 10 | | 1,94E+05 | 4,88E+05 | 3 | 3 | 0 | 1 | 3 | Carbamidomethylation | 39032 | PPTG_08212 | Phytophthora parasitica INRA-310 hypothetical protein (347 aa) |

| | | | | | | | | | | | | | | | | | | | |
|---|---|---|---|---|---|---|---|---|---|---|---|---|---|---|---|---|---|---|---|
| 3122 | 3505 | PPTG_05980 | 50,46 | 11 | 0 | 11 | 4 | | 6,44E+05 | 3,51E+05 | 2 | 1 | 0 | 2 | 1 | Carbamidomethylation | 26064 | PPTG_05980 | Phytophthora parasitica INRA-310 hypothetical protein (244 aa) |
| 3200 | 3460 | PPTG_05608 | 50,45 | 5 | 0 | 5 | 2 | | 8,36E+04 | 2,80E+05 | 2 | 1 | 0 | 2 | 1 | | 51683 | PPTG_05608 | Phytophthora parasitica INRA-310 DEAD/DEAH box RNA helicase (475 aa) |
| 2375 | 2723 | PPTG_19794 | 50,39 | 11 | 0 | 5 | 8 | | 1,17E+05 | 7,11E+05 | 5 | 5 | 0 | 2 | 4 | | 50337 | PPTG_19794 | Phytophthora parasitica INRA-310 hypothetical protein (446 aa) |
| 2529 | 2786 | PPTG_03856 | 50,38 | 16 | 0 | 12 | 4 | | 1,23E+06 | 0 | 4 | 4 | 0 | 4 | 1 | | 37667 | PPTG_03856 | Phytophthora parasitica INRA-310 hypothetical protein (341 aa) |
| 2885 | 3495 | PPTG_10970 | 50,37 | 16 | 0 | 16 | 16 | | 4,49E+05 | 1,84E+06 | 2 | 2 | 0 | 2 | 2 | | 14378 | PPTG_10970 | Phytophthora parasitica INRA-310 hypothetical protein (131 aa) |
| 2438 | 3451 | PPTG_01745 | 50,36 | 19 | 0 | 19 | 19 | | 4,25E+06 | 8,77E+06 | 2 | 2 | 0 | 3 | 3 | Oxidation (M) | 15914 | PPTG_01745 | Phytophthora parasitica INRA-310 hypothetical protein (150 aa) |
| 2905 | 3047 | PPTG_17875 | 50,34 | 10 | 5 | 5 | 10 | 1,69E+06 | 3,19E+06 | 1,51E+07 | 2 | 2 | 1 | 1 | 2 | | 21455 | PPTG_17875 | Phytophthora parasitica INRA-310 protein RER1A (184 aa) |
| 3403 | 3334 | PPTG_11610 | 50,26 | 12 | 0 | 0 | 12 | | | 1,09E+06 | 2 | 2 | 0 | 0 | 2 | | 23721 | PPTG_11610 | Phytophthora parasitica INRA-310 polypeptide deformylase (211 aa) |
| 3425 | 3014 | PPTG_12368 | 50,19 | 4 | 0 | 2 | 2 | | 6,19E+04 | 0 | 2 | 2 | 0 | 1 | 1 | | 84513 | PPTG_12368 | Phytophthora parasitica INRA-310 RNA polymerase-associated protein RTF1 (759 aa) |
| 2035 | 3088 | PPTG_10355 | 50,16 | 4 | 1 | 1 | 4 | | | 7,82E+05 | 2 | 1 | 2 | 2 | 4 | | 62759 | PPTG_10355 | Phytophthora parasitica INRA-310 protein kinase (570 aa) |
| 2435 | 3256 | PPTG_18659 | 50,13 | 7 | 4 | 7 | 7 | 9,70E+04 | 7,85E+05 | 5,47E+05 | 2 | 2 | 1 | 2 | 3 | Oxidation (M); Acetylation (N-term) | 36514 | PPTG_18659 | Phytophthora parasitica INRA-310 translation initiation factor eIF-2B subunit gamma (323 aa) |
| 3591 | 4004 | PPTG_02562 | 50,1 | 7 | 0 | 7 | 7 | | 9,05E+04 | 1,39E+05 | 1 | 1 | 0 | 1 | 1 | | 26136 | PPTG_02562 | Phytophthora parasitica INRA-310 histone H1 (251 aa) |
| 3062 | 2852 | PPTG_12056 | 50,09 | 24 | 12 | 0 | 12 | 1,09E+05 | | 2,13E+06 | 3 | 3 | 1 | 0 | 2 | Carbamidomethylation | 25349 | PPTG_12056 | Phytophthora parasitica INRA-310 hypothetical protein (231 aa) |
| 2715 | 2665 | PPTG_13709 | 50,04 | 2 | 0 | 1 | 1 | | 1,97E+05 | 2,42E+05 | 3 | 3 | 0 | 2 | 2 | | 189119 | PPTG_13709 | Phytophthora parasitica INRA-310 hypothetical protein (1746 aa) |
| 3401 | 3211 | PPTG_10756 | 50,04 | 1 | 0 | 0 | 1 | | | 3,21E+05 | 2 | 2 | 0 | 0 | 2 | | 226169 | PPTG_10756 | Phytophthora parasitica INRA-310 hypothetical protein (2001 aa) |
| 3043 | 3264 | PPTG_09763 | 49,97 | 9 | 0 | 7 | 9 | | 1,55E+05 | 5,15E+05 | 2 | 2 | 0 | 1 | 2 | Carbamidomethylation | 35390 | PPTG_09763 | Phytophthora parasitica INRA-310 alcohol dehydrogenase (333 aa) |
| 2739 | 2625 | PPTG_18080 | 49,91 | 6 | 0 | 2 | 6 | | 6,72E+05 | 2,61E+06 | 3 | 3 | 0 | 1 | 3 | | 54600 | PPTG_18080 | Phytophthora parasitica INRA-310 protein kinase (483 aa) |
| 3232 | 4229 | PPTG_18555 | 49,88 | 12 | 12 | 12 | 12 | 2,18E+05 | 8,50E+05 | 2,10E+06 | 1 | 1 | 1 | 1 | 1 | | 15108 | PPTG_18555 | Phytophthora parasitica INRA-310 proline-rich protein 6 (137 aa) |
| 2416 | 2765 | PPTG_02788 | 49,88 | 12 | 0 | 12 | 4 | | 4,71E+05 | 4,92E+05 | 4 | 4 | 0 | 4 | 2 | | 51222 | PPTG_02788 | Phytophthora parasitica INRA-310 hypothetical protein (458 aa) |
| 2811 | 2868 | PPTG_02023 | 49,86 | 12 | 6 | 6 | 12 | 1,19E+05 | 1,57E+05 | 1,41E+06 | 2 | 2 | 1 | 1 | 2 | | 22595 | PPTG_02023 | Phytophthora parasitica INRA-310 hypothetical protein (203 aa) |
| 3233 | 4230 | PPTG_14680 | 49,78 | 3 | 0 | 3 | 3 | | 7,40E+05 | 4,53E+06 | 1 | 1 | 0 | 1 | 2 | Oxidation (M) | 54794 | PPTG_14680 | Phytophthora parasitica INRA-310 ammonium transporter (Amt) family (524 aa) |
| 3031 | 2880 | PPTG_07745 | 49,77 | 6 | 0 | 6 | 0 | | 1,86E+06 | | 3 | 2 | 0 | 3 | 0 | Carbamidomethylation; Amidation; Carbamylation | 61088 | PPTG_07745 | Phytophthora parasitica INRA-310 crinkler (CRN) family protein pseudogene (541 aa) |
| 2692 | 2543 | PPTG_17933 | 49,75 | 21 | 0 | 6 | 21 | | 3,40E+05 | 1,81E+06 | 3 | 3 | 0 | 1 | 3 | Carbamidomethylation | 19323 | PPTG_17933 | Phytophthora parasitica INRA-310 hypothetical protein (178 aa) |
| 3660 | 3545 | PPTG_13973 | 49,75 | 9 | 0 | 0 | 9 | | | 5,07E+05 | 2 | 2 | 0 | 0 | 2 | | 31084 | PPTG_13973 | Phytophthora parasitica INRA-310 hypothetical protein (269 aa) |
| 3140 | 3068 | PPTG_04701 | 49,74 | 15 | 0 | 11 | 15 | 0 | | 7,75E+05 | 2 | 2 | 0 | 1 | 2 | | 16699 | PPTG_04701 | Phytophthora parasitica INRA-310 hypothetical protein (152 aa) |
| 2922 | 3568 | PPTG_10099 | 49,69 | 26 | 14 | 14 | 26 | 1,28E+06 | 1,55E+06 | 1,89E+07 | 2 | 2 | 1 | 1 | 2 | | 7896 | PPTG_10099 | Phytophthora parasitica INRA-310 hypothetical protein (74 aa) |
| 2909 | 3285 | PPTG_19388 | 49,69 | 2 | 0 | 2 | 2 | | 3,11E+05 | 4,35E+05 | 2 | 2 | 0 | 2 | 2 | Carbamidomethylation | 107019 | PPTG_19388 | Phytophthora parasitica INRA-310 hypothetical protein (970 aa) |
| 3226 | 4090 | PPTG_17614 | 49,65 | 19 | 19 | 19 | 19 | 1,09E+06 | 1,33E+06 | 9,02E+06 | 1 | 1 | 1 | 1 | 1 | | 7066 | PPTG_17614 | Phytophthora parasitica INRA-310 hypothetical protein (65 aa) |
| 3435 | 3292 | PPTG_04991 | 49,62 | 6 | 0 | 6 | 0 | | 2,37E+05 | | 2 | 2 | 0 | 2 | 0 | | 37008 | PPTG_04991 | Phytophthora parasitica INRA-310 hypothetical protein (345 aa) |
| 3349 | 3052 | PPTG_07763 | 49,59 | 1 | 0 | 0 | 1 | | | 7,33E+05 | 2 | 2 | 0 | 0 | 2 | | 181622 | PPTG_07763 | Phytophthora parasitica INRA-310 hypothetical protein (1671 aa) |
| 3422 | 2973 | PPTG_03995 | 49,59 | 4 | 4 | 0 | 4 | 2,23E+04 | | 4,72E+05 | 1 | 1 | 1 | 0 | 1 | Acetylation (N-term) | 38945 | PPTG_03995 | Phytophthora parasitica INRA-310 phosphatidic acid phosphatase (352 aa) |
| 3468 | 2909 | PPTG_13489 | 49,57 | 7 | 0 | 0 | 7 | | | 1,33E+05 | 2 | 2 | 0 | 0 | 2 | | 49042 | PPTG_13489 | Phytophthora parasitica INRA-310 vacuolar protein-sorting-associated protein (440 aa) |
| 3087 | 3978 | PPTG_04643 | 49,54 | 4 | 4 | 4 | 4 | 1,26E+06 | 1,40E+07 | 1,32E+07 | 1 | 1 | 1 | 1 | 1 | | 36627 | PPTG_04643 | Phytophthora parasitica INRA-310 hypothetical protein (350 aa) |
| 3070 | 3223 | PPTG_11826 | 49,47 | 3 | 0 | 2 | 3 | | 9,97E+04 | 3,90E+05 | 2 | 2 | 0 | 1 | 2 | | 97171 | PPTG_11826 | Phytophthora parasitica INRA-310 intraflagellar transporter 88 (867 aa) |
| 3063 | 2915 | PPTG_08300 | 49,44 | 3 | 0 | 1 | 2 | 0 | | 3,16E+05 | 3 | 3 | 0 | 1 | 2 | | 120781 | PPTG_08300 | Phytophthora parasitica INRA-310 hypothetical protein (1080 aa) |
| 2805 | 2764 | PPTG_06083 | 49,32 | 4 | 0 | 1 | 4 | | 6,84E+05 | 1,77E+06 | 3 | 3 | 0 | 1 | 3 | | 76379 | PPTG_06083 | Phytophthora parasitica INRA-310 acetoacetate-CoA ligase (690 aa) |
| 2647 | 3463 | PPTG_09213 | 49,32 | 7 | 4 | 7 | 7 | 8,56E+04 | 1,00E+06 | 2,79E+06 | 2 | 2 | 1 | 2 | 2 | | 31941 | PPTG_09213 | Phytophthora parasitica INRA-310 hypothetical protein (303 aa) |
| 2552 | 2799 | PPTG_14032 | 49,3 | 7 | 0 | 5 | 7 | | 6,66E+05 | 1,40E+06 | 3 | 3 | 0 | 2 | 3 | | 48496 | PPTG_14032 | Phytophthora parasitica INRA-310 hypothetical protein (421 aa) |
| 3411 | 3897 | PPTG_06906 | 49,29 | 4 | 0 | 4 | 4 | | 3,21E+06 | 4,19E+06 | 1 | 1 | 0 | 1 | 1 | Carbamidomethylation | 41224 | PPTG_06906 | Phytophthora parasitica INRA-310 hypothetical protein (391 aa) |
| 3411 | 4011 | PPTG_06911 | 49,29 | 4 | 0 | 4 | 4 | | 3,21E+06 | 4,19E+06 | 1 | 1 | 0 | 1 | 1 | Carbamidomethylation | 39092 | PPTG_06911 | Phytophthora parasitica INRA-310 hypothetical protein (372 aa) |
| 2699 | 2964 | PPTG_10760 | 49,28 | 7 | 2 | 3 | 5 | 1,51E+05 | 4,37E+05 | 2,27E+06 | 3 | 3 | 1 | 1 | 2 | Carbamidomethylation | 48846 | PPTG_10760 | Phytophthora parasitica INRA-310 bystin (423 aa) |
| 2804 | 2736 | PPTG_02471 | 49,27 | 8 | 2 | 2 | 5 | 9,35E+04 | 1,30E+05 | 5,87E+05 | 3 | 3 | 1 | 1 | 2 | | 59599 | PPTG_02471 | Phytophthora parasitica INRA-310 glycosyl transferase (520 aa) |
| 3642 | 3284 | PPTG_02279 | 49,27 | 5 | 0 | 0 | 5 | | | 3,57E+05 | 2 | 2 | 0 | 0 | 2 | | 59439 | PPTG_02279 | Phytophthora parasitica INRA-310 hypothetical protein (528 aa) |
| 2949 | 3623 | PPTG_03083 | 49,25 | 14 | 0 | 14 | 6 | | 1,05E+06 | 3,94E+05 | 2 | 2 | 0 | 3 | 1 | Deamidation (NQ) | 32376 | PPTG_03083 | Phytophthora parasitica INRA-310 chitin-binding protein (311 aa) |
| 2902 | 2803 | PPTG_17931 | 49,19 | 6 | 0 | 0 | 6 | | | 4,38E+05 | 4 | 3 | 0 | 0 | 4 | | 79650 | PPTG_17931 | Phytophthora parasitica INRA-310 voltage-gated Ion Channel (VIC) superfamily (706 aa) |
| 2857 | 2772 | PPTG_18444 | 49,15 | 7 | 0 | 2 | 7 | | 1,57E+05 | 6,25E+05 | 3 | 3 | 0 | 1 | 3 | Carbamidomethylation; Deamidation (NQ) | 51797 | PPTG_18444 | Phytophthora parasitica INRA-310 serine protease family S10 (459 aa) |
| 3334 | 3821 | PPTG_10993 | 49,12 | 4 | 0 | 4 | 4 | | 1,57E+05 | 1,80E+05 | 1 | 1 | 0 | 1 | 1 | | 39761 | PPTG_10993 | Phytophthora parasitica INRA-310 hypothetical protein (355 aa) |
| 3940 | 3643 | PPTG_11845 | 49,12 | 1 | 0 | 0 | 1 | | | 2,68E+05 | 1 | 1 | 0 | 0 | 1 | | 163771 | PPTG_11845 | Phytophthora parasitica INRA-310 ATP-binding Cassette (ABC) superfamily protein (1493 aa) |
| 3940 | 3644 | PPTG_11845 | 49,12 | 1 | 0 | 0 | 1 | | | 2,68E+05 | 1 | 1 | 0 | 0 | 1 | | 163859 | PPTG_11845 | Phytophthora parasitica INRA-310 ATP-binding Cassette (ABC) superfamily protein (1494 aa) |
| 2923 | 3578 | PPTG_01652 | 49,1 | 8 | 0 | 8 | 8 | | 4,90E+05 | 1,07E+06 | 2 | 2 | 0 | 2 | 2 | | 28406 | PPTG_01652 | Phytophthora parasitica INRA-310 myosin-like protein (249 aa) |
| 3184 | 2925 | PPTG_16091 | 49,07 | 15 | 0 | 10 | 5 | | 1,12E+06 | 6,10E+05 | 3 | 3 | 0 | 2 | 1 | | 23323 | PPTG_16091 | Phytophthora parasitica INRA-310 elongation factor P (212 aa) |
| 3433 | 3153 | PPTG_07254 | 49,06 | 6 | 0 | 0 | 6 | | | 6,14E+05 | 2 | 2 | 0 | 0 | 2 | | 39939 | PPTG_07254 | Phytophthora parasitica INRA-310 hypothetical protein (363 aa) |
| 2644 | 2767 | PPTG_13149 | 49,06 | 21 | 0 | 0 | 21 | | | 1,78E+06 | 5 | 5 | 0 | 0 | 5 | Carbamidomethylation | 32393 | PPTG_13149 | Phytophthora parasitica INRA-310 hypothetical protein (293 aa) |
| 3646 | 3327 | PPTG_13881 | 48,92 | 14 | 0 | 14 | 0 | | 3,49E+05 | | 2 | 2 | 0 | 2 | 0 | | 34036 | PPTG_13881 | Phytophthora parasitica INRA-310 hypothetical protein (304 aa) |
| 3100 | 2903 | PPTG_12317 | 48,92 | 6 | 0 | 2 | 6 | | 2,72E+05 | 8,45E+05 | 2 | 2 | 0 | 1 | 2 | | 44227 | PPTG_12317 | Phytophthora parasitica INRA-310 hypothetical protein (403 aa) |
| 2878 | 3305 | PPTG_15191 | 48,88 | 4 | 0 | 4 | 4 | | 2,36E+05 | 3,23E+05 | 2 | 2 | 0 | 2 | 2 | | 43205 | PPTG_15191 | Phytophthora parasitica INRA-310 exodeoxyribonuclease (385 aa) |
| 2600 | 2697 | PPTG_09060 | 48,83 | 5 | 0 | 4 | 5 | | 1,93E+06 | 3,15E+06 | 3 | 3 | 0 | 2 | 3 | | 68334 | PPTG_09060 | Phytophthora parasitica INRA-310 gephyrin-like protein (643 aa) |
| 2746 | 2679 | PPTG_15147 | 48,68 | 7 | 0 | 5 | 4 | | 5,26E+05 | 4,99E+05 | 3 | 3 | 0 | 2 | 2 | Carbamidomethylation | 67913 | PPTG_15147 | Phytophthora parasitica INRA-310 hypothetical protein (600 aa) |
| 2919 | 3500 | PPTG_13390 | 48,65 | 7 | 0 | 7 | 7 | | 4,82E+05 | 1,15E+06 | 2 | 2 | 0 | 2 | 2 | | 32362 | PPTG_13390 | Phytophthora parasitica INRA-310 phosphoglycerate mutase (283 aa) |
| 3015 | 2947 | PPTG_05174 | 48,6 | 5 | 0 | 3 | 5 | | 2,09E+05 | 5,42E+05 | 2 | 2 | 0 | 1 | 2 | Carbamidomethylation; Deamidation (NQ) | 45812 | PPTG_05174 | Phytophthora parasitica INRA-310 ribosomal RNA assembly protein mis3 (398 aa) |
| 3150 | 3313 | PPTG_06874 | 48,58 | 2 | 0 | 1 | 2 | | 2,26E+05 | 2,44E+05 | 2 | 2 | 0 | 1 | 2 | | 102407 | PPTG_06874 | Phytophthora parasitica INRA-310 hypothetical protein (935 aa) |
| 3186 | 2978 | PPTG_09699 | 48,57 | 7 | 0 | 7 | 3 | | 7,31E+05 | 4,36E+05 | 2 | 2 | 0 | 2 | 1 | Oxidation (M) | 36452 | PPTG_09699 | Phytophthora parasitica INRA-310 hypothetical protein (330 aa) |
| 3402 | 3274 | PPTG_14478 | 48,54 | 3 | 0 | 0 | 3 | | | 2,04E+06 | 2 | 2 | 0 | 0 | 2 | | 67807 | PPTG_14478 | Phytophthora parasitica INRA-310 hypothetical protein (618 aa) |
| 2932 | 2862 | PPTG_04250 | 48,52 | 13 | 0 | 4 | 13 | 0 | | 3,94E+05 | 3 | 3 | 0 | 1 | 3 | | 29349 | PPTG_04250 | Phytophthora parasitica INRA-310 ribonuclease P protein subunit p30 (270 aa) |
| 3366 | 2956 | PPTG_03500 | 48,52 | 13 | 0 | 0 | 13 | | | 1,05E+06 | 2 | 2 | 0 | 0 | 2 | Carbamidomethylation | 28769 | PPTG_03500 | Phytophthora parasitica INRA-310 GrxB family glutaredoxin (254 aa) |
| 3008 | 3514 | PPTG_02672 | 48,5 | 2 | 1 | 0 | 2 | 5,37E+04 | | 8,15E+04 | 2 | 2 | 1 | 0 | 2 | | 162185 | PPTG_02672 | Phytophthora parasitica INRA-310 regulatory-associated protein-mTOR (1458 aa) |
| 2643 | 2747 | PPTG_01572 | 48,48 | 16 | 0 | 11 | 16 | | 5,40E+05 | 1,10E+06 | 3 | 3 | 0 | 2 | 3 | | 22514 | PPTG_01572 | Phytophthora parasitica INRA-310 Ras family GTPase (199 aa) |
| 2643 | 2748 | PPTG_01572T2 | 48,48 | 16 | 0 | 11 | 16 | | 5,40E+05 | 1,10E+06 | 3 | 3 | 0 | 2 | 3 | | 22514 | PPTG_01572 | Phytophthora parasitica INRA-310 Ras family GTPase (199 aa) |
| 2643 | 2749 | PPTG_01572 | 48,48 | 16 | 0 | 11 | 16 | | 5,40E+05 | 1,10E+06 | 3 | 3 | 0 | 2 | 3 | | 22514 | PPTG_01572 | Phytophthora parasitica INRA-310 Ras family GTPase (199 aa) |
| 2424 | 3582 | PPTG_10125 | 48,42 | 4 | 2 | 4 | 2 | 2,15E+05 | 2,28E+06 | 5,28E+06 | 2 | 2 | 1 | 3 | 2 | Oxidation (M) | 50031 | PPTG_10125 | Phytophthora parasitica INRA-310 hypothetical protein (477 aa) |
| 3067 | 3123 | PPTG_03899 | 48,41 | 2 | 0 | 2 | 1 | | 6,51E+05 | 2,89E+05 | 2 | 2 | 0 | 2 | 1 | | 83421 | PPTG_03899 | Phytophthora parasitica INRA-310 hypothetical protein (733 aa) |
| 3504 | 3634 | PPTG_20432 | 48,39 | 3 | 0 | 3 | 3 | | 1,84E+05 | 4,71E+05 | 1 | 1 | 0 | 1 | 1 | | 51213 | PPTG_20432 | Phytophthora parasitica INRA-310 hypothetical protein (468 aa) |
| 2501 | 3081 | PPTG_07613 | 48,34 | 6 | 3 | 6 | 6 | 6,56E+04 | 1,22E+06 | 2,57E+06 | 2 | 2 | 1 | 2 | 2 | | 39378 | PPTG_07613 | Phytophthora parasitica INRA-310 hypothetical protein (358 aa) |
| 2821 | 3221 | PPTG_09627 | 48,32 | 7 | 0 | 0 | 7 | | | 8,48E+05 | 2 | 2 | 0 | 0 | 4 | Oxidation (M) | 38406 | PPTG_09627 | Phytophthora parasitica INRA-310 hypothetical protein (336 aa) |
| 4604 | 4231 | PPTG_16270 | 48,27 | 7 | 0 | 0 | 7 | | | 1,95E+05 | 1 | 1 | 0 | 0 | 1 | Carbamidomethylation | 27010 | PPTG_16270 | Phytophthora parasitica INRA-310 hypothetical protein (254 aa) |
| 3653 | 3484 | PPTG_01185 | 48,22 | 10 | 0 | 10 | 0 | | 4,05E+05 | | 2 | 2 | 0 | 2 | 0 | Carbamidomethylation | 31976 | PPTG_01185 | Phytophthora parasitica INRA-310 hypothetical protein (290 aa) |
| 3114 | 3322 | PPTG_09085 | 48,22 | 8 | 0 | 8 | 4 | | 1,16E+06 | 1,07E+06 | 2 | 2 | 0 | 2 | 1 | | 34106 | PPTG_09085 | Phytophthora parasitica INRA-310 sporangia induced parkin coregulated protein (307 aa) |
| 3060 | 2833 | PPTG_08159 | 48,19 | 5 | 0 | 2 | 5 | | 2,36E+05 | 5,58E+05 | 2 | 2 | 0 | 1 | 2 | Carbamidomethylation | 47972 | PPTG_08159 | Phytophthora parasitica INRA-310 chaperone dnaJ (441 aa) |
| 3703 | 4239 | PPTG_08949 | 48,13 | 6 | 0 | 6 | 6 | | 5,94E+06 | 7,22E+06 | 1 | 1 | 0 | 1 | 1 | Carbamidomethylation | 20444 | PPTG_08949 | Phytophthora parasitica INRA-310 hypothetical protein (203 aa) |
| 2756 | 3328 | PPTG_01653 | 48,07 | 0 | 0 | 0 | 0 | | 1,61E+05 | 0 | 2 | 1 | 0 | 2 | 2 | | 464056 | PPTG_01653 | Phytophthora parasitica INRA-310 dynein heavy chain (4128 aa) |
| 2968 | 2683 | PPTG_18637 | 48,05 | 2 | 0 | 1 | 1 | | 4,75E+05 | 1,07E+05 | 3 | 3 | 0 | 2 | 1 | Carbamidomethylation | 213623 | PPTG_18637 | Phytophthora parasitica INRA-310 ubiquitin-specific protease (1909 aa) |
| 2876 | 3278 | PPTG_11280 | 48,03 | 4 | 2 | 2 | 4 | 4,30E+04 | 1,49E+05 | 6,38E+05 | 2 | 2 | 1 | 1 | 2 | | 49050 | PPTG_11280 | Phytophthora parasitica INRA-310 hypothetical protein (438 aa) |
| 2877 | 3296 | PPTG_17325 | 47,99 | 4 | 2 | 4 | 2 | 1,06E+05 | 9,36E+05 | 7,92E+05 | 2 | 2 | 1 | 2 | 1 | | 72061 | PPTG_17325 | Phytophthora parasitica INRA-310 protein arginine N-methyltransferase (630 aa) |
| 2980 | 2604 | PPTG_16564 | 47,99 | 4 | 0 | 0 | 4 | | | 8,69E+05 | 3 | 3 | 0 | 0 | 3 | | 83426 | PPTG_16564 | Phytophthora parasitica INRA-310 protein kinase (758 aa) |
| 3767 | 3494 | PPTG_15927 | 47,98 | 4 | 0 | 0 | 4 | | | 2,72E+05 | 2 | 2 | 0 | 0 | 2 | | 64782 | PPTG_15927 | Phytophthora parasitica INRA-310 choline dehydrogenase (591 aa) |
| 3034 | 2921 | PPTG_00036 | 47,87 | 13 | 0 | 4 | 9 | 0 | | 1,91E+06 | 3 | 3 | 0 | 1 | 2 | Acetylation (N-term) | 36484 | PPTG_00036 | Phytophthora parasitica INRA-310 arsenite-Antimonite (ArsAB) Efflux family (329 aa) |
| 2750 | 2842 | PPTG_10932 | 47,84 | 8 | 0 | 5 | 5 | | 3,64E+05 | 8,57E+05 | 3 | 3 | 0 | 2 | 2 | | 45776 | PPTG_10932 | Phytophthora parasitica INRA-310 cyclin-H (399 aa) |
| 3112 | 3239 | PPTG_01887 | 47,81 | 7 | 4 | 0 | 7 | 0 | | 1,66E+06 | 2 | 2 | 1 | 0 | 2 | | 26759 | PPTG_01887 | Phytophthora parasitica INRA-310 hypothetical protein (264 aa) |
| 3112 | 3240 | PPTG_01890 | 47,81 | 7 | 4 | 0 | 7 | 0 | | 1,66E+06 | 2 | 2 | 1 | 0 | 2 | | 27793 | PPTG_01890 | Phytophthora parasitica INRA-310 hypothetical protein (270 aa) |
| 3112 | 3668 | PPTG_01857 | 47,81 | 7 | 4 | 0 | 7 | 0 | | 1,66E+06 | 2 | 2 | 1 | 0 | 2 | | 25348 | PPTG_01857 | Phytophthora parasitica INRA-310 hypothetical protein (246 aa) |
| 2695 | 2658 | PPTG_14074 | 47,79 | 16 | 0 | 16 | 4 | | 1,62E+06 | 6,27E+05 | 3 | 3 | 0 | 3 | 1 | Carbamidomethylation | 39873 | PPTG_14074 | Phytophthora parasitica INRA-310 serine-pyruvate aminotransferase (371 aa) |
| 2888 | 3652 | PPTG_02202 | 47,75 | 4 | 2 | 2 | 4 | 0 | 5,73E+05 | 5,91E+05 | 2 | 1 | 1 | 1 | 2 | Oxidation (M) | 49948 | PPTG_02202 | Phytophthora parasitica INRA-310 histone-binding protein RBBP4 (447 aa) |
| 2657 | 2754 | PPTG_15502 | 47,72 | 12 | 0 | 8 | 9 | | 1,03E+06 | 2,63E+06 | 3 | 3 | 0 | 2 | 2 | | 31247 | PPTG_15502 | Phytophthora parasitica INRA-310 hypothetical protein (276 aa) |
| 3363 | 2924 | PPTG_06466 | 47,67 | 4 | 0 | 0 | 4 | | | 8,46E+05 | 2 | 2 | 0 | 0 | 2 | | 56954 | PPTG_06466 | Phytophthora parasitica INRA-310 hypothetical protein (513 aa) |
| 3766 | 3492 | PPTG_20231 | 47,62 | 26 | 0 | 0 | 26 | | | 8,35E+05 | 2 | 2 | 0 | 0 | 2 | | 8620 | PPTG_20231 | Phytophthora parasitica INRA-310 hypothetical protein (76 aa) |
| 2860 | 2861 | PPTG_14280 | 47,59 | 2 | 0 | 1 | 2 | | 3,30E+05 | 8,93E+05 | 3 | 3 | 0 | 1 | 3 | | 187873 | PPTG_14280 | Phytophthora parasitica INRA-310 hypothetical protein (1689 aa) |
| 3373 | 3161 | PPTG_08773 | 47,55 | 5 | 0 | 0 | 5 | | | 1,38E+06 | 2 | 2 | 0 | 0 | 2 | | 75868 | PPTG_08773 | Phytophthora parasitica INRA-310 hypothetical protein (695 aa) |
| 3545 | 3056 | PPTG_18669 | 47,53 | 6 | 0 | 3 | 3 | | 6,60E+04 | 0 | 2 | 2 | 0 | 1 | 1 | | 60477 | PPTG_18669 | Phytophthora parasitica INRA-310 serine carboxypeptidase-like family S10 (554 aa) |
| 3185 | 2951 | PPTG_11141 | 47,52 | 17 | 6 | 0 | 17 | 6,44E+04 | | 2,61E+06 | 2 | 2 | 1 | 0 | 2 | | 22918 | PPTG_11141 | Phytophthora parasitica INRA-310 hypothetical protein (204 aa) |
| 3652 | 3480 | PPTG_00577 | 47,49 | 7 | 0 | 4 | 3 | | 2,72E+05 | 2,75E+05 | 2 | 2 | 0 | 1 | 1 | Oxidation (M) | 43773 | PPTG_00577 | Phytophthora parasitica INRA-310 hypothetical protein (404 aa) |
| 2747 | 2770 | PPTG_05147 | 47,48 | 8 | 0 | 3 | 8 | 0 | | 8,01E+05 | 3 | 3 | 0 | 1 | 3 | | 45959 | PPTG_05147 | Phytophthora parasitica INRA-310 hypothetical protein (412 aa) |
| 3688 | 4123 | PPTG_05365 | 47,48 | 2 | 0 | 2 | 2 | | 3,40E+05 | 4,71E+05 | 1 | 1 | 0 | 1 | 1 | | 71062 | PPTG_05365 | Phytophthora parasitica INRA-310 aspartyl protease family A01B (663 aa) |
| 3206 | 3567 | PPTG_16334 | 47,48 | 9 | 0 | 5 | 9 | 0 | | 4,54E+05 | 2 | 2 | 0 | 1 | 2 | | 26855 | PPTG_16334 | Phytophthora parasitica INRA-310 hypothetical protein (248 aa) |
| 3474 | 3030 | PPTG_15316 | 47,47 | 10 | 0 | 0 | 10 | | | 1,72E+06 | 2 | 2 | 0 | 0 | 2 | Carbamidomethylation | 23815 | PPTG_15316 | Phytophthora parasitica INRA-310 hypothetical protein (229 aa) |
| 2999 | 2968 | PPTG_01703 | 47,42 | 6 | 0 | 3 | 6 | | 1,35E+05 | 7,12E+05 | 2 | 2 | 0 | 1 | 2 | Carbamidomethylation | 38675 | PPTG_01703 | Phytophthora parasitica INRA-310 Rab32/38 family GTPase (348 aa) |
| 3643 | 3306 | PPTG_11335 | 47,38 | 14 | 0 | 7 | 7 | 0 | | 1,62E+05 | 2 | 2 | 0 | 1 | 1 | | 26029 | PPTG_11335 | Phytophthora parasitica INRA-310 deoxyuridine 5'-triphosphate nucleotidohydrolase (240 aa) |

| | | | | | | | | | | | | | | | | | | | |
|---|---|---|---|---|---|---|---|---|---|---|---|---|---|---|---|---|---|---|---|
| 2668 | 3608 | PPTG_06704 | 47,37 | 3 | 0 | 1 | 3 | | 9,45E+05 | 2,26E+06 | 2 | 1 | 0 | 2 | 2 | | 82325 | PPTG_06704 | Phytophthora parasitica INRA-310 transmembrane protein (729 aa) |
| 2859 | 2815 | PPTG_08091 | 47,36 | 5 | 0 | 2 | 5 | | 8,34E+04 | 4,80E+05 | 3 | 3 | 0 | 1 | 3 | Carbamidomethylation | 83609 | PPTG_08091 | Phytophthora parasitica INRA-310 hypothetical protein (796 aa) |
| 3657 | 3530 | PPTG_04680 | 47,35 | 7 | 0 | 0 | 7 | | | 5,35E+05 | 2 | 2 | 0 | 0 | 2 | | 51422 | PPTG_04680 | Phytophthora parasitica INRA-310 hypothetical protein (472 aa) |
| 2723 | 2886 | PPTG_12135 | 47,33 | 10 | 3 | 10 | 3 | 2,43E+05 | 2,04E+06 | 1,95E+06 | 2 | 2 | 1 | 2 | 1 | | 26136 | PPTG_12135 | Phytophthora parasitica INRA-310 hypothetical protein (234 aa) |
| 3211 | 3627 | PPTG_06611 | 47,31 | 3 | 0 | 3 | 1 | | 1,65E+05 | 3,65E+05 | 2 | 2 | 0 | 2 | 1 | | 103433 | PPTG_06611 | Phytophthora parasitica INRA-310 hypothetical protein (934 aa) |
| 3106 | 3078 | PPTG_08524 | 47,31 | 7 | 0 | 7 | 3 | | 1,79E+05 | 9,87E+04 | 2 | 2 | 0 | 2 | 1 | | 42635 | PPTG_08524 | Phytophthora parasitica INRA-310 dynein 1b Light Intermediate Chain (383 aa) |
| 3136 | 2913 | PPTG_15754 | 47,2 | 3 | 1 | 0 | 2 | 7,01E+04 | | 5,38E+05 | 3 | 3 | 1 | 0 | 2 | | 116227 | PPTG_15754 | Phytophthora parasitica INRA-310 vacuolar protein sorting-associated protein 41 (1032 aa) |
| 3133 | 2826 | PPTG_04254 | 47,12 | 13 | 0 | 0 | 13 | | | 7,40E+05 | 3 | 3 | 0 | 0 | 3 | Carbamidomethylation | 36557 | PPTG_04254 | Phytophthora parasitica INRA-310 calcineurin-like phosphoesterase (329 aa) |
| 3048 | 3452 | PPTG_13555 | 47,1 | 2 | 0 | 1 | 2 | | 0 | 4,47E+05 | 2 | 2 | 0 | 1 | 2 | Carbamidomethylation | 112567 | PPTG_13555 | Phytophthora parasitica INRA-310 hypothetical protein (993 aa) |
| 4021 | 3762 | PPTG_10111 | 47,1 | 2 | 0 | 0 | 2 | | | 1,68E+05 | 1 | 1 | 0 | 0 | 1 | | 69023 | PPTG_10111 | Phytophthora parasitica INRA-310 hypothetical protein (611 aa) |
| 3151 | 3319 | PPTG_08900 | 47,06 | 3 | 0 | 2 | 3 | | 4,06E+05 | 8,82E+05 | 2 | 2 | 0 | 1 | 2 | | 71867 | PPTG_08900 | Phytophthora parasitica INRA-310 hypothetical protein (657 aa) |
| 3135 | 2850 | PPTG_12756 | 47,02 | 4 | 0 | 3 | 1 | | 9,24E+05 | 4,46E+05 | 3 | 3 | 0 | 2 | 1 | | 98281 | PPTG_12756 | Phytophthora parasitica INRA-310 polyribonucleotide nucleotidyltransferase (907 aa) |
| 3639 | 3060 | PPTG_10445 | 47 | 7 | 0 | 0 | 7 | | | 8,02E+05 | 2 | 2 | 0 | 0 | 2 | Carbamidomethylation | 52458 | PPTG_10445 | Phytophthora parasitica INRA-310 glycoside-Pentoside-Hexuronide (GPH):Cation Symporter family (478 aa) |
| 2911 | 3323 | PPTG_12127 | 47 | 7 | 0 | 7 | 7 | | 4,78E+05 | 5,36E+05 | 2 | 2 | 0 | 2 | 2 | Carbamidomethylation | 49084 | PPTG_12127 | Phytophthora parasitica INRA-310 queuine tRNA-ribosyltransferase domain-containing protein (446 aa) |
| 2635 | 3471 | PPTG_09147 | 46,95 | 3 | 2 | 3 | 3 | 6,15E+04 | 3,36E+05 | 1,07E+06 | 2 | 2 | 1 | 2 | 2 | | 62589 | PPTG_09147 | Phytophthora parasitica INRA-310 hypothetical protein (574 aa) |
| 2994 | 2839 | PPTG_17766 | 46,94 | 22 | 0 | 22 | 0 | | 2,23E+06 | | 3 | 3 | 0 | 3 | 0 | | 14171 | PPTG_17766 | Phytophthora parasitica INRA-310 50S ribosomal protein L27 (132 aa) |
| 3257 | 3490 | PPTG_02216 | 46,92 | 7 | 0 | 7 | 4 | | 5,40E+05 | 8,04E+05 | 2 | 2 | 0 | 2 | 1 | | 27301 | PPTG_02216 | Phytophthora parasitica INRA-310 hypothetical protein (255 aa) |
| 3257 | 3493 | PPTG_19626 | 46,92 | 4 | 0 | 4 | 2 | | 5,40E+05 | 8,04E+05 | 2 | 2 | 0 | 2 | 1 | | 55580 | PPTG_19626 | Phytophthora parasitica INRA-310 hypothetical protein (511 aa) |
| 3501 | 3616 | PPTG_12394 | 46,9 | 11 | 0 | 6 | 5 | | 3,38E+05 | 4,06E+05 | 2 | 2 | 0 | 1 | 1 | | 24921 | PPTG_12394 | Phytophthora parasitica INRA-310 co-chaperone GrpE (222 aa) |
| 2819 | 3178 | PPTG_13104 | 46,88 | 3 | 0 | 3 | 3 | | 1,02E+06 | 2,56E+06 | 2 | 2 | 0 | 2 | 2 | | 89234 | PPTG_13104 | Phytophthora parasitica INRA-310 protein phosphatase 1E (803 aa) |
| 3032 | 2902 | PPTG_09675 | 46,83 | 3 | 0 | 1 | 3 | | 2,22E+05 | 7,68E+05 | 2 | 2 | 0 | 1 | 2 | | 94308 | PPTG_09675 | Phytophthora parasitica INRA-310 transcription initiation factor TFIID subunit (864 aa) |
| 2583 | 3300 | PPTG_01976 | 46,82 | 13 | 5 | 13 | 13 | 9,08E+05 | 4,48E+06 | 8,82E+06 | 2 | 2 | 1 | 2 | 2 | | 16586 | PPTG_01976 | Phytophthora parasitica INRA-310 50S ribosomal protein L17 (147 aa) |
| 3115 | 3352 | PPTG_00583 | 46,73 | 10 | 0 | 10 | 3 | | 1,92E+06 | 7,22E+05 | 2 | 2 | 0 | 2 | 1 | Carbamidomethylation | 27174 | PPTG_00583 | Phytophthora parasitica INRA-310 SCP-like extracellular protein (260 aa) |
| 3115 | 3353 | PPTG_08719 | 46,73 | 10 | 0 | 10 | 3 | | 1,92E+06 | 7,22E+05 | 2 | 2 | 0 | 2 | 1 | Carbamidomethylation | 27301 | PPTG_08719 | Phytophthora parasitica INRA-310 SCP-like extracellular protein (261 aa) |
| 3115 | 3354 | PPTG_00582 | 46,73 | 10 | 0 | 10 | 3 | | 1,92E+06 | 7,22E+05 | 2 | 2 | 0 | 2 | 1 | Carbamidomethylation | 27301 | PPTG_00582 | Phytophthora parasitica INRA-310 SCP-like extracellular protein (261 aa) |
| 3115 | 3355 | PPTG_08720 | 46,73 | 10 | 0 | 10 | 3 | | 1,92E+06 | 7,22E+05 | 2 | 2 | 0 | 2 | 1 | Carbamidomethylation | 27363 | PPTG_08720 | Phytophthora parasitica INRA-310 SCP-like extracellular protein (261 aa) |
| 3053 | 3767 | PPTG_00993 | 46,69 | 2 | 2 | 2 | 2 | 6,55E+04 | 8,28E+04 | 3,94E+05 | 1 | 1 | 1 | 1 | 1 | | 63208 | PPTG_00993 | Phytophthora parasitica INRA-310 4-coumarate-CoA ligase (574 aa) |
| 3053 | 3768 | PPTG_00993 | 46,69 | 2 | 2 | 2 | 2 | 6,55E+04 | 8,28E+04 | 3,94E+05 | 1 | 1 | 1 | 1 | 1 | | 63208 | PPTG_00993 | Phytophthora parasitica INRA-310 4-coumarate-CoA ligase (574 aa) |
| 2634 | 3470 | PPTG_18340 | 46,69 | 13 | 13 | 7 | 13 | 4,04E+06 | 5,66E+06 | 2,61E+07 | 2 | 2 | 2 | 1 | 2 | | 14002 | PPTG_18340 | Phytophthora parasitica INRA-310 mitochondrial protein Translocase (MPT) family (132 aa) |
| 3234 | 4237 | PPTG_14681 | 46,68 | 4 | 4 | 4 | 4 | 3,21E+05 | 3,34E+05 | 1,99E+06 | 1 | 1 | 1 | 1 | 1 | Oxidation (M) | 37013 | PPTG_14681 | Phytophthora parasitica INRA-310 ammonium transporter (Amt) family (350 aa) |
| 3420 | 2916 | PPTG_00365 | 46,53 | 4 | 2 | 0 | 2 | 0 | | 5,75E+05 | 2 | 2 | 1 | 0 | 1 | Oxidation (M) | 75678 | PPTG_00365 | Phytophthora parasitica INRA-310 ATP-binding Cassette (ABC) superfamily (666 aa) |
| 2883 | 3478 | PPTG_13981 | 46,53 | 12 | 4 | 4 | 12 | 1,65E+05 | 8,02E+05 | 2,31E+06 | 2 | 2 | 1 | 1 | 2 | | 22290 | PPTG_13981 | Phytophthora parasitica INRA-310 hypothetical protein (200 aa) |
| 2474 | 2798 | PPTG_09624 | 46,47 | 9 | 3 | 3 | 9 | 5,80E+05 | 1,41E+06 | 5,33E+06 | 3 | 3 | 1 | 1 | 3 | Carbamidomethylation | 37199 | PPTG_09624 | Phytophthora parasitica INRA-310 alcohol dehydrogenase (343 aa) |
| 3375 | 3499 | PPTG_06693 | 46,46 | 7 | 0 | 0 | 7 | | | 3,33E+05 | 2 | 2 | 0 | 0 | 2 | | 34990 | PPTG_06693 | Phytophthora parasitica INRA-310 hypothetical protein (312 aa) |
| 3704 | 4247 | PPTG_11998 | 46,43 | 5 | 0 | 5 | 5 | | 1,94E+05 | 9,81E+04 | 1 | 1 | 0 | 1 | 1 | | 37003 | PPTG_11998 | Phytophthora parasitica INRA-310 hypothetical protein (342 aa) |
| 3475 | 3046 | PPTG_11023 | 46,4 | 5 | 0 | 3 | 2 | | 4,32E+04 | 1,50E+05 | 2 | 2 | 0 | 1 | 1 | | 55672 | PPTG_11023 | Phytophthora parasitica INRA-310 bifunctional dihydrofolate reductase-thymidylate synthase 2 (492 aa) |
| 3329 | 2879 | PPTG_05875 | 46,38 | 3 | 0 | 0 | 3 | | | 1,61E+05 | 2 | 2 | 0 | 0 | 2 | | 116294 | PPTG_05875 | Phytophthora parasitica INRA-310 elongation factor Tu GTP-binding domain-containing protein (1045 aa) |
| 2655 | 2919 | PPTG_05899 | 46,38 | 2 | 0 | 1 | 2 | | 5,18E+05 | 6,22E+05 | 3 | 3 | 0 | 2 | 2 | | 194152 | PPTG_05899 | Phytophthora parasitica INRA-310 hypothetical protein (1754 aa) |
| 3765 | 3477 | PPTG_13133 | 46,36 | 9 | 0 | 0 | 9 | | | 1,30E+05 | 2 | 2 | 0 | 0 | 2 | | 29782 | PPTG_13133 | Phytophthora parasitica INRA-310 hypothetical protein (278 aa) |
| 3145 | 3156 | PPTG_17100 | 46,31 | 7 | 0 | 3 | 7 | | 4,56E+05 | 1,10E+06 | 2 | 2 | 0 | 1 | 2 | | 48038 | PPTG_17100 | Phytophthora parasitica INRA-310 pre-mRNA cleavage complex II protein Clp1 (442 aa) |
| 3161 | 3572 | PPTG_08556 | 46,21 | 22 | 0 | 15 | 22 | | 4,21E+05 | 2,17E+06 | 2 | 2 | 0 | 1 | 2 | | 13865 | PPTG_08556 | Phytophthora parasitica INRA-310 hypothetical protein (128 aa) |
| 2487 | 2831 | PPTG_04724 | 46,05 | 4 | 0 | 3 | 4 | | 6,37E+05 | 1,54E+06 | 3 | 3 | 0 | 2 | 3 | | 85384 | PPTG_04724 | Phytophthora parasitica INRA-310 hypothetical protein (778 aa) |
| 3768 | 3507 | PPTG_13226 | 46,05 | 16 | 0 | 0 | 16 | | | 5,59E+05 | 2 | 2 | 0 | 0 | 2 | Carbamidomethylation | 19823 | PPTG_13226 | Phytophthora parasitica INRA-310 molybdenum cofactor biosynthesis protein C (185 aa) |
| 2934 | 2922 | PPTG_01540 | 46,05 | 5 | 2 | 3 | 2 | 0 | 2,83E+05 | 2,08E+05 | 3 | 3 | 1 | 2 | 1 | | 71860 | PPTG_01540 | Phytophthora parasitica INRA-310 hypothetical protein (637 aa) |
| 2827 | 3318 | PPTG_07605 | 46,04 | 4 | 1 | 1 | 4 | 6,48E+04 | 1,96E+05 | 5,13E+05 | 2 | 2 | 1 | 1 | 2 | | 74990 | PPTG_07605 | Phytophthora parasitica INRA-310 hypothetical protein (681 aa) |
| 3022 | 3465 | PPTG_14722 | 46,04 | 1 | 0 | 0 | 1 | | | 1,04E+06 | 2 | 1 | 0 | 1 | 2 | | 490399 | PPTG_14722 | Phytophthora parasitica INRA-310 dynein heavy chain (4334 aa) |
| 3269 | 3671 | PPTG_08667 | 46,02 | 9 | 0 | 5 | 9 | | 2,62E+05 | 5,20E+05 | 2 | 2 | 0 | 1 | 2 | | 31057 | PPTG_08667 | Phytophthora parasitica INRA-310 haloacid dehalogenase-like hydrolase (280 aa) |
| 3039 | 3079 | PPTG_00354 | 46,02 | 13 | 0 | 13 | 6 | | 6,58E+05 | 2,20E+05 | 2 | 2 | 0 | 2 | 1 | | 25272 | PPTG_00354 | Phytophthora parasitica INRA-310 26S proteasome non-ATPase regulatory subunit (228 aa) |
| 4271 | 4092 | PPTG_11170 | 46,02 | 4 | 0 | 0 | 4 | | | 4,92E+05 | 1 | 1 | 0 | 0 | 1 | | 47143 | PPTG_11170 | Phytophthora parasitica INRA-310 amino Acid/Auxin Permease (AAAP) family (437 aa) |
| 2685 | 2926 | PPTG_03872 | 45,97 | 2 | 0 | 2 | 2 | | 5,63E+05 | 1,23E+06 | 2 | 2 | 0 | 2 | 2 | | 94065 | PPTG_03872 | Phytophthora parasitica INRA-310 N-terminal kinase-like protein (884 aa) |
| 3465 | 2856 | PPTG_07054 | 45,96 | 3 | 0 | 0 | 3 | | | 6,49E+05 | 2 | 2 | 0 | 0 | 2 | | 78797 | PPTG_07054 | Phytophthora parasitica INRA-310 hypothetical protein (713 aa) |
| 2686 | 2935 | PPTG_10166 | 45,94 | 4 | 0 | 4 | 1 | | 7,63E+05 | 1,55E+05 | 3 | 3 | 0 | 3 | 1 | | 89166 | PPTG_10166 | Phytophthora parasitica INRA-310 endo-1 3(4)-beta-glucanase 1 (796 aa) |
| 2686 | 2936 | PPTG_10162 | 45,94 | 4 | 0 | 4 | 1 | | 7,63E+05 | 1,55E+05 | 3 | 3 | 0 | 3 | 1 | | 88965 | PPTG_10162 | Phytophthora parasitica INRA-310 endo-1 3(4)-beta-glucanase 1 (797 aa) |
| 2686 | 2937 | PPTG_04547 | 45,94 | 4 | 0 | 4 | 1 | | 7,63E+05 | 1,55E+05 | 3 | 3 | 0 | 3 | 1 | | 88965 | PPTG_04547 | Phytophthora parasitica INRA-310 endo-1 3(4)-beta-glucanase 1 (796 aa) |
| 3265 | 3575 | PPTG_13890 | 45,87 | 14 | 0 | 14 | 5 | | 3,82E+05 | 3,30E+05 | 2 | 2 | 0 | 2 | 1 | | 28558 | PPTG_13890 | Phytophthora parasitica INRA-310 phosphoribosylformimino-5-aminoimidazole carboxamide ribotide isomerase (261 aa) |
| 3645 | 3320 | PPTG_09947 | 45,87 | 11 | 0 | 0 | 11 | | | 1,15E+05 | 2 | 2 | 0 | 0 | 2 | | 37599 | PPTG_09947 | Phytophthora parasitica INRA-310 hypothetical protein (341 aa) |
| 2662 | 2816 | PPTG_00404 | 45,82 | 2 | 1 | 0 | 1 | 0 | 0 | 0 | 3 | 3 | 1 | 1 | 2 | | 187842 | PPTG_00404 | Phytophthora parasitica INRA-310 ATPase (1660 aa) |
| 3559 | 3394 | PPTG_01180 | 45,81 | 4 | 0 | 0 | 4 | | | 3,18E+05 | 2 | 2 | 0 | 0 | 2 | | 74795 | PPTG_01180 | Phytophthora parasitica INRA-310 voltage-gated Ion Channel (VIC) superfamily (661 aa) |
| 3193 | 3302 | PPTG_16752 | 45,79 | 4 | 0 | 0 | 4 | | | 6,09E+05 | 2 | 2 | 0 | 0 | 3 | | 79252 | PPTG_16752 | Phytophthora parasitica INRA-310 hypothetical protein (710 aa) |
| 3571 | 3659 | PPTG_08737 | 45,78 | 19 | 0 | 10 | 9 | | | 1,02E+05 | 2 | 1 | 0 | 1 | 1 | Carbamidomethylation | 18493 | PPTG_08737 | Phytophthora parasitica INRA-310 SCP-like extracellular protein (170 aa) |
| 4410 | 4242 | PPTG_00697 | 45,78 | 5 | 0 | 0 | 5 | | | 3,75E+05 | 1 | 1 | 0 | 0 | 1 | | 26958 | PPTG_00697 | Phytophthora parasitica INRA-310 hypothetical protein (252 aa) |
| 3207 | 3595 | PPTG_08122 | 45,77 | 5 | 0 | 5 | 1 | | 1,58E+05 | 6,01E+05 | 2 | 2 | 0 | 2 | 1 | | 58764 | PPTG_08122 | Phytophthora parasitica INRA-310 periodic tryptophan protein (537 aa) |
| 3016 | 2986 | PPTG_11413 | 45,75 | 4 | 0 | 1 | 4 | | 3,82E+05 | 8,25E+05 | 2 | 2 | 0 | 1 | 2 | | 61172 | PPTG_11413 | Phytophthora parasitica INRA-310 acyl-CoA synthetase (553 aa) |
| 3649 | 3406 | PPTG_19392 | 45,73 | 2 | 0 | 2 | 1 | | 9,62E+04 | 2,04E+05 | 2 | 2 | 0 | 1 | 1 | | 108536 | PPTG_19392 | Phytophthora parasitica INRA-310 hypothetical protein (981 aa) |
| 3156 | 3449 | PPTG_01518 | 45,71 | 8 | 0 | 8 | 4 | | 2,19E+05 | 3,84E+05 | 2 | 2 | 0 | 2 | 1 | | 29936 | PPTG_01518 | Phytophthora parasitica INRA-310 ribosome biogenesis protein N (261 aa) |
| 2825 | 3293 | PPTG_11847 | 45,68 | 1 | 0 | 1 | 1 | | 8,82E+05 | 1,81E+06 | 2 | 2 | 0 | 2 | 2 | | 183896 | PPTG_11847 | Phytophthora parasitica INRA-310 ubiquitin-specific protease (1650 aa) |
| 4102 | 3995 | PPTG_18174 | 45,67 | 2 | 0 | 0 | 2 | | | 2,39E+05 | 1 | 1 | 0 | 0 | 1 | | 73055 | PPTG_18174 | Phytophthora parasitica INRA-310 Ufm1-specific protease (667 aa) |
| 4167 | 3926 | PPTG_14333 | 45,67 | 3 | 0 | 0 | 3 | | | 3,70E+05 | 1 | 1 | 0 | 0 | 1 | | 51346 | PPTG_14333 | Phytophthora parasitica INRA-310 hypothetical protein (479 aa) |
| 2947 | 3609 | PPTG_20005 | 45,66 | 7 | 0 | 7 | 7 | | 5,92E+05 | 7,29E+05 | 2 | 2 | 0 | 2 | 2 | Carbamidomethylation | 31788 | PPTG_20005 | Phytophthora parasitica INRA-310 hypothetical protein (281 aa) |
| 4411 | 4243 | PPTG_14519 | 45,62 | 4 | 0 | 0 | 4 | | | 1,43E+05 | 1 | 1 | 0 | 0 | 1 | | 47242 | PPTG_14519 | Phytophthora parasitica INRA-310 metal transporter (432 aa) |
| 3787 | 4253 | PPTG_02099 | 45,53 | 7 | 0 | 7 | 7 | | 8,42E+05 | 1,39E+06 | 1 | 1 | 0 | 1 | 1 | | 18157 | PPTG_02099 | Phytophthora parasitica INRA-310 carbohydrate-binding protein (201 aa) |
| 3787 | 4254 | PPTG_02086 | 45,53 | 7 | 0 | 7 | 7 | | 8,42E+05 | 1,39E+06 | 1 | 1 | 0 | 1 | 1 | | 18180 | PPTG_02086 | Phytophthora parasitica INRA-310 carbohydrate-binding protein (201 aa) |
| 3452 | 3919 | PPTG_17873 | 45,5 | 1 | 0 | 1 | 1 | | 3,04E+05 | 4,07E+05 | 1 | 1 | 0 | 1 | 1 | Carbamidomethylation | 118902 | PPTG_17873 | Phytophthora parasitica INRA-310 hypothetical protein (1085 aa) |
| 3477 | 3086 | PPTG_19618 | 45,45 | 2 | 0 | 0 | 2 | | | 2,16E+05 | 2 | 2 | 0 | 0 | 2 | | 157927 | PPTG_19618 | Phytophthora parasitica INRA-310 cation-transporting ATPase (1427 aa) |
| 3577 | 3773 | PPTG_13853 | 45,44 | 2 | 2 | 2 | 0 | 0 | 4,29E+05 | | 1 | 1 | 1 | 1 | 0 | | 71090 | PPTG_13853 | Phytophthora parasitica INRA-310 protein kinase (659 aa) |
| 2553 | 2860 | PPTG_19293 | 45,36 | 21 | 0 | 4 | 21 | | 2,47E+06 | 4,67E+06 | 4 | 4 | 0 | 1 | 4 | Oxidation (M) | 18559 | PPTG_19293 | Phytophthora parasitica INRA-310 hypothetical protein (164 aa) |
| 2553 | 2863 | PPTG_10619 | 45,36 | 24 | 0 | 5 | 24 | | 2,47E+06 | 4,67E+06 | 4 | 4 | 0 | 1 | 4 | Oxidation (M) | 16702 | PPTG_10619 | Phytophthora parasitica INRA-310 hypothetical protein (147 aa) |
| 3072 | 3307 | PPTG_17265 | 45,36 | 3 | 0 | 3 | 1 | | 1,27E+06 | 0 | 2 | 2 | 0 | 2 | 1 | | 84998 | PPTG_17265 | Phytophthora parasitica INRA-310 hypothetical protein (748 aa) |
| 2582 | 2844 | PPTG_19906 | 45,28 | 41 | 0 | 32 | 41 | | 2,16E+05 | 1,29E+06 | 3 | 2 | 0 | 2 | 3 | | 10596 | PPTG_19906 | Phytophthora parasitica INRA-310 hypothetical protein (94 aa) |
| 2701 | 3022 | PPTG_03727 | 45,27 | 1 | 0 | 1 | 1 | | 4,16E+05 | 9,73E+05 | 2 | 2 | 0 | 2 | 2 | | 166299 | PPTG_03727 | Phytophthora parasitica INRA-310 sterol 3-beta-glucosyltransferase (1509 aa) |
| 2944 | 3537 | PPTG_18823 | 45,27 | 5 | 0 | 5 | 5 | | 2,51E+05 | 3,14E+05 | 2 | 2 | 0 | 2 | 2 | Carbamidomethylation | 53309 | PPTG_18823 | Phytophthora parasitica INRA-310 tRNA (adenine-N(1)-)-methyltransferase non-catalytic subunit trm6 (476 aa) |
| 3529 | 4122 | PPTG_01958 | 45,25 | 6 | 0 | 6 | 6 | | 9,16E+04 | 1,69E+05 | 1 | 1 | 0 | 1 | 1 | | 29434 | PPTG_01958 | Phytophthora parasitica INRA-310 hypothetical protein (263 aa) |
| 2979 | 3050 | PPTG_09995 | 45,21 | 4 | 0 | 2 | 4 | | 5,56E+05 | 6,85E+05 | 2 | 2 | 0 | 1 | 2 | | 48256 | PPTG_09995 | Phytophthora parasitica INRA-310 hypothetical protein (446 aa) |
| 3519 | 3922 | PPTG_02037 | 45,15 | 4 | 0 | 4 | 4 | | 3,71E+05 | 8,61E+04 | 1 | 1 | 0 | 1 | 1 | | 39454 | PPTG_02037 | Phytophthora parasitica INRA-310 riboflavin biosynthesis protein ribBA (363 aa) |
| 3456 | 4082 | PPTG_00781 | 45,14 | 3 | 0 | 3 | 3 | | 3,53E+05 | 7,80E+05 | 1 | 1 | 0 | 1 | 1 | | 64654 | PPTG_00781 | Phytophthora parasitica INRA-310 hypothetical protein (587 aa) |
| 3557 | 3382 | PPTG_15780 | 45,14 | 9 | 0 | 5 | 4 | | 6,79E+05 | 1,31E+05 | 2 | 2 | 0 | 1 | 1 | | 27241 | PPTG_15780 | Phytophthora parasitica INRA-310 exosome complex exonuclease MTR3-like protein (253 aa) |
| 3108 | 3158 | PPTG_11771 | 45,09 | 5 | 0 | 2 | 5 | | 1,05E+05 | 1,62E+06 | 2 | 2 | 0 | 1 | 2 | Oxidation (M) | 44382 | PPTG_11771 | Phytophthora parasitica INRA-310 hypothetical protein (401 aa) |
| 3556 | 3336 | PPTG_16315 | 45,07 | 2 | 0 | 0 | 2 | | | 4,56E+05 | 2 | 2 | 0 | 0 | 2 | | 126710 | PPTG_16315 | Phytophthora parasitica INRA-310 voltage-gated Ion Channel (VIC) superfamily (1127 aa) |
| 3336 | 2999 | PPTG_10239 | 45,05 | 12 | 0 | 0 | 12 | | | 9,13E+06 | 2 | 2 | 0 | 0 | 2 | | 25588 | PPTG_10239 | Phytophthora parasitica INRA-310 hypothetical protein (229 aa) |
| 2402 | 3435 | PPTG_18441 | 45,03 | 4 | 2 | 2 | 4 | 8,99E+04 | 4,53E+05 | 1,45E+06 | 2 | 2 | 1 | 2 | 3 | Deamidation (NQ) | 51504 | PPTG_18441 | Phytophthora parasitica INRA-310 phenylalanine-4-hydroxylase henna-like protein (449 aa) |
| 3656 | 3526 | PPTG_16937 | 45,02 | 4 | 0 | 0 | 4 | | | 7,20E+05 | 2 | 2 | 0 | 0 | 2 | | 49457 | PPTG_16937 | Phytophthora parasitica INRA-310 hypothetical protein (451 aa) |
| 2726 | 2998 | PPTG_19124 | 45,01 | 7 | 3 | 7 | 4 | 2,92E+05 | 2,10E+06 | 1,20E+06 | 2 | 2 | 1 | 2 | 1 | | 38779 | PPTG_19124 | Phytophthora parasitica INRA-310 DNA-directed RNA polymerase I and III subunit RPAC1 (345 aa) |
| 3494 | 3559 | PPTG_14651 | 45,01 | 5 | 0 | 0 | 5 | | | 1,67E+06 | 2 | 2 | 0 | 0 | 2 | Carbamidomethylation | 35930 | PPTG_14651 | Phytophthora parasitica INRA-310 hypothetical protein (317 aa) |
| 2833 | 3589 | PPTG_03355 | 44,97 | 5 | 0 | 4 | 5 | | 5,51E+05 | 8,36E+05 | 2 | 2 | 0 | 2 | 2 | | 42048 | PPTG_03355 | Phytophthora parasitica INRA-310 hypothetical protein (370 aa) |
| 3004 | 3141 | PPTG_08370 | 44,95 | 5 | 0 | 2 | 5 | | 1,35E+05 | 6,31E+05 | 2 | 2 | 0 | 1 | 2 | Carbamidomethylation | 72219 | PPTG_08370 | Phytophthora parasitica INRA-310 hypothetical protein (649 aa) |
| 3103 | 2990 | PPTG_13053 | 44,88 | 3 | 1 | 1 | 2 | 1,27E+05 | 2,07E+05 | 4,89E+05 | 2 | 2 | 1 | 1 | 1 | Carbamylation | 70535 | PPTG_13053 | Phytophthora parasitica INRA-310 hypothetical protein (633 aa) |
| 2836 | 3737 | PPTG_17141 | 44,87 | 6 | 6 | 6 | 6 | 1,33E+05 | 1,83E+06 | 9,00E+05 | 1 | 1 | 1 | 1 | 2 | Deamidation (NQ) | 45728 | PPTG_17141 | Phytophthora parasitica INRA-310 glycerophosphoryl diester phosphodiesterase (418 aa) |
| 3258 | 3515 | PPTG_15115 | 44,86 | 17 | 0 | 17 | 6 | | 4,25E+05 | 3,80E+05 | 2 | 2 | 0 | 2 | 1 | | 15826 | PPTG_15115 | Phytophthora parasitica INRA-310 hypothetical protein (146 aa) |
| 3337 | 3036 | PPTG_19145 | 44,86 | 3 | 0 | 0 | 3 | | | 6,91E+05 | 2 | 2 | 0 | 0 | 2 | Carbamidomethylation | 77296 | PPTG_19145 | Phytophthora parasitica INRA-310 myosin-like protein (662 aa) |
| 3515 | 3833 | PPTG_19508 | 44,85 | 5 | 0 | 5 | 5 | | 6,46E+05 | 4,14E+05 | 1 | 1 | 0 | 1 | 1 | | 42229 | PPTG_19508 | Phytophthora parasitica INRA-310 serine protease family S33 (384 aa) |
| 3194 | 3359 | PPTG_18042 | 44,84 | 3 | 0 | 3 | 1 | | 4,59E+05 | 3,59E+05 | 2 | 2 | 0 | 2 | 1 | | 90945 | PPTG_18042 | Phytophthora parasitica INRA-310 hypothetical protein (854 aa) |
| 3644 | 3312 | PPTG_05155 | 44,83 | 5 | 0 | 0 | 5 | | | 1,02E+06 | 2 | 2 | 0 | 0 | 2 | Carbamidomethylation | 65848 | PPTG_05155 | Phytophthora parasitica INRA-310 dolichyl-P-Man:Man(5)GlcNAc(2)-PP-dolichyl mannosyltransferase (573 aa) |
| 2948 | 3611 | PPTG_18344 | 44,83 | 3 | 1 | 3 | 1 | 1,05E+05 | 4,23E+05 | 5,88E+05 | 2 | 2 | 1 | 2 | 1 | | 86063 | PPTG_18344 | Phytophthora parasitica INRA-310 hypothetical protein (755 aa) |
| 2917 | 3445 | PPTG_11156 | 44,73 | 6 | 0 | 6 | 6 | | 3,14E+05 | 4,48E+05 | 2 | 2 | 0 | 2 | 2 | | 49026 | PPTG_11156 | Phytophthora parasitica INRA-310 hypothetical protein (430 aa) |
| 2917 | 3446 | PPTG_11157 | 44,73 | 5 | 0 | 5 | 5 | | 3,14E+05 | 4,48E+05 | 2 | 2 | 0 | 2 | 2 | | 57829 | PPTG_11157 | Phytophthora parasitica INRA-310 hypothetical protein (512 aa) |
| 3484 | 3315 | PPTG_04848 | 44,68 | 5 | 0 | 0 | 5 | | | 9,05E+05 | 2 | 2 | 0 | 0 | 2 | Acetylation (N-term) | 39594 | PPTG_04848 | Phytophthora parasitica INRA-310 hypothetical protein (365 aa) |

| | | | | | | | | | | | | | | | | | | | |
|---|---|---|---|---|---|---|---|---|---|---|---|---|---|---|---|---|---|---|---|
| 4412 | 4245 | PPTG_14799 | 44,66 | 9 | 0 | 0 | 9 | | 0 | | 1 | 1 | 0 | 0 | 1 | | 18241 | PPTG_14799 | Phytophthora parasitica INRA-310 hypothetical protein (170 aa) |
| 4255 | 3998 | PPTG_15368 | 44,64 | 2 | 0 | 2 | 0 | | 2,01E+05 | | 1 | 1 | 0 | 1 | 0 | | 62111 | PPTG_15368 | Phytophthora parasitica INRA-310 hypothetical protein (585 aa) |
| 2807 | 2821 | PPTG_08283 | 44,55 | 4 | 0 | 1 | 4 | | 3,05E+05 | 1,39E+06 | 3 | 2 | 0 | 1 | 3 | | 93557 | PPTG_08283 | Phytophthora parasitica INRA-310 fat-free family protein (823 aa) |
| 3246 | 2846 | PPTG_09298 | 44,49 | 4 | 0 | 0 | 4 | | | 1,12E+06 | 3 | 3 | 0 | 0 | 3 | Carbamidomethylation | 97505 | PPTG_09298 | Phytophthora parasitica INRA-310 voltage-gated Ion Channel (VIC) superfamily (856 aa) |
| 3438 | 3596 | PPTG_14638 | 44,48 | 4 | 0 | 1 | 3 | | 0 | | 2 | 1 | 0 | 1 | 1 | | 58011 | PPTG_14638 | Phytophthora parasitica INRA-310 protein kinase (537 aa) |
| 3471 | 2991 | PPTG_14464 | 44,44 | 3 | 0 | 0 | 3 | | | 1,69E+05 | 2 | 2 | 0 | 0 | 2 | | 90899 | PPTG_14464 | Phytophthora parasitica INRA-310 pre-rRNA-processing protein TSR1 (827 aa) |
| 3259 | 3523 | PPTG_07034 | 44,42 | 2 | 1 | 0 | 2 | 4,00E+04 | | 4,86E+05 | 2 | 2 | 1 | 0 | 2 | | 117029 | PPTG_07034 | Phytophthora parasitica INRA-310 hypothetical protein (1059 aa) |
| 3213 | 3637 | PPTG_04262 | 44,4 | 56 | 0 | 17 | 39 | | 6,82E+05 | 2,09E+05 | 2 | 2 | 0 | 1 | 2 | | 9499 | PPTG_04262 | Phytophthora parasitica INRA-310 ubiquitin-fold modifier 1 (90 aa) |
| 3421 | 2954 | PPTG_08454 | 44,4 | 2 | 0 | 0 | 2 | | | 2,48E+05 | 2 | 2 | 0 | 0 | 2 | | 137288 | PPTG_08454 | Phytophthora parasitica INRA-310 voltage-gated Ion Channel (VIC) superfamily (1220 aa) |
| 3481 | 3253 | PPTG_20387 | 44,38 | 6 | 0 | 0 | 6 | | | 7,07E+05 | 2 | 2 | 0 | 0 | 2 | Carbamidomethylation | 51744 | PPTG_20387 | Phytophthora parasitica INRA-310 hypothetical protein (459 aa) |
| 2996 | 2858 | PPTG_01701 | 44,37 | 21 | 0 | 9 | 21 | 0 | | 1,77E+06 | 2 | 2 | 0 | 1 | 2 | | 11011 | PPTG_01701 | Phytophthora parasitica INRA-310 50S ribosomal protein L23 (91 aa) |
| 2757 | 3439 | PPTG_07191 | 44,32 | 12 | 7 | 7 | 12 | 4,96E+05 | 3,17E+06 | 5,51E+06 | 2 | 2 | 1 | 1 | 2 | | 18123 | PPTG_07191 | Phytophthora parasitica INRA-310 hypothetical protein (162 aa) |
| 3648 | 3381 | PPTG_17604 | 44,18 | 3 | 0 | 0 | 3 | | | 5,18E+05 | 2 | 2 | 0 | 0 | 2 | | 60214 | PPTG_17604 | Phytophthora parasitica INRA-310 hypothetical protein (539 aa) |
| 2556 | 3020 | PPTG_03091 | 44,16 | 7 | 0 | 7 | 7 | | 9,49E+05 | 1,78E+06 | 2 | 2 | 0 | 2 | 2 | | 41348 | PPTG_03091 | Phytophthora parasitica INRA-310 ubiquitin family protein (398 aa) |
| 3772 | 3602 | PPTG_08443 | 44,16 | 4 | 0 | 0 | 4 | | | 5,77E+05 | 2 | 2 | 0 | 0 | 2 | | 56063 | PPTG_08443 | Phytophthora parasitica INRA-310 hypothetical protein (486 aa) |
| 2904 | 2982 | PPTG_08842 | 44,13 | 11 | 0 | 8 | 3 | | 1,65E+06 | 4,89E+05 | 4 | 4 | 0 | 3 | 1 | | 32682 | PPTG_08842 | Phytophthora parasitica INRA-310 hypothetical protein (304 aa) |
| 3046 | 3329 | PPTG_15017 | 44,1 | 6 | 0 | 3 | 3 | | 5,47E+06 | 3,33E+06 | 3 | 3 | 0 | 1 | 2 | Oxidation (M) | 30717 | PPTG_15017 | Phytophthora parasitica INRA-310 hypothetical protein (282 aa) |
| 2998 | 2931 | PPTG_09315 | 44,1 | 1 | 0 | 0 | 1 | | 1,32E+05 | 3,99E+04 | 2 | 2 | 0 | 1 | 2 | | 329594 | PPTG_09315 | Phytophthora parasitica INRA-310 hybrid signal transduction histidine kinase (2989 aa) |
| 2323 | 2716 | PPTG_16042 | 44,06 | 4 | 0 | 4 | 4 | | 1,21E+06 | 1,92E+06 | 3 | 3 | 0 | 3 | 3 | | 73928 | PPTG_16042 | Phytophthora parasitica INRA-310 integrator complex subunit (670 aa) |
| 2323 | 2717 | PPTG_16042 | 44,06 | 4 | 0 | 4 | 4 | | 1,21E+06 | 1,92E+06 | 3 | 3 | 0 | 3 | 3 | | 73928 | PPTG_16042 | Phytophthora parasitica INRA-310 integrator complex subunit (670 aa) |
| 3769 | 3546 | PPTG_05297 | 44,03 | 5 | 0 | 5 | 0 | | 2,34E+05 | | 2 | 2 | 0 | 2 | 0 | | 82614 | PPTG_05297 | Phytophthora parasitica INRA-310 tRNA (cytosine-5-)-methyltransferase NSUN2-like protein (750 aa) |
| 3120 | 3427 | PPTG_07184 | 44,01 | 16 | 0 | 7 | 16 | | 1,18E+06 | 5,46E+06 | 2 | 2 | 0 | 1 | 2 | | 16899 | PPTG_07184 | Phytophthora parasitica INRA-310 hypothetical protein (159 aa) |
| 3661 | 3570 | PPTG_02157 | 43,99 | 2 | 0 | 0 | 2 | | | 5,04E+05 | 2 | 2 | 0 | 0 | 2 | | 107122 | PPTG_02157 | Phytophthora parasitica INRA-310 hypothetical protein (958 aa) |
| 4397 | 4115 | PPTG_18814 | 43,98 | 4 | 0 | 4 | 0 | | 2,88E+05 | | 1 | 1 | 0 | 1 | 0 | Carbamidomethylation | 44306 | PPTG_18814 | Phytophthora parasitica INRA-310 metalloprotease family M12A (397 aa) |
| 3101 | 2955 | PPTG_10952 | 43,92 | 11 | 0 | 6 | 11 | | 1,22E+05 | 6,36E+05 | 2 | 2 | 0 | 1 | 2 | Carbamidomethylation | 28153 | PPTG_10952 | Phytophthora parasitica INRA-310 tryptophan synthase alpha chain (266 aa) |
| 3282 | 4432 | PPTG_13594 | 43,9 | 14 | 0 | 14 | 10 | | 6,99E+05 | 4,20E+05 | 2 | 2 | 0 | 2 | 1 | | 10061 | PPTG_13594 | Phytophthora parasitica INRA-310 hypothetical protein (88 aa) |
| 3470 | 2958 | PPTG_00079 | 43,89 | 5 | 0 | 0 | 5 | | | 4,13E+05 | 2 | 2 | 0 | 0 | 2 | | 43687 | PPTG_00079 | Phytophthora parasitica INRA-310 hypothetical protein (388 aa) |
| 4256 | 4000 | PPTG_15792 | 43,87 | 4 | 0 | 0 | 4 | | | 1,84E+05 | 1 | 1 | 0 | 0 | 1 | | 46792 | PPTG_15792 | Phytophthora parasitica INRA-310 hypothetical protein (414 aa) |
| 3785 | 4249 | PPTG_09901 | 43,85 | 5 | 0 | 5 | 5 | | 4,89E+05 | 1,29E+06 | 1 | 1 | 0 | 1 | 1 | | 31200 | PPTG_09901 | Phytophthora parasitica INRA-310 hypothetical protein (279 aa) |
| 3245 | 2819 | PPTG_04001 | 43,84 | 10 | 0 | 0 | 10 | | | 8,84E+05 | 3 | 3 | 0 | 0 | 3 | | 47436 | PPTG_04001 | Phytophthora parasitica INRA-310 phosphatidic acid phosphatase (422 aa) |
| 3399 | 3126 | PPTG_16450 | 43,82 | 9 | 0 | 0 | 9 | | | 5,61E+05 | 2 | 2 | 0 | 0 | 2 | | 35261 | PPTG_16450 | Phytophthora parasitica INRA-310 hypothetical protein (325 aa) |
| 3665 | 3590 | PPTG_02680 | 43,8 | 2 | 0 | 0 | 2 | | | 3,44E+05 | 2 | 2 | 0 | 0 | 2 | | 190896 | PPTG_02680 | Phytophthora parasitica INRA-310 hypothetical protein (1751 aa) |
| 4170 | 3956 | PPTG_13212 | 43,79 | 3 | 0 | 3 | 0 | | 1,09E+05 | | 1 | 1 | 0 | 1 | 0 | | 52124 | PPTG_13212 | Phytophthora parasitica INRA-310 secreted RxLR effector peptide (460 aa) |
| 3360 | 3877 | PPTG_08415 | 43,78 | 2 | 2 | 0 | 2 | 2,20E+05 | | 2,20E+06 | 1 | 1 | 1 | 0 | 1 | | 64341 | PPTG_08415 | Phytophthora parasitica INRA-310 protein kinase (582 aa) |
| 3538 | 2896 | PPTG_12881 | 43,77 | 3 | 0 | 0 | 3 | | | 3,11E+05 | 2 | 2 | 0 | 0 | 2 | | 73591 | PPTG_12881 | Phytophthora parasitica INRA-310 Solute:Sodium Symporter (SSS) family (667 aa) |
| 3271 | 3693 | PPTG_16573 | 43,75 | 4 | 0 | 4 | 2 | | 2,44E+05 | 3,41E+05 | 2 | 2 | 0 | 2 | 1 | | 62524 | PPTG_16573 | Phytophthora parasitica INRA-310 hypothetical protein (567 aa) |
| 3160 | 3548 | PPTG_11578 | 43,73 | 8 | 0 | 8 | 5 | | 4,88E+05 | 5,70E+05 | 2 | 2 | 0 | 2 | 1 | | 28895 | PPTG_11578 | Phytophthora parasitica INRA-310 hypothetical protein (260 aa) |
| 3076 | 3525 | PPTG_11591 | 43,71 | 21 | 0 | 7 | 21 | | 2,46E+06 | 3,76E+06 | 2 | 2 | 0 | 1 | 2 | Carbamidomethylation | 13436 | PPTG_11591 | Phytophthora parasitica INRA-310 hypothetical protein (123 aa) |
| 3786 | 4250 | PPTG_01171 | 43,66 | 2 | 0 | 2 | 2 | | 1,71E+05 | 5,00E+05 | 1 | 1 | 0 | 1 | 1 | | 71407 | PPTG_01171 | Phytophthora parasitica INRA-310 long-chain-fatty-acid-CoA ligase (649 aa) |
| 2632 | 3411 | PPTG_04781 | 43,6 | 12 | 3 | 12 | 12 | 1,17E+05 | 1,12E+06 | 1,48E+06 | 2 | 2 | 1 | 2 | 2 | | 25775 | PPTG_04781 | Phytophthora parasitica INRA-310 hypothetical protein (231 aa) |
| 3263 | 3561 | PPTG_14387 | 43,57 | 2 | 0 | 1 | 2 | | 2,75E+05 | 7,34E+05 | 2 | 2 | 0 | 1 | 2 | | 97071 | PPTG_14387 | Phytophthora parasitica INRA-310 DEAD/DEAH box RNA helicase (851 aa) |
| 2828 | 3325 | PPTG_15460 | 43,55 | 3 | 0 | 3 | 3 | | 2,75E+05 | 4,23E+05 | 2 | 2 | 0 | 2 | 2 | | 101829 | PPTG_15460 | Phytophthora parasitica INRA-310 hypothetical protein (908 aa) |
| 3480 | 3159 | PPTG_05753 | 43,54 | 3 | 0 | 1 | 2 | | 1,32E+05 | 5,44E+05 | 2 | 2 | 0 | 1 | 1 | | 94139 | PPTG_05753 | Phytophthora parasitica INRA-310 protein kinase (846 aa) |
| 3426 | 3028 | PPTG_04553 | 43,54 | 3 | 0 | 0 | 3 | | | 3,96E+05 | 2 | 2 | 0 | 0 | 2 | | 100108 | PPTG_04553 | Phytophthora parasitica INRA-310 hypothetical protein (905 aa) |
| 2995 | 2849 | PPTG_08334 | 43,54 | 9 | 0 | 0 | 9 | | | 4,88E+05 | 3 | 3 | 0 | 0 | 3 | Carbamidomethylation | 45564 | PPTG_08334 | Phytophthora parasitica INRA-310 cytochrome c/c1 heme lyase (412 aa) |
| 3705 | 4264 | PPTG_05505 | 43,47 | 6 | 0 | 6 | 6 | | 9,41E+05 | 1,30E+06 | 1 | 1 | 0 | 1 | 1 | | 24971 | PPTG_05505 | Phytophthora parasitica INRA-310 casein kinase II subunit beta (217 aa) |
| 3705 | 4265 | PPTG_05505 | 43,47 | 6 | 0 | 6 | 6 | | 9,41E+05 | 1,30E+06 | 1 | 1 | 0 | 1 | 1 | | 28203 | PPTG_05505 | Phytophthora parasitica INRA-310 casein kinase II subunit beta (247 aa) |
| 3770 | 3550 | PPTG_00977 | 43,47 | 5 | 0 | 0 | 5 | | | 3,40E+05 | 2 | 2 | 0 | 0 | 2 | | 35921 | PPTG_00977 | Phytophthora parasitica INRA-310 hypothetical protein (316 aa) |
| 3771 | 3571 | PPTG_02347 | 43,44 | 10 | 0 | 10 | 0 | | 2,14E+05 | | 2 | 2 | 0 | 2 | 0 | | 29605 | PPTG_02347 | Phytophthora parasitica INRA-310 hypothetical protein (267 aa) |
| 3215 | 3660 | PPTG_01430 | 43,42 | 12 | 6 | 6 | 6 | 1,01E+05 | 1,25E+06 | 3,53E+06 | 2 | 2 | 1 | 1 | 1 | Carbamylation | 20276 | PPTG_01430 | Phytophthora parasitica INRA-310 ADP-ribosylation factor 1 (181 aa) |
| 3188 | 3018 | PPTG_17580 | 43,41 | 17 | 0 | 9 | 17 | | 1,12E+06 | 1,00E+07 | 2 | 2 | 0 | 1 | 2 | | 12058 | PPTG_17580 | Phytophthora parasitica INRA-310 hypothetical protein (109 aa) |
| 4173 | 3972 | PPTG_09953 | 43,38 | 4 | 0 | 0 | 4 | | | 4,72E+05 | 1 | 1 | 0 | 0 | 1 | | 41596 | PPTG_09953 | Phytophthora parasitica INRA-310 UDP-N-acetylglucosamine transporter (376 aa) |
| 3121 | 3453 | PPTG_01379 | 43,37 | 12 | 0 | 12 | 4 | | 7,06E+05 | 3,81E+05 | 2 | 2 | 0 | 2 | 1 | | 24681 | PPTG_01379 | Phytophthora parasitica INRA-310 hypothetical protein (230 aa) |
| 3121 | 3454 | PPTG_01379 | 43,37 | 12 | 0 | 12 | 4 | | 7,06E+05 | 3,81E+05 | 2 | 2 | 0 | 2 | 1 | | 24681 | PPTG_01379 | Phytophthora parasitica INRA-310 hypothetical protein (230 aa) |
| 3119 | 3424 | PPTG_02577 | 43,36 | 2 | 0 | 2 | 1 | | 2,32E+05 | 2,29E+05 | 2 | 2 | 0 | 2 | 1 | | 123449 | PPTG_02577 | Phytophthora parasitica INRA-310 hypothetical protein (1098 aa) |
| 2831 | 3486 | PPTG_10330 | 43,34 | 4 | 0 | 3 | 4 | | 1,33E+05 | 1,70E+06 | 2 | 2 | 0 | 1 | 3 | Oxidation (M) | 51273 | PPTG_10330 | Phytophthora parasitica INRA-310 SWI4 1 Peter Pan-like protein suppressor (456 aa) |
| 3209 | 3617 | PPTG_16522 | 43,33 | 5 | 0 | 3 | 5 | | 1,68E+05 | 9,75E+05 | 2 | 2 | 0 | 1 | 2 | | 40151 | PPTG_16522 | Phytophthora parasitica INRA-310 hypothetical protein (351 aa) |
| 3217 | 3666 | PPTG_03700 | 43,33 | 7 | 3 | 0 | 7 | 1,00E+06 | | 7,70E+06 | 2 | 2 | 1 | 0 | 2 | | 29500 | PPTG_03700 | Phytophthora parasitica INRA-310 aquaporin (275 aa) |
| 3486 | 3331 | PPTG_18331 | 43,32 | 3 | 0 | 0 | 3 | | | 5,56E+05 | 2 | 2 | 0 | 0 | 2 | | 60518 | PPTG_18331 | Phytophthora parasitica INRA-310 protein kinase (552 aa) |
| 3264 | 3563 | PPTG_19258 | 43,3 | 24 | 0 | 24 | 0 | | 1,83E+05 | | 2 | 2 | 0 | 3 | 0 | Deamidation (NQ) | 20806 | PPTG_19258 | Phytophthora parasitica INRA-310 hypothetical protein (186 aa) |
| 3005 | 3509 | PPTG_01574 | 43,23 | 2 | 0 | 1 | 2 | | 2,82E+05 | 5,18E+05 | 2 | 2 | 0 | 1 | 2 | | 96297 | PPTG_01574 | Phytophthora parasitica INRA-310 hypothetical protein (934 aa) |
| 2731 | 3376 | PPTG_14006 | 43,22 | 25 | 0 | 15 | 25 | | 9,57E+05 | 1,07E+07 | 2 | 2 | 0 | 1 | 3 | Carbamidomethylation; Dehydration; Deamidation (NQ) | 9124 | PPTG_14006 | Phytophthora parasitica INRA-310 hypothetical protein (81 aa) |
| 2987 | 3106 | PPTG_19296 | 43,21 | 38 | 0 | 38 | 11 | | 4,28E+06 | 4,19E+06 | 2 | 2 | 0 | 2 | 1 | Carbamidomethylation | 9474 | PPTG_19296 | Phytophthora parasitica INRA-310 hypothetical protein (86 aa) |
| 2863 | 2942 | PPTG_13882 | 43,21 | 11 | 4 | 4 | 7 | 0 | 3,14E+05 | 1,61E+06 | 3 | 3 | 1 | 1 | 2 | | 36021 | PPTG_13882 | Phytophthora parasitica INRA-310 GDP-L-fucose synthetase (322 aa) |
| 4250 | 3969 | PPTG_13754 | 43,2 | 2 | 0 | 2 | 0 | | 7,21E+04 | | 1 | 1 | 0 | 1 | 0 | | 117910 | PPTG_13754 | Phytophthora parasitica INRA-310 valyl-tRNA synthetase (1046 aa) |
| 3427 | 3034 | PPTG_16645 | 43,16 | 15 | 0 | 15 | 0 | | 3,15E+05 | | 2 | 2 | 0 | 2 | 0 | | 34377 | PPTG_16645 | Phytophthora parasitica INRA-310 aminoacylase-1 (308 aa) |
| 3124 | 3599 | PPTG_07524 | 43,15 | 19 | 0 | 11 | 19 | | 4,69E+05 | 7,46E+05 | 2 | 2 | 0 | 1 | 2 | | 21318 | PPTG_07524 | Phytophthora parasitica INRA-310 ubiquitin-conjugating enzyme E2 (197 aa) |
| 3154 | 3408 | PPTG_17326 | 43,11 | 5 | 0 | 2 | 5 | 0 | | 4,65E+05 | 2 | 2 | 0 | 1 | 2 | | 49281 | PPTG_17326 | Phytophthora parasitica INRA-310 hypothetical protein (452 aa) |
| 3107 | 3099 | PPTG_07202 | 43 | 6 | 0 | 6 | 3 | | 4,32E+05 | 0 | 2 | 2 | 0 | 2 | 1 | | 45033 | PPTG_07202 | Phytophthora parasitica INRA-310 hypothetical protein (403 aa) |
| 3116 | 3372 | PPTG_03961 | 42,98 | 6 | 0 | 6 | 4 | | 5,06E+05 | 5,30E+05 | 2 | 2 | 0 | 2 | 1 | | 50889 | PPTG_03961 | Phytophthora parasitica INRA-310 sporangia induced mitogen-activated protein kinase (457 aa) |
| 3037 | 3042 | PPTG_01832 | 42,95 | 8 | 0 | 8 | 5 | | 7,33E+05 | 5,04E+05 | 2 | 2 | 0 | 2 | 1 | | 30123 | PPTG_01832 | Phytophthora parasitica INRA-310 thioredoxin H-type (269 aa) |
| 4605 | 4257 | PPTG_02723 | 42,95 | 3 | 0 | 0 | 3 | | | 4,18E+05 | 1 | 1 | 0 | 0 | 1 | | 40307 | PPTG_02723 | Phytophthora parasitica INRA-310 drug/Metabolite transporter (DMT) superfamily (371 aa) |
| 3521 | 3954 | PPTG_08877 | 42,89 | 9 | 0 | 9 | 0 | | 3,20E+05 | | 1 | 1 | 0 | 2 | 0 | Carbamidomethylation | 21938 | PPTG_08877 | Phytophthora parasitica INRA-310 adenine phosphoribosyltransferase (203 aa) |
| 3205 | 3558 | PPTG_10516 | 42,86 | 13 | 0 | 4 | 13 | | 1,15E+06 | 1,51E+06 | 2 | 2 | 0 | 1 | 2 | Acetylation (N-term) | 37955 | PPTG_10516 | Phytophthora parasitica INRA-310 WD repeat protein pop3 (354 aa) |
| 3261 | 3556 | PPTG_10107 | 42,86 | 13 | 0 | 7 | 13 | | 5,64E+05 | 9,67E+05 | 2 | 2 | 0 | 1 | 2 | Carbamidomethylation | 40867 | PPTG_10107 | Phytophthora parasitica INRA-310 COP9 signalosome complex subunit 5 (364 aa) |
| 3197 | 3410 | PPTG_05356 | 42,85 | 19 | 0 | 0 | 19 | | | 5,66E+05 | 2 | 2 | 0 | 0 | 3 | | 15023 | PPTG_05356 | Phytophthora parasitica INRA-310 hypothetical protein (130 aa) |
| 3331 | 3339 | PPTG_16352 | 42,85 | 3 | 0 | 3 | 3 | | 1,81E+05 | 9,15E+05 | 1 | 1 | 0 | 1 | 1 | | 64487 | PPTG_16352 | Phytophthora parasitica INRA-310 protein kinase (582 aa) |
| 3033 | 2918 | PPTG_18701 | 42,85 | 9 | 0 | 9 | 3 | | 9,66E+04 | 1,38E+05 | 2 | 2 | 0 | 2 | 1 | | 45454 | PPTG_18701 | Phytophthora parasitica INRA-310 peroxisomal (S)-2-hydroxy-acid oxidase (420 aa) |
| 3669 | 3664 | PPTG_00193 | 42,8 | 11 | 0 | 0 | 11 | | | 1,25E+06 | 2 | 2 | 0 | 0 | 2 | Carbamidomethylation | 19024 | PPTG_00193 | Phytophthora parasitica INRA-310 hypothetical protein (169 aa) |
| 3567 | 3576 | PPTG_02180 | 42,8 | 4 | 0 | 2 | 2 | | 5,43E+05 | 4,70E+05 | 2 | 2 | 0 | 1 | 1 | | 60912 | PPTG_02180 | Phytophthora parasitica INRA-310 hypothetical protein (554 aa) |
| 3247 | 2966 | PPTG_18426 | 42,78 | 8 | 0 | 8 | 0 | | 3,14E+05 | | 3 | 3 | 0 | 3 | 0 | | 41548 | PPTG_18426 | Phytophthora parasitica INRA-310 hypothetical protein (387 aa) |
| 3247 | 2967 | PPTG_05001 | 42,78 | 5 | 0 | 5 | 0 | | 3,14E+05 | | 3 | 3 | 0 | 3 | 0 | | 75945 | PPTG_05001 | Phytophthora parasitica INRA-310 hypothetical protein (705 aa) |
| 3049 | 3533 | PPTG_07352 | 42,77 | 9 | 0 | 5 | 9 | | 3,88E+05 | 4,61E+05 | 2 | 2 | 0 | 1 | 2 | | 37496 | PPTG_07352 | Phytophthora parasitica INRA-310 hypothetical protein (328 aa) |
| 3489 | 3468 | PPTG_11359 | 42,74 | 4 | 0 | 2 | 2 | | 2,90E+05 | 1,29E+05 | 2 | 2 | 0 | 1 | 1 | | 50341 | PPTG_11359 | Phytophthora parasitica INRA-310 hypothetical protein (461 aa) |
| 2697 | 2870 | PPTG_19531 | 42,71 | 7 | 0 | 4 | 4 | | 6,79E+05 | 9,03E+05 | 3 | 3 | 0 | 2 | 2 | | 54914 | PPTG_19531 | Phytophthora parasitica INRA-310 COP9 signalosome complex subunit (477 aa) |
| 3397 | 3039 | PPTG_08327 | 42,71 | 4 | 0 | 0 | 4 | | | 4,44E+05 | 2 | 2 | 0 | 0 | 2 | Carbamidomethylation | 75305 | PPTG_08327 | Phytophthora parasitica INRA-310 hypothetical protein (666 aa) |
| 3361 | 3866 | PPTG_01310 | 42,68 | 1 | 0 | 1 | 1 | | 7,52E+04 | 7,93E+04 | 1 | 1 | 0 | 1 | 1 | | 109171 | PPTG_01310 | Phytophthora parasitica INRA-310 pre-mRNA-splicing factor ATP-dependent RNA helicase (953 aa) |
| 3361 | 3910 | PPTG_19143 | 42,68 | 2 | 0 | 2 | 2 | | 7,52E+04 | 7,93E+04 | 1 | 1 | 0 | 1 | 1 | | 81060 | PPTG_19143 | Phytophthora parasitica INRA-310 pre-mRNA-splicing factor ATP-dependent RNA helicase (720 aa) |
| 3788 | 4259 | PPTG_08103 | 42,49 | 7 | 0 | 7 | 7 | | 1,06E+05 | 1,95E+05 | 1 | 1 | 0 | 1 | 1 | | 24945 | PPTG_08103 | Phytophthora parasitica INRA-310 hypothetical protein (222 aa) |
| 3364 | 2943 | PPTG_00976 | 42,48 | 5 | 0 | 0 | 5 | | | 5,10E+05 | 2 | 2 | 0 | 0 | 2 | Oxidation (M) | 61463 | PPTG_00976 | Phytophthora parasitica INRA-310 hypothetical protein (544 aa) |
| 3381 | 3812 | PPTG_12017 | 42,46 | 1 | 0 | 1 | 1 | | 1,03E+05 | 9,27E+05 | 1 | 1 | 0 | 1 | 1 | | 138621 | PPTG_12017 | Phytophthora parasitica INRA-310 hypothetical protein (1241 aa) |
| 3064 | 2923 | PPTG_12607 | 42,44 | 2 | 0 | 1 | 2 | | 2,59E+05 | 3,62E+06 | 2 | 2 | 0 | 1 | 2 | | 77914 | PPTG_12607 | Phytophthora parasitica INRA-310 hypothetical protein (706 aa) |
| 2814 | 2946 | PPTG_04810 | 42,43 | 6 | 0 | 6 | 2 | | 3,27E+05 | 6,28E+05 | 3 | 3 | 0 | 3 | 1 | Carbamidomethylation | 58125 | PPTG_04810 | Phytophthora parasitica INRA-310 phosphatidylinositol-3 4 5-trisphosphate 3-phosphatase (514 aa) |
| 3099 | 2889 | PPTG_09052 | 42,42 | 4 | 0 | 1 | 3 | 0 | | 3,85E+05 | 3 | 3 | 0 | 1 | 2 | | 66072 | PPTG_09052 | Phytophthora parasitica INRA-310 4-coumarate-CoA ligase (599 aa) |
| 3045 | 3326 | PPTG_06112 | 42,41 | 3 | 0 | 1 | 3 | | 2,03E+05 | 8,16E+05 | 2 | 2 | 0 | 1 | 2 | | 111653 | PPTG_06112 | Phytophthora parasitica INRA-310 hypothetical protein (1012 aa) |
| 3502 | 3625 | PPTG_08727 | 42,38 | 5 | 0 | 5 | 0 | | 1,09E+05 | | 2 | 2 | 0 | 2 | 0 | | 47742 | PPTG_08727 | Phytophthora parasitica INRA-310 nuclear pore glycoprotein (474 aa) |
| 3326 | 3806 | PPTG_12257 | 42,34 | 0 | 0 | 0 | 0 | | 1,10E+05 | 0 | 1 | 1 | 0 | 1 | 1 | | 374587 | PPTG_12257 | Phytophthora parasitica INRA-310 phosphatidylinositol kinase (3367 aa) |
| 3789 | 4260 | PPTG_00717 | 42,33 | 8 | 0 | 8 | 8 | | 3,40E+05 | 4,57E+05 | 1 | 1 | 0 | 1 | 1 | | 15377 | PPTG_00717 | Phytophthora parasitica INRA-310 hypothetical protein (138 aa) |
| 2584 | 3426 | PPTG_02250 | 42,26 | 6 | 3 | 6 | 6 | 3,17E+05 | 2,36E+05 | 2,03E+06 | 2 | 2 | 1 | 2 | 2 | | 29930 | PPTG_02250 | Phytophthora parasitica INRA-310 hypothetical protein (263 aa) |
| 2858 | 2814 | PPTG_01503 | 42,21 | 5 | 0 | 1 | 5 | | 3,16E+05 | 1,25E+06 | 3 | 3 | 0 | 1 | 3 | | 70062 | PPTG_01503 | Phytophthora parasitica INRA-310 aspartate aminotransferase (640 aa) |
| 4414 | 4261 | PPTG_14808 | 42,21 | 6 | 0 | 0 | 6 | | 0 | | 1 | 1 | 0 | 0 | 1 | | 26265 | PPTG_14808 | Phytophthora parasitica INRA-310 hypothetical protein (236 aa) |
| 4414 | 4262 | PPTG_14808 | 42,21 | 6 | 0 | 0 | 6 | | 0 | | 1 | 1 | 0 | 0 | 1 | | 26265 | PPTG_14808 | Phytophthora parasitica INRA-310 hypothetical protein (236 aa) |
| 4606 | 4263 | PPTG_03981 | 42,19 | 5 | 0 | 0 | 5 | | | 2,10E+05 | 1 | 1 | 0 | 0 | 1 | | 26542 | PPTG_03981 | Phytophthora parasitica INRA-310 hypothetical protein (240 aa) |
| 3446 | 3816 | PPTG_18563 | 42,19 | 3 | 0 | 3 | 3 | 0 | | 1,93E+06 | 1 | 1 | 0 | 1 | 1 | | 40696 | PPTG_18563 | Phytophthora parasitica INRA-310 hypothetical protein (375 aa) |
| 4415 | 4266 | PPTG_13802 | 42,17 | 3 | 0 | 0 | 3 | | | 3,12E+05 | 1 | 1 | 0 | 0 | 1 | | 54990 | PPTG_13802 | Phytophthora parasitica INRA-310 transmembrane protein (489 aa) |
| 3266 | 3603 | PPTG_04224 | 42,15 | 2 | 0 | 1 | 2 | | 1,38E+05 | 8,38E+04 | 2 | 2 | 0 | 1 | 2 | | 131778 | PPTG_04224 | Phytophthora parasitica INRA-310 exportin-4 (1196 aa) |

| | | | | | | | | | | | | | | | | | | | |
|---|---|---|---|---|---|---|---|---|---|---|---|---|---|---|---|---|---|---|---|
| 3440 | 3670 | PPTG_18161 | 42,15 | 2 | 0 | 2 | 0 | | 6,53E+05 | | 2 | 2 | 0 | 2 | 0 | | 144238 | PPTG_18161 | Phytophthora parasitica INRA-310 hypothetical protein (1317 aa) |
| 4091 | 3856 | PPTG_14665 | 42,14 | 10 | 0 | 0 | 10 | | | 7,02E+05 | 1 | 1 | 0 | 0 | 1 | | 24100 | PPTG_14665 | Phytophthora parasitica INRA-310 glutamine amidotransferase subunit pdxT (225 aa) |
| 2780 | 3008 | PPTG_02379 | 42,07 | 5 | 0 | 5 | 5 | | 1,03E+06 | 1,77E+06 | 2 | 2 | 0 | 2 | 2 | | 39860 | PPTG_02379 | Phytophthora parasitica INRA-310 farnesyl pyrophosphate synthetase (346 aa) |
| 2921 | 3543 | PPTG_18951 | 42,06 | 14 | 7 | 14 | 7 | | 0 | 0 | 2 | 1 | 1 | 2 | 1 | | 20397 | PPTG_18951 | Phytophthora parasitica INRA-310 ATP-binding Cassette (ABC) superfamily protein (181 aa) |
| 3777 | 3697 | PPTG_11172 | 42,05 | 8 | 0 | 4 | 4 | | 2,01E+05 | 3,31E+04 | 2 | 2 | 0 | 1 | 1 | | 29895 | PPTG_11172 | Phytophthora parasitica INRA-310 hypothetical protein (270 aa) |
| 4239 | 3710 | PPTG_06295 | 42,03 | 4 | 0 | 4 | 0 | | 1,70E+05 | | 1 | 1 | 0 | 1 | 0 | | 38780 | PPTG_06295 | Phytophthora parasitica INRA-310 FKBP-type peptidyl-prolyl cis-trans isomerase (349 aa) |
| 3196 | 3407 | PPTG_15394 | 41,98 | 6 | 3 | 0 | 6 | 7,95E+04 | | 6,10E+06 | 2 | 2 | 1 | 0 | 2 | Carbamidomethylation | 40020 | PPTG_15394 | Phytophthora parasitica INRA-310 hypothetical protein (366 aa) |
| 3312 | 3016 | PPTG_18838 | 41,95 | 1 | 0 | 1 | 1 | | 0 | 1,20E+05 | 2 | 2 | 0 | 1 | 1 | | 155862 | PPTG_18838 | Phytophthora parasitica INRA-310 myosin-like protein (1378 aa) |
| 3495 | 3562 | PPTG_12605 | 41,91 | 7 | 0 | 0 | 7 | | | 3,09E+05 | 2 | 2 | 0 | 0 | 2 | | 32760 | PPTG_12605 | Phytophthora parasitica INRA-310 hypothetical protein (298 aa) |
| 3017 | 3004 | PPTG_14656 | 41,9 | 2 | 0 | 1 | 2 | | 2,78E+05 | 1,09E+06 | 2 | 2 | 0 | 1 | 2 | | 166655 | PPTG_14656 | Phytophthora parasitica INRA-310 hypothetical protein (1494 aa) |
| 3024 | 3777 | PPTG_13324 | 41,88 | 4 | 0 | 4 | 4 | | 2,01E+05 | 2,53E+05 | 2 | 2 | 0 | 1 | 2 | | 33200 | PPTG_13324 | Phytophthora parasitica INRA-310 pre-mRNA-splicing factor SF2 (292 aa) |
| 2887 | 3594 | PPTG_02172 | 41,87 | 30 | 10 | 10 | 30 | 8,86E+04 | 4,13E+05 | 1,35E+06 | 2 | 2 | 1 | 1 | 2 | | 10789 | PPTG_02172 | Phytophthora parasitica INRA-310 hypothetical protein (91 aa) |
| 3077 | 3592 | PPTG_00538 | 41,87 | 4 | 0 | 2 | 4 | | 3,55E+05 | 4,69E+05 | 2 | 2 | 0 | 1 | 2 | | 59655 | PPTG_00538 | Phytophthora parasitica INRA-310 serine/threonine protein kinase (534 aa) |
| 3791 | 4279 | PPTG_02941 | 41,85 | 4 | 0 | 4 | 4 | | 6,08E+05 | 1,13E+06 | 1 | 1 | 0 | 1 | 1 | | 48862 | PPTG_02941 | Phytophthora parasitica INRA-310 hypothetical protein (451 aa) |
| 3355 | 3665 | PPTG_15575 | 41,83 | 2 | 0 | 2 | 2 | | 8,92E+04 | 1,05E+05 | 1 | 1 | 0 | 1 | 1 | | 83206 | PPTG_15575 | Phytophthora parasitica INRA-310 heat shock protein 101 (753 aa) |
| 3689 | 4129 | PPTG_02020 | 41,82 | 2 | 0 | 2 | 2 | | 1,16E+05 | 4,34E+05 | 1 | 1 | 0 | 1 | 1 | | 72731 | PPTG_02020 | Phytophthora parasitica INRA-310 hypothetical protein (670 aa) |
| 3664 | 3586 | PPTG_03424 | 41,78 | 10 | 0 | 6 | 4 | | 1,42E+05 | 5,92E+05 | 2 | 2 | 0 | 1 | 1 | | 25359 | PPTG_03424 | Phytophthora parasitica INRA-310 hypothetical protein (242 aa) |
| 4607 | 4268 | PPTG_08527 | 41,78 | 4 | 0 | 0 | 4 | | | 2,66E+06 | 1 | 1 | 0 | 0 | 1 | | 42205 | PPTG_08527 | Phytophthora parasitica INRA-310 hypothetical protein (416 aa) |
| 3671 | 3678 | PPTG_01421 | 41,74 | 18 | 0 | 7 | 11 | | 1,42E+05 | 3,87E+05 | 2 | 2 | 0 | 1 | 1 | | 17397 | PPTG_01421 | Phytophthora parasitica INRA-310 protein mago nashi (150 aa) |
| 3706 | 4271 | PPTG_09486 | 41,74 | 2 | 0 | 2 | 2 | | 2,44E+05 | 4,79E+05 | 1 | 1 | 0 | 1 | 1 | | 47792 | PPTG_09486 | Phytophthora parasitica INRA-310 hypothetical protein (438 aa) |
| 3208 | 3607 | PPTG_01183 | 41,64 | 9 | 0 | 5 | 9 | | 7,19E+05 | 5,26E+05 | 2 | 2 | 0 | 1 | 2 | | 28768 | PPTG_01183 | Phytophthora parasitica INRA-310 hypothetical protein (266 aa) |
| 3497 | 3569 | PPTG_16201 | 41,57 | 3 | 0 | 0 | 3 | | | 1,10E+06 | 2 | 1 | 0 | 0 | 2 | | 71852 | PPTG_16201 | Phytophthora parasitica INRA-310 protein kinase (657 aa) |
| 2971 | 2932 | PPTG_08345 | 41,57 | 2 | 0 | 1 | 1 | | 1,17E+05 | 5,05E+05 | 3 | 3 | 0 | 1 | 2 | | 278843 | PPTG_08345 | Phytophthora parasitica INRA-310 autophagy-like protein (2519 aa) |
| 3579 | 3818 | PPTG_19299 | 41,52 | 3 | 0 | 3 | 3 | | 1,56E+05 | 1,25E+05 | 1 | 1 | 0 | 1 | 1 | | 36836 | PPTG_19299 | Phytophthora parasitica INRA-310 inositol 2-dehydrogenase (340 aa) |
| 3579 | 3862 | PPTG_19301 | 41,52 | 4 | 0 | 4 | 4 | | 1,56E+05 | 1,25E+05 | 1 | 1 | 0 | 1 | 1 | | 28136 | PPTG_19301 | Phytophthora parasitica INRA-310 oxidoreductase family protein (255 aa) |
| 2983 | 2928 | PPTG_16030 | 41,48 | 4 | 1 | 1 | 1 | 1,76E+05 | 8,86E+04 | 1,04E+05 | 3 | 3 | 1 | 1 | 1 | Carbamidomethylation | 94506 | PPTG_16030 | Phytophthora parasitica INRA-310 hypothetical protein (860 aa) |
| 3277 | 4281 | PPTG_20381 | 41,41 | 2 | 2 | 2 | 2 | 1,39E+05 | 1,20E+05 | 1,35E+06 | 1 | 1 | 1 | 1 | 1 | Carbamidomethylation | 47811 | PPTG_20381 | Phytophthora parasitica INRA-310 hypothetical protein (439 aa) |
| 2649 | 4288 | PPTG_18273 | 41,41 | 10 | 0 | 10 | 10 | | 8,33E+05 | 6,63E+05 | 1 | 1 | 0 | 3 | 2 | Oxidation (M) | 11714 | PPTG_18273 | Phytophthora parasitica INRA-310 hypothetical protein (107 aa) |
| 3078 | 3692 | PPTG_03435 | 41,4 | 8 | 0 | 8 | 8 | | 4,66E+05 | 8,17E+05 | 1 | 1 | 0 | 1 | 2 | | 19192 | PPTG_03435 | Phytophthora parasitica INRA-310 hypothetical protein (171 aa) |
| 3685 | 4113 | PPTG_12384 | 41,38 | 2 | 0 | 2 | 2 | | 5,79E+04 | 4,36E+04 | 1 | 1 | 0 | 1 | 1 | | 100435 | PPTG_12384 | Phytophthora parasitica INRA-310 cyclin-dependent kinase with F-box domain-containing protein (891 aa) |
| 4398 | 4116 | PPTG_02247 | 41,34 | 1 | 0 | 0 | 1 | | | 3,07E+05 | 1 | 1 | 0 | 0 | 1 | | 151579 | PPTG_02247 | Phytophthora parasitica INRA-310 ATP-binding Cassette (ABC) superfamily (1360 aa) |
| 4398 | 4296 | PPTG_19943 | 41,34 | 1 | 0 | 0 | 1 | | | 3,07E+05 | 1 | 1 | 0 | 0 | 1 | | 100067 | PPTG_19943 | Phytophthora parasitica INRA-310 ATP-binding Cassette (ABC) superfamily (900 aa) |
| 3370 | 3090 | PPTG_05232 | 41,29 | 1 | 0 | 0 | 0 | | 9,91E+04 | 0 | 2 | 2 | 0 | 1 | 1 | | 256380 | PPTG_05232 | Phytophthora parasitica INRA-310 microtubule-associated protein (2344 aa) |
| 3047 | 3423 | PPTG_02808 | 41,22 | 3 | 0 | 1 | 3 | | 9,48E+04 | 2,28E+05 | 2 | 2 | 0 | 1 | 2 | | 85374 | PPTG_02808 | Phytophthora parasitica INRA-310 oligomeric Golgi complex subunit (758 aa) |
| 3678 | 3750 | PPTG_05275 | 41,18 | 4 | 0 | 2 | 3 | | 2,03E+05 | 3,45E+05 | 2 | 2 | 0 | 1 | 1 | | 51002 | PPTG_05275 | Phytophthora parasitica INRA-310 U3 small nucleolar RNA-associated protein 18 (460 aa) |
| 3663 | 3584 | PPTG_09244 | 41,16 | 8 | 0 | 0 | 8 | | | 1,84E+05 | 2 | 2 | 0 | 0 | 2 | | 35735 | PPTG_09244 | Phytophthora parasitica INRA-310 mitochondrial Carrier (MC) family (330 aa) |
| 3638 | 2965 | PPTG_18160 | 41,13 | 8 | 0 | 0 | 8 | | | 2,32E+05 | 2 | 2 | 0 | 0 | 2 | Carbamidomethylation | 37813 | PPTG_18160 | Phytophthora parasitica INRA-310 hypothetical protein (353 aa) |
| 3038 | 3054 | PPTG_05582 | 41,09 | 3 | 0 | 3 | 1 | | 3,85E+05 | 5,05E+05 | 2 | 2 | 0 | 2 | 1 | | 89198 | PPTG_05582 | Phytophthora parasitica INRA-310 DNA mismatch repair protein Msh2 (816 aa) |
| 3073 | 3316 | PPTG_13178 | 41,04 | 5 | 0 | 5 | 2 | | 6,12E+05 | 7,80E+05 | 2 | 2 | 0 | 2 | 1 | | 51184 | PPTG_13178 | Phytophthora parasitica INRA-310 hypothetical protein (455 aa) |
| 3658 | 3541 | PPTG_17127 | 41,02 | 7 | 0 | 0 | 7 | | | 5,00E+05 | 2 | 2 | 0 | 0 | 2 | Carbamidomethylation | 38458 | PPTG_17127 | Phytophthora parasitica INRA-310 hypothetical protein (352 aa) |
| 4043 | 3168 | PPTG_11373 | 41,01 | 4 | 0 | 0 | 4 | | | 5,18E+05 | 1 | 1 | 0 | 0 | 1 | | 32187 | PPTG_11373 | Phytophthora parasitica INRA-310 hypothetical protein (285 aa) |
| 3279 | 4285 | PPTG_15479 | 40,99 | 7 | 7 | 7 | 7 | 1,38E+06 | 1,36E+06 | 5,49E+06 | 1 | 1 | 1 | 1 | 1 | | 15787 | PPTG_15479 | Phytophthora parasitica INRA-310 hypothetical protein (136 aa) |
| 2978 | 2988 | PPTG_16757 | 40,97 | 3 | 0 | 1 | 3 | | 0 | 1,70E+06 | 2 | 2 | 0 | 1 | 2 | | 107545 | PPTG_16757 | Phytophthora parasitica INRA-310 P-type ATPase (990 aa) |
| 4285 | 4274 | PPTG_04623 | 40,97 | 5 | 0 | 0 | 5 | | | 1,52E+05 | 1 | 1 | 0 | 0 | 1 | | 35752 | PPTG_04623 | Phytophthora parasitica INRA-310 hypothetical protein (337 aa) |
| 2951 | 3681 | PPTG_04376 | 40,95 | 5 | 5 | 0 | 5 | 5,82E+05 | | 3,38E+06 | 2 | 2 | 2 | 0 | 2 | | 51983 | PPTG_04376 | Phytophthora parasitica INRA-310 major facilitator superfamily transporter (MFS) (480 aa) |
| 3187 | 3011 | PPTG_09557 | 40,95 | 18 | 0 | 18 | 0 | | 3,12E+05 | | 3 | 3 | 0 | 3 | 0 | Carbamidomethylation | 31733 | PPTG_09557 | Phytophthora parasitica INRA-310 exosome complex exonuclease RRP42-like protein (296 aa) |
| 2925 | 3628 | PPTG_03644 | 40,91 | 7 | 0 | 7 | 7 | | 9,85E+05 | 1,25E+05 | 2 | 2 | 0 | 2 | 2 | Carbamidomethylation | 29927 | PPTG_03644 | Phytophthora parasitica INRA-310 hypothetical protein (266 aa) |
| 3276 | 4275 | PPTG_01698 | 40,84 | 6 | 6 | 6 | 6 | 7,53E+05 | 1,22E+06 | 5,76E+06 | 1 | 1 | 1 | 1 | 1 | | 19586 | PPTG_01698 | Phytophthora parasitica INRA-310 hypothetical protein (176 aa) |
| 4400 | 4131 | PPTG_09687 | 40,81 | 2 | 0 | 0 | 2 | | | 1,57E+05 | 1 | 1 | 0 | 0 | 1 | | 86511 | PPTG_09687 | Phytophthora parasitica INRA-310 importin subunit alpha (805 aa) |
| 4400 | 4132 | PPTG_09687 | 40,81 | 2 | 0 | 0 | 2 | | | 1,57E+05 | 1 | 1 | 0 | 0 | 1 | | 86511 | PPTG_09687 | Phytophthora parasitica INRA-310 importin subunit alpha (805 aa) |
| 3221 | 3751 | PPTG_08375 | 40,74 | 4 | 0 | 4 | 2 | | 5,47E+05 | 6,02E+05 | 2 | 2 | 0 | 2 | 1 | | 71359 | PPTG_08375 | Phytophthora parasitica INRA-310 EF-hand domain-containing protein (622 aa) |
| 3488 | 3391 | PPTG_04964 | 40,71 | 9 | 0 | 0 | 9 | | | 8,91E+05 | 2 | 2 | 0 | 0 | 2 | | 38659 | PPTG_04964 | Phytophthora parasitica INRA-310 hypothetical protein (354 aa) |
| 3790 | 4276 | PPTG_13688 | 40,66 | 6 | 0 | 6 | 6 | | 9,70E+04 | 1,90E+05 | 1 | 1 | 0 | 1 | 1 | | 31572 | PPTG_13688 | Phytophthora parasitica INRA-310 hypothetical protein (278 aa) |
| 3707 | 4277 | PPTG_04876 | 40,61 | 8 | 0 | 8 | 8 | | 3,41E+05 | 0 | 1 | 1 | 0 | 1 | 1 | | 15398 | PPTG_04876 | Phytophthora parasitica INRA-310 hypothetical protein (142 aa) |
| 3686 | 4114 | PPTG_09489 | 40,56 | 2 | 0 | 2 | 2 | | 7,26E+04 | 2,94E+05 | 1 | 1 | 0 | 1 | 1 | | 61705 | PPTG_09489 | Phytophthora parasitica INRA-310 major facilitator superfamily transporter (MFS) (563 aa) |
| 3776 | 3676 | PPTG_17339 | 40,51 | 4 | 0 | 0 | 4 | | | 2,33E+05 | 2 | 2 | 0 | 0 | 2 | | 85844 | PPTG_17339 | Phytophthora parasitica INRA-310 transmembrane protein (778 aa) |
| 3776 | 3677 | PPTG_20270 | 40,51 | 4 | 0 | 0 | 4 | | | 2,33E+05 | 2 | 2 | 0 | 0 | 2 | | 85700 | PPTG_20270 | Phytophthora parasitica INRA-310 transmembrane protein (778 aa) |
| 4182 | 4136 | PPTG_07091 | 40,49 | 10 | 0 | 0 | 10 | | | 5,84E+05 | 1 | 1 | 0 | 0 | 1 | | 18606 | PPTG_07091 | Phytophthora parasitica INRA-310 hypothetical protein (217 aa) |
| 4182 | 4137 | PPTG_07090 | 40,49 | 10 | 0 | 0 | 10 | | | 5,84E+05 | 1 | 1 | 0 | 0 | 1 | | 18576 | PPTG_07090 | Phytophthora parasitica INRA-310 hypothetical protein (217 aa) |
| 3023 | 3508 | PPTG_13842 | 40,43 | 5 | 0 | 3 | 5 | | 8,74E+05 | 8,38E+06 | 2 | 2 | 0 | 1 | 2 | | 38016 | PPTG_13842 | Phytophthora parasitica INRA-310 hypothetical protein (333 aa) |
| 3428 | 3045 | PPTG_06300 | 40,42 | 4 | 0 | 0 | 4 | | | 7,37E+05 | 2 | 2 | 0 | 0 | 2 | | 52270 | PPTG_06300 | Phytophthora parasitica INRA-310 hypothetical protein (466 aa) |
| 3565 | 3552 | PPTG_15117 | 40,4 | 4 | 0 | 0 | 4 | | | 0 | 2 | 2 | 0 | 0 | 2 | | 59171 | PPTG_15117 | Phytophthora parasitica INRA-310 threonine ammonia-lyase biosynthetic (540 aa) |
| 3006 | 3511 | PPTG_03075 | 40,4 | 4 | 0 | 2 | 4 | | 3,59E+05 | 5,22E+05 | 2 | 2 | 0 | 1 | 2 | | 57829 | PPTG_03075 | Phytophthora parasitica INRA-310 SUMO-activating enzyme (SAE) (528 aa) |
| 3216 | 3661 | PPTG_10895 | 40,31 | 7 | 0 | 3 | 7 | | 2,98E+05 | 1,42E+06 | 2 | 2 | 0 | 1 | 2 | | 48132 | PPTG_10895 | Phytophthora parasitica INRA-310 hypothetical protein (437 aa) |
| 3555 | 3304 | PPTG_05708 | 40,3 | 2 | 0 | 0 | 2 | | | 2,21E+05 | 2 | 1 | 0 | 0 | 2 | | 92447 | PPTG_05708 | Phytophthora parasitica INRA-310 HECT E3 ubiquitin ligase (809 aa) |
| 2952 | 3699 | PPTG_03079 | 40,22 | 4 | 2 | 2 | 4 | 1,97E+05 | 1,01E+06 | 1,74E+06 | 2 | 2 | 1 | 1 | 2 | | 52188 | PPTG_03079 | Phytophthora parasitica INRA-310 hypothetical protein (464 aa) |
| 4006 | 2907 | PPTG_10198 | 40,21 | 4 | 0 | 0 | 4 | | | 0 | 1 | 1 | 0 | 0 | 1 | Carbamidomethylation | 50907 | PPTG_10198 | Phytophthora parasitica INRA-310 hypothetical protein (461 aa) |
| 3451 | 3907 | PPTG_01658 | 40,2 | 7 | 0 | 7 | 7 | | 1,87E+05 | 2,06E+05 | 1 | 1 | 0 | 1 | 1 | | 23959 | PPTG_01658 | Phytophthora parasitica INRA-310 maf-like protein (214 aa) |
| 3595 | 4015 | PPTG_00930 | 40,17 | 7 | 0 | 7 | 7 | | 5,02E+05 | 2,49E+05 | 1 | 1 | 0 | 1 | 1 | | 19199 | PPTG_00930 | Phytophthora parasitica INRA-310 monothiol glutaredoxin-5 (172 aa) |
| 3126 | 3621 | PPTG_13601 | 40,16 | 2 | 0 | 1 | 2 | | | 3,31E+05 | 2 | 1 | 0 | 1 | 2 | | 85223 | PPTG_13601 | Phytophthora parasitica INRA-310 protein kinase (763 aa) |
| 3588 | 3996 | PPTG_15640 | 40,15 | 5 | 0 | 5 | 5 | | 4,03E+05 | 7,21E+05 | 1 | 1 | 0 | 1 | 1 | | 29372 | PPTG_15640 | Phytophthora parasitica INRA-310 hypothetical protein (264 aa) |
| 3654 | 3510 | PPTG_16187 | 40,14 | 3 | 0 | 0 | 3 | | | 7,22E+05 | 2 | 2 | 0 | 0 | 2 | | 70838 | PPTG_16187 | Phytophthora parasitica INRA-310 protein kinase (641 aa) |
| 3668 | 3624 | PPTG_17015 | 40,13 | 23 | 0 | 14 | 9 | | 6,05E+05 | 9,19E+04 | 2 | 2 | 0 | 1 | 1 | | 18273 | PPTG_17015 | Phytophthora parasitica INRA-310 hypothetical protein (162 aa) |
| 3174 | 4091 | PPTG_06710 | 40,07 | 8 | 8 | 8 | 8 | 3,50E+05 | 6,23E+05 | 2,96E+06 | 1 | 1 | 1 | 1 | 1 | | 15464 | PPTG_06710 | Phytophthora parasitica INRA-310 hypothetical protein (146 aa) |
| 3500 | 3614 | PPTG_10031 | 40,06 | 24 | 0 | 0 | 24 | | | 6,75E+06 | 2 | 2 | 0 | 0 | 2 | | 8815 | PPTG_10031 | Phytophthora parasitica INRA-310 hypothetical protein (81 aa) |
| 3655 | 3524 | PPTG_09003 | 40,05 | 4 | 0 | 0 | 4 | | | 7,21E+05 | 2 | 2 | 0 | 0 | 2 | | 47901 | PPTG_09003 | Phytophthora parasitica INRA-310 hsp70-like protein (450 aa) |
| 3158 | 3512 | PPTG_06681 | 40,01 | 3 | 0 | 1 | 3 | | 2,13E+05 | 3,22E+05 | 2 | 2 | 0 | 1 | 2 | | 107942 | PPTG_06681 | Phytophthora parasitica INRA-310 hypothetical protein (967 aa) |
| 3539 | 2899 | PPTG_17930 | 39,99 | 2 | 0 | 0 | 2 | | | 5,55E+05 | 2 | 1 | 0 | 0 | 2 | | 82277 | PPTG_17930 | Phytophthora parasitica INRA-310 voltage-gated Ion Channel (VIC) superfamily (735 aa) |
| 2914 | 3375 | PPTG_01151 | 39,97 | 3 | 1 | 1 | 3 | 1,74E+05 | 3,39E+05 | 1,21E+06 | 2 | 2 | 1 | 1 | 2 | Carbamidomethylation | 76716 | PPTG_01151 | Phytophthora parasitica INRA-310 hypothetical protein (687 aa) |
| 3566 | 3574 | PPTG_00656 | 39,97 | 12 | 0 | 8 | 4 | | 9,90E+04 | 4,82E+05 | 2 | 2 | 0 | 1 | 1 | | 26536 | PPTG_00656 | Phytophthora parasitica INRA-310 hypothetical protein (227 aa) |
| 3224 | 4066 | PPTG_02313 | 39,96 | 3 | 3 | 3 | 3 | 1,67E+05 | 1,52E+06 | 1,89E+06 | 1 | 1 | 1 | 1 | 1 | | 35480 | PPTG_02313 | Phytophthora parasitica INRA-310 protein phosphatase (328 aa) |
| 4608 | 4283 | PPTG_15541 | 39,92 | 1 | 0 | 0 | 1 | | | 2,51E+05 | 1 | 1 | 0 | 0 | 1 | | 104744 | PPTG_15541 | Phytophthora parasitica INRA-310 hypothetical protein (1017 aa) |
| 4158 | 3757 | PPTG_10071 | 39,89 | 5 | 0 | 0 | 5 | | | 3,07E+05 | 1 | 1 | 0 | 0 | 1 | Carbamidomethylation | 29278 | PPTG_10071 | Phytophthora parasitica INRA-310 hypothetical protein (250 aa) |
| 3278 | 4284 | PPTG_02147 | 39,88 | 3 | 0 | 3 | 3 | | 1,12E+05 | 2,69E+05 | 1 | 1 | 0 | 1 | 2 | | 69152 | PPTG_02147 | Phytophthora parasitica INRA-310 hypothetical protein (617 aa) |
| 3423 | 2993 | PPTG_09493 | 39,86 | 3 | 0 | 0 | 3 | | | 1,08E+06 | 2 | 2 | 0 | 0 | 2 | Oxidation (M) | 91746 | PPTG_09493 | Phytophthora parasitica INRA-310 hypothetical protein (824 aa) |
| 4152 | 3519 | PPTG_11044 | 39,81 | 4 | 0 | 0 | 4 | | | 6,89E+05 | 1 | 1 | 0 | 0 | 1 | | 46858 | PPTG_11044 | Phytophthora parasitica INRA-310 transmembrane protein (429 aa) |
| 3472 | 2995 | PPTG_18865 | 39,79 | 2 | 0 | 2 | 2 | | 7,02E+04 | 3,44E+05 | 1 | 1 | 0 | 1 | 1 | | 80901 | PPTG_18865 | Phytophthora parasitica INRA-310 hypothetical protein (740 aa) |
| 2920 | 3501 | PPTG_17097 | 39,68 | 3 | 0 | 3 | 3 | | 3,38E+06 | 5,22E+06 | 2 | 2 | 0 | 2 | 2 | | 67363 | PPTG_17097 | Phytophthora parasitica INRA-310 large subunit GTPase 1 (603 aa) |
| 3469 | 2940 | PPTG_04200 | 39,68 | 2 | 0 | 0 | 2 | | | 1,83E+05 | 2 | 2 | 0 | 0 | 2 | Carbamidomethylation | 104593 | PPTG_04200 | Phytophthora parasitica INRA-310 hypothetical protein (936 aa) |
| 3080 | 3698 | PPTG_07253 | 39,65 | 10 | 4 | 0 | 5 | 7,64E+05 | | 1,25E+06 | 2 | 2 | 1 | 0 | 2 | Carbamylation | 19639 | PPTG_07253 | Phytophthora parasitica INRA-310 hypothetical protein (186 aa) |
| 2953 | 4286 | PPTG_01707 | 39,65 | 14 | 0 | 14 | 14 | | 6,06E+05 | 2,05E+06 | 1 | 1 | 0 | 2 | 2 | | 15300 | PPTG_01707 | Phytophthora parasitica INRA-310 vesicle transporter GOT1B-like protein (140 aa) |
| 2834 | 3591 | PPTG_14840 | 39,63 | 12 | 5 | 12 | 7 | 8,70E+04 | 1,12E+06 | 2,58E+05 | 2 | 2 | 1 | 2 | 1 | Carbamidomethylation | 24671 | PPTG_14840 | Phytophthora parasitica INRA-310 hypothetical protein (236 aa) |
| 2759 | 3615 | PPTG_03871 | 39,63 | 8 | 0 | 8 | 5 | | 1,64E+05 | 3,37E+04 | 2 | 2 | 0 | 2 | 2 | Oxidation (M) | 70826 | PPTG_03871 | Phytophthora parasitica INRA-310 ATP-dependent DNA helicase 2 subunit (636 aa) |
| 4009 | 3854 | PPTG_11041 | 39,61 | 2 | 0 | 0 | 2 | | | 4,10E+05 | 1 | 1 | 0 | 0 | 1 | | 98409 | PPTG_11041 | Phytophthora parasitica INRA-310 voltage-gated Ion Channel (VIC) superfamily (887 aa) |
| 3667 | 3622 | PPTG_00389 | 39,61 | 7 | 0 | 5 | 3 | | 9,50E+04 | 3,27E+05 | 2 | 2 | 0 | 1 | 1 | | 36770 | PPTG_00389 | Phytophthora parasitica INRA-310 annexin (Annexin) family (330 aa) |
| 4402 | 4144 | PPTG_05413 | 39,6 | 3 | 0 | 0 | 3 | | | 1,02E+06 | 1 | 1 | 0 | 0 | 1 | | 35843 | PPTG_05413 | Phytophthora parasitica INRA-310 hypothetical protein (328 aa) |
| 3354 | 3286 | PPTG_04038 | 39,56 | 4 | 0 | 4 | 0 | | 2,47E+05 | | 2 | 2 | 0 | 2 | 0 | | 73991 | PPTG_04038 | Phytophthora parasitica INRA-310 pre-mRNA-processing factor 39 (655 aa) |
| 3267 | 3604 | PPTG_11753 | 39,56 | 8 | 0 | 5 | 8 | | 8,32E+04 | 1,47E+06 | 2 | 2 | 0 | 1 | 2 | Oxidation (M) | 46672 | PPTG_11753 | Phytophthora parasitica INRA-310 phosphatidate cytidylyltransferase (409 aa) |
| 2972 | 3023 | PPTG_02449 | 39,54 | 1 | 0 | 0 | 1 | | 3,44E+05 | 1,17E+06 | 2 | 2 | 0 | 1 | 2 | | 449386 | PPTG_02449 | Phytophthora parasitica INRA-310 fatty acid synthase subunit alpha (4129 aa) |
| 3476 | 3070 | PPTG_12411 | 39,52 | 3 | 0 | 0 | 3 | | | 5,93E+05 | 2 | 2 | 0 | 0 | 2 | Oxidation (M) | 96857 | PPTG_12411 | Phytophthora parasitica INRA-310 endoplasmic reticulum metallopeptidase (875 aa) |
| 2788 | 3642 | PPTG_07356 | 39,43 | 6 | 0 | 6 | 6 | | 4,53E+05 | 4,75E+05 | 1 | 1 | 0 | 2 | 2 | Carbamidomethylation | 34155 | PPTG_07356 | Phytophthora parasitica INRA-310 casein kinase II subunit beta (303 aa) |
| 2954 | 4290 | PPTG_01583 | 39,37 | 8 | 0 | 8 | 8 | | 4,79E+05 | 3,54E+05 | 1 | 1 | 0 | 2 | 2 | | 25289 | PPTG_01583 | Phytophthora parasitica INRA-310 protein kinase (223 aa) |
| 3710 | 4299 | PPTG_01721 | 39,37 | 3 | 0 | 3 | 3 | | 1,03E+06 | 7,75E+06 | 1 | 1 | 0 | 1 | 1 | | 29310 | PPTG_01721 | Phytophthora parasitica INRA-310 mannose-P-dolichol utilization defect 1 protein (272 aa) |
| 3001 | 3032 | PPTG_13163 | 39,36 | 16 | 0 | 10 | 6 | | 1,24E+06 | 1,00E+05 | 3 | 3 | 0 | 2 | 1 | Carbamidomethylation | 21574 | PPTG_13163 | Phytophthora parasitica INRA-310 N-terminal acetyltransferase complex ARD1 subunit (190 aa) |
| 3268 | 3650 | PPTG_19161 | 39,35 | 5 | 0 | 5 | 1 | | 2,24E+05 | 2,89E+05 | 2 | 2 | 0 | 2 | 1 | | 53759 | PPTG_19161 | Phytophthora parasitica INRA-310 hypothetical protein (468 aa) |
| 3716 | 4320 | PPTG_01089 | 39,34 | 9 | 0 | 9 | 9 | | 6,87E+05 | 4,54E+05 | 1 | 1 | 0 | 1 | 1 | | 17704 | PPTG_01089 | Phytophthora parasitica INRA-310 hypothetical protein (174 aa) |

| | | | | | | | | | | | | | | | | | |
|---|---|---|---|---|---|---|---|---|---|---|---|---|---|---|---|---|---|
| 4419 | 4291 | PPTG_13697 | 39,34 | 7 | 0 | 0 | 7 | | | 1,64E+05 | 1 | 1 | 0 | 0 | 1 | | 21848 \| PPTG_13697 \| Phytophthora parasitica INRA-310 metalloprotease family M76 (191 aa) |
| 3708 | 4292 | PPTG_05979 | 39,3 | 4 | 0 | 4 | 4 | | 2,06E+05 | 3,42E+05 | 1 | 1 | 0 | 1 | 1 | | 43011 \| PPTG_05979 \| Phytophthora parasitica INRA-310 hypothetical protein (390 aa) |
| 3431 | 3118 | PPTG_20238 | 39,3 | 9 | 3 | 0 | 6 | 1,04E+05 | | 1,81E+05 | 2 | 2 | 1 | 0 | 1 | | 34441 \| PPTG_20238 \| Phytophthora parasitica INRA-310 hypothetical protein (324 aa) |
| 4272 | 4105 | PPTG_20420 | 39,27 | 13 | 0 | 0 | 13 | | | 6,32E+05 | 1 | 1 | 0 | 0 | 1 | | 17186 \| PPTG_20420 \| Phytophthora parasitica INRA-310 hypothetical protein (159 aa) |
| 3330 | 3101 | PPTG_10097 | 39,22 | 0 | 0 | 0 | 0 | | | 1,93E+05 | 2 | 2 | 0 | 0 | 2 | | 507204 \| PPTG_10097 \| Phytophthora parasitica INRA-310 phosphatidylinositol kinase (PIK-L3) (4577 aa) |
| 3365 | 2945 | PPTG_16294 | 39,22 | 4 | 0 | 0 | 4 | | | 6,82E+05 | 2 | 2 | 0 | 0 | 2 | | 44585 \| PPTG_16294 \| Phytophthora parasitica INRA-310 sarcosine oxidase (407 aa) |
| 4609 | 4293 | PPTG_19616 | 39,19 | 8 | 0 | 0 | 8 | | | 3,17E+05 | 1 | 1 | 0 | 0 | 1 | | 18274 \| PPTG_19616 \| Phytophthora parasitica INRA-310 hypothetical protein (168 aa) |
| 3236 | 4323 | PPTG_08003 | 39,18 | 8 | 0 | 8 | 8 | | 8,15E+05 | 4,35E+05 | 1 | 1 | 0 | 2 | 1 | Deamidation (NQ) | 14575 \| PPTG_08003 \| Phytophthora parasitica INRA-310 hypothetical protein (128 aa) |
| 4241 | 3759 | PPTG_11763 | 39,18 | 1 | 0 | 0 | 1 | | | 2,85E+05 | 1 | 1 | 0 | 0 | 1 | | 136531 \| PPTG_11763 \| Phytophthora parasitica INRA-310 fatty acyl-CoA reductase (1217 aa) |
| 3511 | 3763 | PPTG_08835 | 39,18 | 3 | 0 | 3 | 3 | | 3,39E+05 | 3,92E+05 | 1 | 1 | 0 | 1 | 1 | | 34755 \| PPTG_08835 \| Phytophthora parasitica INRA-310 pantoate-beta-alanine ligase (315 aa) |
| 4611 | 4295 | PPTG_08284 | 39,16 | 3 | 0 | 0 | 3 | | | 1,76E+05 | 1 | 1 | 0 | 0 | 1 | | 48600 \| PPTG_08284 \| Phytophthora parasitica INRA-310 drug/Metabolite transporter (DMT) superfamily (430 aa) |
| 3054 | 3889 | PPTG_16215 | 39,16 | 2 | 0 | 2 | 2 | | 1,13E+06 | 3,94E+05 | 1 | 1 | 0 | 2 | 1 | Amidation | 92449 \| PPTG_16215 \| Phytophthora parasitica INRA-310 hypothetical protein (869 aa) |
| 4610 | 4294 | PPTG_14317 | 39,15 | 7 | 0 | 0 | 7 | | | 1,62E+05 | 1 | 1 | 0 | 0 | 1 | | 30681 \| PPTG_14317 \| Phytophthora parasitica INRA-310 inorganic pyrophosphatase (272 aa) |
| 3436 | 3534 | PPTG_00093 | 39,14 | 3 | 0 | 2 | 2 | | 5,72E+04 | 4,19E+05 | 2 | 2 | 0 | 1 | 1 | Carbamidomethylation | 73308 \| PPTG_00093 \| Phytophthora parasitica INRA-310 aspartyl-tRNA synthetase (656 aa) |
| 3691 | 4153 | PPTG_08318 | 39,1 | 6 | 6 | 6 | 0 | 4,45E+05 | 3,09E+06 | | 1 | 1 | 1 | 1 | 0 | | 19810 \| PPTG_08318 \| Phytophthora parasitica INRA-310 coatomer subunit zeta-1 (180 aa) |
| 3164 | 3657 | PPTG_18100 | 39,08 | 8 | 0 | 4 | 8 | | 1,21E+06 | 5,63E+06 | 2 | 2 | 0 | 1 | 2 | Carbamidomethylation | 23989 \| PPTG_18100 \| Phytophthora parasitica INRA-310 hypothetical protein (214 aa) |
| 2752 | 2971 | PPTG_08575 | 39,06 | 3 | 0 | 2 | 2 | | 2,49E+05 | 7,62E+05 | 3 | 3 | 0 | 2 | 2 | | 98534 \| PPTG_08575 \| Phytophthora parasitica INRA-310 nucleolar protein (869 aa) |
| 2997 | 2929 | PPTG_15179 | 39,06 | 7 | 0 | 0 | 7 | | | 7,60E+05 | 2 | 2 | 0 | 0 | 3 | Deamidation (NQ) | 37397 \| PPTG_15179 \| Phytophthora parasitica INRA-310 polygalacturonase (362 aa) |
| 2864 | 3026 | PPTG_03935 | 39,04 | 5 | 3 | 3 | 5 | 1,03E+05 | 1,28E+05 | 7,85E+05 | 2 | 2 | 1 | 1 | 2 | | 40314 \| PPTG_03935 \| Phytophthora parasitica INRA-310 annexin (Annexin) family (352 aa) |
| 3105 | 3048 | PPTG_18668 | 39,01 | 22 | 0 | 22 | 0 | | 5,88E+05 | | 3 | 3 | 0 | 3 | 0 | | 29838 \| PPTG_18668 \| Phytophthora parasitica INRA-310 exosome complex exonuclease RRP43 (284 aa) |
| 2681 | 3647 | PPTG_07912 | 39 | 9 | 0 | 9 | 9 | | 2,41E+06 | 7,17E+06 | 1 | 1 | 0 | 1 | 3 | Deamidation (NQ) | 11970 \| PPTG_07912 \| Phytophthora parasitica INRA-310 hypothetical protein (129 aa) |
| 3123 | 3588 | PPTG_14821 | 38,99 | 2 | 0 | 2 | 1 | | 3,32E+05 | 3,17E+05 | 2 | 2 | 0 | 2 | 1 | | 125289 \| PPTG_14821 \| Phytophthora parasitica INRA-310 hypothetical protein (1146 aa) |
| 4392 | 3945 | PPTG_09108 | 38,99 | 4 | 0 | 0 | 4 | | | 2,45E+05 | 1 | 1 | 0 | 0 | 1 | | 40066 \| PPTG_09108 \| Phytophthora parasitica INRA-310 hypothetical protein (357 aa) |
| 3692 | 4160 | PPTG_02701 | 38,97 | 6 | 0 | 6 | 6 | | 4,85E+05 | 2,20E+05 | 1 | 1 | 0 | 1 | 1 | | 22384 \| PPTG_02701 \| Phytophthora parasitica INRA-310 hypothetical protein (199 aa) |
| 3607 | 4164 | PPTG_08143 | 38,96 | 11 | 0 | 11 | 11 | | 9,71E+05 | 1,36E+06 | 1 | 1 | 0 | 1 | 1 | | 12465 \| PPTG_08143 \| Phytophthora parasitica INRA-310 monooxygenase (107 aa) |
| 2924 | 3626 | PPTG_04261 | 38,94 | 4 | 0 | 4 | 4 | | 1,19E+06 | 9,61E+05 | 2 | 2 | 0 | 2 | 2 | | 55492 \| PPTG_04261 \| Phytophthora parasitica INRA-310 serine protease family S10 (497 aa) |
| 3505 | 3674 | PPTG_15574 | 38,93 | 5 | 0 | 5 | 5 | | 6,60E+04 | 2,92E+05 | 1 | 1 | 0 | 1 | 1 | | 33746 \| PPTG_15574 \| Phytophthora parasitica INRA-310 hypothetical protein (313 aa) |
| 3773 | 3618 | PPTG_06418 | 38,85 | 4 | 0 | 0 | 4 | | | 2,91E+05 | 2 | 2 | 0 | 0 | 2 | | 72117 \| PPTG_06418 \| Phytophthora parasitica INRA-310 hypothetical protein (644 aa) |
| 2639 | 3640 | PPTG_01702 | 38,83 | 8 | 4 | 8 | 8 | 1,01E+05 | 7,57E+05 | 1,26E+06 | 2 | 2 | 1 | 2 | 2 | | 28931 \| PPTG_01702 \| Phytophthora parasitica INRA-310 hypothetical protein (251 aa) |
| 3125 | 3600 | PPTG_11586 | 38,81 | 7 | 0 | 3 | 7 | | 1,75E+05 | 1,42E+05 | 2 | 2 | 0 | 1 | 2 | | 31783 \| PPTG_11586 \| Phytophthora parasitica INRA-310 sepiapterin reductase (287 aa) |
| 3159 | 3532 | PPTG_04845 | 38,81 | 14 | 0 | 4 | 14 | | 6,52E+05 | 1,54E+06 | 2 | 2 | 0 | 1 | 2 | Carbamidomethylation | 26592 \| PPTG_04845 \| Phytophthora parasitica INRA-310 mRNA turnover protein 4 (235 aa) |
| 4615 | 4327 | PPTG_07325 | 38,72 | 19 | 0 | 19 | 0 | | 7,64E+04 | | 1 | 1 | 0 | 1 | 0 | | 9112 \| PPTG_07325 \| Phytophthora parasitica INRA-310 hypothetical protein (85 aa) |
| 3189 | 3057 | PPTG_10637 | 38,67 | 10 | 0 | 4 | 10 | | 7,91E+04 | 8,10E+05 | 2 | 2 | 0 | 1 | 2 | Carbamidomethylation | 36420 \| PPTG_10637 \| Phytophthora parasitica INRA-310 peptidyl-prolyl cis-trans isomerase B (335 aa) |
| 3204 | 3549 | PPTG_12114 | 38,65 | 4 | 0 | 2 | 4 | | 2,63E+05 | 9,49E+05 | 2 | 2 | 0 | 1 | 2 | | 53945 \| PPTG_12114 \| Phytophthora parasitica INRA-310 glycoside hydrolase (591 aa) |
| 3496 | 3566 | PPTG_04945 | 38,64 | 4 | 0 | 0 | 4 | | | 2,03E+05 | 2 | 2 | 0 | 0 | 2 | | 56126 \| PPTG_04945 \| Phytophthora parasitica INRA-310 hypothetical protein (496 aa) |
| 4273 | 4107 | PPTG_00814 | 38,64 | 9 | 0 | 0 | 9 | | | 5,90E+05 | 1 | 1 | 0 | 0 | 1 | | 18699 \| PPTG_00814 \| Phytophthora parasitica INRA-310 AP-2 complex subunit sigma (163 aa) |
| 3603 | 4102 | PPTG_03684 | 38,61 | 2 | 0 | 2 | 2 | | 0 | 3,43E+05 | 1 | 1 | 0 | 1 | 1 | | 77294 \| PPTG_03684 \| Phytophthora parasitica INRA-310 hypothetical protein (697 aa) |
| 3210 | 3619 | PPTG_14841 | 38,6 | 2 | 0 | 2 | 1 | | 2,59E+05 | 2,17E+05 | 2 | 2 | 0 | 2 | 1 | | 115044 \| PPTG_14841 \| Phytophthora parasitica INRA-310 hypothetical protein (1031 aa) |
| 2891 | 4125 | PPTG_01744 | 38,6 | 10 | 0 | 10 | 10 | | 5,68E+05 | 1,75E+06 | 1 | 1 | 0 | 1 | 3 | Oxidation (M) | 18835 \| PPTG_01744 \| Phytophthora parasitica INRA-310 U6 snRNA-associated Sm-like protein (173 aa) |
| 3564 | 3520 | PPTG_12155 | 38,58 | 4 | 0 | 0 | 4 | | | 6,74E+04 | 2 | 2 | 0 | 0 | 2 | | 67361 \| PPTG_12155 \| Phytophthora parasitica INRA-310 cleft lip and palate transmembrane family protein (596 aa) |
| 2832 | 3573 | PPTG_04880 | 38,57 | 4 | 3 | 4 | 3 | 5,86E+04 | 1,25E+06 | 3,26E+05 | 2 | 2 | 1 | 2 | 1 | | 56843 \| PPTG_04880 \| Phytophthora parasitica INRA-310 U3 small nucleolar RNA-associated protein (515 aa) |
| 3137 | 2939 | PPTG_08108 | 38,55 | 1 | 0 | 1 | 1 | | 2,77E+05 | 2,28E+05 | 2 | 2 | 0 | 2 | 1 | | 210606 \| PPTG_08108 \| Phytophthora parasitica INRA-310 hypothetical protein (1902 aa) |
| 3640 | 3071 | PPTG_09646 | 38,54 | 5 | 0 | 0 | 5 | | | 1,22E+06 | 2 | 2 | 0 | 0 | 2 | Carbamidomethylation | 48976 \| PPTG_09646 \| Phytophthora parasitica INRA-310 animal Inward Rectifier K Channel (IRK-C) family (436 aa) |
| 3659 | 3542 | PPTG_03850 | 38,51 | 3 | 0 | 0 | 3 | | | 2,07E+05 | 2 | 2 | 0 | 0 | 2 | | 78017 \| PPTG_03850 \| Phytophthora parasitica INRA-310 hypothetical protein (698 aa) |
| 3487 | 3338 | PPTG_17898 | 38,5 | 7 | 0 | 3 | 4 | | 2,21E+05 | 6,06E+05 | 2 | 2 | 0 | 1 | 1 | Carbamidomethylation | 35582 \| PPTG_17898 \| Phytophthora parasitica INRA-310 hypothetical protein (338 aa) |
| 3280 | 4301 | PPTG_06441 | 38,49 | 8 | 8 | 8 | 8 | 5,07E+05 | 3,31E+05 | 3,46E+06 | 1 | 1 | 1 | 1 | 1 | | 13173 \| PPTG_06441 \| Phytophthora parasitica INRA-310 hypothetical protein (116 aa) |
| 3444 | 3787 | PPTG_18749 | 38,48 | 4 | 0 | 4 | 4 | | 5,03E+05 | 0 | 1 | 1 | 0 | 1 | 1 | | 31301 \| PPTG_18749 \| Phytophthora parasitica INRA-310 phosphatidylinositol transfer protein beta isoform (274 aa) |
| 3711 | 4300 | PPTG_13217 | 38,47 | 14 | 0 | 0 | 14 | | | 3,46E+06 | 1 | 1 | 0 | 0 | 2 | | 10208 \| PPTG_13217 \| Phytophthora parasitica INRA-310 hypothetical protein (92 aa) |
| 3587 | 3990 | PPTG_13888 | 38,38 | 8 | 0 | 8 | 8 | | 4,84E+05 | 4,05E+05 | 1 | 1 | 0 | 1 | 1 | | 25046 \| PPTG_13888 \| Phytophthora parasitica INRA-310 COP9 signalosome complex subunit (227 aa) |
| 4423 | 4328 | PPTG_08956 | 38,33 | 9 | 0 | 9 | 0 | | 2,03E+05 | | 1 | 1 | 0 | 1 | 0 | | 19898 \| PPTG_08956 \| Phytophthora parasitica INRA-310 hypothetical protein (174 aa) |
| 3599 | 4071 | PPTG_12108 | 38,31 | 10 | 0 | 10 | 10 | | 2,17E+05 | 1,09E+06 | 1 | 1 | 0 | 1 | 1 | | 18301 \| PPTG_12108 \| Phytophthora parasitica INRA-310 hypothetical protein (161 aa) |
| 3712 | 4303 | PPTG_17623 | 38,3 | 7 | 0 | 7 | 7 | | 7,85E+05 | 5,11E+06 | 1 | 1 | 0 | 1 | 1 | | 17659 \| PPTG_17623 \| Phytophthora parasitica INRA-310 hypothetical protein (163 aa) |
| 3139 | 2983 | PPTG_12115 | 38,3 | 2 | 0 | 1 | 2 | | 2,58E+05 | 6,21E+05 | 2 | 2 | 0 | 1 | 2 | | 101016 \| PPTG_12115 \| Phytophthora parasitica INRA-310 leucyl-tRNA synthetase (905 aa) |
| 3792 | 4302 | PPTG_01553 | 38,29 | 6 | 0 | 6 | 6 | | 7,99E+04 | 4,74E+05 | 1 | 1 | 0 | 1 | 1 | | 25674 \| PPTG_01553 \| Phytophthora parasitica INRA-310 hypothetical protein (230 aa) |
| 3687 | 4121 | PPTG_18277 | 38,28 | 7 | 0 | 7 | 7 | | 9,00E+04 | 1,12E+05 | 1 | 1 | 0 | 1 | 1 | | 24153 \| PPTG_18277 \| Phytophthora parasitica INRA-310 polyadenylate-binding protein 2 (226 aa) |
| 3405 | 3564 | PPTG_11778 | 38,26 | 12 | 0 | 5 | 7 | | 1,56E+05 | 2,11E+05 | 2 | 2 | 0 | 1 | 1 | | 25071 \| PPTG_11778 \| Phytophthora parasitica INRA-310 hypothetical protein (231 aa) |
| 3273 | 3722 | PPTG_09472 | 38,19 | 3 | 0 | 3 | 2 | | 3,30E+05 | 1,58E+05 | 2 | 2 | 0 | 2 | 1 | | 95616 \| PPTG_09472 \| Phytophthora parasitica INRA-310 replication factor C subunit (862 aa) |
| 4616 | 4329 | PPTG_12117 | 38,19 | 4 | 0 | 4 | 0 | | 3,75E+05 | | 1 | 1 | 0 | 1 | 0 | | 35216 \| PPTG_12117 \| Phytophthora parasitica INRA-310 hypothetical protein (318 aa) |
| 4097 | 3909 | PPTG_04210 | 38,18 | 2 | 0 | 0 | 2 | | | 1,37E+05 | 1 | 1 | 0 | 0 | 1 | | 99408 \| PPTG_04210 \| Phytophthora parasitica INRA-310 GPI inositol-deacylase (906 aa) |
| 3583 | 3938 | PPTG_16557 | 38,12 | 5 | 0 | 5 | 5 | | 2,09E+05 | 4,10E+04 | 1 | 1 | 0 | 1 | 1 | Carbamidomethylation | 39009 \| PPTG_16557 \| Phytophthora parasitica INRA-310 hypothetical protein (361 aa) |
| 4056 | 3911 | PPTG_10502 | 38,11 | 2 | 0 | 0 | 2 | | | 1,61E+05 | 1 | 1 | 0 | 0 | 1 | | 95891 \| PPTG_10502 \| Phytophthora parasitica INRA-310 hypothetical protein (859 aa) |
| 4166 | 3903 | PPTG_15202 | 38,07 | 5 | 0 | 0 | 5 | | | 3,94E+05 | 1 | 1 | 0 | 0 | 1 | | 26169 \| PPTG_15202 \| Phytophthora parasitica INRA-310 hypothetical protein (241 aa) |
| 3415 | 3953 | PPTG_04320 | 38,06 | 2 | 0 | 2 | 2 | | 4,76E+05 | 9,92E+05 | 1 | 1 | 0 | 1 | 1 | | 47156 \| PPTG_04320 \| Phytophthora parasitica INRA-310 hypothetical protein (407 aa) |
| 3774 | 3648 | PPTG_11008 | 38,06 | 5 | 0 | 0 | 5 | | | 1,35E+05 | 2 | 2 | 0 | 0 | 2 | Carbamidomethylation | 64795 \| PPTG_11008 \| Phytophthora parasitica INRA-310 hypothetical protein (594 aa) |
| 3163 | 3656 | PPTG_16637 | 38,02 | 2 | 0 | 2 | 1 | | 4,07E+05 | 4,84E+05 | 2 | 2 | 0 | 2 | 1 | | 76096 \| PPTG_16637 \| Phytophthora parasitica INRA-310 U3 small nucleolar RNA-associated protein 6 (673 aa) |
| 3218 | 3703 | PPTG_12691 | 37,99 | 9 | 0 | 9 | 6 | | 3,58E+05 | 0 | 2 | 2 | 0 | 2 | 1 | | 22068 \| PPTG_12691 \| Phytophthora parasitica INRA-310 hypothetical protein (194 aa) |
| 3516 | 3848 | PPTG_18714 | 37,98 | 5 | 0 | 5 | 5 | | 8,25E+05 | 1,03E+06 | 1 | 1 | 0 | 1 | 1 | | 26384 \| PPTG_18714 \| Phytophthora parasitica INRA-310 hypothetical protein (226 aa) |
| 3590 | 4002 | PPTG_18883 | 37,97 | 3 | 0 | 3 | 3 | | 1,22E+05 | 1,52E+05 | 1 | 1 | 0 | 1 | 1 | | 47899 \| PPTG_18883 \| Phytophthora parasitica INRA-310 hypothetical protein (435 aa) |
| 4420 | 4305 | PPTG_10938 | 37,93 | 2 | 0 | 0 | 2 | | | 0 | 1 | 1 | 0 | 0 | 1 | | 58389 \| PPTG_10938 \| Phytophthora parasitica INRA-310 major facilitator superfamily transporter (MFS) (540 aa) |
| 2778 | 2944 | PPTG_17395 | 37,89 | 5 | 0 | 2 | 5 | | 2,75E+05 | 5,62E+05 | 3 | 3 | 0 | 1 | 3 | | 49827 \| PPTG_17395 \| Phytophthora parasitica INRA-310 hypothetical protein (444 aa) |
| 3003 | 3113 | PPTG_00381 | 37,89 | 3 | 0 | 3 | 1 | | 1,02E+06 | | 2 | 1 | 0 | 2 | 1 | | 64020 \| PPTG_00381 \| Phytophthora parasitica INRA-310 RPS6 protein kinase (567 aa) |
| 3690 | 4141 | PPTG_04428 | 37,83 | 14 | 0 | 14 | 14 | | 3,67E+05 | 2,35E+05 | 1 | 1 | 0 | 1 | 1 | | 10831 \| PPTG_04428 \| Phytophthora parasitica INRA-310 hypothetical protein (99 aa) |
| 3498 | 3581 | PPTG_10288 | 37,81 | 3 | 0 | 0 | 3 | | | 9,91E+05 | 2 | 2 | 0 | 0 | 2 | | 81789 \| PPTG_10288 \| Phytophthora parasitica INRA-310 ATPase AFG2 protein (752 aa) |
| 3066 | 3053 | PPTG_19614 | 37,8 | 4 | 2 | 0 | 4 | 6,68E+04 | | 8,65E+05 | 2 | 2 | 1 | 0 | 2 | | 62071 \| PPTG_19614 \| Phytophthora parasitica INRA-310 protein kinase (565 aa) |
| 4612 | 4312 | PPTG_00878 | 37,73 | 4 | 0 | 0 | 4 | | | 4,22E+05 | 1 | 1 | 0 | 0 | 1 | | 31026 \| PPTG_00878 \| Phytophthora parasitica INRA-310 hypothetical protein (282 aa) |
| 4613 | 4313 | PPTG_09271 | 37,71 | 19 | 0 | 0 | 19 | | | 0 | 1 | 1 | 0 | 0 | 1 | | 7966 \| PPTG_09271 \| Phytophthora parasitica INRA-310 hypothetical protein (70 aa) |
| 3478 | 3105 | PPTG_16594 | 37,68 | 2 | 0 | 0 | 2 | | | 3,54E+05 | 2 | 2 | 0 | 0 | 2 | | 129764 \| PPTG_16594 \| Phytophthora parasitica INRA-310 myotubularin-like protein (1159 aa) |
| 3541 | 2979 | PPTG_08209 | 37,68 | 5 | 0 | 0 | 5 | | | 5,38E+05 | 2 | 2 | 0 | 0 | 2 | | 45009 \| PPTG_08209 \| Phytophthora parasitica INRA-310 protein phosphatase 2C (421 aa) |
| 3270 | 3682 | PPTG_04029 | 37,68 | 8 | 0 | 4 | 8 | | 1,23E+05 | 5,74E+05 | 2 | 2 | 0 | 1 | 2 | Carbamidomethylation | 26500 \| PPTG_04029 \| Phytophthora parasitica INRA-310 phosphoglycerate mutase (234 aa) |
| 3082 | 3760 | PPTG_02648 | 37,66 | 3 | 3 | 3 | 3 | 9,93E+04 | 4,30E+05 | 9,95E+05 | 1 | 1 | 1 | 1 | 1 | | 36054 \| PPTG_02648 \| Phytophthora parasitica INRA-310 hypothetical protein (327 aa) |
| 3713 | 4315 | PPTG_18885 | 37,64 | 3 | 0 | 3 | 3 | | 4,83E+05 | 4,48E+05 | 1 | 1 | 0 | 1 | 1 | | 49689 \| PPTG_18885 \| Phytophthora parasitica INRA-310 hypothetical protein (456 aa) |
| 3568 | 3629 | PPTG_02756 | 37,64 | 6 | 0 | 6 | 0 | | 2,47E+05 | | 2 | 2 | 0 | 2 | 0 | | 38351 \| PPTG_02756 \| Phytophthora parasitica INRA-310 O-sialoglycoprotein endopeptidase (351 aa) |
| 3272 | 3701 | PPTG_19357 | 37,62 | 16 | 0 | 8 | 16 | | 1,20E+06 | 1,95E+06 | 2 | 2 | 0 | 1 | 2 | | 15075 \| PPTG_19357 \| Phytophthora parasitica INRA-310 hypothetical protein (139 aa) |
| 4086 | 3749 | PPTG_01764 | 37,61 | 3 | 0 | 0 | 3 | | | 1,08E+05 | 1 | 1 | 0 | 0 | 1 | | 56574 \| PPTG_01764 \| Phytophthora parasitica INRA-310 hypothetical protein (499 aa) |
| 4161 | 3805 | PPTG_06687 | 37,61 | 3 | 0 | 0 | 3 | | | 2,38E+05 | 1 | 1 | 0 | 0 | 1 | | 43345 \| PPTG_06687 \| Phytophthora parasitica INRA-310 hypothetical protein (389 aa) |
| 3606 | 4158 | PPTG_18705 | 37,6 | 8 | 0 | 8 | 8 | | 6,63E+05 | 7,84E+05 | 1 | 1 | 0 | 1 | 1 | | 15658 \| PPTG_18705 \| Phytophthora parasitica INRA-310 outer Dynein Arm Light Chain 2 (137 aa) |
| 4088 | 3808 | PPTG_15680 | 37,56 | 5 | 0 | 0 | 5 | | | 3,96E+05 | 1 | 1 | 0 | 0 | 1 | | 28116 \| PPTG_15680 \| Phytophthora parasitica INRA-310 hypothetical protein (241 aa) |
| 3429 | 3072 | PPTG_13438 | 37,53 | 5 | 0 | 0 | 5 | | | 6,70E+05 | 2 | 2 | 0 | 0 | 2 | Carbamidomethylation | 49738 \| PPTG_13438 \| Phytophthora parasitica INRA-310 alanyl-tRNA synthetase domain-containing protein 1 (461 aa) |
| 3674 | 3686 | PPTG_17929 | 37,52 | 2 | 0 | 0 | 2 | | | 7,92E+05 | 2 | 1 | 0 | 0 | 2 | | 75693 \| PPTG_17929 \| Phytophthora parasitica INRA-310 voltage-gated Ion Channel (VIC) superfamily (669 aa) |
| 3674 | 3687 | PPTG_17929 | 37,52 | 2 | 0 | 0 | 2 | | | 7,92E+05 | 2 | 1 | 0 | 0 | 2 | | 78197 \| PPTG_17929 \| Phytophthora parasitica INRA-310 voltage-gated Ion Channel (VIC) superfamily (692 aa) |
| 3600 | 4075 | PPTG_12087 | 37,48 | 5 | 0 | 5 | 5 | | 1,31E+05 | 2,45E+05 | 1 | 1 | 0 | 1 | 1 | | 24010 \| PPTG_12087 \| Phytophthora parasitica INRA-310 aldo/keto reductase family protein (216 aa) |
| 3580 | 3844 | PPTG_07056 | 37,48 | 2 | 0 | 2 | 2 | | 1,58E+05 | 2,93E+05 | 1 | 1 | 0 | 1 | 1 | | 73726 \| PPTG_07056 \| Phytophthora parasitica INRA-310 hypothetical protein (661 aa) |
| 4159 | 3778 | PPTG_16498 | 37,47 | 9 | 0 | 0 | 9 | | | 3,31E+05 | 1 | 1 | 0 | 0 | 1 | | 15367 \| PPTG_16498 \| Phytophthora parasitica INRA-310 hypothetical protein (131 aa) |
| 3235 | 4317 | PPTG_02826 | 37,45 | 12 | 0 | 12 | 12 | | 4,40E+05 | 1,83E+06 | 1 | 1 | 0 | 1 | 2 | Oxidation (M) | 10696 \| PPTG_02826 \| Phytophthora parasitica INRA-310 small nuclear ribonucleoprotein E (93 aa) |
| 3715 | 4318 | PPTG_13160 | 37,44 | 2 | 0 | 2 | 2 | | 0 | 1,00E+06 | 1 | 1 | 0 | 1 | 1 | | 51395 \| PPTG_13160 \| Phytophthora parasitica INRA-310 glycoside-Pentoside-Hexuronide (GPH):Cation Symporter family (477 aa) |
| 3371 | 3093 | PPTG_16548 | 37,43 | 3 | 0 | 0 | 3 | | | 8,99E+05 | 2 | 2 | 0 | 0 | 2 | | 77346 \| PPTG_16548 \| Phytophthora parasitica INRA-310 metalloprotease family M20D (698 aa) |
| 3414 | 3952 | PPTG_02611 | 37,42 | 1 | 1 | 0 | 1 | 3,94E+04 | | 3,62E+05 | 1 | 1 | 1 | 0 | 1 | | 65710 \| PPTG_02611 \| Phytophthora parasitica INRA-310 hypothetical protein (587 aa) |
| 3672 | 3679 | PPTG_15648 | 37,42 | 16 | 0 | 0 | 16 | | | 1,47E+05 | 2 | 2 | 0 | 0 | 2 | | 23179 \| PPTG_15648 \| Phytophthora parasitica INRA-310 charged multivesicular body protein 1b-2 (210 aa) |
| 3514 | 3820 | PPTG_13070 | 37,38 | 2 | 0 | 2 | 2 | | 2,72E+05 | 1,17E+06 | 1 | 1 | 0 | 1 | 1 | | 75145 \| PPTG_13070 \| Phytophthora parasitica INRA-310 hypothetical protein (683 aa) |
| 3437 | 3539 | PPTG_13364 | 37,35 | 0 | 0 | 0 | 0 | | | 2,81E+05 | 2 | 2 | 0 | 0 | 2 | | 521073 \| PPTG_13364 \| Phytophthora parasitica INRA-310 vacuolar protein sorting-associated protein (4683 aa) |
| 4422 | 4319 | PPTG_11794 | 37,31 | 4 | 0 | 0 | 4 | | | 4,73E+05 | 1 | 1 | 0 | 0 | 1 | | 29899 \| PPTG_11794 \| Phytophthora parasitica INRA-310 hypothetical protein (313 aa) |
| 3605 | 4145 | PPTG_10191 | 37,31 | 12 | 0 | 12 | 12 | | 2,91E+05 | 2,68E+05 | 1 | 1 | 0 | 1 | 1 | | 12683 \| PPTG_10191 \| Phytophthora parasitica INRA-310 dynein light chain (111 aa) |
| 3719 | 4339 | PPTG_09960 | 37,3 | 4 | 0 | 4 | 4 | | 5,61E+05 | 7,42E+05 | 1 | 1 | 0 | 1 | 1 | | 28943 \| PPTG_09960 \| Phytophthora parasitica INRA-310 hypothetical protein (275 aa) |
| 3569 | 3633 | PPTG_00877 | 37,23 | 3 | 0 | 0 | 3 | | | 2,15E+05 | 2 | 2 | 0 | 0 | 2 | | 98803 \| PPTG_00877 \| Phytophthora parasitica INRA-310 drug/Metabolite transporter (DMT) superfamily (918 aa) |
| 3492 | 3516 | PPTG_16784 | 37,16 | 13 | 0 | 0 | 13 | | | 7,95E+05 | 2 | 2 | 0 | 0 | 2 | | 21989 \| PPTG_16784 \| Phytophthora parasitica INRA-310 hypothetical protein (196 aa) |
| 3696 | 4178 | PPTG_04023 | 37,16 | 1 | 0 | 1 | 1 | | 8,59E+04 | 1,28E+05 | 1 | 1 | 0 | 1 | 1 | | 150863 \| PPTG_04023 \| Phytophthora parasitica INRA-310 DIS3-like exonuclease 1 (1355 aa) |

| | | | | | | | | | | | | | | | | | | | |
|---|---|---|---|---|---|---|---|---|---|---|---|---|---|---|---|---|---|---|---|
| 3666 | 3612 | PPTG_16517 | 37,15 | 2 | 0 | 0 | 2 | | | 3,83E+05 | 2 | 2 | 0 | 0 | 2 | | 110854 | PPTG_16517 | Phytophthora parasitica INRA-310 voltage-gated Ion Channel (VIC) superfamily (1004 aa) |
| 3342 | 3605 | PPTG_18746 | 37,11 | 2 | 0 | 2 | 2 | | 6,96E+05 | 1,67E+06 | 1 | 1 | 0 | 1 | 1 | | 60393 | PPTG_18746 | Phytophthora parasitica INRA-310 sorting nexin (546 aa) |
| 3717 | 4322 | PPTG_10123 | 37,09 | 4 | 0 | 4 | 4 | | 4,28E+05 | 1,65E+06 | 1 | 1 | 0 | 1 | 1 | | 29830 | PPTG_10123 | Phytophthora parasitica INRA-310 hypothetical protein (267 aa) |
| 4620 | 4353 | PPTG_08244 | 37,07 | 4 | 0 | 4 | 0 | | 1,57E+05 | | 1 | 1 | 0 | 1 | 0 | | 35006 | PPTG_08244 | Phytophthora parasitica INRA-310 hypothetical protein (313 aa) |
| 3778 | 3708 | PPTG_13921 | 37,02 | 2 | 0 | 1 | 1 | | 1,72E+05 | 0 | 2 | 2 | 0 | 1 | 1 | | 129236 | PPTG_13921 | Phytophthora parasitica INRA-310 hypothetical protein (1162 aa) |
| 3141 | 3108 | PPTG_11178 | 36,98 | 5 | 0 | 2 | 5 | | 5,20E+05 | 1,27E+06 | 2 | 2 | 0 | 1 | 2 | Carbamidomethylation | 39493 | PPTG_11178 | Phytophthora parasitica INRA-310 hypothetical protein (353 aa) |
| 3675 | 3724 | PPTG_05709 | 36,97 | 7 | 0 | 7 | 0 | | 1,88E+05 | | 2 | 2 | 0 | 2 | 0 | Carbamidomethylation | 42672 | PPTG_05709 | Phytophthora parasitica INRA-310 hypothetical protein (378 aa) |
| 3675 | 3725 | PPTG_05709 | 36,97 | 5 | 0 | 5 | 0 | | 1,88E+05 | | 2 | 2 | 0 | 2 | 0 | Carbamidomethylation | 51972 | PPTG_05709 | Phytophthora parasitica INRA-310 hypothetical protein (472 aa) |
| 3592 | 4009 | PPTG_20410 | 36,89 | 3 | 0 | 3 | 3 | | 1,93E+05 | 2,97E+05 | 1 | 1 | 0 | 1 | 1 | | 41401 | PPTG_20410 | Phytophthora parasitica INRA-310 hypothetical protein (381 aa) |
| 4050 | 3771 | PPTG_01571 | 36,88 | 3 | 0 | 3 | 0 | | 1,08E+05 | | 1 | 1 | 0 | 1 | 0 | Carbamidomethylation | 75471 | PPTG_01571 | Phytophthora parasitica INRA-310 Nup85 Nucleoporin (664 aa) |
| 4184 | 4156 | PPTG_00477 | 36,88 | 10 | 0 | 0 | 10 | | | 5,61E+05 | 1 | 1 | 0 | 0 | 1 | | 14064 | PPTG_00477 | Phytophthora parasitica INRA-310 pre-mRNA branch site p14-like protein (123 aa) |
| 3604 | 4128 | PPTG_01290 | 36,87 | 1 | 0 | 1 | 1 | | 1,68E+05 | 3,67E+05 | 1 | 1 | 0 | 1 | 1 | | 118430 | PPTG_01290 | Phytophthora parasitica INRA-310 serine/threonine protein kinase (1102 aa) |
| 4614 | 4325 | PPTG_02914 | 36,84 | 13 | 0 | 0 | 13 | | | 3,85E+04 | 1 | 1 | 0 | 0 | 1 | Carbamidomethylation | 25097 | PPTG_02914 | Phytophthora parasitica INRA-310 hypothetical protein (229 aa) |
| 3718 | 4326 | PPTG_10532 | 36,84 | 3 | 0 | 3 | 3 | | 1,41E+05 | 3,77E+05 | 1 | 1 | 0 | 1 | 1 | | 44514 | PPTG_10532 | Phytophthora parasitica INRA-310 hypothetical protein (416 aa) |
| 4427 | 4354 | PPTG_14188 | 36,83 | 3 | 0 | 3 | 0 | | 8,43E+04 | | 1 | 1 | 0 | 1 | 0 | | 48645 | PPTG_14188 | Phytophthora parasitica INRA-310 hypothetical protein (432 aa) |
| 3378 | 3620 | PPTG_05486 | 36,78 | 2 | 0 | 1 | 1 | | 2,47E+05 | 0 | 2 | 2 | 0 | 1 | 1 | | 146575 | PPTG_05486 | Phytophthora parasitica INRA-310 hypothetical protein (1307 aa) |
| 4098 | 3914 | PPTG_10754 | 36,78 | 1 | 0 | 0 | 1 | | | 1,11E+05 | 1 | 1 | 0 | 0 | 1 | | 129836 | PPTG_10754 | Phytophthora parasitica INRA-310 patatin like protein (1173 aa) |
| 2927 | 3728 | PPTG_01052 | 36,76 | 5 | 0 | 5 | 5 | | 4,68E+05 | 6,91E+05 | 2 | 2 | 0 | 2 | 2 | | 51699 | PPTG_01052 | Phytophthora parasitica INRA-310 histone deacetylase (462 aa) |
| 3432 | 3144 | PPTG_05324 | 36,75 | 3 | 0 | 0 | 3 | | | 1,77E+05 | 2 | 1 | 0 | 0 | 2 | Carbamidomethylation | 76094 | PPTG_05324 | Phytophthora parasitica INRA-310 HECT E3 ubiquitin ligase (673 aa) |
| 3570 | 3654 | PPTG_16399 | 36,69 | 9 | 0 | 5 | 4 | | 1,30E+05 | 0 | 2 | 2 | 0 | 1 | 1 | Carbamidomethylation | 24863 | PPTG_16399 | Phytophthora parasitica INRA-310 5-formyltetrahydrofolate cyclo-ligase (229 aa) |
| 4026 | 3935 | PPTG_07526 | 36,66 | 3 | 0 | 0 | 3 | | | 1,23E+06 | 1 | 1 | 0 | 0 | 1 | | 44755 | PPTG_07526 | Phytophthora parasitica INRA-310 hypothetical protein (394 aa) |
| 2926 | 3719 | PPTG_09948 | 36,64 | 18 | 7 | 7 | 18 | 1,49E+06 | 2,04E+06 | 1,55E+07 | 2 | 2 | 1 | 1 | 2 | | 13372 | PPTG_09948 | Phytophthora parasitica INRA-310 hypothetical protein (120 aa) |
| 3165 | 3695 | PPTG_04898 | 36,59 | 2 | 0 | 2 | 1 | | 3,97E+05 | 3,80E+05 | 2 | 2 | 0 | 2 | 1 | | 86379 | PPTG_04898 | Phytophthora parasitica INRA-310 hypothetical protein (789 aa) |
| 4183 | 4146 | PPTG_08586 | 36,57 | 4 | 0 | 4 | 0 | | 4,07E+05 | | 1 | 1 | 0 | 1 | 0 | | 35222 | PPTG_08586 | Phytophthora parasitica INRA-310 aldose 1-epimerase (320 aa) |
| 3576 | 3741 | PPTG_18271 | 36,57 | 1 | 0 | 1 | 0 | | 4,78E+04 | 4,06E+05 | 2 | 2 | 0 | 1 | 1 | | 253916 | PPTG_18271 | Phytophthora parasitica INRA-310 phytophthora infestans avirulence-associated protein 3,4F-A (2284 aa) |
| 3129 | 3948 | PPTG_06396 | 36,54 | 7 | 0 | 0 | 7 | | | 2,59E+04 | 1 | 1 | 0 | 0 | 3 | Deamidation (NQ) | 52922 | PPTG_06396 | Phytophthora parasitica INRA-310 hypothetical protein (478 aa) |
| 4403 | 4149 | PPTG_18428 | 36,51 | 3 | 0 | 0 | 3 | | | 0 | 1 | 1 | 0 | 0 | 1 | | 47280 | PPTG_18428 | Phytophthora parasitica INRA-310 hypothetical protein (430 aa) |
| 4289 | 4355 | PPTG_00895 | 36,51 | 4 | 0 | 4 | 0 | | 3,17E+05 | | 1 | 1 | 0 | 1 | 0 | | 28362 | PPTG_00895 | Phytophthora parasitica INRA-310 hypothetical protein (261 aa) |
| 3499 | 3597 | PPTG_09761 | 36,49 | 2 | 0 | 0 | 2 | | | 2,33E+05 | 2 | 2 | 0 | 0 | 2 | | 107840 | PPTG_09761 | Phytophthora parasitica INRA-310 transmembrane protein (964 aa) |
| 3351 | 3134 | PPTG_08611 | 36,45 | 2 | 0 | 1 | 1 | | 4,57E+05 | 7,92E+05 | 2 | 2 | 0 | 1 | 1 | | 64392 | PPTG_08611 | Phytophthora parasitica INRA-310 SNF1-like protein kinase catalytic subunit alpha (572 aa) |
| 4394 | 4016 | PPTG_00304 | 36,41 | 6 | 0 | 6 | 0 | | 6,37E+05 | | 1 | 1 | 0 | 1 | 0 | Carbamidomethylation | 27990 | PPTG_00304 | Phytophthora parasitica INRA-310 hypothetical protein (254 aa) |
| 4051 | 3772 | PPTG_03772 | 36,41 | 3 | 0 | 3 | 0 |  | 1,20E+05 |  | 1 | 1 | 0 | 1 | 0 |  | 53286 | PPTG_03772 | Phytophthora parasitica INRA-310 crinkler (CRN) family protein pseudogene (470 aa) |
| 3795 | 4342 | PPTG_18168 | 36,39 | 7 | 0 | 7 | 7 | | 8,90E+04 | 3,30E+05 | 1 | 1 | 0 | 1 | 1 | | 15357 | PPTG_18168 | Phytophthora parasitica INRA-310 hypothetical protein (137 aa) |
| 3795 | 4343 | PPTG_20245 | 36,39 | 7 | 0 | 7 | 7 | | 8,90E+04 | 3,30E+05 | 1 | 1 | 0 | 1 | 1 | | 15357 | PPTG_20245 | Phytophthora parasitica INRA-310 hypothetical protein (137 aa) |
| 4095 | 3900 | PPTG_05086 | 36,34 | 11 | 0 | 11 | 0 | | 1,75E+05 | | 1 | 1 | 0 | 1 | 0 | | 12497 | PPTG_05086 | Phytophthora parasitica INRA-310 hypothetical protein (113 aa) |
| 3228 | 4150 | PPTG_06642 | 36,3 | 2 | 2 | 2 | 2 | 7,46E+04 | 3,41E+05 | 5,78E+05 | 1 | 1 | 1 | 1 | 1 | | 51513 | PPTG_06642 | Phytophthora parasitica INRA-310 tubulin gamma chain (461 aa) |
| 3560 | 3401 | PPTG_20208 | 36,28 | 13 | 0 | 13 | 13 | | 4,14E+05 | 1,38E+05 | 1 | 1 | 0 | 1 | 1 | | 11710 | PPTG_20208 | Phytophthora parasitica INRA-310 hypothetical protein (108 aa) |
| 3793 | 4332 | PPTG_02273 | 36,25 | 5 | 0 | 5 | 5 | | 1,79E+05 | 3,01E+05 | 1 | 1 | 0 | 1 | 1 | | 32283 | PPTG_02273 | Phytophthora parasitica INRA-310 hypothetical protein (287 aa) |
| 4617 | 4333 | PPTG_06905 | 36,24 | 8 | 0 | 0 | 8 | | | 2,85E+05 | 1 | 1 | 0 | 0 | 1 | Carbamidomethylation | 20555 | PPTG_06905 | Phytophthora parasitica INRA-310 hypothetical protein (178 aa) |
| 3377 | 3613 | PPTG_16086 | 36,22 | 1 | 0 | 0 | 1 | | | 2,71E+05 | 2 | 2 | 0 | 0 | 2 | Carbamidomethylation | 223326 | PPTG_16086 | Phytophthora parasitica INRA-310 hypothetical protein (2042 aa) |
| 3720 | 4340 | PPTG_15958 | 36,17 | 4 | 0 | 4 | 4 | | 8,26E+05 | 1,22E+06 | 1 | 1 | 0 | 1 | 1 | | 30792 | PPTG_15958 | Phytophthora parasitica INRA-310 hypothetical protein (282 aa) |
| 3281 | 4336 | PPTG_12560 | 36,15 | 3 | 0 | 3 | 3 | | 8,08E+05 | 1,10E+06 | 1 | 1 | 0 | 1 | 2 | Carbamidomethylation | 44129 | PPTG_12560 | Phytophthora parasitica INRA-310 hypothetical protein (421 aa) |
| 2758 | 3579 | PPTG_10946 | 36,14 | 3 | 0 | 3 | 3 | | 3,58E+05 | 6,54E+05 | 2 | 2 | 0 | 2 | 2 | | 76568 | PPTG_10946 | Phytophthora parasitica INRA-310 DEAD/DEAH box RNA helicase (687 aa) |
| 3548 | 3109 | PPTG_19170 | 36,1 | 2 | 0 | 0 | 2 | | | 8,50E+05 | 2 | 2 | 0 | 0 | 2 | | 96246 | PPTG_19170 | Phytophthora parasitica INRA-310 hypothetical protein (846 aa) |
| 3724 | 4367 | PPTG_00927 | 36,06 | 3 | 0 | 3 | 3 | | 6,57E+05 | 9,92E+05 | 1 | 1 | 0 | 1 | 1 | | 40521 | PPTG_00927 | Phytophthora parasitica INRA-310 3-deoxy-7-phosphoheptulonate synthase (375 aa) |
| 3598 | 4067 | PPTG_10141 | 36,05 | 2 | 0 | 2 | 2 | | 1,88E+05 | 0 | 1 | 1 | 0 | 1 | 1 | | 71065 | PPTG_10141 | Phytophthora parasitica INRA-310 hypothetical protein (642 aa) |
| 3693 | 4163 | PPTG_12060 | 36,05 | 4 | 0 | 4 | 4 | | 9,53E+04 | 1,39E+05 | 1 | 1 | 0 | 1 | 1 | | 36445 | PPTG_12060 | Phytophthora parasitica INRA-310 hypothetical protein (322 aa) |
| 3416 | 3983 | PPTG_07152 | 36,04 | 1 | 1 | 1 | 0 | 1,34E+05 | 6,53E+05 |  | 1 | 1 | 1 | 1 | 0 | Carbamylation | 64457 | PPTG_07152 | Phytophthora parasitica INRA-310 crinkler (CRN) family protein pseudogene (584 aa) |
| 3416 | 4026 | PPTG_07159 | 36,04 | 2 | 2 | 2 | 0 | 1,34E+05 | 6,53E+05 |  | 1 | 1 | 1 | 1 | 0 | Carbamylation | 35707 | PPTG_07159 | Phytophthora parasitica INRA-310 crinkler (CRN) family protein pseudogene (322 aa) |
| 3491 | 3502 | PPTG_10264 | 36,02 | 1 | 0 | 0 | 1 | | | 4,48E+06 | 2 | 2 | 0 | 0 | 2 | Carbamidomethylation | 323744 | PPTG_10264 | Phytophthora parasitica INRA-310 hypothetical protein (2870 aa) |
| 4424 | 4337 | PPTG_10397 | 36 | 7 | 0 | 0 | 7 | | | 5,12E+05 | 1 | 1 | 0 | 0 | 1 | | 22637 | PPTG_10397 | Phytophthora parasitica INRA-310 YIPF6-like protein (209 aa) |
| 4618 | 4338 | PPTG_11042 | 35,97 | 3 | 0 | 0 | 3 | | | 1,86E+05 | 1 | 1 | 0 | 0 | 1 | | 52668 | PPTG_11042 | Phytophthora parasitica INRA-310 hypothetical protein (503 aa) |
| 4047 | 3714 | PPTG_02903 | 35,96 | 7 | 0 | 0 | 7 | | | 1,55E+05 | 1 | 1 | 0 | 0 | 1 | | 39550 | PPTG_02903 | Phytophthora parasitica INRA-310 hypothetical protein (363 aa) |
| 3522 | 3955 | PPTG_04954 | 35,95 | 1 | 1 | 0 | 1 | 1,69E+05 | | 1,30E+06 | 1 | 1 | 1 | 0 | 1 | | 111206 | PPTG_04954 | Phytophthora parasitica INRA-310 calcium-activated potassium channel subunit alpha-1 (1001 aa) |
| 4080 | 3577 | PPTG_18455 | 35,94 | 6 | 0 | 0 | 6 | | | 3,08E+05 | 1 | 1 | 0 | 0 | 1 | Carbamidomethylation | 43309 | PPTG_18455 | Phytophthora parasitica INRA-310 homoserine kinase (413 aa) |
| 4623 | 4370 | PPTG_05055 | 35,93 | 7 | 0 | 7 | 0 | | 5,72E+04 | | 1 | 1 | 0 | 1 | 0 | | 42751 | PPTG_05055 | Phytophthora parasitica INRA-310 hypothetical protein (399 aa) |
| 4254 | 3992 | PPTG_20482 | 35,89 | 4 | 0 | 0 | 4 | | | 2,32E+05 | 1 | 1 | 0 | 0 | 1 | | 37519 | PPTG_20482 | Phytophthora parasitica INRA-310 hypothetical protein (427 aa) |
| 4624 | 4372 | PPTG_11767 | 35,87 | 4 | 0 | 4 | 0 | | 3,42E+05 | | 1 | 1 | 0 | 1 | 0 | | 31583 | PPTG_11767 | Phytophthora parasitica INRA-310 SCP-like extracellular protein (297 aa) |
| 3143 | 3121 | PPTG_04234 | 35,87 | 6 | 0 | 6 | 3 | | 2,85E+05 | 1,88E+05 | 2 | 2 | 0 | 2 | 1 | | 37417 | PPTG_04234 | Phytophthora parasitica INRA-310 hypothetical protein (323 aa) |
| 4406 | 4185 | PPTG_00761 | 35,86 | 4 | 0 | 4 | 0 | | 7,91E+05 | | 1 | 1 | 0 | 1 | 0 | Carbamylation | 39719 | PPTG_00761 | Phytophthora parasitica INRA-310 hypothetical protein (348 aa) |
| 3728 | 4377 | PPTG_13458 | 35,8 | 7 | 0 | 7 | 7 | | 5,54E+05 | 0 | 1 | 1 | 0 | 1 | 1 | Carbamidomethylation | 20869 | PPTG_13458 | Phytophthora parasitica INRA-310 hypothetical protein (210 aa) |
| 3219 | 3731 | PPTG_17899 | 35,74 | 5 | 0 | 3 | 5 | | 9,07E+04 | 2,41E+05 | 2 | 2 | 0 | 1 | 2 | | 40801 | PPTG_17899 | Phytophthora parasitica INRA-310 choline/ethanolamine kinase (361 aa) |
| 4431 | 4384 | PPTG_18564 | 35,73 | 8 | 0 | 8 | 0 | | 1,04E+05 | | 1 | 1 | 0 | 1 | 0 | Carbamidomethylation | 18748 | PPTG_18564 | Phytophthora parasitica INRA-310 hypothetical protein (173 aa) |
| 3794 | 4341 | PPTG_09816 | 35,73 | 5 | 0 | 0 | 5 | | | 1,89E+05 | 1 | 1 | 0 | 0 | 2 | | 60800 | PPTG_09816 | Phytophthora parasitica INRA-310 hypothetical protein (564 aa) |
| 2865 | 3080 | PPTG_10927 | 35,71 | 7 | 0 | 7 | 7 | | 5,04E+05 | 2,59E+05 | 2 | 2 | 0 | 2 | 2 | | 33831 | PPTG_10927 | Phytophthora parasitica INRA-310 O-methyltransferase (303 aa) |
| 3176 | 4168 | PPTG_02495 | 35,71 | 2 | 2 | 2 | 2 | 3,53E+04 | 2,60E+05 | 2,86E+05 | 1 | 1 | 1 | 1 | 1 | | 66756 | PPTG_02495 | Phytophthora parasitica INRA-310 3-hydroxyisobutyrate dehydrogenase (617 aa) |
| 4024 | 3880 | PPTG_15213 | 35,66 | 1 | 0 | 0 | 1 | | | 0 | 1 | 1 | 0 | 0 | 1 | | 171016 | PPTG_15213 | Phytophthora parasitica INRA-310 hypothetical protein (1536 aa) |
| 3994 | 3112 | PPTG_11310 | 35,65 | 1 | 0 | 0 | 1 | | | 5,81E+05 | 1 | 1 | 0 | 0 | 1 | | 90133 | PPTG_11310 | Phytophthora parasitica INRA-310 hypothetical protein (799 aa) |
| 3220 | 3744 | PPTG_17564 | 35,64 | 14 | 0 | 7 | 14 | | 2,76E+06 | 6,13E+06 | 2 | 2 | 0 | 1 | 2 | | 13945 | PPTG_17564 | Phytophthora parasitica INRA-310 nuclear transcription factor Y subunit B-3 (124 aa) |
| 2732 | 3688 | PPTG_11966 | 35,64 | 5 | 0 | 5 | 5 | | 1,17E+05 | 2,05E+05 | 2 | 2 | 0 | 2 | 2 | | 70029 | PPTG_11966 | Phytophthora parasitica INRA-310 hypothetical protein (666 aa) |
| 4626 | 4388 | PPTG_18857 | 35,62 | 4 | 0 | 4 | 0 | | 2,08E+05 | | 1 | 1 | 0 | 1 | 0 | | 40145 | PPTG_18857 | Phytophthora parasitica INRA-310 hypothetical protein (368 aa) |
| 3525 | 3993 | PPTG_18621 | 35,62 | 1 | 0 | 1 | 1 | | 1,16E+05 | 3,79E+05 | 1 | 1 | 0 | 1 | 1 | | 87986 | PPTG_18621 | Phytophthora parasitica INRA-310 inactive ubiquitin carboxyl-terminal hydrolase (786 aa) |
| 3441 | 3684 | PPTG_00334 | 35,61 | 2 | 0 | 0 | 2 | | | 5,04E+05 | 2 | 2 | 0 | 0 | 2 | | 126400 | PPTG_00334 | Phytophthora parasitica INRA-310 hypothetical protein (1138 aa) |
| 3572 | 3727 | PPTG_05484 | 35,58 | 1 | 0 | 0 | 1 | | | 2,46E+05 | 2 | 2 | 0 | 0 | 2 | | 236288 | PPTG_05484 | Phytophthora parasitica INRA-310 inositol-3 4-bisphosphate 4-phosphatase (2202 aa) |
| 3611 | 4344 | PPTG_15039 | 35,53 | 5 | 0 | 0 | 5 | | | 3,48E+05 | 1 | 1 | 0 | 0 | 2 | Oxidation (M); Deamidation (NQ) | 56052 | PPTG_15039 | Phytophthora parasitica INRA-310 major facilitator superfamily transporter (MFS) (512 aa) |
| 3611 | 4345 | PPTG_15060 | 35,53 | 5 | 0 | 0 | 5 | | | 3,48E+05 | 1 | 1 | 0 | 0 | 2 | Oxidation (M); Deamidation (NQ) | 56616 | PPTG_15060 | Phytophthora parasitica INRA-310 major facilitator superfamily transporter (MFS) (520 aa) |
| 3575 | 3738 | PPTG_14990 | 35,42 | 3 | 0 | 0 | 3 | | | 5,21E+05 | 2 | 2 | 0 | 0 | 2 | | 101269 | PPTG_14990 | Phytophthora parasitica INRA-310 calcineurin-like phosphoesterase (907 aa) |
| 3593 | 4012 | PPTG_09016 | 35,41 | 2 | 0 | 2 | 2 | | 4,98E+04 | 8,35E+04 | 1 | 1 | 0 | 1 | 1 | | 103005 | PPTG_09016 | Phytophthora parasitica INRA-310 hydroxymethylglutaryl-CoA synthase (955 aa) |
| 3493 | 3547 | PPTG_01924 | 35,4 | 22 | 0 | 0 | 22 | | | 1,17E+06 | 2 | 2 | 0 | 0 | 2 | Carbamidomethylation | 16385 | PPTG_01924 | Phytophthora parasitica INRA-310 hypothetical protein (153 aa) |
| 3563 | 3503 | PPTG_11284 | 35,35 | 7 | 0 | 7 | 0 | | 6,02E+04 | | 2 | 2 | 0 | 2 | 0 | | 48450 | PPTG_11284 | Phytophthora parasitica INRA-310 hypothetical protein (423 aa) |
| 4619 | 4351 | PPTG_12904 | 35,35 | 4 | 0 | 0 | 4 | | | 3,07E+05 | 1 | 1 | 0 | 0 | 1 | | 35319 | PPTG_12904 | Phytophthora parasitica INRA-310 hypothetical protein (326 aa) |
| 4253 | 3981 | PPTG_01468 | 35,35 | 3 | 0 | 3 | 0 | | 2,56E+05 | | 1 | 1 | 0 | 1 | 0 | Carbamidomethylation | 53011 | PPTG_01468 | Phytophthora parasitica INRA-310 hypothetical protein (462 aa) |
| 3081 | 3743 | PPTG_06010 | 35,31 | 1 | 0 | 1 | 0 | | 1,73E+05 | 2,76E+05 | 2 | 2 | 0 | 2 | 1 | | 233383 | PPTG_06010 | Phytophthora parasitica INRA-310 cytoskeleton-associated protein (2167 aa) |
| 3051 | 3704 | PPTG_05508 | 35,26 | 3 | 0 | 2 | 3 | | 1,11E+05 | 8,15E+05 | 2 | 2 | 0 | 1 | 2 | Carbamidomethylation | 64600 | PPTG_05508 | Phytophthora parasitica INRA-310 NAD kinase (584 aa) |
| 3430 | 3089 | PPTG_09512 | 35,25 | 4 | 0 | 2 | 1 | | 8,00E+04 | 2,53E+05 | 2 | 2 | 0 | 1 | 1 | Carbamidomethylation | 85839 | PPTG_09512 | Phytophthora parasitica INRA-310 intraflagellar transporter 80 (768 aa) |
| 3367 | 3001 | PPTG_18256 | 35,24 | 3 | 0 | 1 | 2 | | 0 | 0 | 2 | 2 | 0 | 1 | 1 | Carbamidomethylation | 87016 | PPTG_18256 | Phytophthora parasitica INRA-310 hypothetical protein (797 aa) |
| 4621 | 4356 | PPTG_19603 | 35,18 | 9 | 0 | 0 | 9 | | | 2,49E+05 | 1 | 1 | 0 | 0 | 1 | | 13944 | PPTG_19603 | Phytophthora parasitica INRA-310 hypothetical protein (122 aa) |
| 3697 | 4180 | PPTG_11605 | 35,16 | 3 | 0 | 3 | 3 | | 1,35E+05 | 1,84E+05 | 1 | 1 | 0 | 1 | 1 | | 37031 | PPTG_11605 | Phytophthora parasitica INRA-310 hypothetical protein (336 aa) |
| 3697 | 4358 | PPTG_11603 | 35,16 | 2 | 0 | 2 | 2 | | 1,35E+05 | 1,84E+05 | 1 | 1 | 0 | 1 | 1 | | 57165 | PPTG_11603 | Phytophthora parasitica INRA-310 hypothetical protein (518 aa) |
| 3799 | 4393 | PPTG_11085 | 35,15 | 10 | 0 | 10 | 0 | | 2,63E+05 | | 1 | 1 | 0 | 1 | 0 | | 13861 | PPTG_11085 | Phytophthora parasitica INRA-310 hypothetical protein (126 aa) |
| 3779 | 3752 | PPTG_02468 | 35,15 | 14 | 0 | 9 | 4 | | 1,08E+05 | 0 | 2 | 2 | 0 | 1 | 1 | | 28073 | PPTG_02468 | Phytophthora parasitica INRA-310 ribosome-recycling factor (260 aa) |
| 4259 | 4025 | PPTG_05075 | 35,13 | 1 | 0 | 0 | 1 | | | 3,08E+05 | 1 | 1 | 0 | 0 | 1 | | 156667 | PPTG_05075 | Phytophthora parasitica INRA-310 hypothetical protein (1399 aa) |
| 3673 | 3683 | PPTG_04027 | 35,1 | 10 | 0 | 0 | 10 | | | 1,05E+06 | 2 | 2 | 0 | 0 | 2 | | 29466 | PPTG_04027 | Phytophthora parasitica INRA-310 hypothetical protein (263 aa) |
| 3797 | 4375 | PPTG_05142 | 35,09 | 3 | 0 | 3 | 3 | | 5,29E+05 | 3,57E+06 | 1 | 1 | 0 | 1 | 1 | | 28792 | PPTG_05142 | Phytophthora parasitica INRA-310 hypothetical protein (253 aa) |
| 4627 | 4398 | PPTG_00070 | 35,02 | 9 | 0 | 9 | 0 | | 2,07E+05 | | 1 | 1 | 0 | 1 | 0 | | 18711 | PPTG_00070 | Phytophthora parasitica INRA-310 carbonic anhydrase (184 aa) |
| 3698 | 4182 | PPTG_08105 | 35,02 | 1 | 0 | 1 | 1 | | 6,81E+04 | 1,65E+05 | 1 | 1 | 0 | 1 | 1 | Carbamidomethylation | 100641 | PPTG_08105 | Phytophthora parasitica INRA-310 rRNA methyltransferase (883 aa) |
| 4151 | 3261 | PPTG_10202 | 34,99 | 5 | 0 | 5 | 0 | | 1,77E+05 | | 1 | 1 | 0 | 1 | 0 | | 21283 | PPTG_10202 | Phytophthora parasitica INRA-310 anthranilate synthase component 2 (192 aa) |
| 3589 | 3999 | PPTG_11423 | 34,96 | 2 | 0 | 2 | 2 | | 9,23E+04 | 1,96E+05 | 1 | 1 | 0 | 1 | 1 | | 54964 | PPTG_11423 | Phytophthora parasitica INRA-310 general transcription factor IIH subunit (484 aa) |
| 4622 | 4361 | PPTG_15778 | 34,92 | 9 | 0 | 0 | 9 | | | 7,28E+04 | 1 | 1 | 0 | 0 | 1 | | 16362 | PPTG_15778 | Phytophthora parasitica INRA-310 hypothetical protein (140 aa) |
| 4278 | 4176 | PPTG_19565 | 34,9 | 4 | 0 | 4 | 0 | | 1,58E+05 | | 1 | 1 | 0 | 1 | 0 | Carbamidomethylation | 43982 | PPTG_19565 | Phytophthora parasitica INRA-310 hypothetical protein (408 aa) |
| 2789 | 3798 | PPTG_03019 | 34,9 | 3 | 1 | 1 | 3 | | | 9,43E+05 | 2 | 1 | 1 | 1 | 2 | | 53731 | PPTG_03019 | Phytophthora parasitica INRA-310 hypothetical protein (489 aa) |
| 4258 | 4020 | PPTG_13195 | 34,86 | 1 | 0 | 0 | 1 | | | 2,71E+05 | 1 | 1 | 0 | 0 | 1 | | 80835 | PPTG_13195 | Phytophthora parasitica INRA-310 hypothetical protein (712 aa) |
| 3796 | 4362 | PPTG_09952 | 34,82 | 4 | 0 | 0 | 4 | | | 1,39E+05 | 1 | 1 | 0 | 0 | 2 | | 38993 | PPTG_09952 | Phytophthora parasitica INRA-310 protein phosphatase methylesterase (357 aa) |
| 2727 | 3002 | PPTG_10615 | 34,81 | 4 | 2 | 2 | 4 | 2,77E+05 | 8,11E+05 | 2,97E+06 | 2 | 2 | 1 | 1 | 2 | | 61660 | PPTG_10615 | Phytophthora parasitica INRA-310 sphingosine kinase (556 aa) |
| 3398 | 3067 | PPTG_14509 | 34,8 | 7 | 4 | 0 | 4 | 5,42E+04 | | 1,27E+05 | 2 | 2 | 1 | 0 | 1 | | 29931 | PPTG_14509 | Phytophthora parasitica INRA-310 hypothetical protein (273 aa) |
| 3574 | 3735 | PPTG_16190 | 34,8 | 3 | 0 | 0 | 3 | | | 2,92E+05 | 2 | 1 | 0 | 0 | 2 | | 70417 | PPTG_16190 | Phytophthora parasitica INRA-310 protein kinase (642 aa) |

| | | | | | | | | | | | | | | | | | |
|---|---|---|---|---|---|---|---|---|---|---|---|---|---|---|---|---|---|
| 3729 | 4394 | PPTG_03405 | 34,79 | 3 | 0 | 3 | 3 | | 2,23E+06 | 2,15E+06 | 1 | 1 | 0 | 1 | 1 | | 64101 \| PPTG_03405 \| Phytophthora parasitica INRA-310 leishmanolysin-like peptidase (591 aa) |
| 2664 | 3846 | PPTG_19528 | 34,77 | 4 | 0 | 4 | 4 | | 2,61E+05 | 4,30E+06 | 1 | 1 | 0 | 1 | 3 | Oxidation (M) | 29715 \| PPTG_19528 \| Phytophthora parasitica INRA-310 hypothetical protein (261 aa) |
| 2614 | 4460 | PPTG_11487 | 34,77 | 7 | 7 | 7 | 7 | 3,55E+07 | 1,05E+08 | 2,72E+08 | 1 | 1 | 1 | 1 | 3 | Carbamylation | 13309 \| PPTG_11487 \| Phytophthora parasitica INRA-310 50S ribosomal protein L34e (118 aa) |
| 3722 | 4364 | PPTG_17521 | 34,76 | 16 | 0 | 0 | 16 | | | 9,65E+06 | 1 | 1 | 0 | 0 | 2 | | 8837 \| PPTG_17521 \| Phytophthora parasitica INRA-310 hypothetical protein (82 aa) |
| 2837 | 3764 | PPTG_10917 | 34,76 | 4 | 2 | 4 | 2 | 8,24E+04 | 7,41E+05 | 0 | 2 | 2 | 1 | 2 | 1 | | 80921 \| PPTG_10917 \| Phytophthora parasitica INRA-310 hypothetical protein (714 aa) |
| 3343 | 3709 | PPTG_14846 | 34,75 | 3 | 0 | 3 | 3 | | 3,57E+05 | 3,28E+05 | 1 | 1 | 0 | 1 | 1 | | 43872 \| PPTG_14846 \| Phytophthora parasitica INRA-310 cell division protein FtsZ (421 aa) |
| 3507 | 3726 | PPTG_01666 | 34,73 | 1 | 0 | 0 | 1 | | | 2,18E+05 | 2 | 2 | 0 | 0 | 2 | | 157490 \| PPTG_01666 \| Phytophthora parasitica INRA-310 hypothetical protein (1407 aa) |
| 4390 | 3912 | PPTG_13363 | 34,72 | 5 | 0 | 0 | 5 | | | 3,24E+05 | 1 | 1 | 0 | 0 | 1 | | 27570 \| PPTG_13363 \| Phytophthora parasitica INRA-310 Der1-like family protein (251 aa) |
| 3417 | 3988 | PPTG_16704 | 34,71 | 1 | 0 | 1 | 1 | | 1,86E+05 | 1,59E+05 | 1 | 1 | 0 | 1 | 1 | | 159911 \| PPTG_16704 \| Phytophthora parasitica INRA-310 ubiquitin-specific protease (1414 aa) |
| 4433 | 4391 | PPTG_08028 | 34,7 | 2 | 0 | 0 | 2 | | | 2,28E+06 | 1 | 1 | 0 | 0 | 1 | | 41005 \| PPTG_08028 \| Phytophthora parasitica INRA-310 aspartyl protease family A22B (371 aa) |
| 4247 | 3920 | PPTG_15543 | 34,66 | 6 | 0 | 0 | 6 | | | 4,98E+05 | 1 | 1 | 0 | 0 | 1 | | 27330 \| PPTG_15543 \| Phytophthora parasitica INRA-310 hypothetical protein (254 aa) |
| 3780 | 3836 | PPTG_12896 | 34,62 | 3 | 0 | 3 | 0 | | 3,89E+05 | | 2 | 2 | 0 | 2 | 0 | | 80769 \| PPTG_12896 \| Phytophthora parasitica INRA-310 pectin lyase (797 aa) |
| 3725 | 4368 | PPTG_03988 | 34,62 | 7 | 0 | 7 | 7 | | 1,22E+05 | 5,97E+05 | 1 | 1 | 0 | 1 | 1 | | 38670 \| PPTG_03988 \| Phytophthora parasitica INRA-310 hypothetical protein (357 aa) |
| 3453 | 3946 | PPTG_17397 | 34,61 | 4 | 0 | 0 | 4 | | | 3,50E+05 | 1 | 1 | 0 | 0 | 2 | | 48586 \| PPTG_17397 \| Phytophthora parasitica INRA-310 cation Diffusion Facilitator (CDF) family (453 aa) |
| 3167 | 3740 | PPTG_11062 | 34,59 | 5 | 0 | 2 | 5 | | 9,66E+04 | 1,10E+06 | 2 | 2 | 0 | 1 | 2 | | 36595 \| PPTG_11062 \| Phytophthora parasitica INRA-310 tafazzin-like protein (330 aa) |
| 4099 | 3932 | PPTG_19692 | 34,59 | 4 | 0 | 0 | 4 | | | 3,51E+05 | 1 | 1 | 0 | 0 | 1 | | 29236 \| PPTG_19692 \| Phytophthora parasitica INRA-310 hypothetical protein (266 aa) |
| 3374 | 3166 | PPTG_10953 | 34,58 | 3 | 0 | 3 | 3 | | 2,70E+05 | 4,84E+05 | 1 | 1 | 0 | 1 | 1 | | 32196 \| PPTG_10953 \| Phytophthora parasitica INRA-310 general transcription factor IIH subunit (294 aa) |
| 3520 | 3939 | PPTG_06313 | 34,57 | 10 | 0 | 10 | 10 | | 2,63E+05 | 2,14E+05 | 1 | 1 | 0 | 1 | 1 | | 18809 \| PPTG_06313 \| Phytophthora parasitica INRA-310 hypothetical protein (174 aa) |
| 3021 | 3464 | PPTG_15957 | 34,56 | 10 | 2 | 0 | 8 | 1,13E+05 | | 8,01E+05 | 3 | 3 | 1 | 0 | 2 | | 43736 \| PPTG_15957 \| Phytophthora parasitica INRA-310 hypothetical protein (390 aa) |
| 4404 | 4154 | PPTG_10531 | 34,53 | 5 | 0 | 0 | 5 | | | 2,29E+05 | 1 | 1 | 0 | 0 | 1 | | 27835 \| PPTG_10531 \| Phytophthora parasitica INRA-310 hypothetical protein (255 aa) |
| 3800 | 4410 | PPTG_00700 | 34,52 | 6 | 0 | 6 | 6 | | 5,16E+05 | 5,39E+05 | 1 | 1 | 0 | 1 | 1 | | 23680 \| PPTG_00700 \| Phytophthora parasitica INRA-310 fumarylacetoacetate (FAA) hydrolase (222 aa) |
| 3238 | 4412 | PPTG_13483 | 34,48 | 8 | 0 | 8 | 8 | | 2,56E+06 | 1,32E+06 | 1 | 1 | 0 | 2 | 1 | Dehydration; Deamidation (NQ) | 16633 \| PPTG_13483 \| Phytophthora parasitica INRA-310 cellulose binding elicitor lectin (CBEL) (156 aa) |
| 3473 | 3019 | PPTG_05043 | 34,47 | 6 | 0 | 0 | 6 | | | 6,58E+05 | 2 | 2 | 0 | 0 | 2 | | 50895 \| PPTG_05043 \| Phytophthora parasitica INRA-310 protein phosphatase (461 aa) |
| 3726 | 4369 | PPTG_00212 | 34,46 | 8 | 0 | 8 | 8 | | 2,11E+05 | 3,97E+05 | 1 | 1 | 0 | 1 | 1 | | 15166 \| PPTG_00212 \| Phytophthora parasitica INRA-310 hypothetical protein (136 aa) |
| 3594 | 4013 | PPTG_15094 | 34,45 | 1 | 0 | 1 | 1 | | 1,61E+05 | 2,84E+05 | 1 | 1 | 0 | 1 | 1 | | 212281 \| PPTG_15094 \| Phytophthora parasitica INRA-310 chromodomain-helicase-DNA-binding protein (1852 aa) |
| 3670 | 3672 | PPTG_05527 | 34,45 | 5 | 0 | 0 | 5 | | | 5,14E+05 | 2 | 2 | 0 | 0 | 2 | | 93353 \| PPTG_05527 \| Phytophthora parasitica INRA-310 hypothetical protein (824 aa) |
| 4078 | 2989 | PPTG_10967 | 34,44 | 2 | 0 | 0 | 2 | | | 9,07E+05 | 1 | 1 | 0 | 0 | 1 | | 52337 \| PPTG_10967 \| Phytophthora parasitica INRA-310 protein kinase (458 aa) |
| 3166 | 3700 | PPTG_06641 | 34,38 | 11 | 0 | 5 | 11 | | 2,13E+05 | 4,00E+05 | 2 | 2 | 0 | 1 | 2 | | 22143 \| PPTG_06641 \| Phytophthora parasitica INRA-310 trafficking protein particle complex subunit (200 aa) |
| 3052 | 3745 | PPTG_19122 | 34,36 | 4 | 0 | 4 | 2 | | 2,36E+05 | 4,46E+05 | 2 | 2 | 0 | 2 | 1 | | 79256 \| PPTG_19122 \| Phytophthora parasitica INRA-310 protein kinase (721 aa) |
| 3680 | 3927 | PPTG_10818 | 34,32 | 3 | 3 | 0 | 3 | 5,26E+04 | | 4,91E+05 | 1 | 1 | 1 | 0 | 1 | | 40764 \| PPTG_10818 \| Phytophthora parasitica INRA-310 metallophosphoesterase 1 (360 aa) |
| 4401 | 4143 | PPTG_00320 | 34,32 | 2 | 0 | 0 | 2 | | | 6,56E+05 | 1 | 1 | 0 | 0 | 1 | | 67897 \| PPTG_00320 \| Phytophthora parasitica INRA-310 hypothetical protein (602 aa) |
| 4401 | 4162 | PPTG_00294 | 34,32 | 2 | 0 | 0 | 2 | | | 6,56E+05 | 1 | 1 | 0 | 0 | 1 | | 83174 \| PPTG_00294 \| Phytophthora parasitica INRA-310 hypothetical protein (739 aa) |
| 3413 | 3929 | PPTG_13579 | 34,29 | 1 | 0 | 1 | 1 | | 2,06E+04 | 1,64E+05 | 1 | 1 | 0 | 1 | 1 | | 211639 \| PPTG_13579 \| Phytophthora parasitica INRA-310 hypothetical protein (1901 aa) |
| 3413 | 3930 | PPTG_13579 | 34,29 | 1 | 0 | 1 | 1 | | 2,06E+04 | 1,64E+05 | 1 | 1 | 0 | 1 | 1 | | 211710 \| PPTG_13579 \| Phytophthora parasitica INRA-310 hypothetical protein (1902 aa) |
| 3727 | 4373 | PPTG_02724 | 34,2 | 5 | 0 | 5 | 5 | | 3,36E+05 | 3,24E+05 | 1 | 1 | 0 | 1 | 1 | | 23110 \| PPTG_02724 \| Phytophthora parasitica INRA-310 hypothetical protein (220 aa) |
| 3007 | 3513 | PPTG_03429 | 34,18 | 1 | 0 | 1 | 1 | | 2,29E+05 | 5,94E+05 | 2 | 2 | 0 | 1 | 2 | | 139432 \| PPTG_03429 \| Phytophthora parasitica INRA-310 ATP-binding Cassette (ABC) superfamily (1257 aa) |
| 4631 | 4426 | PPTG_15710 | 34,16 | 5 | 0 | 5 | 0 | | 7,76E+04 | | 1 | 1 | 0 | 1 | 0 | Carbamidomethylation | 34995 \| PPTG_15710 \| Phytophthora parasitica INRA-310 arabinan endo-1 5-alpha-L-arabinosidase A (323 aa) |
| 3527 | 4027 | PPTG_17518 | 34,13 | 8 | 0 | 8 | 8 | | 9,32E+05 | 8,64E+05 | 1 | 1 | 0 | 1 | 1 | Acetylation (N-term) | 18542 \| PPTG_17518 \| Phytophthora parasitica INRA-310 hypothetical protein (167 aa) |
| 3325 | 3496 | PPTG_13254 | 34,07 | 1 | 0 | 1 | 1 | | 2,11E+05 | 2,15E+05 | 1 | 1 | 0 | 1 | 1 | | 87536 \| PPTG_13254 \| Phytophthora parasitica INRA-310 formin-likey 2 domain-containing protein (780 aa) |
| 4430 | 4376 | PPTG_00285 | 34,05 | 15 | 0 | 0 | 15 | | | 5,65E+05 | 1 | 1 | 0 | 0 | 1 | | 9261 \| PPTG_00285 \| Phytophthora parasitica INRA-310 hypothetical protein (82 aa) |
| 4251 | 3973 | PPTG_18893 | 34,05 | 5 | 0 | 0 | 5 | | | 1,60E+05 | 1 | 1 | 0 | 0 | 1 | | 32881 \| PPTG_18893 \| Phytophthora parasitica INRA-310 hypothetical protein (292 aa) |
| 3518 | 3894 | PPTG_05170 | 34,01 | 5 | 0 | 5 | 0 | | 4,73E+05 | | 2 | 2 | 0 | 2 | 0 | | 48118 \| PPTG_05170 \| Phytophthora parasitica INRA-310 cyclin-dependent kinase (426 aa) |
| 3127 | 3713 | PPTG_12850 | 33,97 | 1 | 1 | 1 | 1 | 9,68E+04 | 2,17E+05 | 6,56E+05 | 1 | 1 | 1 | 1 | 1 | | 101105 \| PPTG_12850 \| Phytophthora parasitica INRA-310 hypothetical protein (922 aa) |
| 3085 | 3826 | PPTG_13225 | 33,97 | 1 | 0 | 0 | 1 | | 0 | | 2 | 1 | 0 | 1 | 2 | | 318585 \| PPTG_13225 \| Phytophthora parasitica INRA-310 hypothetical protein (2967 aa) |
| 3694 | 4165 | PPTG_08379 | 33,97 | 7 | 0 | 7 | 7 | | 4,88E+04 | 0 | 1 | 1 | 0 | 1 | 1 | | 22571 \| PPTG_08379 \| Phytophthora parasitica INRA-310 hypothetical protein (203 aa) |
| 3079 | 3694 | PPTG_18493 | 33,97 | 9 | 0 | 0 | 9 | | | 1,49E+06 | 2 | 2 | 0 | 0 | 3 | Deamidation (NQ) | 42564 \| PPTG_18493 \| Phytophthora parasitica INRA-310 mitochondrial import inner membrane translocase subunit TIM50 (387 aa) |
| 3348 | 3031 | PPTG_10237 | 33,96 | 2 | 0 | 0 | 2 | | | 1,35E+06 | 2 | 2 | 0 | 0 | 2 | | 84612 \| PPTG_10237 \| Phytophthora parasitica INRA-310 hypothetical protein (788 aa) |
| 4169 | 3947 | PPTG_07049 | 33,93 | 2 | 0 | 0 | 2 | | | 2,62E+05 | 1 | 1 | 0 | 0 | 1 | | 63683 \| PPTG_07049 \| Phytophthora parasitica INRA-310 hypothetical protein (586 aa) |
| 3608 | 4169 | PPTG_14397 | 33,77 | 4 | 0 | 4 | 4 | | 2,14E+05 | 4,85E+05 | 1 | 1 | 0 | 1 | 1 | | 32928 \| PPTG_14397 \| Phytophthora parasitica INRA-310 4-nitrophenylphosphatase (304 aa) |
| 4249 | 3964 | PPTG_04434 | 33,75 | 3 | 0 | 0 | 3 | | | 4,48E+05 | 1 | 1 | 0 | 0 | 1 | | 37184 \| PPTG_04434 \| Phytophthora parasitica INRA-310 hypothetical protein (325 aa) |
| 4625 | 4387 | PPTG_14797 | 33,74 | 3 | 0 | 0 | 3 | | | 8,15E+05 | 1 | 1 | 0 | 0 | 1 | Carbamidomethylation | 51977 \| PPTG_14797 \| Phytophthora parasitica INRA-310 cysteine protease family C01A (483 aa) |
| 4104 | 4021 | PPTG_13623 | 33,74 | 1 | 0 | 0 | 1 | | | 2,95E+05 | 1 | 1 | 0 | 0 | 1 | | 76358 \| PPTG_13623 \| Phytophthora parasitica INRA-310 Poly(A) polymerase (692 aa) |
| 3677 | 3747 | PPTG_05485 | 33,73 | 4 | 0 | 2 | 2 | | 2,27E+05 | 0 | 2 | 2 | 0 | 1 | 1 | | 48378 \| PPTG_05485 \| Phytophthora parasitica INRA-310 chromodomain protein (432 aa) |
| 3338 | 3058 | PPTG_05310 | 33,72 | 5 | 0 | 0 | 5 | | | 1,75E+06 | 2 | 2 | 0 | 0 | 2 | Carbamidomethylation | 46459 \| PPTG_05310 \| Phytophthora parasitica INRA-310 Rab11 family GTPase (418 aa) |
| 3222 | 3823 | PPTG_07263 | 33,68 | 16 | 0 | 16 | 9 | | 4,71E+05 | 4,71E+05 | 2 | 2 | 0 | 2 | 1 | | 19387 \| PPTG_07263 \| Phytophthora parasitica INRA-310 hypothetical protein (174 aa) |
| 2930 | 4959 | PPTG_19126 | 33,67 | 12 | 9 | 9 | 12 | 2,20E+05 | 5,08E+05 | 3,02E+06 | 2 | 2 | 1 | 1 | 2 | | 8511 \| PPTG_19126 \| Phytophthora parasitica INRA-310 hypothetical protein (78 aa) |
| 3586 | 3963 | PPTG_01386 | 33,65 | 3 | 0 | 3 | 3 | | 1,53E+05 | 2,15E+05 | 1 | 1 | 0 | 1 | 1 | | 47913 \| PPTG_01386 \| Phytophthora parasitica INRA-310 aminotransferase (443 aa) |
| 4632 | 4442 | PPTG_02582 | 33,63 | 3 | 0 | 3 | 0 | 0 | | | 1 | 1 | 0 | 1 | 0 | | 47715 \| PPTG_02582 \| Phytophthora parasitica INRA-310 transmembrane protein (435 aa) |
| 3035 | 3021 | PPTG_18249 | 33,57 | 5 | 0 | 3 | 5 | | 5,01E+05 | 5,25E+05 | 2 | 2 | 0 | 1 | 2 | | 30224 \| PPTG_18249 \| Phytophthora parasitica INRA-310 hypothetical protein (270 aa) |
| 3395 | 2972 | PPTG_08525 | 33,57 | 7 | 0 | 0 | 7 | | | 1,29E+06 | 2 | 2 | 0 | 0 | 2 | | 25719 \| PPTG_08525 \| Phytophthora parasitica INRA-310 vesicle-trafficking protein SEC22b (224 aa) |
| 3086 | 3950 | PPTG_16286 | 33,56 | 1 | 1 | 1 | 1 | 1,11E+05 | 4,00E+05 | 7,99E+05 | 1 | 1 | 1 | 1 | 1 | | 128658 \| PPTG_16286 \| Phytophthora parasitica INRA-310 DNA repair protein (1145 aa) |
| 4405 | 4184 | PPTG_03092 | 33,52 | 1 | 0 | 0 | 1 | | | 2,19E+05 | 1 | 1 | 0 | 0 | 1 | | 74912 \| PPTG_03092 \| Phytophthora parasitica INRA-310 hypothetical protein (710 aa) |
| 4082 | 3601 | PPTG_16639 | 33,5 | 2 | 0 | 0 | 2 | | | 2,56E+05 | 1 | 1 | 0 | 0 | 1 | | 59529 \| PPTG_16639 \| Phytophthora parasitica INRA-310 60S ribosomal export protein NMD3 (523 aa) |
| 3227 | 4119 | PPTG_16919 | 33,48 | 4 | 4 | 4 | 4 | 1,85E+05 | 6,63E+05 | 1,51E+06 | 1 | 1 | 1 | 1 | 1 | | 29231 \| PPTG_16919 \| Phytophthora parasitica INRA-310 hypothetical protein (272 aa) |
| 3730 | 4395 | PPTG_16037 | 33,44 | 5 | 0 | 5 | 5 | | 8,71E+05 | 2,04E+06 | 1 | 1 | 0 | 1 | 1 | | 26065 \| PPTG_16037 \| Phytophthora parasitica INRA-310 hypothetical protein (240 aa) |
| 2560 | 4397 | PPTG_13076 | 33,43 | 20 | 20 | 20 | 20 | 1,90E+07 | 6,37E+07 | 2,06E+08 | 1 | 1 | 1 | 2 | 2 | | 6311 \| PPTG_13076 \| Phytophthora parasitica INRA-310 60S ribosomal protein L39 (52 aa) |
| 4058 | 3979 | PPTG_16007 | 33,4 | 3 | 0 | 0 | 3 | | | 2,23E+05 | 1 | 1 | 0 | 0 | 1 | | 58370 \| PPTG_16007 \| Phytophthora parasitica INRA-310 acyltransferase (511 aa) |
| 4185 | 4194 | PPTG_14045 | 33,39 | 4 | 0 | 4 | 0 | | 1,99E+05 | | 1 | 1 | 0 | 1 | 0 | | 39234 \| PPTG_14045 \| Phytophthora parasitica INRA-310 hypothetical protein (343 aa) |
| 3128 | 3789 | PPTG_07776 | 33,35 | 13 | 0 | 13 | 2 | | 8,96E+05 | 0 | 2 | 2 | 0 | 2 | 1 | Carbamidomethylation | 32143 \| PPTG_07776 \| Phytophthora parasitica INRA-310 50S ribosomal protein (300 aa) |
| 3526 | 4005 | PPTG_19147 | 33,34 | 7 | 0 | 7 | 7 | | 3,03E+05 | 0 | 1 | 1 | 0 | 1 | 1 | | 17958 \| PPTG_19147 \| Phytophthora parasitica INRA-310 hypothetical protein (166 aa) |
| 4049 | 3769 | PPTG_14518 | 33,34 | 1 | 0 | 0 | 1 | | | 7,63E+05 | 1 | 1 | 0 | 0 | 1 | | 89558 \| PPTG_14518 \| Phytophthora parasitica INRA-310 hypothetical protein (812 aa) |
| 3513 | 3819 | PPTG_13602 | 33,33 | 2 | 0 | 2 | 2 | | 2,35E+05 | 4,90E+05 | 1 | 1 | 0 | 1 | 1 | | 71715 \| PPTG_13602 \| Phytophthora parasitica INRA-310 kinesin-like protein (643 aa) |
| 4171 | 3966 | PPTG_06556 | 33,3 | 2 | 0 | 2 | 0 | 0 | | | 1 | 1 | 0 | 1 | 0 | | 39097 \| PPTG_06556 \| Phytophthora parasitica INRA-310 hypothetical protein (356 aa) |
| 4442 | 4445 | PPTG_14679 | 33,3 | 11 | 0 | 11 | 0 | | 5,07E+05 | | 1 | 1 | 0 | 1 | 0 | | 26621 \| PPTG_14679 \| Phytophthora parasitica INRA-310 glycine-rich cell wall structural protein (278 aa) |
| 3445 | 3793 | PPTG_02927 | 33,29 | 6 | 0 | 2 | 3 | | 3,87E+05 | 3,22E+05 | 2 | 2 | 0 | 1 | 1 | | 35605 \| PPTG_02927 \| Phytophthora parasitica INRA-310 mitotic checkpoint protein (327 aa) |
| 3445 | 3794 | PPTG_02927 | 33,29 | 6 | 0 | 2 | 3 | | 3,87E+05 | 3,22E+05 | 2 | 2 | 0 | 1 | 1 | | 35605 \| PPTG_02927 \| Phytophthora parasitica INRA-310 mitotic checkpoint protein (327 aa) |
| 4407 | 4196 | PPTG_16570 | 33,22 | 2 | 0 | 2 | 0 | 0 | | | 1 | 1 | 0 | 1 | 0 | | 75041 \| PPTG_16570 \| Phytophthora parasitica INRA-310 NADPH-cytochrome P450 reductase (675 aa) |
| 4434 | 4405 | PPTG_12240 | 33,21 | 2 | 0 | 0 | 2 | | | 3,03E+05 | 1 | 1 | 0 | 0 | 1 | | 67686 \| PPTG_12240 \| Phytophthora parasitica INRA-310 hypothetical protein (614 aa) |
| 4434 | 4406 | PPTG_12223 | 33,21 | 1 | 0 | 0 | 1 | | | 3,03E+05 | 1 | 1 | 0 | 0 | 1 | | 78112 \| PPTG_12223 \| Phytophthora parasitica INRA-310 hypothetical protein (721 aa) |
| 3350 | 3125 | PPTG_13450 | 33,19 | 1 | 0 | 1 | 1 | | 6,78E+04 | 0 | 1 | 1 | 0 | 1 | 1 | | 123762 \| PPTG_13450 \| Phytophthora parasitica INRA-310 hypothetical protein (1092 aa) |
| 3369 | 3084 | PPTG_09312 | 33,18 | 2 | 0 | 1 | 1 | | 1,82E+05 | 2,00E+05 | 2 | 2 | 0 | 1 | 1 | | 135219 \| PPTG_09312 \| Phytophthora parasitica INRA-310 hect domain-containing protein (1197 aa) |
| 4164 | 3863 | PPTG_06613 | 33,17 | 10 | 0 | 10 | 0 | | 3,88E+05 | | 1 | 1 | 0 | 1 | 0 | | 12436 \| PPTG_06613 \| Phytophthora parasitica INRA-310 tubulin binding cofactor A (108 aa) |
| 3230 | 4197 | PPTG_00793 | 33,15 | 1 | 1 | 1 | 1 | 4,61E+04 | 1,99E+05 | 3,78E+05 | 1 | 1 | 1 | 1 | 1 | | 104767 \| PPTG_00793 \| Phytophthora parasitica INRA-310 oligomeric Golgi complex subunit 5 (956 aa) |
| 4399 | 4124 | PPTG_20248 | 33,15 | 3 | 0 | 0 | 3 | | 0 | | 1 | 1 | 0 | 0 | 1 | | 39088 \| PPTG_20248 \| Phytophthora parasitica INRA-310 hypothetical protein (351 aa) |
| 4399 | 4400 | PPTG_17841 | 33,15 | 4 | 0 | 0 | 4 | | 0 | | 1 | 1 | 0 | 0 | 1 | | 36756 \| PPTG_17841 \| Phytophthora parasitica INRA-310 hypothetical protein (335 aa) |
| 3237 | 4399 | PPTG_11226 | 33,15 | 6 | 6 | 6 | 6 | 8,42E+04 | 2,42E+05 | 5,60E+05 | 1 | 1 | 1 | 1 | 1 | | 31182 \| PPTG_11226 \| Phytophthora parasitica INRA-310 hypothetical protein (283 aa) |
| 3531 | 4174 | PPTG_10427 | 33,14 | 3 | 0 | 3 | 3 | | 1,93E+05 | 3,11E+05 | 1 | 1 | 0 | 1 | 1 | | 35465 \| PPTG_10427 \| Phytophthora parasitica INRA-310 hypothetical protein (320 aa) |
| 3347 | 2963 | PPTG_17593 | 33,13 | 2 | 0 | 0 | 2 | | | 5,47E+05 | 2 | 2 | 0 | 0 | 2 | Carbamidomethylation | 115846 \| PPTG_17593 \| Phytophthora parasitica INRA-310 resistance-Nodulation-Cell Division (RND) superfamily (1042 aa) |
| 4101 | 3989 | PPTG_17199 | 33,12 | 1 | 0 | 0 | 1 | | | 1,72E+05 | 1 | 1 | 0 | 0 | 1 | | 84345 \| PPTG_17199 \| Phytophthora parasitica INRA-310 hypothetical protein (759 aa) |
| 3532 | 4186 | PPTG_16620 | 33,1 | 2 | 0 | 2 | 2 | | 2,28E+05 | 1,01E+06 | 1 | 1 | 0 | 1 | 1 | | 68023 \| PPTG_16620 \| Phytophthora parasitica INRA-310 translation initiation factor IF-2 (620 aa) |
| 3524 | 3982 | PPTG_16865 | 33,06 | 5 | 0 | 5 | 5 | | 1,64E+05 | 3,91E+05 | 1 | 1 | 0 | 1 | 1 | | 24631 \| PPTG_16865 \| Phytophthora parasitica INRA-310 hypothetical protein (228 aa) |
| 4094 | 3868 | PPTG_04025 | 33,06 | 2 | 0 | 0 | 2 | | 0 | | 1 | 1 | 0 | 0 | 1 | | 64943 \| PPTG_04025 \| Phytophthora parasitica INRA-310 hypothetical protein (584 aa) |
| 4057 | 3934 | PPTG_17808 | 33,05 | 1 | 0 | 0 | 1 | | | 4,78E+04 | 1 | 1 | 0 | 0 | 1 | | 130414 \| PPTG_17808 \| Phytophthora parasitica INRA-310 pre-mRNA-splicing factor ATP-dependent RNA helicase PRP16 (1147 aa) |
| 3512 | 3803 | PPTG_08121 | 33,05 | 3 | 0 | 3 | 3 | | 2,36E+05 | 3,09E+05 | 1 | 1 | 0 | 1 | 1 | | 59145 \| PPTG_08121 \| Phytophthora parasitica INRA-310 hypothetical protein (532 aa) |
| 3578 | 3784 | PPTG_16057 | 33,04 | 11 | 0 | 4 | 7 | | 5,67E+05 | 1,65E+06 | 2 | 2 | 0 | 1 | 1 | Carbamidomethylation | 28140 \| PPTG_16057 \| Phytophthora parasitica INRA-310 hypothetical protein (242 aa) |
| 4237 | 3024 | PPTG_18926 | 33,01 | 5 | 0 | 0 | 5 | | | 3,76E+05 | 1 | 1 | 0 | 0 | 1 | | 40127 \| PPTG_18926 \| Phytophthora parasitica INRA-310 hypothetical protein (375 aa) |
| 3084 | 3825 | PPTG_10830 | 33 | 3 | 0 | 3 | 1 | | 3,09E+05 | | 2 | 1 | 0 | 2 | 1 | | 72967 \| PPTG_10830 \| Phytophthora parasitica INRA-310 hypothetical protein (671 aa) |
| 3410 | 3875 | PPTG_09670 | 32,99 | 3 | 0 | 3 | 3 | | 1,09E+05 | 2,76E+05 | 1 | 1 | 0 | 1 | 1 | | 41155 \| PPTG_09670 \| Phytophthora parasitica INRA-310 hypothetical protein (379 aa) |
| 3550 | 3151 | PPTG_12036 | 32,93 | 6 | 0 | 0 | 6 | | | 8,52E+05 | 2 | 2 | 0 | 0 | 2 | | 34524 \| PPTG_12036 \| Phytophthora parasitica INRA-310 hypothetical protein (299 aa) |
| 4154 | 3675 | PPTG_00913 | 32,93 | 7 | 0 | 7 | 0 | | 2,28E+05 | | 1 | 1 | 0 | 1 | 0 | | 20677 \| PPTG_00913 \| Phytophthora parasitica INRA-310 hypothetical protein (185 aa) |
| 3353 | 3270 | PPTG_10408 | 32,92 | 3 | 0 | 0 | 3 | | | 1,14E+06 | 2 | 2 | 0 | 0 | 2 | Carbamidomethylation | 83529 \| PPTG_10408 \| Phytophthora parasitica INRA-310 hypothetical protein (749 aa) |
| 3676 | 3742 | PPTG_18741 | 32,92 | 3 | 0 | 0 | 3 | | | 2,01E+05 | 2 | 2 | 0 | 0 | 2 | | 122217 \| PPTG_18741 \| Phytophthora parasitica INRA-310 hypothetical protein (1129 aa) |
| 3424 | 3012 | PPTG_14105 | 32,91 | 2 | 0 | 0 | 2 | | | 4,99E+05 | 2 | 2 | 0 | 0 | 2 | | 101133 \| PPTG_14105 \| Phytophthora parasitica INRA-310 calcium/calmodulin-dependent protein kinase (913 aa) |
| 4629 | 4411 | PPTG_05042 | 32,9 | 3 | 0 | 0 | 3 | | | 2,49E+05 | 1 | 1 | 0 | 0 | 1 | | 66851 \| PPTG_05042 \| Phytophthora parasitica INRA-310 hypothetical protein (579 aa) |
| 4435 | 4413 | PPTG_00244 | 32,87 | 6 | 0 | 0 | 6 | | | 3,38E+05 | 1 | 1 | 0 | 0 | 1 | | 37451 \| PPTG_00244 \| Phytophthora parasitica INRA-310 hypothetical protein (333 aa) |
| 3804 | 4449 | PPTG_05555 | 32,86 | 7 | 0 | 7 | 7 | | 1,77E+05 | 1,95E+05 | 1 | 1 | 0 | 1 | 1 | | 16716 \| PPTG_05555 \| Phytophthora parasitica INRA-310 dynein light chain (149 aa) |

| | | | | | | | | | | | | | | | | | | | |
|---|---|---|---|---|---|---|---|---|---|---|---|---|---|---|---|---|---|---|---|
| 4172 | 3967 | PPTG_01212 | 32,82 | 3 | 0 | 0 | 3 | | 0 | | 1 | 1 | 0 | 0 | 1 | | 43139 | PPTG_01212 | Phytophthora parasitica INRA-310 protein kinase (382 aa) |
| 3801 | 4414 | PPTG_09541 | 32,82 | 3 | 0 | 3 | 3 | | 1,52E+05 | 1,43E+05 | 1 | 1 | 0 | 1 | 1 | | 61547 | PPTG_09541 | Phytophthora parasitica INRA-310 GlcNac transferase (553 aa) |
| 4187 | 4448 | PPTG_09656 | 32,81 | 1 | 0 | 0 | 1 | | | 2,84E+05 | 1 | 1 | 0 | 0 | 1 | | 94634 | PPTG_09656 | Phytophthora parasitica INRA-310 hypothetical protein (872 aa) |
| 3409 | 3859 | PPTG_08434 | 32,76 | 1 | 0 | 1 | 1 | | 9,43E+04 | 2,07E+05 | 1 | 1 | 0 | 1 | 1 | | 190676 | PPTG_08434 | Phytophthora parasitica INRA-310 hypothetical protein (1751 aa) |
| 3546 | 3061 | PPTG_04391 | 32,73 | 4 | 0 | 0 | 4 | | | 2,35E+05 | 2 | 2 | 0 | 0 | 2 | | 54171 | PPTG_04391 | Phytophthora parasitica INRA-310 hypothetical protein (477 aa) |
| 3805 | 4478 | PPTG_12289 | 32,73 | 4 | 0 | 4 | 4 | 0 | 0 | | 1 | 1 | 0 | 1 | 1 | | 27673 | PPTG_12289 | Phytophthora parasitica INRA-310 elicitin-like protein (243 aa) |
| 4630 | 4416 | PPTG_03702 | 32,73 | 4 | 0 | 0 | 4 | | | 1,01E+06 | 1 | 1 | 0 | 0 | 1 | | 29648 | PPTG_03702 | Phytophthora parasitica INRA-310 aquaporin (275 aa) |
| 3339 | 3097 | PPTG_05746 | 32,71 | 2 | 0 | 0 | 2 | | | 1,34E+06 | 2 | 2 | 0 | 0 | 2 | | 102067 | PPTG_05746 | Phytophthora parasitica INRA-310 ribosomal protein S6 kinase (919 aa) |
| 3454 | 3968 | PPTG_18456 | 32,69 | 2 | 1 | 1 | 0 | 1,51E+05 | 1,70E+05 | | 2 | 2 | 1 | 1 | 0 | | 84003 | PPTG_18456 | Phytophthora parasitica INRA-310 DNA topoisomerase (744 aa) |
| 3382 | 3890 | PPTG_05901 | 32,69 | 1 | 0 | 1 | 1 | | 2,61E+05 | 4,00E+05 | 1 | 1 | 0 | 1 | 1 | | 120539 | PPTG_05901 | Phytophthora parasitica INRA-310 exosome complex exonuclease RRP44 (1079 aa) |
| 3802 | 4417 | PPTG_04452 | 32,63 | 3 | 0 | 3 | 3 | | 2,33E+05 | 4,60E+05 | 1 | 1 | 0 | 1 | 1 | | 54415 | PPTG_04452 | Phytophthora parasitica INRA-310 tRNA nucleotidyltransferase (480 aa) |
| 3702 | 4227 | PPTG_11895 | 32,62 | 2 | 0 | 2 | 2 | | 1,07E+05 | 1,09E+05 | 1 | 1 | 0 | 1 | 1 | | 51164 | PPTG_11895 | Phytophthora parasitica INRA-310 hypothetical protein (463 aa) |
| 3274 | 3858 | PPTG_04709 | 32,58 | 7 | 0 | 3 | 7 | | 3,14E+04 | 4,73E+05 | 2 | 2 | 0 | 1 | 2 | | 38008 | PPTG_04709 | Phytophthora parasitica INRA-310 RNA binding protein (344 aa) |
| 3442 | 3705 | PPTG_03901 | 32,53 | 5 | 0 | 5 | 5 | | 3,57E+05 | 3,07E+05 | 1 | 1 | 0 | 1 | 1 | | 38184 | PPTG_03901 | Phytophthora parasitica INRA-310 F-box protein (331 aa) |
| 3490 | 3485 | PPTG_04785 | 32,5 | 3 | 0 | 0 | 3 | | | 1,27E+05 | 2 | 2 | 0 | 0 | 2 | | 120572 | PPTG_04785 | Phytophthora parasitica INRA-310 transient Receptor Ca2 Channel (Trp-CC) family (1109 aa) |
| 3596 | 4028 | PPTG_02794 | 32,5 | 5 | 0 | 5 | 5 | | 9,45E+04 | 1,70E+05 | 1 | 1 | 0 | 1 | 1 | | 33300 | PPTG_02794 | Phytophthora parasitica INRA-310 U3 small nucleolar ribonucleoprotein IMP4 (288 aa) |
| 4637 | 4485 | PPTG_16009 | 32,49 | 3 | 0 | 3 | 0 | | 2,61E+05 | | 1 | 1 | 0 | 1 | 0 | Carbamidomethylation | 49114 | PPTG_16009 | Phytophthora parasitica INRA-310 hypothetical protein (442 aa) |
| 3448 | 3847 | PPTG_08273 | 32,48 | 7 | 0 | 7 | 7 | | 2,34E+06 | 5,70E+06 | 1 | 1 | 0 | 1 | 1 | | 16399 | PPTG_08273 | Phytophthora parasitica INRA-310 ubiquitin-conjugating enzyme E2-16 kDa (148 aa) |
| 4257 | 4008 | PPTG_08908 | 32,48 | 2 | 0 | 0 | 2 | | | 3,18E+05 | 1 | 1 | 0 | 0 | 1 | | 66457 | PPTG_08908 | Phytophthora parasitica INRA-310 hypothetical protein (616 aa) |
| 4176 | 4058 | PPTG_15853 | 32,43 | 3 | 0 | 3 | 0 | | 2,77E+05 | | 1 | 1 | 0 | 1 | 0 | | 45341 | PPTG_15853 | Phytophthora parasitica INRA-310 myb-like DNA-binding protein (415 aa) |
| 4176 | 4486 | PPTG_15852 | 32,43 | 2 | 0 | 2 | 0 | | 2,77E+05 | | 1 | 1 | 0 | 1 | 0 | | 55906 | PPTG_15852 | Phytophthora parasitica INRA-310 myb-like DNA-binding protein (498 aa) |
| 3803 | 4424 | PPTG_17567 | 32,42 | 3 | 0 | 3 | 3 | | 2,83E+05 | 3,11E+05 | 1 | 1 | 0 | 1 | 1 | | 36155 | PPTG_17567 | Phytophthora parasitica INRA-310 hypothetical protein (322 aa) |
| 4264 | 4047 | PPTG_06374 | 32,42 | 1 | 0 | 1 | 0 | | 7,95E+04 | | 1 | 1 | 0 | 1 | 0 | | 123334 | PPTG_06374 | Phytophthora parasitica INRA-310 hypothetical protein (1127 aa) |
| 3699 | 4193 | PPTG_03271 | 32,4 | 11 | 0 | 11 | 11 | | 1,41E+05 | 2,16E+05 | 1 | 1 | 0 | 1 | 1 | Acetylation (N-term) | 15248 | PPTG_03271 | Phytophthora parasitica INRA-310 hypothetical protein (133 aa) |
| 3695 | 4171 | PPTG_10731 | 32,4 | 5 | 0 | 5 | 5 | 0 | 0 | | 1 | 1 | 0 | 1 | 1 | | 27848 | PPTG_10731 | Phytophthora parasitica INRA-310 hypothetical protein (252 aa) |
| 3597 | 4053 | PPTG_01573 | 32,4 | 1 | 0 | 1 | 1 | | 3,46E+05 | 3,82E+05 | 1 | 1 | 0 | 1 | 1 | | 65405 | PPTG_01573 | Phytophthora parasitica INRA-310 hypothetical protein (589 aa) |
| 2866 | 3116 | PPTG_09700 | 32,37 | 29 | 0 | 29 | 8 | | 8,32E+05 | 0 | 2 | 2 | 0 | 3 | 1 | Carbamidomethylation | 12573 | PPTG_09700 | Phytophthora parasitica INRA-310 cytolethal distending toxin A/C family protein (111 aa) |
| 4103 | 4014 | PPTG_06080 | 32,36 | 2 | 0 | 2 | 0 | | 3,83E+05 | | 1 | 1 | 0 | 1 | 0 | | 50889 | PPTG_06080 | Phytophthora parasitica INRA-310 RNA-binding protein (462 aa) |
| 3457 | 4208 | PPTG_19215 | 32,34 | 5 | 0 | 5 | 5 | | 6,82E+05 | 5,69E+05 | 1 | 1 | 0 | 1 | 1 | | 21960 | PPTG_19215 | Phytophthora parasitica INRA-310 cutinase (218 aa) |
| 3457 | 4466 | PPTG_19214 | 32,34 | 5 | 0 | 5 | 5 | | 6,82E+05 | 5,69E+05 | 1 | 1 | 0 | 1 | 1 | | 20984 | PPTG_19214 | Phytophthora parasitica INRA-310 cutinase (202 aa) |
| 3457 | 4467 | PPTG_07182 | 32,34 | 5 | 0 | 5 | 5 | | 6,82E+05 | 5,69E+05 | 1 | 1 | 0 | 1 | 1 | | 21328 | PPTG_07182 | Phytophthora parasitica INRA-310 cutinase (206 aa) |
| 4081 | 3580 | PPTG_04811 | 32,34 | 3 | 0 | 0 | 3 | | 0 | | 1 | 1 | 0 | 0 | 1 | | 45842 | PPTG_04811 | Phytophthora parasitica INRA-310 protein kinase (420 aa) |
| 3573 | 3730 | PPTG_03116 | 32,33 | 2 | 0 | 2 | 2 | | 6,06E+04 | 4,01E+05 | 1 | 1 | 0 | 1 | 1 | | 58576 | PPTG_03116 | Phytophthora parasitica INRA-310 proton-dependent Oligopeptide transporter (POT) family (535 aa) |
| 3573 | 3774 | PPTG_17303 | 32,33 | 2 | 0 | 2 | 2 | | 6,06E+04 | 4,01E+05 | 1 | 1 | 0 | 1 | 1 | | 67077 | PPTG_17303 | Phytophthora parasitica INRA-310 proton-dependent Oligopeptide transporter (POT) family (612 aa) |
| 3573 | 3775 | PPTG_03114 | 32,33 | 2 | 0 | 2 | 2 | | 6,06E+04 | 4,01E+05 | 1 | 1 | 0 | 1 | 1 | | 67823 | PPTG_03114 | Phytophthora parasitica INRA-310 proton-dependent Oligopeptide transporter (POT) family (619 aa) |
| 3573 | 3776 | PPTG_19948 | 32,33 | 2 | 0 | 2 | 2 | | 6,06E+04 | 4,01E+05 | 1 | 1 | 0 | 1 | 1 | | 69403 | PPTG_19948 | Phytophthora parasitica INRA-310 proton-dependent Oligopeptide transporter (POT) family (634 aa) |
| 4413 | 4255 | PPTG_17498 | 32,32 | 3 | 0 | 3 | 0 | | 8,11E+04 | | 1 | 1 | 0 | 1 | 0 | | 59803 | PPTG_17498 | Phytophthora parasitica INRA-310 carbohydrate-binding protein (580 aa) |
| 4450 | 4489 | PPTG_05062 | 32,31 | 1 | 0 | 1 | 0 | | 1,23E+05 | | 1 | 1 | 0 | 1 | 0 | | 138220 | PPTG_05062 | Phytophthora parasitica INRA-310 peptide-N(4)-(N-acetyl-beta-glucosaminyl)asparagine amidase (1257 aa) |
| 3229 | 4187 | PPTG_18928 | 32,3 | 3 | 3 | 3 | 3 | 1,14E+05 | 2,80E+05 | 8,25E+05 | 1 | 1 | 1 | 1 | 1 | Acetylation (N-term) | 32804 | PPTG_18928 | Phytophthora parasitica INRA-310 hypothetical protein (289 aa) |
| 3000 | 2985 | PPTG_15643 | 32,27 | 6 | 6 | 6 | 6 | 0 | 2,21E+05 | 2,26E+05 | 1 | 1 | 1 | 1 | 1 | | 18165 | PPTG_15643 | Phytophthora parasitica INRA-310 hypothetical protein (169 aa) |
| 3102 | 2976 | PPTG_08272 | 32,26 | 4 | 4 | 4 | 4 | 1,63E+05 | 3,08E+05 | 1,56E+06 | 1 | 1 | 1 | 1 | 1 | | 30562 | PPTG_08272 | Phytophthora parasitica INRA-310 syntaxin 7-like protein (272 aa) |
| 4384 | 3770 | PPTG_09703 | 32,25 | 3 | 0 | 0 | 3 | | | 1,05E+05 | 1 | 1 | 0 | 0 | 1 | | 54905 | PPTG_09703 | Phytophthora parasitica INRA-310 hypothetical protein (498 aa) |
| 4639 | 4491 | PPTG_19203 | 32,24 | 6 | 0 | 6 | 0 | | 8,05E+04 | | 1 | 1 | 0 | 1 | 0 | Carbamidomethylation | 31766 | PPTG_19203 | Phytophthora parasitica INRA-310 hypothetical protein (290 aa) |
| 4444 | 4452 | PPTG_13942 | 32,22 | 3 | 0 | 0 | 3 | | 0 | | 1 | 1 | 0 | 0 | 1 | | 34094 | PPTG_13942 | Phytophthora parasitica INRA-310 5'-AMP-activated protein kinase subunit beta (308 aa) |
| 3042 | 3148 | PPTG_11927 | 32,18 | 1 | 0 | 1 | 0 | 6,54E+04 | 4,55E+05 | | 2 | 2 | 1 | 2 | 0 | Carbamidomethylation | 230337 | PPTG_11927 | Phytophthora parasitica INRA-310 hypothetical protein (2046 aa) |
| 4284 | 4272 | PPTG_09672 | 32,18 | 2 | 0 | 2 | 0 | 0 | | | 1 | 1 | 0 | 1 | 0 | | 52889 | PPTG_09672 | Phytophthora parasitica INRA-310 hypothetical protein (477 aa) |
| 3283 | 4443 | PPTG_05435 | 32,15 | 18 | 18 | 18 | 18 | 7,91E+05 | 1,94E+06 | 5,43E+06 | 1 | 1 | 1 | 1 | 1 | Carbamidomethylation | 12478 | PPTG_05435 | Phytophthora parasitica INRA-310 hypothetical protein (109 aa) |
| 3443 | 3755 | PPTG_03873 | 32,14 | 4 | 0 | 0 | 4 | | | 2,47E+06 | 2 | 2 | 0 | 0 | 2 | | 42870 | PPTG_03873 | Phytophthora parasitica INRA-310 CDP-alcohol phosphatidyltransferase (394 aa) |
| 3396 | 3009 | PPTG_14726 | 32,1 | 4 | 0 | 4 | 4 | | 2,87E+05 | 2,92E+05 | 1 | 1 | 0 | 1 | 1 | | 26199 | PPTG_14726 | Phytophthora parasitica INRA-310 hypothetical protein (244 aa) |
| 4106 | 4036 | PPTG_09993 | 32,09 | 1 | 0 | 1 | 0 | | 6,26E+04 | | 1 | 1 | 0 | 1 | 0 | | 125600 | PPTG_09993 | Phytophthora parasitica INRA-310 sulfite reductase [NADPH] flavoprotein component (1149 aa) |
| 4297 | 4495 | PPTG_04777 | 32,08 | 6 | 0 | 6 | 0 | | 1,86E+05 | | 1 | 1 | 0 | 1 | 0 | | 24512 | PPTG_04777 | Phytophthora parasitica INRA-310 hypothetical protein (219 aa) |
| 3479 | 3143 | PPTG_02524 | 32,04 | 3 | 0 | 1 | 2 | | 1,80E+05 | 4,34E+05 | 2 | 2 | 0 | 1 | 1 | | 61209 | PPTG_02524 | Phytophthora parasitica INRA-310 asparaginyl-tRNA synthetase (548 aa) |
| 4633 | 4446 | PPTG_15157 | 32,01 | 3 | 0 | 0 | 3 | | | 1,38E+05 | 1 | 1 | 0 | 0 | 1 | | 52234 | PPTG_15157 | Phytophthora parasitica INRA-310 hypothetical protein (475 aa) |
| 3735 | 4447 | PPTG_06436 | 31,97 | 3 | 0 | 3 | 3 | | 1,15E+05 | 2,69E+05 | 1 | 1 | 0 | 1 | 1 | Carbamidomethylation | 39084 | PPTG_06436 | Phytophthora parasitica INRA-310 hypothetical protein (350 aa) |
| 4385 | 3824 | PPTG_05549 | 31,95 | 3 | 0 | 0 | 3 | | | 9,17E+05 | 1 | 1 | 0 | 0 | 1 | | 37196 | PPTG_05549 | Phytophthora parasitica INRA-310 hypothetical protein (337 aa) |
| 3225 | 4070 | PPTG_10501 | 31,95 | 2 | 2 | 2 | 2 | 1,76E+05 | 1,19E+05 | 1,02E+06 | 1 | 1 | 1 | 1 | 1 | | 72477 | PPTG_10501 | Phytophthora parasitica INRA-310 choline transporter-like protein (654 aa) |
| 4445 | 4462 | PPTG_19333 | 31,93 | 2 | 0 | 0 | 2 | | 0 | | 1 | 1 | 0 | 0 | 1 | | 57326 | PPTG_19333 | Phytophthora parasitica INRA-310 dolichyl-P-Man:Man(7)GlcNAc(2)-PP-dolichyl-alpha-1 6-mannosyltransferase (508 aa) |
| 3700 | 4195 | PPTG_00936 | 31,93 | 4 | 4 | 0 | 4 | 1,40E+05 | | 1,47E+06 | 1 | 1 | 1 | 0 | 1 | | 62779 | PPTG_00936 | Phytophthora parasitica INRA-310 presenilin-like protein (570 aa) |
| 3736 | 4454 | PPTG_08301 | 31,91 | 5 | 0 | 5 | 5 | | 8,74E+04 | 3,17E+05 | 1 | 1 | 0 | 1 | 1 | Acetylation (N-term) | 20002 | PPTG_08301 | Phytophthora parasitica INRA-310 hypothetical protein (179 aa) |
| 2439 | 4498 | PPTG_01489 | 31,9 | 6 | 0 | 6 | 6 | | 8,02E+05 | 2,30E+06 | 1 | 1 | 0 | 3 | 3 | Oxidation (M) | 19199 | PPTG_01489 | Phytophthora parasitica INRA-310 methionine-R-sulfoxide reductase (176 aa) |
| 4634 | 4453 | PPTG_13227 | 31,83 | 2 | 0 | 0 | 2 | | | 8,27E+04 | 1 | 1 | 0 | 0 | 1 | | 70806 | PPTG_13227 | Phytophthora parasitica INRA-310 hypothetical protein (638 aa) |
| 3798 | 4389 | PPTG_11392 | 31,8 | 6 | 0 | 6 | 0 | | 5,86E+05 | | 2 | 2 | 0 | 2 | 0 | | 32942 | PPTG_11392 | Phytophthora parasitica INRA-310 hypothetical protein (295 aa) |
| 4408 | 4200 | PPTG_00738 | 31,8 | 4 | 0 | 0 | 4 | | | 1,67E+05 | 1 | 1 | 0 | 0 | 1 | Carbamidomethylation | 33715 | PPTG_00738 | Phytophthora parasitica INRA-310 hypothetical protein (308 aa) |
| 3408 | 3780 | PPTG_18149 | 31,78 | 5 | 0 | 0 | 5 | | | 8,92E+05 | 2 | 2 | 0 | 0 | 2 | | 50388 | PPTG_18149 | Phytophthora parasitica INRA-310 ATP-binding Cassette (ABC) superfamily protein (463 aa) |
| 4277 | 4173 | PPTG_15893 | 31,72 | 4 | 0 | 0 | 4 | | | 7,49E+05 | 1 | 1 | 0 | 0 | 1 | | 26114 | PPTG_15893 | Phytophthora parasitica INRA-310 Rab28 family GTPase (236 aa) |
| 3547 | 3062 | PPTG_14735 | 31,71 | 5 | 0 | 0 | 5 | | | 1,15E+06 | 2 | 2 | 0 | 0 | 2 | | 45734 | PPTG_14735 | Phytophthora parasitica INRA-310 hypothetical protein (417 aa) |
| 4455 | 4506 | PPTG_11406 | 31,69 | 4 | 0 | 4 | 0 | | 5,51E+04 | | 1 | 1 | 0 | 1 | 0 | Carbamidomethylation | 34458 | PPTG_11406 | Phytophthora parasitica INRA-310 hypothetical protein (311 aa) |
| 4238 | 3593 | PPTG_04579 | 31,68 | 3 | 0 | 0 | 3 | | | 1,02E+05 | 1 | 1 | 0 | 0 | 1 | Oxidation (M); Dehydration; Deamidation (NQ) | 57915 | PPTG_04579 | Phytophthora parasitica INRA-310 major facilitator superfamily transporter (MFS) (534 aa) |
| 2760 | 3814 | PPTG_14028 | 31,63 | 31 | 10 | 31 | 10 | 3,21E+05 | 1,72E+06 | 2,80E+06 | 2 | 2 | 1 | 2 | 1 | Carbamidomethylation | 9359 | PPTG_14028 | Phytophthora parasitica INRA-310 30S ribosomal protein S17 (81 aa) |
| 4265 | 4050 | PPTG_05617 | 31,58 | 5 | 0 | 5 | 0 | | 1,24E+05 | | 1 | 1 | 0 | 1 | 0 | Carbamidomethylation | 46655 | PPTG_05617 | Phytophthora parasitica INRA-310 aminoacylase metalloprotease family M20A (418 aa) |
| 3714 | 4316 | PPTG_09352 | 31,55 | 4 | 0 | 4 | 4 | | 4,60E+05 | 3,31E+05 | 1 | 1 | 0 | 1 | 1 | | 37795 | PPTG_09352 | Phytophthora parasitica INRA-310 WD repeat protein 39 (338 aa) |
| 3709 | 4297 | PPTG_16104 | 31,53 | 3 | 0 | 3 | 3 | | 8,01E+04 | 1,63E+05 | 1 | 1 | 0 | 1 | 1 | | 42050 | PPTG_16104 | Phytophthora parasitica INRA-310 double stranded RNA binding (371 aa) |
| 4295 | 4469 | PPTG_13572 | 31,48 | 2 | 0 | 0 | 2 | | | 3,75E+05 | 1 | 1 | 0 | 0 | 1 | | 53781 | PPTG_13572 | Phytophthora parasitica INRA-310 amino Acid-Polyamine-Organocation (APC) family (494 aa) |
| 4295 | 4470 | PPTG_13571 | 31,48 | 2 | 0 | 0 | 2 | | | 3,75E+05 | 1 | 1 | 0 | 0 | 1 | | 54187 | PPTG_13571 | Phytophthora parasitica INRA-310 amino Acid-Polyamine-Organocation (APC) family (501 aa) |
| 4252 | 3980 | PPTG_08117 | 31,48 | 1 | 0 | 1 | 0 | | 1,64E+05 | | 1 | 1 | 0 | 1 | 0 | | 85644 | PPTG_08117 | Phytophthora parasitica INRA-310 centromere/kinetochore protein (774 aa) |
| 3357 | 3790 | PPTG_07285 | 31,47 | 5 | 5 | 5 | 0 | 2,39E+05 | 1,13E+06 | | 1 | 1 | 1 | 1 | 0 | | 16202 | PPTG_07285 | Phytophthora parasitica INRA-310 hypothetical protein (151 aa) |
| 4261 | 4030 | PPTG_05522 | 31,43 | 2 | 0 | 0 | 2 | | 0 | | 1 | 1 | 0 | 0 | 1 | | 88538 | PPTG_05522 | Phytophthora parasitica INRA-310 voltage-gated Ion Channel (VIC) superfamily (787 aa) |
| 3701 | 4205 | PPTG_12131 | 31,43 | 1 | 0 | 1 | 1 | | 8,40E+04 | 2,35E+05 | 1 | 1 | 0 | 1 | 1 | | 152791 | PPTG_12131 | Phytophthora parasitica INRA-310 hypothetical protein (1344 aa) |
| 3284 | 4471 | PPTG_07060 | 31,41 | 11 | 11 | 11 | 11 | 2,08E+04 | 5,17E+05 | 5,49E+05 | 1 | 1 | 1 | 1 | 1 | | 10142 | PPTG_07060 | Phytophthora parasitica INRA-310 hypothetical protein (94 aa) |
| 4389 | 3864 | PPTG_08196 | 31,38 | 7 | 0 | 0 | 7 | | | 1,57E+05 | 1 | 1 | 0 | 0 | 1 | Carbamidomethylation | 15915 | PPTG_08196 | Phytophthora parasitica INRA-310 hypothetical protein (148 aa) |
| 4281 | 4207 | PPTG_00305 | 31,38 | 26 | 0 | 0 | 26 | | | 7,70E+04 | 1 | 1 | 0 | 0 | 1 | | 8944 | PPTG_00305 | Phytophthora parasitica INRA-310 hypothetical protein (78 aa) |
| 4635 | 4473 | PPTG_19173 | 31,35 | 4 | 0 | 0 | 4 | | | 3,06E+05 | 1 | 1 | 0 | 0 | 1 | | 30352 | PPTG_19173 | Phytophthora parasitica INRA-310 Ras-like protein (270 aa) |
| 4447 | 4474 | PPTG_16747 | 31,32 | 4 | 0 | 0 | 4 | | | 7,59E+05 | 1 | 1 | 0 | 0 | 1 | | 32862 | PPTG_16747 | Phytophthora parasitica INRA-310 mitochondrial Carrier (MC) family (307 aa) |
| 3223 | 3997 | PPTG_03639 | 31,32 | 3 | 2 | 0 | 3 | 1,52E+05 | | 1,60E+06 | 2 | 2 | 1 | 0 | 2 | | 49976 | PPTG_03639 | Phytophthora parasitica INRA-310 dicarboxylate/Amino Acid:Cation (Na or H) Symporter (DAACS) family (459 aa) |
| 4296 | 4475 | PPTG_05832 | 31,29 | 4 | 0 | 0 | 4 | | | 7,58E+05 | 1 | 1 | 0 | 0 | 1 | Oxidation (M); Deamidation (NQ) | 34635 | PPTG_05832 | Phytophthora parasitica INRA-310 zinc (Zn2)-Iron (Fe2) Permease (ZIP) family (327 aa) |
| 3737 | 4477 | PPTG_18310 | 31,26 | 3 | 0 | 3 | 3 | | 2,34E+05 | 7,11E+05 | 1 | 1 | 0 | 1 | 1 | | 38176 | PPTG_18310 | Phytophthora parasitica INRA-310 hypothetical protein (334 aa) |
| 3406 | 3630 | PPTG_00841 | 31,25 | 2 | 0 | 1 | 1 | 0 | | 2,00E+05 | 2 | 2 | 0 | 1 | 1 | Carbamidomethylation | 135675 | PPTG_00841 | Phytophthora parasitica INRA-310 hypothetical protein (1212 aa) |
| 4459 | 4540 | PPTG_08950 | 31,22 | 5 | 0 | 5 | 0 | | 7,22E+05 | | 1 | 1 | 0 | 1 | 0 | | 18295 | PPTG_08950 | Phytophthora parasitica INRA-310 elicitin-like protein (185 aa) |
| 3806 | 4479 | PPTG_00664 | 31,18 | 5 | 0 | 5 | 5 | | 2,54E+05 | 5,58E+05 | 1 | 1 | 0 | 1 | 1 | | 27719 | PPTG_00664 | Phytophthora parasitica INRA-310 hypothetical protein (242 aa) |
| 4105 | 4035 | PPTG_12224 | 31,14 | 1 | 0 | 0 | 1 | | | 2,96E+05 | 1 | 1 | 0 | 0 | 1 | | 85454 | PPTG_12224 | Phytophthora parasitica INRA-310 signal peptide peptidase-like protein (791 aa) |
| 4028 | 3977 | PPTG_17143 | 31,14 | 1 | 0 | 1 | 0 | 0 | | | 1 | 1 | 0 | 1 | 0 | | 111853 | PPTG_17143 | Phytophthora parasitica INRA-310 oxoglutarate dehydrogenase (succinyl-transferring) E1 component (987 aa) |
| 3738 | 4480 | PPTG_19489 | 31,13 | 5 | 0 | 5 | 5 | | 3,66E+05 | 1,82E+05 | 1 | 1 | 0 | 1 | 1 | | 25584 | PPTG_19489 | Phytophthora parasitica INRA-310 hypothetical protein (248 aa) |
| 4246 | 3870 | PPTG_10663 | 31,13 | 2 | 0 | 0 | 2 | | | 1,97E+05 | 1 | 1 | 0 | 0 | 1 | | 47613 | PPTG_10663 | Phytophthora parasitica INRA-310 dTDP-D-glucose 4 6-dehydratase (422 aa) |
| 4287 | 4308 | PPTG_13094 | 31,12 | 3 | 0 | 0 | 3 | | | 4,49E+05 | 1 | 1 | 0 | 0 | 1 | | 64981 | PPTG_13094 | Phytophthora parasitica INRA-310 hypothetical protein (566 aa) |
| 4636 | 4481 | PPTG_12845 | 31,12 | 5 | 0 | 0 | 5 | | | 7,47E+05 | 1 | 1 | 0 | 0 | 1 | | 28106 | PPTG_12845 | Phytophthora parasitica INRA-310 hypothetical protein (267 aa) |
| 4242 | 3785 | PPTG_11203 | 31,06 | 2 | 0 | 0 | 2 | | | 1,24E+05 | 1 | 1 | 0 | 0 | 1 | | 54070 | PPTG_11203 | Phytophthora parasitica INRA-310 protein kinase (484 aa) |
| 4027 | 3965 | PPTG_11287 | 31,06 | 3 | 0 | 3 | 0 | | 2,49E+05 | | 1 | 1 | 0 | 1 | 0 | | 39442 | PPTG_11287 | Phytophthora parasitica INRA-310 WD domain-containing protein (357 aa) |
| 3168 | 3804 | PPTG_04289 | 31,05 | 4 | 0 | 2 | 4 | | 2,58E+05 | 6,49E+05 | 2 | 2 | 0 | 1 | 2 | Oxidation (M) | 63790 | PPTG_04289 | Phytophthora parasitica INRA-310 methionyl-tRNA synthetase (566 aa) |
| 3742 | 4519 | PPTG_14742 | 31,02 | 3 | 0 | 3 | 3 | | 3,41E+05 | 4,16E+05 | 1 | 1 | 0 | 1 | 1 | | 46597 | PPTG_14742 | Phytophthora parasitica INRA-310 hypothetical protein (415 aa) |
| 3407 | 3707 | PPTG_07331 | 31,01 | 4 | 0 | 4 | 4 | | 5,61E+05 | 5,23E+05 | 1 | 1 | 0 | 1 | 1 | | 27940 | PPTG_07331 | Phytophthora parasitica INRA-310 hypothetical protein (261 aa) |
| 3983 | 3458 | PPTG_19137 | 30,96 | 2 | 0 | 0 | 2 | | | 1,47E+05 | 1 | 1 | 0 | 0 | 1 | | 82626 | PPTG_19137 | Phytophthora parasitica INRA-310 patatin-like phospholipase (734 aa) |
| 4449 | 4484 | PPTG_09182 | 30,95 | 3 | 0 | 0 | 3 | | | 9,13E+04 | 1 | 1 | 0 | 0 | 1 | | 47027 | PPTG_09182 | Phytophthora parasitica INRA-310 hypothetical protein (414 aa) |
| 4432 | 4390 | PPTG_04420 | 30,91 | 1 | 0 | 1 | 0 | | 2,01E+05 | | 1 | 1 | 0 | 1 | 0 | | 92952 | PPTG_04420 | Phytophthora parasitica INRA-310 digestive organ expansion factor (808 aa) |
| 4638 | 4487 | PPTG_00170 | 30,89 | 11 | 0 | 0 | 11 | | | 7,67E+04 | 1 | 1 | 0 | 0 | 1 | | 19675 | PPTG_00170 | Phytophthora parasitica INRA-310 alcohol dehydrogenase (189 aa) |
| 3503 | 3632 | PPTG_09089 | 30,88 | 7 | 0 | 0 | 7 | | | 1,21E+06 | 2 | 2 | 0 | 0 | 2 | | 33675 | PPTG_09089 | Phytophthora parasitica INRA-310 hypothetical protein (297 aa) |
| 3809 | 4553 | PPTG_05037 | 30,88 | 3 | 0 | 3 | 3 | | 2,05E+05 | 2,42E+05 | 1 | 1 | 0 | 1 | 1 | Carbamidomethylation | 30361 | PPTG_05037 | Phytophthora parasitica INRA-310 SCP-like extracellular protein (289 aa) |

| | | | | | | | | | | | | | | | | | | | |
|---|---|---|---|---|---|---|---|---|---|---|---|---|---|---|---|---|---|---|---|
| 3739 | 4503 | PPTG_04560 | 30,81 | 6 | 6 | 0 | 6 | 2,84E+05 | | 1,56E+06 | 1 | 1 | 1 | 0 | 1 | | 15986 | PPTG_04560 | Phytophthora parasitica INRA-310 hypothetical protein (143 aa) |
| 4436 | 4415 | PPTG_16439 | 30,79 | 2 | 0 | 0 | 2 | | | 4,68E+05 | 1 | 1 | 0 | 0 | 1 | | 44071 | PPTG_16439 | Phytophthora parasitica INRA-310 hypothetical protein (399 aa) |
| 3447 | 3842 | PPTG_10002 | 30,75 | 4 | 0 | 4 | 4 | | 4,97E+05 | 7,02E+05 | 1 | 1 | 0 | 1 | 1 | Carbamidomethylation | 23381 | PPTG_10002 | Phytophthora parasitica INRA-310 maltose O-acetyltransferase (213 aa) |
| 4452 | 4497 | PPTG_19631 | 30,69 | 9 | 0 | 0 | 9 | | | 4,20E+05 | 1 | 1 | 0 | 0 | 1 | Dehydration; Deamidation (NQ) | 13995 | PPTG_19631 | Phytophthora parasitica INRA-310 hypothetical protein (123 aa) |
| 4416 | 4269 | PPTG_04464 | 30,67 | 3 | 0 | 0 | 3 | | | 3,55E+05 | 1 | 1 | 0 | 0 | 1 | | 46047 | PPTG_04464 | Phytophthora parasitica INRA-310 hypothetical protein (413 aa) |
| 4453 | 4500 | PPTG_10885 | 30,64 | 3 | 0 | 0 | 3 | | | 3,58E+05 | 1 | 1 | 0 | 0 | 1 | | 38058 | PPTG_10885 | Phytophthora parasitica INRA-310 scavenger mRNA-decapping enzyme DcpS (331 aa) |
| 3083 | 3765 | PPTG_19391 | 30,64 | 2 | 2 | 2 | 2 | 1,28E+05 | 5,44E+05 | 1,01E+06 | 1 | 1 | 1 | 1 | 1 | | 62169 | PPTG_19391 | Phytophthora parasitica INRA-310 RIB72 protein (532 aa) |
| 4298 | 4499 | PPTG_14330 | 30,59 | 11 | 0 | 0 | 11 | | | 1,01E+06 | 1 | 1 | 0 | 0 | 1 | Acetylation (N-term) | 15620 | PPTG_14330 | Phytophthora parasitica INRA-310 cytochrome b5 (148 aa) |
| 4055 | 3885 | PPTG_05546 | 30,56 | 1 | 0 | 1 | 0 | 0 | | | 1 | 1 | 0 | 1 | 0 | | 227686 | PPTG_05546 | Phytophthora parasitica INRA-310 hypothetical protein (2062 aa) |
| 4646 | 4586 | PPTG_15918 | 30,54 | 6 | 0 | 6 | 0 | 0 | | | 1 | 1 | 0 | 1 | 0 | | 35354 | PPTG_15918 | Phytophthora parasitica INRA-310 delta-1-pyrroline-5-carboxylate synthetase (325 aa) |
| 4396 | 4062 | PPTG_16012 | 30,5 | 3 | 0 | 0 | 3 | | | 3,16E+05 | 1 | 1 | 0 | 0 | 1 | | 38572 | PPTG_16012 | Phytophthora parasitica INRA-310 hypothetical protein (336 aa) |
| 3807 | 4504 | PPTG_14283 | 30,47 | 3 | 0 | 3 | 3 | | 9,96E+05 | 1,44E+06 | 1 | 1 | 0 | 1 | 1 | | 35515 | PPTG_14283 | Phytophthora parasitica INRA-310 hypothetical protein (312 aa) |
| 3584 | 3940 | PPTG_09582 | 30,38 | 1 | 0 | 0 | 1 | | | 1,94E+05 | 2 | 2 | 0 | 0 | 2 | | 213516 | PPTG_09582 | Phytophthora parasitica INRA-310 voltage-gated Ion Channel (VIC) superfamily (1903 aa) |
| 4107 | 4046 | PPTG_06092 | 30,38 | 1 | 0 | 0 | 1 | | 0 | | 1 | 1 | 0 | 0 | 1 | Amidation | 111764 | PPTG_06092 | Phytophthora parasitica INRA-310 hypothetical protein (995 aa) |
| 3241 | 4590 | PPTG_08736 | 30,38 | 2 | 2 | 2 | 2 | 3,78E+04 | 2,57E+05 | 4,42E+05 | 1 | 1 | 1 | 1 | 1 | | 73525 | PPTG_08736 | Phytophthora parasitica INRA-310 peroxisomal acyl-coenzyme A oxidase (664 aa) |
| 4443 | 4450 | PPTG_12023 | 30,31 | 4 | 0 | 4 | 0 | 0 | | | 1 | 1 | 0 | 1 | 0 | | 36960 | PPTG_12023 | Phytophthora parasitica INRA-310 D-isomer specific 2-hydroxyacid dehydrogenase (334 aa) |
| 3380 | 3691 | PPTG_03167 | 30,23 | 1 | 0 | 1 | 1 | | 1,60E+05 | 2,14E+05 | 1 | 1 | 0 | 1 | 1 | | 154458 | PPTG_03167 | Phytophthora parasitica INRA-310 hypothetical protein (1425 aa) |
| 3458 | 4365 | PPTG_07241 | 30,19 | 10 | 0 | 0 | 10 | | | 2,15E+06 | 1 | 1 | 0 | 0 | 2 | Oxidation (M) | 12825 | PPTG_07241 | Phytophthora parasitica INRA-310 dolichyl-diphosphooligosaccharide-protein glycosyltransferase subunit (117 aa) |
| 4640 | 4520 | PPTG_03831 | 30,13 | 6 | 0 | 0 | 6 | | | 3,03E+05 | 1 | 1 | 0 | 0 | 1 | | 17044 | PPTG_03831 | Phytophthora parasitica INRA-310 hypothetical protein (153 aa) |
| 4640 | 4523 | PPTG_04985 | 30,13 | 4 | 0 | 0 | 4 | | | 3,03E+05 | 1 | 1 | 0 | 0 | 1 | | 24095 | PPTG_04985 | Phytophthora parasitica INRA-310 hypothetical protein (215 aa) |
| 4640 | 4525 | PPTG_08022 | 30,13 | 4 | 0 | 0 | 4 | | | 3,03E+05 | 1 | 1 | 0 | 0 | 1 | | 29081 | PPTG_08022 | Phytophthora parasitica INRA-310 hypothetical protein (257 aa) |
| 4640 | 4526 | PPTG_00643 | 30,13 | 2 | 0 | 0 | 2 | | | 3,03E+05 | 1 | 1 | 0 | 0 | 1 | | 40880 | PPTG_00643 | Phytophthora parasitica INRA-310 hypothetical protein (368 aa) |
| 4425 | 4348 | PPTG_06040 | 30,11 | 4 | 0 | 4 | 0 | | 1,18E+05 | | 1 | 1 | 0 | 1 | 0 | | 43658 | PPTG_06040 | Phytophthora parasitica INRA-310 hypothetical protein (394 aa) |
| 4457 | 4524 | PPTG_10759 | 30,06 | 4 | 0 | 0 | 4 | | | 3,85E+06 | 1 | 1 | 0 | 0 | 1 | | 28136 | PPTG_10759 | Phytophthora parasitica INRA-310 adaptin-like protein (260 aa) |
| 3455 | 4017 | PPTG_14550 | 30,05 | 2 | 0 | 0 | 2 | | | 2,03E+05 | 2 | 2 | 0 | 0 | 2 | | 94509 | PPTG_14550 | Phytophthora parasitica INRA-310 Cullin family protein (824 aa) |
| 3808 | 4549 | PPTG_08399 | 30,03 | 8 | 0 | 8 | 8 | | 1,20E+05 | 3,37E+05 | 1 | 1 | 0 | 1 | 1 | | 13196 | PPTG_08399 | Phytophthora parasitica INRA-310 hypothetical protein (116 aa) |
| 3379 | 3685 | PPTG_06197 | 30,02 | 1 | 0 | 1 | 1 | | 1,80E+05 | 2,90E+05 | 1 | 1 | 0 | 1 | 1 | | 110028 | PPTG_06197 | Phytophthora parasitica INRA-310 hypothetical protein (960 aa) |
| 4109 | 4347 | PPTG_09818 | 30,01 | 1 | 0 | 0 | 1 | | | 5,19E+05 | 1 | 1 | 0 | 0 | 1 | | 105678 | PPTG_09818 | Phytophthora parasitica INRA-310 vacuolar protein sorting-associated protein (952 aa) |
| 3744 | 4537 | PPTG_04114 | 30 | 2 | 0 | 2 | 2 | | 1,90E+05 | 2,76E+05 | 1 | 1 | 0 | 1 | 1 | | 41840 | PPTG_04114 | Phytophthora parasitica INRA-310 hypothetical protein (381 aa) |
| 4641 | 4527 | PPTG_14328 | 29,98 | 8 | 0 | 0 | 8 | | | 2,15E+05 | 1 | 1 | 0 | 0 | 1 | | 17096 | PPTG_14328 | Phytophthora parasitica INRA-310 hypothetical protein (155 aa) |
| 2955 | 4528 | PPTG_02059 | 29,96 | 7 | 0 | 7 | 7 | | 3,51E+05 | 7,71E+05 | 1 | 1 | 0 | 2 | 2 | Oxidation (M) | 19990 | PPTG_02059 | Phytophthora parasitica INRA-310 DNA-directed RNA polymerase II polypeptide (178 aa) |
| 4421 | 4311 | PPTG_14811 | 29,95 | 5 | 0 | 0 | 5 | | | 5,40E+05 | 1 | 1 | 0 | 0 | 1 | | 23631 | PPTG_14811 | Phytophthora parasitica INRA-310 HAD superfamily (subfamily IIIA) phosphatase (212 aa) |
| 4665 | 4715 | PPTG_00444 | 29,94 | 2 | 2 | 0 | 0 | 2,25E+05 | | | 1 | 1 | 1 | 0 | 0 | | 33213 | PPTG_00444 | Phytophthora parasitica INRA-310 hypothetical protein (299 aa) |
| 4155 | 3711 | PPTG_14946 | 29,93 | 3 | 0 | 0 | 3 | | | 7,47E+05 | 1 | 1 | 0 | 0 | 1 | | 51154 | PPTG_14946 | Phytophthora parasitica INRA-310 hypothetical protein (484 aa) |
| 4426 | 4349 | PPTG_09877 | 29,85 | 10 | 0 | 0 | 10 | | | 3,77E+05 | 1 | 1 | 0 | 0 | 1 | | 11343 | PPTG_09877 | Phytophthora parasitica INRA-310 hypothetical protein (105 aa) |
| 3721 | 4346 | PPTG_05973 | 29,83 | 14 | 0 | 14 | 14 | | 5,64E+04 | 3,14E+05 | 1 | 1 | 0 | 1 | 1 | | 7663 | PPTG_05973 | Phytophthora parasitica INRA-310 hypothetical protein (75 aa) |
| 3009 | 4543 | PPTG_11905 | 29,82 | 11 | 0 | 0 | 11 | | | 1,08E+08 | 1 | 1 | 0 | 0 | 3 | Oxidation (M) | 17253 | PPTG_11905 | Phytophthora parasitica INRA-310 V-type proton ATPase proteolipid subunit (172 aa) |
| 3009 | 4544 | PPTG_11908 | 29,82 | 11 | 0 | 0 | 11 | | | 1,08E+08 | 1 | 1 | 0 | 0 | 3 | Oxidation (M) | 17253 | PPTG_11908 | Phytophthora parasitica INRA-310 V-type proton ATPase proteolipid subunit (172 aa) |
| 3462 | 4838 | PPTG_15070 | 29,8 | 4 | 0 | 4 | 0 | | 2,49E+05 | | 2 | 2 | 0 | 2 | 0 | | 56703 | PPTG_15070 | Phytophthora parasitica INRA-310 splicing factor 3A (491 aa) |
| 4007 | 3815 | PPTG_14048 | 29,74 | 1 | 0 | 0 | 1 | | | 2,39E+05 | 1 | 1 | 0 | 0 | 1 | Oxidation (M) | 109470 | PPTG_14048 | Phytophthora parasitica INRA-310 hypothetical protein (985 aa) |
| 3745 | 4563 | PPTG_00025 | 29,74 | 1 | 1 | 1 | 0 | 1,65E+06 | 2,84E+06 | | 1 | 1 | 1 | 1 | 0 | Oxidation (M) | 56705 | PPTG_00025 | Phytophthora parasitica INRA-310 amino Acid/Auxin Permease (AAAP) family (519 aa) |
| 3439 | 3638 | PPTG_19889 | 29,71 | 14 | 0 | 14 | 14 | | 4,65E+05 | 4,78E+05 | 1 | 1 | 0 | 1 | 1 | | 19650 | PPTG_19889 | Phytophthora parasitica INRA-310 hypothetical protein (174 aa) |
| 4032 | 4095 | PPTG_10661 | 29,7 | 1 | 0 | 1 | 0 | | 5,21E+04 | | 1 | 1 | 0 | 1 | 0 | Deamidation (NQ) | 75777 | PPTG_10661 | Phytophthora parasitica INRA-310 urocanate hydratase (691 aa) |
| 4428 | 4357 | PPTG_03920 | 29,7 | 2 | 0 | 0 | 2 | | | 5,10E+05 | 1 | 1 | 0 | 0 | 1 | | 56007 | PPTG_03920 | Phytophthora parasitica INRA-310 serine protease family S33 (498 aa) |
| 4446 | 4465 | PPTG_02380 | 29,68 | 3 | 0 | 3 | 0 | 0 | | | 1 | 1 | 0 | 1 | 0 | | 33493 | PPTG_02380 | Phytophthora parasitica INRA-310 down syndrome critical region protein 3 (298 aa) |
| 3723 | 4366 | PPTG_12837 | 29,63 | 3 | 0 | 3 | 3 | | 1,44E+05 | 2,57E+05 | 1 | 1 | 0 | 1 | 1 | | 35731 | PPTG_12837 | Phytophthora parasitica INRA-310 hypothetical protein (331 aa) |
| 4642 | 4547 | PPTG_09345 | 29,62 | 14 | 0 | 0 | 14 | | | 4,25E+05 | 1 | 1 | 0 | 0 | 1 | Carbamidomethylation | 13588 | PPTG_09345 | Phytophthora parasitica INRA-310 hypothetical protein (120 aa) |
| 4048 | 3717 | PPTG_16344 | 29,53 | 2 | 0 | 2 | 0 | | 1,03E+05 | | 1 | 1 | 0 | 1 | 0 | | 69498 | PPTG_16344 | Phytophthora parasitica INRA-310 hypothetical protein (630 aa) |
| 3581 | 3881 | PPTG_07035 | 29,53 | 5 | 0 | 5 | 5 | | 2,65E+05 | 1,57E+05 | 1 | 1 | 0 | 1 | 1 | | 25091 | PPTG_07035 | Phytophthora parasitica INRA-310 hypothetical protein (230 aa) |
| 4291 | 4408 | PPTG_18657 | 29,53 | 2 | 0 | 0 | 2 | | | 2,08E+05 | 1 | 1 | 0 | 0 | 1 | | 55698 | PPTG_18657 | Phytophthora parasitica INRA-310 glycyl-tRNA synthetase (488 aa) |
| 4267 | 4055 | PPTG_01374 | 29,51 | 2 | 0 | 0 | 2 | | | 3,87E+05 | 1 | 1 | 0 | 0 | 1 | | 67529 | PPTG_01374 | Phytophthora parasitica INRA-310 hypothetical protein (604 aa) |
| 3633 | 4967 | PPTG_05502 | 29,5 | 14 | 0 | 14 | 0 | | 5,97E+05 | | 2 | 2 | 0 | 2 | 0 | | 13598 | PPTG_05502 | Phytophthora parasitica INRA-310 glutaredoxin (126 aa) |
| 4175 | 4039 | PPTG_13421 | 29,47 | 3 | 0 | 0 | 3 | | | 1,16E+06 | 1 | 1 | 0 | 0 | 1 | | 36896 | PPTG_13421 | Phytophthora parasitica INRA-310 protein kinase (328 aa) |
| 3333 | 4018 | PPTG_16574 | 29,41 | 5 | 0 | 5 | 5 | | 1,57E+07 | 3,09E+07 | 1 | 1 | 0 | 1 | 1 | | 15402 | PPTG_16574 | Phytophthora parasitica INRA-310 histone a (137 aa) |
| 3333 | 4019 | PPTG_16571 | 29,41 | 5 | 0 | 5 | 5 | | 1,57E+07 | 3,09E+07 | 1 | 1 | 0 | 1 | 1 | | 15402 | PPTG_16571 | Phytophthora parasitica INRA-310 histone a (137 aa) |
| 4053 | 3829 | PPTG_16997 | 29,41 | 2 | 0 | 2 | 0 | | 1,09E+05 | | 1 | 1 | 0 | 1 | 0 | | 92498 | PPTG_16997 | Phytophthora parasitica INRA-310 myosin-like protein (829 aa) |
| 4437 | 4422 | PPTG_14574 | 29,38 | 2 | 0 | 0 | 2 | | | 2,67E+05 | 1 | 1 | 0 | 0 | 1 | | 48830 | PPTG_14574 | Phytophthora parasitica INRA-310 calcium/calmodulin-dependent protein kinase (448 aa) |
| 4030 | 4038 | PPTG_16499 | 29,36 | 3 | 0 | 3 | 0 | | 1,10E+05 | | 1 | 1 | 0 | 1 | 0 | | 46307 | PPTG_16499 | Phytophthora parasitica INRA-310 hypothetical protein (437 aa) |
| 4648 | 4605 | PPTG_09523 | 29,35 | 8 | 0 | 8 | 0 | | 4,23E+05 | | 1 | 1 | 0 | 1 | 0 | | 13444 | PPTG_09523 | Phytophthora parasitica INRA-310 DNA-directed RNA polymerase II subunit J (120 aa) |
| 4293 | 4428 | PPTG_00949 | 29,33 | 3 | 0 | 0 | 3 | | | 2,75E+05 | 1 | 1 | 0 | 0 | 1 | Carbamidomethylation | 45368 | PPTG_00949 | Phytophthora parasitica INRA-310 E1 family ubiquitin activating enzyme (417 aa) |
| 4293 | 4429 | PPTG_00947 | 29,33 | 3 | 0 | 0 | 3 | | | 2,75E+05 | 1 | 1 | 0 | 0 | 1 | Carbamidomethylation | 45397 | PPTG_00947 | Phytophthora parasitica INRA-310 E1 family ubiquitin activating enzyme (417 aa) |
| 3741 | 4518 | PPTG_01604 | 29,32 | 7 | 0 | 7 | 7 | | 3,65E+05 | 1,45E+05 | 1 | 1 | 0 | 1 | 1 | | 19144 | PPTG_01604 | Phytophthora parasitica INRA-310 hypothetical protein (170 aa) |
| 3679 | 3807 | PPTG_01714 | 29,23 | 6 | 0 | 0 | 6 | | | 1,01E+05 | 2 | 2 | 0 | 0 | 2 | | 37494 | PPTG_01714 | Phytophthora parasitica INRA-310 secreted RxLR effector peptide (340 aa) |
| 3463 | 4865 | PPTG_09791 | 29,22 | 18 | 0 | 18 | 0 | | 5,11E+04 | | 2 | 2 | 0 | 2 | 0 | | 13924 | PPTG_09791 | Phytophthora parasitica INRA-310 hypothetical protein (125 aa) |
| 4649 | 4612 | PPTG_04566 | 29,21 | 5 | 0 | 5 | 0 | | 6,38E+05 | | 1 | 1 | 0 | 1 | 0 | Carbamidomethylation | 36289 | PPTG_04566 | Phytophthora parasitica INRA-310 carbohydrate-binding protein (366 aa) |
| 3050 | 3557 | PPTG_13203 | 29,19 | 4 | 4 | 4 | 4 | 6,51E+04 | 3,45E+05 | 8,92E+05 | 1 | 1 | 1 | 1 | 1 | Amidation | 30569 | PPTG_13203 | Phytophthora parasitica INRA-310 myosin-like protein (275 aa) |
| 3740 | 4505 | PPTG_18602 | 29,16 | 2 | 0 | 2 | 2 | | 2,32E+05 | 5,51E+05 | 1 | 1 | 0 | 1 | 1 | | 46916 | PPTG_18602 | Phytophthora parasitica INRA-310 hypothetical protein (423 aa) |
| 3612 | 4385 | PPTG_10263 | 29,15 | 2 | 2 | 0 | 2 | 3,68E+04 | | 3,20E+05 | 1 | 1 | 1 | 0 | 1 | | 60879 | PPTG_10263 | Phytophthora parasitica INRA-310 elongator complex protein 3 (542 aa) |
| 3613 | 4456 | PPTG_01814 | 29,15 | 7 | 0 | 7 | 7 | | 1,62E+05 0 | | 1 | 1 | 0 | 1 | 1 | | 17093 | PPTG_01814 | Phytophthora parasitica INRA-310 hypothetical protein (151 aa) |
| 3376 | 3610 | PPTG_09322 | 29,1 | 1 | 0 | 1 | 1 | | 3,06E+05 0 | | 1 | 1 | 0 | 1 | 1 | | 77729 | PPTG_09322 | Phytophthora parasitica INRA-310 hypothetical protein (711 aa) |
| 4052 | 3795 | PPTG_07525 | 29,09 | 2 | 0 | 2 | 0 | | 1,37E+05 | | 1 | 1 | 0 | 1 | 0 | Carbamidomethylation | 107430 | PPTG_07525 | Phytophthora parasitica INRA-310 DNA polymerase alpha-associated DNA helicase A (976 aa) |
| 4460 | 4561 | PPTG_09622 | 28,99 | 4 | 0 | 0 | 4 | | 0 | | 1 | 1 | 0 | 0 | 1 | | 46267 | PPTG_09622 | Phytophthora parasitica INRA-310 prenyltransferase-like protein (417 aa) |
| 4301 | 4522 | PPTG_12491 | 28,96 | 0 | 0 | 0 | 0 | | 1,62E+05 | | 1 | 1 | 0 | 1 | 0 | Carbamidomethylation | 375155 | PPTG_12491 | Phytophthora parasitica INRA-310 hypothetical protein (3366 aa) |
| 4268 | 4057 | PPTG_16830 | 28,91 | 5 | 0 | 0 | 5 | | | 1,88E+05 | 1 | 1 | 0 | 0 | 1 | Carbamidomethylation | 35391 | PPTG_16830 | Phytophthora parasitica INRA-310 hypothetical protein (317 aa) |
| 3177 | 4425 | PPTG_05957 | 28,91 | 2 | 2 | 2 | 2 | 4,35E+04 | 1,27E+05 | 4,39E+05 | 1 | 1 | 1 | 1 | 1 | | 59417 | PPTG_05957 | Phytophthora parasitica INRA-310 vacuolar amino acid transporter (532 aa) |
| 3178 | 4730 | PPTG_19627 | 28,88 | 3 | 0 | 2 | 3 | | 1,56E+05 | 1,75E+05 | 2 | 1 | 0 | 1 | 2 | Oxidation (M) | 50609 | PPTG_19627 | Phytophthora parasitica INRA-310 histone-binding protein RBBP7 (452 aa) |
| 4454 | 4502 | PPTG_00921 | 28,83 | 1 | 0 | 0 | 1 | | | 5,99E+05 | 1 | 1 | 0 | 0 | 1 | | 68672 | PPTG_00921 | Phytophthora parasitica INRA-310 hypothetical protein (627 aa) |
| 4441 | 4439 | PPTG_02407 | 28,74 | 1 | 0 | 0 | 1 | | | 1,51E+05 | 1 | 1 | 0 | 0 | 1 | | 114131 | PPTG_02407 | Phytophthora parasitica INRA-310 hypothetical protein (1045 aa) |
| 4269 | 4064 | PPTG_06409 | 28,73 | 2 | 0 | 0 | 2 | | | 2,60E+05 | 1 | 1 | 0 | 0 | 1 | | 67443 | PPTG_06409 | Phytophthora parasitica INRA-310 hypothetical protein (599 aa) |
| 4438 | 4427 | PPTG_03847 | 28,72 | 4 | 0 | 0 | 4 | | 0 | | 1 | 1 | 0 | 0 | 1 | | 64613 | PPTG_03847 | Phytophthora parasitica INRA-310 cell 5A endo-1 4-betaglucanase (578 aa) |
| 4448 | 4476 | PPTG_08102 | 28,71 | 1 | 0 | 0 | 1 | | | 3,62E+05 | 1 | 1 | 0 | 0 | 1 | | 121483 | PPTG_08102 | Phytophthora parasitica INRA-310 kinesin-like protein (1094 aa) |
| 4643 | 4570 | PPTG_19156 | 28,64 | 1 | 0 | 0 | 1 | | | 4,57E+04 | 1 | 1 | 0 | 0 | 1 | | 136383 | PPTG_19156 | Phytophthora parasitica INRA-310 guanylate cyclase (1220 aa) |
| 4439 | 4434 | PPTG_03955 | 28,55 | 2 | 0 | 0 | 2 | | | 2,10E+05 | 1 | 1 | 0 | 0 | 1 | Carbamidomethylation | 58558 | PPTG_03955 | Phytophthora parasitica INRA-310 GPI transamidase component (530 aa) |
| 4266 | 4052 | PPTG_19144 | 28,47 | 2 | 0 | 0 | 2 | | 0 | | 1 | 1 | 0 | 0 | 1 | | 53290 | PPTG_19144 | Phytophthora parasitica INRA-310 DEAD/DEAH box RNA helicase (486 aa) |
| 4179 | 4073 | PPTG_15993 | 28,46 | 5 | 0 | 5 | 0 | | 1,10E+05 | | 1 | 1 | 0 | 1 | 0 | | 34441 | PPTG_15993 | Phytophthora parasitica INRA-310 hypothetical protein (314 aa) |
| 4417 | 4270 | PPTG_10314 | 28,46 | 3 | 0 | 0 | 3 | | | 3,83E+05 | 1 | 1 | 0 | 0 | 1 | | 40612 | PPTG_10314 | Phytophthora parasitica INRA-310 beta-lactamase-like protein (365 aa) |
| 4657 | 4657 | PPTG_16257 | 28,45 | 17 | 0 | 17 | 0 | 0 | | | 1 | 1 | 0 | 1 | 0 | | 10007 | PPTG_16257 | Phytophthora parasitica INRA-310 hypothetical protein (91 aa) |
| 4657 | 4658 | PPTG_16257 | 28,45 | 16 | 0 | 16 | 0 | 0 | | | 1 | 1 | 0 | 1 | 0 | | 10669 | PPTG_16257 | Phytophthora parasitica INRA-310 hypothetical protein (97 aa) |
| 3732 | 4423 | PPTG_16492 | 28,45 | 3 | 0 | 0 | 3 | | | 1,16E+06 | 1 | 1 | 0 | 0 | 2 | Oxidation (M) | 40581 | PPTG_16492 | Phytophthora parasitica INRA-310 N(4)-(Beta-N-acetylglucosaminyl)-L-asparaginase (377 aa) |
| 3734 | 4440 | PPTG_08381 | 28,45 | 2 | 0 | 2 | 2 | | 2,21E+05 | 2,92E+05 | 1 | 1 | 0 | 1 | 1 | | 69138 | PPTG_08381 | Phytophthora parasitica INRA-310 hypothetical protein (619 aa) |
| 4089 | 3813 | PPTG_00383 | 28,45 | 4 | 0 | 0 | 4 | | | 1,44E+06 | 1 | 1 | 0 | 0 | 1 | | 32192 | PPTG_00383 | Phytophthora parasitica INRA-310 hypothetical protein (289 aa) |
| 3601 | 4079 | PPTG_16776 | 28,37 | 4 | 0 | 4 | 4 | | 3,48E+05 | 4,09E+05 | 1 | 1 | 0 | 1 | 1 | | 44507 | PPTG_16776 | Phytophthora parasitica INRA-310 p21-activated protein kinase-interacting protein (405 aa) |
| 3811 | 4588 | PPTG_10748 | 28,36 | 2 | 0 | 2 | 2 | | 1,92E+05 | 1,75E+05 | 1 | 1 | 0 | 1 | 1 | | 36473 | PPTG_10748 | Phytophthora parasitica INRA-310 hypothetical protein (340 aa) |
| 4647 | 4598 | PPTG_06994 | 28,33 | 1 | 0 | 0 | 1 | | | 4,34E+05 | 1 | 1 | 0 | 0 | 1 | | 61911 | PPTG_06994 | Phytophthora parasitica INRA-310 amino Acid-Polyamine-Organocation (APC) family (570 aa) |
| 3242 | 4664 | PPTG_07093 | 28,32 | 11 | 0 | 11 | 11 | | 3,06E+05 | 1,12E+06 | 1 | 1 | 0 | 1 | 2 | Oxidation (M) | 12069 | PPTG_07093 | Phytophthora parasitica INRA-310 dynein light chain 2B (110 aa) |
| 3614 | 4514 | PPTG_11786 | 28,32 | 3 | 0 | 3 | 3 | | 1,05E+05 | 9,40E+04 | 1 | 1 | 0 | 1 | 1 | | 47551 | PPTG_11786 | Phytophthora parasitica INRA-310 crinkler (CRN) family protein (433 aa) |
| 3615 | 4551 | PPTG_06244 | 28,31 | 3 | 0 | 0 | 3 | | | 1,05E+06 | 2 | 2 | 0 | 0 | 2 | | 56611 | PPTG_06244 | Phytophthora parasitica INRA-310 protein kinase (516 aa) |
| 3240 | 4459 | PPTG_07906 | 28,29 | 8 | 8 | 8 | 8 | 2,64E+05 | 8,57E+05 | 1,59E+06 | 1 | 1 | 1 | 1 | 1 | Carbamidomethylation | 15797 | PPTG_07906 | Phytophthora parasitica INRA-310 eukaryotic translation initiation factor 1A (140 aa) |
| 4645 | 4579 | PPTG_14452 | 28,27 | 4 | 0 | 0 | 4 | | | 5,63E+05 | 1 | 1 | 0 | 0 | 1 | | 25983 | PPTG_14452 | Phytophthora parasitica INRA-310 major facilitator superfamily transporter (MFS) (243 aa) |
| 3616 | 4567 | PPTG_00844 | 28,25 | 1 | 0 | 1 | 1 | | 3,66E+05 | 7,18E+05 | 1 | 1 | 0 | 1 | 1 | | 75523 | PPTG_00844 | Phytophthora parasitica INRA-310 phosphatidylinositol-3 4 5-trisphosphate 3-phosphatase (687 aa) |
| 4644 | 4573 | PPTG_18924 | 28,24 | 2 | 0 | 0 | 2 | | | 3,49E+05 | 1 | 1 | 0 | 0 | 1 | | 68413 | PPTG_18924 | Phytophthora parasitica INRA-310 hypothetical protein (615 aa) |
| 4679 | 4766 | PPTG_11248 | 28,21 | 3 | 3 | 0 | 0 | 3,89E+04 | | | 1 | 1 | 1 | 0 | 0 | | 29689 | PPTG_11248 | Phytophthora parasitica INRA-310 3-oxo-5-alpha-steroid 4-dehydrogenase (260 aa) |
| 3509 | 3754 | PPTG_15413 | 28,16 | 4 | 4 | 4 | 0 | 7,31E+04 | 2,23E+05 | | 1 | 1 | 1 | 1 | 0 | | 25312 | PPTG_15413 | Phytophthora parasitica INRA-310 hypothetical protein (230 aa) |
| 4245 | 3810 | PPTG_02831 | 28,12 | 2 | 0 | 0 | 2 | | 0 | | 1 | 1 | 0 | 0 | 1 | | 56744 | PPTG_02831 | Phytophthora parasitica INRA-310 hypothetical protein (512 aa) |
| 3681 | 3976 | PPTG_11893 | 28,09 | 31 | 0 | 0 | 31 | | | 7,87E+04 | 1 | 1 | 0 | 0 | 2 | | 12476 | PPTG_11893 | Phytophthora parasitica INRA-310 hypothetical protein (118 aa) |
| 3528 | 4074 | PPTG_16016 | 28,08 | 1 | 0 | 1 | 1 | | 2,15E+05 | 4,05E+05 | 1 | 1 | 0 | 1 | 1 | | 141154 | PPTG_16016 | Phytophthora parasitica INRA-310 DNA mismatch repair protein Msh6 (1264 aa) |

| | | | | | | | | | | | | | | | | | | | |
|---|---|---|---|---|---|---|---|---|---|---|---|---|---|---|---|---|---|---|---|
| 4461 | 4575 | PPTG_14029 | 28,07 | 6 | 0 | 0 | 6 | | | 8,27E+04 | 1 | 1 | 0 | 0 | 1 | | 25803 | PPTG_14029 | Phytophthora parasitica INRA-310 hypothetical protein (233 aa) |
| 4160 | 3797 | PPTG_03276 | 28,05 | 2 | 0 | 0 | 2 | | | 4,46E+05 | 1 | 1 | 0 | 0 | 1 | | 71339 | PPTG_03276 | Phytophthora parasitica INRA-310 hypothetical protein (623 aa) |
| 4462 | 4576 | PPTG_14404 | 28,04 | 4 | 0 | 0 | 4 | | | 2,25E+05 | 1 | 1 | 0 | 0 | 1 | | 36262 | PPTG_14404 | Phytophthora parasitica INRA-310 hypothetical protein (327 aa) |
| 4153 | 3655 | PPTG_14795 | 28,02 | 2 | 0 | 0 | 2 | | | 1,58E+05 | 1 | 1 | 0 | 0 | 1 | Carbamidomethylation | 53872 | PPTG_14795 | Phytophthora parasitica INRA-310 cysteine protease family C01A (509 aa) |
| 3810 | 4577 | PPTG_17480 | 28 | 6 | 0 | 6 | 6 | | 8,10E+04 | 2,70E+05 | 1 | 1 | 0 | 1 | 1 | | 21085 | PPTG_17480 | Phytophthora parasitica INRA-310 hypothetical protein (195 aa) |
| 3356 | 3781 | PPTG_08293 | 28 | 1 | 0 | 1 | 1 | | 1,06E+05 | 1,57E+05 | 1 | 1 | 0 | 1 | 1 | | 73590 | PPTG_08293 | Phytophthora parasitica INRA-310 hypothetical protein (668 aa) |
| 4387 | 3835 | PPTG_00747 | 27,99 | 1 | 0 | 1 | 0 | | 9,62E+04 | | 1 | 1 | 0 | 1 | 0 | | 118349 | PPTG_00747 | Phytophthora parasitica INRA-310 nardilysin (1048 aa) |
| 3239 | 4458 | PPTG_06873 | 27,99 | 3 | 3 | 3 | 3 | 2,93E+05 | 3,61E+05 | 1,37E+06 | 1 | 1 | 1 | 1 | 1 | | 32864 | PPTG_06873 | Phytophthora parasitica INRA-310 hypothetical protein (289 aa) |
| 3372 | 3136 | PPTG_08621 | 27,98 | 2 | 2 | 0 | 2 | 5,83E+04 | | 3,37E+05 | 1 | 1 | 1 | 0 | 1 | | 46485 | PPTG_08621 | Phytophthora parasitica INRA-310 hypothetical protein (416 aa) |
| 4304 | 4584 | PPTG_15921 | 27,93 | 1 | 0 | 1 | 0 | | 0 | | 1 | 1 | 0 | 1 | 0 | | 138699 | PPTG_15921 | Phytophthora parasitica INRA-310 hypothetical protein (1223 aa) |
| 4463 | 4583 | PPTG_04937 | 27,92 | 3 | 0 | 0 | 3 | | | 3,13E+05 | 1 | 1 | 0 | 0 | 1 | | 34411 | PPTG_04937 | Phytophthora parasitica INRA-310 hypothetical protein (311 aa) |
| 3585 | 3943 | PPTG_05183 | 27,91 | 2 | 0 | 2 | 2 | | 2,24E+05 | 3,80E+05 | 1 | 1 | 0 | 1 | 1 | | 45482 | PPTG_05183 | Phytophthora parasitica INRA-310 hypothetical protein (404 aa) |
| 4282 | 4222 | PPTG_17146 | 27,91 | 5 | 5 | 0 | 0 | 9,77E+04 | | | 1 | 1 | 1 | 0 | 0 | Carbamidomethylation | 25277 | PPTG_17146 | Phytophthora parasitica INRA-310 ER lumen protein retaining receptor (218 aa) |
| 4464 | 4585 | PPTG_11873 | 27,89 | 7 | 0 | 0 | 7 | | | 1,25E+06 | 1 | 1 | 0 | 0 | 1 | | 15754 | PPTG_11873 | Phytophthora parasitica INRA-310 hypothetical protein (142 aa) |
| 3506 | 3689 | PPTG_06562 | 27,88 | 3 | 3 | 3 | 0 | 3,02E+05 | 5,85E+05 | | 1 | 1 | 1 | 1 | 0 | | 27030 | PPTG_06562 | Phytophthora parasitica INRA-310 splicing factor arginine/serine-rich (236 aa) |
| 4451 | 4496 | PPTG_16074 | 27,87 | 2 | 0 | 0 | 2 | | | 3,47E+05 | 1 | 1 | 0 | 0 | 1 | | 49910 | PPTG_16074 | Phytophthora parasitica INRA-310 obg family GTPase CgtA (454 aa) |
| 4658 | 4671 | PPTG_00211 | 27,83 | 3 | 0 | 3 | 0 | | 1,32E+05 | | 1 | 1 | 0 | 1 | 0 | | 38871 | PPTG_00211 | Phytophthora parasitica INRA-310 hypothetical protein (355 aa) |
| 4650 | 4613 | PPTG_02131 | 27,75 | 8 | 0 | 0 | 8 | | | 2,50E+05 | 1 | 1 | 0 | 0 | 1 | | 16919 | PPTG_02131 | Phytophthora parasitica INRA-310 hypothetical protein (160 aa) |
| 4659 | 4675 | PPTG_07666 | 27,73 | 4 | 0 | 4 | 0 | | 1,38E+05 | | 1 | 1 | 0 | 1 | 0 | | 42024 | PPTG_07666 | Phytophthora parasitica INRA-310 glycosyl hydrolase family 10 (384 aa) |
| 3733 | 4431 | PPTG_01815 | 27,72 | 1 | 0 | 1 | 1 | | 1,59E+05 | 2,80E+05 | 1 | 1 | 0 | 1 | 1 | | 61785 | PPTG_01815 | Phytophthora parasitica INRA-310 hypothetical protein (559 aa) |
| 3812 | 4618 | PPTG_16582 | 27,69 | 2 | 0 | 2 | 2 | | 1,25E+05 | 2,26E+05 | 1 | 1 | 0 | 1 | 1 | | 38609 | PPTG_16582 | Phytophthora parasitica INRA-310 hypothetical protein (342 aa) |
| 4276 | 4157 | PPTG_05415 | 27,68 | 1 | 0 | 0 | 1 | | | 2,57E+05 | 1 | 1 | 0 | 0 | 1 | | 107316 | PPTG_05415 | Phytophthora parasitica INRA-310 animal Inward Rectifier K Channel (IRK-C) family (952 aa) |
| 4456 | 4516 | PPTG_08249 | 27,67 | 2 | 0 | 0 | 2 | | | 3,79E+05 | 1 | 1 | 0 | 0 | 1 | | 76774 | PPTG_08249 | Phytophthora parasitica INRA-310 hypothetical protein (709 aa) |
| 4279 | 4181 | PPTG_13373 | 27,61 | 3 | 0 | 0 | 3 | | | 1,57E+05 | 1 | 1 | 0 | 0 | 1 | Carbamidomethylation | 59884 | PPTG_13373 | Phytophthora parasitica INRA-310 hypothetical protein (559 aa) |
| 4178 | 4068 | PPTG_04156 | 27,59 | 2 | 0 | 0 | 2 | | | 8,59E+05 | 1 | 1 | 0 | 0 | 1 | | 51373 | PPTG_04156 | Phytophthora parasitica INRA-310 hypothetical protein (457 aa) |
| 4178 | 4069 | PPTG_04164 | 27,59 | 2 | 0 | 0 | 2 | | | 8,59E+05 | 1 | 1 | 0 | 0 | 1 | | 53638 | PPTG_04164 | Phytophthora parasitica INRA-310 hypothetical protein (474 aa) |
| 4178 | 4512 | PPTG_04163 | 27,59 | 2 | 0 | 0 | 2 | | | 8,59E+05 | 1 | 1 | 0 | 0 | 1 | | 50169 | PPTG_04163 | Phytophthora parasitica INRA-310 hypothetical protein (447 aa) |
| 3508 | 3736 | PPTG_06243 | 27,55 | 3 | 0 | 0 | 3 | | | 3,19E+05 | 1 | 1 | 0 | 0 | 2 | | 56362 | PPTG_06243 | Phytophthora parasitica INRA-310 protein kinase (519 aa) |
| 4651 | 4617 | PPTG_04821 | 27,54 | 7 | 0 | 0 | 7 | | | 1,87E+05 | 1 | 1 | 0 | 0 | 1 | | 23678 | PPTG_04821 | Phytophthora parasitica INRA-310 hypothetical protein (228 aa) |
| 4156 | 3729 | PPTG_03357 | 27,54 | 1 | 0 | 0 | 1 | | | 1,97E+05 | 1 | 1 | 0 | 0 | 1 | | 201464 | PPTG_03357 | Phytophthora parasitica INRA-310 nucleoside diphosphate kinase (1910 aa) |
| 4300 | 4513 | PPTG_14383 | 27,53 | 3 | 0 | 0 | 3 | | | 3,01E+05 | 1 | 1 | 0 | 0 | 1 | | 35490 | PPTG_14383 | Phytophthora parasitica INRA-310 dimethyladenosine transferase (313 aa) |
| 4309 | 4627 | PPTG_19030 | 27,48 | 3 | 0 | 3 | 0 | | 1,44E+05 | | 1 | 1 | 0 | 1 | 0 | | 55069 | PPTG_19030 | Phytophthora parasitica INRA-310 hypothetical protein (480 aa) |
| 4313 | 4689 | PPTG_13387 | 27,45 | 8 | 0 | 8 | 0 | | 1,66E+06 | | 1 | 1 | 0 | 1 | 0 | | 13524 | PPTG_13387 | Phytophthora parasitica INRA-310 hypothetical protein (123 aa) |
| 4180 | 4080 | PPTG_06836 | 27,43 | 2 | 0 | 0 | 2 | | | 0 | 1 | 1 | 0 | 0 | 1 | Carbamidomethylation | 75675 | PPTG_06836 | Phytophthora parasitica INRA-310 sulfate Permease (SulP) family (697 aa) |
| 3530 | 4126 | PPTG_17155 | 27,42 | 2 | 0 | 2 | 2 | | 1,42E+05 | 1,81E+05 | 1 | 1 | 0 | 1 | 1 | | 73007 | PPTG_17155 | Phytophthora parasitica INRA-310 hypothetical protein (642 aa) |
| 3517 | 3857 | PPTG_02513 | 27,4 | 2 | 0 | 2 | 2 | | 0 | 2,55E+05 | 1 | 1 | 0 | 1 | 1 | | 52992 | PPTG_02513 | Phytophthora parasitica INRA-310 dual 3' 5'-cyclic-AMP and-GMP phosphodiesterase (483 aa) |
| 4440 | 4435 | PPTG_09941 | 27,39 | 4 | 0 | 0 | 4 | | | 0 | 1 | 1 | 0 | 0 | 1 | | 24285 | PPTG_09941 | Phytophthora parasitica INRA-310 hypothetical protein (220 aa) |
| 3358 | 3800 | PPTG_02507 | 27,37 | 2 | 0 | 2 | 2 | | 4,86E+04 | 4,14E+04 | 1 | 1 | 0 | 1 | 1 | | 93757 | PPTG_02507 | Phytophthora parasitica INRA-310 hypothetical protein (859 aa) |
| 4307 | 4622 | PPTG_02393 | 27,33 | 7 | 0 | 0 | 7 | | | 8,98E+05 | 1 | 1 | 0 | 0 | 1 | | 16566 | PPTG_02393 | Phytophthora parasitica INRA-310 hypothetical protein (154 aa) |
| 4100 | 3957 | PPTG_04363 | 27,3 | 4 | 0 | 4 | 0 | | 2,66E+05 | | 1 | 1 | 0 | 1 | 0 | | 34141 | PPTG_04363 | Phytophthora parasitica INRA-310 serine protease family S33 (305 aa) |
| 4467 | 4623 | PPTG_10393 | 27,27 | 4 | 0 | 4 | 0 | | 1,55E+05 | | 1 | 1 | 0 | 1 | 0 | | 41524 | PPTG_10393 | Phytophthora parasitica INRA-310 hypothetical protein (376 aa) |
| 4157 | 3732 | PPTG_02818 | 27,21 | 1 | 0 | 0 | 1 | | | 2,28E+05 | 1 | 1 | 0 | 0 | 1 | | 212376 | PPTG_02818 | Phytophthora parasitica INRA-310 hypothetical protein (1919 aa) |
| 3813 | 4633 | PPTG_15946 | 27,19 | 3 | 0 | 0 | 3 | | | 7,81E+05 | 1 | 1 | 0 | 0 | 2 | | 42594 | PPTG_15946 | Phytophthora parasitica INRA-310 hypothetical protein (375 aa) |
| 3747 | 4593 | PPTG_02136 | 27,19 | 1 | 0 | 1 | 1 | | 1,30E+05 | 2,08E+05 | 1 | 1 | 0 | 1 | 1 | | 126486 | PPTG_02136 | Phytophthora parasitica INRA-310 hypothetical protein (1121 aa) |
| 3814 | 4645 | PPTG_02378 | 27,11 | 2 | 0 | 2 | 2 | | 1,82E+05 | 2,20E+05 | 1 | 1 | 0 | 1 | 1 | | 50742 | PPTG_02378 | Phytophthora parasitica INRA-310 hypothetical protein (458 aa) |
| 4393 | 3987 | PPTG_09803 | 27,1 | 2 | 0 | 0 | 2 | | | 1,79E+05 | 1 | 1 | 0 | 0 | 1 | | 58649 | PPTG_09803 | Phytophthora parasitica INRA-310 hypothetical protein (519 aa) |
| 4085 | 3723 | PPTG_02150 | 27,1 | 1 | 0 | 0 | 1 | | | 5,29E+05 | 1 | 1 | 0 | 0 | 1 | | 95178 | PPTG_02150 | Phytophthora parasitica INRA-310 ion channel (860 aa) |
| 4085 | 4640 | PPTG_02149 | 27,1 | 2 | 0 | 0 | 2 | | | 5,29E+05 | 1 | 1 | 0 | 0 | 1 | | 87718 | PPTG_02149 | Phytophthora parasitica INRA-310 ion channel (791 aa) |
| 3285 | 4641 | PPTG_12179 | 27,08 | 16 | 16 | 16 | 16 | 1,14E+06 | 1,70E+06 | 3,18E+06 | 1 | 1 | 1 | 1 | 1 | Carbamidomethylation | 6142 | PPTG_12179 | Phytophthora parasitica INRA-310 hypothetical protein (58 aa) |
| 4652 | 4646 | PPTG_00177 | 26,97 | 7 | 0 | 0 | 7 | | | 1,11E+05 | 1 | 1 | 0 | 0 | 1 | | 18413 | PPTG_00177 | Phytophthora parasitica INRA-310 hypothetical protein (168 aa) |
| 3996 | 3461 | PPTG_13379 | 26,96 | 1 | 0 | 1 | 0 | | 2,09E+05 | | 1 | 1 | 0 | 1 | 0 | | 134739 | PPTG_13379 | Phytophthora parasitica INRA-310 structural maintenance-chromosomes protein (1184 aa) |
| 4429 | 4359 | PPTG_00617 | 26,92 | 3 | 0 | 3 | 0 | | 1,51E+05 | | 1 | 1 | 0 | 1 | 0 | | 43079 | PPTG_00617 | Phytophthora parasitica INRA-310 transglutaminase elicitor-like protein (388 aa) |
| 4429 | 4360 | PPTG_00616 | 26,92 | 2 | 0 | 2 | 0 | | 1,51E+05 | | 1 | 1 | 0 | 1 | 0 | | 48246 | PPTG_00616 | Phytophthora parasitica INRA-310 hypothetical protein (439 aa) |
| 4653 | 4647 | PPTG_06731 | 26,92 | 4 | 0 | 0 | 4 | | | 2,80E+05 | 1 | 1 | 0 | 0 | 1 | | 52932 | PPTG_06731 | Phytophthora parasitica INRA-310 hypothetical protein (452 aa) |
| 4177 | 4061 | PPTG_08311 | 26,92 | 2 | 0 | 0 | 2 | | | 1,92E+05 | 1 | 1 | 0 | 0 | 1 | | 99897 | PPTG_08311 | Phytophthora parasitica INRA-310 hypothetical protein (892 aa) |
| 4465 | 4596 | PPTG_09547 | 26,85 | 3 | 0 | 0 | 3 | | | 6,14E+04 | 1 | 1 | 0 | 0 | 1 | | 43602 | PPTG_09547 | Phytophthora parasitica INRA-310 peroxisome assembly protein (392 aa) |
| 4654 | 4649 | PPTG_04574T2 | 26,83 | 3 | 0 | 0 | 3 | | | 1,35E+05 | 1 | 1 | 0 | 0 | 1 | | 52619 | PPTG_04574 | Phytophthora parasitica INRA-310 hypothetical protein (474 aa) |
| 4654 | 4650 | PPTG_04574 | 26,83 | 3 | 0 | 0 | 3 | | | 1,35E+05 | 1 | 1 | 0 | 0 | 1 | | 52747 | PPTG_04574 | Phytophthora parasitica INRA-310 hypothetical protein (475 aa) |
| 4654 | 4651 | PPTG_04574 | 26,83 | 2 | 0 | 0 | 2 | | | 1,35E+05 | 1 | 1 | 0 | 0 | 1 | | 54914 | PPTG_04574 | Phytophthora parasitica INRA-310 hypothetical protein (495 aa) |
| 3169 | 3809 | PPTG_18494 | 26,82 | 3 | 3 | 3 | 3 | 8,27E+04 | 3,98E+05 | 6,59E+05 | 1 | 1 | 1 | 1 | 1 | | 36320 | PPTG_18494 | Phytophthora parasitica INRA-310 hypothetical protein (321 aa) |
| 4655 | 4652 | PPTG_14648 | 26,78 | 2 | 0 | 0 | 2 | | | 1,39E+05 | 1 | 1 | 0 | 0 | 1 | | 65953 | PPTG_14648 | Phytophthora parasitica INRA-310 hypothetical protein (602 aa) |
| 3621 | 4660 | PPTG_01779 | 26,77 | 10 | 0 | 0 | 10 | | | 1,64E+06 | 1 | 1 | 0 | 0 | 2 | Oxidation (M) | 8657 | PPTG_01779 | Phytophthora parasitica INRA-310 hypothetical protein (74 aa) |
| 3815 | 4654 | PPTG_10936 | 26,77 | 10 | 0 | 10 | 10 | | 7,45E+04 | 4,42E+05 | 1 | 1 | 0 | 1 | 1 | | 9586 | PPTG_10936 | Phytophthora parasitica INRA-310 hypothetical protein (88 aa) |
| 4286 | 4304 | PPTG_03557 | 26,74 | 1 | 0 | 0 | 1 | | | 1,33E+05 | 1 | 1 | 0 | 0 | 1 | | 140861 | PPTG_03557 | Phytophthora parasitica INRA-310 transmembrane protein (1225 aa) |
| 3624 | 4739 | PPTG_19713 | 26,71 | 3 | 0 | 3 | 3 | | 5,42E+05 | 2,68E+06 | 1 | 1 | 0 | 1 | 1 | | 35199 | PPTG_19713 | Phytophthora parasitica INRA-310 ribosomal large subunit pseudouridine synthase F (316 aa) |
| 3523 | 3961 | PPTG_18124 | 26,69 | 4 | 4 | 0 | 4 | 8,71E+04 | | 0 | 1 | 1 | 1 | 0 | 1 | | 27839 | PPTG_18124 | Phytophthora parasitica INRA-310 hypothetical protein (252 aa) |
| 4096 | 3901 | PPTG_09957 | 26,69 | 3 | 0 | 0 | 3 | | | 1,11E+05 | 1 | 1 | 0 | 0 | 1 | | 67594 | PPTG_09957 | Phytophthora parasitica INRA-310 hypothetical protein (618 aa) |
| 3743 | 4530 | PPTG_16376 | 26,68 | 2 | 0 | 2 | 2 | | 3,67E+05 | 5,09E+05 | 1 | 1 | 0 | 1 | 1 | | 35783 | PPTG_16376 | Phytophthora parasitica INRA-310 hypothetical protein (325 aa) |
| 4059 | 3991 | PPTG_18231 | 26,67 | 2 | 0 | 2 | 0 | | 4,71E+04 | | 1 | 1 | 0 | 1 | 0 | | 73525 | PPTG_18231 | Phytophthora parasitica INRA-310 carbohydrate-binding protein (685 aa) |
| 4656 | 4656 | PPTG_14763 | 26,66 | 2 | 0 | 0 | 2 | | | 2,37E+05 | 1 | 1 | 0 | 0 | 1 | | 64716 | PPTG_14763 | Phytophthora parasitica INRA-310 transmembrane protein (571 aa) |
| 4669 | 4744 | PPTG_08592 | 26,63 | 2 | 0 | 2 | 0 | | 4,08E+05 | | 1 | 1 | 0 | 1 | 0 | | 62907 | PPTG_08592 | Phytophthora parasitica INRA-310 hypothetical protein (594 aa) |
| 4669 | 4745 | PPTG_08595 | 26,63 | 2 | 0 | 2 | 0 | | 4,08E+05 | | 1 | 1 | 0 | 1 | 0 | | 62812 | PPTG_08595 | Phytophthora parasitica INRA-310 hypothetical protein (594 aa) |
| 3582 | 3936 | PPTG_19778 | 26,61 | 7 | 0 | 7 | 7 | | 4,90E+05 | 0 | 1 | 1 | 0 | 1 | 1 | | 15969 | PPTG_19778 | Phytophthora parasitica INRA-310 hypothetical protein (141 aa) |
| 3582 | 3937 | PPTG_14455 | 26,61 | 7 | 0 | 7 | 7 | | 4,90E+05 | 0 | 1 | 1 | 0 | 1 | 1 | | 15969 | PPTG_14455 | Phytophthora parasitica INRA-310 hypothetical protein (141 aa) |
| 3089 | 4076 | PPTG_09914 | 26,55 | 2 | 2 | 2 | 2 | 1,54E+05 | 5,29E+05 | 9,08E+05 | 1 | 1 | 1 | 1 | 1 | | 55374 | PPTG_09914 | Phytophthora parasitica INRA-310 hypothetical protein (508 aa) |
| 4302 | 4541 | PPTG_13678 | 26,55 | 2 | 0 | 2 | 0 | | 1,64E+05 | | 1 | 1 | 0 | 1 | 0 | | 90615 | PPTG_13678 | Phytophthora parasitica INRA-310 tRNA (uracil-5-)-methyltransferase (815 aa) |
| 4191 | 4569 | PPTG_18947 | 26,5 | 3 | 0 | 0 | 3 | | | 0 | 1 | 1 | 0 | 0 | 1 | | 42624 | PPTG_18947 | Phytophthora parasitica INRA-310 ATP-binding Cassette (ABC) superfamily protein (377 aa) |
| 4181 | 4099 | PPTG_09954 | 26,47 | 5 | 0 | 0 | 5 | | | 2,71E+05 | 1 | 1 | 0 | 0 | 1 | | 20913 | PPTG_09954 | Phytophthora parasitica INRA-310 vacuolar protein sorting-associated protein 29 (190 aa) |
| 4475 | 4698 | PPTG_02057 | 26,46 | 1 | 0 | 1 | 0 | | 3,67E+04 | | 1 | 1 | 0 | 1 | 0 | | 297393 | PPTG_02057 | Phytophthora parasitica INRA-310 hypothetical protein (2678 aa) |
| 4469 | 4665 | PPTG_01618 | 26,44 | 7 | 0 | 0 | 7 | | | 1,33E+05 | 1 | 1 | 0 | 0 | 1 | | 17586 | PPTG_01618 | Phytophthora parasitica INRA-310 D-tyrosyl-tRNA(Tyr) deacylase (156 aa) |
| 4083 | 3702 | PPTG_15994 | 26,43 | 1 | 0 | 0 | 1 | | | 7,53E+05 | 1 | 1 | 0 | 0 | 1 | | 134207 | PPTG_15994 | Phytophthora parasitica INRA-310 ribosome biogenesis protein BMS1 (1179 aa) |
| 3816 | 4667 | PPTG_06855 | 26,41 | 6 | 0 | 6 | 6 | | 5,05E+05 | 6,13E+05 | 1 | 1 | 0 | 1 | 1 | | 19108 | PPTG_06855 | Phytophthora parasitica INRA-310 hypothetical protein (180 aa) |
| 4470 | 4668 | PPTG_17919 | 26,38 | 4 | 0 | 0 | 4 | | | 7,61E+05 | 1 | 1 | 0 | 0 | 1 | | 43828 | PPTG_17919 | Phytophthora parasitica INRA-310 UDP-N-acetylglucosamine-dolichyl-phosphate N-acetylglucosaminephosphotransferase (396 aa) |
| 3751 | 4686 | PPTG_11856 | 26,32 | 12 | 0 | 0 | 12 | | | 1,10E+06 | 1 | 1 | 0 | 0 | 2 | Oxidation (M) | 6861 | PPTG_11856 | Phytophthora parasitica INRA-310 hypothetical protein (58 aa) |
| 4472 | 4679 | PPTG_12182 | 26,32 | 4 | 0 | 0 | 4 | | | 3,57E+05 | 1 | 1 | 0 | 0 | 1 | | 31548 | PPTG_12182 | Phytophthora parasitica INRA-310 hypothetical protein (277 aa) |
| 4283 | 4256 | PPTG_05063 | 26,3 | 3 | 0 | 3 | 0 | | 4,57E+05 | | 1 | 1 | 0 | 1 | 0 | Carbamidomethylation | 53913 | PPTG_05063 | Phytophthora parasitica INRA-310 hypothetical protein (484 aa) |
| 4660 | 4681 | PPTG_13312 | 26,3 | 3 | 0 | 0 | 3 | | | 6,05E+05 | 1 | 1 | 0 | 0 | 1 | | 49229 | PPTG_13312 | Phytophthora parasitica INRA-310 major facilitator superfamily transporter (MFS) (444 aa) |
| 4018 | 3013 | PPTG_08035 | 26,29 | 1 | 0 | 0 | 1 | | | 8,24E+05 | 1 | 1 | 0 | 0 | 1 | | 147946 | PPTG_08035 | Phytophthora parasitica INRA-310 N-acetylglucosamine-1-phosphotransferase alpha/beta (1311 aa) |
| 4661 | 4684 | PPTG_14401 | 26,24 | 2 | 0 | 0 | 2 | | | 0 | 1 | 1 | 0 | 0 | 1 | | 73243 | PPTG_14401 | Phytophthora parasitica INRA-310 hypothetical protein (661 aa) |
| 4308 | 4625 | PPTG_03549 | 26,22 | 1 | 0 | 0 | 1 | | | 3,75E+05 | 1 | 1 | 0 | 0 | 1 | | 84268 | PPTG_03549 | Phytophthora parasitica INRA-310 hypothetical protein (759 aa) |
