## Supplementary table 2 for "Zoospore-derived extracellular vesicles in the flagellated stramenopile *Phytophthora parasitica*"

**Title : Annotations of identified and selected EV protein candidates in terms of being part of a family, putative EV markers, and gene ontology**

| Gene Id | Uniprot Entry | Protein names | Length | candidates related to EV markers | Protein families | Gene Ontology (biological process) | Gene Ontology (cellular component) | Gene Ontology (molecular function) | Gene Ontology (GO) | Proteomes |
|---|---|---|---|---|---|---|---|---|---|---|
| PPTG_04587 | W2R3C1 | Titin | 10565 | | | | | | | UP000018817 |
| PPTG_06389 | W2QVA6 | Uncharacterized protein | 878 | | | | | | | UP000018817 |
| PPTG_01092 | W2RJZ0 | Fibronectin type-III domain-containing protein | 5778 | Ral et al., 2024 | | | | | | UP000018817 |
| PPTG_11936 | W2Q8W4 | Clathrin heavy chain | 1719 | | Clathrin heavy chain family | intracellular protein transport [GO:0006886 | clathrin coat of coated pit [GO:0030132]; cla | clathrin light chain binding [GO:0032051]; st | clathrin coat of coated pit | UP000018817 |
| PPTG_16264 | W2PQY5 | Spondin-like TSP1 domain-containing protein | 2399 | | | | | | | UP000018817 |
| PPTG_17090 | W2PLY2 | Uncharacterized protein | 1138 | | UDPGP type 1 family; Phosphohexose mutase fam | carbohydrate metabolic process [GO:0005 | cytosol [GO:0005829] | magnesium ion binding [GO:0000287]; phos | cytosol [GO:0005829]; ma | UP000018817 |
| PPTG_01091 | W2RJX2 | Fibronectin type-III domain-containing protein | 5915 | Ral et al., 2024 | | | | | | UP000018817 |
| PPTG_02441 | W2RAR2 | EGF-like domain-containing protein | 597 | Welsh et al., 2024 | | | | | | UP000018817 |
| PPTG_09633 | W2QFM9 | Cation-transporting P-type ATPase N-terminal domain-containing protein | 1343 | | Cation transport ATPase (P-type) (TC 3.A.3) family | intracellular potassium ion homeostasis [G | plasma membrane [GO:0005886] | ATP binding [GO:0005524]; ATP hydrolysis ac | plasma membrane [GO:00 | UP000018817 |
| PPTG_02434 | W2RCV6 | Translation elongation factor aEF-2 | 858 | | | | cytosol [GO:0005829]; ribonucleoprotein con | GTP binding [GO:0005525]; GTPase activity [ | cytosol [GO:0005829]; ribo | UP000018817 |
| PPTG_16242 | W2PNS1 | Uncharacterized protein | 854 | Vinay and Belleannée, 2022 | | | membrane [GO:0016020] | | membrane [GO:0016020] | UP000018817 |
| PPTG_13841 | W2PYH8 | ATP synthase subunit beta (EC 7.1.2.2) | 501 | | ATPase alpha/beta chains family | proton motive force-driven mitochondrial A | mitochondrial inner membrane [GO:000574 | ATP binding [GO:0005524]; proton-transporti | mitochondrial inner memb | UP000018817 |
| PPTG_04733 | W2R3Z5 | Uncharacterized protein | 5308 | | | | membrane [GO:0016020] | calcium ion binding [GO:0005509] | membrane [GO:0016020]; | UP000018817 |
| PPTG_03648 | W2R5N9 | EGF-like domain-containing protein | 393 | Welsh et al., 2024 | | | | | | UP000018817 |
| PPTG_06394 | W2QUN2 | Uncharacterized protein | 1125 | | | | | | | UP000018817 |
| PPTG_05265 | W2QWF0 | Dynein heavy chain, cytoplasmic (Dynein heavy chain, cytosolic) | 4706 | | Dynein heavy chain family | cilium assembly [GO:0060271]; intracellula | axonemal dynein complex [GO:0005858]; cili | ATP binding [GO:0005524]; dynein intermedi | axonemal dynein complex | UP000018817 |
| PPTG_06382 | W2QSV6 | Uncharacterized protein | 1089 | | | | | | | UP000018817 |
| PPTG_02260 | W2RA63 | leucine--tRNA ligase (EC 6.1.1.4) (Leucyl-tRNA synthetase) | 1108 | | Class-I aminoacyl-tRNA synthetase family | leucyl-tRNA aminoacylation [GO:0006429] | | aminoacyl-tRNA deacylase activity [GO:000 | aminoacyl-tRNA deacylas | UP000018817 |
| PPTG_08772 | W2QII2 | Carbamoyl-phosphate synthase, large subunit | 1499 | | | 'de novo' pyrimidine nucleobase biosynthet | carbamoyl-phosphate synthase complex [GO | ATP binding [GO:0005524]; carbamoyl-phos | carbamoyl-phosphate synt | UP000018817 |
| PPTG_14782 | W2PWT7 | B30.2/SPRY domain-containing protein | 5307 | | | | | ubiquitin-protein transferase activity [GO:00 | ubiquitin-protein transfera | UP000018817 |
| PPTG_14544 | W2PXK9 | 26S proteasome non-ATPase regulatory subunit 2 | 897 | | Proteasome subunit S2 family | proteasome-mediated ubiquitin-dependent | nucleus [GO:0005634]; proteasome regulato | enzyme regulator activity [GO:0030234] | nucleus [GO:0005634]; pro | UP000018817 |
| PPTG_20351 | W2P921 | Ricin B lectin domain-containing protein | 466 | | | protein O-linked glycosylation [GO:0006493] | | carbohydrate binding [GO:0030246]; polype | carbohydrate binding [GO: | UP000018817 |
| PPTG_06857 | W2QSZ3 | C2 domain-containing protein | 1764 | | | plasma membrane organization [GO:00070 | membrane [GO:0016020] | | membrane [GO:0016020]; | UP000018817 |
| PPTG_02122 | W2R9I2 | Hsp70-like protein | 657 | Welsh et al., 2024 | Heat shock protein 70 family | | | ATP binding [GO:0005524]; ATP-dependent p | ATP binding [GO:0005524] | UP000018817 |
| PPTG_05028 | W2QVJ2 | Acyl-coenzyme A dehydrogenase (EC 1.3.8.7) (EC 1.3.8.8) | 764 | | Acyl-CoA dehydrogenase family | fatty acid beta-oxidation using acyl-CoA deh | mitochondrion [GO:0005739] | flavin adenine dinucleotide binding [GO:005 | mitochondrion [GO:0005 | UP000018817 |
| PPTG_14525 | W2PVS0 | Carbamoyl phosphate synthase arginine-specific large chain (EC 6.3.4.16) (EC 6.3.5.5) | 1511 | | CarB family | 'de novo' pyrimidine nucleobase biosynthet | carbamoyl-phosphate synthase complex [GO | ATP binding [GO:0005524]; carbamoyl-phos | carbamoyl-phosphate synt | UP000018817 |
| PPTG_06395 | W2QVB8 | EGF-like domain-containing protein | 1089 | Welsh et al., 2024 | | | | | | UP000018817 |
| PPTG_12938 | W2Q3X5 | PH domain-containing protein | 1016 | | | | membrane [GO:0016020] | glycosyltransferase activity [GO:0016757] | membrane [GO:0016020]; | UP000018817 |
| PPTG_12937 | W2Q357 | PH domain-containing protein | 1026 | | | | membrane [GO:0016020] | glycosyltransferase activity [GO:0016757] | membrane [GO:0016020]; | UP000018817 |
| PPTG_02292 | W2RAA2 | HP domain-containing protein | 881 | | Villin/gelsolin family | actin filament capping [GO:0051693]; cytoskeleton organization [GO:0007010] | | actin filament binding [GO:0051015] | actin filament binding [GO | UP000018817 |
| PPTG_03714 | W2R7Y3 | Phosphoglycerate kinase (EC 2.7.2.3) | 414 | | Phosphoglycerate kinase family | gluconeogenesis [GO:0006094]; glycolytic p | cytosol [GO:0005829] | ADP binding [GO:0043531]; ATP binding [GO | cytosol [GO:0005829]; ADI | UP000018817 |
| PPTG_02565 | W2RBF0 | Serine protease (EC 3.4.21.-) | 808 | | Peptidase S1B family | proteolysis [GO:0006508] | | serine-type peptidase activity [GO:0008236] | serine-type peptidase acti | UP000018817 |
| PPTG_12232 | W2Q852 | ABC transporter domain-containing protein | 1948 | | ABC transporter superfamily, ABCA family | | membrane [GO:0016020] | ABC-type transporter activity [GO:0140359]; | membrane [GO:0016020]; | UP000018817 |
| PPTG_01206 | W2RKB5 | Pyruvate carboxylase (EC 6.4.1.1) | 1186 | | | gluconeogenesis [GO:0006094] | cytoplasm [GO:0005737] | ATP binding [GO:0005524]; metal ion binding | cytoplasm [GO:0005737]; | UP000018817 |
| PPTG_19375 | W2PCW7 | Heat shock protein 70 | 809 | Welsh et al., 2024 | | | cytosol [GO:0005829]; nucleus [GO:0005634 | ATP binding [GO:0005524]; ATP-dependent p | cytosol [GO:0005829]; nuc | UP000018817 |
| PPTG_20013 | W2Q2P7 | phosphopyruvate hydratase (EC 4.2.1.11) | 483 | | Enolase family | glycolytic process [GO:0006096] | phosphopyruvate hydratase complex [GO:00 | magnesium ion binding [GO:0000287]; phos | phosphopyruvate hydratas | UP000018817 |
| PPTG_06730 | W2QR91 | EGF-like domain-containing protein | 676 | Welsh et al., 2024 | | | | | | UP000018817 |
| PPTG_08579 | W2QN32 | 1,3-beta-glucan synthase (EC 2.4.1.34) | 2228 | Zhao et al., 2019 | Glycosyltransferase 48 family | (1->3)-beta-D-glucan biosynthetic process [ | 1,3-beta-D-glucan synthase complex [GO:00 | 1,3-beta-D-glucan synthase activity [GO:000 | 1,3-beta-D-glucan synthas | UP000018817 |
| PPTG_07612 | W2QN96 | AP-2 complex subunit alpha | 987 | | Adaptor complexes large subunit family | clathrin-dependent endocytosis [GO:00725 | AP-2 adaptor complex [GO:0030122] | clathrin adaptor activity [GO:0035615] | AP-2 adaptor complex [GO | UP000018817 |
| PPTG_10603 | W2QD70 | Creatine kinase, flagellar | 785 | | ATP:guanido phosphotransferase family | phosphocreatine biosynthetic process [GO: | extracellular space [GO:0005615] | ATP binding [GO:0005524]; creatine kinase a | extracellular space [GO:00 | UP000018817 |
| PPTG_07345 | W2QPP7 | Calcium-transporting ATPase (EC 7.2.2.10) | 1048 | | Cation transport ATPase (P-type) (TC 3.A.3) family | | endomembrane system [GO:0012505]; plasr | ATP binding [GO:0005524]; ATP hydrolysis ac | endomembrane system [G | UP000018817 |
| PPTG_03836 | W2QY41 | glucan endo-1,3-beta-D-glucosidase (EC 3.2.1.39) | 556 | | PGA52 family | cell wall organization [GO:0071555] | | glucan endo-1,3-beta-D-glucosidase activity | glucan endo-1,3-beta-D-gl | UP000018817 |
| PPTG_01093 | W2RI82 | Fibronectin type-III domain-containing protein | 1390 | Ral et al., 2024 | | | | | | UP000018817 |
| PPTG_03121 | W2R438 | Chaperone DnaK | 786 | Welsh et al., 2024 | Heat shock protein 70 family | | | ATP binding [GO:0005524]; ATP-dependent p | ATP binding [GO:0005524] | UP000018817 |
| PPTG_12681 | W2Q2U4 | ABC transporter B family member 11 | 1290 | | ABC transporter superfamily, ABCB family, Multidr | oligopeptide export from mitochondrion [GO | mitochondrial inner membrane [GO:000574 | ABC-type oligopeptide transporter activity [G | mitochondrial inner memb | UP000018817 |
| PPTG_04750 | W2R256 | Peptidase M16 N-terminal domain-containing protein | 447 | | Peptidase M16 family | | membrane [GO:0016020]; mitochondrion [G | metal ion binding [GO:0046872] | membrane [GO:0016020]; | UP000018817 |
| PPTG_19658 | W2PDR0 | Argininosuccinate synthase (EC 6.3.4.5) (Citrulline--aspartate ligase) | 421 | | | argininosuccinate metabolic process [GO:0 | cytoplasm [GO:0005737] | argininosuccinate synthase activity [GO:000 | cytoplasm [GO:0005737]; | UP000018817 |
| PPTG_01676 | W2R8E6 | Guanine nucleotide-binding protein subunit beta-like protein | 317 | | WD repeat G protein beta family, Ribosomal protein RACK1 subfamily | | | ribosome binding [GO:0043022]; translation | ribosome binding [GO:004 | UP000018817 |
| PPTG_16803 | W2PNG4 | H(+)-transporting two-sector ATPase (EC 7.1.2.2) | 618 | | ATPase alpha/beta chains family | ATP metabolic process [GO:0046034] | proton-transporting V-type ATPase, V1 domai | ATP binding [GO:0005524]; ATP hydrolysis ac | proton-transporting V-type | UP000018817 |
| PPTG_13258 | W2Q0U2 | 1,3-beta-glucan synthase (EC 2.4.1.34) | 2286 | Zhao et al., 2019 | Glycosyltransferase 48 family | (1->3)-beta-D-glucan biosynthetic process [ | 1,3-beta-D-glucan synthase complex [GO:00 | 1,3-beta-D-glucan synthase activity [GO:000 | 1,3-beta-D-glucan synthas | UP000018817 |
| PPTG_09208 | W2QHN0 | Glycoside hydrolase family 5 domain-containing protein | 505 | | Glycosyl hydrolase 72 family | cell wall (1->3)-beta-D-glucan biosynthetic | plasma membrane [GO:0005886] | 1,3-beta-glucanosyltransferase activity [GO: | plasma membrane [GO:00 | UP000018817 |
| PPTG_04703 | W2R288 | ADP/ATP translocase (ADP,ATP carrier protein) | 310 | | Mitochondrial carrier (TC 2.A.29) family | mitochondrial ADP transmembrane transpo | mitochondrial inner membrane [GO:000574 | ATP:ADP antiporter activity [GO:0005471] | mitochondrial inner memb | UP000018817 |
| PPTG_07368 | W2QPT4 | Trifunctional enzyme subunit alpha, mitochondrial (EC 1.1.1.211) (EC 4.2.1.17) (Monolysocardiolip | 747 | | Enoyl-CoA hydratase/isomerase family; Enoyl-CoA | fatty acid beta-oxidation [GO:0006635] | mitochondrial fatty acid beta-oxidation multi | enoyl-CoA hydratase activity [GO:0004300]; | mitochondrial fatty acid be | UP000018817 |
| PPTG_19188 | W2PE41 | Heat shock protein 90-2 | 706 | Welsh et al., 2024 | Heat shock protein 90 family | | cytoplasm [GO:0005737] | ATP binding [GO:0005524]; ATP hydrolysis ac | cytoplasm [GO:0005737]; | UP000018817 |
| PPTG_05941 | W2QUI9 | Cation-transporting P-type ATPase N-terminal domain-containing protein | 1076 | | Cation transport ATPase (P-type) (TC 3.A.3) family | intracellular potassium ion homeostasis [G | plasma membrane [GO:0005886] | ATP binding [GO:0005524]; ATP hydrolysis ac | plasma membrane [GO:00 | UP000018817 |
| PPTG_14787 | W2PVX1 | EGF-like domain-containing protein | 627 | Welsh et al., 2024 | Glycosyl hydrolase 72 family | cell wall (1->3)-beta-D-glucan biosynthetic | plasma membrane [GO:0005886] | 1,3-beta-glucanosyltransferase activity [GO: | plasma membrane [GO:00 | UP000018817 |
| PPTG_01216 | W2R636 | alpha-glucosidase (EC 3.2.1.20) (Maltase) | 1142 | | Glycosyl hydrolase 31 family | carbohydrate metabolic process [GO:0005975] | | alpha-glucosidase activity [GO:0090599]; sta | alpha-glucosidase activity | UP000018817 |
| PPTG_12239 | W2Q5I7 | ubiquitinyl hydrolase 1 (EC 3.4.19.12) | 4322 | | | protein deubiquitination involved in ubiquiti | cytoplasm [GO:0005737]; nucleus [GO:0005 | calcium ion binding [GO:0005509]; cysteine- | cytoplasm [GO:0005737]; | UP000018817 |
| PPTG_17884 | W2PI34 | Annexin | 328 | Vinay and Belleannée, 2022 | Annexin family | | cytoplasm [GO:0005737]; plasma membrane | calcium ion binding [GO:0005509]; calcium- | cytoplasm [GO:0005737]; | UP000018817 |
| PPTG_17883 | W2PII0 | Annexin | 328 | Vinay and Belleannée, 2022 | Annexin family | | cytoplasm [GO:0005737]; plasma membrane | calcium ion binding [GO:0005509]; calcium- | cytoplasm [GO:0005737]; | UP000018817 |
| PPTG_16401 | W2PNJ7 | Eukaryotic translation initiation factor 3 subunit A (eIF3a) (Eukaryotic translation initiation factor 3 | 1146 | | EIF-3 subunit A family | formation of cytoplasmic translation initiati | eukaryotic 43S preinitiation complex [GO:00 | mRNA binding [GO:0003729]; translation init | eukaryotic 43S preinitiatio | UP000018817 |
| PPTG_02311 | W2RCG4 | Creatine kinase | 435 | | ATP:guanido phosphotransferase family | phosphocreatine biosynthetic process [GO: | extracellular space [GO:0005615] | ATP binding [GO:0005524]; creatine kinase a | extracellular space [GO:00 | UP000018817 |
| PPTG_10528 | W2QC09 | Malate dehydrogenase (EC 1.1.1.37) | 335 | | LDH/MDH superfamily, MDH type 1 family | malate metabolic process [GO:0006108]; tr | cytoplasm [GO:0005737] | L-malate dehydrogenase (NAD+) activity [GO | cytoplasm [GO:0005737]; | UP000018817 |
| PPTG_15348 | W2PVD2 | Actin-1 | 376 | | Actin family | | | ATP binding [GO:0005524]; hydrolase activity | ATP binding [GO:0005524] | UP000018817 |
| PPTG_06442 | W2QVJ6 | AMP-dependent synthetase/ligase domain-containing protein | 646 | | | | endoplasmic reticulum [GO:0005783]; mem | long-chain fatty acid-CoA ligase activity [GO: | endoplasmic reticulum [G | UP000018817 |
| PPTG_13419 | W2Q329 | Aconitate hydratase, mitochondrial (Aconitase) (EC 4.2.1.3) | 786 | | Aconitase/IPM isomerase family | tricarboxylic acid cycle [GO:0006099] | cytosol [GO:0005829]; mitochondrion [GO:0 | 4 iron, 4 sulfur cluster binding [GO:0051539] | cytosol [GO:0005829]; mit | UP000018817 |
| PPTG_08245 | W2QLK3 | Luminal-binding protein | 658 | | Heat shock protein 70 family | | | ATP binding [GO:0005524]; ATP-dependent p | ATP binding [GO:0005524] | UP000018817 |
| PPTG_18415 | W2PI14 | Coatomer subunit beta (Beta-coat protein) | 993 | | | endoplasmic reticulum to Golgi vesicle-me | COPI vesicle coat [GO:0030126]; Golgi mem | structural molecule activity [GO:0005198] | COPI vesicle coat [GO:003 | UP000018817 |
| PPTG_12072 | W2Q5T5 | B30.2/SPRY domain-containing protein | 4810 | | | protein ubiquitination [GO:0016567] | | metal ion binding [GO:0046872]; ubiquitin-p | metal ion binding [GO:004 | UP000018817 |
| PPTG_11762 | W2QAQ8 | 6-phosphogluconate dehydrogenase, decarboxylating (EC 1.1.1.44) | 489 | | 6-phosphogluconate dehydrogenase family | D-gluconate metabolic process [GO:0019521]; pentose-phosphate shunt [GO:0006098] | | NADP binding [GO:0050661]; phosphogluco | NADP binding [GO:005066 | UP000018817 |
| PPTG_08188 | W2QLD0 | Phosphoribosylaminoimidazolecarboxamide formyltransferase/IMP cyclohydrolase | 616 | | PurH family | 'de novo' IMP biosynthetic process [GO:000 | cytosol [GO:0005829] | IMP cyclohydrolase activity [GO:0003937]; p | cytosol [GO:0005829]; IMF | UP000018817 |
| PPTG_13558 | W2Q2U0 | Elongation factor 3 (Eukaryotic elongation factor 3) | 1038 | | ABC transporter superfamily, ABCF family, EF3 subfamily | | cytoplasm [GO:0005737] | ATP binding [GO:0005524]; ATP hydrolysis ac | cytoplasm [GO:0005737]; | UP000018817 |
| PPTG_04382 | W2R0H5 | Major vault protein | 881 | | | | cytoplasm [GO:0005737]; nucleus [GO:0005634]; ribonucleoprotein complex [GO:199090 | | cytoplasm [GO:0005737]; | UP000018817 |
| PPTG_05087 | W2QVI9 | Kinesin motor domain-containing protein | 1617 | | TRAFAC class myosin-kinesin ATPase superfamily, | transport along microtubule [GO:0010970] | | ATP binding [GO:0005524]; microtubule bind | ATP binding [GO:0005524] | UP000018817 |
| PPTG_01098 | W2RHS3 | Glyceraldehyde-3-phosphate dehydrogenase (EC 1.2.1.12) | 370 | Welsh et al., 2024 | Glyceraldehyde-3-phosphate dehydrogenase famil | glucose metabolic process [GO:0006006]; g | cytosol [GO:0005829] | glyceraldehyde-3-phosphate dehydrogenase | cytosol [GO:0005829]; glyc | UP000018817 |
| PPTG_09519 | W2QG53 | ATP-citrate synthase (EC 2.3.3.8) (ATP-citrate (pro-S-)-lyase) (Citrate cleavage enzyme) | 1105 | | Succinate/malate CoA ligase alpha subunit family; | acetyl-CoA biosynthetic process [GO:00060 | cytosol [GO:0005829] | ATP binding [GO:0005524]; ATP citrate synth | cytosol [GO:0005829]; ATF | UP000018817 |
| PPTG_15414 | W2PV05 | enoyl-[acyl-carrier-protein] reductase (EC 1.3.1.104) | 348 | | Zinc-containing alcohol dehydrogenase family, Qui | fatty acid biosynthetic process [GO:000663 | mitochondrion [GO:0005739] | enoyl-[acyl-carrier-protein] reductase (NADP | mitochondrion [GO:0005 | UP000018817 |
| PPTG_01415 | W2R6W3 | Coatomer subunit alpha | 1243 | | | endoplasmic reticulum to Golgi vesicle-me | COPI vesicle coat [GO:0030126]; Golgi mem | structural molecule activity [GO:0005198] | COPI vesicle coat [GO:003 | UP000018817 |
| PPTG_12134 | W2Q7V6 | Vacuolar proton pump subunit B (V-ATPase subunit B) (Vacuolar proton pump subunit B) | 495 | | ATPase alpha/beta chains family | ATP metabolic process [GO:0046034]; vacu | proton-transporting V-type ATPase, V1 domai | ATP binding [GO:0005524]; proton-transporti | proton-transporting V-type | UP000018817 |
| PPTG_08926 | W2QK47 | Helicase ATP-binding domain-containing protein | 1094 | | | | cytoplasm [GO:0005737] | ATP binding [GO:0005524]; helicase activity | cytoplasm [GO:0005737]; | UP000018817 |
| PPTG_18707 | W2PGV0 | Alanine--tRNA ligase (EC 6.1.1.7) (Alanyl-tRNA synthetase) (AlaRS) | 994 | | Class-II aminoacyl-tRNA synthetase family, Alax-L | mitochondrial alanyl-tRNA aminoacylation [ | mitochondrion [GO:0005739] | alanine-tRNA ligase activity [GO:0004813]; a | mitochondrion [GO:0005 | UP000018817 |
| PPTG_02435 | W2RD85 | Aldehyde dehydrogenase domain-containing protein | 525 | | Aldehyde dehydrogenase family | | | oxidoreductase activity, acting on the aldehy | oxidoreductase activity, ac | UP000018817 |
| PPTG_06139 | W2QSF2 | Uncharacterized protein | 771 | | | | | oxidoreductase activity, acting on the CH-OH | oxidoreductase activity, ac | UP000018817 |
| PPTG_17398 | W2PM22 | glutamate--tRNA ligase (EC 6.1.1.17) (Glutamyl-tRNA synthetase) | 876 | | Class-I aminoacyl-tRNA synthetase family, Glutam | glutamyl-tRNA aminoacylation [GO:000642 | aminoacyl-tRNA synthetase multienzyme co | ATP binding [GO:0005524]; glutamate-tRNA | aminoacyl-tRNA synthetas | UP000018817 |
| PPTG_05924 | W2QWJ0 | V-type proton ATPase subunit a | 869 | | V-ATPase 116 kDa subunit family | vacuolar acidification [GO:0007035] | vacuolar proton-transporting V-type ATPase, | ATPase binding [GO:0051117]; proton-trans | vacuolar proton-transporti | UP000018817 |
| PPTG_11337 | W2QBU7 | ABC transporter domain-containing protein | 1983 | | ABC transporter superfamily, ABCA family | | membrane [GO:0016020] | ABC-type transporter activity [GO:0140359]; | membrane [GO:0016020]; | UP000018817 |
| PPTG_01995 | W2R940 | F-ATPase gamma subunit | 304 | | ATPase gamma chain family | | proton-transporting ATP synthase complex [G | proton-transporting ATP synthase activity, ro | proton-transporting ATP sy | UP000018817 |
| PPTG_02282 | W2RAS2 | Phospholipid-transporting ATPase (EC 7.6.2.1) | 1516 | | Cation transport ATPase (P-type) (TC 3.A.3) family, | phospholipid translocation [GO:0045332] | endomembrane system [GO:0012505]; plasr | ATP binding [GO:0005524]; ATP hydrolysis ac | endomembrane system [G | UP000018817 |
| PPTG_12013 | W2Q5J5 | Piwi domain-containing protein | 1268 | | Argonaute family | | | RNA binding [GO:0003723] | RNA binding [GO:0003723 | UP000018817 |
| PPTG_00791 | W2RGG3 | Peptidase M1 leukotriene A4 hydrolase/aminopeptidase C-terminal domain-containing protein | 1008 | | | | | aminopeptidase activity [GO:0004177]; met | aminopeptidase activity [G | UP000018817 |
| PPTG_15526 | W2PP80 | Large ribosomal subunit protein uL18 C-terminal eukaryotes domain-containing protein | 301 | | Universal ribosomal protein uL18 family | ribosomal large subunit assembly [GO:000 | cytosolic large ribosomal subunit [GO:0022 | 5S rRNA binding [GO:0008097]; structural co | cytosolic large ribosomal s | UP000018817 |
| PPTG_13182 | W2PZY0 | 1,3-beta-glucan synthase (EC 2.4.1.34) | 2447 | Zhao et al., 2019 | Glycosyltransferase 48 family | (1->3)-beta-D-glucan biosynthetic process [ | 1,3-beta-D-glucan synthase complex [GO:00 | 1,3-beta-D-glucan synthase activity [GO:000 | 1,3-beta-D-glucan synthas | UP000018817 |
| PPTG_00110 | W2RG48 | Glutathione peroxidase | 561 | | Glutathione peroxidase family | response to oxidative stress [GO:0006979] | | peroxidase activity [GO:0004601] | peroxidase activity [GO:00 | UP000018817 |
| PPTG_09984 | W2QFA2 | Gamma carbonic anhydrase | 251 | | | | | | | UP000018817 |
| PPTG_04240 | W2R2K0 | 3-ketoacyl-CoA thiolase | 397 | | Thiolase-like superfamily, Thiolase family | fatty acid beta-oxidation [GO:0006635] | mitochondrion [GO:0005739] | acetyl-CoA C-acetyltransferase activity [GO: | mitochondrion [GO:0005 | UP000018817 |
| PPTG_14789 | W2PXW3 | AAA+ ATPase domain-containing protein | 4117 | | Dynein heavy chain family | microtubule-based movement [GO:000701 | axoneme [GO:0005930]; dynein complex [GC | ATP binding [GO:0005524]; dynein intermedi | axoneme [GO:0005930]; d | UP000018817 |
| PPTG_03763 | W2QZZ8 | Transitional endoplasmic reticulum ATPase | 804 | | | autophagosome maturation [GO:0097352]; | cytosol [GO:0005829]; nucleus [GO:0005634 | ATP binding [GO:0005524]; ATP hydrolysis ac | cytosol [GO:0005829]; nuc | UP000018817 |
| PPTG_03165 | W2R490 | Chaperonin GroL | 598 | Welsh et al., 2024 | Chaperonin (HSP60) family | protein refolding [GO:0042026] | | ATP binding [GO:0005524]; ATP-dependent p | ATP binding [GO:0005524] | UP000018817 |

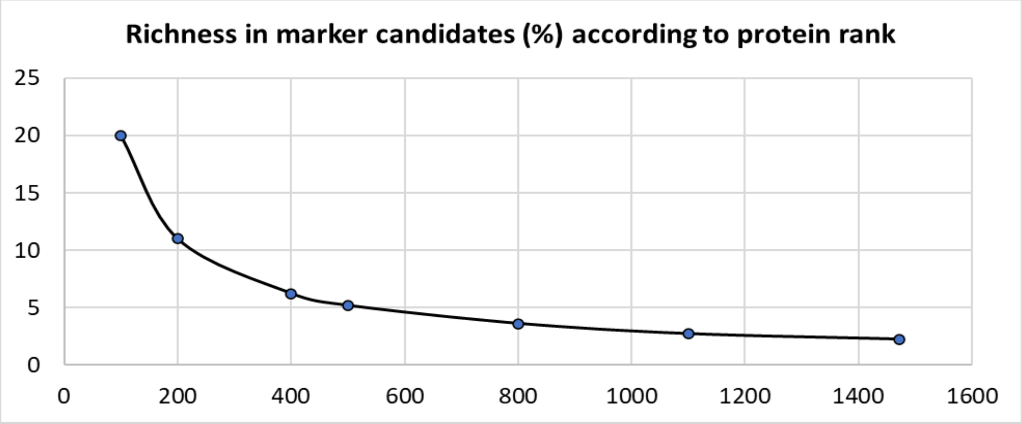

| | | | | | | | | | | |
|---|---|---|---|---|---|---|---|---|---|---|
| PPTG_14992 | W2PTN4 | tyrosine--tRNA ligase (EC 6.1.1.1) (Tyrosyl-tRNA synthetase) | 765 | | Class-I aminoacyl-tRNA synthetase family | tyrosyl-tRNA aminoacylation [GO:0006437] | cytosol [GO:0005829] | ATP binding [GO:0005524]; tyrosine-tRNA lig | cytosol [GO:0005829]; ATP | UP000018817 |
| PPTG_06379 | W2QUL0 | Glutamate/phenylalanine/leucine/valine/L-tryptophan dehydrogenase C-terminal domain-containi | 1057 | | Glu/Leu/Phe/Val dehydrogenases family | L-glutamate catabolic process [GO:000653 | mitochondrion [GO:0005739] | glutamate dehydrogenase (NAD+) activity [G | mitochondrion [GO:00057 | UP000018817 |
| PPTG_13414 | W2Q5J9 | 60S ribosomal protein L3 | 389 | | Universal ribosomal protein uL3 family | translation [GO:0006412] | cytosolic large ribosomal subunit [GO:00226 | RNA binding [GO:0003723]; structural consti | cytosolic large ribosomal s | UP000018817 |
| PPTG_06268 | W2QS57 | Uncharacterized protein | 811 | | CTL (choline transporter-like) family | | membrane [GO:0016020] | transmembrane transporter activity [GO:002 | membrane [GO:0016020]; | UP000018817 |
| PPTG_15113 | W2PW38 | Tubulin beta chain | 446 | | Tubulin family | microtubule-based process [GO:0007017] | microtubule [GO:0005874] | GTP binding [GO:0005525]; GTPase activity [ | microtubule [GO:0005874 | UP000018817 |
| PPTG_01682 | W2R7Z6 | Uncharacterized protein | 183 | | ATPase d subunit family | proton motive force-driven ATP synthesis [G | mitochondrial inner membrane [GO:000574 | proton transmembrane transporter activity [ | mitochondrial inner memb | UP000018817 |
| PPTG_00714 | W2RG42 | Uncharacterized protein | 891 | | | | membrane [GO:0016020] | | membrane [GO:0016020] | UP000018817 |
| PPTG_09527 | W2QG60 | Uncharacterized protein | 997 | | Proteasome subunit S1 family | proteasome-mediated ubiquitin-dependent | nucleus [GO:0005634]; proteasome regulato | enzyme regulator activity [GO:0030234] | nucleus [GO:0005634]; pro | UP000018817 |
| PPTG_19590 | W2PCJ2 | Calreticulin | 458 | | Calreticulin family | ERAD pathway [GO:0036503]; protein foldir | endoplasmic reticulum lumen [GO:0005788] | calcium ion binding [GO:0005509]; carbohyc | endoplasmic reticulum lur | UP000018817 |
| PPTG_08479 | W2QKS8 | methylcrotonoyl-CoA carboxylase (EC 6.4.1.4) (3-methylcrotonyl-CoA carboxylase 2) (3-methylcrot | 567 | | AccD/PCCB family | L-leucine catabolic process [GO:0006552] | methylcrotonoyl-CoA carboxylase complex [ | methylcrotonoyl-CoA carboxylase activity [G | methylcrotonoyl-CoA carb | UP000018817 |
| PPTG_12389 | W2Q3Y1 | Uncharacterized protein | 919 | | | mRNA catabolic process [GO:0006402]; reg | cytosol [GO:0005829]; nucleus [GO:0005634 | nuclease activity [GO:0004518]; RNA binding | cytosol [GO:0005829]; nuc | UP000018817 |
| PPTG_14642 | W2PXZ4 | Glutamine--fructose-6-phosphate aminotransferase [isomerizing] (EC 2.6.1.16) | 701 | | | fructose 6-phosphate metabolic process [GO:0006002]; protein N-linked glycosylation [G | | carbohydrate derivative binding [GO:009736 | carbohydrate derivative bi | UP000018817 |
| PPTG_02473 | W2RDE5 | Aldehyde dehydrogenase domain-containing protein | 525 | | Aldehyde dehydrogenase family | | | oxidoreductase activity, acting on the aldehy | oxidoreductase activity, ac | UP000018817 |
| PPTG_16532 | W2PPX8 | Large ribosomal subunit protein uL4 C-terminal domain-containing protein | 387 | | Universal ribosomal protein uL4 family | translation [GO:0006412] | ribonucleoprotein complex [GO:1990904]; rit | structural constituent of ribosome [GO:0003 | ribonucleoprotein complex | UP000018817 |
| PPTG_00859 | W2RGS8 | Uncharacterized protein | 604 | | | | | | | UP000018817 |
| PPTG_08551 | W2QL14 | 26S proteasome non-ATPase regulatory subunit 3 N-terminal TPR repeats domain-containing prote | 373 | | | ubiquitin-dependent protein catabolic proc | proteasome regulatory particle, lid subcomplex [GO:0008541] | | proteasome regulatory par | UP000018817 |
| PPTG_12096 | W2Q6V4 | Myosin motor domain-containing protein | 1474 | | TRAFAC class myosin-kinesin ATPase superfamily, | actin filament organization [GO:0007015] | cytoplasm [GO:0005737]; membrane [GO:00 | actin filament binding [GO:0051015]; ATP bi | cytoplasm [GO:0005737]; | UP000018817 |
| PPTG_01939 | W2RAZ9 | X8 domain-containing protein | 750 | | Glycosyl hydrolase 5 (cellulase A) family | glucan catabolic process [GO:0009251] | cell surface [GO:0009986]; extracellular regi | beta-glucosidase activity [GO:0008422] | cell surface [GO:0009986] | UP000018817 |
| PPTG_04350 | W2R0C6 | Isocitrate dehydrogenase [NADP] (EC 1.1.1.42) | 427 | | Isocitrate and isopropylmalate dehydrogenases fa | glyoxylate cycle [GO:0006097]; isocitrate m | mitochondrion [GO:0005739] | isocitrate dehydrogenase (NADP+) activity [ | mitochondrion [GO:00057 | UP000018817 |
| PPTG_00228 | W2RE69 | Uncharacterized protein | 612 | | | actin filament depolymerization [GO:00300 | cortical actin cytoskeleton [GO:0030864] | actin filament binding [GO:0051015] | cortical actin cytoskeleton | UP000018817 |
| PPTG_01205 | W2RI21 | Myosin motor domain-containing protein | 1079 | | TRAFAC class myosin-kinesin ATPase superfamily, | actin filament organization [GO:0007015] | cytoplasm [GO:0005737]; membrane [GO:00 | actin filament binding [GO:0051015]; ATP bi | cytoplasm [GO:0005737]; | UP000018817 |
| PPTG_15388 | W2PSU7 | methylmalonate-semialdehyde dehydrogenase (CoA acylating) (EC 1.2.1.27) | 505 | | | L-valine catabolic process [GO:0006574]; thymine catabolic process [GO:0006210] | | methylmalonate-semialdehyde dehydrogena | methylmalonate-semialde | UP000018817 |
| PPTG_14406 | W2PU97 | PDZ domain-containing protein | 1050 | | Peptidase S1C family | proteolysis [GO:0006508] | | serine-type endopeptidase activity [GO:0004 | serine-type endopeptidase | UP000018817 |
| PPTG_00150 | W2RDR6 | Succinate dehydrogenase [ubiquinone] flavoprotein subunit, mitochondrial (EC 1.3.5.1) | 645 | | FAD-dependent oxidoreductase 2 family, FRD/SDH | mitochondrial electron transport, succinate | mitochondrial inner membrane [GO:000574 | electron transfer activity [GO:0009055]; flavi | mitochondrial inner memb | UP000018817 |
| PPTG_10774 | W2QA19 | Eukaryotic translation initiation factor 3 subunit C (eIF3c) (Eukaryotic translation initiation factor 3 | 973 | | EIF-3 subunit C family | formation of cytoplasmic translation initiati | eukaryotic 43S preinitiation complex [GO:00 | RNA binding [GO:0003723]; translation initia | eukaryotic 43S preinitiatio | UP000018817 |
| PPTG_11901 | W2QB87 | Citrate synthase | 464 | | Citrate synthase family | carbohydrate metabolic process [GO:0005 | mitochondrial matrix [GO:0005759] | acyltransferase activity, acyl groups convert | mitochondrial matrix [GO: | UP000018817 |
| PPTG_10291 | W2QEW5 | 5-methyltetrahydropteroyltriglutamate--homocysteine S-methyltransferase (EC 2.1.1.14) | 767 | | Vitamin-B12 independent methionine synthase fa | methionine biosynthetic process [GO:0009086]; methylation [GO:0032259] | | 5-methyltetrahydropteroyltriglutamate-hom | 5-methyltetrahydropteroyl | UP000018817 |
| PPTG_18730 | W2PFQ4 | Calcium-transporting ATPase (EC 7.2.2.10) | 1059 | | Cation transport ATPase (P-type) (TC 3.A.3) family | copper ion transport [GO:0006825] | endomembrane system [GO:0012505]; plasr | ATP binding [GO:0005524]; ATP hydrolysis ac | endomembrane system [G | UP000018817 |
| PPTG_11356 | W2QB22 | 60S acidic ribosomal protein P0 | 317 | | Universal ribosomal protein uL10 family | cytoplasmic translation [GO:0002181]; ribo | cytosolic large ribosomal subunit [GO:00226 | large ribosomal subunit rRNA binding [GO:0 | cytosolic large ribosomal s | UP000018817 |
| PPTG_03117 | W2R3Q4 | subtilisin (EC 3.4.21.62) | 486 | | Peptidase S8 family | proteolysis [GO:0006508] | | serine-type endopeptidase activity [GO:0004 | serine-type endopeptidase | UP000018817 |
| PPTG_13885 | W2Q1C0 | Dihydrodipicolinate reductase | 633 | | | lysine biosynthetic process via diaminopimelate [GO:0009089] | | 2-aminoadipate transaminase activity [GO:0 | 2-aminoadipate transamir | UP000018817 |
| PPTG_10972 | W2QCT9 | mitochondrial processing peptidase (EC 3.4.24.64) (Beta-MPP) | 466 | | Peptidase M16 family | proteolysis [GO:0006508] | mitochondrial matrix [GO:0005759] | metal ion binding [GO:0046872]; metalloenc | mitochondrial matrix [GO: | UP000018817 |
| PPTG_11759 | W2Q908 | Alpha-mannosidase (EC 3.2.1.-) | 1025 | | Glycosyl hydrolase 38 family | mannose metabolic process [GO:0006013] | | alpha-mannosidase activity [GO:0004559]; | alpha-mannosidase activit | UP000018817 |
| PPTG_17902 | W2PJ95 | Cellulose synthase 3 | 1142 | | | cellulose biosynthetic process [GO:003024 | endomembrane system [GO:0012505]; mem | cellulose synthase (UDP-forming) activity [G | endomembrane system [G | UP000018817 |
| PPTG_08190 | W2QKA6 | phosphatidylinositol 3-kinase (EC 2.7.1.137) | 1036 | | | bleb assembly [GO:0032060]; cell migratior | cytoplasm [GO:0005737]; phosphatidylinosit | 1-phosphatidylinositol-3-kinase activity [GO: | cytoplasm [GO:0005737]; | UP000018817 |
| PPTG_16495 | W2PPR1 | E1 ubiquitin-activating enzyme (EC 6.2.1.45) | 1124 | | Ubiquitin-activating E1 family | protein sumoylation [GO:0016925] | cytoplasm [GO:0005737]; SUMO activating e | ATP binding [GO:0005524]; SUMO activating | cytoplasm [GO:0005737]; | UP000018817 |
| PPTG_18870 | W2PFJ5 | ATP synthase subunit b | 308 | | Eukaryotic ATPase B chain family | proton motive force-driven ATP synthesis [G | mitochondrial inner membrane [GO:000574 | proton transmembrane transporter activity [ | mitochondrial inner memb | UP000018817 |
| PPTG_08622 | W2QL97 | Serine hydroxymethyltransferase (EC 2.1.2.1) | 464 | | SHMT family | glycine biosynthetic process from serine [G | mitochondrion [GO:0005739] | glycine hydroxymethyltransferase activity [G | mitochondrion [GO:00057 | UP000018817 |
| PPTG_01466 | W2R9J1 | Eukaryotic translation initiation factor 5A (eIF-5A) | 161 | | EIF-5A family | positive regulation of translational elongati | chloroplast [GO:0009507] | ribosome binding [GO:0043022]; RNA bindin | chloroplast [GO:0009507]; | UP000018817 |
| PPTG_18309 | W2PGM1 | Protein disulfide-isomerase (EC 5.3.4.1) | 521 | | Protein disulfide isomerase family | protein folding [GO:0006457]; response to | endoplasmic reticulum lumen [GO:0005788] | protein disulfide isomerase activity [GO:000 | endoplasmic reticulum lur | UP000018817 |
| PPTG_08986 | W2QIM1 | ABC transporter domain-containing protein | 1945 | | ABC transporter superfamily, ABCA family | | membrane [GO:0016020] | ABC-type transporter activity [GO:0140359]; | membrane [GO:0016020]; | UP000018817 |
| PPTG_08453 | W2QMI3 | Tubulin alpha chain | 454 | | Tubulin family | microtubule-based process [GO:0007017] | microtubule [GO:0005874] | GTP binding [GO:0005525]; hydrolase activit | microtubule [GO:0005874 | UP000018817 |
| PPTG_10444 | W2QEI6 | phospholipase D (EC 3.1.4.4) | 530 | | | phospholipid catabolic process [GO:00093 | plasma membrane [GO:0005886] | phospholipase D activity [GO:0004630] | plasma membrane [GO:00 | UP000018817 |
| PPTG_01189 | W2RIK1 | Secreted protein | 457 | | | | | | | UP000018817 |
| PPTG_17298 | W2PJE4 | Aconitate hydratase | 700 | | | L-amino acid biosynthetic process [GO:0170034]; proteinogenic amino acid biosynthetic | | hydro-lyase activity [GO:0016836]; iron-sulfu | hydro-lyase activity [GO:0 | UP000018817 |
| PPTG_15137 | W2PU56 | Dynamin GTPase | 705 | | TRAFAC class dynamin-like GTPase superfamily, Dynamin/Fzo/YdjA family | | cytoplasm [GO:0005737]; membrane [GO:00 | GTP binding [GO:0005525]; GTPase activity [ | cytoplasm [GO:0005737]; | UP000018817 |
| PPTG_02874 | W2RCX3 | glycerol kinase (EC 2.7.1.30) (ATP:glycerol 3-phosphotransferase) | 552 | | FGGY kinase family | glycerol catabolic process [GO:0019563]; gl | cytosol [GO:0005829] | ATP binding [GO:0005524]; glycerol kinase a | cytosol [GO:0005829]; ATP | UP000018817 |
| PPTG_01022 | W2RHI5 | Glucose-6-phosphate isomerase (EC 5.3.1.9) | 556 | | GPI family | gluconeogenesis [GO:0006094]; glucose 6- | cytosol [GO:0005829] | carbohydrate derivative binding [GO:009736 | cytosol [GO:0005829]; car | UP000018817 |
| PPTG_15898 | W2PSQ3 | C2 domain-containing protein | 1174 | | | | membrane [GO:0016020] | chloride channel activity [GO:0005254] | membrane [GO:0016020]; | UP000018817 |
| PPTG_07665 | W2QN93 | Glycosyl hydrolase family 30 TIM-barrel domain-containing protein | 540 | | Glycosyl hydrolase 30 family | glucosylceramide catabolic process [GO:00 | membrane [GO:0016020] | glucosylceramidase activity [GO:0004348] | membrane [GO:0016020]; | UP000018817 |
| PPTG_01692 | W2R8G7 | Mitochondrial carnitine/acylcarnitine carrier protein | 310 | | Mitochondrial carrier (TC 2.A.29) family | carnitine transmembrane transport [GO:19 | mitochondrial membrane [GO:0031966] | O-acyl-L-carnitine transmembrane transport | mitochondrial membrane | UP000018817 |
| PPTG_11158 | W2QA21 | Elongation factor 1-alpha | 443 | | TRAFAC class translation factor GTPase superfamily, Classic translation factor GTPase family, | | cytoplasm [GO:0005737] | GTP binding [GO:0005525]; GTPase activity [ | cytoplasm [GO:0005737]; | UP000018817 |
| PPTG_08465 | W2QNB5 | Tubulin alpha chain | 452 | | Tubulin family | microtubule-based process [GO:0007017] | microtubule [GO:0005874] | GTP binding [GO:0005525]; hydrolase activit | microtubule [GO:0005874 | UP000018817 |
| PPTG_10588 | W2QD53 | ABC transporter domain-containing protein | 1351 | | ABC transporter superfamily, ABCG family, PDR (TC 3.A.1.205) subfamily | | membrane [GO:0016020] | ABC-type transporter activity [GO:0140359]; | membrane [GO:0016020]; | UP000018817 |
| PPTG_04978 | W2R3G8 | Rab GDP dissociation inhibitor | 462 | | Rab GDI family | protein transport [GO:0015031]; small GTP | cytoplasm [GO:0005737] | Rab GDP-dissociation inhibitor activity [GO: | cytoplasm [GO:0005737]; | UP000018817 |
| PPTG_13413 | W2Q4S5 | Dolichyl-diphosphooligosaccharide--protein glycosyltransferase subunit 1 | 470 | | OST1 family | protein N-linked glycosylation via asparagin | oligosaccharyltransferase complex [GO:0008250] | | oligosaccharyltransferase | UP000018817 |
| PPTG_01563 | W2R7Y2 | Polyadenylate-binding protein (PABP) | 640 | | Polyadenylate-binding protein type-1 family | mRNA processing [GO:0006397] | cytoplasm [GO:0005737]; nucleus [GO:0005 | RNA binding [GO:0003723] | cytoplasm [GO:0005737]; | UP000018817 |
| PPTG_15102 | W2PW23 | Glucose-6-phosphate 1-dehydrogenase (EC 1.1.1.49) | 552 | | Glucose-6-phosphate dehydrogenase family | glucose metabolic process [GO:0006006]; pentose-phosphate shunt, oxidative branch [G | | glucose-6-phosphate dehydrogenase activity | glucose-6-phosphate dehy | UP000018817 |
| PPTG_09470 | W2QH02 | Uncharacterized protein | 2326 | | | fatty acid biosynthetic process [GO:0006633]; malonyl-CoA biosynthetic process [GO:20 | | acetyl-CoA carboxylase activity [GO:000398 | acetyl-CoA carboxylase ac | UP000018817 |
| PPTG_14740 | W2PVT8 | 1,3-beta-glucan synthase (EC 2.4.1.34) | 2040 | Zhao et al., 2019 | Glycosyltransferase 48 family | (1->3)-beta-D-glucan biosynthetic process [ | 1,3-beta-D-glucan synthase complex [GO:00 | 1,3-beta-D-glucan synthase activity [GO:000 | 1,3-beta-D-glucan synthas | UP000018817 |
| PPTG_11194 | W2Q9D3 | EF-hand domain-containing protein | 2480 | | Recoverin family | | | calcium ion binding [GO:0005509] | calcium ion binding [GO:0 | UP000018817 |
| PPTG_17691 | W2PLE5 | Plasma membrane ATPase (EC 7.1.2.1) | 793 | | Cation transport ATPase (P-type) (TC 3.A.3) family, | proton export across plasma membrane [G | plasma membrane [GO:0005886] | ATP binding [GO:0005524]; ATP hydrolysis ac | plasma membrane [GO:00 | UP000018817 |
| PPTG_03625 | W2R5K1 | P-type Ca(2+) transporter (EC 7.2.2.10) | 1045 | | Cation transport ATPase (P-type) (TC 3.A.3) family, Type IIA subfamily | | membrane [GO:0016020] | ATP binding [GO:0005524]; ATP hydrolysis ac | membrane [GO:0016020]; | UP000018817 |
| PPTG_02578 | W2RDH0 | arginine--tRNA ligase (EC 6.1.1.19) (Arginyl-tRNA synthetase) | 815 | | Class-I aminoacyl-tRNA synthetase family | arginyl-tRNA aminoacylation [GO:0006420] | cytoplasm [GO:0005737] | arginine-tRNA ligase activity [GO:0004814]; | cytoplasm [GO:0005737]; | UP000018817 |
| PPTG_09888 | W2QE91 | phosphopyruvate hydratase (EC 4.2.1.11) | 457 | | Enolase family | glycolytic process [GO:0006096] | phosphopyruvate hydratase complex [GO:00 | magnesium ion binding [GO:0000287]; phos | phosphopyruvate hydratas | UP000018817 |
| PPTG_14635 | W2PVE4 | aspartyl aminopeptidase (EC 3.4.11.21) | 462 | | Peptidase M18 family | proteolysis [GO:0006508] | cytoplasm [GO:0005737] | aminopeptidase activity [GO:0004177]; met | cytoplasm [GO:0005737]; | UP000018817 |
| PPTG_00328 | W2RGZ4 | Beta-adaptin appendage C-terminal subdomain domain-containing protein | 915 | | Adaptor complexes large subunit family | intracellular protein transport [GO:0006886 | clathrin adaptor complex [GO:0030131]; cytoplasmic vesicle [GO:0031410]; endomembra | | clathrin adaptor complex [ | UP000018817 |
| PPTG_01949 | W2R9G7 | Adenosylhomocysteinase (EC 3.13.2.1) | 481 | | Adenosylhomocysteinase family | one-carbon metabolic process [GO:000673 | cytosol [GO:0005829] | adenosylhomocysteinase activity [GO:0004 | cytosol [GO:0005829]; ade | UP000018817 |
| PPTG_11937 | W2Q8V7 | C2 domain-containing protein | 1827 | | | plasma membrane organization [GO:00070 | membrane [GO:0016020] | | membrane [GO:0016020]; | UP000018817 |
| PPTG_14774 | W2PYI8 | Malate synthase (EC 2.3.3.9) | 535 | | Malate synthase family | glyoxylate cycle [GO:0006097]; tricarboxylic | cytoplasm [GO:0005737] | malate synthase activity [GO:0004474] | cytoplasm [GO:0005737]; | UP000018817 |
| PPTG_18274 | W2PGE0 | Chaperone dnaK | 648 | | Heat shock protein 70 family | | | ATP binding [GO:0005524]; ATP-dependent | ATP binding [GO:0005524] | UP000018817 |
| PPTG_20398 | W2PBA3 | Malate synthase (EC 2.3.3.9) | 535 | | Malate synthase family | glyoxylate cycle [GO:0006097]; tricarboxylic | cytoplasm [GO:0005737] | malate synthase activity [GO:0004474] | cytoplasm [GO:0005737]; | UP000018817 |
| PPTG_15943 | W2PU40 | Ketol-acid reductoisomerase, chloroplastic (EC 1.1.1.86) (Acetohydroxy-acid reductoisomerase) (A | 515 | | Ketol-acid reductoisomerase family | isoleucine biosynthetic process [GO:00090 | chloroplast [GO:0009507] | isomerase activity [GO:0016853]; ketol-acid | chloroplast [GO:0009507]; | UP000018817 |
| PPTG_04558 | W2R3T9 | Uncharacterized protein | 542 | | | cellulose catabolic process [GO:0030245] | | | cellulose catabolic proces | UP000018817 |
| PPTG_14734 | W2PXM6 | 60S ribosomal protein L7a | 263 | | Eukaryotic ribosomal protein eL8 family | ribosome biogenesis [GO:0042254] | cytosolic large ribosomal subunit [GO:00226 | RNA binding [GO:0003723] | cytosolic large ribosomal s | UP000018817 |
| PPTG_19127 | W2PDR1 | Mitochondrial phosphate carrier protein | 345 | | Mitochondrial carrier (TC 2.A.29) family | mitochondrial phosphate ion transmembra | mitochondrial inner membrane [GO:000574 | phosphate transmembrane transporter activ | mitochondrial inner memb | UP000018817 |
| PPTG_09250 | W2QGX3 | NAD(P)H:quinone oxidoreductase, type IV | 200 | | WrbA family | | membrane [GO:0016020] | FMN binding [GO:0010181]; NAD(P)H dehyd | membrane [GO:0016020]; | UP000018817 |
| PPTG_02536 | W2RDB0 | TOG domain-containing protein | 2759 | | GCN1 family | cellular response to amino acid starvation [ | cytosol [GO:0005829] | protein kinase regulator activity [GO:001988 | cytosol [GO:0005829]; pro | UP000018817 |
| PPTG_00853 | W2RGP1 | inositol-3-phosphate synthase (EC 5.5.1.4) | 517 | | Myo-inositol 1-phosphate synthase family | inositol biosynthetic process [GO:0006021] | cytoplasm [GO:0005737] | inositol-3-phosphate synthase activity [GO:0 | cytoplasm [GO:0005737]; | UP000018817 |
| PPTG_07764 | W2QMB5 | Small ribosomal subunit protein eS1 | 261 | | Eukaryotic ribosomal protein eS1 family | translation [GO:0006412] | cytosolic small ribosomal subunit [GO:00226 | structural constituent of ribosome [GO:0003 | cytosolic small ribosomal | UP000018817 |
| PPTG_08768 | W2QHM7 | ABC transporter domain-containing protein | 1969 | | ABC transporter superfamily, ABCA family | | membrane [GO:0016020] | ABC-type transporter activity [GO:0140359]; | membrane [GO:0016020]; | UP000018817 |
| PPTG_11971 | W2Q5E7 | Uncharacterized protein | 496 | | | | | | | UP000018817 |
| PPTG_13000 | W2Q5X9 | Dihydrolipoyl dehydrogenase (EC 1.8.1.4) | 387 | | Class-I pyridine nucleotide-disulfide oxidoreductas | 2-oxoglutarate metabolic process [GO:0006 | mitochondrion [GO:0005739]; oxoglutarate d | dihydrolipoyl dehydrogenase (NADH) activity | mitochondrion [GO:00057 | UP000018817 |
| PPTG_11916 | W2Q8U0 | Fructose-1,6-bisphosphatase, cytosolic (EC 3.1.3.11) | 333 | | FBPase class 1 family | fructose 1,6-bisphosphate metabolic proce | cytosol [GO:0005829] | fructose 1,6-bisphosphate 1-phosphatase a | cytosol [GO:0005829]; fruc | UP000018817 |
| PPTG_00673 | W2RFX8 | TATA-binding protein interacting (TIP20) domain-containing protein | 1180 | | CAND family | SCF complex assembly [GO:0010265] | | | SCF complex assembly [G | UP000018817 |
| PPTG_09177 | W2QHJ6 | Alpha-L-glutamate ligase-related protein ATP-grasp domain-containing protein | 426 | | | | | | | UP000018817 |
| PPTG_04610 | W2R1L5 | Beta'-coat protein | 1094 | | WD repeat COPB2 family | endoplasmic reticulum to Golgi vesicle-me | COPI vesicle coat [GO:0030126]; Golgi meml | structural molecule activity [GO:0005198] | COPI vesicle coat [GO:003 | UP000018817 |
| PPTG_12917 | W2Q337 | isoleucine--tRNA ligase (EC 6.1.1.5) (Isoleucyl-tRNA synthetase) | 1175 | | Class-I aminoacyl-tRNA synthetase family | isoleucyl-tRNA aminoacylation [GO:0006428] | | aminoacyl-tRNA deacylase activity [GO:000 | aminoacyl-tRNA deacylase | UP000018817 |
| PPTG_17187 | W2PL97 | glucan endo-1,3-beta-D-glucosidase (EC 3.2.1.39) (Endo-1,3-beta-glucanase btgC) (Laminarinase | 458 | | | cell wall organization [GO:0071555]; polysa | plasma membrane [GO:0005886] | glucan endo-1,3-beta-D-glucosidase activity | plasma membrane [GO:00 | UP000018817 |
| PPTG_18619 | W2PIH3 | NADH dehydrogenase [ubiquinone] flavoprotein 1, mitochondrial (EC 7.1.1.2) | 499 | | Complex I 51 kDa subunit family | mitochondrial electron transport, NADH to | mitochondrial inner membrane [GO:000574 | 4 iron, 4 sulfur cluster binding [GO:0051539] | mitochondrial inner memb | UP000018817 |
| PPTG_19136 | W2PDK7 | Acyl-CoA dehydrogenase | 610 | | Acyl-CoA dehydrogenase family | fatty acid metabolic process [GO:0006631] | mitochondrial inner membrane [GO:000574 | acyl-CoA dehydrogenase activity [GO:00039 | mitochondrial inner memb | UP000018817 |
| PPTG_16241 | W2PP34 | 2-oxoglutarate dehydrogenase, mitochondrial (EC 1.2.4.2) (2-oxoglutarate dehydrogenase comple | 1043 | | Alpha-ketoglutarate dehydrogenase family | tricarboxylic acid cycle [GO:0006099] | mitochondrion [GO:0005739]; oxoglutarate d | metal ion binding [GO:0046872]; oxoglutarat | mitochondrion [GO:00057 | UP000018817 |
| PPTG_03132 | W2R452 | Glutamate/phenylalanine/leucine/valine/L-tryptophan dehydrogenase C-terminal domain-containi | 1032 | | Glu/Leu/Phe/Val dehydrogenases family | L-glutamate catabolic process [GO:000653 | mitochondrion [GO:0005739] | glutamate dehydrogenase (NAD+) activity [G | mitochondrion [GO:00057 | UP000018817 |
| PPTG_15793 | W2PPD3 | GB1/RHD3-type G domain-containing protein | 638 | | TRAFAC class dynamin-like GTPase superfamily, GB1/RHD3 GTPase family | | | GTP binding [GO:0005525]; GTPase activity [ | GTP binding [GO:0005525] | UP000018817 |
| PPTG_17946 | W2PIM1 | RNA helicase (EC 3.6.4.13) | 411 | | DEAD box helicase family, eIF4A subfamily | | | ATP binding [GO:0005524]; hydrolase activity | ATP binding [GO:0005524] | UP000018817 |
| PPTG_13850 | W2Q158 | fumarate hydratase (EC 4.2.1.2) | 528 | | Class-II fumarase/aspartase family, Fumarase sub | fumarate metabolic process [GO:0006106]; | mitochondrion [GO:0005739] | fumarate hydratase activity [GO:0004333] | mitochondrion [GO:00057 | UP000018817 |
| PPTG_07238 | W2QPH8 | EGF-like domain-containing protein | 226 | | | heterophilic cell-cell adhesion via plasma membrane cell adhesion molecules [GO:0007 | | cell adhesion molecule binding [GO:005083 | cell adhesion molecule bi | UP000018817 |
| PPTG_16640 | W2PM64 | Aldehyde dehydrogenase domain-containing protein | 547 | | Aldehyde dehydrogenase family | | | aldehyde dehydrogenase (NAD+) activity [G | aldehyde dehydrogenase ( | UP000018817 |
| PPTG_19659 | W2PEB1 | S-(hydroxymethyl)glutathione dehydrogenase (EC 1.1.1.284) | 387 | | Zinc-containing alcohol dehydrogenase family, Cla | formaldehyde catabolic process [GO:00462 | cytosol [GO:0005829] | alcohol dehydrogenase (NAD+) activity [GO: | cytosol [GO:0005829]; alc | UP000018817 |
| PPTG_13436 | W2Q1B4 | Protein transport protein SEC31 (Protein transport protein sec31) | 1175 | | WD repeat SEC31 family | COPII-coated vesicle cargo loading [GO:009 | COPII vesicle coat [GO:0030127]; endoplasm | structural molecule activity [GO:0005198] | COPII vesicle coat [GO:00 | UP000018817 |
| PPTG_07472 | W2QN94 | Uncharacterized protein | 1001 | | Peptidase S8 family | proteolysis [GO:0006508] | cytosol [GO:0005829] | aminopeptidase activity [GO:0004177]; serir | cytosol [GO:0005829]; am | UP000018817 |

| | | | | | | | | | | |
|---|---|---|---|---|---|---|---|---|---|---|
| PPTG_10119 | W2QD88 | Coatomer subunit gamma | 931 | | COPG family | endoplasmic reticulum to Golgi vesicle-me | COPI vesicle coat [GO:0030126]; endoplasm | structural molecule activity [GO:0005198] | COPI vesicle coat [GO:003 | UP000018817 |
| PPTG_01922 | W2RAY1 | phosphoenolpyruvate carboxykinase (ATP) (EC 4.1.1.49) | 569 | | Phosphoenolpyruvate carboxykinase (ATP) family | gluconeogenesis [GO:0006094] | cytosol [GO:0005829] | ATP binding [GO:0005524]; kinase activity [G | cytosol [GO:0005829]; ATF | UP000018817 |
| PPTG_14608 | W2PXW1 | Electron transfer flavoprotein-ubiquinone oxidoreductase (ETF-QO) (EC 1.5.5.1) | 610 | | ETF-QO/FixC family | | mitochondrial inner membrane [GO:000574 | 4 iron, 4 sulfur cluster binding [GO:0051539] | mitochondrial inner memb | UP000018817 |
| PPTG_10900 | W2QAL4 | Peptidase M28 domain-containing protein | 868 | | Peptidase M28 family, M28B subfamily | | | carboxypeptidase activity [GO:0004180] | carboxypeptidase activity [ | UP000018817 |
| PPTG_03857 | W2R0Z2 | Vesicle-fusing ATPase (EC 3.6.4.6) | 765 | | AAA ATPase family | Golgi to plasma membrane protein transpo | Golgi stack [GO:0005795] | ATP binding [GO:0005524]; ATP hydrolysis ac | Golgi stack [GO:0005795]; | UP000018817 |
| PPTG_00165 | W2RDT8 | EGF-like domain-containing protein | 723 | Ral et al., 2024 | | (1->6)-beta-D-glucan biosynthetic process [ | endoplasmic reticulum membrane [GO:0005 | glucosidase activity [GO:0015926] | endoplasmic reticulum me | UP000018817 |
| PPTG_16899 | W2PMI6 | DNA-binding protein, 42 kDa | 393 | | Peptidase M24 family | | | DNA binding [GO:0003677] | DNA binding [GO:0003677 | UP000018817 |
| PPTG_14388 | W2PYI0 | HECT-type E3 ubiquitin transferase (EC 2.3.2.26) | 4149 | | UPL family, TOM1/PTR1 subfamily | mRNA transport [GO:0051028]; protein poly | cytoplasm [GO:0005737]; nucleus [GO:0005 | ubiquitin protein ligase activity [GO:0061630 | cytoplasm [GO:0005737]; | UP000018817 |
| PPTG_16875 | W2PNH8 | Cation-transporting ATPase (EC 7.2.2.-) | 1339 | | Cation transport ATPase (P-type) (TC 3.A.3) family, Type V subfamily | | membrane [GO:0016020] | ATP binding [GO:0005524]; ATP hydrolysis ac | membrane [GO:0016020]; | UP000018817 |
| PPTG_15783 | W2PPP1 | 40S ribosomal protein S7 | 190 | | Eukaryotic ribosomal protein eS7 family | ribosomal small subunit biogenesis [GO:00 | 90S preribosome [GO:0030686]; cytosolic sm | structural constituent of ribosome [GO:0003 | 90S preribosome [GO:003 | UP000018817 |
| PPTG_14116 | W2PXL8 | Peptidase C-terminal archaeal/bacterial domain-containing protein | 247 | | | | | | | UP000018817 |
| PPTG_11874 | W2Q9D6 | Small ribosomal subunit protein uS5 (40S ribosomal protein S2) | 261 | | Universal ribosomal protein uS5 family | translation [GO:0006412] | cytosolic small ribosomal subunit [GO:00226 | RNA binding [GO:0003723]; structural consti | cytosolic small ribosomal | UP000018817 |
| PPTG_15328 | W2PSP8 | Phospholipid-transporting ATPase (EC 7.6.2.1) | 1391 | | Cation transport ATPase (P-type) (TC 3.A.3) family, | phospholipid translocation [GO:0045332] | endomembrane system [GO:0012505]; plasn | ATP binding [GO:0005524]; ATP hydrolysis ac | endomembrane system [G | UP000018817 |
| PPTG_01838 | W2RAJ2 | phosphoribosylaminoimidazole carboxylase (EC 4.1.1.21) (AIR carboxylase) | 595 | | AIR carboxylase family, Class I subfamily | 'de novo' IMP biosynthetic process [GO:0006189] | | ATP binding [GO:0005524]; metal ion binding | ATP binding [GO:0005524] | UP000018817 |
| PPTG_18811 | W2PFW8 | Medium-chain specific acyl-CoA dehydrogenase, mitochondrial | 362 | | Acyl-CoA dehydrogenase family | medium-chain fatty acid catabolic process | mitochondrion [GO:0005739] | flavin adenine dinucleotide binding [GO:005 | mitochondrion [GO:0005 | UP000018817 |
| PPTG_16504 | W2PNQ9 | Isocitrate lyase | 537 | | Isocitrate lyase/PEP mutase superfamily, Isocitrate | glyoxylate cycle [GO:0006097]; tricarboxylic acid cycle [GO:0006099] | | isocitrate lyase activity [GO:0004451]; metal | isocitrate lyase activity [G | UP000018817 |
| PPTG_13500 | W2Q3G6 | cysteine--tRNA ligase (EC 6.1.1.16) (Cysteinyl-tRNA synthetase) | 688 | | Class-I aminoacyl-tRNA synthetase family | cysteinyl-tRNA aminoacylation [GO:000642 | cytoplasm [GO:0005737] | ATP binding [GO:0005524]; cysteine-tRNA lig | cytoplasm [GO:0005737]; | UP000018817 |
| PPTG_14253 | W2PZ52 | Tyrosinase copper-binding domain-containing protein | 631 | | | | | metal ion binding [GO:0046872]; oxidoreduc | metal ion binding [GO:004 | UP000018817 |
| PPTG_06616 | W2QS77 | Adenylosuccinate synthetase (AMPSase) (AdSS) (EC 6.3.4.4) (IMP--aspartate ligase) | 554 | | Adenylosuccinate synthetase family | 'de novo' AMP biosynthetic process [GO:00 | cytoplasm [GO:0005737] | adenylosuccinate synthase activity [GO:000 | cytoplasm [GO:0005737]; | UP000018817 |
| PPTG_09817 | W2QC07 | Thioredoxin domain-containing protein | 469 | | | | COPII-coated ER to Golgi transport vesicle [GO:0030134]; endoplasmic reticulum [GO:000 | | COPII-coated ER to Golgi t | UP000018817 |
| PPTG_11914 | W2QBD5 | Translation elongation factor EF1B beta/delta subunit guanine nucleotide exchange domain-contai | 228 | | EF-1-beta/EF-1-delta family | | cytosol [GO:0005829]; eukaryotic translation | guanyl-nucleotide exchange factor activity [ | cytosol [GO:0005829]; euk | UP000018817 |
| PPTG_15972 | W2PS84 | Aminopeptidase (EC 3.4.11.-) | 884 | | Peptidase M1 family | peptide catabolic process [GO:0043171]; p | cytoplasm [GO:0005737]; extracellular space | metalloaminopeptidase activity [GO:007000 | cytoplasm [GO:0005737]; | UP000018817 |
| PPTG_03240 | W2R6M4 | glutamate synthase (ferredoxin) (EC 1.4.7.1) | 1582 | | Glutamate synthase family | ammonia assimilation cycle [GO:0019676]; glutamate biosynthetic process [GO:000653 | | 3 iron, 4 sulfur cluster binding [GO:0051538] | 3 iron, 4 sulfur cluster bind | UP000018817 |
| PPTG_11537 | W2Q6A3 | Lipase-like C-terminal domain-containing protein | 479 | | | lipid metabolic process [GO:0006629] | extracellular region [GO:0005576] | hydrolase activity [GO:0016787] | extracellular region [GO:0 | UP000018817 |
| PPTG_18378 | W2PH74 | Uncharacterized protein | 344 | | WD repeat G protein beta family | signal transduction [GO:0007165] | | | signal transduction [GO:0 | UP000018817 |
| PPTG_09098 | W2QG96 | Dynein-1, subspecies f | 4551 | | Dynein heavy chain family | cilium movement involved in cell motility [G | inner dynein arm [GO:0036156]; microtubule | ATP binding [GO:0005524]; dynein intermedi | inner dynein arm [GO:003 | UP000018817 |
| PPTG_10590 | W2QC67 | ABC transporter domain-containing protein | 1358 | | ABC transporter superfamily, ABCG family, PDR (TC 3.A.1.205) subfamily | | membrane [GO:0016020] | ABC-type transporter activity [GO:0140359]; | membrane [GO:0016020]; | UP000018817 |
| PPTG_15229 | W2PSK8 | C2 domain-containing protein | 871 | | | | membrane [GO:0016020] | calcium ion binding [GO:0005509] | membrane [GO:0016020]; | UP000018817 |
| PPTG_08506 | W2QKW5 | Tubulin alpha chain | 453 | | Tubulin family | microtubule-based process [GO:0007017] | microtubule [GO:0005874] | GTP binding [GO:0005525]; hydrolase activit | microtubule [GO:0005874 | UP000018817 |
| PPTG_08498 | W2QKW0 | Tubulin alpha chain | 453 | | Tubulin family | microtubule-based process [GO:0007017] | microtubule [GO:0005874] | GTP binding [GO:0005525]; hydrolase activit | microtubule [GO:0005874 | UP000018817 |
| PPTG_02328 | W2RCX4 | NAD-dependent epimerase/dehydratase domain-containing protein | 379 | | | | mitochondrion [GO:0005739] | protein-containing complex binding [GO:004 | mitochondrion [GO:0005 | UP000018817 |
| PPTG_14857 | W2PWB8 | AP-4 complex subunit epsilon-1 C-terminal domain-containing protein | 1103 | | | clathrin-dependent endocytosis [GO:00725 | AP-2 adaptor complex [GO:0030122] | clathrin adaptor activity [GO:0035615] | AP-2 adaptor complex [GO | UP000018817 |
| PPTG_02398 | W2RCS0 | Elongation factor 1-gamma | 406 | | | | cytoplasm [GO:0005737]; nucleus [GO:0005 | translation elongation factor activity [GO:00 | cytoplasm [GO:0005737]; | UP000018817 |
| PPTG_17524 | W2PKD5 | Thioredoxin domain-containing protein | 377 | | Peroxiredoxin family, AhpC/Prx1 subfamily | cell redox homeostasis [GO:0045454]; cell | cytosol [GO:0005829] | thioredoxin peroxidase activity [GO:0008379 | cytosol [GO:0005829]; thic | UP000018817 |
| PPTG_18809 | W2PHP4 | proline--tRNA ligase (EC 6.1.1.15) (Prolyl-tRNA synthetase) | 711 | | | prolyl-tRNA aminoacylation [GO:0006433] | aminoacyl-tRNA synthetase multienzyme co | aminoacyl-tRNA deacylase activity [GO:000 | aminoacyl-tRNA synthetas | UP000018817 |
| PPTG_16306 | W2PRN2 | Uncharacterized protein | 230 | | | | | | | UP000018817 |
| PPTG_02800 | W2RCJ8 | Transaldolase (EC 2.2.1.2) | 334 | | Transaldolase family, Type 1 subfamily | carbohydrate metabolic process [GO:0005 | cytoplasm [GO:0005737] | transaldolase activity [GO:0004801] | cytoplasm [GO:0005737]; | UP000018817 |
| PPTG_16510 | W2PN09 | Inosine-5'-monophosphate dehydrogenase (IMP dehydrogenase) (IMPD) (IMPDH) (EC 1.1.1.205) | 508 | | IMPDH/GMPR family | GMP biosynthetic process [GO:0006177]; G | cytoplasm [GO:0005737] | IMP dehydrogenase activity [GO:0003938]; n | cytoplasm [GO:0005737]; | UP000018817 |
| PPTG_14664 | W2PWA6 | Phosphatase PP2A regulatory subunit A/Splicing factor 3B subunit 1-like HEAT repeat domain-cont | 457 | | Phosphatase 2A regulatory subunit A family | | cytosol [GO:0005829]; nucleus [GO:0005634 | protein phosphatase regulator activity [GO:0 | cytosol [GO:0005829]; nuc | UP000018817 |
| PPTG_20006 | W2P9U9 | Uncharacterized protein | 293 | | SNAP family | intracellular protein transport [GO:0006886 | SNARE complex [GO:0031201]; vacuolar me | soluble NSF attachment protein activity [GO | SNARE complex [GO:0031 | UP000018817 |
| PPTG_00098 | W2RFQ0 | Uncharacterized protein | 317 | | | | | | | UP000018817 |
| PPTG_13005 | W2Q5Y3 | Ribosomal protein | 229 | | Universal ribosomal protein uL1 family | translation [GO:0006412] | large ribosomal subunit [GO:0015934] | RNA binding [GO:0003723]; structural consti | large ribosomal subunit [G | UP000018817 |
| PPTG_13491 | W2Q1K6 | Phospholipase (EC 3.1.4.4) | 1117 | | Phospholipase D family | intracellular signal transduction [GO:0035 | plasma membrane [GO:0005886] | phospholipase D activity [GO:0004630] | plasma membrane [GO:0 | UP000018817 |
| PPTG_03845 | W2QY59 | Glycoside hydrolase family 5 domain-containing protein | 582 | | Glycosyl hydrolase 5 (cellulase A) family | cellulose catabolic process [GO:0030245] | | hydrolase activity, hydrolyzing O-glycosyl co | hydrolase activity, hydroly | UP000018817 |
| PPTG_06869 | W2QT07 | Probable threonine--tRNA ligase, cytoplasmic (EC 6.1.1.3) (Threonyl-tRNA synthetase) | 744 | | Class-II aminoacyl-tRNA synthetase family | threonyl-tRNA aminoacylation [GO:000643 | mitochondrion [GO:0005739] | ATP binding [GO:0005524]; threonine-tRNA l | mitochondrion [GO:0005 | UP000018817 |
| PPTG_18357 | W2PJ83 | ABC transporter domain-containing protein | 1308 | | ABC transporter superfamily, ABCG family, PDR (TC 3.A.1.205) subfamily | | membrane [GO:0016020] | ABC-type transporter activity [GO:0140359]; | membrane [GO:0016020]; | UP000018817 |
| PPTG_05552 | W2QXD3 | fructose-bisphosphate aldolase (EC 4.1.2.13) | 358 | | Class II fructose-bisphosphate aldolase family | gluconeogenesis [GO:0006094]; glycolytic p | cytosol [GO:0005829] | fructose-bisphosphate aldolase activity [GO: | cytosol [GO:0005829]; fruc | UP000018817 |
| PPTG_05553 | W2QXP4 | fructose-bisphosphate aldolase (EC 4.1.2.13) | 358 | | Class II fructose-bisphosphate aldolase family | gluconeogenesis [GO:0006094]; glycolytic p | cytosol [GO:0005829] | fructose-bisphosphate aldolase activity [GO: | cytosol [GO:0005829]; fruc | UP000018817 |
| PPTG_02824 | W2RF50 | Choline/carnitine acyltransferase domain-containing protein | 621 | | Carnitine/choline acetyltransferase family | fatty acid beta-oxidation [GO:0006635] | mitochondrion [GO:0005739] | carnitine O-palmitoyltransferase activity [GC | mitochondrion [GO:0005 | UP000018817 |
| PPTG_12292 | W2Q6M6 | Calcineurin-like phosphoesterase domain-containing protein | 479 | | | | | hydrolase activity [GO:0016787] | hydrolase activity [GO:001 | UP000018817 |
| PPTG_10113 | W2QE49 | Importin N-terminal domain-containing protein | 1079 | | | protein import into nucleus [GO:0006606] | cytoplasm [GO:0005737] | small GTPase binding [GO:0031267] | cytoplasm [GO:0005737]; | UP000018817 |
| PPTG_14687 | W2PY60 | Homoserine dehydrogenase (EC 1.1.1.3) | 436 | | Homoserine dehydrogenase family | methionine biosynthetic process [GO:0009086]; threonine biosynthetic process [GO:000 | | homoserine dehydrogenase activity [GO:000 | homoserine dehydrogenas | UP000018817 |
| PPTG_00180 | W2RGD6 | UDP-glucose 6-dehydrogenase (EC 1.1.1.22) | 475 | | UDP-glucose/GDP-mannose dehydrogenase famil | glycosaminoglycan biosynthetic process [G | nucleus [GO:0005634] | NAD binding [GO:0051287]; UDP-glucose 6- | nucleus [GO:0005634]; NA | UP000018817 |
| PPTG_07998 | W2QLC3 | V-type proton ATPase subunit H | 463 | | V-ATPase H subunit family | | vacuolar proton-transporting V-type ATPase, | proton-transporting ATPase activity, rotation | vacuolar proton-transport | UP000018817 |
| PPTG_06649 | W2QSA7 | Nicastrin | 719 | | Nicastrin family | Notch signaling pathway [GO:0007219]; pro | plasma membrane [GO:0005886] | | plasma membrane [GO:0 | UP000018817 |
| PPTG_00284 | W2REE6 | Cytochrome b-c1 complex subunit Rieske, mitochondrial (EC 7.1.1.8) | 215 | | Rieske iron-sulfur protein family | | mitochondrial inner membrane [GO:000574 | 2 iron, 2 sulfur cluster binding [GO:0051537] | mitochondrial inner memb | UP000018817 |
| PPTG_02671 | W2RCG7 | GST C-terminal domain-containing protein | 354 | | | | cytoplasm [GO:0005737] | glutathione transferase activity [GO:000436 | cytoplasm [GO:0005737]; | UP000018817 |
| PPTG_02226 | W2RC14 | Calcium-transporting ATPase (EC 7.2.2.10) | 1067 | | Cation transport ATPase (P-type) (TC 3.A.3) family, | copper ion transport [GO:0006825] | plasma membrane [GO:0005886] | ATP binding [GO:0005524]; ATP hydrolysis ac | plasma membrane [GO:0 | UP000018817 |
| PPTG_11055 | W2Q752 | Glycerol-3-phosphate dehydrogenase (EC 1.1.5.3) | 615 | | FAD-dependent glycerol-3-phosphate dehydrogena | glycerol-3-phosphate metabolic process [G | mitochondrion [GO:0005739] | glycerol-3-phosphate dehydrogenase (quino | mitochondrion [GO:0005 | UP000018817 |
| PPTG_16969 | W2PNS3 | Eukaryotic translation initiation factor 3 subunit M (eIF3m) | 369 | | CSN7/EIF3M family, CSN7 subfamily; EIF-3 subunit | formation of cytoplasmic translation initiati | eukaryotic 43S preinitiation complex [GO:00 | translation initiation factor activity [GO:0003 | eukaryotic 43S preinitiatio | UP000018817 |
| PPTG_02785 | W2REY1 | H(+)-exporting diphosphatase (EC 7.1.3.1) | 768 | | | | endomembrane system [GO:0012505]; mem | diphosphate hydrolysis-driven proton transm | endomembrane system [G | UP000018817 |
| PPTG_13257 | W2Q066 | Serine hydroxymethyltransferase (EC 2.1.2.1) | 502 | | SHMT family | glycine biosynthetic process from serine [G | mitochondrion [GO:0005739] | glycine hydroxymethyltransferase activity [G | mitochondrion [GO:0005 | UP000018817 |
| PPTG_07317 | W2QPK9 | C-CAP/cofactor C-like domain-containing protein | 461 | | CAP family | actin filament organization [GO:0007015]; | cytoplasm [GO:0005737] | actin binding [GO:0003779]; adenylate cycla | cytoplasm [GO:0005737]; | UP000018817 |
| PPTG_06269 | W2QSE4 | Succinate--CoA ligase [ADP-forming] subunit beta, mitochondrial (EC 6.2.1.5) (Succinyl-CoA synth | 437 | | Succinate/malate CoA ligase beta subunit family | succinyl-CoA metabolic process [GO:00061 | mitochondrion [GO:0005739]; succinate-Co | ATP binding [GO:0005524]; magnesium ion | mitochondrion [GO:0005 | UP000018817 |
| PPTG_03926 | W2QZ71 | aspartate--tRNA ligase (EC 6.1.1.12) (Aspartyl-tRNA synthetase) | 514 | | Class-II aminoacyl-tRNA synthetase family, Type 2 | aspartyl-tRNA aminoacylation [GO:000642 | aminoacyl-tRNA synthetase multienzyme co | aspartate-tRNA ligase activity [GO:0004815] | aminoacyl-tRNA synthetas | UP000018817 |
| PPTG_17667 | W2PIK2 | Glycoside hydrolase family 19 catalytic domain-containing protein | 276 | | | cell wall macromolecule catabolic process [GO:0016998]; chitin catabolic process [GO:0 | | chitinase activity [GO:0004568] | chitinase activity [GO:000 | UP000018817 |
| PPTG_03931 | W2QYV6 | Annexin | 690 | Vinay and Belleannée, 2022 | Annexin family | | cytoplasm [GO:0005737]; plasma membrane | calcium ion binding [GO:0005509]; calcium- | cytoplasm [GO:0005737]; | UP000018817 |
| PPTG_04112 | W2QZ92 | Superoxide dismutase (EC 1.15.1.1) | 250 | | Iron/manganese superoxide dismutase family | | cytoplasm [GO:0005737] | metal ion binding [GO:0046872]; superoxide | cytoplasm [GO:0005737]; | UP000018817 |
| PPTG_15407 | W2PVJ8 | Serine protease | 685 | | | | | | | UP000018817 |
| PPTG_16072 | W2PQA7 | 60S ribosomal protein L6 | 204 | | Eukaryotic ribosomal protein eL6 family | cytoplasmic translation [GO:0002181]; ribo | chloroplast [GO:0009507]; cytosolic large rib | RNA binding [GO:0003723]; structural consti | chloroplast [GO:0009507]; | UP000018817 |
| PPTG_12608 | W2Q1R2 | CSC1/OSCA1-like 7TM region domain-containing protein | 845 | | CSC1 (TC 1.A.17) family | | plasma membrane [GO:0005886] | calcium-activated cation channel activity [G | plasma membrane [GO:0 | UP000018817 |
| PPTG_18605 | W2PFJ7 | YHYH domain-containing protein | 5468 | | | | | | | UP000018817 |
| PPTG_02400 | W2RB59 | protein disulfide-isomerase (EC 5.3.4.1) | 353 | | Protein disulfide isomerase family | protein folding [GO:0006457] | endoplasmic reticulum [GO:0005783] | protein disulfide isomerase activity [GO:000 | endoplasmic reticulum [G | UP000018817 |
| PPTG_08136 | W2QK24 | dolichyl-diphosphooligosaccharide--protein glycotransferase (EC 2.4.99.18) | 885 | | STT3 family | | endomembrane system [GO:0012505]; mem | dolichyl-diphosphooligosaccharide-protein g | endomembrane system [G | UP000018817 |
| PPTG_09870 | W2QD77 | Uncharacterized protein | 2873 | | VPS13 family | protein retention in Golgi apparatus [GO:0045053]; protein targeting to vacuole [GO:0006 | | calcium ion binding [GO:0005509] | calcium ion binding [GO:0 | UP000018817 |
| PPTG_16844 | W2PMJ0 | 2,4-dienoyl-CoA reductase | 322 | | | fatty acid beta-oxidation [GO:0006635] | mitochondrion [GO:0005739] | 2,4-dienoyl-CoA reductase (NADPH) activity | mitochondrion [GO:0005 | UP000018817 |
| PPTG_00768 | W2RIN1 | Coronin | 507 | | WD repeat coronin family | actin filament organization [GO:0007015] | | actin filament binding [GO:0051015] | actin filament binding [GO | UP000018817 |
| PPTG_19399 | W2PDE7 | Peptidyl-prolyl cis-trans isomerase (PPIase) (EC 5.2.1.8) | 171 | | Cyclophilin-type PPIase family | protein folding [GO:0006457] | cytoplasm [GO:0005737] | cyclosporin A binding [GO:0016018]; peptidy | cytoplasm [GO:0005737]; | UP000018817 |
| PPTG_13453 | W2Q1M2 | Guanine nucleotide-binding protein alpha-1 subunit | 355 | | | adenylate cyclase-modulating G protein-co | cytoplasm [GO:0005737]; heterotrimeric G-p | G protein-coupled receptor binding [GO:000 | cytoplasm [GO:0005737]; | UP000018817 |
| PPTG_11336 | W2QB00 | AP complex subunit beta | 837 | | Adaptor complexes large subunit family | intracellular protein transport [GO:0006886 | clathrin adaptor complex [GO:0030131]; end | clathrin binding [GO:0030276] | clathrin adaptor complex [ | UP000018817 |
| PPTG_05088 | W2QVT6 | 5-oxoprolinase | 1292 | | Oxoprolinase family | glutathione metabolic process [GO:000674 | cytosol [GO:0005829] | 5-oxoprolinase (ATP-hydrolyzing) activity [GC | cytosol [GO:0005829]; 5-o | UP000018817 |
| PPTG_15134 | W2PW69 | 40S ribosomal protein S12 | 175 | | Eukaryotic ribosomal protein eS12 family | translation [GO:0006412] | ribonucleoprotein complex [GO:1990904]; rit | structural constituent of ribosome [GO:0003 | ribonucleoprotein comple | UP000018817 |
| PPTG_14749 | W2PYE3 | serine--tRNA ligase (EC 6.1.1.11) (Seryl-tRNA synthetase) | 454 | | Class-II aminoacyl-tRNA synthetase family, Type-1 | seryl-tRNA aminoacylation [GO:0006434] | | ATP binding [GO:0005524]; serine-tRNA liga | ATP binding [GO:0005524] | UP000018817 |
| PPTG_05429 | W2QX24 | cAMP-dependent protein kinase regulatory subunit | 290 | | CAMP-dependent kinase regulatory chain family | | cAMP-dependent protein kinase complex [G | cAMP binding [GO:0030552]; cAMP-depende | cAMP-dependent protein k | UP000018817 |
| PPTG_02700 | W2RCJ5 | CN hydrolase domain-containing protein | 323 | | Carbon-nitrogen hydrolase superfamily, NIT1/NIT2 | asparagine metabolic process [GO:000652 | mitochondrion [GO:0005739] | omega-amidase activity [GO:0050152] | mitochondrion [GO:0005 | UP000018817 |
| PPTG_04052 | W2QZP3 | Trifunctional purine biosynthetic protein adenosine-3 (EC 2.1.2.2) (EC 6.3.3.1) (EC 6.3.4.13) | 1143 | | GART family; GARS family; AIR synthase family | 'de novo' IMP biosynthetic process [GO:000 | cytosol [GO:0005829] | ATP binding [GO:0005524]; IMP cyclohydrola | cytosol [GO:0005829]; ATF | UP000018817 |
| PPTG_12075 | W2Q4W4 | USP domain-containing protein | 2913 | | | protein deubiquitination [GO:0016579]; pro | cytosol [GO:0005829]; nucleus [GO:0005634 | cysteine-type deubiquitinase activity [GO:00 | cytosol [GO:0005829]; nuc | UP000018817 |
| PPTG_16811 | W2PQ87 | glucose-6-phosphate 1-epimerase (EC 5.1.3.15) | 300 | | Glucose-6-phosphate 1-epimerase family | carbohydrate metabolic process [GO:0005 | cytoplasm [GO:0005737] | carbohydrate binding [GO:0030246]; glucose | cytoplasm [GO:0005737]; | UP000018817 |
| PPTG_16755 | W2PPB1 | Uncharacterized protein | 1034 | | | | | actin binding [GO:0003779]; calcium ion bin | actin binding [GO:000377 | UP000018817 |
| PPTG_19423 | W2PCK8 | Calmodulin | 575 | | | endocytosis [GO:0006897]; endosomal tra | plasma membrane [GO:0005886]; recycling | calcium ion binding [GO:0005509]; GTP bind | plasma membrane [GO:0 | UP000018817 |
| PPTG_01358 | W2R6Z9 | Glyceraldehyde-3-phosphate dehydrogenase, type I | 524 | | Glyceraldehyde-3-phosphate dehydrogenase family | | | NAD binding [GO:0051287]; oxidoreductase | NAD binding [GO:0051287 | UP000018817 |
| PPTG_02403 | W2RCS4 | Pre-mRNA-processing-splicing factor 8 | 2359 | | | spliceosomal tri-snRNP complex assembly | catalytic step 2 spliceosome [GO:0071013]; | metallopeptidase activity [GO:0008237]; pre | catalytic step 2 spliceoson | UP000018817 |
| PPTG_08466 | W2QLE9 | Tubulin alpha chain | 452 | | Tubulin family | microtubule-based process [GO:0007017] | microtubule [GO:0005874] | GTP binding [GO:0005525]; hydrolase activit | microtubule [GO:0005874 | UP000018817 |
| PPTG_00417 | W2RFD5 | Phospholipid-transporting ATPase (EC 7.6.2.1) | 1324 | | Cation transport ATPase (P-type) (TC 3.A.3) family, | phospholipid translocation [GO:0045332] | endomembrane system [GO:0012505]; plasn | ATP binding [GO:0005524]; ATP hydrolysis ac | endomembrane system [G | UP000018817 |
| PPTG_02851 | W2RCR9 | E1 ubiquitin-activating enzyme (EC 6.2.1.45) | 1035 | | Ubiquitin-activating E1 family | protein sumoylation [GO:0016925] | cytoplasm [GO:0005737]; SUMO activating e | ATP binding [GO:0005524]; SUMO activating | cytoplasm [GO:0005737]; | UP000018817 |
| PPTG_12936 | W2Q187 | Protein transport protein SEC23 | 861 | | SEC23/SEC24 family, SEC23 subfamily | COPII-coated vesicle cargo loading [GO:00 | COPII vesicle coat [GO:0030127]; endoplasm | GTPase activator activity [GO:0005096]; zinc | COPII vesicle coat [GO:00 | UP000018817 |
| PPTG_05975 | W2QWP6 | Carbonic anhydrase (EC 4.2.1.1) | 272 | | Alpha-carbonic anhydrase family | | | carbonate dehydratase activity [GO:000408 | carbonate dehydratase ac | UP000018817 |
| PPTG_02370 | W2RAI7 | glucan endo-1,3-beta-D-glucosidase (EC 3.2.1.39) | 750 | | Glycosyl hydrolase 81 family | cell wall organization [GO:0071555]; polysaccharide catabolic process [GO:0000272] | | endo-1,3(4)-beta-glucanase activity [GO:00 | endo-1,3(4)-beta-glucanas | UP000018817 |
| PPTG_18566 | W2PH71 | Cytoplasmic dynein 2 heavy chain 1 | 4396 | | Dynein heavy chain family | cilium assembly [GO:0060271]; intracellula | ciliary membrane [GO:0060170]; dynein com | ATP binding [GO:0005524]; dynein intermedi | ciliary membrane [GO:006 | UP000018817 |
| PPTG_06281 | W2QSV9 | Propionyl-CoA carboxylase alpha chain, mitochondrial (EC 6.4.1.3) (Propanoyl-CoA:carbon dioxide | 764 | | | lipid catabolic process [GO:0016042] | mitochondrial matrix [GO:0005759] | ATP binding [GO:0005524]; metal ion binding | mitochondrial matrix [GO: | UP000018817 |
| PPTG_09222 | W2QGU1 | CAMK/CAMK1 protein kinase | 385 | | Protein kinase superfamily, Ser/Thr protein kinase | glycogen metabolic process [GO:0005977] | phosphorylase kinase complex [GO:0005964 | ATP binding [GO:0005524]; calmodulin bindi | phosphorylase kinase com | UP000018817 |

| | | | | | | | | | | |
|---|---|---|---|---|---|---|---|---|---|---|
| PPTG_08673 | W2QLJ3 | SEC7 domain-containing protein | 2046 | | | protein transport [GO:0015031]; regulation | cytoplasm [GO:0005737]; membrane [GO:00 | guanyl-nucleotide exchange factor activity [G | cytoplasm [GO:0005737]; | UP000018817 |
| PPTG_00390 | W2REP0 | FAD/NAD(P)-binding domain-containing protein | 559 | | FAD-dependent oxidoreductase family | mitochondrial respiratory chain complex as | mitochondrion [GO:0005739] | FAD binding [GO:0071949]; NAD(P)H oxidase | mitochondrion [GO:00057 | UP000018817 |
| PPTG_06192 | W2QUI4 | 40S ribosomal protein S3 | 246 | | Universal ribosomal protein uS3 family | translation [GO:0006412] | cytosolic small ribosomal subunit [GO:00226 | RNA binding [GO:0003723]; structural consti | cytosolic small ribosomal s | UP000018817 |
| PPTG_10783 | W2QBT9 | Jacalin-type lectin domain-containing protein | 1075 | | | | | | | UP000018817 |
| PPTG_18587 | W2PHU9 | cellulose 1,4-beta-cellobiosidase (non-reducing end) (EC 3.2.1.91) | 521 | | Glycosyl hydrolase 7 (cellulase C) family | cellulose catabolic process [GO:0030245] | | cellulose 1,4-beta-cellobiosidase activity [G | cellulose 1,4-beta-cellobic | UP000018817 |
| PPTG_16769 | W2PNB6 | EGF-like domain-containing protein | 1060 | Ral et al., 2024 | | | | | | UP000018817 |
| PPTG_02259 | W2RAA1 | protein-synthesizing GTPase (EC 3.6.5.3) | 462 | | TRAFAC class translation factor GTPase superfamil | formation of translation preinitiation compl | cytosol [GO:0005829]; eukaryotic translation | GTP binding [GO:0005525]; GTPase activity [ | cytosol [GO:0005829]; euk | UP000018817 |
| PPTG_08401 | W2QKH4 | PCI domain-containing protein | 264 | | Proteasome subunit S14 family | proteasome-mediated ubiquitin-dependent | cytosol [GO:0005829]; nucleus [GO:0005634]; proteasome regulatory particle, lid subcom | | cytosol [GO:0005829]; nuc | UP000018817 |
| PPTG_12440 | W2Q6M8 | Uncharacterized protein | 329 | | | | | oxidoreductase activity [GO:0016491] | oxidoreductase activity [G | UP000018817 |
| PPTG_00500 | W2RH96 | Electron transfer flavoprotein subunit beta (Beta-ETF) | 251 | | ETF beta-subunit/FixA family | carboxylic acid catabolic process [GO:0046 | mitochondrial matrix [GO:0005759] | electron transfer activity [GO:0009055] | mitochondrial matrix [GO: | UP000018817 |
| PPTG_13924 | W2Q0S7 | Prolyl endopeptidase (EC 3.4.21.-) | 752 | | Peptidase S9A family | proteolysis [GO:0006508] | cytosol [GO:0005829] | oligopeptidase activity [GO:0070012]; serine | cytosol [GO:0005829]; olig | UP000018817 |
| PPTG_09717 | W2QGS2 | ABC transmembrane type-1 domain-containing protein | 1011 | | | | membrane [GO:0016020] | ABC-type transporter activity [GO:0140359]; | membrane [GO:0016020]; | UP000018817 |
| PPTG_07447 | W2QPG4 | Uncharacterized protein | 930 | | Argonaute family | | | RNA binding [GO:0003723] | RNA binding [GO:0003723 | UP000018817 |
| PPTG_06246 | W2QSA5 | TKL protein kinase | 674 | | | | | ATP binding [GO:0005524]; protein serine/th | ATP binding [GO:0005524] | UP000018817 |
| PPTG_14424 | W2PX30 | Large ribosomal subunit protein uL6 alpha-beta domain-containing protein | 189 | | Universal ribosomal protein uL6 family | cytoplasmic translation [GO:0002181] | cytosolic large ribosomal subunit [GO:00226 | rRNA binding [GO:0019843]; structural const | cytosolic large ribosomal s | UP000018817 |
| PPTG_02597 | W2RBQ4 | Importin N-terminal domain-containing protein | 858 | | Importin beta family, Importin beta-1 subfamily | protein import into nucleus [GO:0006606] | cytoplasm [GO:0005737] | small GTPase binding [GO:0031267] | cytoplasm [GO:0005737]; | UP000018817 |
| PPTG_06597 | W2QQ58 | Uncharacterized protein | 381 | | | | | | | UP000018817 |
| PPTG_14738 | W2PYD0 | MIT domain-containing protein | 135 | | | | | | | UP000018817 |
| PPTG_18479 | W2PGF2 | 6-phosphogluconolactonase | 360 | | Cycloisomerase 2 family | | | 6-phosphogluconolactonase activity [GO:00 | 6-phosphogluconolactona | UP000018817 |
| PPTG_10646 | W2QBJ2 | 40S ribosomal protein S4 | 261 | | Eukaryotic ribosomal protein eS4 family | translation [GO:0006412] | cytosolic small ribosomal subunit [GO:00226 | rRNA binding [GO:0019843]; structural const | cytosolic small ribosomal s | UP000018817 |
| PPTG_19949 | W2PAP8 | subtilisin (EC 3.4.21.62) | 487 | | Peptidase S8 family | proteolysis [GO:0006508] | | serine-type endopeptidase activity [GO:0004 | serine-type endopeptidase | UP000018817 |
| PPTG_16088 | W2PS91 | Pyruvate dehydrogenase E1 component subunit alpha (EC 1.2.4.1) | 402 | | | pyruvate decarboxylation to acetyl-CoA [GO:0006086] | | pyruvate dehydrogenase (acetyl-transferring | pyruvate dehydrogenase (a | UP000018817 |
| PPTG_13499 | W2Q1L6 | 60S ribosomal protein L6 | 233 | | Eukaryotic ribosomal protein eL6 family | cytoplasmic translation [GO:0002181]; ribo | chloroplast [GO:0009507]; cytosolic large rib | RNA binding [GO:0003723]; structural consti | chloroplast [GO:0009507]; | UP000018817 |
| PPTG_11926 | W2Q8U3 | C2 NT-type domain-containing protein | 794 | | | | | | | UP000018817 |
| PPTG_00140 | W2RFV5 | Glycoside hydrolase | 460 | | | cellulose catabolic process [GO:0030245] | | hydrolase activity, hydrolyzing O-glycosyl cor | hydrolase activity, hydrolyz | UP000018817 |
| PPTG_09673 | W2QFQ0 | Uncharacterized protein | 342 | | | | | phosphatidylinositol binding [GO:0035091] | phosphatidylinositol bindir | UP000018817 |
| PPTG_13069 | W2Q664 | MARVEL domain-containing protein | 357 | Breen et al., 2025 | | | membrane [GO:0016020] | | membrane [GO:0016020] | UP000018817 |
| PPTG_11437 | W2Q9L5 | malate dehydrogenase (EC 1.1.1.37) | 336 | | LDH/MDH superfamily, MDH type 2 family | malate metabolic process [GO:0006108] | | L-malate dehydrogenase (NAD+) activity [GC | L-malate dehydrogenase ( | UP000018817 |
| PPTG_04024 | W2QYX2 | CAMK/CDPK protein kinase | 555 | | Protein kinase superfamily, Ser/Thr protein kinase family, CDPK subfamily | | | ATP binding [GO:0005524]; calcium ion bindi | ATP binding [GO:0005524] | UP000018817 |
| PPTG_09495 | W2QF83 | Glutamate dehydrogenase | 494 | | Glu/Leu/Phe/Val dehydrogenases family | glutamate biosynthetic process [GO:00065 | cytosol [GO:0005829] | glutamate dehydrogenase (NADP+) activity [ | cytosol [GO:0005829]; glut | UP000018817 |
| PPTG_02716 | W2RC68 | ribose-phosphate diphosphokinase (EC 2.7.6.1) | 475 | | Ribose-phosphate pyrophosphokinase family | 5-phosphoribose 1-diphosphate biosyntheti | cytoplasm [GO:0005737]; ribose phosphate c | ATP binding [GO:0005524]; kinase activity [G | cytoplasm [GO:0005737]; | UP000018817 |
| PPTG_03875 | W2QYB8 | EF-hand domain-containing protein | 2895 | | VPS13 family | protein retention in Golgi apparatus [GO:0045053]; protein targeting to vacuole [GO:0006 | | calcium ion binding [GO:0005509] | calcium ion binding [GO:0 | UP000018817 |
| PPTG_10537 | W2QC18 | Phospholipid-transporting ATPase (EC 7.6.2.1) | 1291 | | Cation transport ATPase (P-type) (TC 3.A.3) family, | phospholipid translocation [GO:0045332] | plasma membrane [GO:0005886] | ATP binding [GO:0005524]; ATP hydrolysis ac | plasma membrane [GO:00 | UP000018817 |
| PPTG_00340 | W2REP6 | Dynein gamma chain, flagellar outer arm | 4622 | | Dynein heavy chain family | microtubule-based movement [GO:00701 | axonemal dynein complex [GO:0005858]; mi | ATP binding [GO:0005524]; dynein intermedi | axonemal dynein complex | UP000018817 |
| PPTG_03645 | W2R7M2 | Phosphoenolpyruvate carboxykinase (ATP) | 379 | | | gluconeogenesis [GO:0006094] | cytosol [GO:0005829] | ATP binding [GO:0005524]; phosphoenolpyru | cytosol [GO:0005829]; AT | UP000018817 |
| PPTG_17319 | W2PJ20 | EIF3d | 551 | | | | eukaryotic translation initiation factor 3 com | RNA binding [GO:0003723]; translation initia | eukaryotic translation initia | UP000018817 |
| PPTG_12256 | W2Q886 | Protein transporter Sec24 | 1119 | | SEC23/SEC24 family, SEC24 subfamily | COPII-coated vesicle cargo loading [GO:009 | COPII vesicle coat [GO:0030127]; endoplasm | SNARE binding [GO:0000149]; zinc ion bindir | COPII vesicle coat [GO:00 | UP000018817 |
| PPTG_11360 | W2Q9A0 | SMP-30/Gluconolactonase/LRE-like region domain-containing protein | 690 | | | | | hydrolase activity [GO:0016787] | hydrolase activity [GO:001 | UP000018817 |
| PPTG_08833 | W2QJT0 | EGF-like domain-containing protein | 679 | | | | | | | UP000018817 |
| PPTG_15911 | W2PRV1 | Calnexin | 557 | | Calreticulin family | ERAD pathway [GO:0036503]; protein foldir | endoplasmic reticulum membrane [GO:0005 | calcium ion binding [GO:0005509]; unfolded | endoplasmic reticulum me | UP000018817 |
| PPTG_19077 | W2PGG5 | Small ribosomal subunit protein uS2 | 282 | | Universal ribosomal protein uS2 family | ribosomal small subunit assembly [GO:000 | cytosolic small ribosomal subunit [GO:00226 | structural constituent of ribosome [GO:0003 | cytosolic small ribosomal s | UP000018817 |
| PPTG_07737 | W2QLK1 | CCT-beta | 526 | | TCP-1 chaperonin family | | chaperonin-containing T-complex [GO:00058 | ATP binding [GO:0005524]; ATP hydrolysis ac | chaperonin-containing T-c | UP000018817 |
| PPTG_13082 | W2Q6B1 | NADH dehydrogenase [ubiquinone] flavoprotein 2, mitochondrial | 271 | | Complex I 24 kDa subunit family | mitochondrial electron transport, NADH to u | catalytic complex [GO:1902494]; membrane | 2 iron, 2 sulfur cluster binding [GO:0051537] | catalytic complex [GO:190 | UP000018817 |
| PPTG_15631 | W2PTN3 | Voltage-dependent anion-selective channel protein | 282 | | Eukaryotic mitochondrial porin family | | mitochondrial outer membrane [GO:000574 | porin activity [GO:0015288]; voltage-gated m | mitochondrial outer memb | UP000018817 |
| PPTG_12116 | W2Q6X8 | V-type proton ATPase subunit | 394 | | V-ATPase V0D/AC39 subunit family | | proton-transporting V-type ATPase, V0 domai | proton-transporting ATPase activity, rotation | proton-transporting V-type | UP000018817 |
| PPTG_00991 | W2RHE9 | dihydroxy-acid dehydratase (EC 4.2.1.9) | 596 | | IlvD/Edd family | isoleucine biosynthetic process [GO:0009097]; L-valine biosynthetic process [GO:00090 | | 2 iron, 2 sulfur cluster binding [GO:0051537] | 2 iron, 2 sulfur cluster bind | UP000018817 |
| PPTG_10595 | W2QBA2 | ABC transporter domain-containing protein | 1354 | | ABC transporter superfamily, ABCG family, PDR (TC 3.A.1.205) subfamily | | membrane [GO:0016020] | ABC-type transporter activity [GO:0140359]; | membrane [GO:0016020]; | UP000018817 |
| PPTG_13968 | W2Q0Y9 | Ubiquinone biosynthesis monooxygenase COQ6, mitochondrial (EC 1.14.15.45) (2-methoxy-6-poly | 482 | | UbiH/COQ6 family | | extrinsic component of mitochondrial inner m | 2-methoxy-6-polyprenolphenol 4-hydroxylase | extrinsic component of mi | UP000018817 |
| PPTG_08403 | W2QL58 | Xylose isomerase (EC 5.3.1.5) | 452 | | Xylose isomerase family | D-xylose metabolic process [GO:0042732] | | metal ion binding [GO:0046872]; xylose isom | metal ion binding [GO:004 | UP000018817 |
| PPTG_18230 | W2PJ23 | T-complex protein 1 subunit delta | 533 | | TCP-1 chaperonin family | | cytoplasm [GO:0005737] | ATP binding [GO:0005524]; ATP hydrolysis ac | cytoplasm [GO:0005737]; | UP000018817 |
| PPTG_03709 | W2R8F7 | Pyruvate kinase (EC 2.7.1.40) | 505 | | Pyruvate kinase family | response to stress [GO:0006950] | | ATP binding [GO:0005524]; kinase activity [G | ATP binding [GO:0005524] | UP000018817 |
| PPTG_16236 | W2PPS8 | Transglutaminase-like domain-containing protein | 530 | | | | | aminoacyltransferase activity [GO:0016755] | aminoacyltransferase acti | UP000018817 |
| PPTG_02106 | W2RBI2 | Xaa-Pro dipeptidyl-peptidase C-terminal domain-containing protein | 740 | | | | | dipeptidyl-peptidase activity [GO:0008239] | dipeptidyl-peptidase activi | UP000018817 |
| PPTG_06377 | W2QSL7 | Aminopeptidase (EC 3.4.11.-) | 902 | | Peptidase M1 family | peptide catabolic process [GO:0043171]; pr | cytoplasm [GO:0005737]; extracellular space | metalloaminopeptidase activity [GO:007000 | cytoplasm [GO:0005737]; | UP000018817 |
| PPTG_11378 | W2QB69 | Eukaryotic translation initiation factor 3 subunit L (eIF3l) | 490 | | EIF-3 subunit L family | formation of cytoplasmic translation initiati | eukaryotic 43S preinitiation complex [GO:00 | translation initiation factor activity [GO:0003 | eukaryotic 43S preinitiatio | UP000018817 |
| PPTG_02894 | W2RFJ7 | Probable enoyl-CoA hydratase, mitochondrial (EC 4.2.1.17) | 279 | | Enoyl-CoA hydratase/isomerase family | fatty acid beta-oxidation [GO:0006635] | mitochondrion [GO:0005739] | enoyl-CoA hydratase activity [GO:0004300] | mitochondrion [GO:00057 | UP000018817 |
| PPTG_03669 | W2R5S3 | 40S ribosomal protein | 196 | | Universal ribosomal protein uS7 family | translation [GO:0006412] | small ribosomal subunit [GO:0015935] | RNA binding [GO:0003723]; structural consti | small ribosomal subunit [G | UP000018817 |
| PPTG_06725 | W2QQI2 | ABC transporter domain-containing protein | 1349 | | ABC transporter superfamily, ABCG family, PDR (TC 3.A.1.205) subfamily | | membrane [GO:0016020] | ABC-type transporter activity [GO:0140359]; | membrane [GO:0016020]; | UP000018817 |
| PPTG_04949 | W2R387 | Pyrroline-5-carboxylate reductase catalytic N-terminal domain-containing protein | 272 | | Pyrroline-5-carboxylate reductase family | L-proline biosynthetic process [GO:0055129] | | pyrroline-5-carboxylate reductase activity [G | pyrroline-5-carboxylate rec | UP000018817 |
| PPTG_17401 | W2PJW0 | Uncharacterized protein | 416 | | | | | | | UP000018817 |
| PPTG_18213 | W2PJ01 | Importin N-terminal domain-containing protein | 1076 | | Exportin family | protein export from nucleus [GO:0006611]; | cytoplasm [GO:0005737]; nucleus [GO:0005 | nuclear export signal receptor activity [GO:0 | cytoplasm [GO:0005737]; | UP000018817 |
| PPTG_07817 | W2QPI2 | Uncharacterized protein | 506 | | | | | | | UP000018817 |
| PPTG_18731 | W2PF41 | Poly [ADP-ribose] polymerase (PARP) (EC 2.4.2.-) | 2885 | | | | | NAD+ poly-ADP-ribosyltransferase activity [G | NAD+ poly-ADP-ribosyltran | UP000018817 |
| PPTG_07085 | W2QP05 | 40S ribosomal protein S17 | 131 | | Eukaryotic ribosomal protein eS17 family | translation [GO:0006412] | cytosol [GO:0005829]; ribonucleoprotein con | structural constituent of ribosome [GO:0003 | cytosol [GO:0005829]; ribc | UP000018817 |
| PPTG_11500 | W2QCD3 | FYVE-type domain-containing protein | 558 | | | | | | | UP000018817 |
| PPTG_02443 | W2RDA0 | Uncharacterized protein | 801 | | | proteolysis [GO:0006508] | | dipeptidyl-peptidase activity [GO:0008239]; | dipeptidyl-peptidase activi | UP000018817 |
| PPTG_15878 | W2PSD0 | Ribonucleoside-diphosphate reductase (EC 1.17.4.1) | 792 | | Ribonucleoside diphosphate reductase large chain | deoxyribonucleotide biosynthetic process [G | ribonucleoside-diphosphate reductase comp | ATP binding [GO:0005524]; ribonucleoside-d | ribonucleoside-diphospha | UP000018817 |
| PPTG_11894 | W2QB77 | 26S protease regulatory subunit 7 | 438 | | AAA ATPase family | proteolysis [GO:0006508] | cytoplasm [GO:0005737]; nucleus [GO:0005 | ATP binding [GO:0005524]; ATP hydrolysis ac | cytoplasm [GO:0005737]; | UP000018817 |
| PPTG_16668 | W2PPP5 | PCI domain-containing protein | 383 | | Proteasome subunit S11 family | ubiquitin-dependent protein catabolic proc | cytosol [GO:0005829]; nucleus [GO:0005634 | structural molecule activity [GO:0005198] | cytosol [GO:0005829]; nuc | UP000018817 |
| PPTG_00631 | W2RI00 | Chaperone DnaJ | 418 | | | protein folding [GO:0006457]; response to heat [GO:0009408] | | ATP binding [GO:0005524]; Hsp70 protein bi | ATP binding [GO:0005524] | UP000018817 |
| PPTG_06749 | W2QQK3 | Uncharacterized protein | 341 | | RNase T2 family | RNA catabolic process [GO:0006401] | extracellular region [GO:0005576] | ribonuclease T2 activity [GO:0033897]; RNA | extracellular region [GO:0 | UP000018817 |
| PPTG_12038 | W2Q6J8 | Serine/threonine-protein phosphatase (EC 3.1.3.16) | 319 | | PPP phosphatase family | | | metal ion binding [GO:0046872]; protein seri | metal ion binding [GO:004 | UP000018817 |
| PPTG_05659 | W2QVF4 | Apple domain-containing protein | 781 | | | proteolysis [GO:0006508] | extracellular region [GO:0005576] | flavin adenine dinucleotide binding [GO:005 | extracellular region [GO:0 | UP000018817 |
| PPTG_12822 | W2Q0U7 | Uncharacterized protein | 456 | | | | | | | UP000018817 |
| PPTG_13120 | W2Q4N9 | Annexin | 329 | Vinay and Belleannée, 2022 | Annexin family | | cytoplasm [GO:0005737]; plasma membrane | calcium ion binding [GO:0005509]; calcium- | cytoplasm [GO:0005737]; | UP000018817 |
| PPTG_02295 | W2RAT4 | MPN domain-containing protein | 324 | | Peptidase M67A family | proteasome-mediated ubiquitin-dependent | proteasome regulatory particle [GO:0005838 | metallopeptidase activity [GO:0008237] | proteasome regulatory par | UP000018817 |
| PPTG_01302 | W2R6B5 | Uncharacterized protein | 512 | | | | | | | UP000018817 |
| PPTG_07760 | W2QLN1 | Phosphoacetylglucosamine mutase (PAGM) (EC 5.4.2.3) (Acetylglucosamine phosphomutase) (N-a | 561 | | Phosphohexose mutase family | carbohydrate metabolic process [GO:0005975]; UDP-N-acetylglucosamine biosynthetic p | | metal ion binding [GO:0046872]; phosphoac | metal ion binding [GO:004 | UP000018817 |
| PPTG_05022 | W2QVI7 | Aspartate aminotransferase (EC 2.6.1.1) | 426 | | Class-I pyridoxal-phosphate-dependent aminotran | amino acid metabolic process [GO:000652 | mitochondrion [GO:0005739] | L-aspartate:2-oxoglutarate aminotransferase | mitochondrion [GO:00057 | UP000018817 |
| PPTG_09874 | W2QD82 | Glutamine synthetase (EC 6.3.1.2) | 402 | | Glutamine synthetase family | glutamine biosynthetic process [GO:00065 | cytoplasm [GO:0005737] | ATP binding [GO:0005524]; glutamine synthe | cytoplasm [GO:0005737]; | UP000018817 |
| PPTG_02839 | W2RCP4 | Replication protein A subunit | 628 | | Replication factor A protein 1 family | DNA recombination [GO:0006310]; DNA rep | nucleus [GO:0005634] | DNA binding [GO:0003677]; zinc ion binding | nucleus [GO:0005634]; DN | UP000018817 |
| PPTG_00924 | W2RJB4 | glycine--tRNA ligase (EC 6.1.1.14) (Diadenosine tetraphosphate synthetase) | 681 | | Class-II aminoacyl-tRNA synthetase family | mitochondrial glycyl-tRNA aminoacylation [ | mitochondrion [GO:0005739] | ATP binding [GO:0005524]; glycine-tRNA liga | mitochondrion [GO:00057 | UP000018817 |
| PPTG_16233 | W2PP25 | Transglutaminase elicitor | 910 | | | regulation of growth [GO:0040008] | | aminoacyltransferase activity [GO:0016755] | aminoacyltransferase acti | UP000018817 |
| PPTG_18330 | W2PII3 | Nucleosome assembly protein 1-like 1 | 368 | | Nucleosome assembly protein (NAP) family | nucleosome assembly [GO:0006334] | nucleus [GO:0005634] | | nucleus [GO:0005634]; nu | UP000018817 |
| PPTG_08378 | W2QKC6 | PCI domain-containing protein | 416 | | | proteasome-mediated ubiquitin-dependent | proteasome complex [GO:0000502] | | proteasome complex [GO: | UP000018817 |
| PPTG_12430 | W2Q5T7 | DUSP domain-containing protein | 637 | | | | | cysteine-type deubiquitinase activity [GO:00 | cysteine-type deubiquitina | UP000018817 |
| PPTG_05337 | W2QZ60 | Glutamate decarboxylase (EC 4.1.1.15) | 493 | | Group II decarboxylase family | L-glutamate catabolic process [GO:00065 | cytosol [GO:0005829] | glutamate decarboxylase activity [GO:00043 | cytosol [GO:0005829]; glut | UP000018817 |
| PPTG_03719 | W2R8G6 | Myosin motor domain-containing protein | 1535 | | TRAFAC class myosin-kinesin ATPase superfamily, | actin filament organization [GO:0007015] | cytoplasm [GO:0005737]; membrane [GO:00 | actin filament binding [GO:0051015]; ATP bi | cytoplasm [GO:0005737]; | UP000018817 |
| PPTG_15739 | W2PRG8 | S5 DRBM domain-containing protein | 312 | | Universal ribosomal protein uS5 family | translation [GO:0006412] | ribonucleoprotein complex [GO:1990904]; rib | RNA binding [GO:0003723]; structural consti | ribonucleoprotein complex | UP000018817 |
| PPTG_08167 | W2QJJ1 | thioredoxin-dependent peroxiredoxin (EC 1.11.1.24) (Nuclear thiol peroxidase) (Thioredoxin peroxi | 367 | | Peroxiredoxin family, BCP/PrxQ subfamily | cell redox homeostasis [GO:0045454]; cellu | cytoplasm [GO:0005737]; nucleus [GO:0005 | thioredoxin peroxidase activity [GO:0008379 | cytoplasm [GO:0005737]; | UP000018817 |
| PPTG_03886 | W2R0F4 | Proteasome subunit alpha type | 244 | | Peptidase T1A family | proteasome-mediated ubiquitin-dependent | cytosol [GO:0005829]; nucleus [GO:0005634]; proteasome core complex, alpha-subunit c | | cytosol [GO:0005829]; nuc | UP000018817 |
| PPTG_05574 | W2QY66 | Band 7 domain-containing protein | 376 | | Band 7/mec-2 family | mitochondrion organization [GO:0007005] | membrane [GO:0016020]; mitochondrion [GO:0005739] | | membrane [GO:0016020]; | UP000018817 |
| PPTG_16051 | W2PPW8 | Anaphase-promoting complex subunit 4 WD40 domain-containing protein | 1507 | | | intraciliary retrograde transport [GO:00357 | axoneme [GO:0005930]; ciliary basal body [GO:0036064]; intraciliary transport particle A [ | | axoneme [GO:0005930]; c | UP000018817 |
| PPTG_05308 | W2QYJ3 | T-complex protein 1 subunit alpha (CCT-alpha) | 546 | | TCP-1 chaperonin family | | cytoplasm [GO:0005737] | ATP binding [GO:0005524]; ATP hydrolysis ac | cytoplasm [GO:0005737]; | UP000018817 |
| PPTG_05779 | W2QUG7 | Acyl-CoA dehydrogenase family member 11 | 801 | | Acyl-CoA dehydrogenase family | fatty acid beta-oxidation using acyl-CoA deh | mitochondrial membrane [GO:0031966]; per | acyl-CoA dehydrogenase activity [GO:00039 | mitochondrial membrane | UP000018817 |
| PPTG_04484 | W2R3J2 | 14-3-3-like protein | 249 | | 14-3-3 family | | | | | UP000018817 |
| PPTG_14511 | W2PWR6 | Importin N-terminal domain-containing protein | 1129 | | | protein import into nucleus [GO:0006606] | cytoplasm [GO:0005737]; nucleus [GO:0005634] | | cytoplasm [GO:0005737]; | UP000018817 |
| PPTG_11181 | W2QA57 | TKL protein kinase | 359 | | | | | ATP binding [GO:0005524]; protein serine/th | ATP binding [GO:0005524] | UP000018817 |
| PPTG_08153 | W2QJH9 | 26S protease regulatory subunit 8 | 401 | | AAA ATPase family | proteolysis [GO:0006508] | cytoplasm [GO:0005737]; nucleus [GO:0005 | ATP binding [GO:0005524]; ATP hydrolysis ac | cytoplasm [GO:0005737]; | UP000018817 |
| PPTG_07818 | W2QMI4 | Glucosidase II subunit alpha | 1022 | | Glycosyl hydrolase 31 family | carbohydrate metabolic process [GO:00059 | endoplasmic reticulum [GO:0005783] | alpha-glucosidase activity [GO:0090599]; ca | endoplasmic reticulum [G | UP000018817 |
| PPTG_01506 | W2R9T5 | Prephenate dehydratase | 1028 | | Class-I pyridoxal-phosphate-dependent aminotran | L-phenylalanine biosynthetic process [GO:0009094]; tyrosine biosynthetic process [GO:0 | | aspartate-prephenate aminotransferase act | aspartate-prephenate ami | UP000018817 |

| | | | | | | | | | | |
|---|---|---|---|---|---|---|---|---|---|---|
| PPTG_02078 | W2R9F0 | Dolichyl-diphosphooligosaccharide--protein glycosyltransferase 48 kDa subunit (Oligosaccharyl tra | 448 | | DDOST 48 kDa subunit family | protein N-linked glycosylation via asparagin | oligosaccharyltransferase complex [GO:0008250] | | oligosaccharyltransferase | UP000018817 |
| PPTG_09099 | W2QI29 | ornithine carbamoyltransferase (EC 2.1.3.3) | 392 | | Aspartate/ornithine carbamoyltransferase superfa | citrulline biosynthetic process [GO:001924 | cytoplasm [GO:0005737] | amino acid binding [GO:0016597]; ornithine | cytoplasm [GO:0005737]; | UP000018817 |
| PPTG_10597 | W2QDW8 | GST C-terminal domain-containing protein | 251 | | | | cytoplasm [GO:0005737] | glutathione transferase activity [GO:000436 | cytoplasm [GO:0005737]; | UP000018817 |
| PPTG_04475 | W2R2W9 | NAC-A/B domain-containing protein | 191 | | | | nascent polypeptide-associated complex [GO:0005854] | | nascent polypeptide-asso | UP000018817 |
| PPTG_00941 | W2RHM7 | Acetyl-CoA C-acetyltransferase | 439 | | Thiolase-like superfamily, Thiolase family | fatty acid beta-oxidation [GO:0006635] | mitochondrion [GO:0005739] | acetyl-CoA C-acetyltransferase activity [GO: | mitochondrion [GO:0005 | UP000018817 |
| PPTG_15403 | W2PSV8 | FYVE-type domain-containing protein | 394 | | | | | fatty-acyl-CoA binding [GO:0000062]; zinc io | fatty-acyl-CoA binding [GO | UP000018817 |
| PPTG_02533 | W2RD98 | 4-hydroxy-tetrahydrodipicolinate synthase (EC 4.3.3.7) | 292 | | DapA family | diaminopimelate biosynthetic process [GO:0019877]; lysine biosynthetic process via dia | 4-hydroxy-tetrahydrodipicolinate synthase a | 4-hydroxy-tetrahydrodipic | | UP000018817 |
| PPTG_12671 | W2Q0E7 | Proteasome subunit alpha type-2 | 236 | | Peptidase T1A family | ubiquitin-dependent protein catabolic proc | cytoplasm [GO:0005737]; nucleus [GO:0005634]; proteasome core complex, alpha-subur | | cytoplasm [GO:0005737]; | UP000018817 |
| PPTG_12747 | W2Q0R0 | Uncharacterized protein | 501 | | | | | | | UP000018817 |
| PPTG_06713 | W2QSH4 | Catalase (EC 1.11.1.6) | 522 | | Catalase family | hydrogen peroxide catabolic process [GO:0 | mitochondrion [GO:0005739]; peroxisome [G | catalase activity [GO:0004096]; heme bindir | mitochondrion [GO:0005 | UP000018817 |
| PPTG_05211 | W2QW96 | CCT-theta | 545 | | TCP-1 chaperonin family | | cytoplasm [GO:0005737] | ATP binding [GO:0005524]; ATP hydrolysis ac | cytoplasm [GO:0005737]; | UP000018817 |
| PPTG_05481 | W2QXA2 | Cullin family profile domain-containing protein | 748 | | Cullin family | ubiquitin-dependent protein catabolic proc | cullin-RING ubiquitin ligase complex [GO:00 | ubiquitin protein ligase binding [GO:003162 | cullin-RING ubiquitin ligas | UP000018817 |
| PPTG_17323 | W2PL80 | tRNA-binding domain-containing protein | 177 | | | | | tRNA binding [GO:0000049] | tRNA binding [GO:000004 | UP000018817 |
| PPTG_08306 | W2QK27 | pyridoxal 5'-phosphate synthase (glutamine hydrolyzing) (EC 4.3.3.6) | 310 | | PdxS/SNZ family | amino acid metabolic process [GO:0006520]; pyridoxal phosphate biosynthetic process [ | | pyridoxal 5'-phosphate synthase (glutamine | pyridoxal 5'-phosphate syn | UP000018817 |
| PPTG_04194 | W2QZW6 | EGF-like domain-containing protein | 340 | | | | | Notch binding [GO:0005112] | Notch binding [GO:000511 | UP000018817 |
| PPTG_18839 | W2PGN0 | EGF-like domain-containing protein | 682 | | | | | | | UP000018817 |
| PPTG_00911 | W2RJ64 | Transketolase (EC 2.2.1.1) | 696 | | Transketolase family | pentose-phosphate shunt [GO:0006098] | cytosol [GO:0005829] | metal ion binding [GO:0046872]; transketola | cytosol [GO:0005829]; me | UP000018817 |
| PPTG_17585 | W2PJX5 | 26S proteasome regulatory subunit RPN10 | 356 | | Proteasome subunit S5A family | proteasome-mediated ubiquitin-dependent | cytosol [GO:0005829]; nucleus [GO:0005634 | polyubiquitin modification-dependent protei | cytosol [GO:0005829]; nuc | UP000018817 |
| PPTG_02266 | W2RCQ8 | cGMP-dependent protein kinase (EC 2.7.11.12) | 810 | | Protein kinase superfamily, AGC Ser/Thr protein kinase family, cGMP subfamily | | cAMP-dependent protein kinase complex [G | ATP binding [GO:0005524]; cAMP-dependen | cAMP-dependent protein k | UP000018817 |
| PPTG_17912 | W2PI90 | Hsp70-Hsp90 organising protein (Stress-inducible protein 1) | 335 | | | | cytoplasm [GO:0005737] | Hsp90 protein binding [GO:0051879] | cytoplasm [GO:0005737]; | UP000018817 |
| PPTG_10741 | W2QBN2 | Calcineurin-like phosphoesterase domain-containing protein | 503 | | | | | hydrolase activity [GO:0016787] | hydrolase activity [GO:001 | UP000018817 |
| PPTG_14168 | W2PWU8 | Eukaryotic translation initiation factor 3 subunit B (eIF3b) (Eukaryotic translation initiation factor 3 | 696 | | EIF-3 subunit B family | formation of cytoplasmic translation initiati | eukaryotic 43S preinitiation complex [GO:00 | RNA binding [GO:0003723]; translation initia | eukaryotic 43S preinitiatio | UP000018817 |
| PPTG_09054 | W2QG34 | Eukaryotic peptide chain release factor subunit 1 | 407 | | Eukaryotic release factor 1 family | | cytoplasm [GO:0005737] | translation release factor activity [GO:00037 | cytoplasm [GO:0005737]; | UP000018817 |
| PPTG_07024 | W2QRV7 | ATP synthase F1, delta subunit | 231 | | ATPase delta chain family | | membrane [GO:0016020] | proton-transporting ATP synthase activity, ro | membrane [GO:0016020]; | UP000018817 |
| PPTG_10267 | W2QFV1 | PCI domain-containing protein | 446 | | Proteasome subunit S9 family | | proteasome complex [GO:0000502] | | proteasome complex [GO: | UP000018817 |
| PPTG_04806 | W2R483 | Serine/threonine-protein phosphatase (EC 3.1.3.16) | 323 | | PPP phosphatase family, PP-1 subfamily | | cytoplasm [GO:0005737]; nucleus [GO:0005 | metal ion binding [GO:0046872]; protein seri | cytoplasm [GO:0005737]; | UP000018817 |
| PPTG_06393 | W2QSX1 | Uncharacterized protein | 754 | | | | | | | UP000018817 |
| PPTG_02872 | W2REZ0 | Uncharacterized protein | 266 | | | | | | | UP000018817 |
| PPTG_17688 | W2PKS8 | 3-phosphoshikimate 1-carboxyvinyltransferase (EC 2.5.1.19) | 744 | | EPSP synthase family | amino acid biosynthetic process [GO:0008 | cytoplasm [GO:0005737] | 3-dehydroquinate synthase activity [GO:000 | cytoplasm [GO:0005737]; | UP000018817 |
| PPTG_14636 | W2PV88 | Propionyl-CoA carboxylase beta chain, mitochondrial (EC 6.4.1.3) (Propanoyl-CoA:carbon dioxide l | 542 | | AccD/PCCB family | fatty acid biosynthetic process [GO:00066 | acetyl-CoA carboxylase complex [GO:00093 | acetyl-CoA carboxylase activity [GO:000398 | acetyl-CoA carboxylase co | UP000018817 |
| PPTG_05533 | W2QY16 | Multiple inositol polyphosphate phosphatase 1 (EC 3.1.3.62) (EC 3.1.3.80) (2,3-bisphosphoglycera | 488 | | Histidine acid phosphatase family, MINPP1 subfamily | | membrane [GO:0016020] | acid phosphatase activity [GO:0003993]; bis | membrane [GO:0016020]; | UP000018817 |
| PPTG_15376 | W2PUW6 | Actin-2 | 375 | | Actin family | | | ATP binding [GO:0005524]; hydrolase activity | ATP binding [GO:0005524] | UP000018817 |
| PPTG_09844 | W2QC73 | 1,3-beta-glucanosyltransferase | 502 | | Glycosyl hydrolase 72 family | cell wall (1->3)-beta-D-glucan biosynthetic | plasma membrane [GO:0005886] | 1,3-beta-glucanosyltransferase activity [GO: | plasma membrane [GO:0 | UP000018817 |
| PPTG_04186 | W2R1R0 | 60S ribosomal protein L7 | 300 | | Universal ribosomal protein uL30 family | maturation of LSU-rRNA from tricistronic rR | cytosolic large ribosomal subunit [GO:0022 | RNA binding [GO:0003723]; structural consti | cytosolic large ribosomal s | UP000018817 |
| PPTG_10299 | W2QGK3 | Proteasome activator PA28 C-terminal domain-containing protein | 243 | | PA28 family | regulation of G1/S transition of mitotic cell | cytoplasm [GO:0005737]; nucleoplasm [GO: | endopeptidase activator activity [GO:00611 | cytoplasm [GO:0005737]; | UP000018817 |
| PPTG_18768 | W2PGE9 | MHD domain-containing protein | 491 | | Adaptor complexes medium subunit family | intracellular protein transport [GO:0006886 | clathrin adaptor complex [GO:0030131]; endomembrane system [GO:0012505] | | clathrin adaptor complex [ | UP000018817 |
| PPTG_04013 | W2QZ62 | Protein transporter Sec24 | 1024 | | SEC23/SEC24 family, SEC24 subfamily | COPII-coated vesicle cargo loading [GO:00 | COPII vesicle coat [GO:0030127]; endoplasm | SNARE binding [GO:0000149]; zinc ion bindi | COPII vesicle coat [GO:00 | UP000018817 |
| PPTG_05029 | W2QW42 | ATP-dependent (S)-NAD(P)H-hydrate dehydratase (EC 4.2.1.93) (ATP-dependent NAD(P)HX dehydr | 241 | | NnrD/CARKD family | metabolite repair [GO:0110051]; nicotinamide nucleotide metabolic process [GO:00464 | | ATP binding [GO:0005524]; ATP-dependent | ATP binding [GO:0005524] | UP000018817 |
| PPTG_09645 | W2QI94 | glyceraldehyde-3-phosphate dehydrogenase (phosphorylating) (EC 1.2.1.12) | 602 | | Glyceraldehyde-3-phosphate dehydrogenase famil | glucose metabolic process [GO:0006006]; | cytosol [GO:0005829] | glyceraldehyde-3-phosphate dehydrogenase | cytosol [GO:0005829]; glyc | UP000018817 |
| PPTG_17956 | W2PJS8 | 60S ribosomal protein L10 | 208 | | Universal ribosomal protein uL16 family | translation [GO:0006412] | ribonucleoprotein complex [GO:1990904]; ri | structural constituent of ribosome [GO:0003 | ribonucleoprotein comple | UP000018817 |
| PPTG_18089 | W2PID6 | Dihydrolipoamide acetyltransferase component of pyruvate dehydrogenase complex (EC 2.3.1.-) | 437 | | 2-oxoacid dehydrogenase family | pyruvate decarboxylation to acetyl-CoA [GO | mitochondrion [GO:0005739]; pyruvate dehy | acyltransferase activity [GO:0016746] | mitochondrion [GO:0005 | UP000018817 |
| PPTG_00539 | W2RHE0 | Choline/carnitine acyltransferase domain-containing protein | 640 | | Carnitine/choline acetyltransferase family | fatty acid beta-oxidation [GO:0006635] | mitochondrion [GO:0005739] | carnitine O-palmitoyltransferase activity [GC | mitochondrion [GO:0005 | UP000018817 |
| PPTG_13332 | W2Q4G3 | Peptidase C1A papain C-terminal domain-containing protein | 535 | | Peptidase C1 family | proteolysis [GO:0006508] | | cysteine-type peptidase activity [GO:000823 | cysteine-type peptidase a | UP000018817 |
| PPTG_00044 | W2RDY8 | CAMK protein kinase | 471 | | Protein kinase superfamily | | | ATP binding [GO:0005524]; protein serine/th | ATP binding [GO:0005524] | UP000018817 |
| PPTG_16444 | W2PNP8 | Phosphoglycerate mutase (EC 5.4.2.11) | 260 | | Phosphoglycerate mutase family, BPG-dependent | glycolytic process [GO:0006096] | | phosphoglycerate mutase activity [GO:0004 | phosphoglycerate mutase | UP000018817 |
| PPTG_01765 | W2R897 | Uncharacterized protein | 772 | | | | | | | UP000018817 |
| PPTG_07933 | W2QQ00 | NADP-dependent oxidoreductase domain-containing protein | 355 | | Shaker potassium channel beta subunit family | | | oxidoreductase activity [GO:0016491] | oxidoreductase activity [G | UP000018817 |
| PPTG_11925 | W2Q8U8 | PDZ domain-containing protein | 564 | | | | | | | UP000018817 |
| PPTG_00307 | W2RGP0 | Peptidase M20 dimerisation domain-containing protein | 477 | | | | | hydrolase activity [GO:0016787]; metal ion b | hydrolase activity [GO:001 | UP000018817 |
| PPTG_03063 | W2R611 | Uncharacterized protein | 410 | | | | | | | UP000018817 |
| PPTG_00731 | W2RG20 | Succinate-semialdehyde dehydrogenase, mitochondrial (EC 1.2.1.24) (NAD(+)-dependent succinic | 507 | | Aldehyde dehydrogenase family | gamma-aminobutyric acid catabolic process [GO:0009450] | | succinate-semialdehyde dehydrogenase (N | succinate-semialdehyde d | UP000018817 |
| PPTG_20079 | W2PA93 | Tripeptidyl peptidase II second Ig-like domain-containing protein | 125 | | | | | | | UP000018817 |
| PPTG_11779 | W2Q868 | Carnitine O-acetyltransferase, mitochondrial | 633 | | Carnitine/choline acetyltransferase family | fatty acid metabolic process [GO:0006631] | mitochondrial inner membrane [GO:000574 | carnitine O-acetyltransferase activity [GO:0 | mitochondrial inner memb | UP000018817 |
| PPTG_03934 | W2R0R7 | Annexin | 334 | Vinay and Belleannée, 2022 | Annexin family | | cytoplasm [GO:0005737]; plasma membrane | calcium ion binding [GO:0005509]; calcium- | cytoplasm [GO:0005737]; | UP000018817 |
| PPTG_06662 | W2QQL1 | Histidinol dehydrogenase | 421 | | Histidinol dehydrogenase family | L-histidine biosynthetic process [GO:00001 | cytosol [GO:0005829] | histidinol dehydrogenase activity [GO:00043 | cytosol [GO:0005829]; his | UP000018817 |
| PPTG_11134 | W2Q9Y5 | PTHB1 N-terminal domain-containing protein | 916 | | | cilium assembly [GO:0060271] | BBSome [GO:0034464]; membrane [GO:0016020] | | BBSome [GO:0034464]; m | UP000018817 |
| PPTG_08520 | W2QNL0 | DNA damage-binding protein 1 | 1197 | | | | nucleus [GO:0005634] | nucleic acid binding [GO:0003676] | nucleus [GO:0005634]; nu | UP000018817 |
| PPTG_07827 | W2QPJ2 | L-type lectin-like domain-containing protein | 414 | | | endoplasmic reticulum to Golgi vesicle-me | COPII-coated ER to Golgi transport vesicle [G | D-mannose binding [GO:0005537] | COPII-coated ER to Golgi t | UP000018817 |
| PPTG_19695 | W2PDI9 | Uncharacterized protein | 585 | | | cell adhesion [GO:0007155] | | | cell adhesion [GO:000715 | UP000018817 |
| PPTG_09068 | W2QIT1 | PKD/REJ-like domain-containing protein | 7040 | | | | membrane [GO:0016020] | | membrane [GO:0016020] | UP000018817 |
| PPTG_08708 | W2QHE9 | Thaumatin-like protein | 384 | | | | | | | UP000018817 |
| PPTG_19359 | W2PCU8 | DNA-directed RNA polymerase subunit (EC 2.7.7.6) | 1859 | | RNA polymerase beta' chain family | transcription by RNA polymerase II [GO:000 | RNA polymerase II, core complex [GO:0005 | DNA binding [GO:0003677]; DNA-directed R | RNA polymerase II, core c | UP000018817 |
| PPTG_02269 | W2RA80 | Large ribosomal subunit protein uL15/eL18 domain-containing protein | 192 | | Eukaryotic ribosomal protein eL18 family | translation [GO:0006412] | cytosolic large ribosomal subunit [GO:0022 | RNA binding [GO:0003723]; structural consti | cytosolic large ribosomal s | UP000018817 |
| PPTG_14777 | W2PVV6 | Uncharacterized protein | 1081 | | | | | | | UP000018817 |
| PPTG_05019 | W2QVI1 | Isovaleryl-CoA dehydrogenase | 425 | | Acyl-CoA dehydrogenase family | L-leucine catabolic process [GO:0006552] | | 3-methylbutanoyl-CoA dehydrogenase activi | 3-methylbutanoyl-CoA de | UP000018817 |
| PPTG_10204 | W2QDQ1 | ribose-phosphate diphosphokinase (EC 2.7.6.1) | 378 | | Ribose-phosphate pyrophosphokinase family | 5-phosphoribose 1-diphosphate biosyntheti | cytoplasm [GO:0005737]; ribose phosphate | ATP binding [GO:0005524]; kinase activity [G | cytoplasm [GO:0005737]; | UP000018817 |
| PPTG_03523 | W2R578 | GrxB family glutaredoxin | 227 | | | | cytosol [GO:0005829] | | cytosol [GO:0005829] | UP000018817 |
| PPTG_13439 | W2Q1J8 | Phosphofructokinase domain-containing protein | 504 | | | fructose 6-phosphate metabolic process [G | cytoplasm [GO:0005737] | 6-phosphofructokinase activity [GO:000387 | cytoplasm [GO:0005737]; | UP000018817 |
| PPTG_04138 | W2QZC4 | rRNA 2'-O-methyltransferase fibrillarin (Histone-glutamine methyltransferase) | 318 | | Methyltransferase superfamily, Fibrillarin family | box C/D sno(s)RNA 3'-end processing [GO: | box C/D methylation guide snoRNP complex | histone H2AQ104 methyltransferase activity | box C/D methylation guide | UP000018817 |
| PPTG_06991 | W2QRS6 | inorganic diphosphatase (EC 3.6.1.1) | 557 | | PPase family | phosphate-containing compound metaboli | cytoplasm [GO:0005737] | calcium ion binding [GO:0005509]; inorganic | cytoplasm [GO:0005737]; | UP000018817 |
| PPTG_01960 | W2RBH6 | DUF4833 domain-containing protein | 167 | | | | | | | UP000018817 |
| PPTG_12168 | W2Q7Y2 | Elongation factor Tu | 416 | | TRAFAC class translation factor GTPase superfami | mitochondrial translational elongation [GO: | mitochondrion [GO:0005739]; plastid [GO:00 | GTP binding [GO:0005525]; GTPase activity [ | mitochondrion [GO:0005 | UP000018817 |
| PPTG_04619 | W2R3H0 | Uncharacterized protein | 276 | | | | | | | UP000018817 |
| PPTG_04276 | W2R048 | Uncharacterized protein | 763 | | | | | | | UP000018817 |
| PPTG_09278 | W2QIS0 | Uncharacterized protein | 1218 | | | intraciliary retrograde transport [GO:00357 | intraciliary transport particle A [GO:0030991]; non-motile cilium [GO:0097730] | | intraciliary transport partic | UP000018817 |
| PPTG_06297 | W2QS85 | Enoyl reductase (ER) domain-containing protein | 354 | | Zinc-containing alcohol dehydrogenase family | | | oxidoreductase activity, acting on the CH-O | oxidoreductase activity, ac | UP000018817 |
| PPTG_19366 | W2PDE4 | Malic enzyme | 538 | | Malic enzymes family | malate metabolic process [GO:0006108] | | malate dehydrogenase (decarboxylating) (N | malate dehydrogenase (de | UP000018817 |
| PPTG_04879 | W2R2P0 | Eukaryotic translation initiation factor 3 subunit K (eIF3k) (eIF-3 p25) | 229 | | EIF-3 subunit K family | formation of cytoplasmic translation initiati | eukaryotic 43S preinitiation complex [GO:00 | ribosome binding [GO:0043022]; RNA bindin | eukaryotic 43S preinitiatio | UP000018817 |
| PPTG_06519 | W2QTU0 | Peptidase M1 leukotriene A4 hydrolase/aminopeptidase C-terminal domain-containing protein | 638 | | Peptidase M1 family | proteolysis [GO:0006508] | cytoplasm [GO:0005737] | metalloaminopeptidase activity [GO:007000 | cytoplasm [GO:0005737]; | UP000018817 |
| PPTG_08490 | W2QKV3 | Myosin motor domain-containing protein | 1464 | | TRAFAC class myosin-kinesin ATPase superfamily, | actin filament organization [GO:0007015] | cytoplasm [GO:0005737]; membrane [GO:00 | actin filament binding [GO:0051015]; ATP bi | cytoplasm [GO:0005737]; | UP000018817 |
| PPTG_01365 | W2R8J3 | T-complex protein 1 subunit gamma | 530 | | TCP-1 chaperonin family | | chaperonin-containing T-complex [GO:0058 | ATP binding [GO:0005524]; ATP hydrolysis ac | chaperonin-containing T-c | UP000018817 |
| PPTG_06818 | W2QQT8 | Methyltransferase type 11 domain-containing protein | 206 | | Methyltransferase superfamily | methylation [GO:0032259] | | S-adenosylmethionine-dependent methyltra | S-adenosylmethionine-de | UP000018817 |
| PPTG_00479 | W2RHF8 | Apple domain-containing protein | 384 | | | carbohydrate metabolic process [GO:0005 | extracellular region [GO:0005576] | cellulose binding [GO:0030248] | extracellular region [GO:0 | UP000018817 |
| PPTG_16449 | W2PPE0 | Pyruvate kinase (EC 2.7.1.40) | 522 | | Pyruvate kinase family | | | ATP binding [GO:0005524]; kinase activity [G | ATP binding [GO:0005524] | UP000018817 |
| PPTG_17103 | W2PM15 | factor independent urate hydroxylase (EC 1.7.3.3) (Urate oxidase) | 361 | | Uricase family | purine nucleobase catabolic process [GO:0 | peroxisome [GO:0005777] | urate oxidase activity [GO:0004846] | peroxisome [GO:0005777] | UP000018817 |
| PPTG_15650 | W2PR72 | TCTP domain-containing protein | 180 | | TCTP family | | cytoplasm [GO:0005737] | calcium ion binding [GO:0005509] | cytoplasm [GO:0005737]; | UP000018817 |
| PPTG_02615 | W2RDT4 | Uncharacterized protein | 811 | | | | | | | UP000018817 |
| PPTG_19557 | W2PCB0 | methionine--tRNA ligase (EC 6.1.1.10) (Methionyl-tRNA synthetase) | 727 | | Class-I aminoacyl-tRNA synthetase family | methionyl-tRNA aminoacylation [GO:0006431] | | ATP binding [GO:0005524]; methionine-tRN | ATP binding [GO:0005524] | UP000018817 |
| PPTG_14256 | W2PXC1 | Tyrosinase copper-binding domain-containing protein | 649 | | | | | metal ion binding [GO:0046872]; oxidoreduc | metal ion binding [GO:004 | UP000018817 |
| PPTG_10870 | W2QBC6 | phenylalanine--tRNA ligase (EC 6.1.1.20) (Phenylalanyl-tRNA synthetase beta subunit) | 614 | | Phenylalanyl-tRNA synthetase beta subunit family, | phenylalanyl-tRNA aminoacylation [GO:000 | phenylalanine-tRNA ligase complex [GO:000 | ATP binding [GO:0005524]; magnesium ion | phenylalanine-tRNA ligase | UP000018817 |
| PPTG_16154 | W2PRD3 | Proteasome subunit beta | 295 | | Peptidase T1B family | proteolysis involved in protein catabolic pro | cytoplasm [GO:0005737]; nucleus [GO:0005 | threonine-type endopeptidase activity [GO:0 | cytoplasm [GO:0005737]; | UP000018817 |
| PPTG_09863 | W2QE60 | ABC transporter domain-containing protein | 1352 | | ABC transporter superfamily, ABCG family, PDR (TC 3.A.1.205) subfamily | | membrane [GO:0016020] | ABC-type transporter activity [GO:0140359]; | membrane [GO:0016020]; | UP000018817 |
| PPTG_02668 | W2RE19 | Uncharacterized protein | 518 | | | RNA processing [GO:0006396] | cytoplasm [GO:0005737]; nucleus [GO:0005 | RNA binding [GO:0003723] | cytoplasm [GO:0005737]; | UP000018817 |
| PPTG_04900 | W2R323 | Ras-like protein 3 | 196 | | Small GTPase superfamily, Ras family | signal transduction [GO:0007165] | plasma membrane [GO:0005886] | GTP binding [GO:0005525]; GTPase activity [ | plasma membrane [GO:0 | UP000018817 |
| PPTG_01105 | W2RHQ0 | 60S ribosomal protein L23 | 141 | | Universal ribosomal protein uL14 family | translation [GO:0006412] | cytosolic large ribosomal subunit [GO:0022 | large ribosomal subunit rRNA binding [GO:0 | cytosolic large ribosomal s | UP000018817 |
| PPTG_00178 | W2RGD1 | Cilia- and flagella-associated protein 61 N-terminal domain-containing protein | 925 | | Adenylate kinase family | | | AMP kinase activity [GO:0004017]; ATP bindi | AMP kinase activity [GO:0 | UP000018817 |
| PPTG_11108 | W2Q858 | Uncharacterized protein | 1029 | | | | | | | UP000018817 |
| PPTG_00091 | W2RE33 | coproporphyrinogen oxidase (EC 1.3.3.3) | 402 | | Aerobic coproporphyrinogen-III oxidase family | protoporphyrinogen IX biosynthetic process | cytoplasm [GO:0005737] | coproporphyrinogen oxidase activity [GO:00 | cytoplasm [GO:0005737]; | UP000018817 |
| PPTG_12649 | W2Q0X0 | Uncharacterized protein | 484 | | STXBP/unc-18/SEC1 family | vesicle-mediated transport [GO:0016192] | | | vesicle-mediated transpor | UP000018817 |
| PPTG_01826 | W2RAX8 | Uncharacterized protein | 218 | | Small GTPase superfamily, Rab family | | endomembrane system [GO:0012505] | GTP binding [GO:0005525]; GTPase activity [ | endomembrane system [G | UP000018817 |
| PPTG_15085 | W2PWM4 | D-3-phosphoglycerate dehydrogenase (EC 1.1.1.95) | 979 | | D-isomer specific 2-hydroxyacid dehydrogenase fa | L-serine biosynthetic process [GO:0006564] | | NAD binding [GO:0051287]; O-phospho-L-se | NAD binding [GO:0051287 | UP000018817 |

| | | | | | | | | | | |
|---|---|---|---|---|---|---|---|---|---|---|
| PPTG_08927 | W2QKY8 | ribose-phosphate diphosphokinase (EC 2.7.6.1) | 396 | | Ribose-phosphate pyrophosphokinase family | 5-phosphoribose 1-diphosphate biosyntheti | cytoplasm [GO:0005737]; ribose phosphate ( | ATP binding [GO:0005524]; kinase activity [G | cytoplasm [GO:0005737]; | UP000018817 |
| PPTG_05778 | W2QVU1 | Short-chain specific acyl-CoA dehydrogenase, mitochondrial (EC 1.3.8.1) (Butyryl-CoA dehydrogen | 413 | | Acyl-CoA dehydrogenase family | butyrate catabolic process [GO:0046359]; fatty acid beta-oxidation using acyl-CoA dehyd | | flavin adenine dinucleotide binding [GO:005( | flavin adenine dinucleotide | UP000018817 |
| PPTG_00719 | W2RG07 | Myosin motor domain-containing protein | 1257 | | TRAFAC class myosin-kinesin ATPase superfamily, | actin filament organization [GO:0007015] | cytoplasm [GO:0005737]; membrane [GO:00 | actin filament binding [GO:0051015]; ATP bir | cytoplasm [GO:0005737]; | UP000018817 |
| PPTG_12003 | W2Q4R7 | Proteasome subunit beta | 288 | | Peptidase T1B family | proteolysis involved in protein catabolic pro | cytoplasm [GO:0005737]; nucleus [GO:0005( | threonine-type endopeptidase activity [GO:0 | cytoplasm [GO:0005737]; | UP000018817 |
| PPTG_11311 | W2Q943 | Peptidase C1A papain C-terminal domain-containing protein | 355 | | Peptidase C1 family | proteolysis [GO:0006508] | | cysteine-type peptidase activity [GO:000823 | cysteine-type peptidase ac | UP000018817 |
| PPTG_15540 | W2PQA9 | Dihydroorotase, mitochondrial (EC 3.5.2.3) | 386 | | Metallo-dependent hydrolases superfamily, DHOas | 'de novo' pyrimidine nucleobase biosyntheti | cytoplasm [GO:0005737] | dihydroorotase activity [GO:0004151]; metal | cytoplasm [GO:0005737]; | UP000018817 |
| PPTG_15776 | W2PQC2 | Uncharacterized protein | 480 | | NADH dehydrogenase family | | mitochondrion [GO:0005739] | NADH dehydrogenase activity [GO:0003954] | mitochondrion [GO:0005] | UP000018817 |
| PPTG_00961 | W2RHQ1 | 26S protease regulatory subunit 6A | 433 | | AAA ATPase family | proteolysis [GO:0006508] | cytoplasm [GO:0005737]; nucleus [GO:0005( | ATP binding [GO:0005524]; ATP hydrolysis ac | cytoplasm [GO:0005737]; | UP000018817 |
| PPTG_13432 | W2Q5Q3 | ADP-ribosylglycohydrolase | 364 | | | | | metal ion binding [GO:0046872] | metal ion binding [GO:004 | UP000018817 |
| PPTG_04728 | W2R3Y6 | 60S ribosomal protein L13 | 205 | | Eukaryotic ribosomal protein eL13 family | translation [GO:0006412] | cytosolic large ribosomal subunit [GO:00226 | RNA binding [GO:0003723]; structural consti | cytosolic large ribosomal s | UP000018817 |
| PPTG_09333 | W2QH87 | Uncharacterized protein | 174 | | | | | GTP binding [GO:0005525]; GTPase activity [ | GTP binding [GO:0005525] | UP000018817 |
| PPTG_12682 | W2Q131 | beta-glucosidase (EC 3.2.1.21) | 775 | | Glycosyl hydrolase 3 family | glucan catabolic process [GO:0009251] | | beta-glucosidase activity [GO:0008422] | beta-glucosidase activity [( | UP000018817 |
| PPTG_02166 | W2R9R7 | Anoctamin transmembrane domain-containing protein | 934 | | | | membrane [GO:0016020] | chloride channel activity [GO:0005254] | membrane [GO:0016020]; | UP000018817 |
| PPTG_08089 | W2QLP3 | Carbonic anhydrase (EC 4.2.1.1) (Carbonate dehydratase) | 324 | | Beta-class carbonic anhydrase family | carbon utilization [GO:0015976] | | carbonate dehydratase activity [GO:000408( | carbonate dehydratase ac | UP000018817 |
| PPTG_09295 | W2QH23 | GPI inositol-deacylase | 542 | | | | | | | UP000018817 |
| PPTG_09326 | W2QH71 | Tryptophan--tRNA ligase, cytoplasmic (EC 6.1.1.2) (Tryptophanyl-tRNA synthetase) | 537 | | Class-I aminoacyl-tRNA synthetase family | tryptophanyl-tRNA aminoacylation [GO:000 | cytoplasm [GO:0005737] | ATP binding [GO:0005524]; tryptophan-tRNA | cytoplasm [GO:0005737]; | UP000018817 |
| PPTG_04562 | W2R3U4 | glutamine--tRNA ligase (EC 6.1.1.18) | 669 | | Class-I aminoacyl-tRNA synthetase family | glutaminyl-tRNA aminoacylation [GO:0006 | cytosol [GO:0005829] | ATP binding [GO:0005524]; glutamine-tRNA | cytosol [GO:0005829]; ATF | UP000018817 |
| PPTG_10943 | W2QAW3 | 26S protease regulatory subunit 10B, variant | 288 | | AAA ATPase family | proteolysis [GO:0006508] | cytoplasm [GO:0005737]; nucleus [GO:0005( | ATP binding [GO:0005524]; ATP hydrolysis ac | cytoplasm [GO:0005737]; | UP000018817 |
| PPTG_14471 | W2PUM6 | Proteasome alpha-type subunits domain-containing protein | 258 | | Peptidase T1A family | ubiquitin-dependent protein catabolic proc | cytoplasm [GO:0005737]; nucleus [GO:0005634]; proteasome core complex, alpha-subur | | cytoplasm [GO:0005737]; | UP000018817 |
| PPTG_07011 | W2QRU6 | GST C-terminal domain-containing protein | 338 | | | | cytoplasm [GO:0005737] | glutathione transferase activity [GO:00436 | cytoplasm [GO:0005737]; | UP000018817 |
| PPTG_05351 | W2QWH0 | Ketoreductase domain-containing protein | 289 | | Short-chain dehydrogenases/reductases (SDR) family | | | oxidoreductase activity [GO:0016491] | oxidoreductase activity [G( | UP000018817 |
| PPTG_18320 | W2PHC0 | PB1 domain-containing protein | 282 | | | | | | | UP000018817 |
| PPTG_18967 | W2PDU7 | peptidylprolyl isomerase (EC 5.2.1.8) | 478 | | | protein folding [GO:0006457] | cytoplasm [GO:0005737] | cyclosporin A binding [GO:0016018]; peptidy | cytoplasm [GO:0005737]; | UP000018817 |
| PPTG_08523 | W2QL17 | Proteasome subunit beta | 197 | | Peptidase T1B family | proteasomal protein catabolic process [GO | cytoplasm [GO:0005737]; nucleus [GO:0005634]; proteasome core complex [GO:000583( | | cytoplasm [GO:0005737]; | UP000018817 |
| PPTG_09291 | W2QH19 | ALA-interacting subunit | 395 | | CDC50/LEM3 family | | endoplasmic reticulum [GO:0005783]; Golgi apparatus [GO:0005794]; plasma membrane | | endoplasmic reticulum [G( | UP000018817 |
| PPTG_05391 | W2QWP5 | Uncharacterized protein | 803 | | | | | | | UP000018817 |
| PPTG_11162 | W2Q998 | FYVE-type domain-containing protein | 834 | | | | | GTPase activator activity [GO:0005096]; sma | GTPase activator activity [( | UP000018817 |
| PPTG_17268 | W2PLN8 | Uncharacterized protein | 133 | | | mitochondrial electron transport, cytochron | mitochondrial envelope [GO:0005740]; respi | metal ion binding [GO:0046872] | mitochondrial envelope [G | UP000018817 |
| PPTG_00034 | W2RDX2 | Argininosuccinate lyase | 461 | | Lyase 1 family, Argininosuccinate lyase subfamily | L-arginine biosynthetic process via ornithine | cytosol [GO:0005829] | argininosuccinate lyase activity [GO:000405( | cytosol [GO:0005829]; argi | UP000018817 |
| PPTG_09542 | W2QFD6 | Proteasome activator Blm10 mid region domain-containing protein | 1915 | | BLM10 family | DNA repair [GO:0006281]; proteasomal ubi | cytosol [GO:0005829]; nucleus [GO:0005634 | peptidase activator activity [GO:0016504]; p | cytosol [GO:0005829]; nuc | UP000018817 |
| PPTG_06371 | W2QUJ2 | 60S ribosomal protein L12 | 165 | | Universal ribosomal protein uL11 family | translation [GO:0006412] | cytosolic large ribosomal subunit [GO:00226 | large ribosomal subunit rRNA binding [GO:00 | cytosolic large ribosomal s | UP000018817 |
| PPTG_16130 | W2PSG3 | 60S ribosomal protein L27 | 144 | | Eukaryotic ribosomal protein eL27 family | translation [GO:0006412] | chloroplast [GO:0009507]; ribonucleoprotein | structural constituent of ribosome [GO:0003 | chloroplast [GO:0009507]; | UP000018817 |
| PPTG_01517 | W2R7C5 | acetyl-CoA C-acyltransferase (EC 2.3.1.16) | 446 | | Thiolase-like superfamily, Thiolase family | fatty acid beta-oxidation [GO:0006635] | mitochondrion [GO:0005739] | acetyl-CoA C-acetyltransferase activity [GO: | mitochondrion [GO:0005] | UP000018817 |
| PPTG_02652 | W2RBV8 | Histidine kinase/HSP90-like ATPase domain-containing protein | 834 | | Heat shock protein 90 family | | | ATP binding [GO:0005524]; ATP hydrolysis ac | ATP binding [GO:0005524] | UP000018817 |
| PPTG_15537 | W2PPL3 | 50S ribosomal protein L5, chloroplastic | 178 | | Universal ribosomal protein uL5 family | translation [GO:0006412] | cytoplasm [GO:0005737]; nucleus [GO:0005( | rRNA binding [GO:0019843]; structural const | cytoplasm [GO:0005737]; | UP000018817 |
| PPTG_09814 | W2QEL6 | GCK domain-containing protein | 320 | | | | | | | UP000018817 |
| PPTG_14746 | W2PWM0 | Glucose 1-dehydrogenase | 282 | | Short-chain dehydrogenases/reductases (SDR) family | | | oxidoreductase activity, acting on the CH-OH | oxidoreductase activity, ac | UP000018817 |
| PPTG_11404 | W2QBA4 | thioredoxin-dependent peroxiredoxin (EC 1.11.1.24) | 208 | | Peroxiredoxin family, AhpC/Prx1 subfamily | cell redox homeostasis [GO:0045454]; cellu | cytosol [GO:0005829] | thioredoxin peroxidase activity [GO:0008379 | cytosol [GO:0005829]; thic | UP000018817 |
| PPTG_01586 | W2R7M4 | Lipase | 426 | | AB hydrolase superfamily, Lipase family | lipid catabolic process [GO:0016042] | | hydrolase activity, acting on ester bonds [GO | hydrolase activity, acting o | UP000018817 |
| PPTG_04494 | W2R1D8 | cytochrome-b5 reductase (EC 1.6.2.2) | 296 | | | | membrane [GO:0016020] | cytochrome-b5 reductase activity, acting on | membrane [GO:0016020]; | UP000018817 |
| PPTG_05682 | W2QTG8 | 60S ribosomal protein L18a | 176 | | Eukaryotic ribosomal protein eL20 family | translation [GO:0006412] | ribonucleoprotein complex [GO:1990904]; rit | structural constituent of ribosome [GO:0003 | ribonucleoprotein complex | UP000018817 |
| PPTG_08132 | W2QJC9 | Urease (EC 3.5.1.5) (Urea amidohydrolase) | 842 | | | urea catabolic process [GO:0043419] | urease complex [GO:0035550] | nickel cation binding [GO:0016151]; urease a | urease complex [GO:0035 | UP000018817 |
| PPTG_10455 | W2QEJ5 | Translation elongation factor EF1B beta/delta subunit guanine nucleotide exchange domain-contai | 229 | | EF-1-beta/EF-1-delta family | | cytosol [GO:0005829]; eukaryotic translation | guanyl-nucleotide exchange factor activity [( | cytosol [GO:0005829]; euk | UP000018817 |
| PPTG_08989 | W2QL70 | ABC transporter domain-containing protein | 1782 | | ABC transporter superfamily, ABCA family | | membrane [GO:0016020] | ABC-type transporter activity [GO:0140359]; | membrane [GO:0016020]; | UP000018817 |
| PPTG_14633 | W2PXY5 | 3-hydroxyisobutyryl-CoA hydrolase (EC 3.1.2.4) | 427 | | | L-valine catabolic process [GO:0006574] | | 3-hydroxyisobutyryl-CoA hydrolase activity [( | 3-hydroxyisobutyryl-CoA hy | UP000018817 |
| PPTG_00353 | W2RH27 | 40S ribosomal protein S14 | 150 | | Universal ribosomal protein uS11 family | translation [GO:0006412] | chloroplast [GO:0009507]; ribonucleoprotein | structural constituent of ribosome [GO:0003 | chloroplast [GO:0009507]; | UP000018817 |
| PPTG_06980 | W2QS79 | AGC-kinase C-terminal domain-containing protein | 260 | | | | | | | UP000018817 |
| PPTG_06424 | W2QT16 | Chloride channel protein | 705 | | Chloride channel (TC 2.A.49) family | | membrane [GO:0016020] | voltage-gated chloride channel activity [GO:( | membrane [GO:0016020]; | UP000018817 |
| PPTG_08929 | W2QID6 | non-specific serine/threonine protein kinase (EC 2.7.11.1) | 464 | | Protein kinase superfamily, CAMK Ser/Thr protein k | signal transduction [GO:0007165] | | ATP binding [GO:0005524]; protein serine/th | ATP binding [GO:0005524] | UP000018817 |
| PPTG_06270 | W2QU59 | ADP-ribosylation factor-like protein 3 | 183 | Vinay and Belleannée, 2022 | Small GTPase superfamily, Arf family | protein transport [GO:0015031] | Golgi apparatus [GO:0005794] | GTP binding [GO:0005525]; GTPase activity [ | Golgi apparatus [GO:0005] | UP000018817 |
| PPTG_17617 | W2PME4 | Eukaryotic translation initiation factor 3 subunit E (eIF3e) (Eukaryotic translation initiation factor 3 s | 447 | | EIF-3 subunit E family | formation of cytoplasmic translation initiati | eukaryotic 43S preinitiation complex [GO:00 | translation initiation factor activity [GO:0003 | eukaryotic 43S preinitiatio | UP000018817 |
| PPTG_04043 | W2R161 | C2 domain-containing protein | 721 | | Copine family | cellular response to calcium ion [GO:00712 | plasma membrane [GO:0005886] | calcium-dependent phospholipid binding [G( | plasma membrane [GO:00 | UP000018817 |
| PPTG_06413 | W2QUR3 | Dimethylmenaquinone methyltransferase | 224 | | | | | 4-hydroxy-4-methyl-2-oxoglutarate aldolase | 4-hydroxy-4-methyl-2-oxog | UP000018817 |
| PPTG_12286 | W2Q8B8 | Triosephosphate isomerase (EC 5.3.1.1) | 250 | | Triosephosphate isomerase family | gluconeogenesis [GO:0006094]; glyceraldel | cytosol [GO:0005829] | triose-phosphate isomerase activity [GO:00( | cytosol [GO:0005829]; trio | UP000018817 |
| PPTG_03968 | W2R1G1 | Biotin-[acetyl-CoA-carboxylase] ligase | 311 | | Biotin--protein ligase family | | cytoplasm [GO:0005737] | biotin--[biotin carboxyl-carrier protein] ligase | cytoplasm [GO:0005737]; | UP000018817 |
| PPTG_20487 | W2PA86 | Annexin | 198 | Vinay and Belleannée, 2022 | Annexin family | | cytoplasm [GO:0005737]; plasma membrane | calcium ion binding [GO:0005509]; calcium- | cytoplasm [GO:0005737]; | UP000018817 |
| PPTG_07079 | W2QNS1 | Tryptophan synthase beta chain-like PALP domain-containing protein | 254 | | Cysteine synthase/cystathionine beta-synthase far | cysteine biosynthetic process from serine [GO:0006535] | | transferase activity [GO:0016740] | transferase activity [GO:00 | UP000018817 |
| PPTG_08135 | W2QLY9 | Proteasome subunit alpha type | 249 | | Peptidase T1A family | ubiquitin-dependent protein catabolic proc | cytoplasm [GO:0005737]; nucleus [GO:0005634]; proteasome core complex, alpha-subur | | cytoplasm [GO:0005737]; | UP000018817 |
| PPTG_09268 | W2QGX6 | Endoribonuclease L-PSP/chorismate mutase-like domain-containing protein | 172 | | | | | | | UP000018817 |
| PPTG_17620 | W2PME9 | Ketoreductase domain-containing protein | 421 | | Short-chain dehydrogenases/reductases (SDR) family | | peroxisome [GO:0005777] | oxidoreductase activity [GO:0016491] | peroxisome [GO:0005777] | UP000018817 |
| PPTG_14823 | W2PYR6 | Ras-like GTP-binding protein YPT1 | 201 | | Small GTPase superfamily, Rab family | protein transport [GO:0015031] | | GTP binding [GO:0005525]; GTPase activity [ | GTP binding [GO:0005525] | UP000018817 |
| PPTG_12673 | W2Q2T2 | non-specific serine/threonine protein kinase (EC 2.7.11.1) | 370 | | Protein kinase superfamily, Ser/Thr protein kinase family, CDPK subfamily | | | ATP binding [GO:0005524]; metal ion binding | ATP binding [GO:0005524] | UP000018817 |
| PPTG_14768 | W2PVU6 | RuvB-like helicase (EC 3.6.4.12) | 454 | | RuvB family | | nucleus [GO:0005634] | ATP binding [GO:0005524]; ATP hydrolysis ac | nucleus [GO:0005634]; AT | UP000018817 |
| PPTG_09986 | W2QCS5 | ABC transporter E family member 2 | 626 | | | regulation of macromolecule metabolic pro | cytoplasm [GO:0005737] | ATP binding [GO:0005524]; ATP hydrolysis ac | cytoplasm [GO:0005737]; | UP000018817 |
| PPTG_01994 | W2R963 | EF-hand domain-containing protein | 499 | | Mitochondrial carrier (TC 2.A.29) family | | mitochondrial inner membrane [GO:000574 | calcium ion binding [GO:0005509]; succinat | mitochondrial inner memb | UP000018817 |
| PPTG_08075 | W2QJR4 | Plasma membrane ATPase (EC 7.1.2.1) | 965 | | Cation transport ATPase (P-type) (TC 3.A.3) family, | proton export across plasma membrane [G( | plasma membrane [GO:0005886] | ATP binding [GO:0005524]; ATP hydrolysis ac | plasma membrane [GO:00 | UP000018817 |
| PPTG_10776 | W2QBT0 | Peptidase C1A papain C-terminal domain-containing protein | 425 | | Peptidase C1 family | proteolysis [GO:0006508] | | cysteine-type peptidase activity [GO:000823 | cysteine-type peptidase ac | UP000018817 |
| PPTG_03960 | W2R0U8 | 26S protease regulatory subunit 4 | 445 | | AAA ATPase family | proteolysis [GO:0006508] | cytoplasm [GO:0005737]; nucleus [GO:0005( | ATP binding [GO:0005524]; ATP hydrolysis ac | cytoplasm [GO:0005737]; | UP000018817 |
| PPTG_16547 | W2PN75 | aspartate carbamoyltransferase (EC 2.1.3.2) | 282 | | Aspartate/ornithine carbamoyltransferase superfa | 'de novo' pyrimidine nucleobase biosynthetic process [GO:0006207]; 'de novo' UMP biosy | | amino acid binding [GO:0016597]; aspartate | amino acid binding [GO:00 | UP000018817 |
| PPTG_18963 | W2PEF4 | HECT domain-containing protein | 5279 | | | protein ubiquitination [GO:0016567] | | metal ion binding [GO:0046872]; thiosulfate- | metal ion binding [GO:004 | UP000018817 |
| PPTG_19379 | W2PCX2 | histidine--tRNA ligase (EC 6.1.1.21) | 885 | | Class-II aminoacyl-tRNA synthetase family | histidyl-tRNA aminoacylation [GO:0006427 | cytosol [GO:0005829]; mitochondrion [GO:0( | ATP binding [GO:0005524]; histidine-tRNA lig | cytosol [GO:0005829]; mit | UP000018817 |
| PPTG_20004 | W2PCS4 | Ran GTPase-activating protein 1 | 587 | | | | | GTPase activator activity [GO:0005096] | GTPase activator activity [( | UP000018817 |
| PPTG_08537 | W2QNM8 | PPM-type phosphatase domain-containing protein | 395 | | | | | protein serine/threonine phosphatase activit | protein serine/threonine pl | UP000018817 |
| PPTG_03687 | W2R7S6 | Glycoside hydrolase family 31 N-terminal domain-containing protein | 805 | | Glycosyl hydrolase 31 family | carbohydrate metabolic process [GO:0005975] | | carbohydrate binding [GO:0030246]; hydrola | carbohydrate binding [GO: | UP000018817 |
| PPTG_00563 | W2RHH6 | SCP domain-containing protein | 183 | | | | | | | UP000018817 |
| PPTG_14118 | W2PWP0 | tRNA pseudouridine(55) synthase | 484 | | Pseudouridine synthase TruB family | box H/ACA sno(s)RNA 3'-end processing [G( | box H/ACA snoRNP complex [GO:0031429] | pseudouridine synthase activity [GO:000998 | box H/ACA snoRNP comple | UP000018817 |
| PPTG_03917 | W2QYQ7 | V-type proton ATPase subunit C | 415 | | V-ATPase C subunit family | | vacuolar proton-transporting V-type ATPase, ' | proton-transporting ATPase activity, rotation | vacuolar proton-transporti | UP000018817 |
| PPTG_13404 | W2Q5I0 | Anaphase-promoting complex subunit 4 WD40 domain-containing protein | 1421 | | | cilium assembly [GO:0060271]; intraciliary | cilium [GO:0005929]; intraciliary transport particle A [GO:0030991] | | cilium [GO:0005929]; intra | UP000018817 |
| PPTG_14247 | W2PX46 | SAC domain-containing protein | 611 | | | phosphatidylinositol dephosphorylation [GC | endoplasmic reticulum [GO:0005783] | phosphatidylinositol-4-phosphate phosphata | endoplasmic reticulum [G( | UP000018817 |
| PPTG_03627 | W2R5K8 | Actin-related protein 2 | 389 | | Actin family, ARP2 subfamily | actin filament organization [GO:0007015] | Arp2/3 protein complex [GO:0005885] | actin binding [GO:0003779]; ATP binding [GC | Arp2/3 protein complex [G( | UP000018817 |
| PPTG_05618 | W2QXM1 | Enoyl-CoA delta isomerase 1, mitochondrial (3,2-trans-enoyl-CoA isomerase) | 314 | | Enoyl-CoA hydratase/isomerase family | fatty acid beta-oxidation [GO:0006635] | mitochondrial matrix [GO:0005759] | delta(3)-delta(2)-enoyl-CoA isomerase activ | mitochondrial matrix [GO:( | UP000018817 |
| PPTG_13487 | W2Q3E9 | Ras-related protein Rab-1 (Small GTP-binding protein rab1) | 202 | | Small GTPase superfamily, Rab family | protein transport [GO:0015031] | | GTP binding [GO:0005525]; GTPase activity [ | GTP binding [GO:0005525] | UP000018817 |
| PPTG_02695 | W2RCJ0 | Nucleolar protein 56 | 518 | | NOP5/NOP56 family | ribosome biogenesis [GO:0042254] | box C/D methylation guide snoRNP complex | | snoRNA binding [GO:0030515] | box C/D methylation guide | UP000018817 |
| PPTG_11354 | W2Q9B2 | Regulator of microtubule dynamics protein 1 (Protein FAM82B) | 296 | | | | cytoplasm [GO:0005737]; mitotic spindle pol | microtubule binding [GO:0008017] | cytoplasm [GO:0005737]; | UP000018817 |
| PPTG_18664 | W2PEU9 | Temptin Cys/Cys disulfide domain-containing protein | 207 | | | | | | | UP000018817 |
| PPTG_04844 | W2R2A0 | Eukaryotic translation initiation factor 3 subunit H (eIF3h) | 366 | | EIF-3 subunit H family | formation of cytoplasmic translation initiati | eukaryotic 43S preinitiation complex [GO:00 | metallopeptidase activity [GO:0008237]; tra | eukaryotic 43S preinitiatio | UP000018817 |
| PPTG_08288 | W2QKN0 | PPM-type phosphatase domain-containing protein | 421 | | | | | protein serine/threonine phosphatase activit | protein serine/threonine pl | UP000018817 |
| PPTG_07329 | W2QQB1 | Protein disulfide-isomerase domain | 363 | | Protein disulfide isomerase family | protein folding [GO:0006457] | endoplasmic reticulum [GO:0005783] | protein disulfide isomerase activity [GO:000 | endoplasmic reticulum [G( | UP000018817 |
| PPTG_01070 | W2RJV0 | Dihydroorotate dehydrogenase (quinone), mitochondrial (DHOdehase) (EC 1.3.5.2) | 424 | | Dihydroorotate dehydrogenase family, Type 2 subfa | 'de novo' pyrimidine nucleobase biosyntheti | mitochondrial inner membrane [GO:000574 | dihydroorotate dehydrogenase (quinone) act | mitochondrial inner memb | UP000018817 |
| PPTG_05057 | W2QVN0 | 6-phosphogluconolactonase | 397 | | Cycloisomerase 2 family | | | 6-phosphogluconolactonase activity [GO:00 | 6-phosphogluconolactona | UP000018817 |
| PPTG_02563 | W2RDT7 | Serine protease (EC 3.4.21.-) | 404 | | Peptidase S1B family | proteolysis [GO:0006508] | | serine-type peptidase activity [GO:0008236] | serine-type peptidase acti | UP000018817 |
| PPTG_09997 | W2QDM4 | assimilatory sulfite reductase (NADPH) (EC 1.8.1.2) | 1503 | | Nitrite and sulfite reductase 4Fe-4S domain family | sulfate assimilation [GO:0000103] | sulfite reductase complex (NADPH) [GO:000 | 4 iron, 4 sulfur cluster binding [GO:0051539] | sulfite reductase complex | UP000018817 |
| PPTG_05621 | W2R0D6 | RNA helicase (EC 3.6.4.13) | 638 | | DEAD box helicase family | | | ATP binding [GO:0005524]; hydrolase activity | ATP binding [GO:0005524] | UP000018817 |
| PPTG_04070 | W2QZ41 | Iron-sulfur clusters transporter atm1, mitochondrial | 697 | | | intracellular iron ion homeostasis [GO:0006 | mitochondrial inner membrane [GO:000574 | ABC-type transporter activity [GO:0140359]; | mitochondrial inner memb | UP000018817 |
| PPTG_03675 | W2R673 | Grx4 family monothiol glutaredoxin | 448 | | | intracellular iron ion homeostasis [GO:0006 | cytosol [GO:0005829]; nucleus [GO:0005634 | iron-sulfur cluster binding [GO:0051536]; me | cytosol [GO:0005829]; nuc | UP000018817 |
| PPTG_13222 | W2Q0P6 | Uncharacterized protein | 169 | | | mitochondrial electron transport, NADH to u | mitochondrial inner membrane [GO:0005743]; respiratory chain complex I [GO:0045271] | | mitochondrial inner memb | UP000018817 |
| PPTG_06420 | W2QSS5 | Nucleoside diphosphate kinase (EC 2.7.4.6) | 225 | | NDK family | CTP biosynthetic process [GO:0006241]; GTP biosynthetic process [GO:0006183]; UTP bi | | ATP binding [GO:0005524]; nucleoside diphc | ATP binding [GO:0005524] | UP000018817 |
| PPTG_08967 | W2QIH3 | Gamma-soluble NSF attachment protein (N-ethylmaleimide-sensitive factor attachment protein ga | 508 | | SNAP family | intracellular protein transport [GO:0006886 | SNARE complex [GO:0031201]; vacuolar mer | soluble NSF attachment protein activity [GO: | SNARE complex [GO:0031 | UP000018817 |
| PPTG_03938 | W2QYM4 | Delta-1-pyrroline-5-carboxylate synthase [Includes: Glutamate 5-kinase (GK) (EC 2.7.2.11) (Gamm | 757 | | Gamma-glutamyl phosphate reductase family; Glu | L-proline biosynthetic process [GO:005512( | cytoplasm [GO:0005737] | ATP binding [GO:0005524]; glutamate 5-kina | cytoplasm [GO:0005737]; | UP000018817 |
| PPTG_01107 | W2RI96 | Dynactin subunit 6 | 236 | | | | | | | UP000018817 |
| PPTG_14719 | W2PVQ2 | Importin N-terminal domain-containing protein | 969 | | XPO2/CSE1 family | protein export from nucleus [GO:0006611]; | cytosol [GO:0005829]; nuclear envelope [GO | nuclear export signal receptor activity [GO:0( | cytosol [GO:0005829]; nuc | UP000018817 |

| | | | | | | | | | | |
|---|---|---|---|---|---|---|---|---|---|---|
| PPTG_12276 | W2Q5N6 | ABC transporter domain-containing protein | 1353 | | ABC transporter superfamily, ABCG family, PDR (TC 3.A.1.205) subfamily | | membrane [GO:0016020] | ABC-type transporter activity [GO:0140359]; | membrane [GO:0016020]; | UP000018817 |
| PPTG_09168 | W2QGK0 | Cyclic nucleotide-binding domain-containing protein | 849 | | Metallo-beta-lactamase superfamily, cNMP phosphodiesterase family | | cAMP-dependent protein kinase complex [GC | cAMP binding [GO:0030552]; cAMP-depende | cAMP-dependent protein k | UP000018817 |
| PPTG_04886 | W2R2P9 | Tubby C-terminal domain-containing protein | 543 | | TUB family | | | | | UP000018817 |
| PPTG_15080 | W2PTX9 | RING-type domain-containing protein | 457 | | | protein ubiquitination [GO:0016567] | nucleus [GO:0005634] | ubiquitin-protein transferase activity [GO:00 | nucleus [GO:0005634]; ub | UP000018817 |
| PPTG_13264 | W2PYE9 | Eukaryotic translation initiation factor 2A (eIF-2A) | 619 | | WD repeat EIF2A family | regulation of translation [GO:0006417] | cytosolic small ribosomal subunit [GO:00226 | mRNA binding [GO:0003729]; ribosome bind | cytosolic small ribosomal | UP000018817 |
| PPTG_07724 | W2QR51 | glucan endo-1,3-beta-D-glucosidase (EC 3.2.1.39) (Endo-1,3-beta-glucanase btgC) (Laminarinase | 298 | | | cell wall organization [GO:0071555]; polysa | plasma membrane [GO:0005886] | glucan endo-1,3-beta-D-glucosidase activity | plasma membrane [GO:00 | UP000018817 |
| PPTG_14474 | W2PUJ2 | Uncharacterized protein | 161 | | Small GTPase superfamily, Rab family | | | GTP binding [GO:0005525]; GTPase activity [ | GTP binding [GO:0005525] | UP000018817 |
| PPTG_17566 | W2PJU9 | 60S ribosomal protein L32 | 135 | | Eukaryotic ribosomal protein eL32 family | translation [GO:0006412] | cytosolic large ribosomal subunit [GO:00226 | structural constituent of ribosome [GO:0003 | cytosolic large ribosomal s | UP000018817 |
| PPTG_03990 | W2QZ39 | Serine/threonine-protein kinase TOR (EC 2.7.11.1) | 2653 | | PI3/PI4-kinase family | negative regulation of macroautophagy [GO | cytoplasm [GO:0005737]; nucleus [GO:0005 | ATP binding [GO:0005524]; protein serine kir | cytoplasm [GO:0005737]; | UP000018817 |
| PPTG_05288 | W2QYH1 | Ubiquitin carboxyl-terminal hydrolase (EC 3.4.19.12) | 851 | | Peptidase C19 family | protein deubiquitination [GO:0016579]; pro | cytosol [GO:0005829]; nucleus [GO:0005634 | cysteine-type deubiquitinase activity [GO:00 | cytosol [GO:0005829]; nuc | UP000018817 |
| PPTG_19371 | W2PC88 | Mitogen-activated protein kinase (EC 2.7.11.24) | 1195 | | Protein kinase superfamily, Ser/Thr protein kinase family, MAP kinase subfamily | | | ATP binding [GO:0005524]; MAP kinase activ | ATP binding [GO:0005524] | UP000018817 |
| PPTG_17616 | W2PLR9 | Uncharacterized protein | 577 | | | | | | | UP000018817 |
| PPTG_05149 | W2QWN9 | AGC/PKA protein kinase | 330 | | Protein kinase superfamily | anatomical structure morphogenesis [GO:0 | cAMP-dependent protein kinase complex [GC | ATP binding [GO:0005524]; cAMP-dependen | cAMP-dependent protein k | UP000018817 |
| PPTG_19704 | W2PDB0 | Cysteine synthase (EC 2.5.1.47) | 355 | | Cysteine synthase/cystathionine beta-synthase far | cysteine biosynthetic process from serine [( | cytoplasm [GO:0005737] | cysteine synthase activity [GO:0004124] | cytoplasm [GO:0005737]; | UP000018817 |
| PPTG_14362 | W2Q053 | DnaJ homologue subfamily C GRV2/DNAJC13 N-terminal domain-containing protein | 2503 | | | endosome organization [GO:0007032]; rece | endosome membrane [GO:0010008] | | endosome membrane [GC | UP000018817 |
| PPTG_00739 | W2RGP3 | Uncharacterized protein | 593 | | | | | | | UP000018817 |
| PPTG_09530 | W2QH80 | Vacuolar protein sorting-associated protein 45 | 625 | | STXBP/unc-18/SEC1 family | vesicle-mediated transport [GO:0016192] | | | vesicle-mediated transpor | UP000018817 |
| PPTG_09215 | W2QJE7 | Glycosyl hydrolase family 30 TIM-barrel domain-containing protein | 635 | | Glycosyl hydrolase 30 family | glucosylceramide catabolic process [GO:00 | membrane [GO:0016020] | glucosylceramidase activity [GO:0004348] | membrane [GO:0016020]; | UP000018817 |
| PPTG_13582 | W2Q4N4 | 2-isopropylmalate synthase (EC 2.3.3.13) | 576 | | Alpha-IPM synthase/homocitrate synthase family, I | L-leucine biosynthetic process [GO:0009098] | | 2-isopropylmalate synthase activity [GO:000 | 2-isopropylmalate synthas | UP000018817 |
| PPTG_19098 | W2PFP4 | Apple domain-containing protein | 612 | | | proteolysis [GO:0006508] | extracellular region [GO:0005576] | | extracellular region [GO:0( | UP000018817 |
| PPTG_10213 | W2QFN1 | Calx-beta domain-containing protein | 3722 | Welsh et al., 2024 | | cell communication [GO:0007154] | membrane [GO:0016020] | | membrane [GO:0016020]; | UP000018817 |
| PPTG_08413 | W2QN40 | Uncharacterized protein | 199 | | Synaptobrevin family | endoplasmic reticulum to Golgi vesicle-me | Golgi apparatus [GO:0005794]; membrane [( | SNAP receptor activity [GO:0005484] | Golgi apparatus [GO:0005 | UP000018817 |
| PPTG_06879 | W2QT21 | Small ribosomal subunit protein mS29 | 407 | | Mitochondrion-specific ribosomal protein mS29 family | | mitochondrial small ribosomal subunit [GO:0 | structural constituent of ribosome [GO:0003 | mitochondrial small ribosc | UP000018817 |
| PPTG_15291 | W2PU12 | CSC1/OSCA1-like cytosolic domain-containing protein | 535 | | | | plasma membrane [GO:0005886] | calcium-activated cation channel activity [G | plasma membrane [GO:00 | UP000018817 |
| PPTG_19125 | W2PG46 | Cytochrome c domain-containing protein | 270 | | Cytochrome c family | mitochondrial electron transport, ubiquinol | mitochondrial inner membrane [GO:000574 | electron transfer activity [GO:0009055]; hem | mitochondrial inner memb | UP000018817 |
| PPTG_07145 | W2QP63 | Protein kinase domain-containing protein | 229 | | | | | ATP binding [GO:0005524]; protein kinase ac | ATP binding [GO:0005524] | UP000018817 |
| PPTG_13946 | W2PZ59 | EB1 C-terminal domain-containing protein | 309 | | MAPRE family | cell division [GO:0051301] | microtubule [GO:0005874] | microtubule binding [GO:0008017] | microtubule [GO:0005874 | UP000018817 |
| PPTG_01700 | W2R8H7 | Nascent polypeptide-associated complex subunit beta | 162 | | NAC-beta family | | | | | UP000018817 |
| PPTG_01954 | W2R9H3 | Ubiquitin-40S ribosomal protein S27a | 156 | | Eukaryotic ribosomal protein eS31 family; Ubiquitir | translation [GO:0006412] | cytoplasm [GO:0005737]; ribonucleoprotein | structural constituent of ribosome [GO:0003 | cytoplasm [GO:0005737]; | UP000018817 |
| PPTG_01436 | W2R7D1 | Intraflagellar transport protein | 1771 | | IFT172 family | intraciliary transport [GO:0042073] | axoneme [GO:0005930]; ciliary basal body [GO:0036064]; intraciliary transport particle B [ | | axoneme [GO:0005930]; c | UP000018817 |
| PPTG_12624 | W2PZY1 | Glycine cleavage system P protein (EC 1.4.4.2) | 998 | | GcvP family | glycine decarboxylation via glycine cleavage | glycine cleavage complex [GO:0005960]; mit | glycine binding [GO:0016594]; glycine dehyd | glycine cleavage complex [ | UP000018817 |
| PPTG_16129 | W2PQ86 | UDENN domain-containing protein | 1328 | | | regulation of Rab protein signal transductio | cytoplasmic vesicle [GO:0031410] | GTPase activator activity [GO:0005096]; zinc | cytoplasmic vesicle [GO:0 | UP000018817 |
| PPTG_15127 | W2PWS9 | Proteasome subunit alpha type | 250 | | Peptidase T1A family | ubiquitin-dependent protein catabolic proc | cytoplasm [GO:0005737]; nucleus [GO:0005634]; proteasome core complex, alpha-subur | | cytoplasm [GO:0005737]; | UP000018817 |
| PPTG_04830 | W2R2G2 | Carrier domain-containing protein | 1311 | | | | | | | UP000018817 |
| PPTG_06464 | W2QV13 | PEBP family protein | 247 | | | | | | | UP000018817 |
| PPTG_12760 | W2Q363 | Uncharacterized protein | 798 | | | | | | | UP000018817 |
| PPTG_06756 | W2QSL8 | Short chain dehydrogenase | 242 | | Short-chain dehydrogenases/reductases (SDR) family | | cytoplasm [GO:0005737] | oxidoreductase activity [GO:0016491] | cytoplasm [GO:0005737]; | UP000018817 |
| PPTG_02151 | W2RC33 | J domain-containing protein | 501 | | | | | | | UP000018817 |
| PPTG_15545 | W2PPA6 | FYVE-type domain-containing protein | 409 | | | | | | | UP000018817 |
| PPTG_14334 | W2Q023 | 3-hydroxyacyl-CoA dehydrogenase (EC 1.1.1.35) | 324 | | 3-hydroxyacyl-CoA dehydrogenase family | fatty acid beta-oxidation [GO:0006635] | mitochondrial matrix [GO:0005759] | (3S)-3-hydroxyacyl-CoA dehydrogenase (NAI | mitochondrial matrix [GO: | UP000018817 |
| PPTG_04856 | W2R2C5 | glucan endo-1,3-beta-D-glucosidase (EC 3.2.1.39) (Endo-1,3-beta-glucanase btgC) (Laminarinase | 381 | | | cell wall organization [GO:0071555]; polysa | plasma membrane [GO:0005886] | glucan endo-1,3-beta-D-glucosidase activity | plasma membrane [GO:00 | UP000018817 |
| PPTG_04854 | W2R2K8 | Uncharacterized protein | 603 | | | non-motile cilium assembly [GO:1905515]; | axoneme [GO:0005930]; BBSome [GO:0034 | patched binding [GO:0005113]; smoothenec | axoneme [GO:0005930]; B | UP000018817 |
| PPTG_14842 | W2PWB7 | Transmembrane 9 superfamily member | 645 | | Nonaspanin (TM9SF) (TC 9.A.2) family | protein localization to membrane [GO:0072 | cytoplasm [GO:0005737]; membrane [GO:0016020] | | cytoplasm [GO:0005737]; | UP000018817 |
| PPTG_02161 | W2RA51 | GTP-binding protein YPTC4 | 210 | | Small GTPase superfamily, Rab family | | endomembrane system [GO:0012505] | GTP binding [GO:0005525]; GTPase activity [ | endomembrane system [G | UP000018817 |
| PPTG_02090 | W2R9E6 | Aldehyde dehydrogenase | 541 | | Aldehyde dehydrogenase family | aldehyde metabolic process [GO:0006081] | cytoplasm [GO:0005737] | aldehyde dehydrogenase (NAD+) activity [GC | cytoplasm [GO:0005737]; | UP000018817 |
| PPTG_15632 | W2PR52 | Voltage-dependent anion-selective channel protein | 282 | | Eukaryotic mitochondrial porin family | | mitochondrial outer membrane [GO:000574 | porin activity [GO:0015288]; voltage-gated m | mitochondrial outer memb | UP000018817 |
| PPTG_02396 | W2RAR3 | Uncharacterized protein | 229 | | GST superfamily | translational elongation [GO:0006414] | cytoplasm [GO:0005737]; nucleus [GO:0005634] | | cytoplasm [GO:0005737]; | UP000018817 |
| PPTG_03260 | W2R4A2 | acetylornithine transaminase (EC 2.6.1.11) | 408 | | Class-III pyridoxal-phosphate-dependent aminotra | L-arginine biosynthetic process [GO:00065 | mitochondrion [GO:0005739] | identical protein binding [GO:0042802]; N2-a | mitochondrion [GO:0005 | UP000018817 |
| PPTG_16923 | W2PLH3 | Uncharacterized protein | 465 | | | glutathione metabolic process [GO:00067 | cytoplasm [GO:0005737] | glutathione transferase activity [GO:00043 | cytoplasm [GO:0005737]; | UP000018817 |
| PPTG_09690 | W2QIE5 | methylmalonate-semialdehyde dehydrogenase (CoA acylating) (EC 1.2.1.27) | 550 | | Aldehyde dehydrogenase family | L-valine catabolic process [GO:0006574]; tl | mitochondrion [GO:0005739] | methylmalonate-semialdehyde dehydrogen | mitochondrion [GO:0005 | UP000018817 |
| PPTG_06388 | W2QUM0 | Tyrosinase copper-binding domain-containing protein | 524 | | | | | metal ion binding [GO:0046872]; oxidoreduc | metal ion binding [GO:004 | UP000018817 |
| PPTG_10122 | W2QDB7 | Hsp70-like protein | 569 | | Heat shock protein 70 family | | | ATP binding [GO:0005524]; ATP-dependent p | ATP binding [GO:0005524] | UP000018817 |
| PPTG_01761 | W2R892 | Nucleoside diphosphate kinase (EC 2.7.4.6) | 151 | | NDK family | CTP biosynthetic process [GO:0006241]; GTP biosynthetic process [GO:0006183]; UTP bi | | ATP binding [GO:0005524]; nucleoside diphc | ATP binding [GO:0005524] | UP000018817 |
| PPTG_08347 | W2QMV5 | NADH dehydrogenase [ubiquinone] 1 alpha subcomplex subunit 12 | 226 | | Complex I NDUFA12 subunit family | response to oxidative stress [GO:0006979] | mitochondrial inner membrane [GO:0005743]; respiratory chain complex I [GO:0045271] | | mitochondrial inner memb | UP000018817 |
| PPTG_14112 | W2PWU7 | 60S ribosomal protein | 256 | | Universal ribosomal protein uL2 family | cytoplasmic translation [GO:0002181] | chloroplast [GO:0009507]; cytosolic large rib | RNA binding [GO:0003723]; structural consti | chloroplast [GO:0009507]; | UP000018817 |
| PPTG_18243 | W2PGT8 | 4-aminobutyrate--2-oxoglutarate transaminase (EC 2.6.1.19) (GABA aminotransferase) (Gamma-a | 503 | | Class-III pyridoxal-phosphate-dependent aminotra | gamma-aminobutyric acid catabolic proces | mitochondrion [GO:0005739] | 4-aminobutyrate:2-oxoglutarate transamina | mitochondrion [GO:0005 | UP000018817 |
| PPTG_09562 | W2QI00 | SAC domain-containing protein | 741 | | | phosphatidylinositol dephosphorylation [GC | endoplasmic reticulum [GO:0005783] | phosphatidylinositol-4-phosphate phosphata | endoplasmic reticulum [G | UP000018817 |
| PPTG_14139 | W2PWS0 | Methyltransferase domain-containing protein | 310 | | | | | methyltransferase activity [GO:0008168] | methyltransferase activity | UP000018817 |
| PPTG_03631 | W2R7K7 | Eukaryotic translation initiation factor 3 subunit F (eIF3f) (Eukaryotic translation initiation factor 3 s | 279 | | EIF-3 subunit F family | formation of cytoplasmic translation initiati | eukaryotic 43S preinitiation complex [GO:00 | metallopeptidase activity [GO:0008237]; tra | eukaryotic 43S preinitiatio | UP000018817 |
| PPTG_11136 | W2Q7F3 | Carboxypeptidase (EC 3.4.16.-) | 554 | | Peptidase S10 family | proteolysis [GO:0006508] | | serine-type carboxypeptidase activity [GO:0( | serine-type carboxypeptid | UP000018817 |
| PPTG_16288 | W2PRG2 | Nicotinate-nucleotide pyrophosphorylase [carboxylating] (EC 2.4.2.19) (Quinolinate phosphoribosy | 301 | | NadC/ModD family | NAD+ biosynthetic process [GO:0009435]; | cytoplasm [GO:0005737] | nicotinate-nucleotide diphosphorylase (carb | cytoplasm [GO:0005737]; | UP000018817 |
| PPTG_02623 | W2REA5 | Galactonolactone dehydrogenase | 560 | | | | membrane [GO:0016020] | D-arabinono-1,4-lactone oxidase activity [G | membrane [GO:0016020]; | UP000018817 |
| PPTG_14269 | W2PZ72 | Tyrosinase copper-binding domain-containing protein | 570 | | | | | metal ion binding [GO:0046872]; oxidoreduc | metal ion binding [GO:004 | UP000018817 |
| PPTG_12599 | W2PZV1 | Amidase domain-containing protein | 618 | | Amidase family | | | catalytic activity [GO:0003824] | catalytic activity [GO:0003 | UP000018817 |
| PPTG_06287 | W2QS75 | ABC transporter domain-containing protein | 1934 | | ABC transporter superfamily, ABCA family | | membrane [GO:0016020] | ABC-type transporter activity [GO:0140359]; | membrane [GO:0016020]; | UP000018817 |
| PPTG_00525 | W2RHC0 | GB1/RHD3-type G domain-containing protein | 620 | | TRAFAC class dynamin-like GTPase superfamily, GB1/RHD3 GTPase family | | | GTP binding [GO:0005525]; GTPase activity [ | GTP binding [GO:0005525] | UP000018817 |
| PPTG_10954 | W2QDG9 | glucan endo-1,3-beta-D-glucosidase (EC 3.2.1.39) (Endo-1,3-beta-glucanase btgC) (Laminarinase | 394 | | Glycosyl hydrolase 17 family | cell wall organization [GO:0071555]; polysa | plasma membrane [GO:0005886] | glucan endo-1,3-beta-D-glucosidase activity | plasma membrane [GO:00 | UP000018817 |
| PPTG_18968 | W2PG99 | DNA-directed RNA polymerase subunit (EC 2.7.7.6) | 1811 | | RNA polymerase beta' chain family | DNA-templated transcription [GO:0006351] | RNA polymerase I complex [GO:0005736] | DNA binding [GO:0003677]; DNA-directed RI | RNA polymerase I complex | UP000018817 |
| PPTG_05082 | W2QVS4 | PCI domain-containing protein | 437 | | Proteasome subunit p55 family | | cytoplasm [GO:0005737]; nucleus [GO:0005634]; proteasome regulatory particle, lid subc | | cytoplasm [GO:0005737]; | UP000018817 |
| PPTG_15753 | W2PTH4 | Ribosomal protein S4 | 192 | | Universal ribosomal protein uS4 family | ribosomal small subunit biogenesis [GO:00 | cytosolic small ribosomal subunit [GO:00226 | rRNA binding [GO:0019843]; structural const | cytosolic small ribosomal | UP000018817 |
| PPTG_14025 | W2Q1X9 | 60S acidic ribosomal protein P2 | 111 | | Eukaryotic ribosomal protein P1/P2 family | cytoplasmic translational elongation [GO:0( | cytosolic large ribosomal subunit [GO:00226 | structural constituent of ribosome [GO:0003 | cytosolic large ribosomal s | UP000018817 |
| PPTG_13761 | W2Q0N0 | Guanylate cyclase domain-containing protein | 883 | | | cyclic nucleotide biosynthetic process [GO: | membrane [GO:0016020] | monoatomic ion channel activity [GO:00052 | membrane [GO:0016020]; | UP000018817 |
| PPTG_16174 | W2PNH9 | START domain-containing protein | 693 | | | | endoplasmic reticulum [GO:0005783] | lipid binding [GO:0008289] | endoplasmic reticulum [G | UP000018817 |
| PPTG_02205 | W2R9Y0 | Phospholipid-transporting ATPase (EC 7.6.2.1) | 1069 | | Cation transport ATPase (P-type) (TC 3.A.3) family, | phospholipid translocation [GO:0045332] | plasma membrane [GO:0005886] | ATP binding [GO:0005524]; ATP hydrolysis ac | plasma membrane [GO:00 | UP000018817 |
| PPTG_06550 | W2QVB3 | Myosin motor domain-containing protein | 1157 | | TRAFAC class myosin-kinesin ATPase superfamily, | actin filament organization [GO:0007015]; e | cytoplasm [GO:0005737]; myosin complex [( | actin filament binding [GO:0051015]; ATP bir | cytoplasm [GO:0005737]; | UP000018817 |
| PPTG_04831 | W2R4A9 | Acetyl-CoA carboxylase, biotin carboxylase subunit | 702 | | | | mitochondrial matrix [GO:0005759] | ATP binding [GO:0005524]; metal ion binding | mitochondrial matrix [GO: | UP000018817 |
| PPTG_00384 | W2REN0 | C2 domain-containing protein | 1973 | | | plasma membrane organization [GO:00070 | membrane [GO:0016020] | | membrane [GO:0016020]; | UP000018817 |
| PPTG_17386 | W2PK79 | Putative auto-transporter adhesin head GIN domain-containing protein | 545 | | | | | | | UP000018817 |
| PPTG_02222 | W2RC07 | AP-1 complex subunit gamma | 849 | | Adaptor complexes large subunit family | intracellular protein transport [GO:0006886 | AP-1 adaptor complex [GO:0030121] | | AP-1 adaptor complex [GO | UP000018817 |
| PPTG_13130 | W2PZM2 | Calmodulin | 149 | | Calmodulin family | | cytoplasm [GO:0005737]; myosin II complex | calcium ion binding [GO:0005509] | cytoplasm [GO:0005737]; | UP000018817 |
| PPTG_02310 | W2RAB8 | T-complex protein 1 subunit eta (TCP-1-eta) (CCT-eta) | 545 | | TCP-1 chaperonin family | | chaperonin-containing T-complex [GO:00058 | ATP binding [GO:0005524]; ATP hydrolysis ac | chaperonin-containing T-c | UP000018817 |
| PPTG_13124 | W2PYI5 | T-complex protein 1 subunit epsilon (CCT-epsilon) | 536 | | TCP-1 chaperonin family | | chaperonin-containing T-complex [GO:00058 | ATP binding [GO:0005524]; ATP hydrolysis ac | chaperonin-containing T-c | UP000018817 |
| PPTG_16034 | W2PSK9 | Hyaluronan/mRNA-binding protein domain-containing protein | 278 | | | | cytoplasm [GO:0005737]; nucleus [GO:0005 | RNA binding [GO:0003723] | cytoplasm [GO:0005737]; | UP000018817 |
| PPTG_07334 | W2QQB4 | Expansin-like EG45 domain-containing protein | 663 | | | | | | | UP000018817 |
| PPTG_00570 | W2RFW1 | SCP domain-containing protein | 212 | | | | | | | UP000018817 |
| PPTG_14524 | W2PXI7 | N-acetyl-gamma-glutamyl-phosphate reductase | 717 | | NAGSA dehydrogenase family; Acetylglutamate kin | L-arginine biosynthetic process [GO:00065 | mitochondrion [GO:0005739] | acetylglutamate kinase activity [GO:000399 | mitochondrion [GO:0005 | UP000018817 |
| PPTG_10325 | W2QF04 | phenylalanine--tRNA ligase (EC 6.1.1.20) (Phenylalanyl-tRNA synthetase alpha subunit) | 502 | | Class-II aminoacyl-tRNA synthetase family, Phe-tR | phenylalanyl-tRNA aminoacylation [GO:000 | cytosol [GO:0005829]; phenylalanine-tRNA li | ATP binding [GO:0005524]; metal ion binding | cytosol [GO:0005829]; phe | UP000018817 |
| PPTG_05245 | W2QWD1 | T-complex protein 1, zeta subunit | 539 | | TCP-1 chaperonin family | | cytoplasm [GO:0005737] | ATP binding [GO:0005524]; ATP hydrolysis ac | cytoplasm [GO:0005737]; | UP000018817 |
| PPTG_11884 | W2QB62 | Uncharacterized protein | 2275 | | | cell morphogenesis [GO:0000902] | cell cortex [GO:0005938]; site of polarized growth [GO:0030427] | | cell cortex [GO:0005938]; | UP000018817 |
| PPTG_08060 | W2QJ11 | 2,4-dienoyl-CoA reductase | 675 | | NADH:flavin oxidoreductase/NADH oxidase family | | | FMN binding [GO:0010181]; iron-sulfur clust | FMN binding [GO:0010181 | UP000018817 |
| PPTG_13260 | W2PYE4 | AMP-dependent synthetase/ligase domain-containing protein | 307 | | | fatty acid metabolic process [GO:0006631] | | medium-chain fatty acid-CoA ligase activity | medium-chain fatty acid-C | UP000018817 |
| PPTG_00988 | W2RHW9 | Ribosomal protein L13 | 203 | | Universal ribosomal protein uL13 family | negative regulation of translation [GO:0017 | cytosolic large ribosomal subunit [GO:00226 | mRNA binding [GO:0003729]; structural cons | cytosolic large ribosomal s | UP000018817 |
| PPTG_00112 | W2RDQ5 | Mitochondrial 2-oxoglutarate/malate carrier protein | 306 | | Mitochondrial carrier (TC 2.A.29) family | transmembrane transport [GO:0055085] | membrane [GO:0016020] | | membrane [GO:0016020]; | UP000018817 |
| PPTG_12927 | W2Q1G2 | ADF-H domain-containing protein | 143 | | Actin-binding proteins ADF family | actin filament depolymerization [GO:00300 | actin cytoskeleton [GO:0015629] | actin binding [GO:0003779] | actin cytoskeleton [GO:00 | UP000018817 |
| PPTG_02509 | W2RB71 | Adenylosuccinate lyase (ASL) (EC 4.3.2.2) (Adenylosuccinase) | 480 | | Lyase 1 family, Adenylosuccinate lyase subfamily | 'de novo' AMP biosynthetic process [GO:00 | cytosol [GO:0005829] | (S)-2-(5-amino-1-(5-phospho-D-ribosyl)imid | cytosol [GO:0005829]; (S)- | UP000018817 |
| PPTG_11897 | W2QB83 | 40S ribosomal protein S8 | 205 | | Eukaryotic ribosomal protein eS8 family | translation [GO:0006412] | ribonucleoprotein complex [GO:1990904]; rit | structural constituent of ribosome [GO:0003 | ribonucleoprotein complex | UP000018817 |
| PPTG_12407 | W2Q3V4 | Upf1 domain-containing protein | 1052 | | DNA2/NAM7 helicase family | nuclear-transcribed mRNA catabolic proces | cytoplasm [GO:0005737] | ATP binding [GO:0005524]; hydrolase activity | cytoplasm [GO:0005737]; | UP000018817 |
| PPTG_14134 | W2PZE1 | Anaphase-promoting complex subunit 4 WD40 domain-containing protein | 980 | | Glycosyltransferase 31 family | endoplasmic reticulum to Golgi vesicle-me | endoplasmic reticulum membrane [GO:0005 | guanyl-nucleotide exchange factor activity [ | endoplasmic reticulum me | UP000018817 |
| PPTG_02042 | W2R985 | 40S ribosomal protein S16 | 163 | | Universal ribosomal protein uS9 family | maturation of SSU-rRNA from tricistronic rR | cytosolic small ribosomal subunit [GO:00226 | RNA binding [GO:0003723]; structural consti | cytosolic small ribosomal | UP000018817 |
| PPTG_12191 | W2Q802 | Uncharacterized protein | 672 | | | | | | | UP000018817 |
| PPTG_13482 | W2Q1I5 | Apple domain-containing protein | 339 | | | carbohydrate metabolic process [GO:0005 | extracellular region [GO:0005576] | cellulose binding [GO:0030248] | extracellular region [GO:00 | UP000018817 |

| | | | | | | | | | | |
|---|---|---|---|---|---|---|---|---|---|---|
| PPTG_10070 | W2QDW6 | Intraflagellar transport protein 122 homolog | 1287 | | | intraciliary retrograde transport [GO:003572 | intraciliary transport particle A [GO:0030991]; non-motile cilium [GO:0097730] | | intraciliary transport partic | UP000018817 |
| PPTG_07910 | W2QPY1 | Proteasome subunit beta | 218 | | Peptidase T1B family | proteolysis involved in protein catabolic pro | cytoplasm [GO:0005737]; nucleus [GO:0005€ | threonine-type endopeptidase activity [GO:0 | cytoplasm [GO:0005737]; | UP000018817 |
| PPTG_10908 | W2QBI5 | Plectin/eS10 N-terminal domain-containing protein | 144 | | Eukaryotic ribosomal protein eS10 family | | cytosolic small ribosomal subunit [GO:00226 | RNA binding [GO:0003723]; structural consti | cytosolic small ribosomal | UP000018817 |
| PPTG_17135 | W2PN86 | CCR4-NOT transcription complex subunit 1 | 2372 | | | negative regulation of translation [GO:0017 | CCR4-NOT core complex [GO:0030015]; nucl | molecular adaptor activity [GO:0060090] | CCR4-NOT core complex [ | UP000018817 |
| PPTG_01560 | W2R7G5 | chorismate mutase (EC 5.4.99.5) | 558 | | | chorismate metabolic process [GO:004641 | cytoplasm [GO:0005737] | chorismate mutase activity [GO:0004106]; p | cytoplasm [GO:0005737]; | UP000018817 |
| PPTG_13586 | W2Q222 | Anoctamin transmembrane domain-containing protein | 1180 | | | | plasma membrane [GO:0005886] | calcium-activated cation channel activity [G | plasma membrane [GO:00 | UP000018817 |
| PPTG_08286 | W2QLR0 | Cytochrome c peroxidase, mitochondrial (EC 1.11.1.5) | 337 | | Peroxidase family | cellular response to oxidative stress [GO:00 | mitochondrial intermembrane space [GO:00 | cytochrome-c peroxidase activity [GO:00041 | mitochondrial intermembr | UP000018817 |
| PPTG_12903 | W2Q1C1 | Ribosomal protein/NADH dehydrogenase domain-containing protein | 148 | | Complex I NDUFA2 subunit family | | mitochondrial inner membrane [GO:0005743] | | mitochondrial inner memb | UP000018817 |
| PPTG_06552 | W2QTY4 | Eukaryotic translation initiation factor 6 (eIF-6) | 245 | | EIF-6 family | cytosolic ribosome assembly [GO:0042256] | cytoplasm [GO:0005737]; nucleolus [GO:000 | ribosomal large subunit binding [GO:004302 | cytoplasm [GO:0005737]; | UP000018817 |
| PPTG_02620 | W2RBU7 | aldehyde dehydrogenase (NAD(+)) (EC 1.2.1.3) | 535 | | Aldehyde dehydrogenase family | | | aldehyde dehydrogenase (NAD+) activity [GC | aldehyde dehydrogenase ( | UP000018817 |
| PPTG_10186 | W2QEG1 | Uncharacterized protein | 270 | | | | | | | UP000018817 |
| PPTG_09270 | W2QHU1 | DNA/RNA-binding protein Alba-like domain-containing protein | 319 | | | deoxyribonucleotide catabolic process [GO:0009264] | | 5'-nucleotidase activity [GO:0008253]; nucle | 5'-nucleotidase activity [G | UP000018817 |
| PPTG_01706 | W2R821 | Orotidine 5'-phosphate decarboxylase (EC 4.1.1.23) (OMP decarboxylase) | 391 | | OMP decarboxylase family, Type 2 subfamily | 'de novo' pyrimidine nucleobase biosynthetic process [GO:0006207]; 'de novo' UMP biosy | | orotate phosphoribosyltransferase activity [ | orotate phosphoribosyltrar | UP000018817 |
| PPTG_16727 | W2PPV9 | Hsp70-like protein | 217 | | | | | ATP binding [GO:0005524]; ATP-dependent p | ATP binding [GO:0005524] | UP000018817 |
| PPTG_08316 | W2QLX1 | Mitochondrial DNA replication protein YHM2 | 292 | | Mitochondrial carrier (TC 2.A.29) family | alpha-ketoglutarate transport [GO:0015742 | membrane [GO:0016020]; mitochondrion [G | tricarboxylate secondary active transmembr | membrane [GO:0016020]; | UP000018817 |
| PPTG_18250 | W2PJ44 | Casein kinase I (EC 2.7.11.1) | 402 | | Protein kinase superfamily, CK1 Ser/Thr protein kinase family, Casein kinase I subfamily | | | ATP binding [GO:0005524]; protein serine/th | ATP binding [GO:0005524] | UP000018817 |
| PPTG_11450 | W2QAJ9 | FYVE-type domain-containing protein | 597 | | | | | zinc ion binding [GO:0008270] | zinc ion binding [GO:00082 | UP000018817 |
| PPTG_06467 | W2QT19 | Lysine--tRNA ligase (EC 6.1.1.6) (Lysyl-tRNA synthetase) | 652 | | Class-II aminoacyl-tRNA synthetase family | lysyl-tRNA aminoacylation [GO:0006430] | cytosol [GO:0005829] | ATP binding [GO:0005524]; lysine-tRNA ligas | cytosol [GO:0005829]; ATF | UP000018817 |
| PPTG_06758 | W2QRB5 | Short chain dehydrogenase | 242 | | Short-chain dehydrogenases/reductases (SDR) family | | cytoplasm [GO:0005737] | oxidoreductase activity [GO:0016491] | cytoplasm [GO:0005737]; | UP000018817 |
| PPTG_12759 | W2Q0H9 | Uncharacterized protein | 1046 | | | | | | | UP000018817 |
| PPTG_09393 | W2QGN0 | Helicase | 1464 | | | nuclear-transcribed mRNA catabolic proces | Ski complex [GO:0055087] | ATP binding [GO:0005524]; helicase activity | Ski complex [GO:0055087 | UP000018817 |
| PPTG_14103 | W2PWM3 | SCP2 domain-containing protein | 249 | | | | cytosol [GO:0005829] | | cytosol [GO:0005829] | UP000018817 |
| PPTG_02850 | W2RCR5 | Interferon-related developmental regulator N-terminal domain-containing protein | 333 | | IFRD family | | | | | UP000018817 |
| PPTG_09918 | W2QCI2 | mannose-1-phosphate guanylyltransferase (EC 2.7.7.13) | 359 | | Transferase hexapeptide repeat family | GDP-mannose biosynthetic process [GO:0009298] | | GTP binding [GO:0005525]; mannose-1-phos | GTP binding [GO:0005525] | UP000018817 |
| PPTG_16221 | W2PP13 | Myosin motor domain-containing protein | 1259 | | TRAFAC class myosin-kinesin ATPase superfamily, | actin filament organization [GO:0007015] | cytoplasm [GO:0005737]; membrane [GO:00 | actin filament binding [GO:0051015]; ATP bir | cytoplasm [GO:0005737]; | UP000018817 |
| PPTG_11383 | W2QB73 | Prohibitin | 275 | | Prohibitin family | mitochondrion organization [GO:0007005] | mitochondrial inner membrane [GO:0005743] | | mitochondrial inner memb | UP000018817 |
| PPTG_10318 | W2QG07 | Large ribosomal subunit protein eL22 (60S ribosomal protein L22) | 125 | | Eukaryotic ribosomal protein eL22 family | cytoplasmic translation [GO:0002181] | cytoplasm [GO:0005737]; ribonucleoprotein | RNA binding [GO:0003723]; structural consti | cytoplasm [GO:0005737]; | UP000018817 |
| PPTG_00652 | W2RHU5 | Uncharacterized protein | 190 | | | P450-containing electron transport chain [G | mitochondrion [GO:0005739] | 2 iron, 2 sulfur cluster binding [GO:0051537] | mitochondrion [GO:00057 | UP000018817 |
| PPTG_07419 | W2QPZ1 | PDZ domain-containing protein | 1226 | | | | | lipid binding [GO:0008289] | lipid binding [GO:0008289 | UP000018817 |
| PPTG_11004 | W2Q6Z3 | 40S ribosomal protein S24 | 137 | | Eukaryotic ribosomal protein eS24 family | translation [GO:0006412] | ribonucleoprotein complex [GO:1990904]; rib | structural constituent of ribosome [GO:0003 | ribonucleoprotein comple | UP000018817 |
| PPTG_03815 | W2R0S0 | Glycine amidinotransferase, mitochondrial (EC 2.1.4.1) (L-arginine:glycine amidinotransferase) (Tr | 396 | | Amidinotransferase family | creatine biosynthetic process [GO:0006601 | mitochondrial inner membrane [GO:000574 | glycine amidinotransferase activity [GO:001 | mitochondrial inner memb | UP000018817 |
| PPTG_09934 | W2QF21 | Glycosyl hydrolase family 32 N-terminal domain-containing protein | 620 | | Glycosyl hydrolase 32 family | sucrose catabolic process [GO:0005987] | cytoplasm [GO:0005737] | sucrose alpha-glucosidase activity [GO:0004 | cytoplasm [GO:0005737]; | UP000018817 |
| PPTG_17633 | W2PKT7 | Ribosomal protein L25, Ctc-form | 251 | | | translation [GO:0006412] | cytosolic large ribosomal subunit [GO:00226 | 5S rRNA binding [GO:0008097]; structural co | cytosolic large ribosomal s | UP000018817 |
| PPTG_19057 | W2PFT4 | Aminomethyltransferase (EC 2.1.2.10) (Glycine cleavage system T protein) | 391 | | GcvT family | glycine catabolic process [GO:0006546] | glycine cleavage complex [GO:0005960]; mit | aminomethyltransferase activity [GO:00040 | glycine cleavage complex [ | UP000018817 |
| PPTG_18216 | W2PH65 | Cilia- and flagella-associated protein 52 | 702 | | CFAP52 family | | axoneme [GO:0005930]; motile cilium [GO:0031514] | | axoneme [GO:0005930]; n | UP000018817 |
| PPTG_05195 | W2QVY7 | J domain-containing protein | 606 | | | post-translational protein targeting to endo | Sec62/Sec63 complex [GO:0031207] | protein transmembrane transporter activity [ | Sec62/Sec63 complex [GC | UP000018817 |
| PPTG_11829 | W2Q8E8 | Tetratricopeptide repeat protein | 1287 | | TTC21 family | intraciliary retrograde transport [GO:003572 | cilium [GO:0005929]; intraciliary transport particle A [GO:0030991] | | cilium [GO:0005929]; intra | UP000018817 |
| PPTG_08040 | W2QJM4 | Acetyl-coenzyme A synthetase (EC 6.2.1.1) | 595 | | ATP-dependent AMP-binding enzyme family | acetyl-CoA biosynthetic process from acetate [GO:0019427] | | acetate-CoA ligase activity [GO:0003987]; A | acetate-CoA ligase activity | UP000018817 |
| PPTG_12094 | W2Q556 | Beta-mannosidase B (EC 3.2.1.25) (Mannanase B) | 937 | | Glycosyl hydrolase 2 family, Beta-mannosidase B s | carbohydrate metabolic process [GO:0005 | extracellular region [GO:0005576] | beta-mannosidase activity [GO:0004567] | extracellular region [GO:00 | UP000018817 |
| PPTG_18748 | W2PEL2 | HID1 domain-containing protein | 740 | | | | Golgi medial cisterna [GO:0005797]; Golgi trans cisterna [GO:0000138]; membrane [GO:0 | | Golgi medial cisterna [GO: | UP000018817 |
| PPTG_10656 | W2QBK0 | Elicitor-like transglutaminase | 654 | | | | | aminoacyltransferase activity [GO:0016755] | aminoacyltransferase acti | UP000018817 |
| PPTG_02675 | W2RC29 | Glutamine-dependent NAD(+) synthetase (EC 6.3.5.1) (NAD(+) synthase [glutamine-hydrolyzing]) | 734 | | NAD synthetase family | NAD+ biosynthetic process [GO:0009435] | cytoplasm [GO:0005737] | ATP binding [GO:0005524]; glutaminase acti | cytoplasm [GO:0005737]; | UP000018817 |
| PPTG_15910 | W2PRY4 | Proteasome subunit beta | 231 | | Peptidase T1B family | proteolysis involved in protein catabolic pro | cytoplasm [GO:0005737]; nucleus [GO:0005634]; proteasome core complex, beta-subuni | | cytoplasm [GO:0005737]; | UP000018817 |
| PPTG_06210 | W2QRY0 | Propionyl-CoA synthetase | 606 | | | | | propionate-CoA ligase activity [GO:0050218 | propionate-CoA ligase acti | UP000018817 |
| PPTG_05918 | W2QUG2 | 3-isopropylmalate dehydratase large subunit | 709 | | | L-amino acid biosynthetic process [GO:0170034]; proteinogenic amino acid biosynthetic | | hydro-lyase activity [GO:0016836]; iron-sulfu | hydro-lyase activity [GO:00 | UP000018817 |
| PPTG_02297 | W2RAA6 | SCP domain-containing protein | 152 | | | | | | | UP000018817 |
| PPTG_11571 | W2Q7B0 | Transcriptional repressor Tup1 N-terminal domain-containing protein | 725 | | | | | | | UP000018817 |
| PPTG_15123 | W2PWS4 | Uncharacterized protein | 546 | | | | endomembrane system [GO:0012505]; membrane [GO:0016020] | | endomembrane system [G | UP000018817 |
| PPTG_09469 | W2QF18 | Phospholipid/glycerol acyltransferase domain-containing protein | 728 | | | phospholipid biosynthetic process [GO:0008654] | | glycerol-3-phosphate O-acyltransferase acti | glycerol-3-phosphate O-ac | UP000018817 |
| PPTG_15525 | W2PPJ4 | Uncharacterized protein | 136 | | Eukaryotic ribosomal protein eL14 family | ribosomal large subunit biogenesis [GO:004 | chloroplast [GO:0009507]; cytosolic large rib | RNA binding [GO:0003723]; structural consti | chloroplast [GO:0009507]; | UP000018817 |
| PPTG_16775 | W2PML7 | Adenosine kinase (AK) (EC 2.7.1.20) (Adenosine 5'-phosphotransferase) | 368 | | Carbohydrate kinase PfkB family | AMP salvage [GO:0044209]; purine nucleob | cytosol [GO:0005829]; nucleus [GO:0005634 | adenosine kinase activity [GO:0004001]; ATF | cytosol [GO:0005829]; nuc | UP000018817 |
| PPTG_13862 | W2Q180 | V-type ATPase, D subunit | 260 | | V-ATPase D subunit family | | | proton-transporting ATPase activity, rotation | proton-transporting ATPas | UP000018817 |
| PPTG_07617 | W2QNA1 | Tyrosinase copper-binding domain-containing protein | 544 | | | | | metal ion binding [GO:0046872]; oxidoreduc | metal ion binding [GO:004 | UP000018817 |
| PPTG_13731 | W2Q0I7 | 1-phosphatidylinositol 4-kinase (EC 2.7.1.67) | 801 | | | phosphatidylinositol phosphate biosyntheti | cytoplasm [GO:0005737]; membrane [GO:00 | 1-phosphatidylinositol 4-kinase activity [GO: | cytoplasm [GO:0005737]; | UP000018817 |
| PPTG_18717 | W2PFP3 | 40S ribosomal protein S23 | 145 | | Universal ribosomal protein uS12 family | translation [GO:0006412] | small ribosomal subunit [GO:0015935] | structural constituent of ribosome [GO:0003 | small ribosomal subunit [G | UP000018817 |
| PPTG_03967 | W2R0V7 | Large ribosomal subunit protein uL1c | 492 | | Universal ribosomal protein uL1 family | translation [GO:0006412] | large ribosomal subunit [GO:0015934] | rRNA binding [GO:0019843]; structural const | large ribosomal subunit [G | UP000018817 |
| PPTG_09242 | W2QIN1 | NADH dehydrogenase [ubiquinone] iron-sulfur protein 4, mitochondrial | 242 | | Complex I NDUFS4 subunit family | electron transport chain [GO:0022900] | mitochondrial inner membrane [GO:0005743] | | mitochondrial inner memb | UP000018817 |
| PPTG_15752 | W2PRD0 | Prohibitin | 299 | | Prohibitin family | mitochondrion organization [GO:0007005] | mitochondrial inner membrane [GO:0005743] | | mitochondrial inner memb | UP000018817 |
| PPTG_11140 | W2Q962 | Importin N-terminal domain-containing protein | 1114 | | Exportin family | protein export from nucleus [GO:0006611] | cytoplasm [GO:0005737]; nuclear pore [GO:0 | nuclear export signal receptor activity [GO:0 | cytoplasm [GO:0005737]; | UP000018817 |
| PPTG_16175 | W2PQM0 | Magnesium transporter protein 1 | 357 | | OST3/OST6 family | protein N-linked glycosylation via asparagin | oligosaccharyltransferase complex [GO:0008250] | | oligosaccharyltransferase | UP000018817 |
| PPTG_11732 | W2Q8W3 | Orn/DAP/Arg decarboxylase 2 N-terminal domain-containing protein | 527 | | | lysine biosynthetic process via diaminopimelate [GO:0009089]; polyamine biosynthetic p | | diaminopimelate decarboxylase activity [GO | diaminopimelate decarbo | UP000018817 |
| PPTG_15500 | W2PP26 | Serine-threonine kinase receptor-associated protein | 341 | | WD repeat STRAP family | spliceosomal snRNP assembly [GO:000038 | SMN complex [GO:0032797] | RNA binding [GO:0003723] | SMN complex [GO:003279 | UP000018817 |
| PPTG_09132 | W2QJ39 | 40S ribosomal protein S13 | 151 | | Universal ribosomal protein uS15 family | translation [GO:0006412] | cytosolic small ribosomal subunit [GO:00226 | small ribosomal subunit rRNA binding [GO:0 | cytosolic small ribosomal | UP000018817 |
| PPTG_16307 | W2PQW0 | RGS domain-containing protein | 339 | | | | | | | UP000018817 |
| PPTG_16153 | W2PT70 | 40S ribosomal protein S18 | 152 | | Universal ribosomal protein uS13 family | translation [GO:0006412] | cytosol [GO:0005829]; small ribosomal subu | RNA binding [GO:0003723]; structural consti | cytosol [GO:0005829]; sm | UP000018817 |
| PPTG_19298 | W2PDK5 | 26S proteasome non-ATPase regulatory subunit 14 | 311 | | | proteolysis [GO:0006508] | proteasome complex [GO:0000502] | metal ion binding [GO:0046872]; metallope | proteasome complex [GO: | UP000018817 |
| PPTG_02483 | W2RB30 | Uncharacterized protein | 379 | | WD repeat PROPPIN family | | cytoplasm [GO:0005737] | | cytoplasm [GO:0005737] | UP000018817 |
| PPTG_05693 | W2QTR9 | Pre-mRNA-splicing helicase BRR2 | 2292 | | | mRNA processing [GO:0006397] | nucleus [GO:0005634] | ATP binding [GO:0005524]; helicase activity | nucleus [GO:0005634]; AT | UP000018817 |
| PPTG_17494 | W2PJ62 | 60S ribosomal protein L30 | 112 | | Eukaryotic ribosomal protein eL30 family | | cytosolic large ribosomal subunit [GO:00226 | RNA binding [GO:0003723]; structural consti | cytosolic large ribosomal s | UP000018817 |
| PPTG_03256 | W2R454 | Cytochrome c oxidase subunit | 84 | | | | mitochondrion [GO:0005739]; respiratory chain complex IV [GO:0045277] | | mitochondrion [GO:00057 | UP000018817 |
| PPTG_18612 | W2PGS0 | Trehalase (EC 3.2.1.28) (Alpha-trehalose glucohydrolase) | 638 | | Glycosyl hydrolase 37 family | trehalose catabolic process [GO:0005993] | | alpha,alpha-trehalase activity [GO:0004555] | alpha,alpha-trehalase acti | UP000018817 |
| PPTG_08373 | W2QL13 | S1 motif domain-containing protein | 332 | | EIF-2-alpha family | | eukaryotic 48S preinitiation complex [GO:00 | ribosome binding [GO:0043022]; RNA bindin | eukaryotic 48S preinitiatio | UP000018817 |
| PPTG_03695 | W2R7W2 | Uncharacterized protein | 723 | | | | plasma membrane [GO:0005886] | chloride channel activity [GO:0005254]; prot | plasma membrane [GO:00 | UP000018817 |
| PPTG_10236 | W2QDS3 | Carrier domain-containing protein | 712 | | ATP-dependent AMP-binding enzyme family | fatty acid metabolic process [GO:0006631]; lipid biosynthetic process [GO:0008610] | | ligase activity [GO:0016874]; phosphopante | ligase activity [GO:001687 | UP000018817 |
| PPTG_15937 | W2PU35 | beta-glucosidase (EC 3.2.1.21) | 514 | | Glycosyl hydrolase 3 family | glucan catabolic process [GO:0009251] | | beta-glucosidase activity [GO:0008422] | beta-glucosidase activity [ | UP000018817 |
| PPTG_02290 | W2RAS9 | Uncharacterized protein | 129 | | | | | | | UP000018817 |
| PPTG_08145 | W2QM10 | 4-aminobutyrate transaminase | 420 | | Class-III pyridoxal-phosphate-dependent aminotransferase family | | | identical protein binding [GO:0042802]; pyric | identical protein binding [G | UP000018817 |
| PPTG_10143 | W2QDC4 | Uncharacterized protein | 1014 | | Saccharopine dehydrogenase family | L-lysine catabolic process to acetyl-CoA via | cytoplasm [GO:0005737] | saccharopine dehydrogenase activity [GO:00 | cytoplasm [GO:0005737]; | UP000018817 |
| PPTG_00708 | W2RGK8 | Importin subunit alpha | 532 | | Importin alpha family | protein import into nucleus [GO:0006606] | cytoplasm [GO:0005737]; nucleus [GO:0005 | nuclear import signal receptor activity [GO:0 | cytoplasm [GO:0005737]; | UP000018817 |
| PPTG_09327 | W2QIZ7 | Delta-aminolevulinic acid dehydratase (EC 4.2.1.24) | 327 | | ALAD family | protoporphyrinogen IX biosynthetic process | cytosol [GO:0005829] | porphobilinogen synthase activity [GO:0004 | cytosol [GO:0005829]; por | UP000018817 |
| PPTG_00418 | W2REZ3 | Uncharacterized protein | 438 | | | | | | | UP000018817 |
| PPTG_00608 | W2RFL0 | asparagine synthase (glutamine-hydrolyzing) (EC 6.3.5.4) | 563 | | | asparagine biosynthetic process [GO:0006 | cytosol [GO:0005829] | asparagine synthase (glutamine-hydrolyzing) | cytosol [GO:0005829]; asp | UP000018817 |
| PPTG_16326 | W2PQR0 | Tryptophan biosynthesis protein TRP1 | 561 | | | L-tryptophan biosynthetic process [GO:0000162] | | indole-3-glycerol-phosphate synthase activit | indole-3-glycerol-phospha | UP000018817 |
| PPTG_18241 | W2PHW9 | Inositol polyphosphate-related phosphatase domain-containing protein | 515 | | | phosphatidylinositol dephosphorylation [GO:0046856] | | phosphatidylinositol-4,5-bisphosphate 5-ph | phosphatidylinositol-4,5-b | UP000018817 |
| PPTG_16234 | W2PQU8 | Transglutaminase elicitor | 768 | | | | | aminoacyltransferase activity [GO:0016755] | aminoacyltransferase acti | UP000018817 |
| PPTG_19310 | W2PCG6 | Uncharacterized protein | 118 | | | | | | | UP000018817 |
| PPTG_06097 | W2QV93 | Dipeptidase | 662 | | Peptidase C69 family, Secernin subfamily | proteolysis [GO:0006508] | | cysteine-type exopeptidase activity [GO:007 | cysteine-type exopeptidas | UP000018817 |
| PPTG_06606 | W2QQG1 | Uncharacterized protein | 466 | | | | | | | UP000018817 |
| PPTG_13151 | W2PYN4 | Transmembrane 9 superfamily member | 643 | | Nonaspanin (TM9SF) (TC 9.A.2) family | protein localization to membrane [GO:0072 | membrane [GO:0016020] | | membrane [GO:0016020]; | UP000018817 |
| PPTG_14216 | W2PX04 | Proteasome subunit beta type-3 | 191 | | | proteasome-mediated ubiquitin-dependent | cytoplasm [GO:0005737]; nucleus [GO:0005634]; proteasome core complex, beta-subuni | | cytoplasm [GO:0005737]; | UP000018817 |
| PPTG_03220 | W2R402 | EGF-like domain-containing protein | 536 | | Glycosyl hydrolase 5 (cellulase A) family | cellulose catabolic process [GO:0030245] | | hydrolase activity, hydrolyzing O-glycosyl cor | hydrolase activity, hydrolyz | UP000018817 |
| PPTG_13960 | W2PZ75 | AP-3 complex subunit delta | 1004 | | Adaptor complexes large subunit family | Golgi to vacuole transport [GO:0006896]; pr | AP-3 adaptor complex [GO:0030123]; endosome membrane [GO:0010008]; Golgi apparat | | AP-3 adaptor complex [GO | UP000018817 |
| PPTG_02092 | W2RBW8 | Peptidyl-prolyl cis-trans isomerase (PPIase) (EC 5.2.1.8) | 206 | | Cyclophilin-type PPIase family | protein folding [GO:0006457] | cytoplasm [GO:0005737] | cyclosporin A binding [GO:0016018]; peptidy | cytoplasm [GO:0005737]; | UP000018817 |
| PPTG_02293 | W2RCE6 | Malic enzyme | 622 | | Malic enzymes family | malate metabolic process [GO:0006108] | mitochondrion [GO:0005739] | malate dehydrogenase (decarboxylating) (N | mitochondrion [GO:00057 | UP000018817 |
| PPTG_04798 | W2R2N4 | FAD/NAD(P)-binding domain-containing protein | 394 | | FAD-dependent oxidoreductase family | | cytoplasm [GO:0005737] | electron-transferring-flavoprotein dehydroge | cytoplasm [GO:0005737]; | UP000018817 |
| PPTG_04800 | W2R474 | FAD/NAD(P)-binding domain-containing protein | 394 | | FAD-dependent oxidoreductase family | | cytoplasm [GO:0005737] | electron-transferring-flavoprotein dehydroge | cytoplasm [GO:0005737]; | UP000018817 |
| PPTG_02740 | W2REQ2 | Glutathione S-transferase | 230 | | | glutathione metabolic process [GO:0006749] | | glutathione transferase activity [GO:000436 | glutathione transferase ac | UP000018817 |
| PPTG_02823 | W2RER0 | EGF-like domain-containing protein | 407 | Welsh et al., 2024 | | | | | | UP000018817 |
| PPTG_16771 | W2PM91 | Cystathionine beta-synthase (EC 4.2.1.22) | 513 | | Cysteine synthase/cystathionine beta-synthase fa | cysteine biosynthetic process from serine [ | cytoplasm [GO:0005737]; nucleus [GO:0005 | cystathionine beta-synthase activity [GO:00 | cytoplasm [GO:0005737]; | UP000018817 |
| PPTG_00422 | W2REZ8 | J domain-containing protein | 420 | | | 'de novo' cotranslational protein folding [GC | cytosol [GO:0005829] | Hsp70 protein binding [GO:0030544]; riboso | cytosol [GO:0005829]; Hsp | UP000018817 |
| PPTG_02494 | W2RD39 | 2-hydroxy-3-oxopropionate reductase | 300 | | | | | NAD binding [GO:0051287]; NADP binding [G | NAD binding [GO:0051287 | UP000018817 |

| | | | | | | | | | |
|---|---|---|---|---|---|---|---|---|---|
| PPTG_17678 | W2PLC5 | Croquemort-like mating protein M82 | 724 | CD36 family | | cytoplasm [GO:0005737]; membrane [GO:00 | scavenger receptor activity [GO:0005044] | cytoplasm [GO:0005737]; | UP000018817 |
| PPTG_08439 | W2QMF0 | Guanylate cyclase domain-containing protein | 2628 | | cyclic nucleotide biosynthetic process [GO: | plasma membrane [GO:0005886] | ATP binding [GO:0005524]; ATP hydrolysis ac | plasma membrane [GO:00 | UP000018817 |
| PPTG_00131 | W2RDP2 | Large ribosomal subunit protein mL46 | 297 | Mitochondrion-specific ribosomal protein mL46 family | | mitochondrial inner membrane [GO:000574 | structural constituent of ribosome [GO:0003 | mitochondrial inner memb | UP000018817 |
| PPTG_18713 | W2PFN8 | Phosphodiesterase (EC 3.1.4.-) | 1031 | Cyclic nucleotide phosphodiesterase family | signal transduction [GO:0007165] | | 3',5'-cyclic-nucleotide phosphodiesterase ac | 3',5'-cyclic-nucleotide pho | UP000018817 |
| PPTG_04233 | W2R0F2 | UDP-glucose:glycoprotein glucosyltransferase | 1553 | | ERAD pathway [GO:0036503]; protein N-linl | endoplasmic reticulum lumen [GO:0005788] | UDP-glucose:glycoprotein glucosyltransferas | endoplasmic reticulum lur | UP000018817 |
| PPTG_11812 | W2Q8B0 | Chromodomain-helicase-DNA-binding protein 7 | 2105 | | | chromatin [GO:0000785]; nucleus [GO:0005 | ATP binding [GO:0005524]; ATP hydrolysis ac | chromatin [GO:0000785]; | UP000018817 |
| PPTG_01724 | W2RA45 | TKL protein kinase | 734 | | | | ATP binding [GO:0005524]; protein serine/th | ATP binding [GO:0005524] | UP000018817 |
| PPTG_09026 | W2QHT8 | Mitochondrial carrier protein | 304 | Mitochondrial carrier (TC 2.A.29) family | | mitochondrial inner membrane [GO:000574 | guanine nucleotide transmembrane transpo | mitochondrial inner memb | UP000018817 |
| PPTG_15024 | W2PWF7 | Lysosomal Pro-X carboxypeptidase | 602 | Peptidase S28 family | proteolysis [GO:0006508] | | dipeptidyl-peptidase activity [GO:0008239]; | dipeptidyl-peptidase activi | UP000018817 |
| PPTG_02253 | W2RCN5 | Glutamate/phenylalanine/leucine/valine/L-tryptophan dehydrogenase C-terminal domain-containi | 1054 | Glu/Leu/Phe/Val dehydrogenases family | L-glutamate catabolic process [GO:000653 | mitochondrion [GO:0005739] | glutamate dehydrogenase (NAD+) activity [G | mitochondrion [GO:00057 | UP000018817 |
| PPTG_04743 | W2R2E7 | Saposin B-type domain-containing protein | 959 | | | | | | UP000018817 |
| PPTG_04076 | W2R1U5 | FYVE-type domain-containing protein | 446 | | | | zinc ion binding [GO:0008270] | zinc ion binding [GO:0008 | UP000018817 |
| PPTG_14756 | W2PVS9 | Uncharacterized protein | 838 | | | | | | UP000018817 |
| PPTG_17882 | W2PJ70 | Dynamin N-terminal domain-containing protein | 527 | | | | | | UP000018817 |
| PPTG_09467 | W2QFX4 | Serine/threonine-protein phosphatase (EC 3.1.3.16) | 309 | PPP phosphatase family | | cytoplasm [GO:0005737]; nucleus [GO:0005 | metal ion binding [GO:0046872]; protein seri | cytoplasm [GO:0005737]; | UP000018817 |
| PPTG_08321 | W2QLX5 | EF-hand domain-containing protein | 263 | | calcium-mediated signaling [GO:0019722] | | calcium ion binding [GO:0005509]; kinase bi | calcium ion binding [GO:0 | UP000018817 |
| PPTG_13691 | W2Q257 | Peptidase M13 C-terminal domain-containing protein | 1179 | Peptidase M13 family | protein processing [GO:0016485] | plasma membrane [GO:0005886] | metal ion binding [GO:0046872]; metalloenc | plasma membrane [GO:00 | UP000018817 |
| PPTG_02703 | W2REK3 | 3-hydroxyacyl-CoA dehydrogenase type-2 (EC 1.1.1.53) (3-hydroxyacyl-CoA dehydrogenase type II) | 256 | Short-chain dehydrogenases/reductases (SDR) family | | | (3S)-3-hydroxyacyl-CoA dehydrogenase (NAI | (3S)-3-hydroxyacyl-CoA de | UP000018817 |
| PPTG_16216 | W2PPP8 | PLC-like phosphodiesterase | 909 | | lipid metabolic process [GO:0006629] | membrane [GO:0016020] | phosphoric diester hydrolase activity [GO:00 | membrane [GO:0016020]; | UP000018817 |
| PPTG_12683 | W2Q0G2 | Serine/threonine protein kinase | 594 | | | extracellular region [GO:0005576]; host cell [ | ATP binding [GO:0005524]; protein serine/th | extracellular region [GO:00 | UP000018817 |
| PPTG_07242 | W2QPI3 | Electron transfer flavoprotein subunit alpha (Alpha-ETF) | 330 | ETF alpha-subunit/FixB family | fatty acid beta-oxidation using acyl-CoA del | mitochondrial matrix [GO:0005759] | electron transfer activity [GO:0009055]; flavi | mitochondrial matrix [GO: | UP000018817 |
| PPTG_11618 | W2Q6J2 | alanine--glyoxylate transaminase (EC 2.6.1.44) | 469 | Class-V pyridoxal-phosphate-dependent aminotrar | glycine biosynthetic process, by transamina | peroxisome [GO:0005777] | alanine-glyoxylate transaminase activity [GC | peroxisome [GO:0005777] | UP000018817 |
| PPTG_18786 | W2PE21 | Methyltransferase type 11 domain-containing protein | 279 | | | | S-adenosylmethionine-dependent methyltra | S-adenosylmethionine-de | UP000018817 |
| PPTG_06692 | W2QSE9 | Bacterial Pleckstrin homology domain-containing protein | 210 | | | | | | UP000018817 |
| PPTG_09412 | W2QGQ8 | Hexose transporter 1 | 511 | Major facilitator superfamily, Sugar transporter (TC 2.A.1.1) family | | membrane [GO:0016020] | carbohydrate:proton symporter activity [GO: | membrane [GO:0016020]; | UP000018817 |
| PPTG_02446 | W2RAR6 | Proteasome subunit alpha type | 253 | Peptidase T1A family | ubiquitin-dependent protein catabolic proc | cytoplasm [GO:0005737]; nucleus [GO:0005634]; proteasome core complex, alpha-subur | | cytoplasm [GO:0005737]; | UP000018817 |
| PPTG_15671 | W2PT91 | Nop domain-containing protein | 496 | NOP5/NOP56 family | ribosome biogenesis [GO:0042254] | box C/D methylation guide snoRNP complex | | snoRNA binding [GO:0030515] | box C/D methylation guide | UP000018817 |
| PPTG_12288 | W2Q5U7 | Uncharacterized protein | 416 | | | | | | UP000018817 |
| PPTG_05319 | W2QZ40 | Ribosomal protein L15 | 206 | Eukaryotic ribosomal protein eL15 family | cytoplasmic translation [GO:0002181] | cytosolic large ribosomal subunit [GO:0022 | RNA binding [GO:0003723]; structural consti | cytosolic large ribosomal s | UP000018817 |
| PPTG_05362 | W2QYR0 | CTP synthase (EC 6.3.4.2) (UTP--ammonia ligase) | 580 | CTP synthase family | 'de novo' CTP biosynthetic process [GO:0044210]; pyrimidine nucleobase biosynthetic pr | | ATP binding [GO:0005524]; CTP synthase ac | ATP binding [GO:0005524] | UP000018817 |
| PPTG_17182 | W2PL95 | Chlorophyll synthesis pathway protein BchC | 359 | Zinc-containing alcohol dehydrogenase family | sorbitol catabolic process [GO:0006062] | | L-iditol 2-dehydrogenase (NAD+) activity [GC | L-iditol 2-dehydrogenase ( | UP000018817 |
| PPTG_05388 | W2QYU8 | Protein-serine/threonine kinase (EC 2.7.11.-) | 380 | PDK/BCKDK protein kinase family | regulation of glucose metabolic process [G | mitochondrial matrix [GO:0005759] | ATP binding [GO:0005524]; pyruvate dehydro | mitochondrial matrix [GO: | UP000018817 |
| PPTG_07843 | W2QM28 | Uncharacterized protein | 359 | | | | | | UP000018817 |
| PPTG_15023 | W2PTU0 | UTP--glucose-1-phosphate uridylyltransferase (EC 2.7.7.9) | 455 | UDPGP type 1 family | UDP-alpha-D-glucose metabolic process [GO:0006011] | | UTP:glucose-1-phosphate uridylyltransferase | UTP:glucose-1-phosphate | UP000018817 |
| PPTG_00887 | W2RJ71 | DNA damage-binding protein 1 | 1148 | DDB1 family | | nucleus [GO:0005634] | nucleic acid binding [GO:0003676] | nucleus [GO:0005634]; nu | UP000018817 |
| PPTG_14718 | W2PWH7 | Amidophosphoribosyltransferase (ATase) (EC 2.4.2.14) (Glutamine phosphoribosylpyrophosphate | 519 | Purine/pyrimidine phosphoribosyltransferase famil | 'de novo' IMP biosynthetic process [GO:0006189]; purine nucleobase biosynthetic proces | | amidophosphoribosyltransferase activity [G | amidophosphoribosyltrans | UP000018817 |
| PPTG_10266 | W2QDW4 | Letm1 RBD domain-containing protein | 333 | | intracellular monoatomic cation homeosta | mitochondrial inner membrane [GO:000574 | ribosome binding [GO:0043022] | mitochondrial inner memb | UP000018817 |
| PPTG_15563 | W2PRM0 | glucan endo-1,3-beta-D-glucosidase (EC 3.2.1.39) (Endo-1,3-beta-glucanase btgC) (Laminarinase | 384 | Glycosyl hydrolase 17 family | cell wall organization [GO:0071555]; polysa | plasma membrane [GO:0005886] | glucan endo-1,3-beta-D-glucosidase activity | plasma membrane [GO:00 | UP000018817 |
| PPTG_16453 | W2PR79 | FYVE-type domain-containing protein | 395 | | | | | | UP000018817 |
| PPTG_18886 | W2PHZ4 | fructokinase (EC 2.7.1.4) | 294 | ROK (NagC/XylR) family | | | ATP binding [GO:0005524]; fructokinase acti | ATP binding [GO:0005524] | UP000018817 |
| PPTG_15025 | W2PUR2 | Ribophorin II (Ribophorin-2) | 393 | SWP1 family | protein N-linked glycosylation [GO:0006487 | oligosaccharyltransferase complex [GO:0008250] | | oligosaccharyltransferase | UP000018817 |
| PPTG_10392 | W2QF76 | MIF4G domain-containing protein | 893 | | mRNA export from nucleus [GO:0006406]; r | nuclear cap binding complex [GO:0005846]; | mRNA binding [GO:0003729]; RNA cap bindi | nuclear cap binding compl | UP000018817 |
| PPTG_14382 | W2PXM1 | C2 domain-containing protein | 383 | | | | | | UP000018817 |
| PPTG_08786 | W2QKE1 | ER membrane protein complex subunit 1 | 994 | EMC1 family | protein folding in endoplasmic reticulum [G | EMC complex [GO:0072546] | | EMC complex [GO:007254 | UP000018817 |
| PPTG_09275 | W2QHU6 | NAD(P)H:quinone oxidoreductase, type IV | 200 | WrbA family | | membrane [GO:0016020] | FMN binding [GO:0010181]; NAD(P)H dehydı | membrane [GO:0016020]; | UP000018817 |
| PPTG_17513 | W2PJN4 | RNA helicase (EC 3.6.4.13) | 411 | DEAD box helicase family, DDX6/DHH1 subfamily | | P-body [GO:0000932] | ATP binding [GO:0005524]; hydrolase activity | P-body [GO:0000932]; ATP | UP000018817 |
| PPTG_02681 | W2RE38 | Alpha-amylase (EC 3.2.1.1) | 865 | Glycosyl hydrolase 13 family | carbohydrate metabolic process [GO:0005975] | | alpha-amylase activity [GO:0004556]; cation | alpha-amylase activity [GC | UP000018817 |
| PPTG_04127 | W2R1G2 | TKL/DRK protein kinase | 952 | | | | ATP binding [GO:0005524]; protein serine/th | ATP binding [GO:0005524] | UP000018817 |
| PPTG_09787 | W2QDT4 | Glutathione reductase (EC 1.8.1.7) | 493 | Class-I pyridine nucleotide-disulfide oxidoreductas | cell redox homeostasis [GO:0045454]; cellu | cytosol [GO:0005829]; mitochondrion [GO:0( | flavin adenine dinucleotide binding [GO:005( | cytosol [GO:0005829]; mit | UP000018817 |
| PPTG_02066 | W2R9U3 | TKL protein kinase | 305 | | | | ATP binding [GO:0005524]; protein serine/th | ATP binding [GO:0005524] | UP000018817 |
| PPTG_09121 | W2QJ22 | NAD(P)(+) transhydrogenase (Si-specific) (EC 1.6.1.1) (NAD(P)(+) transhydrogenase [B-specific]) | 520 | Class-I pyridine nucleotide-disulfide oxidoreductas | 2-oxoglutarate metabolic process [GO:0006 | cytosol [GO:0005829] | dihydrolipoyl dehydrogenase (NADH) activity | cytosol [GO:0005829]; dihy | UP000018817 |
| PPTG_08095 | W2QJ64 | Carbonic anhydrase (EC 4.2.1.1) (Carbonate dehydratase) | 317 | Beta-class carbonic anhydrase family | carbon utilization [GO:0015976] | | carbonate dehydratase activity [GO:000408 | carbonate dehydratase ac | UP000018817 |
| PPTG_13316 | W2Q2N8 | Complex 1 LYR protein domain-containing protein | 128 | Complex I LYR family | response to oxidative stress [GO:0006979] | mitochondrial inner membrane [GO:0005743]; respiratory chain complex I [GO:0045271] | | mitochondrial inner memb | UP000018817 |
| PPTG_11616 | W2Q7I0 | RING-type domain-containing protein | 517 | TUB family | | | zinc ion binding [GO:0008270] | zinc ion binding [GO:0008 | UP000018817 |
| PPTG_07985 | W2QQ53 | Uncharacterized protein | 204 | | | | | | UP000018817 |
| PPTG_00778 | W2RGA2 | Enoyl reductase (ER) domain-containing protein | 330 | | | | oxidoreductase activity [GO:0016491]; zinc i | oxidoreductase activity [G( | UP000018817 |
| PPTG_00066 | W2RFL9 | Uncharacterized protein | 329 | | | | | | UP000018817 |
| PPTG_05956 | W2QVB0 | DNA sliding clamp PCNA | 259 | PCNA family | leading strand elongation [GO:0006272]; m | PCNA complex [GO:0043626] | DNA binding [GO:0003677]; DNA polymerase | PCNA complex [GO:00436 | UP000018817 |
| PPTG_03693 | W2R5V9 | protein-serine/threonine phosphatase (EC 3.1.3.16) | 1138 | PP2C family | | cAMP-dependent protein kinase complex [G | ATP binding [GO:0005524]; cAMP-dependen | cAMP-dependent protein k | UP000018817 |
| PPTG_05343 | W2QWG2 | Uncharacterized protein | 251 | | | | | | UP000018817 |
| PPTG_13440 | W2Q1B9 | Aminotransferase class I/classII large domain-containing protein | 499 | Class-I pyridoxal-phosphate-dependent aminotran | L-alanine catabolic process [GO:0042853] | | L-alanine:2-oxoglutarate aminotransferase a | L-alanine:2-oxoglutarate a | UP000018817 |
| PPTG_09762 | W2QEF4 | Transmembrane protein | 945 | | | | | | UP000018817 |
| PPTG_13259 | W2PZ37 | HECT domain-containing protein | 5009 | | | | ubiquitin-protein transferase activity [GO:00 | ubiquitin-protein transfera | UP000018817 |
| PPTG_02677 | W2REI0 | Calcium-transporting ATPase (EC 7.2.2.10) | 1040 | Cation transport ATPase (P-type) (TC 3.A.3) family | | endomembrane system [GO:0012505]; plasr | ATP binding [GO:0005524]; ATP hydrolysis ac | endomembrane system [G | UP000018817 |
| PPTG_04373 | W2R0P5 | Phosphoribosyltransferase domain-containing protein | 308 | | | | | | UP000018817 |
| PPTG_18069 | W2PJH1 | Ribosomal protein S6 | 194 | Bacterial ribosomal protein bS6 family | translation [GO:0006412] | cytoplasm [GO:0005737]; ribosome [GO:000 | small ribosomal subunit rRNA binding [GO:0 | cytoplasm [GO:0005737]; | UP000018817 |
| PPTG_05458 | W2QZN5 | 60S ribosomal protein L23a | 150 | Universal ribosomal protein uL23 family | translation [GO:0006412] | ribonucleoprotein complex [GO:1990904]; rit | rRNA binding [GO:0019843]; structural const | ribonucleoprotein comple | UP000018817 |
| PPTG_18551 | W2PER3 | Uncharacterized protein | 785 | | | cytoplasm [GO:0005737] | | cytoplasm [GO:0005737] | UP000018817 |
| PPTG_03134 | W2R3T1 | Acetyltransferase component of pyruvate dehydrogenase complex (EC 2.3.1.12) | 480 | 2-oxoacid dehydrogenase family | pyruvate decarboxylation to acetyl-CoA [GO | mitochondrion [GO:0005739]; pyruvate dehy | dihydrolipoyllysine-residue acetyltransferase | mitochondrion [GO:00057 | UP000018817 |
| PPTG_07468 | W2QN89 | Serine/threonine protein kinase | 525 | | cell surface receptor signaling pathway [GO:0007166] | | ATP binding [GO:0005524]; protein serine/th | ATP binding [GO:0005524] | UP000018817 |
| PPTG_16075 | W2PST1 | Phospholipid-transporting ATPase (EC 7.6.2.1) | 1201 | Cation transport ATPase (P-type) (TC 3.A.3) family, | phospholipid translocation [GO:0045332] | endomembrane system [GO:0012505]; plasr | ATP binding [GO:0005524]; ATP hydrolysis ac | endomembrane system [G | UP000018817 |
| PPTG_14405 | W2PUB8 | Superkiller protein 3 | 1325 | | RNA catabolic process [GO:0006401] | Ski complex [GO:0055087] | | Ski complex [GO:0055087] | UP000018817 |
| PPTG_06596 | W2QQV3 | Acylamino-acid-releasing enzyme (EC 3.4.19.1) | 785 | Peptidase S9C family | proteolysis [GO:0006508] | cytoplasm [GO:0005737] | omega peptidase activity [GO:0008242]; ser | cytoplasm [GO:0005737]; | UP000018817 |
| PPTG_03858 | W2QYW8 | Peptidase M3A/M3B catalytic domain-containing protein | 706 | Peptidase M3 family | peptide metabolic process [GO:0006518]; proteolysis [GO:0006508] | | metal ion binding [GO:0046872]; metalloenc | metal ion binding [GO:004 | UP000018817 |
| PPTG_13152 | W2PXT3 | C2 domain-containing protein | 337 | | | | | | UP000018817 |
| PPTG_01646 | W2R7V3 | Peptidyl-prolyl cis-trans isomerase (EC 5.2.1.8) | 113 | | | cytosol [GO:0005829]; host cell cytoplasm [G | peptidyl-prolyl cis-trans isomerase activity [( | cytosol [GO:0005829]; hos | UP000018817 |
| PPTG_05104 | W2QVK8 | Potassium/sodium efflux P-type ATPase, fungal-type | 1201 | | intracellular potassium ion homeostasis [G | plasma membrane [GO:0005886] | ATP binding [GO:0005524]; ATP hydrolysis ac | plasma membrane [GO:00 | UP000018817 |
| PPTG_00758 | W2RGB5 | AB hydrolase-1 domain-containing protein | 340 | | | lysosome [GO:0005764] | thiolester hydrolase activity [GO:0016790] | lysosome [GO:0005764]; t | UP000018817 |
| PPTG_05278 | W2QYZ1 | EF-hand domain-containing protein | 563 | | intracellular calcium ion homeostasis [GO: | endomembrane system [GO:0012505]; vacu | calcium ion binding [GO:0005509]; calcium: | endomembrane system [G | UP000018817 |
| PPTG_00969 | W2RJG9 | methylmalonyl-CoA mutase (EC 5.4.99.2) | 760 | Methylmalonyl-CoA mutase family | propionate metabolic process, methylmalo | mitochondrion [GO:0005739] | cobalamin binding [GO:0031419]; metal ion | mitochondrion [GO:00057 | UP000018817 |
| PPTG_02501 | W2RB13 | GTP cyclohydrolase II | 459 | | | | | | UP000018817 |
| PPTG_13814 | W2PYE8 | Long-chain-fatty-acid--CoA ligase (EC 6.2.1.3) | 666 | ATP-dependent AMP-binding enzyme family | | endoplasmic reticulum [GO:0005783]; meml | ATP binding [GO:0005524]; long-chain fatty a | endoplasmic reticulum [G | UP000018817 |
| PPTG_09261 | W2QJJ8 | Pentacotripeptide-repeat region of PRORP domain-containing protein | 278 | | | | | | UP000018817 |
| PPTG_16135 | W2PQA5 | PCI domain-containing protein | 263 | CSN7/EIF3M family, CSN7 subfamily | | COP9 signalosome [GO:0008180] | | COP9 signalosome [GO:00 | UP000018817 |
| PPTG_15564 | W2PTK9 | glucan endo-1,3-beta-D-glucosidase (EC 3.2.1.39) (Endo-1,3-beta-glucanase btgC) (Laminarinase | 366 | Glycosyl hydrolase 17 family | cell wall organization [GO:0071555]; polysa | plasma membrane [GO:0005886] | glucan endo-1,3-beta-D-glucosidase activity | plasma membrane [GO:00 | UP000018817 |
| PPTG_09146 | W2QGI3 | protein-tyrosine-phosphatase (EC 3.1.3.48) | 422 | Protein-tyrosine phosphatase family, Non-receptor | cell division [GO:0051301]; cellular respons | cytoplasm [GO:0005737]; cytoskeleton [GO: | protein tyrosine phosphatase activity [GO:00 | cytoplasm [GO:0005737]; | UP000018817 |
| PPTG_04333 | W2R0A8 | Uncharacterized protein | 1518 | Non-repetitive/WGA-negative nucleoporin family | protein import into nucleus [GO:0006606]; | nuclear pore inner ring [GO:0044611] | structural constituent of nuclear pore [GO:0 | nuclear pore inner ring [GC | UP000018817 |
| PPTG_14242 | W2PZR0 | Uncharacterized protein | 488 | | | | | | UP000018817 |
| PPTG_13847 | W2PYQ2 | Uncharacterized protein | 812 | | | | | | UP000018817 |
| PPTG_09264 | W2QIQ9 | Choline/carnitine acyltransferase domain-containing protein | 634 | Carnitine/choline acetyltransferase family | | | acyltransferase activity [GO:0016746] | acyltransferase activity [G | UP000018817 |
| PPTG_13910 | W2Q0Q5 | NADH dehydrogenase [ubiquinone] iron-sulfur protein 8, mitochondrial | 211 | Complex I 23 kDa subunit family | mitochondrial electron transport, NADH to u | membrane [GO:0016020]; mitochondrion [G | 4 iron, 4 sulfur cluster binding [GO:0051539] | membrane [GO:0016020]; | UP000018817 |
| PPTG_06372 | W2QV75 | EF-hand domain-containing protein | 537 | DMRL synthase family | riboflavin biosynthetic process [GO:000923 | riboflavin synthase complex [GO:0009349] | calcium ion binding [GO:0005509]; transfera | riboflavin synthase comple | UP000018817 |
| PPTG_06696 | W2QQP1 | Uncharacterized protein | 713 | | | | | | UP000018817 |
| PPTG_16595 | W2PQW1 | Fumarylacetoacetase-like C-terminal domain-containing protein | 305 | FAH family | oxaloacetate metabolic process [GO:0006107] | | acetylpyruvate hydrolase activity [GO:00187 | acetylpyruvate hydrolase a | UP000018817 |
| PPTG_11756 | W2Q847 | serine C-palmitoyltransferase (EC 2.3.1.50) | 533 | Class-II pyridoxal-phosphate-dependent aminotrar | ceramide biosynthetic process [GO:004651 | endoplasmic reticulum [GO:0005783]; meml | pyridoxal phosphate binding [GO:0030170]; | endoplasmic reticulum [G | UP000018817 |
| PPTG_12099 | W2Q537 | non-specific serine/threonine protein kinase (EC 2.7.11.1) | 519 | | signal transduction [GO:0007165] | | ATP binding [GO:0005524]; protein serine/th | ATP binding [GO:0005524] | UP000018817 |
| PPTG_18781 | W2PE19 | ABC transporter domain-containing protein | 613 | ABC transporter superfamily, ABCG family, Eye pigment precursor importer (TC 3.A.1.204) subf | | membrane [GO:0016020] | ABC-type transporter activity [GO:0140359]; | membrane [GO:0016020]; | UP000018817 |
| PPTG_02730 | W2RCM9 | Uncharacterized protein | 445 | | | | | | UP000018817 |
| PPTG_16799 | W2PMQ4 | Thioredoxin domain-containing protein | 496 | | | COPII-coated ER to Golgi transport vesicle [GO:0030134]; endoplasmic reticulum [GO:000 | | COPII-coated ER to Golgi t | UP000018817 |
| PPTG_12536 | W2Q568 | Peptidyl-prolyl cis-trans isomerase (EC 5.2.1.8) | 148 | PpiC/parvulin rotamase family | | cytoplasm [GO:0005737] | peptidyl-prolyl cis-trans isomerase activity [( | cytoplasm [GO:0005737]; | UP000018817 |
| PPTG_01690 | W2RA07 | Phosphoribosylaminoimidazole-succinocarboxamide synthase, chloroplastic (EC 6.3.2.6) (SAICAR | 341 | SAICAR synthetase family | 'de novo' IMP biosynthetic process [GO:000 | cytoplasm [GO:0005737] | ATP binding [GO:0005524]; phosphoribosyla | cytoplasm [GO:0005737]; | UP000018817 |

| | | | | | | | | | |
|---|---|---|---|---|---|---|---|---|---|
| PPTG_06370 | W2QST5 | 6,7-dimethyl-8-ribityllumazine synthase (DMRL synthase) (EC 2.5.1.78) | 220 | DMRL synthase family | riboflavin biosynthetic process [GO:000923 | riboflavin synthase complex [GO:0009349] | 6,7-dimethyl-8-ribityllumazine synthase acti | riboflavin synthase comple | UP000018817 |
| PPTG_19149 | W2PE68 | Dolichol-phosphate mannosyltransferase subunit 1 (EC 2.4.1.83) | 240 | Glycosyltransferase 2 family | alcohol metabolic process [GO:0006066]; d | endoplasmic reticulum membrane [GO:0005 | dolichyl-phosphate beta-D-mannosyltransfe | endoplasmic reticulum me | UP000018817 |
| PPTG_00875 | W2RHD9 | CNH domain-containing protein | 1034 | | autophagy [GO:0006914]; endosomal vesic | cytoplasm [GO:0005737]; membrane [GO:0016020] | | cytoplasm [GO:0005737]; | UP000018817 |
| PPTG_19189 | W2PDJ6 | Importin N-terminal domain-containing protein | 1030 | | protein import into nucleus [GO:0006606] | cytosol [GO:0005829]; nuclear envelope [GO | small GTPase binding [GO:0031267] | cytosol [GO:0005829]; nuc | UP000018817 |
| PPTG_01496 | W2R762 | DUF445 domain-containing protein | 561 | | | | | | UP000018817 |
| PPTG_00442 | W2RH45 | Uncharacterized protein | 224 | | superoxide metabolic process [GO:0006801] | | metal ion binding [GO:0046872] | metal ion binding [GO:004 | UP000018817 |
| PPTG_17984 | W2PJV6 | Pyruvate dehydrogenase E1 component subunit beta (EC 1.2.4.1) | 359 | | pyruvate decarboxylation to acetyl-CoA [GO | mitochondrion [GO:0005739] | metal ion binding [GO:0046872]; pyruvate de | mitochondrion [GO:00057 | UP000018817 |
| PPTG_13362 | W2Q2Q6 | DNA-directed RNA polymerase RpoA/D/Rpb3-type domain-containing protein | 240 | Archaeal Rpo3/eukaryotic RPB3 RNA polymerase s | transcription by RNA polymerase II [GO:000 | RNA polymerase II, core complex [GO:00056 | DNA binding [GO:0003677]; DNA-directed RI | RNA polymerase II, core cc | UP000018817 |
| PPTG_08473 | W2QKP6 | TKL protein kinase | 473 | | | | ATP binding [GO:0005524]; protein serine/th | ATP binding [GO:0005524] | UP000018817 |
| PPTG_06915 | W2QR67 | superoxide dismutase (EC 1.15.1.1) | 155 | Cu-Zn superoxide dismutase family | | | copper ion binding [GO:0005507]; superoxid | copper ion binding [GO:00 | UP000018817 |
| PPTG_07301 | W2QS63 | 2,4-dienoyl-CoA reductase [(3E)-enoyl-CoA-producing] (EC 1.3.1.124) | 297 | | fatty acid catabolic process [GO:0009062] | peroxisome [GO:0005777] | 2,4-dienoyl-CoA reductase (NADPH) activity | peroxisome [GO:0005777] | UP000018817 |
| PPTG_06943 | W2QS05 | C2 DOCK-type domain-containing protein | 1912 | DOCK family | small GTPase-mediated signal transduction [GO:0007264] | | guanyl-nucleotide exchange factor activity [G | guanyl-nucleotide exchang | UP000018817 |
| PPTG_07225 | W2QRW3 | asparagine--tRNA ligase (EC 6.1.1.22) | 495 | Class-II aminoacyl-tRNA synthetase family | asparaginyl-tRNA aminoacylation [GO:0006 | mitochondrion [GO:0005739] | asparagine-tRNA ligase activity [GO:000481 | mitochondrion [GO:00057 | UP000018817 |
| PPTG_19545 | W2PDZ7 | SnoaL-like domain-containing protein | 184 | | | | | | UP000018817 |
| PPTG_10533 | W2QB70 | Eukaryotic translation initiation factor 3 subunit I (eIF3i) | 327 | WD repeat STRAP family; EIF-3 subunit I family | formation of cytoplasmic translation initiati | eukaryotic 43S preinitiation complex [GO:00 | RNA binding [GO:0003723]; translation initia | eukaryotic 43S preinitiatio | UP000018817 |
| PPTG_08766 | W2QIH1 | Mitogen-activated protein kinase (EC 2.7.11.24) | 663 | OSBP family; Protein kinase superfamily, Ser/Thr p | lipid transport [GO:0006869] | | ATP binding [GO:0005524]; MAP kinase activ | ATP binding [GO:0005524] | UP000018817 |
| PPTG_05903 | W2QUC8 | catechol O-methyltransferase (EC 2.1.1.6) | 266 | Class I-like SAM-binding methyltransferase superfa | catecholamine metabolic process [GO:0006584]; methylation [GO:0032259] | | catechol O-methyltransferase activity [GO:0 | catechol O-methyltransfer | UP000018817 |
| PPTG_02084 | W2R9D3 | ER membrane protein complex subunit 10 | 285 | | | | | | UP000018817 |
| PPTG_14972 | W2PTL1 | Uncharacterized protein | 805 | | | | | | UP000018817 |
| PPTG_13317 | W2Q3H5 | FYVE-type domain-containing protein | 664 | | | | zinc ion binding [GO:0008270] | zinc ion binding [GO:0008 | UP000018817 |
| PPTG_01338 | W2R6X3 | GST C-terminal domain-containing protein | 176 | | | aminoacyl-tRNA synthetase multienzyme complex [GO:0017101]; cytoplasm [GO:000573 | | aminoacyl-tRNA synthetas | UP000018817 |
| PPTG_10200 | W2QDM3 | Isochorismatase-like domain-containing protein | 209 | Isochorismatase family | | | | | UP000018817 |
| PPTG_14788 | W2PVX5 | Obg-like ATPase 1 | 390 | TRAFAC class OBG-HflX-like GTPase superfamily, ( | response to stress [GO:0006950] | cytosol [GO:0005829] | ATP binding [GO:0005524]; ATP hydrolysis ac | cytosol [GO:0005829]; ATF | UP000018817 |
| PPTG_18280 | W2PHH8 | Uncharacterized protein | 384 | | endoplasmic reticulum unfolded protein res | endoplasmic reticulum chaperone complex [ | ATP binding [GO:0005524]; ATP-dependent p | endoplasmic reticulum ch | UP000018817 |
| PPTG_12069 | W2Q509 | Carboxypeptidase (EC 3.4.16.-) | 496 | Peptidase S10 family | proteolysis [GO:0006508] | | serine-type carboxypeptidase activity [GO:0 | serine-type carboxypeptid | UP000018817 |
| PPTG_09767 | W2QBX5 | Uracil catabolism protein 4 | 443 | | | | | | UP000018817 |
| PPTG_16014 | W2PQ00 | Ribosomal protein L22 | 190 | Universal ribosomal protein uL22 family | cytoplasmic translation [GO:0002181] | cytosolic large ribosomal subunit [GO:00226 | structural constituent of ribosome [GO:0003 | cytosolic large ribosomal s | UP000018817 |
| PPTG_10319 | W2QGN3 | 60S acidic ribosomal protein P2 | 113 | Eukaryotic ribosomal protein P1/P2 family | cytoplasmic translational elongation [GO:0 | cytosolic large ribosomal subunit [GO:00226 | structural constituent of ribosome [GO:0003 | cytosolic large ribosomal s | UP000018817 |
| PPTG_08251 | W2QJU3 | protein-disulfide reductase (EC 1.8.1.8) | 146 | Nucleoredoxin family | | | protein-disulfide reductase [NAD(P)H] activi | protein-disulfide reductas | UP000018817 |
| PPTG_15152 | W2PU74 | non-specific serine/threonine protein kinase (EC 2.7.11.1) | 348 | Protein kinase superfamily | regulation of cell cycle [GO:0051726] | cytosol [GO:0005829]; nucleus [GO:0005634 | ATP binding [GO:0005524]; protein serine/th | cytosol [GO:0005829]; nuc | UP000018817 |
| PPTG_18704 | W2PFI0 | ATP synthase F1 complex delta/epsilon subunit N-terminal domain-containing protein | 169 | ATPase epsilon chain family | | mitochondrial inner membrane [GO:000574 | proton-transporting ATP synthase activity, ro | mitochondrial inner memb | UP000018817 |
| PPTG_00928 | W2RJ86 | Chorismate synthase (EC 4.2.3.5) | 456 | Chorismate synthase family | amino acid biosynthetic process [GO:0008 | cytosol [GO:0005829] | chorismate synthase activity [GO:0004107]; | cytosol [GO:0005829]; cho | UP000018817 |
| PPTG_06939 | W2QTY6 | short-chain 2-methylacyl-CoA dehydrogenase (EC 1.3.8.5) | 423 | Acyl-CoA dehydrogenase family | fatty acid metabolic process [GO:0006631] | mitochondrion [GO:0005739] | flavin adenine dinucleotide binding [GO:005 | mitochondrion [GO:00057 | UP000018817 |
| PPTG_10094 | W2QD36 | Serine/threonine-protein phosphatase (EC 3.1.3.16) | 308 | PPP phosphatase family, PP-4 (PP-X) subfamily | | | metal ion binding [GO:0046872]; protein seri | metal ion binding [GO:004 | UP000018817 |
| PPTG_00108 | W2RDL1 | Glutathione peroxidase | 288 | Glutathione peroxidase family | response to oxidative stress [GO:0006979] | | peroxidase activity [GO:0004601] | peroxidase activity [GO:00 | UP000018817 |
| PPTG_06221 | W2QUM3 | Uncharacterized protein | 315 | Small GTPase superfamily, Rab family | | | GTP binding [GO:0005525]; GTPase activity [ | GTP binding [GO:0005525] | UP000018817 |
| PPTG_04178 | W2R2A9 | Stealth protein CR2 conserved region 2 domain-containing protein | 515 | Stealth family | | Golgi apparatus [GO:0005794] | transferase activity, transferring phosphorus | Golgi apparatus [GO:0005 | UP000018817 |
| PPTG_11749 | W2QAN3 | Uncharacterized protein | 1775 | VPS8 family | endosomal vesicle fusion [GO:0034058]; pr | HOPS complex [GO:0030897]; late endosome [GO:0005770] | | HOPS complex [GO:00308 | UP000018817 |
| PPTG_01494 | W2PAX2 | Thioredoxin | 501 | | | | | | UP000018817 |
| PPTG_12028 | W2Q5M4 | S-adenosyl-L-homocysteine hydrolase NAD binding domain-containing protein | 352 | D-isomer specific 2-hydroxyacid dehydrogenase family | | | NAD binding [GO:0051287]; oxidoreductase | NAD binding [GO:0051287 | UP000018817 |
| PPTG_09959 | W2QCL9 | Exocyst complex subunit Exo70 C-terminal domain-containing protein | 542 | EXO70 family | exocytosis [GO:0006887] | exocyst [GO:0000145] | phosphatidylinositol-4,5-bisphosphate bindi | exocyst [GO:0000145]; pho | UP000018817 |
| PPTG_08368 | W2QMY2 | Betaine aldehyde dehydrogenase | 491 | Aldehyde dehydrogenase family | | | oxidoreductase activity, acting on the aldehy | oxidoreductase activity, ac | UP000018817 |
| PPTG_01003 | W2RHY4 | Arp2/3 complex 34 kDa subunit | 233 | ARPC2 family | actin filament polymerization [GO:0030041 | Arp2/3 protein complex [GO:0005885] | actin filament binding [GO:0051015]; structu | Arp2/3 protein complex [G | UP000018817 |
| PPTG_18442 | W2PI57 | UV excision repair protein Rad23 | 467 | RAD23 family | nucleotide-excision repair [GO:0006289]; p | cytosol [GO:0005829]; nucleoplasm [GO:000 | damaged DNA binding [GO:0003684]; polyul | cytosol [GO:0005829]; nuc | UP000018817 |
| PPTG_00246 | W2RGL8 | MHD domain-containing protein | 437 | Adaptor complexes medium subunit family | endocytosis [GO:0006897]; intracellular prc | clathrin adaptor complex [GO:0030131]; clathrin-coated pit [GO:0005905]; plasma memb | | clathrin adaptor complex [ | UP000018817 |
| PPTG_17096 | W2PKV5 | non-specific serine/threonine protein kinase (EC 2.7.11.1) | 379 | Protein kinase superfamily | | | ATP binding [GO:0005524]; protein serine kir | ATP binding [GO:0005524] | UP000018817 |
| PPTG_05079 | W2QWC8 | glucan endo-1,3-beta-D-glucosidase (EC 3.2.1.39) (Endo-1,3-beta-glucanase btgC) (Laminarinase | 447 | | cell wall organization [GO:0071555]; polysa | plasma membrane [GO:0005886] | glucan endo-1,3-beta-D-glucosidase activity | plasma membrane [GO:00 | UP000018817 |
| PPTG_14299 | W2PXB8 | NAD(P)-binding domain-containing protein | 223 | | import into nucleus [GO:0051170] | cytoplasm [GO:0005737] | | cytoplasm [GO:0005737]; | UP000018817 |
| PPTG_13136 | W2PXW0 | Uncharacterized protein | 861 | | cyclic nucleotide biosynthetic process [GO: | membrane [GO:0016020] | | membrane [GO:0016020]; | UP000018817 |
| PPTG_02717 | W2RC51 | Hsp90-like protein | 687 | Heat shock protein 90 family | | | ATP binding [GO:0005524]; ATP hydrolysis ac | ATP binding [GO:0005524] | UP000018817 |
| PPTG_19449 | W2PCP1 | Protein transporter Sec61 subunit alpha isoform 2 | 474 | SecY/SEC61-alpha family | protein transport [GO:0015031] | endoplasmic reticulum membrane [GO:0005789] | | endoplasmic reticulum me | UP000018817 |
| PPTG_02437 | W2RAQ5 | protein-serine/threonine phosphatase (EC 3.1.3.16) | 344 | PP2C family | | membrane [GO:0016020] | metal ion binding [GO:0046872]; protein seri | membrane [GO:0016020]; | UP000018817 |
| PPTG_09868 | W2QCA5 | glucan endo-1,3-beta-D-glucosidase (EC 3.2.1.39) | 635 | Glycosyl hydrolase 81 family | cell wall organization [GO:0071555]; polysaccharide catabolic process [GO:0000272] | | endo-1,3(4)-beta-glucanase activity [GO:005 | endo-1,3(4)-beta-glucanas | UP000018817 |
| PPTG_04696 | W2R1Y7 | Xaa-Pro aminopeptidase | 633 | Peptidase M24B family | | cytoplasm [GO:0005737] | metal ion binding [GO:0046872]; metalloam | cytoplasm [GO:0005737]; | UP000018817 |
| PPTG_19282 | W2PDI4 | ABC transporter family G domain-containing protein | 530 | | | | | | UP000018817 |
| PPTG_06658 | W2QSB7 | Uncharacterized protein | 1592 | | | | | | UP000018817 |
| PPTG_06981 | W2QRH2 | Fibronectin type-III domain-containing protein | 369 | | | | | | UP000018817 |
| PPTG_02605 | W2RBN5 | Transcription elongation factor spt6 | 1526 | SPT6 family | nucleosome organization [GO:0034728]; tr | chromosome [GO:0005694]; transcription el | DNA binding [GO:0003677]; histone binding | chromosome [GO:00056 | UP000018817 |
| PPTG_05233 | W2QYB1 | Translation initiation factor SUI1 | 109 | SUI1 family | | | translation initiation factor activity [GO:0003 | translation initiation factor | UP000018817 |
| PPTG_02739 | W2REA4 | Glutathione S-transferase | 200 | | glutathione metabolic process [GO:0006749] | | glutathione transferase activity [GO:000436 | glutathione transferase ac | UP000018817 |
| PPTG_15683 | W2PTV6 | 40S ribosomal protein S21 | 79 | Eukaryotic ribosomal protein eS21 family | translation [GO:0006412] | ribonucleoprotein complex [GO:1990904]; rib | structural constituent of ribosome [GO:0003 | ribonucleoprotein comple | UP000018817 |
| PPTG_05330 | W2QWE8 | FYVE-type domain-containing protein | 1703 | | | | phosphatidylinositol binding [GO:0035091]; | phosphatidylinositol bindi | UP000018817 |
| PPTG_04097 | W2QZH4 | Transmembrane protein | 696 | | | | | | UP000018817 |
| PPTG_13095 | W2Q6C7 | USP domain-containing protein | 2984 | | protein deubiquitination [GO:0016579]; pro | cytosol [GO:0005829]; nucleus [GO:0005634 | cysteine-type deubiquitinase activity [GO:00 | cytosol [GO:0005829]; nuc | UP000018817 |
| PPTG_05122 | W2QYH3 | Secreted protein | 197 | | | | | | UP000018817 |
| PPTG_12093 | W2Q6T6 | t-SNARE coiled-coil homology domain-containing protein | 301 | Syntaxin family | intracellular protein transport [GO:0006886 | Golgi membrane [GO:0000139]; SNARE com | SNAP receptor activity [GO:0005484]; SNARI | Golgi membrane [GO:000 | UP000018817 |
| PPTG_00371 | W2RET1 | NADP-dependent oxidoreductase domain-containing protein | 363 | Shaker potassium channel beta subunit family | | | oxidoreductase activity [GO:0016491] | oxidoreductase activity [G | UP000018817 |
| PPTG_15675 | W2PR38 | LarA-like N-terminal domain-containing protein | 470 | | | | lactate racemase activity [GO:0050043] | lactate racemase activity [ | UP000018817 |
| PPTG_06654 | W2QSY4 | Adenylate kinase active site lid domain-containing protein | 241 | Adenylate kinase family | | | AMP kinase activity [GO:0004017]; ATP bindi | AMP kinase activity [GO:0 | UP000018817 |
| PPTG_15477 | W2PRS9 | Interferon-related developmental regulator N-terminal domain-containing protein | 495 | IFRD family | | | | | UP000018817 |
| PPTG_03066 | W2R3J5 | C2 domain-containing protein | 382 | | positive regulation of gene expression [GO:0010628] | | | positive regulation of gene | UP000018817 |
| PPTG_15304 | W2PT43 | Glycerol-3-phosphate dehydrogenase [NAD(+)] (EC 1.1.1.8) | 269 | NAD-dependent glycerol-3-phosphate dehydrogen | carbohydrate metabolic process [GO:0005 | cytosol [GO:0005829] | glycerol-3-phosphate dehydrogenase (NAD+ | cytosol [GO:0005829]; glyc | UP000018817 |
| PPTG_02025 | W2R988 | Hsp70-Hsp90 organising protein (Stress-inducible protein 1) | 263 | | | cytoplasm [GO:0005737] | Hsp90 protein binding [GO:0051879] | cytoplasm [GO:0005737]; | UP000018817 |
| PPTG_00954 | W2RJF1 | Protein disulfide-isomerase domain | 210 | | response to endoplasmic reticulum stress [ | endoplasmic reticulum lumen [GO:0005788] | isomerase activity [GO:0016853]; protein-di | endoplasmic reticulum lu | UP000018817 |
| PPTG_19262 | W2PF53 | EF-hand domain-containing protein | 552 | | | | | | UP000018817 |
| PPTG_13508 | W2Q4B4 | Uncharacterized protein | 171 | | | | | | UP000018817 |
| PPTG_09255 | W2QGY0 | Vacuolar protein sorting-associated protein 11 homolog | 1008 | VPS11 family | endosome organization [GO:0007032]; intr | endosome [GO:0005768]; HOPS complex [G | protein-macromolecule adaptor activity [GO | endosome [GO:0005768]; | UP000018817 |
| PPTG_17901 | W2PKD3 | Uncharacterized protein | 359 | | | | | | UP000018817 |
| PPTG_00641 | W2RFK3 | EF-hand domain-containing protein | 2421 | WD repeat EMAP family | | | calcium ion binding [GO:0005509]; microtub | calcium ion binding [GO:0 | UP000018817 |
| PPTG_04185 | W2QZV7 | Uncharacterized protein | 1514 | | | | | | UP000018817 |
| PPTG_00550 | W2RF80 | PH domain-containing protein | 1009 | | | | | | UP000018817 |
| PPTG_01937 | W2R901 | V-type proton ATPase subunit E | 185 | V-ATPase E subunit family | | proton-transporting two-sector ATPase comp | proton-transporting ATPase activity, rotation | proton-transporting two-se | UP000018817 |
| PPTG_14107 | W2PYN7 | Uncharacterized protein | 2106 | NUP186/NUP192/NUP205 family | | nuclear pore [GO:0005643] | | nuclear pore [GO:0005643 | UP000018817 |
| PPTG_02660 | W2RBY0 | Thioredoxin domain-containing protein | 106 | | | | | | UP000018817 |
| PPTG_05866 | W2QU39 | Crinkler effector protein N-terminal domain-containing protein | 358 | | | extracellular region [GO:0005576]; host cell [GO:0043657] | | extracellular region [GO:0 | UP000018817 |
| PPTG_01055 | W2RJU9 | Uncharacterized protein | 366 | | | | | | UP000018817 |
| PPTG_08494 | W2QKZ0 | Polymer-forming cytoskeletal protein | 203 | | | | | | UP000018817 |
| PPTG_02405 | W2RD56 | NTF2 domain-containing protein | 500 | | | cytosol [GO:0005829]; ribonucleoprotein con | mRNA binding [GO:0003729] | cytosol [GO:0005829]; ribc | UP000018817 |
| PPTG_07323 | W2QS92 | Anoctamin transmembrane domain-containing protein | 1875 | | | membrane [GO:0016020] | chloride channel activity [GO:0005254] | membrane [GO:0016020]; | UP000018817 |
| PPTG_16825 | W2PPK9 | Myosin motor domain-containing protein | 1303 | TRAFAC class myosin-kinesin ATPase superfamily, | actin filament organization [GO:0007015] | cytoplasm [GO:0005737]; membrane [GO:00 | actin filament binding [GO:0051015]; ATP bi | cytoplasm [GO:0005737]; | UP000018817 |
| PPTG_06806 | W2QRH6 | WLGC domain-containing protein | 809 | | | | | | UP000018817 |
| PPTG_08695 | W2QP55 | Replication protein A C-terminal domain-containing protein | 280 | Replication factor A protein 2 family | DNA replication [GO:0006260]; double-stra | chromosome, telomeric region [GO:0000781 | single-stranded DNA binding [GO:0003697] | chromosome, telomeric re | UP000018817 |
| PPTG_01465 | W2R912 | ADF-H domain-containing protein | 165 | Actin-binding proteins ADF family | actin filament depolymerization [GO:00300 | actin cytoskeleton [GO:0015629] | actin binding [GO:0003779] | actin cytoskeleton [GO:00 | UP000018817 |
| PPTG_06104 | W2QVA7 | Proline iminopeptidase (EC 3.4.11.5) | 382 | Peptidase S33 family | proteolysis [GO:0006508] | cytoplasm [GO:0005737] | aminopeptidase activity [GO:0004177] | cytoplasm [GO:0005737]; | UP000018817 |
| PPTG_08333 | W2QK70 | 40S ribosomal protein S6 | 103 | Eukaryotic ribosomal protein eS6 family | translation [GO:0006412] | ribonucleoprotein complex [GO:1990904]; rib | structural constituent of ribosome [GO:0003 | ribonucleoprotein comple | UP000018817 |
| PPTG_09698 | W2QHN8 | Phospholipase A-2-activating protein | 777 | | proteasome-mediated ubiquitin-dependent | cytoplasm [GO:0005737]; nucleus [GO:0005 | ubiquitin binding [GO:0043130] | cytoplasm [GO:0005737]; | UP000018817 |
| PPTG_07986 | W2QN48 | Uncharacterized protein | 278 | Diacylglycerol acyltransferase family | | | O-acyltransferase activity [GO:0008374] | O-acyltransferase activity | UP000018817 |
| PPTG_17400 | W2PKB9 | MHD domain-containing protein | 425 | Adaptor complexes medium subunit family | intracellular protein transport [GO:0006886 | clathrin adaptor complex [GO:0030131]; endomembrane system [GO:0012505] | | clathrin adaptor complex [ | UP000018817 |
| PPTG_00686 | W2RFW0 | Threonine synthase | 532 | Threonine synthase family | threonine biosynthetic process [GO:0009088] | | threonine synthase activity [GO:0004795] | threonine synthase activity | UP000018817 |
| PPTG_06672 | W2QQL9 | G-protein coupled receptors family 3 profile domain-containing protein | 890 | | | G protein-coupled receptor heterodimeric co | G protein-coupled GABA receptor activity [G | G protein-coupled recepto | UP000018817 |
| PPTG_09306 | W2QJR0 | PH domain-containing protein | 836 | | | | | | UP000018817 |

| | | | | | | | | | |
|---|---|---|---|---|---|---|---|---|---|
| PPTG_02890 | W2RD06 | CAMK/CAMK1 protein kinase | 366 | Protein kinase superfamily | | | ATP binding [GO:0005524]; protein serine/th | ATP binding [GO:0005524] | UP000018817 |
| PPTG_18123 | W2PIH1 | Sulfite oxidase (EC 1.8.3.1) | 587 | | sulfur compound metabolic process [GO:00 | mitochondrial intermembrane space [GO:00 | heme binding [GO:0020037]; molybdenum i | mitochondrial intermembr | UP000018817 |
| PPTG_03585 | W2R7C7 | Ornithine aminotransferase (EC 2.6.1.13) | 432 | Class-III pyridoxal-phosphate-dependent aminotra | L-arginine catabolic process to L-glutamate | cytoplasm [GO:0005737] | identical protein binding [GO:0042802]; orni | cytoplasm [GO:0005737]; | UP000018817 |
| PPTG_15689 | W2PR58 | N-acetyltransferase domain-containing protein | 980 | MDM20/NAA25 family | | NatB complex [GO:0031416] | | NatB complex [GO:003141 | UP000018817 |
| PPTG_11307 | W2Q939 | Ubiquitin-like 1-activating enzyme E1A | 320 | Ubiquitin-activating E1 family | protein sumoylation [GO:0016925] | cytoplasm [GO:0005737]; SUMO activating e | SUMO activating enzyme activity [GO:00199 | cytoplasm [GO:0005737]; | UP000018817 |
| PPTG_11789 | W2Q875 | Uncharacterized protein | 1692 | | | | | | UP000018817 |
| PPTG_04051 | W2R1R6 | 40S ribosomal protein S20 | 147 | Universal ribosomal protein uS10 family | translation [GO:0006412] | small ribosomal subunit [GO:0015935] | RNA binding [GO:0003723]; structural consti | small ribosomal subunit [G | UP000018817 |
| PPTG_06888 | W2QR31 | ATP-grasp domain-containing protein | 1015 | | | | ATP binding [GO:0005524]; ligase activity [G | ATP binding [GO:0005524] | UP000018817 |
| PPTG_13126 | W2PZL7 | Wntless-like transmembrane domain-containing protein | 425 | | | membrane [GO:0016020] | toxic substance binding [GO:0015643] | membrane [GO:0016020]; | UP000018817 |
| PPTG_06837 | W2QR59 | Tyrosinase copper-binding domain-containing protein | 523 | | | | metal ion binding [GO:0046872]; oxidoreduc | metal ion binding [GO:004 | UP000018817 |
| PPTG_13472 | W2Q1H6 | Acid phosphatase | 464 | Histidine acid phosphatase family | | | phosphatase activity [GO:0016791] | phosphatase activity [GO: | UP000018817 |
| PPTG_02241 | W2RA12 | Coatomer subunit delta | 401 | Adaptor complexes medium subunit family, Delta- | endoplasmic reticulum to Golgi vesicle-me | COPI vesicle coat [GO:0030126]; Golgi membrane [GO:0000139] | | COPI vesicle coat [GO:003 | UP000018817 |
| PPTG_05501 | W2QXW8 | Acetolactate synthase (EC 2.2.1.6) | 648 | TPP enzyme family | isoleucine biosynthetic process [GO:00090 | acetolactate synthase complex [GO:000594 | acetolactate synthase activity [GO:0003984 | acetolactate synthase con | UP000018817 |
| PPTG_12273 | W2Q8A4 | Caltractin | 172 | Centrin family | | myosin II complex [GO:0016460] | calcium ion binding [GO:0005509] | myosin II complex [GO:001 | UP000018817 |
| PPTG_16550 | W2PQP5 | glucan endo-1,3-beta-D-glucosidase (EC 3.2.1.39) | 415 | PGA52 family | cell wall organization [GO:0071555] | | glucan endo-1,3-beta-D-glucosidase activity | glucan endo-1,3-beta-D-gl | UP000018817 |
| PPTG_13535 | W2Q1S2 | protein disulfide-isomerase (EC 5.3.4.1) | 446 | Protein disulfide isomerase family | response to endoplasmic reticulum stress [ | endoplasmic reticulum lumen [GO:0005788] | protein disulfide isomerase activity [GO:000 | endoplasmic reticulum lur | UP000018817 |
| PPTG_02871 | W2RCU3 | Sodium/calcium exchanger membrane region domain-containing protein | 554 | Ca(2+):cation antiporter (CaCA) (TC 2.A.19) family | intracellular calcium ion homeostasis [GO: | plasma membrane [GO:0005886] | calcium channel activity [GO:0005262]; calc | plasma membrane [GO:00 | UP000018817 |
| PPTG_14724 | W2PYA7 | 60S acidic ribosomal protein P1 | 117 | Eukaryotic ribosomal protein P1/P2 family | | ribonucleoprotein complex [GO:1990904]; ribosome [GO:0005840] | | ribonucleoprotein comple | UP000018817 |
| PPTG_18087 | W2PJJ0 | IFT81 calponin homology domain-containing protein | 702 | IFT81 family | cilium assembly [GO:0060271]; intraciliary | ciliary basal body [GO:0036064]; intraciliary t | tubulin binding [GO:0015631] | ciliary basal body [GO:003 | UP000018817 |
| PPTG_10402 | W2QEC2 | Methionine-tRNA ligase, beta subunit | 227 | | | | ligase activity [GO:0016874]; tRNA binding [ | ligase activity [GO:001687 | UP000018817 |
| PPTG_10282 | W2QFW8 | RanBD1 domain-containing protein | 204 | | nucleocytoplasmic transport [GO:0006913] | cytoplasm [GO:0005737]; nuclear pore [GO: | GTPase activator activity [GO:0005096] | cytoplasm [GO:0005737]; | UP000018817 |
| PPTG_18912 | W2PF54 | P-type ATPase A domain-containing protein | 1402 | Cation transport ATPase (P-type) (TC 3.A.3) family, Type V subfamily | | membrane [GO:0016020] | ATP binding [GO:0005524]; ATP hydrolysis ac | membrane [GO:0016020]; | UP000018817 |
| PPTG_16818 | W2PPJ9 | 40S ribosomal protein S25 | 111 | Eukaryotic ribosomal protein eS25 family | | ribonucleoprotein complex [GO:1990904]; ribosome [GO:0005840] | | ribonucleoprotein comple | UP000018817 |
| PPTG_10118 | W2QDB2 | Histone H2B | 115 | Histone H2B family | | nucleosome [GO:0000786]; nucleus [GO:000 | DNA binding [GO:0003677]; protein heterodi | nucleosome [GO:0000786 | UP000018817 |
| PPTG_13452 | W2Q2B2 | Histone H2B | 117 | Histone H2B family | | nucleosome [GO:0000786]; nucleus [GO:000 | DNA binding [GO:0003677]; protein heterodi | nucleosome [GO:0000786 | UP000018817 |
| PPTG_10079 | W2QEY3 | Aminotransferase class I/classII large domain-containing protein | 491 | Class-I pyridoxal-phosphate-dependent aminotran | L-alanine catabolic process [GO:0042853] | | L-alanine:2-oxoglutarate aminotransferase a | L-alanine:2-oxoglutarate a | UP000018817 |
| PPTG_18702 | W2PHY0 | FMN hydroxy acid dehydrogenase domain-containing protein | 382 | FMN-dependent alpha-hydroxy acid dehydrogenase family | | cytoplasm [GO:0005737] | FMN binding [GO:0010181]; oxidoreductase | cytoplasm [GO:0005737]; | UP000018817 |
| PPTG_18295 | W2PJC2 | MYG1 protein | 365 | MYG1 family | | cytoplasm [GO:0005737]; nucleus [GO:0005634] | | cytoplasm [GO:0005737]; | UP000018817 |
| PPTG_07315 | W2QS83 | Cyclic nucleotide-binding domain-containing protein | 407 | | | cytosol [GO:0005829] | DNA-binding transcription factor activity [GO | cytosol [GO:0005829]; DN | UP000018817 |
| PPTG_00975 | W2RJH7 | BEACH domain-containing protein | 3318 | | | | | | UP000018817 |
| PPTG_07498 | W2QPM1 | 3-isopropylmalate dehydrogenase (EC 1.1.1.85) | 361 | Isocitrate and isopropylmalate dehydrogenases fa | L-leucine biosynthetic process [GO:000909 | cytosol [GO:0005829] | 3-isopropylmalate dehydrogenase activity [G | cytosol [GO:0005829]; 3-is | UP000018817 |
| PPTG_11731 | W2Q9T0 | GOLD domain-containing protein | 369 | EMP24/GP25L family | | membrane [GO:0016020] | | membrane [GO:0016020] | UP000018817 |
| PPTG_04179 | W2R068 | Exopolysaccharide phosphotransferase | 477 | Stealth family | | Golgi apparatus [GO:0005794] | transferase activity, transferring phosphorus | Golgi apparatus [GO:0005 | UP000018817 |
| PPTG_01569 | W2RA05 | W2 domain-containing protein | 445 | EIF-2-beta/eIF-5 family | formation of cytoplasmic translation initiati | cytosol [GO:0005829] | eukaryotic initiation factor eIF2 binding [GO: | cytosol [GO:0005829]; euk | UP000018817 |
| PPTG_11388 | W2QAC8 | VWFA domain-containing protein | 223 | | protein ubiquitination [GO:0016567] | nucleus [GO:0005634] | ubiquitin-protein transferase activity [GO:00 | nucleus [GO:0005634]; ub | UP000018817 |
| PPTG_18395 | W2PG49 | Expansin-like EG45 domain-containing protein | 411 | | | | | | UP000018817 |
| PPTG_04003 | W2QYU9 | Knr4/Smi1-like domain-containing protein | 174 | | | | | | UP000018817 |
| PPTG_19383 | W2PFA3 | Corrinoid adenosyltransferase MMAB (ATP:co(I)rrinoid adenosyltransferase MMAB) | 224 | Cob(I)alamin adenosyltransferase family | cobalamin metabolic process [GO:0009235] | | ATP binding [GO:0005524]; corrinoid adenos | ATP binding [GO:0005524] | UP000018817 |
| PPTG_18258 | W2PHH6 | Imidazole glycerol phosphate synthase hisHF [Includes: Glutaminase (EC 3.5.1.2); Cyclase] | 548 | HisA/HisF family; HisA/HisF family | L-histidine biosynthetic process [GO:0000105] | | glutaminase activity [GO:0004359]; imidazol | glutaminase activity [GO:0 | UP000018817 |
| PPTG_07422 | W2QQ19 | SSD domain-containing protein | 1492 | Patched family | sterol transport [GO:0015918] | membrane [GO:0016020] | sterol binding [GO:0032934] | membrane [GO:0016020]; | UP000018817 |
| PPTG_16766 | W2PM86 | Uncharacterized protein | 298 | | fatty acid beta-oxidation [GO:0006635] | peroxisome [GO:0005777] | (3S)-3-hydroxyacyl-CoA dehydrogenase (NA | peroxisome [GO:0005777] | UP000018817 |
| PPTG_08738 | W2QHK3 | SCP domain-containing protein | 173 | | | | | | UP000018817 |
| PPTG_01811 | W2R8F4 | Phosphoribosyl-ATP diphosphatase | 477 | | L-histidine biosynthetic process [GO:0000105] | | ATP binding [GO:0005524]; phosphoribosyl-A | ATP binding [GO:0005524] | UP000018817 |
| PPTG_02424 | W2RCU6 | Succinate dehydrogenase [ubiquinone] iron-sulfur subunit, mitochondrial (EC 1.3.5.1) | 271 | Succinate dehydrogenase/fumarate reductase iron | respiratory electron transport chain [GO:00 | mitochondrial inner membrane [GO:000574 | 2 iron, 2 sulfur cluster binding [GO:0051537] | mitochondrial inner memb | UP000018817 |
| PPTG_19378 | W2PDG1 | Inactive glycoside hydrolase XLP1 (Glycoside hydrolase family 12 protein XLP1) (GH12 protein XLP1 | 238 | Glycosyl hydrolase 12 (cellulase H) family | polysaccharide catabolic process [GO:0000 | extracellular region [GO:0005576] | cellulase activity [GO:0008810] | extracellular region [GO:00 | UP000018817 |
| PPTG_02217 | W2RC03 | glutaryl-CoA dehydrogenase (ETF) (EC 1.3.8.6) | 420 | Acyl-CoA dehydrogenase family | fatty acid beta-oxidation using acyl-CoA deh | mitochondrial matrix [GO:0005759] | fatty-acyl-CoA binding [GO:0000062]; flavin a | mitochondrial matrix [GO: | UP000018817 |
| PPTG_04264 | W2R032 | Sidoreflexin | 325 | Sideroflexin family | serine import into mitochondrion [GO:0140 | mitochondrial inner membrane [GO:000574 | monoatomic ion transmembrane transporte | mitochondrial inner memb | UP000018817 |
| PPTG_06564 | W2QVZ8 | Uncharacterized protein | 2376 | | endocytosis [GO:0006897]; intracellular pro | cytosol [GO:0005829]; endocytic vesicle [GO:0030139]; Golgi apparatus [GO:0005794]; m | | cytosol [GO:0005829]; enc | UP000018817 |
| PPTG_07614 | W2QQR6 | 3-oxo-5-alpha-steroid 4-dehydrogenase C-terminal domain-containing protein | 282 | Steroid 5-alpha reductase family | very long-chain fatty acid biosynthetic proce | endoplasmic reticulum membrane [GO:0005 | oxidoreductase activity, acting on the CH-CH | endoplasmic reticulum me | UP000018817 |
| PPTG_14384 | W2Q080 | Kinesin-like protein | 976 | TRAFAC class myosin-kinesin ATPase superfamily, | microtubule-based movement [GO:000701 | microtubule [GO:0005874] | ATP binding [GO:0005524]; microtubule bind | microtubule [GO:0005874 | UP000018817 |
| PPTG_16603 | W2PN30 | CRAL-TRIO domain-containing protein | 630 | | | cytoplasm [GO:0005737] | | cytoplasm [GO:0005737] | UP000018817 |
| PPTG_16493 | W2PMY5 | Serine/threonine-protein phosphatase (EC 3.1.3.16) | 320 | PPP phosphatase family, PP-4 (PP-X) subfamily | | | metal ion binding [GO:0046872]; protein seri | metal ion binding [GO:004 | UP000018817 |
| PPTG_13449 | W2Q1C8 | Serine/threonine-protein phosphatase (EC 3.1.3.16) | 513 | PPP phosphatase family, PP-5 (PP-T) subfamily | | cytoplasm [GO:0005737] | metal ion binding [GO:0046872]; protein seri | cytoplasm [GO:0005737]; | UP000018817 |
| PPTG_07816 | W2QNT0 | Ras-related protein Rab-21 | 277 | Small GTPase superfamily, Rab family | protein transport [GO:0015031]; Rab protei | endomembrane system [GO:0012505] | GTP binding [GO:0005525]; GTPase activity [ | endomembrane system [G | UP000018817 |
| PPTG_11612 | W2Q6M3 | CAMK/CDPK protein kinase | 581 | Protein kinase superfamily, Ser/Thr protein kinase family, CDPK subfamily | | | ATP binding [GO:0005524]; calcium ion bind | ATP binding [GO:0005524] | UP000018817 |
| PPTG_17147 | W2PL44 | DRTGG domain-containing protein | 413 | | | | | | UP000018817 |
| PPTG_15953 | W2PU58 | Uncharacterized protein | 357 | | | | | | UP000018817 |
| PPTG_06099 | W2QVT4 | S-formylglutathione hydrolase (EC 3.1.2.12) | 282 | Esterase D family | formaldehyde catabolic process [GO:00462 | cytosol [GO:0005829] | carboxylic ester hydrolase activity [GO:0052 | cytosol [GO:0005829]; car | UP000018817 |
| PPTG_17439 | W2PMT4 | NAD(P)-binding domain-containing protein | 317 | | | | | | UP000018817 |
| PPTG_05462 | W2QZ43 | DnaJ homologue subfamily C GRV2/DNAJC13 N-terminal domain-containing protein | 1540 | | endosome organization [GO:0007032]; rece | endosome membrane [GO:0010008] | | endosome membrane [GC | UP000018817 |
| PPTG_19332 | W2PDU0 | Choline transporter-like protein | 745 | CTL (choline transporter-like) family | | plasma membrane [GO:0005886] | transmembrane transporter activity [GO:002 | plasma membrane [GO:00 | UP000018817 |
| PPTG_07828 | W2QMJ3 | L-type lectin-like domain-containing protein | 417 | | endoplasmic reticulum to Golgi vesicle-me | COPII-coated ER to Golgi transport vesicle [G | D-mannose binding [GO:0005537] | COPII-coated ER to Golgi t | UP000018817 |
| PPTG_11903 | W2Q8P7 | RuvB-like helicase (EC 3.6.4.12) | 463 | RuvB family | DNA repair [GO:0006281] | nucleus [GO:0005634] | ATP binding [GO:0005524]; ATP hydrolysis ac | nucleus [GO:0005634]; AT | UP000018817 |
| PPTG_13854 | W2Q161 | E2 ubiquitin-conjugating enzyme (EC 2.3.2.23) | 149 | Ubiquitin-conjugating enzyme family | | | ATP binding [GO:0005524]; ubiquitin conjuga | ATP binding [GO:0005524] | UP000018817 |
| PPTG_05322 | W2QWL5 | Eukaryotic translation initiation factor 4E | 196 | Eukaryotic initiation factor 4E family | regulation of translation [GO:0006417] | eukaryotic translation initiation factor 4F com | RNA 7-methylguanosine cap binding [GO:00 | eukaryotic translation initia | UP000018817 |
| PPTG_08558 | W2QLT7 | Uncharacterized protein | 275 | | | | | | UP000018817 |
| PPTG_00147 | W2RG96 | Enoyl reductase (ER) domain-containing protein | 327 | | | | oxidoreductase activity [GO:0016491] | oxidoreductase activity [G | UP000018817 |
| PPTG_11462 | W2QC95 | Branched-chain-amino-acid aminotransferase (EC 2.6.1.42) | 390 | Class-IV pyridoxal-phosphate-dependent aminotra | amino acid biosynthetic process [GO:0008652]; branched-chain amino acid biosynthetic | | L-isoleucine-2-oxoglutarate transaminase ac | L-isoleucine-2-oxoglutarat | UP000018817 |
| PPTG_00848 | W2RGQ3 | Peptidyl-prolyl cis-trans isomerase (PPIase) (EC 5.2.1.8) | 203 | Cyclophilin-type PPIase family | protein folding [GO:0006457] | cytoplasm [GO:0005737] | cyclosporin A binding [GO:0016018]; peptidy | cytoplasm [GO:0005737]; | UP000018817 |
| PPTG_05032 | W2QVK2 | Xanthine dehydrogenase, molybdopterin binding subunit | 1448 | Xanthine dehydrogenase family | | | 2 iron, 2 sulfur cluster binding [GO:0051537] | 2 iron, 2 sulfur cluster bind | UP000018817 |
| PPTG_03089 | W2R3M3 | Condensation domain-containing protein | 567 | | | | | | UP000018817 |
| PPTG_08068 | W2QJQ3 | Importin N-terminal domain-containing protein | 485 | | protein import into nucleus [GO:0006606] | cytosol [GO:0005829]; nuclear envelope [GO | small GTPase binding [GO:0031267] | cytosol [GO:0005829]; nuc | UP000018817 |
| PPTG_12648 | W2Q2M3 | Uncharacterized protein | 326 | | | | | | UP000018817 |
| PPTG_15104 | W2PU36 | BRO1 domain-containing protein | 883 | | protein transport to vacuole involved in ubiq | endosome [GO:0005768] | | endosome [GO:0005768]; | UP000018817 |
| PPTG_00604 | W2RFF4 | DNA-directed RNA polymerase subunit beta (EC 2.7.7.6) | 1202 | RNA polymerase beta chain family | DNA-templated transcription [GO:0006351 | DNA-directed RNA polymerase complex [GO: | DNA binding [GO:0003677]; DNA-directed R | DNA-directed RNA polyme | UP000018817 |
| PPTG_15284 | W2PT24 | Tubby C-terminal domain-containing protein | 391 | TUB family | | | | | UP000018817 |
| PPTG_15508 | W2PQ53 | AP-3 complex subunit beta | 1079 | Adaptor complexes large subunit family | intracellular protein transport [GO:0006886 | AP-3 adaptor complex [GO:0030123]; endomembrane system [GO:0012505] | | AP-3 adaptor complex [GO | UP000018817 |
| PPTG_18060 | W2PK06 | Phosphomannomutase (EC 5.4.2.8) | 247 | Eukaryotic PMM family | GDP-mannose biosynthetic process [GO:00 | cytosol [GO:0005829] | metal ion binding [GO:0046872]; phosphom | cytosol [GO:0005829]; me | UP000018817 |
| PPTG_05912 | W2QWH9 | Tetrahydrofolate dehydrogenase/cyclohydrolase NAD(P)-binding domain-containing protein | 317 | | purine nucleobase biosynthetic process [G | cytosol [GO:0005829] | methenyltetrahydrofolate cyclohydrolase ac | cytosol [GO:0005829]; me | UP000018817 |
| PPTG_06085 | W2QX31 | Pyridoxal phosphate phosphatase | 793 | | | | metal ion binding [GO:0046872]; phosphata | metal ion binding [GO:004 | UP000018817 |
| PPTG_03164 | W2R6C7 | Phospholipid:diacylglycerol acyltransferase | 674 | | lipid metabolic process [GO:0006629] | | O-acyltransferase activity [GO:0008374] | O-acyltransferase activity | UP000018817 |
| PPTG_05414 | W2QX04 | RNA helicase (EC 3.6.4.13) | 543 | DEAD box helicase family | | | ATP binding [GO:0005524]; hydrolase activity | ATP binding [GO:0005524] | UP000018817 |
| PPTG_10426 | W2QGE3 | Uncharacterized protein | 783 | | | | | | UP000018817 |
| PPTG_12299 | W2Q7J5 | Fimbrin | 931 | | actin filament bundle assembly [GO:00510 | actin filament [GO:0005884]; actin filament | actin filament binding [GO:0051015]; calciu | actin filament [GO:000588 | UP000018817 |
| PPTG_08876 | W2QI75 | Succinyl-CoA:3-ketoacid-coenzyme A transferase (EC 2.8.3.5) | 499 | 3-oxoacid CoA-transferase family | ketone body catabolic process [GO:004695 | mitochondrion [GO:0005739] | succinyl-CoA:3-oxo-acid CoA-transferase ac | mitochondrion [GO:00057 | UP000018817 |
| PPTG_13677 | W2Q0E0 | Anaphase-promoting complex subunit 4 WD40 domain-containing protein | 392 | | ribosomal large subunit biogenesis [GO:004 | nucleolus [GO:0005730]; preribosome, large subunit precursor [GO:0030687] | | nucleolus [GO:0005730]; | UP000018817 |
| PPTG_06719 | W2QT46 | Exocyst complex component Sec3 PIP2-binding N-terminal domain-containing protein | 959 | SEC3 family | exocytosis [GO:0006887]; Golgi to plasma r | exocyst [GO:0000145]; plasma membrane [G | phosphatidylinositol-4,5-bisphosphate bindi | exocyst [GO:0000145]; pla | UP000018817 |
| PPTG_17429 | W2PM53 | Uncharacterized protein | 936 | | | | | | UP000018817 |
| PPTG_07053 | W2QPB9 | Mitochondrial pyruvate carrier | 126 | Mitochondrial pyruvate carrier (MPC) (TC 2.A.105) | pyruvate import into mitochondria [GO:000 | mitochondrial inner membrane [GO:0005743] | | mitochondrial inner memb | UP000018817 |
| PPTG_04280 | W2R0M6 | Plant heme peroxidase family profile domain-containing protein | 690 | | cellular response to hydrogen peroxide [GO: | cytosol [GO:0005829] | catalase activity [GO:0004096]; heme bindir | cytosol [GO:0005829]; cat | UP000018817 |
| PPTG_08092 | W2QJU1 | PHD-type domain-containing protein | 541 | | | | zinc ion binding [GO:0008270] | zinc ion binding [GO:0008 | UP000018817 |
| PPTG_09748 | W2QDM7 | tRNA ligase phosphodiesterase domain-containing protein | 1194 | | tRNA splicing, via endonucleolytic cleavage and ligation [GO:0006388] | | ATP binding [GO:0005524]; RNA ligase (ATP) | ATP binding [GO:0005524] | UP000018817 |
| PPTG_08041 | W2QIY3 | acetate--CoA ligase (EC 6.2.1.1) | 281 | ATP-dependent AMP-binding enzyme family | acetyl-CoA biosynthetic process [GO:0006085] | | acetate-CoA ligase activity [GO:0003987] | acetate-CoA ligase activity | UP000018817 |
| PPTG_09057 | W2QH05 | E1-like protein-activating enzyme Gsa7p/Apg7p | 471 | | autophagosome assembly [GO:0000045]; a | phagophore assembly site [GO:0000407] | Atg12 activating enzyme activity [GO:00197 | phagophore assembly site | UP000018817 |
| PPTG_18680 | W2PG33 | EF-hand domain-containing protein | 1997 | | | | calcium ion binding [GO:0005509] | calcium ion binding [GO:0 | UP000018817 |
| PPTG_09944 | W2QF31 | MI domain-containing protein | 1424 | Eukaryotic initiation factor 4G family | | eukaryotic translation initiation factor 4F com | mRNA binding [GO:0003729]; translation init | eukaryotic translation initia | UP000018817 |
| PPTG_13996 | W2Q1S8 | Ribosomal protein L15 | 251 | Universal ribosomal protein uL15 family | translation [GO:0006412] | mitochondrial large ribosomal subunit [GO:0 | structural constituent of ribosome [GO:0003 | mitochondrial large riboso | UP000018817 |
| PPTG_19075 | W2PDH6 | ADF-H domain-containing protein | 308 | Actin-binding proteins ADF family, Twinfilin subfam | actin filament depolymerization [GO:00300 | actin filament [GO:0005884]; cytoplasm [GO | actin filament binding [GO:0051015]; actin n | actin filament [GO:000588 | UP000018817 |
| PPTG_05936 | W2QUI5 | Lon protease homolog, mitochondrial (EC 3.4.21.53) | 915 | Peptidase S16 family | cellular response to oxidative stress [GO:00 | mitochondrial matrix [GO:0005759] | ATP binding [GO:0005524]; ATP hydrolysis ac | mitochondrial matrix [GO: | UP000018817 |
| PPTG_09259 | W2QGW9 | Phosphoglycerate mutase (EC 5.4.2.11) | 292 | Phosphoglycerate mutase family, BPG-dependent | glycolytic process [GO:0006096] | | phosphoglycerate mutase activity [GO:0004 | phosphoglycerate mutase | UP000018817 |
| PPTG_03299 | W2R4T4 | Bacterial Pleckstrin homology domain-containing protein | 241 | | | | | | UP000018817 |

| | | | | | | | | | | |
|---|---|---|---|---|---|---|---|---|---|---|
| PPTG_09277 | W2QGY6 | Uncharacterized protein | 876 | | | | | | | UP000018817 |
| PPTG_08475 | W2QMM1 | J domain-containing protein | 476 | | | clathrin coat disassembly [GO:0072318]; cl | cytoplasm [GO:0005737]; vesicle [GO:0031 | clathrin binding [GO:0030276] | cytoplasm [GO:0005737]; | UP000018817 |
| PPTG_11247 | W2Q9S3 | Eukaryotic translation initiation factor 5B (EC 3.6.5.3) (Translation initiation factor IF-2) | 973 | | TRAFAC class translation factor GTPase superfamily, Classic translation factor GTPase family, | | mitochondrion [GO:0005739] | GTP binding [GO:0005525]; GTPase activity [ | mitochondrion [GO:0005 | UP000018817 |
| PPTG_19151 | W2PFF6 | Translation initiation factor eIF2B subunit epsilon (eIF2B GDP-GTP exchange factor subunit epsilon | 741 | | EIF-2B gamma/epsilon subunits family | | cytosol [GO:0005829]; eukaryotic translation | guanyl-nucleotide exchange factor activity [ | cytosol [GO:0005829]; euk | UP000018817 |
| PPTG_15073 | W2PWK8 | TNFR-Cys domain-containing protein | 378 | | | | | | | UP000018817 |
| PPTG_09418 | W2QHF3 | ABC transporter domain-containing protein | 1363 | | ABC transporter superfamily, ABCG family, PDR (TC 3.A.1.205) subfamily | | membrane [GO:0016020] | ABC-type transporter activity [GO:0140359]; | membrane [GO:0016020]; | UP000018817 |
| PPTG_04251 | W2R213 | 40S ribosomal protein S15 | 184 | | Universal ribosomal protein uS19 family | ribosomal small subunit assembly [GO:000 | cytosolic small ribosomal subunit [GO:00226 | RNA binding [GO:0003723]; structural consti | cytosolic small ribosomal | UP000018817 |
| PPTG_13875 | W2Q0K2 | 50S ribosomal protein L19, chloroplastic | 178 | | Bacterial ribosomal protein bL19 family | translation [GO:0006412] | chloroplast [GO:0009507]; mitochondrial lar | structural constituent of ribosome [GO:0003 | chloroplast [GO:0009507]; | UP000018817 |
| PPTG_08965 | W2QL34 | C3H1-type domain-containing protein | 433 | | | | | | | UP000018817 |
| PPTG_14832 | W2PW99 | L-type lectin-like domain-containing protein | 415 | | | endoplasmic reticulum to Golgi vesicle-me | COPII-coated ER to Golgi transport vesicle [G | D-mannose binding [GO:0005537] | COPII-coated ER to Golgi t | UP000018817 |
| PPTG_05877 | W2QUV8 | beta-glucosidase (EC 3.2.1.21) | 788 | | Glycosyl hydrolase 3 family | glucan catabolic process [GO:0009251] | | beta-glucosidase activity [GO:0008422] | beta-glucosidase activity [ | UP000018817 |
| PPTG_04788 | W2R2L9 | Arginine biosynthesis bifunctional protein ArgJ, mitochondrial [Cleaved into: Arginine biosynthesis | 465 | | ArgJ family | L-arginine biosynthetic process [GO:00065 | mitochondrial matrix [GO:0005759] | L-glutamate N-acetyltransferase activity [GC | mitochondrial matrix [GO: | UP000018817 |
| PPTG_00515 | W2RF93 | Anoctamin transmembrane domain-containing protein | 1044 | | | | membrane [GO:0016020] | chloride channel activity [GO:0005254] | membrane [GO:0016020]; | UP000018817 |
| PPTG_13113 | W2Q3Q5 | DNA polymerase (EC 2.7.7.7) | 1089 | | DNA polymerase type-B family | base-excision repair, gap-filling [GO:00062 | delta DNA polymerase complex [GO:004362 | 3'-5'-DNA exonuclease activity [GO:0008296 | delta DNA polymerase con | UP000018817 |
| PPTG_14629 | W2PW67 | PH domain-containing protein | 394 | | | | | | | UP000018817 |
| PPTG_00742 | W2RIA7 | EGF-like domain-containing protein | 796 | | | | | | | UP000018817 |
| PPTG_08947 | W2QL11 | Uncharacterized protein | 955 | | | | | glycosyltransferase activity [GO:0016757] | glycosyltransferase activity | UP000018817 |
| PPTG_01013 | W2RJM4 | homogentisate 1,2-dioxygenase (EC 1.13.11.5) | 440 | | Homogentisate dioxygenase family | L-phenylalanine catabolic process [GO:000 | cytoplasm [GO:0005737] | homogentisate 1,2-dioxygenase activity [GO | cytoplasm [GO:0005737]; | UP000018817 |
| PPTG_01711 | W2R828 | ATP phosphoribosyltransferase (EC 2.4.2.17) | 299 | | ATP phosphoribosyltransferase family | L-histidine biosynthetic process [GO:00001 | cytoplasm [GO:0005737] | ATP binding [GO:0005524]; ATP phosphoribo | cytoplasm [GO:0005737]; | UP000018817 |
| PPTG_14623 | W2PV73 | Fibronectin type III-like domain-containing protein | 813 | | | arabinan catabolic process [GO:0031222]; xylan catabolic process [GO:0045493] | | alpha-L-arabinofuranosidase activity [GO:00 | alpha-L-arabinofuranosida | UP000018817 |
| PPTG_03736 | W2R6G5 | Endoplasmic reticulum vesicle transporter N-terminal domain-containing protein | 292 | | ERGIC family | | COPII-coated ER to Golgi transport vesicle [GO:0030134]; endoplasmic reticulum [GO:00 | | COPII-coated ER to Golgi t | UP000018817 |
| PPTG_03562 | W2R7A1 | Probable pectate lyase F (EC 4.2.2.2) | 261 | | Polysaccharide lyase 3 family | pectin catabolic process [GO:0045490] | extracellular region [GO:0005576] | pectate lyase activity [GO:0030570] | extracellular region [GO:0 | UP000018817 |
| PPTG_14270 | W2PY30 | Tyrosinase copper-binding domain-containing protein | 546 | | | | | metal ion binding [GO:0046872]; oxidoreduc | metal ion binding [GO:004 | UP000018817 |
| PPTG_07051 | W2QQL4 | Cytochrome c | 112 | | Cytochrome c family | | mitochondrial intermembrane space [GO:00 | electron transfer activity [GO:0009055]; hem | mitochondrial intermembr | UP000018817 |
| PPTG_14386 | W2PZK7 | Fibronectin type III-like domain-containing protein | 772 | | Glycosyl hydrolase 3 family | arabinan catabolic process [GO:0031222]; xylan catabolic process [GO:0045493] | | alpha-L-arabinofuranosidase activity [GO:00 | alpha-L-arabinofuranosida | UP000018817 |
| PPTG_12471 | W2Q503 | GH16 domain-containing protein | 754 | | | (1->6)-beta-D-glucan biosynthetic process [ | endoplasmic reticulum membrane [GO:0005 | glucosidase activity [GO:0015926] | endoplasmic reticulum me | UP000018817 |
| PPTG_17152 | W2PM98 | Ribonucleoprotein-associated protein | 128 | | Eukaryotic ribosomal protein eL8 family | mRNA splicing, via spliceosome [GO:00003 | nucleolus [GO:0005730]; spliceosomal comp | RNA binding [GO:0003723] | nucleolus [GO:0005730]; s | UP000018817 |
| PPTG_00308 | W2RGX0 | Peptidase M20 dimerisation domain-containing protein | 464 | | | | | hydrolase activity [GO:0016787]; metal ion b | hydrolase activity [GO:001 | UP000018817 |
| PPTG_14391 | W2PUA5 | beta-glucosidase (EC 3.2.1.21) | 774 | | Glycosyl hydrolase 3 family | glucan catabolic process [GO:0009251] | | beta-glucosidase activity [GO:0008422] | beta-glucosidase activity [ | UP000018817 |
| PPTG_14802 | W2PW48 | Acetolactate synthase, small subunit, variant | 241 | | Acetolactate synthase small subunit family | isoleucine biosynthetic process [GO:00090 | cytosol [GO:0005829] | acetolactate synthase activity [GO:0003984 | cytosol [GO:0005829]; ace | UP000018817 |
| PPTG_04184 | W2QZM3 | Uncharacterized protein | 1083 | | | | | | | UP000018817 |
| PPTG_09348 | W2QJZ7 | Phospholipid/glycerol acyltransferase domain-containing protein | 423 | | | phosphatidic acid biosynthetic process [GO:0006654] | | 1-acylglycerol-3-phosphate O-acyltransferas | 1-acylglycerol-3-phosphat | UP000018817 |
| PPTG_10374 | W2QEB1 | Complex III subunit VII | 103 | | UQCRB/QCR7 family | mitochondrial electron transport, ubiquinol | mitochondrial inner membrane [GO:0005743]; respiratory chain complex III [GO:004527 | | mitochondrial inner memb | UP000018817 |
| PPTG_19740 | W2PDC9 | AB hydrolase-1 domain-containing protein | 361 | | AB hydrolase superfamily, Lipase family | lipid catabolic process [GO:0016042] | | hydrolase activity, acting on ester bonds [GO | hydrolase activity, acting o | UP000018817 |
| PPTG_17649 | W2PII4 | V-SNARE coiled-coil homology domain-containing protein | 1289 | | WD repeat L(2)GL family | exocytosis [GO:0006887]; Golgi to plasma r | cytoplasm [GO:0005737]; plasma membrane | GTPase activator activity [GO:0005096]; myc | cytoplasm [GO:0005737]; | UP000018817 |
| PPTG_11061 | W2Q8W7 | Serine/threonine-protein phosphatase (EC 3.1.3.16) | 312 | | PPP phosphatase family | | | metal ion binding [GO:0046872]; protein seri | metal ion binding [GO:004 | UP000018817 |
| PPTG_07092 | W2QNT7 | Importin subunit alpha | 559 | | Importin alpha family | protein import into nucleus [GO:0006606] | cytoplasm [GO:0005737] | nuclear import signal receptor activity [GO:0 | cytoplasm [GO:0005737]; | UP000018817 |
| PPTG_16032 | W2PSK4 | 1,3-beta-glucan synthase (EC 2.4.1.34) | 2647 | Zhao et al., 2019 | SKN1/KRE6 family; Glycosyltransferase 48 family | (1->3)-beta-D-glucan biosynthetic process [ | 1,3-beta-D-glucan synthase complex [GO:00 | 1,3-beta-D-glucan synthase activity [GO:000 | 1,3-beta-D-glucan synthas | UP000018817 |
| PPTG_04970 | W2R332 | Histone H2A | 140 | | Histone H2A family | | nucleosome [GO:0000786]; nucleus [GO:000 | DNA binding [GO:0003677]; protein heterodi | nucleosome [GO:0000786 | UP000018817 |
| PPTG_15078 | W2PUW4 | Uncharacterized protein | 328 | | Short-chain dehydrogenases/reductases (SDR) family | | endoplasmic reticulum [GO:0005783] | oxidoreductase activity [GO:0016491] | endoplasmic reticulum [G | UP000018817 |
| PPTG_04665 | W2R1M1 | Glutaredoxin | 104 | | | cellular response to oxidative stress [GO:00 | cytoplasm [GO:0005737] | glutathione disulfide oxidoreductase activity | cytoplasm [GO:0005737]; | UP000018817 |
| PPTG_05471 | W2QWY7 | SEC7 domain-containing protein | 1540 | | | regulation of ARF protein signal transductio | ESCRT I complex [GO:0000813] | guanyl-nucleotide exchange factor activity [ | ESCRT I complex [GO:000 | UP000018817 |
| PPTG_16150 | W2PRC3 | Glutaredoxin domain-containing protein | 127 | | | | | | | UP000018817 |
| PPTG_09516 | W2QG51 | GTP-binding nuclear protein | 214 | | Small GTPase superfamily, Ran family | protein import into nucleus [GO:0006606]; | cytoplasm [GO:0005737]; nucleus [GO:0005 | GTP binding [GO:0005525]; GTPase activity [ | cytoplasm [GO:0005737]; | UP000018817 |
| PPTG_00774 | W2RIH1 | Actin-2 | 403 | | Actin family, ARP1 subfamily | | cytoskeleton [GO:0005856] | ATP binding [GO:0005524]; hydrolase activity | cytoskeleton [GO:000585 | UP000018817 |
| PPTG_00206 | W2RDZ5 | Uncharacterized protein | 181 | | | | | | | UP000018817 |
| PPTG_18654 | W2PFD0 | Protein disulfide-isomerase domain | 422 | | Protein disulfide isomerase family | response to endoplasmic reticulum stress [ | endoplasmic reticulum lumen [GO:0005788] | protein disulfide isomerase activity [GO:000 | endoplasmic reticulum lu | UP000018817 |
| PPTG_02889 | W2RDG6 | glucan endo-1,3-beta-D-glucosidase (EC 3.2.1.39) (Endo-1,3-beta-glucanase btgC) (Laminarinase | 586 | | | cell wall organization [GO:0071555]; polysa | plasma membrane [GO:0005886] | glucan endo-1,3-beta-D-glucosidase activity | plasma membrane [GO:0 | UP000018817 |
| PPTG_11245 | W2QAP6 | Uncharacterized protein | 125 | | | | | | | UP000018817 |
| PPTG_01321 | W2R6D6 | VOC domain-containing protein | 353 | | | | | | | UP000018817 |
| PPTG_19559 | W2PE44 | FYVE-type domain-containing protein | 774 | | | | | zinc ion binding [GO:0008270] | zinc ion binding [GO:0008 | UP000018817 |
| PPTG_00569 | W2RHT1 | Uncharacterized protein | 488 | | | | | | | UP000018817 |
| PPTG_03020 | W2R5W0 | NAD(P)H:quinone oxidoreductase, type IV | 200 | | WrbA family | | membrane [GO:0016020] | FMN binding [GO:0010181]; NAD(P)H dehyd | membrane [GO:0016020]; | UP000018817 |
| PPTG_00045 | W2RDI2 | DUSP domain-containing protein | 519 | | | | | cysteine-type deubiquitinase activity [GO:00 | cysteine-type deubiquitina | UP000018817 |
| PPTG_02518 | W2RB34 | Magnesium transporter | 462 | | CorA metal ion transporter (MIT) (TC 1.A.35) family | | mitochondrial inner membrane [GO:000574 | magnesium ion transmembrane transporter | mitochondrial inner memb | UP000018817 |
| PPTG_03859 | W2QY84 | 6-phosphogluconolactonase (6PGL) (EC 3.1.1.31) | 245 | | Glucosamine/galactosamine-6-phosphate isomer | carbohydrate metabolic process [GO:0005 | cytoplasm [GO:0005737] | 6-phosphogluconolactonase activity [GO:00 | cytoplasm [GO:0005737]; | UP000018817 |
| PPTG_06866 | W2QRP3 | Catalase (EC 1.11.1.6) | 515 | | Catalase family | hydrogen peroxide catabolic process [GO:0 | mitochondrion [GO:0005739]; peroxisome [G | catalase activity [GO:0004096]; heme bindir | mitochondrion [GO:0005 | UP000018817 |
| PPTG_01694 | W2R807 | Ribosomal protein L19 | 185 | | Eukaryotic ribosomal protein eL19 family | translation [GO:0006412] | cytosolic large ribosomal subunit [GO:0022 | RNA binding [GO:0003723]; structural consti | cytosolic large ribosomal s | UP000018817 |
| PPTG_01865 | W2R955 | Tyrosinase copper-binding domain-containing protein | 1493 | | | | | metal ion binding [GO:0046872]; oxidoreduc | metal ion binding [GO:004 | UP000018817 |
| PPTG_13874 | W2PYP1 | Signal recognition particle subunit SRP72 | 622 | | SRP72 family | SRP-dependent cotranslational protein targ | signal recognition particle, endoplasmic retic | 7S RNA binding [GO:0008312]; ribosome bin | signal recognition particle, | UP000018817 |
| PPTG_02812 | W2REN6 | EcxA zinc-binding domain-containing protein | 907 | | | | | metallopeptidase activity [GO:0008237] | metallopeptidase activity [ | UP000018817 |
| PPTG_04177 | W2QZU3 | STE/STE20/FRAY protein kinase | 377 | | Protein kinase superfamily, STE Ser/Thr protein kinase family, STE20 subfamily | | | ATP binding [GO:0005524]; protein kinase ac | ATP binding [GO:0005524] | UP000018817 |
| PPTG_08508 | W2QMU7 | Uncharacterized protein | 673 | | Glycosyl hydrolase 30 family | glucosylceramide catabolic process [GO:00 | membrane [GO:0016020] | glucosylceramidase activity [GO:0004348] | membrane [GO:0016020]; | UP000018817 |
| PPTG_02167 | W2R9P4 | Nucleolar GTP-binding protein 1 | 664 | | TRAFAC class OBG-HflX-like GTPase superfamily, ( | ribosome biogenesis [GO:0042254] | nucleolus [GO:0005730] | GTP binding [GO:0005525] | nucleolus [GO:0005730]; ( | UP000018817 |
| PPTG_07734 | W2QLN4 | TKL protein kinase | 625 | | | | | ATP binding [GO:0005524]; protein serine/th | ATP binding [GO:0005524] | UP000018817 |
| PPTG_15854 | W2PPV6 | peptidylprolyl isomerase (EC 5.2.1.8) | 326 | | | | | peptidyl-prolyl cis-trans isomerase activity [ | peptidyl-prolyl cis-trans isc | UP000018817 |
| PPTG_13548 | W2Q3N0 | Glutaredoxin | 458 | | | cellular response to oxidative stress [GO:00 | cytoplasm [GO:0005737] | glutathione disulfide oxidoreductase activity | cytoplasm [GO:0005737]; | UP000018817 |
| PPTG_01312 | W2R6E3 | Multidrug resistance-associated protein 1 | 1338 | | ABC transporter superfamily, ABCC family, Conjugate transporter (TC 3.A.1.208) subfamily | | vacuolar membrane [GO:0005774] | ABC-type transporter activity [GO:0140359]; | vacuolar membrane [GO:0 | UP000018817 |
| PPTG_01305 | W2R8C8 | Nuclear segregation protein Bfr1 | 502 | | | intracellular mRNA localization [GO:00082 | endoplasmic reticulum [GO:0005783]; nucle | mRNA binding [GO:0003729] | endoplasmic reticulum [G | UP000018817 |
| PPTG_01148 | W2RIE8 | SCP domain-containing protein | 295 | | | | | | | UP000018817 |
| PPTG_16931 | W2PML9 | RING-type E3 ubiquitin transferase (EC 2.3.2.27) | 719 | | | proteasome-mediated ubiquitin-dependent | endomembrane system [GO:0012505] | ubiquitin protein ligase activity [GO:0061630 | endomembrane system [G | UP000018817 |
| PPTG_15232 | W2PV59 | Aldose 1-epimerase | 330 | | | galactose catabolic process via UDP-galactose, Leloir pathway [GO:0033499]; glucose m | | aldose 1-epimerase activity [GO:0004034]; | aldose 1-epimerase activit | UP000018817 |
| PPTG_11267 | W2Q8Z4 | glucan endo-1,3-beta-D-glucosidase (EC 3.2.1.39) (Endo-1,3-beta-glucanase btgC) (Laminarinase | 292 | | Glycosyl hydrolase 17 family | cell wall organization [GO:0071555]; polysa | plasma membrane [GO:0005886] | glucan endo-1,3-beta-D-glucosidase activity | plasma membrane [GO:0 | UP000018817 |
| PPTG_18987 | W2PDK2 | Uncharacterized protein | 1296 | | | microtubule cytoskeleton organization [GO:0000226]; post-chaperonin tubulin folding pa | | beta-tubulin binding [GO:0048487]; GTPase | beta-tubulin binding [GO:0 | UP000018817 |
| PPTG_00440 | W2RF17 | Uncharacterized protein | 244 | | | superoxide metabolic process [GO:0006801] | | metal ion binding [GO:0046872] | metal ion binding [GO:004 | UP000018817 |
| PPTG_03835 | W2QYT4 | Multidrug resistance protein | 1285 | | ABC transporter superfamily, ABCB family, Multidr | oligopeptide export from mitochondrion [G | mitochondrial inner membrane [GO:000574 | ABC-type oligopeptide transporter activity [G | mitochondrial inner memb | UP000018817 |
| PPTG_15925 | W2PSU1 | Choline dehydrogenase (EC 1.1.99.1) | 587 | | GMC oxidoreductase family | glycine betaine biosynthetic process from choline [GO:0019285] | | choline dehydrogenase activity [GO:000881 | choline dehydrogenase ac | UP000018817 |
| PPTG_18935 | W2PG47 | Glucokinase | 398 | | | glycolytic process [GO:0006096] | | ATP binding [GO:0005524]; D-glucose bindin | ATP binding [GO:0005524] | UP000018817 |
| PPTG_07606 | W2QQQ7 | Transmembrane protein | 320 | | | | | | | UP000018817 |
| PPTG_04973 | W2R3G5 | Uncharacterized protein | 750 | | Dynein intermediate chain family | cilium movement [GO:0003341]; outer dyn | microtubule [GO:0005874]; outer dynein arm | dynein heavy chain binding [GO:0045504]; d | microtubule [GO:0005874 | UP000018817 |
| PPTG_09002 | W2QL82 | ER membrane protein complex subunit 2 | 297 | | EMC2 family | | EMC complex [GO:0072546] | | EMC complex [GO:007254 | UP000018817 |
| PPTG_12231 | W2Q7B9 | Diaminopimelate decarboxylase | 422 | | | lysine biosynthetic process via diaminopimelate [GO:0009089] | | diaminopimelate decarboxylase activity [GO | diaminopimelate decarbo | UP000018817 |
| PPTG_05598 | W2QZR0 | Phosphatidylinositol-3,4,5-trisphosphate 3-phosphatase | 355 | | | lipid metabolic process [GO:0006629] | cytosol [GO:0005829] | phosphatidylinositol-3,4,5-trisphosphate 3- | cytosol [GO:0005829]; ph | UP000018817 |
| PPTG_04169 | W2R1N5 | phosphatidylinositol 3-kinase (EC 2.7.1.137) | 1373 | | PI3/PI4-kinase family | bleb assembly [GO:0032060]; cell migratio | cytoplasm [GO:0005737]; phosphatidylinosit | 1-phosphatidylinositol-3-kinase activity [GO: | cytoplasm [GO:0005737]; | UP000018817 |
| PPTG_18422 | W2PG71 | Uncharacterized protein | 702 | | | | cytosol [GO:0005829]; nucleus [GO:0005634 | ATP binding [GO:0005524]; ATP-dependent | cytosol [GO:0005829]; nuc | UP000018817 |
| PPTG_11457 | W2QBG5 | Uncharacterized protein | 410 | | | | | | | UP000018817 |
| PPTG_01546 | W2R7W4 | Uncharacterized protein | 452 | | | | membrane [GO:0016020] | | membrane [GO:0016020] | UP000018817 |
| PPTG_15717 | W2PTE0 | NADH dehydrogenase [ubiquinone] 1 beta subcomplex subunit 9 (Complex I-B22) (NADH-ubiquino | 162 | | Complex I LYR family | mitochondrial electron transport, NADH to | mitochondrial inner membrane [GO:0005743] | | mitochondrial inner memb | UP000018817 |
| PPTG_04091 | W2QZG3 | folate gamma-glutamyl hydrolase (EC 3.4.19.9) | 317 | | Peptidase C26 family | tetrahydrofolylpolyglutamate metabolic pro | extracellular region [GO:0005576]; vacuole [ | gamma-glutamyl-peptidase activity [GO:003 | extracellular region [GO:0 | UP000018817 |
| PPTG_14741 | W2PVP0 | Uncharacterized protein | 1398 | | SAPS family | | | protein phosphatase binding [GO:0019903] | protein phosphatase bindi | UP000018817 |
| PPTG_18531 | W2PH27 | Condensation domain-containing protein | 501 | | | | | | | UP000018817 |
| PPTG_20156 | W2P9C1 | Uncharacterized protein | 341 | | STXBP/unc-18/SEC1 family | vesicle-mediated transport [GO:0016192] | | | vesicle-mediated transpor | UP000018817 |
| PPTG_02905 | W2RFK8 | t-SNARE coiled-coil homology domain-containing protein | 316 | | Syntaxin family | exocytosis [GO:0006887]; intracellular prot | endomembrane system [GO:0012505]; plasn | SNAP receptor activity [GO:0005484]; SNARI | endomembrane system [G | UP000018817 |
| PPTG_11081 | W2Q9S0 | Strictosidine synthase conserved region domain-containing protein | 416 | | Strictosidine synthase family | | endomembrane system [GO:0012505] | hydrolase activity [GO:0016787] | endomembrane system [G | UP000018817 |
| PPTG_02580 | W2RBD5 | Beta-adaptin appendage C-terminal subdomain domain-containing protein | 444 | | | intracellular protein transport [GO:0006886 | clathrin adaptor complex [GO:0030131] | | clathrin adaptor complex [ | UP000018817 |
| PPTG_00630 | W2RHR4 | 60S ribosomal protein L31 | 120 | | Eukaryotic ribosomal protein eL31 family | cytoplasmic translation [GO:0002181] | cytosolic large ribosomal subunit [GO:0022 | structural constituent of ribosome [GO:0003 | cytosolic large ribosomal s | UP000018817 |
| PPTG_13102 | W2Q3P5 | Transportin-1 (Importin beta-2) (Karyopherin beta-2) | 903 | | | protein import into nucleus [GO:0006606] | cytoplasm [GO:0005737]; nucleus [GO:0005 | small GTPase binding [GO:0031267] | cytoplasm [GO:0005737]; | UP000018817 |
| PPTG_06127 | W2PHM4 | NADH:ubiquinone oxidoreductase-like 20kDa subunit domain-containing protein | 205 | | Complex I 20 kDa subunit family | aerobic respiration [GO:0009060]; electron | mitochondrion [GO:0005739]; respiratory ch | 4 iron, 4 sulfur cluster binding [GO:0051539] | mitochondrion [GO:0005 | UP000018817 |
| PPTG_13276 | W2PZ49 | ABC transporter domain-containing protein | 1933 | | ABC transporter superfamily, ABCA family | | membrane [GO:0016020] | ABC-type transporter activity [GO:0140359]; | membrane [GO:0016020]; | UP000018817 |
| PPTG_02754 | W2RC91 | Vesicle transport v-SNARE N-terminal domain-containing protein | 219 | | VTI1 family | intracellular protein transport [GO:0006886 | endoplasmic reticulum membrane [GO:0005 | SNAP receptor activity [GO:0005484]; SNARI | endoplasmic reticulum me | UP000018817 |
| PPTG_15143 | W2PV42 | Histidine acid phosphatase | 389 | | Histidine acid phosphatase family | | | phosphatase activity [GO:0016791] | phosphatase activity [GO: | UP000018817 |

| | | | | | | | | | |
|---|---|---|---|---|---|---|---|---|---|
| PPTG_17118 | W2PNT2 | 26S protease regulatory subunit 6B | 405 | AAA ATPase family | proteolysis [GO:0006508] | cytoplasm [GO:0005737]; nucleus [GO:0005 | ATP binding [GO:0005524]; ATP hydrolysis ac | cytoplasm [GO:0005737]; | UP000018817 |
| PPTG_13520 | W2Q3J3 | 2-amino-3-ketobutyrate coenzyme A ligase, mitochondrial (EC 2.3.1.29) (Aminoacetone synthase) | 413 | Class-II pyridoxal-phosphate-dependent aminotrar | L-threonine catabolic process [GO:0006567 | mitochondrion [GO:0005739] | glycine C-acetyltransferase activity [GO:000 | mitochondrion [GO:0005 | UP000018817 |
| PPTG_19393 | W2PCT2 | Dihydrolipoamide acetyltransferase component of pyruvate dehydrogenase complex (EC 2.3.1.-) | 544 | 2-oxoacid dehydrogenase family | tricarboxylic acid cycle [GO:0006099] | mitochondrion [GO:0005739] | dihydrolipoyllysine-residue succinyltransfera | mitochondrion [GO:0005 | UP000018817 |
| PPTG_09216 | W2QHN7 | Glycosyl hydrolase family 30 TIM-barrel domain-containing protein | 542 | Glycosyl hydrolase 30 family | glucosylceramide catabolic process [GO:00 | membrane [GO:0016020] | glucosylceramidase activity [GO:0004348] | membrane [GO:0016020]; | UP000018817 |
| PPTG_09260 | W2QIQ6 | Uncharacterized protein | 243 | | | | | | UP000018817 |
| PPTG_00519 | W2RF98 | Peptidase M13 C-terminal domain-containing protein | 778 | Peptidase M13 family | protein processing [GO:0016485] | plasma membrane [GO:0005886] | metal ion binding [GO:0046872]; metalloenc | plasma membrane [GO:00 | UP000018817 |
| PPTG_05916 | W2QUN6 | F-actin-capping protein subunit alpha | 288 | F-actin-capping protein alpha subunit family | actin cytoskeleton organization [GO:003003 | cortical cytoskeleton [GO:0030863]; F-actin c | actin filament binding [GO:0051015] | cortical cytoskeleton [GO: | UP000018817 |
| PPTG_02506 | W2RB17 | Magnesium transporter | 456 | CorA metal ion transporter (MIT) (TC 1.A.35) family | | mitochondrial inner membrane [GO:000574 | magnesium ion transmembrane transporter | mitochondrial inner memb | UP000018817 |
| PPTG_19427 | W2PC03 | FACT complex subunit SSRP1 | 533 | SSRP1 family | DNA repair [GO:0006281]; DNA replication | | FACT complex [GO:0035101] | DNA binding [GO:0003677]; histone binding | | FACT complex [GO:00351 | UP000018817 |
| PPTG_12022 | W2Q4U4 | S-adenosyl-L-homocysteine hydrolase NAD binding domain-containing protein | 371 | D-isomer specific 2-hydroxyacid dehydrogenase family | | | NAD binding [GO:0051287]; oxidoreductase | NAD binding [GO:0051287 | UP000018817 |
| PPTG_16172 | W2PPJ2 | Superoxide dismutase copper/zinc binding domain-containing protein | 273 | | superoxide metabolic process [GO:0006801] | | metal ion binding [GO:0046872] | metal ion binding [GO:004 | UP000018817 |
| PPTG_19428 | W2PEF1 | GPI-anchored leucine-rich lipoprotein | 453 | | | | hydrolase activity [GO:0016787] | hydrolase activity [GO:001 | UP000018817 |
| PPTG_17926 | W2PKA3 | 3-deoxy-7-phosphoheptulonate synthase (EC 2.5.1.54) (3-deoxy-D-arabino-heptulosonate 7-phosp | 441 | Class-I DAHP synthase family | amino acid biosynthetic process [GO:00086 | cytoplasm [GO:0005737] | 3-deoxy-7-phosphoheptulonate synthase ac | cytoplasm [GO:0005737]; | UP000018817 |
| PPTG_00934 | W2RJC5 | PPM-type phosphatase domain-containing protein | 297 | | | | protein serine/threonine phosphatase activit | protein serine/threonine p | UP000018817 |
| PPTG_03992 | W2R1J0 | cytochrome-b5 reductase (EC 1.6.2.2) | 316 | Flavoprotein pyridine nucleotide cytochrome reductase family | | membrane [GO:0016020]; mitochondrion [G | cytochrome-b5 reductase activity, acting on | membrane [GO:0016020]; | UP000018817 |
| PPTG_05790 | W2QVV8 | TatD family hydrolase | 332 | Metallo-dependent hydrolases superfamily, TatD-type hydrolase family | | cytosol [GO:0005829] | metal ion binding [GO:0046872]; single-strar | cytosol [GO:0005829]; me | UP000018817 |
| PPTG_16268 | W2PQZ2 | PH domain-containing protein | 1019 | | cellulose biosynthetic process [GO:003024 | endomembrane system [GO:0012505]; mem | cellulose synthase (UDP-forming) activity [G | endomembrane system [G | UP000018817 |
| PPTG_04098 | W2R1D1 | Transmembrane protein | 683 | | | | | | UP000018817 |
| PPTG_07911 | W2QMW9 | AAA+ ATPase domain-containing protein | 1163 | AAA ATPase family | | | ATP binding [GO:0005524]; ATP hydrolysis ac | ATP binding [GO:0005524] | UP000018817 |
| PPTG_01133 | W2RID1 | Phospholipid-transporting ATPase (EC 7.6.2.1) | 1114 | Cation transport ATPase (P-type) (TC 3.A.3) family, | phospholipid translocation [GO:0045332] | plasma membrane [GO:0005886] | ATP binding [GO:0005524]; ATP hydrolysis ac | plasma membrane [GO:00 | UP000018817 |
| PPTG_13977 | W2PZZ3 | Intraflagellar transport protein 52 | 501 | | cilium assembly [GO:0060271]; intraciliary | centriole [GO:0005814]; cilium [GO:0005929]; intraciliary transport particle B [GO:00309 | | centriole [GO:0005814]; ci | UP000018817 |
| PPTG_04242 | W2R2K5 | Vacuolar protein sorting-associated protein 13 second N-terminal domain-containing protein | 1025 | | | | | | UP000018817 |
| PPTG_07651 | W2QN78 | Uncharacterized protein | 502 | | | | | | UP000018817 |
| PPTG_13825 | W2PZ98 | ALA-interacting subunit | 462 | CDC50/LEM3 family | | endoplasmic reticulum [GO:0005783]; Golgi apparatus [GO:0005794]; plasma membrane | | endoplasmic reticulum [G | UP000018817 |
| PPTG_11458 | W2QC92 | Tudor domain-containing protein | 1002 | | | | | | UP000018817 |
| PPTG_11171 | W2Q7G5 | Uncharacterized protein | 365 | | | | | | UP000018817 |
| PPTG_02406 | W2RB67 | t-SNARE coiled-coil homology domain-containing protein | 223 | Syntaxin family | intracellular protein transport [GO:0006886 | endomembrane system [GO:0012505]; SNAF | SNAP receptor activity [GO:0005484]; SNARI | endomembrane system [G | UP000018817 |
| PPTG_04139 | W2QZC9 | PA14 domain-containing protein | 1783 | | actin cytoskeleton organization [GO:003003 | membrane [GO:0016020] | actin filament binding [GO:0051015] | membrane [GO:0016020]; | UP000018817 |
| PPTG_08625 | W2QM06 | Cystathionine beta-synthase, variant | 374 | | | | | | UP000018817 |
| PPTG_11447 | W2Q9M6 | Serine aminopeptidase S33 domain-containing protein | 423 | | | | | | UP000018817 |
| PPTG_19415 | W2PCR8 | Expansin-like EG45 domain-containing protein | 552 | | | | | | UP000018817 |
| PPTG_17308 | W2PJ08 | FZ domain-containing protein | 598 | | | membrane [GO:0016020] | magnesium ion transmembrane transporter | membrane [GO:0016020]; | UP000018817 |
| PPTG_01819 | W2R8G4 | Proteasome subunit beta | 247 | | proteolysis involved in protein catabolic pro | cytoplasm [GO:0005737]; nucleus [GO:0005634]; proteasome core complex [GO:000583 | | cytoplasm [GO:0005737]; | UP000018817 |
| PPTG_07200 | W2QPC5 | Mitochondrial Carrier (MC) Family | 337 | Mitochondrial carrier (TC 2.A.29) family | | mitochondrial membrane [GO:0031966] | succinate:fumarate antiporter activity [GO:0 | mitochondrial membrane | UP000018817 |
| PPTG_19709 | W2PAU5 | Beta-galactosidase | 589 | Glycosyl hydrolase 1 family | carbohydrate metabolic process [GO:0005975] | | beta-glucosidase activity [GO:0008422] | beta-glucosidase activity [ | UP000018817 |
| PPTG_17411 | W2PL20 | Uncharacterized protein | 389 | | | | | | UP000018817 |
| PPTG_08066 | W2QKR8 | SnoaL-like domain-containing protein | 153 | | | | | | UP000018817 |
| PPTG_13613 | W2Q4Y4 | Uncharacterized protein | 1561 | RICTOR family | TORC2 signaling [GO:0038203] | TORC2 complex [GO:0031932] | | TORC2 complex [GO:0031 | UP000018817 |
| PPTG_10401 | W2QEE6 | Oxidoreductase | 317 | | | | oxidoreductase activity [GO:0016491] | oxidoreductase activity [G | UP000018817 |
| PPTG_12908 | W2Q144 | Uncharacterized protein | 543 | Peptidase S28 family | proteolysis [GO:0006508] | | dipeptidyl-peptidase activity [GO:0008239]; | dipeptidyl-peptidase activi | UP000018817 |
| PPTG_01130 | W2RK20 | Serine/threonine-protein phosphatase (EC 3.1.3.16) | 935 | PPP phosphatase family | | | calcium ion binding [GO:0005509]; protein s | calcium ion binding [GO:0 | UP000018817 |
| PPTG_15118 | W2PWR3 | RRM domain-containing protein | 355 | | | | RNA binding [GO:0003723] | RNA binding [GO:0003723 | UP000018817 |
| PPTG_02655 | W2RCB6 | Translocation protein SEC62 | 334 | SEC62 family | post-translational protein targeting to mem | endoplasmic reticulum membrane [GO:0005789] | | endoplasmic reticulum me | UP000018817 |
| PPTG_13672 | W2PZP4 | cystathionine gamma-lyase (EC 4.4.1.1) (Gamma-cystathionase) | 408 | Trans-sulfuration enzymes family | cysteine biosynthetic process via cystathior | cytoplasm [GO:0005737] | cystathionine gamma-lyase activity [GO:000 | cytoplasm [GO:0005737]; | UP000018817 |
| PPTG_05094 | W2QYA9 | 6-phosphogluconolactonase | 385 | Cycloisomerase 2 family | | | 6-phosphogluconolactonase activity [GO:00 | 6-phosphogluconolacton | UP000018817 |
| PPTG_19880 | W2PCN8 | Phosphatase PP2A regulatory subunit A/Splicing factor 3B subunit 1-like HEAT repeat domain-cont | 606 | Phosphatase 2A regulatory subunit A family | | cytosol [GO:0005829]; nucleus [GO:0005634 | protein phosphatase regulator activity [GO:0 | cytosol [GO:0005829]; nuc | UP000018817 |
| PPTG_18716 | W2PHH5 | Thioredoxin domain-containing protein | 755 | | | | | | UP000018817 |
| PPTG_07062 | W2QRD8 | Protein transporter SEC13 | 246 | WD repeat SEC13 family | COPII-coated vesicle budding [GO:0090114 | COPII vesicle coat [GO:0030127]; nuclear po | structural molecule activity [GO:0005198] | COPII vesicle coat [GO:00 | UP000018817 |
| PPTG_11270 | W2Q9V3 | cGMP-dependent protein kinase | 1851 | | | cAMP-dependent protein kinase complex [G | ATP binding [GO:0005524]; cAMP-dependen | cAMP-dependent protein k | UP000018817 |
| PPTG_20054 | W2PCP7 | Aminotransferase class I/classII large domain-containing protein | 367 | | | | pyridoxal phosphate binding [GO:0030170] | pyridoxal phosphate bindi | UP000018817 |
| PPTG_13262 | W2Q071 | Enoyl-CoA hydratase | 269 | | | | | | UP000018817 |
| PPTG_15656 | W2PR81 | Pentacotripeptide-repeat region of PRORP domain-containing protein | 269 | GET4 family | protein insertion into ER membrane [GO:00 | cytosol [GO:0005829] | | cytosol [GO:0005829]; pro | UP000018817 |
| PPTG_17510 | W2PJM8 | Uncharacterized protein | 633 | | | cytoplasm [GO:0005737] | RNA binding [GO:0003723] | cytoplasm [GO:0005737]; | UP000018817 |
| PPTG_17122 | W2PN64 | RxLR effector protein | 204 | | | | | | UP000018817 |
| PPTG_10740 | W2Q9V0 | RRM domain-containing protein | 262 | | | | RNA binding [GO:0003723] | RNA binding [GO:0003723 | UP000018817 |
| PPTG_08097 | W2QKW4 | Uncharacterized protein | 743 | | | | | | UP000018817 |
| PPTG_09481 | W2QH22 | Nudix hydrolase domain-containing protein | 316 | | | | coenzyme A diphosphatase activity [GO:001 | coenzyme A diphosphatas | UP000018817 |
| PPTG_08464 | W2QKN2 | Vesicle transport v-SNARE N-terminal domain-containing protein | 233 | VTI1 family | intracellular protein transport [GO:0006886 | endoplasmic reticulum membrane [GO:0005 | SNAP receptor activity [GO:0005484]; SNARI | endoplasmic reticulum me | UP000018817 |
| PPTG_01600 | W2R7P8 | CAP-Gly domain-containing protein | 1336 | | | dynein complex [GO:0030286]; microtubule [GO:0005874]; spindle [GO:0005819] | | dynein complex [GO:0030 | UP000018817 |
| PPTG_11855 | W2QB19 | Uncharacterized protein | 351 | | | | GTP binding [GO:0005525]; GTPase activity [ | GTP binding [GO:0005525] | UP000018817 |
| PPTG_10539 | W2QB49 | Translation initiation factor eIF2B subunit delta (eIF2B GDP-GTP exchange factor subunit delta) | 538 | EIF-2B alpha/beta/delta subunits family | | cytosol [GO:0005829] | translation initiation factor activity [GO:0003 | cytosol [GO:0005829]; tra | UP000018817 |
| PPTG_06152 | W2QRQ8 | Insulin-degrading enzyme | 785 | Peptidase M16 family | peptide catabolic process [GO:0043171]; p | cytosol [GO:0005829]; mitochondrion [GO:0 | metal ion binding [GO:0046872]; metalloenc | cytosol [GO:0005829]; mit | UP000018817 |
| PPTG_05144 | W2QVT2 | NADP-dependent oxidoreductase domain-containing protein | 277 | Aldo/keto reductase family | | | oxidoreductase activity [GO:0016491] | oxidoreductase activity [G | UP000018817 |
| PPTG_09688 | W2QFS0 | 40S ribosomal protein S27 | 82 | Eukaryotic ribosomal protein eS27 family | translation [GO:0006412] | ribonucleoprotein complex [GO:1990904]; rit | structural constituent of ribosome [GO:0003 | ribonucleoprotein comple | UP000018817 |
| PPTG_09435 | W2QHH2 | Sec1 family protein | 653 | STXBP/unc-18/SEC1 family | vesicle-mediated transport [GO:0016192] | | | vesicle-mediated transpor | UP000018817 |
| PPTG_10199 | W2QEH7 | Thaumatin-like protein | 297 | | | | | | UP000018817 |
| PPTG_02795 | W2RCJ1 | Myosin-8 | 1861 | TRAFAC class myosin-kinesin ATPase superfamily, | actin filament organization [GO:0007015] | cytoplasm [GO:0005737]; membrane [GO:00 | actin filament binding [GO:0051015]; ATP bi | cytoplasm [GO:0005737]; | UP000018817 |
| PPTG_18767 | W2PE04 | Small ribosomal subunit protein mS35 mitochondrial conserved domain-containing protein | 222 | | mitochondrial translation [GO:0032543] | mitochondrial small ribosomal subunit [GO:0 | structural constituent of ribosome [GO:0003 | mitochondrial small ribos | UP000018817 |
| PPTG_18525 | W2PH16 | Uncharacterized protein | 193 | | | | | | UP000018817 |
| PPTG_17174 | W2PK51 | CSC1/OSCA1-like 7TM region domain-containing protein | 1454 | | | plasma membrane [GO:0005886] | calcium-activated cation channel activity [G | plasma membrane [GO:00 | UP000018817 |
| PPTG_01288 | W2R8A8 | Nicotinate phosphoribosyltransferase (EC 6.3.4.21) | 533 | NAPRTase family | NAD+ biosynthetic process via the salvage | cytosol [GO:0005829] | glycosyltransferase activity [GO:0016757]; n | cytosol [GO:0005829]; gly | UP000018817 |
| PPTG_17496 | W2PLZ6 | glucan endo-1,3-beta-D-glucosidase (EC 3.2.1.39) | 567 | PGA52 family | cell wall organization [GO:0071555] | | glucan endo-1,3-beta-D-glucosidase activity | glucan endo-1,3-beta-D-gl | UP000018817 |
| PPTG_03239 | W2R625 | Uncharacterized protein | 377 | Peptidase C1 family | proteolysis [GO:0006508] | | cysteine-type peptidase activity [GO:000823 | cysteine-type peptidase a | UP000018817 |
| PPTG_07347 | W2QRP0 | Acetyl-CoA acetyltransferase | 421 | Thiolase-like superfamily, Thiolase family | fatty acid beta-oxidation [GO:0006635] | mitochondrion [GO:0005739] | acetyl-CoA C-acetyltransferase activity [GO: | mitochondrion [GO:0005 | UP000018817 |
| PPTG_16802 | W2PQ76 | Serine/threonine-protein phosphatase (EC 3.1.3.16) | 772 | PPP phosphatase family | | | calcium ion binding [GO:0005509]; protein s | calcium ion binding [GO:0 | UP000018817 |
| PPTG_14661 | W2PY14 | B box-type domain-containing protein | 597 | | | | zinc ion binding [GO:0008270] | zinc ion binding [GO:0008 | UP000018817 |
| PPTG_08182 | W2QJK9 | Formiminoglutamate deiminase | 863 | Peptidase M20A family | | | hydrolase activity, acting on carbon-nitrogen | hydrolase activity, acting o | UP000018817 |
| PPTG_09875 | W2QCB2 | Very long-chain fatty acid transport protein (Very-long-chain acyl-CoA synthetase) | 656 | ATP-dependent AMP-binding enzyme family | long-chain fatty acid import into cell [GO:00 | peroxisomal membrane [GO:0005778]; plasr | ATP binding [GO:0005524]; long-chain fatty a | peroxisomal membrane [G | UP000018817 |
| PPTG_08785 | W2QJG0 | PH domain-containing protein | 944 | | lipid transport [GO:0006869] | cytosol [GO:0005829]; endoplasmic reticulur | sterol binding [GO:0032934] | cytosol [GO:0005829]; enc | UP000018817 |
| PPTG_15547 | W2PS32 | FYVE-type domain-containing protein | 408 | | | | zinc ion binding [GO:0008270] | zinc ion binding [GO:0008 | UP000018817 |
| PPTG_13569 | W2Q238 | Pyruvate, phosphate dikinase (EC 2.7.9.1) | 900 | PEP-utilizing enzyme family | | | ATP binding [GO:0005524]; kinase activity [G | ATP binding [GO:0005524] | UP000018817 |
| PPTG_01292 | W2R6C2 | Protein phosphatase 1 regulatory subunit 7 | 353 | SDS22 family | | nucleus [GO:0005634] | | nucleus [GO:0005634] | UP000018817 |
| PPTG_06643 | W2QQJ6 | Ferrochelatase, mitochondrial (EC 4.98.1.1) (Heme synthase) (Protoheme ferro-lyase) | 369 | Ferrochelatase family | heme biosynthetic process [GO:0006783] | mitochondrial inner membrane [GO:000574 | ferrochelatase activity [GO:0004325] | mitochondrial inner memb | UP000018817 |
| PPTG_18636 | W2PFM9 | Beta-adaptin appendage C-terminal subdomain domain-containing protein | 771 | | intracellular protein transport [GO:0006886 | clathrin adaptor complex [GO:0030131] | | clathrin adaptor complex [ | UP000018817 |
| PPTG_18599 | W2PFI8 | 60S ribosomal protein L27a | 148 | Universal ribosomal protein uL15 family | translation [GO:0006412] | cytosolic large ribosomal subunit [GO:0022 | structural constituent of ribosome [GO:0003 | cytosolic large ribosomal s | UP000018817 |
| PPTG_01118 | W2RHT9 | Protein transport protein sec16 | 1621 | SEC16 family | Golgi organization [GO:0007030]; protein lo | endoplasmic reticulum exit site [GO:0070971]; ER to Golgi transport vesicle membrane [G | | endoplasmic reticulum exi | UP000018817 |
| PPTG_18019 | W2PHJ4 | non-specific serine/threonine protein kinase (EC 2.7.11.1) | 874 | Protein kinase superfamily, AGC Ser/Thr protein kinase family | | | ATP binding [GO:0005524]; protein serine kir | ATP binding [GO:0005524] | UP000018817 |
| PPTG_09030 | W2QHU2 | ACT domain-containing protein | 893 | Homoserine dehydrogenase family; Aspartokinase | homoserine biosynthetic process [GO:0009090]; threonine biosynthetic process [GO:000 | | aspartate kinase activity [GO:0004072]; ATP | aspartate kinase activity [ | UP000018817 |
| PPTG_18756 | W2PF66 | NADH dehydrogenase [ubiquinone] iron-sulfur protein 7, mitochondrial | 218 | Complex I 20 kDa subunit family | aerobic respiration [GO:0009060]; electron | mitochondrion [GO:0005739]; respiratory ch | 4 iron, 4 sulfur cluster binding [GO:0051539] | mitochondrion [GO:0005 | UP000018817 |
| PPTG_01488 | W2R752 | Signal recognition particle 54 kDa protein | 511 | GTP-binding SRP family, SRP54 subfamily | SRP-dependent cotranslational protein targ | cytosol [GO:0005829]; endoplasmic reticulur | 7S RNA binding [GO:0008312]; endoplasmic | cytosol [GO:0005829]; enc | UP000018817 |
| PPTG_16777 | W2PMM2 | Calponin-homology (CH) domain-containing protein | 443 | | actin filament organization [GO:0007015] | actin cytoskeleton [GO:0015629] | actin filament binding [GO:0051015] | actin cytoskeleton [GO:00 | UP000018817 |
| PPTG_10101 | W2QD65 | RNA helicase (EC 3.6.4.13) | 344 | DEAD box helicase family, DECD subfamily | | nucleus [GO:0005634] | ATP binding [GO:0005524]; hydrolase activity | nucleus [GO:0005634]; AT | UP000018817 |
| PPTG_11582 | W2Q6C5 | Uncharacterized protein | 3554 | Dynein heavy chain family | microtubule-based movement [GO:00070 | axoneme [GO:0005930]; dynein complex [GC | ATP binding [GO:0005524]; dynein intermedi | axoneme [GO:0005930]; d | UP000018817 |
| PPTG_18389 | W2PGJ7 | Uncharacterized protein | 136 | | | | | | UP000018817 |
| PPTG_15462 | W2PRR4 | Uncharacterized protein | 1585 | | | nuclear pore [GO:0005643] | structural constituent of nuclear pore [GO:00 | nuclear pore [GO:0005643 | UP000018817 |
| PPTG_00783 | W2RIJ5 | Peptidase M1 leukotriene A4 hydrolase/aminopeptidase C-terminal domain-containing protein | 806 | | | | aminopeptidase activity [GO:0004177]; met | aminopeptidase activity [G | UP000018817 |
| PPTG_10959 | W2QAY1 | Uncharacterized protein | 158 | | | mitochondrial large ribosomal subunit [GO:0005762] | | mitochondrial large riboso | UP000018817 |
| PPTG_02654 | W2RED4 | Dienelactone hydrolase domain-containing protein | 362 | | | | hydrolase activity [GO:0016787] | hydrolase activity [GO:001 | UP000018817 |
| PPTG_10769 | W2QCJ3 | SBDS family rRNA metabolism protein | 349 | SDO1/SBDS family | cytosolic ribosome assembly [GO:0042256] | cytoplasm [GO:0005737]; nucleus [GO:0005634] | | cytoplasm [GO:0005737]; | UP000018817 |
| PPTG_00702 | W2RID3 | GMP synthase [glutamine-hydrolyzing] (EC 6.3.5.2) (GMP synthetase) (Glutamine amidotransferas | 528 | | | cytosol [GO:0005829] | ATP binding [GO:0005524]; GMP synthase ac | cytosol [GO:0005829]; ATF | UP000018817 |
| PPTG_06256 | W2QUS0 | Chitin-binding type-4 domain-containing protein | 318 | | | | | | UP000018817 |

| | | | | | | | | | | |
|---|---|---|---|---|---|---|---|---|---|---|
| PPTG_15785 | W2PRJ1 | Uncharacterized protein | 989 | | | | | | | UP000018817 |
| PPTG_10116 | W2QFU4 | Histone H4 | 103 | | Histone H4 family | | nucleosome [GO:0000786]; nucleus [GO:000 | DNA binding [GO:0003677]; protein heterodi | nucleosome [GO:0000786 | UP000018817 |
| PPTG_11020 | W2Q6Y5 | Histone deacetylase interacting domain-containing protein | 1598 | | | negative regulation of transcription by RNA | chromatin [GO:0000785]; histone deacetylas | transcription corepressor activity [GO:00037 | chromatin [GO:0000785]; | UP000018817 |
| PPTG_10296 | W2QE22 | Bardet-Biedl syndrome 7 protein | 748 | | | cilium assembly [GO:0060271]; intracellula | axoneme [GO:0005930]; BBSome [GO:0034464]; ciliary basal body [GO:0036064]; memb | | axoneme [GO:0005930]; B | UP000018817 |
| PPTG_15051 | W2PTX6 | EF-hand domain-containing protein | 401 | | | stabilization of membrane potential [GO:00 | cytoplasm [GO:0005737]; plasma membrane | calcium ion binding [GO:0005509]; outward | cytoplasm [GO:0005737]; | UP000018817 |
| PPTG_11045 | W2Q743 | short-chain 2-methylacyl-CoA dehydrogenase (EC 1.3.8.5) | 411 | | Acyl-CoA dehydrogenase family | fatty acid metabolic process [GO:0006631] | mitochondrion [GO:0005739] | flavin adenine dinucleotide binding [GO:005 | mitochondrion [GO:0005 | UP000018817 |
| PPTG_00602 | W2RG05 | Tetraspanin | 372 | Vinay and Belleannée, 2022 | | auxin-activated signaling pathway [GO:000 | membrane [GO:0016020] | | membrane [GO:0016020]; | UP000018817 |
| PPTG_02143 | W2RA31 | Uncharacterized protein | 1444 | | | | membrane [GO:0016020] | ATP binding [GO:0005524]; ATPase-coupled | membrane [GO:0016020]; | UP000018817 |
| PPTG_01894 | W2R8V6 | Cytochrome b-c1 complex subunit 8 (Complex III subunit 8) | 124 | | UQCRQ/QCR8 family | mitochondrial electron transport, ubiquinol | mitochondrial inner membrane [GO:0005743]; respiratory chain complex III [GO:0045275 | | mitochondrial inner memb | UP000018817 |
| PPTG_00771 | W2RGV1 | serine C-palmitoyltransferase (EC 2.3.1.50) | 483 | | Class-II pyridoxal-phosphate-dependent aminotran | ceramide biosynthetic process [GO:004651 | membrane [GO:0016020]; serine palmitoyltra | pyridoxal phosphate binding [GO:0030170]; | membrane [GO:0016020]; | UP000018817 |
| PPTG_11225 | W2Q7S8 | WASH complex subunit 4 | 1114 | | | endosomal transport [GO:0016197]; endos | endosome [GO:0005768]; WASH complex [GO:0071203] | | endosome [GO:0005768]; | UP000018817 |
| PPTG_14825 | W2PYR9 | Anaphase-promoting complex subunit 4 WD40 domain-containing protein | 335 | | WD repeat SEC13 family | COPII-coated vesicle budding [GO:0090114 | COPII vesicle coat [GO:0030127]; nuclear po | structural molecule activity [GO:0005198] | COPII vesicle coat [GO:00 | UP000018817 |
| PPTG_01392 | W2R759 | EF-hand domain-containing protein | 325 | | | | | calcium ion binding [GO:0005509] | calcium ion binding [GO:0 | UP000018817 |
| PPTG_03305 | W2R4J2 | Uncharacterized protein | 1594 | | 5'-3' exonuclease family | nuclear-transcribed mRNA catabolic proces | nucleus [GO:0005634] | 5'-3' RNA exonuclease activity [GO:0004534 | nucleus [GO:0005634]; 5'- | UP000018817 |
| PPTG_04732 | W2R236 | Cytosol aminopeptidase domain-containing protein | 549 | | Peptidase M17 family | proteolysis [GO:0006508] | cytoplasm [GO:0005737] | manganese ion binding [GO:0030145]; meta | cytoplasm [GO:0005737]; | UP000018817 |
| PPTG_04396 | W2R120 | Uncharacterized protein | 206 | | | | | | | UP000018817 |
| PPTG_18146 | W2PGY0 | ABC transporter B family member 2 | 1070 | | ABC transporter superfamily, ABCB family, Multidr | oligopeptide export from mitochondrion [G | mitochondrial inner membrane [GO:000574 | ABC-type oligopeptide transporter activity [G | mitochondrial inner memb | UP000018817 |
| PPTG_00987 | W2RJM3 | SCY1 protein kinase | 928 | | | | | ATP binding [GO:0005524]; protein kinase ac | ATP binding [GO:0005524] | UP000018817 |
| PPTG_15231 | W2PUM5 | Uncharacterized protein | 258 | | Necrosis inducing protein (NPP1) family | | extracellular region [GO:0005576] | | extracellular region [GO:0 | UP000018817 |
| PPTG_19697 | W2PCA1 | Fumarylacetoacetase (EC 3.7.1.2) (Fumarylacetoacetate hydrolase) | 425 | | FAH family | homogentisate catabolic process [GO:1902000]; L-phenylalanine catabolic process [GO: | | fumarylacetoacetase activity [GO:0004334]; | fumarylacetoacetase activ | UP000018817 |
| PPTG_13738 | W2PZM7 | DNA-directed RNA polymerase subunit (EC 2.7.7.6) | 1534 | | RNA polymerase beta' chain family | DNA-templated transcription [GO:0006351] | DNA-directed RNA polymerase complex [GO: | DNA binding [GO:0003677]; DNA-directed R | DNA-directed RNA polyme | UP000018817 |
| PPTG_00662 | W2RHW6 | Clustered mitochondria protein homolog | 1304 | | CLU family | mitochondrion organization [GO:0007005] | cytoplasm [GO:0005737] | RNA binding [GO:0003723] | cytoplasm [GO:0005737]; | UP000018817 |
| PPTG_13532 | W2Q1X1 | EF-hand domain-containing protein | 508 | | | cilium assembly [GO:0060271] | motile cilium [GO:0031514] | | motile cilium [GO:003151 | UP000018817 |
| PPTG_01026 | W2RHG0 | Intraflagellar transport protein 56 | 560 | | IFT56 family | intraciliary anterograde transport [GO:0035 | ciliary basal body [GO:0036064]; ciliary base | intraciliary transport particle B binding [GO: | ciliary basal body [GO:003 | UP000018817 |
| PPTG_06992 | W2QTI8 | PIPK domain-containing protein | 650 | | | phosphatidylinositol phosphate biosyntheti | plasma membrane [GO:0005886] | 1-phosphatidylinositol-4-phosphate 5-kinase | plasma membrane [GO:00 | UP000018817 |
| PPTG_05312 | W2QYJ6 | CSC1/OSCA1-like 7TM region domain-containing protein | 1059 | | CSC1 (TC 1.A.17) family | | plasma membrane [GO:0005886] | calcium-activated cation channel activity [G | plasma membrane [GO:00 | UP000018817 |
| PPTG_02401 | W2RAR8 | TKL protein kinase | 400 | | Protein kinase superfamily | | | ATP binding [GO:0005524]; protein serine/th | ATP binding [GO:0005524] | UP000018817 |
| PPTG_13428 | W2Q4V7 | PDZ domain-containing protein | 3747 | | | | | | | UP000018817 |
| PPTG_03813 | W2QYA1 | HECT-type E3 ubiquitin transferase (EC 2.3.2.26) | 683 | | | protein ubiquitination [GO:0016567]; ubiqui | cytoplasm [GO:0005737] | ubiquitin protein ligase activity [GO:0061630 | cytoplasm [GO:0005737]; | UP000018817 |
| PPTG_17805 | W2PI88 | 10 kDa chaperonin | 99 | | GroES chaperonin family | | mitochondrion [GO:0005739] | ATP binding [GO:0005524]; metal ion binding | mitochondrion [GO:0005 | UP000018817 |
| PPTG_01387 | W2R753 | Large ribosomal subunit protein eL24-related N-terminal domain-containing protein | 151 | | Eukaryotic ribosomal protein eL24 family | cytoplasmic translation [GO:0002181] | cytosolic large ribosomal subunit [GO:0022 | mRNA binding [GO:0003729]; structural con | cytosolic large ribosomal s | UP000018817 |
| PPTG_19290 | W2PD05 | 3-methyl-2-oxobutanoate hydroxymethyltransferase (EC 2.1.2.11) | 370 | | PanB family | methylation [GO:0032259]; pantothenate b | mitochondrion [GO:0005739] | 3-methyl-2-oxobutanoate hydroxymethyltran | mitochondrion [GO:0005 | UP000018817 |
| PPTG_18068 | W2PH82 | 26S proteasome complex subunit DSS1 | 168 | | DSS1/SEM1 family | double-strand break repair via homologous | proteasome regulatory particle, lid subcomplex [GO:0008541] | | proteasome regulatory par | UP000018817 |
| PPTG_01276 | W2R6Q0 | Uncharacterized protein | 762 | | SAPS family | | | protein phosphatase binding [GO:0019903]; | protein phosphatase bindi | UP000018817 |
| PPTG_05217 | W2QWV9 | 3-dehydrosphinganine reductase (EC 1.1.1.102) | 337 | | | 3-keto-sphinganine metabolic process [GO: | endoplasmic reticulum membrane [GO:0005 | 3-dehydrosphinganine reductase activity [G | endoplasmic reticulum me | UP000018817 |
| PPTG_04952 | W2R5D7 | phytol kinase (EC 2.7.1.182) | 176 | | Polyprenol kinase family | | chloroplast [GO:0009507]; membrane [GO:0 | phytol kinase activity [GO:0010276]; zinc ion | chloroplast [GO:0009507]; | UP000018817 |
| PPTG_18841 | W2PFG0 | Laminin IV type A domain-containing protein | 363 | | | | | | | UP000018817 |
| PPTG_16625 | W2PN57 | PH domain-containing protein | 589 | | | | | | | UP000018817 |
| PPTG_02638 | W2RBU0 | PDZ domain-containing protein | 1267 | | | | cytosol [GO:0005829]; nucleus [GO:0005634 | protein phosphatase regulator activity [GO:0 | cytosol [GO:0005829]; nuc | UP000018817 |
| PPTG_09093 | W2QIW7 | Uncharacterized protein | 337 | | ATG3 family | autophagosome assembly [GO:0000045]; a | cytosol [GO:0005829]; phagophore assembly | Atg8-family ligase activity [GO:0019776] | cytosol [GO:0005829]; pha | UP000018817 |
| PPTG_17690 | W2PIM9 | Uncharacterized protein | 424 | | | | | | | UP000018817 |
| PPTG_11613 | W2Q6I6 | Intraflagellar Transport Protein 72/74 | 773 | | | intraciliary transport involved in cilium asse | cilium [GO:0005929]; intraciliary transport pa | beta-tubulin binding [GO:0048487] | cilium [GO:0005929]; intra | UP000018817 |
| PPTG_12918 | W2Q3V1 | Mitochondrial carrier protein | 307 | | Mitochondrial carrier (TC 2.A.29) family | | mitochondrial inner membrane [GO:000574 | transmembrane transporter activity [GO:002 | mitochondrial inner memb | UP000018817 |
| PPTG_12021 | W2Q6H4 | S-adenosyl-L-homocysteine hydrolase NAD binding domain-containing protein | 333 | | D-isomer specific 2-hydroxyacid dehydrogenase family | | | NAD binding [GO:0051287]; oxidoreductase | NAD binding [GO:0051287 | UP000018817 |
| PPTG_00628 | W2RFN9 | PX domain-containing protein | 300 | | | | | phosphatidylinositol binding [GO:0035091] | phosphatidylinositol bindir | UP000018817 |
| PPTG_15160 | W2PSD7 | Pre-mRNA-processing factor 19 (EC 2.3.2.27) | 525 | | WD repeat PRP19 family | DNA repair [GO:0006281]; mRNA splicing, v | cytoplasm [GO:0005737]; nucleoplasm [GO: | ubiquitin protein ligase activity [GO:0061630 | cytoplasm [GO:0005737]; | UP000018817 |
| PPTG_09272 | W2QGY2 | PX domain-containing protein | 303 | | | endocytosis [GO:0006897]; endosomal tran | cytoplasmic vesicle [GO:0031410]; plasma m | phosphatidylinositol binding [GO:0035091] | cytoplasmic vesicle [GO:0 | UP000018817 |
| PPTG_15088 | W2PTZ2 | COP9 signalosome complex subunit 3 | 465 | | CSN3 family | ubiquitin-dependent protein catabolic proc | COP9 signalosome [GO:0008180]; cytoplasm [GO:0005737] | | COP9 signalosome [GO:00 | UP000018817 |
| PPTG_02622 | W2RDV5 | Dienelactone hydrolase domain-containing protein | 251 | | | | | hydrolase activity [GO:0016787] | hydrolase activity [GO:001 | UP000018817 |
| PPTG_03369 | W2R4L9 | Glycosyl hydrolase family 30 TIM-barrel domain-containing protein | 539 | | Glycosyl hydrolase 30 family | glucosylceramide catabolic process [GO:00 | membrane [GO:0016020] | glucosylceramidase activity [GO:0004348] | membrane [GO:0016020]; | UP000018817 |
| PPTG_10317 | W2QE23 | non-specific serine/threonine protein kinase (EC 2.7.11.1) | 366 | | Protein kinase superfamily, AGC Ser/Thr protein kir | intracellular signal transduction [GO:0035556] | | ATP binding [GO:0005524]; protein serine/th | ATP binding [GO:0005524] | UP000018817 |
| PPTG_18933 | W2PE99 | Glucokinase | 351 | | | glycolytic process [GO:0006096] | | ATP binding [GO:0005524]; D-glucose bindin | ATP binding [GO:0005524] | UP000018817 |
| PPTG_18927 | W2PGP4 | Glucokinase | 382 | | | glycolytic process [GO:0006096] | | ATP binding [GO:0005524]; D-glucose bindin | ATP binding [GO:0005524] | UP000018817 |
| PPTG_15364 | W2PUV0 | Histidine ammonia-lyase | 599 | | PAL/histidase family | | | ammonia-lyase activity [GO:0016841] | ammonia-lyase activity [G | UP000018817 |
| PPTG_03064 | W2R3X2 | Uncharacterized protein | 446 | | | | | | | UP000018817 |
| PPTG_02204 | W2RAB7 | PX domain-containing protein | 203 | | | | endosome [GO:0005768] | phosphatidylinositol binding [GO:0035091] | endosome [GO:0005768]; | UP000018817 |
| PPTG_12102 | W2Q543 | Beta-glucosidase | 601 | | Glycosyl hydrolase 1 family | carbohydrate metabolic process [GO:0005975] | | beta-glucosidase activity [GO:0008422] | beta-glucosidase activity [ | UP000018817 |
| PPTG_07417 | W2QQ71 | SSD domain-containing protein | 1476 | | Patched family | sterol transport [GO:0015918] | membrane [GO:0016020] | sterol binding [GO:0032934] | membrane [GO:0016020]; | UP000018817 |
| PPTG_12242 | W2Q864 | D-3-phosphoglycerate dehydrogenase | 603 | | | amino acid biosynthetic process [GO:0008652] | | NAD binding [GO:0051287]; phosphoglycera | NAD binding [GO:0051287 | UP000018817 |
| PPTG_17994 | W2PHQ0 | RRM domain-containing protein | 467 | | | regulation of translation [GO:0006417] | | mRNA binding [GO:0003729] | mRNA binding [GO:00037 | UP000018817 |
| PPTG_13689 | W2PZH8 | DUF4097 domain-containing protein | 717 | | | | | | | UP000018817 |
| PPTG_10517 | W2QCZ5 | EF-hand domain-containing protein | 166 | | | | | calcium ion binding [GO:0005509] | calcium ion binding [GO:0 | UP000018817 |
| PPTG_18898 | W2PHG0 | AGC/RSK/RSKP90 protein kinase (AGC/RSK/RSKP90 protein kinase, variant 2) (AGC/RSK/RSKP90 | 1559 | | Protein kinase superfamily, AGC Ser/Thr protein kinase family, RAC subfamily | | | ATP binding [GO:0005524]; protein serine/th | ATP binding [GO:0005524] | UP000018817 |
| PPTG_11742 | W2Q812 | Uncharacterized protein | 243 | | | | | | | UP000018817 |
| PPTG_11824 | W2Q8E4 | Enoyl reductase (ER) domain-containing protein | 348 | | | | | oxidoreductase activity [GO:0016491] | oxidoreductase activity [G | UP000018817 |
| PPTG_09033 | W2QFZ3 | CAAX prenyl protease (EC 3.4.24.84) | 484 | | Peptidase M48A family | CAAX-box protein processing [GO:0071586] | endoplasmic reticulum membrane [GO:0005 | metal ion binding [GO:0046872]; metalloenc | endoplasmic reticulum me | UP000018817 |
| PPTG_04171 | W2QZJ2 | CRAL-TRIO domain-containing protein | 656 | | | | | | | UP000018817 |
| PPTG_06043 | W2QXG0 | Ribosomal protein L37a | 95 | | Eukaryotic ribosomal protein eL43 family | translation [GO:0006412] | ribonucleoprotein complex [GO:1990904]; rit | structural constituent of ribosome [GO:0003 | ribonucleoprotein complex | UP000018817 |
| PPTG_15678 | W2PTA0 | NADH dehydrogenase [ubiquinone] 1 alpha subcomplex subunit 5 | 116 | | Complex I NDUFA5 subunit family | respiratory electron transport chain [GO:00 | mitochondrial inner membrane [GO:0005743] | | mitochondrial inner memb | UP000018817 |
| PPTG_20426 | W2P8A2 | Cytochrome b | 157 | | | mitochondrial electron transport, ubiquinol | mitochondrial inner membrane [GO:000574 | metal ion binding [GO:0046872]; oxidoreduc | mitochondrial inner memb | UP000018817 |
| PPTG_01253 | W2R8P0 | Tr-type G domain-containing protein | 986 | | | mRNA splicing, via spliceosome [GO:00003 | cytosol [GO:0005829]; U2-type catalytic step | GTP binding [GO:0005525]; GTPase activity [ | cytosol [GO:0005829]; U2- | UP000018817 |
| PPTG_08421 | W2QKJ6 | PLAC8 family protein | 160 | | | | | | | UP000018817 |
| PPTG_17908 | W2PKZ9 | TOG domain-containing protein | 712 | | | | | | | UP000018817 |
| PPTG_11868 | W2QAA8 | Uncharacterized protein | 312 | | Mitochondrial carrier (TC 2.A.29) family | fatty acid beta-oxidation [GO:0006635]; per | peroxisomal membrane [GO:0005778] | ADP transmembrane transporter activity [G | peroxisomal membrane [G | UP000018817 |
| PPTG_11871 | W2Q8I8 | Uncharacterized protein | 209 | | | | | | | UP000018817 |
| PPTG_08002 | W2QLC7 | Protein root UVB sensitive/RUS domain-containing protein | 492 | | RUS1 family | | | | | UP000018817 |
| PPTG_19369 | W2PDE9 | Malic enzyme | 596 | | Malic enzymes family | malate metabolic process [GO:0006108] | | malate dehydrogenase (decarboxylating) (N | malate dehydrogenase (de | UP000018817 |
| PPTG_05510 | W2QXY0 | Ribosomal RNA small subunit methyltransferase B | 608 | | Class I-like SAM-binding methyltransferase superfa | maturation of LSU-rRNA [GO:0000470]; rRN | nucleolus [GO:0005730] | RNA binding [GO:0003723]; rRNA (cytosine- | nucleolus [GO:0005730]; | UP000018817 |
| PPTG_07371 | W2QQH5 | Uncharacterized protein | 1268 | | ABC transporter superfamily, ABCB family, Multidr | oligopeptide export from mitochondrion [G | mitochondrial inner membrane [GO:000574 | ABC-type oligopeptide transporter activity [G | mitochondrial inner memb | UP000018817 |
| PPTG_19683 | W2PCC7 | Proteasome subunit alpha type | 149 | | Peptidase T1A family | ubiquitin-dependent protein catabolic proc | cytoplasm [GO:0005737]; nucleus [GO:0005634]; proteasome core complex, alpha-subur | | cytoplasm [GO:0005737]; | UP000018817 |
| PPTG_14695 | W2PVH8 | Uncharacterized protein | 217 | | | | | | | UP000018817 |
| PPTG_01441 | W2R7D6 | NodB homology domain-containing protein | 340 | | | carbohydrate metabolic process [GO:0005975] | | hydrolase activity, acting on carbon-nitrogen | hydrolase activity, acting o | UP000018817 |
| PPTG_05377 | W2QWM2 | START domain-containing protein | 240 | | | | cytoplasm [GO:0005737] | lipid binding [GO:0008289] | cytoplasm [GO:0005737]; | UP000018817 |
| PPTG_15677 | W2PR44 | U6 snRNA-associated Sm-like protein LSm1 | 102 | | SnRNP Sm proteins family | deadenylation-dependent decapping of nuc | Lsm1-7-Pat1 complex [GO:1990726]; P-body | mRNA binding [GO:0003729] | Lsm1-7-Pat1 complex [GO | UP000018817 |
| PPTG_06146 | W2QRY7 | RNA helicase (EC 3.6.4.13) | 458 | | | | | ATP binding [GO:0005524]; hydrolase activity | ATP binding [GO:0005524] | UP000018817 |
| PPTG_08964 | W2QK91 | type II protein arginine methyltransferase (EC 2.1.1.320) | 698 | | NDUFAF7 family | methylation [GO:0032259] | mitochondrion [GO:0005739] | protein-arginine omega-N symmetric methyl | mitochondrion [GO:0005 | UP000018817 |
| PPTG_19041 | W2PDA0 | Probable acetate kinase (EC 2.7.2.1) (Acetokinase) | 428 | | Acetokinase family | acetate metabolic process [GO:0006083]; acetyl-CoA biosynthetic process [GO:0006085 | | acetate kinase activity [GO:0008776]; ATP bi | acetate kinase activity [GC | UP000018817 |
| PPTG_10899 | W2QBH4 | Ryanodine-inositol 1,4,5-triphosphate receptor Ca2 channel (RIR-CaC) family protein | 1386 | | Polycystin family | | membrane [GO:0016020] | calcium ion binding [GO:0005509] | membrane [GO:0016020]; | UP000018817 |
| PPTG_09548 | W2QFB9 | Sm protein B | 262 | | SnRNP SmB/SmN family | mRNA splicing, via spliceosome [GO:00003 | catalytic step 2 spliceosome [GO:0071013]; | RNA binding [GO:0003723]; snRNP binding [ | catalytic step 2 spliceoson | UP000018817 |
| PPTG_17414 | W2PM38 | Uncharacterized protein | 444 | | | | | | | UP000018817 |
| PPTG_16503 | W2PQH7 | Peptidyl-prolyl cis-trans isomerase (PPIase) (EC 5.2.1.8) | 226 | | Cyclophilin-type PPIase family | protein folding [GO:0006457] | cytoplasm [GO:0005737] | cyclosporin A binding [GO:0016018]; peptidy | cytoplasm [GO:0005737]; | UP000018817 |
| PPTG_05609 | W2QXK5 | Uncharacterized protein | 319 | | GTR/RAG GTP-binding protein family | cellular response to starvation [GO:000926 | endomembrane system [GO:0012505]; Gtr1- | GTP binding [GO:0005525]; GTPase activity [ | endomembrane system [G | UP000018817 |
| PPTG_04797 | W2R4Q2 | FAD/NAD(P)-binding domain-containing protein | 393 | | FAD-dependent oxidoreductase family | | cytoplasm [GO:0005737] | electron-transferring-flavoprotein dehydroge | cytoplasm [GO:0005737]; | UP000018817 |
| PPTG_05256 | W2QWZ9 | Mitochondrial proton/calcium exchanger protein (Leucine zipper-EF-hand-containing transmembra | 795 | | LETM1 family | intracellular monoatomic cation homeosta | mitochondrial inner membrane [GO:000574 | antiporter activity [GO:0015297]; calcium io | mitochondrial inner memb | UP000018817 |
| PPTG_04751 | W2R1X7 | DnaJ homologue subfamily C GRV2/DNAJC13 N-terminal domain-containing protein | 2795 | | | endosome organization [GO:0007032]; rece | endosome membrane [GO:0010008] | | endosome membrane [GC | UP000018817 |
| PPTG_12540 | W2Q4F4 | Expansin-like EG45 domain-containing protein | 384 | | | | | | | UP000018817 |
| PPTG_13651 | W2PZM0 | Coatomer subunit epsilon | 292 | | COPE family | endoplasmic reticulum to Golgi vesicle-me | COPI vesicle coat [GO:0030126]; Golgi meml | structural molecule activity [GO:0005198] | COPI vesicle coat [GO:003 | UP000018817 |
| PPTG_10197 | W2QDJ2 | 40S ribosomal protein S28 | 67 | | Eukaryotic ribosomal protein eS28 family | maturation of SSU-rRNA [GO:0030490]; rib | cytosolic small ribosomal subunit [GO:00226 | structural constituent of ribosome [GO:0003 | cytosolic small ribosomal | UP000018817 |
| PPTG_16312 | W2PQX2 | Tr-type G domain-containing protein | 563 | | TRAFAC class translation factor GTPase superfamily, Classic translation factor GTPase family, | | cytoplasm [GO:0005737] | GTP binding [GO:0005525]; GTPase activity [ | cytoplasm [GO:0005737]; | UP000018817 |
| PPTG_18989 | W2PGK0 | PH domain-containing protein | 1213 | | | | cytoplasm [GO:0005737] | phosphatidylinositol-3,4-bisphosphate 4-ph | cytoplasm [GO:0005737]; | UP000018817 |
| PPTG_06494 | W2QV37 | Vacuolar protein 8 | 270 | | Beta-catenin family | nucleus-vacuole junction assembly [GO:00 | vacuolar membrane [GO:0005774] | protein-membrane adaptor activity [GO:004 | vacuolar membrane [GO:0 | UP000018817 |

| | | | | | | | | | |
|---|---|---|---|---|---|---|---|---|---|
| PPTG_18900 | W2PF40 | Ubiquitin carboxyl-terminal hydrolase (EC 3.4.19.12) | 260 | Peptidase C12 family | protein deubiquitination [GO:0016579]; ubi | cytoplasm [GO:0005737] | cysteine-type deubiquitinase activity [GO:00 | cytoplasm [GO:0005737]; | UP000018817 |
| PPTG_00858 | W2RHB3 | NADH dehydrogenase [ubiquinone] 1 alpha subcomplex subunit 3 | 73 | | | | | | UP000018817 |
| PPTG_10233 | W2QDU2 | TIL domain-containing protein | 146 | | | | | | UP000018817 |
| PPTG_13578 | W2Q3T3 | Uncharacterized protein | 690 | | | | | | UP000018817 |
| PPTG_19674 | W2PE72 | Transglutaminase elicitor | 579 | | | | aminoacyltransferase activity [GO:0016755] | aminoacyltransferase acti | UP000018817 |
| PPTG_04926 | W2R356 | Ribosomal eL28/Mak16 domain-containing protein | 139 | Eukaryotic ribosomal protein eL28 family | translation [GO:0006412] | ribonucleoprotein complex [GO:1990904]; rib | structural constituent of ribosome [GO:0003 | ribonucleoprotein complex | UP000018817 |
| PPTG_13556 | W2Q1W8 | WASH complex subunit strumpellin | 1022 | Strumpellin family | actin filament polymerization [GO:0030041 | endosome [GO:0005768]; WASH complex [GO:0071203] | | endosome [GO:0005768]; | UP000018817 |
| PPTG_06328 | W2QSM0 | Vacuolar protein 14 C-terminal Fig4-binding domain-containing protein | 788 | VAC14 family | phosphatidylinositol biosynthetic process [( | endosome membrane [GO:0010008]; PAS complex [GO:0070772] | | endosome membrane [GC | UP000018817 |
| PPTG_10842 | W2QC49 | Sodium/hydrogen exchanger | 517 | Monovalent cation:proton antiporter 1 (CPA1) trans | regulation of intracellular pH [GO:0051453] | Golgi membrane [GO:0000139]; plasma men | potassium:proton antiporter activity [GO:001 | Golgi membrane [GO:000( | UP000018817 |
| PPTG_01294 | W2R6C5 | Mitogen-activated protein kinase (EC 2.7.11.24) | 374 | Protein kinase superfamily, Ser/Thr protein kinase family, MAP kinase subfamily | | | ATP binding [GO:0005524]; MAP kinase activ | ATP binding [GO:0005524] | UP000018817 |
| PPTG_15779 | W2PQC6 | Signal recognition particle subunit SRP68 (SRP68) | 614 | SRP68 family | SRP-dependent cotranslational protein targ | cytosol [GO:0005829]; endoplasmic reticulur | 7S RNA binding [GO:0008312]; endoplasmic | cytosol [GO:0005829]; enc | UP000018817 |
| PPTG_02908 | W2RDL2 | Developmentally-regulated GTP-binding protein 1 | 369 | | | | GTP binding [GO:0005525]; GTPase activity [ | GTP binding [GO:0005525] | UP000018817 |
| PPTG_15692 | W2PS44 | Uncharacterized protein | 1174 | MYBBP1A family | regulation of DNA-templated transcription [ | nucleolus [GO:0005730] | DNA binding [GO:0003677] | nucleolus [GO:0005730]; [ | UP000018817 |
| PPTG_14255 | W2PY13 | Uncharacterized protein | 268 | | | | | | UP000018817 |
| PPTG_07069 | W2QPD9 | Ribosomal protein S9 | 322 | Universal ribosomal protein uS9 family | translation [GO:0006412] | cytosolic small ribosomal subunit [GO:00226 | RNA binding [GO:0003723]; structural consti | cytosolic small ribosomal | UP000018817 |
| PPTG_02357 | W2RB05 | PWWP domain-containing protein | 482 | | | | | | UP000018817 |
| PPTG_11928 | W2Q9P5 | Succinate--CoA ligase [ADP-forming] subunit alpha, mitochondrial (EC 6.2.1.5) (Succinyl-CoA syntl | 318 | Succinate/malate CoA ligase alpha subunit family | tricarboxylic acid cycle [GO:0006099] | mitochondrion [GO:0005739]; succinate-CoA | nucleotide binding [GO:0000166]; succinate | mitochondrion [GO:0005 | UP000018817 |
| PPTG_11770 | W2Q859 | Sensor domain-containing protein | 328 | | | | | | UP000018817 |
| PPTG_13437 | W2Q289 | Acid phosphatase | 480 | Histidine acid phosphatase family | | | phosphatase activity [GO:0016791] | phosphatase activity [GO:( | UP000018817 |
| PPTG_17517 | W2PLE8 | Thioredoxin domain-containing protein | 190 | | photoreceptor cell maintenance [GO:0045494] | | | photoreceptor cell mainte | UP000018817 |
| PPTG_08457 | W2QNA3 | acetyl-CoA C-acyltransferase (EC 2.3.1.16) | 411 | Thiolase-like superfamily, Thiolase family | fatty acid beta-oxidation [GO:0006635]; phe | peroxisome [GO:0005777] | acetyl-CoA C-acyltransferase activity [GO:00 | peroxisome [GO:0005777] | UP000018817 |
| PPTG_12597 | W2Q2G9 | Calpain catalytic domain-containing protein | 1272 | Peptidase C2 family | proteolysis [GO:0006508] | | calcium-dependent cysteine-type endopepti | calcium-dependent cystei | UP000018817 |
| PPTG_12781 | W2Q0V3 | protein-serine/threonine phosphatase (EC 3.1.3.16) | 1102 | PP2C family | | cAMP-dependent protein kinase complex [GC | ATP binding [GO:0005524]; cAMP-dependen | cAMP-dependent protein k | UP000018817 |
| PPTG_05384 | W2QYU4 | DUF3160 domain-containing protein | 789 | | | | | | UP000018817 |
| PPTG_11486 | W2QBJ4 | non-specific serine/threonine protein kinase (EC 2.7.11.1) | 3242 | PI3/PI4-kinase family | negative regulation of macroautophagy [GO | cytoplasm [GO:0005737]; nucleus [GO:0005( | ATP binding [GO:0005524]; protein serine/th | cytoplasm [GO:0005737]; | UP000018817 |
| PPTG_01179 | W2RHY6 | Dynein light intermediate chain | 435 | Dynein light intermediate chain family | microtubule cytoskeleton organization [GO: | centrosome [GO:0005813]; cytoplasmic dyne | ATP binding [GO:0005524]; dynein heavy cha | centrosome [GO:0005813 | UP000018817 |
| PPTG_10723 | W2QCK8 | ABC transmembrane type-1 domain-containing protein | 810 | ABC transporter superfamily, ABCB family, Multidr | oligopeptide export from mitochondrion [GC | mitochondrial inner membrane [GO:000574 | ABC-type oligopeptide transporter activity [G | mitochondrial inner memb | UP000018817 |
| PPTG_13081 | W2Q5G8 | Cytochrome b-c1 complex subunit 6 | 67 | UQCRH/QCR6 family | mitochondrial electron transport, ubiquinol | mitochondrial inner membrane [GO:0005743] | | mitochondrial inner memb | UP000018817 |
| PPTG_03685 | W2R5U0 | Protein MON2 homolog | 1720 | | | | | | UP000018817 |
| PPTG_04035 | W2R1P0 | ADP-ribosylation factor | 215 | Small GTPase superfamily, Arf family | protein transport [GO:0015031]; vesicle-me | Golgi apparatus [GO:0005794] | GTP binding [GO:0005525]; GTPase activity [ | Golgi apparatus [GO:0005 | UP000018817 |
| PPTG_18224 | W2PHU6 | Dynein assembly factor with WD repeat domains 1 | 417 | | axonemal dynein complex assembly [GO:0070286] | | | axonemal dynein complex | UP000018817 |
| PPTG_18700 | W2PEZ4 | Uncharacterized protein | 1893 | | mRNA export from nucleus [GO:0006406]; p | nuclear pore [GO:0005643] | structural constituent of nuclear pore [GO:00 | nuclear pore [GO:0005643 | UP000018817 |
| PPTG_01192 | W2RK99 | Uncharacterized protein | 857 | | | | calcium ion binding [GO:0005509] | calcium ion binding [GO:0( | UP000018817 |
| PPTG_08290 | W2QLR5 | ACB domain-containing protein | 88 | ACBP family | fatty acid metabolic process [GO:0006631] | | fatty-acyl-CoA binding [GO:0000062] | fatty-acyl-CoA binding [GO | UP000018817 |
| PPTG_11875 | W2Q8K1 | AAA+ ATPase domain-containing protein | 873 | Peptidase M41 family; AAA ATPase family | mitochondrial protein processing [GO:0034 | m-AAA complex [GO:0005745] | ATP binding [GO:0005524]; ATP hydrolysis ac | m-AAA complex [GO:0005 | UP000018817 |
| PPTG_08250 | W2QJS2 | Ribosomal protein L22 | 201 | Universal ribosomal protein uL22 family | translation [GO:0006412] | mitochondrial large ribosomal subunit [GO:0( | structural constituent of ribosome [GO:0003 | mitochondrial large riboso | UP000018817 |
| PPTG_17872 | W2PIG4 | RRM domain-containing protein | 206 | | | | RNA binding [GO:0003723] | RNA binding [GO:0003723 | UP000018817 |
| PPTG_09950 | W2QF40 | THO complex subunit 2 | 1711 | THOC2 family | mRNA export from nucleus [GO:0006406]; r | THO complex part of transcription export com | mRNA binding [GO:0003729] | THO complex part of trans | UP000018817 |
| PPTG_09135 | W2QGH1 | Phosphoglucomutase-2 | 609 | Phosphohexose mutase family | glucose metabolic process [GO:0006006]; p | cytoplasm [GO:0005737]; nucleus [GO:0005( | magnesium ion binding [GO:0000287]; phos | cytoplasm [GO:0005737]; | UP000018817 |
| PPTG_12930 | W2Q3W5 | C2 domain-containing protein | 328 | | | | | | UP000018817 |
| PPTG_10878 | W2QAG6 | Translation initiation factor beta propellor-like domain-containing protein | 303 | | | Cdc73/Paf1 complex [GO:0016593] | | Cdc73/Paf1 complex [GO:( | UP000018817 |
| PPTG_09573 | W2QFG6 | C3H1-type domain-containing protein | 404 | | | | zinc ion binding [GO:0008270] | zinc ion binding [GO:0008 | UP000018817 |
| PPTG_00139 | W2RDQ1 | Enoyl reductase (ER) domain-containing protein | 332 | | | | oxidoreductase activity [GO:0016491] | oxidoreductase activity [G( | UP000018817 |
| PPTG_11476 | W2QBI4 | N-acetyltransferase domain-containing protein | 472 | PhzF family | | cytoplasm [GO:0005737] | acyltransferase activity, transferring groups ( | cytoplasm [GO:0005737]; | UP000018817 |
| PPTG_08193 | W2QLD5 | Uroporphyrinogen decarboxylase (EC 4.1.1.37) | 408 | Uroporphyrinogen decarboxylase family | protoporphyrinogen IX biosynthetic process | cytosol [GO:0005829] | uroporphyrinogen decarboxylase activity [GC | cytosol [GO:0005829]; uro | UP000018817 |
| PPTG_18647 | W2PFZ5 | Glycylpeptide N-tetradecanoyltransferase (EC 2.3.1.97) | 372 | NMT family | | cytoplasm [GO:0005737] | glycylpeptide N-tetradecanoyltransferase ac | cytoplasm [GO:0005737]; | UP000018817 |
| PPTG_18888 | W2PFJ4 | Vacuolar protein sorting-associated protein 33A | 629 | STXBP/unc-18/SEC1 family | vesicle-mediated transport [GO:0016192] | | | vesicle-mediated transpor | UP000018817 |
| PPTG_01356 | W2R8I4 | 6,7-dimethyl-8-ribityllumazine synthase (DMRL synthase) (EC 2.5.1.78) | 215 | DMRL synthase family | riboflavin biosynthetic process [GO:000923 | riboflavin synthase complex [GO:0009349] | 6,7-dimethyl-8-ribityllumazine synthase acti | riboflavin synthase comple | UP000018817 |
| PPTG_02902 | W2RD51 | Uncharacterized protein | 276 | | | | | | UP000018817 |
| PPTG_16621 | W2PN53 | Uncharacterized protein | 302 | Mitochondrial carrier (TC 2.A.29) family | mitochondrial ADP transmembrane transpc | mitochondrial inner membrane [GO:000574 | ATP:ADP antiporter activity [GO:0005471] | mitochondrial inner memb | UP000018817 |
| PPTG_12174 | W2Q698 | Uncharacterized protein | 319 | | | membrane [GO:0016020] | | membrane [GO:0016020] | UP000018817 |
| PPTG_14016 | W2PZ76 | ELMO domain-containing protein | 938 | | | | | | UP000018817 |
| PPTG_01821 | W2RAX1 | Sec20 C-terminal domain-containing protein | 254 | | protein transport [GO:0015031]; vesicle fusi | endoplasmic reticulum membrane [GO:0005 | SNAP receptor activity [GO:0005484]; SNARI | endoplasmic reticulum me | UP000018817 |
| PPTG_11095 | W2Q907 | Probable pectate lyase F (EC 4.2.2.2) | 249 | Polysaccharide lyase 3 family | pectin catabolic process [GO:0045490] | extracellular region [GO:0005576] | pectate lyase activity [GO:0030570] | extracellular region [GO:00 | UP000018817 |
| PPTG_06763 | W2QQL7 | Uncharacterized protein | 396 | Sodium:solute symporter (SSF) (TC 2.A.21) family | | plasma membrane [GO:0005886] | spermidine transmembrane transporter acti | plasma membrane [GO:00 | UP000018817 |
| PPTG_05390 | W2QXG2 | Sphingomyelin synthase-like domain-containing protein | 342 | Sphingomyelin synthase family | ceramide biosynthetic process [GO:004651 | endoplasmic reticulum membrane [GO:0005 | ceramide cholinephosphotransferase activit | endoplasmic reticulum me | UP000018817 |
| PPTG_08933 | W2QJ51 | Polycystin cation channel PKD1/PKD2 domain-containing protein | 700 | | | membrane [GO:0016020] | | membrane [GO:0016020] | UP000018817 |
| PPTG_13987 | W2Q004 | Golgi apparatus membrane protein TVP15 | 208 | | | membrane [GO:0016020] | | membrane [GO:0016020] | UP000018817 |
| PPTG_13455 | W2Q1F0 | SRP54-type proteins GTP-binding domain-containing protein | 601 | GTP-binding SRP family | intracellular protein transport [GO:0006886 | signal recognition particle receptor complex [ | GTP binding [GO:0005525]; GTPase activity [ | signal recognition particle | UP000018817 |
