## Supplementary table 3 for "Zoospore-derived extracellular vesicles in the flagellated stramenopile *Phytophthora parasitica*"

**Title: Orthologs of Evs-related proteins between *P.parasitica* and *P. infestans***

| Uniprot Accession *P. parasitica* | Uniprot Accession *P. infestans* | BlastP e-Value |
|---|---|---|
| W2PR52 | D0NYH9 | 1E-180 |
| W2PT43 | D0N0G8 | 2E-180 |
| W2PTN3 | D0P2Y0 | 6E-180 |
| W2QI94 | D0NSR2 | 1E-179 |
| W2R9T5 | D0MUV7 | 3E-179 |
| W2R7K7 | D0NBC1 | 3E-179 |
| W2QEH7 | D0P3W0 | 6E-179 |
| W2Q4N9 | D0P3M8 | 2E-178 |
| W2QMB5 | D0NIH7 | 3E-178 |
| W2R387 | D0MYT8 | 1E-177 |
| W2PWM0 | D0N0B9 | 2E-177 |
| W2R5N9 | D0NBD7 | 2E-177 |
| W2PM53 | D0NPZ0 | 7E-177 |
| W2QVT2 | D0MWA2 | 2E-176 |
| W2PWU7 | D0N4E3 | 2E-176 |
| W2PII3 | D0N1E5 | 1E-175 |
| W2PNP8 | D0N8B4 | 1E-175 |
| W2PDI9 | D0NCV5 | 1E-175 |
| W2RJP0 | D0MU84 | 6E-175 |
| W2RA05 | D0MV15 | 3E-173 |
| W2R988 | D0NPD3 | 3E-173 |
| W2PXM6 | D0MXP8 | 2E-172 |
| W2QVB0 | D0NPR2 | 4E-172 |
| W2RAR6 | D0NFS0 | 1E-171 |
| W2QJJ8 | D0NMP9 | 1E-170 |
| W2QCJ3 | D0MQP9 | 4E-170 |
| W2Q071 | D0N4V4 | 4E-170 |
| W2QA46 | D0NRU9 | 8E-170 |
| W2QV37 | D0NCY9 | 1E-169 |
| W2RDV5 | D0N2V9 | 4E-169 |
| W2RFJ7 | D0NW02 | 3E-167 |
| W2R0V7 | D0NJ99 | 4E-167 |
| W2REK3 | D0N327 | 6E-167 |
| W2PP34 | D0MR35 | 3E-166 |
| W2QFA2 | D0NZL4 | 3E-166 |
| W2QKH4 | D0NA40 | 5E-166 |
| W2R8G4 | D0NCM1 | 2E-165 |
| W2QPD9 | D0NKZ4 | 2E-165 |
| W2Q180 | D0N597 | 3E-165 |
| W2QWG2 | D0P351 | 4E-164 |
| W2R2C4 | D0N7N5 | 5E-163 |
| W2QTY4 | D0MYJ9 | 3E-162 |
| W2Q8B8 | D0NKB9 | 8E-162 |
| W2PZ59 | D0NQL6 | 9E-161 |
| W2PE68 | D0NDW3 | 1E-160 |
| W2PRY4 | D0P074 | 2E-160 |
| W2RI96 | D0MUH0 | 2E-159 |
| W2PFF6 | D0NDW0 | 4E-159 |
| W2PCU8 | D0P371 | 4E-159 |
| W2PKT7 | D0P070 | 1E-158 |
| W2R7Y3 | D0NBK1 | 2E-158 |
| W2QY84 | D0NB06 | 3E-158 |
| W2RH96 | D0NF02 | 6E-158 |
| W2PI57 | D0MVR9 | 2E-157 |
| W2Q0E7 | D0NRZ7 | 3E-157 |
| W2Q1S8 | D0P1Y0 | 1E-156 |
| W2RA54 | D0NG89 | 2E-155 |
| W2QWM2 | D0MWU3 | 1E-153 |
| W2Q812 | D0NE34 | 1E-153 |
| W2PRM0 | D0N0M4 | 3E-153 |
| W2QEC2 | D0MS46 | 8E-153 |
| W2QLY9 | D0MRA7 | 6E-151 |
| W2QIQ6 | D0NMP8 | 6E-151 |
| W2PQH7 | D0N727 | 2E-150 |
| W2PUM5 | D0NUA4 | 2E-150 |
| W2QZ92 | D0N7U0 | 3E-150 |
| W2QFB9 | D0NZ60 | 1E-149 |

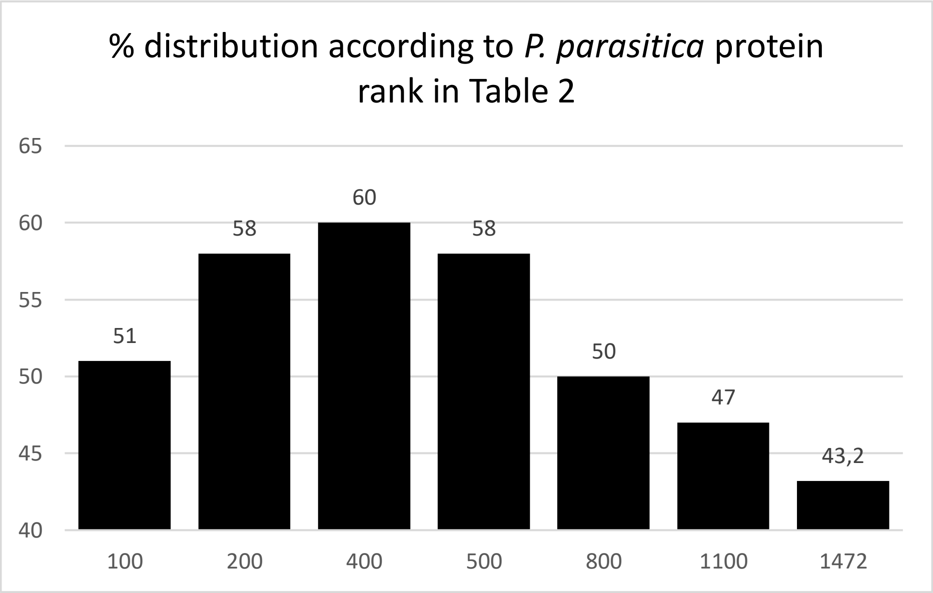


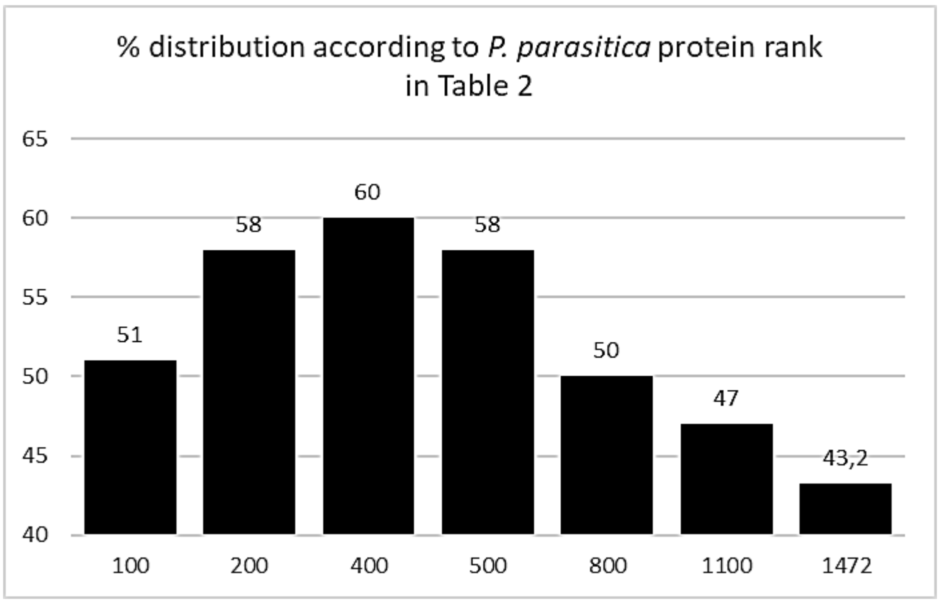

| | | |
|---|---|---|
| W2Q2M3 | D0NRX5 | 3E-149 |
| W2R578 | D0NB62 | 5E-149 |
| W2PW48 | D0N068 | 3E-148 |
| W2R3T1 | D0NHJ8 | 7E-148 |
| W2QJ22 | D0MU25 | 1E-147 |
| W2RAI7 | D0MXT1 | 2E-146 |
| W2QPY1 | D0NI70 | 1E-145 |
| W2QBA4 | D0NRE5 | 2E-145 |
| W2R4H7 | D0N242 | 3E-145 |
| W2QZ40 | D0MWP5 | 4E-145 |
| W2QQK3 | D0NSN5 | 6E-145 |
| W2QRV7 | D0NT87 | 4E-144 |
| W2RA51 | D0NGF6 | 7E-144 |
| W2QGK3 | D0RLZ3 | 9E-144 |
| W2PLH3 | D0NK11 | 2E-143 |
| W2PJS8 | D0NYW1 | 5E-143 |
| W2RFQ0 | D0NXV2 | 7E-143 |
| W2R9F4 | D0NCU9 | 1E-142 |
| W2QK24 | D0MRA6 | 1E-141 |
| W2R2P0 | D0MYZ2 | 2E-141 |
| W2PQU8 | D0N444 | 4E-141 |
| W2PX63 | D0N9R0 | 4E-140 |
| W2RAX8 | D0P4J4 | 5E-140 |
| W2RHW6 | D0P366 | 8E-139 |
| W2QSS5 | D0ND55 | 1E-137 |
| W2QQ00 | D0MYC1 | 2E-137 |
| W2PQY3 | D0MZV7 | 3E-137 |
| W2PR58 | D0NC11 | 2E-136 |
| W2PPB1 | D0NLH6 | 2E-136 |
| W2RGQ3 | D0MTV1 | 3E-136 |
| W2RJF1 | D0MU36 | 3E-136 |
| W2RHW9 | D0MU72 | 8E-136 |
| W2PYR6 | D0N050 | 9E-136 |
| W2RG96 | D0P2X8 | 2E-135 |
| W2QN94 | D0N1B1 | 3E-135 |
| W2R3Y6 | D0N258 | 6E-135 |
| W2QJ05 | D0NMU9 | 6E-135 |
| W2PE04 | D0NES3 | 1E-134 |
| W2PX94 | D0MZH3 | 4E-133 |
| W2QB83 | D0MUS0 | 9E-133 |
| W2QS85 | D0NUP0 | 2E-132 |
| W2QL17 | D0N999 | 4E-132 |
| W2QHU6 | D0NMR1 | 1E-131 |
| W2QFW8 | D0MSE7 | 4E-131 |
| W2PWA4 | D0P415 | 7E-131 |
| W2QN40 | D0NA49 | 8E-131 |
| W2R323 | D0MYX2 | 1E-129 |
| W2PTH4 | D0NCA2 | 2E-129 |
| W2PJH1 | D0N1D6 | 5E-129 |
| W2QA19 | D0MQP5 | 1E-128 |
| W2QWL5 | D0MWP7 | 1E-128 |
| W2PS44 | D0NC13 | 1E-128 |
| W2QGX3 | D0NMN8 | 3E-128 |
| W2PQ00 | D0N014 | 4E-128 |
| W2PN09 | D0MY66 | 8E-128 |
| W2R5S3 | D0NBG4 | 4E-127 |
| W2RA80 | D0NG62 | 3E-126 |
| W2PSK9 | D0MZZ4 | 9E-126 |
| W2PQA5 | D0MZQ2 | 1E-125 |
| W2RC07 | D0NGA4 | 1E-125 |
| W2RCV6 | D0N2P0 | 2E-124 |
| W2QD77 | D0NNT2 | 4E-124 |
| W2RAB7 | D0NGC4 | 5E-124 |
| W2RKB5 | D0N4W2 | 6E-124 |
| W2Q5Y3 | D0RLW4 | 7E-124 |
| W2PCT2 | D0NSD5 | 2E-123 |
| W2PPP1 | D0NP68 | 7E-123 |
| W2Q7Y2 | D0NUD9 | 6E-122 |
| W2QTG8 | D0NTC7 | 1E-121 |
| W2R807 | D0NC86 | 1E-120 |
| W2PRN2 | D0NHQ2 | 1E-118 |

| | | | |
|---|---|---|---|
| W2PH27 | D0P0S3 | | 4E-118 |
| W2PDE7 | D0N1F9 | | 2E-117 |
| W2PDZ7 | D0NLV3 | | 2E-115 |
| W2Q8C4 | D0NVP0 | | 2E-115 |
| W2RDL2 | D0N3B4 | | 9E-115 |
| W2R1P0 | D0NJH9 | | 1E-114 |
| W2R5D7 | D0MYT5 | | 3E-114 |
| W2PMJ0 | D0NLR7 | | 1E-113 |
| W2REA4 | D0N362 | | 2E-113 |
| W2QG53 | D0NZ33 | | 3E-113 |
| W2QYZ1 | D0MWK2 | | 1E-112 |
| W2R7Z6 | D0NC90 | | 1E-112 |
| W2PL80 | D0NKM8 | | 1E-112 |
| W2PG71 | D0MVW0 | | 4E-112 |
| W2PKV5 | D0MVC2 | | 9E-112 |
| W2PXC1 | D0N9N0 | | 1E-111 |
| W2R3H0 | D0N2F7 | | 4E-111 |
| W2Q2I1 | D0NS64 | | 3E-110 |
| W2RBH6 | D0NEJ2 | | 7E-110 |
| W2Q1L6 | D0MY70 | | 3E-109 |
| W2R6X3 | D0NV58 | | 4E-108 |
| W2QX04 | D0MT79 | | 2E-107 |
| W2Q0K2 | D0N5B1 | | 2E-107 |
| W2PH16 | D0N3Z7 | | 8E-107 |
| W2PPD3 | D0NP43 | | 1E-105 |
| W2PT70 | D0MZN9 | | 3E-105 |
| W2PA86 | D0NIJ7 | | 8E-105 |
| W2REY1 | D0N7Z1 | | 2E-104 |
| W2QUJ2 | D0ND91 | | 4E-104 |
| W2PFI0 | D0N9U2 | | 1E-103 |
| W2QYT4 | D0N3T6 | | 5E-103 |
| W2PFI8 | D0N851 | | 3E-102 |
| W2R892 | D0NCD0 | | 3E-102 |
| W2R8H7 | D0NC73 | | 5E-101 |
| W2R213 | D0N7G4 | | 7E-101 |
| W2PU56 | D0MQ53 | 0.0 | |
| W2PW79 | D0MQ59 | 0.0 | |
| W2PU74 | D0MQ64 | 0.0 | |
| W2QCT9 | D0MQ72 | 0.0 | |
| W2QDG9 | D0MQ88 | 0.0 | |
| W2QBR2 | D0MQ98 | 0.0 | |
| W2QAG6 | D0MQF8 | 0.0 | |
| W2QBC6 | D0MQG5 | 0.0 | |
| W2QBT9 | D0MQJ7 | 0.0 | |
| W2QK27 | D0MQW5 | 0.0 | |
| W2QLD0 | D0MR53 | 0.0 | |
| W2QJJ1 | D0MR68 | 0.0 | |
| W2QM10 | D0MR97 | 0.0 | |
| W2QLP3 | D0MRF1 | 0.0 | |
| W2QJR4 | D0MRG2 | 0.0 | |
| W2QJQ3 | D0MRG4 | 0.0 | |
| W2QJM4 | D0MRI8 | 0.0 | |
| W2QLC3 | D0MRN0 | 0.0 | |
| W2PJ23 | D0MRQ9 | 0.0 | |
| W2QB73 | D0MRS4 | 0.0 | |
| W2PKD5 | D0MRU3 | 0.0 | |
| W2PJN4 | D0MRV6 | 0.0 | |
| W2PJM8 | D0MRW0 | 0.0 | |
| W2QEI6 | D0MS08 | 0.0 | |
| W2QF76 | D0MS56 | 0.0 | |
| W2QF04 | D0MSA5 | 0.0 | |
| W2QEZ6 | D0MSB1 | 0.0 | |
| W2QEW5 | D0MSD8 | 0.0 | |
| W2QEQ2 | D0MSJ1 | 0.0 | |
| W2PPP5 | D0MSN1 | 0.0 | |
| W2PFJ5 | D0MST8 | 0.0 | |
| W2PWM4 | D0MSZ0 | 0.0 | |
| W2PW23 | D0MT03 | 0.0 | |
| W2PWR3 | D0MT20 | 0.0 | |
| W2PWS4 | D0MT25 | 0.0 | |
| W2Q543 | D0MT52 | 0.0 | |

| | | |
|---|---|---|
| W2Q5J5 | D0MT54 | 0.0 |
| W2Q5M4 | D0MT62 | 0.0 |
| W2Q6J8 | D0MT70 | 0.0 |
| W2Q5T5 | D0MTA1 | 0.0 |
| W2Q6V4 | D0MTC5 | 0.0 |
| W2RGE9 | D0MTE8 | 0.0 |
| W2RG07 | D0MTH1 | 0.0 |
| W2RG20 | D0MTI1 | 0.0 |
| W2RIN1 | D0MTM0 | 0.0 |
| W2RIH1 | D0MTM6 | 0.0 |
| W2RGG3 | D0MTN1 | 0.0 |
| W2RGP1 | D0MTV6 | 0.0 |
| W2RHD9 | D0MTX5 | 0.0 |
| W2RJ64 | D0MU04 | 0.0 |
| W2RJB4 | D0MU13 | 0.0 |
| W2RHM7 | D0MU28 | 0.0 |
| W2RHQ1 | D0MU42 | 0.0 |
| W2RJH7 | D0MU57 | 0.0 |
| W2RHE9 | D0MU74 | 0.0 |
| W2RHI5 | D0MU98 | 0.0 |
| W2RHS3 | D0MUG4 | 0.0 |
| W2RHY6 | D0MUL8 | 0.0 |
| W2Q8W4 | D0MUN7 | 0.0 |
| W2Q9P5 | D0MUP3 | 0.0 |
| W2Q8U0 | D0MUQ5 | 0.0 |
| W2Q8P7 | D0MUR4 | 0.0 |
| W2QB87 | D0MUR6 | 0.0 |
| W2QHE9 | D0MUS3 | 0.0 |
| W2R752 | D0MUT9 | 0.0 |
| W2PAX2 | D0MUU6 | 0.0 |
| W2PDC9 | D0MUV1 | 0.0 |
| W2R7C5 | D0MUX1 | 0.0 |
| W2R7Y2 | D0MV08 | 0.0 |
| W2R7M4 | D0MV35 | 0.0 |
| W2PXY5 | D0MVI4 | 0.0 |
| W2PVE4 | D0MVI6 | 0.0 |
| W2PXD4 | D0MVK6 | 0.0 |
| W2PY60 | D0MVP3 | 0.0 |
| W2QVI7 | D0MVV7 | 0.0 |
| W2PI14 | D0MVW8 | 0.0 |
| W2QVJ2 | D0MVY1 | 0.0 |
| W2QVN0 | D0MW10 | 0.0 |
| W2QVS4 | D0MW28 | 0.0 |
| W2QVT6 | D0MW33 | 0.0 |
| W2QVK8 | D0MW53 | 0.0 |
| W2QVY7 | D0MWE3 | 0.0 |
| W2QW96 | D0MWF5 | 0.0 |
| W2QWD1 | D0MWH7 | 0.0 |
| W2QWZ9 | D0MWI5 | 0.0 |
| W2QWF0 | D0MWJ6 | 0.0 |
| W2QYJ3 | D0MWN4 | 0.0 |
| W2QWE8 | D0MWQ3 | 0.0 |
| W2QZ60 | D0MWR1 | 0.0 |
| W2QWH0 | D0MWS1 | 0.0 |
| W2QYR0 | D0MWT2 | 0.0 |
| W2QYU8 | D0MWV1 | 0.0 |
| W2QXA2 | D0MX24 | 0.0 |
| W2QXW8 | D0MX38 | 0.0 |
| W2QXP4 | D0MX79 | 0.0 |
| W2QXM1 | D0MXD7 | 0.0 |
| W2R0D6 | D0MXE0 | 0.0 |
| W2PU40 | D0MXI4 | 0.0 |
| W2PSU1 | D0MXL8 | 0.0 |
| W2PVT8 | D0MXP4 | 0.0 |
| W2PVQ2 | D0MXR3 | 0.0 |
| W2PWH7 | D0MXR4 | 0.0 |
| W2QDB7 | D0MXW0 | 0.0 |
| W2QF54 | D0MXW3 | 0.0 |
| W2QE49 | D0MXW9 | 0.0 |
| W2QE09 | D0MXY1 | 0.0 |
| W2Q284 | D0MY05 | 0.0 |

| | | |
|---|---|---|
| W2Q1J8 | D0MY08 | 0.0 |
| W2PD93 | D0MY63 | 0.0 |
| W2Q3G6 | D0MY71 | 0.0 |
| W2Q3J3 | D0MY93 | 0.0 |
| W2Q1S2 | D0MY99 | 0.0 |
| W2Q2U0 | sp\|D0M | 0.0 |
| W2Q238 | D0MYC5 | 0.0 |
| W2Q3T3 | D0MYD5 | 0.0 |
| W2Q4N4 | D0MYD7 | 0.0 |
| W2Q4Y4 | D0MYG6 | 0.0 |
| W2QVB3 | D0MYK1 | 0.0 |
| W2QCQ7 | D0MYM3 | 0.0 |
| W2PDU7 | D0MYP9 | 0.0 |
| W2R3G8 | D0MYR5 | 0.0 |
| W2Q4G3 | D0MZ63 | 0.0 |
| W2Q3H5 | D0MZ74 | 0.0 |
| W2PXK9 | D0MZB9 | 0.0 |
| W2PVS0 | D0MZD0 | 0.0 |
| W2PXI7 | D0MZD2 | 0.0 |
| W2PWR6 | D0MZE3 | 0.0 |
| W2QW86 | D0MZG4 | 0.0 |
| W2P9U9 | D0MZR7 | 0.0 |
| W2PS91 | D0MZU0 | 0.0 |
| W2PSK4 | D0MZZ6 | 0.0 |
| W2PWB8 | D0N021 | 0.0 |
| W2PWB7 | D0N036 | 0.0 |
| W2PVX5 | D0N081 | 0.0 |
| W2PYI8 | D0N096 | 0.0 |
| W2PVU6 | D0N0A1 | 0.0 |
| W2PYE3 | D0N0B6 | 0.0 |
| W2PL97 | D0N0L9 | 0.0 |
| W2QR51 | D0N0P5 | 0.0 |
| W2QN78 | D0N0X7 | 0.0 |
| W2QPM1 | D0N196 | 0.0 |
| W2QPG4 | D0N1B9 | 0.0 |
| W2QA72 | D0N1M0 | 0.0 |
| W2QA57 | D0N1M5 | 0.0 |
| W2Q962 | D0N1Q6 | 0.0 |
| W2R4A9 | D0N1Q9 | 0.0 |
| W2R2G2 | D0N1R2 | 0.0 |
| W2R483 | D0N1T1 | 0.0 |
| W2R256 | D0N1W9 | 0.0 |
| W2R288 | D0N283 | 0.0 |
| W2R1L5 | D0N2G3 | 0.0 |
| W2R3U4 | D0N2K2 | 0.0 |
| W2RDH0 | D0N2R1 | 0.0 |
| W2RCG7 | D0N305 | 0.0 |
| W2RCJ0 | D0N319 | 0.0 |
| W2RCJ5 | D0N324 | 0.0 |
| W2RCM9 | D0N344 | 0.0 |
| W2R0M6 | D0N359 | 0.0 |
| W2RCJ1 | D0N3A7 | 0.0 |
| W2RCJ8 | D0N3B2 | 0.0 |
| W2RER0 | D0N3D4 | 0.0 |
| W2RF96 | D0N3G2 | 0.0 |
| W2R0S0 | D0N3L1 | 0.0 |
| W2QY41 | D0N3P2 | 0.0 |
| W2PJ44 | D0N3V9 | 0.0 |
| W2PQX2 | D0N412 | 0.0 |
| W2PQR0 | D0N427 | 0.0 |
| W2PF54 | D0N447 | 0.0 |
| W2RD98 | D0N460 | 0.0 |
| W2RDB0 | D0N464 | 0.0 |
| W2PEB1 | D0N470 | 0.0 |
| W2PDR0 | D0N471 | 0.0 |
| W2R6E3 | D0N4B7 | 0.0 |
| W2PWP0 | D0N4G5 | 0.0 |
| W2PYI5 | D0N4K6 | 0.0 |
| W2Q066 | D0N4U8 | 0.0 |
| W2PYE4 | D0N4V1 | 0.0 |
| W2PYE9 | D0N4V7 | 0.0 |

| | | |
|---|---|---|
| W2RI21 | D0N4W3 | 0.0 |
| W2PWU8 | D0N4Y5 | 0.0 |
| W2PYE8 | D0N554 | 0.0 |
| W2PYH8 | D0N577 | 0.0 |
| W2Q158 | D0N586 | 0.0 |
| W2PYU6 | D0N5B0 | 0.0 |
| W2QQB1 | D0N5D0 | 0.0 |
| W2QQB4 | D0N5D5 | 0.0 |
| W2QPT4 | D0N5G7 | 0.0 |
| W2RE69 | D0N5N5 | 0.0 |
| W2RGP0 | D0N5V2 | 0.0 |
| W2RGZ4 | D0N5X1 | 0.0 |
| W2PSD0 | D0N666 | 0.0 |
| W2PGP4 | D0N683 | 0.0 |
| W2PID6 | D0N6E8 | 0.0 |
| W2Q7V6 | D0N6F5 | 0.0 |
| W2QV93 | D0N6P9 | 0.0 |
| W2QVT4 | D0N6Q1 | 0.0 |
| W2PN57 | D0N6R6 | 0.0 |
| W2PQP5 | D0N6Y2 | 0.0 |
| W2PPX8 | D0N700 | 0.0 |
| W2PNQ9 | D0N726 | 0.0 |
| W2PPR1 | D0N735 | 0.0 |
| W2R0H5 | D0N745 | 0.0 |
| W2R0C6 | D0N755 | 0.0 |
| W2R2K0 | D0N7H7 | 0.0 |
| W2R2J0 | D0N7I0 | 0.0 |
| W2QZW6 | D0N7M0 | 0.0 |
| W2R2A9 | D0N7P3 | 0.0 |
| W2QZU3 | D0N7P4 | 0.0 |
| W2R1N5 | D0N7Q2 | 0.0 |
| W2QVF4 | D0N7R8 | 0.0 |
| W2QZC4 | D0N7S3 | 0.0 |
| W2QZG3 | D0N7W0 | 0.0 |
| W2Q752 | D0N7X9 | 0.0 |
| W2Q743 | D0N7Z2 | 0.0 |
| W2Q6Y5 | D0N809 | 0.0 |
| W2PHU9 | D0N841 | 0.0 |
| W2RFD5 | D0N8F9 | 0.0 |
| W2QLJ3 | D0N8N6 | 0.0 |
| W2QF83 | D0N8V3 | 0.0 |
| W2QGQ8 | D0N8Y2 | 0.0 |
| W2QGN0 | D0N907 | 0.0 |
| W2QNA3 | D0N949 | 0.0 |
| W2QNB5 | D0N956 | 0.0 |
| W2QKS8 | D0N969 | 0.0 |
| W2QKV3 | D0N977 | 0.0 |
| W2QKW0 | D0N984 | 0.0 |
| W2QKW5 | D0N985 | 0.0 |
| W2QMU7 | D0N988 | 0.0 |
| W2QNL0 | D0N996 | 0.0 |
| W2Q053 | D0N9B2 | 0.0 |
| W2RFF4 | D0N9M5 | 0.0 |
| W2PZ52 | D0N9N3 | 0.0 |
| W2REZ8 | D0N9S9 | 0.0 |
| W2PHY0 | D0N9T9 | 0.0 |
| W2PE44 | D0N9U5 | 0.0 |
| W2PCB0 | D0N9U7 | 0.0 |
| W2PFD0 | D0N9Y6 | 0.0 |
| W2QMY2 | D0NA18 | 0.0 |
| W2QL13 | D0NA23 | 0.0 |
| W2QKC6 | D0NA25 | 0.0 |
| W2PNS3 | D0NAD8 | 0.0 |
| W2PZM7 | D0NAG7 | 0.0 |
| W2Q257 | D0NAK5 | 0.0 |
| W2PZP4 | D0NAM5 | 0.0 |
| W2PZM0 | D0NAN9 | 0.0 |
| W2Q0U7 | D0NAQ9 | 0.0 |
| W2Q337 | D0NAY8 | 0.0 |
| W2R0Z2 | D0NB08 | 0.0 |
| W2QY59 | D0NB21 | 0.0 |

| | | |
|---|---|---|
| W2R7C7 | D0NBA5 | 0.0 |
| W2R5K8 | D0NBB6 | 0.0 |
| W2R7M2 | D0NBD5 | 0.0 |
| W2R673 | D0NBG6 | 0.0 |
| W2R8F7 | D0NBJ9 | 0.0 |
| W2R8G6 | D0NBK3 | 0.0 |
| W2PT91 | D0NBZ5 | 0.0 |
| W2PRG8 | D0NC61 | 0.0 |
| W2RA07 | D0NC82 | 0.0 |
| W2R8E6 | D0NC99 | 0.0 |
| W2R829 | D0NCB2 | 0.0 |
| W2RAJ2 | D0NCN7 | 0.0 |
| W2R955 | D0NCR4 | 0.0 |
| W2PCP7 | D0NCT6 | 0.0 |
| W2RAZ9 | D0NCV1 | 0.0 |
| W2PDI4 | D0NCV7 | 0.0 |
| W2PDK5 | D0NCX3 | 0.0 |
| W2QT19 | D0ND10 | 0.0 |
| W2QVJ6 | D0ND38 | 0.0 |
| W2QV75 | D0ND90 | 0.0 |
| W2QSE4 | D0NDE7 | 0.0 |
| W2QRY7 | D0NDP1 | 0.0 |
| W2QUC6 | D0NDP8 | 0.0 |
| W2PDK7 | D0NDY1 | 0.0 |
| W2Q357 | D0NDY8 | 0.0 |
| W2PU97 | D0NE02 | 0.0 |
| W2QAN3 | D0NE39 | 0.0 |
| W2Q847 | D0NE45 | 0.0 |
| W2Q908 | D0NE48 | 0.0 |
| W2QAQ8 | D0NE49 | 0.0 |
| W2QB77 | D0NEH7 | 0.0 |
| W2R9G7 | D0NEK2 | 0.0 |
| W2RAY1 | D0NEL2 | 0.0 |
| W2QDQ1 | D0NEM6 | 0.0 |
| W2PHP4 | D0NEQ1 | 0.0 |
| W2PHP9 | D0NEQ3 | 0.0 |
| W2PGE9 | D0NES4 | 0.0 |
| W2RHC0 | D0NF22 | 0.0 |
| W2RFD6 | D0NF39 | 0.0 |
| W2QII2 | D0NFC4 | 0.0 |
| W2QJG0 | D0NFD3 | 0.0 |
| W2QI75 | D0NFM9 | 0.0 |
| W2RAR2 | D0NFS5 | 0.0 |
| W2RD85 | D0NFT1 | 0.0 |
| W2RD56 | D0NFV0 | 0.0 |
| W2RCS4 | D0NFV2 | 0.0 |
| W2RB59 | D0NFV4 | 0.0 |
| W2RCS0 | D0NFV6 | 0.0 |
| W2RB05 | D0NFY8 | 0.0 |
| W2RCG4 | D0NG20 | 0.0 |
| W2RAG6 | D0NG21 | 0.0 |
| W2RAT4 | D0NG36 | 0.0 |
| W2RAA2 | D0NG40 | 0.0 |
| W2RAA1 | D0NG72 | 0.0 |
| W2RCP5 | D0NG73 | 0.0 |
| W2RCN5 | D0NG78 | 0.0 |
| W2RC03 | D0NGA5 | 0.0 |
| W2R9P4 | D0NGE9 | 0.0 |
| W2RC33 | D0NGG3 | 0.0 |
| W2Q3V4 | D0NH56 | 0.0 |
| W2Q3Y1 | D0NH63 | 0.0 |
| W2PJX5 | D0NHB2 | 0.0 |
| W2PME4 | D0NHD9 | 0.0 |
| W2R3J5 | D0NHE0 | 0.0 |
| W2R438 | D0NHI7 | 0.0 |
| W2R402 | D0NHQ0 | 0.0 |
| W2R6M4 | D0NHS1 | 0.0 |
| W2R4L7 | D0NHT5 | 0.0 |
| W2RBI2 | D0NI42 | 0.0 |
| W2QMW9 | D0NI69 | 0.0 |
| W2PFP4 | D0NIB9 | 0.0 |

| | | |
|---|---|---|
| W2QM28 | D0NIC1 | 0.0 |
| W2QMI4 | D0NID9 | 0.0 |
| W2QLN1 | D0NIH8 | 0.0 |
| W2Q7B0 | D0NIR3 | 0.0 |
| W2Q979 | D0NIV4 | 0.0 |
| W2QYQ7 | D0NJ50 | 0.0 |
| W2QZ71 | D0NJ60 | 0.0 |
| W2QYV6 | D0NJ65 | 0.0 |
| W2R0R7 | D0NJ67 | 0.0 |
| W2QYM4 | D0NJ70 | 0.0 |
| W2R0U8 | D0NJ92 | 0.0 |
| W2QZ62 | D0NJF3 | 0.0 |
| W2QYX2 | D0NJG4 | 0.0 |
| W2R161 | D0NJI7 | 0.0 |
| W2QZP3 | D0NJJ2 | 0.0 |
| W2R1U5 | D0NJL6 | 0.0 |
| W2RFW0 | D0NJP0 | 0.0 |
| W2Q7B9 | D0NJS3 | 0.0 |
| W2PHH5 | D0NJX9 | 0.0 |
| W2PGV0 | D0NJY9 | 0.0 |
| W2PMI6 | D0NK33 | 0.0 |
| W2PL77 | D0NKM5 | 0.0 |
| W2QHU2 | D0NKQ0 | 0.0 |
| W2QG34 | D0NKS2 | 0.0 |
| W2PLC5 | D0NKT0 | 0.0 |
| W2PDB0 | D0NL09 | 0.0 |
| W2QRW3 | D0NLB9 | 0.0 |
| W2QPI3 | D0NLD7 | 0.0 |
| W2PM91 | D0NLJ6 | 0.0 |
| W2PML7 | D0NLJ9 | 0.0 |
| W2PQ87 | D0NLM4 | 0.0 |
| W2PNG4 | D0NLM9 | 0.0 |
| W2PPK9 | D0NLP7 | 0.0 |
| W2RI00 | D0NLT6 | 0.0 |
| W2RFL0 | D0NLW5 | 0.0 |
| W2PHJ4 | D0NLX8 | 0.0 |
| W2QBJ2 | D0NM53 | 0.0 |
| W2QC67 | D0NMB1 | 0.0 |
| W2QB70 | D0NMD0 | 0.0 |
| W2QB49 | D0NMD7 | 0.0 |
| W2Q664 | D0NMH7 | 0.0 |
| W2QGY0 | D0NMP3 | 0.0 |
| W2QHU1 | D0NMQ5 | 0.0 |
| W2QH71 | D0NMW4 | 0.0 |
| W2R8P0 | D0NN25 | 0.0 |
| W2PER3 | D0NN38 | 0.0 |
| W2R6Q0 | D0NN88 | 0.0 |
| W2R8A8 | D0NN99 | 0.0 |
| W2R6B5 | D0NNB0 | 0.0 |
| W2R8C8 | D0NNB3 | 0.0 |
| W2R6D6 | D0NNC7 | 0.0 |
| W2QC07 | D0NNN5 | 0.0 |
| W2QC73 | D0NNQ8 | 0.0 |
| W2QCC9 | D0NNT0 | 0.0 |
| W2QCI2 | D0NNX5 | 0.0 |
| W2PC03 | D0NNZ6 | 0.0 |
| W2QF31 | D0NP07 | 0.0 |
| W2QF40 | D0NP13 | 0.0 |
| W2PQC6 | D0NP72 | 0.0 |
| W2QWJ0 | D0NPN2 | 0.0 |
| W2PM22 | D0NPW4 | 0.0 |
| W2PKB9 | D0NPW9 | 0.0 |
| W2PM38 | D0NPX5 | 0.0 |
| W2PMT4 | D0NPZ7 | 0.0 |
| W2QSH4 | D0NQ89 | 0.0 |
| W2PFT4 | D0NQF6 | 0.0 |
| W2Q0Y9 | D0NQJ6 | 0.0 |
| W2PZ75 | D0NQK5 | 0.0 |
| W2Q0S7 | D0NQN0 | 0.0 |
| W2Q1C0 | D0NQR5 | 0.0 |
| W2QL97 | D0NQV6 | 0.0 |

| | | |
|---|---|---|
| W2QNX4 | D0NQV7 | 0.0 |
| W2QN32 | D0NQY4 | 0.0 |
| W2PCJ2 | D0NR25 | 0.0 |
| W2Q5X9 | D0NR26 | 0.0 |
| W2QCD3 | D0NR57 | 0.0 |
| W2QC95 | D0NRA1 | 0.0 |
| W2Q9L5 | D0NRC1 | 0.0 |
| W2PV05 | D0NRI6 | 0.0 |
| W2Q1R2 | D0NRV5 | 0.0 |
| W2PZY1 | D0NRV9 | 0.0 |
| W2Q2G9 | D0NS69 | 0.0 |
| W2QS63 | D0NS79 | 0.0 |
| W2PQ53 | D0NSH0 | 0.0 |
| W2PHH8 | D0NSI1 | 0.0 |
| W2PP26 | D0NSJ0 | 0.0 |
| W2PGE0 | D0NSJ5 | 0.0 |
| W2QHN8 | D0NSL9 | 0.0 |
| W2QIE5 | D0NSM6 | 0.0 |
| W2QFQ0 | D0NSP0 | 0.0 |
| W2QRS6 | D0NT24 | 0.0 |
| W2QTY6 | D0NT67 | 0.0 |
| W2QRU6 | D0NT91 | 0.0 |
| W2QTR9 | D0NTD6 | 0.0 |
| W2QUG7 | D0NTR1 | 0.0 |
| W2PTN4 | D0NU10 | 0.0 |
| W2PUQ9 | D0NU40 | 0.0 |
| W2PUR2 | D0NU46 | 0.0 |
| W2PSK8 | D0NUA2 | 0.0 |
| W2PP25 | D0NUG8 | 0.0 |
| W2PPS8 | D0NUH2 | 0.0 |
| W2PQZ2 | D0NUJ1 | 0.0 |
| W2PJ95 | D0NUK6 | 0.0 |
| W2PIN3 | D0NUL2 | 0.0 |
| W2PI34 | D0NUM9 | 0.0 |
| W2QGS2 | D0NUX3 | 0.0 |
| W2R6W3 | D0NUY3 | 0.0 |
| W2R8J3 | D0NV29 | 0.0 |
| W2R6Z9 | D0NV36 | 0.0 |
| W2PDR1 | D0NVB4 | 0.0 |
| W2QLS7 | D0NVC3 | 0.0 |
| W2QQG1 | D0NVH0 | 0.0 |
| W2QS77 | sp\|D0NT | 0.0 |
| W2QB69 | D0NVK9 | 0.0 |
| W2QB22 | D0NVM9 | 0.0 |
| W2QB00 | D0NVQ0 | 0.0 |
| W2RCX3 | D0NW28 | 0.0 |
| W2RGK8 | D0NW33 | 0.0 |
| W2RDX2 | D0NW98 | 0.0 |
| W2Q6X8 | D0NWC2 | 0.0 |
| W2QJE7 | D0NWF9 | 0.0 |
| W2QGU1 | D0NWG4 | 0.0 |
| W2R4J2 | D0NWL1 | 0.0 |
| W2PP13 | D0NWM7 | 0.0 |
| W2PLY2 | D0NWZ2 | 0.0 |
| W2PM15 | D0NX08 | 0.0 |
| W2PNV8 | D0NX32 | 0.0 |
| W2PLE5 | D0NXA0 | 0.0 |
| W2PKS8 | D0NXA2 | 0.0 |
| W2RB13 | D0NXG3 | 0.0 |
| W2RD39 | D0NXG9 | 0.0 |
| W2PRS9 | D0NXN9 | 0.0 |
| W2PDA0 | D0NXQ0 | 0.0 |
| W2RG48 | D0NXV5 | 0.0 |
| W2RDL1 | D0NXV6 | 0.0 |
| W2RDP2 | D0NXW8 | 0.0 |
| W2RDR6 | D0NXY8 | 0.0 |
| W2RGD6 | D0NY08 | 0.0 |
| W2Q329 | D0NY26 | 0.0 |
| W2Q5J9 | D0NY29 | 0.0 |
| W2Q4S5 | D0NY30 | 0.0 |
| W2PCP1 | D0NY60 | 0.0 |

| | | |
|---|---|---|
| W2QWH9 | D0NYB0 | 0.0 |
| W2PTR5 | D0NYG9 | 0.0 |
| W2Q6M6 | D0NYL6 | 0.0 |
| W2R1D8 | D0NYS4 | 0.0 |
| W2PIM1 | D0NYW9 | 0.0 |
| W2PJV6 | D0NYZ7 | 0.0 |
| W2QH02 | D0NZ18 | 0.0 |
| W2QG60 | D0NZ40 | 0.0 |
| W2QCS5 | D0NZL3 | 0.0 |
| W2QT21 | D0NZU1 | 0.0 |
| W2PME9 | D0NZV4 | 0.0 |
| W2Q6C7 | D0P055 | 0.0 |
| W2R9F0 | D0P0B8 | 0.0 |
| W2PSD7 | D0P0H8 | 0.0 |
| W2PK79 | D0P0X0 | 0.0 |
| W2QQ19 | D0P181 | 0.0 |
| W2Q9S3 | D0P1T1 | 0.0 |
| W2QHJ6 | D0P1T4 | 0.0 |
| W2PRG2 | D0P1T9 | 0.0 |
| W2RID3 | D0P1V5 | 0.0 |
| W2PP80 | D0P1Y9 | 0.0 |
| W2Q3Q5 | D0P270 | 0.0 |
| W2PFJ4 | D0P2F7 | 0.0 |
| W2PHZ4 | D0P2F9 | 0.0 |
| W2PCA1 | D0P2S2 | 0.0 |
| W2R2A0 | D0P2T1 | 0.0 |
| W2QT07 | D0P3H8 | 0.0 |
| W2PNJ7 | D0P3U0 | 0.0 |
| W2QID6 | D0P3V4 | 0.0 |
| W2QK47 | D0P3V7 | 0.0 |
| W2PQW1 | D0P4H2 | 0.0 |
| W2PCX2 | D0RLR0 | 0.0 |
