## Supplementary table 4 for "Zoospore-derived extracellular vesicles in the flagellated stramenopile *Phytophthora parasitica*"

Title : List of *P. parasitica* proteins exhibiting or not significant similarity (BlastP e-Value < E-6) with known EV markers in other species

| EV MARKERS | | | FEATURES OF PROTEIN SEQUENCES IDENTIFIED IN THE UNIPROT *P. PARASITICA* PROTEOME (ID UP000018817) | | | | | TRANSCRIPTOMIC & PROTEOMIC ZOOSPORE DATA (Bassani et al. 2020b) (Lupatelli et al., 2025) | | |
|---|---|---|---|---|---|---|---|---|---|---|
| Reference | EV marker | Query sequence submitted to UniProt BLASTP search | UniProt Id | Gene name | Protein name | Subcellular location | BlastP e-Value < E-6 | mRNA expression measurement (FPKM) | Protein expression measurement (Sum of peak intensities) | Protein relative abundance in Flagella versus Cell body (FDR <0,01) |
| **Transmembrane (or GPI-anchored) proteins associated with plasma membrane and/or endosomes** | | | ***P. parasitica* candidates** | | | | | | | |
| Welsh et al., 2024 | CD63 | P08962 · CD63_HUMAN | W2PLH7 | *PPTG_16927* | Tetraspannin | Membrane | | 126,5 | 2,40E+03 | |
| | CD9 | P21926 · CD9_HUMAN | W2PLH7 | *PPTG_16928* | Tetraspannin | Membrane | | 10,6 | 1,83E+04 | |
| | CD81 | P60033 · CD81_HUMAN | W2QZG1 | *PPTG_03988* | Tetraspannin | Membrane | | 190,2 | 3,59E+03 | |
| | CD47 | Q08722 · CD47_HUMAN | W2QYJ7 | *PPTG_03767* | Hexose transporter 1 | Membrane | | 4,7 | 9,62E+03 | |
| | TSAP6 | Q658P3 · STEA3_HUMAN | W2QNZ5 | *PPTG_07685* | Uncharacterized protein | Membrane | | 0,8 | 6,68E+03 | |
| | MHCI, II | P30511 · HLAF_HUMAN | W2QWP5 | *PPTG_05391* | Uncharacterized protein | Cytoplasm | | 0,5 | 1,29E+04 | |
| | ITGA | P13612 · ITA4_HUMAN | W2QFN1 | *PPTG_10213* | Calx-beta domain-containing protein | Secreted | | 4,2 | 4,88E+02 | |
| | ITGB | P05556 · ITB1_HUMAN | W2RER0 | *PPTG_02823* | EGF-like domain-containing protein | Secreted (GPI site380) | 1,20E-10 | 427,0 | 1,44E+05 | |
| | TRF2 | Q9UP52 · TFR2_HUMAN | W2QRQ0 | *PPTG_07176* | TFR Glutamate carboxypeptidase | Membrane | 1,40E-56 | 5,8 | *UR* | |
| | LAMP1/2 | P11279 · LAMP1_HUMAN | W2PF19 | *PPTG_19330* | Uncharacterized protein | Secreted | | 0,0 | 3,37E+03 | |
| | SDC | P18828 · SDC1_MOUSE | W2PXY9 | *PPTG_14637* | TKL protein kinase | Membrane | | 0,0 | *UR* | |
| | EMMPRIN (BSG) | Q54A51 · Q54A51_HUMAN | W2QS46 | *PPTG_06204* | RxLR effector protein | Secreted | | 0,0 | *UR* | |
| | ADAM10 | O14672 · ADA10_HUMAN | W2PG38 | *PPTG_18685* | VWFA domain-containing protein | Cytosol/Nucleus | | 0,0 | *UR* | |
| | GPCI | P35052 · GPC1_HUMAN | W2Q363 | *PPTG_12760* | Uncharacterized protein | Secreted | | 0,9 | 3,89E+05 | |
| | CD73 | P21589 · 5NTD_HUMAN | W2QEP4 | *PPTG_09834* | 5'-nucleotidase | Secreted | 2,30E-34 | 12,3 | *UR* | |
| | CD59 | P13987 · CD59_HUMAN | W2RCH7 | *PPTG_02678* | Ubiquitin carboxyl-terminal hydrolase | Cytosol/Nucleus | | 68,6 | 1,34E+03 | |
| Zhao et al., 2019 | FKS1 | P38631 · FKS1_YEAST | W2PVT8 | *PPTG_14740* | 1,3-beta-glucan synthase | Membrane | 3,00E-87 | 80,2 | 2,37E+04 | 0,21 |
| | CHS3 | P29465 · CHS3_YEAST | W2PDG1 | *PPTG_19378* | Inactive glycoside hydrolase | Secreted | | 0,5 | *UR* | |
| Breen et al., 2025 | PiMDP1 | D0NMH7_PHYIT | W2Q664 | *PPTG_13069* | MARVEL domain-containing protein | Membrane | 0 | 0,0 | 2,49E+07 | 64 |
| | PiMDP2 | D0NMH8_PHYIT PITG_13661 | W2QPB2 | *PPTG_07253* | MARVEL domain-containing protein | Membrane | 0 | 347,3 | 1,45E+06 | 3,26 |
| **Cytosolic proteins** | | | ***P. parasitica* candidates** | | | | | | | |
| Welsh et al., 2024 | ESCRT-I | F5H442_HUMAN | W2QVL7 | *PPTG_06448* | UEV domain-containing protein | Endosome | 6,30E-49 | 7,2 | UR | |
| | ESCRT-II | Q86VN1 · VPS36_HUMAN | W2Q1R3 | *PPTG_13489* | ESCRT-II complex subunit VPS36 | Endosome/cytosol | 9,60E-64 | 13,0 | 8,07E+04 | |
| | ESCRT-III | O43633 · CHM2A_HUMAN | W2PE75 | *PPTG_19536* | Charged multivesicular body protein 2a | Cytoplasm | 1,00E-65 | 118,1 | 1,71E+04 | |
| | ALIX (PDCD6IP) | Q8WUM4 · PDC6I_HUMAN | W2PU36 | *PPTG_15104* | BRO1 domain-containing protein | Endosome/cytosol | 1,40E-71 | 0,3 | 4,30E+04 | |
| | VPS4A/B | Q9UN37 · VPS4A_HUMAN | W2RID8 | *PPTG_00705* | Vesicle-fusing ATPase | Endosome mb | 4,30E-155 | 14,2 | 1,85E+04 | |
| | ARRDC1 | Q8N5I2 · ARRD1_HUMAN | W2QL75 | *PPTG_08604* | AAA+ ATPase domain-containing protein | Secreted | 3,00E-31 | 8,8 | *UR* | |
| | FLOT1 | O75955 · FLOT1_HUMAN | W2PPI7 | *PPTG_23955* | Cilia- and flagella-associated protein 45 | Cytoplasm | 1,30E-07 | 0 | *UR* | |
| | caveolins (CAV*) | Q03135 · CAV1_HUMAN | W2QDI2 | *PPTG_09961* | tRNA N(3)-methylcytidine methyltransferase | Cytoplasm | | 11,8 | *UR* | |
| | syntenin (SDCBP) | O00560 · SDCB1_HUMAN | W2R406 | *PPTG_03092* | PDZ domain-containing protein | Cytosol/Nucleus | | 157,8 | *UR* | |
| | HSC70 (HSPA8) | P11142 · HSP7C_HUMAN | W2R9I2 | *PPTG_02122* | Hsp70-like protein | Cytoplasm | 0 | 560,0 | 6,30E+06 | |
| | HSP84 | P11499 · HS90B_MOUSE | W2PE41 | *PPTG_19188* | Heat shock protein 90-2 | Cytoplasm | 0 | 154,6 | 7,44E+06 | |
| | GAPDH | O14556 · G3PT_HUMAN | W2RHS3 | *PPTG_01098* | Glyceraldehyde-3-phosphate dehydrogenase | Cytoplasm | 1,4E-151 | 2,1 | 1,52E+05 | |
| **Ciliary proteins** | | | ***P. parasitica* candidates** | | | | | | | |
| Vinay and Belleannée, 2022 | Prominin-1 | A0A329SL01_9STRA | W2PNS1 | *PPTG_16242* | Uncharacterized protein - Prominin-1 (mammals) | Membrane | 7,00E-07 | 3,10 | 1,53E+07 | 11,4 |
| | PC1 | PKD1_HUMAN | W2R048 | *PPTG_04276* | Polycystin domain-containing protein | Membrane | 0 | 0,5 | 1,35E+06 | 6,3 |
| | PC2 | PKD2_HUMAN | W2QKI5 | *PPTG_08411* | PKD2 domain-containing protein | Membrane | 2,70E-38 | 8,2 | 0,00E+00 | |
| | Arl3b | Q1MTE5 · ARL3_DANRE | W2QU59 | *PPTG_06270* | ADP-ribosylation factor-like protein 3 | Golgi apparatus | 1,5E-91 | 2,3 | 4,78E+06 | 0,1 |
| | POR | P37040 · NCPR_MOUSE | W2Q8L5 | *PPTG_11883* | NADPH--cytochrome P450 reductase | Endoplasmic reticulum mb | 3,80E-147 | 9,0 | 1,45E+05 | |
| | Annexins | ANX13_HUMAN | W2Q4N9 | *PPTG_17884* | Annexin | Membrane | 2,90E-99 | 85,35 | 2,58E+06 | 21,0 |
| | Nup155 | NU155_HUMAN | W2R0A8 | *PPTG_04333* | nuclear pore protein Nup155 (1519 aa) | Nucleus | 2,70E-72 | 37,0 | 8,92E+04 | |
| | Septin-10 | SEP10_HUMAN | W2PPI7 | *PPTG_09144* | Cilia- and flagella-associated protein 45 | Flagella | | 26,8 | 9,45E+03 | |
| | CD151 | CD151_HUMAN | W2RHM8 | *PPTG_00602* | Tetraspannin | Membrane | | 8,5 | 5,09E+04 | |
| | Keratin-19 | K1C19_HUMAN | W2QC84 | *PPTG_11449* | LTD domain-containing protein | Nucleus | 2,8E-13 | 14,4 | 1,45E+04 | |
