## Supplementary table 5 for "Zoospore-derived extracellular vesicles in the flagellated stramenopile *Phytophthora parasitica*"

**Title: GO terms analysis with g:Profiler for enrichment of TOP400 proteins in molecular functions, cellular components, and biological processes**

| source | term_name | term_id | highlighted | adjusted_p_value | negative_log10_of_adjusted_p_value | term_size | query_size | intersection_size | effective_domain_size | intersections |
|---|---|---|---|---|---|---|---|---|---|---|
| GO:MF | small molecule binding | GO:0036094 | VRAI | 1.8028278776315696e-248 | 247.74404573493212 | 3526 | 7905 | 3523 | 9001 | F443_00056,F443_03595,F443_03607,[illegible] |
| GO:MF | catalytic activity | GO:0003824 | VRAI | 1.247575731570904e-182 | 181.90393308193296 | 4908 | 7905 | 4744 | 9001 | F443_00158,F443_03595,F443_03607,[illegible] |
| GO:MF | nucleic acid binding | GO:0003676 | VRAI | 8.681384954270599e-93 | 92.06141098565098 | 1510 | 7905 | 1510 | 9001 | F443_02217,F443_04303,F443_05119,[illegible] |
| GO:MF | structural molecule activity | GO:0005198 | VRAI | 3.632783061572907e-9 | 8.43976053606926 | 179 | 7905 | 179 | 9001 | F443_00048,F443_00056,F443_00158,[illegible] |
| GO:MF | phospholipid binding | GO:0005543 | VRAI | 3.7349491683782796e-7 | 6.4277153044357425 | 144 | 7905 | 144 | 9001 | F443_02069,F443_03196,F443_04004,[illegible] |
| GO:MF | cytoskeletal motor activity | GO:0003774 | VRAI | 0.00003301571439840411 | 4.481279300967738 | 110 | 7905 | 110 | 9001 | F443_12488,F443_17918,F443_00424,[illegible] |
| GO:MF | protein dimerization activity | GO:0046983 | VRAI | 0.0000945256225993776 | 4.0244504534461605 | 102 | 7905 | 102 | 9001 | F443_05119,F443_06642,F443_06745,[illegible] |
| GO:MF | cytoskeletal protein binding | GO:0008092 | VRAI | 0.00009520041350608968 | 4.021361165239234 | 143 | 7905 | 141 | 9001 | F443_01444,F443_03233,F443_12488,[illegible] |
| GO:BP | transmembrane transport | GO:0055085 | VRAI | 2.9924887440451655e-9 | 8.523967474474857 | 749 | 5412 | 749 | 5567 | F443_00104,F443_00135,F443_00307,[illegible] |
| GO:BP | response to stimulus | GO:0050896 | VRAI | 7.04260335225796e-7 | 6.152266770897252 | 579 | 5412 | 579 | 5567 | F443_03584,F443_11394,F443_21452,[illegible] |
| GO:BP | proteolysis | GO:0006508 | VRAI | 0.00007242453099489981 | 4.14011430850924 | 430 | 5412 | 430 | 5567 | F443_00635,F443_00778,F443_00887,[illegible] |
| GO:CC | membrane | GO:0016020 | VRAI | 3.591045080466643e-17 | 16.444779142889796 | 3635 | 5888 | 3635 | 5929 | F443_02217,F443_06470,F443_07033,[illegible] |
| GO:MF | ion binding | GO:0043167 | FAUX | 2.7842181806181446e-232 | 231.55529673496457 | 3355 | 7905 | 3352 | 9001 | F443_03595,F443_03607,F443_03609,[illegible] |
| GO:MF | hydrolase activity | GO:0016787 | FAUX | 2.195478245416048e-124 | 123.6584708618812 | 1955 | 7905 | 1955 | 9001 | F443_00158,F443_03595,F443_03607,[illegible] |
| GO:MF | cation binding | GO:0043169 | FAUX | 4.6899478222814924e-119 | 118.32883198897389 | 1882 | 7905 | 1882 | 9001 | F443_04109,F443_05515,F443_21596,[illegible] |
| GO:MF | metal ion binding | GO:0046872 | FAUX | 1.784915903234062e-118 | 117.74838224096455 | 1874 | 7905 | 1874 | 9001 | F443_04109,F443_05515,F443_21596,[illegible] |
| GO:MF | nucleotide binding | GO:0000166 | FAUX | 8.742887436369071e-114 | 113.05834511306517 | 1809 | 7905 | 1809 | 9001 | F443_03595,F443_03607,F443_03609,[illegible] |
| GO:MF | heterocyclic compound binding | GO:1901363 | FAUX | 1.896351528788384e-113 | 112.72208115388501 | 1899 | 7905 | 1896 | 9001 | F443_03595,F443_03607,F443_03609,[illegible] |
| GO:MF | nucleoside phosphate binding | GO:1901265 | FAUX | 1.6049223512892454e-107 | 106.7945459746126 | 1816 | 7905 | 1813 | 9001 | F443_03595,F443_03607,F443_03609,[illegible] |
| GO:MF | transferase activity | GO:0016740 | FAUX | 6.454305597518219e-100 | 99.19015047553363 | 1613 | 7905 | 1613 | 9001 | F443_00767,F443_01750,F443_02151,[illegible] |
| GO:MF | anion binding | GO:0043168 | FAUX | 1.2068309314262313e-99 | 98.91835356726106 | 1704 | 7905 | 1701 | 9001 | F443_03595,F443_03607,F443_03609,[illegible] |
| GO:MF | purine nucleotide binding | GO:0017076 | FAUX | 7.383054016553193e-97 | 96.13176395414219 | 1569 | 7905 | 1569 | 9001 | F443_03595,F443_03607,F443_03609,[illegible] |
| GO:MF | carbohydrate derivative binding | GO:0097367 | FAUX | 2.0941467515122457e-95 | 94.67899288753836 | 1548 | 7905 | 1548 | 9001 | F443_03595,F443_03607,F443_03609,[illegible] |
| GO:MF | ribonucleotide binding | GO:0032553 | FAUX | 2.2697969128610902e-94 | 93.64401299900486 | 1533 | 7905 | 1533 | 9001 | F443_03595,F443_03607,F443_03609,[illegible] |
| GO:MF | purine ribonucleotide binding | GO:0032555 | FAUX | 6.162016171211292e-91 | 90.21027716628221 | 1483 | 7905 | 1483 | 9001 | F443_03595,F443_03607,F443_03609,[illegible] |
| GO:MF | purine ribonucleoside triphosphate binding | GO:0035639 | FAUX | 1.8555434509116097e-90 | 89.73152887140401 | 1476 | 7905 | 1476 | 9001 | F443_03595,F443_03607,F443_03609,[illegible] |
| GO:MF | adenyl nucleotide binding | GO:0030554 | FAUX | 2.4054991909801455e-84 | 83.61879478482373 | 1386 | 7905 | 1386 | 9001 | F443_00460,F443_00508,F443_00730,[illegible] |
| GO:MF | adenyl ribonucleotide binding | GO:0032559 | FAUX | 1.4114797490689874e-78 | 77.85032534842793 | 1300 | 7905 | 1300 | 9001 | F443_00508,F443_00730,F443_01322,[illegible] |
| GO:MF | ATP binding | GO:0005524 | FAUX | 2.2368735147045357e-78 | 77.65035857263959 | 1297 | 7905 | 1297 | 9001 | F443_00508,F443_00730,F443_01322,[illegible] |
| GO:MF | catalytic activity, acting on a protein | GO:0140096 | FAUX | 7.001336703285349e-70 | 69.15481903606572 | 1338 | 7905 | 1332 | 9001 | F443_03554,F443_00085,F443_00086,[illegible] |
| GO:MF | transition metal ion binding | GO:0046914 | FAUX | 4.15746062919133e-47 | 46.38117185482183 | 812 | 7905 | 812 | 9001 | F443_04109,F443_21242,F443_00071,[illegible] |
| GO:MF | transmembrane transporter activity | GO:0022857 | FAUX | 2.6601067484382623e-45 | 44.57510093505034 | 783 | 7905 | 783 | 9001 | F443_00508,F443_00730,F443_01322,[illegible] |
| GO:MF | transferase activity, transferring phosphorus-containing groups | GO:0016772 | FAUX | 9.638532407532475e-45 | 44.0159890880714 | 774 | 7905 | 774 | 9001 | F443_00767,F443_01750,F443_02151,[illegible] |
| GO:MF | transporter activity | GO:0005215 | FAUX | 9.01768827616406e-44 | 43.04490478148029 | 792 | 7905 | 791 | 9001 | F443_00508,F443_00730,F443_01322,[illegible] |
| GO:MF | oxidoreductase activity | GO:0016491 | FAUX | 1.3357888323596353e-40 | 39.87426219170289 | 707 | 7905 | 707 | 9001 | F443_17902,F443_00460,F443_02816,[illegible] |
| GO:MF | zinc ion binding | GO:0008270 | FAUX | 3.0051954392192494e-39 | 38.52212727893098 | 685 | 7905 | 685 | 9001 | F443_04109,F443_21242,F443_00071,[illegible] |
| GO:MF | kinase activity | GO:0016301 | FAUX | 6.539144078518115e-35 | 34.184479093618116 | 614 | 7905 | 614 | 9001 | F443_01750,F443_02151,F443_03554,[illegible] |
| GO:MF | DNA binding | GO:0003677 | FAUX | 7.52198855329265e-35 | 34.123667331807745 | 613 | 7905 | 613 | 9001 | F443_05119,F443_06454,F443_06642,[illegible] |
| GO:MF | phosphotransferase activity, alcohol group as acceptor | GO:0016773 | FAUX | 1.518560416005515e-32 | 31.818567924973337 | 575 | 7905 | 575 | 9001 | F443_01750,F443_02151,F443_03554,[illegible] |
| GO:MF | hydrolase activity, acting on acid anhydrides | GO:0016817 | FAUX | 9.848772898834965e-31 | 30.00661787675751 | 545 | 7905 | 545 | 9001 | F443_03595,F443_03607,F443_03609,[illegible] |
| GO:MF | hydrolase activity, acting on acid anhydrides, in phosphorus-containing anhydrides | GO:0016818 | FAUX | 1.2999705202087603e-30 | 29.886066496197646 | 543 | 7905 | 543 | 9001 | F443_03595,F443_03607,F443_03609,[illegible] |
| GO:MF | pyrophosphatase activity | GO:0016462 | FAUX | 4.529357504655997e-30 | 29.343963398864023 | 534 | 7905 | 534 | 9001 | F443_03595,F443_03607,F443_03609,[illegible] |
| GO:MF | ribonucleoside triphosphate phosphatase activity | GO:0017111 | FAUX | 1.0929699291585785e-28 | 27.96139178661402 | 511 | 7905 | 511 | 9001 | F443_03595,F443_03607,F443_03609,[illegible] |
| GO:MF | protein kinase activity | GO:0004672 | FAUX | 7.865618174816779e-27 | 26.10426713961899 | 480 | 7905 | 480 | 9001 | F443_03554,F443_00085,F443_00086,[illegible] |
| GO:MF | hydrolase activity, acting on ester bonds | GO:0016788 | FAUX | 1.6192109015525338e-25 | 24.79069658088515 | 458 | 7905 | 458 | 9001 | F443_03548,F443_03554,F443_03555,[illegible] |
| GO:MF | peptidase activity | GO:0008233 | FAUX | 8.399802665174286e-25 | 24.075730916608663 | 446 | 7905 | 446 | 9001 | F443_00635,F443_00662,F443_00778,[illegible] |
| GO:MF | ATP-dependent activity | GO:0140657 | FAUX | 1.6797959844182155e-24 | 23.774743461263242 | 627 | 7905 | 619 | 9001 | F443_00508,F443_00730,F443_01322,[illegible] |
| GO:MF | catalytic activity, acting on a nucleic acid | GO:0140640 | FAUX | 1.4889380697129012e-23 | 22.827123365740967 | 425 | 7905 | 425 | 9001 | F443_04687,F443_07870,F443_07886,[illegible] |
| GO:MF | protein serine/threonine kinase activity | GO:0004674 | FAUX | 7.794152203758756e-22 | 21.1082311175827 | 396 | 7905 | 396 | 9001 | F443_03554,F443_00085,F443_00086,[illegible] |
| GO:MF | RNA binding | GO:0003723 | FAUX | 1.5617943371873872e-19 | 18.80637615618552 | 357 | 7905 | 357 | 9001 | F443_04303,F443_06182,F443_07644,[illegible] |
| GO:MF | ATP hydrolysis activity | GO:0016887 | FAUX | 2.682142218678362e-18 | 17.57151819771869 | 336 | 7905 | 336 | 9001 | F443_00508,F443_00730,F443_01322,[illegible] |
| GO:MF | acyltransferase activity | GO:0016746 | FAUX | 1.1858614204068838e-17 | 16.925966059596565 | 325 | 7905 | 325 | 9001 | F443_06332,F443_06347,F443_00210,[illegible] |
| GO:MF | active transmembrane transporter activity | GO:0022804 | FAUX | 3.995411670680822e-17 | 16.39843846613872 | 316 | 7905 | 316 | 9001 | F443_00508,F443_00730,F443_01322,[illegible] |
| GO:MF | isomerase activity | GO:0016853 | FAUX | 3.0163037988633305e-16 | 15.520524918927796 | 301 | 7905 | 301 | 9001 | F443_04899,F443_18309,F443_01119,[illegible] |
| GO:MF | monoatomic ion transmembrane transporter activity | GO:0015075 | FAUX | 5.167871840286959e-16 | 15.286688265087726 | 297 | 7905 | 297 | 9001 | F443_00508,F443_00730,F443_01322,[illegible] |
| GO:MF | primary active transmembrane transporter activity | GO:0015399 | FAUX | 3.702649950144889e-13 | 12.431487344366024 | 248 | 7905 | 248 | 9001 | F443_00508,F443_00730,F443_01322,[illegible] |
| GO:MF | phosphoric ester hydrolase activity | GO:0042578 | FAUX | 4.838245672027087e-13 | 12.315312083205047 | 246 | 7905 | 246 | 9001 | F443_03548,F443_03554,F443_03555,[illegible] |
| GO:MF | catalytic activity, acting on RNA | GO:0140098 | FAUX | 5.530506410479005e-13 | 12.257235099941756 | 245 | 7905 | 245 | 9001 | F443_04687,F443_07870,F443_07886,[illegible] |
| GO:MF | monoatomic cation transmembrane transporter activity | GO:0008324 | FAUX | 6.321713175377199e-13 | 12.199165212573755 | 244 | 7905 | 244 | 9001 | F443_00508,F443_00730,F443_01322,[illegible] |
| GO:MF | hydrolase activity, acting on glycosyl bonds | GO:0016798 | FAUX | 7.225993734005943e-13 | 12.141102419364707 | 243 | 7905 | 243 | 9001 | F443_00166,F443_00481,F443_00485,[illegible] |
| GO:MF | hydrolase activity, hydrolyzing O-glycosyl compounds | GO:0004553 | FAUX | 3.141380817209629e-12 | 11.502879412602528 | 232 | 7905 | 232 | 9001 | F443_00166,F443_00481,F443_00485,[illegible] |
| GO:MF | calcium ion binding | GO:0005509 | FAUX | 3.141380817209629e-12 | 11.502879412602528 | 232 | 7905 | 232 | 9001 | F443_01899,F443_00017,F443_00224,[illegible] |
| GO:MF | ATPase-coupled transmembrane transporter activity | GO:0042626 | FAUX | 3.141380817209629e-12 | 11.502879412602528 | 232 | 7905 | 232 | 9001 | F443_00508,F443_00730,F443_01322,[illegible] |
| GO:MF | inorganic molecular entity transmembrane transporter activity | GO:0015318 | FAUX | 3.141380817209629e-12 | 11.502879412602528 | 232 | 7905 | 232 | 9001 | F443_00365,F443_00611,F443_00837,[illegible] |
| GO:MF | transferase activity, transferring one-carbon groups | GO:0016741 | FAUX | 4.688487578069277e-12 | 11.328967230297554 | 229 | 7905 | 229 | 9001 | F443_14475,F443_01142,F443_01853,[illegible] |
| GO:MF | methyltransferase activity | GO:0008168 | FAUX | 1.7794298568151576e-11 | 10.749719126737695 | 219 | 7905 | 219 | 9001 | F443_14475,F443_01142,F443_01853,[illegible] |
| GO:MF | phosphatase activity | GO:0016791 | FAUX | 1.7115333783012e-10 | 9.766614626838301 | 202 | 7905 | 202 | 9001 | F443_03548,F443_03554,F443_03555,[illegible] |
| GO:MF | guanyl nucleotide binding | GO:0019001 | FAUX | 4.958154896991857e-10 | 9.304679909625879 | 194 | 7905 | 194 | 9001 | F443_03595,F443_03607,F443_03609,[illegible] |
| GO:MF | guanyl ribonucleotide binding | GO:0032561 | FAUX | 4.958154896991857e-10 | 9.304679909625879 | 194 | 7905 | 194 | 9001 | F443_03595,F443_03607,F443_03609,[illegible] |
| GO:MF | macromolecular conformation isomerase activity | GO:0120543 | FAUX | 5.662788943556031e-10 | 9.246969624556959 | 193 | 7905 | 193 | 9001 | F443_17918,F443_00424,F443_00539,[illegible] |
| GO:MF | inorganic cation transmembrane transporter activity | GO:0022890 | FAUX | 6.46745855098992e-10 | 9.189266345933266 | 192 | 7905 | 192 | 9001 | F443_00365,F443_00837,F443_00952,[illegible] |
| GO:MF | GTP binding | GO:0005525 | FAUX | 2.1367226441919535e-9 | 8.670251847472969 | 183 | 7905 | 183 | 9001 | F443_03595,F443_03607,F443_03609,[illegible] |
| GO:MF | S-adenosylmethionine-dependent methyltransferase activity | GO:0008757 | FAUX | 3.1814647397089452e-9 | 8.497372885702845 | 180 | 7905 | 180 | 9001 | F443_14475,F443_01142,F443_01853,[illegible] |
| GO:MF | endopeptidase activity | GO:0004175 | FAUX | 3.632783061572907e-9 | 8.43976053606926 | 179 | 7905 | 179 | 9001 | F443_00635,F443_00662,F443_00948,[illegible] |
| GO:MF | GTPase activity | GO:0003924 | FAUX | 4.736343774570724e-9 | 8.324556783062858 | 177 | 7905 | 177 | 9001 | F443_03595,F443_03607,F443_03609,[illegible] |
| GO:MF | acyltransferase activity, transferring groups other than amino-acyl groups | GO:0016747 | FAUX | 6.174745366231928e-9 | 8.209380947108057 | 175 | 7905 | 175 | 9001 | F443_06332,F443_06347,F443_00224,[illegible] |
| GO:MF | cysteine-type peptidase activity | GO:0008234 | FAUX | 1.0492694487340097e-8 | 7.979112972176439 | 171 | 7905 | 171 | 9001 | F443_04142,F443_07066,F443_07768,[illegible] |
| GO:MF | ABC-type transporter activity | GO:0140359 | FAUX | 2.6520868259648276e-8 | 7.576412261783046 | 164 | 7905 | 164 | 9001 | F443_00104,F443_00335,F443_00336,[illegible] |
| GO:MF | glycosyltransferase activity | GO:0016757 | FAUX | 1.1368799053721537e-7 | 6.944285409708803 | 153 | 7905 | 153 | 9001 | F443_10266,F443_00260,F443_02016,[illegible] |
| GO:MF | channel activity | GO:0015267 | FAUX | 2.5127948543511314e-7 | 6.599842965947818 | 147 | 7905 | 147 | 9001 | F443_01128,F443_01407,F443_01677,[illegible] |
| GO:MF | passive transmembrane transporter activity | GO:0022803 | FAUX | 2.5127948543511314e-7 | 6.599842965947818 | 147 | 7905 | 147 | 9001 | F443_01128,F443_01407,F443_01677,[illegible] |
| GO:MF | catalytic activity, acting on DNA | GO:0140097 | FAUX | 9.411820902898575e-7 | 6.026326345586849 | 137 | 7905 | 137 | 9001 | F443_11394,F443_21452,F443_00885,[illegible] |
| GO:MF | metal ion transmembrane transporter activity | GO:0046873 | FAUX | 0.0000013982788238940545 | 5.854406219431067 | 134 | 7905 | 134 | 9001 | F443_00837,F443_00952,F443_01128,[illegible] |
| GO:MF | phosphatidylinositol binding | GO:0035091 | FAUX | 0.0000040155373052297 | 5.396256335046853 | 126 | 7905 | 126 | 9001 | F443_02069,F443_03196,F443_04004,[illegible] |
| GO:MF | monoatomic ion channel activity | GO:0005216 | FAUX | 0.000008852511457545232 | 5.052933502474912 | 120 | 7905 | 120 | 9001 | F443_01128,F443_01407,F443_01677,[illegible] |
| GO:MF | aminoacyltransferase activity | GO:0016755 | FAUX | 0.000008852511457545232 | 5.052933502474912 | 120 | 7905 | 120 | 9001 | F443_00870,F443_00871,F443_00872,[illegible] |
| GO:MF | structural constituent of ribosome | GO:0003735 | FAUX | 0.000011519996692400719 | 4.938547645606585 | 118 | 7905 | 118 | 9001 | F443_00048,F443_00158,F443_00432,[illegible] |
| GO:MF | enzyme regulator activity | GO:0030234 | FAUX | 0.00001314119807592296 | 4.881365038578142 | 117 | 7905 | 117 | 9001 | F443_00752,F443_10655,F443_10956,[illegible] |
| GO:MF | nuclease activity | GO:0004518 | FAUX | 0.000028945843407441226 | 4.538413791688425 | 111 | 7905 | 111 | 9001 | F443_04687,F443_11394,F443_20738,[illegible] |
| GO:MF | nucleotidyltransferase activity | GO:0016779 | FAUX | 0.000042950584730846445 | 4.36703091929138 | 108 | 7905 | 108 | 9001 | F443_01750,F443_18309,F443_21452,[illegible] |
| GO:MF | nucleic acid conformation isomerase activity | GO:0120545 | FAUX | 0.000042950584730846445 | 4.36703091929138 | 108 | 7905 | 108 | 9001 | F443_00539,F443_00805,F443_01498,[illegible] |
| GO:MF | phosphoprotein phosphatase activity | GO:0004721 | FAUX | 0.00008288385584256027 | 4.0815300533011225 | 103 | 7905 | 103 | 9001 | F443_00122,F443_00226,F443_00989,[illegible] |
| GO:MF | oxidoreductase activity, acting on CH-OH group of donors | GO:0016614 | FAUX | 0.00008288385584256027 | 4.0815300533011225 | 103 | 7905 | 103 | 9001 | F443_00460,F443_02816,F443_06332,[illegible] |
| GO:MF | helicase activity | GO:0004386 | FAUX | 0.00008288385584256027 | 4.0815300533011225 | 103 | 7905 | 103 | 9001 | F443_00539,F443_00805,F443_01498,[illegible] |
| GO:BP | establishment of localization | GO:0051234 | FAUX | 0.00001739981645246068 | 4.759455332987741 | 1195 | 5412 | 1184 | 5567 | F443_02217,F443_02234,F443_03209,[illegible] |
| GO:BP | transport | GO:0006810 | FAUX | 0.000029078448296500892 | 4.536428772305687 | 1174 | 5412 | 1163 | 5567 | F443_02217,F443_02234,F443_03209,[illegible] |
| GO:BP | cellular response to stimulus | GO:0051716 | FAUX | 0.000029712851645999937 | 4.527055665439997 | 459 | 5412 | 459 | 5567 | F443_03584,F443_11394,F443_21452,[illegible] |
| GO:BP | localization | GO:0051179 | FAUX | 0.000043467572636197326 | 4.361834611480181 | 1209 | 5412 | 1197 | 5567 | F443_02217,F443_02234,F443_03209,[illegible] |
| GO:CC | cellular anatomical structure | GO:0110165 | FAUX | 2.863615027900356e-10 | 9.543085367124062 | 5822 | 5888 | 5792 | 5929 | F443_00048,F443_00432,F443_00749,[illegible] |
