## Supplementary table 6 for "Zoospore-derived extracellular vesicles in the flagellated stramenopile *Phytophthora parasitica*"

**Title : Protein family expansion and subcellular localisation of the TOP400 proteins**

| Gene Id | Uniprot Entry | -10lgP | Protein names | Length | Protein families | Transmembrane domains | Signal P | GPI-anchor site | Localizations | Signals | Gene Ontology (biological process) | Gene Ontology (cellular component) | Gene Ontology (molecular function) | Gene Ontology (GO) | Proteomes |
|---|---|---|---|---|---|---|---|---|---|---|---|---|---|---|---|
| **Transmembrane proteins** | | | | | | | | | | | | | | | |
| PPTG_12232 | W2Q852 | 248,74 | ABC transporter domain-containing protein | 1948 | ABC transporter superfamily, ABCA family | 14 | 0 | 0 | Cell membrane \| Lysosome/Vacuole | Transmembrane domain | | membrane [GO:001 | ABC-type transporter act | membrane [G | UP000018817 |
| PPTG_11337 | W2QBU7 | 221,19 | ABC transporter domain-containing protein | 1983 | ABC transporter superfamily, ABCA family | 11 | 0 | 0 | Cell membrane \| Lysosome/Vacuole | Transmembrane domain | | membrane [GO:001 | ABC-type transporter act | membrane [G | UP000018817 |
| PPTG_08986 | W2QIM1 | 210,01 | ABC transporter domain-containing protein | 1945 | ABC transporter superfamily, ABCA family | 12 | 0 | 0 | Cell membrane \| Lysosome/Vacuole | Transmembrane domain | | membrane [GO:001 | ABC-type transporter act | membrane [G | UP000018817 |
| PPTG_08768 | W2QHM7 | 201,54 | ABC transporter domain-containing protein | 1969 | ABC transporter superfamily, ABCA family | 13 | 0 | 0 | Cell membrane \| Lysosome/Vacuole | Transmembrane domain | | membrane [GO:001 | ABC-type transporter act | membrane [G | UP000018817 |
| PPTG_12681 | W2Q2U4 | 240,96 | ABC transporter B family member 11 | 1290 | ABC transporter superfamily, ABCB family, Multidrug resistanc | 9 | 1 | 1 | Cell membrane \| Lysosome/Vacuole | Transmembrane domain | oligopeptide export from m | mitochondrial inner | ABC-type oligopeptide tr | mitochondrial | UP000018817 |
| PPTG_10588 | W2QD53 | 206,91 | ABC transporter domain-containing protein | 1351 | ABC transporter superfamily, ABCG family, PDR (TC 3.A.1.205) | 11 | 0 | 0 | Cell membrane \| Lysosome/Vacuole | Transmembrane domain | | membrane [GO:001 | ABC-type transporter act | membrane [G | UP000018817 |
| PPTG_10590 | W2QC67 | 192,26 | ABC transporter domain-containing protein | 1358 | ABC transporter superfamily, ABCG family, PDR (TC 3.A.1.205) | 11 | 0 | 0 | Cell membrane \| Lysosome/Vacuole | Transmembrane domain | | membrane [GO:001 | ABC-type transporter act | membrane [G | UP000018817 |
| PPTG_18357 | W2PJ83 | 189,26 | ABC transporter domain-containing protein | 1308 | ABC transporter superfamily, ABCG family, PDR (TC 3.A.1.205) | 11 | 0 | 0 | Cell membrane | Transmembrane domain | | membrane [GO:001 | ABC-type transporter act | membrane [G | UP000018817 |
| PPTG_10595 | W2QBA2 | 177,66 | ABC transporter domain-containing protein | 1354 | ABC transporter superfamily, ABCG family, PDR (TC 3.A.1.205) | 10 | 0 | 0 | Cell membrane \| Lysosome/Vacuole | Transmembrane domain | | membrane [GO:001 | ABC-type transporter act | membrane [G | UP000018817 |
| PPTG_06725 | W2QQI2 | 176,51 | ABC transporter domain-containing protein | 1349 | ABC transporter superfamily, ABCG family, PDR (TC 3.A.1.205) | 11 | 0 | 0 | Cell membrane \| Lysosome/Vacuole | Transmembrane domain | | membrane [GO:001 | ABC-type transporter act | membrane [G | UP000018817 |
| PPTG_15911 | W2PRV1 | 177,87 | Calnexin | 557 | Calreticulin family | 1 | 1 | 0 | Endoplasmic reticulum | Signal peptide \| Transmembrane domain | ERAD pathway [GO:00365 | endoplasmic reticu | calcium ion binding [GO: | endoplasmic r | UP000018817 |
| PPTG_07345 | W2QPP7 | 244,2 | Calcium-transporting ATPase (EC 7.2.2.10) | 1048 | Cation transport ATPase (P-type) (TC 3.A.3) family | 8 | 0 | 0 | Cell membrane \| Lysosome/Vacuole | Transmembrane domain | | endomembrane sys | ATP binding [GO:000552 | endomembra | UP000018817 |
| PPTG_18730 | W2PFQ4 | 212,59 | Calcium-transporting ATPase (EC 7.2.2.10) | 1059 | Cation transport ATPase (P-type) (TC 3.A.3) family | 7 | 0 | 0 | Cell membrane \| Lysosome/Vacuole | Transmembrane domain | copper ion transport [GO:0 | endomembrane sys | ATP binding [GO:000552 | endomembra | UP000018817 |
| PPTG_09633 | W2QFM9 | 297,63 | Cation-transporting P-type ATPase N-terminal domain-containing protein | 1343 | Cation transport ATPase (P-type) (TC 3.A.3) family | 10 | 1 | 0 | Cell membrane | Transmembrane domain | intracellular potassium ior | plasma membrane | ATP binding [GO:000552 | plasma memb | UP000018817 |
| PPTG_05941 | W2QUI9 | 233,12 | Cation-transporting P-type ATPase N-terminal domain-containing protein | 1076 | Cation transport ATPase (P-type) (TC 3.A.3) family | 8 | 0 | 0 | Lysosome/Vacuole | Signal peptide \| Transmembrane domain | intracellular potassium ior | plasma membrane | ATP binding [GO:000552 | plasma memb | UP000018817 |
| PPTG_03625 | W2R5K1 | 203,87 | P-type Ca(2+) transporter (EC 7.2.2.10) | 1045 | Cation transport ATPase (P-type) (TC 3.A.3) family, Type IIA subf | 7 | 0 | 0 | Lysosome/Vacuole | Signal peptide \| Transmembrane domain | | membrane [GO:001 | ATP binding [GO:000552 | membrane [G | UP000018817 |
| PPTG_02226 | W2RC14 | 187,37 | Calcium-transporting ATPase (EC 7.2.2.10) | 1067 | Cation transport ATPase (P-type) (TC 3.A.3) family, Type IIB sub | 8 | 0 | 1 | Cell membrane \| Lysosome/Vacuole | Transmembrane domain | copper ion transport [GO:0 | plasma membrane | ATP binding [GO:000552 | plasma memb | UP000018817 |
| PPTG_17691 | W2PLE5 | 203,88 | Plasma membrane ATPase (EC 7.1.2.1) | 793 | Cation transport ATPase (P-type) (TC 3.A.3) family, Type IIIA sub | 4 | 0 | 0 | Cell membrane | Transmembrane domain | proton export across plasn | plasma membrane | ATP binding [GO:000552 | plasma memb | UP000018817 |
| PPTG_02282 | W2RAS2 | 220,97 | Phospholipid-transporting ATPase (EC 7.6.2.1) | 1516 | Cation transport ATPase (P-type) (TC 3.A.3) family, Type IV subf | 8 | 0 | 0 | Lysosome/Vacuole | Signal peptide \| Transmembrane domain | phospholipid translocation | endomembrane sys | ATP binding [GO:000552 | endomembra | UP000018817 |
| PPTG_15328 | W2PSP8 | 195,86 | Phospholipid-transporting ATPase (EC 7.6.2.1) | 1391 | Cation transport ATPase (P-type) (TC 3.A.3) family, Type IV subf | 10 | 0 | 0 | Lysosome/Vacuole | Signal peptide \| Transmembrane domain | phospholipid translocation | endomembrane sys | ATP binding [GO:000552 | endomembra | UP000018817 |
| PPTG_00417 | W2RFD5 | 182,66 | Phospholipid-transporting ATPase (EC 7.6.2.1) | 1324 | Cation transport ATPase (P-type) (TC 3.A.3) family, Type IV subf | 9 | 0 | 0 | Lysosome/Vacuole \| Golgi apparatus | Signal peptide \| Transmembrane domain | phospholipid translocation | endomembrane sys | ATP binding [GO:000552 | endomembra | UP000018817 |
| PPTG_10537 | W2QC18 | 178,59 | Phospholipid-transporting ATPase (EC 7.6.2.1) | 1291 | Cation transport ATPase (P-type) (TC 3.A.3) family, Type IV subf | 10 | 0 | 0 | Lysosome/Vacuole | Signal peptide \| Transmembrane domain | phospholipid translocation | plasma membrane | ATP binding [GO:000552 | plasma memb | UP000018817 |
| PPTG_16875 | W2PNH8 | 197,28 | Cation-transporting ATPase (EC 7.2.2.-) | 1339 | Cation transport ATPase (P-type) (TC 3.A.3) family, Type V subfa | 4 | 0 | 0 | Lysosome/Vacuole | Signal peptide \| Transmembrane domain | | membrane [GO:001 | ATP binding [GO:000552 | membrane [G | UP000018817 |
| PPTG_12608 | W2Q1R2 | 184,97 | CSC1/OSCA1-like 7TM region domain-containing protein | 845 | CSC1 (TC 1.A.17) family | 11 | 0 | 0 | Cell membrane | Transmembrane domain | | plasma membrane | calcium-activated cation | plasma memb | UP000018817 |
| PPTG_06268 | W2QS57 | 218,25 | Uncharacterized protein | 811 | CTL (choline transporter-like) family | 10 | 0 | 0 | Cell membrane \| Lysosome/Vacuole | Transmembrane domain | | membrane [GO:001 | transmembrane transpo | membrane [G | UP000018817 |
| PPTG_03845 | W2QY59 | 189,77 | Glycoside hydrolase family 5 domain-containing protein | 582 | Glycosyl hydrolase 5 (cellulase A) family | 1 | 0 | 0 | Cell membrane | Signal peptide | cellulose catabolic process [GO:0030245] | | hydrolase activity, hydrol | hydrolase acti | UP000018817 |
| PPTG_01939 | W2RAZ9 | 215,53 | X8 domain-containing protein | 750 | Glycosyl hydrolase 5 (cellulase A) family | 1 | 1 | 0 | Lysosome/Vacuole | Signal peptide \| Transmembrane domain | glucan catabolic process [ | cell surface [GO:00 | beta-glucosidase activity | cell surface [G | UP000018817 |
| PPTG_14787 | W2PVX1 | 232,91 | EGF-like domain-containing protein | 627 | Glycosyl hydrolase 72 family | 1 | 0 | 0 | Extracellular | Signal peptide | cell wall (1->3)-beta-D-gluc | plasma membrane | 1,3-beta-glucanosyltrans | plasma memb | UP000018817 |
| PPTG_08579 | W2QN32 | 245,01 | 1,3-beta-glucan synthase (EC 2.4.1.34) | 2228 | Glycosyltransferase 48 family | 25 | 0 | 0 | Cell membrane | Transmembrane domain | (1->3)-beta-D-glucan biosy | 1,3-beta-D-glucan s | 1,3-beta-D-glucan syntha | 1,3-beta-D-glu | UP000018817 |
| PPTG_13258 | W2Q0U2 | 236,43 | 1,3-beta-glucan synthase (EC 2.4.1.34) | 2286 | Glycosyltransferase 48 family | 25 | 0 | 0 | Cell membrane | Transmembrane domain | (1->3)-beta-D-glucan biosy | 1,3-beta-D-glucan s | 1,3-beta-D-glucan syntha | 1,3-beta-D-glu | UP000018817 |
| PPTG_13182 | W2PZY0 | 219,29 | 1,3-beta-glucan synthase (EC 2.4.1.34) | 2447 | Glycosyltransferase 48 family | 26 | 0 | 0 | Cell membrane | Transmembrane domain | (1->3)-beta-D-glucan biosy | 1,3-beta-D-glucan s | 1,3-beta-D-glucan syntha | 1,3-beta-D-glu | UP000018817 |
| PPTG_14740 | W2PVT8 | 204,39 | 1,3-beta-glucan synthase (EC 2.4.1.34) | 2040 | Glycosyltransferase 48 family | 11 | 0 | 0 | Cell membrane | Signal peptide \| Transmembrane domain | (1->3)-beta-D-glucan biosy | 1,3-beta-D-glucan s | 1,3-beta-D-glucan syntha | 1,3-beta-D-glu | UP000018817 |
| PPTG_11437 | W2Q9L5 | 179,13 | malate dehydrogenase (EC 1.1.1.37) | 336 | LDH/MDH superfamily, MDH type 2 family | 1 | 0 | 0 | Cytoplasm | | malate metabolic process [GO:0006108] | | L-malate dehydrogenase | L-malate dehy | UP000018817 |
| PPTG_04703 | W2R288 | 235,49 | ADP/ATP translocase (ADP,ATP carrier protein) | 310 | Mitochondrial carrier (TC 2.A.29) family | 3 | 0 | 0 | Mitochondrion | Transmembrane domain | mitochondrial ADP transm | mitochondrial inner | ATP:ADP antiporter activ | mitochondrial | UP000018817 |
| PPTG_01692 | W2R8G7 | 207,27 | Mitochondrial carnitine/acylcarnitine carrier protein | 310 | Mitochondrial carrier (TC 2.A.29) family | 2 | 0 | 0 | Mitochondrion | Transmembrane domain | carnitine transmembrane | mitochondrial mem | O-acyl-L-carnitine transr | mitochondrial | UP000018817 |
| PPTG_06649 | W2QSA7 | 188,16 | Nicastrin | 719 | Nicastrin family | 1 | 1 | 0 | Lysosome/Vacuole | Signal peptide \| Transmembrane domain | Notch signaling pathway [G | plasma membrane [GO:0005886] | | plasma memb | UP000018817 |
| PPTG_13413 | W2Q4S5 | 206,39 | Dolichyl-diphosphooligosaccharide--protein glycosyltransferase subunit 1 | 470 | OST1 family | 1 | 1 | 0 | Endoplasmic reticulum | Signal peptide \| Transmembrane domain | protein N-linked glycosylat | oligosaccharyltransferase complex [GO:0008 | | oligosaccharyl | UP000018817 |
| PPTG_10900 | W2QAL4 | 198,15 | Peptidase M28 domain-containing protein | 868 | Peptidase M28 family, M28B subfamily | 1 | 0 | 0 | Lysosome/Vacuole \| Golgi apparatus | Signal peptide \| Transmembrane domain | | | carboxypeptidase activit | carboxypeptic | UP000018817 |
| PPTG_08136 | W2QK24 | 184,8 | dolichyl-diphosphooligosaccharide--protein glycotransferase (EC 2.4.99.18) | 885 | STT3 family | 13 | 0 | 0 | Endoplasmic reticulum | Signal peptide \| Transmembrane domain | | endomembrane sys | dolichyl-diphosphooligos | endomembra | UP000018817 |
| PPTG_05924 | W2QWJ0 | 221,54 | V-type proton ATPase subunit a | 869 | V-ATPase 116 kDa subunit family | 6 | 0 | 0 | Lysosome/Vacuole | Transmembrane domain | vacuolar acidification [GO: | vacuolar proton-tra | ATPase binding [GO:005 | vacuolar proto | UP000018817 |
| PPTG_09717 | W2QGS2 | 180,51 | ABC transmembrane type-1 domain-containing protein | 1011 | | 9 | 0 | 0 | Cell membrane | Signal peptide \| Transmembrane domain | | membrane [GO:001 | ABC-type transporter act | membrane [G | UP000018817 |
| PPTG_09177 | W2QHJ6 | 200,83 | Alpha-L-glutamate ligase-related protein ATP-grasp domain-containing protein | 426 | | 1 | 0 | 0 | Cytoplasm \| Endoplasmic reticulum | | | | | | UP000018817 |
| PPTG_06857 | W2QSZ3 | 256,69 | C2 domain-containing protein | 1764 | | 1 | 0 | 0 | Cell membrane | Signal peptide \| Transmembrane domain | plasma membrane organiz | membrane [GO:0016020] | | membrane [G | UP000018817 |
| PPTG_15898 | W2PSQ3 | 207,7 | C2 domain-containing protein | 1174 | | 7 | 0 | 0 | Endoplasmic reticulum | Transmembrane domain | | membrane [GO:001 | chloride channel activity | membrane [G | UP000018817 |
| PPTG_12292 | W2Q6M6 | 188,48 | Calcineurin-like phosphoesterase domain-containing protein | 479 | | 1 | 0 | 0 | Extracellular \| Cell membrane | Signal peptide | | | hydrolase activity [GO:0 | hydrolase acti | UP000018817 |
| PPTG_17902 | W2PJ95 | 210,81 | Cellulose synthase 3 | 1142 | | 14 | 0 | 0 | Golgi apparatus | Signal peptide \| Transmembrane domain | cellulose biosynthetic proc | endomembrane sys | cellulose synthase (UDP- | endomembra | UP000018817 |
| PPTG_06395 | W2QVB8 | 250,16 | EGF-like domain-containing protein | 1089 | | 1 | 1 | 0 | Lysosome/Vacuole \| Golgi apparatus | Signal peptide \| Transmembrane domain | | | | | UP000018817 |
| PPTG_00165 | W2RDT8 | 197,84 | EGF-like domain-containing protein | 723 | | 1 | 1 | 0 | Cell membrane \| Lysosome/Vacuole \| Golgi apparatus | Signal peptide \| Transmembrane domain | (1->6)-beta-D-glucan biosy | endoplasmic reticu | glucosidase activity [GO: | endoplasmic r | UP000018817 |
| PPTG_02785 | W2REY1 | 186,9 | H(+)-exporting diphosphatase (EC 7.1.3.1) | 768 | | 12 | 0 | 0 | Cell membrane \| Lysosome/Vacuole | Transmembrane domain | | endomembrane sys | diphosphate hydrolysis-c | endomembra | UP000018817 |
| PPTG_13069 | W2Q664 | 179,41 | MARVEL domain-containing protein | 357 | | 8 | 0 | 0 | Lysosome/Vacuole | | | membrane [GO:0016020] | | membrane [G | UP000018817 |
| PPTG_12938 | W2Q3X5 | 250,14 | PH domain-containing protein | 1016 | | 8 | 0 | 0 | Golgi apparatus | Signal peptide \| Transmembrane domain | | membrane [GO:001 | glycosyltransferase activ | membrane [G | UP000018817 |
| PPTG_12937 | W2Q357 | 249,82 | PH domain-containing protein | 1026 | | 7 | 0 | 0 | Lysosome/Vacuole \| Golgi apparatus | Signal peptide \| Transmembrane domain | | membrane [GO:001 | glycosyltransferase activ | membrane [G | UP000018817 |
| PPTG_09817 | W2QC07 | 193,89 | Thioredoxin domain-containing protein | 469 | | 2 | 0 | 0 | Endoplasmic reticulum \| Golgi apparatus | Signal peptide \| Transmembrane domain | | COPII-coated ER to Golgi transport vesicle [G | | COPII-coated | UP000018817 |
| PPTG_06246 | W2QSA5 | 180,31 | TKL protein kinase | 674 | | 1 | 1 | 0 | Cell membrane \| Lysosome/Vacuole | Signal peptide \| Transmembrane domain | | | ATP binding [GO:000552 | ATP binding [G | UP000018817 |
| PPTG_06389 | W2QVA6 | 324,08 | Uncharacterized protein | 878 | | 1 | 1 | 0 | Lysosome/Vacuole | Signal peptide \| Transmembrane domain | | | | | UP000018817 |
| PPTG_16242 | W2PNS1 | 292,42 | Uncharacterized protein | 854 | | 5 | 1 | 0 | Cell membrane \| Lysosome/Vacuole | Signal peptide \| Transmembrane domain | | membrane [GO:0016020] | | membrane [G | UP000018817 |
| PPTG_06394 | W2QUN2 | 265,09 | Uncharacterized protein | 1125 | | 1 | 1 | 0 | Golgi apparatus | Signal peptide \| Transmembrane domain | | | | | UP000018817 |
| PPTG_06382 | W2QSV6 | 260,17 | Uncharacterized protein | 1089 | | 1 | 1 | 0 | Golgi apparatus | Signal peptide \| Transmembrane domain | | | | | UP000018817 |
| PPTG_00714 | W2RG42 | 217,72 | Uncharacterized protein | 891 | | 5 | 0 | 0 | Cell membrane | Signal peptide \| Transmembrane domain | | membrane [GO:0016020] | | membrane [G | UP000018817 |
| PPTG_00859 | W2RGS8 | 215,93 | Uncharacterized protein | 604 | | 1 | 0 | 0 | Cell membrane | Signal peptide \| Transmembrane domain | | | | | UP000018817 |
| PPTG_04558 | W2R3T9 | 202,49 | Uncharacterized protein | 542 | | 1 | 0 | 0 | Cell membrane | Signal peptide \| Transmembrane domain | cellulose catabolic process [GO:0030245] | | | cellulose cata | UP000018817 |
| PPTG_12822 | W2Q0U7 | 174,7 | Uncharacterized protein | 456 | | 1 | 0 | 0 | Extracellular | Signal peptide | | | | | UP000018817 |
| **Extracellular proteins** | | | | | | | | | | | | | | | |
| PPTG_05975 | W2QWP6 | 182,47 | Carbonic anhydrase (EC 4.2.1.1) | 272 | Alpha-carbonic anhydrase family | 0 | 1 | 0 | Extracellular | Signal peptide | | | carbonate dehydratase a | carbonate deh | UP000018817 |
| PPTG_07665 | W2QN93 | 207,54 | Glycosyl hydrolase family 30 TIM-barrel domain-containing protein | 540 | Glycosyl hydrolase 30 family | 0 | 1 | 0 | Extracellular | Signal peptide | glucosylceramide cataboli | membrane [GO:001 | glucosylceramidase acti | membrane [G | UP000018817 |
| PPTG_11759 | W2Q908 | 210,83 | Alpha-mannosidase (EC 3.2.1.-) | 1025 | Glycosyl hydrolase 38 family | 0 | 1 | 0 | Extracellular | Signal peptide | mannose metabolic process [GO:0006013] | | alpha-mannosidase acti | alpha-mannos | UP000018817 |
| PPTG_01216 | W2R636 | 232,64 | alpha-glucosidase (EC 3.2.1.20) (Maltase) | 1142 | Glycosyl hydrolase 31 family | 0 | 1 | 0 | Extracellular | Signal peptide | carbohydrate metabolic process [GO:0005975 | | alpha-glucosidase activi | alpha-glucosid | UP000018817 |
| PPTG_18587 | W2PHU9 | 181,72 | cellulose 1,4-beta-cellobiosidase (non-reducing end) (EC 3.2.1.91) | 521 | Glycosyl hydrolase 7 (cellulase C) family | 0 | 1 | 0 | Extracellular | Signal peptide | cellulose catabolic process [GO:0030245] | | cellulose 1,4-beta-cellob | cellulose 1,4-b | UP000018817 |
| PPTG_09208 | W2QHN0 | 235,72 | Glycoside hydrolase family 5 domain-containing protein | 505 | Glycosyl hydrolase 72 family | 0 | 1 | 0 | Extracellular | Signal peptide | cell wall (1->3)-beta-D-gluc | plasma membrane | 1,3-beta-glucanosyltrans | plasma memb | UP000018817 |
| PPTG_08245 | W2QLK3 | 227,82 | Luminal-binding protein | 658 | Heat shock protein 70 family | 0 | 1 | 0 | Endoplasmic reticulum | Signal peptide | | | ATP binding [GO:000552 | ATP binding [G | UP000018817 |
| PPTG_02565 | W2RBF0 | 248,99 | Serine protease (EC 3.4.21.-) | 808 | Peptidase S1B family | 0 | 1 | 0 | Extracellular | Signal peptide | proteolysis [GO:0006508] | | serine-type peptidase ac | serine-type pe | UP000018817 |
| PPTG_03117 | W2R3Q4 | 212,29 | subtilisin (EC 3.4.21.62) | 486 | Peptidase S8 family | 0 | 1 | 0 | Extracellular \| Lysosome/Vacuole | Signal peptide | proteolysis [GO:0006508] | | serine-type endopeptida | serine-type er | UP000018817 |
| PPTG_19949 | W2PAP8 | 179,71 | subtilisin (EC 3.4.21.62) | 487 | Peptidase S8 family | 0 | 1 | 0 | Extracellular \| Lysosome/Vacuole | Signal peptide | proteolysis [GO:0006508] | | serine-type endopeptida | serine-type er | UP000018817 |
| PPTG_03836 | W2QY41 | 243 | glucan endo-1,3-beta-D-glucosidase (EC 3.2.1.39) | 556 | PGA52 family | 0 | 1 | 0 | Extracellular | Signal peptide | cell wall organization [GO:0071555] | | glucan endo-1,3-beta-D- | glucan endo-1 | UP000018817 |
| PPTG_02370 | W2RAI7 | 182,3 | glucan endo-1,3-beta-D-glucosidase (EC 3.2.1.39) | 750 | Glycosyl hydrolase 81 family | 0 | 1 | 0 | Extracellular | Signal peptide | cell wall organization [GO:0071555]; polysacch | | endo-1,3(4)-beta-glucan | endo-1,3(4)-b | UP000018817 |
| PPTG_18309 | W2PGM1 | 210,04 | Protein disulfide-isomerase (EC 5.3.4.1) | 521 | Protein disulfide isomerase family | 0 | 1 | 0 | Endoplasmic reticulum | Signal peptide | protein folding [GO:00064 | endoplasmic reticu | protein disulfide isomera | endoplasmic r | UP000018817 |
| PPTG_06749 | W2QQK3 | 175,1 | Uncharacterized protein | 341 | RNase T2 family | 0 | 1 | 0 | Extracellular | Signal peptide | RNA catabolic process [GC | extracellular region | ribonuclease T2 activity [ | extracellular r | UP000018817 |
| PPTG_01676 | W2R8E6 | 236,92 | Guanine nucleotide-binding protein subunit beta-like protein | 317 | WD repeat G protein beta family, Ribosomal protein RACK1 sub | 0 | 1 | 0 | Cytoplasm \| Nucleus | Nuclear localization signal \| Nuclear export signal | | | ribosome binding [GO:0 | ribosome binc | UP000018817 |
| PPTG_02441 | W2RAR2 | 298,82 | EGF-like domain-containing protein | 597 | Mastigoneme protein family | 0 | 1 | 0 | Endoplasmic reticulum | Signal peptide | | | | | UP000018817 |
| PPTG_03648 | W2R5N9 | 271,67 | EGF-like domain-containing protein | 393 | Mastigoneme protein family | 0 | 1 | 0 | Endoplasmic reticulum | Signal peptide | | | | | UP000018817 |
| PPTG_07238 | W2QPH8 | 199,45 | EGF-like domain-containing protein | 226 | Mastigoneme protein family | 0 | 1 | 0 | Extracellular | Signal peptide | heterophilic cell-cell adhesion via plasma men | | cell adhesion molecule b | cell adhesion | UP000018817 |
| PPTG_06730 | W2QR91 | 245,82 | EGF-like domain-containing protein | 676 | | 0 | 1 | 0 | Extracellular | Signal peptide | | | | | UP000018817 |

| | | | | | | | | | | | | | | | |
|---|---|---|---|---|---|---|---|---|---|---|---|---|---|---|---|
| PPTG_16769 | W2PNB6 | 181,62 | EGF-like domain-containing protein | 1060 | | 0 | 1 | 0 | Extracellular | Signal peptide | | | | | UP000018817 |
| PPTG_08833 | W2QJT0 | 177,88 | EGF-like domain-containing protein | 679 | | 0 | 1 | 0 | Extracellular | Signal peptide | | | | | UP000018817 |
| PPTG_05659 | W2QVF4 | 174,75 | Apple domain-containing protein | 781 | | 0 | 1 | 0 | Extracellular | Signal peptide | proteolysis [GO:0006508] | extracellular region | flavin adenine dinucleoti | extracellular r | UP000018817 |
| PPTG_01093 | W2RI82 | 242,25 | Fibronectin type-III domain-containing protein | 1390 | | 0 | 1 | 0 | Extracellular | Signal peptide | | | | | UP000018817 |
| PPTG_17187 | W2PL97 | 200,46 | glucan endo-1,3-beta-D-glucosidase (EC 3.2.1.39) (Endo-1,3-beta-glucanase btg | 458 | | 0 | 1 | 0 | Extracellular | Signal peptide | cell wall organization [GO: | plasma membrane | glucan endo-1,3-beta-D- | plasma memb | UP000018817 |
| PPTG_00140 | W2RFV5 | 179,55 | Glycoside hydrolase | 460 | | 0 | 1 | 0 | Extracellular | Signal peptide | cellulose catabolic process [GO:0030245] | | hydrolase activity, hydrol | hydrolase acti | UP000018817 |
| PPTG_10783 | W2QBT9 | 181,75 | Jacalin-type lectin domain-containing protein | 1075 | | 0 | 1 | 0 | Extracellular | Signal peptide | | | | | UP000018817 |
| PPTG_11537 | W2Q6A3 | 193,04 | Lipase-like C-terminal domain-containing protein | 479 | | 0 | 1 | 0 | Extracellular | Signal peptide | lipid metabolic process [G | extracellular region | hydrolase activity [GO:0 | extracellular r | UP000018817 |
| PPTG_14116 | W2PXL8 | 196,72 | Peptidase C-terminal archaeal/bacterial domain-containing protein | 247 | | 0 | 1 | 0 | Extracellular | Signal peptide | | | | | UP000018817 |
| PPTG_01189 | W2RIK1 | 209,88 | Secreted protein | 457 | | 0 | 1 | 1 | Extracellular \| Cell membrane | | | | | | UP000018817 |
| PPTG_11360 | W2Q9A0 | 177,93 | SMP-30/Gluconolactonase/LRE-like region domain-containing protein | 690 | | 0 | 1 | 0 | Extracellular | Signal peptide | | | hydrolase activity [GO:0 | hydrolase acti | UP000018817 |
| PPTG_16264 | W2PQY5 | 314,02 | Spondin-like TSP1 domain-containing protein | 2399 | | 0 | 1 | 0 | Extracellular | Signal peptide | | | | | UP000018817 |
| PPTG_16236 | W2PPS8 | 176,87 | Transglutaminase-like domain-containing protein | 530 | | 0 | 1 | 0 | Extracellular | Signal peptide | | | aminoacyltransferase ac | aminoacyltrar | UP000018817 |
| PPTG_14253 | W2PZ52 | 194,17 | Tyrosinase copper-binding domain-containing protein | 631 | | 0 | 1 | 0 | Extracellular | Signal peptide | | | metal ion binding [GO:00 | metal ion binc | UP000018817 |
| PPTG_07817 | W2QPI2 | 176,11 | Uncharacterized protein | 506 | | 0 | 1 | 1 | Extracellular | | | | | | UP000018817 |
| PPTG_01302 | W2R6B5 | 174,2 | Uncharacterized protein | 512 | | 0 | 1 | 0 | Extracellular | Signal peptide | | | | | UP000018817 |
| PPTG_18605 | W2PFJ7 | 184,94 | YHYH domain-containing protein | 5468 | | 0 | 1 | 0 | Cell membrane | Signal peptide | | | | | UP000018817 |

**Cytoplasmic proteins**

| | | | | | | | | | | | | | | | |
|---|---|---|---|---|---|---|---|---|---|---|---|---|---|---|---|
| PPTG_04484 | W2R3J2 | 172,52 | 14-3-3-like protein | 249 | 14-3-3 family | 0 | 0 | 0 | Cytoplasm | Nuclear export signal | | | | | UP000018817 |
| PPTG_11762 | W2QAQ8 | 227,1 | 6-phosphogluconate dehydrogenase, decarboxylating (EC 1.1.1.44) | 489 | 6-phosphogluconate dehydrogenase family | 0 | 0 | 0 | Peroxisome | Peroxisomal targeting signal | D-gluconate metabolic process [GO:0019521]; | | NADP binding [GO:0050 | NADP binding | UP000018817 |
| PPTG_11894 | W2QB77 | 175,73 | 26S protease regulatory subunit 7 | 438 | AAA ATPase family | 0 | 0 | 0 | Cytoplasm | Nuclear export signal | proteolysis [GO:0006508] | cytoplasm [GO:000 | ATP binding [GO:00552 | cytoplasm [G | UP000018817 |
| PPTG_03857 | W2R0Z2 | 198,12 | Vesicle-fusing ATPase (EC 3.6.4.6) | 765 | AAA ATPase family | 0 | 0 | 0 | Cytoplasm \| Nucleus | | Golgi to plasma membran | Golgi stack [GO:00 | ATP binding [GO:00552 | Golgi stack [G | UP000018817 |
| PPTG_13558 | W2Q2U0 | 225,42 | Elongation factor 3 (Eukaryotic elongation factor 3) | 1038 | ABC transporter superfamily, ABCF family, EF3 subfamily | 0 | 0 | 0 | Cytoplasm | Nuclear localization signal | | cytoplasm [GO:000 | ATP binding [GO:00552 | cytoplasm [G | UP000018817 |
| PPTG_08479 | W2QKS8 | 217,19 | methylcrotonoyl-CoA carboxylase (EC 6.4.1.4) (3-methylcrotonyl-CoA carboxylas | 567 | AccD/PCCB family | 0 | 0 | 0 | Mitochondrion | Mitochondrial transit peptide | L-leucine catabolic proces | methylcrotonoyl-C | methylcrotonoyl-CoA ca | methylcroton | UP000018817 |
| PPTG_13419 | W2Q329 | 227,93 | Aconitate hydratase, mitochondrial (Aconitase) (EC 4.2.1.3) | 786 | Aconitase/IPM isomerase family | 0 | 0 | 0 | Mitochondrion | Mitochondrial transit peptide | tricarboxylic acid cycle [G | cytosol [GO:00058 | 4 iron, 4 sulfur cluster bi | cytosol [GO:0 | UP000018817 |
| PPTG_15348 | W2PVD2 | 229,14 | Actin-1 | 376 | Actin family | 0 | 0 | 0 | Cytoplasm \| Nucleus | Nuclear export signal | | | ATP binding [GO:00552 | ATP binding [G | UP000018817 |
| PPTG_19136 | W2PDK7 | 200,29 | Acyl-CoA dehydrogenase | 610 | Acyl-CoA dehydrogenase family | 0 | 0 | 0 | Mitochondrion | Mitochondrial transit peptide | fatty acid metabolic proce | mitochondrial inne | acyl-CoA dehydrogenase | mitochondrial | UP000018817 |
| PPTG_05779 | W2QUG7 | 172,7 | Acyl-CoA dehydrogenase family member 11 | 801 | Acyl-CoA dehydrogenase family | 0 | 0 | 0 | Peroxisome | Peroxisomal targeting signal | fatty acid beta-oxidation u | mitochondrial mem | acyl-CoA dehydrogenase | mitochondrial | UP000018817 |
| PPTG_05028 | W2QVJ2 | 254,74 | Acyl-coenzyme A dehydrogenase (EC 1.3.8.7) (EC 1.3.8.8) | 764 | Acyl-CoA dehydrogenase family | 0 | 0 | 0 | Mitochondrion | Mitochondrial transit peptide | fatty acid beta-oxidation u | mitochondrion [GO | flavin adenine dinucleoti | mitochondrio | UP000018817 |
| PPTG_18811 | W2PFW8 | 195,47 | Medium-chain specific acyl-CoA dehydrogenase, mitochondrial | 362 | Acyl-CoA dehydrogenase family | 0 | 0 | 0 | Cytoplasm | Nuclear export signal | medium-chain fatty acid c | mitochondrion [GO | flavin adenine dinucleoti | mitochondrio | UP000018817 |
| PPTG_11336 | W2QB00 | 183,7 | AP complex subunit beta | 837 | Adaptor complexes large subunit family | 0 | 0 | 0 | Cytoplasm | Nuclear localization signal | intracellular protein trans | clathrin adaptor co | clathrin binding [GO:003 | clathrin adapt | UP000018817 |
| PPTG_07612 | W2QN96 | 244,51 | AP-2 complex subunit alpha | 987 | Adaptor complexes large subunit family | 0 | 0 | 0 | Cytoplasm \| Lysosome/Vacuole \| Golgi apparatus | | clathrin-dependent endoc | AP-2 adaptor comp | clathrin adaptor activity [ | AP-2 adaptor | UP000018817 |
| PPTG_00328 | W2RGZ4 | 203,57 | Beta-adaptin appendage C-terminal subdomain domain-containing protein | 915 | Adaptor complexes large subunit family | 0 | 0 | 0 | Cytoplasm \| Lysosome/Vacuole \| Golgi apparatus | | intracellular protein trans | clathrin adaptor complex [GO:0030131]; cyt | | clathrin adapt | UP000018817 |
| PPTG_01949 | W2R9G7 | 203,49 | Adenosylhomocysteinase (EC 3.13.2.1) | 481 | Adenosylhomocysteinase family | 0 | 0 | 0 | Cytoplasm | Nuclear localization signal | one-carbon metabolic pro | cytosol [GO:00058 | adenosylhomocysteinas | cytosol [GO:0 | UP000018817 |
| PPTG_06616 | W2QS77 | 193,99 | Adenylosuccinate synthetase (AMPSase) (AdSS) (EC 6.3.4.4) (IMP--aspartate liga | 554 | Adenylosuccinate synthetase family | 0 | 0 | 0 | Mitochondrion | Mitochondrial transit peptide | 'de novo' AMP biosynthetic | cytoplasm [GO:000 | adenylosuccinate syntha | cytoplasm [G | UP000018817 |
| PPTG_01838 | W2RAJ2 | 195,48 | phosphoribosylaminoimidazole carboxylase (EC 4.1.1.21) (AIR carboxylase) | 595 | AIR carboxylase family, Class I subfamily | 0 | 0 | 0 | Cytoplasm | Peroxisomal targeting signal | 'de novo' IMP biosynthetic process [GO:00061 | | ATP binding [GO:00552 | ATP binding [G | UP000018817 |
| PPTG_16640 | W2PM64 | 199,29 | Aldehyde dehydrogenase domain-containing protein | 547 | Aldehyde dehydrogenase family | 0 | 0 | 0 | Mitochondrion | Mitochondrial transit peptide | | | aldehyde dehydrogenase | aldehyde deh | UP000018817 |
| PPTG_02473 | W2RDE5 | 215,97 | Aldehyde dehydrogenase domain-containing protein | 525 | Aldehyde dehydrogenase family | 0 | 0 | 0 | Mitochondrion | Mitochondrial transit peptide | | | oxidoreductase activity, | oxidoreductas | UP000018817 |
| PPTG_02435 | W2RD85 | 222,21 | Aldehyde dehydrogenase domain-containing protein | 525 | Aldehyde dehydrogenase family | 0 | 0 | 0 | Mitochondrion | Mitochondrial transit peptide | | | oxidoreductase activity, | oxidoreductas | UP000018817 |
| PPTG_16241 | W2PP34 | 200,16 | 2-oxoglutarate dehydrogenase, mitochondrial (EC 1.2.4.2) (2-oxoglutarate dehyd | 1043 | Alpha-ketoglutarate dehydrogenase family | 0 | 0 | 0 | Mitochondrion | Mitochondrial transit peptide | tricarboxylic acid cycle [G | mitochondrion [GO | metal ion binding [GO:00 | mitochondrio | UP000018817 |
| PPTG_17884 | W2PI34 | 230,79 | Annexin | 328 | Annexin family | 0 | 0 | 0 | Cytoplasm \| Nucleus | Nuclear localization signal | | cytoplasm [GO:000 | calcium ion binding [GO: | cytoplasm [G | UP000018817 |
| PPTG_17883 | W2PII0 | 230,35 | Annexin | 328 | Annexin family | 0 | 0 | 0 | Cytoplasm \| Nucleus | Nuclear localization signal | | cytoplasm [GO:000 | calcium ion binding [GO: | cytoplasm [G | UP000018817 |
| PPTG_03931 | W2QYV6 | 185,33 | Annexin | 690 | Annexin family | 0 | 0 | 0 | Cytoplasm | | | cytoplasm [GO:000 | calcium ion binding [GO: | cytoplasm [G | UP000018817 |
| PPTG_13120 | W2Q4N9 | 174,49 | Annexin | 329 | Annexin family | 0 | 0 | 0 | Cytoplasm \| Nucleus | Nuclear localization signal \| Nuclear export signal | | cytoplasm [GO:000 | calcium ion binding [GO: | cytoplasm [G | UP000018817 |
| PPTG_12013 | W2Q5J5 | 220,58 | Piwi domain-containing protein | 1268 | Argonaute family | 0 | 0 | 0 | Cytoplasm \| Nucleus | Nuclear localization signal | | | RNA binding [GO:000372 | RNA binding [ | UP000018817 |
| PPTG_07447 | W2QPG4 | 180,4 | Uncharacterized protein | 930 | Argonaute family | 0 | 0 | 0 | Cytoplasm | Nuclear localization signal | | | RNA binding [GO:000372 | RNA binding [ | UP000018817 |
| PPTG_02311 | W2RCG4 | 229,72 | Creatine kinase | 435 | ATP:guanido phosphotransferase family | 0 | 0 | 0 | Mitochondrion | Mitochondrial transit peptide | phosphocreatine biosynth | extracellular space | ATP binding [GO:00552 | extracellular s | UP000018817 |
| PPTG_10603 | W2QD70 | 244,39 | Creatine kinase, flagellar | 785 | ATP:guanido phosphotransferase family | 0 | 0 | 0 | Cytoplasm | | phosphocreatine biosynth | extracellular space | ATP binding [GO:00552 | extracellular s | UP000018817 |
| PPTG_13841 | W2PYH8 | 283,63 | ATP synthase subunit beta (EC 7.1.2.2) | 501 | ATPase alpha/beta chains family | 0 | 0 | 0 | Mitochondrion | Mitochondrial transit peptide | proton motive force-driven | mitochondrial inne | ATP binding [GO:00552 | mitochondrial | UP000018817 |
| PPTG_16803 | W2PNG4 | 236,48 | H(+)-transporting two-sector ATPase (EC 7.1.2.2) | 618 | ATPase alpha/beta chains family | 0 | 0 | 0 | Cytoplasm | Nuclear export signal | ATP metabolic process [G | proton-transporting | ATP binding [GO:00552 | proton-transp | UP000018817 |
| PPTG_12134 | W2Q7V6 | 223,27 | Vacuolar proton pump subunit B (V-ATPase subunit B) (Vacuolar proton pump su | 495 | ATPase alpha/beta chains family | 0 | 0 | 0 | Cytoplasm | | ATP metabolic process [G | proton-transporting | ATP binding [GO:00552 | proton-transp | UP000018817 |
| PPTG_01682 | W2R7Z6 | 217,92 | Uncharacterized protein | 183 | ATPase d subunit family | 0 | 0 | 0 | Mitochondrion | Mitochondrial transit peptide | proton motive force-driven | mitochondrial inne | proton transmembrane t | mitochondrial | UP000018817 |
| PPTG_01995 | W2R940 | 221,18 | F-ATPase gamma subunit | 304 | ATPase gamma chain family | 0 | 0 | 0 | Mitochondrion | Mitochondrial transit peptide | | proton-transporting | proton-transporting ATP | proton-transp | UP000018817 |
| PPTG_05574 | W2QY66 | 172,95 | Band 7 domain-containing protein | 376 | Band 7/mec-2 family | 0 | 0 | 0 | Mitochondrion | Mitochondrial transit peptide | mitochondrion organizatio | membrane [GO:0016020]; mitochondrion [G | | membrane [G | UP000018817 |
| PPTG_19590 | W2PCJ2 | 217,31 | Calreticulin | 458 | Calreticulin family | 0 | 0 | 0 | Endoplasmic reticulum | Signal peptide | ERAD pathway [GO:00365 | endoplasmic reticu | calcium ion binding [GO: | endoplasmic | UP000018817 |
| PPTG_05429 | W2QX24 | 183,35 | cAMP-dependent protein kinase regulatory subunit | 290 | CAMP-dependent kinase regulatory chain family | 0 | 0 | 0 | Cytoplasm \| Nucleus | Nuclear localization signal \| Nuclear export signal | | cAMP-dependent p | cAMP binding [GO:0030 | cAMP-depend | UP000018817 |
| PPTG_00673 | W2RFX8 | 200,91 | TATA-binding protein interacting (TIP20) domain-containing protein | 1180 | CAND family | 0 | 0 | 0 | Cytoplasm \| Nucleus | Nuclear localization signal | SCF complex assembly [GO:0010265] | | | SCF complex | UP000018817 |
| PPTG_07317 | W2QPK9 | 186,66 | C-CAP/cofactor C-like domain-containing protein | 461 | CAP family | 0 | 0 | 0 | Cytoplasm | | actin filament organizatio | cytoplasm [GO:000 | actin binding [GO:00037 | cytoplasm [G | UP000018817 |
| PPTG_14525 | W2PVS0 | 253,13 | Carbamoyl phosphate synthase arginine-specific large chain (EC 6.3.4.16) (EC 6. | 1511 | CarB family | 0 | 0 | 0 | Mitochondrion | Mitochondrial transit peptide | 'de novo' pyrimidine nucle | carbamoyl-phospha | ATP binding [GO:00552 | carbamoyl-ph | UP000018817 |
| PPTG_02700 | W2RCJ5 | 183,35 | CN hydrolase domain-containing protein | 323 | Carbon-nitrogen hydrolase superfamily, NIT1/NIT2 family | 0 | 0 | 0 | Mitochondrion | Mitochondrial transit peptide | asparagine metabolic pro | mitochondrion [GO | omega-amidase activity | mitochondrio | UP000018817 |
| PPTG_02824 | W2RF50 | 189 | Choline/carnitine acyltransferase domain-containing protein | 621 | Carnitine/choline acetyltransferase family | 0 | 0 | 0 | Mitochondrion | Mitochondrial transit peptide | fatty acid beta-oxidation [ | mitochondrion [GO | carnitine O-palmitoyltra | mitochondrio | UP000018817 |
| PPTG_11901 | W2QB87 | 212,82 | Citrate synthase | 464 | Citrate synthase family | 0 | 0 | 0 | Mitochondrion | Mitochondrial transit peptide | carbohydrate metabolic p | mitochondrial mat | acyltransferase activity, | mitochondrial | UP000018817 |
| PPTG_05552 | W2QXD3 | 189,09 | fructose-bisphosphate aldolase (EC 4.1.2.13) | 358 | Class II fructose-bisphosphate aldolase family | 0 | 0 | 0 | Cytoplasm \| Nucleus | Nuclear export signal | gluconeogenesis [GO:000 | cytosol [GO:00058 | fructose-bisphosphate a | cytosol [GO:0 | UP000018817 |
| PPTG_05553 | W2QXP4 | 189,09 | fructose-bisphosphate aldolase (EC 4.1.2.13) | 358 | Class II fructose-bisphosphate aldolase family | 0 | 0 | 0 | Cytoplasm \| Nucleus | Nuclear export signal | gluconeogenesis [GO:000 | cytosol [GO:00058 | fructose-bisphosphate a | cytosol [GO:0 | UP000018817 |
| PPTG_02578 | W2RDH0 | 203,84 | arginine--tRNA ligase (EC 6.1.1.19) (Arginyl-tRNA synthetase) | 815 | Class-I aminoacyl-tRNA synthetase family | 0 | 0 | 0 | Cytoplasm | Nuclear localization signal | arginyl-tRNA aminoacylati | cytoplasm [GO:000 | arginine-tRNA ligase acti | cytoplasm [G | UP000018817 |
| PPTG_13500 | W2Q3G6 | 194,8 | cysteine--tRNA ligase (EC 6.1.1.16) (Cysteinyl-tRNA synthetase) | 688 | Class-I aminoacyl-tRNA synthetase family | 0 | 0 | 0 | Cytoplasm | Nuclear localization signal | cysteinyl-tRNA aminoacyla | cytoplasm [GO:000 | ATP binding [GO:00552 | cytoplasm [G | UP000018817 |
| PPTG_12917 | W2Q337 | 200,59 | isoleucine--tRNA ligase (EC 6.1.1.5) (Isoleucyl-tRNA synthetase) | 1175 | Class-I aminoacyl-tRNA synthetase family | 0 | 0 | 0 | Cytoplasm | | isoleucyl-tRNA aminoacylation [GO:0006428] | | aminoacyl-tRNA deacyla | aminoacyl-tR | UP000018817 |
| PPTG_02260 | W2RA63 | 259,97 | leucine--tRNA ligase (EC 6.1.1.4) (Leucyl-tRNA synthetase) | 1108 | Class-I aminoacyl-tRNA synthetase family | 0 | 0 | 0 | Cytoplasm | Nuclear localization signal | leucyl-tRNA aminoacylation [GO:0006429] | | aminoacyl-tRNA deacyla | aminoacyl-tR | UP000018817 |
| PPTG_14992 | W2PTN4 | 218,68 | tyrosine--tRNA ligase (EC 6.1.1.1) (Tyrosyl-tRNA synthetase) | 765 | Class-I aminoacyl-tRNA synthetase family | 0 | 0 | 0 | Cytoplasm | | tyrosyl-tRNA aminoacylati | cytosol [GO:00058 | ATP binding [GO:00552 | cytosol [GO:0 | UP000018817 |
| PPTG_17398 | W2PM22 | 221,83 | glutamate--tRNA ligase (EC 6.1.1.17) (Glutamyl-tRNA synthetase) | 876 | Class-I aminoacyl-tRNA synthetase family, Glutamate--tRNA li | 0 | 0 | 0 | Cytoplasm | Nuclear localization signal | glutamyl-tRNA aminoacyla | aminoacyl-tRNA sy | ATP binding [GO:00552 | aminoacyl-tR | UP000018817 |
| PPTG_13000 | W2Q5X9 | 201,52 | Dihydrolipoyl dehydrogenase (EC 1.8.1.4) | 387 | Class-I pyridine nucleotide-disulfide oxidoreductase family | 0 | 0 | 0 | Cytoplasm | Nuclear localization signal \| Nuclear export signal | 2-oxoglutarate metabolic | mitochondrion [GO | dihydrolipoyl dehydroge | mitochondrio | UP000018817 |
| PPTG_05022 | W2QVI7 | 174,07 | Aspartate aminotransferase (EC 2.6.1.1) | 426 | Class-I pyridoxal-phosphate-dependent aminotransferase fami | 0 | 0 | 0 | Mitochondrion | Mitochondrial transit peptide | amino acid metabolic pro | mitochondrion [GO | L-aspartate:2-oxoglutara | mitochondrio | UP000018817 |
| PPTG_00924 | W2RJB4 | 173,9 | glycine--tRNA ligase (EC 6.1.1.14) (Diadenosine tetraphosphate synthetase) | 681 | Class-II aminoacyl-tRNA synthetase family | 0 | 0 | 0 | Cytoplasm | | mitochondrial glycyl-tRNA | mitochondrion [GO | ATP binding [GO:00552 | mitochondrio | UP000018817 |
| PPTG_06869 | W2QT07 | 189,53 | Probable threonine--tRNA ligase, cytoplasmic (EC 6.1.1.3) (Threonyl-tRNA synthe | 744 | Class-II aminoacyl-tRNA synthetase family | 0 | 0 | 0 | Cytoplasm | | threonyl-tRNA aminoacyla | mitochondrion [GO | ATP binding [GO:00552 | mitochondrio | UP000018817 |
| PPTG_18707 | W2PGV0 | 222,51 | Alanine--tRNA ligase (EC 6.1.1.7) (Alanyl-tRNA synthetase) (AlaRS) | 994 | Class-II aminoacyl-tRNA synthetase family, Alax-L subfamily | 0 | 0 | 0 | Cytoplasm | Nuclear localization signal | mitochondrial alanyl-tRNA | mitochondrion [GO | alanine-tRNA ligase acti | mitochondrio | UP000018817 |
| PPTG_03926 | W2QZ71 | 186,29 | aspartate--tRNA ligase (EC 6.1.1.12) (Aspartyl-tRNA synthetase) | 514 | Class-II aminoacyl-tRNA synthetase family, Type 2 subfamily | 0 | 0 | 0 | Cytoplasm | Nuclear localization signal | aspartyl-tRNA aminoacyla | aminoacyl-tRNA sy | aspartate-tRNA ligase ac | aminoacyl-tR | UP000018817 |
| PPTG_14749 | W2PYE3 | 183,53 | serine--tRNA ligase (EC 6.1.1.11) (Seryl-tRNA synthetase) | 454 | Class-II aminoacyl-tRNA synthetase family, Type-1 seryl-tRNA s | 0 | 0 | 0 | Cytoplasm | Nuclear localization signal | seryl-tRNA aminoacylation [GO:0006434] | | ATP binding [GO:00552 | ATP binding [G | UP000018817 |
| PPTG_13850 | W2Q158 | 199,58 | fumarate hydratase (EC 4.2.1.2) | 528 | Class-II fumarase/aspartase family, Fumarase subfamily | 0 | 0 | 0 | Mitochondrion | Mitochondrial transit peptide | fumarate metabolic proce | mitochondrion [GO | fumarate hydratase acti | mitochondrio | UP000018817 |
| PPTG_11936 | W2Q8W4 | 317,04 | Clathrin heavy chain | 1719 | Clathrin heavy chain family | 0 | 0 | 0 | Cytoplasm | Nuclear localization signal | intracellular protein trans | clathrin coat of coa | clathrin light chain bindi | clathrin coat | UP000018817 |
| PPTG_13082 | W2Q6B1 | 177,78 | NADH dehydrogenase [ubiquinone] flavoprotein 2, mitochondrial | 271 | Complex I 24 kDa subunit family | 0 | 0 | 0 | Mitochondrion | Mitochondrial transit peptide | mitochondrial electron tra | catalytic complex [ | 2 iron, 2 sulfur cluster bi | catalytic comp | UP000018817 |
| PPTG_18619 | W2PIH3 | 200,3 | NADH dehydrogenase [ubiquinone] flavoprotein 1, mitochondrial (EC 7.1.1.2) | 499 | Complex I 51 kDa subunit family | 0 | 0 | 0 | Mitochondrion | Mitochondrial transit peptide | mitochondrial electron tra | mitochondrial inne | 4 iron, 4 sulfur cluster bi | mitochondrial | UP000018817 |
| PPTG_10119 | W2QD88 | 198,56 | Coatomer subunit gamma | 931 | COPG family | 0 | 0 | 0 | Cytoplasm | Nuclear export signal | endoplasmic reticulum to | COPI vesicle coat [ | structural molecule acti | COPI vesicle | UP000018817 |
| PPTG_16969 | W2PNS3 | 186,93 | Eukaryotic translation initiation factor 3 subunit M (eIF3m) | 369 | CSN7/EIF3M family, CSN7 subfamily; EIF-3 subunit M family | 0 | 0 | 0 | Cytoplasm | Nuclear localization signal | formation of cytoplasmic t | eukaryotic 43S prei | translation initiation fact | eukaryotic 43 | UP000018817 |
| PPTG_18479 | W2PGF2 | 179,8 | 6-phosphogluconolactonase | 360 | Cycloisomerase 2 family | 0 | 0 | 0 | Cytoplasm | Nuclear export signal | | | 6-phosphogluconolacto | 6-phosphoglu | UP000018817 |
| PPTG_19399 | W2PDE7 | 184,25 | Peptidyl-prolyl cis-trans isomerase (PPIase) (EC 5.2.1.8) | 171 | Cyclophilin-type PPIase family | 0 | 0 | 0 | Cytoplasm | | protein folding [GO:00064 | cytoplasm [GO:000 | cyclosporin A binding [G | cytoplasm [G | UP000018817 |
| PPTG_17946 | W2PIM1 | 199,8 | RNA helicase (EC 3.6.4.13) | 411 | DEAD box helicase family, eIF4A subfamily | 0 | 0 | 0 | Cytoplasm \| Nucleus | Nuclear localization signal \| Nuclear export signal | | | ATP binding [GO:00552 | ATP binding [G | UP000018817 |
| PPTG_14789 | W2PXW3 | 218,89 | AAA+ ATPase domain-containing protein | 4117 | Dynein heavy chain family | 0 | 0 | 0 | Cytoplasm | | microtubule-based moven | axoneme [GO:0005 | ATP binding [GO:00552 | axoneme [GO | UP000018817 |
| PPTG_18566 | W2PH71 | 182,21 | Cytoplasmic dynein 2 heavy chain 1 | 4396 | Dynein heavy chain family | 0 | 0 | 0 | Cytoplasm | | cilium assembly [GO:0060 | ciliary membrane [ | ATP binding [GO:00552 | ciliary membr | UP000018817 |

| | | | | | | | | | | | | | | | |
|---|---|---|---|---|---|---|---|---|---|---|---|---|---|---|---|
| PPTG_00340 | W2REP6 | 178,57 | Dynein gamma chain, flagellar outer arm | 4622 | Dynein heavy chain family | 0 | 0 | 0 | Cytoplasm | | microtubule-based moven | axonemal dynein cc | ATP binding [GO:00055 | axonemal dyn | UP000018817 |
| PPTG_05265 | W2QWF0 | 264,3 | Dynein heavy chain, cytoplasmic (Dynein heavy chain, cytosolic) | 4706 | Dynein heavy chain family | 0 | 0 | 0 | Cytoplasm | | cilium assembly [GO:0060 | axonemal dynein cc | ATP binding [GO:00055 | axonemal dyn | UP000018817 |
| PPTG_09098 | W2QG96 | 192,36 | Dynein-1, subspecies f | 4551 | Dynein heavy chain family | 0 | 0 | 1 | Cytoplasm | | cilium movement involved | inner dynein arm [G | ATP binding [GO:00055 | inner dynein a | UP000018817 |
| PPTG_11914 | W2QBD5 | 193,79 | Translation elongation factor EF1B beta/delta subunit guanine nucleotide exchan | 228 | EF-1-beta/EF-1-delta family | 0 | 0 | 0 | Cytoplasm | Nuclear localization signal | | cytosol [GO:000582 | guanyl-nucleotide excha | cytosol [GO:0 | UP000018817 |
| PPTG_16401 | W2PNJ7 | 229,83 | Eukaryotic translation initiation factor 3 subunit A (eIF3a) (Eukaryotic translation | 1146 | EIF-3 subunit A family | 0 | 0 | 0 | Cytoplasm \| Nucleus | Nuclear localization signal | formation of cytoplasmic t | eukaryotic 43S prei | mRNA binding [GO:0003 | eukaryotic 43 | UP000018817 |
| PPTG_10774 | W2QA19 | 213,3 | Eukaryotic translation initiation factor 3 subunit C (eIF3c) (Eukaryotic translation | 973 | EIF-3 subunit C family | 0 | 0 | 0 | Cytoplasm \| Nucleus | Nuclear localization signal | formation of cytoplasmic t | eukaryotic 43S prei | RNA binding [GO:000372 | eukaryotic 43 | UP000018817 |
| PPTG_11378 | W2QB69 | 176,65 | Eukaryotic translation initiation factor 3 subunit L (eIF3l) | 490 | EIF-3 subunit L family | 0 | 0 | 0 | Cytoplasm \| Nucleus | Nuclear localization signal | formation of cytoplasmic t | eukaryotic 43S prei | translation initiation fact | eukaryotic 43 | UP000018817 |
| PPTG_01466 | W2R9J1 | 210,13 | Eukaryotic translation initiation factor 5A (eIF-5A) | 161 | EIF-5A family | 0 | 0 | 0 | Cytoplasm \| Nucleus | Nuclear localization signal \| Nuclear export signal | positive regulation of trans | chloroplast [GO:00 | ribosome binding [GO:00 | chloroplast [G | UP000018817 |
| PPTG_09888 | W2QE91 | 203,74 | phosphopyruvate hydratase (EC 4.2.1.11) | 457 | Enolase family | 0 | 0 | 0 | Mitochondrion | Mitochondrial transit peptide | glycolytic process [GO:000 | phosphopyruvate hy | magnesium ion binding [ | phosphopyruv | UP000018817 |
| PPTG_20013 | W2Q2P7 | 246,11 | phosphopyruvate hydratase (EC 4.2.1.11) | 483 | Enolase family | 0 | 0 | 0 | Cytoplasm \| Nucleus | | glycolytic process [GO:000 | phosphopyruvate hy | magnesium ion binding [ | phosphopyruv | UP000018817 |
| PPTG_02894 | W2RFJ7 | 176,63 | Probable enoyl-CoA hydratase, mitochondrial (EC 4.2.1.17) | 279 | Enoyl-CoA hydratase/isomerase family | 0 | 0 | 0 | Mitochondrion | Mitochondrial transit peptide | fatty acid beta-oxidation [G | mitochondrion [GO: | enoyl-CoA hydratase act | mitochondrion | UP000018817 |
| PPTG_07368 | W2QPT4 | 234,9 | Trifunctional enzyme subunit alpha, mitochondrial (EC 1.1.1.211) (EC 4.2.1.17) (I | 747 | Enoyl-CoA hydratase/isomerase family; Enoyl-CoA hydratase/is | 0 | 0 | 0 | Mitochondrion | Mitochondrial transit peptide | fatty acid beta-oxidation [G | mitochondrial fatty | enoyl-CoA hydratase act | mitochondrial | UP000018817 |
| PPTG_00500 | W2RH96 | 180,74 | Electron transfer flavoprotein subunit beta (Beta-ETF) | 251 | ETF beta-subunit/FixA family | 0 | 0 | 0 | Cytoplasm | | carboxylic acid catabolic p | mitochondrial matr | electron transfer activity | mitochondrial | UP000018817 |
| PPTG_14608 | W2PXW1 | 198,27 | Electron transfer flavoprotein-ubiquinone oxidoreductase (ETF-QO) (EC 1.5.5.1) | 610 | ETF-QO/FixC family | 0 | 0 | 0 | Mitochondrion | Mitochondrial transit peptide | | mitochondrial inner | 4 iron, 4 sulfur cluster bir | mitochondrial | UP000018817 |
| PPTG_18870 | W2PFJ5 | 210,31 | ATP synthase subunit b | 308 | Eukaryotic ATPase B chain family | 0 | 0 | 0 | Mitochondrion | Mitochondrial transit peptide | proton motive force-driven | mitochondrial inner | proton transmembrane t | mitochondrial | UP000018817 |
| PPTG_15631 | W2PTN3 | 177,78 | Voltage-dependent anion-selective channel protein | 282 | Eukaryotic mitochondrial porin family | 0 | 0 | 0 | Cytoplasm | | | mitochondrial outer | porin activity [GO:00152 | mitochondrial | UP000018817 |
| PPTG_13499 | W2Q1L6 | 179,62 | 60S ribosomal protein L6 | 233 | Eukaryotic ribosomal protein eL6 family | 0 | 0 | 0 | Cytoplasm | Nuclear localization signal | cytoplasmic translation [G | chloroplast [GO:00 | RNA binding [GO:000372 | chloroplast [G | UP000018817 |
| PPTG_16072 | W2PQA7 | 185,23 | 60S ribosomal protein L6 | 204 | Eukaryotic ribosomal protein eL6 family | 0 | 0 | 0 | Cytoplasm | Nuclear localization signal | cytoplasmic translation [G | chloroplast [GO:00 | RNA binding [GO:000372 | chloroplast [G | UP000018817 |
| PPTG_14734 | W2PXM6 | 202,3 | 60S ribosomal protein L7a | 263 | Eukaryotic ribosomal protein eL8 family | 0 | 0 | 0 | Cytoplasm | Nuclear localization signal | ribosome biogenesis [GO: | cytosolic large ribos | RNA binding [GO:000372 | cytosolic large | UP000018817 |
| PPTG_07764 | W2QMB5 | 201,85 | Small ribosomal subunit protein eS1 | 261 | Eukaryotic ribosomal protein eS1 family | 0 | 0 | 0 | Cytoplasm | Nuclear localization signal | translation [GO:0006412] | cytosolic small ribo | structural constituent of | cytosolic sma | UP000018817 |
| PPTG_15134 | W2PW69 | 183,55 | 40S ribosomal protein S12 | 175 | Eukaryotic ribosomal protein eS12 family | 0 | 0 | 0 | Cytoplasm | | translation [GO:0006412] | ribonucleoprotein c | structural constituent of | ribonucleopro | UP000018817 |
| PPTG_07085 | W2QP05 | 175,88 | 40S ribosomal protein S17 | 131 | Eukaryotic ribosomal protein eS17 family | 0 | 0 | 0 | Cytoplasm | Nuclear localization signal | translation [GO:0006412] | cytosol [GO:000582 | structural constituent of | cytosol [GO:0 | UP000018817 |
| PPTG_10646 | W2QBJ2 | 179,75 | 40S ribosomal protein S4 | 261 | Eukaryotic ribosomal protein eS4 family | 0 | 0 | 0 | Cytoplasm | Nuclear localization signal | translation [GO:0006412] | cytosolic small ribo | rRNA binding [GO:00198 | cytosolic sma | UP000018817 |
| PPTG_15783 | W2PPP1 | 196,89 | 40S ribosomal protein S7 | 190 | Eukaryotic ribosomal protein eS7 family | 0 | 0 | 0 | Cytoplasm | | ribosomal small subunit bi | 90S preribosome [G | structural constituent of | 90S preriboso | UP000018817 |
| PPTG_18213 | W2PJ01 | 176,14 | Importin N-terminal domain-containing protein | 1076 | Exportin family | 0 | 0 | 0 | Cytoplasm | Nuclear export signal | protein export from nucleu | cytoplasm [GO:000 | nuclear export signal rec | cytoplasm [G | UP000018817 |
| PPTG_11055 | W2Q752 | 187,06 | Glycerol-3-phosphate dehydrogenase (EC 1.1.5.3) | 615 | FAD-dependent glycerol-3-phosphate dehydrogenase family | 0 | 0 | 0 | Mitochondrion | Mitochondrial transit peptide | glycerol-3-phosphate meta | mitochondrion [GO: | glycerol-3-phosphate del | mitochondrion | UP000018817 |
| PPTG_00150 | W2RDR6 | 214,07 | Succinate dehydrogenase [ubiquinone] flavoprotein subunit, mitochondrial (EC 1 | 645 | FAD-dependent oxidoreductase 2 family, FRD/SDH subfamily | 0 | 0 | 0 | Mitochondrion | Mitochondrial transit peptide | mitochondrial electron tra | mitochondrial inner | electron transfer activity | mitochondrial | UP000018817 |
| PPTG_00390 | W2REP0 | 181,96 | FAD/NAD(P)-binding domain-containing protein | 559 | FAD-dependent oxidoreductase family | 0 | 0 | 0 | Mitochondrion | Mitochondrial transit peptide | mitochondrial respiratory | mitochondrion [GO: | FAD binding [GO:007194 | mitochondrion | UP000018817 |
| PPTG_11916 | W2Q8U0 | 201,43 | Fructose-1,6-bisphosphatase, cytosolic (EC 3.1.3.11) | 333 | FBPase class 1 family | 0 | 0 | 0 | Cytoplasm \| Nucleus | Nuclear localization signal \| Nuclear export signal | fructose 1,6-bisphosphate | cytosol [GO:000582 | fructose 1,6-bisphospha | cytosol [GO:0 | UP000018817 |
| PPTG_02874 | W2RCX3 | 208,72 | glycerol kinase (EC 2.7.1.30) (ATP:glycerol 3-phosphotransferase) | 552 | FGGY kinase family | 0 | 0 | 0 | Mitochondrion | Mitochondrial transit peptide | glycerol catabolic process | cytosol [GO:000582 | ATP binding [GO:00055 | cytosol [GO:0 | UP000018817 |
| PPTG_04052 | W2QZP3 | 183,32 | Trifunctional purine biosynthetic protein adenosine-3 (EC 2.1.2.2) (EC 6.3.3.1) (E | 1143 | GART family; GARS family; AIR synthase family | 0 | 0 | 0 | Cytoplasm | | 'de novo' IMP biosynthetic | cytosol [GO:000582 | ATP binding [GO:00055 | cytosol [GO:0 | UP000018817 |
| PPTG_02536 | W2RDB0 | 202,07 | TOG domain-containing protein | 2759 | GCN1 family | 0 | 0 | 0 | Cytoplasm | Nuclear localization signal | cellular response to amino | cytosol [GO:000582 | protein kinase regulator | cytosol [GO:0 | UP000018817 |
| PPTG_03132 | W2R452 | 199,92 | Glutamate/phenylalanine/leucine/valine/L-tryptophan dehydrogenase C-termina | 1032 | Glu/Leu/Phe/Val dehydrogenases family | 0 | 0 | 0 | Cytoplasm | Nuclear localization signal | L-glutamate catabolic proc | mitochondrion [GO: | glutamate dehydrogenas | mitochondrion | UP000018817 |
| PPTG_06379 | W2QUL0 | 218,56 | Glutamate/phenylalanine/leucine/valine/L-tryptophan dehydrogenase C-termina | 1057 | Glu/Leu/Phe/Val dehydrogenases family | 0 | 0 | 0 | Mitochondrion | Mitochondrial transit peptide | L-glutamate catabolic proc | mitochondrion [GO: | glutamate dehydrogenas | mitochondrion | UP000018817 |
| PPTG_09495 | W2QF83 | 178,94 | Glutamate dehydrogenase | 494 | Glu/Leu/Phe/Val dehydrogenases family | 0 | 0 | 0 | Mitochondrion | Mitochondrial transit peptide | glutamate biosynthetic pro | cytosol [GO:000582 | glutamate dehydrogenas | cytosol [GO:0 | UP000018817 |
| PPTG_16811 | W2PQ87 | 183,27 | glucose-6-phosphate 1-epimerase (EC 5.1.3.15) | 300 | Glucose-6-phosphate 1-epimerase family | 0 | 0 | 0 | Cytoplasm | Nuclear localization signal | carbohydrate metabolic pr | cytoplasm [GO:000 | carbohydrate binding [G | cytoplasm [G | UP000018817 |
| PPTG_15102 | W2PW23 | 204,81 | Glucose-6-phosphate 1-dehydrogenase (EC 1.1.1.49) | 552 | Glucose-6-phosphate dehydrogenase family | 0 | 0 | 0 | Cytoplasm | Peroxisomal targeting signal | glucose metabolic process [GO:0006006]; pen | | glucose-6-phosphate del | glucose-6-pho | UP000018817 |
| PPTG_03240 | W2R6M4 | 193,16 | glutamate synthase (ferredoxin) (EC 1.4.7.1) | 1582 | Glutamate synthase family | 0 | 0 | 0 | Mitochondrion | Mitochondrial transit peptide | ammonia assimilation cycle [GO:0019676]; glu | | 3 iron, 4 sulfur cluster bir | 3 iron, 4 sulfur | UP000018817 |
| PPTG_09874 | W2QD82 | 173,98 | Glutamine synthetase (EC 6.3.1.2) | 402 | Glutamine synthetase family | 0 | 0 | 0 | Cytoplasm | | glutamine biosynthetic pro | cytoplasm [GO:000 | ATP binding [GO:00055 | cytoplasm [G | UP000018817 |
| PPTG_00110 | W2RG48 | 219,22 | Glutathione peroxidase | 561 | Glutathione peroxidase family | 0 | 0 | 0 | Cytoplasm | Nuclear localization signal | response to oxidative stress [GO:0006979] | | peroxidase activity [GO: | peroxidase ac | UP000018817 |
| PPTG_01098 | W2RHS3 | 224,63 | Glyceraldehyde-3-phosphate dehydrogenase (EC 1.2.1.12) | 370 | Glyceraldehyde-3-phosphate dehydrogenase family | 0 | 0 | 0 | Cytoplasm | | glucose metabolic process | cytosol [GO:000582 | glyceraldehyde-3-phospl | cytosol [GO:0 | UP000018817 |
| PPTG_01358 | W2R6Z9 | 182,84 | Glyceraldehyde-3-phosphate dehydrogenase, type I | 524 | Glyceraldehyde-3-phosphate dehydrogenase family | 0 | 0 | 0 | Cytoplasm | Peroxisomal targeting signal | | | NAD binding [GO:00512 | NAD binding [ | UP000018817 |
| PPTG_01022 | W2RHI5 | 208,62 | Glucose-6-phosphate isomerase (EC 5.3.1.9) | 556 | GPI family | 0 | 0 | 0 | Cytoplasm | Nuclear localization signal | gluconeogenesis [GO:000 | cytosol [GO:000582 | carbohydrate derivative | cytosol [GO:0 | UP000018817 |
| PPTG_05337 | W2QZ60 | 173,53 | Glutamate decarboxylase (EC 4.1.1.15) | 493 | Group II decarboxylase family | 0 | 0 | 0 | Cytoplasm \| Nucleus | Nuclear localization signal \| Nuclear export signal | L-glutamate catabolic proc | cytosol [GO:000582 | glutamate decarboxylase | cytosol [GO:0 | UP000018817 |
| PPTG_00631 | W2RI00 | 175,33 | Chaperone DnaJ | 418 | Heat shock protein 40 family | 0 | 0 | 0 | Cytoplasm \| Nucleus | Nuclear localization signal \| Nuclear export signal | protein folding [GO:0006457]; response to hea | | ATP binding [GO:00055 | ATP binding [G | UP000018817 |
| PPTG_18274 | W2PGE0 | 202,99 | Heat shock chaperone dnaK | 648 | Heat shock protein 70 family | 0 | 0 | 0 | Mitochondrion | Mitochondrial transit peptide | | | ATP binding [GO:00055 | ATP binding [G | UP000018817 |
| PPTG_03121 | W2R438 | 241,12 | Heat shock chaperone DnaK | 786 | Heat shock protein 70 family | 0 | 0 | 0 | Cytoplasm \| Nucleus | Nuclear localization signal | | | ATP binding [GO:00055 | ATP binding [G | UP000018817 |
| PPTG_02122 | W2R9I2 | 254,77 | Hsp70-like protein | 657 | Heat shock protein 70 family | 0 | 0 | 0 | Cytoplasm | Nuclear export signal | | | ATP binding [GO:00055 | ATP binding [G | UP000018817 |
| PPTG_19375 | W2PCW7 | 246,52 | Heat shock protein 70 | 809 | Heat shock protein 70 family | 0 | 0 | 0 | Cytoplasm | Nuclear localization signal | | cytosol [GO:000582 | ATP binding [GO:00055 | cytosol [GO:0 | UP000018817 |
| PPTG_03165 | W2R490 | 218,68 | Chaperonin GroL | 598 | Heat shock protein 70 family-Chaperonin | 0 | 0 | 0 | Mitochondrion | Mitochondrial transit peptide | protein refolding [GO:0042026] | | ATP binding [GO:00055 | ATP binding [G | UP000018817 |
| PPTG_19188 | W2PE41 | 233,36 | Heat shock protein 90-2 | 706 | Heat shock protein 90 family | 0 | 0 | 0 | Cytoplasm | | | cytoplasm [GO:000 | ATP binding [GO:00055 | cytoplasm [G | UP000018817 |
| PPTG_14687 | W2PY60 | 188,39 | Homoserine dehydrogenase (EC 1.1.1.3) | 436 | Homoserine dehydrogenase family | 0 | 0 | 0 | Cytoplasm | | methionine biosynthetic process [GO:0009086 | | homoserine dehydrogen | homoserine d | UP000018817 |
| PPTG_00991 | W2RHE9 | 177,73 | dihydroxy-acid dehydratase (EC 4.2.1.9) | 596 | IlvD/Edd family | 0 | 0 | 0 | Mitochondrion | Mitochondrial transit peptide | isoleucine biosynthetic process [GO:0009097] | | 2 iron, 2 sulfur cluster bir | 2 iron, 2 sulfur | UP000018817 |
| PPTG_16510 | W2PN09 | 190,79 | Inosine-5'-monophosphate dehydrogenase (IMP dehydrogenase) (IMPD) (IMPDH | 508 | IMPDH/GMPR family | 0 | 0 | 0 | Cytoplasm \| Nucleus | Nuclear localization signal | GMP biosynthetic process | cytoplasm [GO:000 | IMP dehydrogenase activ | cytoplasm [G | UP000018817 |
| PPTG_02597 | W2RBQ4 | 180,18 | Importin N-terminal domain-containing protein | 858 | Importin beta family, Importin beta-1 subfamily | 0 | 0 | 0 | Cytoplasm \| Nucleus | Nuclear localization signal \| Nuclear export signal | protein import into nucleus | cytoplasm [GO:000 | small GTPase binding [G | cytoplasm [G | UP000018817 |
| PPTG_04112 | W2QZ92 | 185,29 | Superoxide dismutase (EC 1.15.1.1) | 250 | Iron/manganese superoxide dismutase family | 0 | 0 | 0 | Cytoplasm | Mitochondrial transit peptide | | cytoplasm [GO:000 | metal ion binding [GO:00 | cytoplasm [G | UP000018817 |
| PPTG_04350 | W2R0C6 | 215,23 | Isocitrate dehydrogenase [NADP] (EC 1.1.1.42) | 427 | Isocitrate and isopropylmalate dehydrogenases family | 0 | 0 | 0 | Mitochondrion | Mitochondrial transit peptide | glyoxylate cycle [GO:0006 | mitochondrion [GO: | isocitrate dehydrogenas | mitochondrion | UP000018817 |
| PPTG_16504 | W2PNQ9 | 194,8 | Isocitrate lyase | 537 | Isocitrate lyase/PEP mutase superfamily, Isocitrate lyase family | 0 | 0 | 0 | Cytoplasm | Peroxisomal targeting signal | glyoxylate cycle [GO:0006097]; tricarboxylic ac | | isocitrate lyase activity [ | isocitrate lyas | UP000018817 |
| PPTG_15943 | W2PU40 | 202,64 | Ketol-acid reductoisomerase, chloroplastic (EC 1.1.1.86) (Acetohydroxy-acid red | 515 | Ketol-acid reductoisomerase family | 0 | 0 | 0 | Mitochondrion | Mitochondrial transit peptide | isoleucine biosynthetic pro | chloroplast [GO:00 | isomerase activity [GO:0 | chloroplast [G | UP000018817 |
| PPTG_10528 | W2QC09 | 229,48 | Malate dehydrogenase (EC 1.1.1.37) | 335 | LDH/MDH superfamily, MDH type 1 family | 0 | 0 | 0 | Mitochondrion | Mitochondrial transit peptide | malate metabolic process | cytoplasm [GO:000 | L-malate dehydrogenase | cytoplasm [G | UP000018817 |
| PPTG_20398 | W2PBA3 | 202,78 | Malate synthase (EC 2.3.3.9) | 535 | Malate synthase family | 0 | 0 | 0 | Cytoplasm | Peroxisomal targeting signal | glyoxylate cycle [GO:0006 | cytoplasm [GO:000 | malate synthase activity | cytoplasm [G | UP000018817 |
| PPTG_14774 | W2PYI8 | 203,06 | Malate synthase (EC 2.3.3.9) | 535 | Malate synthase family | 0 | 0 | 0 | Cytoplasm | Peroxisomal targeting signal | glyoxylate cycle [GO:0006 | cytoplasm [GO:000 | malate synthase activity | cytoplasm [G | UP000018817 |
| PPTG_19127 | W2PDR1 | 202,21 | Mitochondrial phosphate carrier protein | 345 | Mitochondrial carrier (TC 2.A.29) family | 0 | 0 | 0 | Mitochondrion | | mitochondrial phosphate i | mitochondrial inner | phosphate transmembra | mitochondrial | UP000018817 |
| PPTG_00853 | W2RGP1 | 201,96 | inositol-3-phosphate synthase (EC 5.5.1.4) | 517 | Myo-inositol 1-phosphate synthase family | 0 | 0 | 0 | Cytoplasm | Peroxisomal targeting signal | inositol biosynthetic proce | cytoplasm [GO:000 | inositol-3-phosphate syn | cytoplasm [G | UP000018817 |
| PPTG_18330 | W2PII3 | 173,71 | Nucleosome assembly protein 1-like 1 | 368 | Nucleosome assembly protein (NAP) family | 0 | 0 | 0 | Cytoplasm | Nuclear localization signal | nucleosome assembly [GO | nucleus [GO:0005634] | | nucleus [GO: | UP000018817 |
| PPTG_05088 | W2QVT6 | 183,61 | 5-oxoprolinase | 1292 | Oxoprolinase family | 0 | 0 | 0 | Cytoplasm | Nuclear export signal | glutathione metabolic proc | cytosol [GO:000582 | 5-oxoprolinase (ATP-hyd | cytosol [GO:0 | UP000018817 |
| PPTG_06377 | W2QSL7 | 176,75 | Aminopeptidase (EC 3.4.11.-) | 902 | Peptidase M1 family | 0 | 0 | 0 | Cytoplasm | Nuclear localization signal \| Nuclear export signal | peptide catabolic process | cytoplasm [GO:000 | metalloaminopeptidase | cytoplasm [G | UP000018817 |
| PPTG_15972 | W2PS84 | 193,17 | Aminopeptidase (EC 3.4.11.-) | 884 | Peptidase M1 family | 0 | 0 | 0 | Cytoplasm | Nuclear localization signal \| Nuclear export signal | peptide catabolic process | cytoplasm [GO:000 | metalloaminopeptidase | cytoplasm [G | UP000018817 |
| PPTG_10972 | W2QCT9 | 210,97 | mitochondrial processing peptidase (EC 3.4.24.64) (Beta-MPP) | 466 | Peptidase M16 family | 0 | 0 | 0 | Mitochondrion | Mitochondrial transit peptide | proteolysis [GO:0006508] | mitochondrial matr | metal ion binding [GO:00 | mitochondrial | UP000018817 |
| PPTG_04750 | W2R256 | 240,21 | Peptidase M16 N-terminal domain-containing protein | 447 | Peptidase M16 family | 0 | 0 | 0 | Mitochondrion | Mitochondrial transit peptide | | membrane [GO:001 | metal ion binding [GO:00 | membrane [G | UP000018817 |
| PPTG_14635 | W2PVE4 | 203,71 | aspartyl aminopeptidase (EC 3.4.11.21) | 462 | Peptidase M18 family | 0 | 0 | 0 | Cytoplasm \| Nucleus | | proteolysis [GO:0006508] | cytoplasm [GO:000 | aminopeptidase activity | cytoplasm [G | UP000018817 |
| PPTG_16899 | W2PMI6 | 197,65 | DNA-binding protein, 42 kDa | 393 | Peptidase M24 family | 0 | 0 | 0 | Cytoplasm | Nuclear localization signal | | | DNA binding [GO:000367 | DNA binding [ | UP000018817 |
| PPTG_02295 | W2RAT4 | 174,22 | MPN domain-containing protein | 324 | Peptidase M67A family | 0 | 0 | 0 | Cytoplasm | Nuclear export signal | proteasome-mediated ubi | proteasome regulat | metallopeptidase activit | proteasome r | UP000018817 |
| PPTG_14406 | W2PU97 | 214,11 | PDZ domain-containing protein | 1050 | Peptidase S1C family | 0 | 0 | 0 | Mitochondrion | Mitochondrial transit peptide | proteolysis [GO:0006508] | | serine-type endopeptida | serine-type er | UP000018817 |
| PPTG_07472 | W2QN94 | 198,57 | Uncharacterized protein | 1001 | Peptidase S8 family | 0 | 0 | 0 | Cytoplasm | Nuclear localization signal | proteolysis [GO:0006508] | cytosol [GO:000582 | aminopeptidase activity | cytosol [GO:0 | UP000018817 |
| PPTG_13924 | W2Q0S7 | 180,6 | Prolyl endopeptidase (EC 3.4.21.-) | 752 | Peptidase S9A family | 0 | 0 | 0 | Cytoplasm | | proteolysis [GO:0006508] | cytosol [GO:000582 | oligopeptidase activity [ | cytosol [GO:0 | UP000018817 |
| PPTG_03886 | W2R0F4 | 173,05 | Proteasome subunit alpha type | 244 | Peptidase T1A family | 0 | 0 | 0 | Cytoplasm \| Nucleus | Nuclear export signal | proteasome-mediated ubi | cytosol [GO:0005829]; nucleus [GO:0005634 | | cytosol [GO:0 | UP000018817 |
| PPTG_17524 | W2PKD5 | 191,22 | Thioredoxin domain-containing protein | 377 | Peroxiredoxin family, AhpC/Prx1 subfamily | 0 | 0 | 0 | Mitochondrion | Mitochondrial transit peptide | cell redox homeostasis [G | cytosol [GO:000582 | thioredoxin peroxidase a | cytosol [GO:0 | UP000018817 |
| PPTG_08167 | W2QJJ1 | 173,06 | thioredoxin-dependent peroxiredoxin (EC 1.11.1.24) (Nuclear thiol peroxidase) (T | 367 | Peroxiredoxin family, BCP/PrxQ subfamily | 0 | 0 | 0 | Nucleus | | cell redox homeostasis [G | cytoplasm [GO:000 | thioredoxin peroxidase a | cytoplasm [G | UP000018817 |
| PPTG_14664 | W2PWA6 | 190,72 | Phosphatase PP2A regulatory subunit A/Splicing factor 3B subunit 1-like HEAT rep | 457 | Phosphatase 2A regulatory subunit A family | 0 | 0 | 0 | Cytoplasm \| Nucleus | Nuclear export signal | | cytosol [GO:000582 | protein phosphatase reg | cytosol [GO:0 | UP000018817 |
| PPTG_01922 | W2RAY1 | 198,34 | phosphoenolpyruvate carboxykinase (ATP) (EC 4.1.1.49) | 569 | Phosphoenolpyruvate carboxykinase (ATP) family | 0 | 0 | 0 | Mitochondrion | Mitochondrial transit peptide | gluconeogenesis [GO:000 | cytosol [GO:000582 | ATP binding [GO:00055 | cytosol [GO:0 | UP000018817 |
| PPTG_03714 | W2R7Y3 | 249,4 | Phosphoglycerate kinase (EC 2.7.2.3) | 414 | Phosphoglycerate kinase family | 0 | 0 | 0 | Cytoplasm | Nuclear localization signal | gluconeogenesis [GO:000 | cytosol [GO:000582 | ADP binding [GO:004353 | cytosol [GO:0 | UP000018817 |
| PPTG_07760 | W2QLN1 | 174,2 | Phosphoacetylglucosamine mutase (PAGM) (EC 5.4.2.3) (Acetylglucosamine pho | 561 | Phosphohexose mutase family | 0 | 0 | 0 | Cytoplasm | Nuclear localization signal | carbohydrate metabolic process [GO:0005975 | | metal ion binding [GO:00 | metal ion bind | UP000018817 |
| PPTG_13491 | W2Q1K6 | 190,08 | Phospholipase (EC 3.1.4.4) | 1117 | Phospholipase D family | 0 | 0 | 0 | Cytoplasm \| Lysosome/Vacuole | | intracellular signal transdu | plasma membrane | phospholipase D activity | plasma memb | UP000018817 |
| PPTG_01563 | W2R7Y2 | 206,19 | Polyadenylate-binding protein (PABP) | 640 | Polyadenylate-binding protein type-1 family | 0 | 0 | 0 | Cytoplasm \| Nucleus | Nuclear localization signal | mRNA processing [GO:000 | cytoplasm [GO:000 | RNA binding [GO:000372 | cytoplasm [G | UP000018817 |
| PPTG_12038 | W2Q6J8 | 174,88 | Serine/threonine-protein phosphatase (EC 3.1.3.16) | 319 | PPP phosphatase family | 0 | 0 | 0 | Cytoplasm \| Nucleus | Nuclear export signal | | | metal ion binding [GO:00 | metal ion bind | UP000018817 |
| PPTG_09527 | W2QG60 | 217,72 | Uncharacterized protein | 997 | Proteasome subunit S1 family | 0 | 0 | 0 | Cytoplasm | | proteasome-mediated ubi | nucleus [GO:00056 | enzyme regulator activity | nucleus [GO: | UP000018817 |
| PPTG_16668 | W2PPP5 | 175,42 | PCI domain-containing protein | 383 | Proteasome subunit S11 family | 0 | 0 | 0 | Cytoplasm \| Nucleus | Nuclear export signal | ubiquitin-dependent prote | cytosol [GO:000582 | structural molecule activ | cytosol [GO:0 | UP000018817 |

| | | | | | | | | | | | | | | | |
|---|---|---|---|---|---|---|---|---|---|---|---|---|---|---|---|
| PPTG_08401 | W2QKH4 | 181,16 | PCI domain-containing protein | 264 | Proteasome subunit S14 family | 0 | 0 | 0 | Nucleus | Nuclear export signal | proteasome-mediated ubi | cytosol [GO:0005829]; nucleus [GO:0005634 | | cytosol [GO:0 | UP000018817 |
| PPTG_14544 | W2PXK9 | 257,61 | 26S proteasome non-ATPase regulatory subunit 2 | 897 | Proteasome subunit S2 family | 0 | 0 | 0 | Cytoplasm | Nuclear export signal | proteasome-mediated ubi | nucleus [GO:00056 | enzyme regulator activity | nucleus [GO:0 | UP000018817 |
| PPTG_02400 | W2RB59 | 184,88 | protein disulfide-isomerase (EC 5.3.4.1) | 353 | Protein disulfide isomerase family | 0 | 0 | 0 | Endoplasmic reticulum | Signal peptide | protein folding [GO:00064 | endoplasmic reticu | protein disulfide isomera | endoplasmic r | UP000018817 |
| PPTG_09222 | W2QGU1 | 182,13 | CAMK/CAMK1 protein kinase | 385 | Protein kinase superfamily, Ser/Thr protein kinase family, CDPK | 0 | 0 | 0 | Cytoplasm | | glycogen metabolic proces | phosphorylase kina | ATP binding [GO:000552 | phosphorylase | UP000018817 |
| PPTG_04024 | W2QYX2 | 179,07 | CAMK/CDPK protein kinase | 555 | Protein kinase superfamily, Ser/Thr protein kinase family, CDPK | 0 | 0 | 0 | Cytoplasm \| Lysosome/Vacuole | | | | ATP binding [GO:000552 | ATP binding [G | UP000018817 |
| PPTG_08188 | W2QLD0 | 226,48 | Phosphoribosylaminoimidazolecarboxamide formyltransferase/IMP cyclohydrol | 616 | PurH family | 0 | 0 | 0 | Cytoplasm | Nuclear localization signal | 'de novo' IMP biosynthetic | cytosol [GO:000582 | IMP cyclohydrolase activ | cytosol [GO:0 | UP000018817 |
| PPTG_04949 | W2R387 | 176,34 | Pyrroline-5-carboxylate reductase catalytic N-terminal domain-containing protei | 272 | Pyrroline-5-carboxylate reductase family | 0 | 0 | 0 | Cytoplasm | Peroxisomal targeting signal | L-proline biosynthetic process [GO:0055129] | | pyrroline-5-carboxylate r | pyrroline-5-ca | UP000018817 |
| PPTG_03709 | W2R8F7 | 177,1 | Pyruvate kinase (EC 2.7.1.40) | 505 | Pyruvate kinase family | 0 | 0 | 0 | Cytoplasm | Nuclear localization signal \| Nuclear export signal | response to stress [GO:0006950] | | ATP binding [GO:000552 | ATP binding [G | UP000018817 |
| PPTG_04978 | W2R3G8 | 206,69 | Rab GDP dissociation inhibitor | 462 | Rab GDI family | 0 | 0 | 0 | Cytoplasm | Nuclear localization signal | protein transport [GO:001 | cytoplasm [GO:000 | Rab GDP-dissociation in | cytoplasm [G | UP000018817 |
| PPTG_11194 | W2Q9D3 | 203,95 | EF-hand domain-containing protein | 2480 | Recoverin family | 0 | 0 | 0 | Cytoplasm | Nuclear localization signal | | | calcium ion binding [GO: | calcium ion bi | UP000018817 |
| PPTG_02839 | W2RCP4 | 173,95 | Replication protein A subunit | 628 | Replication factor A protein 1 family | 0 | 0 | 0 | Cytoplasm \| Nucleus | Nuclear localization signal | DNA recombination [GO:0 | nucleus [GO:00056 | DNA binding [GO:000367 | nucleus [GO:0 | UP000018817 |
| PPTG_15878 | W2PSD0 | 175,8 | Ribonucleoside-diphosphate reductase (EC 1.17.4.1) | 792 | Ribonucleoside diphosphate reductase large chain family | 0 | 0 | 0 | Cytoplasm | | deoxyribonucleotide biosy | ribonucleoside-dipl | ATP binding [GO:000552 | ribonucleosid | UP000018817 |
| PPTG_02716 | W2RC68 | 178,92 | ribose-phosphate diphosphokinase (EC 2.7.6.1) | 475 | Ribose-phosphate pyrophosphokinase family | 0 | 0 | 0 | Mitochondrion | Mitochondrial transit peptide | 5-phosphoribose 1-diphos | cytoplasm [GO:000 | ATP binding [GO:000552 | cytoplasm [G | UP000018817 |
| PPTG_00284 | W2REE6 | 187,99 | Cytochrome b-c1 complex subunit Rieske, mitochondrial (EC 7.1.1.8) | 215 | Rieske iron-sulfur protein family | 0 | 0 | 0 | Mitochondrion | Mitochondrial transit peptide | | mitochondrial inner | 2 iron, 2 sulfur cluster bir | mitochondrial | UP000018817 |
| PPTG_12936 | W2Q187 | 182,58 | Protein transport protein SEC23 | 861 | SEC23/SEC24 family, SEC23 subfamily | 0 | 0 | 0 | Cytoplasm | Nuclear localization signal | COPII-coated vesicle carg | COPII vesicle coat [ | GTPase activator activity | COPII vesicle | UP000018817 |
| PPTG_12256 | W2Q886 | 178,16 | Protein transporter Sec24 | 1119 | SEC23/SEC24 family, SEC24 subfamily | 0 | 0 | 0 | Cytoplasm | Nuclear localization signal | COPII-coated vesicle carg | COPII vesicle coat [ | SNARE binding [GO:000 | COPII vesicle | UP000018817 |
| PPTG_13257 | W2Q066 | 186,83 | Serine hydroxymethyltransferase (EC 2.1.2.1) | 502 | SHMT family | 0 | 0 | 0 | Mitochondrion | Mitochondrial transit peptide | glycine biosynthetic proce | mitochondrion [GO | glycine hydroxymethyltra | mitochondrion | UP000018817 |
| PPTG_08622 | W2QL97 | 210,16 | Serine hydroxymethyltransferase (EC 2.1.2.1) | 464 | SHMT family | 0 | 0 | 0 | Cytoplasm | Nuclear localization signal \| Nuclear export signal | glycine biosynthetic proce | mitochondrion [GO | glycine hydroxymethyltra | mitochondrion | UP000018817 |
| PPTG_20006 | W2P9U9 | 190,44 | Uncharacterized protein | 293 | SNAP family | 0 | 0 | 0 | Cytoplasm \| Nucleus | Nuclear localization signal \| Nuclear export signal | intracellular protein transp | SNARE complex [G | soluble NSF attachment | SNARE compl | UP000018817 |
| PPTG_09519 | W2QG53 | 223,85 | ATP-citrate synthase (EC 2.3.3.8) (ATP-citrate (pro-S-)-lyase) (Citrate cleavage er | 1105 | Succinate/malate CoA ligase alpha subunit family; Succinate/n | 0 | 0 | 0 | Cytoplasm | | acetyl-CoA biosynthetic pr | cytosol [GO:000582 | ATP binding [GO:000552 | cytosol [GO:0 | UP000018817 |
| PPTG_06269 | W2QSE4 | 186,34 | Succinate--CoA ligase [ADP-forming] subunit beta, mitochondrial (EC 6.2.1.5) (Su | 437 | Succinate/malate CoA ligase beta subunit family | 0 | 0 | 0 | Mitochondrion | Mitochondrial transit peptide | succinyl-CoA metabolic pr | mitochondrion [GO | ATP binding [GO:000552 | mitochondrion | UP000018817 |
| PPTG_07737 | W2QLK1 | 177,83 | CCT-beta | 526 | TCP-1 chaperonin family | 0 | 0 | 0 | Cytoplasm | Nuclear localization signal | | chaperonin-contain | ATP binding [GO:000552 | chaperonin-co | UP000018817 |
| PPTG_05308 | W2QYJ3 | 172,87 | T-complex protein 1 subunit alpha (CCT-alpha) | 546 | TCP-1 chaperonin family | 0 | 0 | 0 | Cytoplasm | Nuclear localization signal \| Nuclear export signal | | cytoplasm [GO:000 | ATP binding [GO:000552 | cytoplasm [G | UP000018817 |
| PPTG_18230 | W2PJ23 | 177,15 | T-complex protein 1 subunit delta | 533 | TCP-1 chaperonin family | 0 | 0 | 0 | Cytoplasm | Nuclear export signal | | cytoplasm [GO:000 | ATP binding [GO:000552 | cytoplasm [G | UP000018817 |
| PPTG_04240 | W2R2K0 | 219,04 | 3-ketoacyl-CoA thiolase | 397 | Thiolase-like superfamily, Thiolase family | 0 | 0 | 0 | Cytoplasm \| Mitochondrion | Mitochondrial transit peptide | fatty acid beta-oxidation [G | mitochondrion [GO | acetyl-CoA C-acetyltrans | mitochondrion | UP000018817 |
| PPTG_15137 | W2PU56 | 209,12 | Dynamin GTPase | 705 | TRAFAC class dynamin-like GTPase superfamily, Dynamin/Fzo/ | 0 | 0 | 0 | Cytoplasm | | | cytoplasm [GO:000 | GTP binding [GO:000552 | cytoplasm [G | UP000018817 |
| PPTG_15793 | W2PPD3 | 199,89 | GB1/RHD3-type G domain-containing protein | 638 | TRAFAC class dynamin-like GTPase superfamily, GB1/RHD3 GT | 0 | 0 | 0 | Cytoplasm \| Golgi apparatus | | | | GTP binding [GO:000552 | GTP binding [G | UP000018817 |
| PPTG_05087 | W2QVI9 | 224,84 | Kinesin motor domain-containing protein | 1617 | TRAFAC class myosin-kinesin ATPase superfamily, Kinesin fami | 0 | 0 | 0 | Cytoplasm | | transport along microtubule [GO:0010970] | | ATP binding [GO:000552 | ATP binding [G | UP000018817 |
| PPTG_03719 | W2R8G6 | 173,35 | Myosin motor domain-containing protein | 1535 | TRAFAC class myosin-kinesin ATPase superfamily, Myosin fami | 0 | 0 | 0 | Cytoplasm | | actin filament organizatior | cytoplasm [GO:000 | actin filament binding [G | cytoplasm [G | UP000018817 |
| PPTG_01205 | W2RI21 | 215,2 | Myosin motor domain-containing protein | 1079 | TRAFAC class myosin-kinesin ATPase superfamily, Myosin fami | 0 | 0 | 0 | Cytoplasm | | actin filament organizatior | cytoplasm [GO:000 | actin filament binding [G | cytoplasm [G | UP000018817 |
| PPTG_12096 | W2Q6V4 | 215,57 | Myosin motor domain-containing protein | 1474 | TRAFAC class myosin-kinesin ATPase superfamily, Myosin fami | 0 | 0 | 0 | Cytoplasm | | actin filament organizatior | cytoplasm [GO:000 | actin filament binding [G | cytoplasm [G | UP000018817 |
| PPTG_11158 | W2QA21 | 207,08 | Elongation factor 1-alpha | 443 | TRAFAC class translation factor GTPase superfamily, Classic tr | 0 | 0 | 0 | Cytoplasm | Nuclear localization signal | | cytoplasm [GO:000 | GTP binding [GO:000552 | cytoplasm [G | UP000018817 |
| PPTG_02259 | W2RAA1 | 181,54 | protein-synthesizing GTPase (EC 3.6.5.3) | 462 | TRAFAC class translation factor GTPase superfamily, Classic tr | 0 | 0 | 0 | Cytoplasm \| Nucleus | Nuclear export signal | formation of translation pr | cytosol [GO:000582 | GTP binding [GO:000552 | cytosol [GO:0 | UP000018817 |
| PPTG_02800 | W2RCJ8 | 190,79 | Transaldolase (EC 2.2.1.2) | 334 | Transaldolase family, Type 1 subfamily | 0 | 0 | 0 | Nucleus | Nuclear localization signal | carbohydrate metabolic pr | cytoplasm [GO:000 | transaldolase activity [G | cytoplasm [G | UP000018817 |
| PPTG_08466 | W2QLE9 | 182,76 | Tubulin alpha chain | 452 | Tubulin family | 0 | 0 | 0 | Cytoplasm | Nuclear export signal | microtubule-based proces | microtubule [GO:00 | GTP binding [GO:000552 | microtubule [ | UP000018817 |
| PPTG_08498 | W2QKW0 | 191,83 | Tubulin alpha chain | 453 | Tubulin family | 0 | 0 | 0 | Cytoplasm | Nuclear export signal | microtubule-based proces | microtubule [GO:00 | GTP binding [GO:000552 | microtubule [ | UP000018817 |
| PPTG_08506 | W2QKW5 | 191,83 | Tubulin alpha chain | 453 | Tubulin family | 0 | 0 | 0 | Cytoplasm | Nuclear export signal | microtubule-based proces | microtubule [GO:00 | GTP binding [GO:000552 | microtubule [ | UP000018817 |
| PPTG_08465 | W2QNB5 | 207,01 | Tubulin alpha chain | 452 | Tubulin family | 0 | 0 | 0 | Cytoplasm | Nuclear export signal | microtubule-based proces | microtubule [GO:00 | GTP binding [GO:000552 | microtubule [ | UP000018817 |
| PPTG_08453 | W2QMI3 | 209,97 | Tubulin alpha chain | 454 | Tubulin family | 0 | 0 | 0 | Cytoplasm | Nuclear export signal | microtubule-based proces | microtubule [GO:00 | GTP binding [GO:000552 | microtubule [ | UP000018817 |
| PPTG_15113 | W2PW38 | 218,06 | Tubulin beta chain | 446 | Tubulin family | 0 | 0 | 0 | Cytoplasm | Nuclear export signal | microtubule-based proces | microtubule [GO:00 | GTP binding [GO:000552 | microtubule [ | UP000018817 |
| PPTG_13968 | W2Q0Y9 | 177,59 | Ubiquinone biosynthesis monooxygenase COQ6, mitochondrial (EC 1.14.15.45) | 482 | UbiH/COQ6 family | 0 | 0 | 0 | Mitochondrion | Mitochondrial transit peptide | | extrinsic componen | 2-methoxy-6-polyprenol | extrinsic comp | UP000018817 |
| PPTG_02851 | W2RCR9 | 182,6 | E1 ubiquitin-activating enzyme (EC 6.2.1.45) | 1035 | Ubiquitin-activating E1 family | 0 | 0 | 0 | Cytoplasm | Nuclear localization signal | protein sumoylation [GO:0 | cytoplasm [GO:000 | ATP binding [GO:000552 | cytoplasm [G | UP000018817 |
| PPTG_16495 | W2PPR1 | 210,53 | E1 ubiquitin-activating enzyme (EC 6.2.1.45) | 1124 | Ubiquitin-activating E1 family | 0 | 0 | 0 | Cytoplasm | Nuclear localization signal | protein sumoylation [GO:0 | cytoplasm [GO:000 | ATP binding [GO:000552 | cytoplasm [G | UP000018817 |
| PPTG_00180 | W2RGD6 | 188,39 | UDP-glucose 6-dehydrogenase (EC 1.1.1.22) | 475 | UDP-glucose/GDP-mannose dehydrogenase family | 0 | 0 | 0 | Cytoplasm | Nuclear localization signal \| Nuclear export signal | glycosaminoglycan biosyn | nucleus [GO:00056 | NAD binding [GO:00512 | nucleus [GO:0 | UP000018817 |
| PPTG_17090 | W2PLY2 | 301,43 | Uncharacterized protein | 1138 | UDPGP type 1 family; Phosphohexose mutase family | 0 | 0 | 0 | Cytoplasm | | carbohydrate metabolic pr | cytosol [GO:000582 | magnesium ion binding [ | cytosol [GO:0 | UP000018817 |
| PPTG_13005 | W2Q5Y3 | 190,35 | Ribosomal protein | 229 | Universal ribosomal protein uL1 family | 0 | 0 | 0 | Cytoplasm | Nuclear localization signal | translation [GO:0006412] | large ribosomal sub | RNA binding [GO:000372 | large ribosom | UP000018817 |
| PPTG_11356 | W2QB22 | 212,59 | 60S acidic ribosomal protein P0 | 317 | Universal ribosomal protein uL10 family | 0 | 0 | 0 | Cytoplasm | Nuclear localization signal | cytoplasmic translation [G | cytosolic large ribos | large ribosomal subunit r | cytosolic large | UP000018817 |
| PPTG_15526 | W2PP80 | 219,43 | Large ribosomal subunit protein uL18 C-terminal eukaryotes domain-containing | 301 | Universal ribosomal protein uL18 family | 0 | 0 | 0 | Cytoplasm | Nuclear localization signal | ribosomal large subunit as | cytosolic large ribos | 5S rRNA binding [GO:000 | cytosolic large | UP000018817 |
| PPTG_13414 | W2Q5J9 | 218,49 | 60S ribosomal protein L3 | 389 | Universal ribosomal protein uL3 family | 0 | 0 | 0 | Cytoplasm | Nuclear localization signal | translation [GO:0006412] | cytosolic large ribos | RNA binding [GO:000372 | cytosolic large | UP000018817 |
| PPTG_16532 | W2PPX8 | 215,96 | Large ribosomal subunit protein uL4 C-terminal domain-containing protein | 387 | Universal ribosomal protein uL4 family | 0 | 0 | 0 | Cytoplasm | Nuclear localization signal | translation [GO:0006412] | ribonucleoprotein c | structural constituent of | ribonucleopro | UP000018817 |
| PPTG_14424 | W2PX30 | 180,3 | Large ribosomal subunit protein uL6 alpha-beta domain-containing protein | 189 | Universal ribosomal protein uL6 family | 0 | 0 | 0 | Cytoplasm | Nuclear localization signal | cytoplasmic translation [G | cytosolic large ribos | rRNA binding [GO:00198 | cytosolic large | UP000018817 |
| PPTG_19077 | W2PGG5 | 177,87 | Small ribosomal subunit protein uS2 | 282 | Universal ribosomal protein uS2 family | 0 | 0 | 0 | Cytoplasm | | ribosomal small subunit as | cytosolic small ribo | structural constituent of | cytosolic sma | UP000018817 |
| PPTG_06192 | W2QUI4 | 181,92 | 40S ribosomal protein S3 | 246 | Universal ribosomal protein uS3 family | 0 | 0 | 0 | Cytoplasm | Nuclear localization signal | translation [GO:0006412] | cytosolic small ribo | RNA binding [GO:000372 | cytosolic sma | UP000018817 |
| PPTG_15739 | W2PRG8 | 173,13 | S5 DRBM domain-containing protein | 312 | Universal ribosomal protein uS5 family | 0 | 0 | 0 | Mitochondrion | Mitochondrial transit peptide | translation [GO:0006412] | ribonucleoprotein c | RNA binding [GO:000372 | ribonucleopro | UP000018817 |
| PPTG_11874 | W2Q9D6 | 196,67 | Small ribosomal subunit protein uS5 (40S ribosomal protein S2) | 261 | Universal ribosomal protein uS5 family | 0 | 0 | 0 | Cytoplasm \| Nucleus | Nuclear localization signal | translation [GO:0006412] | cytosolic small ribo | RNA binding [GO:000372 | cytosolic sma | UP000018817 |
| PPTG_03669 | W2R5S3 | 176,54 | 40S ribosomal protein | 196 | Universal ribosomal protein uS7 family | 0 | 0 | 0 | Cytoplasm \| Nucleus | Nuclear localization signal | translation [GO:0006412] | small ribosomal sul | RNA binding [GO:000372 | small ribosom | UP000018817 |
| PPTG_14388 | W2PYI0 | 197,36 | HECT-type E3 ubiquitin transferase (EC 2.3.2.26) | 4149 | UPL family, TOM1/PTR1 subfamily | 0 | 0 | 0 | Nucleus | Nuclear localization signal \| Nuclear export signal | mRNA transport [GO:0051 | cytoplasm [GO:000 | ubiquitin protein ligase a | cytoplasm [G | UP000018817 |
| PPTG_07998 | W2QLC3 | 188,28 | V-type proton ATPase subunit H | 463 | V-ATPase H subunit family | 0 | 0 | 0 | Cytoplasm | Nuclear localization signal \| Nuclear export signal | | vacuolar proton-tra | proton-transporting ATPa | vacuolar proto | UP000018817 |
| PPTG_12116 | W2Q6X8 | 177,75 | V-type proton ATPase subunit | 394 | V-ATPase V0D/AC39 subunit family | 0 | 0 | 0 | Cytoplasm \| Lysosome/Vacuole | Nuclear localization signal | | proton-transporting | proton-transporting ATPa | proton-transp | UP000018817 |
| PPTG_02292 | W2RAA2 | 249,71 | HP domain-containing protein | 881 | Villin/gelsolin family | 0 | 0 | 0 | Cytoplasm | | actin filament capping [GO:0051693]; cytoskel | | actin filament binding [G | actin filament | UP000018817 |
| PPTG_10291 | W2QEW5 | 212,81 | 5-methyltetrahydropteroyltriglutamate--homocysteine S-methyltransferase (EC | 767 | Vitamin-B12 independent methionine synthase family | 0 | 0 | 0 | Cytoplasm | Nuclear localization signal | methionine biosynthetic process [GO:0009086 | | 5-methyltetrahydroptero | 5-methyltetra | UP000018817 |
| PPTG_03875 | W2QYB8 | 178,79 | EF-hand domain-containing protein | 2895 | VPS13 family | 0 | 0 | 0 | Lysosome/Vacuole | Transmembrane domain | protein retention in Golgi apparatus [GO:0045 | | calcium ion binding [GO: | calcium ion bi | UP000018817 |
| PPTG_09870 | W2QD77 | 184,67 | Uncharacterized protein | 2873 | VPS13 family | 0 | 0 | 0 | Lysosome/Vacuole | Transmembrane domain | protein retention in Golgi apparatus [GO:0045 | | calcium ion binding [GO: | calcium ion bi | UP000018817 |
| PPTG_04610 | W2R1L5 | 200,62 | Beta'-coat protein | 1094 | WD repeat COPB2 family | 0 | 0 | 0 | Cytoplasm | Nuclear localization signal \| Nuclear export signal | endoplasmic reticulum to | COPI vesicle coat [ | structural molecule activ | COPI vesicle c | UP000018817 |
| PPTG_00768 | W2RIN1 | 184,26 | Coronin | 507 | WD repeat coronin family | 0 | 0 | 0 | Cytoplasm | | actin filament organization [GO:0007015] | | actin filament binding [G | actin filament | UP000018817 |
| PPTG_18378 | W2PH74 | 192,71 | Uncharacterized protein | 344 | WD repeat G protein beta family | 0 | 0 | 0 | Cytoplasm | Nuclear localization signal | signal transduction [GO:0007165] | | | signal transdu | UP000018817 |
| PPTG_13436 | W2Q1B4 | 198,67 | Protein transport protein SEC31 (Protein transport protein sec31) | 1175 | WD repeat SEC31 family | 0 | 0 | 0 | Cytoplasm | Nuclear localization signal | COPII-coated vesicle carg | COPII vesicle coat [ | structural molecule activ | COPII vesicle | UP000018817 |
| PPTG_09250 | W2QGX3 | 202,17 | NAD(P)H:quinone oxidoreductase, type IV | 200 | WrbA family | 0 | 0 | 0 | Cytoplasm | | | membrane [GO:001 | FMN binding [GO:00101 | membrane [G | UP000018817 |
| PPTG_08403 | W2QL58 | 177,48 | Xylose isomerase (EC 5.3.1.5) | 452 | Xylose isomerase family | 0 | 0 | 0 | Cytoplasm \| Nucleus | Nuclear localization signal \| Nuclear export signal | D-xylose metabolic process [GO:0042732] | | metal ion binding [GO:00 | metal ion bind | UP000018817 |
| PPTG_19659 | W2PEB1 | 199,2 | S-(hydroxymethyl)glutathione dehydrogenase (EC 1.1.1.284) | 387 | Zinc-containing alcohol dehydrogenase family, Class-III subfam | 0 | 0 | 0 | Cytoplasm | Nuclear localization signal | formaldehyde catabolic pr | cytosol [GO:000582 | alcohol dehydrogenase ( | cytosol [GO:0 | UP000018817 |
| PPTG_15414 | W2PV05 | 223,82 | enoyl-[acyl-carrier-protein] reductase (EC 1.3.1.104) | 348 | Zinc-containing alcohol dehydrogenase family, Quinone oxidore | 0 | 0 | 0 | Mitochondrion | Mitochondrial transit peptide | fatty acid biosynthetic proc | mitochondrion [GO | enoyl-[acyl-carrier-prote | mitochondrion | UP000018817 |
| PPTG_16844 | W2PMJ0 | 184,4 | 2,4-dienoyl-CoA reductase | 322 | | 0 | 0 | 0 | Mitochondrion | Mitochondrial transit peptide | fatty acid beta-oxidation [G | mitochondrion [GO | 2,4-dienoyl-CoA reducta | mitochondrion | UP000018817 |
| PPTG_08551 | W2QL14 | 215,69 | 26S proteasome non-ATPase regulatory subunit 3 N-terminal TPR repeats domai | 373 | | 0 | 0 | 0 | Nucleus | Nuclear localization signal | ubiquitin-dependent prote | proteasome regulatory particle, lid subcompl | | proteasome re | UP000018817 |
| PPTG_17298 | W2PJE4 | 209,2 | Aconitate hydratase | 700 | | 0 | 0 | 0 | Cytoplasm | | L-amino acid biosynthetic process [GO:01700 | | hydro-lyase activity [GO: | hydro-lyase ac | UP000018817 |
| PPTG_06442 | W2QVJ6 | 228,1 | AMP-dependent synthetase/ligase domain-containing protein | 646 | | 0 | 0 | 0 | Cytoplasm | Nuclear localization signal | | endoplasmic reticu | long-chain fatty acid-Co/ | endoplasmic r | UP000018817 |
| PPTG_16051 | W2PPW8 | 172,9 | Intraflagellar transport protein | 1507 | | 0 | 0 | 0 | Cytoplasm | Nuclear localization signal | intraciliary retrograde tran | axoneme [GO:0005930]; ciliary basal body [G | | axoneme [GO | UP000018817 |
| PPTG_14857 | W2PWB8 | 191,75 | AP-4 complex subunit epsilon-1 C-terminal domain-containing protein | 1103 | | 0 | 0 | 0 | Cytoplasm | Nuclear localization signal | clathrin-dependent endoc | AP-2 adaptor compl | clathrin adaptor activity [ | AP-2 adaptor c | UP000018817 |
| PPTG_19658 | W2PDR0 | 238,19 | Argininosuccinate synthase (EC 6.3.4.5) (Citrulline--aspartate ligase) | 421 | | 0 | 0 | 0 | Cytoplasm | Nuclear localization signal | argininosuccinate metabo | cytoplasm [GO:000 | argininosuccinate syntha | cytoplasm [G | UP000018817 |
| PPTG_12072 | W2Q5T5 | 227,12 | B30.2/SPRY domain-containing protein | 4810 | | 0 | 0 | 0 | Cytoplasm \| Nucleus | Nuclear localization signal \| Nuclear export signal | protein ubiquitination [GO:0016567] | | metal ion binding [GO:00 | metal ion bind | UP000018817 |
| PPTG_14782 | W2PWT7 | 258,46 | B30.2/SPRY domain-containing protein | 5307 | | 0 | 0 | 0 | Cytoplasm | Nuclear export signal | | | ubiquitin-protein transfe | ubiquitin-prot | UP000018817 |
| PPTG_15229 | W2PSK8 | 192,07 | C2 domain-containing protein | 871 | | 0 | 0 | 0 | Cytoplasm \| Lysosome/Vacuole | | | membrane [GO:001 | calcium ion binding [GO: | membrane [G | UP000018817 |
| PPTG_11937 | W2Q8V7 | 203,41 | C2 domain-containing protein | 1827 | | 0 | 0 | 0 | Cell membrane \| Golgi apparatus | Signal peptide \| Transmembrane domain | plasma membrane organiz | membrane [GO:0016020] | | membrane [G | UP000018817 |
| PPTG_11926 | W2Q8U3 | 179,61 | C2 NT-type domain-containing protein | 794 | | 0 | 0 | 0 | Cytoplasm | | | | | | UP000018817 |
| PPTG_19423 | W2PCK8 | 182,87 | Calmodulin | 575 | | 0 | 0 | 0 | Cytoplasm | | endocytosis [GO:0006897 | plasma membrane | calcium ion binding [GO: | plasma memb | UP000018817 |
| PPTG_08772 | W2QII2 | 259,32 | Carbamoyl-phosphate synthase, large subunit | 1499 | | 0 | 0 | 0 | Cytoplasm | Peroxisomal targeting signal | 'de novo' pyrimidine nucle | carbamoyl-phospha | ATP binding [GO:000552 | carbamoyl-ph | UP000018817 |
| PPTG_01415 | W2R6W3 | 223,76 | Coatomer subunit alpha | 1243 | | 0 | 0 | 0 | Cytoplasm | Nuclear export signal | endoplasmic reticulum to | COPI vesicle coat [ | structural molecule activ | COPI vesicle c | UP000018817 |
| PPTG_18415 | W2PI14 | 227,37 | Coatomer subunit beta (Beta-coat protein) | 993 | | 0 | 0 | 0 | Cytoplasm \| Golgi apparatus | | endoplasmic reticulum to | COPI vesicle coat [ | structural molecule activ | COPI vesicle c | UP000018817 |
| PPTG_13885 | W2Q1C0 | 211 | Dihydrodipicolinate reductase | 633 | | 0 | 0 | 0 | Cytoplasm | Nuclear localization signal | lysine biosynthetic process via diaminopimelat | | 2-aminoadipate transam | 2-aminoadipa | UP000018817 |

| | | | | | | | | | | | | | | |
|---|---|---|---|---|---|---|---|---|---|---|---|---|---|---|
| PPTG_12430 | W2Q5T7 | 173,55 | DUSP domain-containing protein | 637 | 0 | 0 | 0 | Cytoplasm \| Lysosome/Vacuole | | | | cysteine-type deubiquiti | cysteine-type | UP000018817 |
| PPTG_17319 | W2PJ20 | 178,23 | EIF3d | 551 | 0 | 0 | 0 | Cytoplasm \| Nucleus | Nuclear localization signal | | eukaryotic translati | RNA binding [GO:000372 | eukaryotic tra | UP000018817 |
| PPTG_02398 | W2RCS0 | 191,43 | Elongation factor 1-gamma | 406 | 0 | 0 | 0 | Cytoplasm | Nuclear localization signal | | cytoplasm [GO:000 | translation elongation fa | cytoplasm [G | UP000018817 |
| PPTG_01091 | W2RJX2 | 299,77 | Fibronectin type-III domain-containing protein | 5915 | 0 | 0 | 0 | Cell membrane | Transmembrane domain | | | | | UP000018817 |
| PPTG_01092 | W2RJZ0 | 323,63 | Fibronectin type-III domain-containing protein | 5778 | 0 | 0 | 0 | Cell membrane | Transmembrane domain | | | | | UP000018817 |
| PPTG_11500 | W2QCD3 | 175,84 | FYVE-type domain-containing protein | 558 | 0 | 0 | 0 | Cytoplasm | | | | | | UP000018817 |
| PPTG_09984 | W2QFA2 | 219,22 | Gamma carbonic anhydrase | 251 | 0 | 0 | 0 | Mitochondrion | Mitochondrial transit peptide | | | | | UP000018817 |
| PPTG_14642 | W2PXZ4 | 216,13 | Glutamine--fructose-6-phosphate aminotransferase [isomerizing] (EC 2.6.1.16) | 701 | 0 | 0 | 0 | Mitochondrion | Mitochondrial transit peptide | fructose 6-phosphate metabolic process [GO: | | carbohydrate derivative | carbohydrate | UP000018817 |
| PPTG_17667 | W2PIK2 | 185,69 | Glycoside hydrolase family 19 catalytic domain-containing protein | 276 | 0 | 0 | 0 | Extracellular | Signal peptide | cell wall macromolecule catabolic process [G | | chitinase activity [GO:00 | chitinase activ | UP000018817 |
| PPTG_02671 | W2RCG7 | 187,68 | GST C-terminal domain-containing protein | 354 | 0 | 0 | 0 | Cytoplasm | | | cytoplasm [GO:000 | glutathione transferase a | cytoplasm [G | UP000018817 |
| PPTG_13453 | W2Q1M2 | 183,84 | Guanine nucleotide-binding protein alpha-1 subunit | 355 | 0 | 0 | 0 | Cytoplasm \| Cell membrane | | adenylate cyclase-modula | cytoplasm [GO:000 | G protein-coupled recep | cytoplasm [G | UP000018817 |
| PPTG_08926 | W2QK47 | 222,79 | Helicase ATP-binding domain-containing protein | 1094 | 0 | 0 | 0 | Nucleus | Nuclear localization signal | | cytoplasm [GO:000 | ATP binding [GO:000552 | cytoplasm [G | UP000018817 |
| PPTG_14511 | W2PWR6 | 172,2 | Importin N-terminal domain-containing protein | 1129 | 0 | 0 | 0 | Cytoplasm \| Nucleus | Nuclear localization signal \| Nuclear export signal | protein import into nucleus | cytoplasm [GO:0005737]; nucleus [GO:0005 | | cytoplasm [G | UP000018817 |
| PPTG_10113 | W2QE49 | 188,42 | Importin N-terminal domain-containing protein | 1079 | 0 | 0 | 0 | Cytoplasm \| Nucleus | Nuclear localization signal | protein import into nucleus | cytoplasm [GO:000 | small GTPase binding [G | cytoplasm [G | UP000018817 |
| PPTG_04382 | W2R0H5 | 225,19 | Major vault protein | 881 | 0 | 0 | 0 | Cytoplasm | | | cytoplasm [GO:0005737]; nucleus [GO:0005 | | cytoplasm [G | UP000018817 |
| PPTG_15388 | W2PSU7 | 214,98 | methylmalonate-semialdehyde dehydrogenase (CoA acylating) (EC 1.2.1.27) | 505 | 0 | 0 | 0 | Cytoplasm \| Nucleus | Nuclear localization signal | L-valine catabolic process [GO:0006574]; thym | | methylmalonate-semial | methylmalona | UP000018817 |
| PPTG_14738 | W2PYD0 | 179,81 | MIT domain-containing protein | 135 | 0 | 0 | 0 | Cytoplasm | | | | | | UP000018817 |
| PPTG_02328 | W2RCX4 | 191,76 | NAD-dependent epimerase/dehydratase domain-containing protein | 379 | 0 | 0 | 0 | Mitochondrion | Mitochondrial transit peptide | | mitochondrion [GO: | protein-containing comp | mitochondrio | UP000018817 |
| PPTG_08378 | W2QKC6 | 173,69 | PCI domain-containing protein | 416 | 0 | 0 | 0 | Cytoplasm \| Nucleus | Nuclear export signal | proteasome-mediated ubi | proteasome complex [GO:0000502] | | proteasome c | UP000018817 |
| PPTG_00791 | W2RGG3 | 220,51 | Peptidase M1 leukotriene A4 hydrolase/aminopeptidase C-terminal domain-con | 1008 | 0 | 0 | 0 | Cytoplasm | Nuclear localization signal | | | aminopeptidase activity | aminopeptida | UP000018817 |
| PPTG_08190 | W2QKA6 | 210,62 | phosphatidylinositol 3-kinase (EC 2.7.1.137) | 1036 | 0 | 0 | 0 | Cytoplasm \| Lysosome/Vacuole | | bleb assembly [GO:00320 | cytoplasm [GO:000 | 1-phosphatidylinositol-3- | cytoplasm [G | UP000018817 |
| PPTG_03645 | W2R7M2 | 178,44 | Phosphoenolpyruvate carboxykinase (ATP) | 379 | 0 | 0 | 0 | Mitochondrion | Mitochondrial transit peptide | gluconeogenesis [GO:000 | cytosol [GO:000582 | ATP binding [GO:000552 | cytosol [GO:0 | UP000018817 |
| PPTG_10444 | W2QEI6 | 209,94 | phospholipase D (EC 3.1.4.4) | 530 | 0 | 0 | 0 | Cytoplasm | | phospholipid catabolic pro | plasma membrane | phospholipase D activity | plasma memb | UP000018817 |
| PPTG_18731 | W2PF41 | 176,08 | Poly [ADP-ribose] polymerase (PARP) (EC 2.4.2.-) | 2885 | 0 | 0 | 0 | Nucleus | Nuclear localization signal | | | NAD+ poly-ADP-ribosyltr | NAD+ poly-AD | UP000018817 |
| PPTG_02403 | W2RCS4 | 182,79 | Pre-mRNA-processing-splicing factor 8 | 2359 | 0 | 0 | 0 | Nucleus | Nuclear export signal | spliceosomal tri-snRNP co | catalytic step 2 spli | metallopeptidase activity | catalytic step | UP000018817 |
| PPTG_18809 | W2PHP4 | 191,07 | proline--tRNA ligase (EC 6.1.1.15) (Prolyl-tRNA synthetase) | 711 | 0 | 0 | 0 | Cytoplasm | Nuclear localization signal | prolyl-tRNA aminoacylatio | aminoacyl-tRNA syr | aminoacyl-tRNA deacyla | aminoacyl-tR | UP000018817 |
| PPTG_06281 | W2QSV9 | 182,17 | Propionyl-CoA carboxylase alpha chain, mitochondrial (EC 6.4.1.3) (Propanoyl-C | 764 | 0 | 0 | 0 | Mitochondrion | Mitochondrial transit peptide | lipid catabolic process [G | mitochondrial matr | ATP binding [GO:000552 | mitochondrial | UP000018817 |
| PPTG_01206 | W2RKB5 | 246,97 | Pyruvate carboxylase (EC 6.4.1.1) | 1186 | 0 | 0 | 0 | Mitochondrion | Mitochondrial transit peptide | gluconeogenesis [GO:000 | cytoplasm [GO:000 | ATP binding [GO:000552 | cytoplasm [G | UP000018817 |
| PPTG_16088 | W2PS91 | 179,68 | Pyruvate dehydrogenase E1 component subunit alpha (EC 1.2.4.1) | 402 | 0 | 0 | 0 | Mitochondrion | Mitochondrial transit peptide | pyruvate decarboxylation to acetyl-CoA [GO:0 | | pyruvate dehydrogenase | pyruvate dehy | UP000018817 |
| PPTG_20351 | W2P921 | 256,78 | Ricin B lectin domain-containing protein | 466 | 0 | 0 | 0 | Cytoplasm | | protein O-linked glycosylation [GO:0006493] | | carbohydrate binding [G | carbohydrate | UP000018817 |
| PPTG_08673 | W2QLJ3 | 181,99 | SEC7 domain-containing protein | 2046 | 0 | 0 | 0 | Cytoplasm \| Lysosome/Vacuole | | protein transport [GO:001 | cytoplasm [GO:000 | guanyl-nucleotide excha | cytoplasm [G | UP000018817 |
| PPTG_15407 | W2PVJ8 | 185,27 | Serine protease | 685 | 0 | 0 | 0 | Lysosome/Vacuole | Signal peptide | | | | | UP000018817 |
| PPTG_04587 | W2R3C1 | 402,1 | Titin | 10565 | 0 | 0 | 0 | | | | | | | UP000018817 |
| PPTG_11181 | W2QA57 | 172,14 | TKL protein kinase | 359 | 0 | 0 | 0 | Cytoplasm \| Cell membrane | Nuclear localization signal | | | ATP binding [GO:000552 | ATP binding [G | UP000018817 |
| PPTG_16233 | W2PP25 | 173,73 | Transglutaminase elicitor | 910 | 0 | 0 | 0 | Extracellular | Signal peptide | regulation of growth [GO:0040008] | | aminoacyltransferase ac | aminoacyltrar | UP000018817 |
| PPTG_03763 | W2QZZ8 | 218,89 | Transitional endoplasmic reticulum ATPase | 804 | 0 | 0 | 0 | Cytoplasm \| Nucleus | Nuclear localization signal \| Nuclear export signal | autophagosome maturatic | cytosol [GO:000582 | ATP binding [GO:000552 | cytosol [GO:0 | UP000018817 |
| PPTG_02434 | W2RCV6 | 295,49 | Translation elongation factor aEF-2 | 858 | 0 | 0 | 0 | Cytoplasm | Nuclear localization signal \| Nuclear export signal | | cytosol [GO:000582 | GTP binding [GO:000552 | cytosol [GO:0 | UP000018817 |
| PPTG_12239 | W2Q5I7 | 231,76 | ubiquitinyl hydrolase 1 (EC 3.4.19.12) | 4322 | 0 | 0 | 0 | Cytoplasm | | protein deubiquitination in | cytoplasm [GO:000 | calcium ion binding [GO: | cytoplasm [G | UP000018817 |
| PPTG_02443 | W2RDA0 | 175,82 | Uncharacterized protein | 801 | 0 | 0 | 0 | Cytoplasm | | proteolysis [GO:0006508] | | dipeptidyl-peptidase acti | dipeptidyl-pep | UP000018817 |
| PPTG_17401 | W2PJW0 | 176,31 | Uncharacterized protein | 416 | 0 | 0 | 0 | Mitochondrion | Mitochondrial transit peptide | | | | | UP000018817 |
| PPTG_09673 | W2QFQ0 | 179,55 | Uncharacterized protein | 342 | 0 | 0 | 0 | Cytoplasm \| Lysosome/Vacuole | | | | phosphatidylinositol binc | phosphatidyli | UP000018817 |
| PPTG_06597 | W2QQ58 | 179,98 | Uncharacterized protein | 381 | 0 | 0 | 0 | Cytoplasm \| Nucleus | Nuclear localization signal | | | | | UP000018817 |
| PPTG_12440 | W2Q6M8 | 180,84 | Uncharacterized protein | 329 | 0 | 0 | 0 | Cytoplasm | | | | oxidoreductase activity [ | oxidoreductas | UP000018817 |
| PPTG_16755 | W2PPB1 | 182,93 | Uncharacterized protein | 1034 | 0 | 0 | 0 | Cytoplasm | Nuclear localization signal | | | actin binding [GO:00037 | actin binding [ | UP000018817 |
| PPTG_00098 | W2RFQ0 | 190,42 | Uncharacterized protein | 317 | 0 | 0 | 0 | Cytoplasm | | | | | | UP000018817 |
| PPTG_16306 | W2PRN2 | 191,03 | Uncharacterized protein | 230 | 0 | 0 | 0 | Cytoplasm | | | | | | UP000018817 |
| PPTG_11971 | W2Q5E7 | 201,53 | Uncharacterized protein | 496 | 0 | 0 | 0 | Cytoplasm | | | | | | UP000018817 |
| PPTG_09470 | W2QH02 | 204,64 | Uncharacterized protein | 2326 | 0 | 0 | 0 | Cytoplasm | | fatty acid biosynthetic process [GO:0006633]; | | acetyl-CoA carboxylase a | acetyl-CoA ca | UP000018817 |
| PPTG_00228 | W2RE69 | 215,22 | Uncharacterized protein | 612 | 0 | 0 | 0 | Cytoplasm | Nuclear localization signal | actin filament depolymeriz | cortical actin cytosk | actin filament binding [G | cortical actin | UP000018817 |
| PPTG_12389 | W2Q3Y1 | 216,78 | Uncharacterized protein | 919 | 0 | 0 | 0 | Cytoplasm | Nuclear localization signal | mRNA catabolic process [ | cytosol [GO:000582 | nuclease activity [GO:00 | cytosol [GO:0 | UP000018817 |
| PPTG_06139 | W2QSF2 | 222,03 | Uncharacterized protein | 771 | 0 | 0 | 0 | Cytoplasm | Peroxisomal targeting signal | | | oxidoreductase activity, | oxidoreductas | UP000018817 |
| PPTG_04733 | W2R3Z5 | 280,93 | Uncharacterized protein | 5308 | 0 | 0 | 0 | Cytoplasm | Nuclear localization signal | | membrane [GO:001 | calcium ion binding [GO: | membrane [G | UP000018817 |
| PPTG_12075 | W2Q4W4 | 183,31 | USP domain-containing protein | 2913 | 0 | 0 | 0 | Cytoplasm \| Nucleus | Nuclear export signal | protein deubiquitination [G | cytosol [GO:000582 | cysteine-type deubiquiti | cytosol [GO:0 | UP000018817 |
| PPTG_02106 | W2RBI2 | 176,86 | Xaa-Pro dipeptidyl-peptidase C-terminal domain-containing protein | 740 | 0 | 0 | 0 | Endoplasmic reticulum | Signal peptide | | | dipeptidyl-peptidase acti | dipeptidyl-pep | UP000018817 |
