## Supplementary table 7 for "Zoospore-derived extracellular vesicles in the flagellated stramenopile *Phytophthora parasitica*"

### Title : An overlap of detected membrane associated proteins between this study and Lupatelli et al., 2025

| Gene ID | UniPROT ID | Protein name | Localisation | Localisation |
|---|---|---|---|---|
| **Transmembrane proteins associated with plasma membrane and/or endosomes** | | | | |
| PPTG_00859 | W2RGS8 | Uncharacterized protein | Transmembrane (1TM) | This study |
| PPTG_12681 | W2Q2U4 | ; transporter B family membe | Transmembrane (9TM) | This study |
| PPTG_13069 | W2Q664 | RVEL domain-containing prc | Transmembrane (8TM) | This study |
| PPTG_16242 | W2PNS1 | iaracterized protein prominir | Transmembrane (5TM) | This study |
| PPTG_08986 | W2QIM1 | Na+/K+-ATPase α-subunit | Transmembrane (12TM) | Lupatelli et al., 2025 |
| **GPI-anchored or secreted proteins associated with plasma and/or flagellar membrane** | | | | |
| PPTG_01189 | W2RIK1 | Secreted protein | GPI anchored | This study |
| PPTG_02441 | W2RAR2 | PnMas1 | Mastigoneme | Hee & Blackman, 2019 |
| PPTG_03648 | W2R5N9 | PnMas2 | Mastigoneme | Hee & Blackman, 2020 |
| PPTG_07238 | W2QPH8 | PnMas3 | Mastigoneme | Hee & Blackman, 2021 |
| PPTG_11537 | W2Q6A3 | Lipase-like protein | Extracellular | This study |
| PPTG_11759 | W2Q908 | Alpha-mannosidase | Extracellular | This study |
