## Supplementary table 8 for "Zoospore-derived extracellular vesicles in the flagellated stramenopile *Phytophthora parasitica*"

**Title: Total number and lipid class distribution of annotated lipids in cell bodies (CB), flagella (FL), and extracellular vesicles (EV).**

| RT | CCS ($Å^2$) | ΔCCS [%] | m/z meas. | M meas. | Ions | MS/MS | Name | Molecular Formula | Annotations | AQ | Annotation Source | Boxplot | Include | Blanc_Elut_pos_1_1_8771 | Blanc_Ext_1_pos_1_1_8772 | Blanc_Ext_2_pos_2_1_8773 | Colonne1 | Colonne2 | QC_3_pos_3_1_8776 | QC_4_pos_3_1_8777 | QC_5_pos_3_1_8781 | QC_6_pos_3_1_8786 | QC_7_pos_3_1_8791 | CV | EV1-2_pos_4_1_8778 | EV3-4_pos_5_1_8779 | EV3p4_pos_6_1_8780 | FL1_pos_7_1_8782 | FL2_pos_8_1_8783 | FL3_pos_9_1_8784 | FL4_pos_10_1_8785 | CB1_pos_11_1_8787 | CB2_pos_12_1_8788 | CB3_pos_13_1_8789 | CB4_pos_14_1_8790 |
|---|---|---|---|---|---|---|---|---|---|---|---|---|---|---|---|---|---|---|---|---|---|---|---|---|---|---|---|---|---|---|---|---|---|---|---|
| 15,09 | 286,5 | -0,5 | 780,55442 | 779,54665 | [M+H]+, [M+N | true | PC 36:5 | C44H78NO8P | Spectral Library - | 20222 | | 6172848 | true | 72 | 773 | 334 | | | 6172848 | 6253435 | 6208006 | 6324567 | 6290821 | 0,97831222 | 492707 | 63529 | 483105 | 408563 | 539414 | 502304 | 251529 | 15459442 | 13024176 | 17847758 | 11733074 |
| 6,24 | 230,5 | -0,5 | 520,33973 | 519,33245 | [M+H]+ | true | LPC 18:2 | C26H50NO7P | SmartFormula - r | 20222 | | 976819,875 | true | 0 | 0 | 0 | | | 986462 | 999726 | 1001627 | 999306 | 976820 | 1,0845567 | 45930 | 2676 | 11425 | 34958 | 408144 | 405521 | 378144 | 2759058 | 1716917 | 2307680 | 3999326 |
| 14,62 | 279,9 | -0,7 | 730,53787 | 729,53059 | [M+H]+ | true | PC 32:2 | C40H76NO8P | Spectral Library - | 20222 | | 619630,813 | true | 0 | 111 | 0 | | | 633983 | 623622 | 619631 | 628605 | 635934 | 1,09014109 | 61014 | 7520 | 56311 | 42523 | 70219 | 51471 | 27080 | 1437569 | 1087262 | 1483596 | 1165387 |
| 15,07 | 293,3 | -0,2 | 830,56877 | 829,56149 | [M+H]+ | true | PC 40:8 | C48H80NO8P | Spectral Library - | 20222 | | 1087267 | true | 0 | 203 | 144 | | | 1120874 | 1137254 | 1087267 | 1099054 | 1117835 | 1,75738684 | 132846 | 15219 | 118978 | 164589 | 263557 | 277454 | 199612 | 2851474 | 1992089 | 2853110 | 2212000 |
| 15,88 | 290,2 | -0,6 | 784,58454 | 783,57726 | [M+H]+ | true | PC 36:3 | C44H82NO8P | Spectral Library - | 20222 | | 743214,188 | true | 0 | 0 | 0 | | | 765131 | 750788 | 778402 | 755736 | 743214 | 1,79371065 | 89366 | 13055 | 91285 | 58532 | 105965 | 85391 | 60455 | 2269708 | 2052508 | 2558934 | 1302479 |
| 15,52 | 294,6 | -0,5 | 832,58428 | 831,577 | [M+H]+ | true | PC 40:7 | C48H82NO8P | Spectral Library - | 20222 | | 219561,266 | true | 0 | 0 | 0 | | | 219561 | 227670 | 220056 | 224560 | 228086 | 1,80989104 | 25251 | 3224 | 24214 | 26719 | 44835 | 39391 | 33810 | 696623 | 547461 | 736415 | 420580 |
| 14,5 | 291,2 | 0,5 | 828,5546 | 827,5458 | [M+H]+, [M+K] | true | PC20:4_20:5 | C48H78NO8P | Spectral Library - | 20122 | LipidBlast-VS69-Pos.ms | 6796564,5 | true | 0 | 597 | 421 | | | 6808029 | 7035813 | 7019573 | 6796565 | 7039679 | 1,81463347 | 627395 | 68427 | 553944 | 781666 | 1241154 | 1238680 | 991346 | 16278916 | 13449767 | 17550754 | 13532773 |
| 16,18 | 297 | -0,3 | 834,60041 | 833,59313 | [M+H]+ | true | PC 40:6 | C48H84NO8P | Spectral Library - | 20212 | | 65300,1797 | true | 0 | 0 | 0 | | | 67600 | 65867 | 68475 | 65300 | 67305 | 1,94311222 | 15464 | 2198 | 14476 | 10820 | 19026 | 15198 | 12357 | 213026 | 181645 | 224390 | 98904 |
| 15,69 | 286,1 | -0,6 | 758,57021 | 757,56256 | [M+H]+, [M+N | true | PC 34:2 | C42H80NO8P | Spectral Library - | 20222 | | 4755270,5 | true | 0 | 593 | 349 | | | 4992416 | 4948143 | 5026956 | 4925480 | 4755271 | 2,13178842 | 593527 | 74229 | 620478 | 358671 | 477618 | 461259 | 215820 | 13923117 | 10429422 | 14930295 | 8439548 |
| 15,81 | 293,4 | -0,2 | 808,5846 | 807,57732 | [M+H]+ | true | PC 38:5 | C46H82NO8P | Spectral Library - | 20222 | | 505772,125 | true | 0 | 119 | 0 | | | 524979 | 534593 | 528990 | 505772 | 529924 | 2,13401698 | 89588 | 12254 | 102780 | 98564 | 185739 | 160290 | 116340 | 1387423 | 1190446 | 1506634 | 839090 |
| 16,61 | 262,5 | 0,8 | 576,53503 | 575,52784 | [M+H]+, [M+N | true | Cer19:3;2O/18:0 | C37H69NO3 | SmartFormula - r | 20212 | LipidBlast-VS69-Pos.ms | 36655,5352 | true | 0 | 0 | 0 | | | 43503 | 45242 | 43305 | 42765 | 43407 | 2,14852765 | 36656 | 4598 | 32465 | 22145 | 24542 | 33970 | 31730 | 86980 | 79958 | 204241 | 34162 |
| 17,08 | 265,9 | 1,2 | 592,56664 | 591,55936 | [M+H]+, [M+H | true | Cer 16:1;O2/22:1 | C38H73NO3 | Lipid Species, Lip | 20221 | | 748904,438 | true | 0 | 294 | 247 | | | 748904 | 753729 | 788012 | 757073 | 777404 | 2,19931498 | 1568823 | 192324 | 1246224 | 740779 | 997347 | 1028788 | 956223 | 439662 | 525668 | 707563 | 145868 |
| 14,35 | 281,4 | 0,8 | 742,53739 | 741,53011 | [M+H]+ | true | PC9:0_24:3 | C41H76NO8P | SmartFormula - r | 20222 | LipidBlast-VS69-Pos.ms | 1733,71118 | true | 0 | 0 | 0 | | | 39493 | 37311 | 38926 | 37718 | 38538 | 2,30797089 | 0 | 1734 | 6944 | 0 | 36180 | 0 | 1152 | 0 | 300305 | 0 | 0 |
| 7,28 | 231,9 | -0,1 | 496,34024 | 495,33272 | [M+H]+, [M+N | true | LPC 16:0 | C24H50NO7P | SmartFormula - r | 20212 | | 127684,297 | true | 0 | 278 | 372 | | | 129537 | 133872 | 128944 | 127684 | 134556 | 2,36121376 | 4961 | 637 | 4928 | 19142 | 9706 | 8941 | 6913 | 369357 | 281777 | 478940 | 487858 |
| 14,98 | 288,5 | 0,1 | 792,55325 | 791,54597 | [M+H]+ | true | PC9:0_28:6 | C45H78NO8P | Spectral Library - | 20222 | LipidBlast-VS69-Pos.ms | 197021,375 | true | 0 | 0 | 0 | | | 366230 | 382295 | 391156 | 375374 | 376867 | 2,42758429 | 11006 | 7154 | 42294 | 13389 | 197021 | 17293 | 11883 | 276634 | 3138826 | 239708 | 128592 |
| 16,98 | 264,1 | -0,9 | 658,54063 | 640,50676 | [M+NH4]+, [M | true | DG 18:2_20:4 | C41H68O5 | Lipid Species, Sm | 20212 | | 151505,813 | true | 0 | 0 | 0 | | | 188194 | 192361 | 199840 | 192581 | 198336 | 2,45490402 | 46215 | 5535 | 40403 | 96582 | 151506 | 144792 | 137149 | 728382 | 349778 | 681148 | 93357 |
| 4,12 | 209,4 | 2,8 | 372,31114 | 371,30387 | [M+H]+ | true | C14Tetradecanoyl-L-Carnitine | C21H41NO4 | SmartFormula - r | 20211 | ALL_GNPS_NO_PROPO | 10461,4434 | true | 0 | 0 | 0 | | | 11008 | 11195 | 10822 | 10461 | 10953 | 2,51345382 | 718 | 0 | 242 | 431 | 474 | 385 | 426 | 35089 | 22952 | 35547 | 35701 |
| 15,12 | 280,3 | 0,2 | 706,53727 | 705,53 | [M+H]+ | true | PC 30:0 | C38H76NO8P | Lipid Species, Sp | 20222 | | 62587,625 | true | 0 | 0 | 0 | | | 62588 | 65435 | 63301 | 66411 | 63264 | 2,54717591 | 7828 | 916 | 6865 | 5481 | 6759 | 6806 | 5701 | 169175 | 91393 | 188113 | 129708 |
| 16,14 | 257,9 | 0,8 | 630,50925 | 612,47519 | [M+NH4]+, [M | true | DG 16:1_20:5 | C39H64O5 | Lipid Species, Lip | 20222 | | 32428,0684 | true | 0 | 0 | 0 | | | 44860 | 44781 | 47132 | 46707 | 44635 | 2,62096929 | 9153 | 675 | 7304 | 11353 | 22762 | 23271 | 19324 | 200104 | 75559 | 197223 | 32428 |
| 14,63 | 278,8 | -0,2 | 786,50658 | 785,4993 | [M+H]+ | true | PE20:4_20:5 | C45H72NO8P | CCS-Predict, Sm | 20222 | LipidBlast-VS69-Pos.ms | 1323982,88 | true | 0 | 183 | 117 | | | 1323983 | 1405263 | 1415830 | 1387942 | 1407545 | 2,6840784 | 283344 | 36452 | 287772 | 261859 | 539426 | 484475 | 380857 | 3485779 | 3537110 | 3677852 | 1664304 |
| 15,01 | 284,1 | 0,7 | 768,55354 | 767,54626 | [M+H]+ | true | PC17:2_18:2 | C43H78NO8P | SmartFormula - r | 20222 | LipidBlast-VS69-Pos.ms | 316405,781 | true | 0 | 99 | 0 | | | 626046 | 653885 | 660940 | 622016 | 650597 | 2,72328806 | 25741 | 13832 | 83566 | 27904 | 316406 | 37184 | 27114 | 555034 | 5120632 | 473357 | 253350 |
| 5,84 | 212,8 | -2 | 500,2775 | 499,27005 | [M+H]+, [M+N | true | LPE20:5 | C25H42NO7P | Spectral Library - | 20221 | LipidBlast-VS69-Pos.ms | 88375,3359 | true | 0 | 0 | 0 | | | 88375 | 92453 | 89587 | 90310 | 94773 | 2,78026063 | 971 | 188 | 0 | 12259 | 3796 | 2919 | 3156 | 292801 | 243026 | 341112 | 376306 |
| 6,92 | 235,1 | -0,1 | 546,35532 | 545,34779 | [M+H]+, [M+N | true | LPC 20:3 | C28H52NO7P | SmartFormula - r | 20222 | | 38854,3086 | true | 0 | 0 | 0 | | | 38854 | 40204 | 41019 | 39024 | 41302 | 2,79185966 | 2377 | 333 | 1086 | 2025 | 19307 | 16363 | 17619 | 113476 | 74797 | 96435 | 179316 |
| 8,23 | 244,5 | -0,7 | 598,38689 | 597,37961 | [M+H]+ | true | PCO-24:5 | C32H56NO7P | SmartFormula - r | 20222 | LipidBlast-VS69-Pos.ms | 2422,14355 | true | 0 | 0 | 0 | | | 2615 | 2511 | 2556 | 2510 | 2422 | 2,80754188 | 247 | 0 | 172 | 251 | 1545 | 1581 | 1592 | 6308 | 4296 | 5695 | 9293 |
| 15,43 | 285,1 | 0,6 | 744,5531 | 743,54582 | [M+H]+ | true | PC14:0_19:2 | C41H78NO8P | SmartFormula - r | 20222 | LipidBlast-VS69-Pos.ms | 71186,5547 | true | 0 | 0 | 0 | | | 72447 | 71187 | 76632 | 73315 | 72292 | 2,83648723 | 6274 | 1542 | 7977 | 5971 | 20467 | 7103 | 4389 | 218873 | 379264 | 186639 | 110458 |
| 15,94 | 292,3 | 1,3 | 772,58453 | 771,57726 | [M+H]+ | true | PC17:1_18:1 | C43H82NO8P | SmartFormula - r | 20221 | LipidBlast-VS69-Pos.ms | 5097,44531 | true | 86 | 0 | 0 | | | 134456 | 144915 | 142874 | 139475 | 139127 | 2,85474443 | 0 | 5097 | 27341 | 0 | 88730 | 0 | 0 | 0 | 1516886 | 0 | 0 |
| 7,87 | 237,8 | -0,6 | 548,37072 | 547,36344 | [M+H]+ | true | LPC 20:2 | C28H54NO7P | Lipid Species, Sp | 20222 | | 3588,00146 | true | 0 | 0 | 0 | | | 3588 | 3747 | 3753 | 3870 | 3831 | 2,88210494 | 520 | 0 | 247 | 481 | 3096 | 2637 | 2431 | 7868 | 7199 | 8085 | 12282 |
| 14,56 | 287,4 | -0,9 | 804,55438 | 803,5471 | [M+H]+ | true | PC 38:7 | C46H78NO8P | Spectral Library - | 20222 | | 6276045,5 | true | 0 | 616 | 329 | | | 6276046 | 6435791 | 6532073 | 6511033 | 6815905 | 3,01427064 | 584529 | 67230 | 552346 | 660006 | 1125910 | 1002782 | 771100 | 14687626 | 12932643 | 15379366 | 12327986 |
| 13,17 | 280,6 | -0,4 | 774,50599 | 773,49872 | [M+H]+ | true | PC 36:8 | C44H72NO8P | Spectral Library - | 20222 | | 5725,84619 | true | 0 | 0 | 0 | | | 5726 | 6012 | 5935 | 6154 | 5754 | 3,03031794 | 0 | 0 | 384 | 466 | 523 | 517 | 536 | 19307 | 11517 | 19260 | 17169 |
| 13,63 | 265,6 | 0,7 | 720,45957 | 719,45215 | [M+H]+, [M+N | true | PE9:0_26:7 | C40H66NO8P | SmartFormula - r | 20222 | LipidBlast-VS69-Pos.ms | 7150,8418 | true | 0 | 0 | 0 | | | 9216 | 9815 | 9903 | 9945 | 9823 | 3,06143391 | 642 | 426 | 2061 | 788 | 7151 | 1398 | 924 | 11561 | 83623 | 12485 | 4790 |
| 17,8 | 269,5 | 1,7 | 594,58234 | 593,57507 | [M+H]+ | true | Cer 16:1;O2/22:0 | C38H75NO3 | Spectral Library - | 20221 | | 270444,563 | true | 0 | 481 | 0 | | | 258318 | 272590 | 270438 | 254239 | 270445 | 3,13817195 | 356051 | 39830 | 238977 | 242480 | 329278 | 322955 | 305272 | 370150 | 327600 | 437625 | 105084 |
| 15,86 | 262,3 | 0,9 | 678,50942 | 660,47536 | [M+NH4]+, [M | true | DG20:5_20:5 | C43H64O5 | SmartFormula - r | 20222 | LipidBlast-VS69-Pos.ms | 1095599,88 | true | 0 | 143 | 0 | | | 1044675 | 1095600 | 1107234 | 1090090 | 1141542 | 3,18396656 | 395817 | 21654 | 328028 | 870544 | 1827133 | 1561670 | 1457490 | 2189577 | 1525071 | 2231971 | 544500 |
| 7,3 | 236,5 | 0,3 | 522,35555 | 521,34824 | [M+H]+, [M+N | true | LPC 18:1 | C26H52NO7P | Spectral Library - | 20222 | | 338561,469 | true | 0 | 0 | 0 | | | 338561 | 367327 | 348391 | 342614 | 354230 | 3,21098114 | 26840 | 1900 | 8802 | 17103 | 232246 | 167636 | 175098 | 743980 | 690709 | 756087 | 1304738 |
| 15,36 | 283 | -0,3 | 732,55364 | 731,54445 | [M+H]+, [M+N | true | PC 32:1 | C40H78NO8P | Spectral Library - | 20222 | | 784976,938 | true | 0 | 151 | 127 | | | 832906 | 855002 | 807636 | 784977 | 814205 | 3,23022185 | 68522 | 11964 | 71416 | 43262 | 68926 | 57904 | 31146 | 2398701 | 1785824 | 2694374 | 1774371 |
| 6,25 | 233,6 | 1 | 544,34 | 543,33272 | [M+H]+ | true | LPC 20:4 | C28H50NO7P | Spectral Library - | 20222 | | 518589,594 | true | 0 | 0 | 0 | | | 518590 | 542683 | 544500 | 533866 | 566643 | 3,2337714 | 13247 | 1147 | 5758 | 14972 | 151404 | 139612 | 148085 | 1597636 | 970456 | 1317064 | 2438467 |
| 5,41 | 230,2 | 0,6 | 542,32402 | 541,31675 | [M+H]+ | true | PCO-20:5 | C28H48NO7P | SmartFormula - r | 20122 | LipidBlast-VS69-Pos.ms | 1773566,63 | true | 0 | 0 | 0 | | | 1773567 | 1893052 | 1911366 | 1804052 | 1884668 | 3,27300796 | 28675 | 2031 | 10337 | 34237 | 386193 | 334058 | 349761 | 4986477 | 3344870 | 4300196 | 8143588 |
| 17,42 | 261,2 | 0,4 | 610,54066 | 592,50618 | [M+NH4]+, [M | true | DG 16:0_18:2 | C37H68O5 | SmartFormula - r | 20222 | | 304304,406 | true | 0 | 0 | 416 | | | 481561 | 491756 | 493358 | 482658 | 521541 | 3,27414774 | 124478 | 12738 | 108320 | 153184 | 260767 | 304304 | 232456 | 1911085 | 1310870 | 2008229 | 243729 |
| 16,44 | 258,8 | 2,5 | 540,53522 | 539,52693 | [M+H]+, [M+K] | true | Cer 16:0;O2/18:0 | C34H69NO3 | SmartFormula - r | 20221 | | 10644,2588 | true | 0 | 0 | 0 | | | 10644 | 11255 | 10744 | 10272 | 10677 | 3,28241925 | 28264 | 2925 | 18350 | 9694 | 14986 | 12244 | 14636 | 2412 | 5798 | 4185 | 0 |
| 6,73 | 237,6 | 0,1 | 570,35546 | 569,34801 | [M+H]+, [M+N | true | PCO-22:5 | C30H52NO7P | Spectral Library - | 20222 | LipidBlast-VS69-Pos.ms | 23298,3965 | true | 0 | 0 | 0 | | | 23298 | 25182 | 24599 | 23757 | 24948 | 3,290881 | 992 | 203 | 506 | 0 | 8330 | 8156 | 7419 | 73295 | 44711 | 60584 | 103600 |
| 4,79 | 223,4 | -0,3 | 492,30866 | 491,30169 | [M+H]+, [M+N | true | LPC 16:2 | C24H46NO7P | SmartFormula - r | 20222 | | 6627,53662 | true | 0 | 0 | 0 | | | 6930 | 6628 | 6923 | 6737 | 7242 | 3,39215462 | 294 | 0 | 111 | 262 | 1578 | 1890 | 1551 | 18704 | 9401 | 16402 | 30355 |
| 6,73 | 216,1 | -0,9 | 502,29303 | 501,28561 | [M+H]+, [M+N | true | LPE 20:4 | C25H44NO7P | SmartFormula - r | 20212 | | 29771,8379 | true | 0 | 0 | 0 | | | 29772 | 31642 | 31228 | 31298 | 32770 | 3,42489614 | 713 | 159 | 609 | 9534 | 2401 | 2107 | 2184 | 92129 | 75674 | 109179 | 111338 |
| 15,99 | 256,5 | 3 | 604,49358 | 586,45977 | [M+NH4]+, [M | true | DG 14:0_20:5 | C37H62O5 | SmartFormula - r | 20211 | | 147394,969 | true | 0 | 0 | 0 | | | 176345 | 193490 | 188339 | 190925 | 189381 | 3,53499003 | 23052 | 1621 | 19693 | 50087 | 80017 | 85914 | 74591 | 797240 | 404585 | 792056 | 147395 |
| 15,04 | 249,6 | 2,8 | 510,48866 | 509,48138 | [M+H]+ | true | Cer 16:1;O2/16:0 | C32H63NO3 | Lipid Species, Lip | 20221 | | 211282,234 | true | 0 | 0 | 0 | | | 226687 | 245853 | 242652 | 237356 | 228849 | 3,54507185 | 464094 | 51902 | 380993 | 151318 | 238454 | 211282 | 203149 | 89892 | 85218 | 83832 | 56259 |
| 16,39 | 264,8 | 1,1 | 680,52503 | 662,49087 | [M+NH4]+, [M | true | DG20:4_20:5 | C43H66O5 | SmartFormula - r | 20221 | LipidBlast-VS69-Pos.ms | 877022,313 | true | 0 | 128 | 0 | | | 837453 | 877022 | 900951 | 899511 | 921673 | 3,61133694 | 370553 | 20635 | 316668 | 820777 | 1633949 | 1508491 | 1370889 | 1617630 | 1166502 | 1813809 | 305991 |
| 16,91 | 263,4 | 1,5 | 632,52507 | 614,49106 | [M+NH4]+, [M | true | DG 16:0_20:5 | C39H66O5 | SmartFormula - r | 20211 | | 261534,766 | true | 0 | 0 | 0 | | | 392972 | 404264 | 404799 | 403878 | 432873 | 3,64871443 | 69190 | 8904 | 58756 | 97542 | 174506 | 185154 | 149795 | 1881645 | 1173950 | 1992841 | 261535 |
| 13,25 | 277,9 | -0,7 | 750,50623 | 749,49895 | [M+H]+ | true | PC 34:6 | C42H72NO8P | Lipid Species, Sm | 20222 | | 449,891357 | true | 0 | 0 | 0 | | | 3185 | 3126 | 3287 | 2973 | 3104 | 3,66791483 | 0 | 0 | 0 | 450 | 0 | 434 | 0 | 11160 | 0 | 10000 | 8390 |
| 18,58 | 329,4 | 2,9 | 876,67057 | 858,63673 | [M+NH4]+, [M | true | ST29:2;O;Hex;FA20:5 | C55H86O7 | SmartFormula - r | 20121 | LipidBlast-VS69-Pos.ms | 29915,4609 | true | 0 | 0 | 0 | | | 30307 | 31539 | 28630 | 30338 | 31186 | 3,69993048 | 20754 | 2991 | 16554 | 24203 | 37375 | 37482 | 30520 | 70937 | 66505 | 81738 | 4991 |
| 17,38 | 266,3 | 1,1 | 634,54048 | 616,50655 | [M+NH4]+, [M | true | DG 16:0_20:4 | C39H68O5 | SmartFormula - r | 20211 | | 79788,7109 | true | 0 | 335 | 259 | | | 133867 | 139916 | 143016 | 146021 | 147008 | 3,73999763 | 30198 | 3085 | 24501 | 45199 | 72337 | 79789 | 66359 | 577770 | 354467 | 597255 | 68774 |
| 11,48 | 215,9 | 2,8 | 413,26629 | 390,27707 | [M+Na]+ | true | Spectral Match to Dioctyl phthalate from NIST14 | C24H38O4 | SmartFormula - r | 20221 | ALL_GNPS_NO_PROPO | 5750,70605 | true | 1178 | 2693 | 3581 | | | 6668 | 7361 | 7102 | 6995 | 7237 | 3,74611502 | 7601 | 4942 | 11097 | 5751 | 5687 | 4814 | 4734 | 2898 | 3842 | 2920 | 15704 |
| 13,49 | 272,2 | 2 | 688,49159 | 687,48431 | [M+H]+ | true | PC13:0_16:2 | C37H70NO8P | Spectral Library - | 20121 | LipidBlast-VS69-Pos.ms | 2636,21899 | true | 0 | 0 | 0 | | | 2692 | 2926 | 2796 | 2935 | 2923 | 3,76155834 | 0 | 0 | 341 | 0 | 1147 | 330 | 0 | 4752 | 24268 | 4680 | 2636 |
| 17,84 | 280,5 | 1 | 668,61883 | 667,6111 | [M+H]+, [M+N | true | Cer18:1;3O/23:0;(2OH) | C41H81NO5 | Spectral Library - | 20212 | LipidBlast-VS69-Pos.ms | 12178,1357 | true | 0 | 0 | 0 | | | 12570 | 12178 | 12119 | 13180 | 12996 | 3,76871231 | 36906 | 6276 | 24044 | 12048 | 17139 | 18066 | 16567 | 2754 | 1894 | 2367 | 0 |
| 16,29 | 291,6 | | 782,56669 | 759,57749 | [M+Na]+, [M+H | true | PC 34:1 | C42H82NO8P | SmartFormula - r | 20110 | | 85443,6953 | true | 0 | 0 | 0 | | | 86789 | 86073 | 93698 | 88548 | 85444 | 3,78207315 | 18838 | 0 | 18056 | 0 | 13888 | 11126 | 0 | 316803 | 211006 | 261578 | 178089 |
| 17,61 | 301,3 | -0,9 | 840,64847 | 839,6412 | [M+H]+ | true | PC 40:3 | C48H90NO8P | SmartFormula - r | 20122 | | 11337,2129 | true | 0 | 0 | 0 | | | 12276 | 11476 | 11337 | 11494 | 12208 | 3,79695896 | 0 | 0 | 3618 | 0 | 0 | 0 | 0 | 46386 | 26886 | 43978 | 11548 |
| 17,01 | 262,6 | 2 | 566,55097 | 565,5437 | [M+H]+, [M+H | true | Cer 16:1;O2/20:0 | C36H71NO3 | Lipid Species, Sm | 20221 | | 159795,891 | true | 0 | 0 | 0 | | | 152920 | 151996 | 166026 | 158043 | 162623 | 3,83069352 | 305099 | 38108 | 211305 | 134263 | 208859 | 180387 | 176144 | 143883 | 172286 | 162274 | 41319 |
| 16,08 | 256,7 | 2,6 | 538,51973 | 537,51212 | [M+H]+, [M+H | true | Cer 16:1;O2/18:0 | C34H67NO3 | SmartFormula - r | 20211 | | 28725,1426 | true | 0 | 0 | 0 | | | 28725 | 27997 | 30008 | 29829 | 27426 | 3,90429101 | 56192 | 4262 | 41751 | 25205 | 42638 | 36913 | 36480 | 25492 | 31817 | 27029 | 8956 |
| 15,61 | 282,2 | 0,6 | 790,53728 | 789,53 | [M+H]+ | true | PE 40:7 | C45H76NO8P | Spectral Library - | 20222 | | 36556,8555 | true | 0 | 0 | 0 | | | 46262 | 42186 | 43111 | 45902 | 44098 | 3,96355452 | 11478 | 1602 | 13972 | 13979 | 29136 | 25526 | 23169 | 127904 | 131181 | 125750 | 36557 |
| 14,18 | 283,9 | -0,7 | 778,53777 | 777,5304 | [M+H]+, [M+K] | true | PC 36:6 | C44H76NO8P | SmartFormula - r | 20222 | | 282739,531 | true | 0 | 0 | 0 | | | 289578 | 310699 | 282740 | 287483 | 303271 | 3,98105219 | 17392 | 2306 | 14495 | 22379 | 32453 | 34407 | 26144 | 844701 | 642612 | 1034809 | 716789 |
| 15,21 | 290,9 | -0,3 | 806,56973 | 805,56168 | [M+H]+, [M+N | true | PC 38:6 | C46H80NO8P | Lipid Species, Sp | 20222 | | 2668659,5 | true | 0 | 351 | 222 | | | 2703923 | 2922329 | 2814610 | 2892266 | 2668660 | 3,99929779 | 333416 | 46100 | 325116 | 391550 | 683811 | 657481 | 547723 | 6998292 | 5795910 | 7199451 | 5253519 |
| 15,33 | 297 | 0 | 856,58393 | 855,57665 | [M+H]+ | true | PC16:3_26:6 | C50H82NO8P | Spectral Library - | 20222 | LipidBlast-VS69-Pos.ms | 115143,453 | true | 0 | 0 | 0 | | | 121564 | 124750 | 128624 | 115143 | 122572 | 4,03068607 | 14795 | 2012 | 13335 | 16993 | 25437 | 25685 | 21676 | 320730 | 280199 | 329357 | 203530 |
| 5,54 | 224,4 | -0,1 | 468,30878 | 467,3015 | [M+H]+ | true | LPC 14:0 | C22H46NO7P | Spectral Library - | 20222 | | 12880,083 | true | 0 | 0 | 0 | | | 13897 | 14640 | 14484 | 14025 | 13192 | 4,05298598 | 495 | 0 | 556 | 3015 | 0 | 1227 | 750 | 43875 | 30391 | 46360 | 52492 |
| 1,9 | 155,1 | -0,8 | 268,10418 | 267,09691 | [M+H]+ | true | Adenosine | C10H13N5O4 | Spectral Library - | 20222 | ALL_GNPS_NO_PROPO | 284,541229 | true | 0 | 0 | 0 | | | 2406 | 2282 | 2290 | 2237 | 2463 | 4,05400189 | 260 | 0 | 285 | 0 | 0 | 0 | 0 | 4129 | 6713 | 6732 | 10963 |
| 14,13 | 281,1 | 0,8 | 764,52165 | 763,51438 | [M+H]+ | true | PC11:0_24:6 | C43H74NO8P | Spectral Library - | 20222 | LipidBlast-VS69-Pos.ms | 50204,1641 | true | 0 | 0 | 0 | | | 63220 | 64941 | 63952 | 60368 | 67503 | 4,05658094 | 2277 | 1355 | 6452 | 3684 | 31817 | 4776 | 3640 | 84457 | 513681 | 71932 | 50204 |
| 17,95 | 264,7 | 0,3 | 612,55612 | 594,52225 | [M+NH4]+, [M | true | DG 16:0_18:1 | C37H70O5 | SmartFormula - r | 20222 | | 163508,422 | true | 0 | 0 | 1327 | | | 200331 | 212980 | 200143 | 198264 | 216405 | 4,08748378 | 79676 | 9872 | 65728 | 96053 | 149728 | 163508 | 141979 | 612984 | 451518 | 717001 | 97045 |
| 16,54 | 254,6 | 2 | 582,50948 | 564,47527 | [M+NH4]+, [M | true | DG 14:0_18:2 | C35H64O5 | SmartFormula - r | 20221 | | 130176,086 | true | 0 | 254 | 0 | | | 146229 | 160438 | 157484 | 154228 | 162560 | 4,09172183 | 24620 | 3043 | 21336 | 34094 | 77755 | 67130 | 70464 | 606676 | 369240 | 643554 | 130176 |
| 14,8 | 248,1 | 1,7 | 588,46229 | 570,42826 | [M+NH4]+, [M | true | DG13:1_20:5 | C36H58O5 | Spectral Library - | 20211 | LipidBlast-VS69-Pos.ms | 3126,02441 | true | 0 | 0 | 0 | | | 27593 | 27726 | 27490 | 30170 | 27329 | 4,23189967 | 1700 | 3126 | 14882 | 592 | 116260 | 1641 | 578 | 0 | 138854 | 0 | 0 |
| 11,47 | 211,7 | 2,7 | 391,28456 | 390,27729 | [M+H]+ | true | Spectral Match to Diisooctyl phthalate from NIST14 | C24H38O4 | SmartFormula - r | 20221 | ALL_GNPS_NO_PROPO | 7029,02295 | true | 0 | 3190 | 4620 | | | 8986 | 9277 | 8430 | 9292 | 9355 | 4,23528159 | 10333 | 7029 | 13754 | 6595 | 6857 | 6683 | 6364 | 3685 | 5422 | 4143 | 20100 |
| 16,34 | 289,4 | -0,2 | 760,5859 | 759,57863 | [M+H]+ | true | PC 16:0_18:1 | C42H82NO8P | Spectral Library - | 20222 | | 2069267,63 | true | 0 | 500 | 274 | | | 2069268 | 2328885 | 2189296 | 2190065 | 2162716 | 4,25006575 | 328071 | 46521 | 344470 | 153069 | 294033 | 238947 | 165745 | 6982505 | 5798093 | 7455634 | 3189154 |
| 16,27 | 260,3 | 2,1 | 642,50891 | 624,47499 | [M+NH4]+, [M | true | DG17:2_20:5 | C40H64O5 | SmartFormula - r | 20221 | LipidBlast-VS69-Pos.ms | 71377,9922 | true | 0 | 0 | 0 | | | 78133 | 77165 | 83572 | 75485 | 81931 | 4,26178335 | 7506 | 2523 | 16903 | 7082 | 130126 | 12587 | 11095 | 81684 | 664401 | 71378 | 17675 |
| 13,87 | 288,3 | -0,2 | 826,53903 | 825,53175 | [M+H]+ | true | PC20:5_20:5 | C48H76NO8P | SmartFormula - r | 20122 | LipidBlast-VS69-Pos.ms | 10291447 | true | 0 | 1012 | 0 | | | 10291447 | 11246172 | 10767153 | 11055517 | 11512035 | 4,2711847 | 739181 | 95009 | 663434 | 963127 | 1668442 | 1434745 | 1201282 | 22516808 | 22114040 | 25008760 | 21657594 |
| 15,69 | 288,1 | 0,4 | 770,56872 | 769,56186 | [M+H]+, [M+N | true | PC11:0_24:3 | C43H80NO8P | Spectral Library - | 20222 | LipidBlast-VS69-Pos.ms | 158363,359 | true | 0 | 0 | 0 | | | 374649 | 373784 | 372417 | 342514 | 348788 | 4,27603489 | 10656 | 9171 | 60463 | 13275 | 244574 | 20545 | 14181 | 169629 | 3135502 | 158363 | 93390 |
| 13,06 | 282,9 | 0 | 798,50598 | 797,50052 | [M+H]+, [M+N | true | PC18:5_20:5 | C46H72NO8P | SmartFormula - r | 20222 | LipidBlast-VS69-Pos.ms | 6859,96338 | true | 0 | 0 | 0 | | | 7104 | 6860 | 6887 | 7569 | 7362 | 4,28240207 | 301 | 0 | 335 | 416 | 890 | 482 | 498 | 20913 | 22393 | 25428 | 18104 |
| 13,98 | 279,9 | -0,7 | 752,52194 | 751,51467 | [M+H]+ | true | PC 34:5 | C42H74NO8P | SmartFormula - r | 20222 | | 873333,188 | true | 0 | 141 | 0 | | | 873333 | 963042 | 904605 | 949613 | 962299 | 4,28967913 | 51847 | 6777 | 43624 | 53837 | 74178 | 66547 | 39642 | 2337952 | 1878681 | 2455393 | 2024778 |
| 12,62 | 276,7 | 2,1 | 736,49103 | 735,48375 | [M+H]+ | true | PC13:1_20:5 | C41H70NO8P | SmartFormula - r | 20121 | LipidBlast-VS69-Pos.ms | 177,286758 | true | 0 | 0 | 0 | | | 4447 | 4282 | 4580 | 4793 | 4416 | 4,29375328 | 0 | 177 | 628 | 0 | 3364 | 0 | 0 | 0 | 44807 | 0 | 0 |
| 16,19 | 286,8 | 0,3 | 734,56864 | 733,56136 | [M+H]+ | true | PC 32:0 | C40H80NO8P | Spectral Library - | 20222 | | 19023,3828 | true | 0 | 0 | 0 | | | 19023 | 21071 | 19062 | 19354 | 19414 | 4,33271363 | 6888 | 579 | 5323 | 0 | 0 | 0 | 2604 | 73981 | 62506 | 73240 | 33157 |
| 15,42 | 253,3 | 3,4 | 512,50385 | 511,49657 | [M+H]+ | true | Cer 16:0;O2/16:0 | C32H65NO3 | Lipid Species, Lip | 20220 | | 115278,195 | true | 0 | 0 | 0 | | | 111740 | 119928 | 125314 | 119570 | 123740 | 4,38033859 | 353606 | 48480 | 265278 | 96195 | 122305 | 127601 | 115278 | 18657 | 19881 | 16325 | 6999 |
| 15,28 | 288,1 | 0,9 | 770,56898 | 769,5617 | [M+H]+ | true | PC17:1_18:2 | C43H80NO8P | Spectral Library - | 20222 | LipidBlast-VS69-Pos.ms | 5534,6582 | true | 0 | 0 | 0 | | | 183270 | 193007 | 184795 | 174520 | 194722 | 4,38153499 | 2557 | 5535 | 26102 | 0 | 100773 | 0 | 0 | 0 | 1745619 | 0 | 0 |
| 5,61 | 228 | 0 | 494,32442 | 493,31729 | [M+H]+, [M+N | true | LPC 16:1 | C24H48NO7P | Spectral Library - | 20222 | | 65913,6016 | true | 0 | 0 | 0 | | | 65914 | 70211 | 74526 | 69296 | 70112 | 4,38686228 | 4155 | 368 | 909 | 2554 | 23908 | 22699 | 21697 | 195293 | 107312 | 177982 | 316619 |
| 5,52 | 216,4 | 2,9 | 400,34227 | 399,335 | [M+H]+ | true | C16Hexadecanoyl-L-Carnitine | C23H45NO4 | Spectral Library - | 20211 | ALL_GNPS_NO_PROPO | 33896,6523 | true | 0 | 0 | 0 | | | 33897 | 35862 | 37756 | 34109 | 35624 | 4,39910936 | 1214 | 258 | 1107 | 1367 | 2341 | 1449 | 1538 | 115927 | 90002 | 112698 | 108678 |
| 7,45 | 220,4 | 0,7 | 480,30859 | 479,30132 | [M+H]+ | true | LPE 18:1 | C23H46NO7P | Spectral Library - | 20222 | | 4812,67773 | true | 0 | 0 | 0 | | | 4911 | 5365 | 4813 | 4935 | 4882 | 4,40400478 | 640 | 134 | 511 | 412 | 3855 | 2975 | 2847 | 10938 | 12008 | 12542 | 15445 |
| 18,46 | 288,2 | | 684,65006 | 683,64262 | [M+H]+, [M+K] | true | Cer 18:0;O3/24:0;O | C42H85NO5 | SmartFormula - r | 20220 | | 28784,6895 | true | 0 | 846 | 569 | | | 28785 | 30441 | 27498 | 27307 | 29387 | 4,57696674 | 46313 | 33047 | 35159 | 41720 | 41968 | 47290 | 53546 | 3899 | 6223 | 6925 | 1490 |
| 14,03 | 285,9 | -0,7 | 802,53771 | 801,53044 | [M+H]+ | true | PC 38:8 | C46H76NO8P | Spectral Library - | 20222 | | 184595,469 | true | 0 | 0 | 0 | | | 204382 | 207817 | 184595 | 202494 | 204582 | 4,60441534 | 10306 | 2024 | 12126 | 14918 | 28995 | 21349 | 18328 | 477041 | 518357 | 482076 | 429020 |
| 15,65 | 289,8 | 0 | 782,56949 | 781,56221 | [M+H]+ | true | PC 36:4 | C44H80NO8P | Spectral Library - | 20222 | | 1900827,75 | true | 0 | 0 | 148 | | | 1940586 | 2076513 | 2023955 | 1900828 | 2125373 | 4,61966508 | 181392 | 22908 | 196796 | 135397 | 175814 | 178067 | 92003 | 5889146 | 4224482 | 6015931 | 3265988 |
| 16,67 | 298,2 | -0,6 | 836,61653 | 835,60926 | [M+H]+ | true | PC 40:5 | C48H86NO8P | Spectral Library - | 20212 | | 23529,4219 | true | 0 | 0 | 0 | | | 26314 | 23529 | 24880 | 24633 | 26219 | 4,65165791 | 0 | 0 | 0 | 4630 | 8388 | 6190 | 8199 | 88497 | 74056 | 94568 | 37689 |
| 15,76 | 288,3 | 0,8 | 746,56876 | 745,56148 | [M+H]+ | true | PC14:0_19:1 | C41H80NO8P | SmartFormula - r | 20222 | LipidBlast-VS69-Pos.ms | 18603,0762 | true | 0 | 0 | 0 | | | 68196 | 74525 | 77047 | 74477 | 71333 | 4,67203491 | 1925 | 2890 | 14514 | 0 | 28989 | 1690 | 0 | 0 | 579185 | 35377 | 18603 |
| 15,28 | 278,2 | 0,7 | 764,52214 | 763,5151 | [M+H]+, [M+N | true | PE 38:6 | C43H74NO8P | Spectral Library - | 20222 | | 129026,766 | true | 0 | 0 | 0 | | | 133729 | 140474 | 132263 | 129027 | 144914 | 4,750881 | 32918 | 5029 | 33482 | 28336 | 62045 | 53517 | 44036 | 387169 | 350338 | 470832 | 146768 |
| 14,15 | 279,9 | 0 | 728,52249 | 727,51521 | [M+H]+ | true | PC 32:3 | C40H74NO8P | Spectral Library - | 20222 | | 25829,9531 | true | 0 | 0 | 0 | | | 28403 | 25830 | 28464 | 27981 | 25992 | 4,80541273 | 1230 | 475 | 2554 | 1254 | 10501 | 0 | 1034 | 42993 | 180407 | 55726 | 40538 |
| 17,15 | 257,3 | 1,6 | 584,52484 | 566,49108 | [M+NH4]+, [M | true | DG 14:0_18:1 | C35H66O5 | Spectral Library - | 20211 | | 75356,4531 | true | 0 | 741 | 0 | | | 106204 | 115697 | 115567 | 119361 | 120307 | 4,83230161 | 29646 | 4478 | 27581 | 37207 | 71180 | 66698 | 74408 | 375827 | 258839 | 430947 | 75356 |
| 14,38 | 285 | 0,8 | 790,53867 | 789,5314 | [M+H]+ | true | PC17:2_20:5 | C45H76NO8P | SmartFormula - r | 20122 | LipidBlast-VS69-Pos.ms | 591352 | true | 0 | 0 | 0 | | | 1155819 | 1221621 | 1245888 | 1297451 | 1303816 | 4,87135976 | 23147 | 18217 | 94734 | 33841 | 591352 | 48266 | 34579 | 860434 | 11003098 | 781210 | 460319 |
| 17,3 | 265,9 | 2,4 | 568,56664 | 567,5593 | [M+H]+, [M+N | true | Cer 16:0;O2/20:0 | C36H73NO3 | Lipid Species, Sm | 20221 | | 20109,9473 | true | 0 | 0 | 0 | | | 18987 | 20445 | 20322 | 19975 | 21740 | 4,88089419 | 65682 | 8329 | 42113 | 20110 | 22118 | 25367 | 24272 | 3309 | 6117 | 3779 | 0 |
| 16,89 | 267,5 | -0,8 | 682,54031 | 664,50637 | [M+NH4]+, [M | true | DG 20:4/20:4 | C43H68O5 | Lipid Species, Sp | 20212 | | 183445,25 | true | 0 | 0 | 0 | | | 171082 | 180193 | 187458 | 183445 | 195398 | 4,89189696 | 85834 | 6877 | 69326 | 174599 | 283856 | 278861 | 247228 | 415572 | 219256 | 368747 | 48145 |
| 16,82 | 258,4 | 1,8 | 620,52467 | 602,49075 | [M+NH4]+, [M | true | DG17:2_18:2 | C38H66O5 | SmartFormula - r | 20211 | LipidBlast-VS69-Pos.ms | 87468,8906 | true | 0 | 0 | 0 | | | 87469 | 96738 | 90690 | 92651 | 98945 | 4,93854661 | 7557 | 4252 | 21979 | 7685 | 150869 | 13616 | 12032 | 95524 | 942130 | 91868 | 12405 |
| 15,95 | 277,2 | 0,6 | 742,53726 | 741,52998 | [M+H]+ | true | PE 36:3 | C41H76NO8P | Spectral Library - | 20222 | | 66494,25 | true | 0 | 0 | 0 | | | 76619 | 80352 | 87671 | 81735 | 83419 | 4,95264619 | 9947 | 1142 | 11994 | 6174 | 20609 | 9925 | 7447 | 238454 | 312346 | 325932 | 66494 |
| 15,29 | 274 | 0,1 | 740,52223 | 739,51495 | [M+H]+ | true | PE 36:4 | C41H74NO8P | Spectral Library - | 20222 | | 127173,797 | true | 0 | 0 | 0 | | | 143902 | 133647 | 141014 | 127174 | 132689 | 4,96428367 | 10012 | 1336 | 9430 | 6925 | 12089 | 9013 | 6733 | 461388 | 371127 | 523529 | 169458 |
| 16,47 | 262 | 0,6 | 590,54987 | 589,5426 | [M+H]+ | true | Cer 16:1;O2/22:2 | C38H71NO3 | Lipid Species, Lip | 20222 | | 7481,73438 | true | 0 | 0 | 0 | | | 7421 | 8329 | 7968 | 7398 | 7917 | 5,07067829 | 10111 | 1507 | 7482 | 6865 | 14001 | 10087 | 9091 | 6722 | 10589 | 13050 | 3538 |
| 15,74 | 274,2 | 1,1 | 716,52241 | 715,51513 | [M+H]+ | true | PE 34:2 | C39H74NO8P | SmartFormula - r | 20221 | | 58961,957 | true | 0 | 0 | 0 | | | 61231 | 58962 | 61911 | 61278 | 67458 | 5,08657903 | 20563 | 2529 | 21005 | 9940 | 17345 | 16332 | 11341 | 208196 | 143157 | 216221 | 69842 |
| 14,62 | 282,8 | -0,4 | 754,53738 | 753,53011 | [M+H]+ | true | PC 34:4 | C42H76NO8P | SmartFormula - r | 20222 | | 26437,9414 | true | 0 | 0 | 0 | | | 268910 | 305711 | 277996 | 279760 | 294205 | 5,10492465 | 26438 | 2820 | 22590 | 20427 | 0 | 23876 | 17097 | 636600 | 0 | 623916 | 530487 |
| 13,87 | 289,9 | -0,3 | 848,5196 | 825,53038 | [M+Na]+ | true | PC40:10 | C48H76NO8P | SmartFormula - r | 20212 | LipidBlast-VS69-Pos.ms | 436135,938 | true | 0 | 0 | 0 | | | 456352 | 436136 | 492050 | 436559 | 445087 | 5,11899213 | 22900 | 2895 | 20816 | 32327 | 51674 | 46631 | 38918 | 530798 | 528082 | 532727 | 514163 |
| 15,64 | 256 | 0,7 | 548,50407 | 547,49669 | [M+H]+, [M+H | true | Cer19:3;2O/16:0 | C35H65NO3 | Spectral Library - | 20212 | LipidBlast-VS69-Pos.ms | 248132,922 | true | 0 | 0 | 0 | | | 364525 | 377508 | 416477 | 395392 | 401096 | 5,19952604 | 176977 | 16045 | 167790 | 128302 | 241908 | 245174 | 230648 | 589102 | 648859 | 1008289 | 248133 |
| 14,04 | 276,1 | -0,3 | 784,49166 | 783,4839 | [M+H]+, [M+N | true | PE20:5_20:5 | C45H70NO8P | SmartFormula - r | 20122 | LipidBlast-VS69-Pos.ms | 1625185,5 | true | 0 | 239 | 0 | | | 1625880 | 1695938 | 1667263 | 1625186 | 1839052 | 5,21368227 | 250845 | 33953 | 288103 | 263300 | 598706 | 485326 | 395328 | 4840161 | 5880526 | 5690068 | 2435183 |
| 14,28 | 264,4 | 0,1 | 698,47399 | 697,46671 | [M+H]+ | true | PE17:0_16:4 | C38H68NO8P | SmartFormula - r | 10122 | LipidBlast-VS69-Pos.ms | 7254,66602 | true | 0 | 0 | 0 | | | 9818 | 10944 | 10502 | 10831 | 11296 | 5,22813511 | 727 | 512 | 2381 | 536 | 7255 | 902 | 652 | 14430 | 97271 | 14723 | 6205 |
| 16,79 | 295,3 | -0,9 | 812,6155 | 811,60822 | [M+H]+ | true | PC 38:3 | C46H86NO8P | SmartFormula - r | 20222 | | 24769,5371 | true | 0 | 0 | 0 | | | 25427 | 28399 | 24770 | 26534 | 25983 | 5,26833981 | 10292 | 0 | 7644 | 0 | 6091 | 6236 | 5036 | 90930 | 70395 | 87605 | 34699 |
| 15,2 | 281 | 0,9 | 788,52181 | 787,51453 | [M+H]+ | true | PE 40:8 | C45H74NO8P | Spectral Library - | 20212 | | 286370,094 | true | 0 | 0 | 0 | | | 306137 | 322801 | 286370 | 291140 | 318310 | 5,27353801 | 74606 | 10173 | 78467 | 71980 | 136309 | 127777 | 102652 | 911250 | 777464 | 873515 | 366275 |
| 6,38 | 214 | -1,2 | 478,29299 | 477,28572 | [M+H]+ | true | LPE 18:2 | C23H44NO7P | Spectral Library - | 20221 | | 10041,085 | true | 0 | 0 | 0 | | | 10041 | 11421 | 10730 | 11010 | 11433 | 5,27861405 | 1031 | 124 | 422 | 0 | 6965 | 5877 | 5761 | 28882 | 22025 | 31806 | 36509 |
| 13,56 | 282,1 | -0,6 | 776,52181 | 775,51453 | [M+H]+ | true | PC 36:7 | C44H74NO8P | SmartFormula - r | 20222 | | 37586,8281 | true | 0 | 0 | 0 | | | 37587 | 39597 | 42836 | 40203 | 42424 | 5,31246546 | 1790 | 286 | 1734 | 2724 | 3851 | 3889 | 3101 | 121709 | 93079 | 138892 | 113794 |
| 14,24 | 277 | -0,3 | 704,52216 | 703,51488 | [M+H]+ | true | PC 30:1 | C38H74NO8P | Spectral Library - | 20222 | | 50574,9102 | true | 0 | 0 | 0 | | | 51365 | 57409 | 50575 | 52325 | 55208 | 5,34975195 | 2621 | 466 | 2856 | 2190 | 3033 | 0 | 1779 | 166025 | 107323 | 199953 | 152740 |
| 15,42 | 270,2 | 1,3 | 690,50661 | 689,49933 | [M+H]+ | true | PE 32:1 | C37H72NO8P | Spectral Library - | 20221 | | 9299 | true | 0 | 0 | 0 | | | 9785 | 10546 | 9299 | 9978 | 10584 | 5,38742932 | 4579 | 854 | 3157 | 1523 | 2533 | 2227 | 1757 | 30309 | 26917 | 39181 | 15700 |
| 5,92 | 219,2 | | 426,35826 | 425,35099 | [M+H]+ | true | Massbank:RP025101 Oleoyl-L-Carnitine\|(3R)-3-[(Z)-octadec-9-e | C25H47NO4 | SmartFormula - r | 20210 | ALL_GNPS_NO_PROPO | 7526,59229 | true | 0 | 0 | 0 | | | 7527 | 8518 | 8671 | 8091 | 8277 | 5,41639268 | 422 | 0 | 328 | 614 | 740 | 570 | 459 | 26768 | 22405 | 31510 | 25374 |
| 18,08 | 266,7 | -1,5 | 638,57151 | 620,5381 | [M+NH4]+, [M | true | DG 18:1/18:1 | C39H72O5 | Lipid Species, Sp | 20221 | | 26001,0371 | true | 0 | 2198 | 1579 | | | 29743 | 32718 | 33055 | 30473 | 33784 | 5,47130395 | 0 | 7257 | 13034 | 16103 | 17988 | 23413 | 26001 | 97817 | 43733 | 119135 | 12901 |
| 14,67 | 275,5 | 0,5 | 762,50637 | 761,49909 | [M+H]+ | true | PE 38:7 | C43H72NO8P | Spectral Library - | 20222 | | 249673,625 | true | 0 | 0 | 0 | | | 257899 | 282107 | 282494 | 249674 | 273628 | 5,48518913 | 51489 | 7303 | 53328 | 44788 | 94739 | 83098 | 64690 | 706526 | 707559 | 773321 | 354076 |
| 17,38 | 269,1 | 1,6 | 594,58185 | 593,57428 | [M+H]+, [M+N | true | Cer 16:0;O2/22:1 | C38H75NO3 | Lipid Species, Lip | 20211 | | 49658,0391 | true | 0 | 0 | 0 | | | 46830 | 50154 | 54392 | 50119 | 52068 | 5,49961829 | 129359 | 20715 | 105131 | 49180 | 73814 | 62339 | 62165 | 0 | 0 | 0 | 0 |
| 13,42 | 284,5 | 0,3 | 800,52205 | 799,51478 | [M+H]+ | true | PC18:4_20:5 | C46H74NO8P | SmartFormula - r | 20222 | LipidBlast-VS69-Pos.ms | 21583,0195 | true | 0 | 0 | 0 | | | 21583 | 22698 | 23348 | 23210 | 25172 | 5,61014669 | 1041 | 168 | 959 | 1653 | 2594 | 1998 | 1737 | 78611 | 56499 | 87660 | 60628 |
| 12,49 | 275,5 | -0,8 | 748,49054 | 747,48327 | [M+H]+ | true | PC 34:7 | C42H70NO8P | Spectral Library - | 20222 | | 1059,8031 | true | 0 | 0 | 0 | | | 1091 | 1213 | 1260 | 1136 | 1157 | 5,64739706 | 0 | 0 | 0 | 155 | 0 | 0 | 142 | 4393 | 1327 | 3502 | 3240 |
| 8,61 | 204,5 | -0,9 | 401,34156 | 400,334 | [M+H]+, [M+H | true | 7-Oxocholesterol | C27H44O2 | Spectral Library - | 20112 | MoNA-export-LC-MS-MS | 392,097443 | true | 0 | 0 | 0 | | | 522 | 469 | 529 | 470 | 498 | 5,64957293 | 604 | 161 | 397 | 253 | 1826 | 1169 | 0 | 0 | 134 | 112 | 0 |
| 14,74 | 294,9 | 0,1 | 854,56872 | 853,56144 | [M+H]+ | true | PC18:5_24:5 | C50H80NO8P | Spectral Library - | 20222 | LipidBlast-VS69-Pos.ms | 216717,641 | true | 0 | 0 | 0 | | | 243857 | 242375 | 216718 | 245420 | 252203 | 5,66760712 | 20985 | 2659 | 19590 | 21718 | 41301 | 37271 | 29112 | 646168 | 574854 | 691187 | 496008 |
| 5,41 | 233,4 | 0,3 | 564,30583 | 541,31661 | [M+Na]+ | true | LPC20:5 | C28H48NO7P | SmartFormula - r | 20222 | LipidBlast-VS69-Pos.ms | 94603,7344 | true | 0 | 0 | 0 | | | 105043 | 109160 | 97536 | 94604 | 99859 | 5,77753373 | 1355 | 208 | 633 | 1733 | 18444 | 16699 | 16488 | 388782 | 206542 | 272176 | 561120 |
| 14,27 | 267,8 | 1,3 | 722,47423 | 721,46695 | [M+H]+ | true | PE9:0_26:6 | C40H68NO8P | SmartFormula - r | 20111 | LipidBlast-VS69-Pos.ms | 6689,4458 | true | 0 | 0 | 0 | | | 8049 | 8602 | 8457 | 8093 | 7385 | 5,8171381 | 628 | 427 | 2024 | 489 | 6689 | 908 | 664 | 11542 | 70181 | 13343 | 5981 |
| 17,05 | 299,2 | -0,4 | 838,63133 | 837,62405 | [M+H]+ | true | PC24:0_16:4 | C48H88NO8P | SmartFormula - r | 20212 | LipidBlast-VS69-Pos.ms | 11993,585 | true | 0 | 0 | 0 | | | 13455 | 13432 | 13476 | 14102 | 11994 | 5,85298236 | 3944 | 762 | 3728 | 2194 | 3343 | 3715 | 3001 | 44579 | 26137 | 51542 | 17537 |
| 16,14 | 302,5 | 0,2 | 884,61579 | 883,60852 | [M+H]+ | true | PC14:1_30:8 | C52H86NO8P | SmartFormula - r | 20122 | LipidBlast-VS69-Pos.ms | 9011,18555 | true | 0 | 0 | 0 | | | 9011 | 9732 | 10169 | 10400 | 9284 | 5,99261553 | 3254 | 534 | 3310 | 3010 | 4287 | 4294 | 3538 | 29663 | 21737 | 29069 | 11187 |
| 17,85 | 272,9 | 1 | 620,59759 | 619,59025 | [M+H]+, [M+N | true | Cer 16:1;O2/24:1 | C40H77NO3 | SmartFormula - r | 20221 | | 48945,1523 | true | 0 | 0 | 0 | | | 46039 | 47344 | 45413 | 48945 | 52962 | 6,26489858 | 93776 | 11774 | 69359 | 49328 | 73959 | 66022 | 61792 | 36526 | 37890 | 49481 | 12084 |
| 14,73 | 267,6 | 1,1 | 688,49305 | 687,48578 | [M+H]+ | true | PE 32:2 | C37H70NO8P | Spectral Library - | 10121 | | 11060,8291 | true | 0 | 0 | 0 | | | 12364 | 13071 | 11705 | 11061 | 12429 | 6,32496659 | 2769 | 367 | 2193 | 1835 | 2073 | 2503 | 1078 | 37658 | 29159 | 44748 | 18286 |
| 6,37 | 234,6 | 1,8 | 508,33997 | 507,33269 | [M+H]+ | true | PCO-17:1 | C25H50NO7P | SmartFormula - r | 20221 | LipidBlast-VS69-Pos.ms | 1801,22107 | true | 0 | 0 | 0 | | | 10547 | 11249 | 10667 | 9639 | 11314 | 6,32648174 | 227 | 164 | 703 | 0 | 24959 | 556 | 474 | 1574 | 97754 | 1801 | 3808 |
| 15,89 | 280,7 | 1 | 766,53754 | 765,53026 | [M+H]+ | true | PE 38:5 | C43H76NO8P | Spectral Library - | 20222 | | 20336,1855 | true | 0 | 0 | 0 | | | 22589 | 22896 | 24154 | 21791 | 25614 | 6,39735151 | 9185 | 1445 | 9909 | 9647 | 15950 | 14780 | 12271 | 62300 | 59567 | 74739 | 20336 |
| 13,06 | 260,3 | 2,2 | 656,42897 | 655,42169 | [M+H]+ | true | PE 15:2/15:2 | C35H62NO8P | Spectral Library - | 20221 | | 475,016785 | true | 0 | 0 | 0 | | | 941 | 964 | 908 | 1075 | 963 | 6,47687924 | 0 | 0 | 484 | 0 | 1072 | 0 | 0 | 475 | 7167 | 407 | 208 |
| 18,46 | 281,8 | | 664,6239 | 663,61659 | [M+H]+, [M+N | true | Cer 18:2;O2/24:0;O | C42H81NO4 | Lipid Species, Lip | 20110 | | 6176,01465 | true | 0 | 0 | 0 | | | 6355 | 6913 | 5978 | 5886 | 6176 | 6,49640952 | 16493 | 2996 | 10995 | 7234 | 9967 | 11652 | 9906 | 300 | 0 | 0 | 0 |
| 12,48 | 278 | 1 | 772,49023 | 771,48296 | [M+H]+ | true | PC16:4_20:5 | C44H70NO8P | Spectral Library - | 20221 | LipidBlast-VS69-Pos.ms | 3058,94092 | true | 0 | 0 | 0 | | | 3081 | 3561 | 3059 | 3400 | 3399 | 6,67293094 | 0 | 0 | 0 | 321 | 0 | 409 | 245 | 11534 | 6007 | 11057 | 9462 |
| 2,83 | 187,2 | 4,3 | 288,28982 | 287,28254 | [M+H]+ | true | Spectral Match to Heptadecasphinganine from NIST14 | C17H37NO2 | SmartFormula - r | 20220 | ALL_GNPS_NO_PROPO | 1237,10144 | true | 0 | 0 | 287 | | | 3439 | 4098 | 3672 | 3928 | 3884 | 6,68713064 | 804 | 710 | 2362 | 1237 | 1207 | 942 | 4659 | 2183 | 341 | 0 | 989 |
| 4,94 | 227,8 | 1,1 | 480,30862 | 479,30134 | [M+H]+ | true | PCO-15:1 | C23H46NO7P | SmartFormula - r | 20121 | LipidBlast-VS69-Pos.ms | 173,830078 | true | 0 | 0 | 0 | | | 1078 | 1265 | 1286 | 1198 | 1218 | 6,72319724 | 0 | 0 | 0 | 0 | 1772 | 174 | 142 | 0 | 8586 | 725 | 0 |
| 2,48 | 216,6 | | 448,34197 | 447,33469 | [M+H]+ | true | N-((4R)-4-((3R,5S,9S,10S,13R,14S,17R)-3-hydroxy-10,13-dimeth | C27H45NO4 | Spectral Library - | 20110 | ALL_GNPS_NO_PROPO | 478,788452 | true | 0 | 0 | 0 | | | 479 | 518 | 493 | 432 | 462 | 6,78863135 | 501 | 186 | 607 | 492 | 248 | 280 | 176 | 621 | 741 | 930 | 813 |
| 18,08 | 271,3 | 1,5 | 596,5972 | 595,58993 | [M+H]+ | true | Cer 16:0;O2/22:0 | C38H77NO3 | Lipid Species, Lip | 20221 | | 25276,4668 | true | 0 | 0 | 0 | | | 22669 | 25276 | 27124 | 25621 | 23963 | 6,78879139 | 65052 | 10841 | 48052 | 31501 | 33154 | 39670 | 34409 | 11004 | 12430 | 0 | 0 |
| 18,85 | 275,6 | -4,3 | 692,61892 | 674,58402 | [M+NH4]+, [M | true | DG 18:2_22:1 | C43H78O5 | Lipid Species, Sm | 20210 | | 7282,07715 | true | 0 | 0 | 0 | | | 6713 | 7662 | 7798 | 7951 | 7945 | 6,79644308 | 3782 | 0 | 2629 | 5918 | 5597 | 8187 | 8727 | 23857 | 9015 | 25733 | 2354 |
| 9,34 | 205,4 | | 357,30017 | 356,29289 | [M+H]+ | true | Monoolein CollisionEnergy:102040 | C21H40O4 | Spectral Library - | 20210 | ALL_GNPS_NO_PROPO | 3582,89575 | true | 300 | 270 | 0 | | | 3814 | 4248 | 4046 | 3583 | 3706 | 6,89050643 | 0 | 0 | 283 | 0 | 2750 | 1970 | 2292 | 9532 | 6910 | 9541 | 10579 |
| 13,51 | 268,5 | | 606,54561 | 605,53833 | [M+H]+ | true | Cer 16:1;O2/22:2;O | C38H71NO4 | Lipid Species, Lip | 20210 | | 27189,8574 | true | 0 | 0 | 0 | | | 24726 | 27573 | 28736 | 25383 | 28767 | 6,97936334 | 58736 | 12673 | 53823 | 27190 | 41025 | 40639 | 35252 | 988 | 819 | 1127 | 650 |
| 16,46 | 260,3 | -1 | 656,52482 | 638,49093 | [M+NH4]+, [M | true | DG 18:2_20:5 | C41H66O5 | Lipid Species, Lip | 20212 | | 380920,344 | true | 0 | 0 | 0 | | | 445329 | 507227 | 470653 | 507329 | 533824 | 7,06210693 | 103799 | 7549 | 87380 | 187607 | 363519 | 380920 | 331651 | 1770277 | 992230 | 1749558 | 315795 |
| 8,99 | 242,7 | -0,2 | 550,38685 | 549,37957 | [M+H]+ | true | LPC 20:1 | C28H56NO7P | SmartFormula - r | 20222 | | 2982,71533 | true | 0 | 0 | 0 | | | 2750 | 3351 | 3050 | 2983 | 3010 | 7,08973922 | 676 | 142 | 464 | 0 | 3792 | 2572 | 2598 | 5724 | 5793 | 6083 | 8537 |
| 13,6 | 261,5 | 1,4 | 646,44434 | 645,43706 | [M+H]+ | true | PE10:0_19:2 | C34H64NO8P | SmartFormula - r | 20221 | LipidBlast-VS69-Pos.ms | 1438,89136 | true | 0 | 0 | 0 | | | 1976 | 2094 | 1830 | 2005 | 2217 | 7,09197015 | 285 | 172 | 546 | 172 | 1190 | 294 | 0 | 3518 | 17856 | 3612 | 1439 |
| 5,74 | 211,7 | -0,4 | 452,27755 | 451,27027 | [M+H]+ | true | LPE 16:1 | C21H42NO7P | SmartFormula - r | 20222 | | 1205,42749 | true | 0 | 0 | 0 | | | 1339 | 1471 | 1438 | 1256 | 1258 | 7,37364028 | 155 | 0 | 0 | 0 | 646 | 481 | 497 | 3720 | 2110 | 4242 | 5459 |
| 7,07 | 219,5 | -0,4 | 504,3083 | 503,30102 | [M+H]+ | true | LPE 20:3 | C25H46NO7P | Spectral Library - | 20222 | | 1480,49353 | true | 0 | 0 | 0 | | | 1480 | 1679 | 1494 | 1538 | 1746 | 7,46996625 | 237 | 0 | 0 | 210 | 1325 | 1352 | 1195 | 3842 | 2675 | 3670 | 4015 |
| 7,64 | 240,3 | 0,6 | 572,37102 | 571,36374 | [M+H]+ | true | LPC 22:4 | C30H54NO7P | Spectral Library - | 20222 | | 9156,2793 | true | 0 | 0 | 0 | | | 9156 | 10375 | 9467 | 9271 | 10825 | 7,52717863 | 527 | 129 | 283 | 643 | 5007 | 4664 | 4015 | 25786 | 16587 | 20255 | 35129 |
| 5,48 | 227,7 | 0 | 518,32437 | 517,31709 | [M+H]+ | true | LPC 18:3 | C26H48NO7P | Lipid Species, Sp | 20222 | | 18766,0781 | true | 0 | 0 | 0 | | | 20032 | 21224 | 18766 | 21481 | 22942 | 7,53855249 | 708 | 74 | 291 | 618 | 6068 | 4386 | 5298 | 63642 | 42607 | 46899 | 81015 |
| 18,49 | 286,5 | 0,7 | 696,65004 | 695,64252 | [M+H]+, [M+N | true | Cer20:1;3O/23:0;(2OH) | C43H85NO5 | Spectral Library - | 20212 | LipidBlast-VS69-Pos.ms | 15834,9346 | true | 0 | 142 | 0 | | | 16994 | 15835 | 15844 | 14658 | 17885 | 7,60140853 | 36015 | 10081 | 23688 | 21644 | 27621 | 27412 | 26519 | 1289 | 1648 | 2412 | 0 |
| 13,54 | 273,2 | -0,9 | 702,50614 | 701,49887 | [M+H]+ | true | PC 30:2 | C38H72NO8P | SmartFormula - r | 20222 | | 6,43E+03 | true | 0 | 0 | 0 | | | 6838 | 6427 | 7606 | 6898 | 7693 | 7,62478526 | 351 | 0 | 0 | 506 | 0 | 0 | 0 | 23382 | 18804 | 25291 | 18023 |
| 15,86 | 298,1 | -0,6 | 858,59955 | 857,59227 | [M+H]+ | true | PC 42:8 | C50H84NO8P | Spectral Library - | 20222 | | 37261,1992 | true | 0 | 0 | 0 | | | 43862 | 45345 | 44269 | 37261 | 40862 | 7,75164692 | 7125 | 868 | 6631 | 7454 | 11696 | 12623 | 10418 | 126610 | 78002 | 128912 | 60521 |
| 12,73 | 273,7 | -0,7 | 724,49024 | 723,48296 | [M+H]+ | true | PC 32:5 | C40H70NO8P | Lipid Species, Sm | 20222 | | 1137,72864 | true | 0 | 0 | 0 | | | 1159 | 1363 | 1162 | 1138 | 1269 | 7,85680067 | 150 | 0 | 0 | 0 | 0 | 149 | 0 | 3563 | 2375 | 4286 | 3128 |
| 15,2 | 275 | 1,2 | 738,50582 | 737,49855 | [M+H]+ | true | PE 36:5 | C41H72NO8P | SmartFormula - r | 20121 | | 42404,3633 | true | 0 | 0 | 0 | | | 42404 | 46910 | 53213 | 47062 | 48090 | 8,11275456 | 11133 | 1644 | 10813 | 5431 | 8831 | 8423 | 6330 | 133892 | 111089 | 159092 | 66109 |
| 16,36 | 276,8 | 1,4 | 718,53816 | 717,53089 | [M+H]+ | true | PE 34:1 | C39H76NO8P | Spectral Library - | 20211 | | 29606,9102 | true | 0 | 0 | 105 | | | 31392 | 38232 | 33760 | 31468 | 33943 | 8,23209102 | 19062 | 3260 | 16238 | 6050 | 11534 | 10301 | 7226 | 84813 | 75512 | 116348 | 29607 |
| 17,08 | 302 | -0,7 | 862,63143 | 861,62415 | [M+H]+ | true | PC 42:6 | C50H88NO8P | SmartFormula - r | 20222 | | 16612,6895 | true | 0 | 0 | 0 | | | 16613 | 19504 | 20113 | 17201 | 17359 | 8,52063078 | 5209 | 0 | 0 | 3850 | 6399 | 7287 | 4312 | 66254 | 43601 | 62203 | 16724 |
| 14,09 | 269 | 1,3 | 710,47521 | 709,46794 | [M+H]+ | true | PE 34:5 | C39H68NO8P | Spectral Library - | 20121 | | 11496,2227 | true | 0 | 0 | 0 | | | 13342 | 14127 | 12231 | 13969 | 11496 | 8,72646264 | 2103 | 0 | 1652 | 1527 | 2210 | 2375 | 1747 | 37679 | 29033 | 39029 | 22034 |
| 18,75 | 291,1 | 1,9 | 698,66544 | 697,6582 | [M+H]+, [M+N | true | Cer18:0;3O/25:0;(2OH) | C43H87NO5 | Spectral Library - | 20211 | LipidBlast-VS69-Pos.ms | 12146,748 | true | 0 | 605 | 451 | | | 11901 | 12794 | 11351 | 13503 | 14069 | 8,77233416 | 17781 | 13906 | 13842 | 19005 | 17095 | 19795 | 24071 | 2345 | 2037 | 3860 | 0 |
| 18,59 | 326 | 2,4 | 864,67044 | 846,63668 | [M+NH4]+, [M | true | ST28:1;O;Hex;FA20:5 | C54H86O7 | SmartFormula - r | 20221 | LipidBlast-VS69-Pos.ms | 7776,38916 | true | 0 | 0 | 0 | | | 7156 | 7992 | 7578 | 7875 | 9103 | 9,13591165 | 5861 | 1006 | 5364 | 6470 | 10270 | 10175 | 8967 | 15226 | 13808 | 20602 | 1533 |
| 16,24 | 250,1 | 3 | 556,49362 | 538,45967 | [M+NH4]+, [M | true | DG 14:0_16:1 | C33H62O5 | SmartFormula - r | 20220 | | 24682,3301 | true | 0 | 417 | 236 | | | 25004 | 29962 | 28365 | 24682 | 29688 | 9,21459936 | 5072 | 1113 | 4346 | 5544 | 12197 | 10473 | 10987 | 111094 | 55769 | 112997 | 32014 |
| 4,75 | 225,7 | 0,8 | 516,30836 | 515,30108 | [M+H]+ | true | LPC 18:4 | C26H46NO7P | Lipid Species, Sm | 20222 | | 1673,63062 | true | 0 | 0 | 0 | | | 2004 | 1952 | 2062 | 1722 | 1674 | 9,23929605 | 0 | 0 | 0 | 0 | 448 | 379 | 426 | 5113 | 2782 | 4146 | 7784 |
| 15,11 | 272,1 | 0,4 | 726,50654 | 725,49926 | [M+H]+ | true | PE19:0_16:4 | C40H72NO8P | Spectral Library - | 20222 | LipidBlast-VS69-Pos.ms | 10305,0176 | true | 0 | 0 | 0 | | | 19691 | 19311 | 16175 | 20812 | 18353 | 9,24450282 | 1388 | 697 | 3184 | 1027 | 10305 | 1522 | 939 | 23269 | 116167 | 18273 | 5588 |
| 19,1 | 276,6 | -2,9 | 668,61827 | 650,5844 | [M+NH4]+, [M | true | DG 16:0_22:1 | C41H78O5 | Spectral Library - | 20221 | | 5110,11572 | true | 0 | 300 | 0 | | | 5110 | 6160 | 4925 | 5958 | 5519 | 9,57191993 | 4658 | 964 | 2914 | 4551 | 4574 | 5852 | 6045 | 14913 | 10265 | 16214 | 2577 |

| | | | | | | | | | | | | | | | | | | | | | | | | | | | | | | | | | |
|---|---|---|---|---|---|---|---|---|---|---|---|---|---|---|---|---|---|---|---|---|---|---|---|---|---|---|---|---|---|---|---|---|---|
| 2,75 | 184,6 | 2 | 274,27426 | 273,26698 | [M+H]+ | true | Lauryldiethanolamine | C16H35NO2 | Spectral Library - | 20211 | MoNA-export-LC-MS-MS | 1411,95923 | true | 0 | 0 | 0 | 1491 | 1616 | 1821 | 1543 | 1420 | 9,72636331 | 2259 | 1412 | 1804 | 436 | 713 | 424 | 315 | 7696 | 682 | 822 | 478 |
| 8,48 | 241,5 | -0,5 | 574,3863 | 573,37903 | [M+H]+ | true | LPC 22:3 | C30H56NO7P | Spectral Library - | 20222 | | 1707,17798 | true | 0 | 0 | 0 | 2028 | 2083 | 2065 | 2219 | 1687 | 9,80905512 | 314 | 0 | 0 | 283 | 1239 | 1412 | 1707 | 4121 | 2690 | 3765 | 6263 |
| 7,12 | 247,5 | -0,9 | 594,37662 | 593,36935 | [M+H]+ | true | Spectral Match to 2-(5-Oxovaleryl)phosphatidylcholine from NIS | C29H56NO9P | SmartFormula - r | 20222 | ALL_GNPS_NO_PROPO( | 3305,33521 | true | 0 | 149 | 57 | 7061 | 9071 | 7367 | 8237 | 8074 | 9,89721003 | 628 | 216 | 451 | 367 | 376 | 285 | 0 | 6401 | 3305 | 4294 | 4696 |
| 17,79 | 282,1 | | 656,61877 | 655,61142 | [M+H]+, [M+N | true | Cer 18:0;O3/22:0;O | C40H81NO5 | Lipid Species, Sp | 20220 | | 14513,2988 | true | 0 | 156 | 179 | 14513 | 16225 | 15275 | 12470 | 15773 | 9,93108172 | 45050 | 7778 | 26875 | 16615 | 16828 | 20010 | 20453 | 1348 | 1549 | 1524 | 0 |
| 18,79 | 289,4 | | 710,66557 | 709,65733 | [M+H]+, [M+K] | true | Cer 18:1;O3/26:0;O | C44H87NO5 | Lipid Species, Sm | 20210 | | 14620,9561 | true | 0 | 0 | 0 | 13484 | 14858 | 13907 | 14905 | 17397 | 10,1865191 | 36067 | 9092 | 22038 | 18941 | 26176 | 26435 | 25228 | 1041 | 1402 | 1912 | 0 |
| 7,21 | 223,9 | 3,2 | 428,37357 | 427,36629 | [M+H]+ | true | C18Octadecanoyl-L-Carnitine | C25H49NO4 | Spectral Library - | 20210 | ALL_GNPS_NO_PROPO( | 3674,68628 | true | 0 | 0 | 0 | 3675 | 4546 | 4724 | 4141 | 4687 | 10,2123756 | 346 | 232 | 198 | 106 | 206 | 187 | 161 | 13618 | 14577 | 13885 | 13723 |
| 14,27 | 247 | 4 | 484,47268 | 483,46547 | [M+H]+, [M+N | true | Cer 16:0;O2/14:0 | C30H61NO3 | Lipid Species, Lip | 20220 | | 3597,30127 | true | 0 | 0 | 0 | 4437 | 3597 | 4013 | 4134 | 4730 | 10,2734563 | 8438 | 1818 | 7007 | 2723 | 4314 | 3939 | 3547 | 0 | 0 | 0 | 0 |
| 12,92 | 269,5 | -0,8 | 698,47522 | 697,46794 | [M+H]+ | true | PC 30:4 | C38H68NO8P | SmartFormula - r | 20122 | | 0 | true | 0 | 0 | 0 | 840 | 751 | 809 | 972 | 929 | 10,4282028 | 0 | 0 | 0 | 0 | 938 | 0 | 0 | 722 | 6657 | 0 | 0 |
| 14,98 | 269,8 | 0,2 | 714,5057 | 713,49842 | [M+H]+ | true | PE 34:3 | C39H72NO8P | Spectral Library - | 20222 | | 15177,7822 | true | 0 | 0 | 0 | 19142 | 15178 | 15639 | 18869 | 17405 | 10,4983843 | 1649 | 276 | 1280 | 708 | 0 | 950 | 847 | 55484 | 50033 | 73347 | 25753 |
| 15,61 | 300 | 0,3 | 882,60088 | 881,5936 | [M+H]+ | true | PC22:5_22:5 | C52H84NO8P | SmartFormula - r | 20122 | LipidBlast-VS69-Pos.ms | 11874,5039 | true | 0 | 0 | 0 | 13670 | 15251 | 12122 | 14151 | 11875 | 10,5568421 | 2918 | 352 | 2906 | 3617 | 5445 | 4758 | 4626 | 33329 | 37770 | 33681 | 21464 |
| 16,39 | 299,3 | -0,9 | 860,61529 | 859,60802 | [M+H]+ | true | PC 42:7 | C50H86NO8P | SmartFormula - r | 20122 | | 11532,71 | true | 0 | 0 | 0 | 11533 | 11757 | 12272 | 13971 | 14582 | 10,6899325 | 2595 | 352 | 2392 | 2648 | 4637 | 0 | 4071 | 45119 | 19556 | 39282 | 17439 |
| 15,62 | 260,7 | | 554,51403 | 553,50675 | [M+H]+ | true | Cer 18:1;O2/16:0;O | C34H67NO4 | Lipid Species, Sm | 20220 | | 5069,58984 | true | 0 | 0 | 0 | 5070 | 4728 | 6157 | 5640 | 5938 | 10,834397 | 13441 | 1123 | 9125 | 4833 | 5115 | 6980 | 7461 | 0 | 0 | 0 | 0 |
| 14,37 | 275,3 | | 634,57681 | 633,56936 | [M+H]+, [M+N | true | Cer 16:1;O2/24:2;O | C40H75NO4 | Lipid Species, Sm | 20210 | | 2532,6123 | true | 0 | 0 | 0 | 2966 | 3334 | 3638 | 3099 | 3850 | 10,8779099 | 3744 | 943 | 3773 | 1607 | 2659 | 2699 | 2349 | 0 | 0 | 0 | 0 |
| 14,08 | 266,1 | 2 | 684,45909 | 683,45181 | [M+H]+ | true | PE 32:4 | C37H66NO8P | Spectral Library - | 20221 | | 253,176788 | true | 0 | 0 | 0 | 5532 | 6088 | 4609 | 5837 | 4941 | 11,405447 | 178 | 253 | 1243 | 0 | 4451 | 223 | 0 | 0 | 45597 | 0 | 0 |
| 9,1 | 239,3 | -0,1 | 524,37063 | 523,36336 | [M+H]+ | true | LPC 18:0 | C26H54NO7P | SmartFormula - r | 20112 | | 765,78833 | true | 0 | 0 | 0 | 871 | 1052 | 766 | 888 | 912 | 11,4357704 | 172 | 143 | 163 | 376 | 277 | 319 | 192 | 1788 | 1525 | 2756 | 2626 |
| 18,18 | 282,9 | | 682,63405 | 681,62677 | [M+H]+, [M+N | true | Cer 18:1;O3/24:0;O | C42H83NO5 | Spectral Library - | 20220 | | 50508,6133 | true | 0 | 0 | 0 | 50509 | 59004 | 46546 | 49359 | 60252 | 11,5122127 | 137048 | 40275 | 86495 | 68753 | 71622 | 84635 | 87055 | 2964 | 3324 | 4483 | 0 |
| 17,47 | 277,4 | | 654,60302 | 653,59574 | [M+H]+ | true | Cer 18:1;O3/22:0;O | C40H79NO5 | SmartFormula - r | 20220 | | 12292,0313 | true | 0 | 0 | 0 | 12831 | 15032 | 12292 | 12207 | 15582 | 11,7634709 | 40166 | 5636 | 24465 | 11622 | 17419 | 17644 | 15556 | 0 | 0 | 0 | 0 |
| 15,23 | 291,1 | 0 | 818,56755 | 817,56028 | [M+H]+ | true | PC9:0_30:7 | C47H80NO8P | Spectral Library - | 10112 | LipidBlast-VS69-Pos.ms | 1778,9519 | true | 0 | 0 | 0 | 20030 | 20830 | 22439 | 16168 | 19105 | 11,8188627 | 0 | 563 | 2743 | 0 | 9819 | 1779 | 1220 | 0 | 148213 | 0 | 0 |
| 14,28 | 283,8 | 1,7 | 714,5512 | 713,54402 | [M+H]+, [M+N | true | HexCer 18:2;O2/16:0;O | C40H75NO9 | Spectral Library - | 20121 | | 1657,56335 | true | 0 | 0 | 0 | 2726 | 2100 | 2470 | 2864 | 2776 | 11,9544288 | 4987 | 1462 | 4165 | 1348 | 2493 | 2747 | 1658 | 0 | 0 | 0 | 0 |
| 18,43 | 274,3 | -3,5 | 690,6026 | 672,56862 | [M+NH4]+, [M | true | DG 18:2_22:2 | C43H76O5 | Lipid Species, Lip | 20210 | | 3240,41479 | true | 0 | 0 | 0 | 4347 | 3107 | 3610 | 3641 | 3792 | 12,0153115 | 1226 | 0 | 939 | 2485 | 2248 | 3240 | 3278 | 10835 | 5347 | 18203 | 1489 |
| 8,19 | 200,3 | 4,4 | 355,28427 | 354,27699 | [M+H]+ | true | Spectral Match to 2-Linoleoylglycerol from NIST14 | C21H38O4 | Spectral Library - | 20210 | ALL_GNPS_NO_PROPO( | 2380,59375 | true | 0 | 0 | 0 | 2919 | 3323 | 2381 | 3013 | 2720 | 12,1859544 | 0 | 0 | 0 | 208 | 1180 | 886 | 1097 | 7357 | 4586 | 9721 | 6238 |
| 7,02 | 248 | -1,1 | 610,37133 | 609,36405 | [M+H]+ | true | Spectral Match to 1-Palmitoyl-2-glutaryl phosphatidylcholine fro | C29H56NO10P | SmartFormula - r | 20221 | ALL_GNPS_NO_PROPO( | 504,536652 | true | 0 | 0 | 0 | 1461 | 1574 | 1688 | 1604 | 2004 | 12,3385559 | 314 | 94 | 209 | 125 | 87 | 121 | 0 | 1525 | 505 | 1008 | 711 |
| 14,48 | 274,5 | 0,9 | 748,49056 | 747,48329 | [M+H]+ | true | PE9:0_28:7 | C42H70NO8P | SmartFormula - r | 20122 | LipidBlast-VS69-Pos.ms | 17895,2656 | true | 0 | 0 | 0 | 27356 | 24608 | 22072 | 22912 | 19563 | 12,4720937 | 2272 | 677 | 3569 | 2098 | 12308 | 3490 | 2413 | 40675 | 111364 | 50416 | 17895 |
| 15,75 | 250,6 | 1,8 | 544,4573 | 526,42327 | [M+NH4]+, [M | true | TG8:0_10:0_10:0 | C31H58O6 | SmartFormula - r | 20211 | LipidBlast-VS69-Pos.ms | 11898,7949 | true | 0 | 6504 | 3404 | 14886 | 13248 | 14132 | 10519 | 12669 | 12,7387538 | 9529 | 12289 | 12648 | 9178 | 9188 | 11899 | 24195 | 6025 | 15332 | 0 | 11280 |
| 7,4 | 216,2 | 0,6 | 454,29332 | 453,28605 | [M+H]+ | true | LPE 16:0 | C21H44NO7P | Lipid Species, Sm | 20222 | | 1227,65137 | true | 0 | 0 | 0 | 1228 | 1468 | 1485 | 1593 | 1774 | 13,1861774 | 195 | 0 | 178 | 276 | 238 | 184 | 212 | 3932 | 2669 | 5509 | 4752 |
| 18,98 | 293,9 | | 712,68068 | 711,67373 | [M+H]+, [M+N | true | Cer 18:0;O3/26:0;O | C44H89NO5 | Lipid Species, Sp | 20220 | | 9330,2168 | true | 0 | 1316 | 1261 | 8762 | 8088 | 9359 | 9330 | 11490 | 13,5558835 | 13370 | 11774 | 11398 | 13162 | 15145 | 16969 | 17953 | 2805 | 2518 | 4030 | 981 |
| 14,32 | 272,3 | 0,9 | 736,49128 | 735,48401 | [M+H]+ | true | PE 36:6 | C41H70NO8P | SmartFormula - r | 20222 | | 6277,27734 | true | 0 | 0 | 0 | 8643 | 8346 | 9361 | 6277 | 8402 | 14,0330772 | 1348 | 374 | 1236 | 1397 | 2360 | 2393 | 1901 | 24772 | 23743 | 27556 | 13388 |
| 15,75 | 276,2 | 1,5 | 728,52131 | 727,51403 | [M+H]+ | true | PE15:1_20:2 | C40H74NO8P | SmartFormula - r | 20211 | LipidBlast-VS69-Pos.ms | 3131,33569 | true | 0 | 0 | 0 | 11797 | 9356 | 8303 | 8397 | 9226 | 15,0064229 | 0 | 550 | 3131 | 1007 | 9627 | 1292 | 1260 | 4948 | 67980 | 0 | 0 |
| 17 | 276,3 | | 628,58739 | 627,58011 | [M+H]+ | true | Cer 18:0;O3/20:0;O | C38H77NO5 | Lipid Species, Sm | 20210 | | 2633,12549 | true | 0 | 0 | 0 | 2091 | 2639 | 2799 | 3195 | 3020 | 15,4336705 | 8133 | 936 | 5571 | 2216 | 2824 | 3327 | 2957 | 0 | 0 | 0 | 0 |
| 10,9 | 145,9 | 17,4 | 223,06368 | 222,0564 | [M+H]+ | true | Spectral Match to Cyclotrisiloxane, hexamethyl- from NIST14 | C6H18O3Si3 | Spectral Library - | 20110 | ALL_GNPS_NO_PROPO( | 0 | true | 0 | 0 | 0 | 1302 | 1772 | 1174 | 1652 | 1490 | 16,5815628 | 325 | 0 | 0 | 0 | 0 | 0 | 0 | 0 | 0 | 0 | 0 |
| 9,13 | 247,2 | 0,9 | 600,40245 | 599,39518 | [M+H]+ | true | LPC24:4-SN2 | C32H58NO7P | SmartFormula - r | 20222 | LipidBlast-VS69-Pos.ms | 1123,42896 | true | 0 | 0 | 0 | 1155 | 1189 | 1220 | 1123 | 1638 | 16,731744 | 218 | 0 | 124 | 214 | 1120 | 959 | 928 | 2947 | 1992 | 2203 | 3772 |
| 14,63 | 260 | | 570,50989 | 569,50253 | [M+H]+, [M+N | true | Cer 18:1;O3/16:0;O | C34H67NO5 | Spectral Library - | 20220 | | 5548,18896 | true | 0 | 0 | 0 | 8354 | 8581 | 10510 | 6709 | 7505 | 17,1049649 | 12315 | 1596 | 9581 | 4117 | 6251 | 5548 | 4909 | 0 | 0 | 0 | 0 |
| 19,09 | 272,6 | -2 | 642,60303 | 624,56922 | [M+NH4]+, [M | true | DG 18:0/18:0 | C39H76O5 | Spectral Library - | 20221 | | 3736,88477 | true | 0 | 2385 | 2019 | 4252 | 3763 | 3133 | 4145 | 5053 | 17,2779442 | 5466 | 5110 | 3737 | 5159 | 4975 | 3078 | 9147 | 3326 | 3147 | 0 | 2058 |
| 11,78 | 272,7 | -1,1 | 746,47558 | 745,4683 | [M+H]+ | true | PC 34:8 | C42H68NO8P | SmartFormula - r | 20121 | | 268,16748 | true | 0 | 0 | 0 | 268 | 410 | 285 | 383 | 318 | 18,5123956 | 0 | 0 | 0 | 0 | 0 | 0 | 0 | 1138 | 416 | 859 | 896 |
| 13,95 | 276,3 | -0,5 | 726,50686 | 725,49958 | [M+H]+ | true | PC 32:4 | C40H72NO8P | Lipid Species, Sp | 20222 | | 3122,79395 | true | 0 | 0 | 0 | 27731 | 25015 | 26056 | 20458 | 34147 | 18,6207941 | 693 | 695 | 3123 | 0 | 14263 | 0 | 0 | 16226 | 140528 | 11677 | 0 |
| 15,74 | 259,6 | 1,9 | 590,45488 | 589,4476 | [M+H]+ | true | CerP 16:1;O2/16:0 | C32H64NO6P | Lipid Species, Sm | 20221 | | 4699,62549 | true | 0 | 0 | 0 | 9075 | 8156 | 9498 | 11447 | 6831 | 18,9660996 | 21537 | 3205 | 19657 | 2811 | 5107 | 4700 | 3435 | 0 | 0 | 0 | 0 |
| 11,39 | 266,3 | -1,3 | 696,45981 | 695,45254 | [M+H]+ | true | PC 30:5 | C38H66NO8P | Spectral Library - | 20121 | | 379,335663 | true | 0 | 0 | 0 | 414 | 570 | 515 | 666 | 384 | 22,5904212 | 0 | 0 | 0 | 0 | 0 | 0 | 0 | 1108 | 991 | 1249 | 1594 |
| 18,98 | 287,1 | | 692,65435 | 691,64708 | [M+H]+ | true | Cer 18:2;O2/26:0;O | C44H85NO4 | Lipid Species, Lip | 20120 | | 2453,29126 | true | 0 | 0 | 0 | 2733 | 4032 | 2219 | 2315 | 2589 | 26,3156354 | 5691 | 1252 | 3748 | 3054 | 3916 | 4766 | 3964 | 0 | 0 | 0 | 0 |
