## Supplementary table 9 for "Zoospore-derived extracellular vesicles in the flagellated stramenopile *Phytophthora parasitica*"

**Title: : LION-term enrichment analysis of EV samples**

| Term ID | Discription | Annotated | Significant | Expected | p-value | FDR q-value |
|---|---|---|---|---|---|---|
| LION:0012441 | N-acylsphingosines (ceramides) [SP0201] | 28 | 26 | 5.18 | 6.0e-23 | 4.56e-21 |
| LION:0000004 | sphingolipids [SP] | 29 | 26 | 5.36 | 5.5e-22 | 1.05e-20 |
| LION:0000077 | ceramides [SP02] | 29 | 26 | 5.36 | 5.5e-22 | 1.05e-20 |
| LION:0012082 | plasma membrane | 29 | 26 | 5.36 | 5.5e-22 | 1.05e-20 |
| LION:0002948 | fatty acid with 16-18 carbons | 61 | 27 | 11.28 | 3.0e-10 | 4.56e-09 |
| LION:0002966 | fatty acid with less than 2 double bonds | 62 | 27 | 11.47 | 5.1e-10 | 6.46e-09 |
| LION:0000464 | negative intrinsic curvature | 78 | 29 | 14.43 | 4.2e-09 | 4.56e-08 |
| LION:0000100 | fatty acid with 18 carbons or less | 69 | 27 | 12.76 | 1.5e-08 | 1.42e-07 |
| LION:0012009 | lipid-mediated signalling | 75 | 27 | 13.87 | 1.9e-07 | 1.60e-06 |
| LION:0002955 | fatty acid with 16 carbons | 28 | 15 | 5.18 | 3.3e-06 | 2.51e-05 |
| LION:0002900 | C16:1 | 14 | 9 | 2.59 | 9.2e-05 | 6.36e-04 |
| LION:0002969 | monounsaturated fatty acid | 35 | 14 | 6.47 | 0.00063 | 3.99e-03 |
| LION:0002921 | C18:0 | 6 | 5 | 1.11 | 0.00086 | 5.03e-03 |
| LION:0002968 | saturated fatty acid | 33 | 13 | 6.1 | 0.00135 | 7.33e-03 |
| LION:0002957 | fatty acid with 18 carbons | 32 | 12 | 5.92 | 0.00387 | 1.96e-02 |
| LION:0002882 | C16:0 | 14 | 6 | 2.59 | 0.02519 | 1.20e-01 |
| LION:0002922 | C18:1 | 12 | 5 | 2.22 | 0.04782 | 2.14e-01 |
| LION:0012080 | endoplasmic reticulum (ER) | 147 | 29 | 27.19 | 0.24355 | 1.00e+00 |
| LION:0002951 | fatty acid with more than 24 carbons | 4 | 1 | 0.74 | 0.56225 | 1.00e+00 |
| LION:0001737 | average transition temperature | 13 | 2 | 2.4 | 0.73449 | 1.00e+00 |
| LION:0002923 | C18:2 | 13 | 2 | 2.4 | 0.73449 | 1.00e+00 |
| LION:0080970 | average bilayer thickness | 15 | 2 | 2.77 | 0.80807 | 1.00e+00 |
| LION:0002970 | fatty acid with 2 double bonds | 19 | 2 | 3.51 | 0.90442 | 1.00e+00 |
| LION:0002950 | fatty acid with 22-24 carbons | 16 | 1 | 2.96 | 0.96798 | 1.00e+00 |
| LION:0080978 | average lateral diffusion | 17 | 1 | 3.14 | 0.97450 | 1.00e+00 |
| LION:0000038 | diacylglycerophosphoethanolamines [GP0201] | 27 | 2 | 4.99 | 0.97979 | 1.00e+00 |
| LION:0000607 | diacylglycerols [GL0201] | 22 | 1 | 4.07 | 0.99205 | 1.00e+00 |
| LION:0000094 | headgroup with neutral charge | 23 | 1 | 4.25 | 0.99373 | 1.00e+00 |
| LION:0000011 | glycerophosphoethanolamines [GP02] | 34 | 2 | 6.29 | 0.99554 | 1.00e+00 |
| LION:0012081 | mitochondrion | 34 | 2 | 6.29 | 0.99554 | 1.00e+00 |
| LION:0080979 | high lateral diffusion | 25 | 1 | 4.62 | 0.99613 | 1.00e+00 |
| LION:0001736 | low transition temperature | 32 | 1 | 5.92 | 0.99932 | 1.00e+00 |
| LION:0080968 | very low bilayer thickness | 41 | 1 | 7.58 | 0.99994 | 1.00e+00 |
| LION:0080980 | very high lateral diffusion | 43 | 1 | 7.95 | 0.99996 | 1.00e+00 |
| LION:0002945 | fatty acid with more than 18 carbons | 56 | 2 | 10.36 | 0.99998 | 1.00e+00 |
| LION:0002967 | polyunsaturated fatty acid | 58 | 2 | 10.73 | 0.99999 | 1.00e+00 |
| LION:0080982 | above average lateral diffusion | 64 | 2 | 11.84 | 1.00000 | 1.00e+00 |
| LION:0000030 | diacylglycerophosphocholines [GP0101] | 62 | 1 | 11.47 | 1.00000 | 1.00e+00 |
| LION:0080973 | below average bilayer thickness | 62 | 1 | 11.47 | 1.00000 | 1.00e+00 |
| LION:0001741 | below average transition temperature | 64 | 1 | 11.84 | 1.00000 | 1.00e+00 |
| LION:0000465 | neutral intrinsic curvature | 67 | 1 | 12.39 | 1.00000 | 1.00e+00 |
| LION:0000010 | glycerophosphocholines [GP01] | 84 | 1 | 15.54 | 1.00000 | 1.00e+00 |
| LION:0000095 | headgroup with positive charge / zwitter-ion | 118 | 3 | 21.83 | 1.00000 | 1.00e+00 |
| LION:0012010 | membrane component | 140 | 4 | 25.9 | 1.00000 | 1.00e+00 |
| LION:0000031 | 1-alkyl,2-acylglycerophosphocholines [GP0102] | 5 | 0 | 0.92 | 1.00000 | 1.00e+00 |
| LION:0000034 | monoacylglycerophosphocholines [GP0105] | 17 | 0 | 3.14 | 1.00000 | 1.00e+00 |
| LION:0000042 | monoacylglycerophosphoethanolamines [GP0205] | 7 | 0 | 1.29 | 1.00000 | 1.00e+00 |

| Discription | -LOG(FDR q-value) | FDR q-value |
|---|---|---|
| C18:1 | 0,669586227 | 2,14E-01 |
| C16:0 | 0,920818754 | 1,20E-01 |
| FA with 18 carbons | 1,707743929 | 1,96E-02 |
| saturated FA | 2,134896025 | 7,33E-03 |
| C18:0 | 2,298432015 | 5,03E-03 |
| monounsaturated FA | 2,399027104 | 3,99E-03 |
| C16:1 | 3,196542884 | 6,36E-04 |
| FA with 16 carbons | 4,600326279 | 2,51E-05 |
| FA with 18 carbons or less | 6,847711656 | 1,42E-07 |
| FA with less than 2 double bonds | 8,189767482 | 6,46E-09 |
| FA with 16-18 carbons | 8,341035157 | 4,56E-09 |
| sphingolipids [SP] | 19,9788107 | 1,05E-20 |
| ceramides [SP02] | 19,9788107 | 1,05E-20 |
| N-acylsphingosines (ceramides) [SP0201] | 20,34103516 | 4,56E-21 |
| lipid-mediated signalling | 5,795880017 | 1,60E-06 |
| negative intrinsic curvature | 7,341035157 | 4,56E-08 |
| plasma membrane | 19,9788107 | 1,05E-20 |

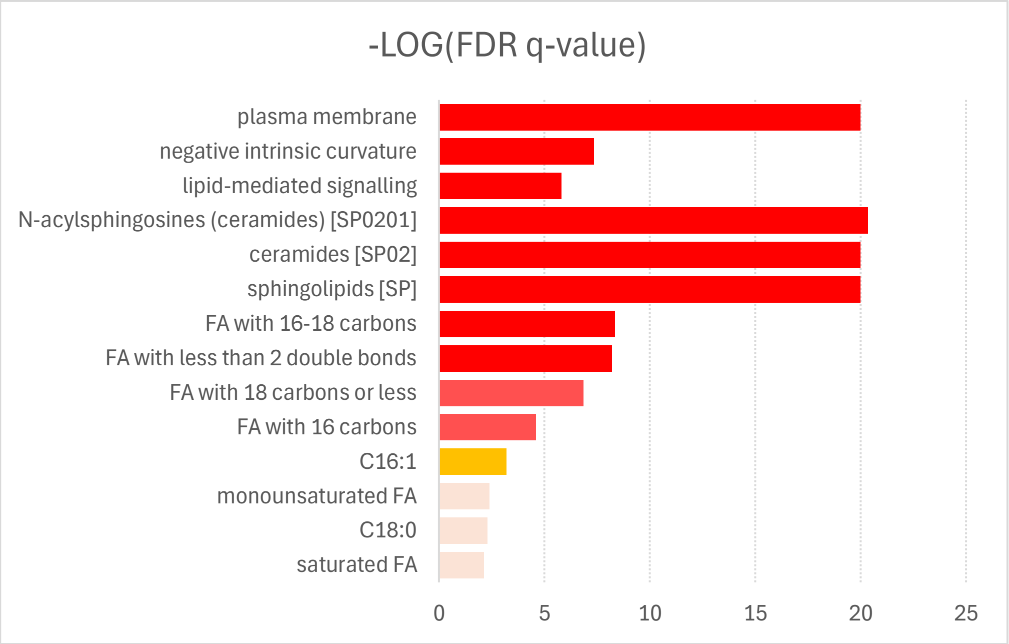

| | | | | | | |
|---|---|---|---|---|---|---|
| LION:0000259 | C14:0 | 7 | 0 | 1.29 | 1.00000 | 1.00e+00 |
| LION:0000466 | positive intrinsic curvature | 24 | 0 | 4.44 | 1.00000 | 1.00e+00 |
| LION:0000467 | contains ether-bond | 5 | 0 | 0.92 | 1.00000 | 1.00e+00 |
| LION:0000599 | lysoglycerophospholipids | 24 | 0 | 4.44 | 1.00000 | 1.00e+00 |
| LION:0001735 | very low transition temperature | 32 | 0 | 5.92 | 1.00000 | 1.00e+00 |
| LION:0001738 | high transition temperature | 3 | 0 | 0.55 | 1.00000 | 1.00e+00 |
| LION:0002929 | C20:4 | 8 | 0 | 1.48 | 1.00000 | 1.00e+00 |
| LION:0002930 | C20:5 | 20 | 0 | 3.7 | 1.00000 | 1.00e+00 |
| LION:0002947 | fatty acid with 13-15 carbons | 11 | 0 | 2.03 | 1.00000 | 1.00e+00 |
| LION:0002949 | fatty acid with 19-21 carbons | 36 | 0 | 6.66 | 1.00000 | 1.00e+00 |
| LION:0002953 | fatty acid with 14 carbons | 8 | 0 | 1.48 | 1.00000 | 1.00e+00 |
| LION:0002956 | fatty acid with 17 carbons | 8 | 0 | 1.48 | 1.00000 | 1.00e+00 |
| LION:0002958 | fatty acids with 19 carbons | 6 | 0 | 1.11 | 1.00000 | 1.00e+00 |
| LION:0002959 | fatty acid with 20 carbons | 30 | 0 | 5.55 | 1.00000 | 1.00e+00 |
| LION:0002961 | fatty acid with 22 carbons | 7 | 0 | 1.29 | 1.00000 | 1.00e+00 |
| LION:0002963 | fatty acid with 24 carbons | 7 | 0 | 1.29 | 1.00000 | 1.00e+00 |
| LION:0002965 | fatty acid with 26 carbons | 3 | 0 | 0.55 | 1.00000 | 1.00e+00 |
| LION:0002971 | fatty acid with 3 double bonds | 8 | 0 | 1.48 | 1.00000 | 1.00e+00 |
| LION:0002972 | fatty acid with 4 double bonds | 10 | 0 | 1.85 | 1.00000 | 1.00e+00 |
| LION:0002973 | fatty acid with 5 double bonds | 24 | 0 | 4.44 | 1.00000 | 1.00e+00 |
| LION:0002974 | fatty acid with 6 double bonds | 3 | 0 | 0.55 | 1.00000 | 1.00e+00 |
| LION:0002976 | fatty acid with more than 3 double bonds | 43 | 0 | 7.95 | 1.00000 | 1.00e+00 |
| LION:0002977 | fatty acid with 3-5 double bonds | 39 | 0 | 7.21 | 1.00000 | 1.00e+00 |
| LION:0002978 | fatty acid with more than 5 double bonds | 4 | 0 | 0.74 | 1.00000 | 1.00e+00 |
| LION:0022234 | C17:1 | 3 | 0 | 0.55 | 1.00000 | 1.00e+00 |
| LION:0022236 | C17:2 | 4 | 0 | 0.74 | 1.00000 | 1.00e+00 |
| LION:0080969 | low bilayer thickness | 22 | 0 | 4.07 | 1.00000 | 1.00e+00 |
| LION:0080971 | high bilayer thickness | 3 | 0 | 0.55 | 1.00000 | 1.00e+00 |
| LION:0080977 | low lateral diffusion | 4 | 0 | 0.74 | 1.00000 | 1.00e+00 |
