## Supplementary table 10 for "Zoospore-derived extracellular vesicles in the flagellated stramenopile *Phytophthora parasitica*"

**Title: Univariate analysis of lipid species intensity (P < 0.05)**

| Lipids | f.value | p.value | -log10(p) | FDR | Tukey's HSD |
|---|---|---|---|---|---|
| Cer 16:0;O2/14:0 | 4575.2 | 5.8222e-13 | 12.235 | 1.8534e-11 | EV-CB; FL-CB; FL-EV |
| Cer 16:0;O2/16:0 | 567.3 | 2.4031e-09 | 8.6192 | 3.0599e-08 | EV-CB; FL-CB; FL-EV |
| Cer 16:0;O2/18:0 | 9.8792 | 0.0068989 | 2.1612 | 0.015502 | EV-CB; FL-CB |
| Cer 16:0;O2/20:0 | 12.115 | 0.0037961 | 2.4207 | 0.0092956 | EV-CB; FL-CB |
| Cer 16:0;O2/22:0 | 10.244 | 0.0062189 | 2.2063 | 0.014141 | EV-CB; FL-CB |
| Cer 16:0;O2/22:1 | 16334 | 3.5929e-15 | 14.445 | 2.2875e-13 | EV-CB; FL-CB; FL-EV |
| Cer 16:1;O2/16:0 | 2057.8 | 1.4167e-11 | 10.849 | 2.7059e-10 | EV-CB; FL-CB; FL-EV |
| Cer 16:1;O2/18:0 | 130.11 | 7.9129e-07 | 6.1017 | 7.197e-06 | EV-CB; FL-CB |
| Cer 16:1;O2/20:0 | 111.69 | 1.4291e-06 | 5.8449 | 1.1868e-05 | EV-CB; FL-CB; FL-EV |
| Cer 16:1;O2/22:0 | 79.502 | 5.2658e-06 | 5.2785 | 3.4682e-05 | EV-CB; FL-CB |
| Cer 16:1;O2/22:1 | 137.83 | 6.3259e-07 | 6.1989 | 6.0412e-06 | EV-CB; FL-CB |
| Cer 16:1;O2/22:2 | 121.12 | 1.0444e-06 | 5.9811 | 9.0677e-06 | EV-CB; FL-CB |
| Cer 16:1;O2/22:2;O | 1949.1 | 1.7595e-11 | 10.755 | 2.8005e-10 | EV-CB; FL-CB; FL-EV |
| Cer 16:1;O2/24:1 | 161.13 | 3.4431e-07 | 6.463 | 3.4613e-06 | EV-CB; FL-CB |
| Cer 16:1;O2/24:2;O | 3278.5 | 2.2051e-12 | 11.657 | 6.0168e-11 | EV-CB; FL-CB; FL-EV |
| Cer 18:0;O3/20:0;O | 5460.5 | 2.8711e-13 | 12.542 | 1.0968e-11 | EV-CB; FL-CB; FL-EV |
| Cer 18:0;O3/22:0;O | 19.96 | 0.00077678 | 3.1097 | 0.0026029 | EV-CB; FL-CB |
| Cer 18:0;O3/24:0;O | 83.744 | 4.3188e-06 | 5.3646 | 3.0551e-05 | EV-CB; FL-CB |
| Cer 18:0;O3/26:0;O | 67.516 | 9.7865e-06 | 5.0094 | 5.6643e-05 | EV-CB; FL-CB |
| Cer 18:1;O2/16:0;O | 2287.4 | 9.2862e-12 | 11.032 | 1.9707e-10 | EV-CB; FL-CB; FL-EV |
| Cer 18:1;O3/16:0;O | 18750 | 2.0695e-15 | 14.684 | 1.9764e-13 | EV-CB; FL-CB; FL-EV |
| Cer 18:1;O3/22:0;O | 7576.3 | 7.7536e-14 | 13.11 | 3.7024e-12 | EV-CB; FL-CB; FL-EV |
| Cer 18:1;O3/24:0;O | 17.982 | 0.0010964 | 2.96 | 0.0033775 | EV-CB; FL-CB |
| Cer 18:1;O3/26:0;O | 20.731 | 0.00068431 | 3.1647 | 0.002334 | EV-CB; FL-CB |
| Cer 18:2;O2/24:0;O | 69.561 | 8.7429e-06 | 5.0583 | 5.3063e-05 | EV-CB; FL-CB |
| Cer 18:2;O2/26:0;O | 2001 | 1.5841e-11 | 10.8 | 2.7506e-10 | EV-CB; FL-CB; FL-EV |
| Cer18:0;3O/25:0;(2OH) | 12.783 | 0.0032265 | 2.4913 | 0.0081086 | EV-CB; FL-CB |
| Cer18:1;3O/23:0;(2OH) | 14.866 | 0.0020206 | 2.6945 | 0.0055133 | EV-CB; FL-CB |
| Cer19:3;2O/18:0 | 18.22 | 0.0010502 | 2.9787 | 0.0033432 | EV-CB; FL-CB |
| Cer20:1;3O/23:0;(2OH) | 18.791 | 0.00094876 | 3.0228 | 0.0030714 | EV-CB; FL-CB |
| CerP 16:1;O2/16:0 | 51145 | 3.7402e-17 | 16.427 | 7.1438e-15 | EV-CB; FL-CB; FL-EV |
| HexCer 18:2;O2/16:0;O | 1638.9 | 3.5141e-11 | 10.454 | 5.163e-10 | EV-CB; FL-CB; FL-EV |
| DG 16:0_22:1 | 17.474 | 0.0012038 | 2.9194 | 0.0036444 | EV-CB; FL-CB |
| DG 20:4/20:4 | 22.492 | 0.00051973 | 3.2842 | 0.0018383 | EV-CB; FL-CB |
| DG20:4_20:5 | 29.008 | 0.00021566 | 3.6662 | 0.00095792 | EV-CB; FL-CB; FL-EV |
| DG20:5_20:5 | 33.024 | 0.00013625 | 3.8657 | 0.00063472 | EV-CB; FL-CB; FL-EV |
| LPC 16:0 | 14.114 | 0.002378 | 2.6238 | 0.0063084 | EV-CB; FL-CB |
| LPC 18:3 | 10.901 | 0.0051923 | 2.2846 | 0.012244 | EV-CB; FL-EV |
| LPC 20:4 | 13.085 | 0.0030045 | 2.5222 | 0.0076513 | EV-CB; FL-EV |
| LPC20:5 | 17.319 | 0.0012392 | 2.9068 | 0.0036444 | EV-CB; FL-EV |
| LPE 20:4 | 9.4645 | 0.007789 | 2.1085 | 0.016906 | EV-CB |
| PC 16:0_18:1 | 40.455 | 6.5547e-05 | 4.1834 | 0.00033836 | FL-CB; FL-EV |
| PC 30:0 | 11.573 | 0.0043527 | 2.3612 | 0.010392 | FL-CB |
| PC 30:5 | 852.88 | 4.7485e-10 | 9.3234 | 6.4783e-09 | EV-CB; FL-CB |
| PC 32:1 | 49.708 | 3.0767e-05 | 4.5119 | 0.00016323 | EV-CB; FL-CB; FL-EV |
| PC 32:2 | 19.563 | 0.00083048 | 3.0807 | 0.0027349 | FL-CB; FL-EV |
| PC 34:2 | 23.254 | 0.00046397 | 3.3335 | 0.0016907 | FL-CB; FL-EV |
| PC 34:5 | 55.962 | 1.9803e-05 | 4.7033 | 0.00010807 | EV-CB; FL-CB |
| PC 34:8 | 359.73 | 1.4625e-08 | 7.8349 | 1.6674e-07 | EV-CB; FL-CB |
| PC 36:3 | 63.139 | 1.2599e-05 | 4.8997 | 7.0778e-05 | EV-CB; FL-CB; FL-EV |
| PC 36:4 | 27.853 | 0.00024869 | 3.6043 | 0.0010556 | EV-CB; FL-CB; FL-EV |
| PC 36:5 | 31.27 | 0.00016543 | 3.7814 | 0.0007523 | EV-CB; FL-CB |
| PC 36:6 | 75.845 | 6.2985e-06 | 5.2008 | 4.0101e-05 | EV-CB; FL-CB |
| PC 36:7 | 91.223 | 3.1137e-06 | 5.5067 | 2.3789e-05 | EV-CB; FL-CB |
| PC 38:6 | 14.535 | 0.0021692 | 2.6637 | 0.0058354 | EV-CB; FL-EV |
| PC 38:7 | 36.188 | 9.8139e-05 | 4.0082 | 0.00046861 | EV-CB; FL-CB |
| PC 38:8 | 69.254 | 8.8901e-06 | 5.0511 | 5.3063e-05 | EV-CB; FL-CB |
| PC 40:5 | 2524.4 | 6.2645e-12 | 11.203 | 1.4957e-10 | EV-CB; FL-EV |
| PC 40:7 | 36.637 | 9.3879e-05 | 4.0274 | 0.00045977 | EV-CB; FL-CB |
| PC14:1_30:8 | 14.954 | 0.0019833 | 2.7026 | 0.0054901 | EV-CB; FL-CB |
| PC16:3_26:6 | 13.393 | 0.0027974 | 2.5533 | 0.0072202 | EV-CB; FL-CB |
| PC16:4_20:5 | 11.884 | 0.0040217 | 2.3956 | 0.0097234 | EV-CB; FL-EV |
| PC18:4_20:5 | 108.96 | 1.5722e-06 | 5.8035 | 1.2512e-05 | EV-CB; FL-CB |
| PC18:5_24:5 | 210.47 | 1.21e-07 | 6.9172 | 1.284e-06 | EV-CB; FL-CB; FL-EV |
| PC20:4_20:5 | 28.405 | 0.00023217 | 3.6342 | 0.0010078 | EV-CB; FL-CB; FL-EV |
| PC20:5_20:5 | 82.82 | 4.5057e-06 | 5.3462 | 3.0735e-05 | EV-CB; FL-CB; FL-EV |
| PC22:5_22:5 | 12.668 | 0.0033164 | 2.4793 | 0.0082263 | FL-CB |
| PC40:10 | 22.219 | 0.0005417 | 3.2662 | 0.0018812 | EV-CB; FL-EV |

| | | | | | |
|---|---|---|---|---|---|
| PCO-20:5 | 17.315 | 0.0012403 | 2.9065 | 0.0036444 | EV-CB; FL-EV |
| PE 32:1 | 26.177 | 0.0003087 | 3.5105 | 0.0012818 | EV-CB; FL-EV |
| PE 34:1 | 23.393 | 0.00045464 | 3.3423 | 0.0016907 | EV-CB; FL-EV |
| PE 36:4 | 37.062 | 9.0047e-05 | 4.0455 | 0.0004526 | EV-CB; FL-CB |
| PE 36:5 | 23.179 | 0.00046916 | 3.3287 | 0.0016907 | EV-CB; FL-CB; FL-EV |
| PE 38:5 | 18.127 | 0.001068 | 2.9714 | 0.0033442 | EV-CB; FL-CB |
| ST28:1;O;Hex;FA20:5 | 16.286 | 0.0015117 | 2.8205 | 0.0043749 | EV-CB; FL-CB |
| Lauryldiethanolamine | 15.009 | 0.0019605 | 2.7076 | 0.0054901 | EV-CB; FL-EV |
| 2-Linoleoylglycerol | 358.41 | 1.4841e-08 | 7.8285 | 1.6674e-07 | EV-CB; FL-EV |
| Adenosine | 13.864 | 0.0025137 | 2.5997 | 0.006577 | FL-CB; FL-EV |
| C16Hexadecanoyl-L-Carnitine | 90.157 | 3.257e-06 | 5.4872 | 2.3927e-05 | EV-CB; FL-CB |
| C18Octadecanoyl-L-Carnitine | 15.722 | 0.001692 | 2.7716 | 0.0048235 | FL-CB; FL-EV |
